# A statistical framework for system-level synthesis

**DOI:** 10.64898/2026.10.19.760244

**Authors:** M. Senthil Kumar, Filip Garbicz, Rubén Carrasco, Rafael A. Irizarry

## Abstract

Simultaneous profiling of thousands of biological features is now routine. Yet transforming the data into interpretable models of system behavior remains a fundamental bottleneck as current analyses generally identify feature-wise associations or extract latent structure without anchoring it to the variables that define the scientific question. Here we propose a framework for system-level synthesis that learns and predicts overarching processes, and reliably identifies participating features without data reuse. We develop Smoothing Spline Analysis of Variance–Singular Value Decomposition to learn overarching processes as smooth, flexible functions of nominal, ordinal and continuous covariates and their interactions. This unlocks prediction from covariates alone, while reserving measurements for conditional inference of feature participation. In applications, the framework recovered established biology while generating predictions and testable hypotheses. By linking high-dimensional variation to covariates, the framework advances analysis beyond feature-wise associations and unsupervised patterns, toward interpretable system-level models that vary with biological and experimental context.

## 1 Introduction

Characterizing the overarching processes that govern the dynamics of complex biological systems is central to mechanistic understanding, accurate prediction, and ultimately, informed control. These are fundamental objectives of systems biology^1–5^. Continued advancements in measurement technologies^6–9^, together with the growing availability of large-scale data repositories^10–16^, have enabled the simultaneous profiling of thousands of biological features in diverse molecular, cellular, organismal and ecological systems, alongside covariates such as treatment, disease status, age and environmental exposures. Yet our ability to measure biological systems at ever greater scale has outpaced our ability to transform the resulting data into a holistic understanding of their behavior.

Two broad paradigms have played central roles in the analysis of high-dimensional biological data over the last three decades: feature-wise regressions^17–20^ and methods for characterizing joint variation among features, including matrix factorization^21–25^. But the bottleneck to synthesis persists because the former studies one feature at a time, while the latter generally extracts dominant patterns of variation without directly tying them to the variables defining the scientific question.

*Feature-wise methods* offer interpretation, prediction and statistical inference by modeling individual features with respect to study design, but characterize the system one feature at a time, ignoring between-feature correlations. Enrichment analyses can organize these feature-wise associations into established biological programs by testing predefined sets of features for coordinated association^26,27^, but do not characterize the role of these programs, or subsets of their features, in the dominant higher-order processes that drive the joint behavior of the measured features. Their interpretation is also constrained by the scope and resolution of the knowledge bases used to define these feature sets. On the other hand, ANOVA-simultaneous component analysis^28,29^ applies dimension reduction to fitted effect matrices obtained from feature-wise regressions or ANOVA. However, separate factorizations can fragment an overarching process among individual effects. Moreover, factorizing fitted effect matrices shifts the objective from explaining joint variation in the observed measurements to explaining variation in estimated effects, allowing estimation variability, including that of null features, to influence the resulting components. These methods also provide limited flexibility for capturing smooth nonlinear functions of continuous covariates.

Conversely, *matrix factorization* techniques leverage joint variation among features to recover low-dimensional representations that capture dominant variation in the measurements. This objective has important consequences for modern high-throughput profiling technologies, which are deliberately designed to measure features broadly rather than restrict attention to those already known to participate in a process of interest. Consequently, in many studies only a small fraction of the measured features may participate in the biological process under investigation. In such settings, the coordinated signal among the relevant features may be overwhelmed by variation among the many unrelated features. Sparse matrix factorization^30^ and nonnegative matrix factorization^22,25^ seek more interpretable representations through sparsity of feature loadings or non-negativity, respectively, but neither constraint establishes the biological meaning of a process or prevents components from capturing technical variation, the effect sizes of which can rival or even exceed biological variation in high-throughput measurements^31^. Other multivariate approaches incorporate covariates more directly. Canonical correlation analysis and partial least squares regression^32,33^ associate linear combinations of covariates with linear combinations of measured features, but because both sides of the representation are transformed, the resulting covariate directions may themselves be difficult to interpret. Covariate-aware factorization methods such as MEFISTO^34^ incorporate observed covariates into latent-factor estimation, but accommodate more restricted covariate structures and do not necessarily render the resulting factors interpretable in terms of experimental or observational design. Finally, existing matrix factorization methods generally do not support prediction and inference of feature loadings under general study designs while avoiding data reuse that can invalidate downstream inference. Post-selection inference methods, including jackstraw^35^, data thinning^36^ and data fission^37^, enable inference following data-driven discovery, but primarily address inference after a representation has been learned rather than the interpretable construction and transfer of the underlying processes themselves.

Thus, achieving mechanistic understanding of complex biological systems continues to be constrained by the lack of analytical frameworks capable of delivering system-level synthesis from multivariate biological data. By *system-level synthesis*, we refer to reconstructing overarching processes interpretably and performing statistical inference on the features that participate in them. Both attributes are essential. Without clear interpretation of the underlying processes, statistical identification of participating features does not provide system-level synthesis.

We are interested in variables that provide reliable context for biological interpretation. Interpretation based on measured feature abundances is complicated for two reasons. First, abundance does not necessarily identify biological function. In molecular systems, for example, function can depend on molecular state, interacting partners, regulators and subcellular localization^38–40^. Second, measured abundances can be sensitive to unforeseen technical sources of structured variation introduced during an experiment^31,41–43^. Thus, even when features vary together, the biological meaning of the resulting system-level process may remain ambiguous from the measurements alone.

Study covariates, however, provide context for interpretation. Variables such as treatment, disease status, time and environmental exposures encode expert knowledge about each study unit and are defined independently of the high-throughput measurements. They therefore provide reliable coordinates for describing biological variation. Indeed, practitioners routinely interpret individual features and prioritize hypotheses for further investigation by examining their behavior relative to these covariates. Extending this familiar principle from individual features to system-level processes provides a natural approach to building interpretable system-level models.

We therefore seek to build system-level models in which overarching processes, parameterized by covariates, capture dominant joint variation among the measured features. The properties of the covariates inform the structure and regularization of the learned processes themselves. For example, continuous covariates such as time can induce smoothness across nearby observations, helping distinguish coherent process-level variation from local stochastic fluctuations. At the same time, this smoothness can accommodate flexible nonlinear trajectories and interactions without requiring prespecified functional forms.

Finally, statistically modeling system-level processes as smooth functions of covariates provides a natural bridge to prediction and inference. Once learned, the processes/functions can be evaluated for covariates from new, not yet seen observations, thereby allowing reconstruction of processes in external data from their covariates alone. Leaving the corresponding biological measurements unused during process reconstruction reserves them for conditional inference of feature participation without data reuse. Thus, feature participation can be tested against processes that were defined independently of the measurements used for inference. This separation between process prediction and feature-level inference has not previously been leveraged for conditional inference of feature participation without data reuse.

Based on the above considerations, we propose an analytical paradigm for high dimensional data that explicitly targets system-level synthesis, establish a three-step framework for achieving it (**Fig. 1A**), and develop Smoothing Spline Analysis of Variance–Singular Value Decomposition (SSANOVA-SVD) (**Fig. 1B-C**) to reconstruct interpretable system-level processes from high-dimensional measurements. We apply the three-step framework and SSANOVA-SVD to data from diverse molecular, cellular, population and ecological systems. The results recapitulated established biological phenomena, supported statistical inference on feature participation using independent measurements, while revealing previously underappreciated structure that generated new, testable hypotheses. Our proposals thus provide a data-driven route to system-level models, complementing knowledge-driven mechanistic models^1,40,44,45^.

**Figure 1:**
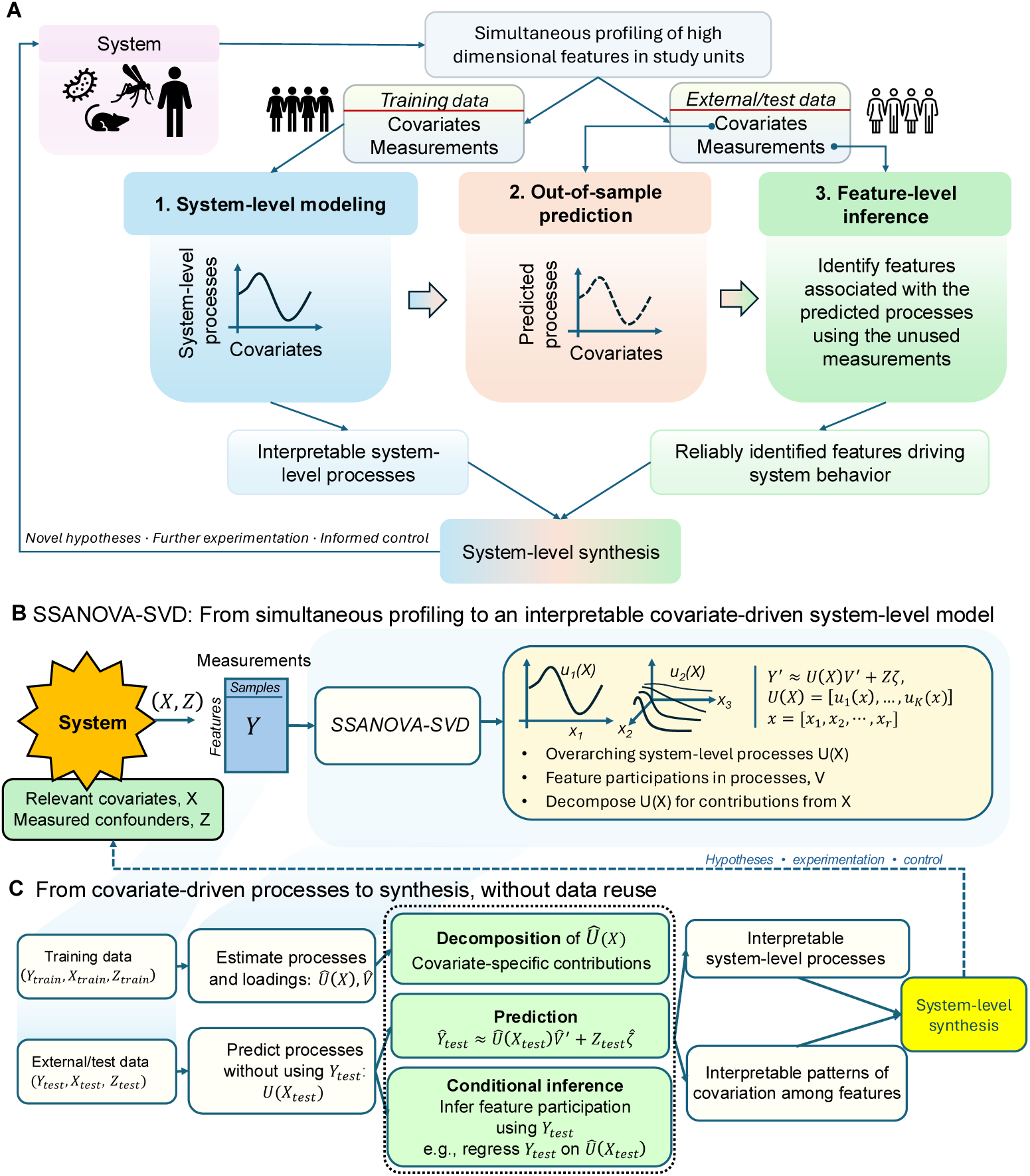
A three-step framework for system-level synthesis, and SSANOVA-SVD for learning interpretable overarching processes. **A**. Simultaneous measurements and their covariates are divided into training and external/test data. (1) The system-level modeling step uses the training data to learn an interpretable representation of system-level processes as smooth functions of covariates. (2) The learned models are used to predict the processes in external data from covariates alone, leaving the corresponding biological measurements unused. (3) The unused measurements are then used for conditional inference of feature participation in the predicted processes without double dipping. The collection of interpretable system-level processes and reliable information on the features driving system behavior provides system-level synthesis, generating new hypotheses for further experimentation, model refinement, and enabling informed control over the system. **B**. SSANOVA-SVD serves the system-level modeling step by reconstructing overarching processes as smooth, flexible functions of relevant covariates X, while accounting for measured nuisance variation Z. Each reconstructed process describes coordinated variation among the measured features through its corresponding feature loadings. The processes can be decomposed into contributions due to covariates and their interactions. **C**. Processes learned from the training data are used to predict processes in external/test data from covariates alone. The predicted processes can be combined with the learned feature loadings to predict external measurements, while the unused measurements support conditional inference of feature participation. This separation of process reconstruction from feature-level inference avoids data reuse, and provides the complementary information required for system-level synthesis, supporting hypothesis generation, further experimentation, and informed control.

## 2 Results

### 2.1 A three-step framework for system-level synthesis

We propose an analytical paradigm that explicitly targets system-level synthesis, by building system-level models in which overarching processes are reconstructed interpretably, and statistical inference can be performed on the features participating in them. Our proposed framework operates in three steps (**Fig. 1A**). First, system-level models are learned from the data, representing overarching processes as interpretable smooth functions of nominal, ordinal and continuous covariates and their interactions. Second, the processes are predicted in external data from their covariates alone. Third, the unused measurements from external data are used for conditional inference of feature participation, thereby enabling statistical inference without reusing the measurements that defined or predicted the processes. We developed SSANOVA-SVD to serve the first step of the framework (**Fig. 1B,C**).

#### Overview of SSANOVA-SVD

SSANOVA-SVD is designed to build the system-level models required by the first step of our framework: a small number of interpretable overarching processes that capture coordinated variation among many measured features while remaining interpretable with respect to the experimental or observational variables that define the scientific question. A central assumption is that biologically meaningful system-level processes vary in a structured manner over these covariates. For continuous or ordered covariates, such as time, age, disease progression, or spatial location, we therefore favor processes that vary smoothly across nearby observations, while allowing flexible nonlinear trajectories and interactions. This distinguishes coherent process-level variation from local fluctuations without requiring the functional form of the process to be specified in advance. Unlike feature-wise methods, which characterize these relationships one feature at a time, and conventional matrix factorization, which captures joint variation without necessarily targeting variation associated with the covariates of interest, SSANOVA-SVD directly reconstructs joint variation as smooth, flexible functions of those covariates.

We consider simultaneously profiled data from *n* study units and *p* features, represented by an *n* × *p* measurement matrix *Y*, whose rows correspond to study units and columns to measured features. For each study unit *i*, let *x_i_* denote the set of *b* covariates with respect to which the system is to be characterized, such as treatment, time, disease stage or spatial location, and let *z_i_* denote the set of *m* measured variables representing nuisance variation, such as batch effects. The covariates *x_i_* provide the coordinates for interpreting system behavior, whereas *z_i_* allows measured nuisance variation to be accounted for separately rather than incorporated into the reconstructed processes.

Rather than assigning an unconstrained latent score to each study unit, we represent system-level process *k* by a function *u_k_*(·) of the original covariates. Evaluating this function at the observed covariates {*x*_1_*, … , x_n_*} yields the *n*-dimensional vector

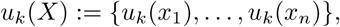

whose entries represent the value of process *k* for each study unit. These vectors form the columns of the *n* × *K* matrix *U* (*X*). Parameterizing each process directly by the original covariates is central to the framework. It makes the reconstructed processes interpretable in terms of the study design, permits their decomposition into contributions from individual covariates and their interactions, and allows the processes to be evaluated for study units not used to fit the model.

Each process has a corresponding *p*-dimensional feature-loading vector *v_k_*, whose entries describe how strongly each of the measured features participates in that process. The loading vectors form the columns of the *p* × *K* matrix *V* . Thus, *U* (*X*) describes how a small number of overarching processes vary across the biological or experimental conditions represented by *X*, whereas *V* connects those processes back to the individual measured features. These two levels provide the complementary information required for system-level synthesis: an interpretable representation of system behavior and information about the features participating in it.

The measurements are then represented as

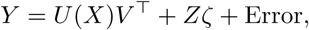

where *Z* is an *n* × *m* design matrix constructed from the nuisance variables and *ζ* is an *m* × *p* matrix containing their feature-specific effects. The term *U* (*X*)*V ^⊤^* therefore captures coordinated feature variation associated with the covariates used to characterize the system, while *Zζ* accounts separately for measured nuisance variation. This separation helps prevent known technical or other unwanted sources of variation from determining the dominant reconstructed processes.

Together, the functions 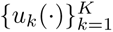 and their corresponding feature loadings 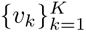 define the ***system-level model*** . We estimate this model by balancing faithful reconstruction of joint variation in the measurements with regularization of the system-level processes. The SSANOVA^46^ regularization encodes the smoothness and structural assumptions described above according to the properties of the covariates, allowing nominal, ordinal and continuous covariates, and their interactions, to be represented within the same framework. In particular, continuous effects can vary smoothly and nonlinearly without requiring their functional forms to be specified in advance. Thus, rather than first estimating feature-specific effects and subsequently searching for multivariate structure among them, SSANOVA-SVD learns the overarching processes directly from the joint measurements while constraining their structure through the covariates that provide their scientific interpretation.

The functions *u_k_*(·) are in general infinite-dimensional. For computational efficiency, we compute the rank-*K* factorization by sequentially solving constrained rank-one optimization problems, enforcing orthogonality to previously estimated processes at each step. A representer theorem reduces each rank-one optimization to a finite-dimensional problem (**Supplementary Methods: SSANOVA-SVD**).

The number of processes, *K*, is selected by cross-validation based on the ability of the learned system-level model 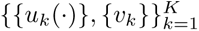 to predict measurements in held-out observations from their covariates alone. This criterion aligns model selection with the intended practical use of the representation: processes are retained when the covariate-parameterized model captures structure that transfers beyond the observations used for estimation, rather than simply improving reconstruction of the training measurements.

Importantly, the ability to evaluate *u_k_*(·) in new study units also connects system-level modeling to the subsequent statistical inference step of our framework. For a new study unit, the processes can be predicted from its covariates without using its high-dimensional measurements. Those measurements therefore remain unused and can subsequently support conditional statistical inference on feature participation without reusing the data that defined or predicted the processes. Thus, the same covariate-parameterized representation that provides interpretation also enables out-of-sample prediction and separates process reconstruction from subsequent feature-level inference.

Details of the model, estimation algorithm and theoretical results are provided in **Supplementary Methods: SSANOVA-SVD**.

#### Enhanced interpretation via decompositions

Because each process is defined directly in relation to the original covariates, its variation can be decomposed into contributions due to individual covariates and their interactions. For example, a process modeled as a function of treatment, time and their interaction can be decomposed into treatment, time and treatment-by-time contributions. The processes can also be evaluated throughout their covariate domains rather than only at values represented in the observed data. Together, these properties directly characterize reconstructed processes in relation to experimentally or observationally defined characteristics.

#### Prediction and inference without data reuse

We estimate the system-level model, comprising the processes and feature loadings, using training data (*Y*_train_*, X*_train_*, Z*_train_). The learned processes can then be evaluated at *X*_test_ to obtain predicted values of the processes *U* (*X*_test_), without using the measurements *Y*_test_. Together with the feature loadings learned from the training data, the predicted processes can be used to predict the external measurements,

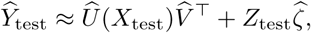

providing an out-of-sample evaluation of the reconstructed system-level model. Crucially, because *Y*_test_ was not used to reconstruct or predict the processes, these measurements remain available for conditional inference of feature participation without double dipping. This inference step can use statistical models appropriate for the measurements at hand. In the applications presented here, we use linear models. The resulting feature associations can subsequently be examined individually or through standard enrichment analyses.

Together, these capabilities enable SSANOVA-SVD to reconstruct overarching processes as interpretable smooth functions of covariates, characterize their covariate-specific behavior, predict them in external data, and enable conditional inference of feature participation using independent measurements. Thus, SSANOVA-SVD unlocks the three-step framework for system-level synthesis in practice.

### 2.2 SSANOVA-SVD and the three-step framework unlock system-level synthesis from high dimensional, simultaneously profiled snapshots

We have proposed a general three-step framework for system-level synthesis (**Fig. 1A**) and developed SSANOVA-SVD to build the system-level models required for this purpose (**Fig. 1B-C**). We therefore evaluated the proposed approach along the main capabilities required for system-level synthesis: *reconstruction* of the underlying system-level processes and feature signals, *interpretation* of those processes with respect to study covariates, *prediction* in unseen observations from covariates alone, and *statistical inference* on the features participating in the reconstructed processes.

We first assessed these capabilities in controlled simulations, where the underlying processes and participating features are known. We then selected real-world datasets spanning distinct biological scales, study designs, data types, and covariate structures, including temporal, developmental, disease-associated, and spatial variation. Additional applications evaluating the framework in longitudinal immune-response and global epidemiological settings are provided in the supplements. The applications were chosen to assess whether the framework could recover interpretable system-level structure across substantially different settings, rather than being tailored to a particular assay, biological system, or type of covariate.

The system-level modeling component of the three-step framework (via SSANOVA-SVD) leverages a natural factorization of the data (Section **2.1**). In this representation, each measured feature is represented as a superposition of overarching processes that, when evaluated at the observed covariates, yield the corresponding component vectors. We therefore assessed the advantages of the proposed three-step framework over contemporary matrix factorization methods^21,22,30,33^. Unlike conventional matrix factorization methods, SSANOVA-SVD enables prediction of overarching processes in unseen samples from covariates alone and decomposition of the processes into smooth, potentially nonlinear covariate-specific contributions. Within the proposed three-step framework, these predictions further enable conditional inference of feature participation without data reuse. Consequently, prediction and inference metrics, as well as certain assessments of interpretability, are available only for the proposed framework. Results for PCA^21^ and NMF^22^ are presented in the main text, while comparisons with PLS^33^, sparse PCA^30^, sparse NMF, and the covariate-aware MEFISTO^34^ are provided in the supplement, yielding broadly similar conclusions.

#### 2.2.1 Validation of system-level reconstruction and inference in simulations

We evaluated the ability of SSANOVA-SVD coupled with the three-step framework to deliver reliable system-level synthesis in controlled simulations, compared with existing matrix factorization methods ( **Supplementary Methods: Simulations**, **Supplementary Note 1**). We considered a challenging high-dimensional setting in which only 2.5% of 5000 features were driven by the simulated biological processes. To avoid confounding performance comparisons with the number of components, all competing methods were evaluated at the common rank, *K*, selected by cross-validation using SSANOVA-SVD. SSANOVA-SVD more accurately prioritized participating features than competing matrix factorization methods and produced low-rank approximations with reconstruction accuracy comparable to the best-performing matrix factorization method (**Supplementary Note 1: Figs. 1A, 1C, 2A, 2C**). The three-step framework for synthesis additionally unlocked capabilities unavailable to matrix factorization under general conditions, by reliably identifying participating features at the desired false discovery rate with increasing power as sample sizes increased (**Supplementary Note 1: Figs. 1B, 2B**). Finally, under dense sampling of this sparse-signal regime (**Supplementary Note 1: Fig. 3**), the smooth processes learned by SSANOVA-SVD explained more than 99.9% of the variation in the simulated processes (*R*^2^ ≥ 0.999 for each process, **Supplementary Note 1: Figs. 4, 5A–D**) and predicted feature signals in held-out samples with very high accuracy (*R*^2^ = 0.994, **Supplementary Note 1: Fig. 5E**), while their highly interpretable decomposition into covariate-specific contributions and interactions revealed the underlying structure of the simulated system-level processes (**Supplementary Note 1: Figs. 4, 6–9**).

#### 2.2.2 Interpretable reconstruction, prediction and inference of circadian dynamics

To test whether SSANOVA-SVD could recover smooth temporal structure without prespecifying periodicity, we analyzed a circadian transcriptomic study^47^ that profiled gene expression every 2 hours for 3 days in mice synchronized to light–dark cycles. We modeled the system-level processes as smooth functions of time while accounting for tissue variation (**Methods**, **Fig. 2, Supplementary Note 2**). Without imposing periodicity, SSANOVA-SVD reconstructed two smooth processes oscillating with 24-hour periods and distinct phases (**Fig. 2A**). In contrast, the dominant components generated by matrix factorization methods were substantially less interpretable (**Fig. 2E–F**; **Supplementary Note 2: Figs. 17–53**). Although some later components showed oscillatory structure, they exhibited substantially greater variability within and between tissues (**Supplementary Note 2: Figs. 17, 22**).

**Figure 2:**
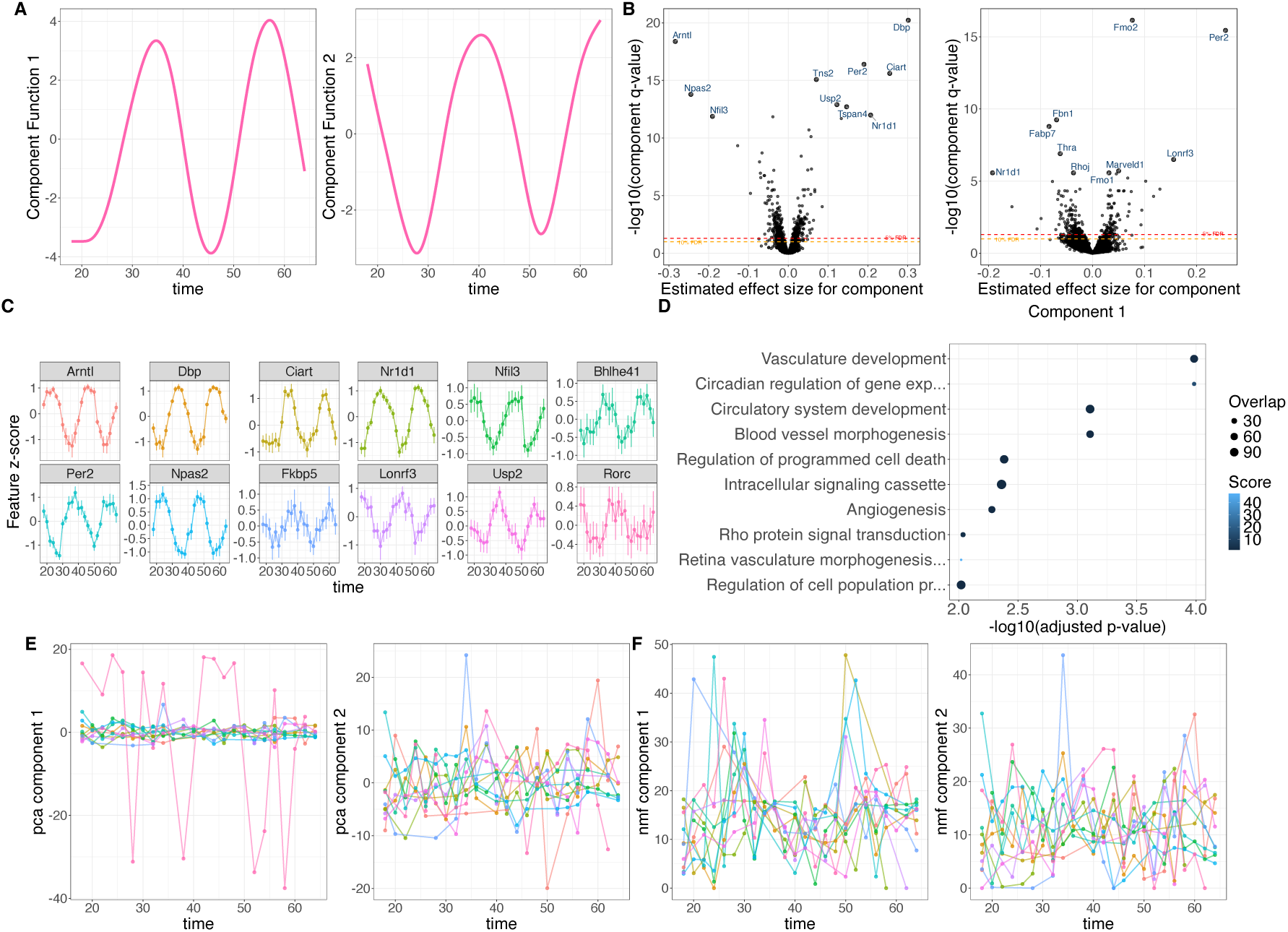
System-level synthesis of circadian rhythms. **A**. Reconstructed processes for the circadian transcriptomic data by SSANOVA-SVD. **B**. Conditional inference of feature loadings for the processes from unseen test data. Volcano plots show estimated component-specific feature effect sizes and corresponding q-values; dashed lines denote 5% and 10% false discovery rate (FDR) thresholds. Top 10 genes are labeled. **C**. Standardized expression profiles of associated genes *Arntl* and *Per2* across time. Points and error bars summarize expression at each sampled time point across tissues. **D**. Gene Ontology Biological Process enrichment among features associated with the components; point size indicates gene-set overlap and color indicates enrichment score. **E**. The first two components obtained using principal component analysis (PCA). **F**. The first two components obtained using non-negative matrix factorization (NMF).

The three-step framework additionally enabled statistical inference on feature participation and prediction in held-out mice without data reuse. Using measurements from the held-out mice, conditional inference independently linked the reconstructed processes to established circadian molecular circuitry and physiological programs. Significantly associated genes included the core regulators *Arntl*, *Per2*, *Npas2*, *Nr1d1*, *Dbp*, *Nfil3*, and *Bhlhe41* ^48,49^ (**Fig. 2B–C**; **Supplementary Note 2: Figs. 11–14**), and were enriched for established circadian, vascular^50^ and extracellular-matrix programs^51,52^ (**Fig. 2D**; **Supplementary Note 2: Fig. 14**).

Importantly, these results largely recapitulate well-established circadian biology. Their value here is not the novelty of the individual associations, but that the framework recovered them directly from the high-dimensional measurements and study covariates without requiring prior specification of the underlying circadian circuitry or programs. In a less well-characterized system, the same analysis would provide an immediate set of data-driven hypotheses about the overarching processes and the features participating in them.

The trained system-level model also predicted expression of the same 5000 genes in held-out mice from their sampling times, explaining 95.4% of the variation in their gene expression measurements.

Thus, SSANOVA-SVD reconstructed an interpretable covariate-parameterized model of circadian system-level variation, while the three-step framework independently recovered known molecular circuitry and physiological programs and accurately predicted gene expression in unseen mice, illustrating how the same analysis could rapidly generate hypotheses in less well-characterized systems.

#### 2.2.3 Persistent and time-dependent processes distinguish Immune Checkpoint Blockade responders

To evaluate the framework in a multifactorial setting involving both group and time-dependent responses, we analyzed a bilateral mouse model study^53^ that profiled 144 AB1 mesothelioma and Renca renal cell carcinomas before and at three time points following immune checkpoint blockade (ICB), with mice classified as responders or nonresponders according to their subsequent treatment response.

We modeled the system-level processes as smooth functions of time, responder status, and their interaction (**Methods**, **Fig. 3, Supplementary Note 3**). SSANOVA-SVD revealed two distinct processes underlying ICB response: a persistent expression difference between responders and nonresponders, and a differential temporal process in which the two groups progressively diverged (**Fig. 3A–B**). In contrast, dominant components generated by matrix factorization methods were substantially less interpretable and were not clearly associated with the experimental design (**Fig. 3E–F**; **Supplementary Note 3: Figs. 61–97**). The three-step framework enabled conditional inference of feature participation without data reuse. Using measurements from held-out mice, this step linked the persistent response-associated process to established immune programs, including lymphocyte and T-cell activation and differentiation, adaptive immune responses, immune-effector processes, and type-II interferon production (**Fig. 3B–D**; **Supplementary Note 3: Figs. 55–58**). Associated genes included *Cd3d*, *Cd4*, *Cd28* ^54^, *Cd2*, *Itk* ^55^, and *Slamf1* (**Fig. 3C**). The second process was associated with genes exhibiting pronounced differential temporal trajectories, particularly in AB1 tumors, including *Podxl2*, *Mdk* ^56^, *Clec11a*^57^, *Mamdc2*, and *Unc5c*. Although these genes did not show significant pathway enrichment, several have established roles in extracellular signaling, tumor–microenvironment interactions, or tissue remodeling, while associations with *Apol9b*^58^, *Apol6*, and *Tgtp1* were consistent with interferon-associated host-response biology.

**Figure 3:**
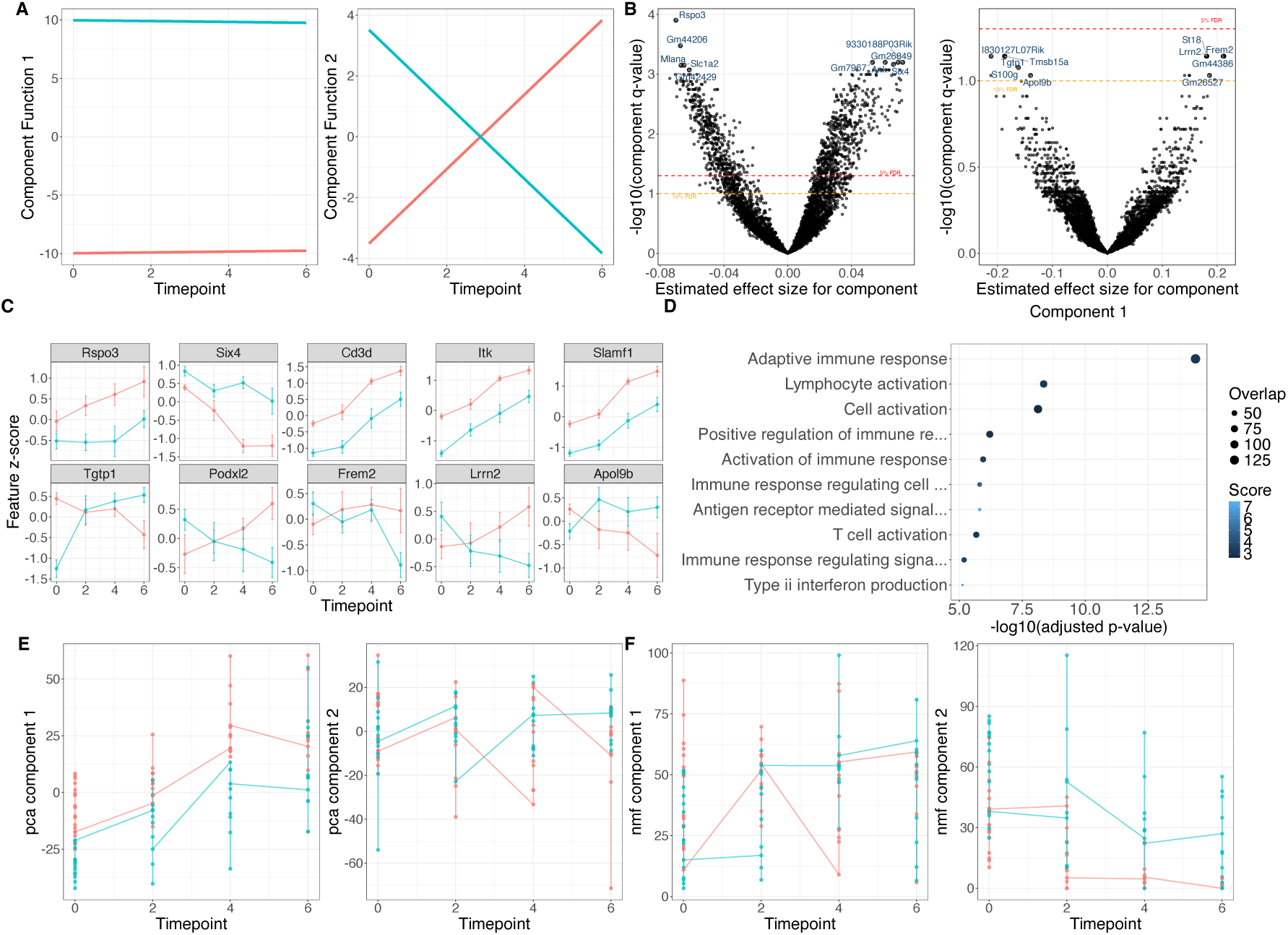
System-level synthesis of immune checkpoint blockade in bilateral mouse models. **A**. Reconstructed processes for the bilateral mouse model data by SSANOVA-SVD. **B**. Conditional inference of feature loadings for the processes from unseen test data. Volcano plots show estimated component-specific feature effect sizes and corresponding q-values; dashed lines denote 5% and 10% false discovery rate (FDR) thresholds. Top 10 genes are labeled. **C**. Standardized expression profiles of associated genes across time for component 1 (top row), and component 2 (bottom row). Points and error bars summarize expression at each sampled time point across tissues. **D**. Gene Ontology Biological Process enrichment among features associated with component 1; point size indicates gene-set overlap and color indicates enrichment score. **E**. The first two components obtained using principal component analysis (PCA). **F**. The first two components obtained using non-negative matrix factorization (NMF).

Limited temporal sampling in this experiment may favor relatively simple reconstructed processes, leaving finer temporal variation between sampled time points unresolved. Nevertheless, associations with molecular features in held-out mice provided independent support for the dominant processes captured by the model.

The original study^53^ resolved temporal gene programs and identified transient IFN*β* signaling as a determinant of ICB response. SSANOVA-SVD provided a succinct, complementary system-level model that isolated persistent immune differences from differential temporal responses in responders and nonresponders, while the three-step framework independently identified the molecular features participating in these processes.

#### 2.2.4 Compact synthesis of a malarial parasite’s life cycle from a multifactorial single-cell atlas

To assess whether SSANOVA-SVD could organize a complex developmental system with many discrete biological states, we analyzed the Malaria Cell Atlas^15^, which profiled single *Plasmodium berghei* parasites throughout development in mammalian hosts and mosquito vectors.

We modeled the system-level processes as discrete functions of life-cycle stage, with smoothness encoded through regularization of stage effects (**Methods**, **Fig. 4, Supplementary Note 4**). SSANOVA-SVD revealed 11 overarching processes that organized the parasite life cycle into broad biological programs while resolving finer developmental transitions (**Fig. 4A**; **Supplementary Note 4: Figs. 98–103**). These included processes associated with gametocytes (C2), male gametogenesis (C4), schizont-associated merozoite maturation (C6), ookinete development and invasion (C3, C7), oocyst development (C1, C9), and sporozoite and liver-stage biology (C5, C11). Distinct processes further separated general gametocyte biology from male gametogenesis, mature ookinete invasion from the subsequent ookinete-to-oocyst transition, trophozoites from exoerythrocytic forms, and salivary-gland from post-inoculation sporozoites.

**Figure 4:**
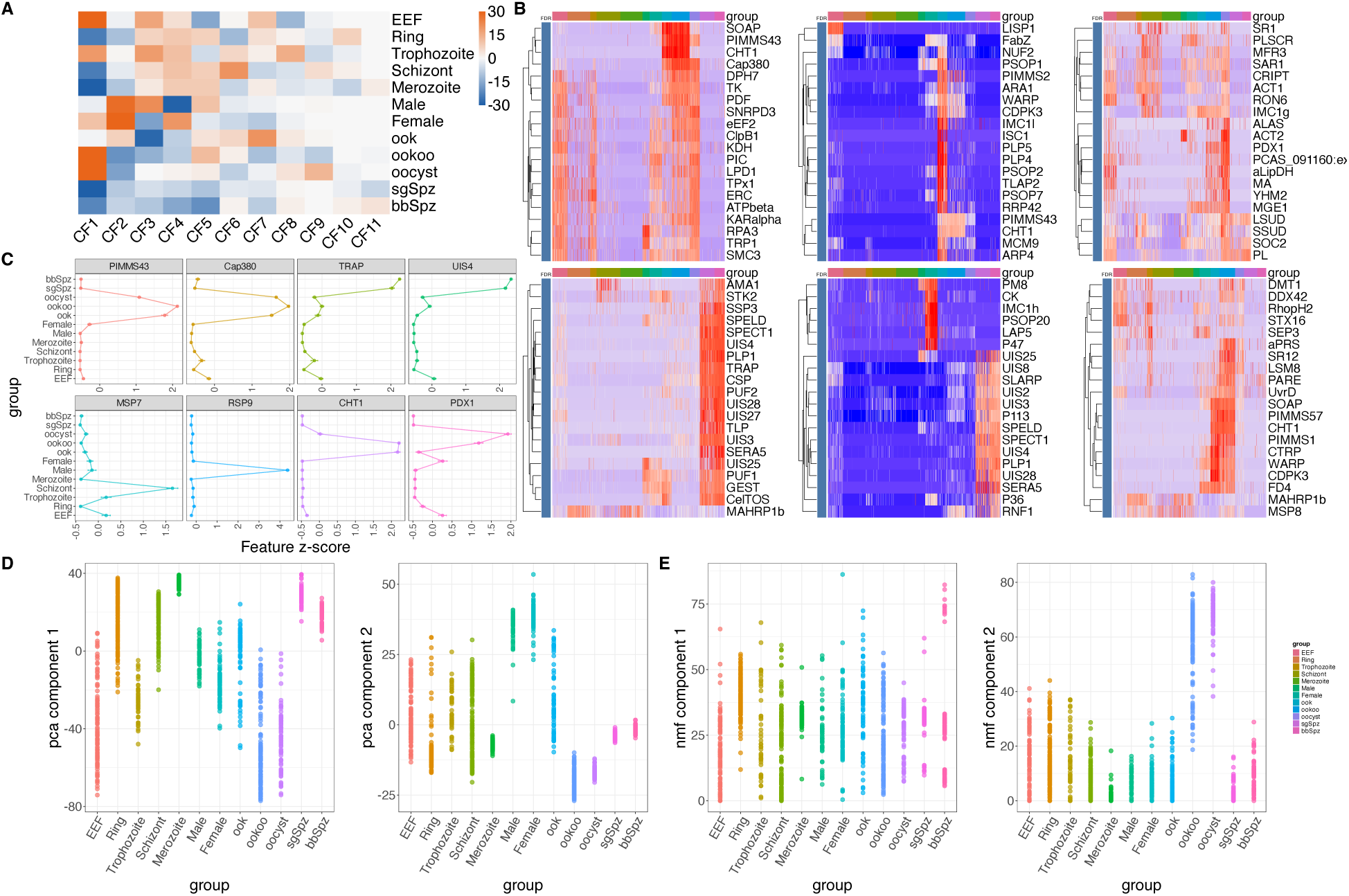
System-level synthesis of malarial parasite’s life cycle. **A**. Reconstructed processes for the Malaria data by SSANOVA-SVD. Values outside limits are clipped for better visualization. **B**. Conditional inference in unseen test data. Heatmaps of z-scores of top 20 positively (top row) and negatively (bottom row) associated genes for components 1(left), 7 (center) and 9 (right), excluding PBANKA genes without known gene symbols. Blue annotation indicates the gene’s *FDR* ≤ .1 for the component. **C**. Standardized expression profiles of representative associations, also marking life stages. Points and error bars summarize expression at each sampled stage. **D**. The first two components obtained using principal component analysis (PCA). **E**. The first two components obtained using non-negative matrix factorization (NMF).

In contrast, although matrix factorization methods also recovered life-cycle-associated variation, their components exhibited substantially greater within-stage heterogeneity and combined multiple biological contrasts, making the dominant components less readily interpretable (**Fig. 4D–E**; **Supplementary Note 4: Figs. 134–176**).

The three-step framework unlocked conditional inference of feature participation using measurements from held-out cells. Despite incomplete functional annotation of the *P. berghei* genome, independently associated genes closely reflected the reconstructed processes (**Fig. 4B–C**; **Supplementary Note 4: Figs. 104–122**). The gametocyte process (C2) was associated with *Mdv1* and *Nek3*, whereas male gametogenesis (C4) was associated with *Rsp9* ^59^, *P230* ^60^, and *Hap2* ^61^. The schizont-associated process (C6) identified *Msp*, *Sera*, *RhopH*, and *RON* -family proteins involved in merozoite maturation and erythrocyte invasion^62^, while an ookinete-specific process (C7) identified *Cht1*, *Warp*^63^, *Soap*, and *Cdpk3* and was enriched for host-entry functions. A distinct process marking the transition from invading ookinetes/early oocysts to established oocysts (C9) was associated with *Pimms57* ^64,65^, together with *Cht1*, *Warp*, *Soap*^66^, *Cdpk3* ^67^, and *Ctrp*. Sporozoite and liver-stage processes were similarly associated with *Csp*, *Trap*^68^, *Spect1* ^69^, *CelTOS* ^70^, *UIS* -family genes, *Slarp*, and *Lisp1/2* ^71^.

Whereas the original Malaria Cell Atlas characterized developmental trajectories and stage-associated gene expression throughout the parasite life cycle^15^, SSANOVA-SVD provided a succinct, complementary system-level model that separated broad life-cycle programs from finer developmental transitions, and the three-step framework independently identified the molecular features participating in these processes.

#### 2.2.5 System-level model of multiple myeloma progression generates an experimentally validated hypothesis

To evaluate the framework in a disease-progression setting and its potential for hypothesis generation, we analyzed purified plasma cells from 32 individuals^72^ spanning four stages of disease progression: normal bone marrow, monoclonal gammopathy of uncertain significance (MGUS), smoldering multiple myeloma (SMM), and multiple myeloma (MM). Given the limited sample size, we restricted this application to learning and interpreting the system-level model.

We modeled the system-level processes as discrete functions of disease stage, with smoothness encoded through regularization of stage effects (**Methods**, **Fig. 5, Supplementary Note 5**). SSANOVA-SVD revealed a single dominant process that closely tracked progression from healthy bone marrow through MGUS and SMM to overt multiple myeloma (**Fig. 5A**). In contrast, the leading components recovered by matrix factorization methods (**Fig. 5E-F**, **Supplementary Note 5: Figs. 181–221**) exhibited substantially greater within-stage variability, and did not provide an equally succinct representation of disease progression. Genes with strong loadings on the reconstructed process included several previously implicated in myeloma biology (**Fig. 5B-C**, **Supplementary Note 5: Figs. 178–179**). These included *ST3GAL6* ^73^, *HGF*, and *CD81* ^74^, as well as potentially actionable features; notably, *CD274* (*PD-L1*) and *KCNMA1* are targets of FDA-approved drugs. Genes with high absolute loadings were enriched for adaptive and B-cell-mediated immune responses, lymphocyte-mediated immunity, and immune-effector processes (**Supplementary Note 5: Fig. 179**). Several interferon-responsive genes, including *XAF1* and *MX1*, increased with disease progression, whereas *CD27* ^75^, whose loss has been associated with progression toward malignant plasma-cell phenotypes, decreased. Hypoxia-associated genes^76^, including *SLC2A3*, *HIF1A*, and *BNIP3*, similarly exhibited pronounced disease-associated changes.

**Figure 5:**
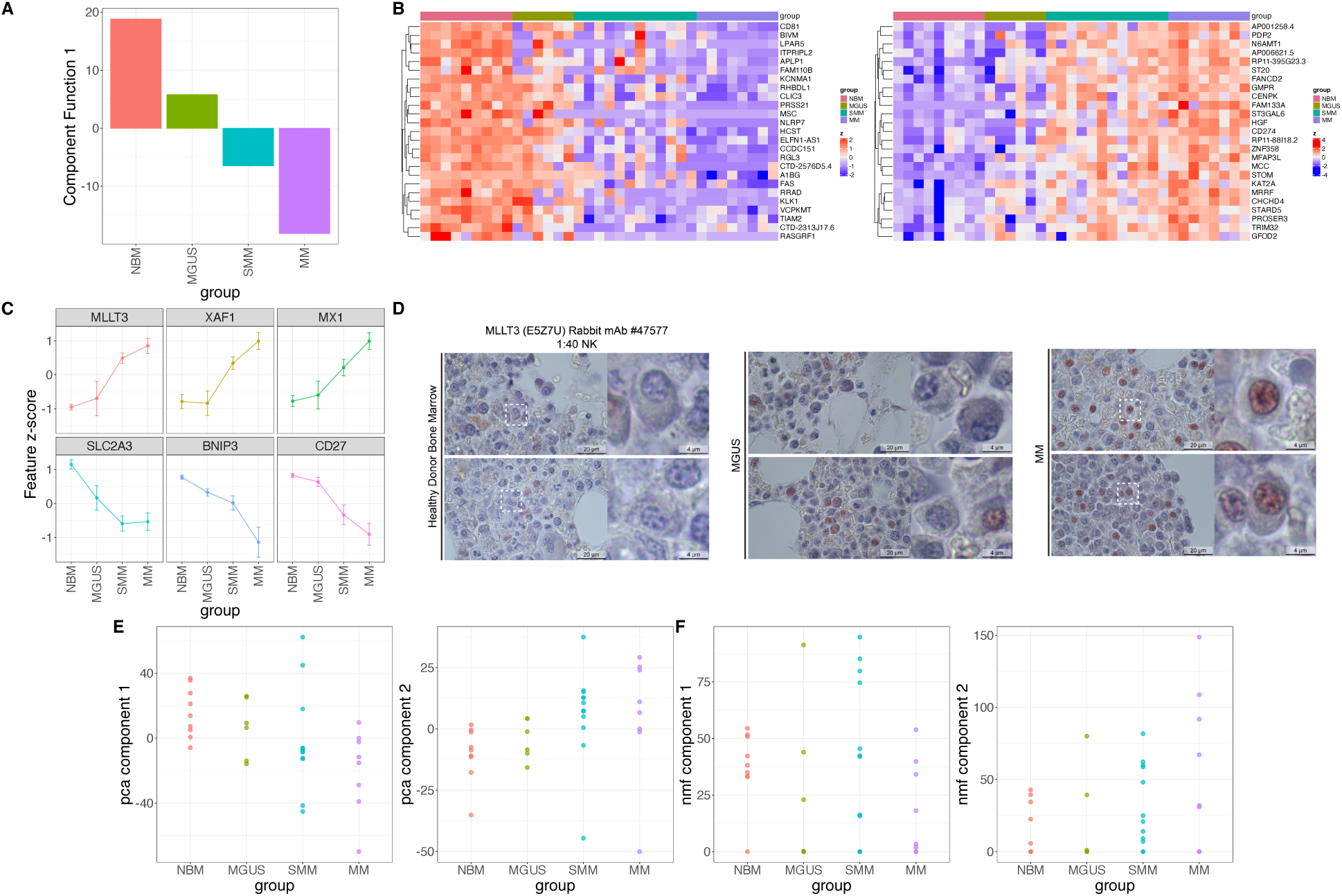
System-level synthesis of Multiple Myeloma progression. **A**. Reconstructed processes for the multiple myeloma data by SSANOVA-SVD. **B**. Heatmaps of the top 25 positively (left) and negatively (right) loading genes, excluding immunoglobulin genes. **C**. Standardized expression profiles of representative genes across stages for component 1. Points and error bars summarize expression at each sampled stage. **D**. Immunohistochemistry of MLLT3 in bone marrow samples isolated from healthy, MGUS and MM patients. **E**. The first two components obtained using principal component analysis (PCA). **F**. The first two components obtained using non-negative matrix factorization (NMF).

Among the hypotheses thus generated from the system-level model learned by SSANOVA-SVD, we examined *MLLT3* ^77,78^ because of its myeloma-specific expression pattern in the Human Protein Atlas^79^. The learned model indicated progressively increasing *MLLT3* expression across disease stages. Immuno-histochemical staining of independent bone-marrow specimens from healthy individuals, MGUS, and MM patients confirmed this increase(**Fig. 5D**), providing independent experimental support for the hypothesis generated by SSANOVA-SVD.

#### 2.2.6 Smooth spatial processes reproducibly capture complex bird assemblages at continental scale

To test the generality of the framework beyond molecular measurements in a system with nonlinear spatial structure, we analyzed the North American Breeding Bird Survey^80^, which profiles the abundances of hundreds of bird species throughout North America. These data exhibit rich spatial structure^81,82^, allowing us to assess whether SSANOVA-SVD could recover interpretable geographic processes and reproducible species assemblages.

We modeled system-level processes as smooth functions of latitude, longitude, and their interaction while accounting for year-to-year variation (**Methods**, **Fig. 6, Supplementary Note 6**). SSANOVA-SVD revealed a hierarchy of spatial processes, ranging from broad east–west and north–south contrasts to localized and increasingly complex spatial patterns (**Fig. 6**, **Supplementary Note 6: Figs. 222–229**). As expected from the strong spatial structure in these data, conventional matrix factorization recovered broadly similar but noisier versions of these spatial components (**Supplementary Note 6: Figs. 295–328**).

**Figure 6:**
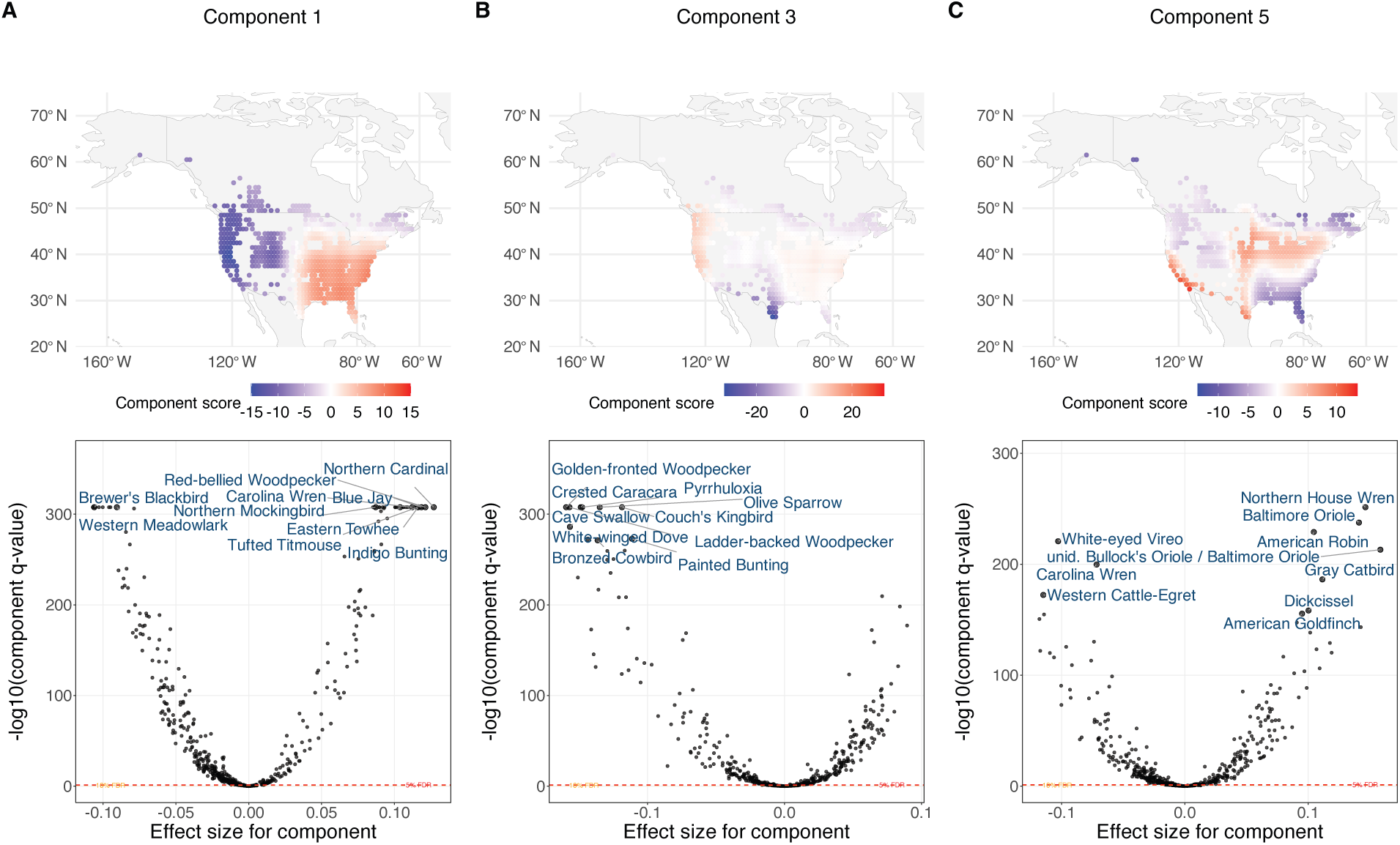
Smooth spatial processes capture bird assemblages in the North American Breeding Bird Survey reproducibly. **A**. Spatial distribution of process 1 (top), capturing the dominant continental-scale contrast in bird-community composition, and conditional inference for individual species associated with the component (bottom). **B**. Spatial distribution of process 3 (top), identifying a localized South Texas–Mexico border assemblage, and corresponding conditional species inference (bottom). **C**. Spatial distribution of process 5 (top), illustrating a more spatially complex pattern of bird-community variation, and corresponding conditional species inference (bottom). In the spatial maps, processes are evaluated at observed BBS locations, with blue and red indicating negative and positive component scores, respectively. Dashed lines in the conditional-inference plots indicate the 5% and 10% false discovery rate (FDR) thresholds.

Crucially, the three-step framework established that the learned smooth spatial processes were reproducibly associated with coherent bird assemblages (**Fig. 6)**, even for continental-scale processes linking geographically distinct regions (**Fig. 6C**, **Supplementary Note 6: Figs. 230–264**). For example, the fifth process jointly captured distinct northeastern, southeastern, Californian, and southwestern assemblages. Process values were first predicted from geographic coordinates in held-out data, without using the corresponding bird measurements, which were subsequently used for conditional inference of species participation. Simpler spatial processes showed similarly reproducible associations. Indeed, the dominant east–west process was associated with characteristic eastern species, including Northern Cardinal, Carolina Wren, and Red-bellied Woodpecker, and western species including Western Tanager, Western Wood-Pewee, and Spotted Towhee (**Fig. 6A**). A localized South Texas process was independently associated with Golden-fronted Woodpecker, Crested Caracara, Olive Sparrow, Pyrrhuloxia, and Couch’s Kingbird (**Fig. 6B**). More broadly, species associated with the learned processes were enriched for ecological attributes including habitat, migration, diet, nesting and foraging strategies (**Supplementary Note 6: Figs. 253–264**).

Thus, beyond recovering established spatial organization, SSANOVA-SVD synthesized complex spatial variation into a handful of smooth geographic processes whose associations with coherent bird assemblages were independently reproduced through the three-step framework.

#### 2.2.7 Additional applications

Additional applications of the framework to global epidemiological data and longitudinal immune-response studies are presented in **Supplementary Notes 7-9**. The resulting synthesis recovered previously established seasonal and climate-associated structure in global dengue incidence (**Supplementary Note 7**), highlighted a previously identified experimental irregularity underlying dominant variation in a longitudinal immune-response study (**Supplementary Note 8**), and resolved compartmental and temporal processes during the early T-cell response (**Supplementary Note 9**). Thus, unlike conventional matrix factorization, the framework directly linked dominant processes to study covariates, making them highly interpretable, and, where sample size permitted, identified participating features through conditional inference.

## 3 Discussion

Advances in experimental and observational technologies have transformed the study of complex systems by enabling the simultaneous profiling of thousands of features, accompanied by a rich set of temporal, spatial, and other contextual covariates. For several decades, analysis of high-dimensional biological data has largely followed two complementary analytical traditions^83^, with different primary objectives: modeling individual features as functions of biological variables (e.g., feature-wise regressions^84^), and extracting joint structure from the measurements through clustering and latent representations^85^. While these approaches have advanced our understanding of complex systems, they do not directly address the challenge of synthesizing multivariate observations into coherent, interpretable models of system behavior. Such system-level models unite overarching processes with the features participating in them, providing a strong foundation for mechanistic understanding, prediction, and hypothesis generation^4,44,86^. Thus, achieving system-level synthesis from simultaneous profiled data remains an important and unresolved statistical challenge.

Classical art offers a highly instructive analogy. For centuries, painters have first established the overall composition and blocked in the major forms and relationships before progressively resolving finer details. Beginning instead with fine details—a jewel in a portrait or a stalk of grass in a landscape—provides little assurance that they will ultimately assemble into a coherent representation of the whole. Over three decades of omics, much of feature-wise analysis has proceeded in the latter direction: individual features are characterized first and subsequently organized, for example through gene-set or pathway enrichment, in an effort to obtain a broader biological picture. We believe the arts offer a useful principle for understanding complex systems, wherein a cohesive picture is more readily constructed by first establishing the broad strokes and then resolving finer details within the context of the whole.

Inspired by this classical strategy, we propose a new analytical paradigm for high dimensional data that explicitly targets system-level synthesis – in which interpretable system-level models are built to capture overarching processes of a simultaneously profiled system and features participating in them are reliably identified. To this end, we establish a three-step framework: (1) using training data, learn system-level models where overarching processes are expressed as interpretable functions of covariates of study units, (2) predict these processes in external data from their covariates alone, and (3) use the unused measurements from these data for conditional inference of feature participation in the processes. We developed SSANOVA-SVD to serve step 1 of the framework. It learns system-level processes as smooth flexible functions of nominal, ordinal, and continuous covariates and their interactions. Their dependence on covariates enables the reconstructed processes to be interpreted, decomposed into covariate-specific contributions, and transferred to external data, naturally separating process reconstruction from subsequent feature-level inference. Thus, beyond its role in enabling system-level synthesis, SSANOVA-SVD extends existing matrix factorization methodology with an interpretable, covariate-parameterized decomposition that supports prediction and downstream inference under general study designs. In diverse real-world systems, the proposed framework coupled with SSANOVA-SVD recovered interpretable overarching processes, enabled prediction in unseen samples, and detected feature associations that recapitulated known biology and generated new hypotheses.

Systems biology has a rich tradition of constructing system-level models^1,44^ (e.g., metabolic models^87^, dynamical models of regulatory and signaling networks^45,88,89^, logical network models^90^, and agent-based or multiscale models^91,92^) that often encode biological knowledge about interactions, constraints and governing mechanisms to characterize the behavior of complex systems. We complement this ***knowledge-driven*** route to system-level synthesis with a ***data-driven*** approach that takes advantage of high-dimensional simultaneous measurements^93,94^ and their covariates, to learn interpretable system-level models. Rather than requiring the underlying biological interactions or governing mechanisms to be specified in advance, our proposal reconstructs overarching processes manifested in the observed system. By revealing underappreciated structure in these processes and identifying features associated with them, data-driven system-level models can generate hypotheses about biological features and their interactions for subsequent experimental and modeling investigation. The resulting biological knowledge can, in turn, inform the construction and refinement of mechanistic system-level models, connecting data-driven discovery with a holistic mechanistic understanding of a target system.

Unlike existing data analysis approaches including feature-wise enrichment, matrix factorization, and supervised multivariate methods, the proposed approach learns system-level models in which overarching processes are represented directly as flexible functions of the original study covariates while capturing dominant joint variation in the measurements. This representation retains experimental or observational covariates as the coordinates of interpretation, enables the learned processes to be predicted in external data from covariates alone, and allows unused measurements to support conditional inference of feature participation without data reuse. Post-selection inference methods^35–37^ can address data reuse following data-driven discovery^95^, but do not themselves provide interpretable process reconstruction or out-of-sample prediction. When external data are unavailable, SSANOVA-SVD can instead be combined with data thinning or fission to separate process reconstruction from subsequent inference within the same dataset.

The applications considered here illustrate the breadth of system behavior that can be represented within the proposed framework. Without prespecifying their form, SSANOVA-SVD reconstructed processes spanning temporal, developmental, disease-associated, and spatial structure. Where sample sizes permitted inference, features associated with these processes recovered established biological programs and generated new hypotheses. Importantly, the framework is not restricted to omics measurements, as demonstrated by its application to continental-scale bird abundance data, where high-dimensional features are jointly observed alongside informative covariates.

This generality also means that there is no single visualization that is optimal for every reconstructed system-level model. Because the processes are represented directly over their covariate domains, their most informative summaries depend on the structure of those covariates and on the scientific question. Temporal processes may be most naturally displayed as trajectories, developmental or ordinal processes as stage-specific profiles, spatial processes as maps, and feature participation through volcano plots, heatmaps, enrichment summaries, or other domain-appropriate displays. Thus, SSANOVA-SVD provides a common statistical representation while allowing visualization and interpretation to remain adapted to the structure of the system under study.

The identification and subsequent experimental validation of MLLT3’s association with multiple myeloma progression illustrate the utility of building interpretable system-level models for generating hypothesis and driving experimental discovery. MLLT3 is a chromatin-associated transcriptional regulator involved in maintaining hematopoietic stem-cell identity and self-renewal^78^. While its functional significance in multiple myeloma remains unclear, our findings motivate the hypothesis that MLLT3 participates in transcriptional programs associated with plasma-cell malignancy progression, warranting further functional investigation.

The following limitations warrant consideration. First, the covariate-dependent formulation that provides interpretability also defines the scope of the processes that can be reconstructed. System-level variation unrelated to, or inadequately represented by, the specified covariates may remain unexplained, while unmeasured nuisance variation may complicate interpretation. As in conventional regression analysis, examination of residual structure^41^ can therefore reveal unexplained variation and motivate refinement of the model. Second, the reconstructed processes and inferred feature participation only represent statistical dependencies and cannot, without additional experimental evidence, be interpreted as establishing causal or mechanistic relationships. Third, inference of feature participation requires sufficient observations to separate process estimation from inference with adequate statistical power; in the smaller applications considered here, we therefore restricted analysis to reconstruction and interpretation of the overarching processes. When independent data are not feasible, post-selection inference approaches such as data thinning or fission may provide alternative strategies, although partitioning information within a small dataset may lead to poorly selected models and reduce statistical power. Finally, the optimization problem is non-convex and the alternating estimation procedure can converge to local optima, as is common in matrix factorization problems.

Several methodological extensions would further broaden the scope of the proposed framework. These include integrative analyses involving multiple data modalities, extensions to additional observation models, and formal uncertainty quantification for the estimated overarching processes. Several of these extensions are already available in the accompanying software package and will be described in forthcoming publications. Although simulations under densely sampled regimes demonstrate accurate recovery of the underlying overarching processes by SSANOVA-SVD, formal guarantees for process recovery have yet to be established. Finally, our current estimation algorithm constructs the factorization sequentially rather than jointly optimizing all components. Characterizing the relationship between the sequential estimator and solutions to the joint optimization problem remains an interesting theoretical direction.

Taken together, these results support system-level synthesis as a general objective for the analysis of high-dimensional data and establish a three-step framework for connecting interpretable system-level modeling to out-of-sample prediction and statistical inference on feature participation using independent measurements. SSANOVA-SVD provides a flexible statistical learning approach for reconstructing overarching processes as smooth functions of nominal, ordinal, and continuous covariates and their interactions, enabling their interpretation, decomposition, and prediction in external data. In diverse biological and ecological systems, the framework recovered established system-level phenomena while revealing previously unrecognized structure and generating new hypotheses. More broadly, analytical frameworks explicitly designed for system-level synthesis provide a route for transforming increasingly high-dimensional multivariate measurements into interpretable models that connect system-level behavior with the individual features that participate in it.

## 4 Methods

In this section, we outline the three-step framework, describe SSANOVA-SVD and its utility in carrying out step 1 of the framework, and the analysis of real world datasets.

### 4.1 Three-step framework for system-level synthesis

The proposed framework for system-level synthesis operates in three steps (**Fig. 1**). First, using training data, system-level models are learned, in which the overarching processes are represented interpretably, parameterized by sample covariates and their interactions. Second, the learned component functions are evaluated at the covariates of external samples to reconstruct overarching processes in the external samples, without using their measurements. Finally, the unused measurements in the external samples are used for conditional inference of feature participation in the reconstructed processes. Thus, parameterization with covariates enables both out-of-sample prediction and subsequent feature-level inference without double dipping.

We developed SSANOVA-SVD to serve the first step of the three-step framework, which in turn unlocks steps 2 and 3. Step 3 can use any inference methodology appropriate for the data. In this paper, we used standard linear models to implement step 3. The individual steps are described below.

### 4.2 Step 1: SSANOVA-SVD

Using training data, we learn the system-level processes as flexible, interpretable functions of sample covariates and their interactions.

Briefly, a given *n* samples x *p* features data matrix *Y* is decomposed with a factor model, *Y_n×p_* = *Ũ V ^′^*+*Zζ* +*Error*. Here *Ũ_n×K_* = (*ũ*_1_*, … , ũ_K_*) is the components matrix obtained by evaluating the respective component functions *U* = {*u*_1_(·), · · · *, u_K_*(·)} at sample covariates (*x*_1_*, … , x_n_*). *V_p×K_* = (*v*_1_ *… v_K_*) is the feature loadings matrix, *Z_n×m_* is a nuisance design matrix (e.g., batch effects) and *ζ_m×p_* is the corresponding coefficients matrix. The set of component functions *U*, and the feature loadings matrix *V*, are obtained by solving a penalized optimization problem that trades off reduction in the reconstruction error of *Y* by the factor model, for smoothness in the reconstructed functions. The component functions are estimated from a smoothing spline ANOVA function space H defined over the sample covariates *X*^46^ (**Supplementary Methods: SSANOVA-SVD**). The optimization problem is:

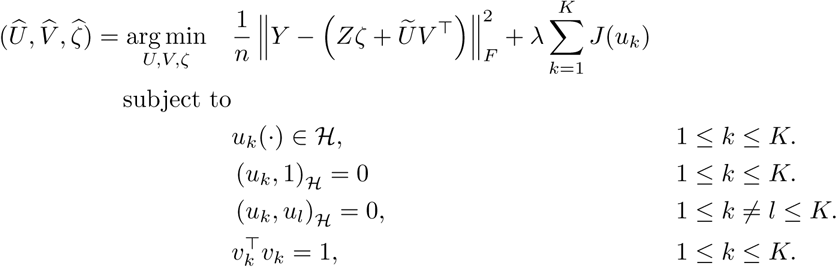

(·, ·)*_H_* is an inner product for the space of functions H, and *J* (·) is a roughness penalty that enforces smoothness in the *u_k_*(·). The second constraint removes the constant component from *u_k_*(·), allowing feature-specific baselines to be identified via *Z*. The third enforces orthogonality among the component functions. The final constraint enables the identification of the scale of *u_k_*(·).

Numerical estimation is performed using alternating minimization as follows. To compute the rank-*K* factorization, we sequentially solve constrained rank-one optimization problems, enforcing orthogonality against previously estimated component functions at each step (Section 4.2.1). A representer theorem reduces each rank-one optimization to a finite-dimensional problem. We sketch the estimation algorithm below. Complete derivations are presented in **Supplementary Methods: SSANOVA-SVD**.

#### 4.2.1 Sequential estimation algorithm for the SSANOVA-SVD problem

To fit the component functions *U*, we consider an efficient sequential estimation algorithm, similar to fast low rank matrix factorization algorithms, in which we estimate *u_k_* given the previously estimated functions *û_t_*(·), feature loadings *û_t_*, *t < k*. Under this estimation strategy, we have the rank-1 SSANOVA-SVD problem that fits one component function *u*(·) at a time via:

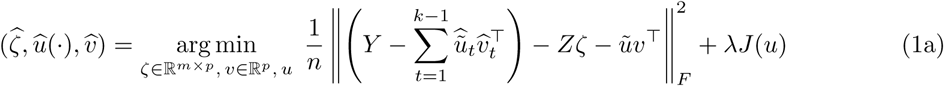

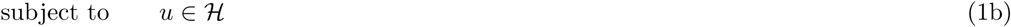

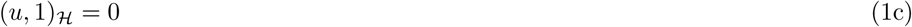

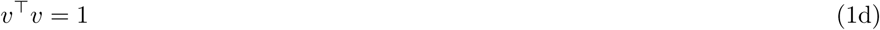

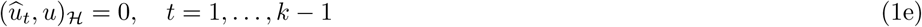

The problem is non-convex. An alternating minimization algorithm is used for estimation:

1. Fit *ζ*, given *û*(·), *v̂*. The problem reduces to a standard least squares problem. Solution is obtained as 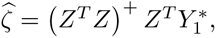 where 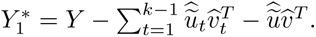
2. Fit *u*, given 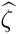 and 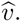 For a suitable choice of *J* (*u*), the problem reduces to fitting a smoothing spline ANOVA model^46^ for the working responses 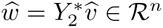 under additional constraints, where 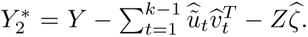 In general, optimizing over the function space is an infinite dimensional problem. Theorem 2 in **Supplementary Methods: SSANOVA-SVD**, proves that the solution to this problem is obtained in finite dimensions as : 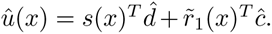 Here, *s*(*x*) contains evaluations of simple, interpretable basis functions at *x* (e.g., constant and linear functions), while *r̃*_1_(*x*) represents higher-order variation through kernel basis functions. The vectors *d* and *c* are the corresponding coefficients. The penalty *J* (*u*) controls the roughness of *u* by penalizing its component in the space represented by *r̃*(·). Similar results also hold for the specific case of estimating the first component function, *u*_1_(·). Complete details are presented in **Supplementary Methods: SSANOVA-SVD**.
3. Fit *v*, given 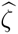 and *û*(·). The solution is obtained as 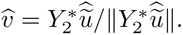

**Initialization, and termination** 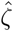 are initialized via least squares as 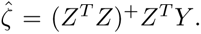 *U* and *V* are initialized via a fast low rank singular value decomposition^96^ of *Y* . Optimization terminates when either the change in the objective or the parameters are below convergence tolerances, or when the maximum number of coordinate-wise iterations is reached.

#### 4.2.2 Cross-validation

The set of hyperparameters for SSANOVA-SVD include the number of component functions *K*, the smoothing penalty *λ*, and an additional set of scaling parameters that parameterize *r̃*_1_(·) and the inner product *J* (*u*). The ability to predict system-level processes in unseen samples, as explained in the next subsection, enables these hyperparameters to be selected by cross-validation based on out-of-sample predictive performance. For a given number of folds, and number of components *K*, SSANOVA-SVD is trained on one fold to build the system-level model {*U*^^^*, V*^^^ } where 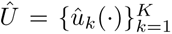 and *V*^^^ = [*v̂*_1_*, … , v̂_K_*], along with nuisance coefficients *ζ̂*. For a test fold with covariates *X_test_*and nuisance design *Z_test_*, the system-level model can be used to predict the test data matrix from the covariates alone as 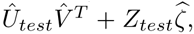 where *U*^^^*test* is the predicted components matrix for the test fold. Cross-validation numerically optimizes over the mean squared error calculated between the observed and the predicted matrix. Complete details are presented in Section 5 of **Supplementary Methods: SSANOVA-SVD**.

### 4.3 Step 2: Prediction of overarching processes

Application of SSANOVA-SVD to training data yields a system-level model {*U*^^^*, V*^^^ }, where 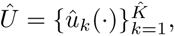 and *V*^^^ = [*v̂*_1_*, … , v̂_K_*]. For an external observation with covariates *x_i_*, its corresponding components matrix can be predicted as [*û*_1_(*x_i_*)*, … , û_k_*(*x_i_*)], with *u_k_*(*x_i_*) = *s*(*x_i_*)*^T^ d*^^^*k* + *r̃*_1_(*x_i_*)*^T^ ĉ_k_*. Evaluating the component functions in this way for all external observations at their respective covariates, and stacking them, we obtain *U_test_*, the components matrix for the external/test data. Thus, only the covariates, and not the actual measurements from external data, is used in reconstructing the components *U_test_*.

### 4.4 Step 3: Conditional inference of feature participation

For both simulated and real world datasets, where sample sizes did not preclude inference, we used 70% of the observations, *Y_train_*, to train a system-level model via SSANOVA-SVD. The remaining 30%, *Y_test_*, was subjected to steps 2 and 3 of the framework for inference of feature participation.

The conditional inference step of the framework aims to infer feature participation in the overarching processes built in step 1. This step can in principle use any inference methodology appropriate for the data. For the datasets in this study, we used limma^17^, to infer feature participations based on the linear model of the data assumed during training, on the adjusted observations *Y_test_* − *Z_test_ζ*^^^ = *U_test_β* + *Error*. If the measured confounder design, *Z_test_* is distinct to training, one may use *Y_test_* = *U_test_β* + *Z_test_ζ* + *Error* for inference of feature participation *β*.

Together, Sections 4.2-4.4 complete the description and implementation of the 3-step framework and SSANOVA-SVD.

### 4.5 Software and code availability

An R package svdx implements SSANOVA-SVD, and the three-step framework. It is available under: https://github.com/mskb01/svdx. The function svdx() implements SSANOVA-SVD. It takes as input training measurements, its metadata, an R formula for relevant design formula, and an R formula for nuisance variation zformula. Once trained, the processes and loadings are available for prediction on external data using the predict() interface, which takes the covariates of the external data as input. The predicted components can then be used for conditional inference of feature participation - e.g., as a design matrix for feature-wise regressions in linear models^17^. Together, this chain implements the proposed three-step framework, with SSANOVA-SVD serving the modeling step.

Code for the analysis presented in the paper is available under: https://github.com/mskb01/svdxPaperAnalysisCode, which makes use of https://github.com/mskb01/svdxBenchmark.

### 4.6 Simulations

We evaluated SSANOVA-SVD in high-dimensional simulations with known system-level processes and feature participation. We simulated 5,000 features over continuous and discrete covariates, with four biological processes and nuisance batch variation. Of the measured features, 125 (2.5%) participated in the biological processes, with the remaining features serving as biological nulls. We varied sample size, confounding between biological group and batch, and overlap between biological and batch-associated features. Performance was evaluated by reconstruction of the simulated biological signal, prioritization of participating features from the estimated loadings, recovery and prediction of the simulated processes, and conditional inference of feature participation in held-out data. Inference was evaluated using power, type-I error, false discovery proportion and discrimination of participating from null features. Complete simulation specifications and evaluation metrics are provided in **Supplementary Methods: Simulations**.

### 4.7 Analysis of real-world datasets and data preparation

In this section, we describe the real-world datasets analyzed in this study, including their sources, preprocessing, and preparation for statistical analysis. For all datasets, 5000 highly variable features were retained for analysis, and their expression values were standardized. For both simulated and real-world datasets, where sample sizes permitted inference, we used 70% of the observations to train a system-level model using SSANOVA-SVD. The remaining 30% was reserved for process prediction and conditional inference of feature participation, corresponding to steps 2 and 3 of the framework.

#### 4.7.1 Circadian Gene Expression

The circadian gene-expression data were obtained from Zhang et al.^47^. Gene expression was profiled in multiple tissues every 2 hours over three days in synchronized mice.

Samples were randomly divided, without stratification, into training (70%) and test (30%) sets. Sampling time was used as the relevant biological covariate of interest, and tissue type was used as an adjustment covariate. Accordingly, SSANOVA-SVD (svdx) was fitted with xformula = ∼time and zformula = ∼tissue_type. Conditional inference of feature participation using the held-out test data was performed using limma^17^. Over-representation analysis was performed using Fisher’s exact test with Gene Ontology Biological Process (BP) gene sets from the Mouse Molecular Signatures Database (MSigDB) M5 collection^97^. P-values were adjusted for multiple testing using the Benjamini–Hochberg procedure^98^, with statistical significance defined at a false discovery rate of 10%.

#### 4.7.2 Bilateral Mouse Models

Bulk RNA sequencing data for the bilateral tumor model of Zemek et al.^53^ for both AB1 mesothelioma and Renca renal cell carcinoma were obtained from Gene Expression Omnibus accession GSE153941. RNA-sequencing counts were normalized for library size and log-transformed prior to analysis.

Samples were randomly divided into training (70%) and test (30%) sets. Sampling time and response group were used as the biological covariates of interest, with tumor type included as an adjustment covariate. Accordingly, SSANOVA-SVD (svdx) was fitted with xformula = ∼Group + Group:Timepoint and zformula = ∼TumorType. Conditional inference of feature participation from the held-out test data was performed using limma^17^. Over-representation analysis was performed using Fisher’s exact test with Gene Ontology Biological Process (BP) gene sets from the Mouse Molecular Signatures Database (MSigDB) M5 collection^97^.P-values were adjusted for multiple testing using the Benjamini-Hoschberg procedure^98^.

#### 4.7.3 Malaria Life-Cycle: Single-parasite RNA sequencing data

Single-cell RNA-sequencing data for *Plasmodium berghei* were obtained from Howick et al.^15^ through the Malaria Cell Atlas. Cells were classified into 12 life-cycle stages: EEF, ring, trophozoite, schizont, merozoite, male gametocyte, female gametocyte, ookinete, ookoo, oocyst, salivary-gland sporozoite, and blood-bolus sporozoite. Gene annotations were obtained from PlasmoDB; where a gene symbol was unavailable, the corresponding PlasmoDB feature identifier was retained as the feature label. RNA sequencing counts were normalized for library size and log-transformed prior to analysis.

Cells were divided into training (70%) and test (30%) sets, with sampling performed separately within each life-cycle stage. Life-cycle stage was used as the biological covariate of interest, with no additional adjustment covariates. Accordingly, SSANOVA-SVD (svdx) was fitted with xformula = ∼group and zformula = 1. Conditional feature-participation inference was performed on the held-out test data using limma^17^.

Gene sets for over-representation analysis were constructed from curated Gene Ontology Biological Process annotations obtained from PlasmoDB^99^. Gene Ontology terms containing fewer than five annotated genes among the analyzed features were excluded. Over-representation analysis was performed using Fisher’s exact test, with enrichment P-values adjusted for multiple testing using the Benjamini–Hochberg procedure.

#### 4.7.4 Multiple Myeloma Progression

Single-cell RNA-sequencing data from the multiple myeloma progression study of Boiarsky et al.^72^ were obtained from Gene Expression Omnibus accession GSE193531. UMI counts were aggregated within each donor to construct pseudobulk expression profiles. Genes with zero counts across all pseudobulk samples were removed. Donors were classified into normal bone marrow (NBM), monoclonal gammopathy of undetermined significance (MGUS), smoldering multiple myeloma (SMM), and multiple myeloma (MM).

Sequencing-batch information samples was obtained from the accompanying sample annotations and matched to the pseudobulk donor profiles. Pseudobulk counts were normalized for library size and log-transformed prior to analysis.

Disease stage was used as the biological covariate of interest, with sequencing batch included as an adjustment covariate. Accordingly, SSANOVA-SVD (svdx) was fitted with xformula = ∼group and zformula = ∼batch. Because of the limited number of donor-level samples, this dataset was used for process reconstruction without sample splitting or conditional inference of feature participation.

#### 4.7.5 North American Breeding Bird Survey

Bird abundance data were obtained from the 2025 release of the North American Breeding Bird Survey (BBS), maintained by the U.S. Geological Survey^80^. Data from the United States and Canada between 2010 and 2024 were retained. BBS routes were assigned to 1*^◦^* ×1*^◦^* geographic grid cells, and grid-year combinations containing fewer than three surveyed routes were excluded. Species abundance within each grid-year was calculated as the total number of individuals observed divided by the number of surveyed routes, thereby treating non-detection on surveyed routes as zero. Species detected in fewer than 1% of grid-year samples or with fewer than 10 total observations were excluded. Grid-year samples lacking breeding-season temperature or precipitation measurements were also excluded.

Mean species-abundance measurements were log-transformed prior to analysis. Latitude and longitude of the grid-cell center were used as the spatial covariates of interest, including their interaction, while year was included as a categorical adjustment covariate. Accordingly, SSANOVA-SVD (svdx) was fitted with xformula = ∼lat_center*lon_center and zformula = ∼yearFactor.

Conditional inference was performed for each reconstructed component to identify individual bird species associated with the corresponding spatial process. Statistical significance was assessed at a false discovery rate (FDR) of 10%. Species-level associations were interpreted together with the sign of the estimated association, allowing species associated with the positive and negative directions of each spatial process to be distinguished.

To characterize the ecological attributes of species associated with each reconstructed process, inference-based enrichment analyses were performed using species annotations derived from AVONET^100^ and additional nesting-trait annotations^101^. Species sets represented taxonomic and ecological characteristics, including habitat association, diet, foraging behavior, migratory behavior, and nesting characteristics. Enrichment was evaluated using Fisher’s teset among species identified by conditional inference for each component, with multiple-testing correction applied across the tested species sets using the Benjamini-Hochberg procedure.

#### 4.7.6 Dengue Incidence

Processed monthly dengue incidence data were obtained from the original study^102^. The data contained longitudinal dengue profiles for each country over the study period. The Indian Ocean Basin-Wide (IOBW) climate index was obtained from the monthly ocean-index data provided by the NCC, as in the original study. The complete historical IOBW series was linearly detrended and standardized, after which lagged three-month averages were calculated. For each month, the IOBW 1–3-month lag was defined as the mean standardized detrended IOBW value over the preceding one to three months. Climate-index preprocessing was performed using the complete historical record before restricting the data to the study period.

Calendar month and the 1–3-month lagged IOBW index were used as covariates of interest, including their interaction. The marginal effects of both covariates were modeled using cubic smooths. Accordingly, SSANOVA-SVD (svdx) was fitted with xformula = ∼month*iobw_lag_1_3 and zformula = ∼1. Cubic smooths spanning the observed range of each covariate were specified via

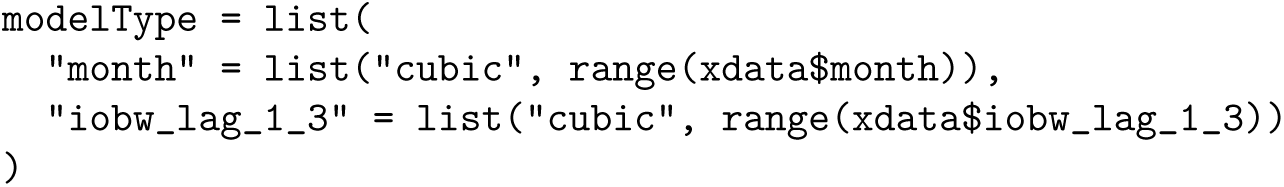

Because of the limited number of observations, this dataset was used for process reconstruction without sample splitting or conditional feature-participation inference.

#### 4.7.7 Longitudinal Anti-PD-1 Single-Cell RNA Sequencing

Single-cell RNA-sequencing data from the longitudinal anti-PD-1 experiments of Barboy et al.^103^ were obtained from Gene Expression Omnibus accession GSE249630. Raw count matrices and accompanying cell- and sample-level metadata were used to construct pseudobulk expression profiles for two longitudinal experiments. PD1 time course 2 corresponds to the longer time course, and PD1 time course 6 corresponds to the experiment focussed on the early phase following adoptive transfer. Supplementary Notes 8 and 9 present the analysis of the respective datasets.

Preprocessing followed the procedure in the original study. Cells with fewer than 450 or more than 2^1^5 total UMIs, or with a mitochondrial read fraction of at least 0.4, were excluded. For PD1 time course 6, cells were additionally required to be annotated as single cells in the accompanying metadata. Counts from retained cells were summed within biological pseudobulks, with counts from technical plates corresponding to the same biological pseudobulk combined.

For the longer PD1 time course 2, tumor-derived OT1 CD8+ T cells were analyzed and counts were aggregated by mouse. The dataset prior to the exclusion of eight mice reported in the original study was used. Treatment and sampling day, including their interaction, were modeled as the biological covariates of interest, with no additional adjustment covariates. Accordingly, SSANOVA-SVD (svdx) was fitted with xformula = ∼treatment*day and zformula = ∼1. Training and test sets were constructed at the mouse level, with one mouse from each observed treatment-by-day combination randomly assigned to the test set, thereby retaining every treatment-by-day combination in both the training and test sets.

For the shorter PD1 time course 6 that focussed on the early phase following adoptive transfer, counts were aggregated by mouse, tissue, and cell type, and OT1 CD8+ T-cell pseudobulks were retained for analysis. Previously reported exclusions of OT1 cells from six mice in the spleen and lymph node compartments were applied while retaining tumor-derived OT1 cells from these mice. Treatment, tissue, sampling day, and their interactions were modeled as the biological covariates of interest, with no additional adjustment covariates. Accordingly, SSANOVA-SVD (svdx) was fitted with xformula = ∼treatmenttissueday and zformula = ∼1. Training and test sets were again constructed at the mouse level. Approximately 30% of mice were assigned to the test set, with held-out mice distributed across treatment-by-day combinations while preserving representation of every observed combination in the training data.

Conditional inference for feature loadings in the held-out test data was performed using limma^17^. Over-representation analysis was performed using Fisher’s exact test with Gene Ontology Biological Process (BP) gene sets from the Mouse Molecular Signatures Database (MSigDB) M5 collection^97^. Enrichment P-values were adjusted for multiple testing using the Benjamini–Hochberg procedure^98^.

### 4.8 Benchmark methods

We compared the three-step framework coupled with SSANOVA-SVD with a representative set of matrix factorization methods. These include PCA^21^ (R’s implementation: stats::prcomp), NMF^22^(scikit-learn), PLS(mixOmics::pls^33^), sparse PCA^30^(mixOmics::spca), sparse NMF (scikit-learn), and the covariate-aware factorization method MEFISTO^34^(mefisto). For methods unable to model measured nuisance co-variates directly, the data was input after adjustment for the known nuisance design. To avoid confounding reconstruction with number of components, all methods were evaluated using a common rank *K*, selected by cross-validation with SSANOVA-SVD. Factorization performance was evaluated by reconstruction error for biological signal and null features, and by the ability of feature loadings to distinguish biological signal from null and batch-associated features. Method-specific preprocessing, software implementations, tuning parameters and additional benchmark details are provided in the **Supplementary Methods**.

### 4.9 MLLT3 immunohistochemistry

To assess MLLT3 expression in plasma cells, we performed immunohistochemistry on bone marrow biopsies from 10 normal bone marrow (NBM), 15 monoclonal gammopathy of undetermined significance (MGUS) and 20 multiple myeloma (MM) cases. Archival, anonymized samples were retrieved from the Department of Pathology at Brigham and Women’s Hospital (BWH) under IRB protocol 01-206.

Formalin-fixed, paraffin-embedded tissue was cut into 4-*µ*m sections. Sections were baked at 60*^◦^*C for 1 h, deparaffinized in xylene (4 changes, 3 min each), rehydrated in 100% ethanol (4 changes, 3 min each) and rinsed in running water for 5 min. Endogenous peroxidase activity was blocked with 3% hydrogen peroxide in methanol for 10 min, followed by a 5-min rinse in running water. Heat-induced antigen retrieval was performed in a pressure cooker (Biocare Medical) at 120*^◦^*C in citrate buffer (Dako Target Retrieval Solution, S1699). Slides were then cooled for 15 min and transferred to Tris-buffered saline (TBS).

Sections were incubated with an anti-MLLT3 antibody (clone E5Z7U, Cell Signaling Technology; 1:40 dilution) for 40 min. Bound antibody was detected with the Novolink Polymer Detection System (Leica Biosystems, RE7260-K), with 30-min incubations each in Post Primary and Polymer reagents. All incubations were performed at room temperature in a humidified chamber, with TBS rinses between steps. Staining was developed with 3,3*^′^*-diaminobenzidine (DAB) as the chromogen, and sections were counterstained with Mayer’s hematoxylin. IHC images were acquired using Leica DM2000 microscope (Leica Microsystems).

## Supporting information

Supplementary Note 9 : OT1 Early Phase Time Course

Supplementary Note 8 : OT1 Longer Time Course

Supplementary Note 7 : Global dengue

Supplementary Note 6 : Breeding Bird Survey

Supplementary Note 5 : Multiple Myeloma

Supplementary Note 4 : Malaria

Supplementary Note 3 : Bilateral Mouse Models

Supplementary Note 2 : Circadian Rhythms

## 5 Acknowledgements

We thank Prof. Eric V. Slud and Dr. Jared Brown for insightful discussions and valuable comments on the manuscript. This work was supported by NIH R35GM131802.

