## Supplementary Note 9 : OT1 Early Phase Time Course for "A statistical framework for system-level synthesis"

### Compartmental and temporal processes dominate early OT1 T-cell response

We analyzed the early phase of the longitudinal profiling of tumor-antigen-specific OT1 CD8 T cells following adoptive transfer into B16-OVA-bearing mice receiving anti-PD1 or control treatment[1]. OT1 cells were profiled in tumor, lymph node and spleen over the first six days following transfer.

We modeled the system-level processes as smooth functions of time, tissue and treatment (**Methods**). SSANOVA-SVD revealed three processes resolving distinct tissue- and time-dependent responses (**Supplementary Figs. 478A, 479-480**). The dominant process identified a persistent difference in expression between tumor-derived OT1 cells and those from lymph node and spleen. The second revealed a switch-like temporal transition in tumor OT1 cells while the lymphoid compartments remained comparatively stable. A third, weaker process captured a distinct temporal pattern in which lymph node and spleen OT1 cells progressively converged toward the state observed in tumor-derived cells. Unlike the longer time course analyzed in the previous section, the processes did not show a pronounced anti-PD1-associated response during this early time course. Although conventional matrix factorization also isolated the dominant anatomical separation, subsequent components were noisier and less clearly related to the experimental design (**Supplementary Figs. 478E-F, 497–535**).

Conditional inference using held-out measurements linked the dominant process to distinct T-cell states in the tumor and lymphoid compartments (**Supplementary Figs. 478B–D,481–487**). Positively associated genes, exhibiting higher expression in tumors, included those involved in T-cell activation and effector function, including *Il2ra*, *Tnfrsf9*, *Ifng*, *Gzmb*, and *Prf1*, together with metabolic and proliferative programs. Negatively associated genes, exhibiting higher expression in lymph node and spleen, included *Klf2*, *S1pr1*, *Sell*, and *Il7r*, consistent with lymphoid trafficking and a less activated T-cell state[2]. Thus, the dominant process distinguished activated, effector and proliferative tumor-associated OT1 cells from a distinct, less activated state in the lymphoid compartments.

Genes positively associated with the second process, corresponding to its early tumor-associated state, included growth, activation and metabolic regulators such as *Myc*, *Pim1*, *Ppargc1a*, *Kdm6b*, and *Cish*[3]. Genes negatively associated with the process, corresponding to its later tumor-associated state, included extensive mitotic machinery together with T-cell-state regulators such as *Tox* and *Il10ra*[4,5]. The process therefore captured a transition from an early growth, activation and metabolic program toward a later program dominated by cell-division machinery and accompanied by changes in T-cell state.

The third process had substantially weaker feature-level support, including an association with *Adora2a*[6].

Together, the dominant processes revealed that early OT1 T-cell responses were structured primarily by anatomical compartment and temporal changes in tumor-associated T-cell state.

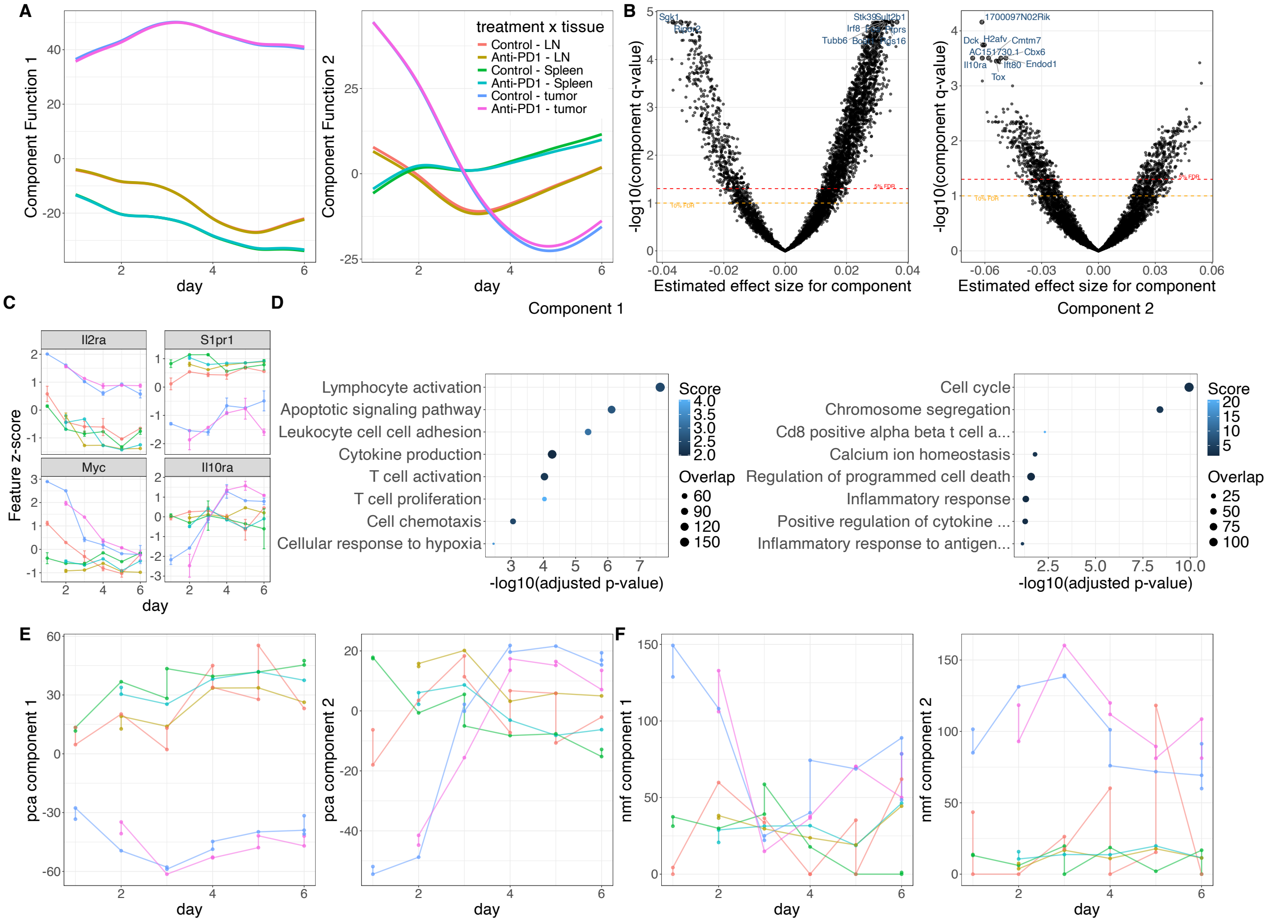


**Supplementary Figure 478: System-level synthesis of OT1 CD8 T-cells in the early phase of immune checkpoint blockade (time course 6).** A. Reconstructed component functions for time course six of the OT1 CD8 T-cells data, by SSANOVA-SVD. Anti-PD-L1 samples were not available at day 1, evaluation for visualization here is therefore an extrapolation. B. Conditional inference of feature loadings for the components from unseen test data. Volcano plots show estimated component-specific feature effect sizes and corresponding q-values; dashed lines denote 5% and 10% false discovery rate (FDR) thresholds. Top 10 genes are labeled. C. Standardized expression profiles of associated genes across time for component 1 (top row), and component 2 (bottom row). Points and error bars summarize expression at each sampled time point across tissues. D. Gene Ontology Biological Process enrichment among features associated with component 1; point size indicates gene-set overlap and color indicates enrichment score. E. The first two components obtained using principal component analysis (PCA). F. The first two components obtained using non-negative matrix factorization (NMF).

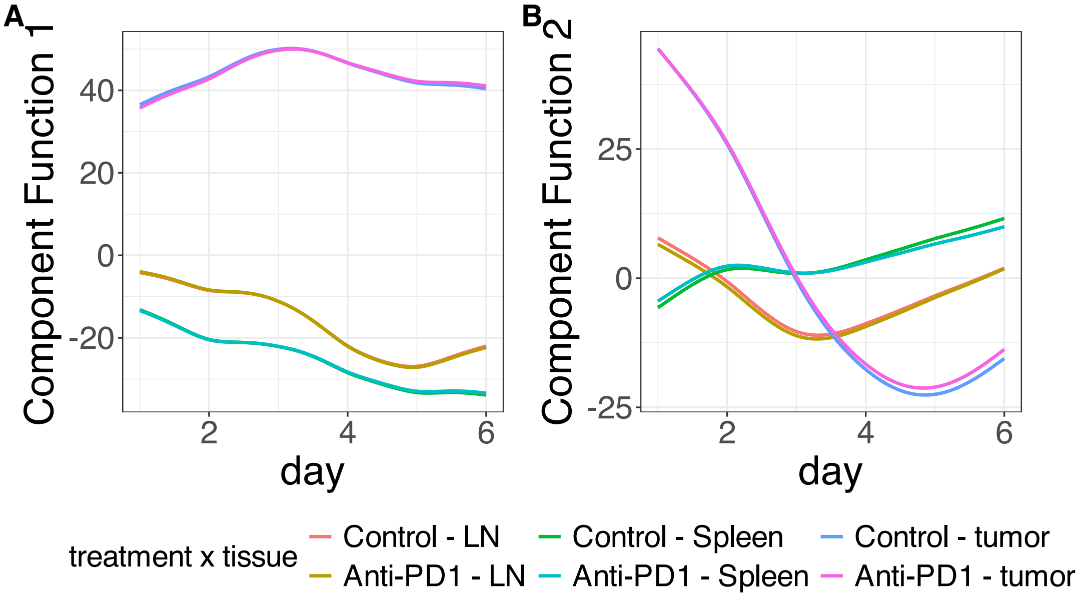


**Supplementary Figure 479.** SSANOVA-SVD reconstructed component functions.(A) Component 1. (B) Component 2.

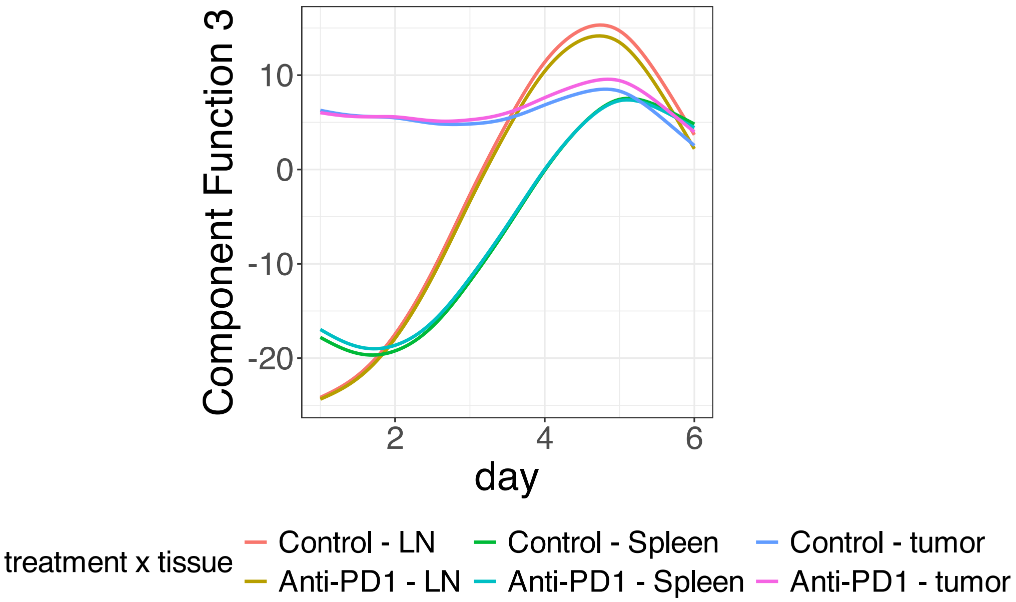


**Supplementary Figure 480.** SSANOVA-SVD reconstructed component functions.Component 3.

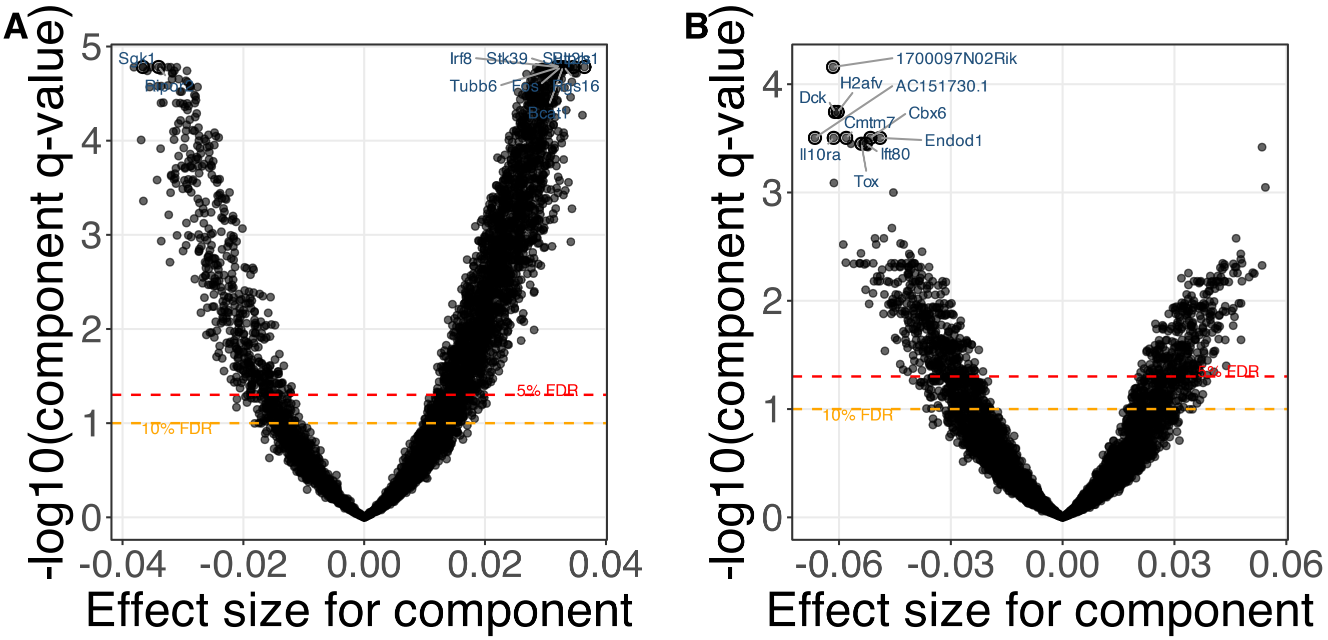


**Supplementary Figure 481.** SSANOVA-SVD based conditional inference for feature participation. (A) Component 1. (B) Component 2.

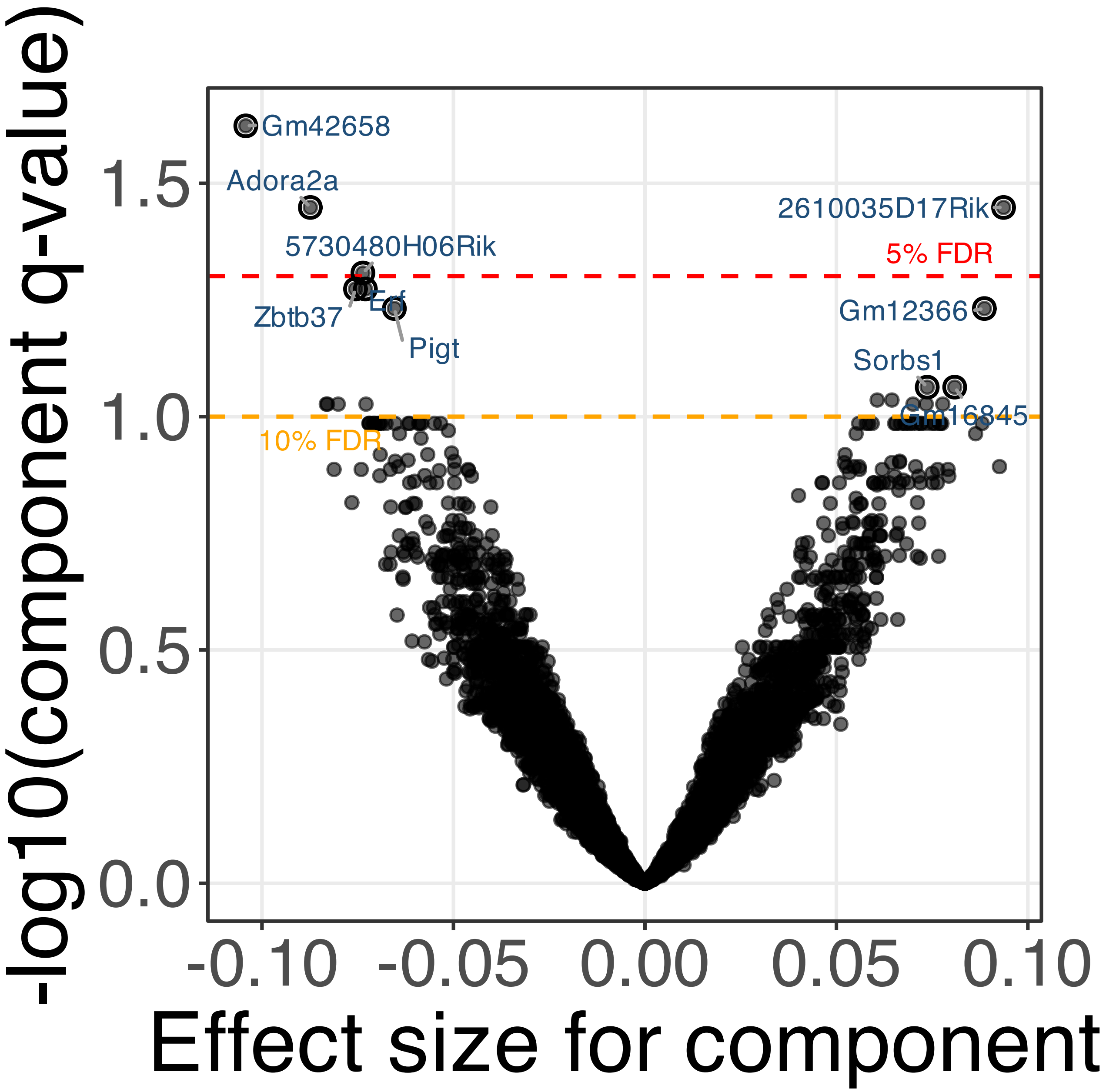


**Supplementary Figure 482.** SSANOVA-SVD based conditional inference for feature participation. Component 3.

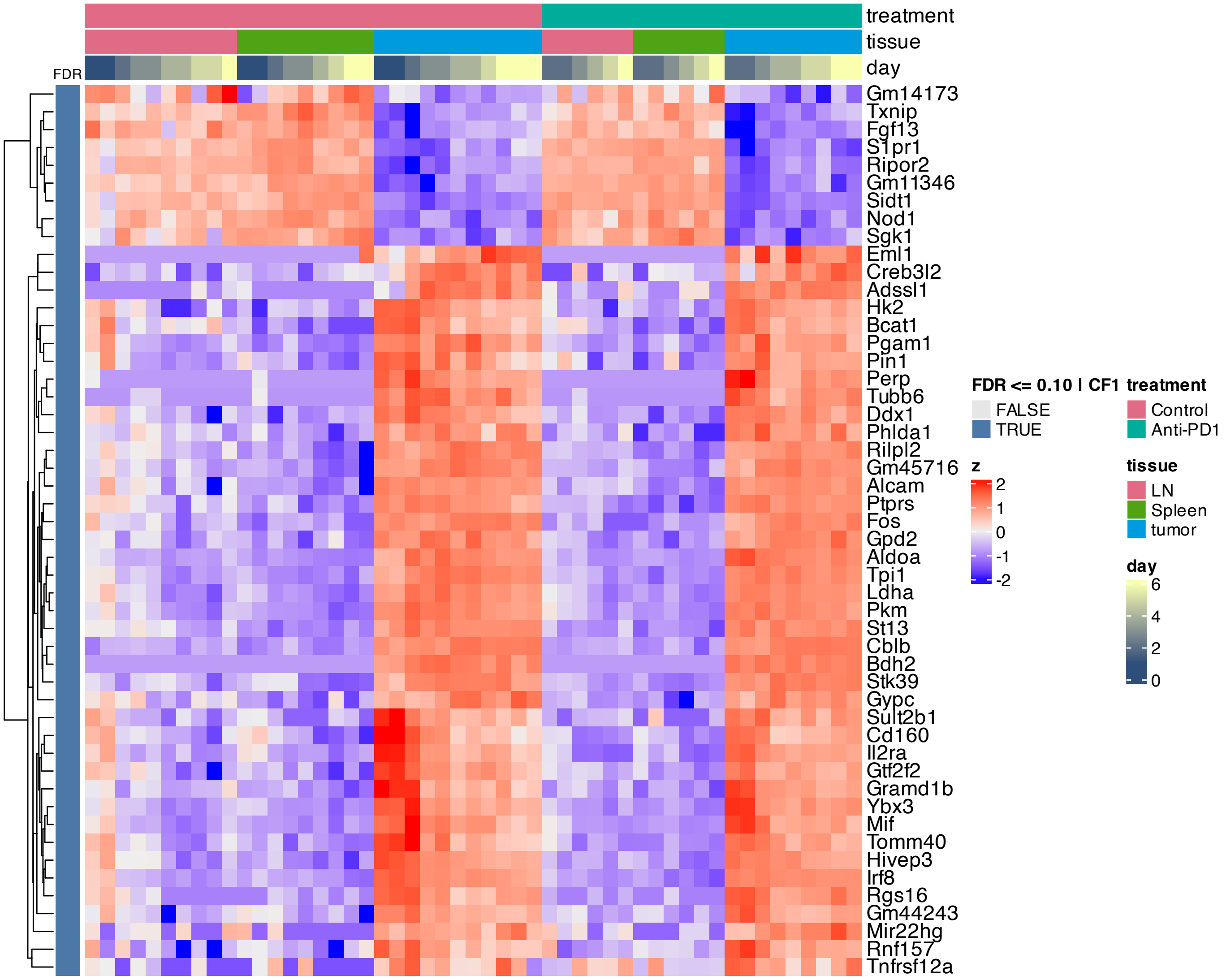


**Supplementary Figure 483.** Heatmap of features inferred to be associated with component 1 by SSANOVA-SVD.

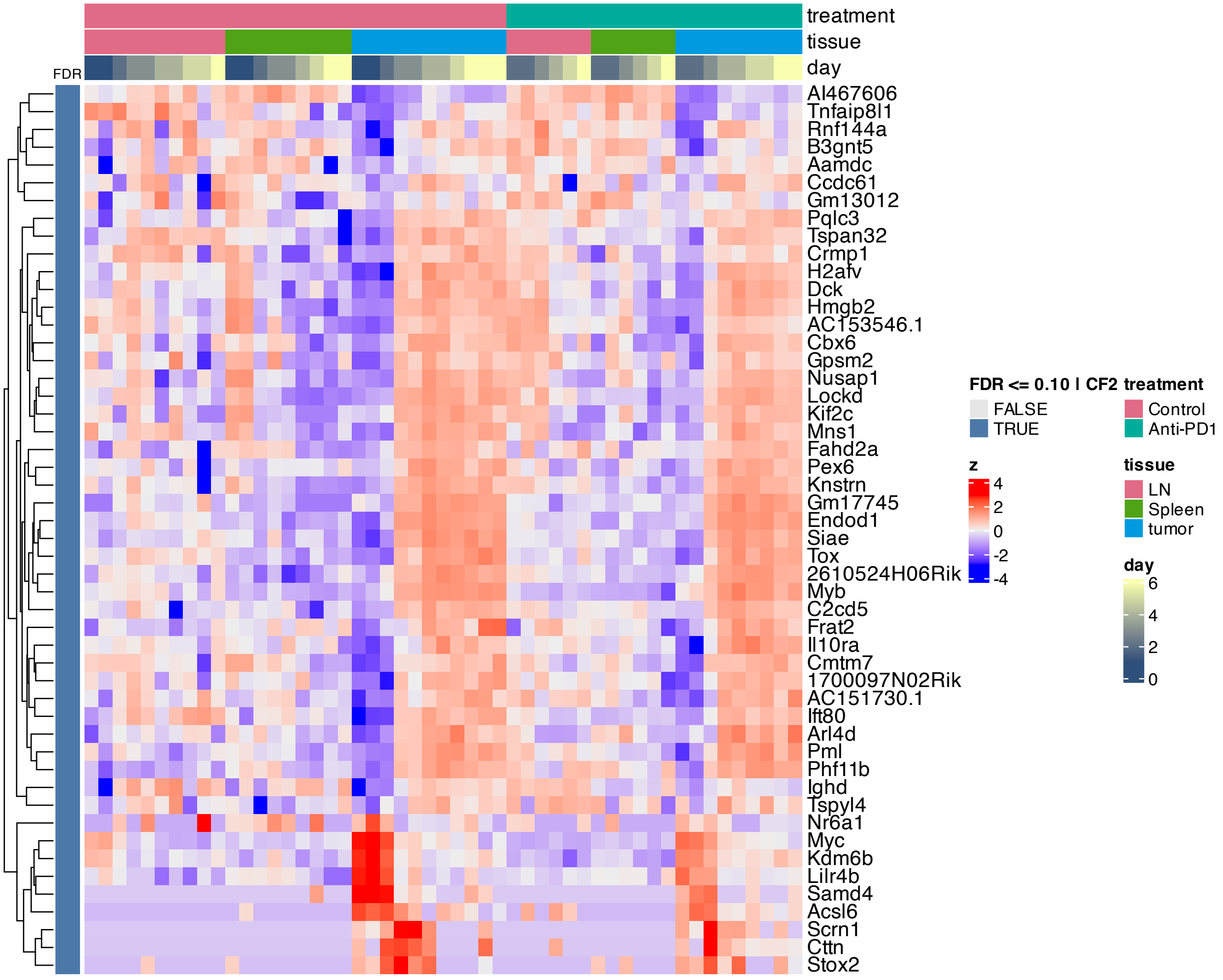


**Supplementary Figure 484.** Heatmap of features inferred to be associated with component 2 by SSANOVA-SVD.

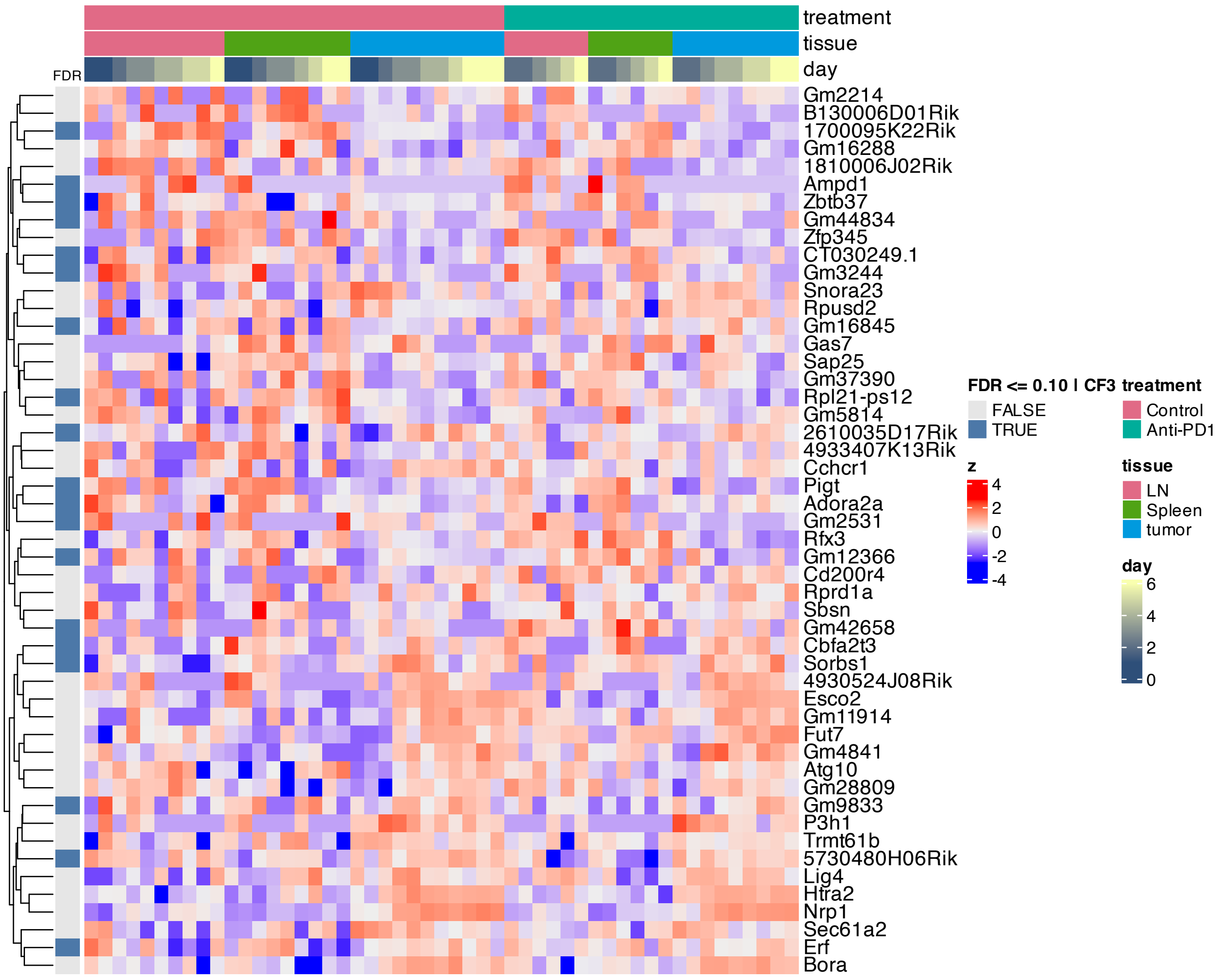


**Supplementary Figure 485.** Heatmap of features inferred to be associated with component 3 by SSANOVA-SVD.

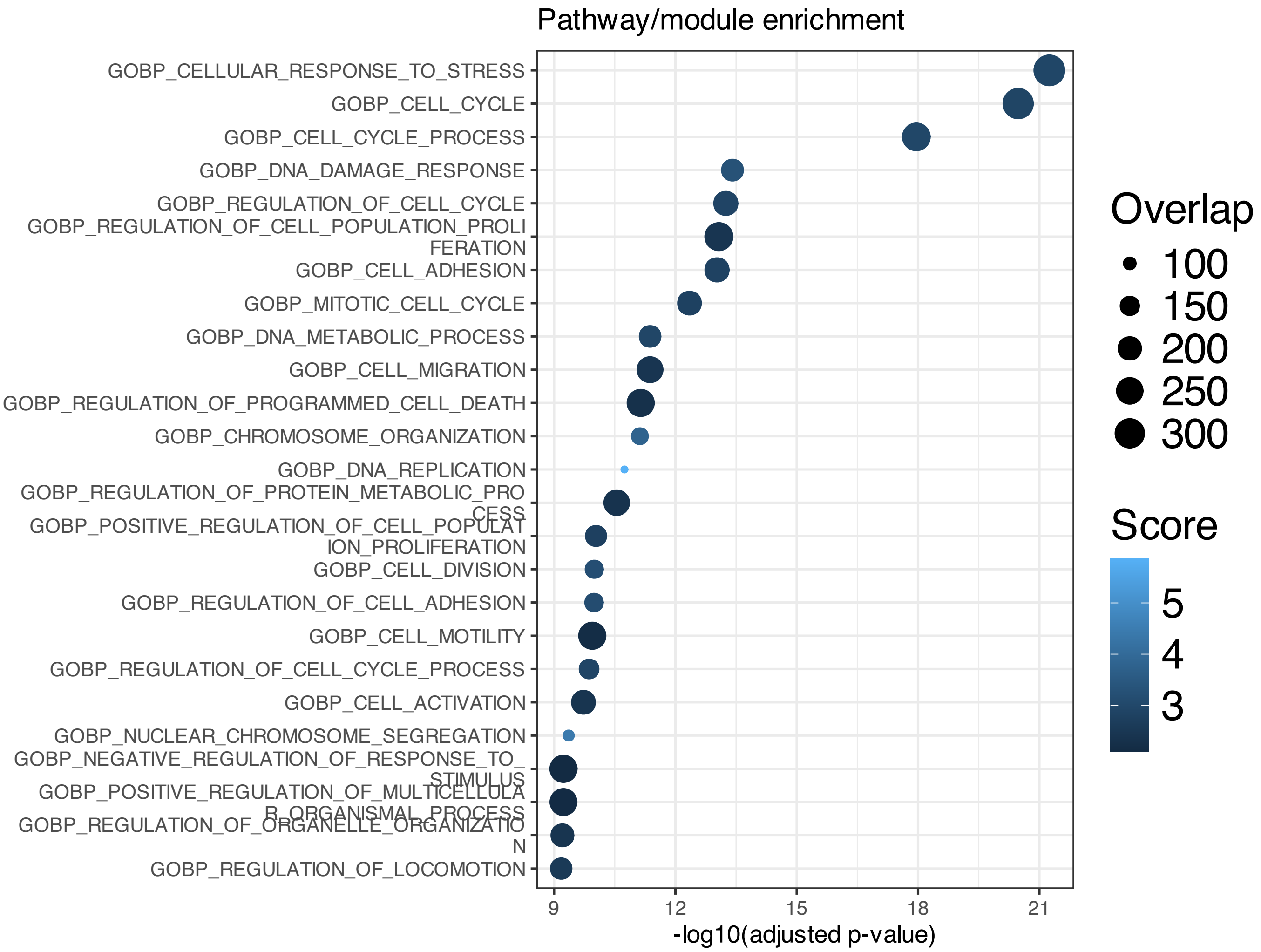


**Supplementary Figure 486.** Enrichment among features inferred to be associated with SSANOVA-SVD component 1; only terms with adjusted P $\leq 0.1$ are shown.

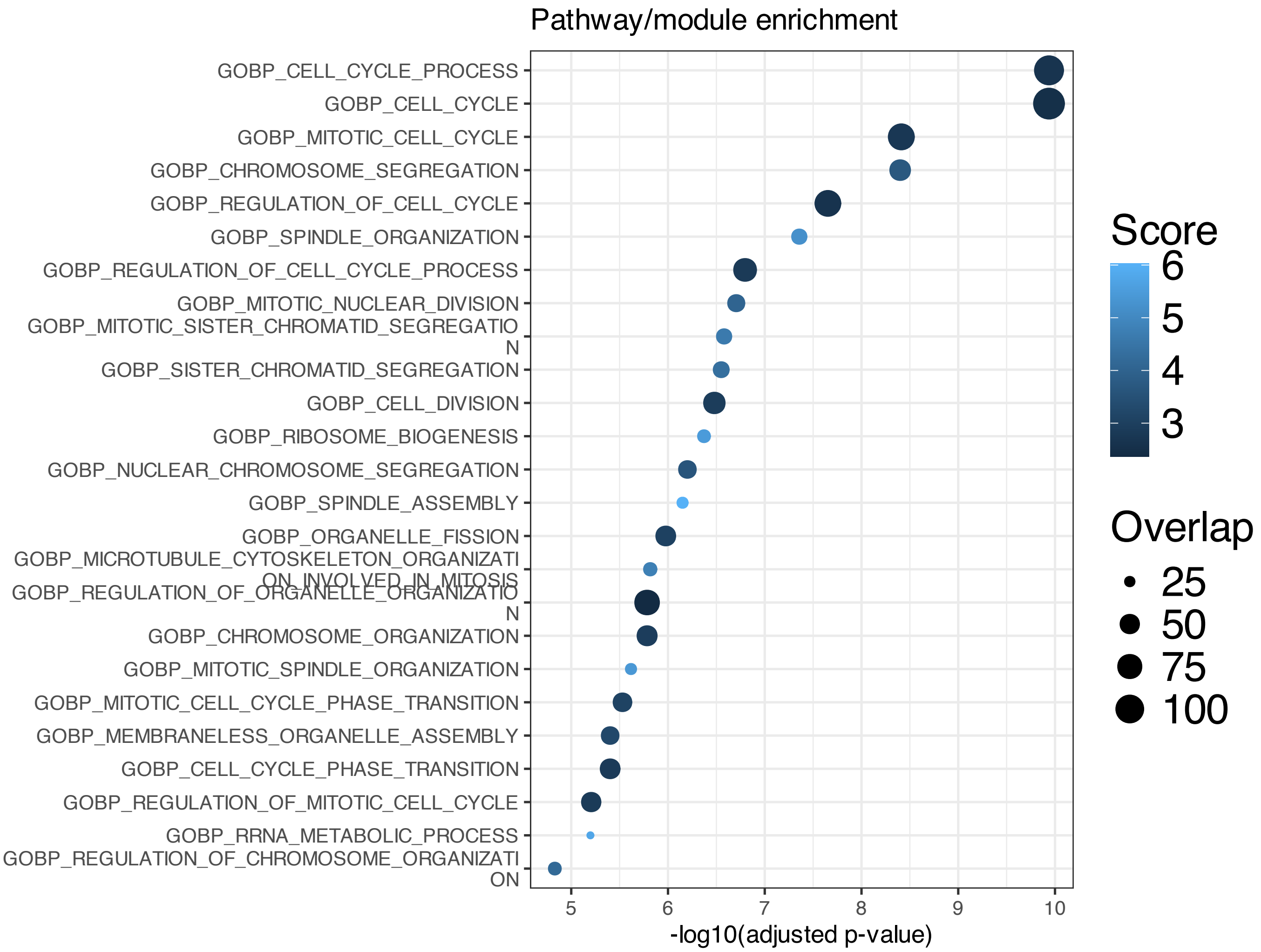


**Supplementary Figure 487.** Enrichment among features inferred to be associated with SSANOVA-SVD component 2; only terms with adjusted P $\leq 0.1$ are shown.

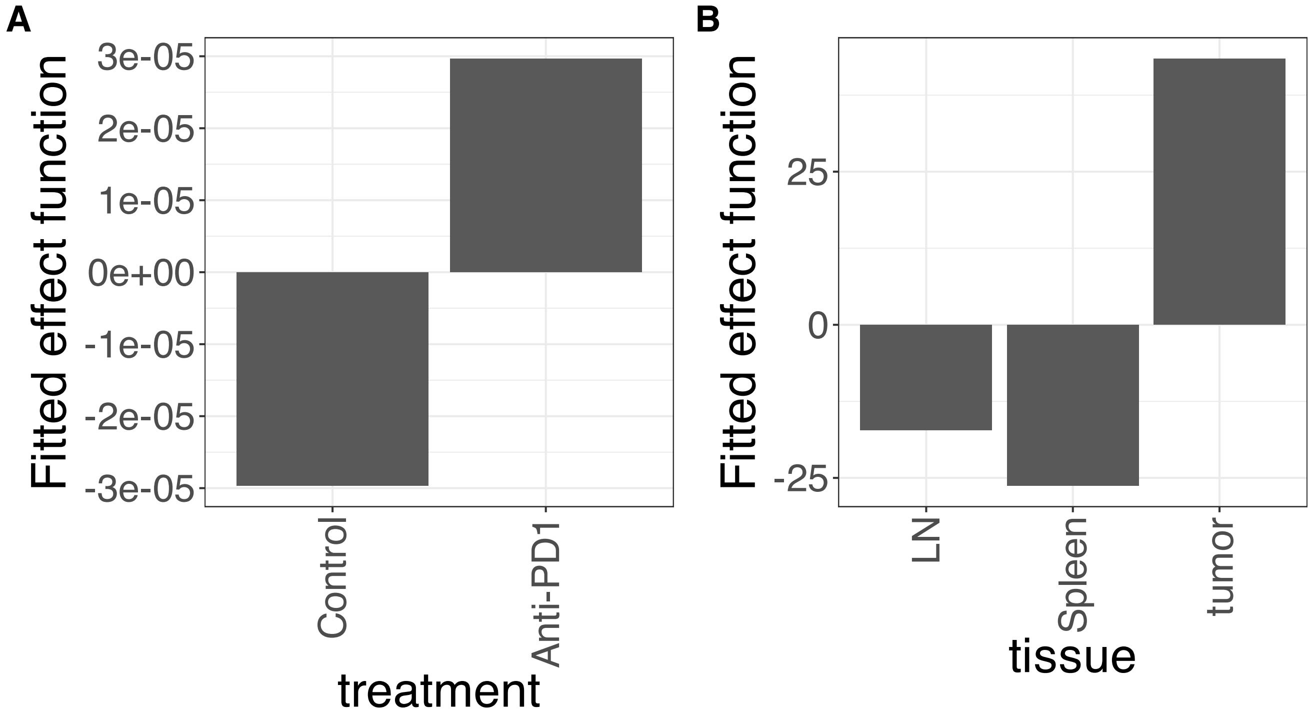


**Supplementary Figure 488.** Decomposition of reconstructed SSANOVA-SVD component function 1. (A) treatment. (B) tissue.

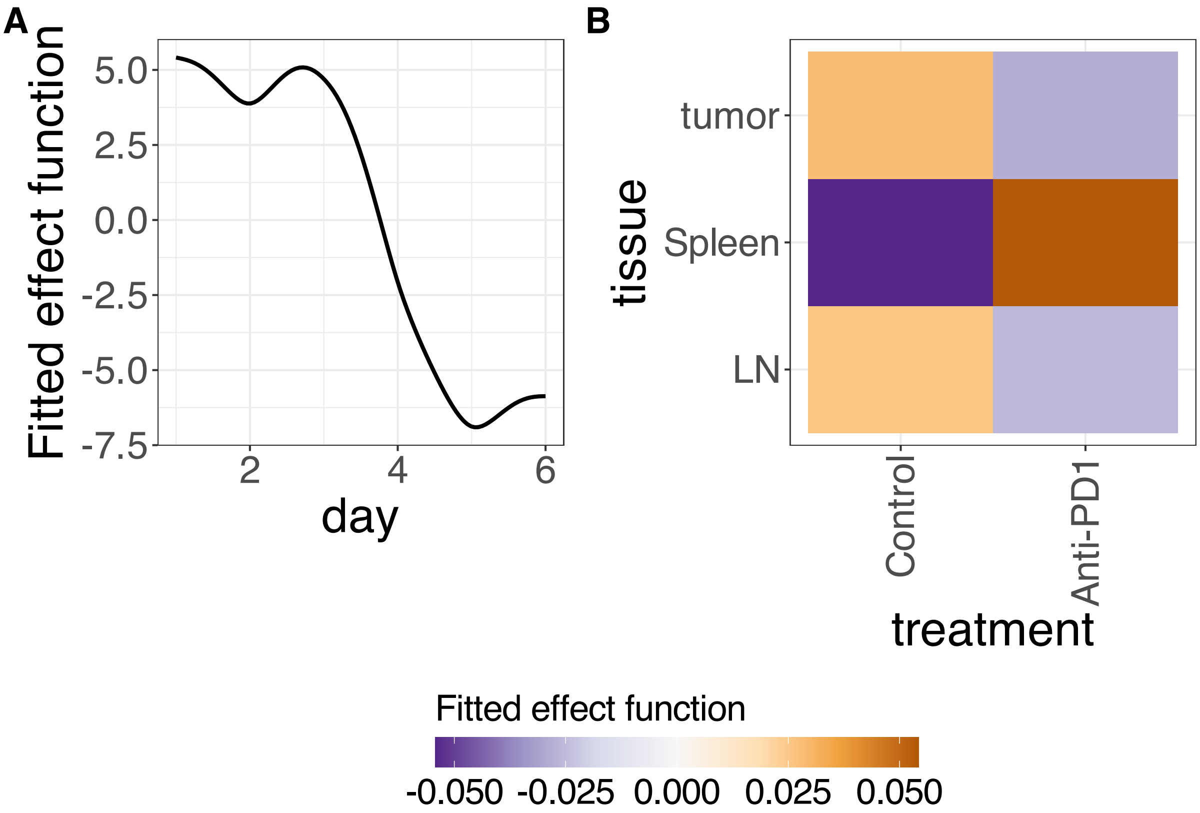


**Supplementary Figure 489.** Decomposition of reconstructed SSANOVA-SVD component function 1. (A) day. (B) treatment:tissue.

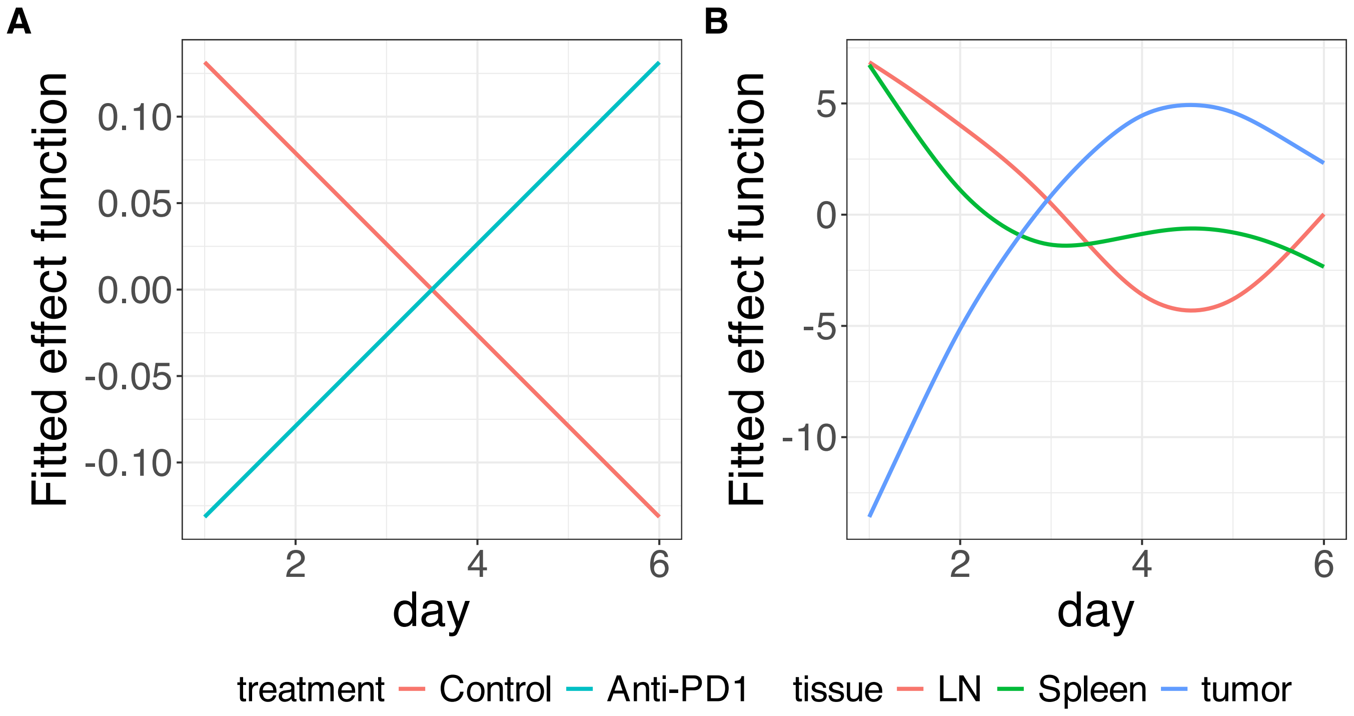


**Supplementary Figure 490.** Decomposition of reconstructed SSANOVA-SVD component function 1. (A) treatment:day. (B) tissue:day.

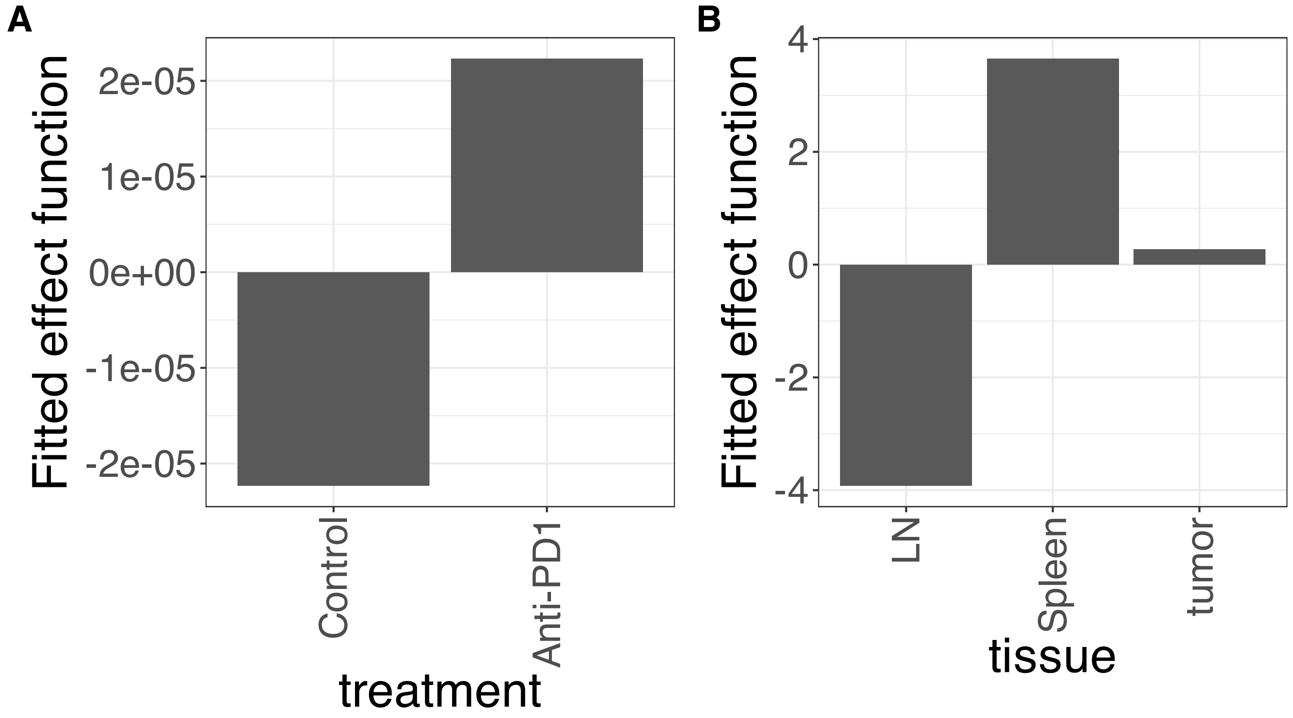


**Supplementary Figure 491.** Decomposition of reconstructed SSANOVA-SVD component function 2. (A) treatment. (B) tissue.

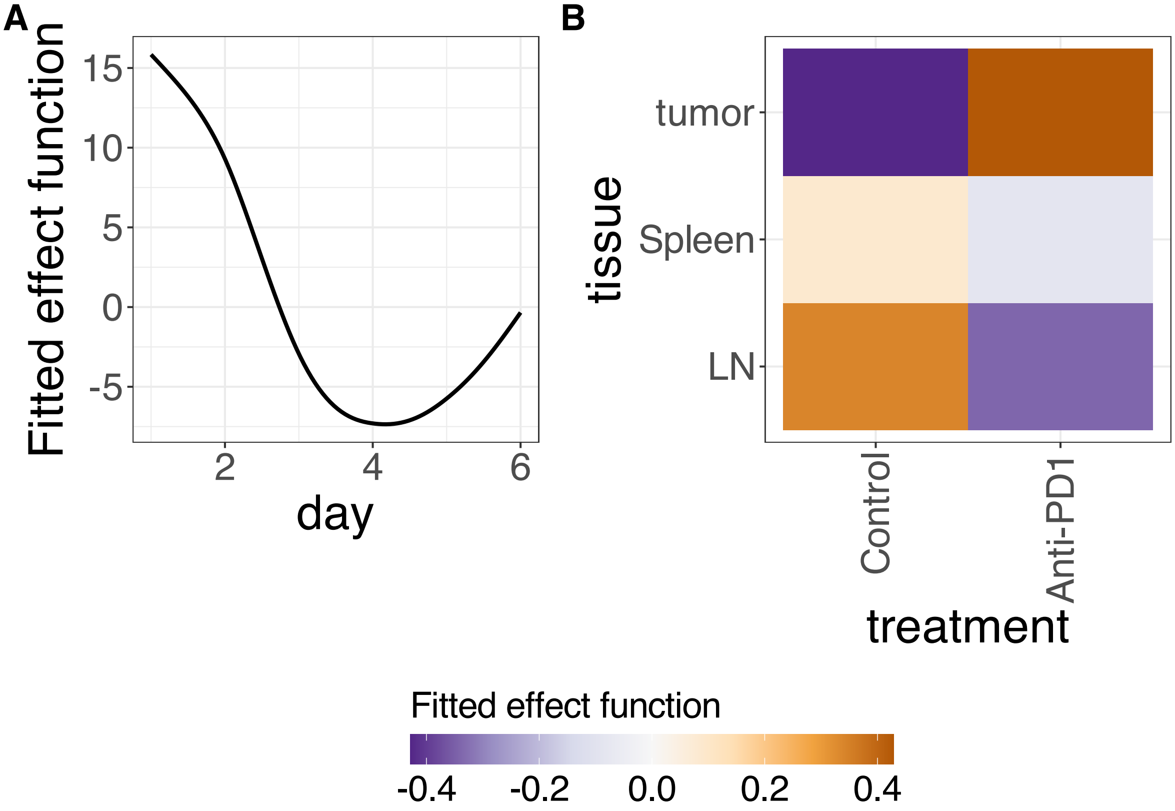


**Supplementary Figure 492.** Decomposition of reconstructed SSANOVA-SVD component function 2. (A) day. (B) treatment:tissue.

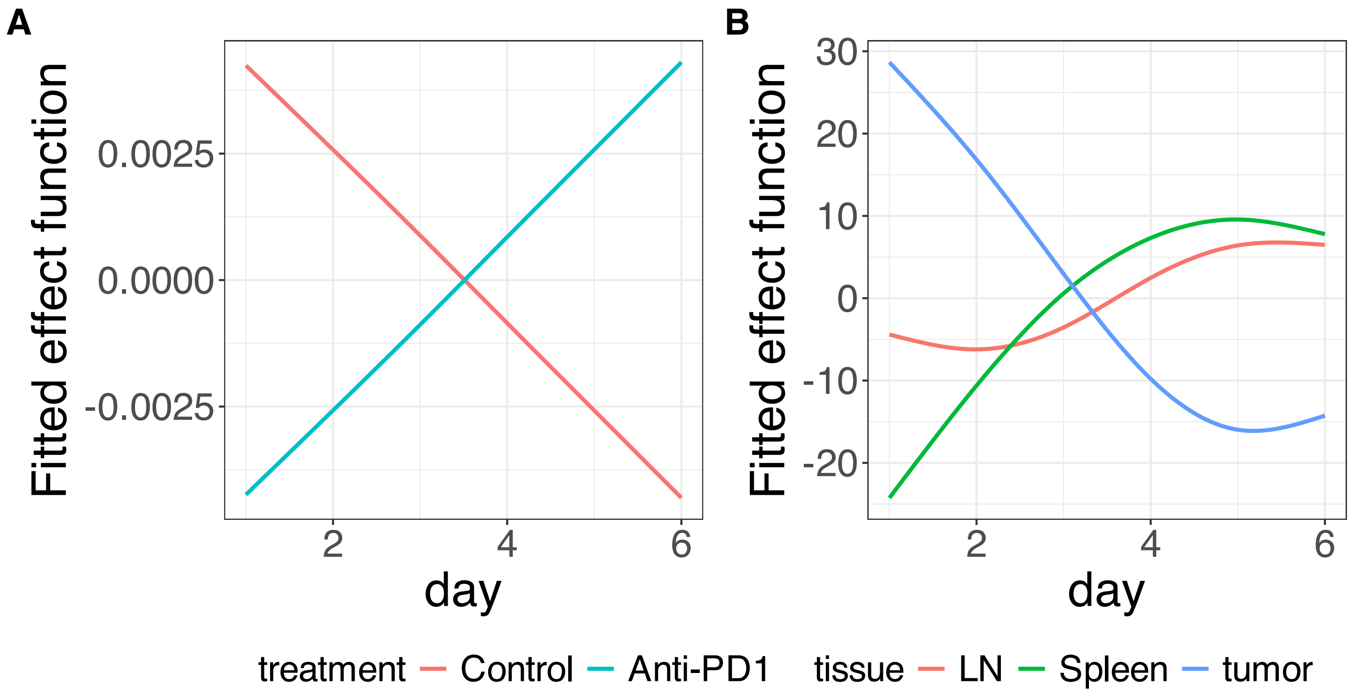


**Supplementary Figure 493.** Decomposition of reconstructed SSANOVA-SVD component function 2. (A) treatment:day. (B) tissue:day.

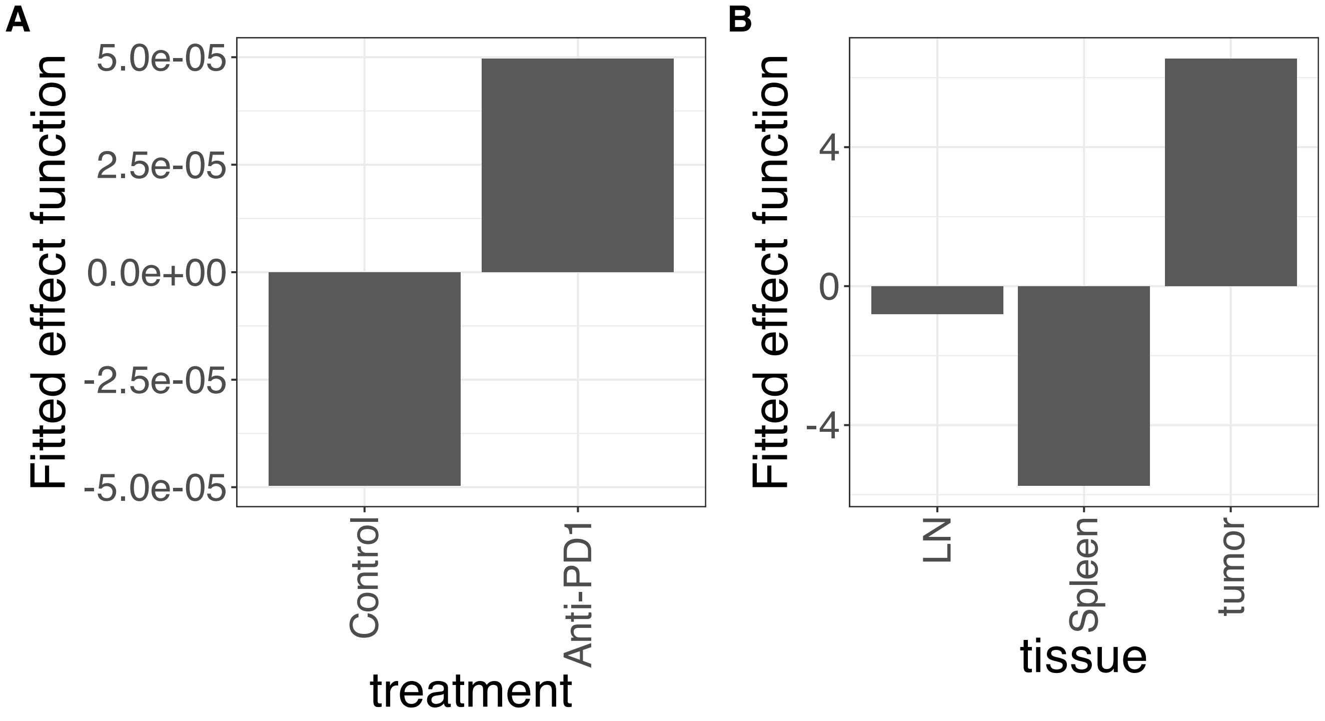


**Supplementary Figure 494.** Decomposition of reconstructed SSANOVA-SVD component function 3. (A) treatment. (B) tissue.

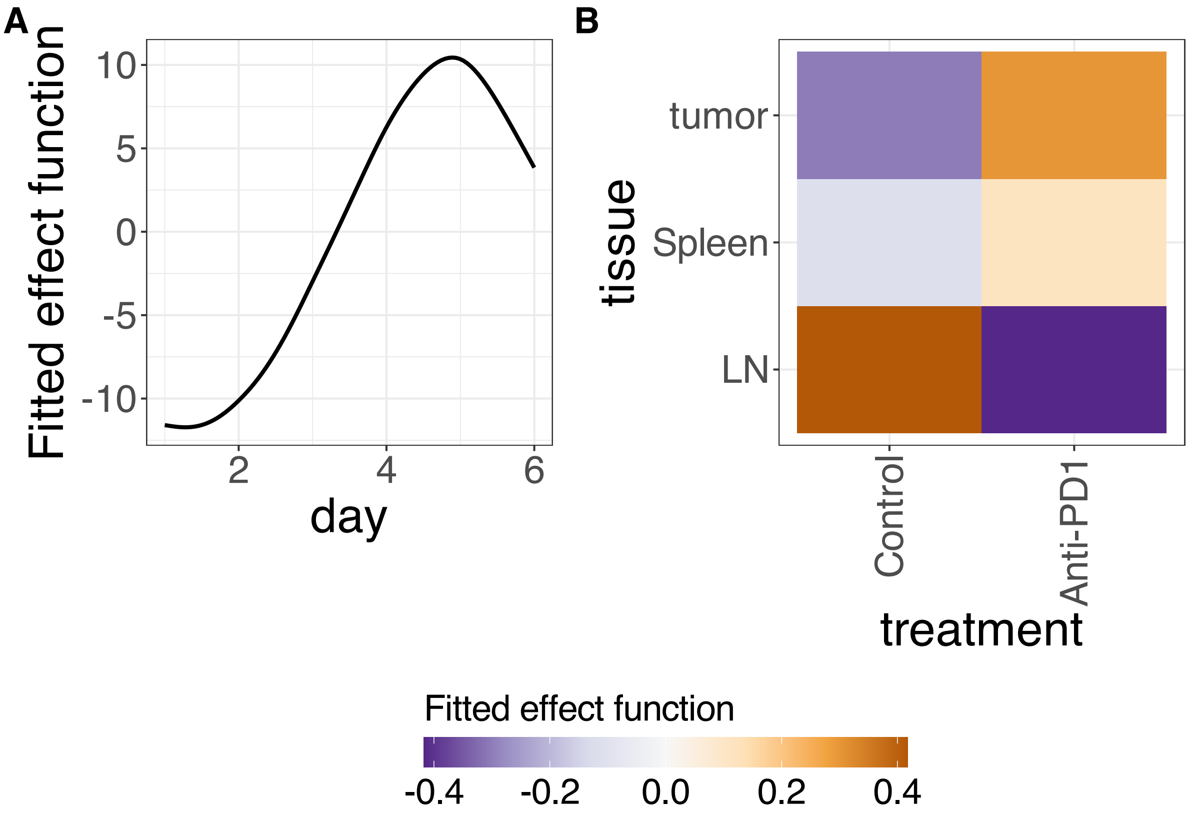


**Supplementary Figure 495.** Decomposition of reconstructed SSANOVA-SVD component function 3. (A) day. (B) treatment:tissue.

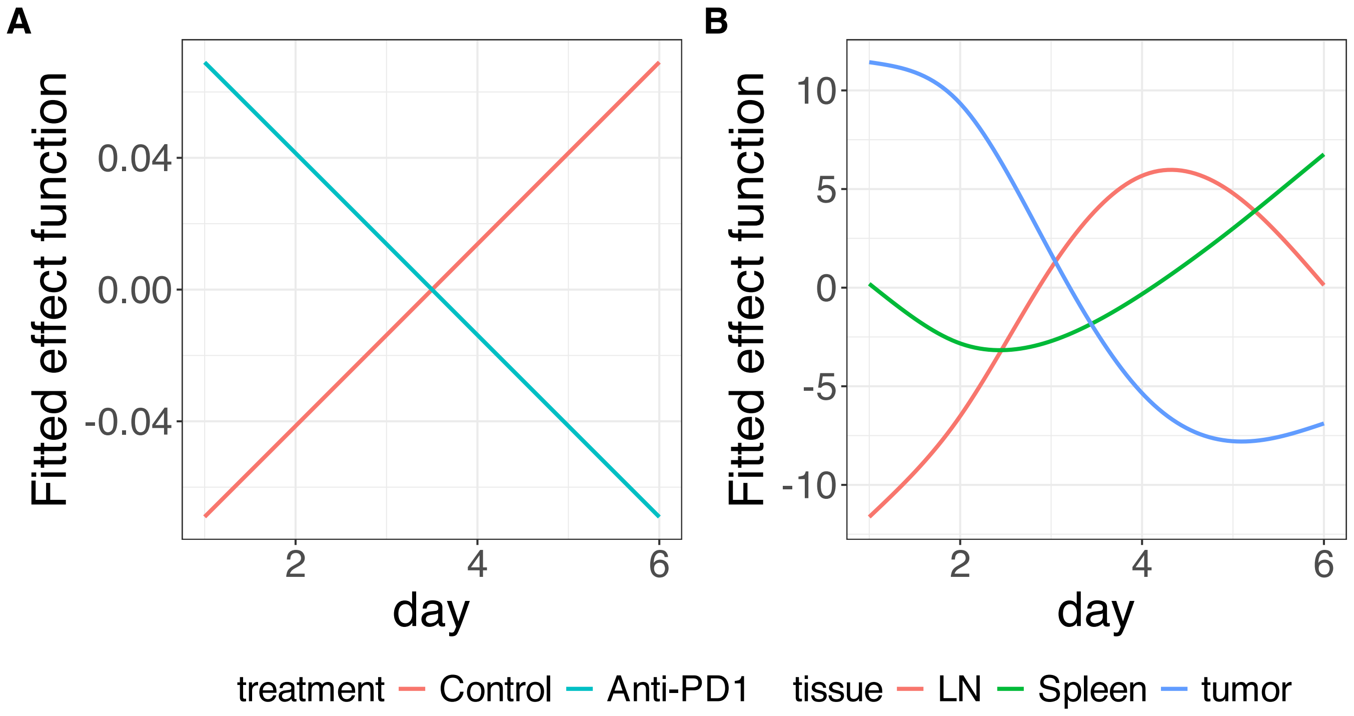


**Supplementary Figure 496.** Decomposition of reconstructed SSANOVA-SVD component function 3. (A) treatment:day. (B) tissue:day.

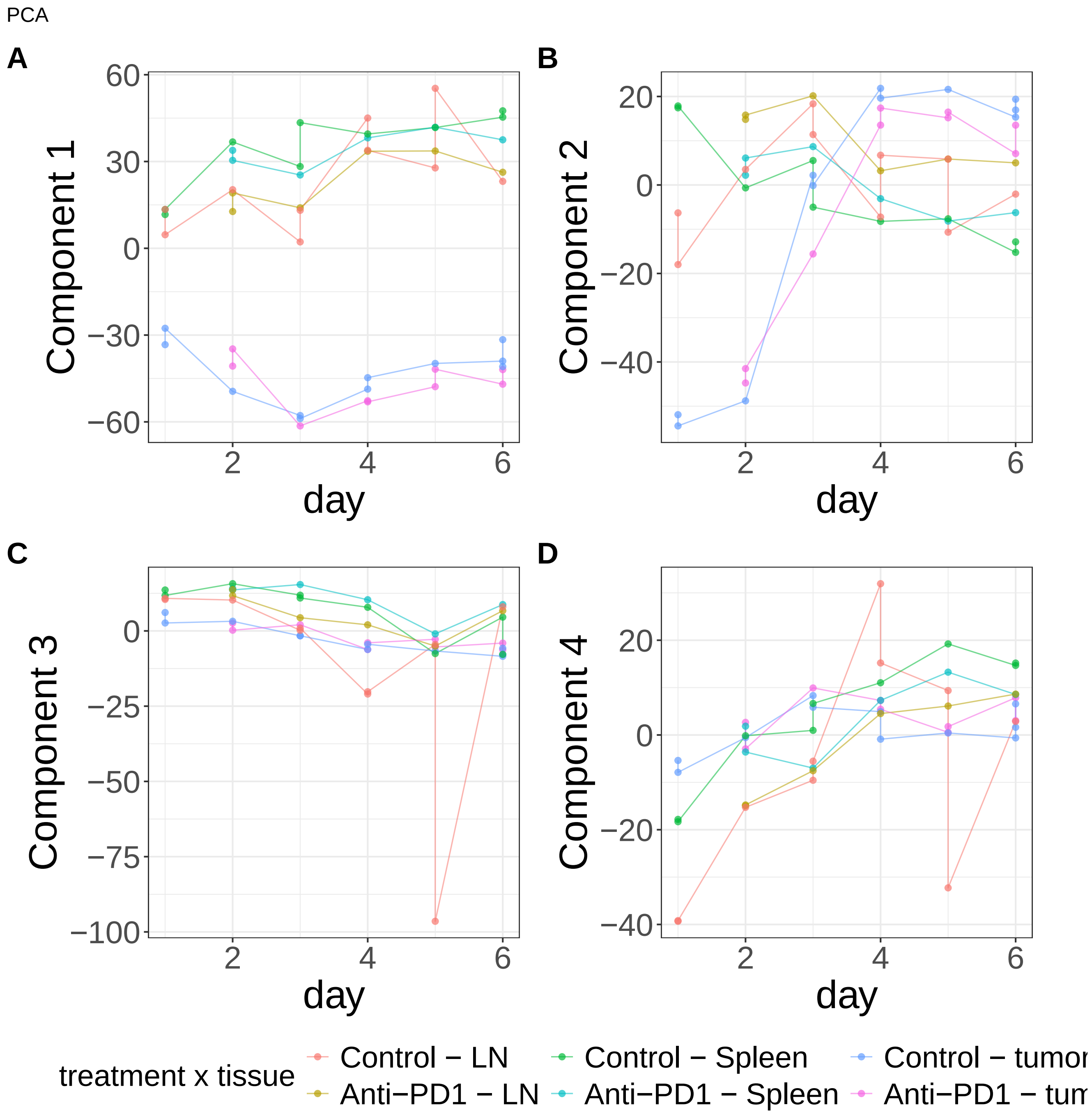


**Supplementary Figure 497.** Components for PCA. (A) Component 1. (B) Component 2. (C) Component 3. (D) Component 4.

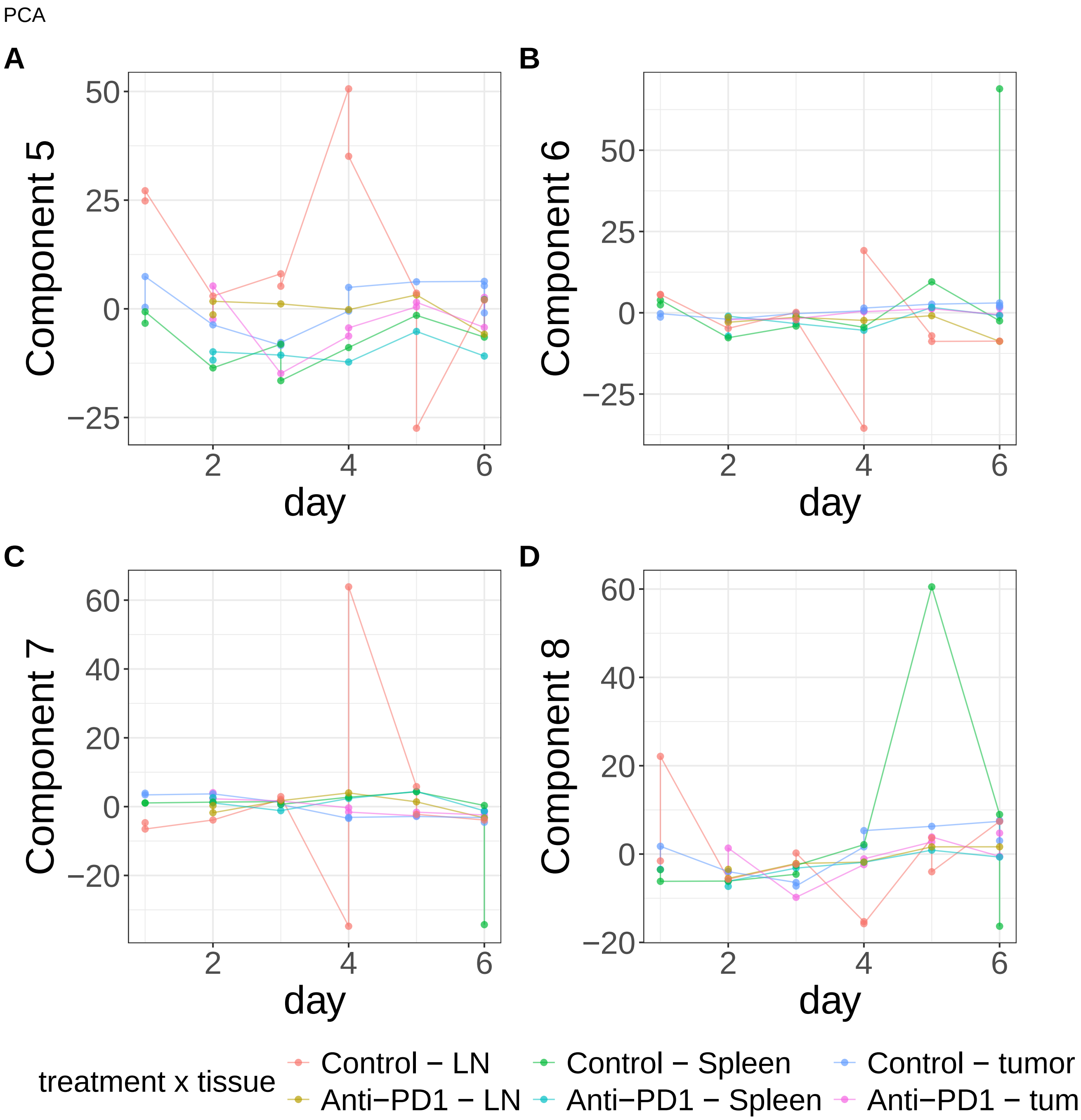


**Supplementary Figure 498.** Components for PCA. (A) Component 5. (B) Component 6. (C) Component 7. (D) Component 8.

PCA

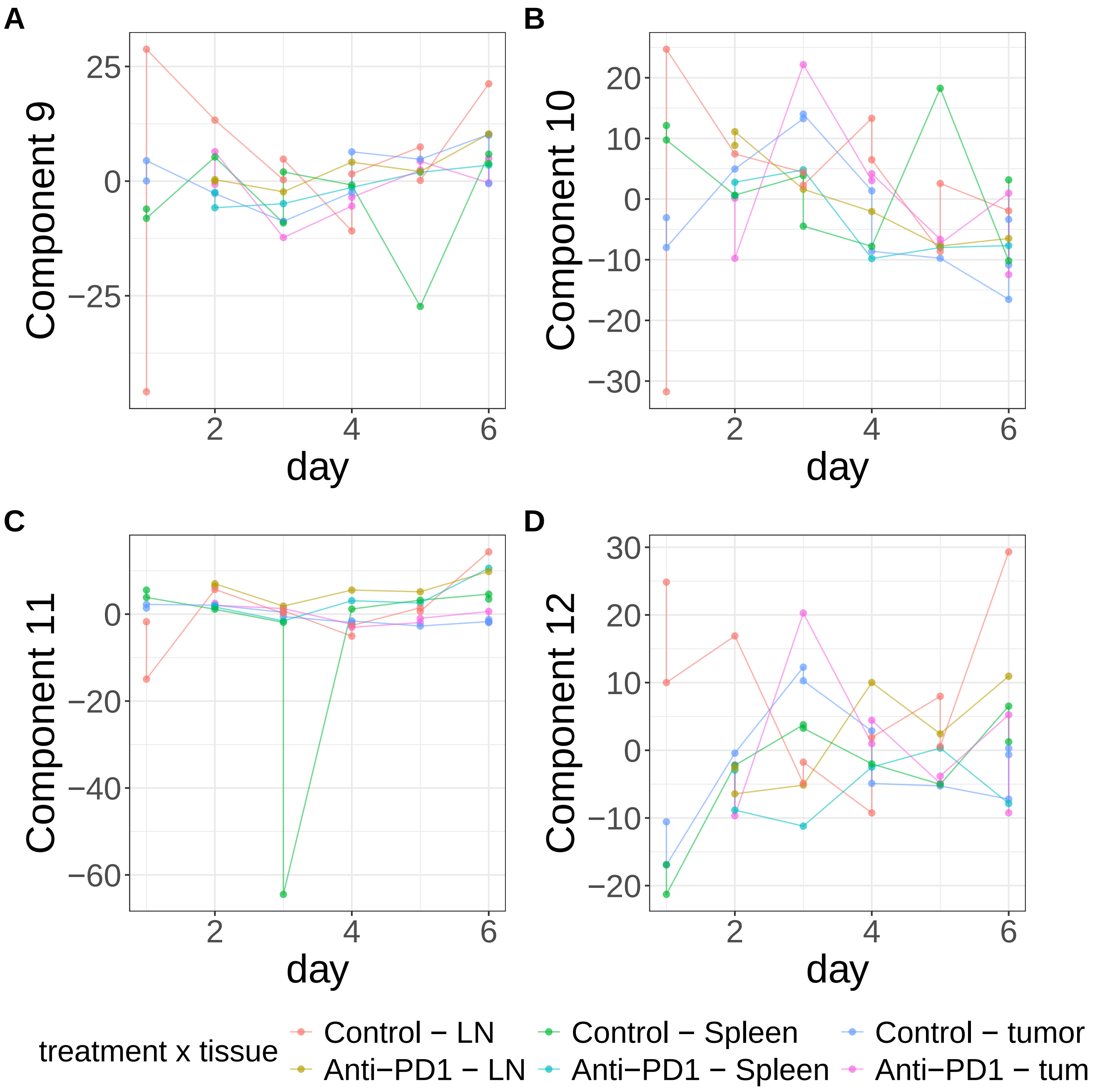


**Supplementary Figure 499.** Components for PCA. (A) Component 9. (B) Component 10. (C) Component 11. (D) Component 12.

PCA

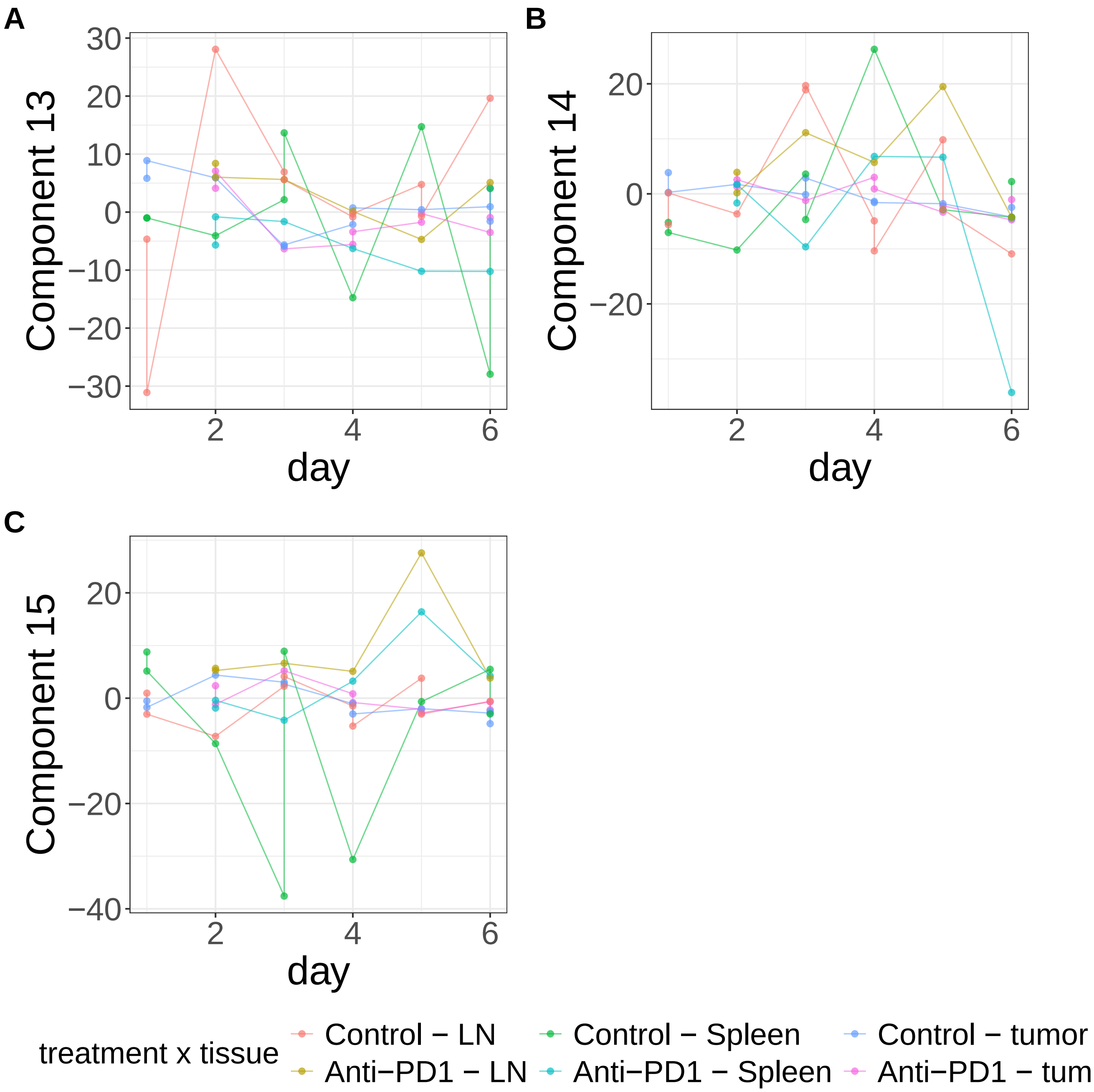


**Supplementary Figure 500.** Components for PCA. (A) Component 13. (B) Component 14. (C) Component 15.

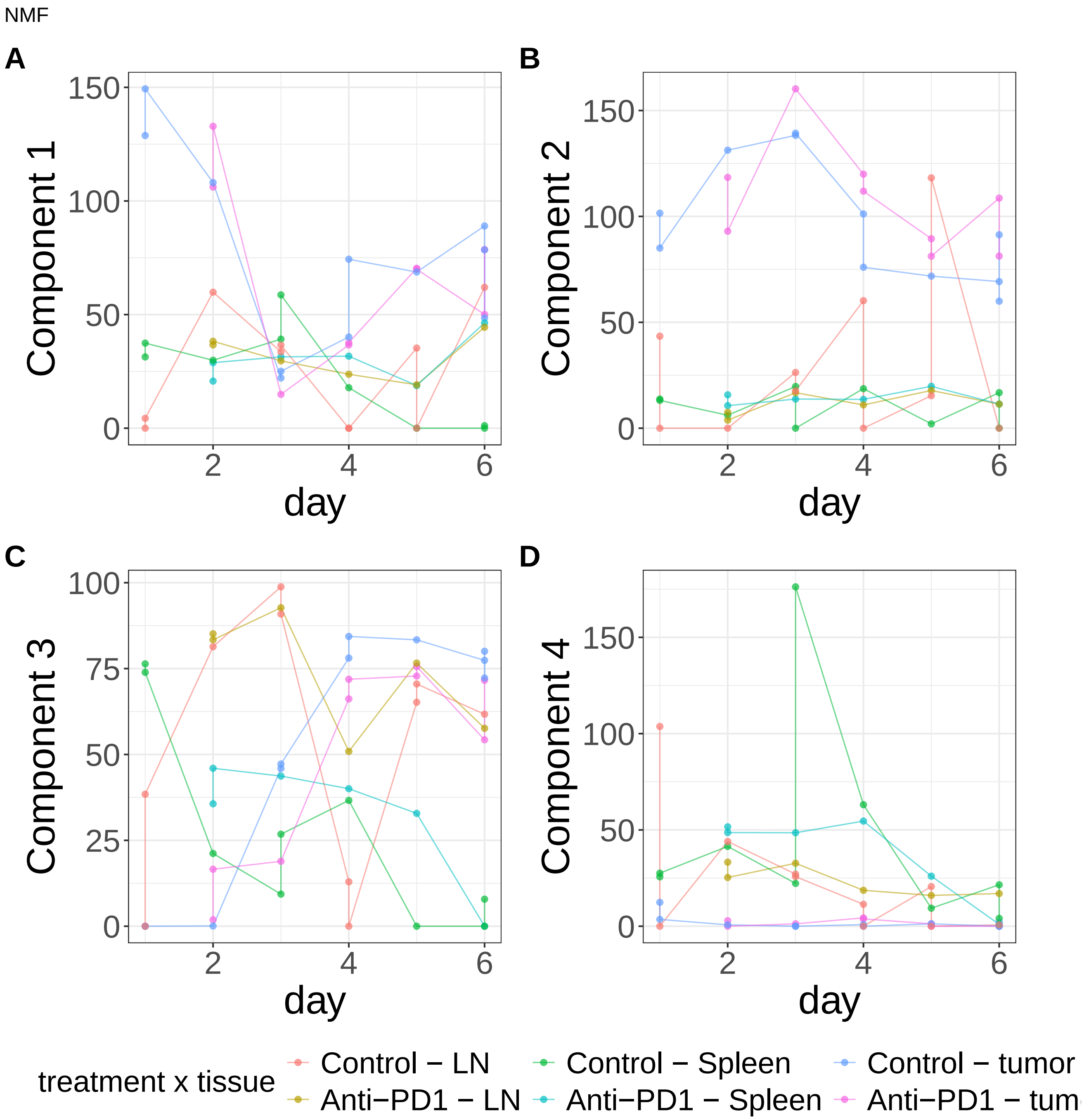


**Supplementary Figure 501.** Components for NMF. (A) Component 1. (B) Component 2. (C) Component 3. (D) Component 4.

NMF

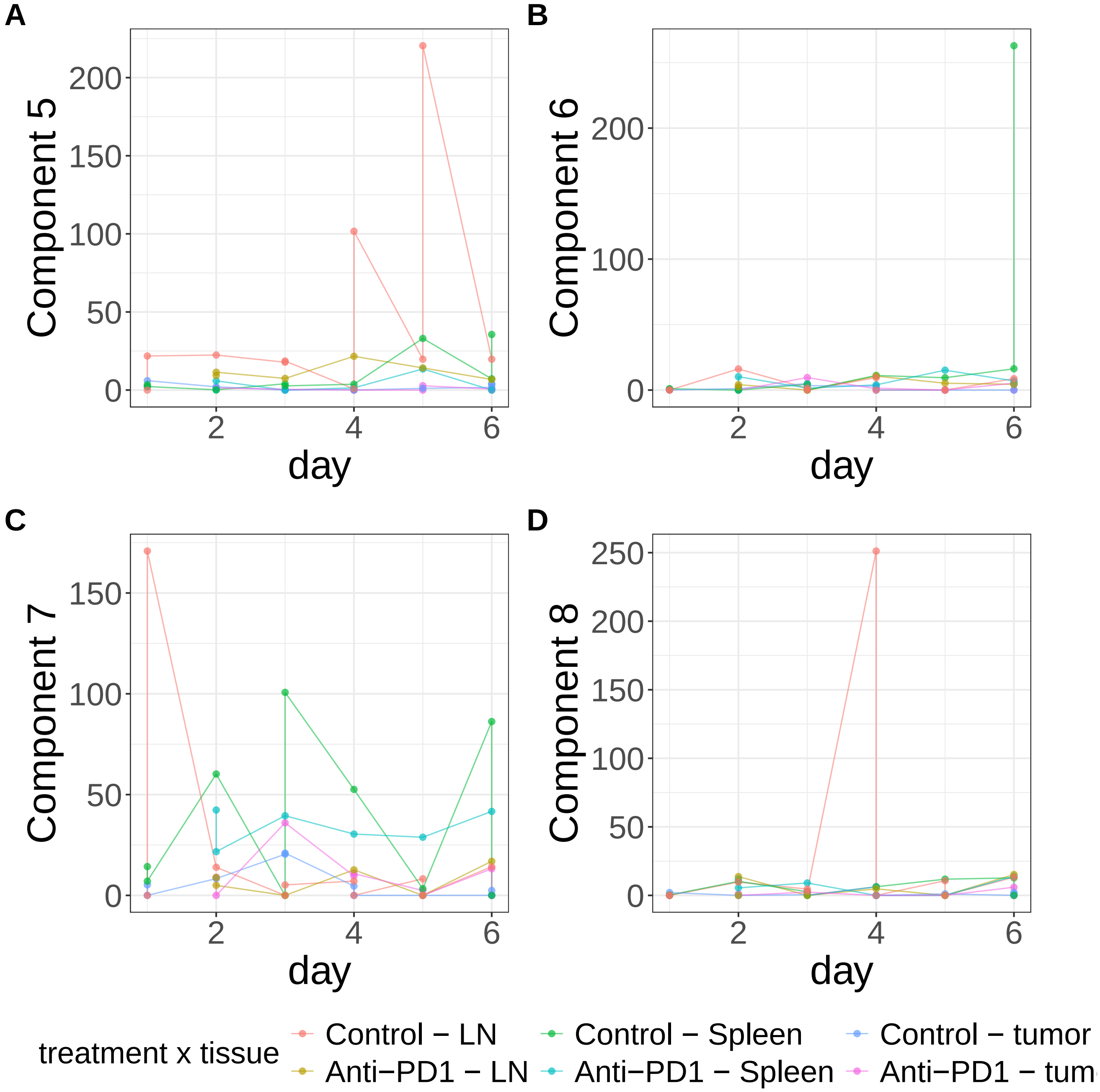


**Supplementary Figure 502.** Components for NMF. (A) Component 5. (B) Component 6. (C) Component 7. (D) Component 8.

NMF

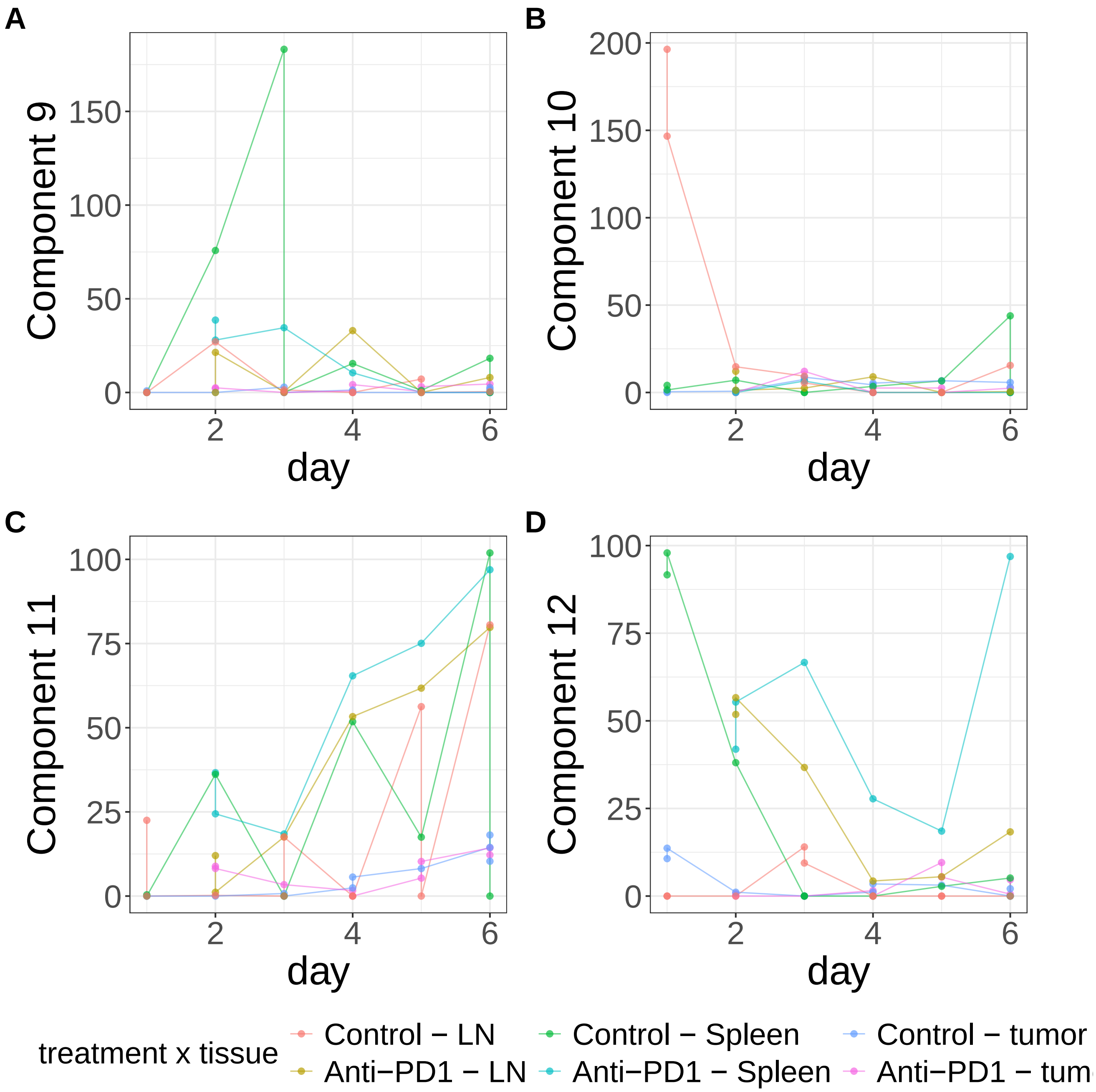


**Supplementary Figure 503.** Components for NMF. (A) Component 9. (B) Component 10. (C) Component 11. (D) Component 12.

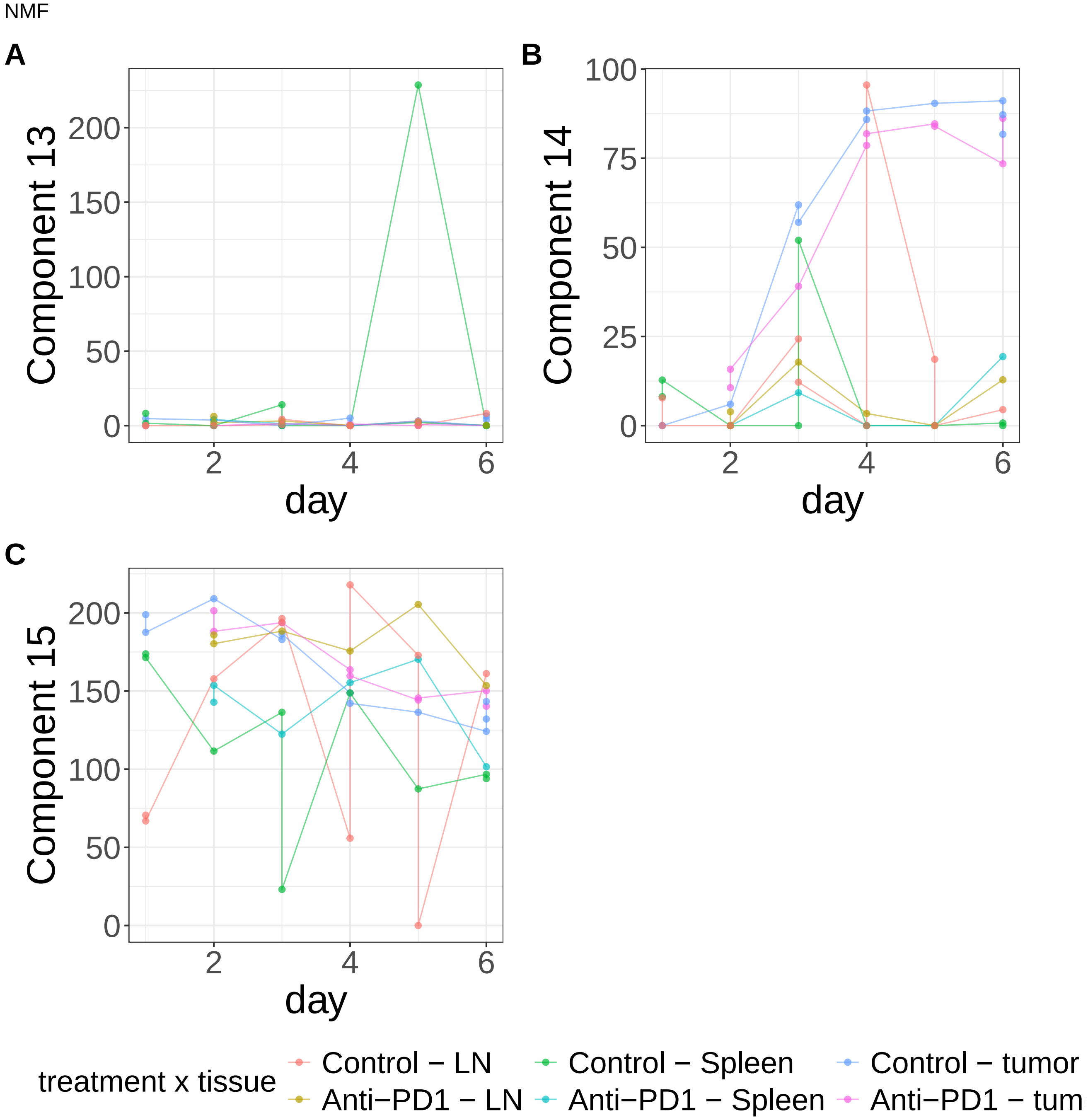


**Supplementary Figure 504.** Components for NMF. (A) Component 13. (B) Component 14. (C) Component 15.

PLS

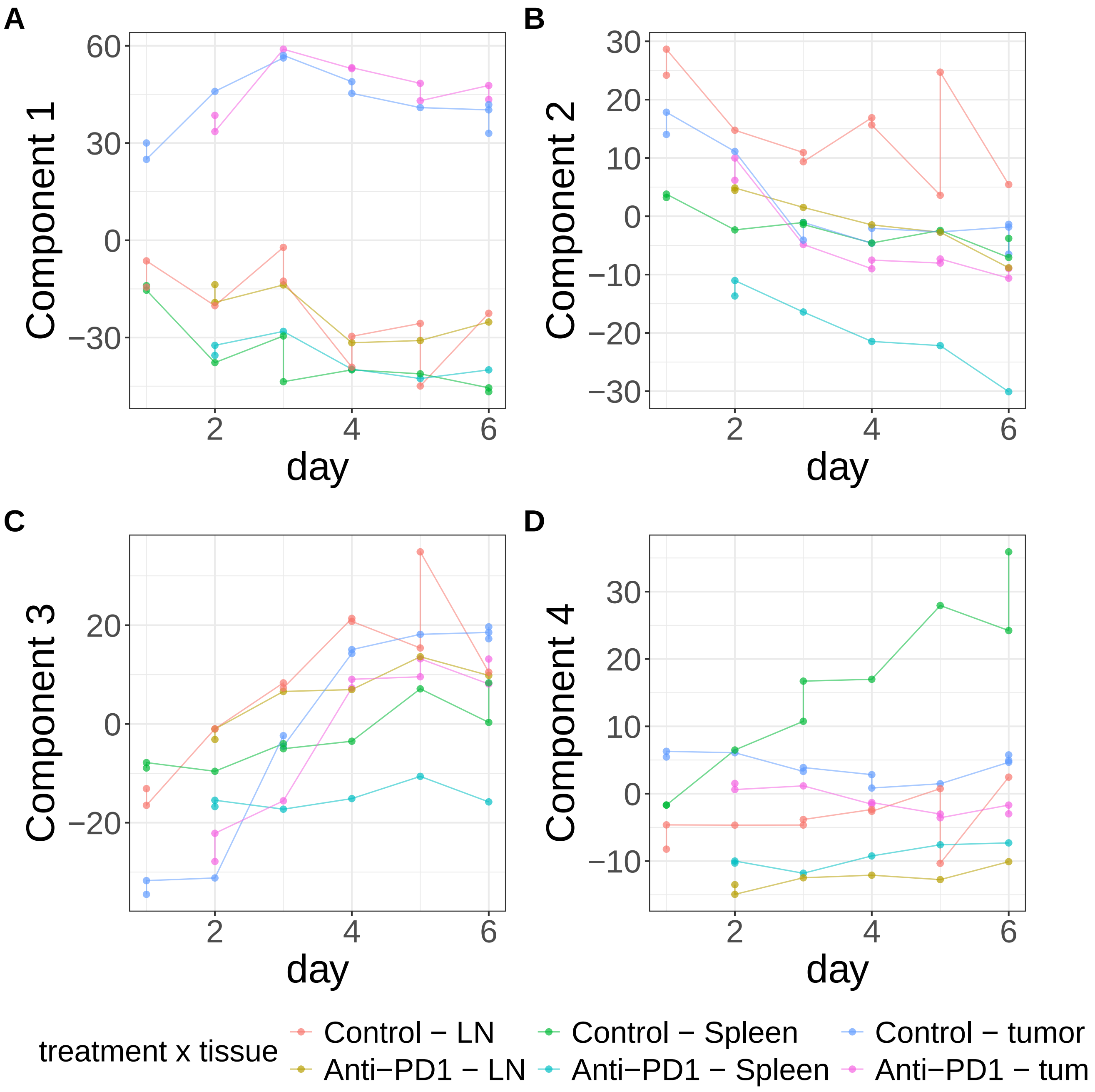


**Supplementary Figure 505.** Components for PLS. (A) Component 1. (B) Component 2. (C) Component 3. (D) Component 4.

PLS

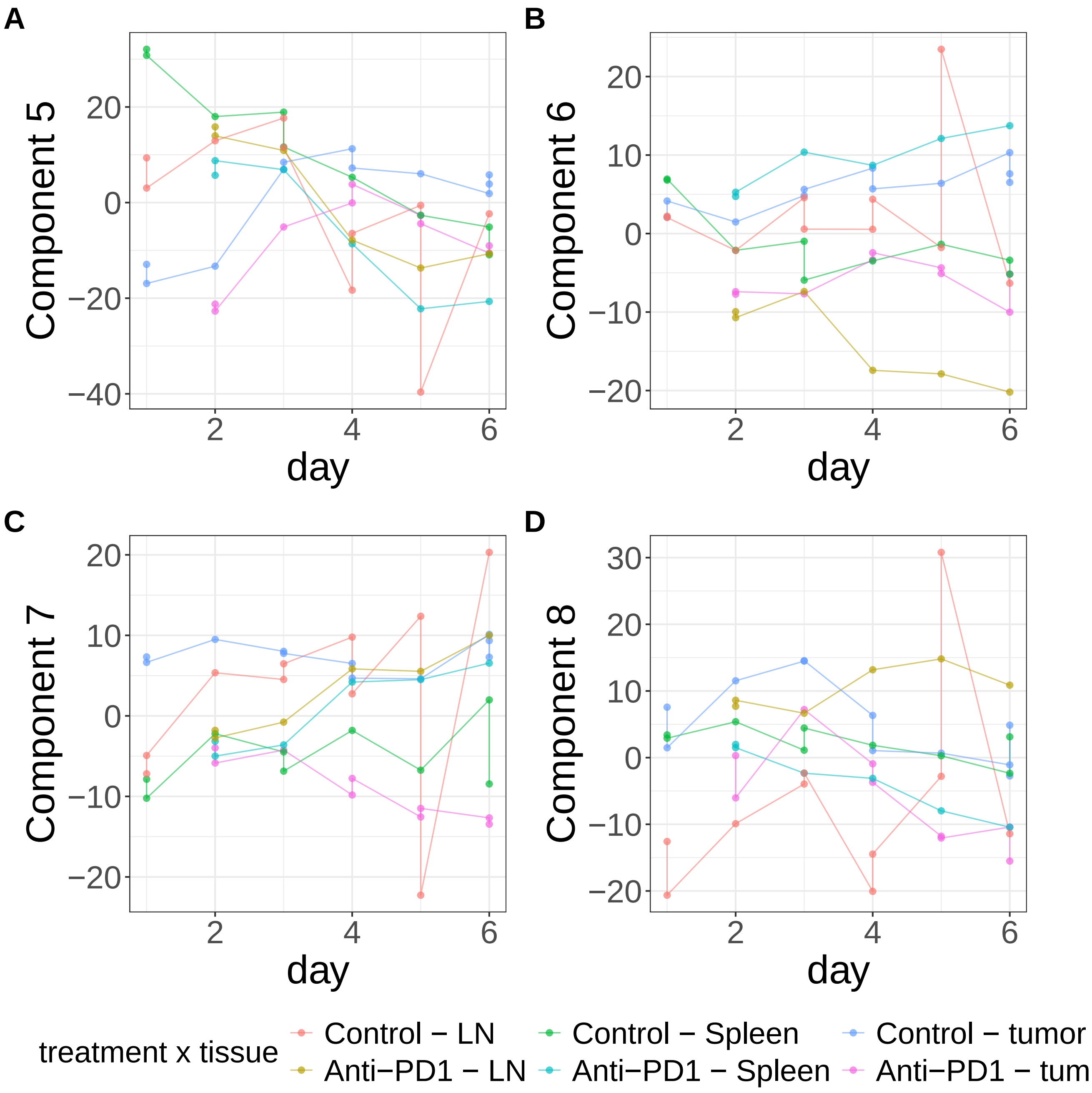


**Supplementary Figure 506.** Components for PLS. (A) Component 5. (B) Component 6. (C) Component 7. (D) Component 8.

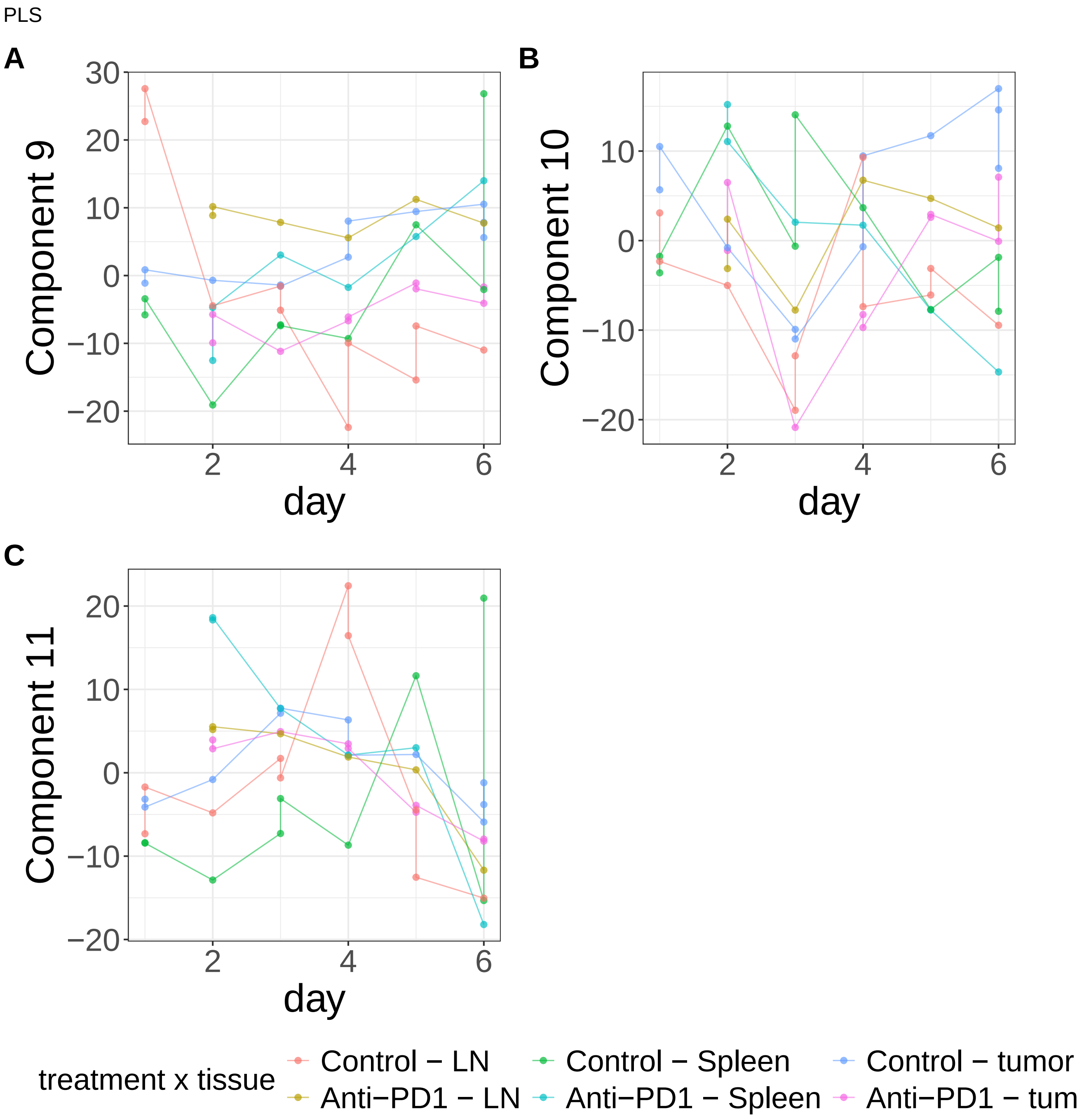


**Supplementary Figure 507.** Components for PLS. (A) Component 9. (B) Component 10. (C) Component 11.

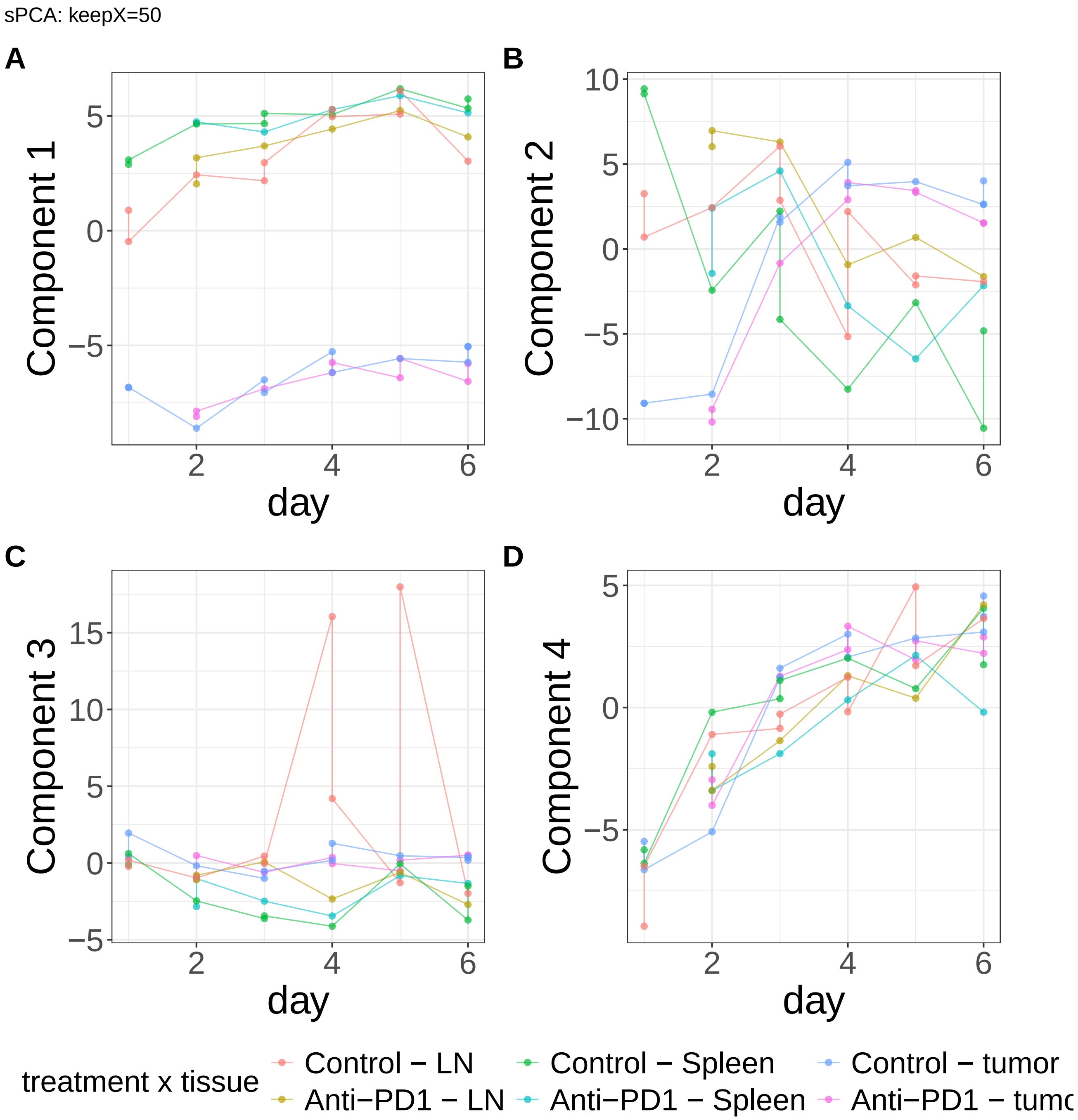


**Supplementary Figure 508.** Components for sPCA: keepX=50. (A) Component 1. (B) Component 2. (C) Component 3. (D) Component 4.

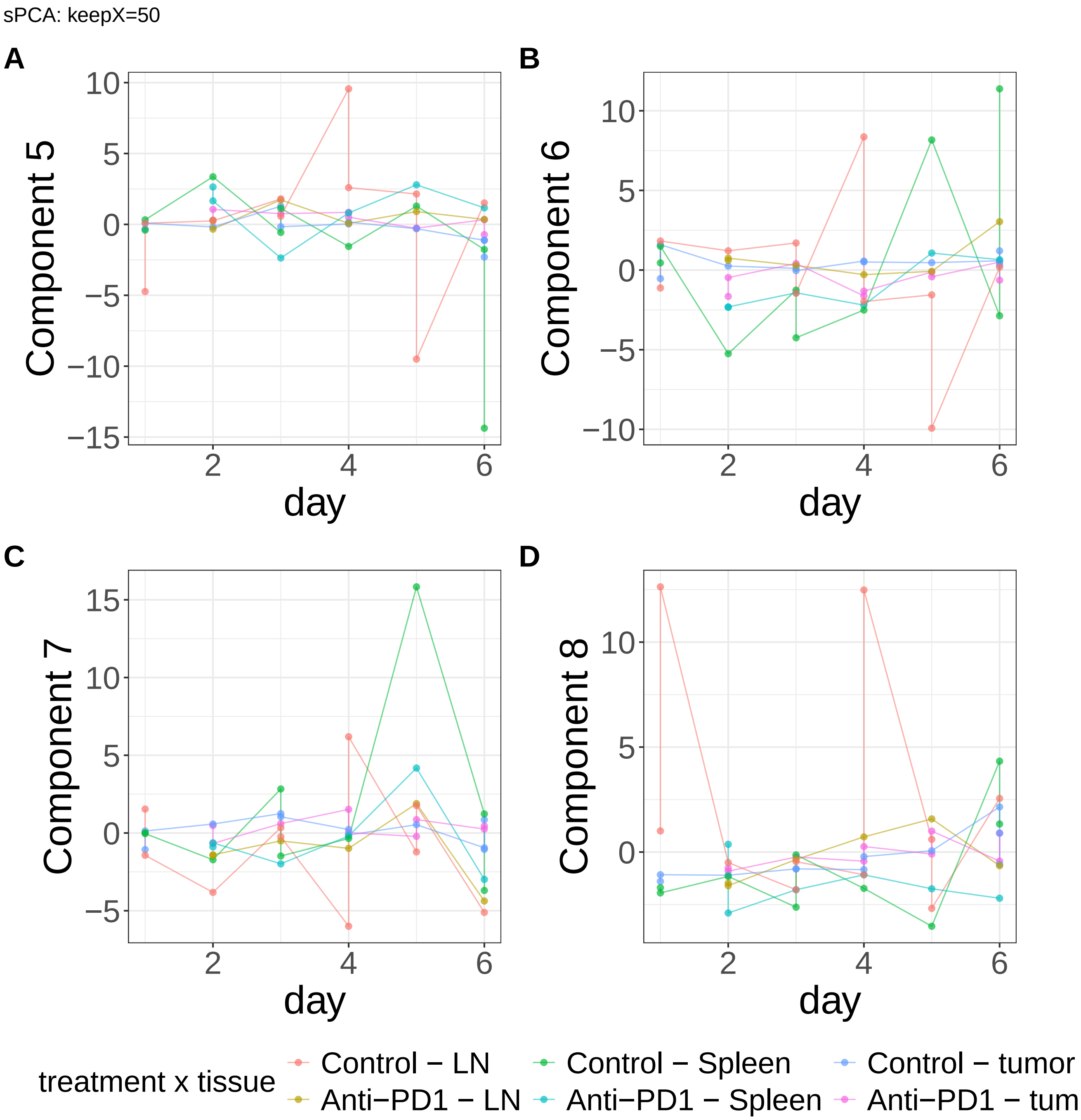


**Supplementary Figure 509.** Components for sPCA: keepX=50. (A) Component 5. (B) Component 6. (C) Component 7. (D) Component 8.

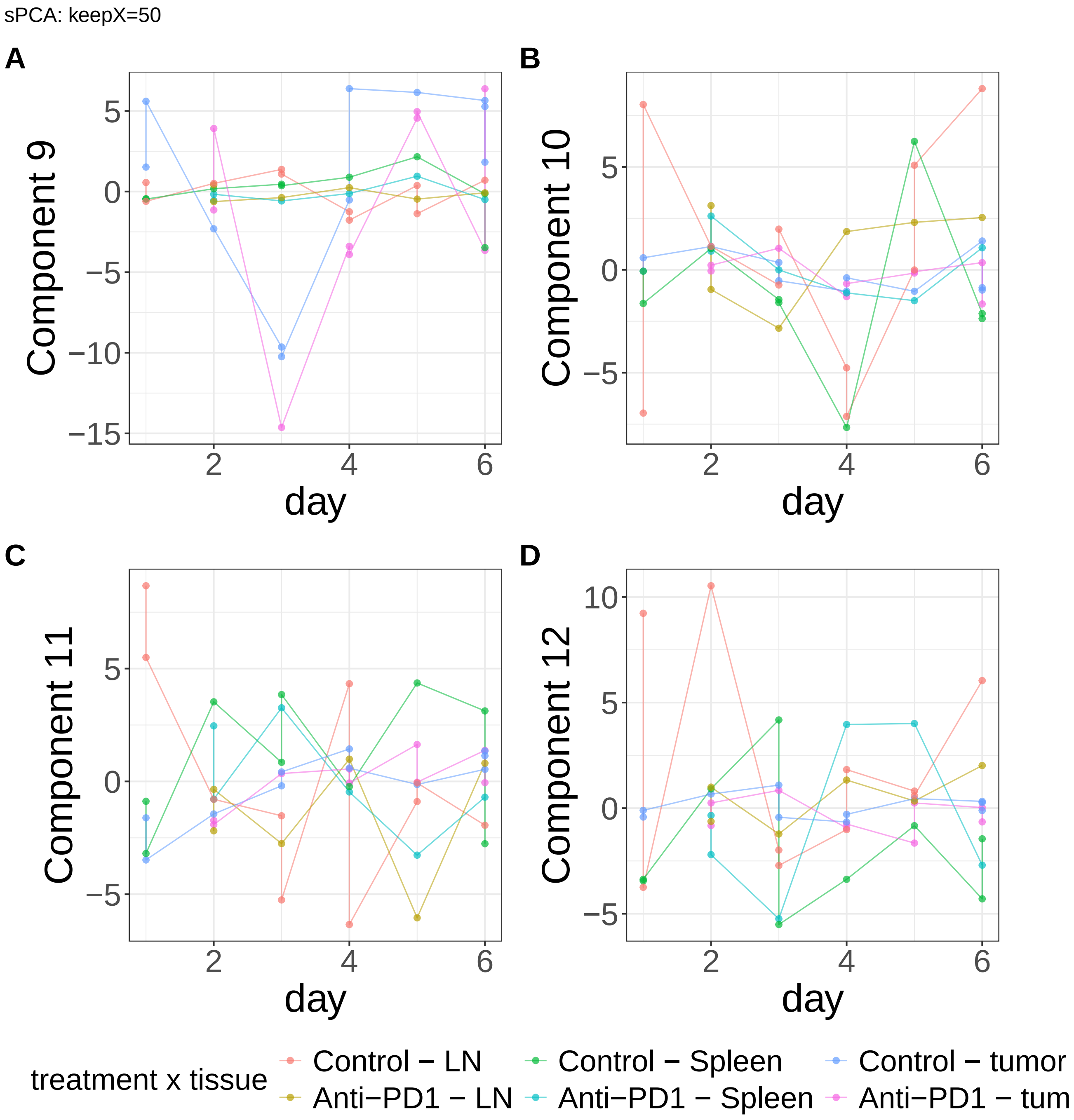


**Supplementary Figure 510.** Components for sPCA: keepX=50. (A) Component 9. (B) Component 10. (C) Component 11. (D) Component 12.

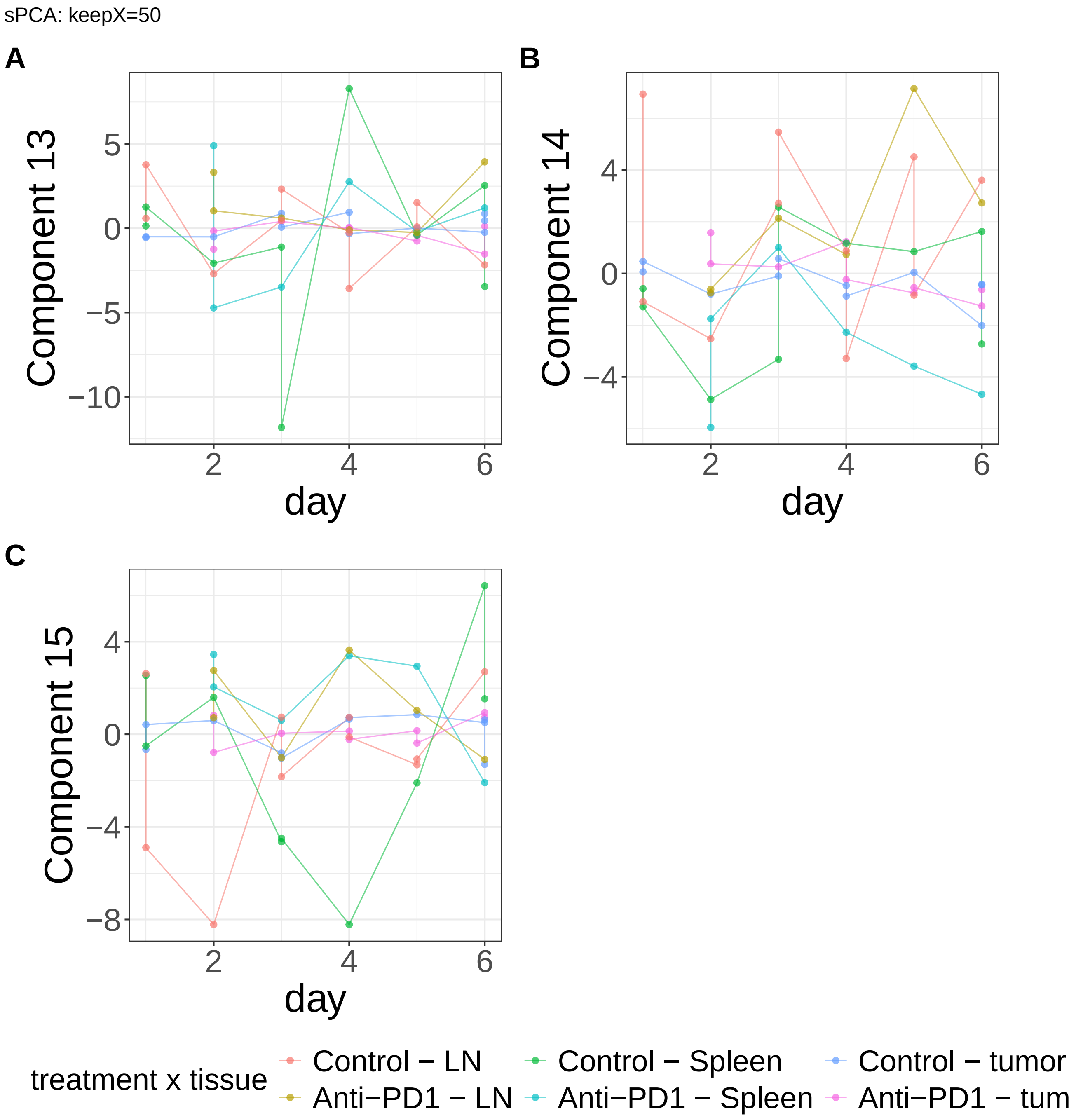


**Supplementary Figure 511.** Components for sPCA: keepX=50. (A) Component 13. (B) Component 14. (C) Component 15.

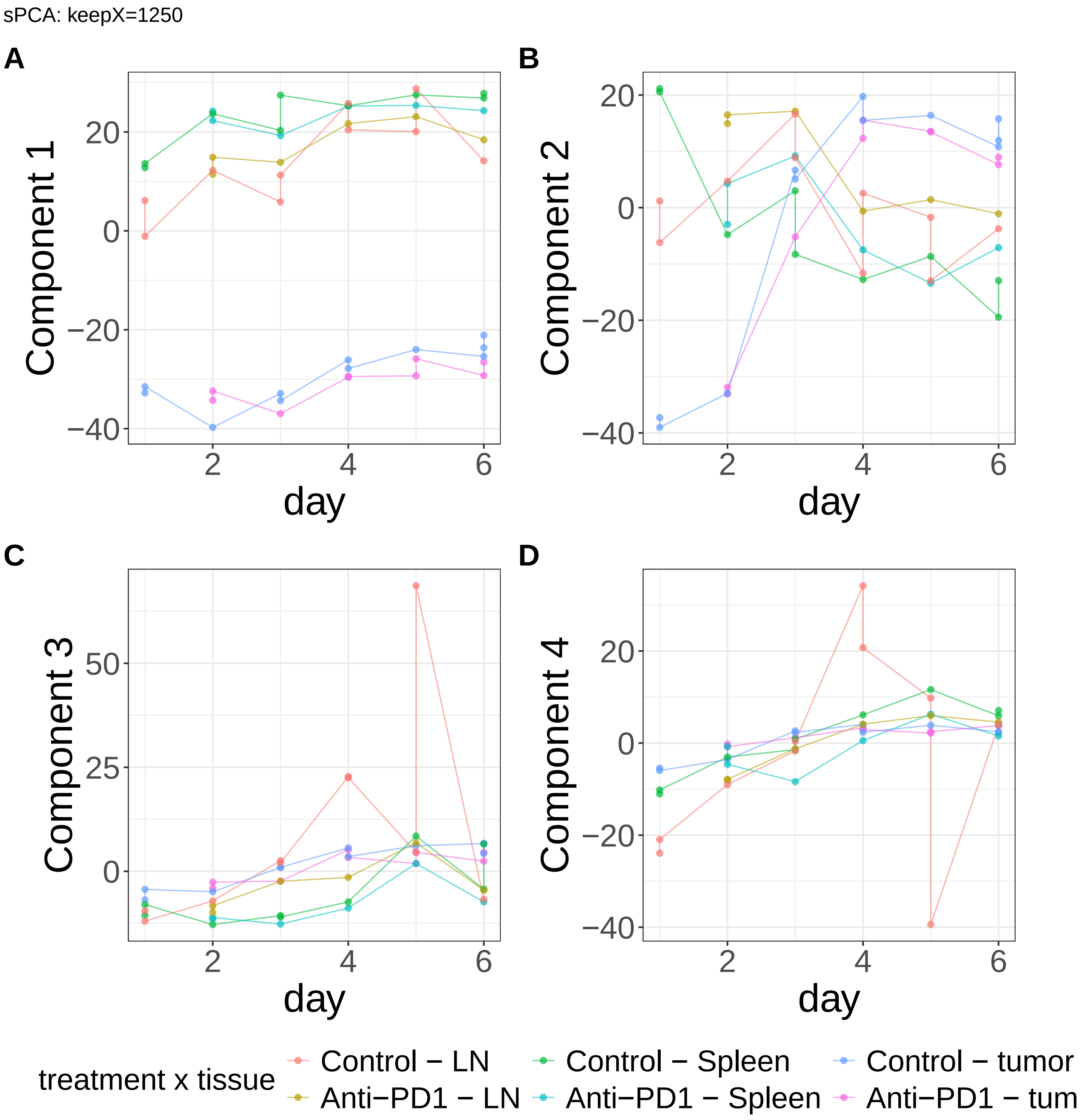


**Supplementary Figure 512.** Components for sPCA: keepX=1250. (A) Component 1. (B) Component 2. (C) Component 3. (D) Component 4.

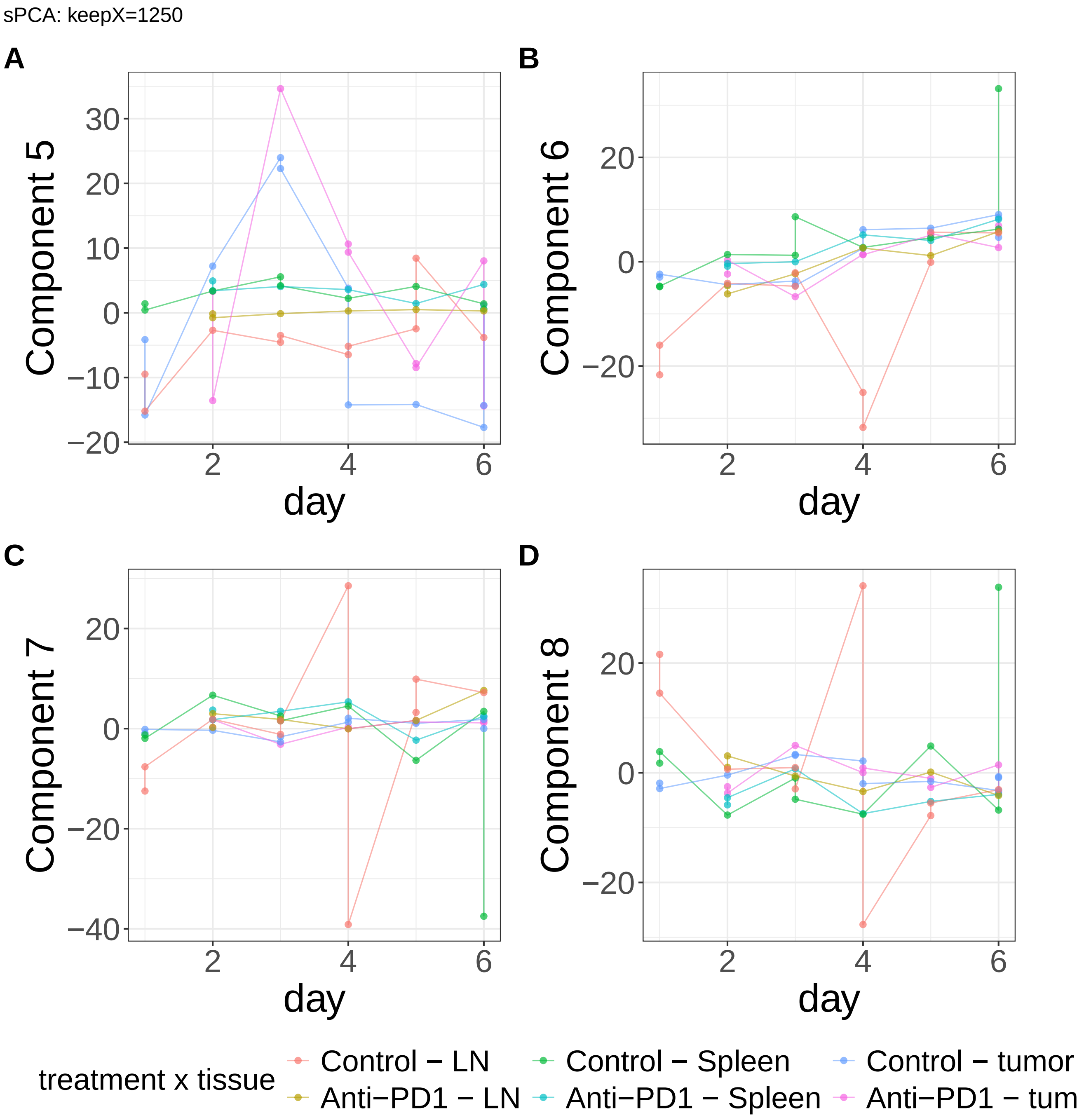


**Supplementary Figure 513.** Components for sPCA: keepX=1250. (A) Component 5. (B) Component 6. (C) Component 7. (D) Component 8.

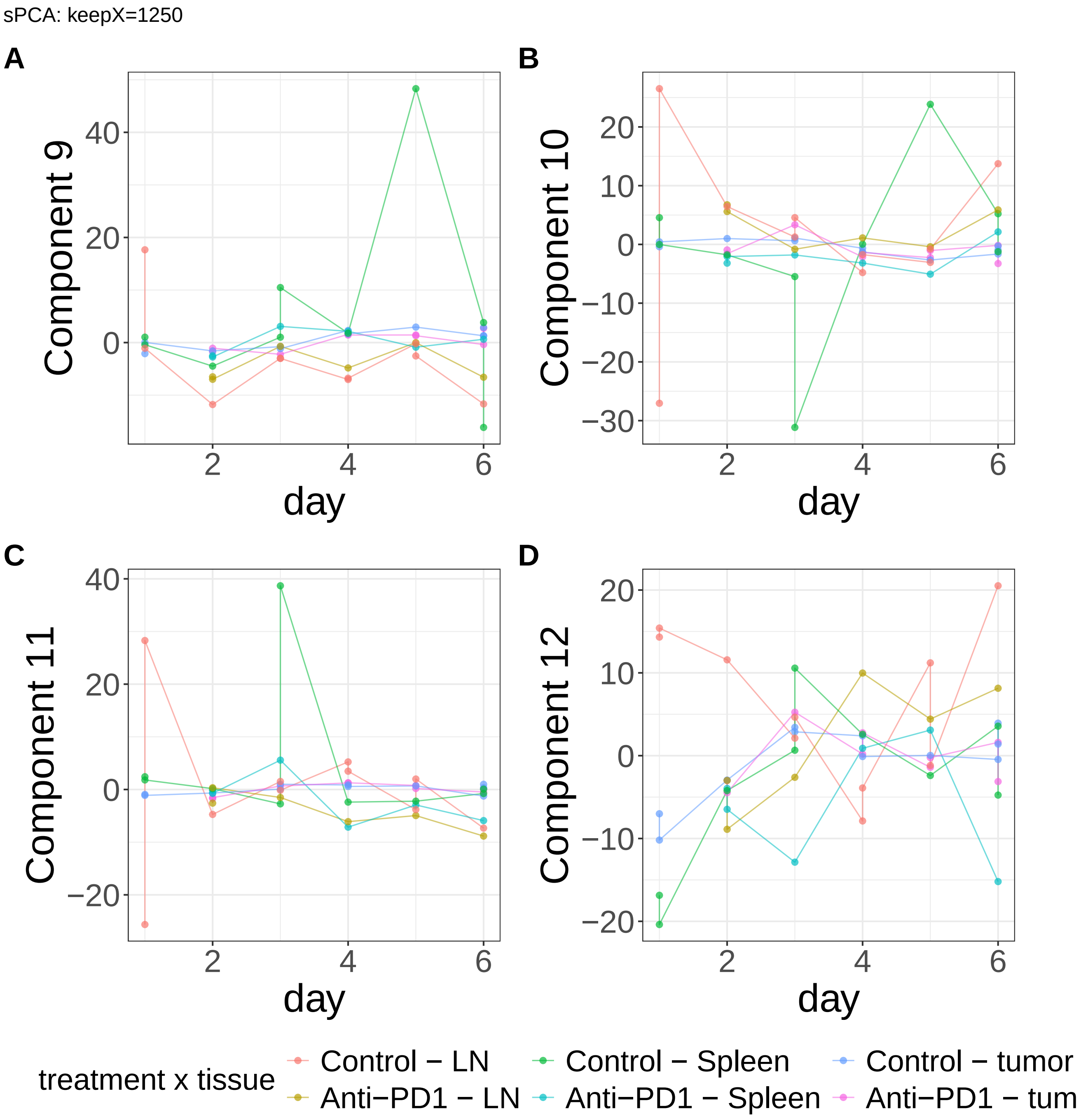


**Supplementary Figure 514.** Components for sPCA: keepX=1250. (A) Component 9. (B) Component 10. (C) Component 11. (D) Component 12.

sPCA: keepX=1250

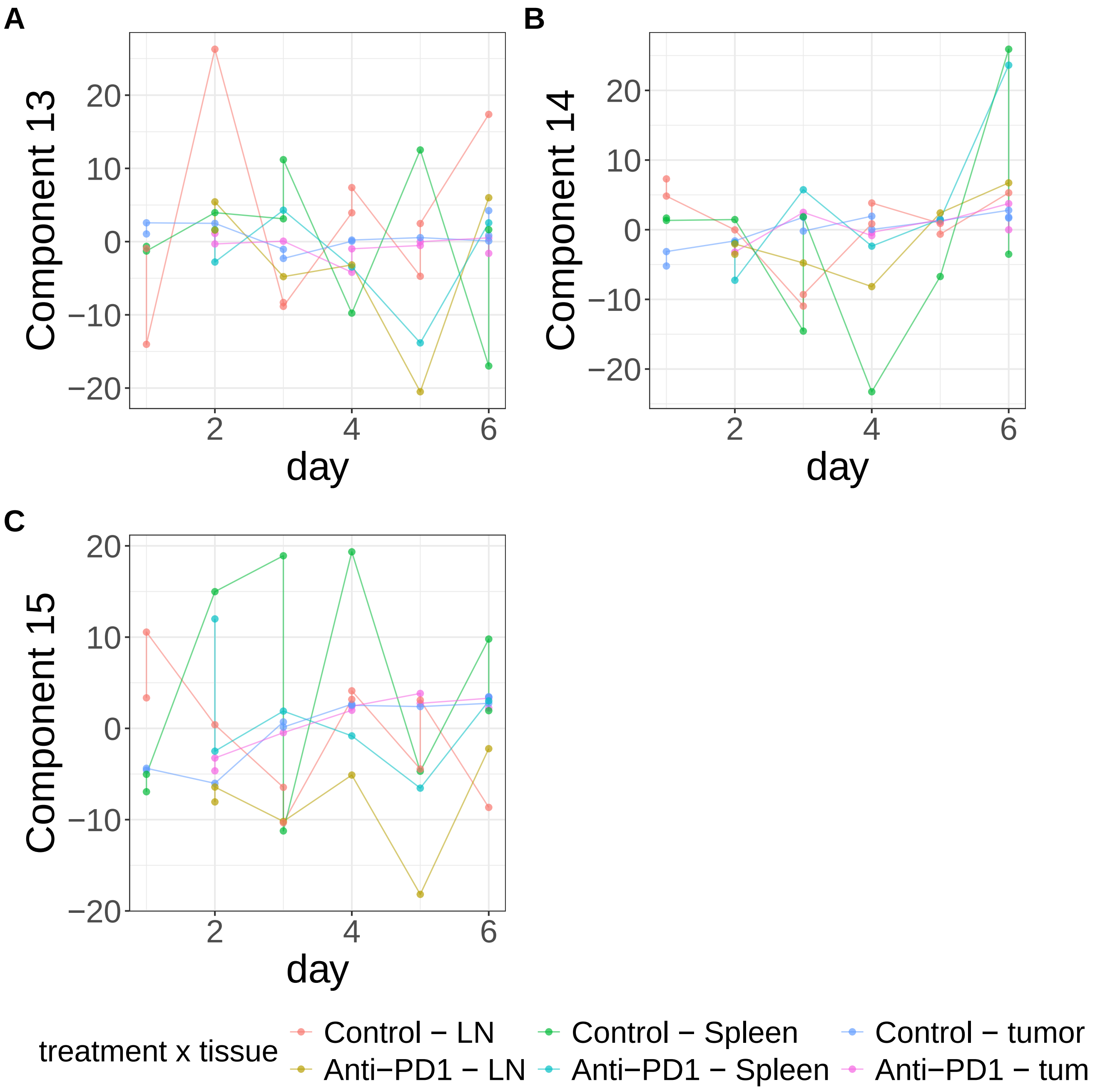


**Supplementary Figure 515.** Components for sPCA: keepX=1250. (A) Component 13. (B) Component 14. (C) Component 15.

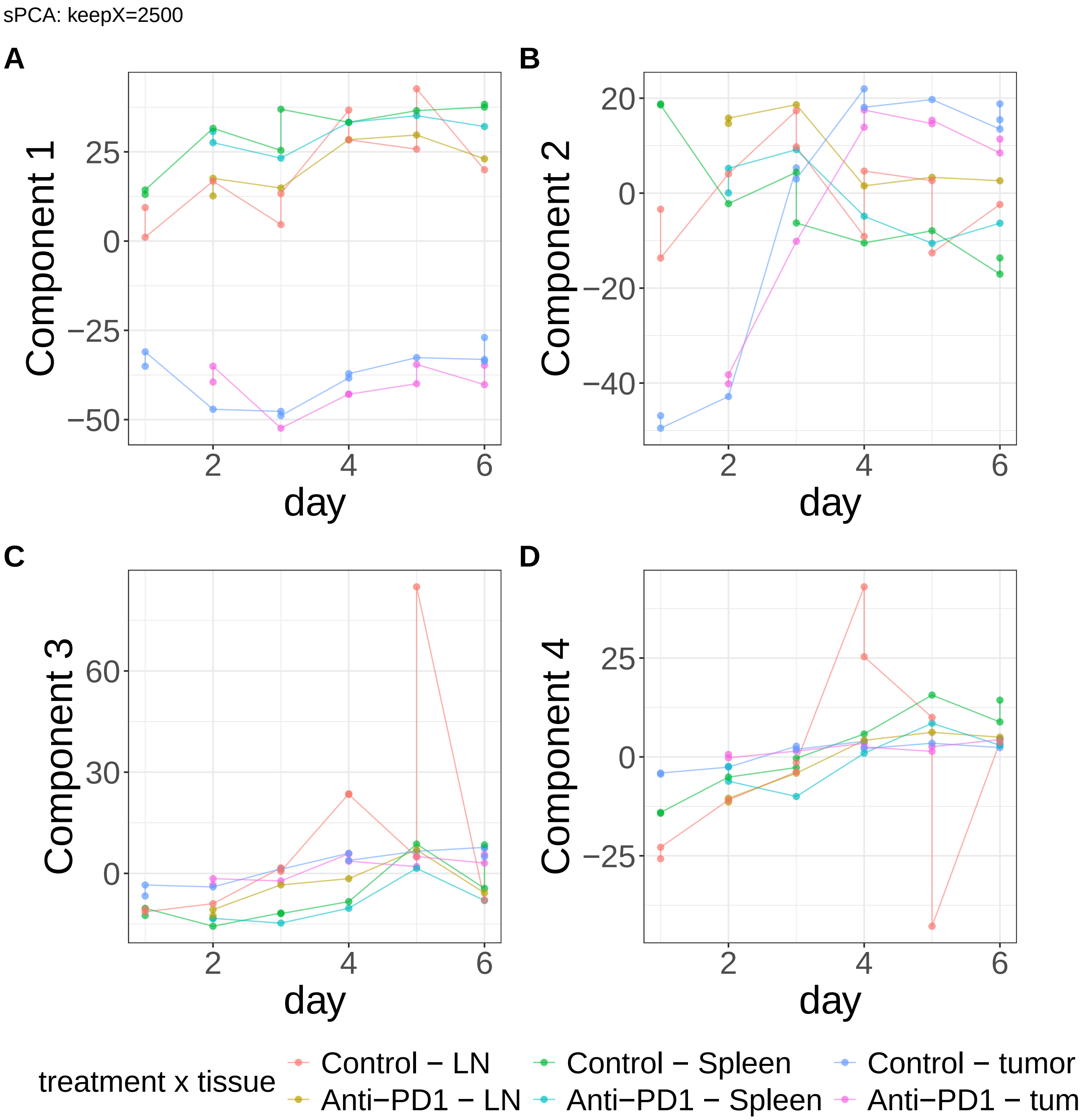


**Supplementary Figure 516.** Components for sPCA: keepX=2500. (A) Component 1. (B) Component 2. (C) Component 3. (D) Component 4.

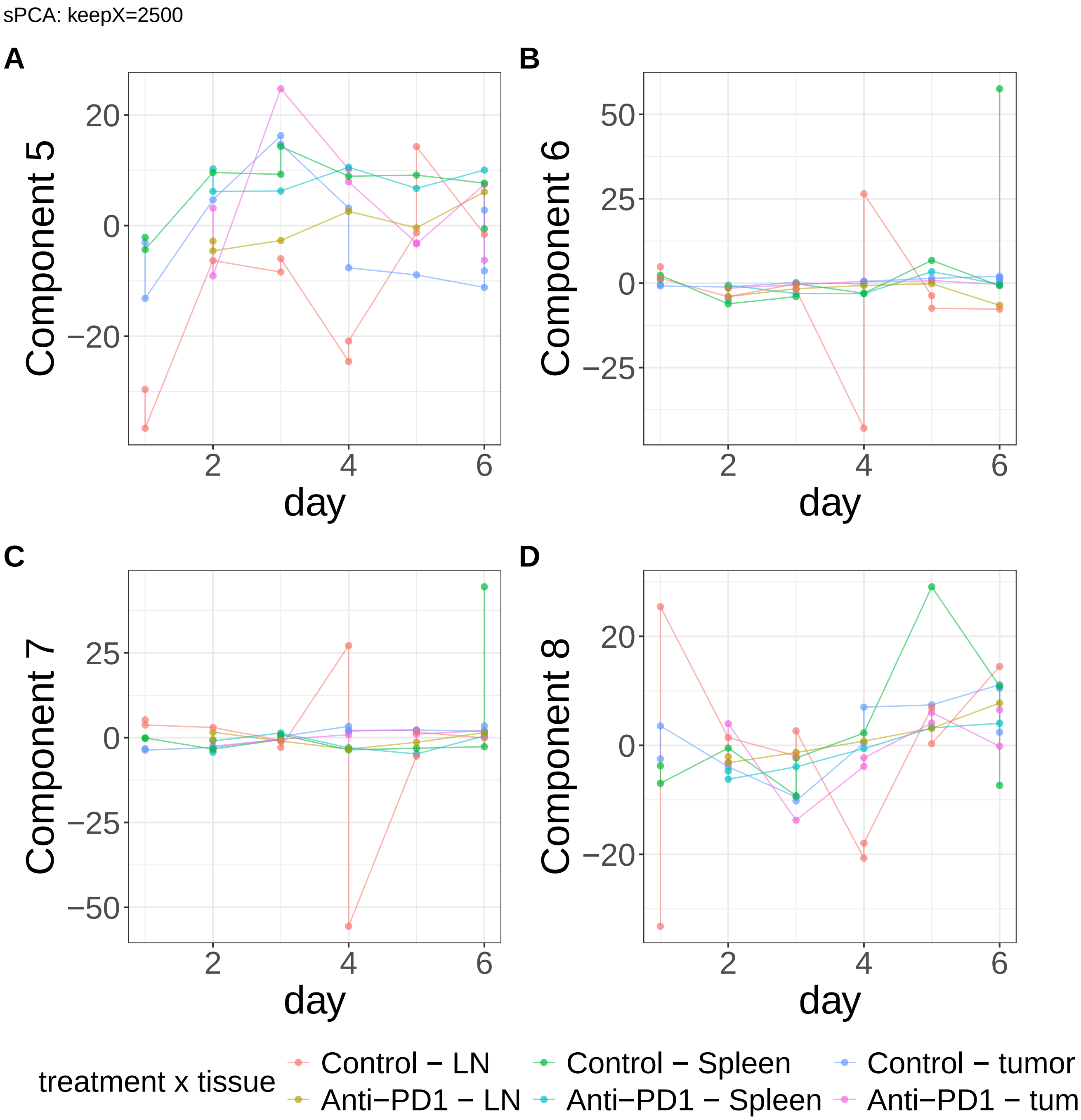


**Supplementary Figure 517.** Components for sPCA: keepX=2500. (A) Component 5. (B) Component 6. (C) Component 7. (D) Component 8.

sPCA: keepX=2500

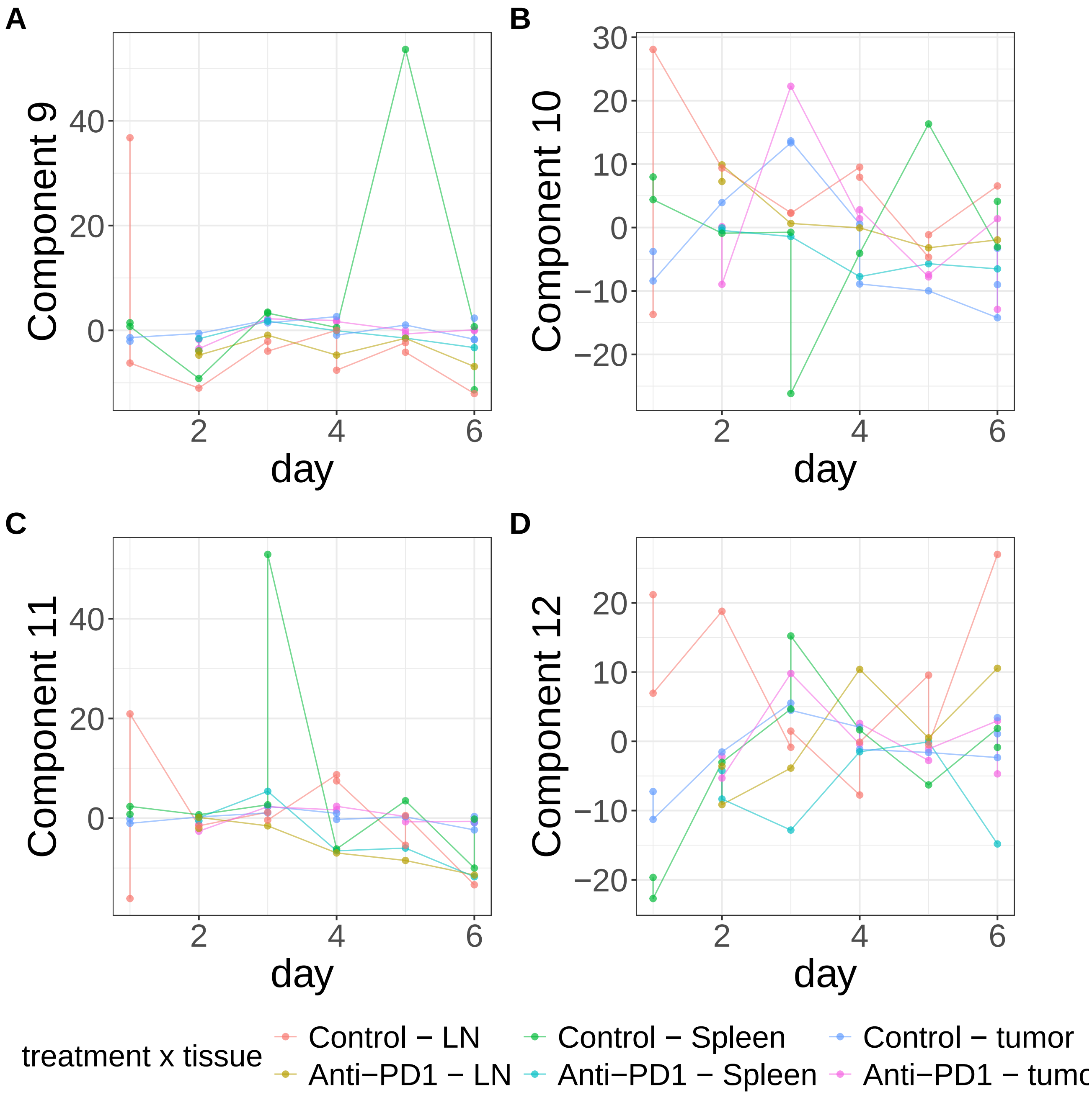


**Supplementary Figure 518.** Components for sPCA: keepX=2500. (A) Component 9. (B) Component 10. (C) Component 11. (D) Component 12.

sPCA: keepX=2500

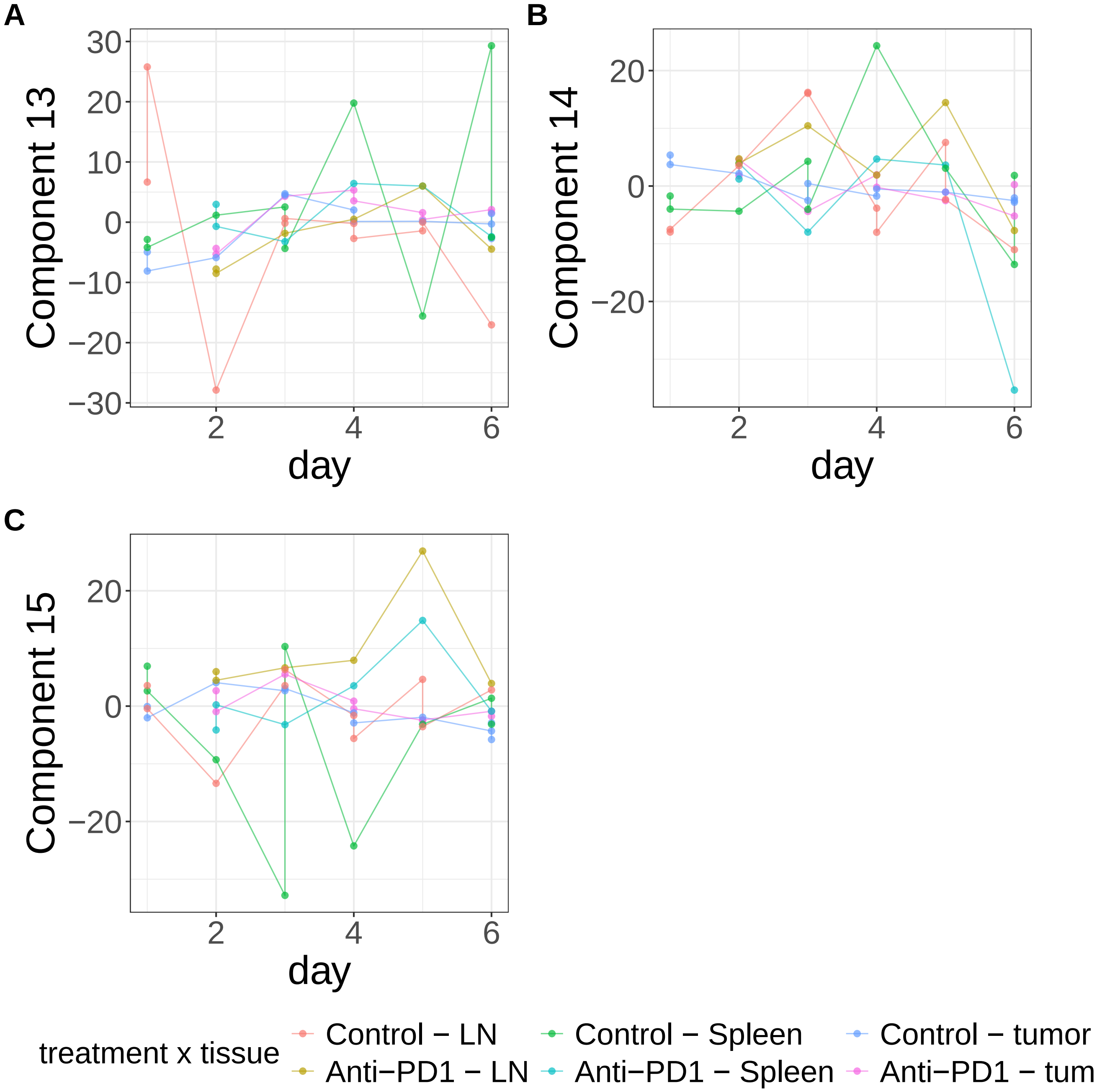


**Supplementary Figure 519.** Components for sPCA: keepX=2500. (A) Component 13. (B) Component 14. (C) Component 15.

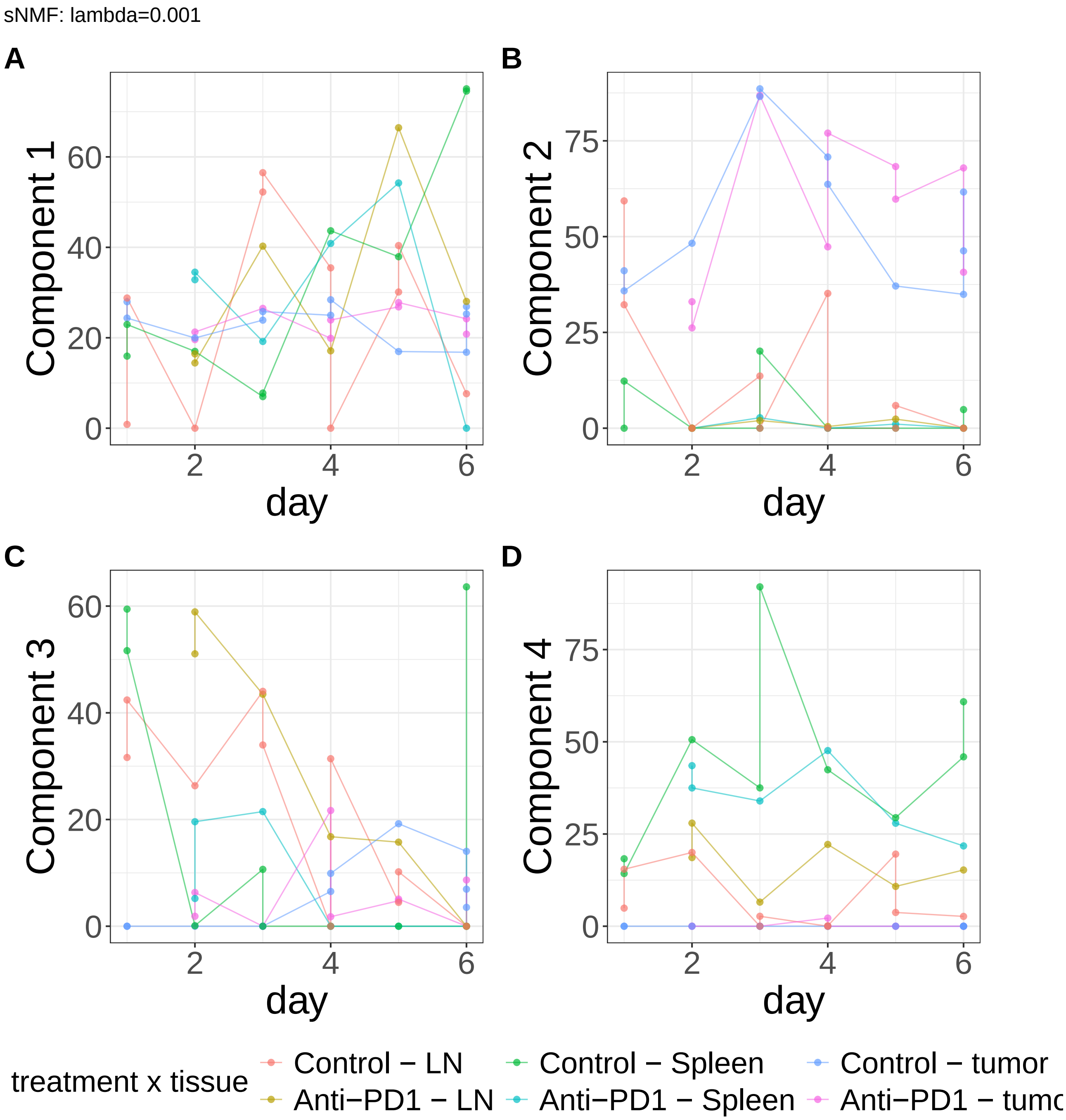


**Supplementary Figure 520.** Components for sNMF: lambda=0.001. (A) Component 1. (B) Component 2. (C) Component 3. (D) Component 4.

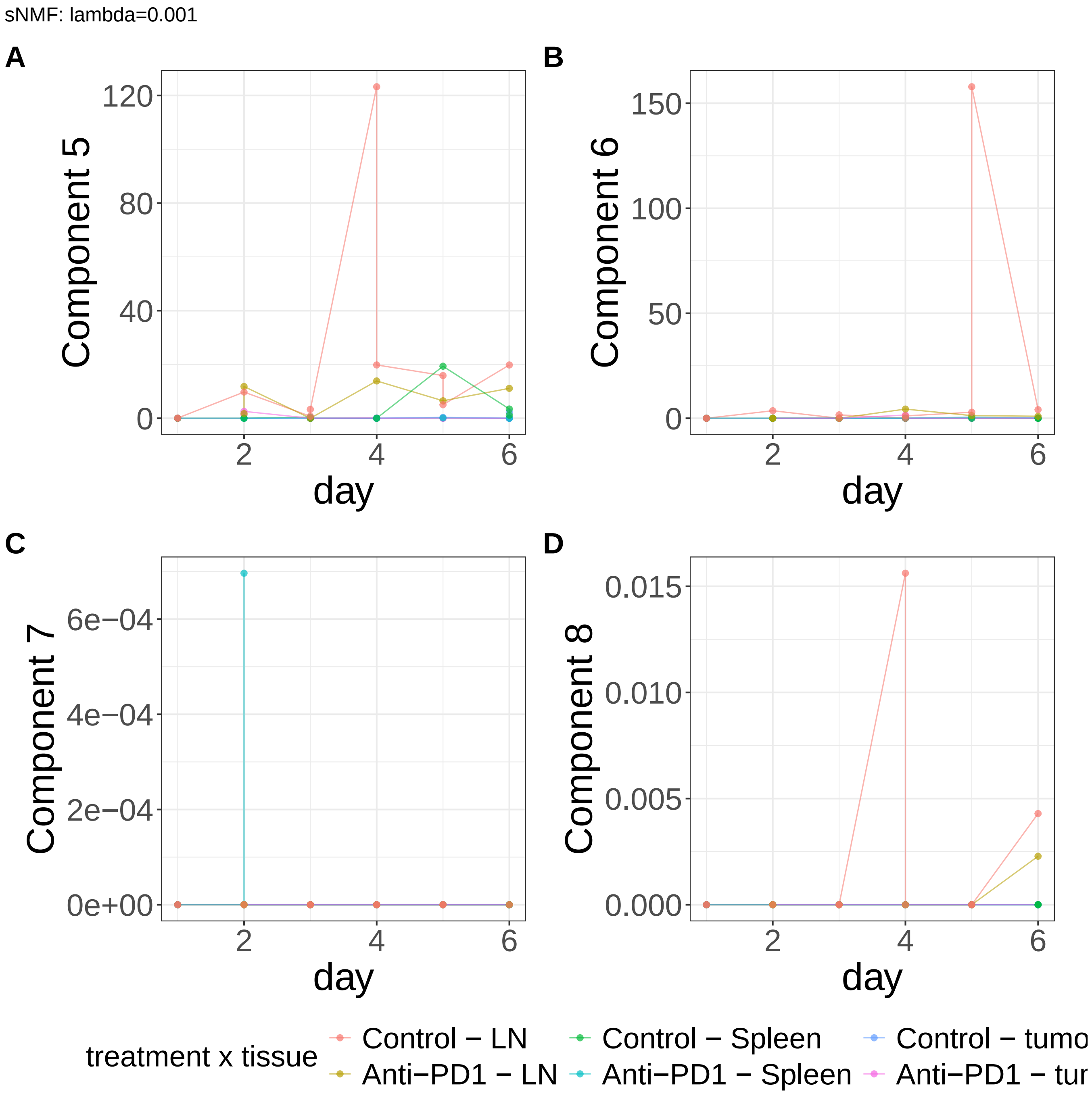


**Supplementary Figure 521.** Components for sNMF: lambda=0.001. (A) Component 5. (B) Component 6. (C) Component 7. (D) Component 8.

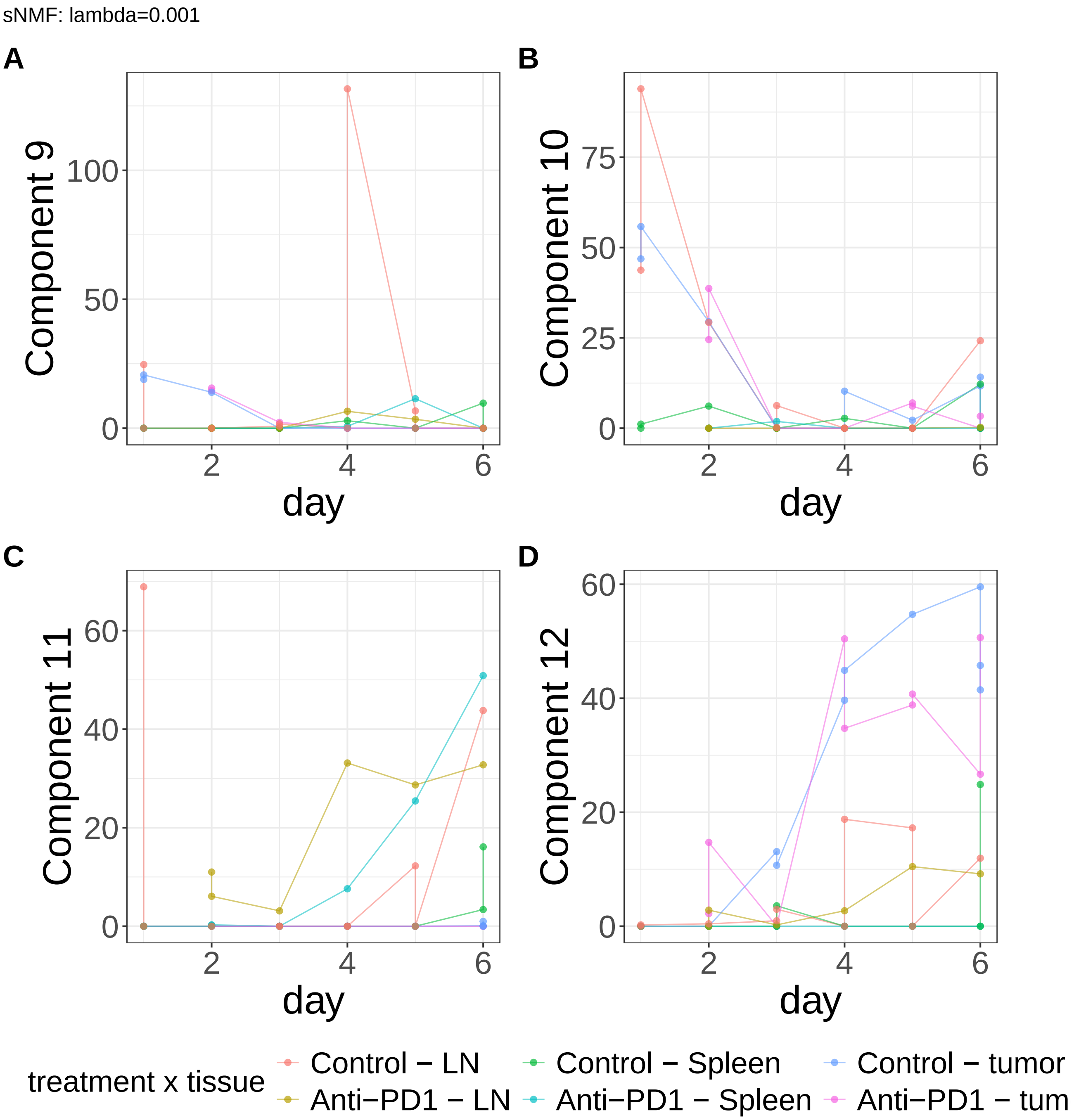


**Supplementary Figure 522.** Components for sNMF: lambda=0.001. (A) Component 9. (B) Component 10. (C) Component 11. (D) Component 12.

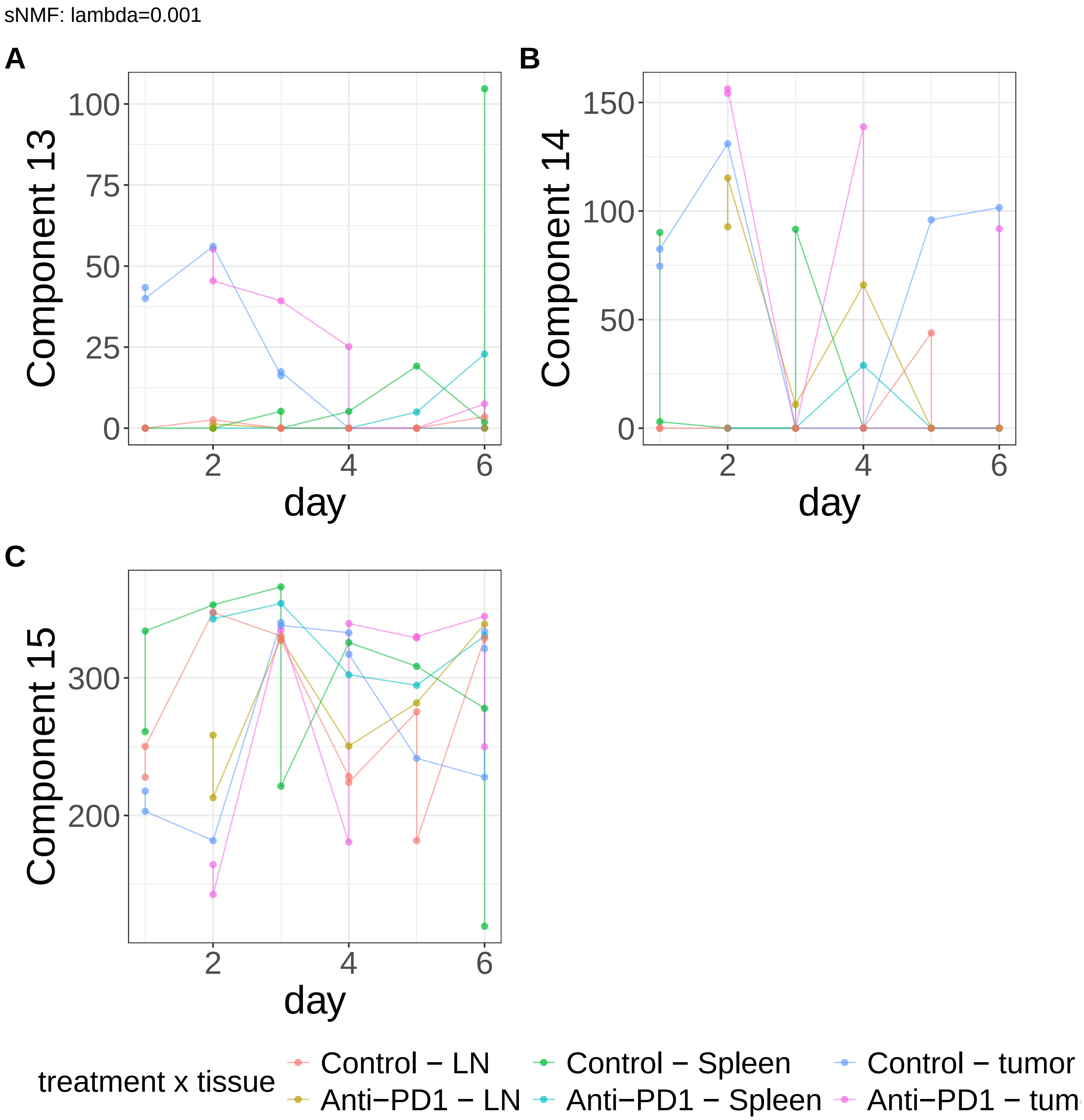


**Supplementary Figure 523.** Components for sNMF: lambda=0.001. (A) Component 13. (B) Component 14. (C) Component 15.

sNMF: lambda=0.01

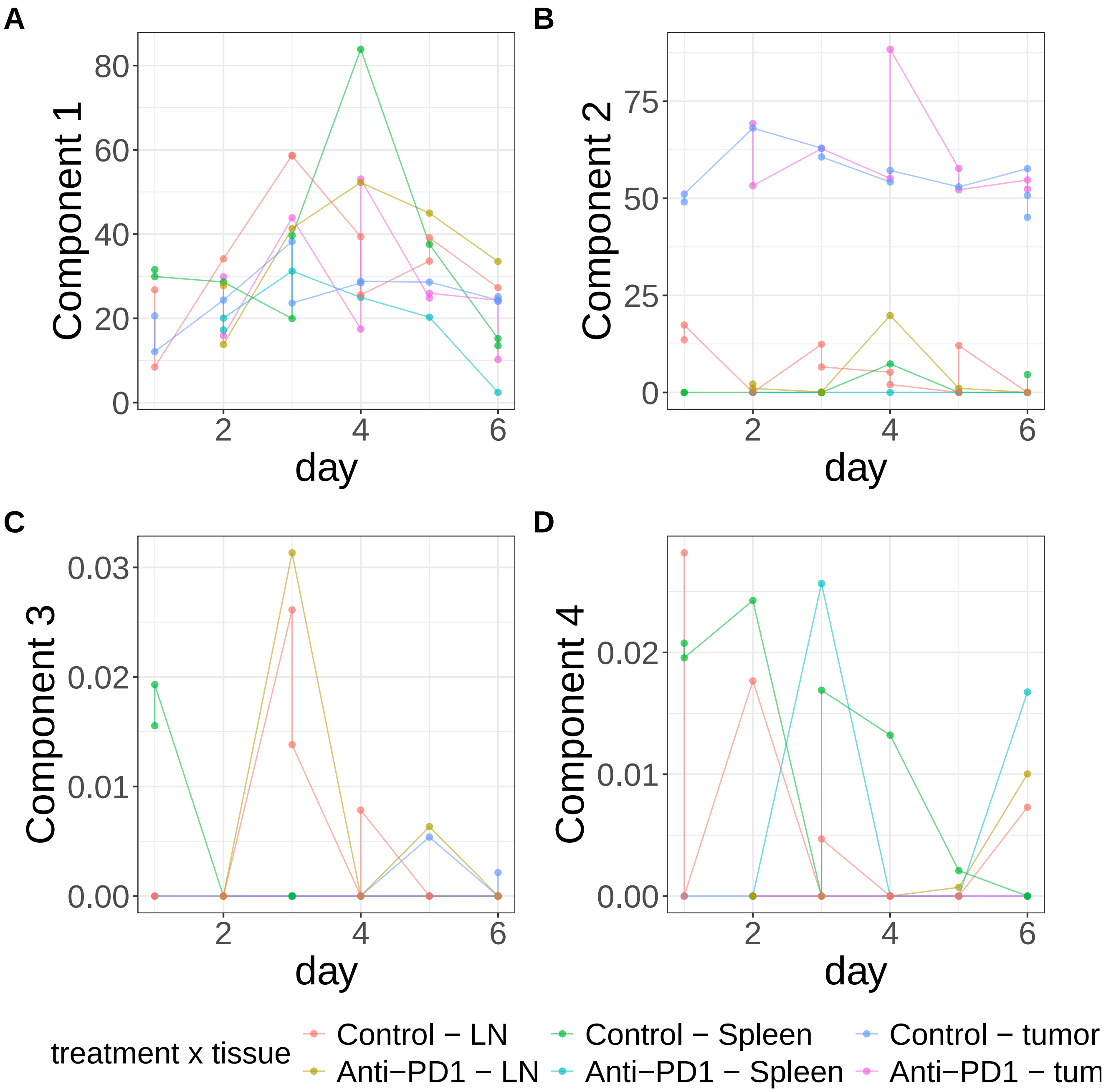


**Supplementary Figure 524.** Components for sNMF: lambda=0.01. (A) Component 1. (B) Component 2. (C) Component 3. (D) Component 4.

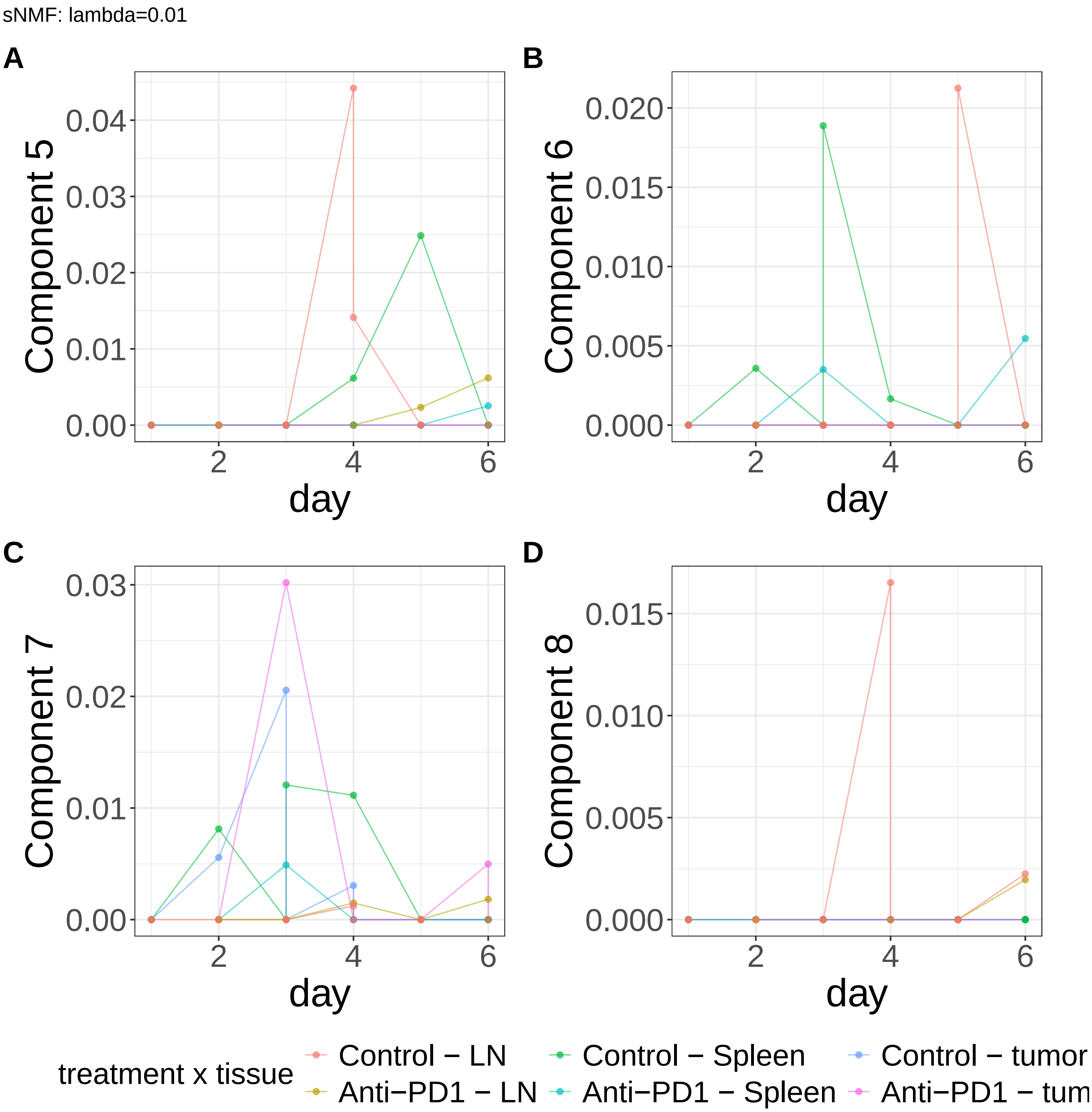


**Supplementary Figure 525.** Components for sNMF: lambda=0.01. (A) Component 5. (B) Component 6. (C) Component 7. (D) Component 8.

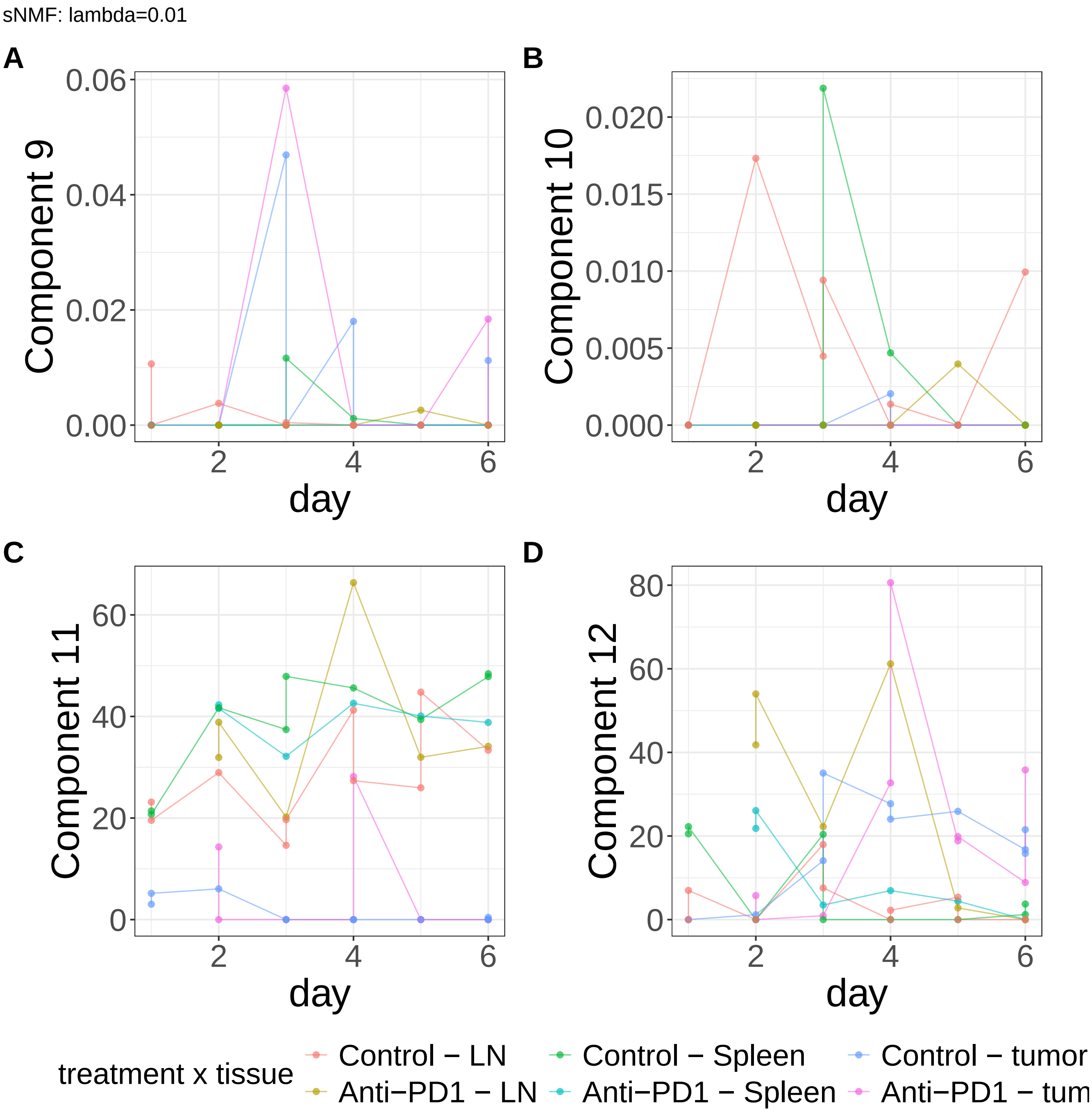


**Supplementary Figure 526.** Components for sNMF: lambda=0.01. (A) Component 9. (B) Component 10. (C) Component 11. (D) Component 12.

sNMF: lambda=0.01

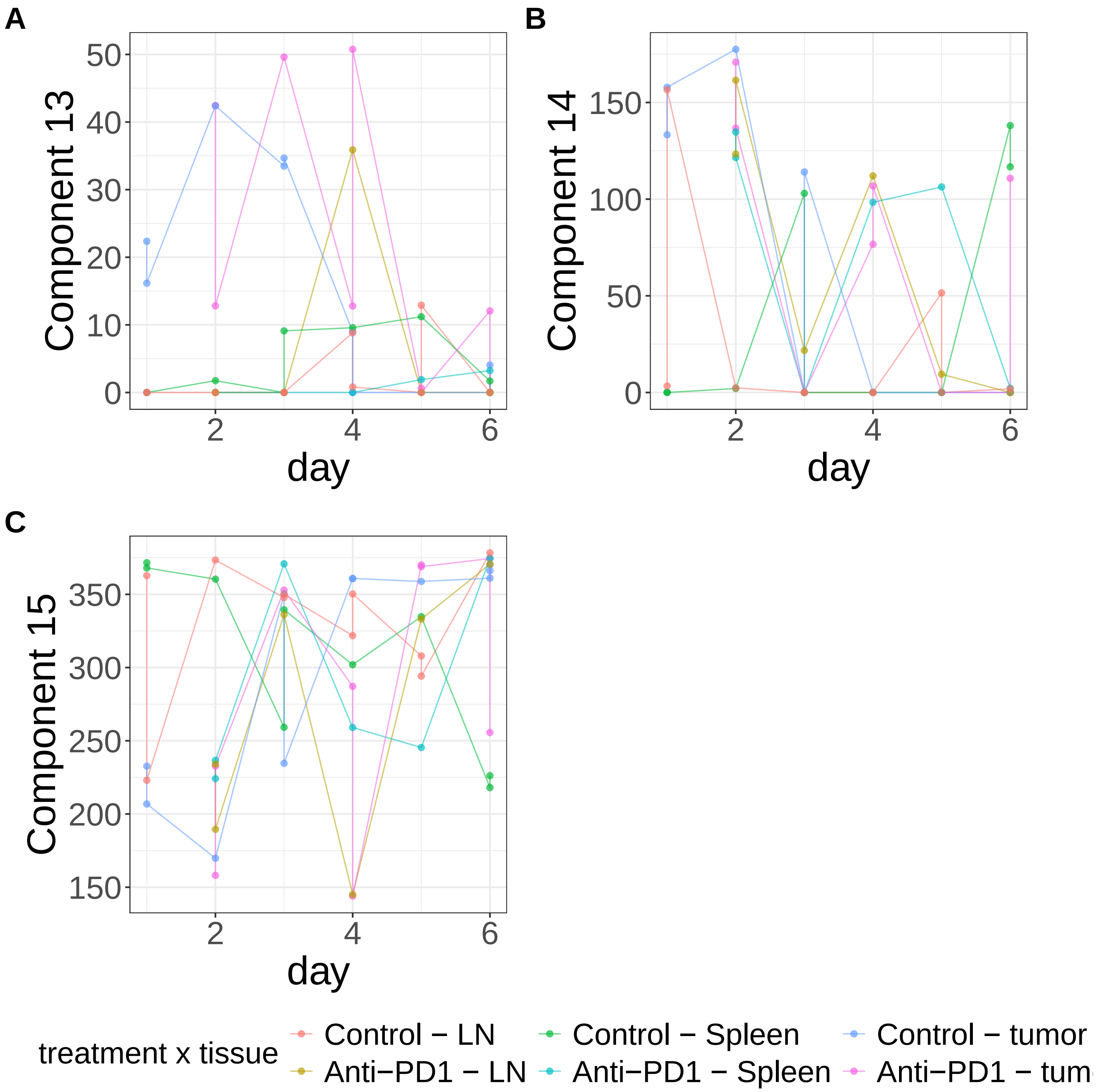


**Supplementary Figure 527.** Components for sNMF: lambda=0.01. (A) Component 13. (B) Component 14. (C) Component 15.

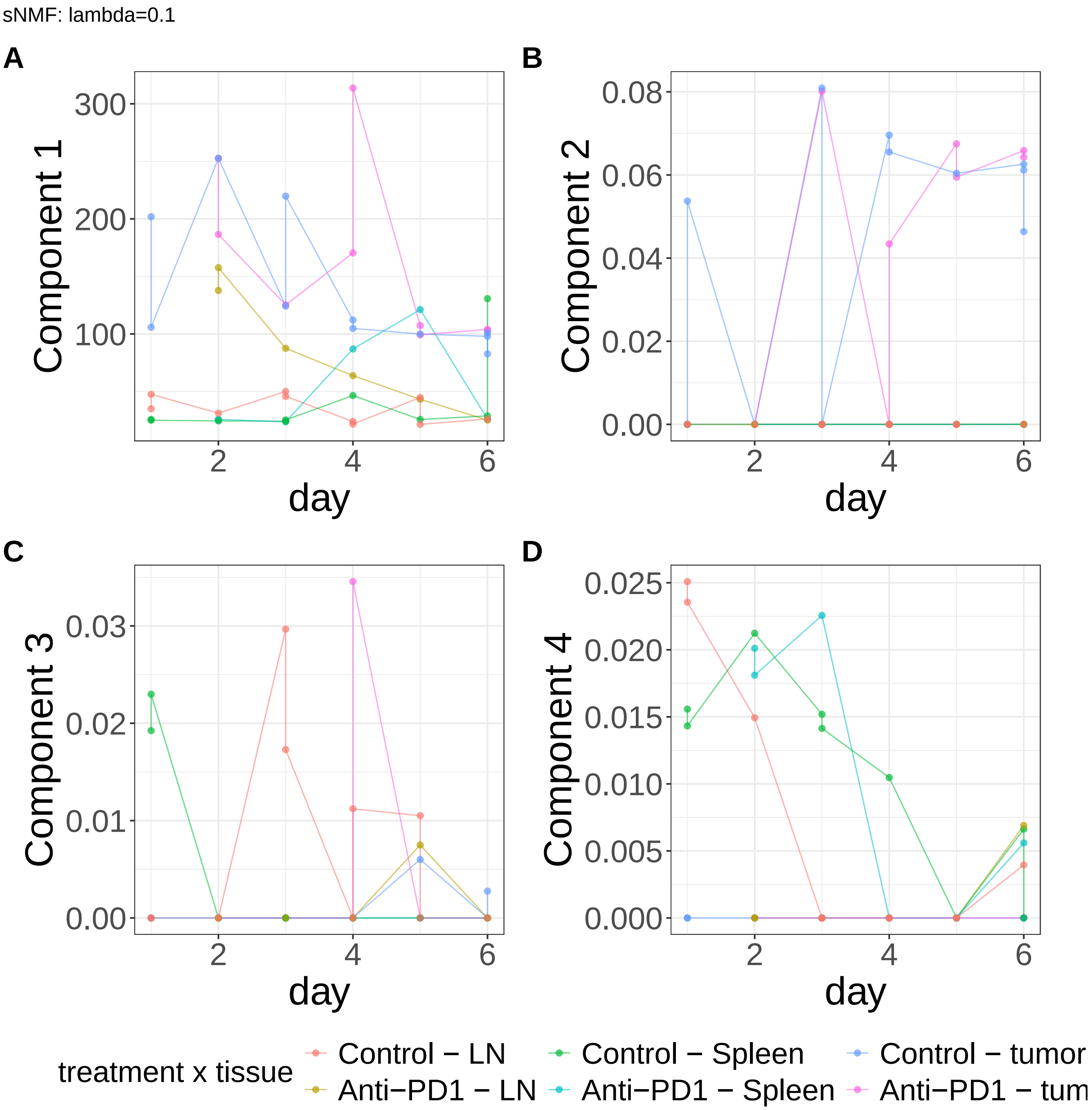


**Supplementary Figure 528.** Components for sNMF: lambda=0.1. (A) Component 1. (B) Component 2. (C) Component 3. (D) Component 4.

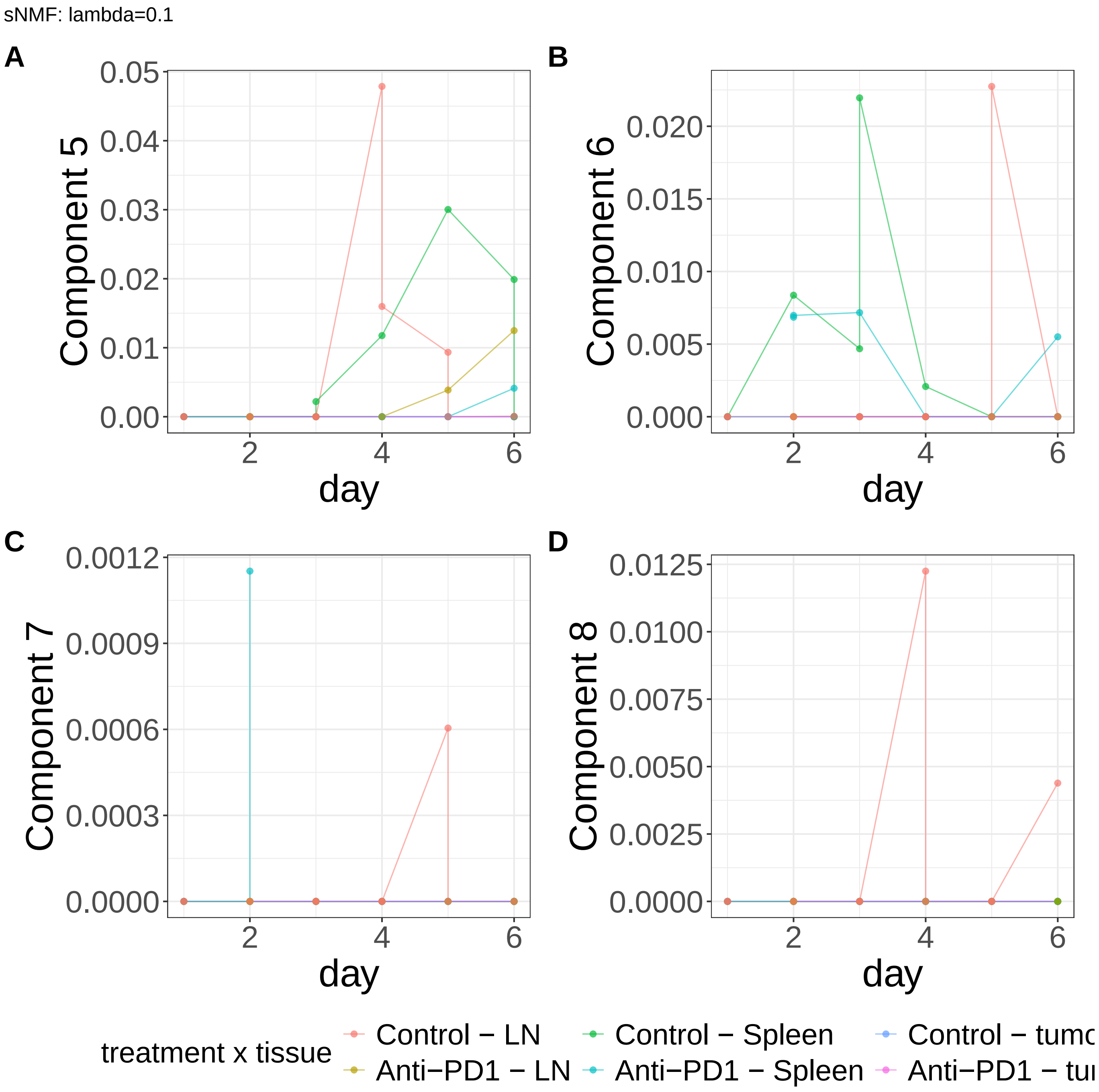


**Supplementary Figure 529.** Components for sNMF: lambda=0.1. (A) Component 5. (B) Component 6. (C) Component 7. (D) Component 8.

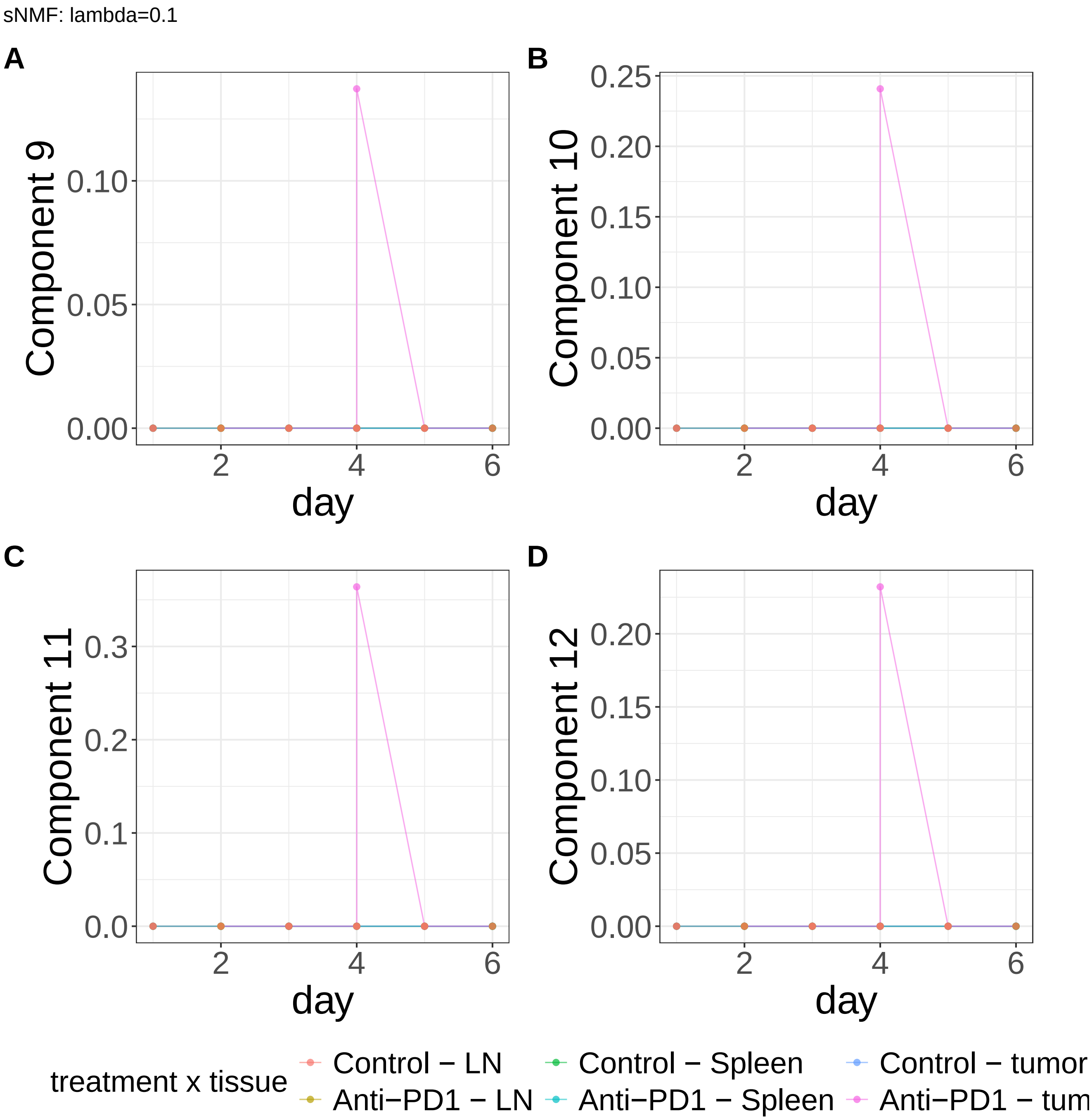


**Supplementary Figure 530.** Components for sNMF: lambda=0.1. (A) Component 9. (B) Component 10. (C) Component 11. (D) Component 12.

sNMF: lambda=0.1

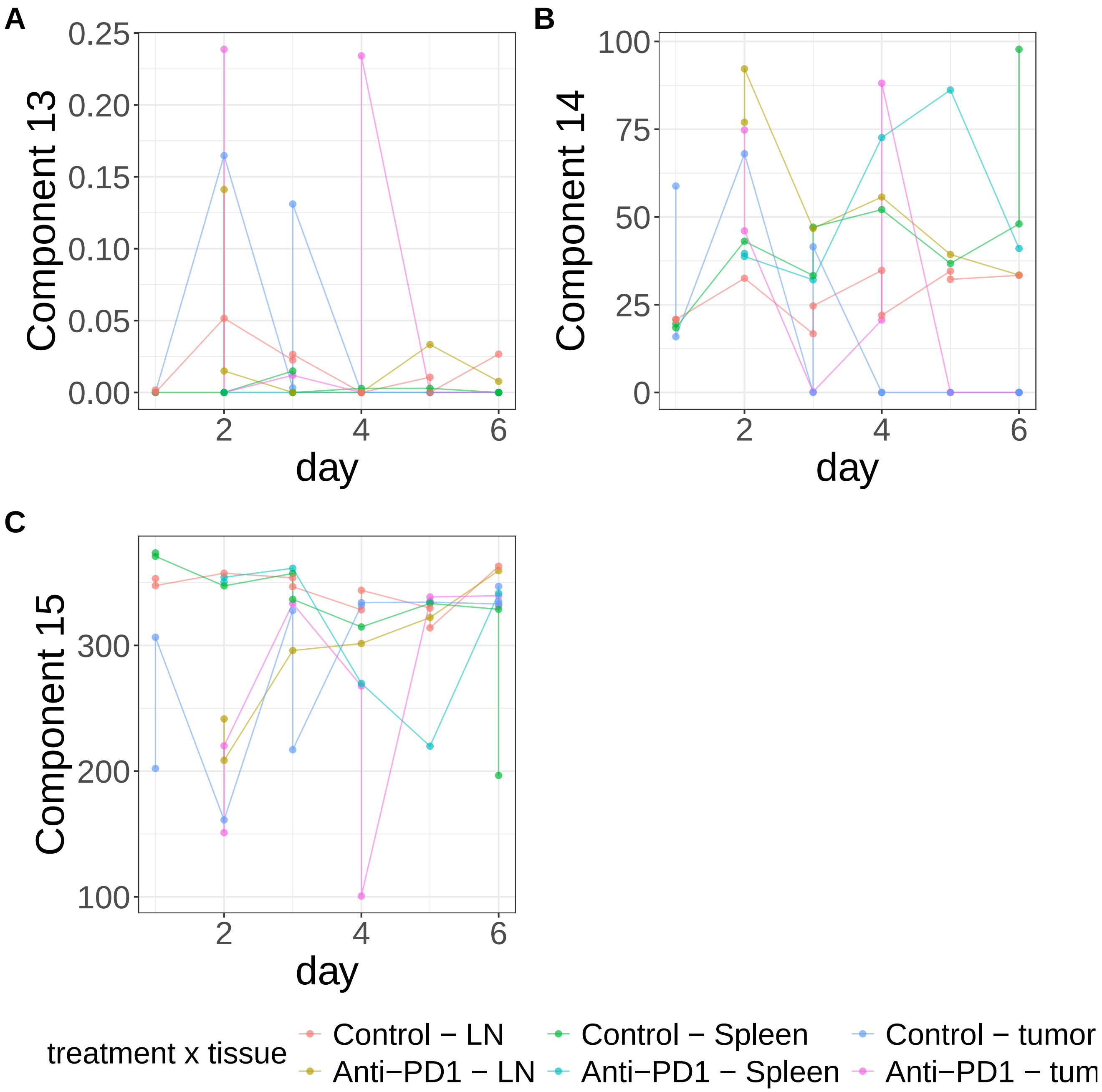


**Supplementary Figure 531.** Components for sNMF: lambda=0.1. (A) Component 13. (B) Component 14. (C) Component 15.

MEFISTO: duplicate views

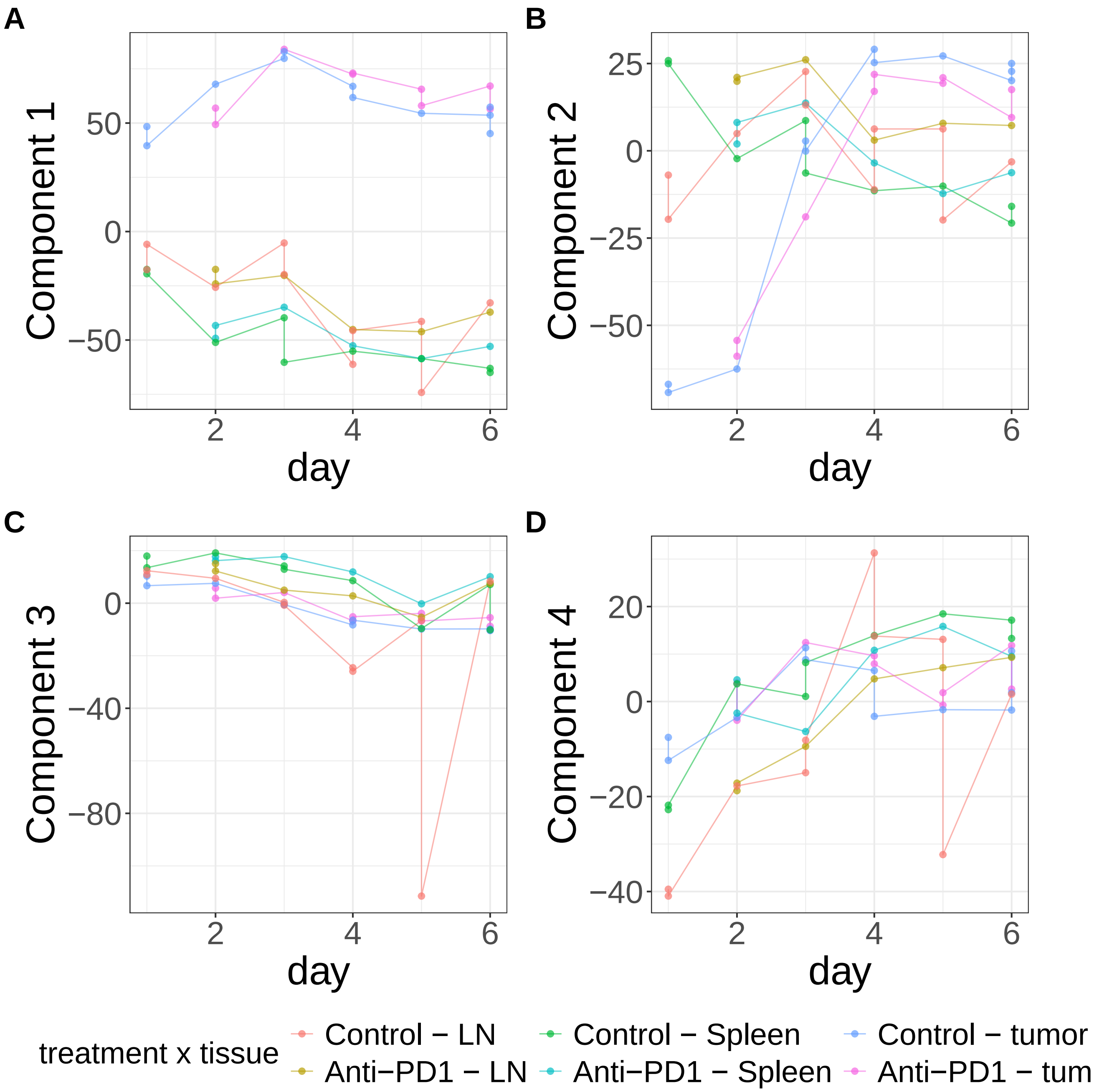


**Supplementary Figure 532.** Components for MEFISTO: duplicate views. (A) Component 1. (B) Component 2. (C) Component 3. (D) Component 4.

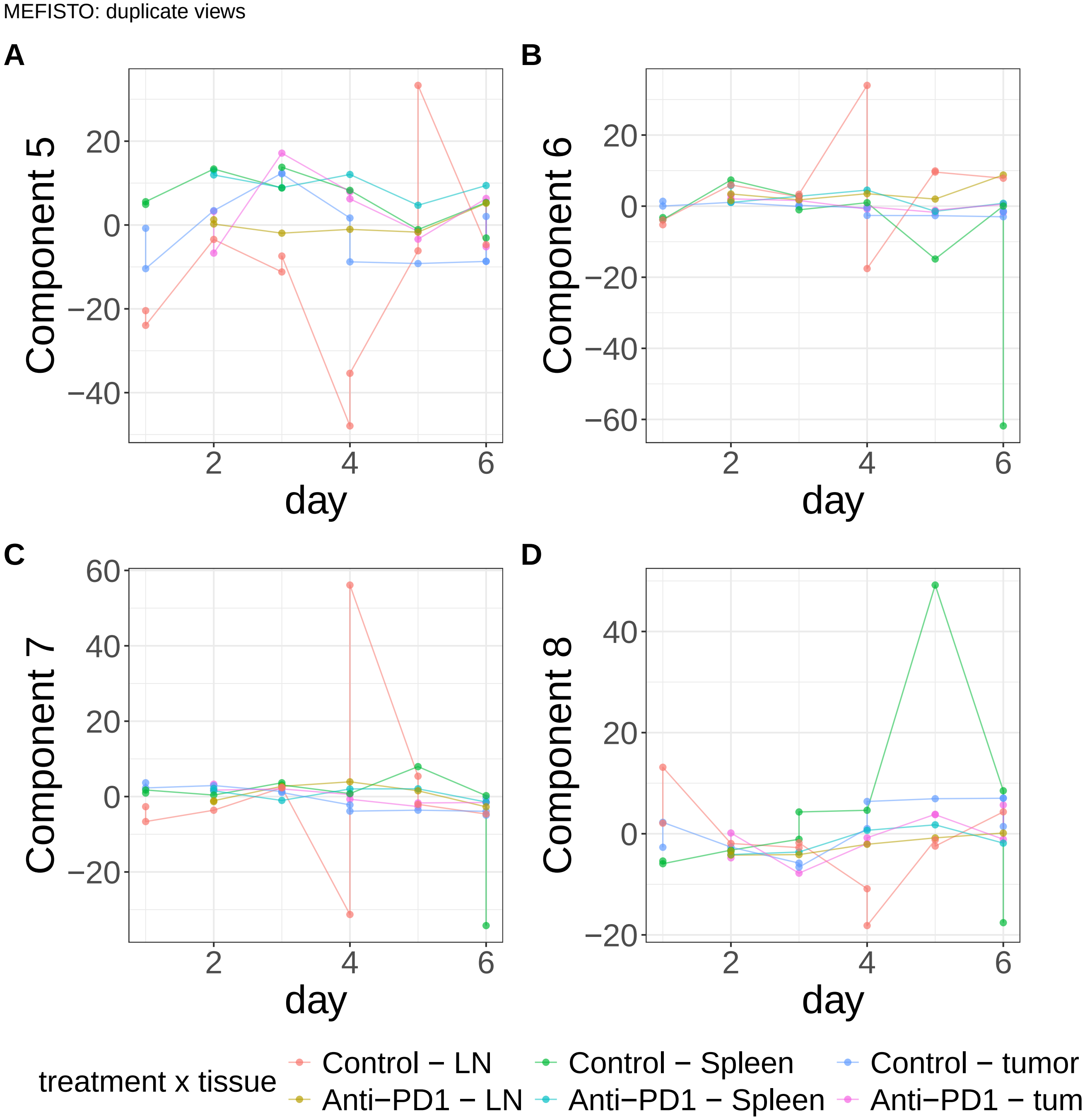


**Supplementary Figure 533.** Components for MEFISTO: duplicate views. (A) Component 5. (B) Component 6. (C) Component 7. (D) Component 8.

MEFISTO: duplicate views

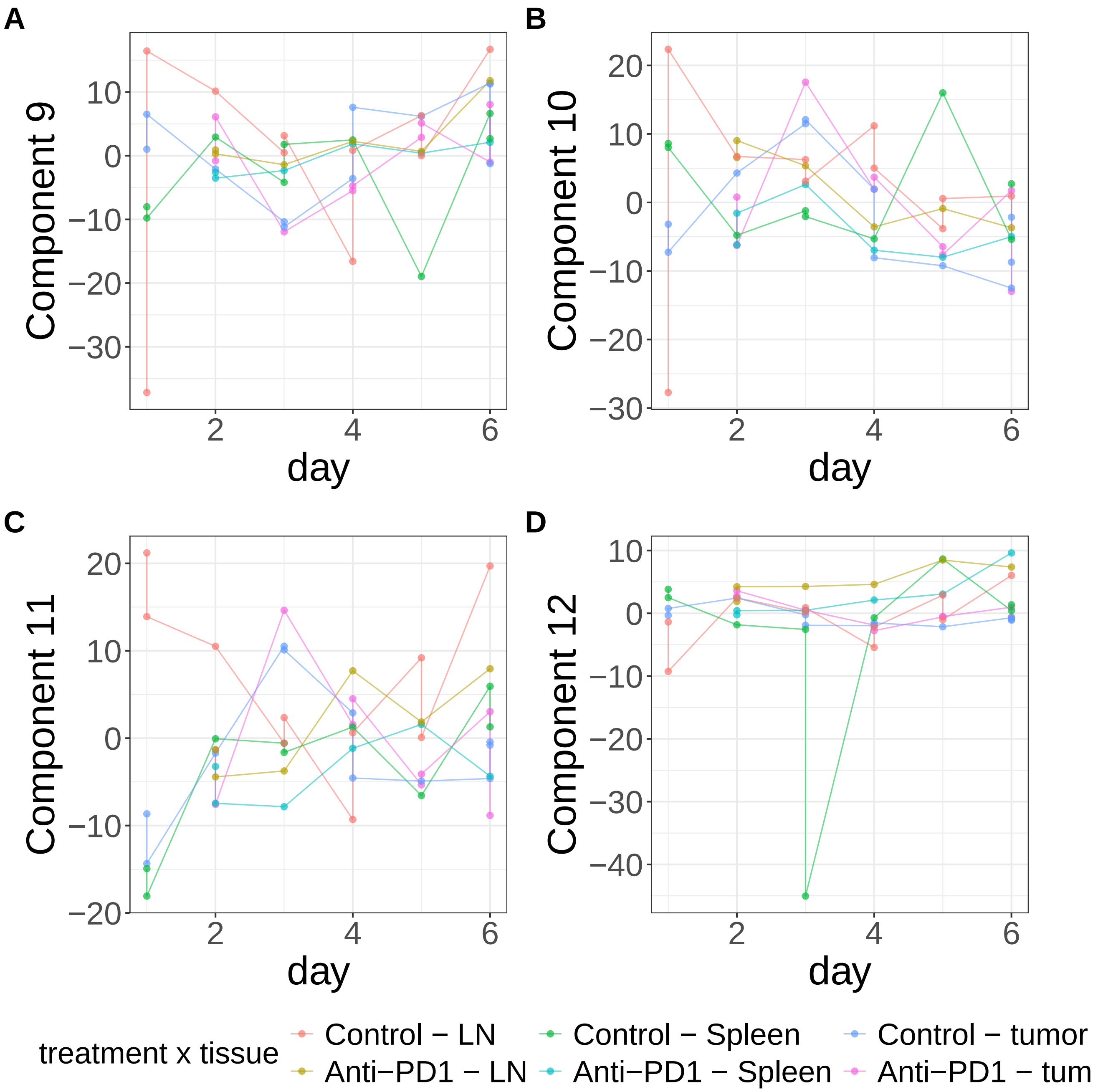


**Supplementary Figure 534.** Components for MEFISTO: duplicate views. (A) Component 9. (B) Component 10. (C) Component 11. (D) Component 12.

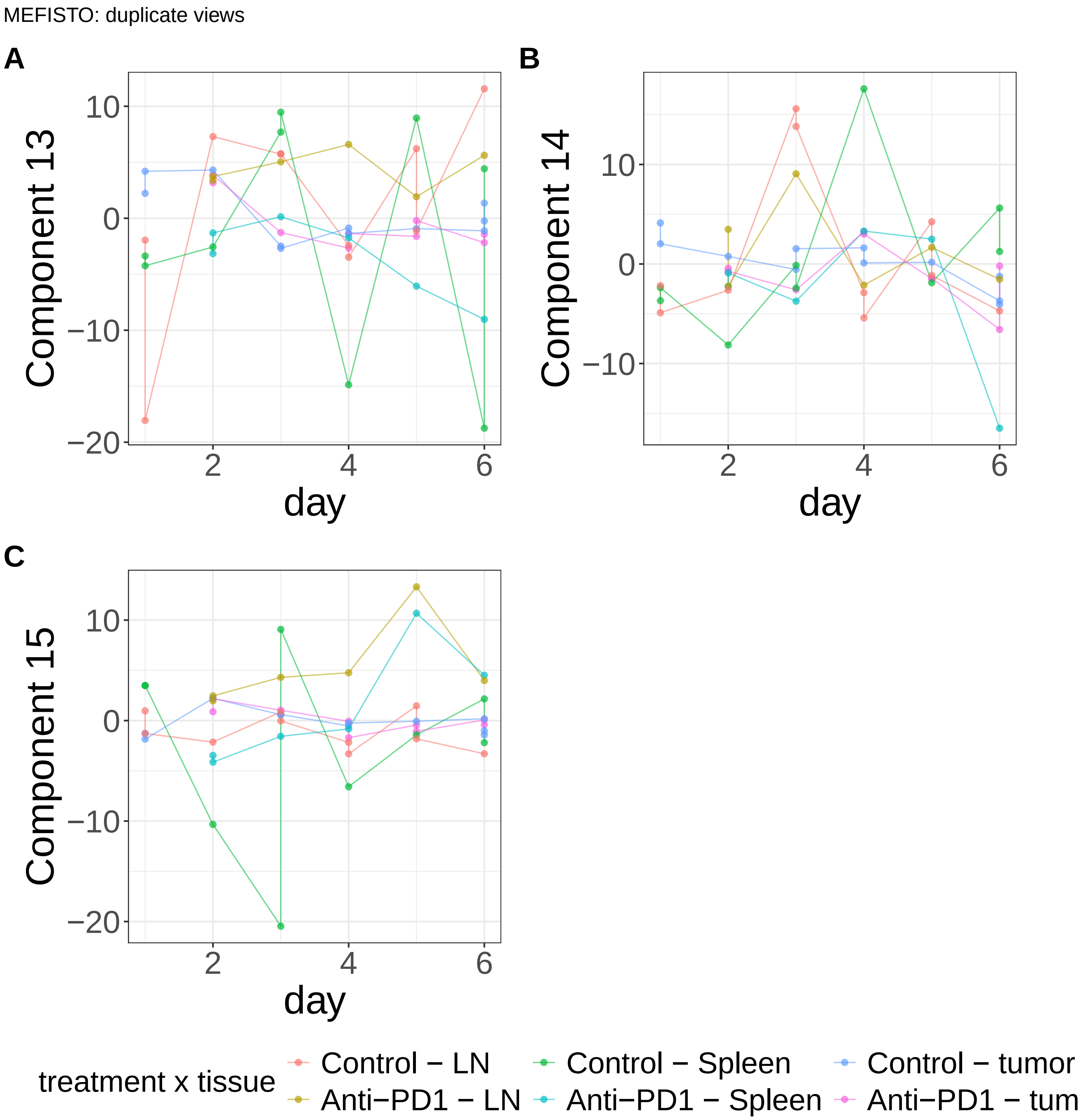


**Supplementary Figure 535.** Components for MEFISTO: duplicate views. (A) Component 13. (B) Component 14. (C) Component 15.
