## Supplementary Note 8 : OT1 Longer Time Course for "A statistical framework for system-level synthesis"

### System-level synthesis exposes an experimental anomaly

We analyzed data from longitudinal profiling of tumor-antigen-specific OT1 CD8 T cells following adoptive transfer into B16-OVA-bearing mice receiving anti-PD1 or control treatment[1]. Cells were collected from tumors for up to 15 days following transfer.

We modeled the system-level processes as smooth functions of time, treatment, and their interaction (**Methods**, **Supplementary Figs. 426-477**). SSANOVA-SVD reconstructed three processes with distinct treatment-dependent temporal trajectories (**Supplementary Figs. 426A, 427-428**). The first showed a striking progressive increase in control animals that was absent following anti-PD1 treatment. The second showed the converse pattern, with a progressive change following anti-PD1 treatment while the control group remained relatively stable over time. The third exhibited comparatively smaller effects. Classical matrix factorization methods such as PCA similarly prioritized the dominant treatment-associated variation, but subsequent components were less readily interpretable (**Supplementary Figs. 441–477**).

Conditional inference using held-out measurements revealed an unexpected explanation for the dominant process. The associated features included the myeloid genes *Arg1*, *Ccl12*, *Msr1*, *Lyz2*, *C1qb*, *C1qc*, *Dab2*, and *Fcgr4*, despite the experiment targeting OT1 T cells (**Supplementary Figs. 426B–C, 429–434**). Examination of the underlying single-cell measurements confirmed substantial populations of myeloid-like cells in several late control samples, and the samples from the eight mice identified through this analysis coincided with mice independently excluded by the original investigators. Removing these mice eliminated all control observations at days 12 and 15, leaving treatment-by-time combinations unobserved and precluding reconstruction of the complete longitudinal design. Thus, the dominant apparent treatment-associated process reflected an experimental compositional irregularity rather than the intended OT1 T-cell response.

In contrast, the second process was independently associated with *Prdm1*, *Cd244*, *Cx3cr1*, *Entpd1*, *Gzmk*, and *Zfp683*, consistent with a progressive anti-PD1-associated transition toward differentiated effector T-cell states[2–4]. This behavior was concordant with the original single-cell analysis, which identified PD-1-blockade-associated changes in the differentiation of tumor-specific activated precursor cells. Notably, *Cx3cr1* was also independently identified in the dominant ICB-response process in the bilateral mouse model, providing a potential molecular connection between the responses reconstructed in the two studies. The third process had comparatively weak feature-level support and retained associations with myeloid genes, preventing similarly clear interpretation.

Together, these results illustrate that system-level synthesis can expose experimental artifacts underlying dominant variation that might otherwise be interpreted as a treatment-associated response.

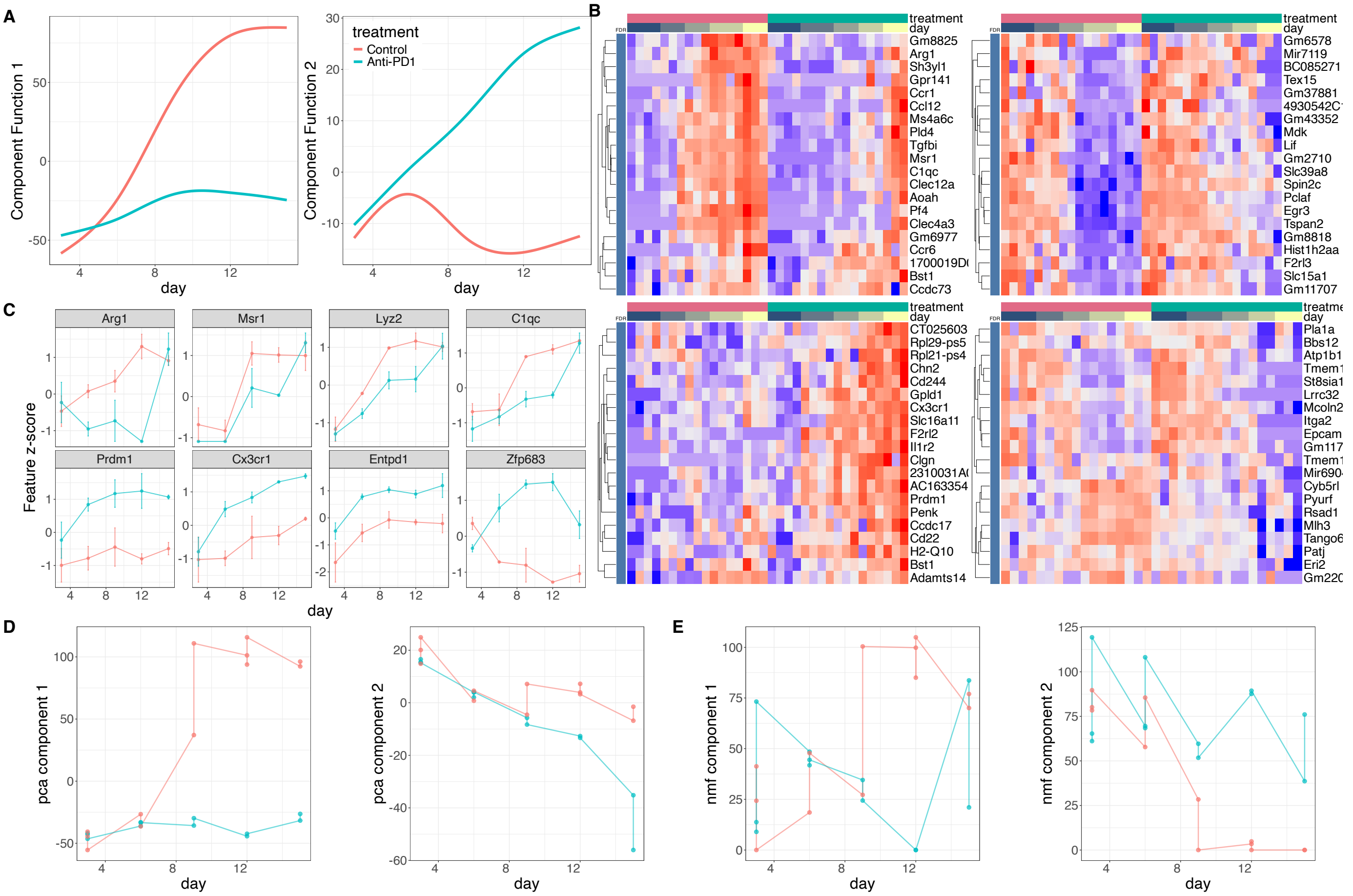


**Supplementary Figure 426: System-level synthesis of tumor tissue derived OT1 CD8 T-cells during immune checkpoint blockade (time course 2).** A. Reconstructed component functions for time course two of the OT1 CD8 T-cells data, by SSANOVA-SVD. B. Conditional inference in unseen test data. Heatmaps of z-scores of top 20 positively (top row) and negatively (bottom row) associated genes for components 1(left), 7 (center) and 9 (right), excluding PBANKA genes without known gene symbols. Blue annotation indicates the gene's $FDR \leq .1$ for the component. C. Standardized expression profiles of associated genes across time for component 1 (top row), and component 2 (bottom row). Points and error bars summarize expression at each sampled time point across tissues. D. D. The first two components obtained using principal component analysis (PCA). E. The first two components obtained using non-negative matrix factorization (NMF).

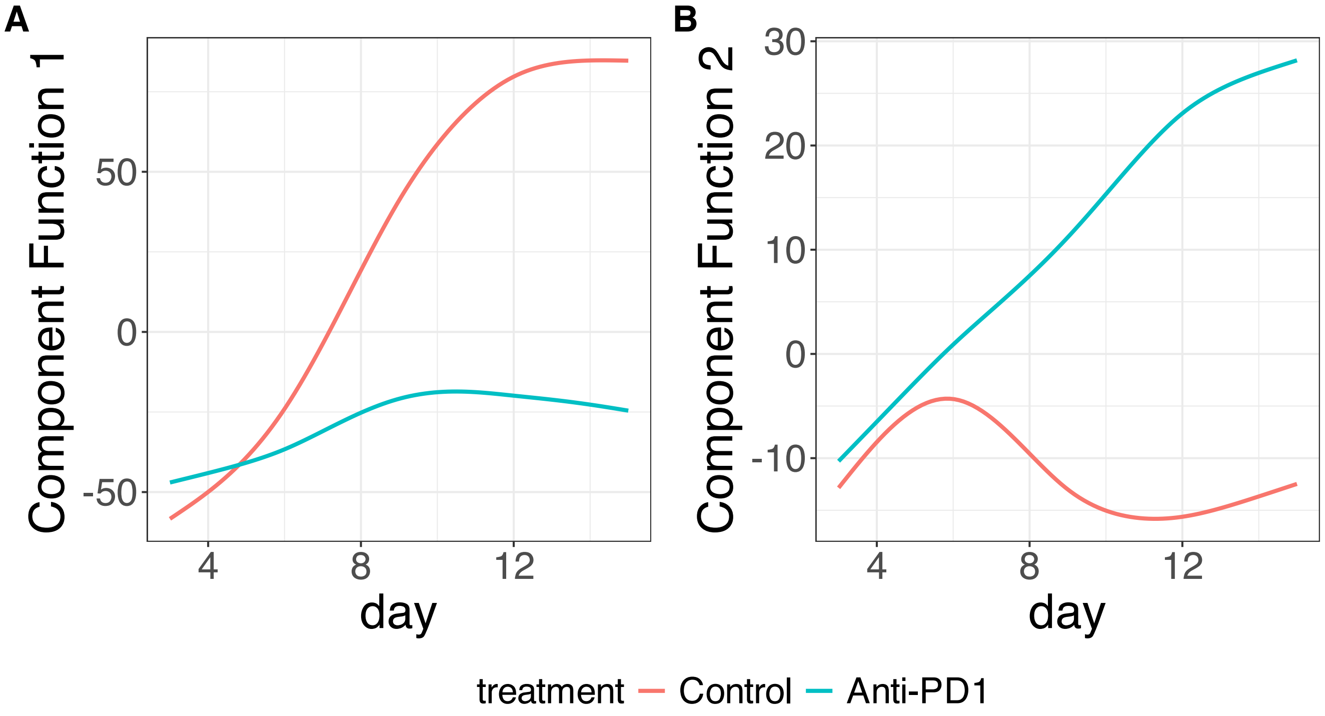


**Supplementary Figure 427.** SSANOVA-SVD reconstructed component functions.(A) Component 1. (B) Component 2.

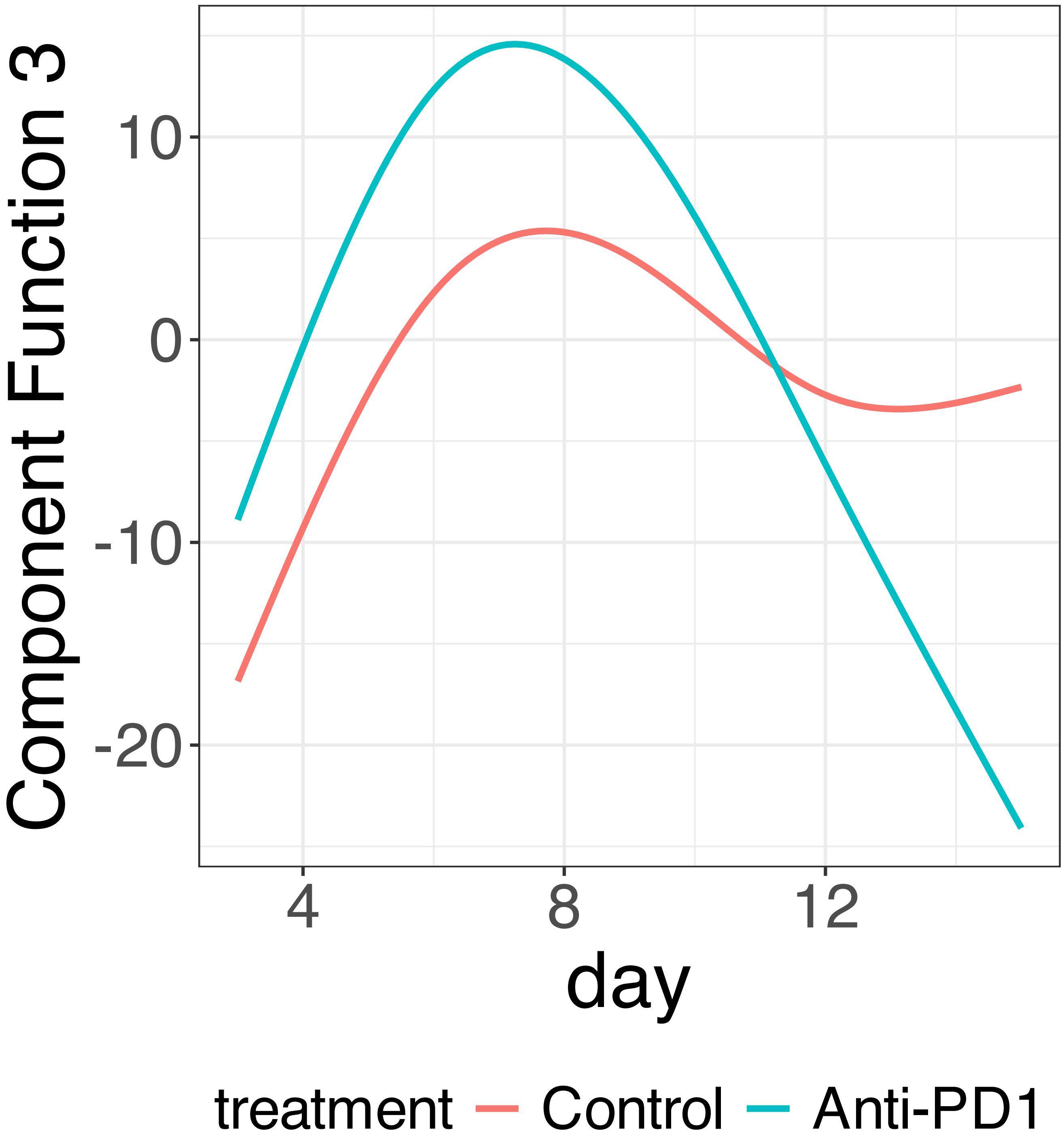


**Supplementary Figure 428.** SSANOVA-SVD reconstructed component functions.Component 3.

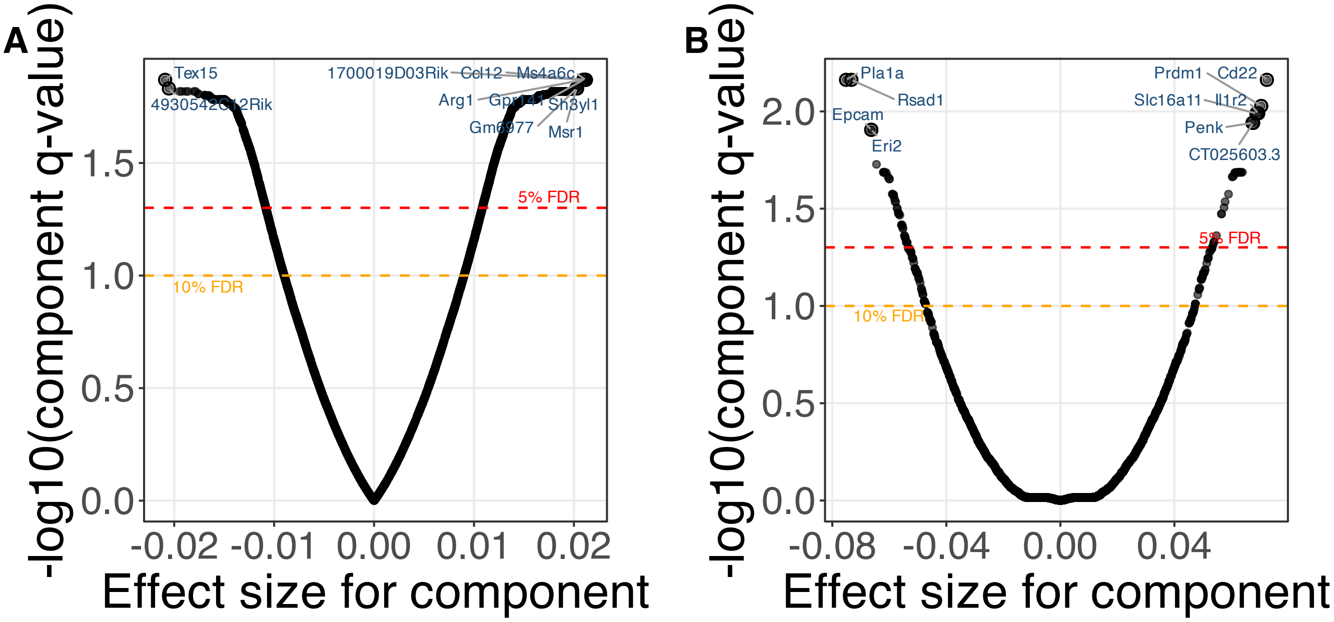


**Supplementary Figure 429.** SSANOVA-SVD based conditional inference for feature participation. (A) Component 1. (B) Component 2.

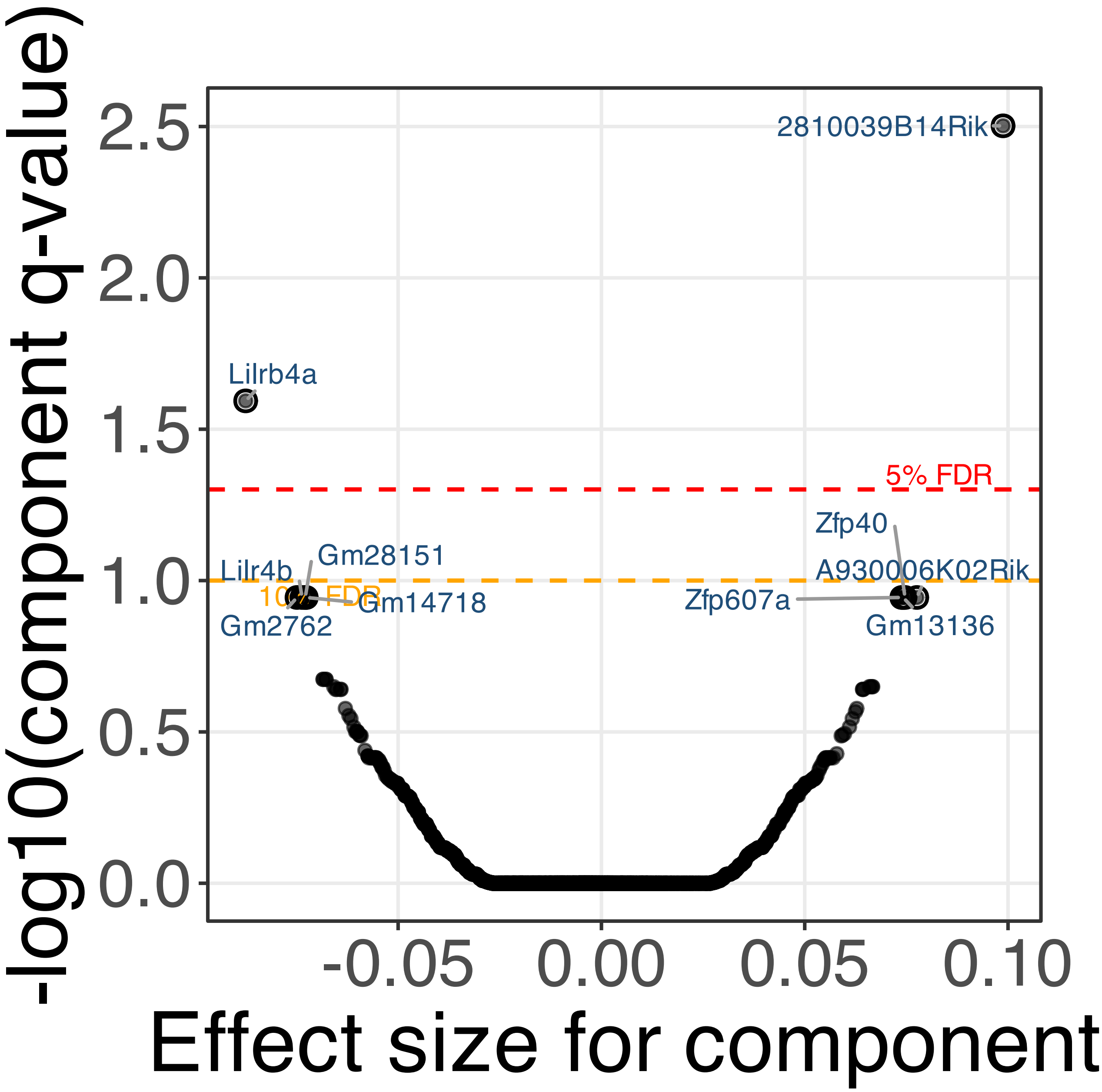


**Supplementary Figure 430.** SSANOVA-SVD based conditional inference for feature participation. Component 3.

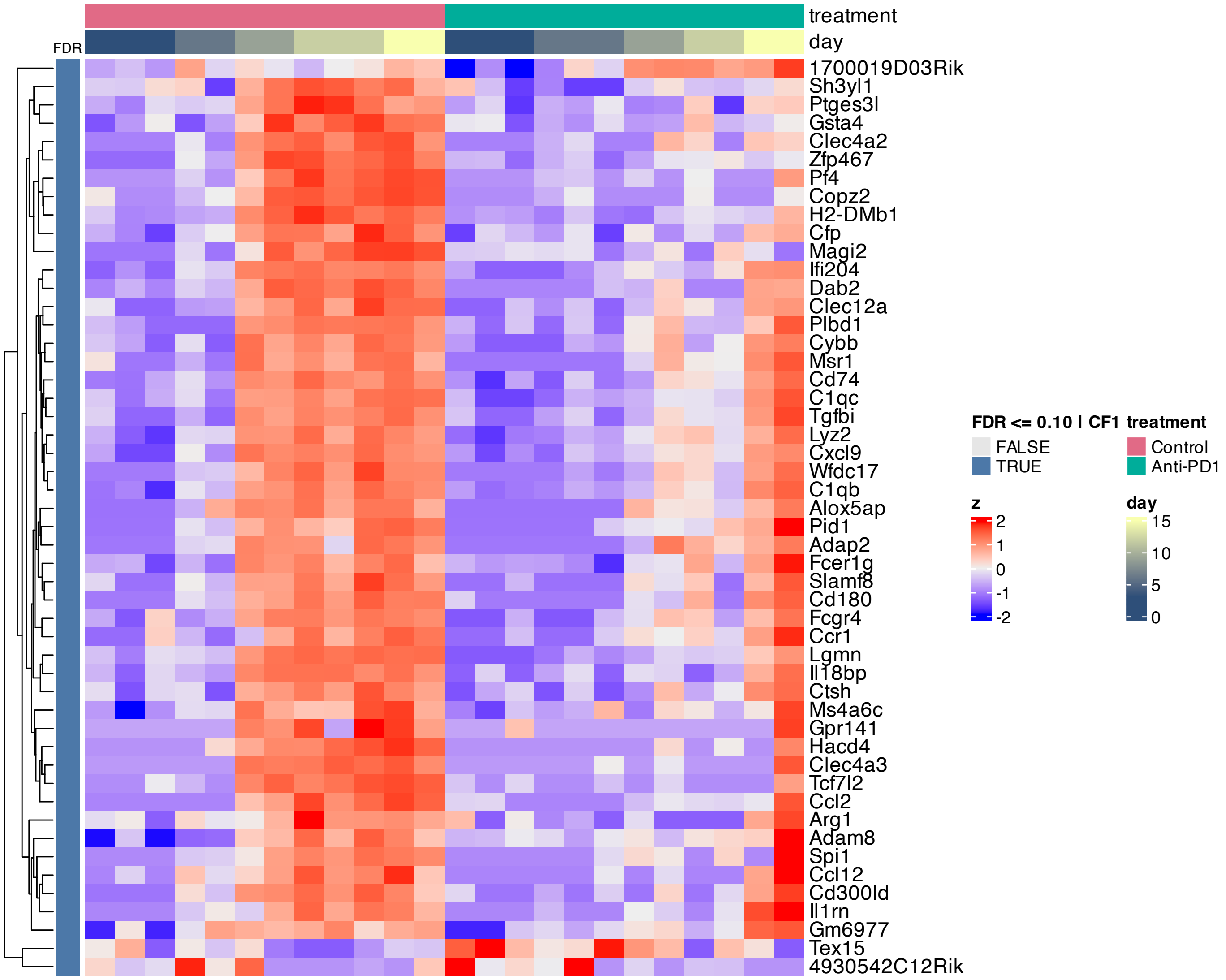


**Supplementary Figure 431.** Heatmap of features inferred to be associated with component 1 by SSANOVA-SVD.

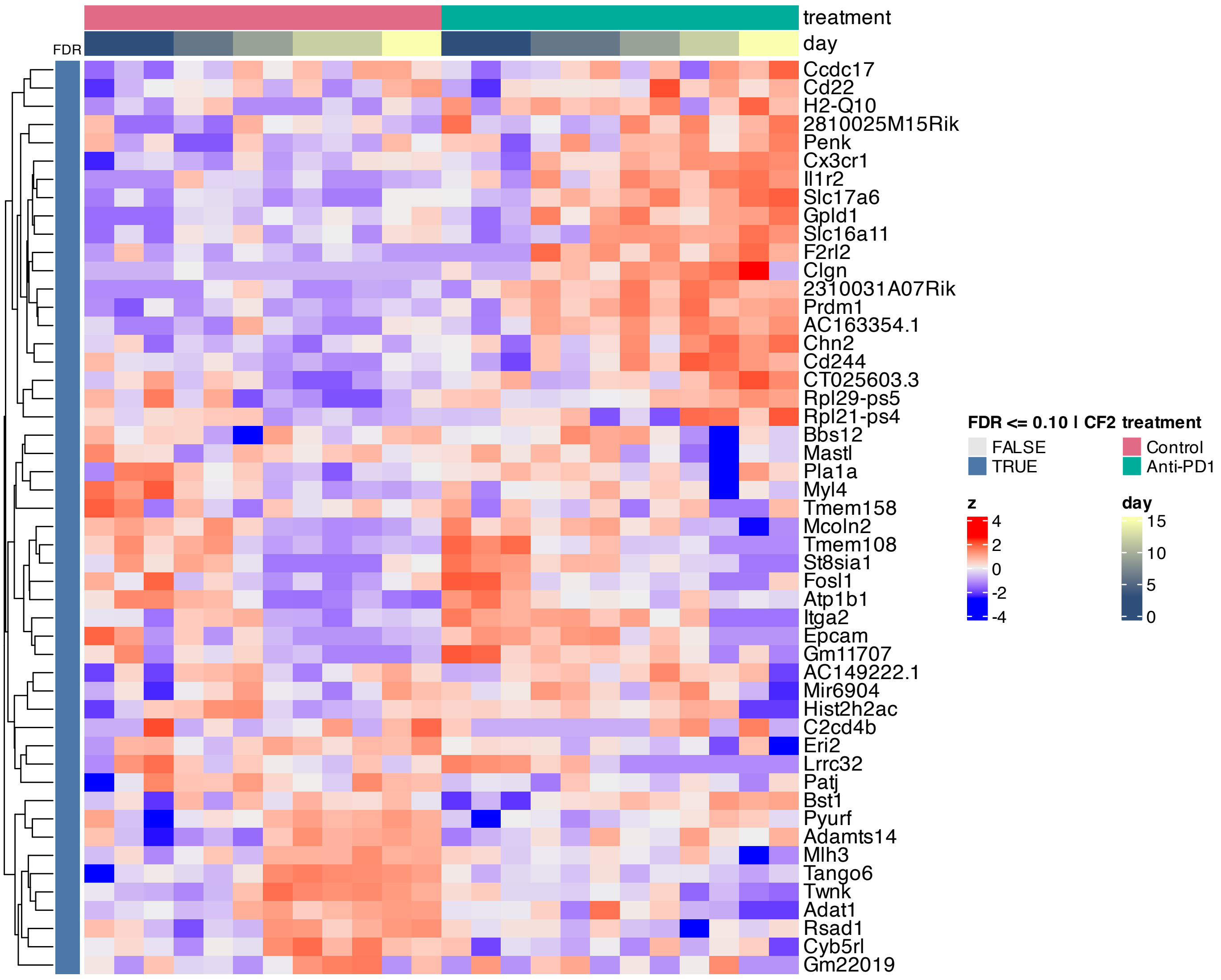


**Supplementary Figure 432.** Heatmap of features inferred to be associated with component 2 by SSANOVA-SVD.

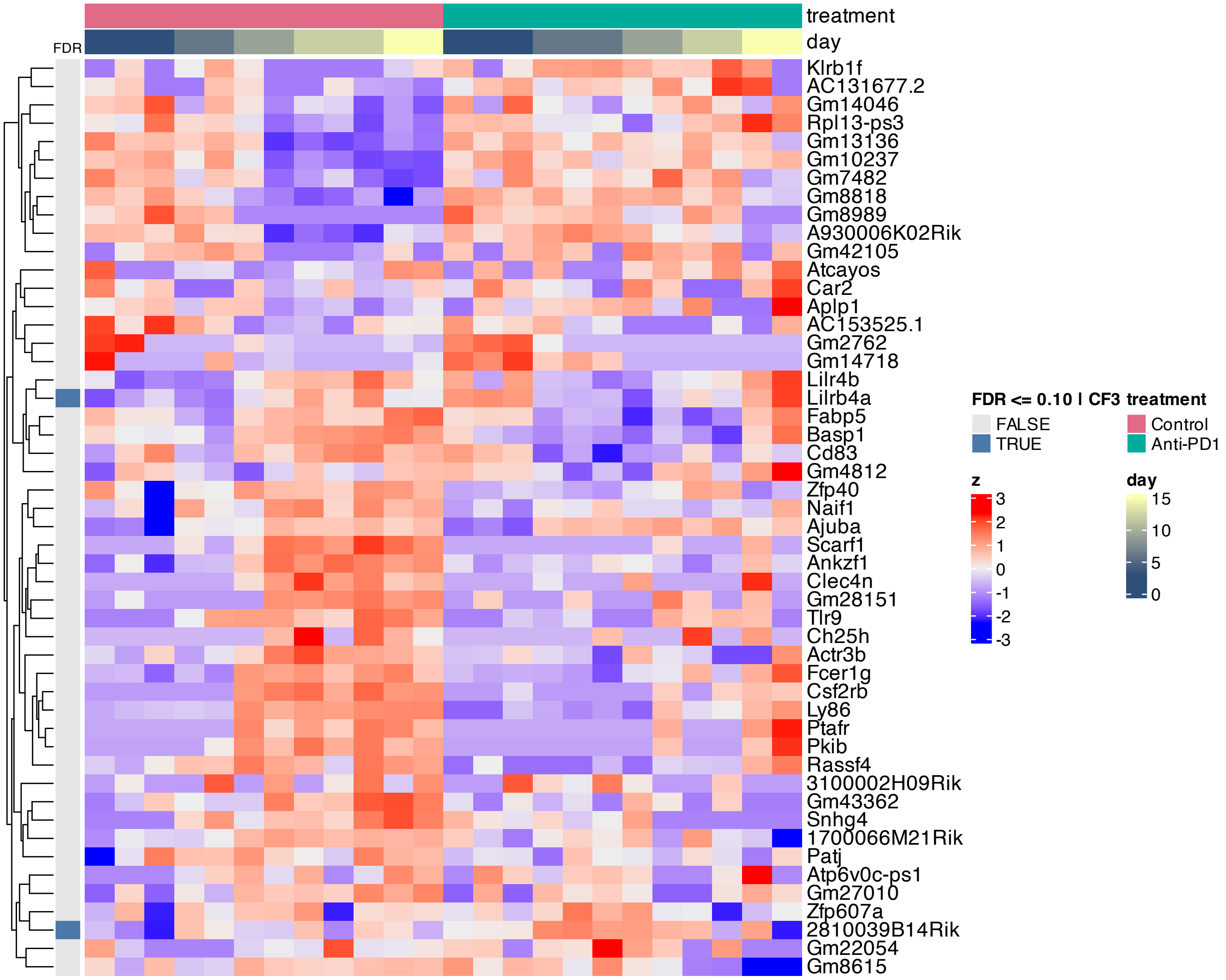


**Supplementary Figure 433.** Heatmap of features inferred to be associated with component 3 by SSANOVA-SVD.

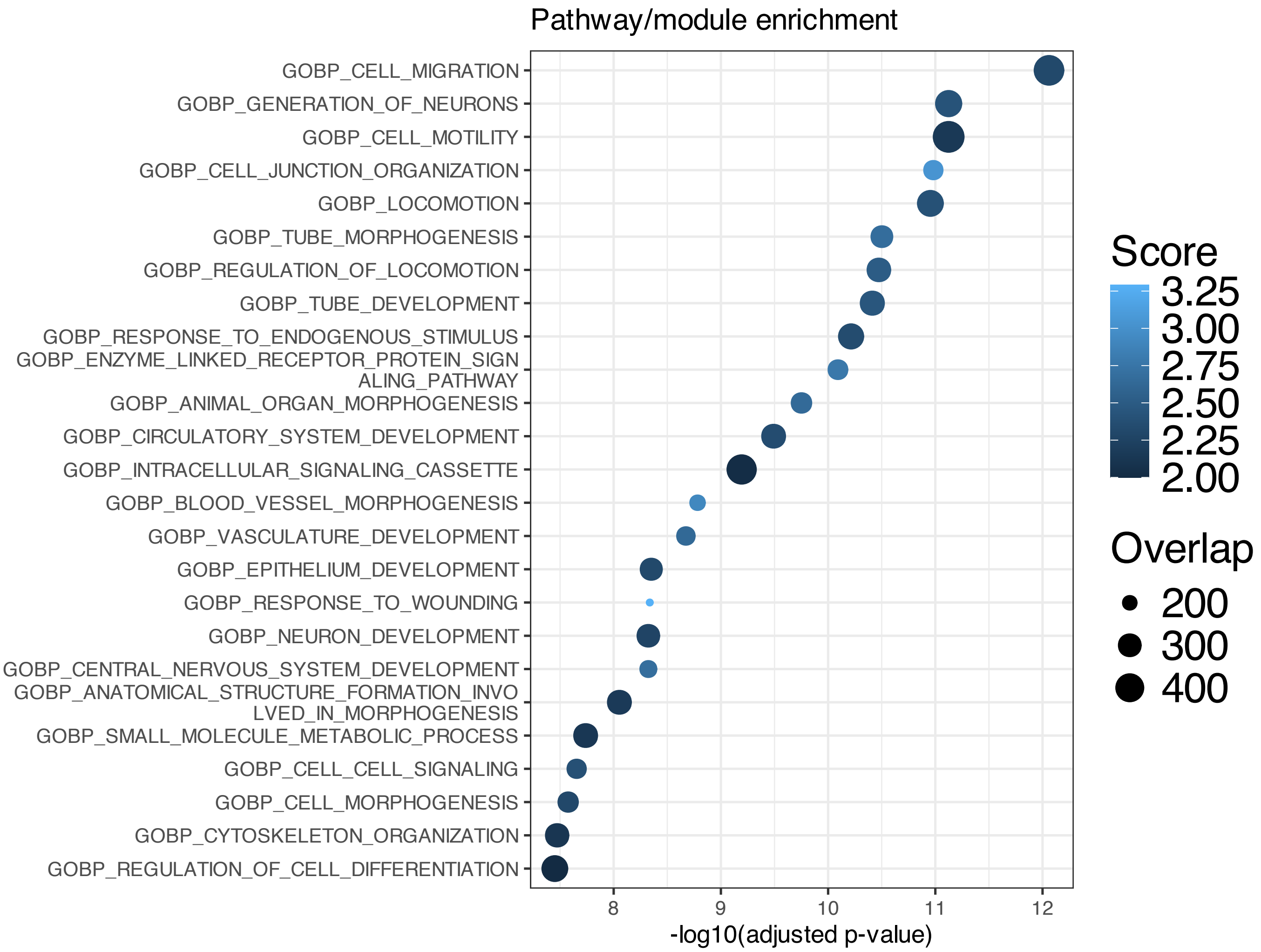


**Supplementary Figure 434.** Enrichment among features inferred to be associated with SSANOVA-SVD component 1; only terms with adjusted P $\leq 0.1$ are shown.

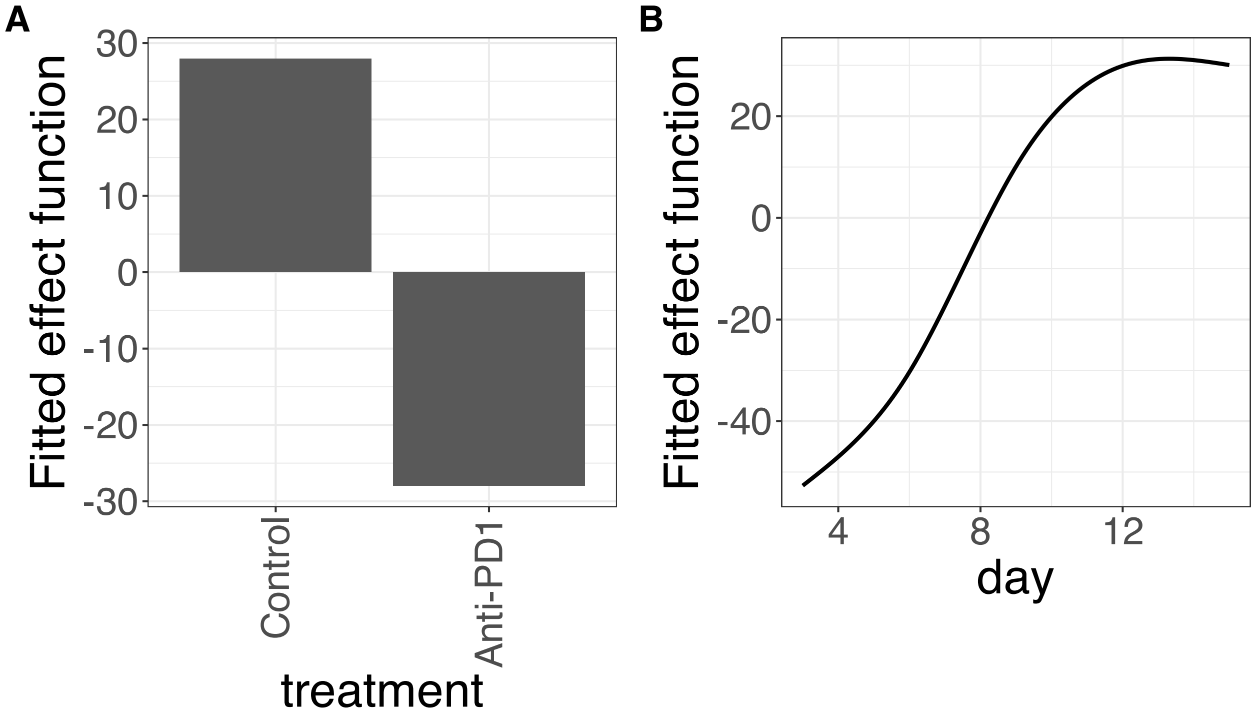


**Supplementary Figure 435.** Decomposition of reconstructed SSANOVA-SVD component function 1. (A) treatment. (B) day.

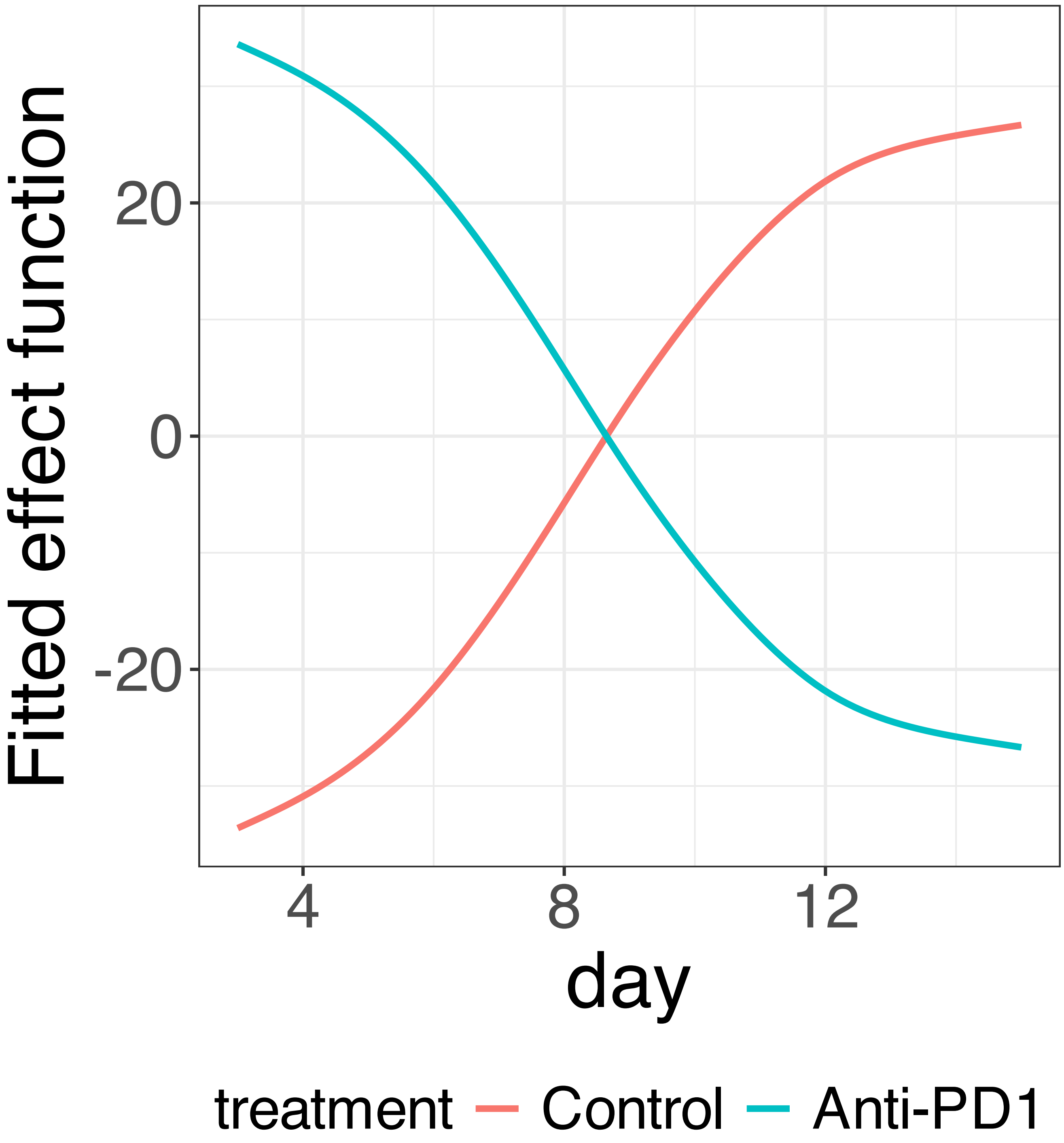


**Supplementary Figure 436.** Decomposition of reconstructed SSANOVA-SVD component function 1.

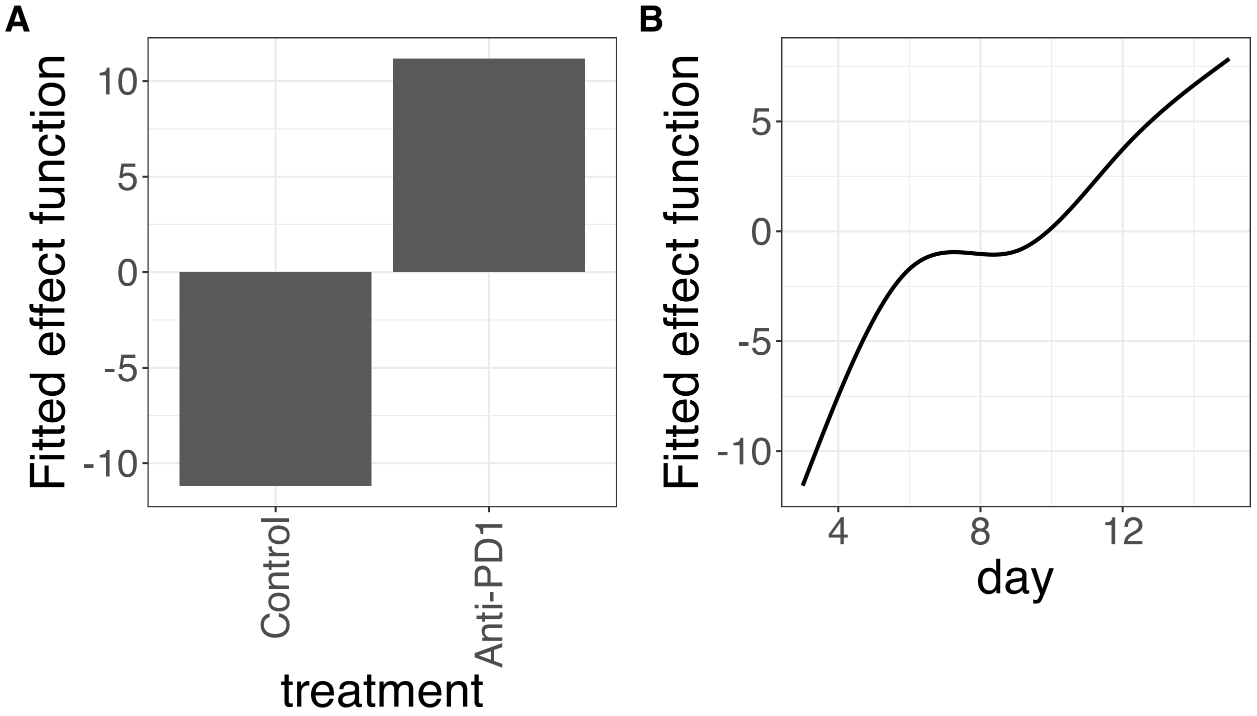


**Supplementary Figure 437.** Decomposition of reconstructed SSANOVA-SVD component function 2. (A) treatment. (B) day.

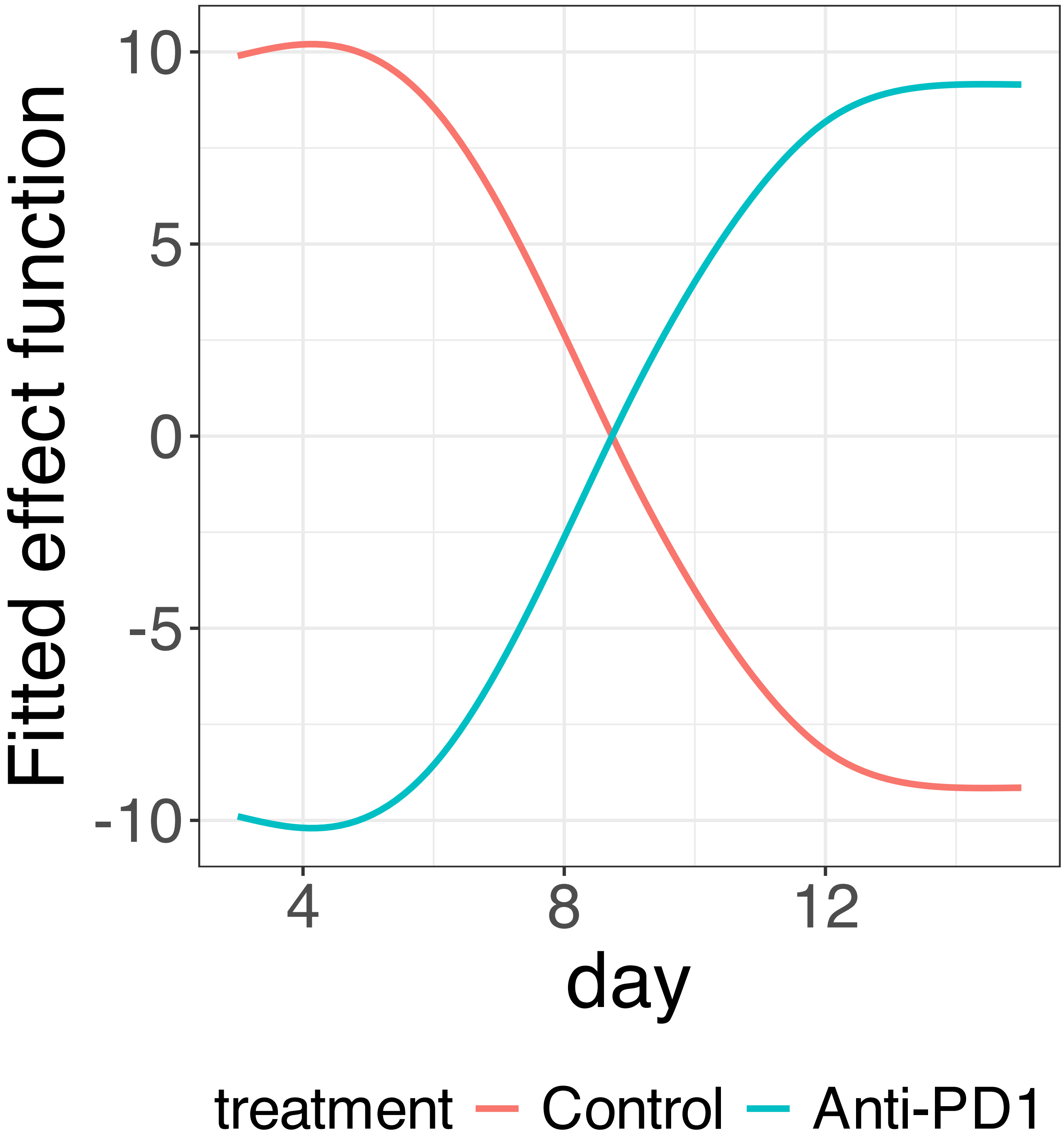


**Supplementary Figure 438.** Decomposition of reconstructed SSANOVA-SVD component function 2.

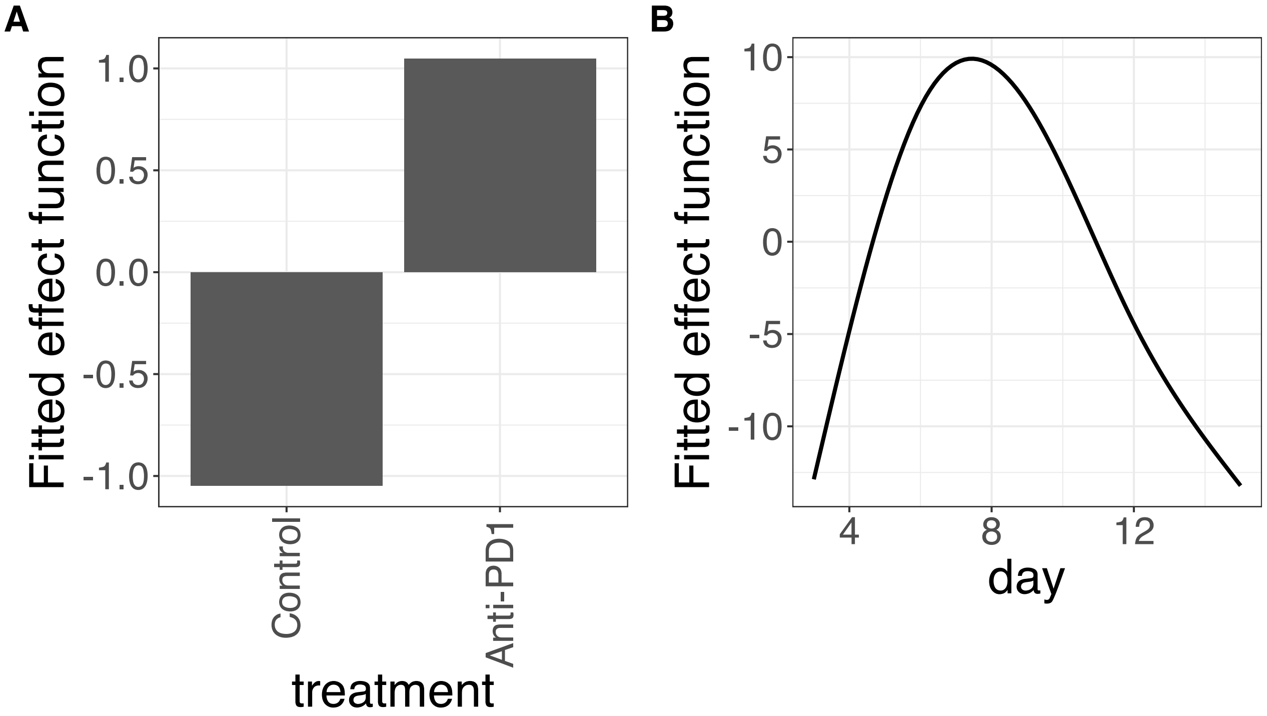


**Supplementary Figure 439.** Decomposition of reconstructed SSANOVA-SVD component function 3. (A) treatment. (B) day.

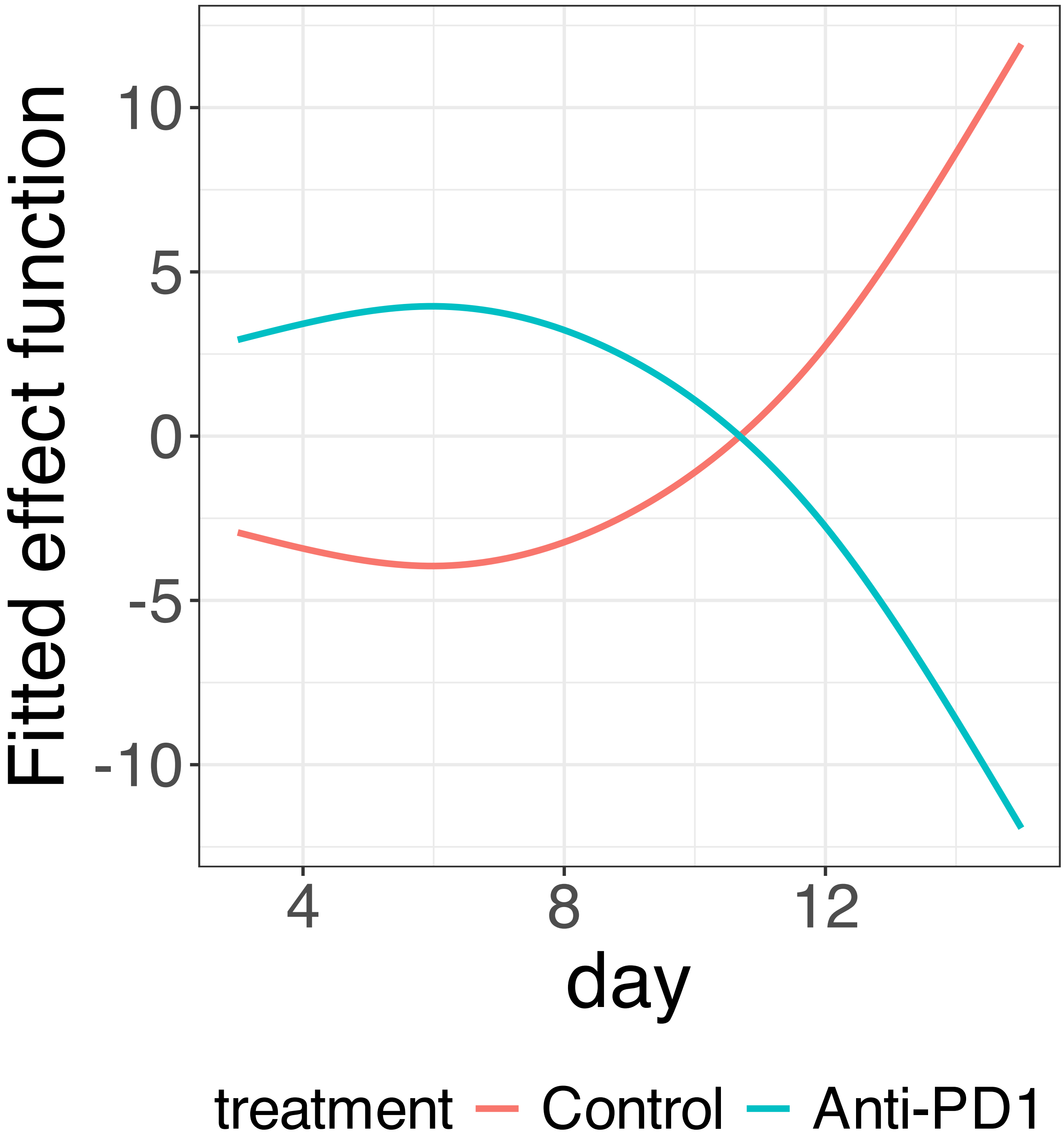


**Supplementary Figure 440.** Decomposition of reconstructed SSANOVA-SVD component function 3.

PCA

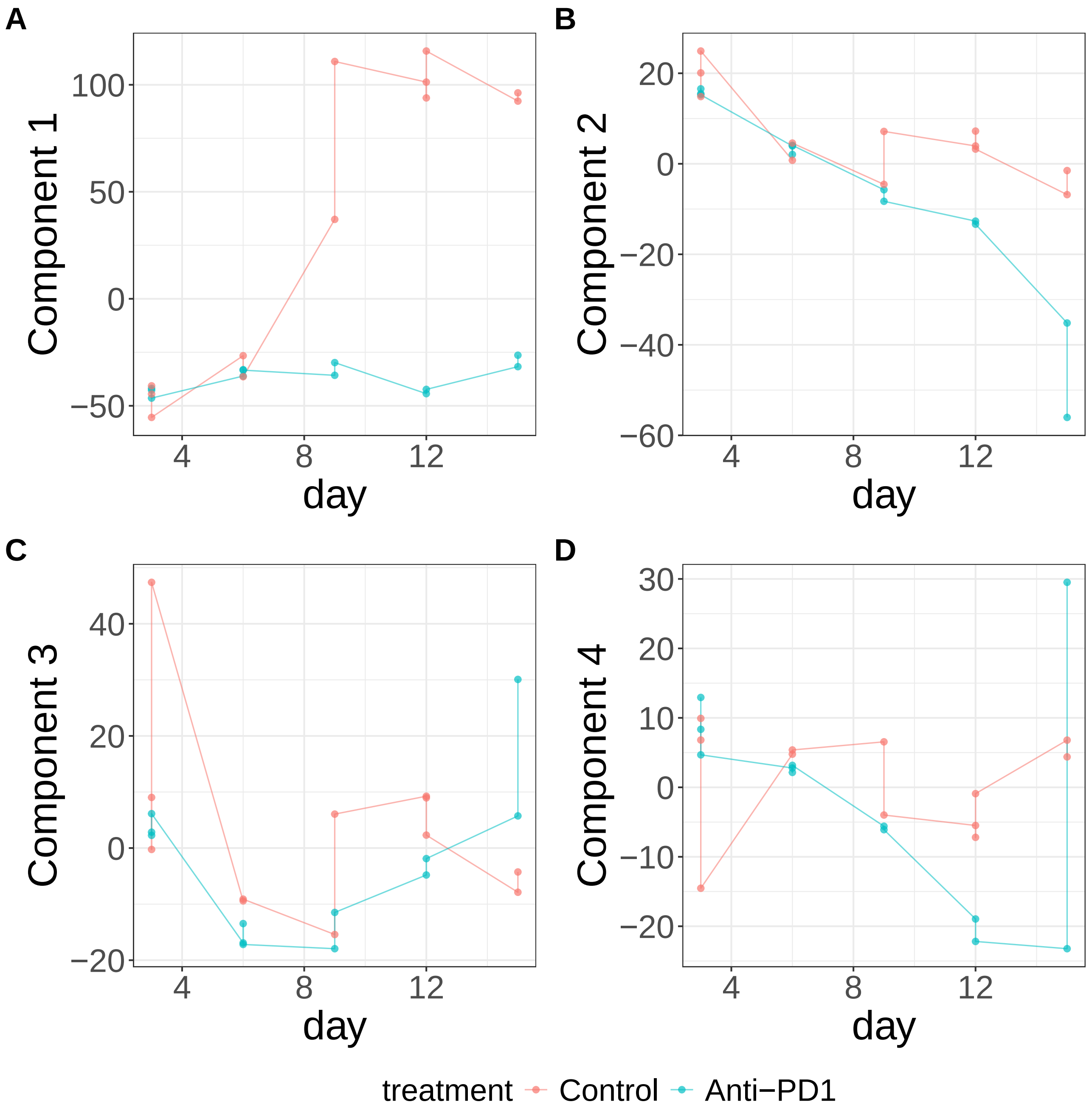


**Supplementary Figure 441.** Components for PCA. (A) Component 1. (B) Component 2. (C) Component 3. (D) Component 4.

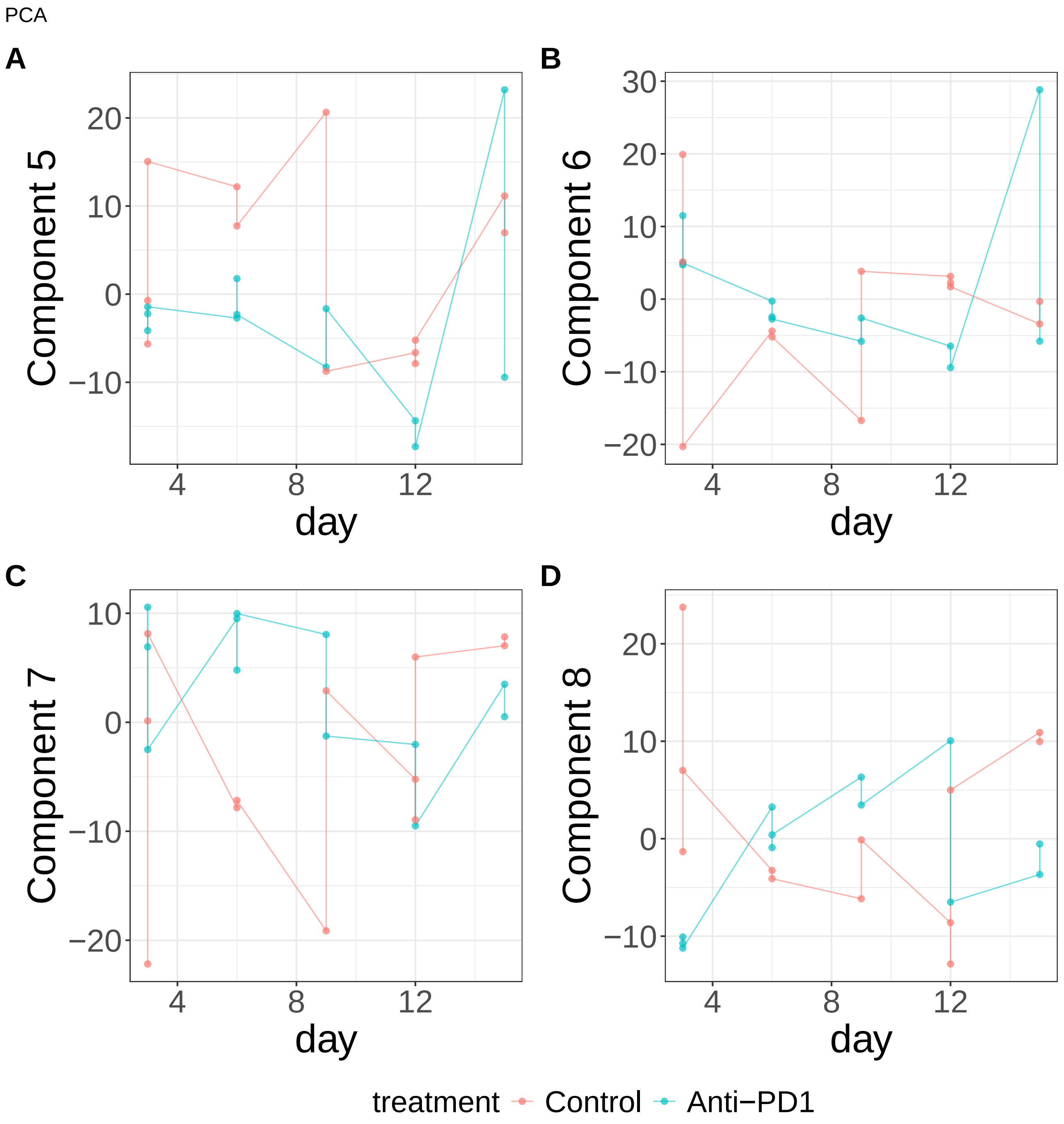


**Supplementary Figure 442.** Components for PCA. (A) Component 5. (B) Component 6. (C) Component 7. (D) Component 8.

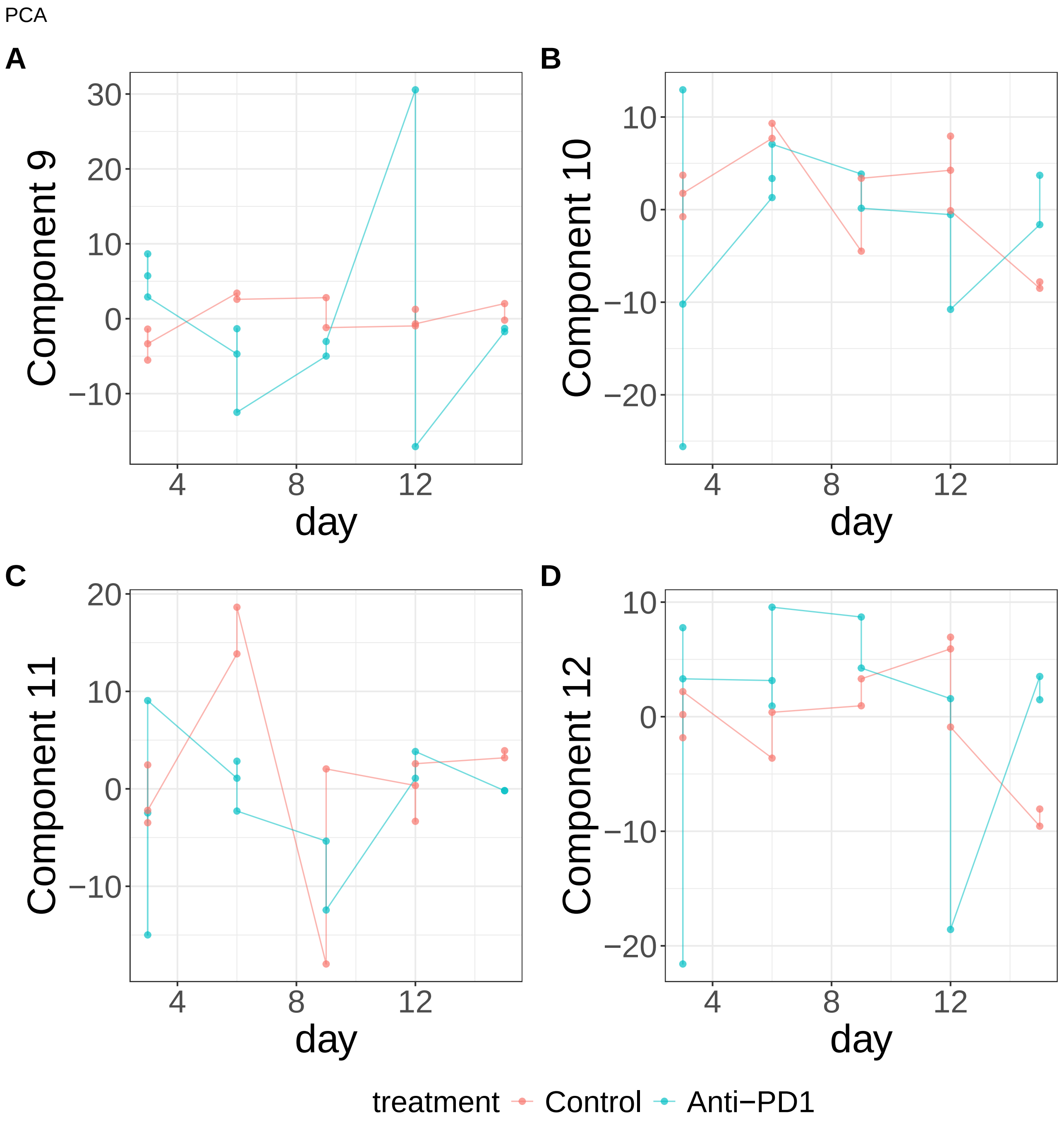


**Supplementary Figure 443.** Components for PCA. (A) Component 9. (B) Component 10. (C) Component 11. (D) Component 12.

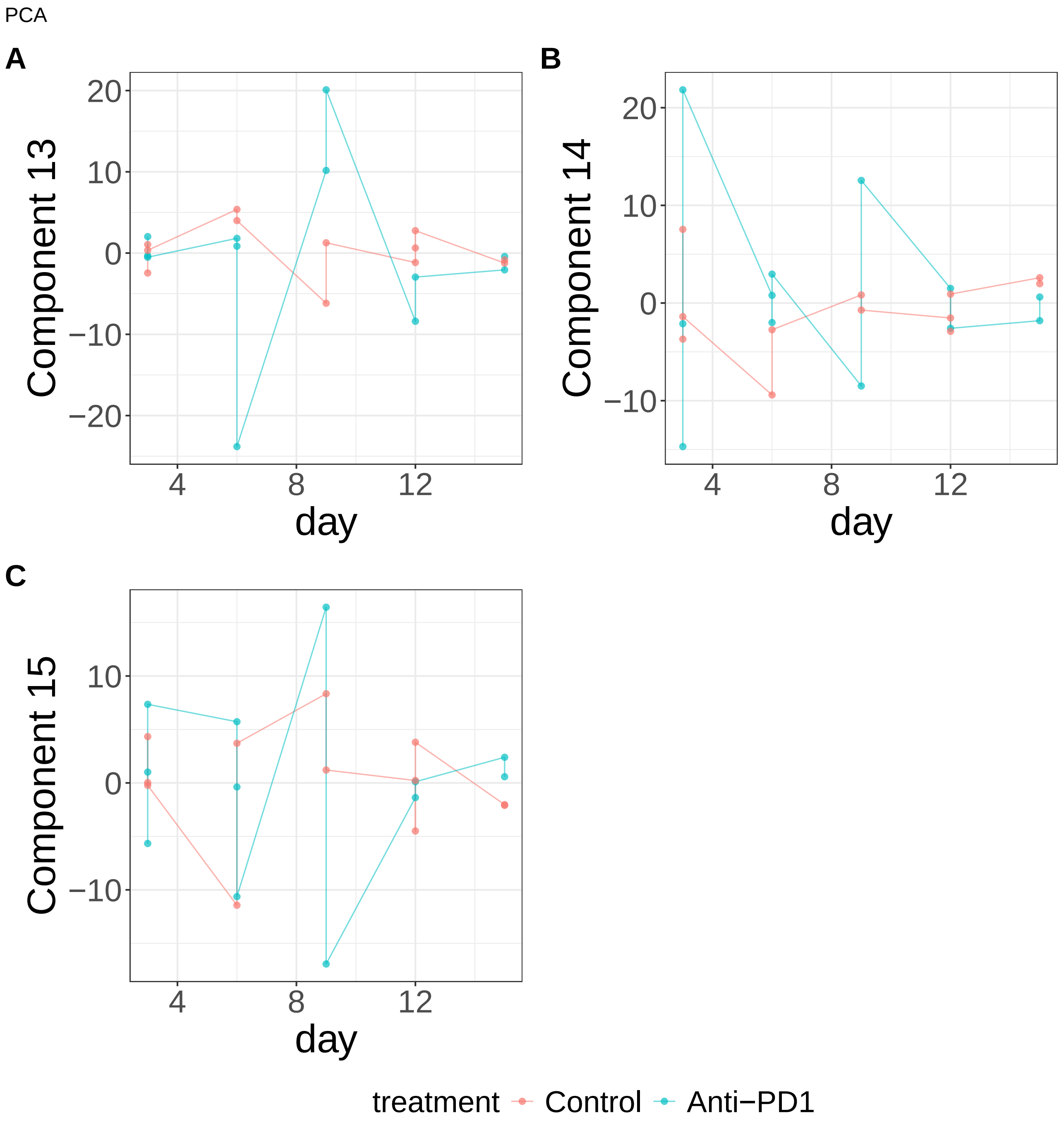


**Supplementary Figure 444.** Components for PCA. (A) Component 13. (B) Component 14. (C) Component 15.

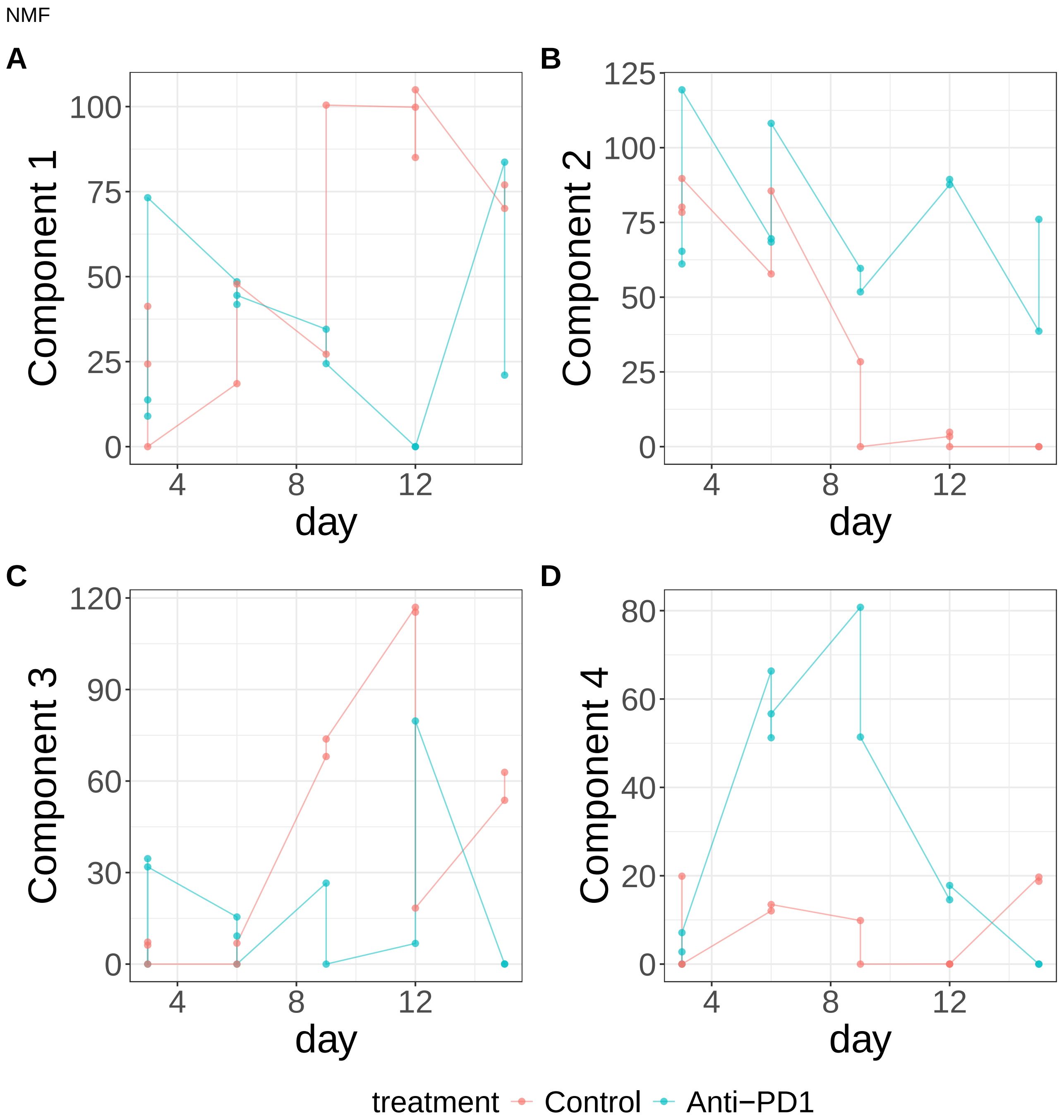


**Supplementary Figure 445.** Components for NMF. (A) Component 1. (B) Component 2. (C) Component 3. (D) Component 4.

NMF

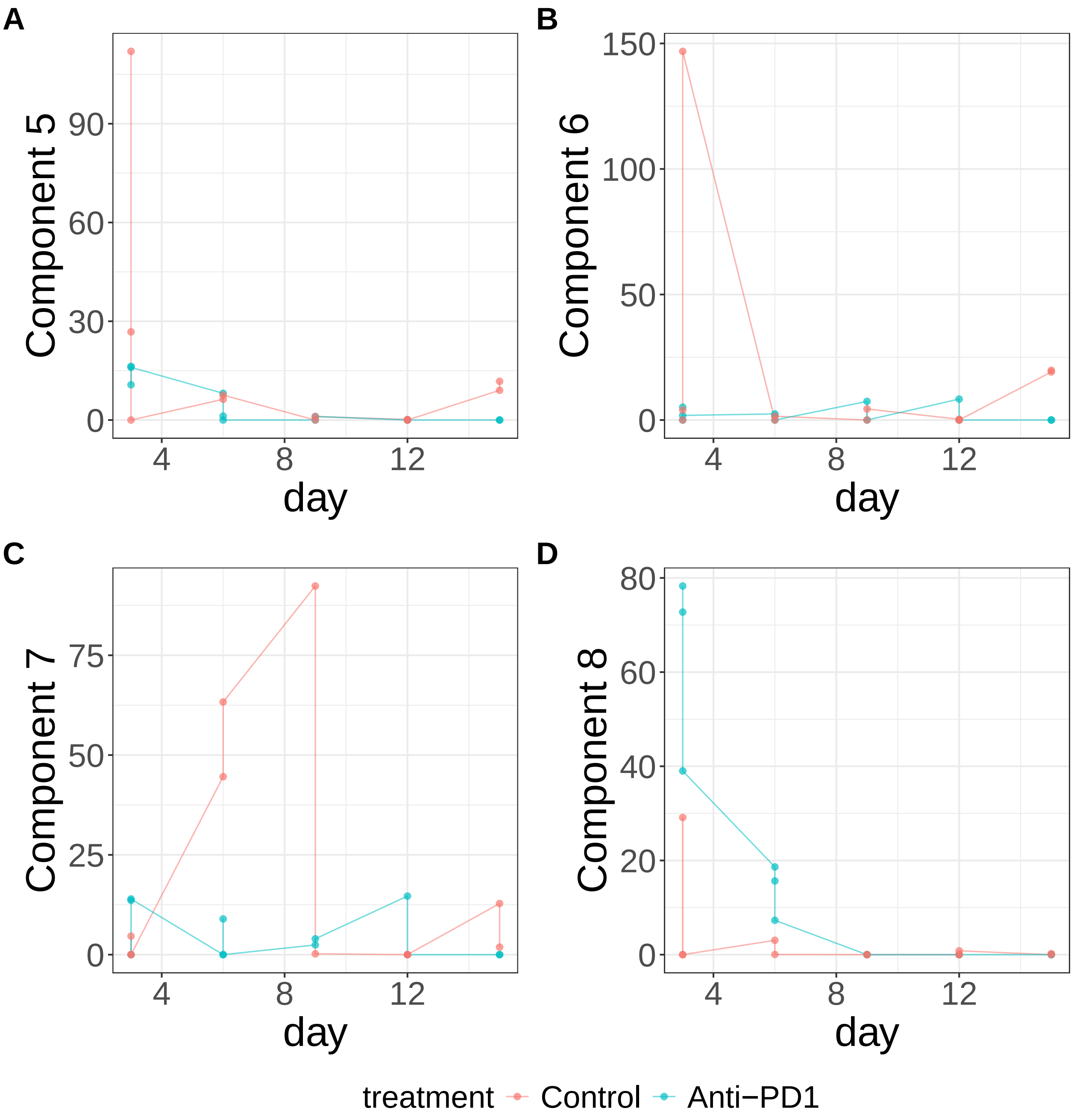


**Supplementary Figure 446.** Components for NMF. (A) Component 5. (B) Component 6. (C) Component 7. (D) Component 8.

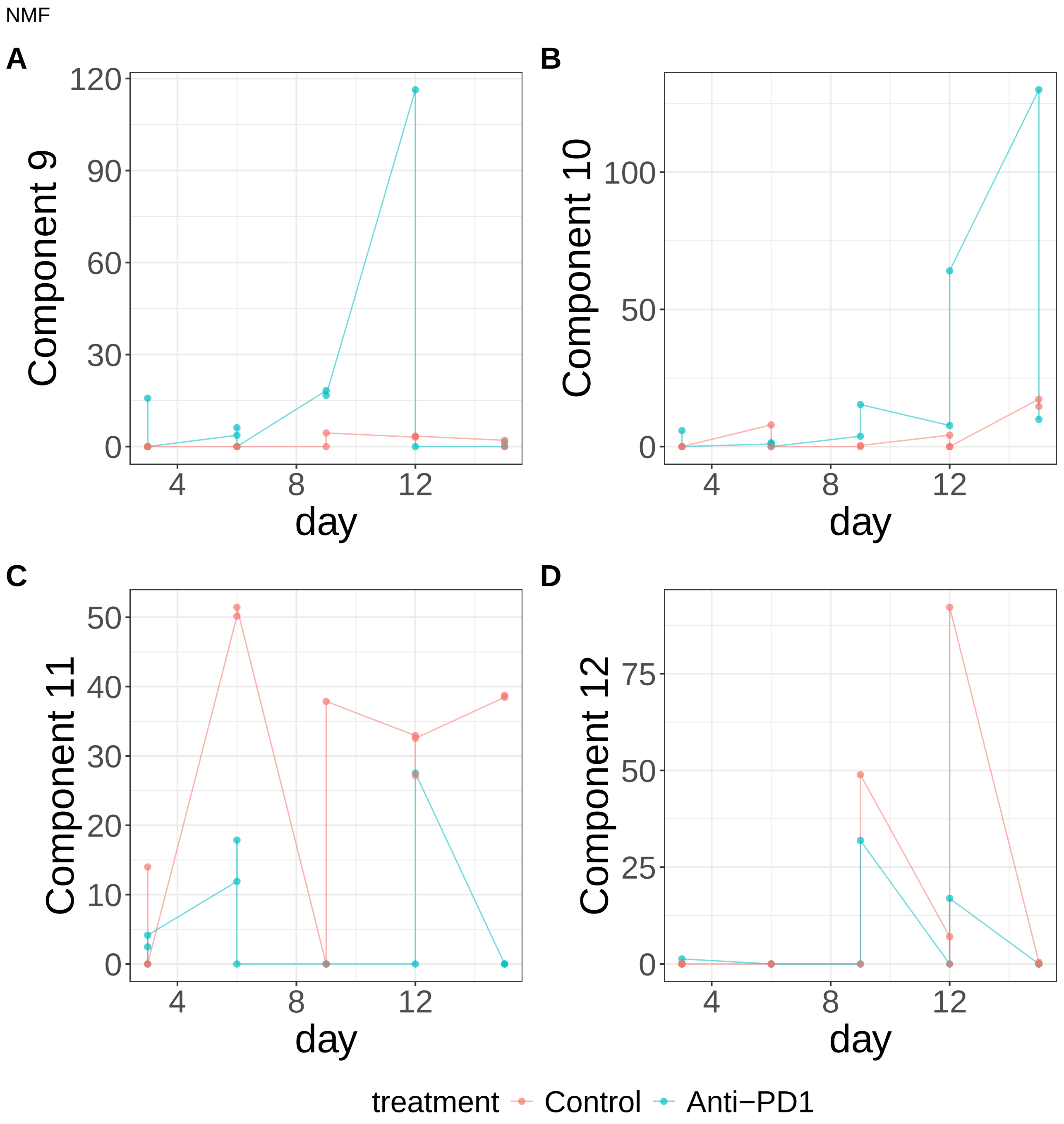


**Supplementary Figure 447.** Components for NMF. (A) Component 9. (B) Component 10. (C) Component 11. (D) Component 12.

NMF

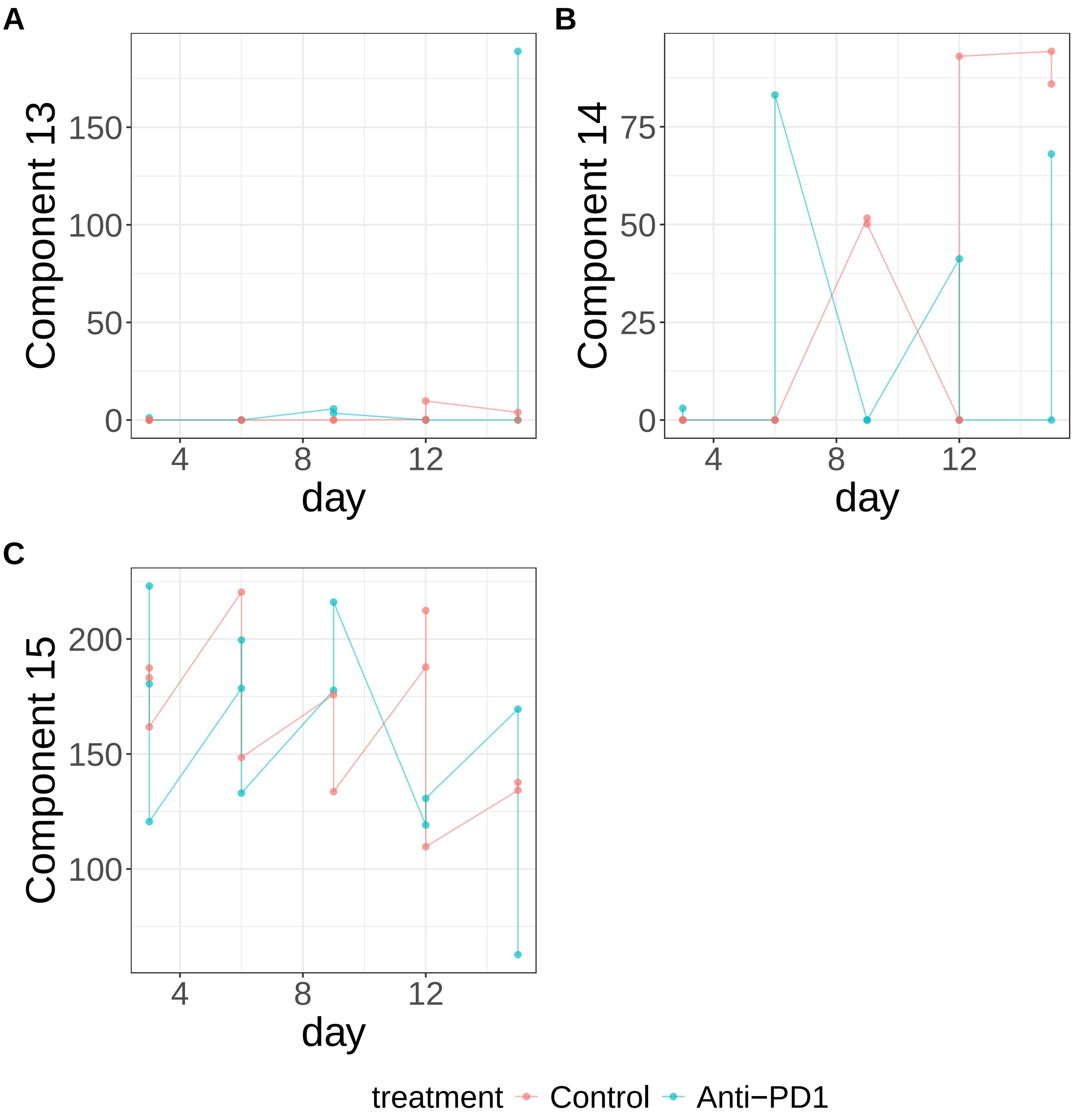


**Supplementary Figure 448.** Components for NMF. (A) Component 13. (B) Component 14. (C) Component 15.

PLS

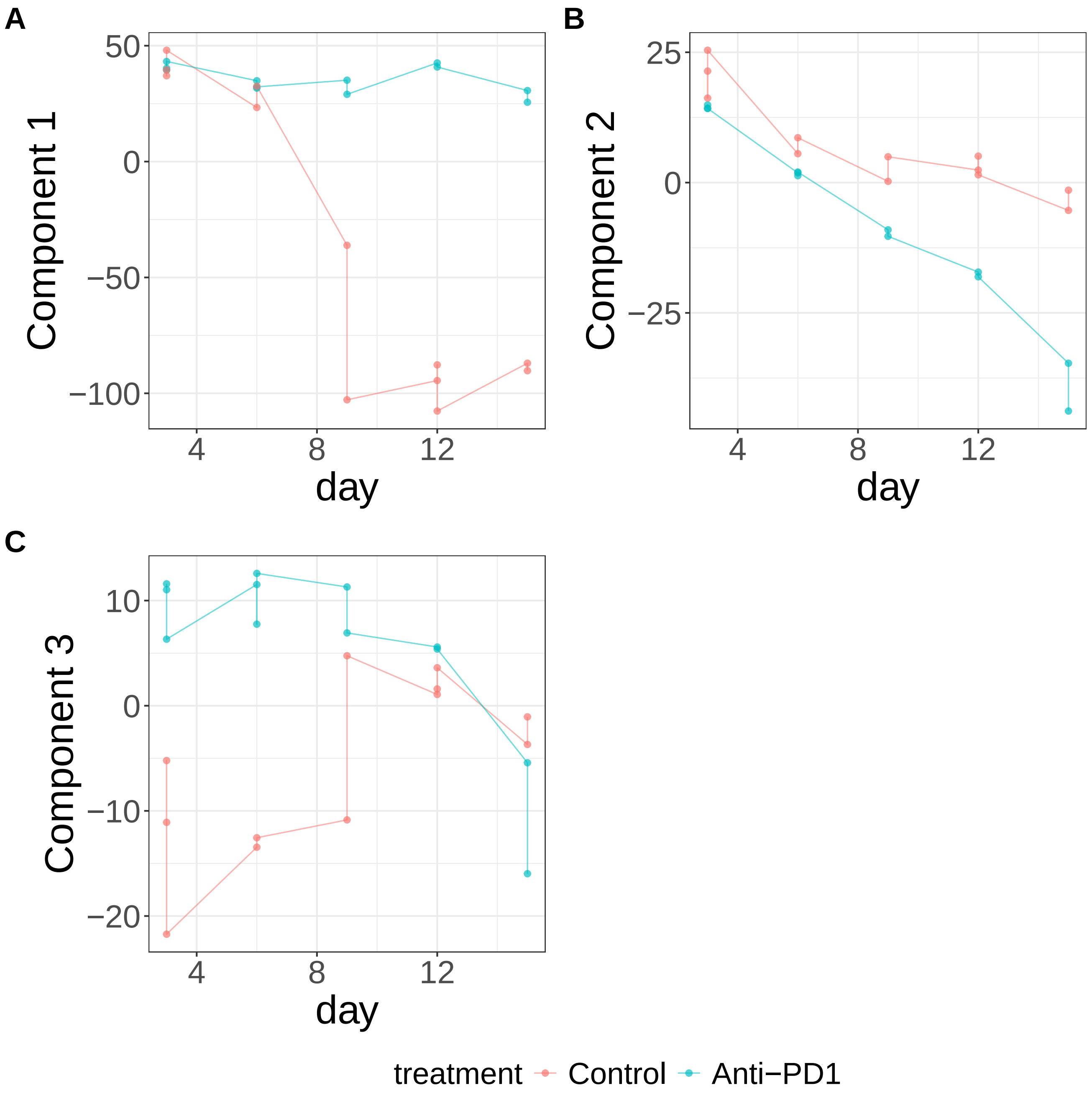


**Supplementary Figure 449.** Components for PLS. (A) Component 1. (B) Component 2. (C) Component 3.

sPCA: keepX=50

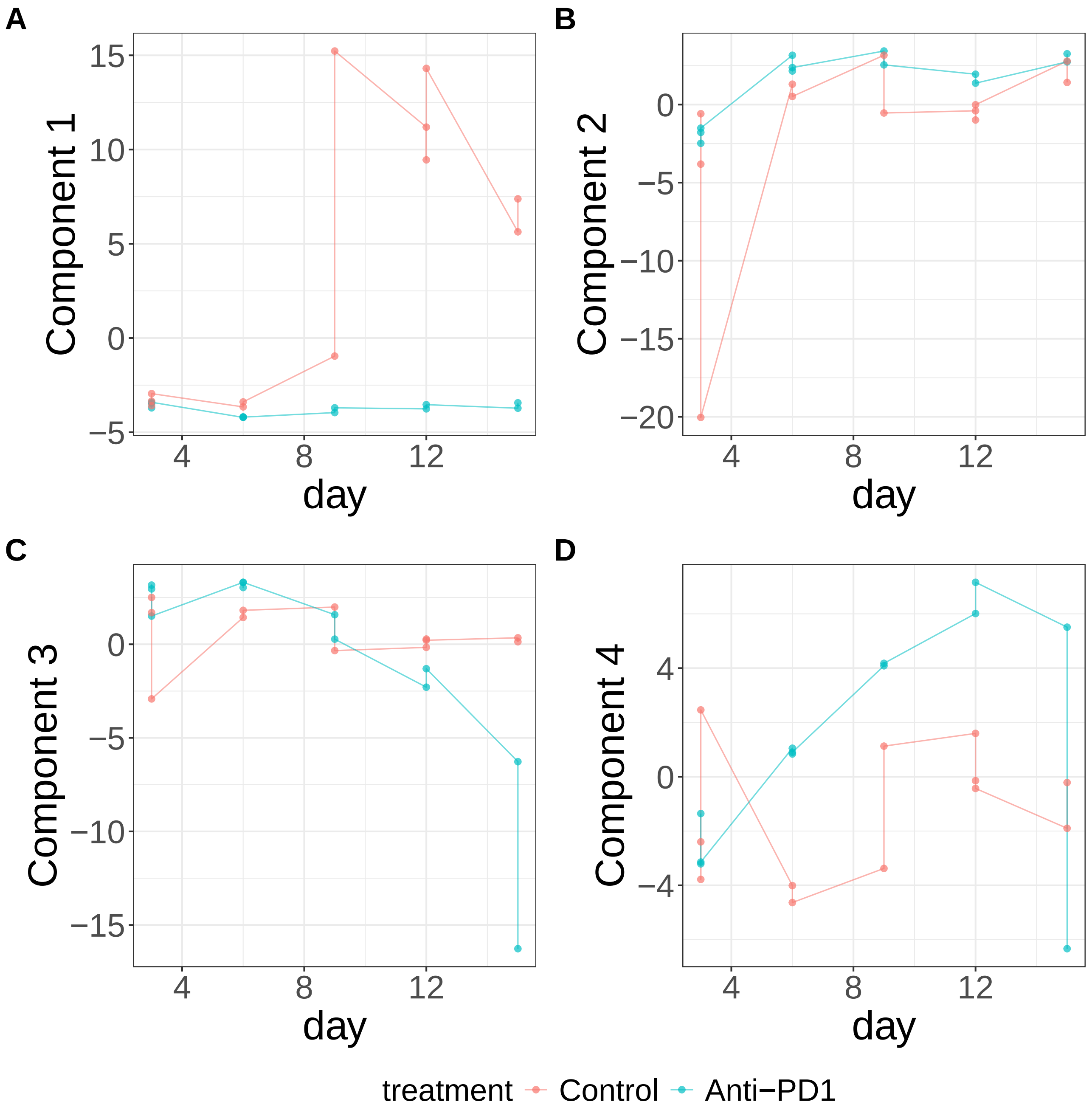


**Supplementary Figure 450.** Components for sPCA: keepX=50. (A) Component 1. (B) Component 2. (C) Component 3. (D) Component 4.

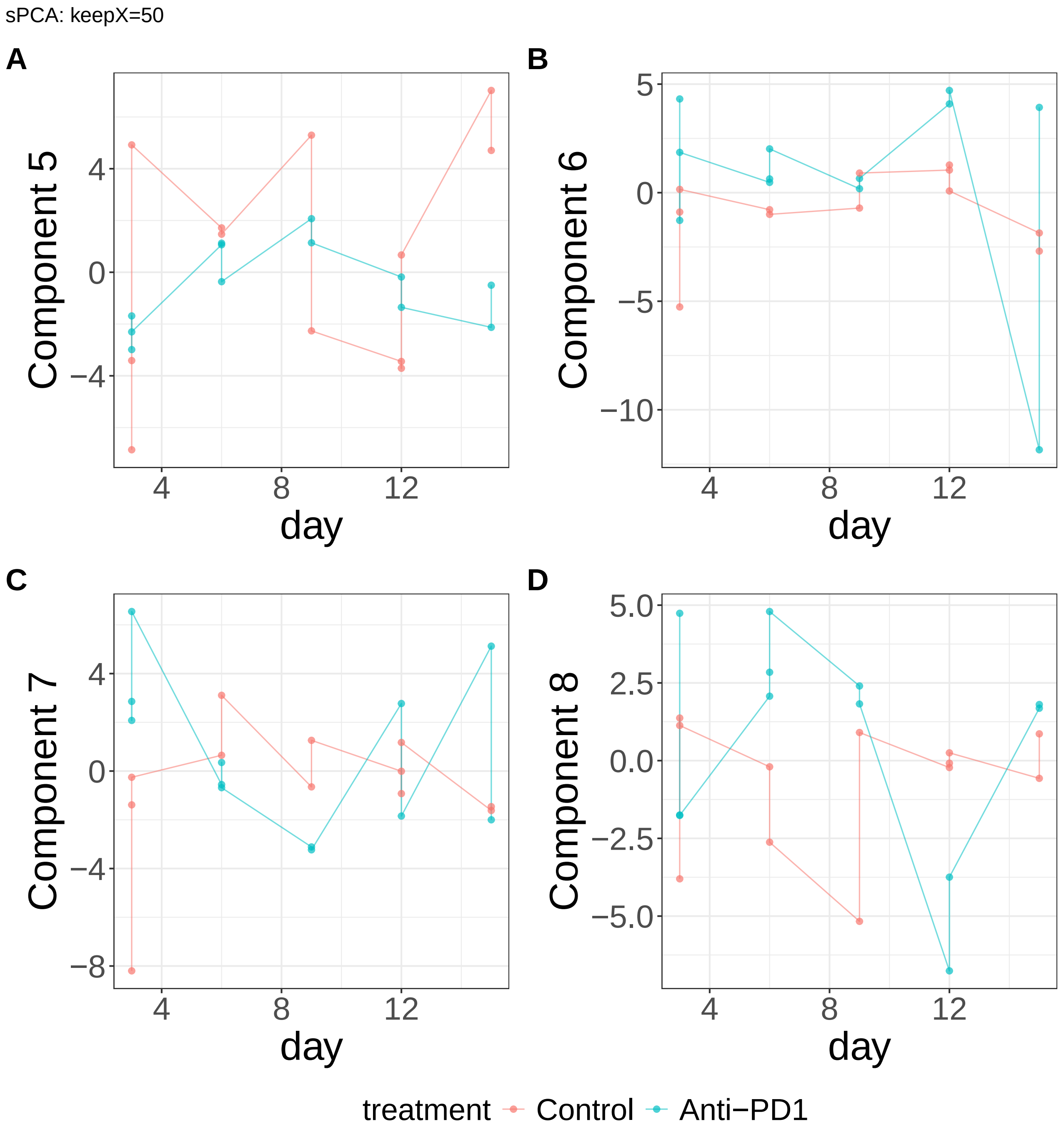


**Supplementary Figure 451.** Components for sPCA: keepX=50. (A) Component 5. (B) Component 6. (C) Component 7. (D) Component 8.

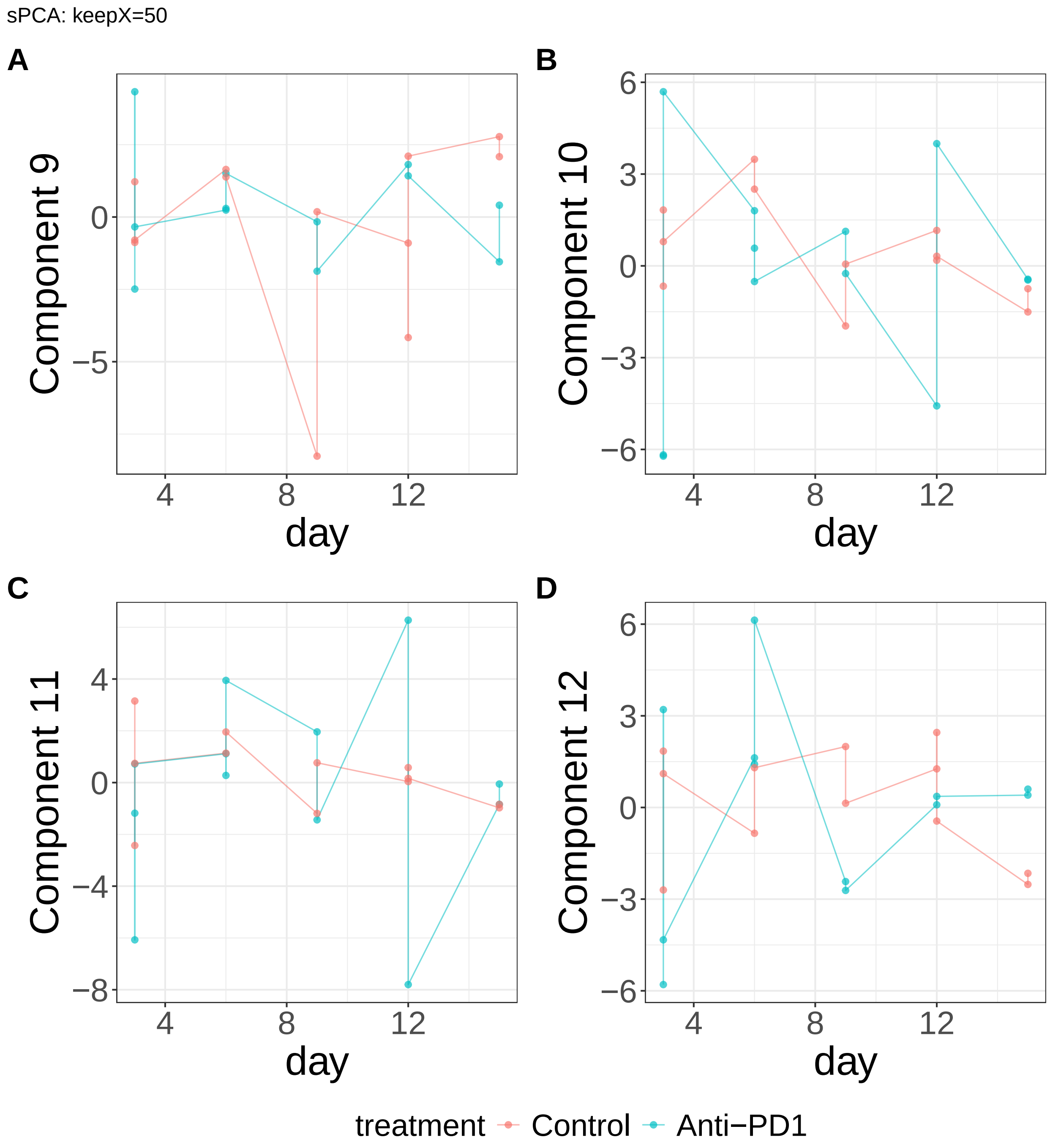


**Supplementary Figure 452.** Components for sPCA: keepX=50. (A) Component 9. (B) Component 10. (C) Component 11. (D) Component 12.

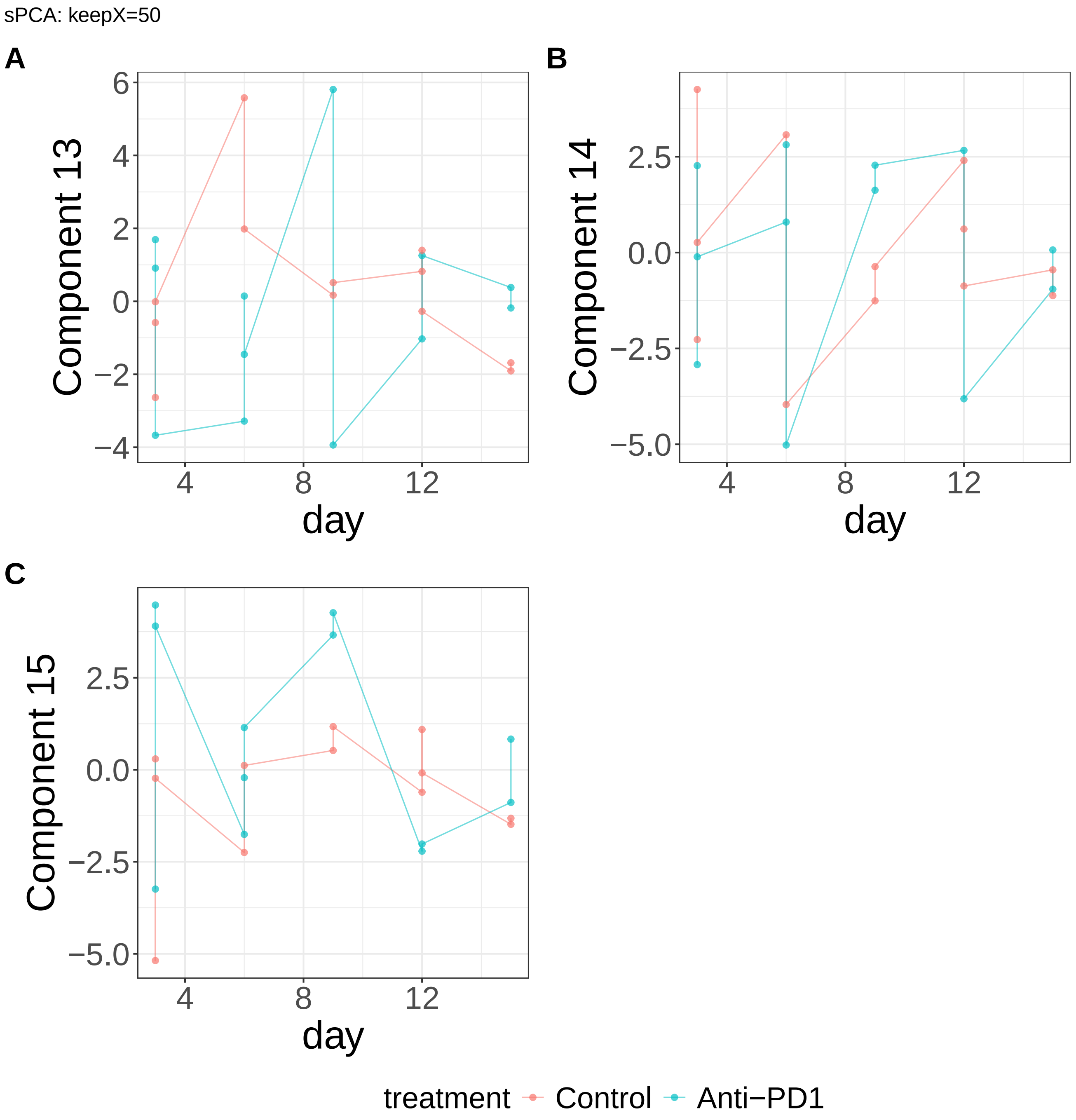


**Supplementary Figure 453.** Components for sPCA: keepX=50. (A) Component 13. (B) Component 14. (C) Component 15.

sPCA: keepX=1250

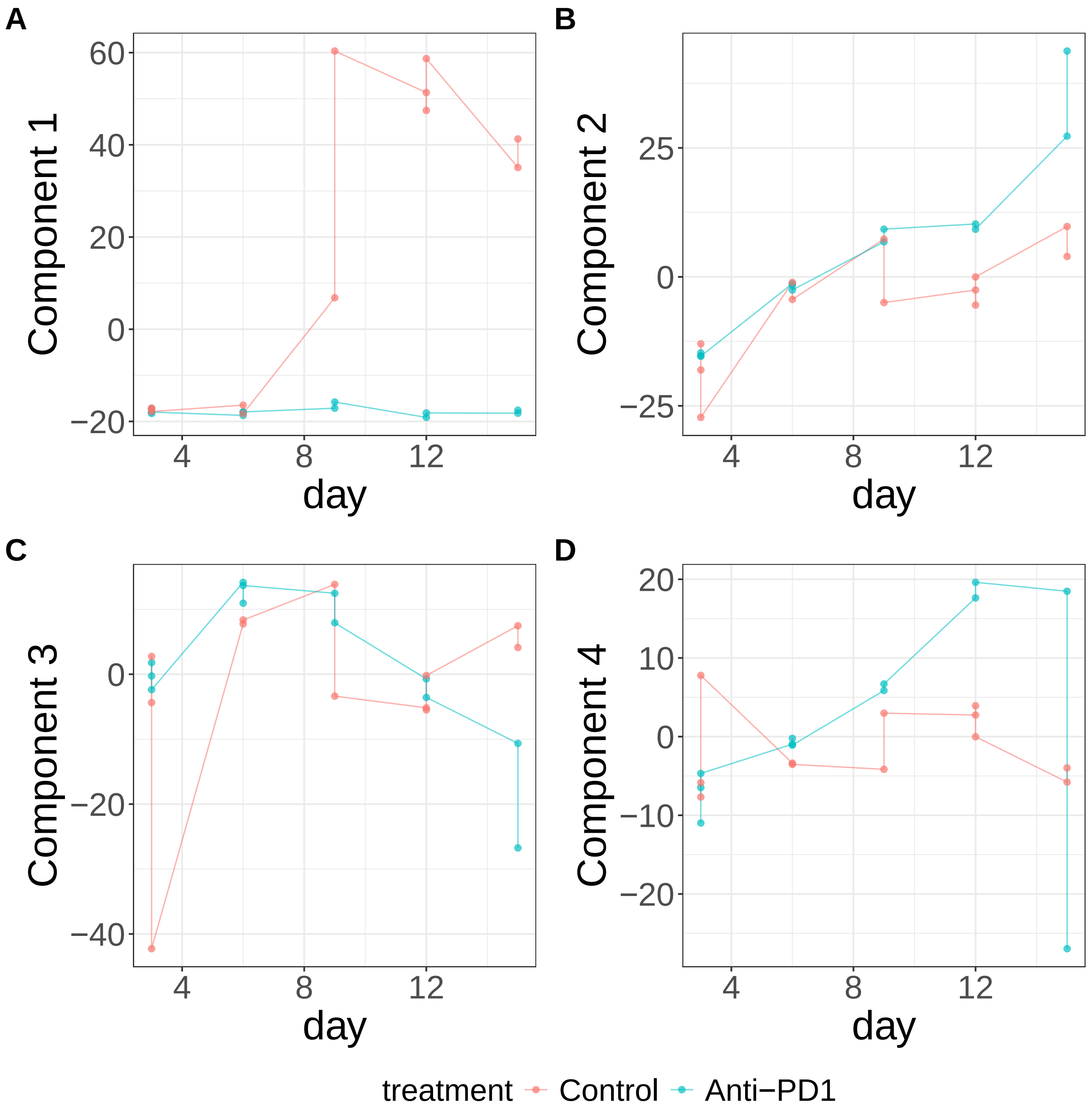


**Supplementary Figure 454.** Components for sPCA: keepX=1250. (A) Component 1. (B) Component 2. (C) Component 3. (D) Component 4.

sPCA: keepX=1250

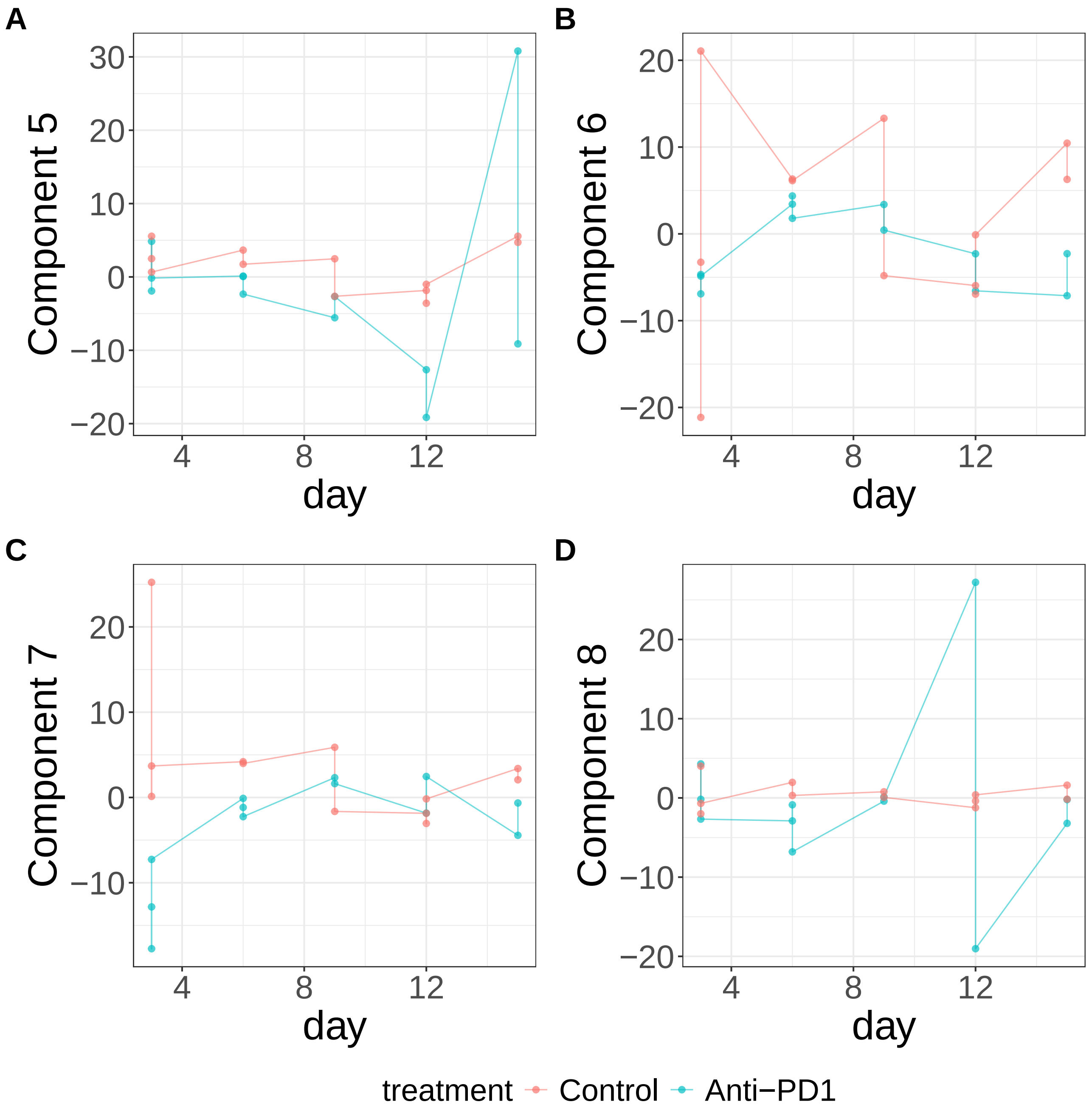


**Supplementary Figure 455.** Components for sPCA: keepX=1250. (A) Component 5. (B) Component 6. (C) Component 7. (D) Component 8.

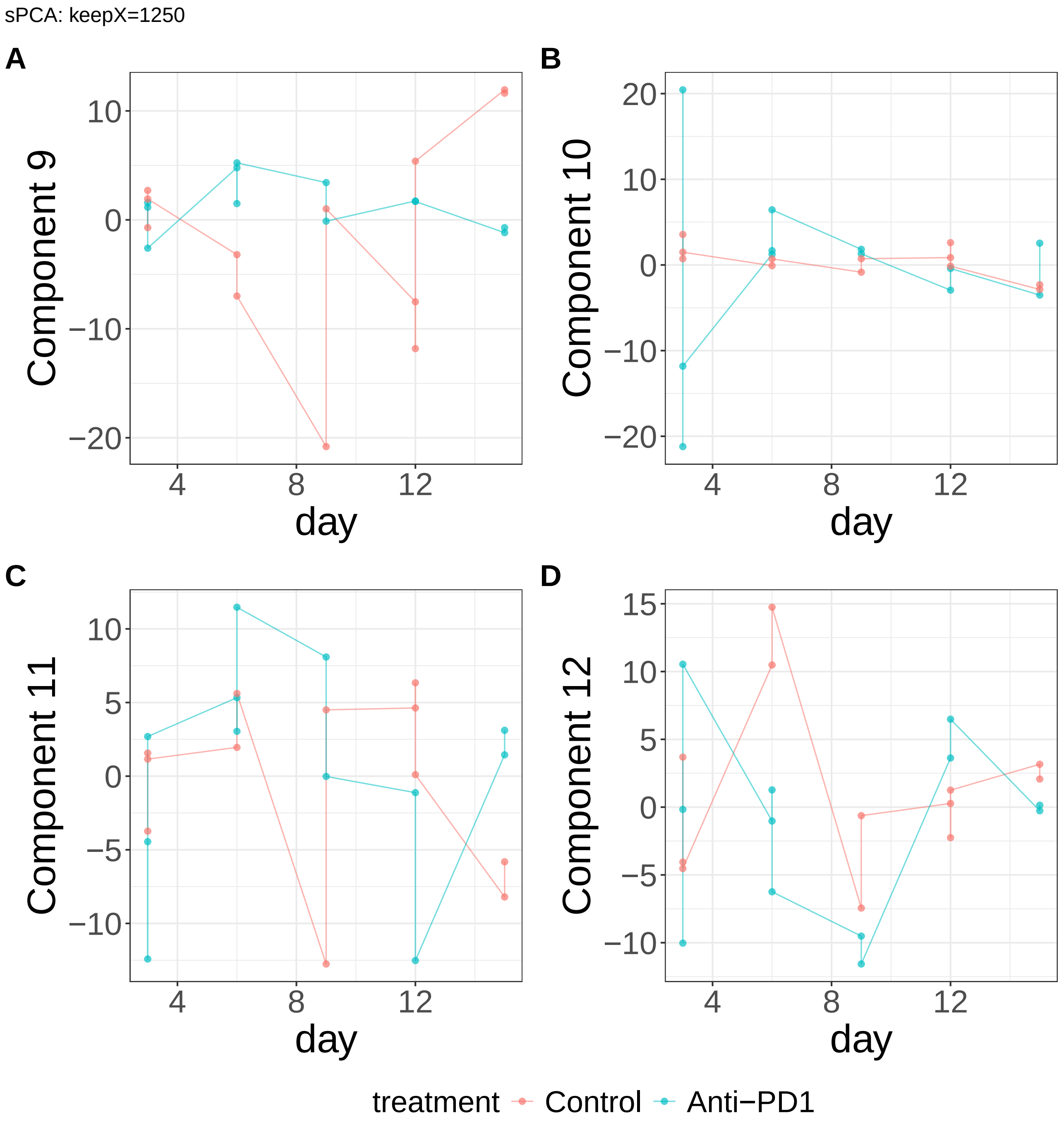


**Supplementary Figure 456.** Components for sPCA: keepX=1250. (A) Component 9. (B) Component 10. (C) Component 11. (D) Component 12.

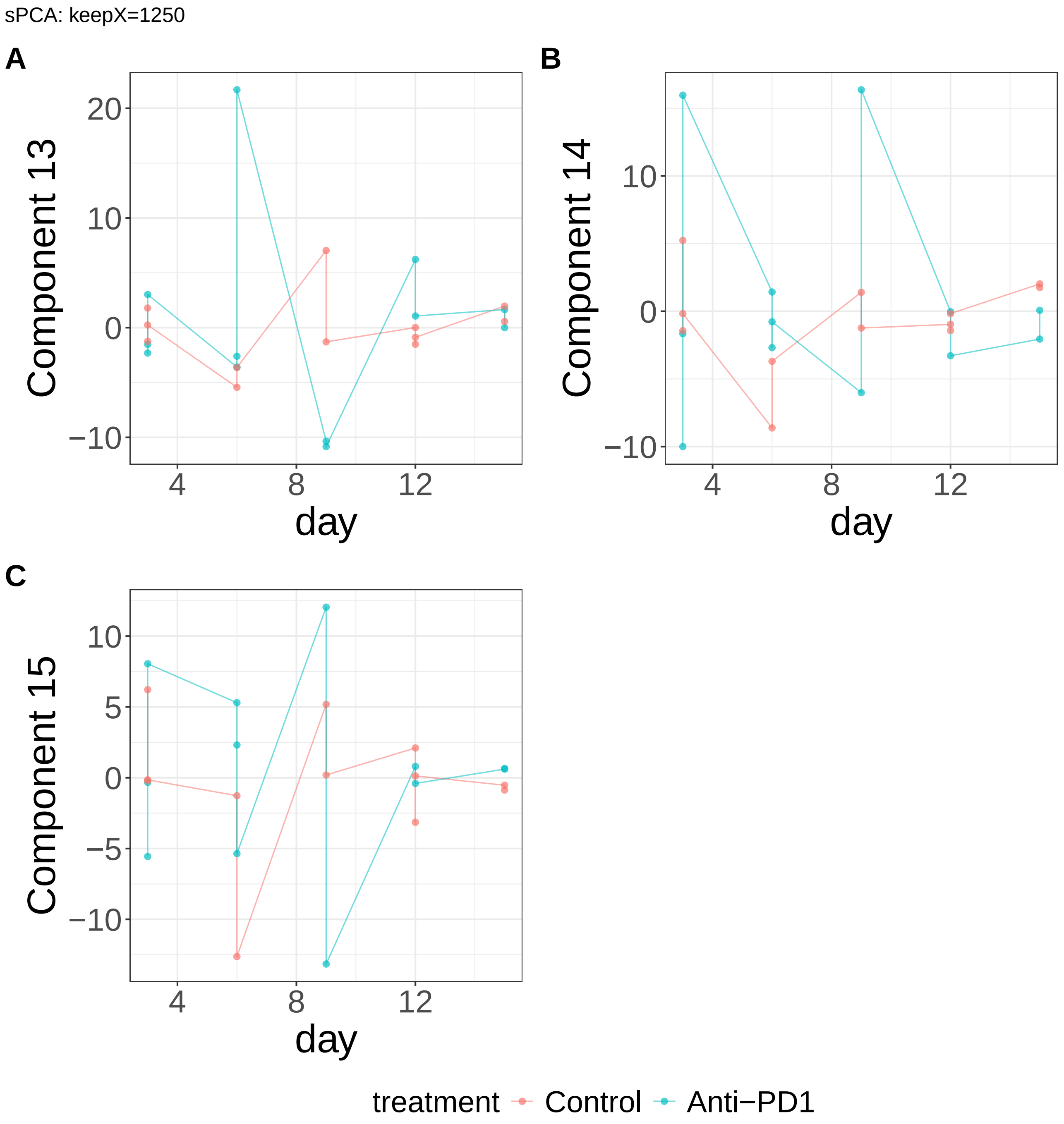


**Supplementary Figure 457.** Components for sPCA: keepX=1250. (A) Component 13. (B) Component 14. (C) Component 15.

sPCA: keepX=2500

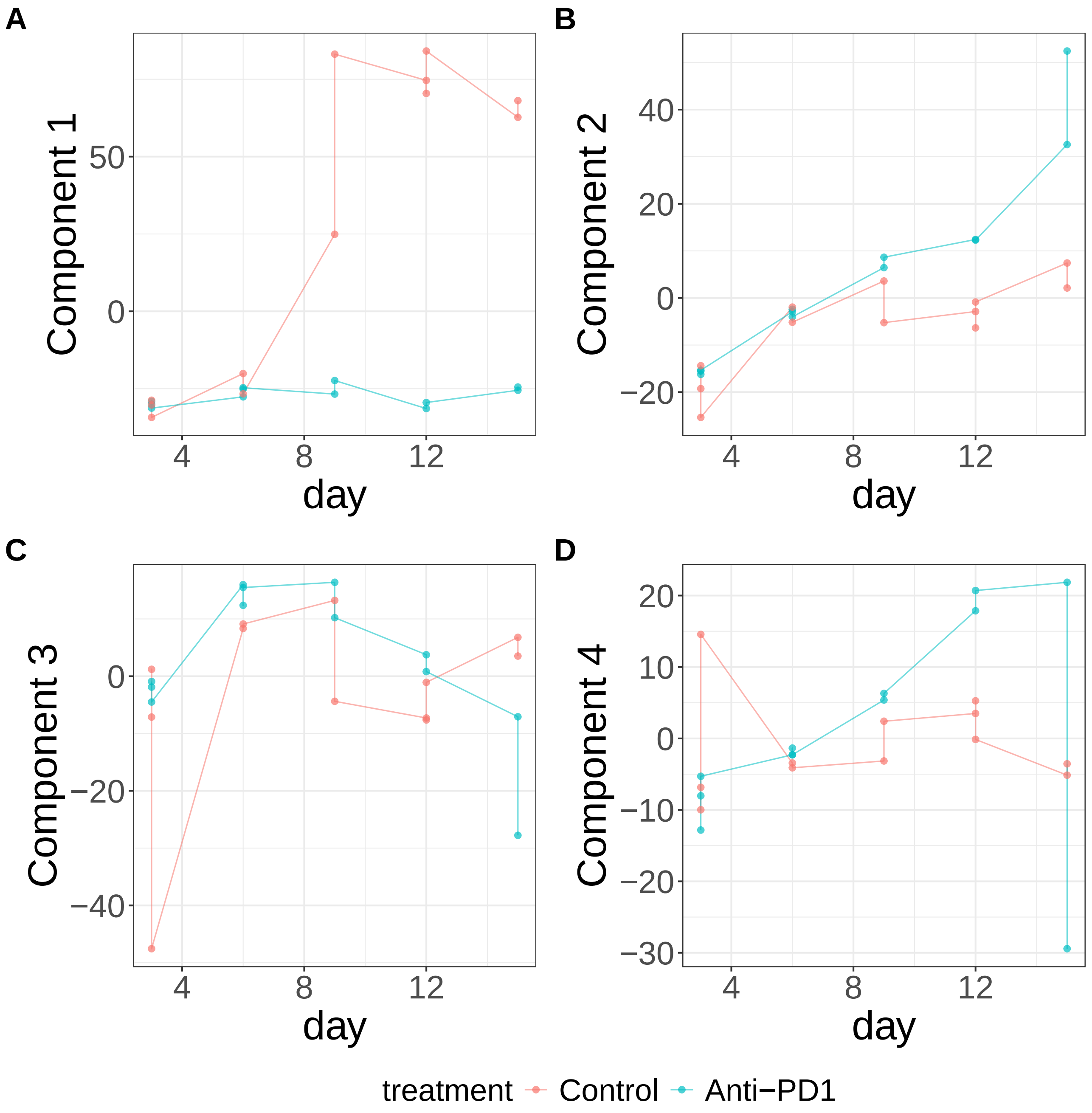


**Supplementary Figure 458.** Components for sPCA: keepX=2500. (A) Component 1. (B) Component 2. (C) Component 3. (D) Component 4.

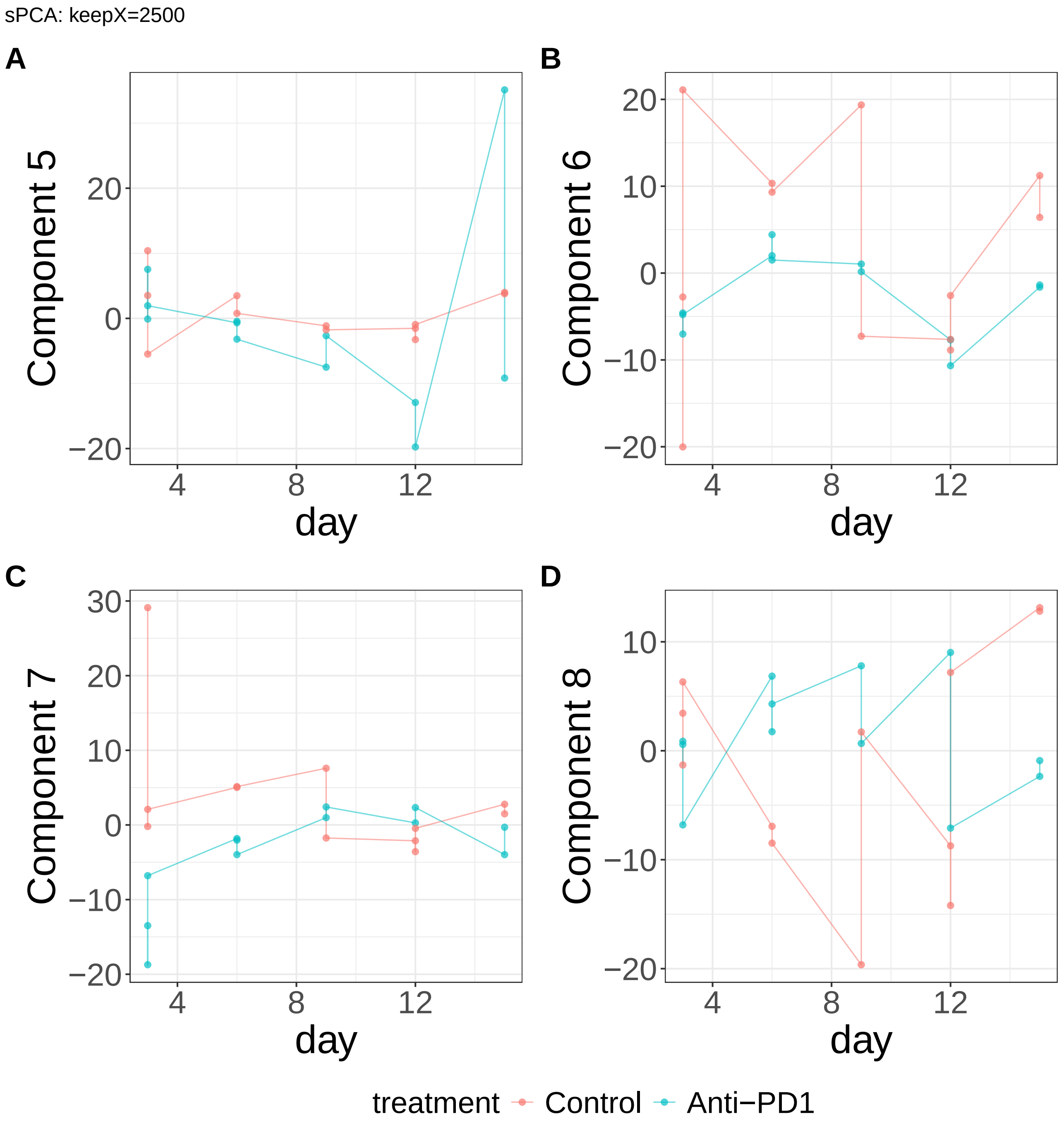


**Supplementary Figure 459.** Components for sPCA: keepX=2500. (A) Component 5. (B) Component 6. (C) Component 7. (D) Component 8.

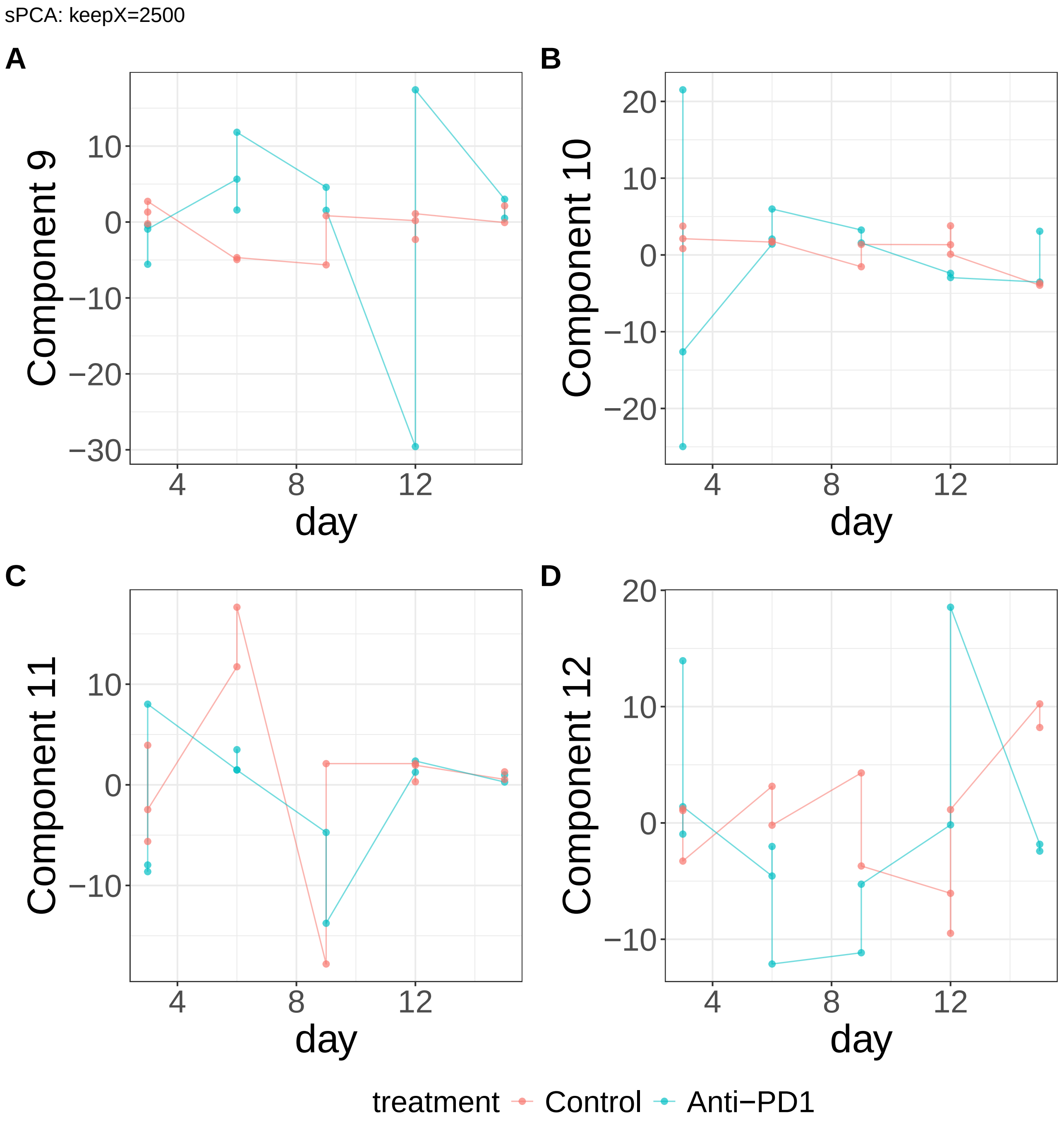


**Supplementary Figure 460.** Components for sPCA: keepX=2500. (A) Component 9. (B) Component 10. (C) Component 11. (D) Component 12.

sPCA: keepX=2500

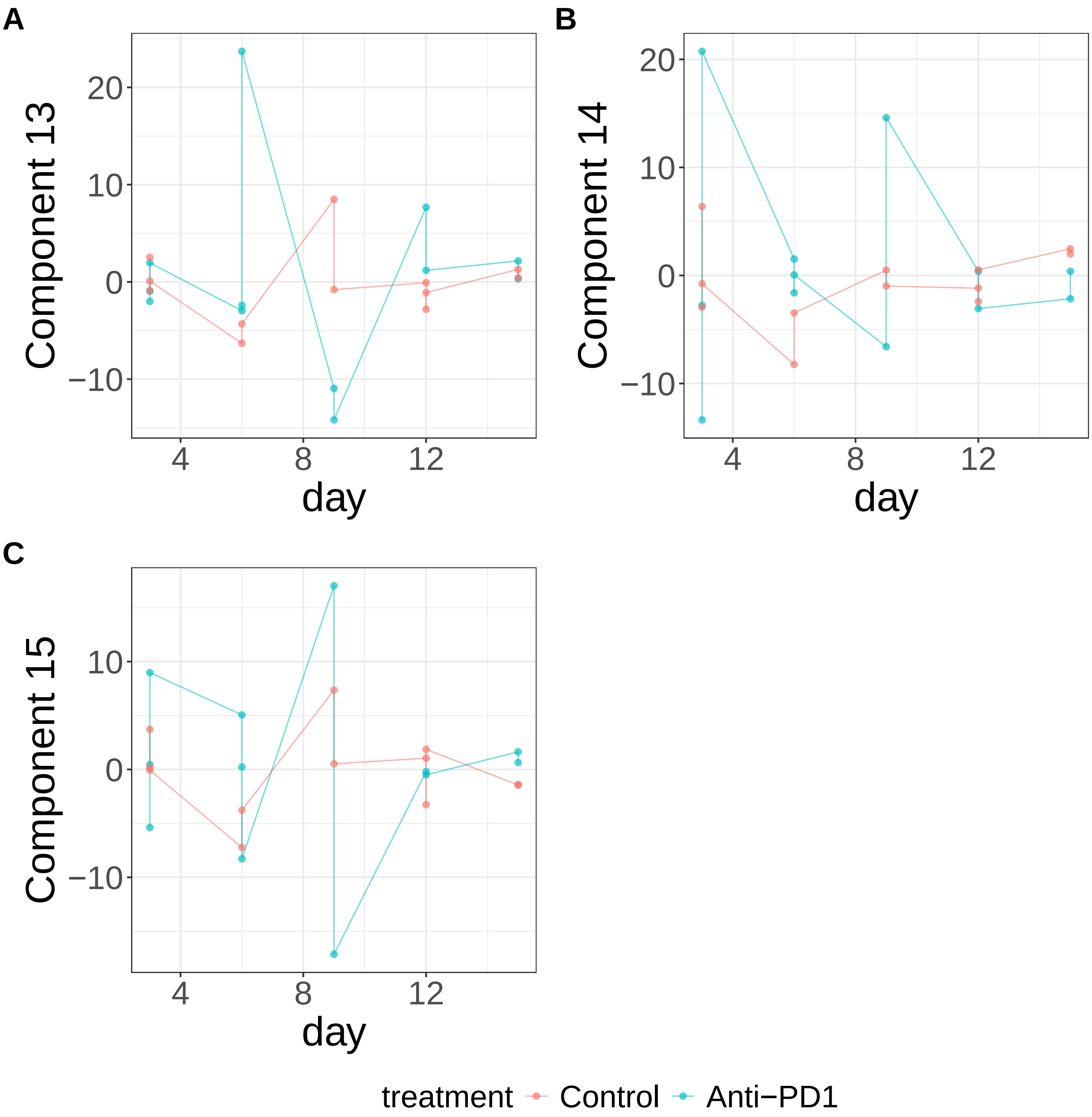


**Supplementary Figure 461.** Components for sPCA: keepX=2500. (A) Component 13. (B) Component 14. (C) Component 15.

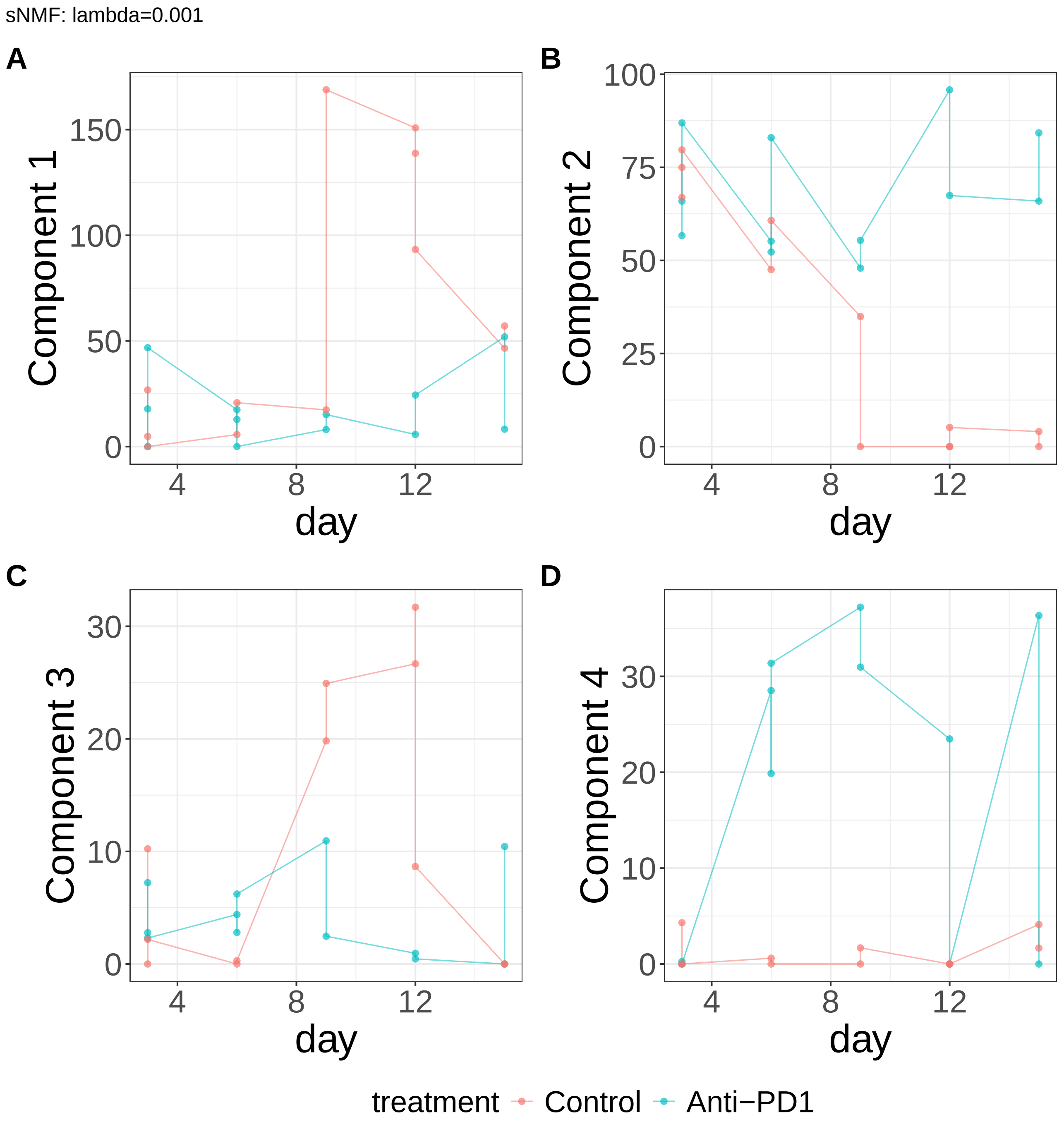


**Supplementary Figure 462.** Components for sNMF: lambda=0.001. (A) Component 1. (B) Component 2. (C) Component 3. (D) Component 4.

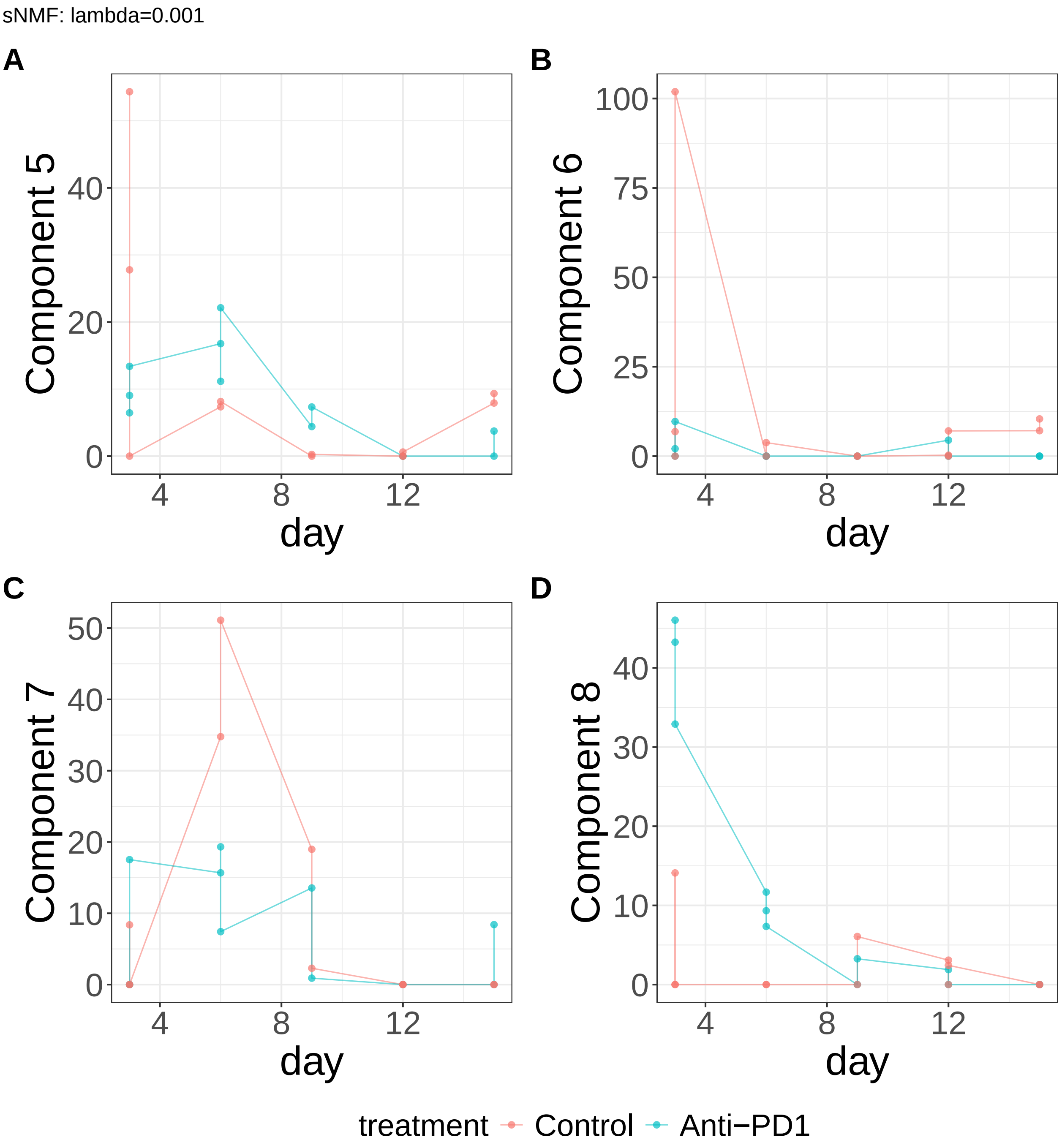


**Supplementary Figure 463.** Components for sNMF: lambda=0.001. (A) Component 5. (B) Component 6. (C) Component 7. (D) Component 8.

sNMF: lambda=0.001

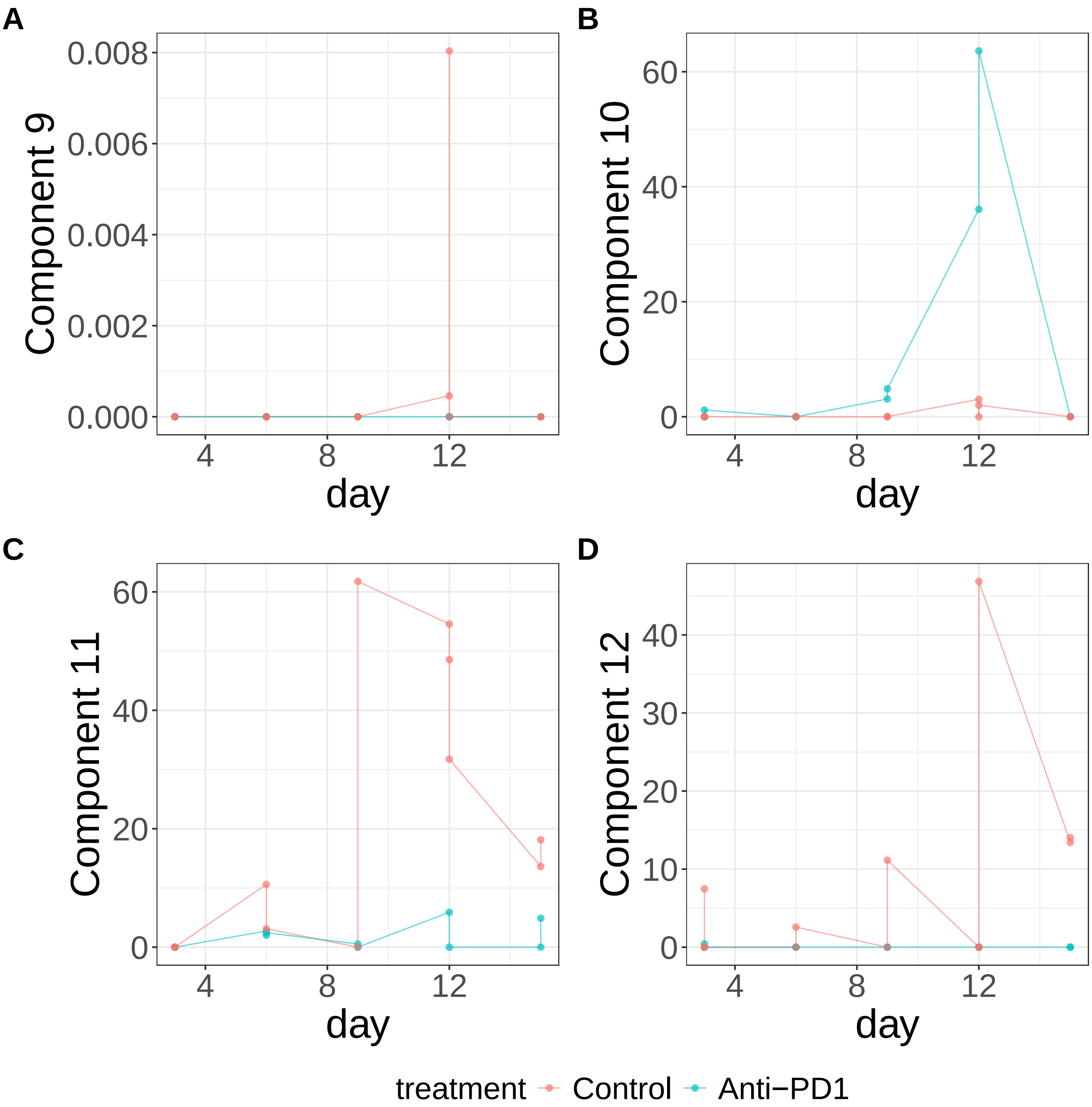


**Supplementary Figure 464.** Components for sNMF: lambda=0.001. (A) Component 9. (B) Component 10. (C) Component 11. (D) Component 12.

sNMF: lambda=0.001

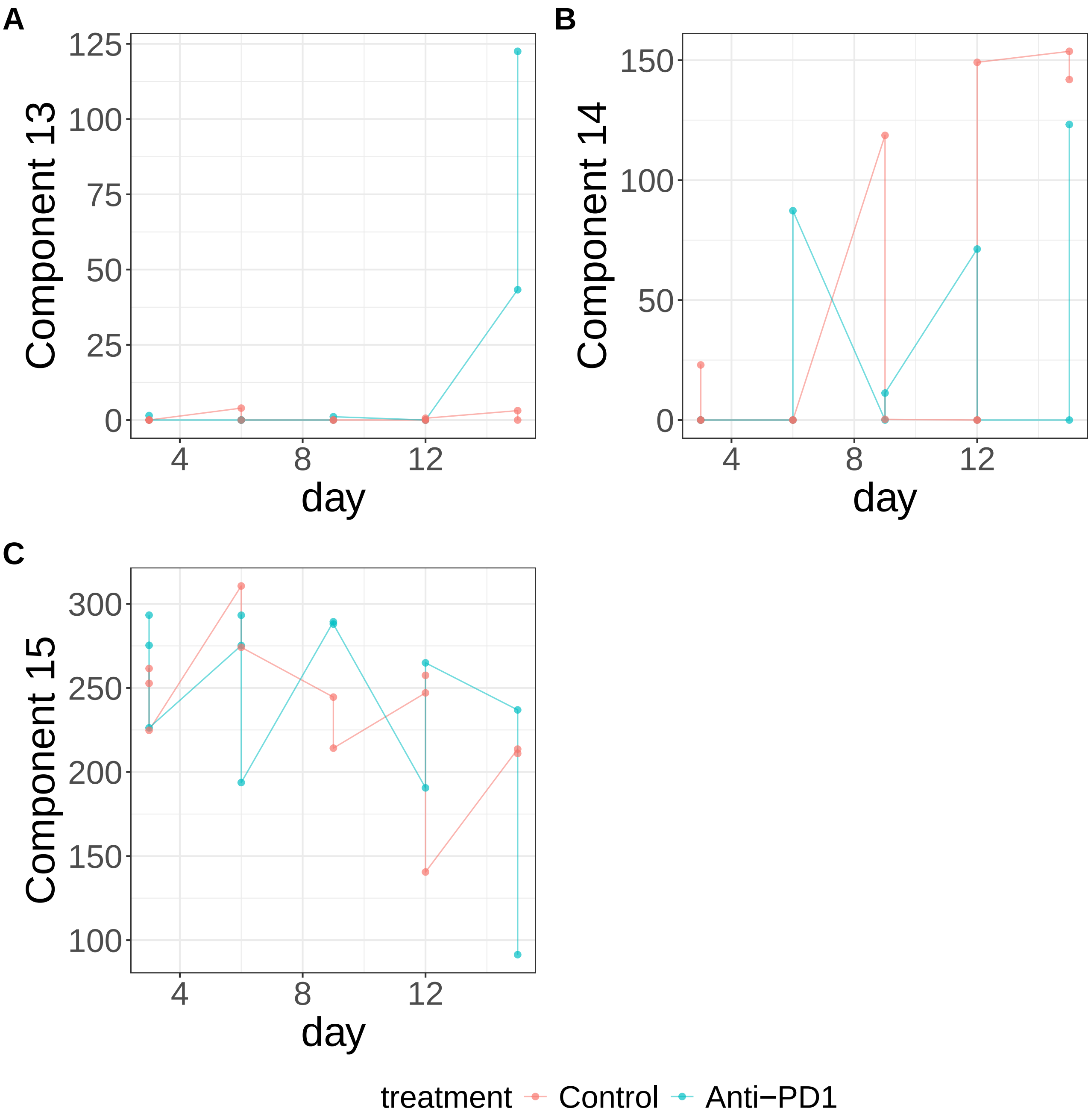


**Supplementary Figure 465.** Components for sNMF: lambda=0.001. (A) Component 13. (B) Component 14. (C) Component 15.

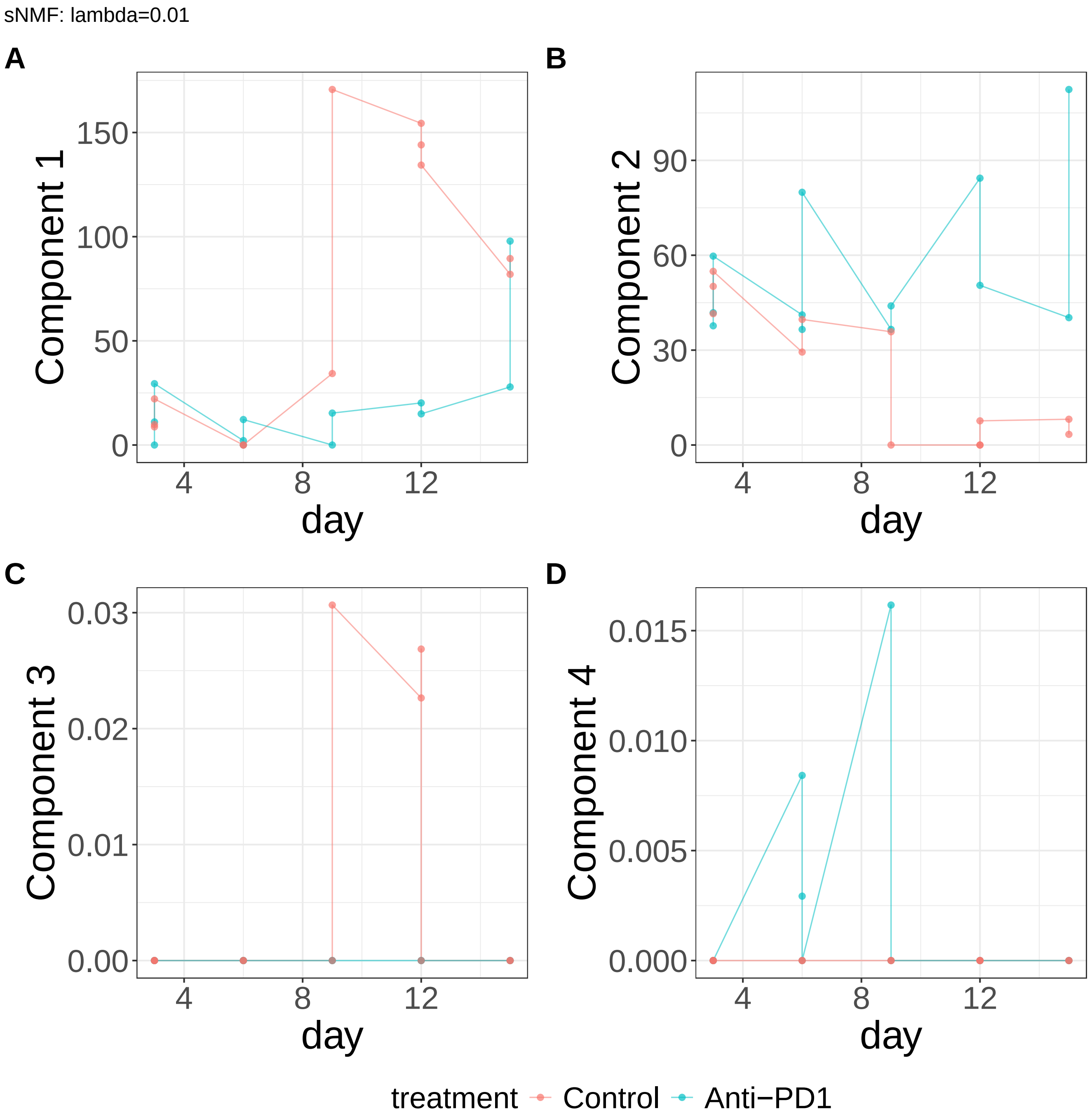


**Supplementary Figure 466.** Components for sNMF: lambda=0.01. (A) Component 1. (B) Component 2. (C) Component 3. (D) Component 4.

sNMF: lambda=0.01

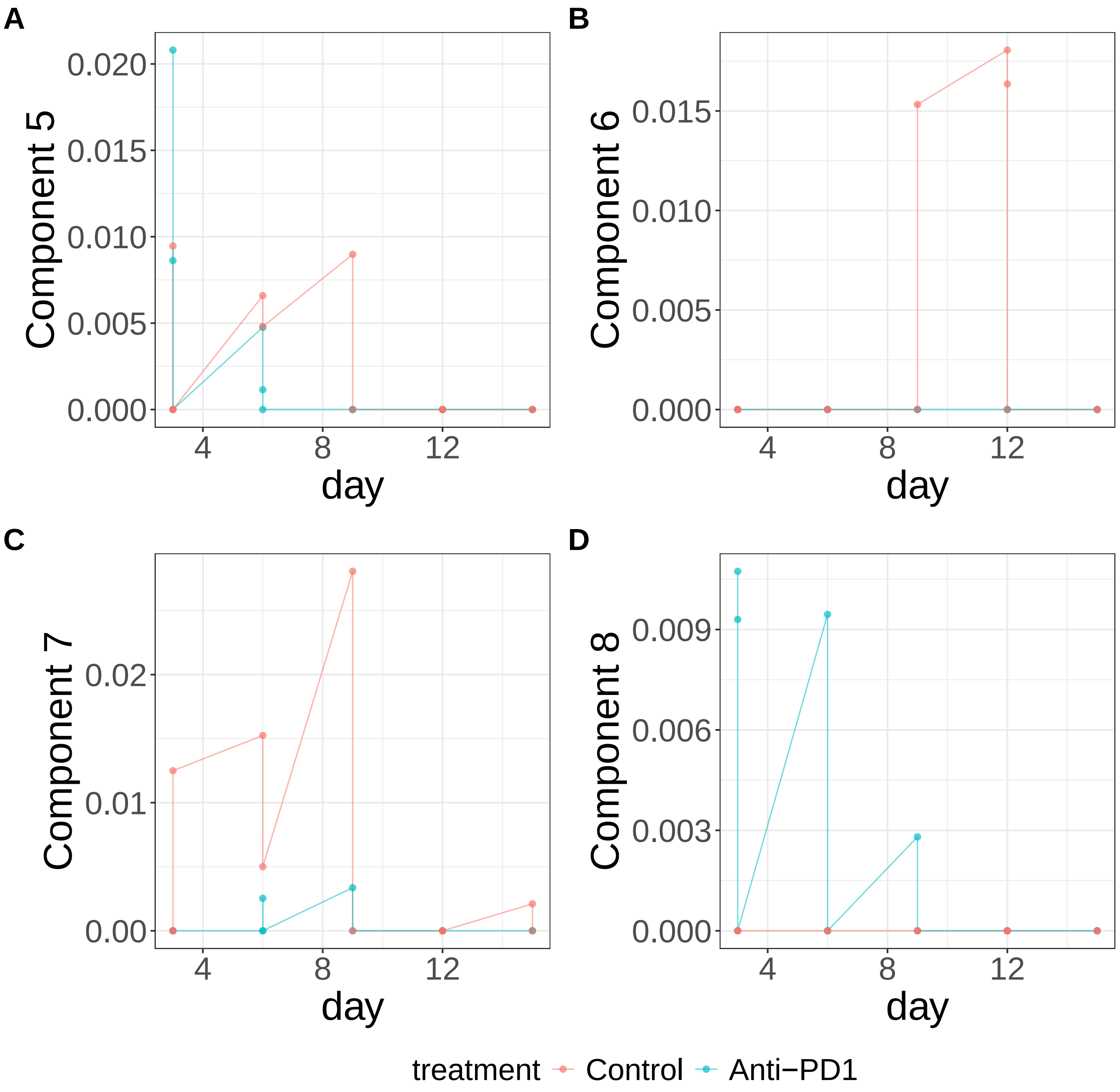


**Supplementary Figure 467.** Components for sNMF: lambda=0.01. (A) Component 5. (B) Component 6. (C) Component 7. (D) Component 8.

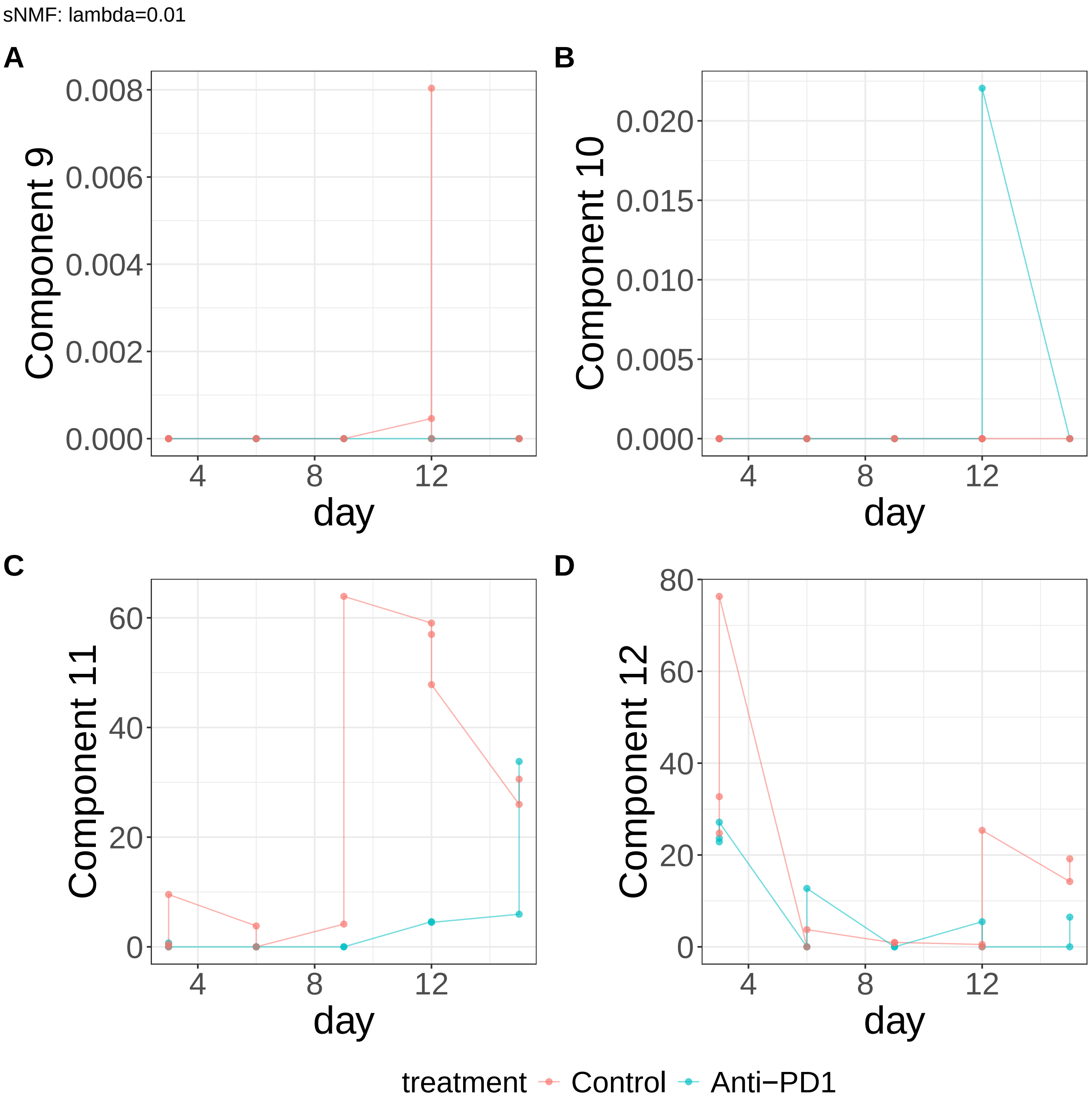


**Supplementary Figure 468.** Components for sNMF: lambda=0.01. (A) Component 9. (B) Component 10. (C) Component 11. (D) Component 12.

sNMF: lambda=0.01

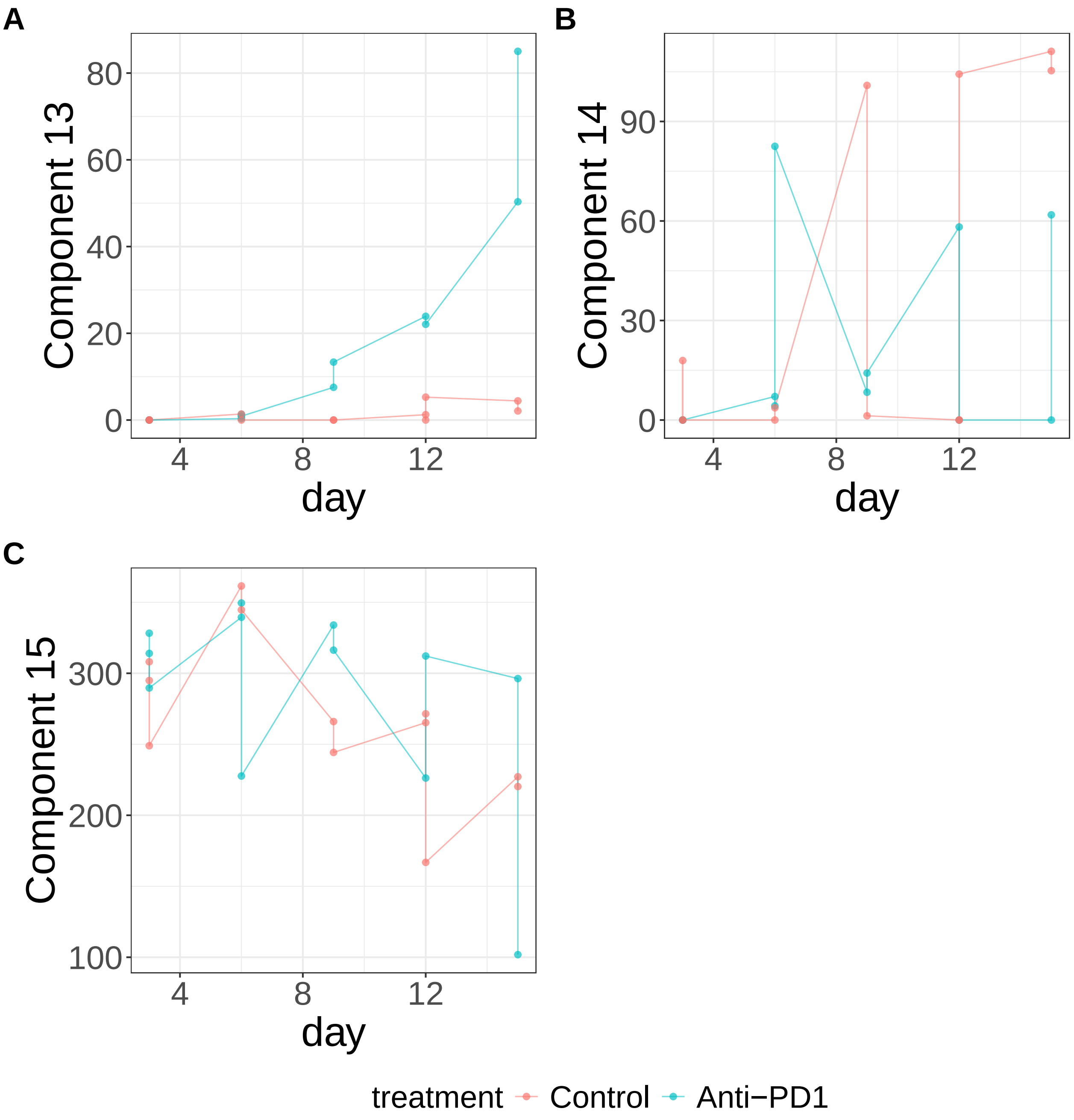


**Supplementary Figure 469.** Components for sNMF: lambda=0.01. (A) Component 13. (B) Component 14. (C) Component 15.

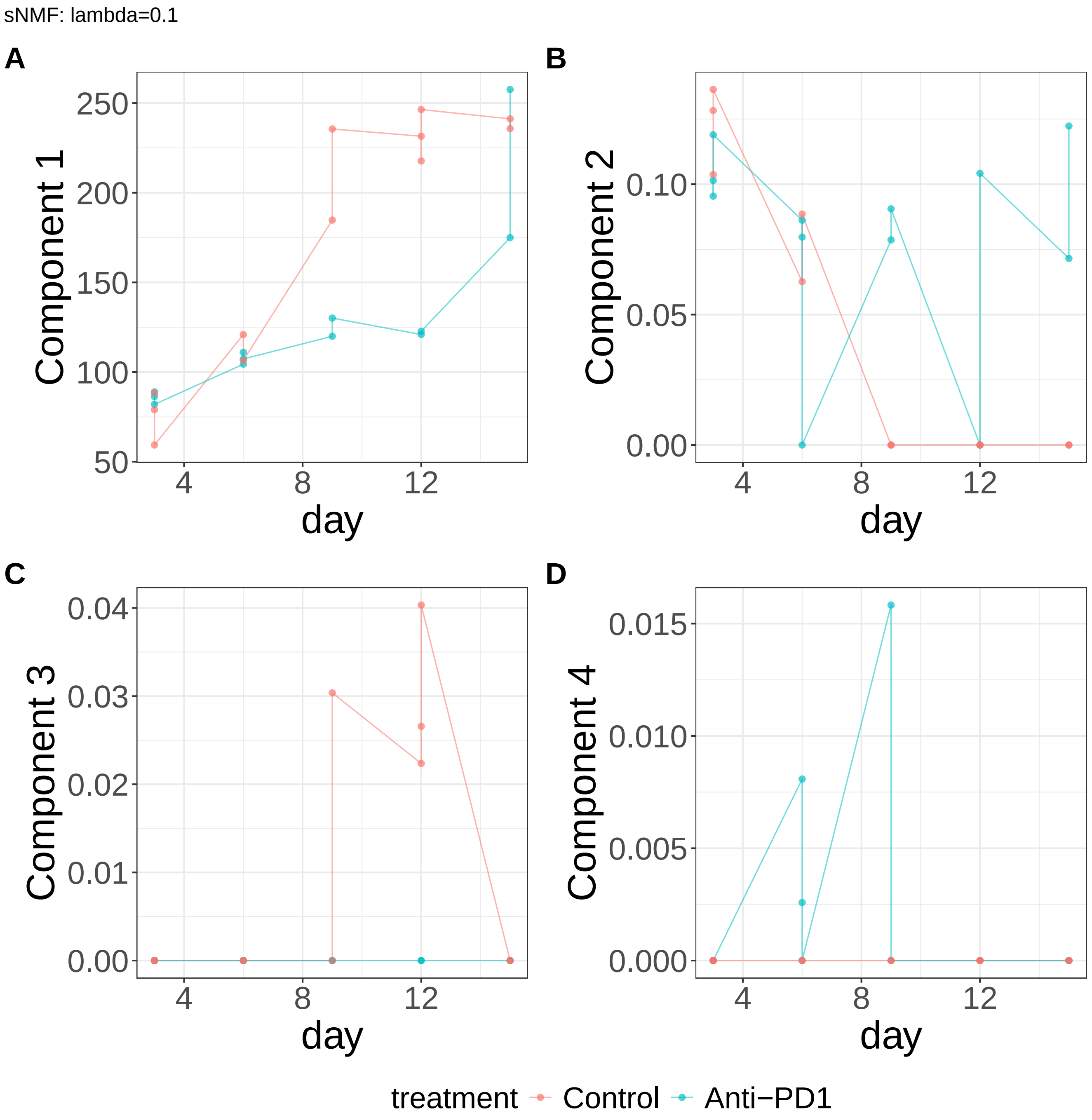


**Supplementary Figure 470.** Components for sNMF: lambda=0.1. (A) Component 1. (B) Component 2. (C) Component 3. (D) Component 4.

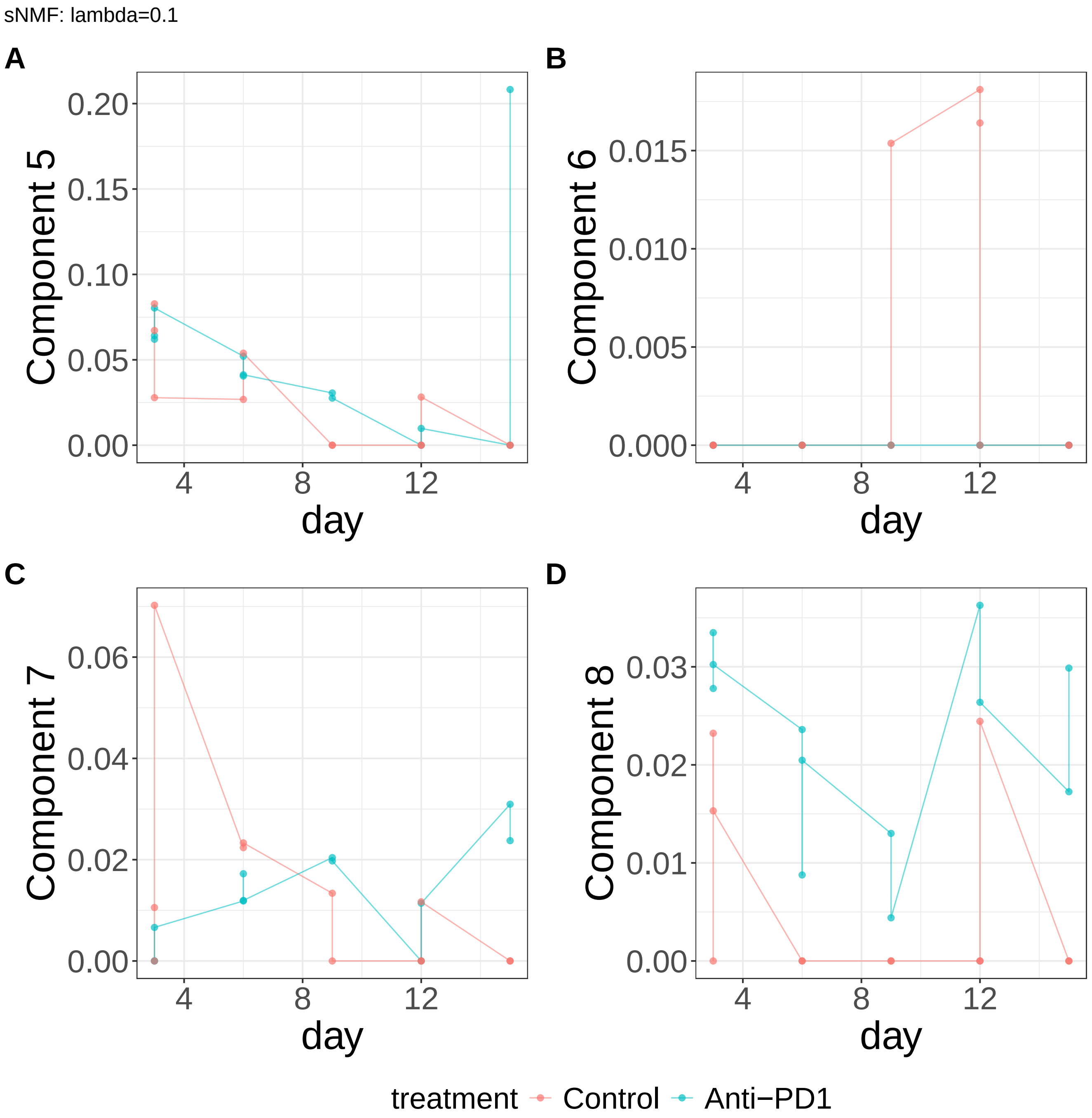


**Supplementary Figure 471.** Components for sNMF: lambda=0.1. (A) Component 5. (B) Component 6. (C) Component 7. (D) Component 8.

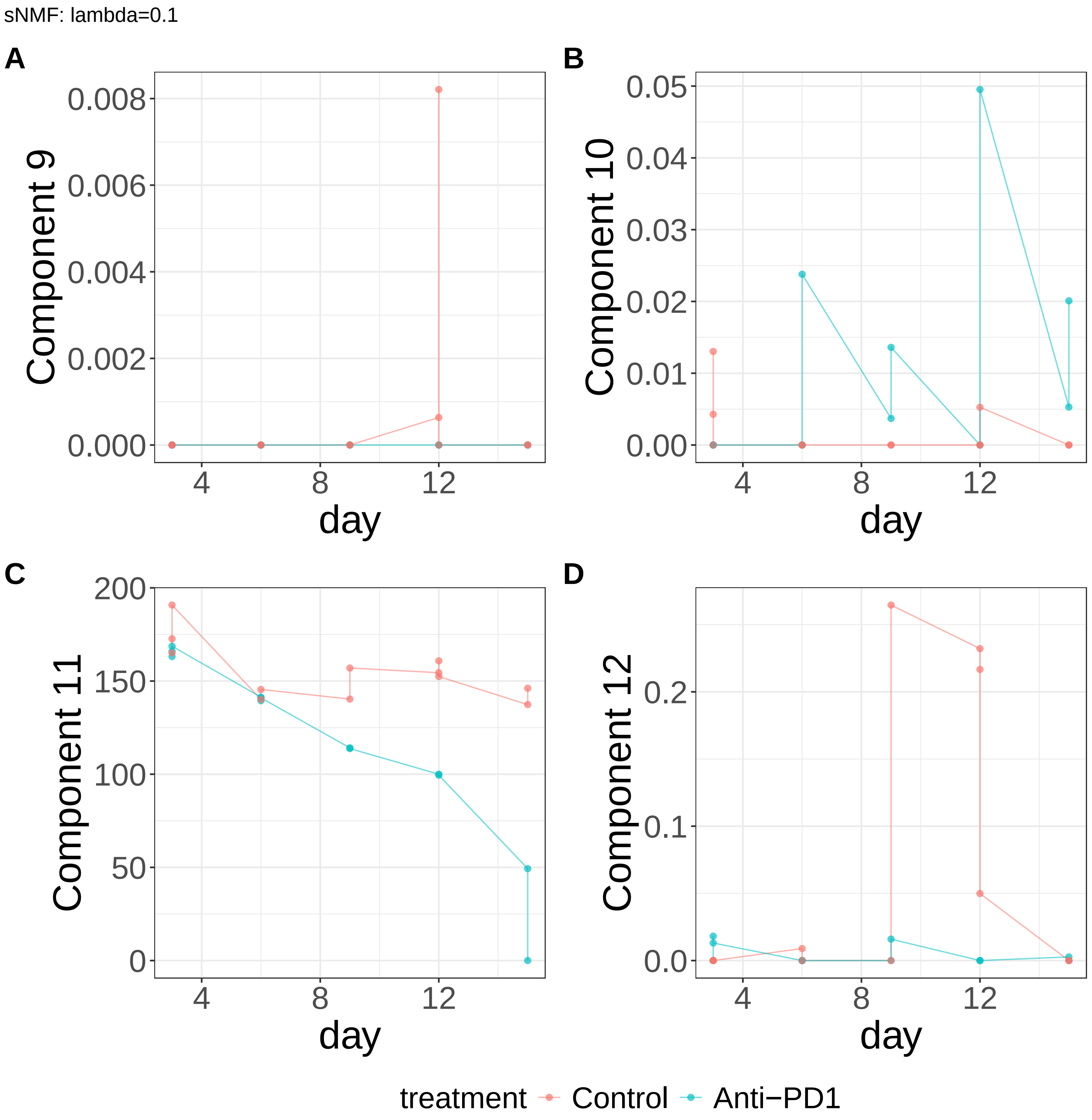


**Supplementary Figure 472.** Components for sNMF: lambda=0.1. (A) Component 9. (B) Component 10. (C) Component 11. (D) Component 12.

sNMF: lambda=0.1

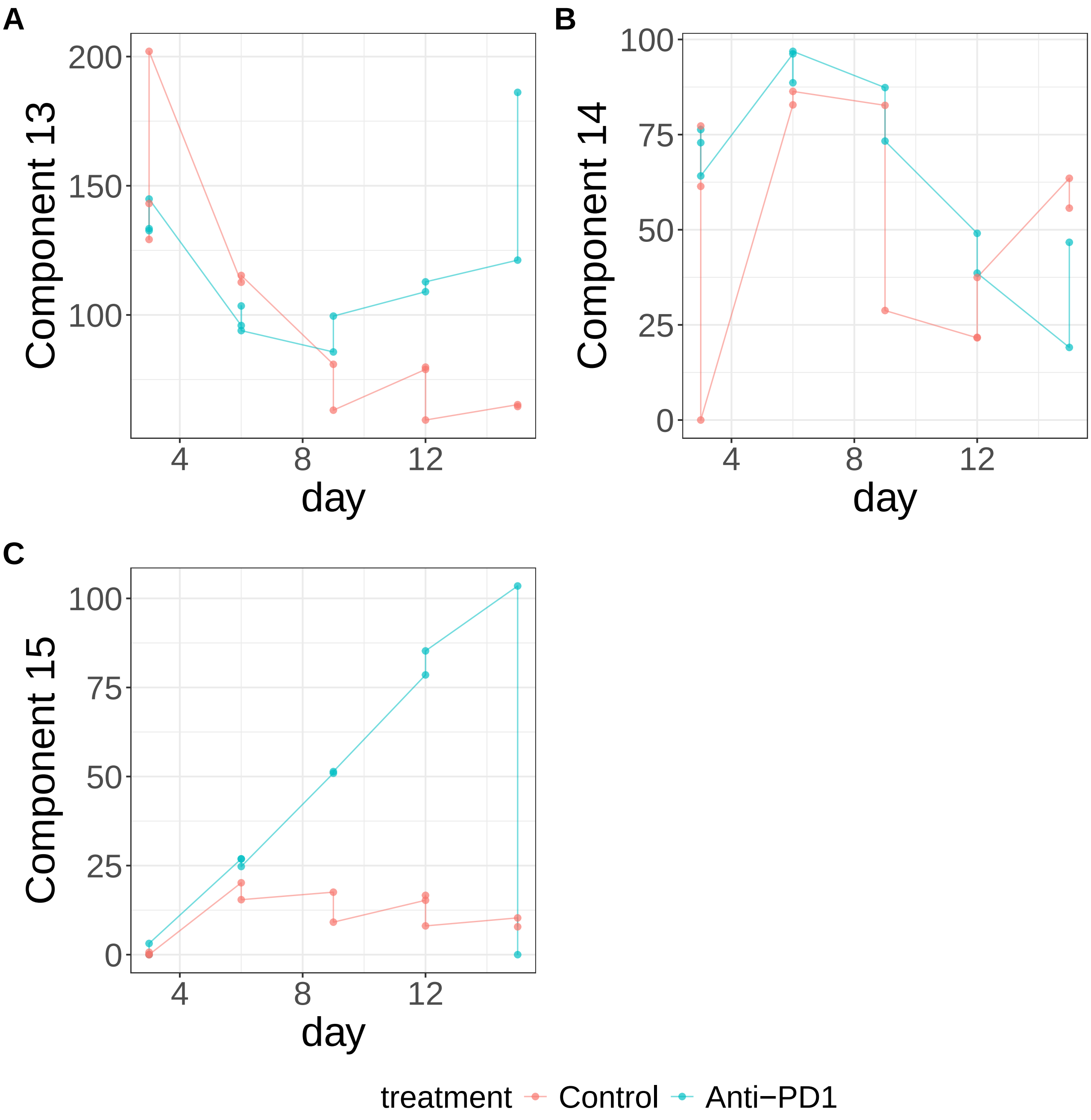


**Supplementary Figure 473.** Components for sNMF: lambda=0.1. (A) Component 13. (B) Component 14. (C) Component 15.

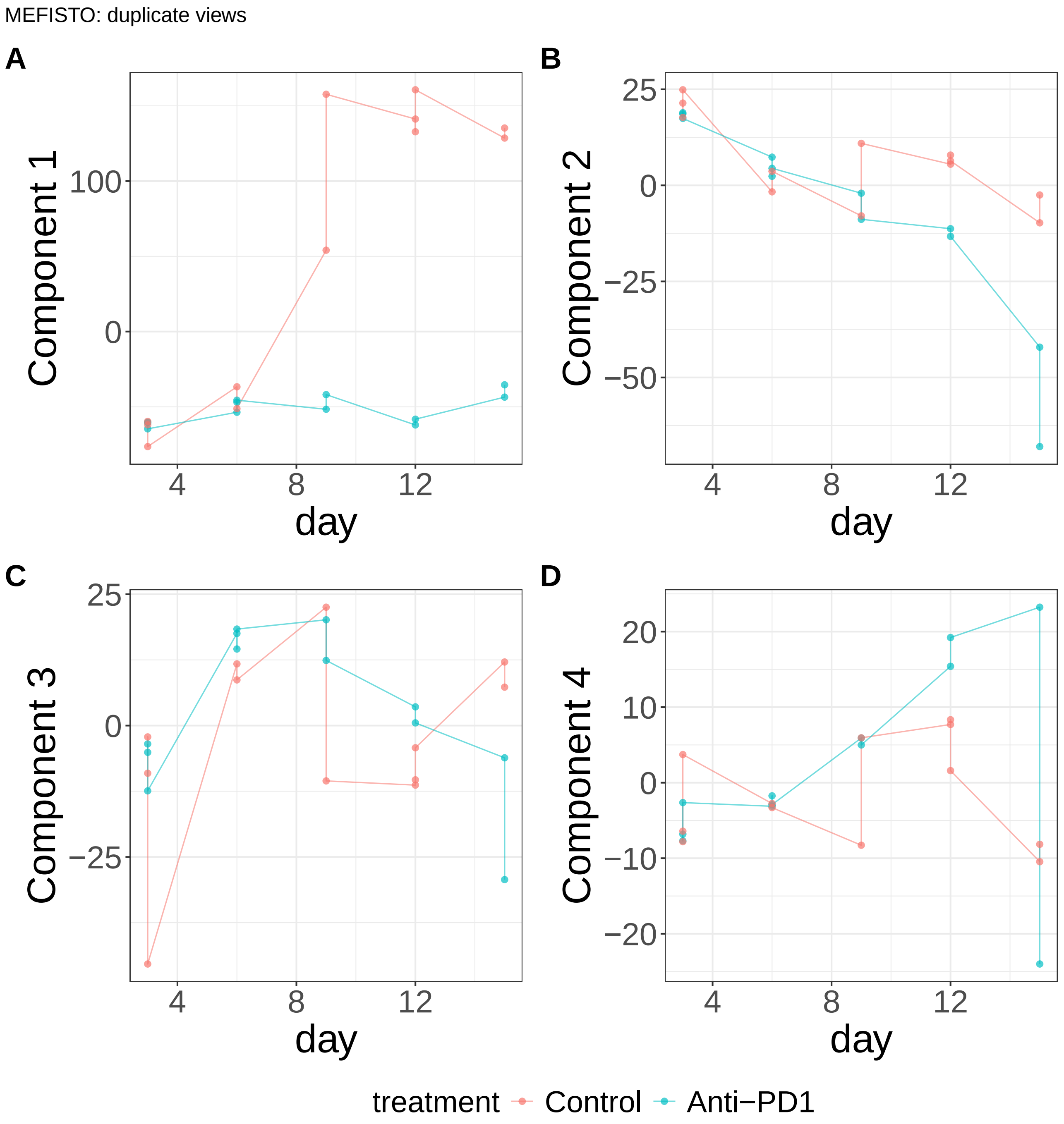


**Supplementary Figure 474.** Components for MEFISTO: duplicate views. (A) Component 1. (B) Component 2. (C) Component 3. (D) Component 4.

MEFISTO: duplicate views

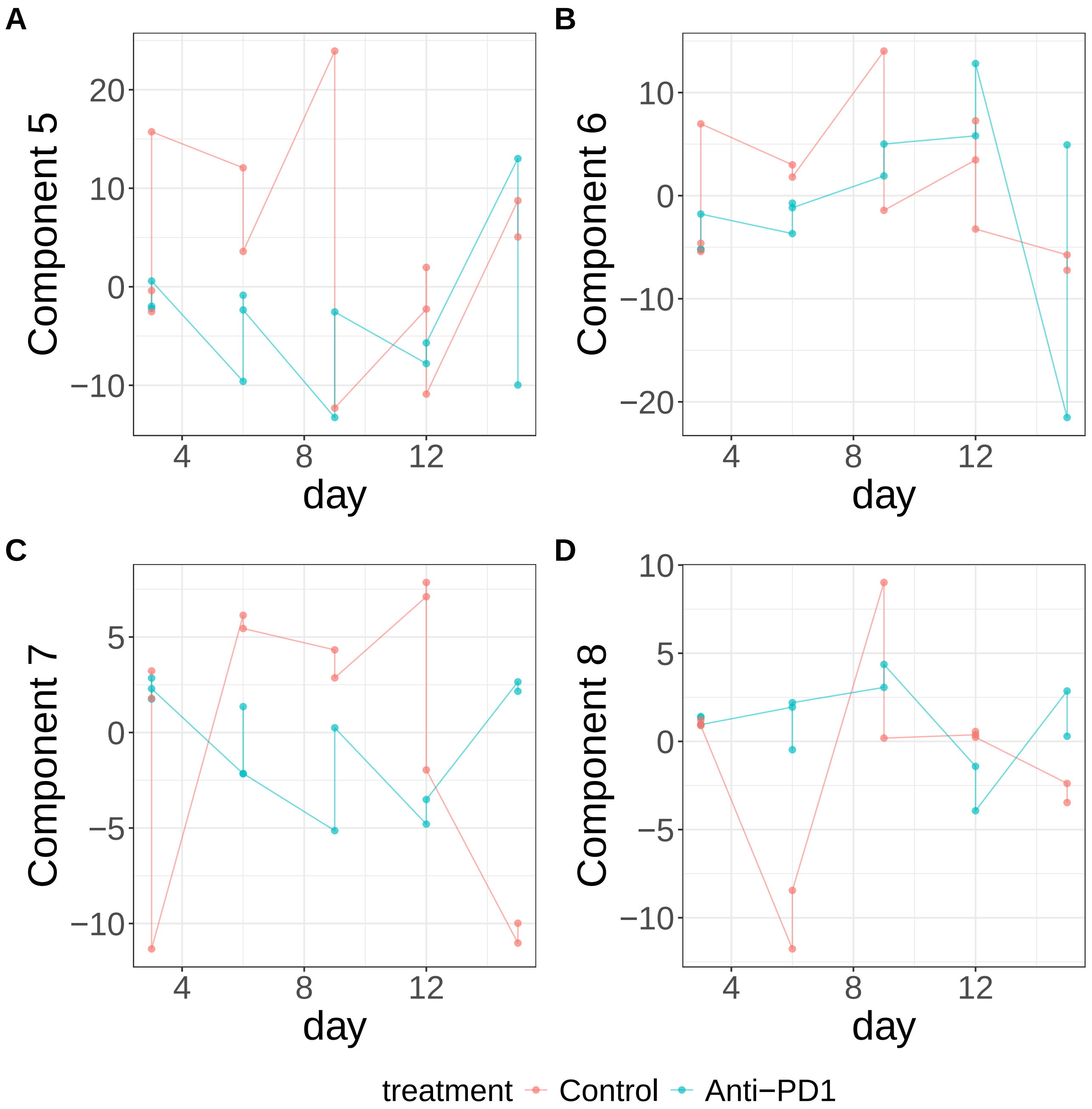


**Supplementary Figure 475.** Components for MEFISTO: duplicate views. (A) Component 5. (B) Component 6. (C) Component 7. (D) Component 8.

MEFISTO: duplicate views

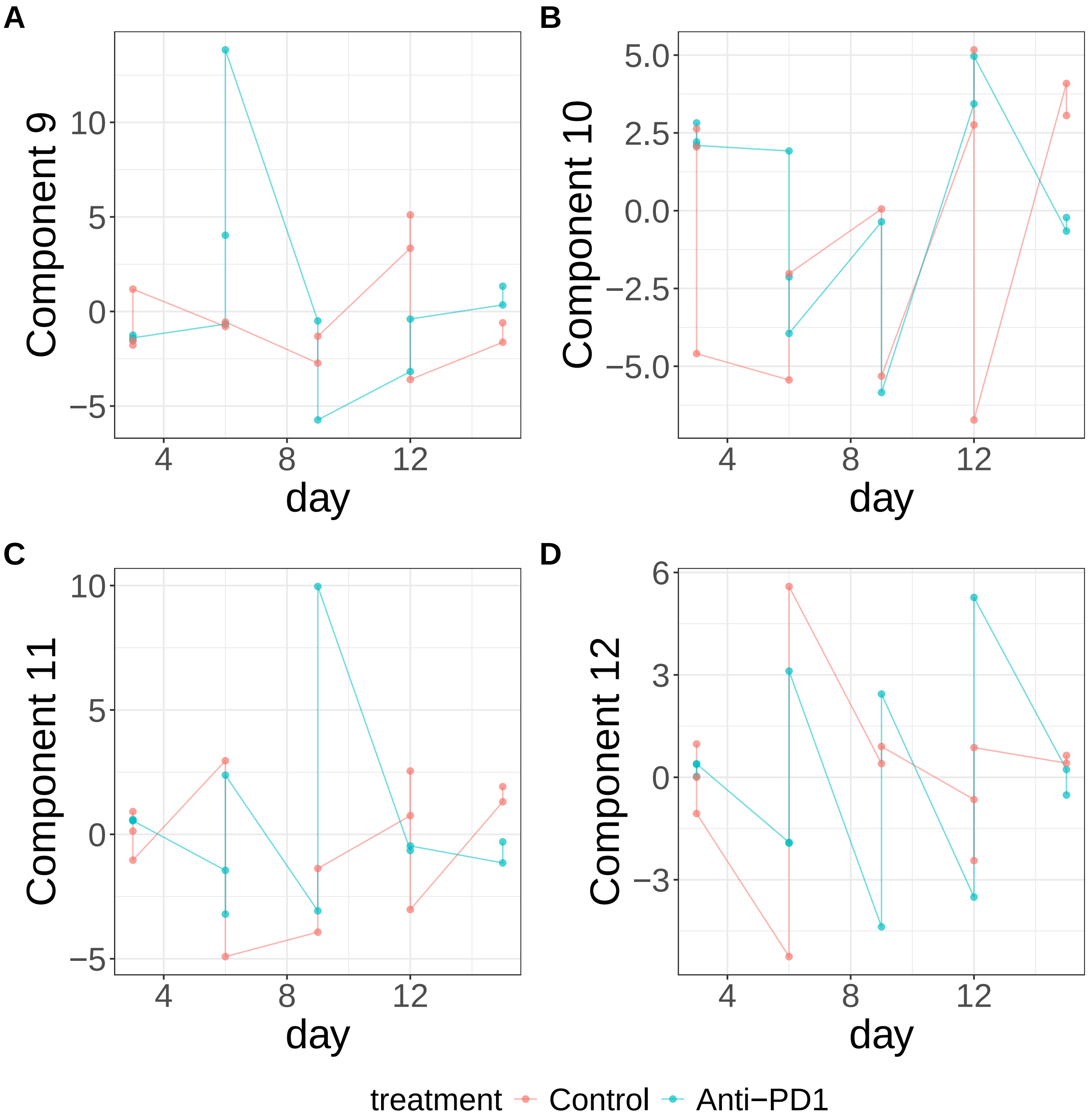


**Supplementary Figure 476.** Components for MEFISTO: duplicate views. (A) Component 9. (B) Component 10. (C) Component 11. (D) Component 12.

MEFISTO: duplicate views

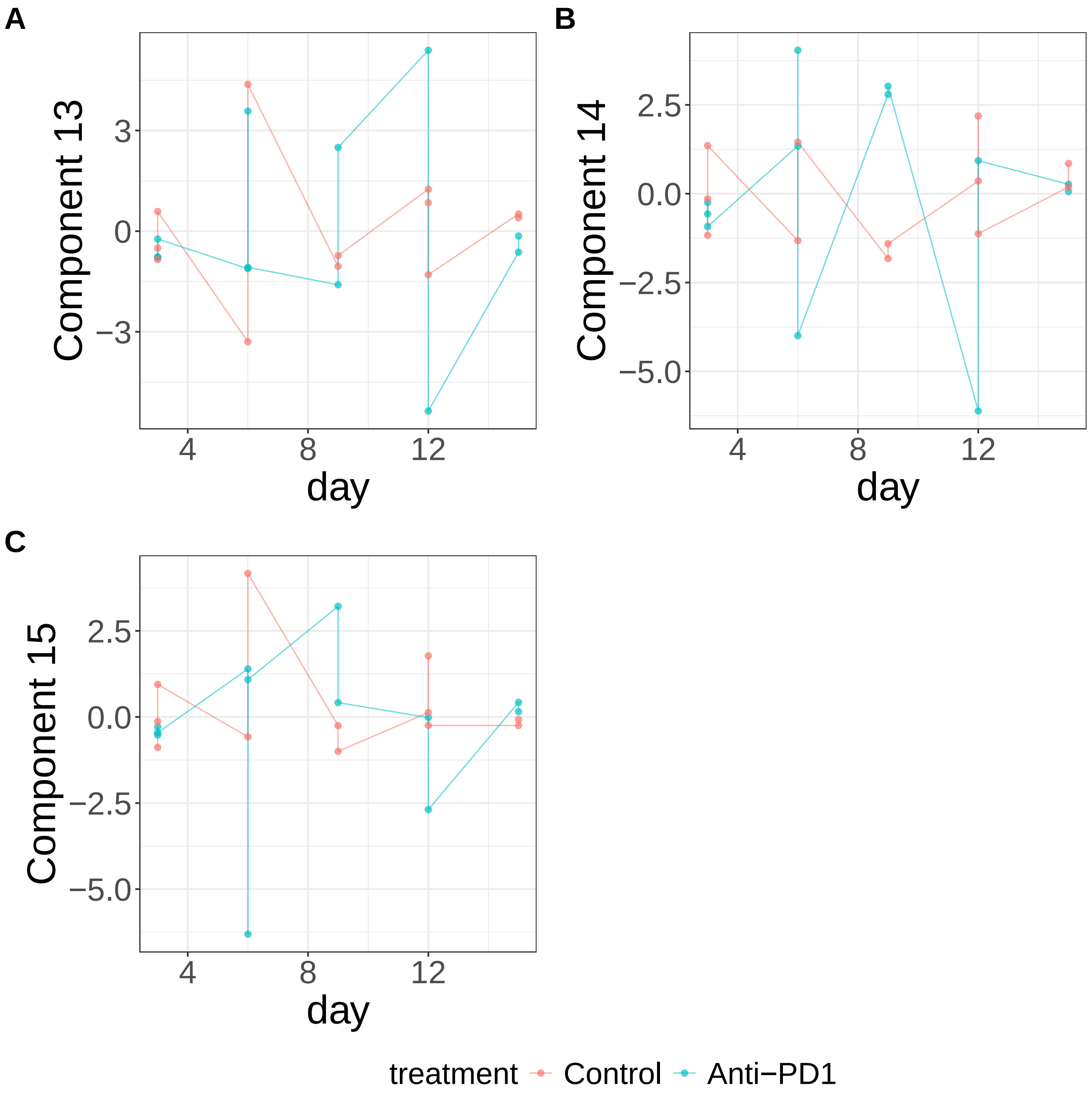


**Supplementary Figure 477.** Components for MEFISTO: duplicate views. (A) Component 13. (B) Component 14. (C) Component 15.
