## Supplementary Note 7 : Global dengue for "A statistical framework for system-level synthesis"

### Seasonal and climate-associated structure in global dengue incidence

Global dengue incidence exhibits complex temporal patterns that have been linked to large-scale climate variability, including Indian Ocean sea surface temperature anomalies represented by the Indian Ocean Basin-Wide (IOBW) index[1]. Given the small sample size (72 standardized monthly dengue incidence observations spanning 6 years), we restricted the analysis to learning and interpreting the system-level model and did not perform conditional inference.

We modeled the system-level processes as smooth functions of month, IOBW, and their interaction (**Methods**, **Supplementary Figs. 329-425**). Despite imposing no periodicity, decomposition of the first two dominant processes with respect to month revealed the characteristic dengue seasons of the Northern and Southern Hemispheres, demonstrating that SSANOVA-SVD reconstructed known epidemiological seasonality without prespecifying its form (**Supplementary Figs. 329**A–B, 330). Decompositions with respect to IOBW revealed two distinct climate-associated patterns(**Supplementary Fig. 329**A-B). The first exhibited peak incidence at higher IOBW values, while the second showed a monotonic increase across the observed IOBW range. Although these climate-associated patterns were consistent with representative country-level trajectories (e.g., Mexico and Colombia)(**Supplementary Fig. 329**C-D), their epidemiological interpretation remains unclear. Additional data and more informative covariates could help disentangle these effects. Beyond component 3, the fitted scores were small, and given the small sample size with several outliers, likely a result of overfitting (**Supplementary Figs. 331-337, 359-382**).

Collectively, these results demonstrate that SSANOVA-SVD reconstructs coherent system-level processes underlying dengue seasonality while revealing climate-associated structure that warrants further investigation. In contrast, conventional matrix factorization did not provide comparably interpretable representations of the seasonal and climate-associated structure(**Supplementary Figs. 383-425**).

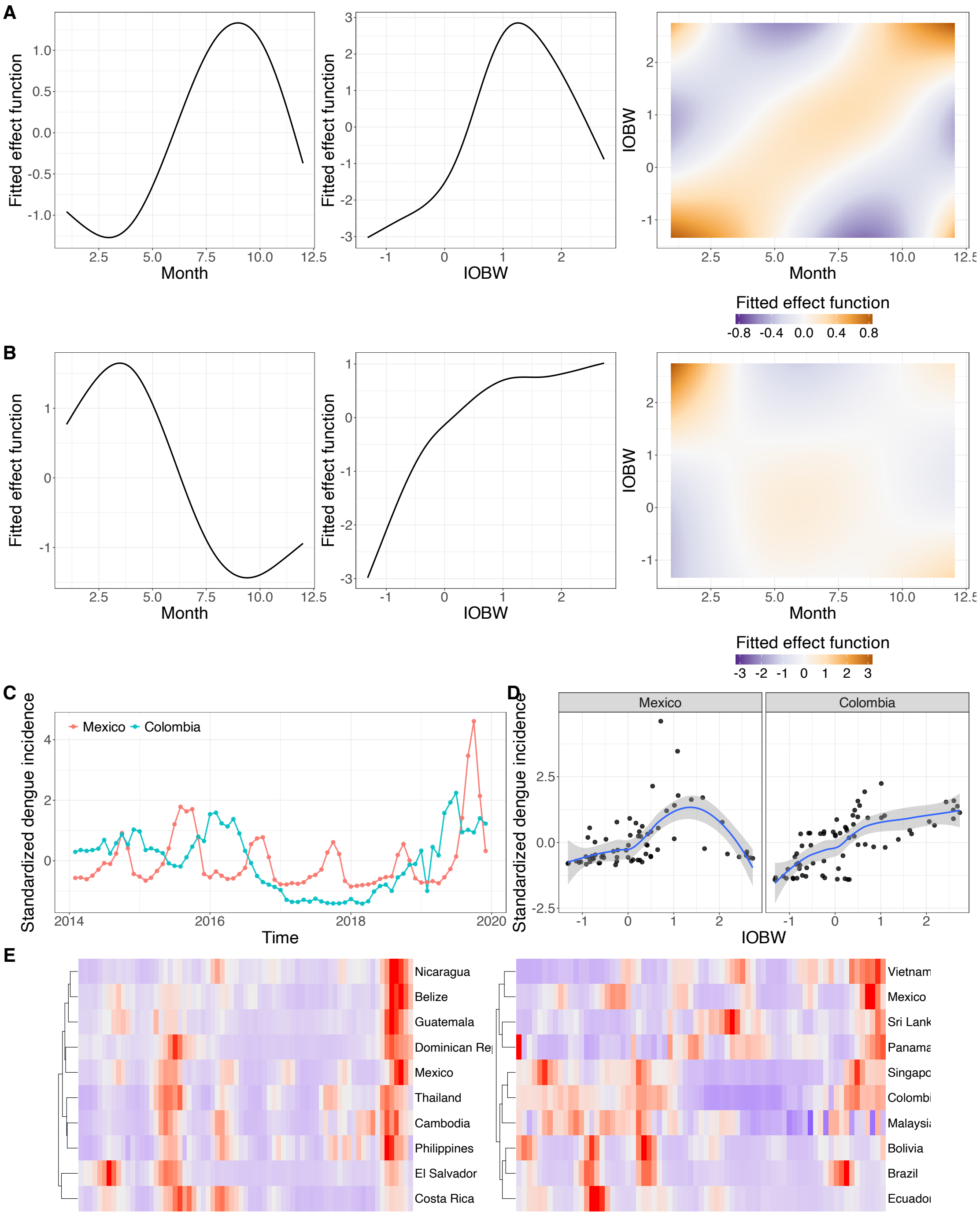


**Supplementary Figure 329: System-level synthesis of global dengue incidence.** A. Decomposition of the first reconstructed component function by SSANOVA-SVD into the fitted effects of month, Indian Ocean Basin-Wide (IOBW) precipitation, and their interaction. B. Corresponding decomposition of the second reconstructed component function into the fitted effects of month, IOBW precipitation, and their interaction. C. Standardized dengue incidence over time for two representative countries, Mexico and Colombia. D. Standardized dengue incidence as a function of IOBW precipitation for Mexico and Colombia. Curves and shaded regions represent LOESS smooths and their standard errors, respectively. E. Heatmaps of standardized dengue incidence for the countries with the largest absolute loadings on reconstructed components 1 (left) and 2 (right).

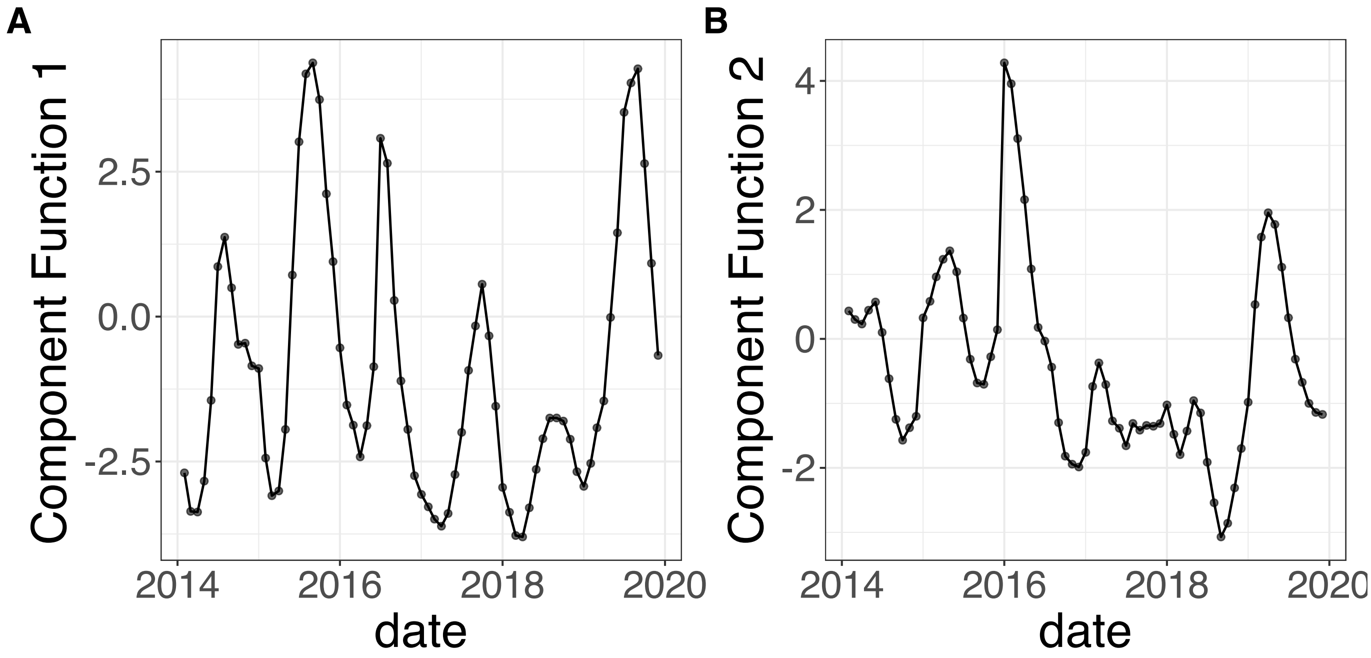


**Supplementary Figure 330.** SSANOVA-SVD reconstructed component functions.(A) Component 1. (B) Component 2.

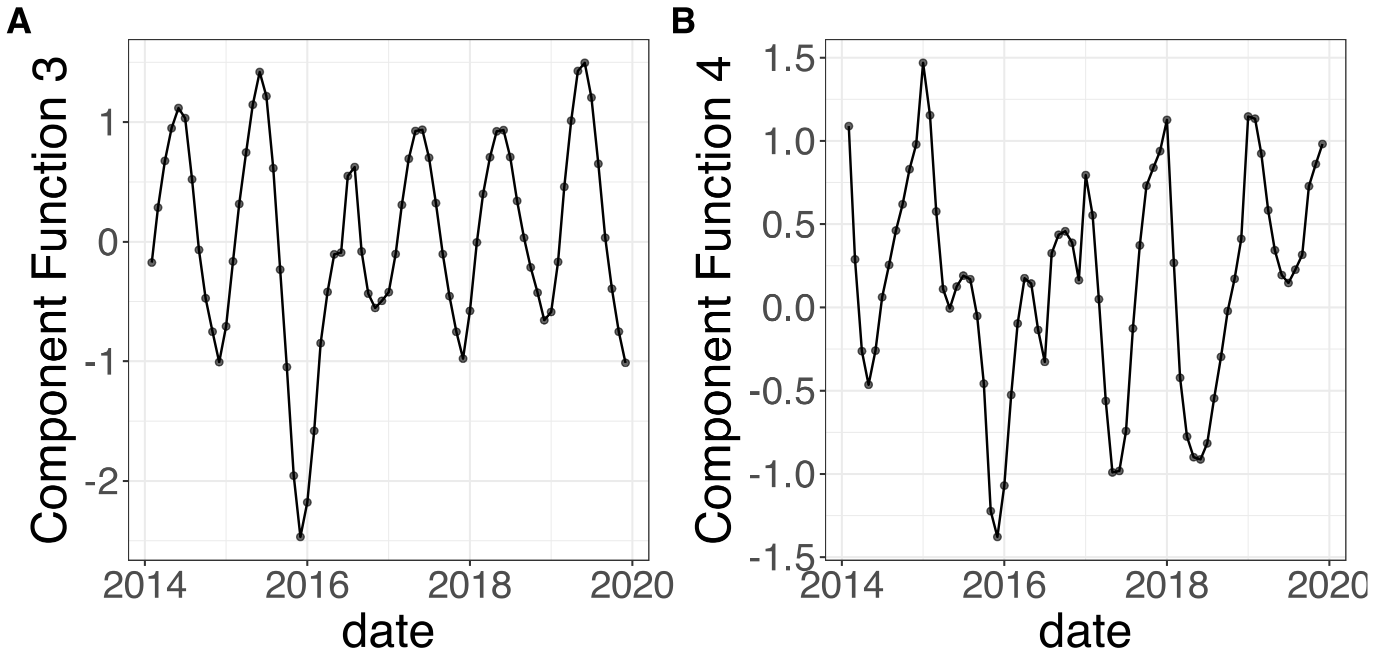


**Supplementary Figure 331.** SSANOVA-SVD reconstructed component functions.(A) Component 3. (B) Component 4.

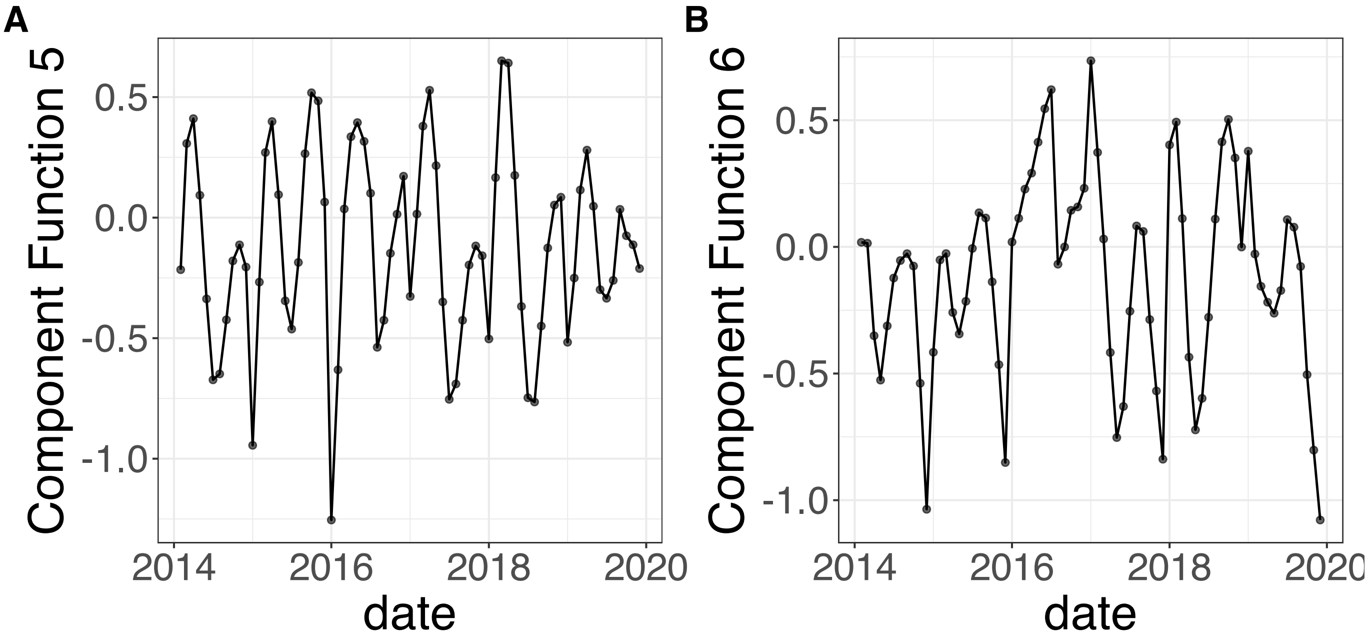


**Supplementary Figure 332.** SSANOVA-SVD reconstructed component functions.(A) Component 5. (B) Component 6.

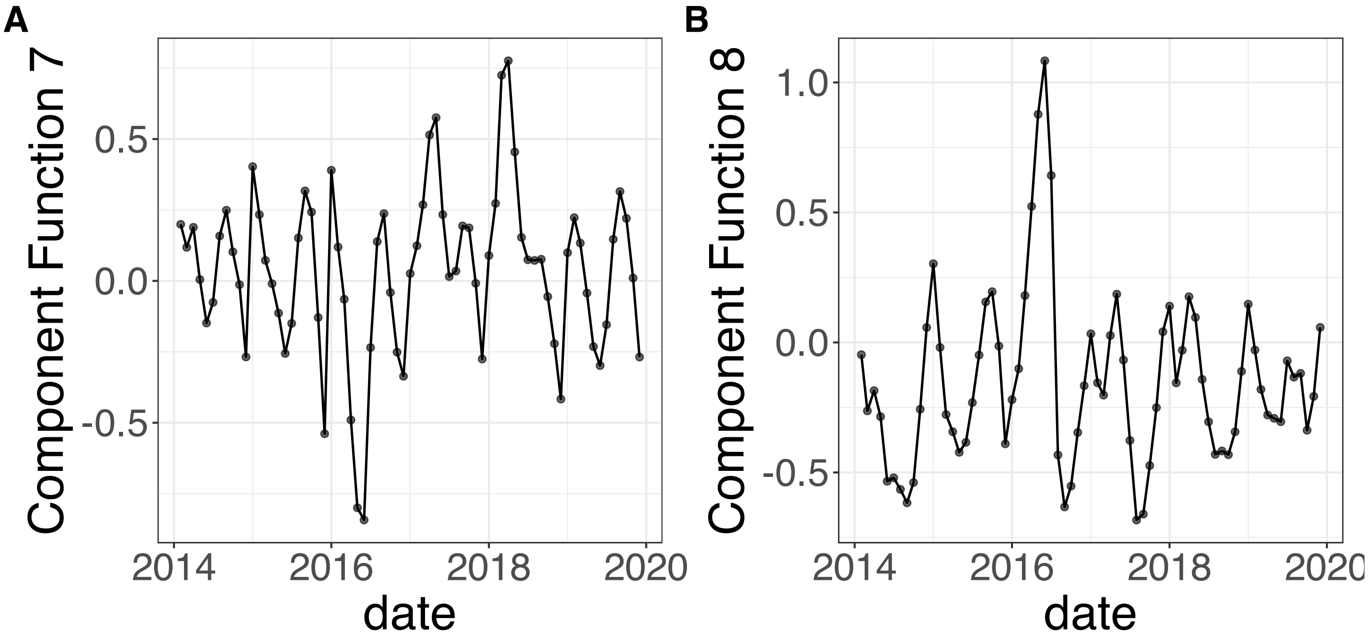


**Supplementary Figure 333.** SSANOVA-SVD reconstructed component functions.(A) Component 7. (B) Component 8.

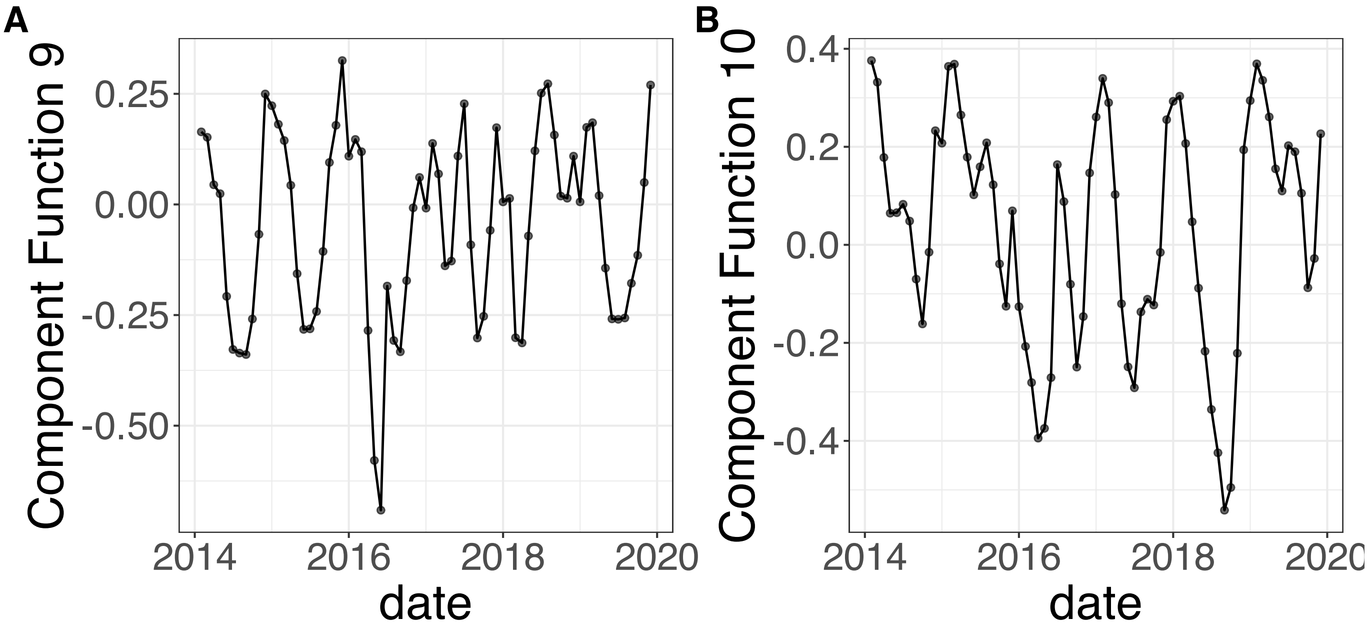


**Supplementary Figure 334.** SSANOVA-SVD reconstructed component functions.(A) Component 9. (B) Component 10.

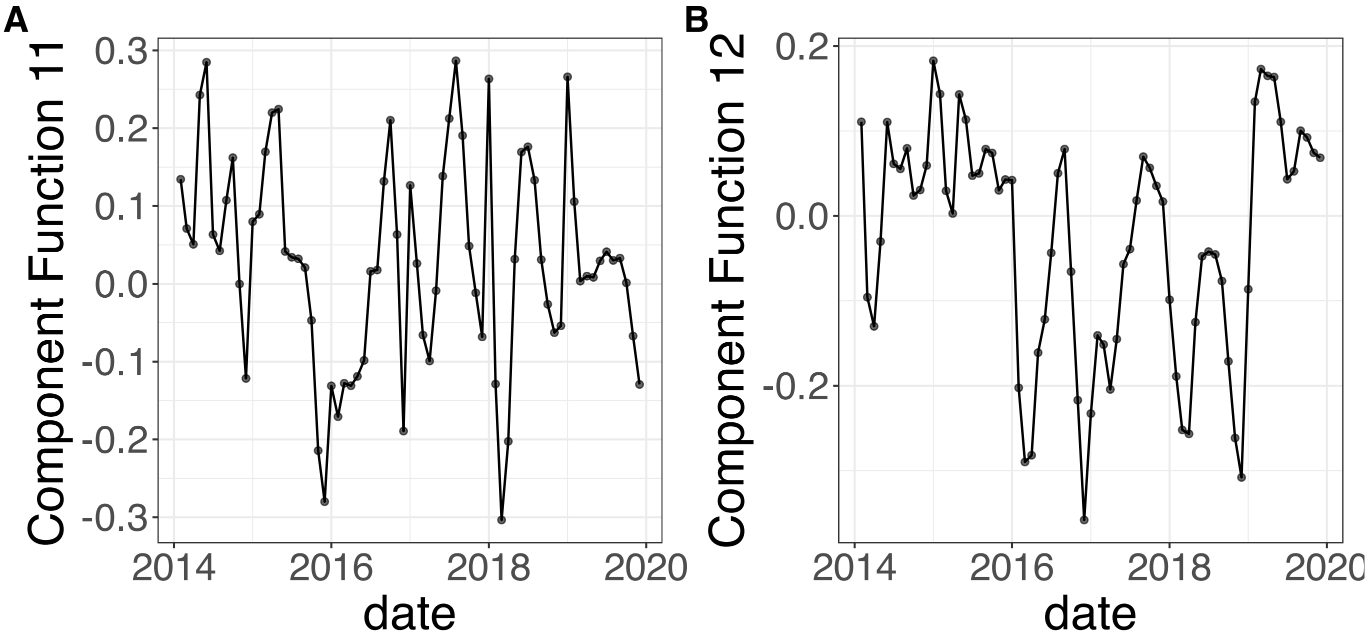


**Supplementary Figure 335.** SSANOVA-SVD reconstructed component functions.(A) Component 11. (B) Component 12.

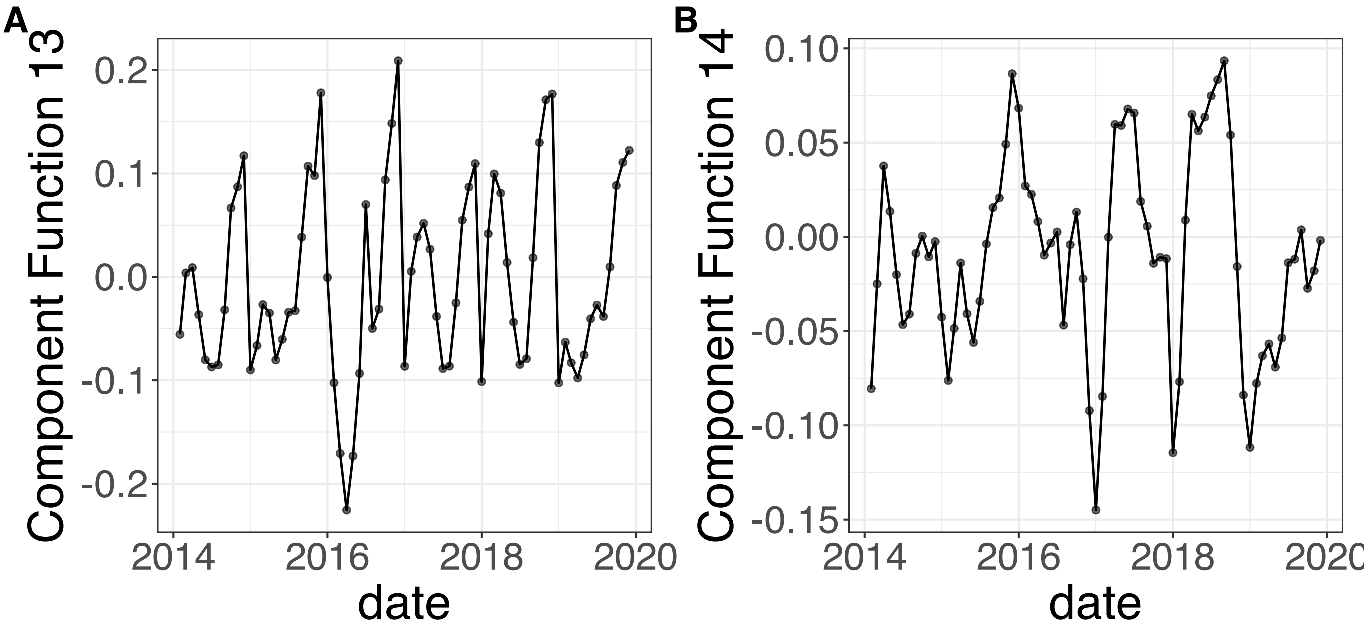


**Supplementary Figure 336.** SSANOVA-SVD reconstructed component functions.(A) Component 13. (B) Component 14.

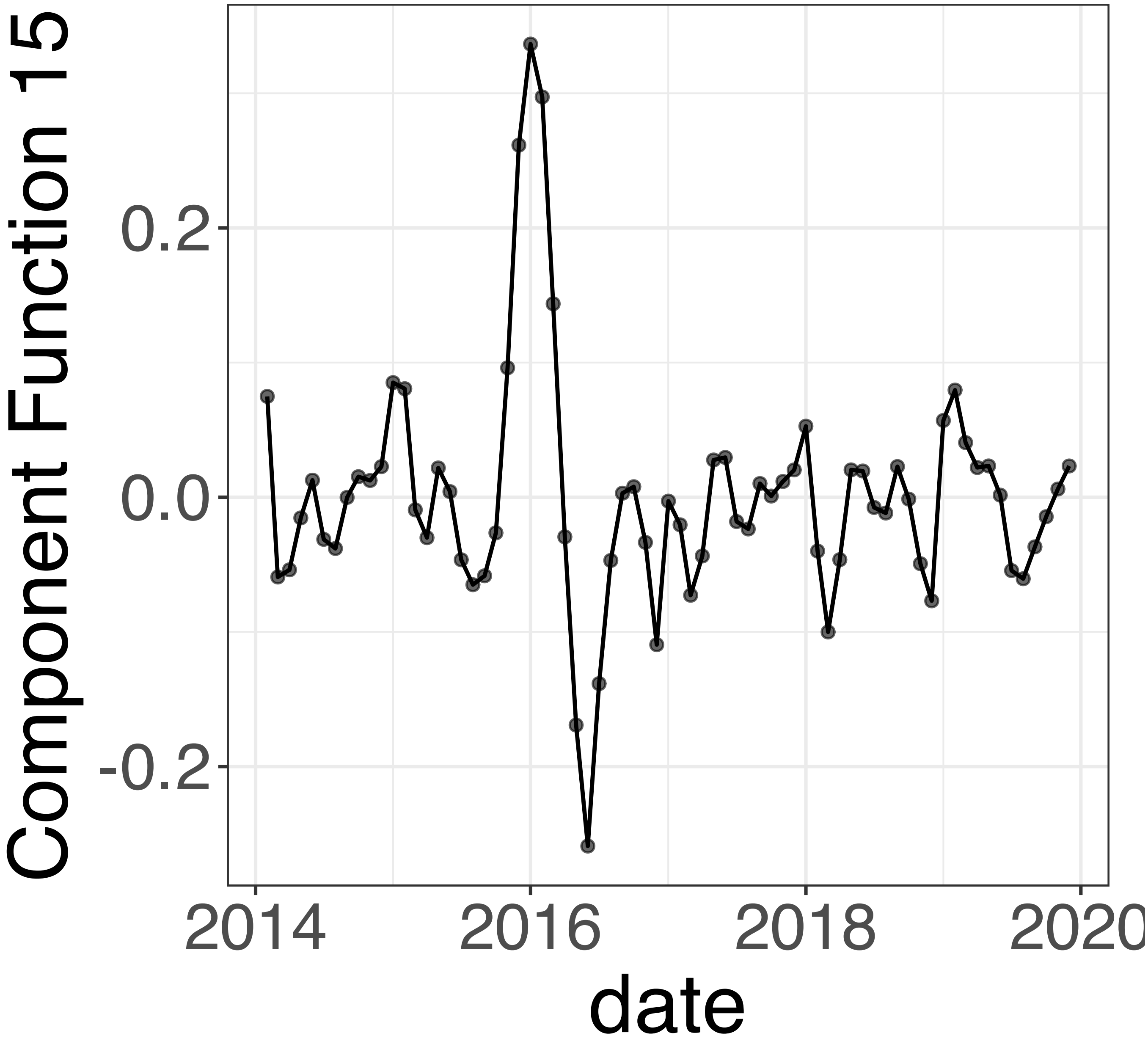


**Supplementary Figure 337.** SSANOVA-SVD reconstructed component functions.Component 15.

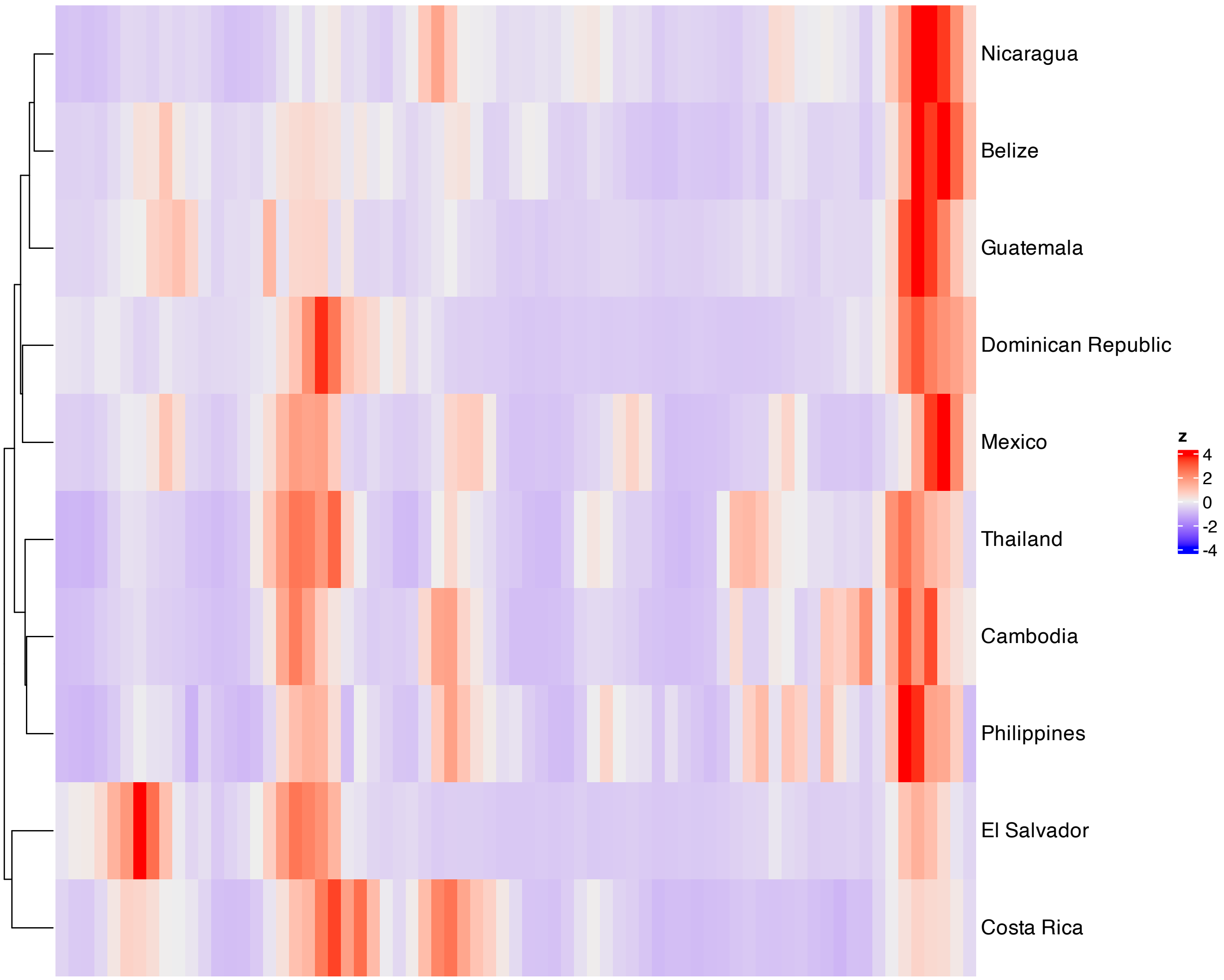


**Supplementary Figure 338.** Heatmap of features with the largest absolute loadings for SSANOVA-SVD component 1.

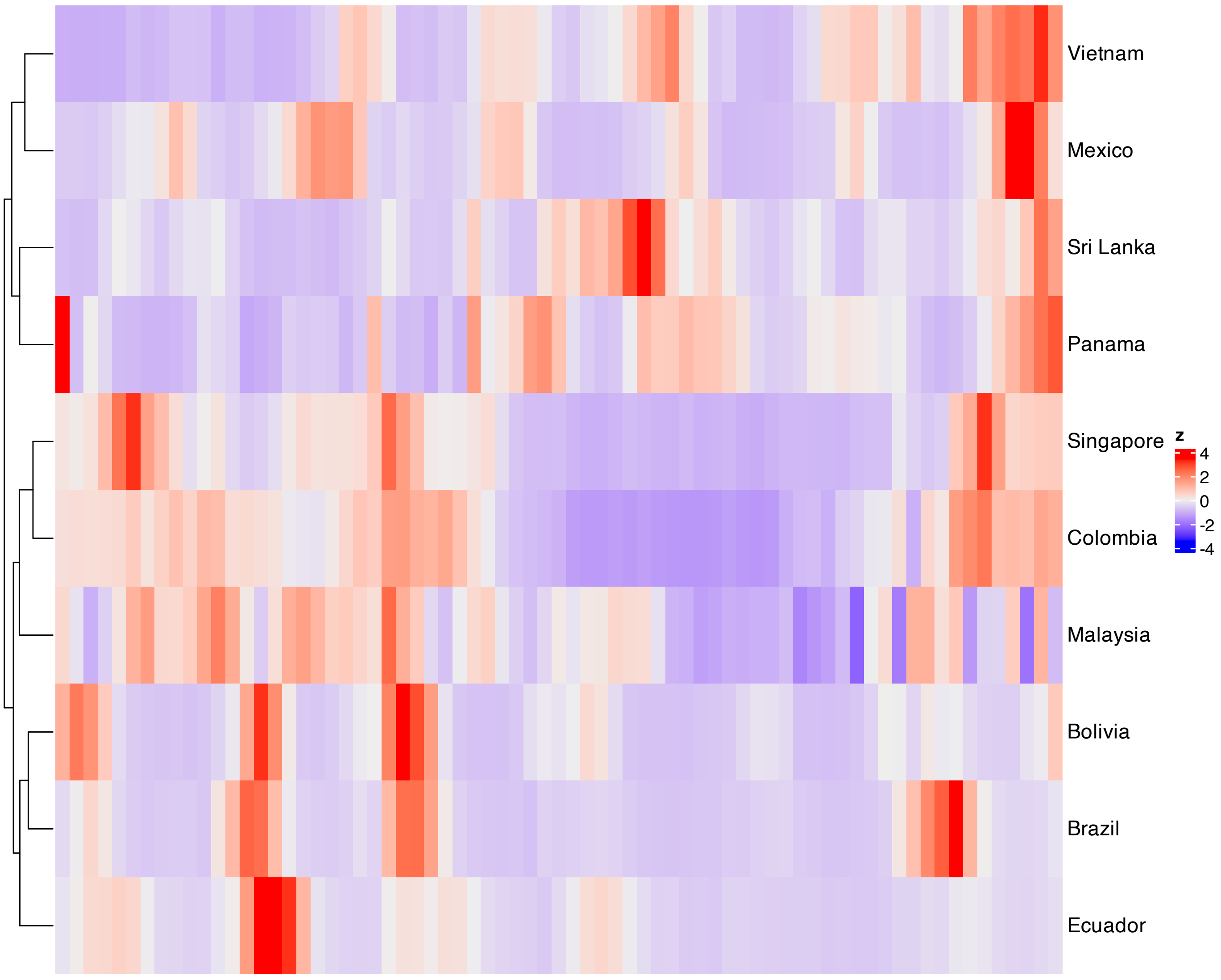


**Supplementary Figure 339.** Heatmap of features with the largest absolute loadings for SSANOVA-SVD component 2.

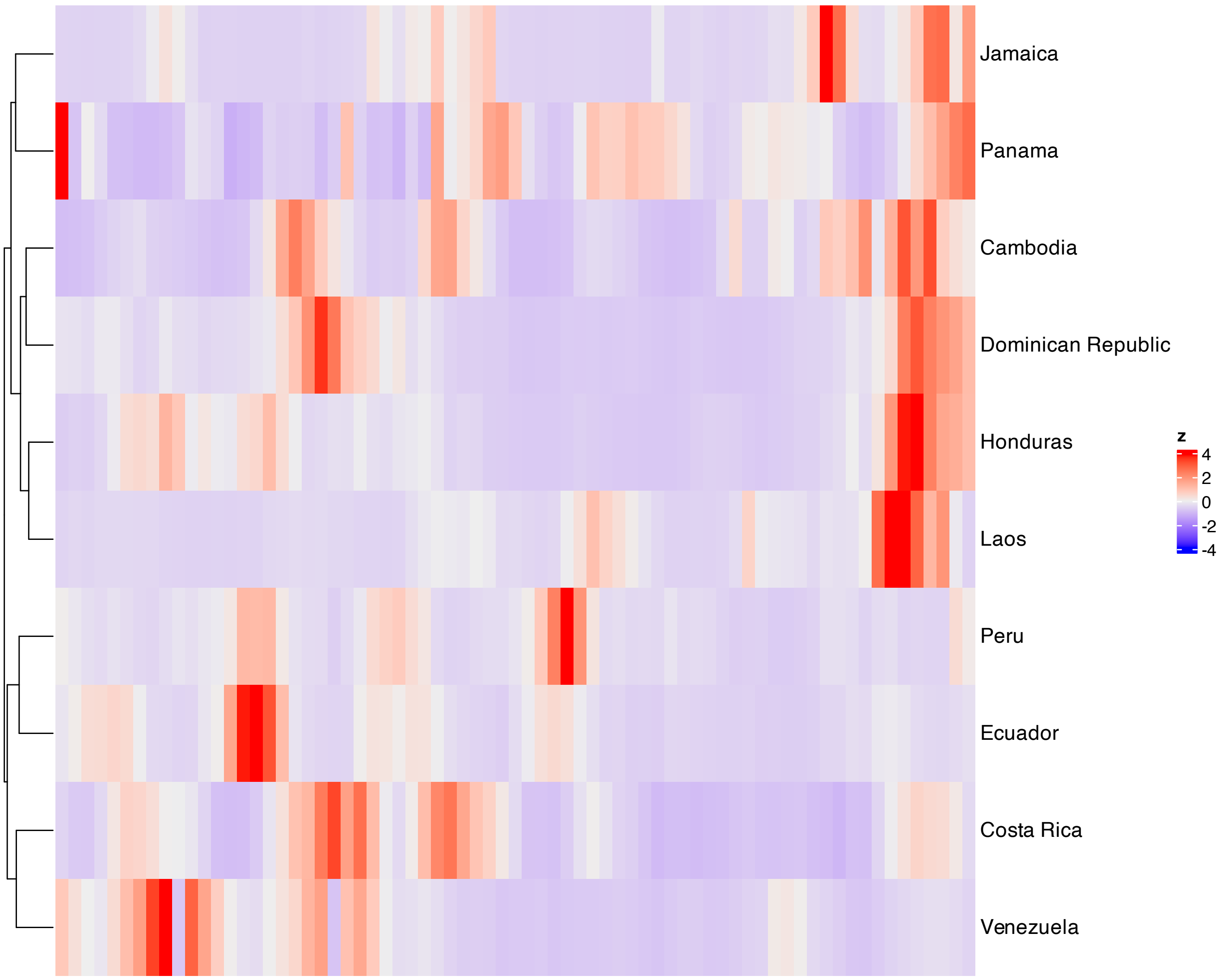


**Supplementary Figure 340.** Heatmap of features with the largest absolute loadings for SSANOVA-SVD component 3.

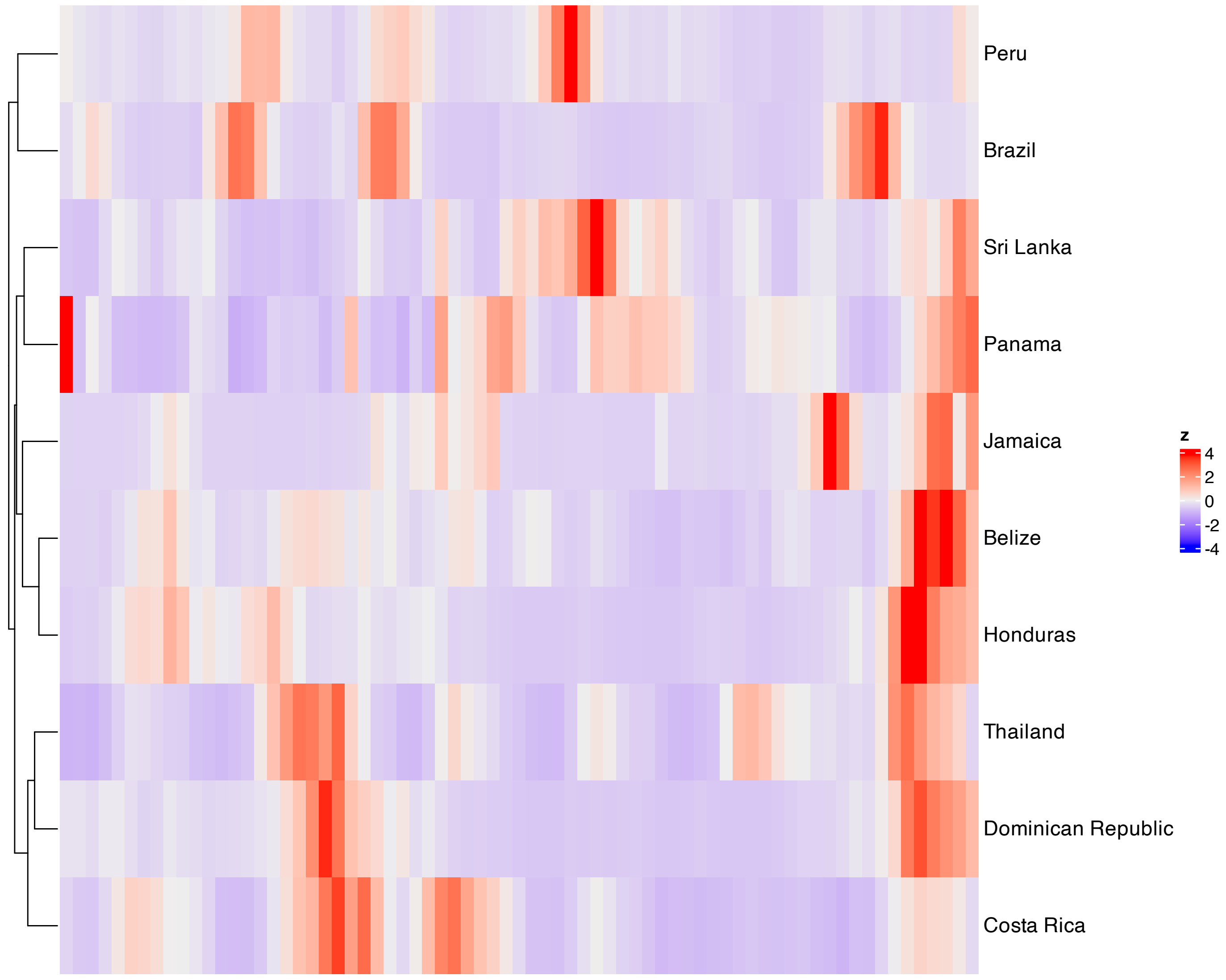


**Supplementary Figure 341.** Heatmap of features with the largest absolute loadings for SSANOVA-SVD component 4.

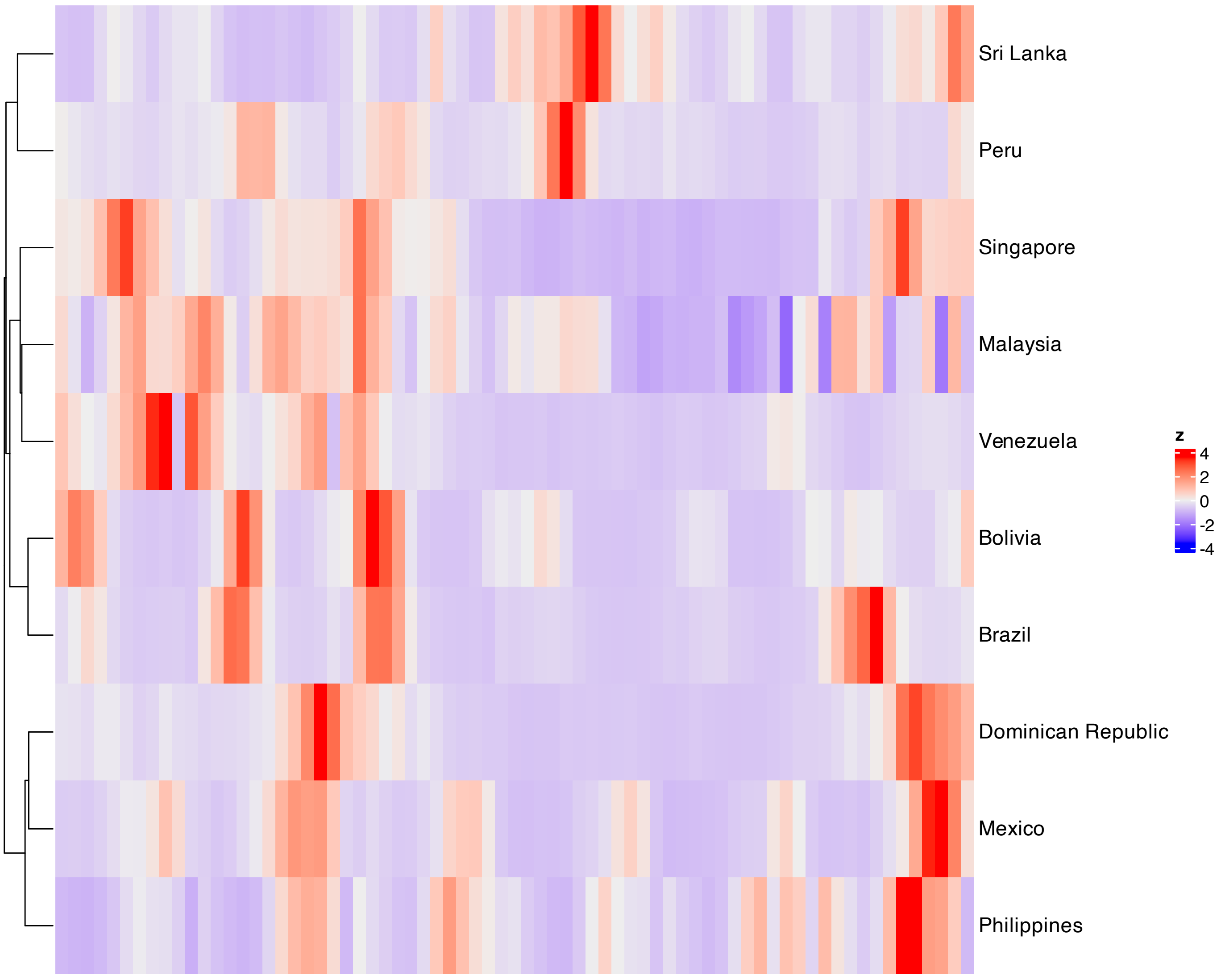


**Supplementary Figure 342.** Heatmap of features with the largest absolute loadings for SSANOVA-SVD component 5.

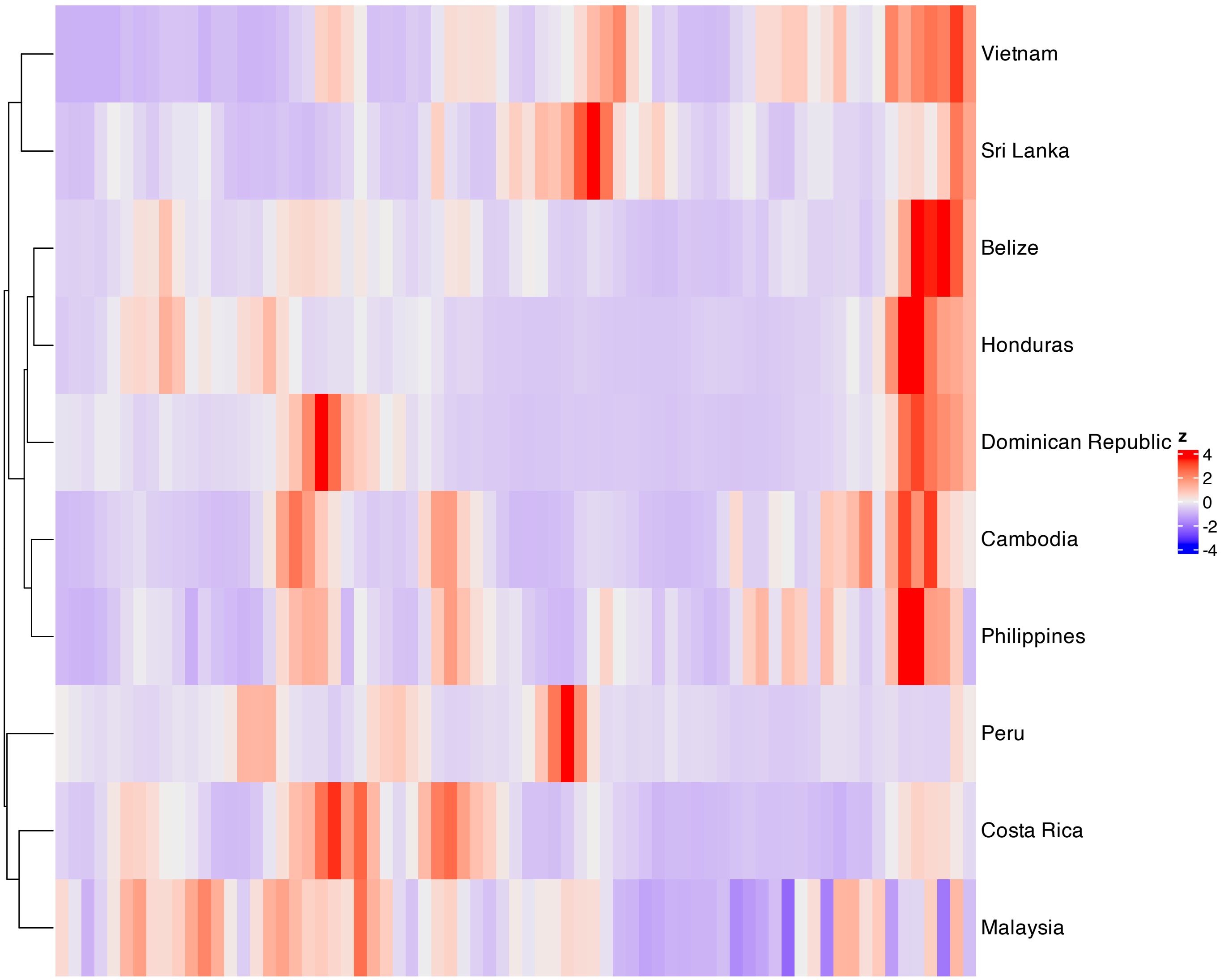


**Supplementary Figure 343.** Heatmap of features with the largest absolute loadings for SSANOVA-SVD component 6.

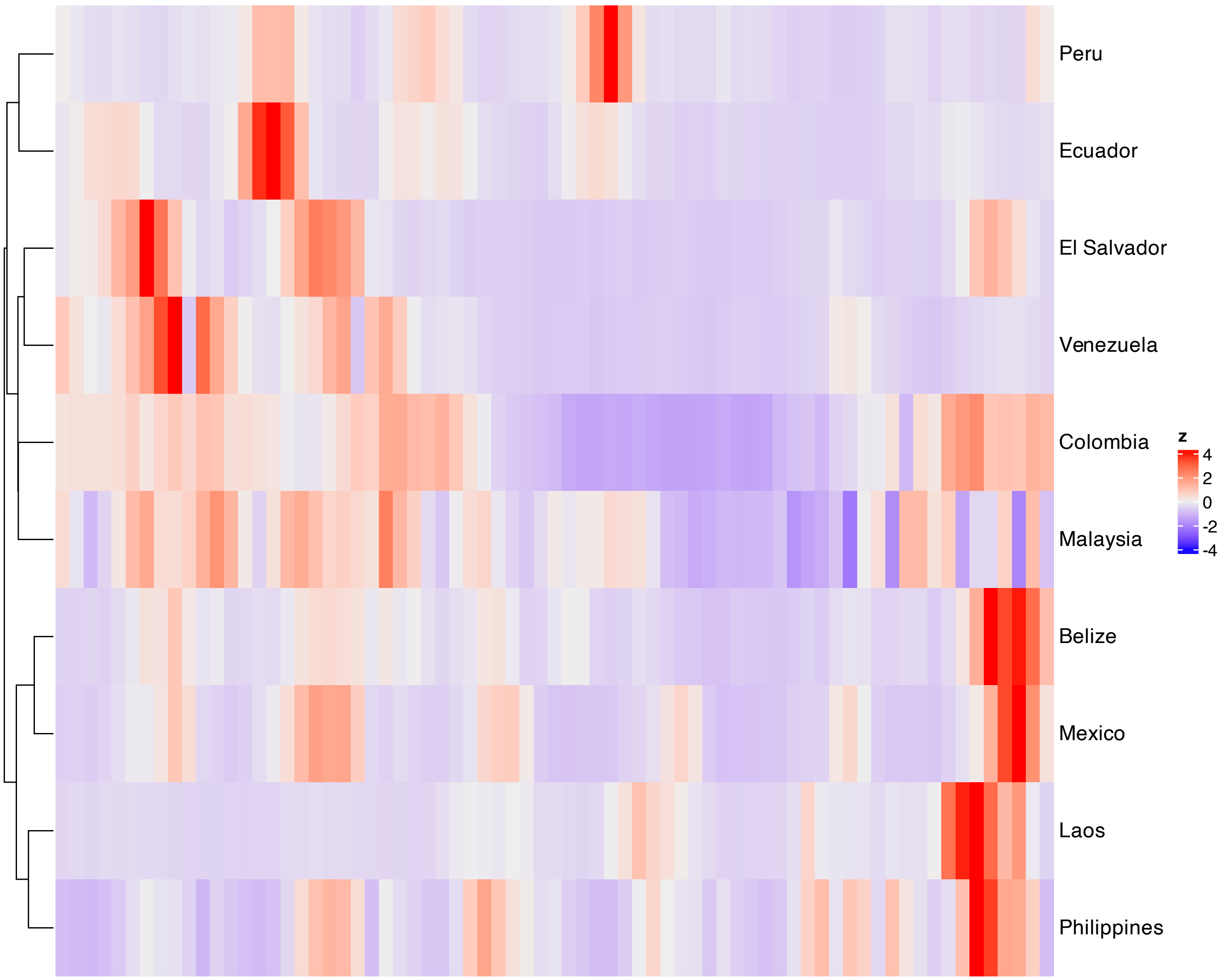


**Supplementary Figure 344.** Heatmap of features with the largest absolute loadings for SSANOVA-SVD component 7.

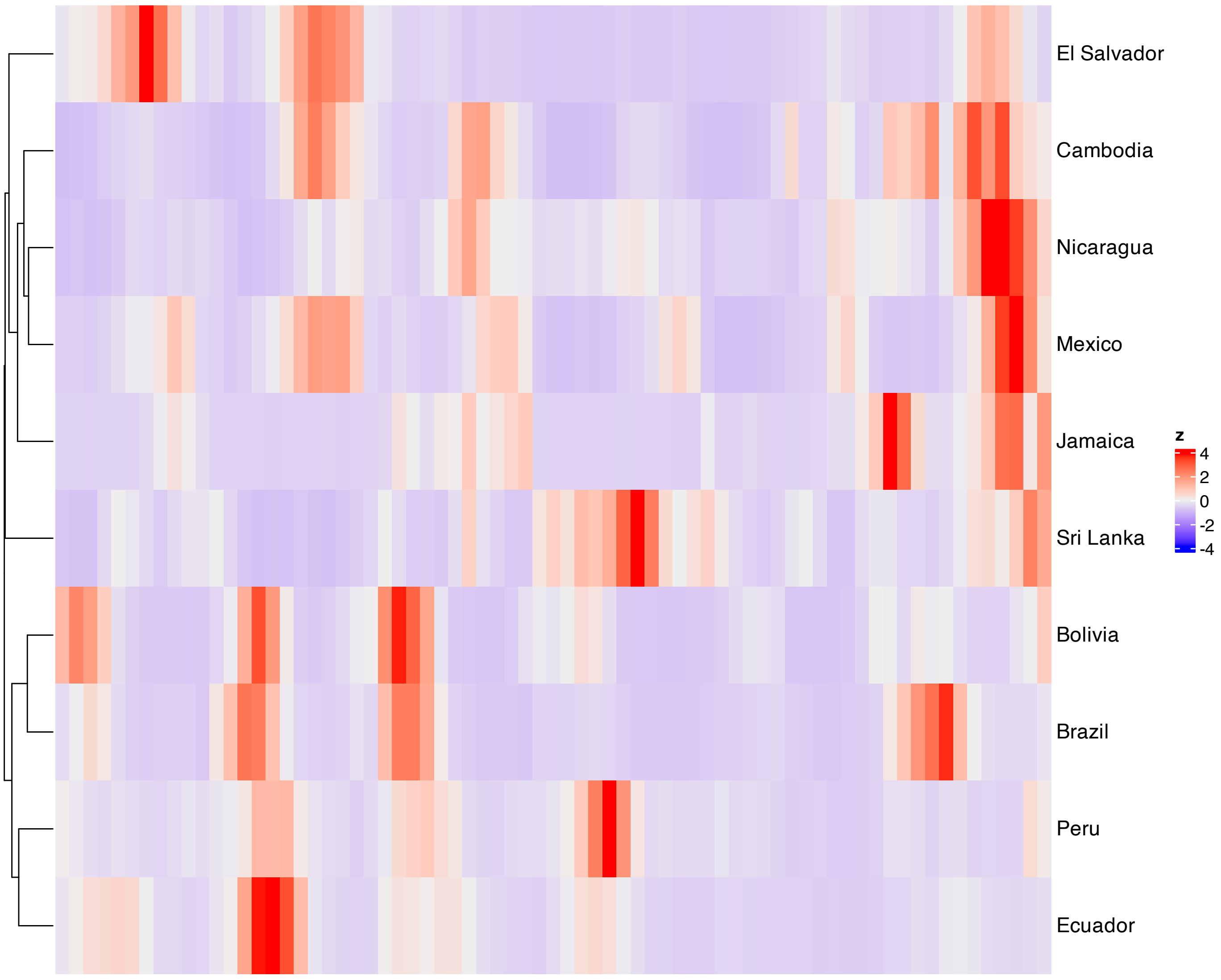


**Supplementary Figure 345.** Heatmap of features with the largest absolute loadings for SSANOVA-SVD component 8.

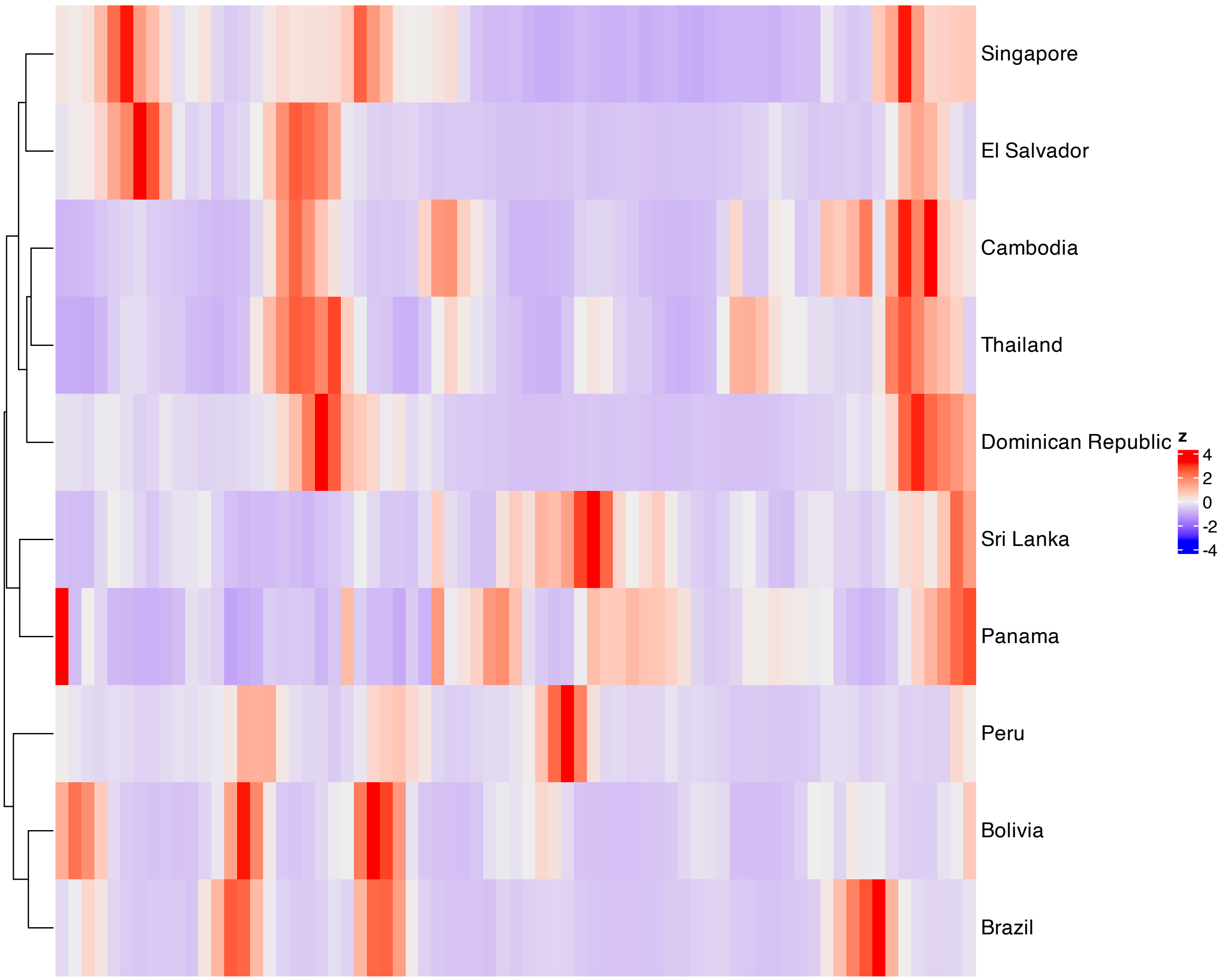


**Supplementary Figure 346.** Heatmap of features with the largest absolute loadings for SSANOVA-SVD component 9.

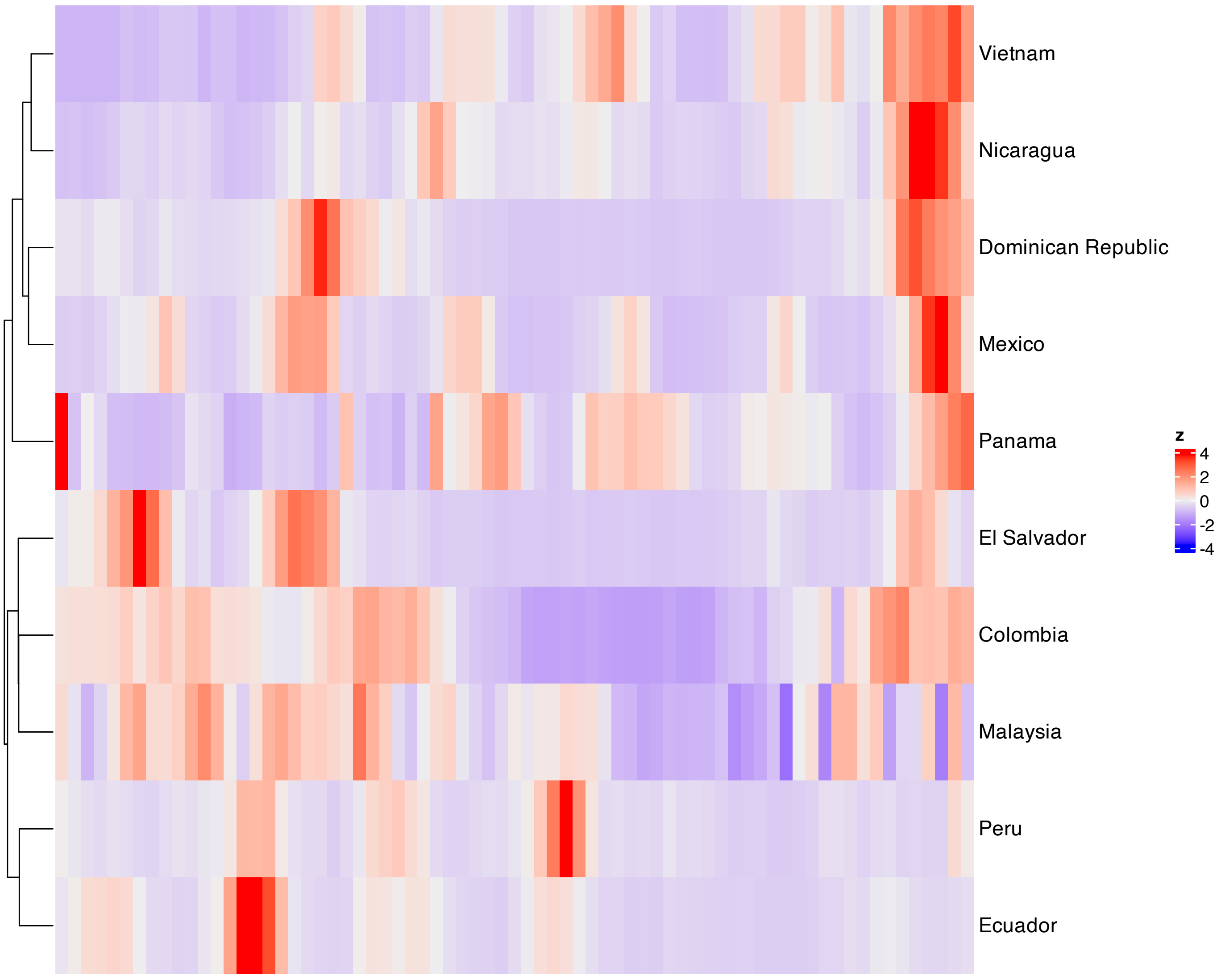


**Supplementary Figure 347.** Heatmap of features with the largest absolute loadings for SSANOVA-SVD component 10.

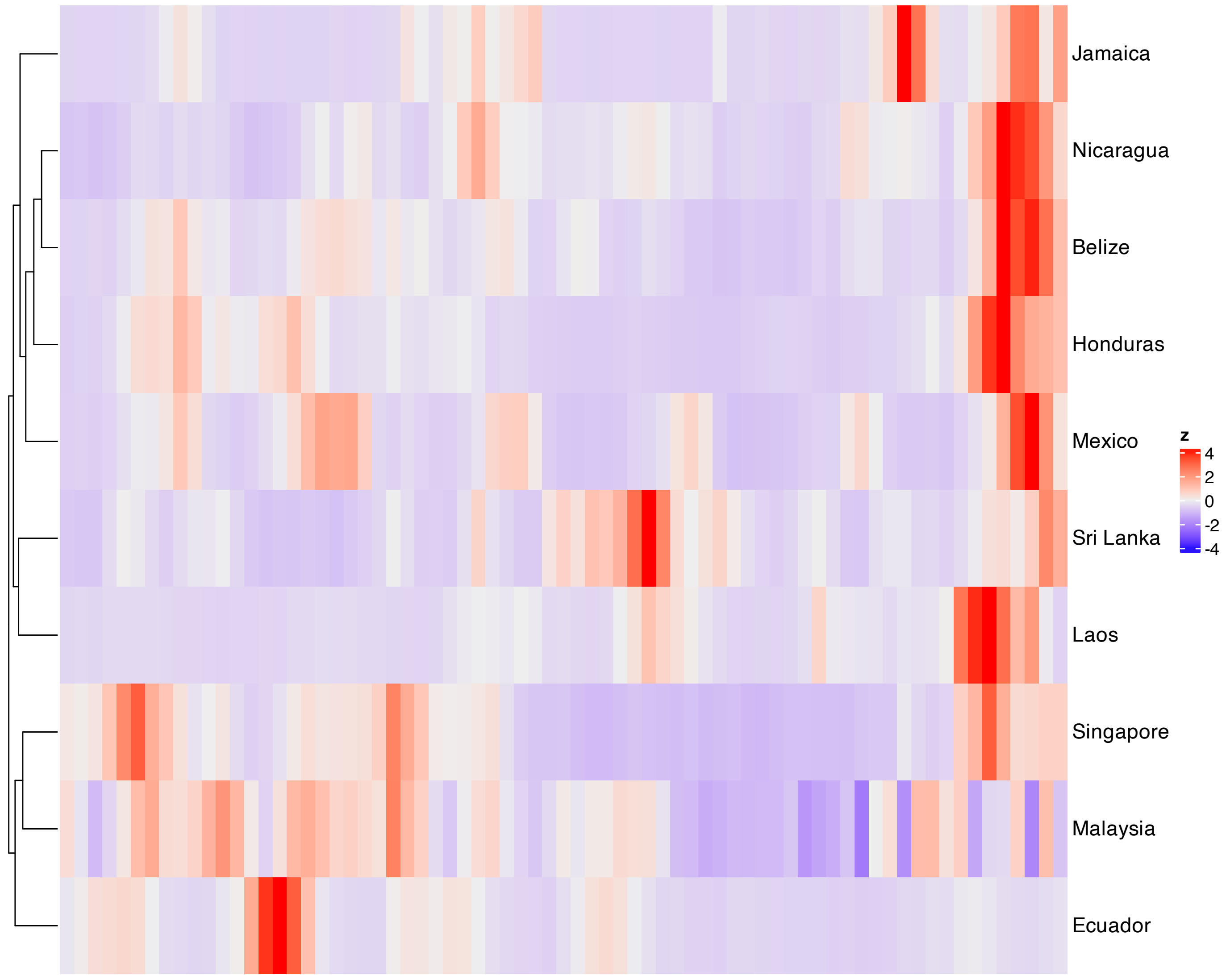


**Supplementary Figure 348.** Heatmap of features with the largest absolute loadings for SSANOVA-SVD component 11.

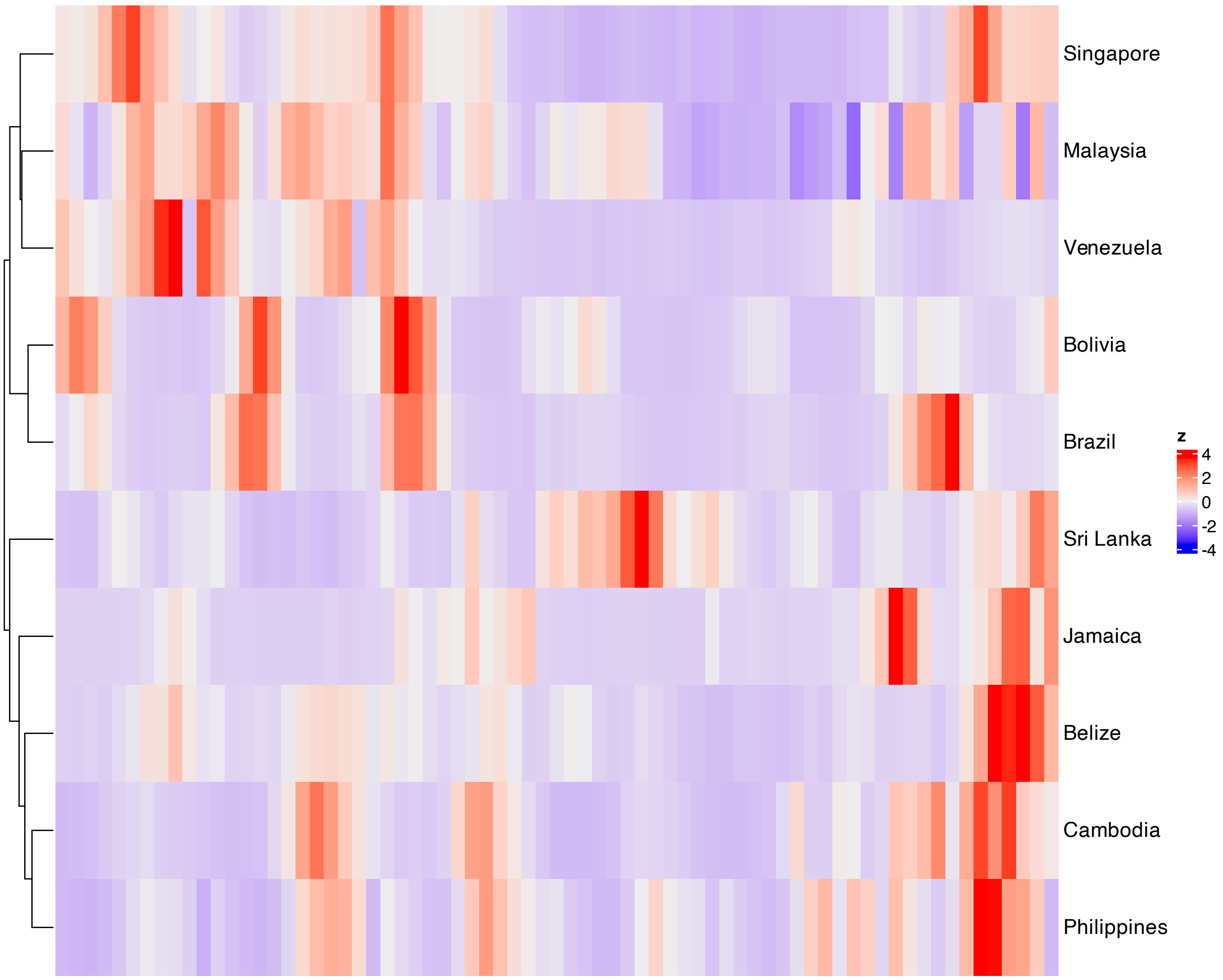


**Supplementary Figure 349.** Heatmap of features with the largest absolute loadings for SSANOVA-SVD component 12.

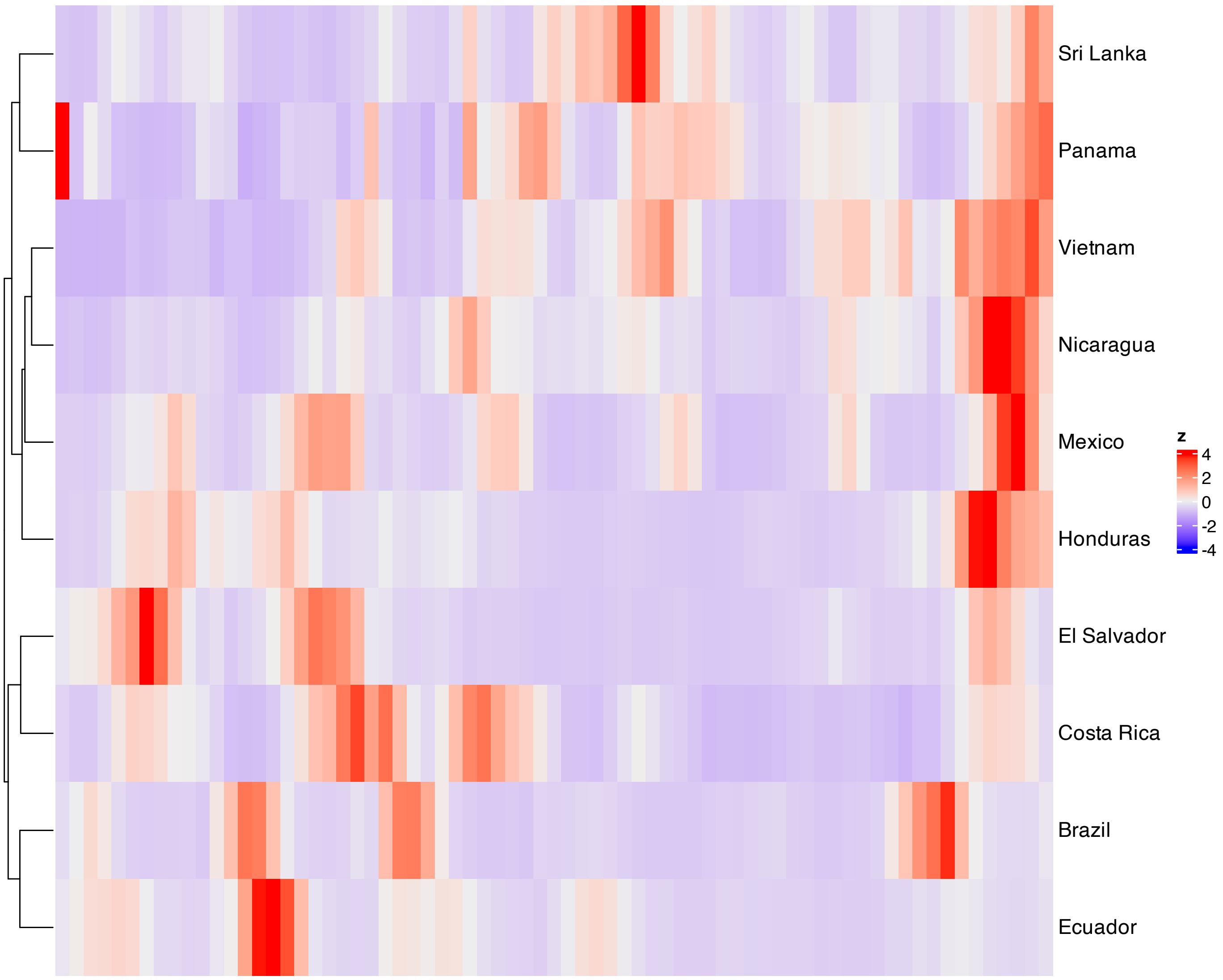


**Supplementary Figure 350.** Heatmap of features with the largest absolute loadings for SSANOVA-SVD component 13.

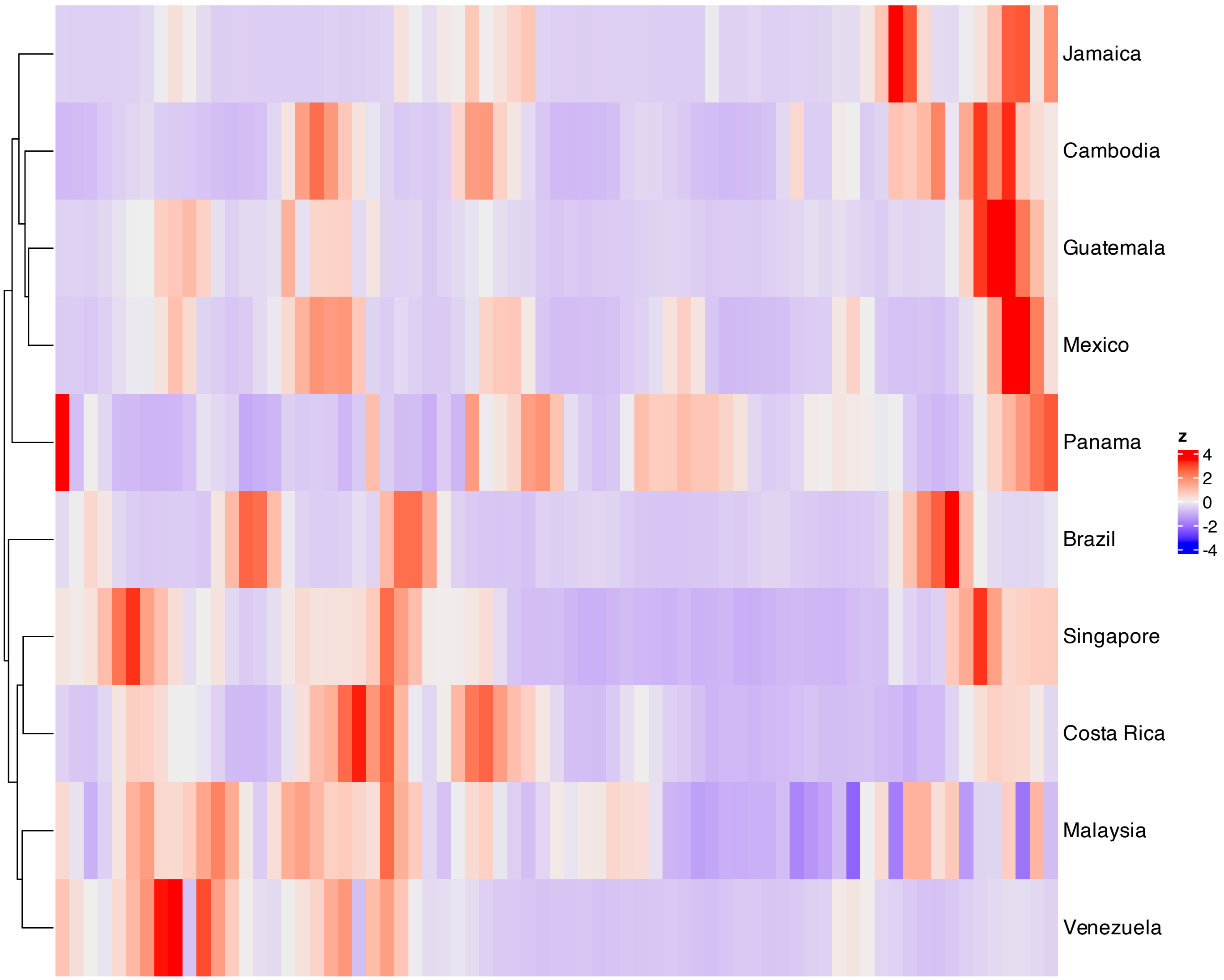


**Supplementary Figure 351.** Heatmap of features with the largest absolute loadings for SSANOVA-SVD component 14.

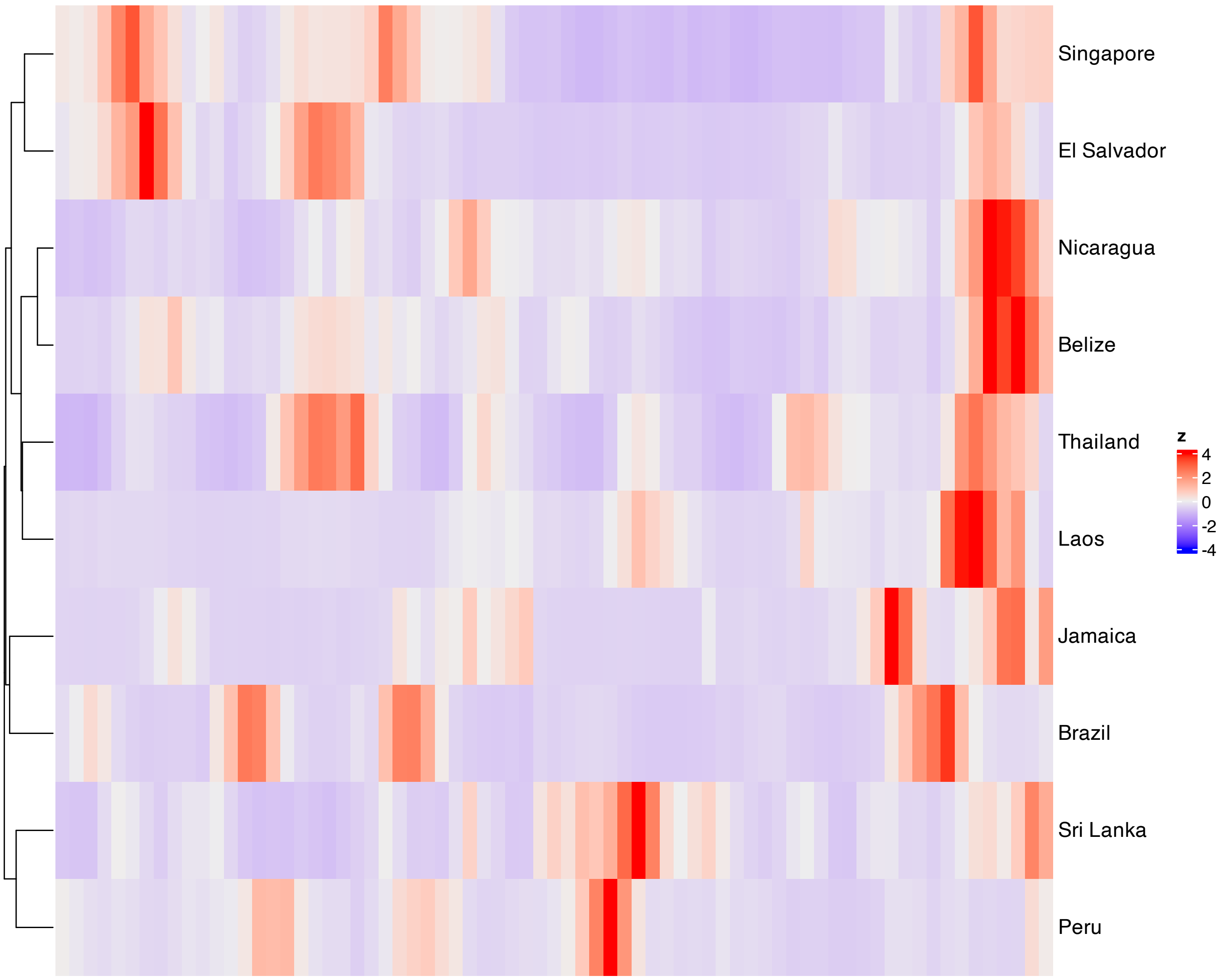


**Supplementary Figure 352.** Heatmap of features with the largest absolute loadings for SSANOVA-SVD component 15.

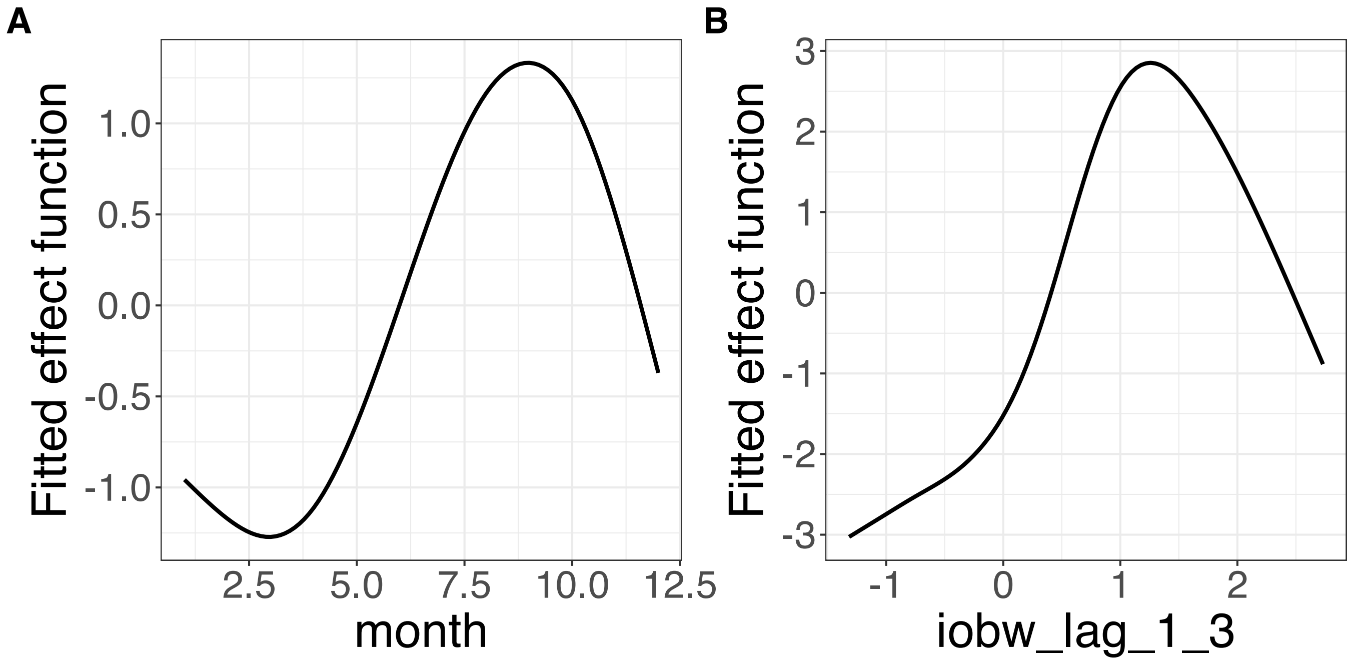


**Supplementary Figure 353.** Decomposition of reconstructed SSANOVA-SVD component function 1. (A) month. (B) iobw_lag_1_3.

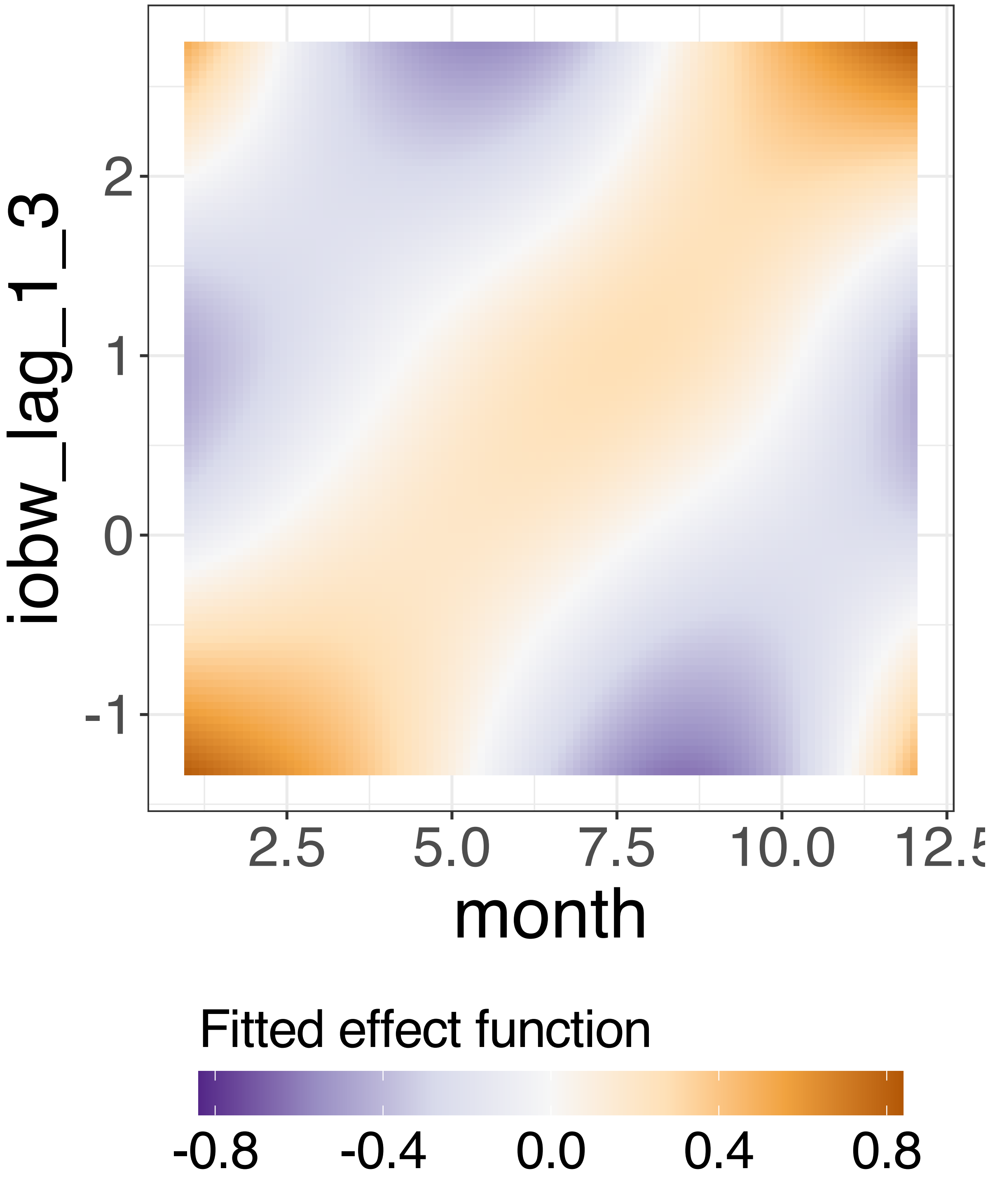


**Supplementary Figure 354.** Decomposition of reconstructed SSANOVA-SVD component function 1.

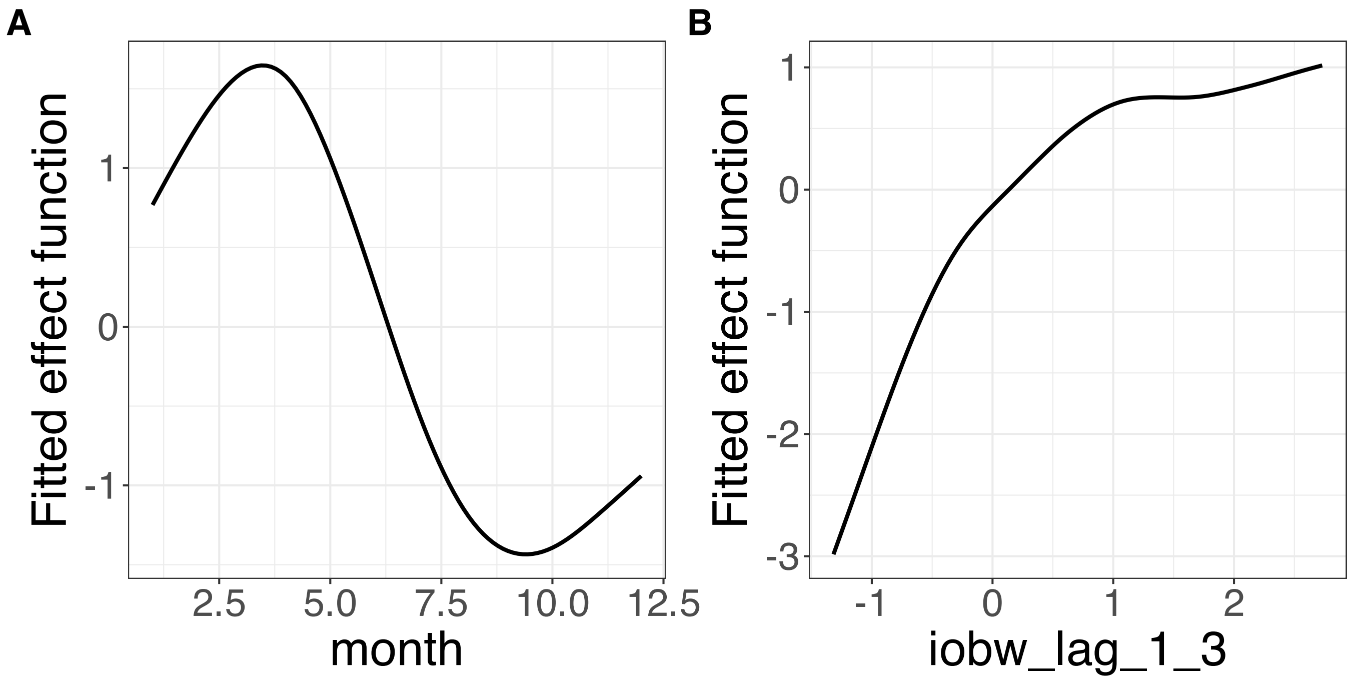


**Supplementary Figure 355.** Decomposition of reconstructed SSANOVA-SVD component function 2. (A) month. (B) iobw_lag_1_3.

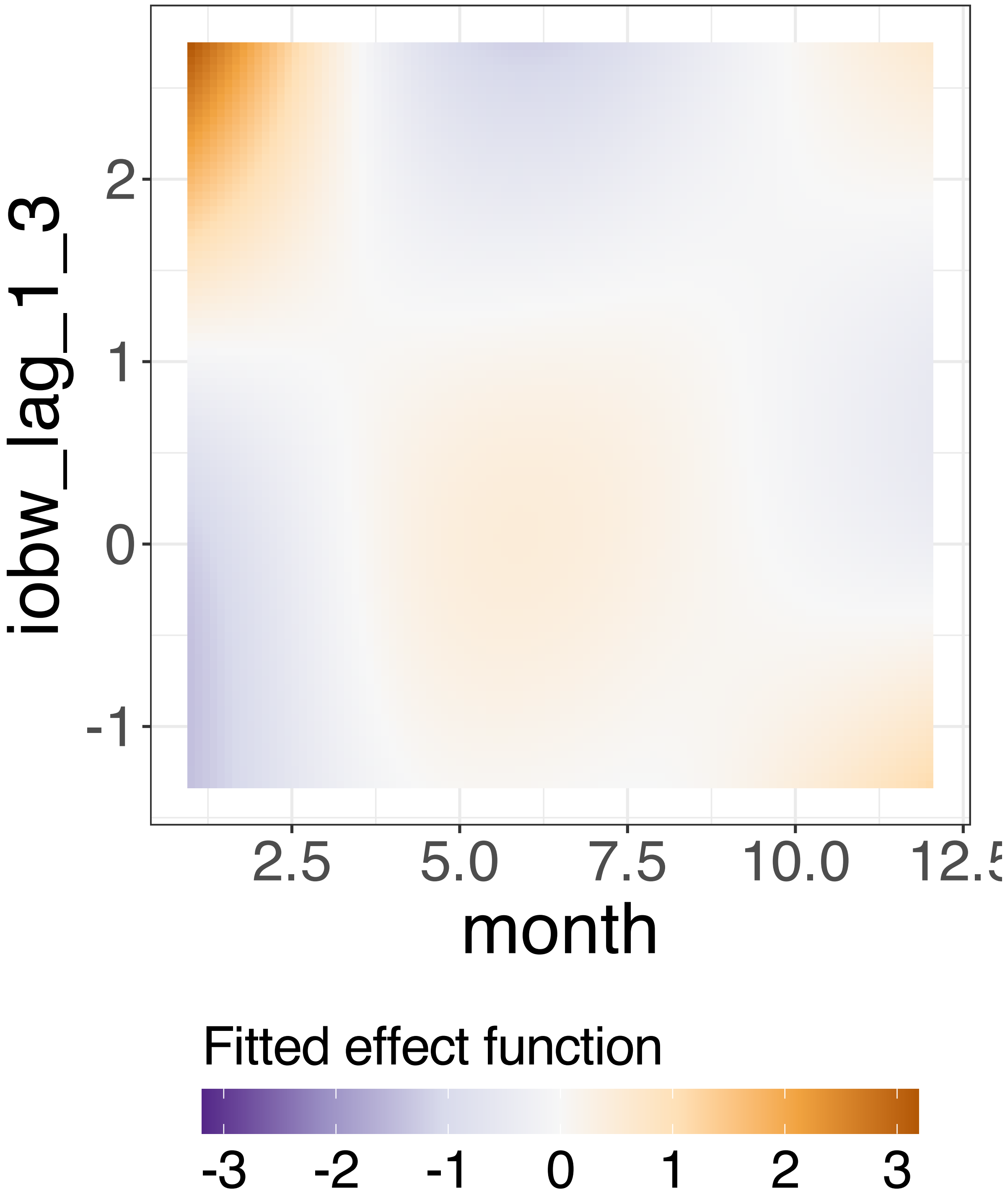


**Supplementary Figure 356.** Decomposition of reconstructed SSANOVA-SVD component function 2.

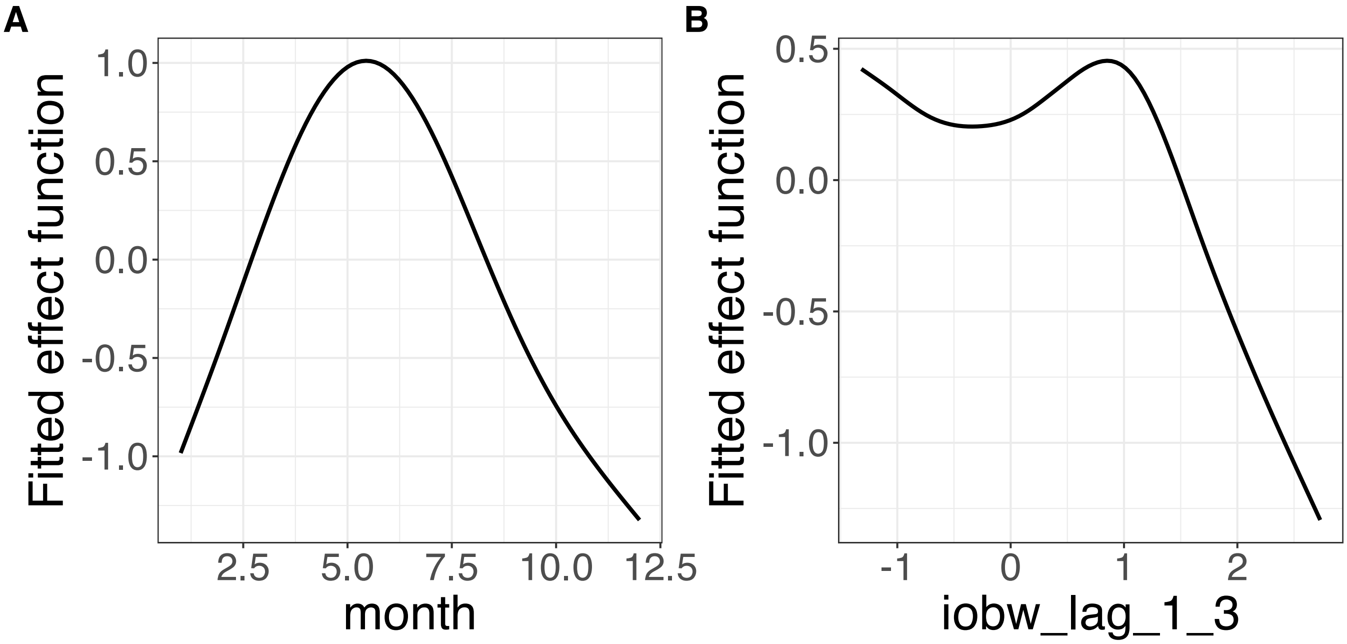


**Supplementary Figure 357.** Decomposition of reconstructed SSANOVA-SVD component function 3. (A) month. (B) iobw_lag_1_3.

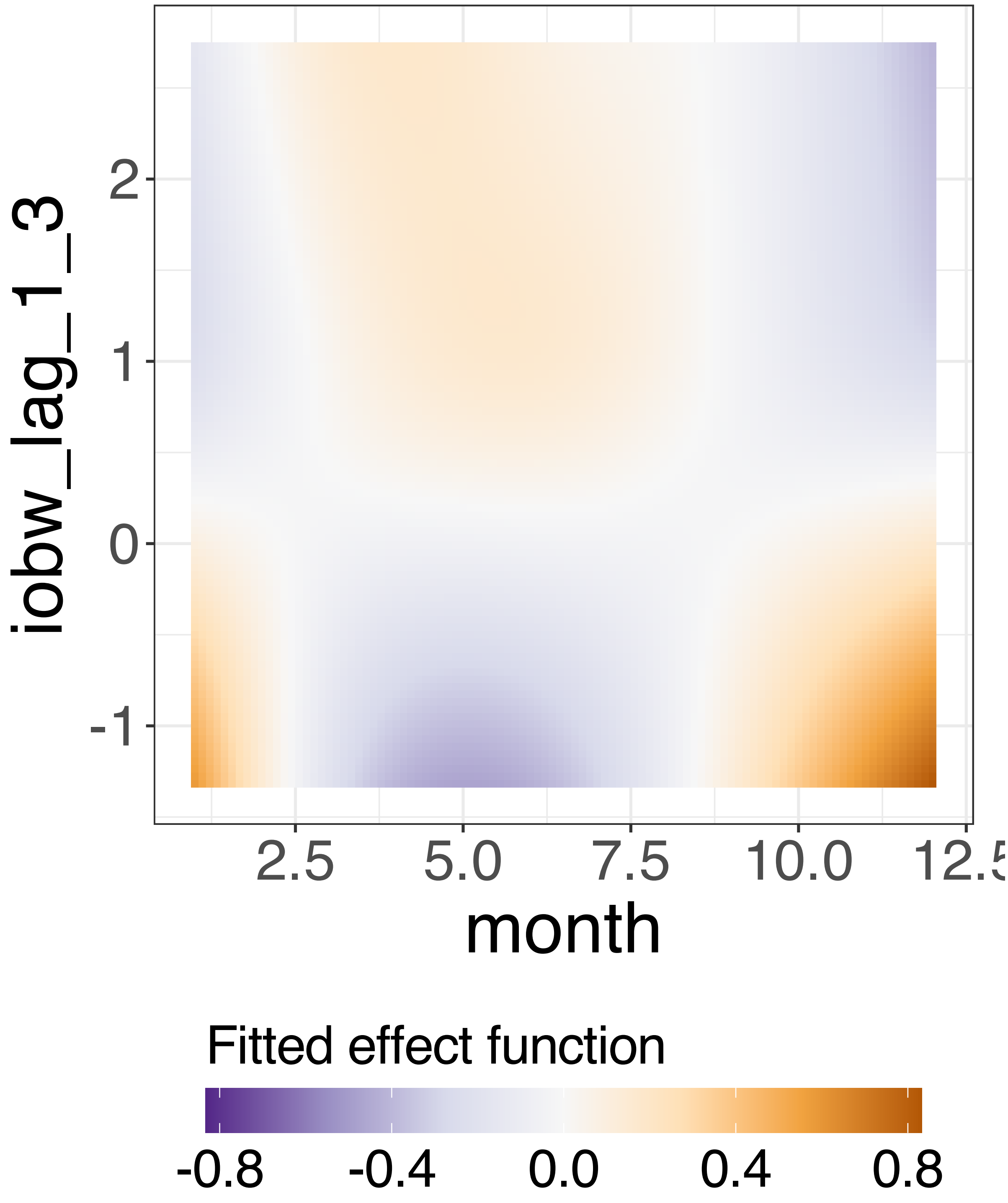


**Supplementary Figure 358.** Decomposition of reconstructed SSANOVA-SVD component function 3.

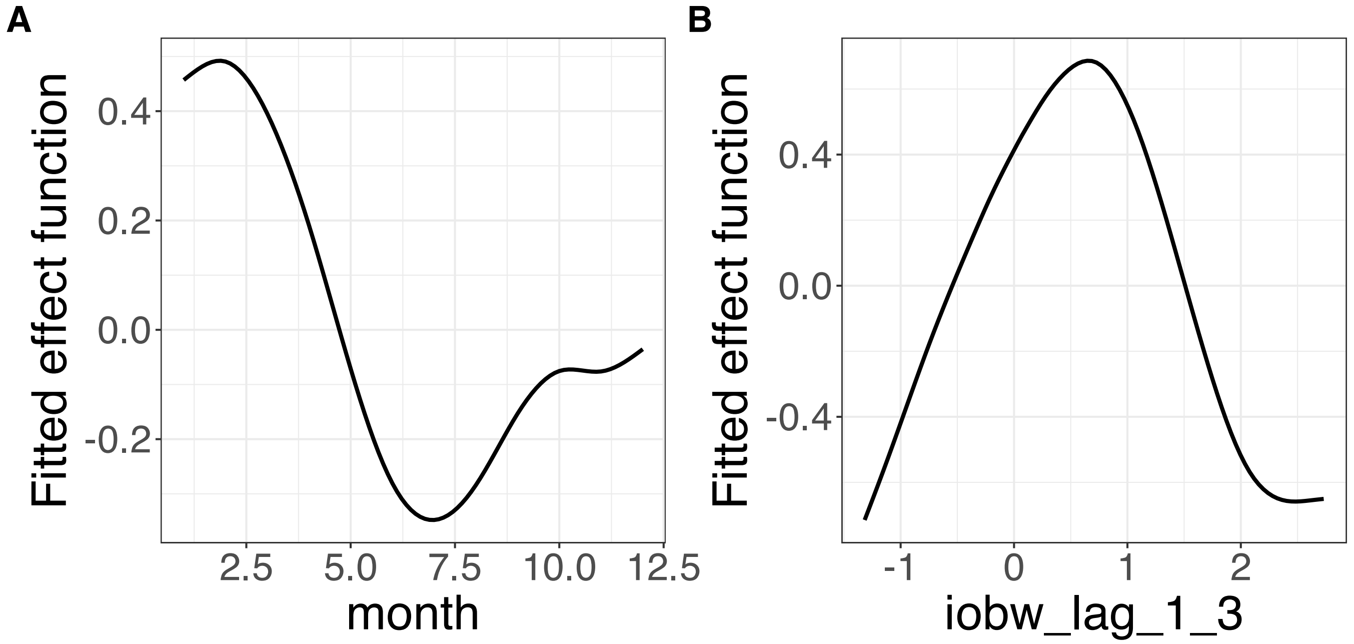


**Supplementary Figure 359.** Decomposition of reconstructed SSANOVA-SVD component function 4. (A) month. (B) iobw_lag_1_3.

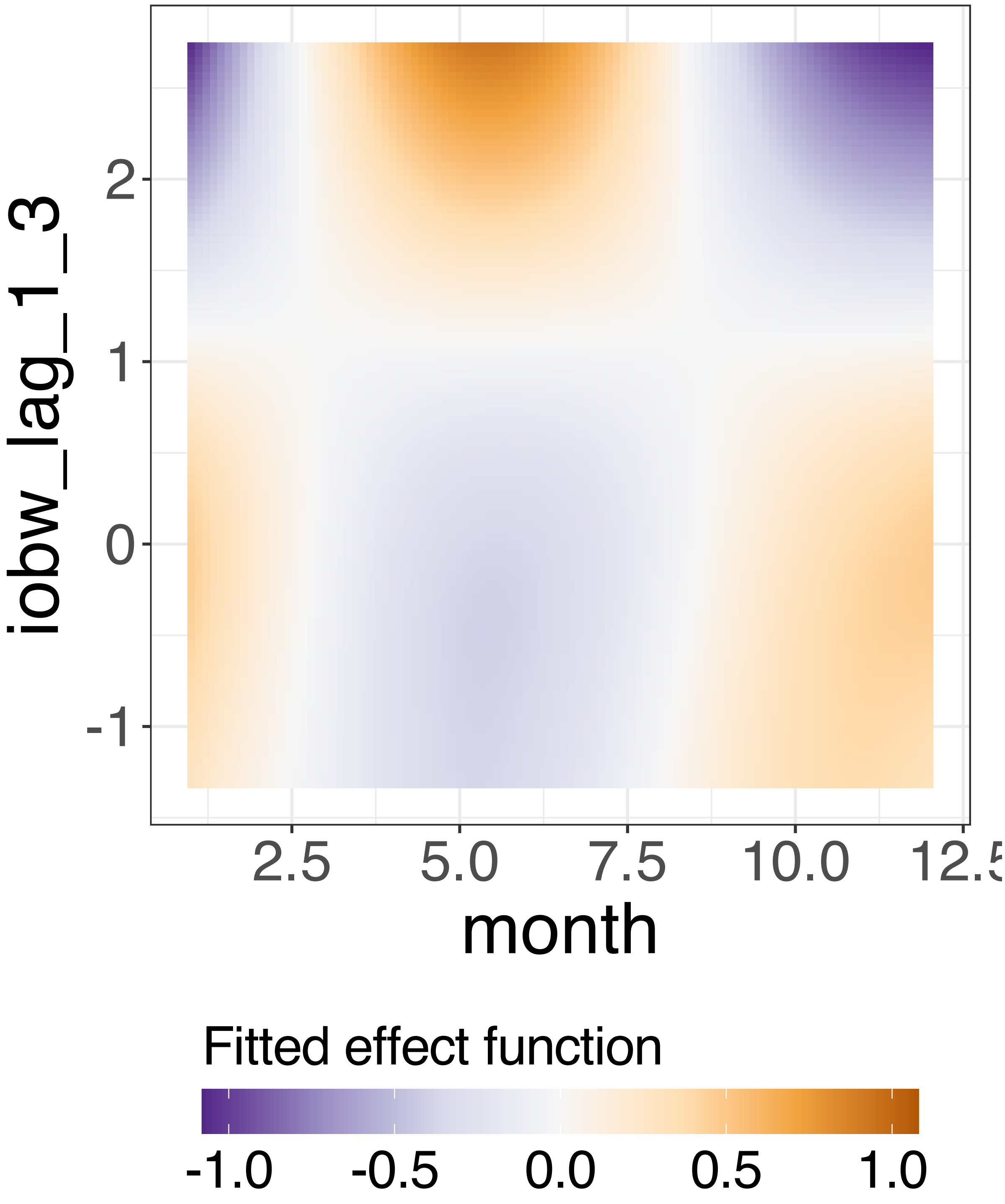


**Supplementary Figure 360.** Decomposition of reconstructed SSANOVA-SVD component function 4.

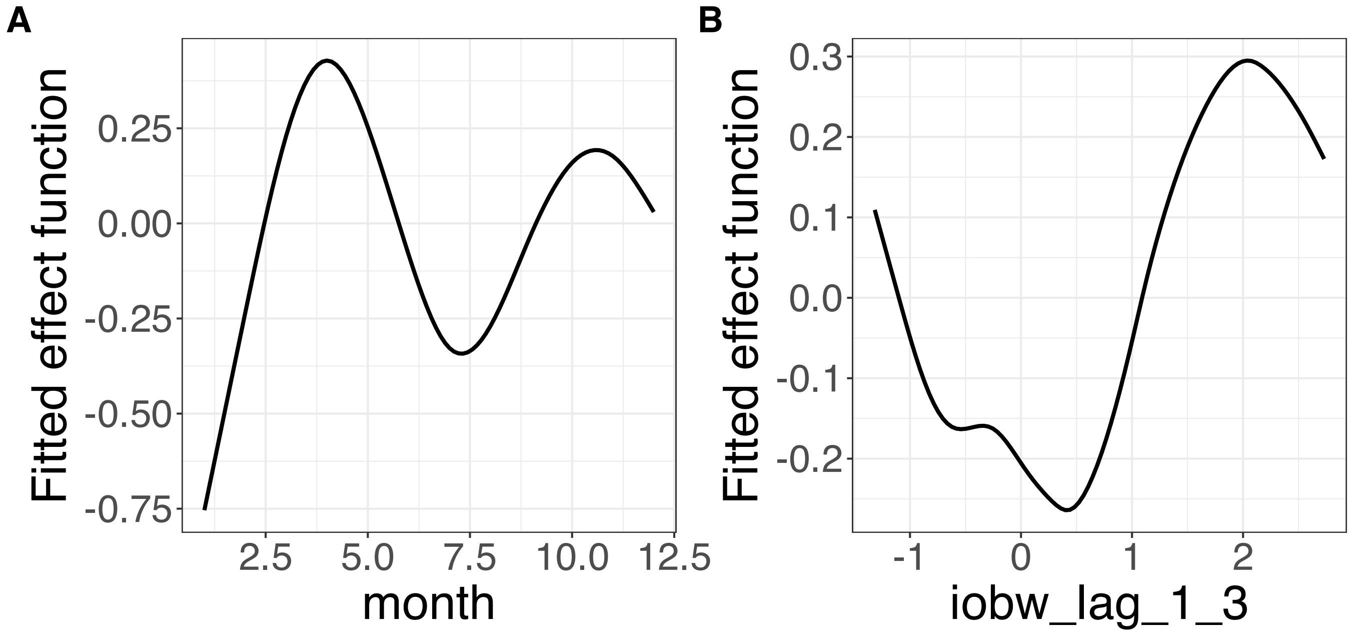


**Supplementary Figure 361.** Decomposition of reconstructed SSANOVA-SVD component function 5. (A) month. (B) iobw_lag_1_3.

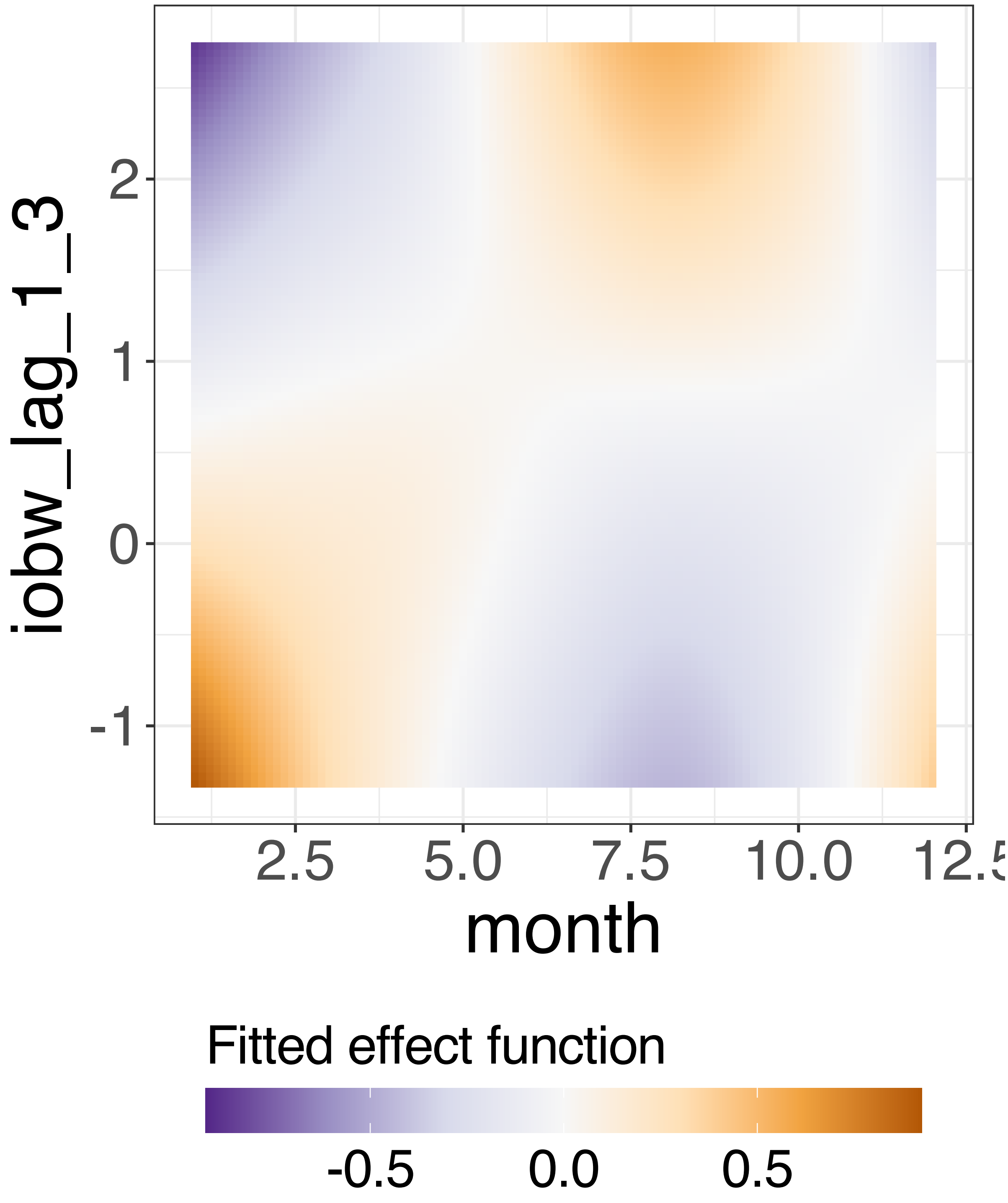


**Supplementary Figure 362.** Decomposition of reconstructed SSANOVA-SVD component function 5.

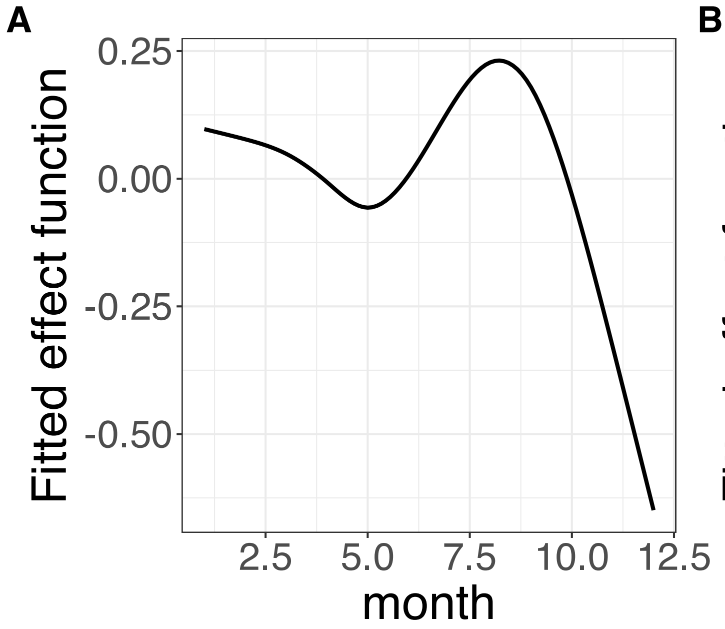


**Supplementary Figure 363.** Decomposition of reconstructed SSANOVA-SVD component function 6. (A) month. (B) iobw_lag_1_3.

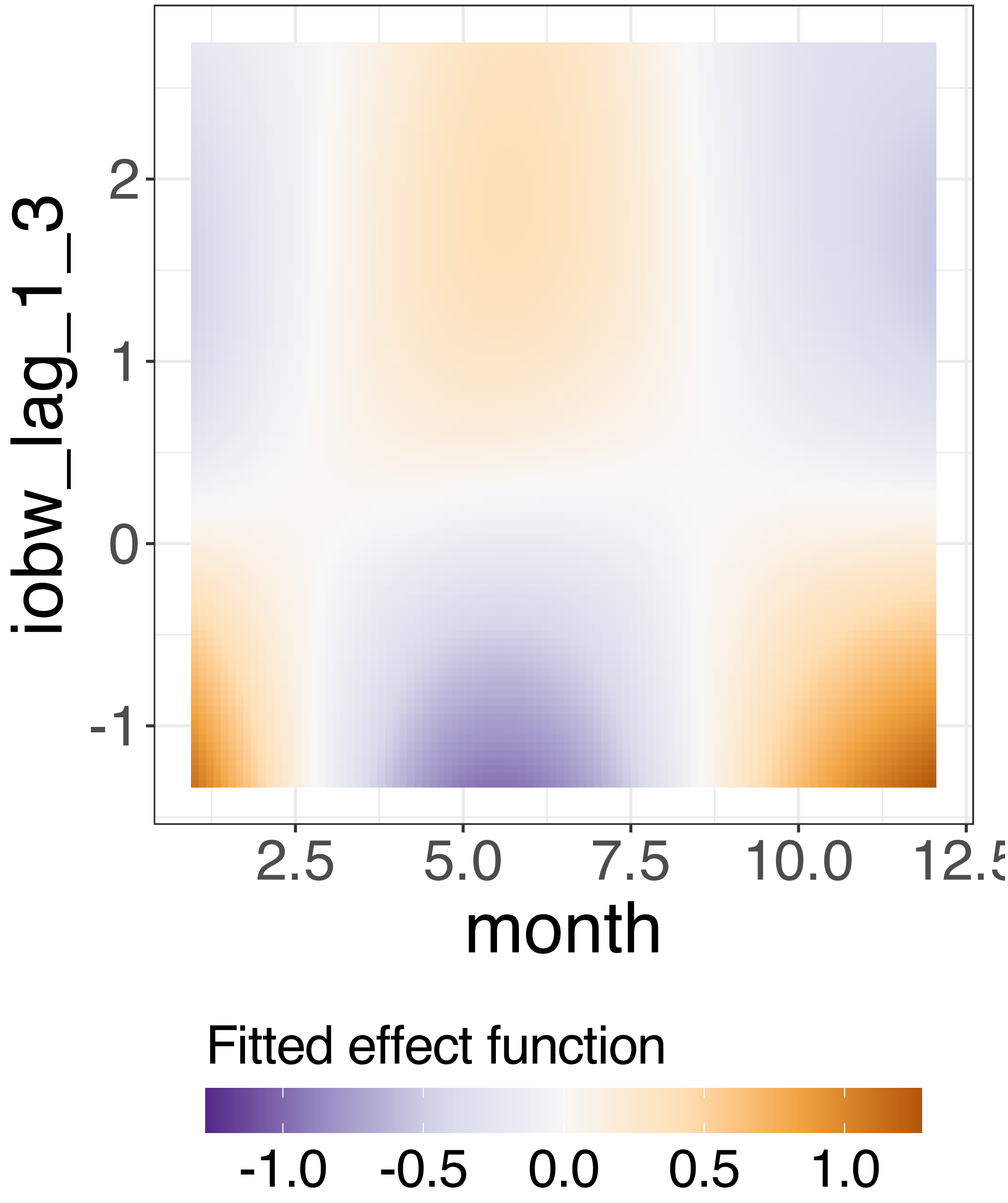


**Supplementary Figure 364.** Decomposition of reconstructed SSANOVA-SVD component function 6.

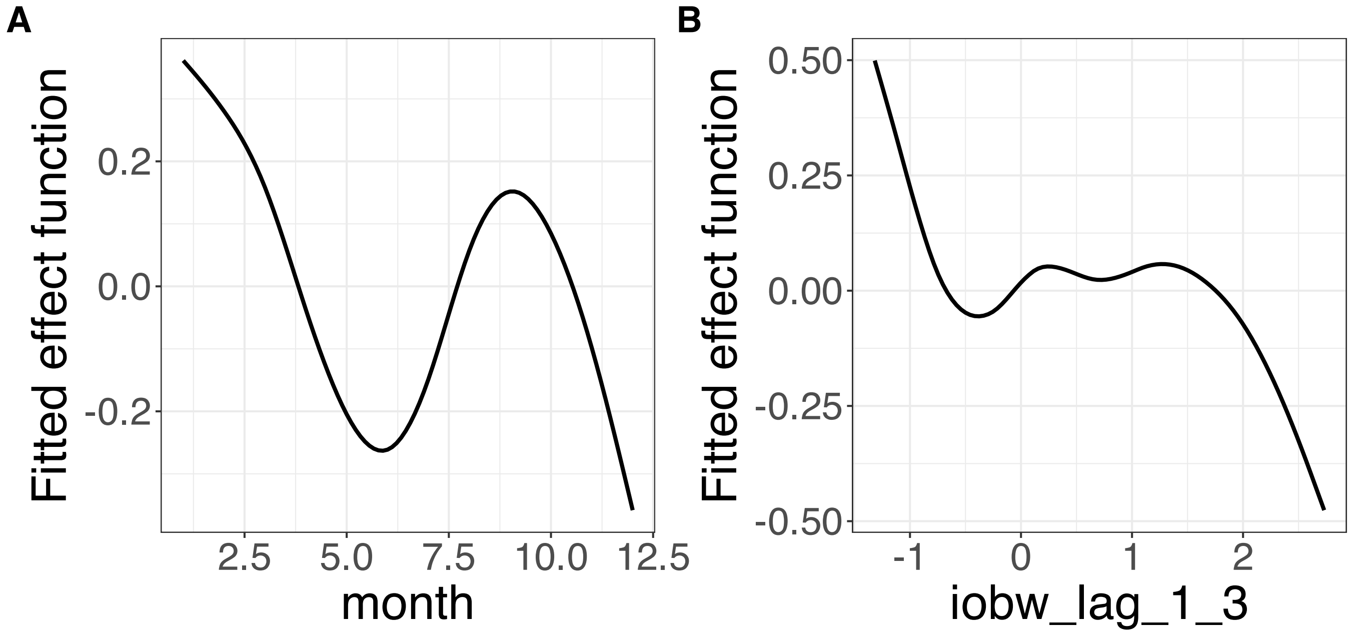


**Supplementary Figure 365.** Decomposition of reconstructed SSANOVA-SVD component function 7. (A) month. (B) iobw_lag_1_3.

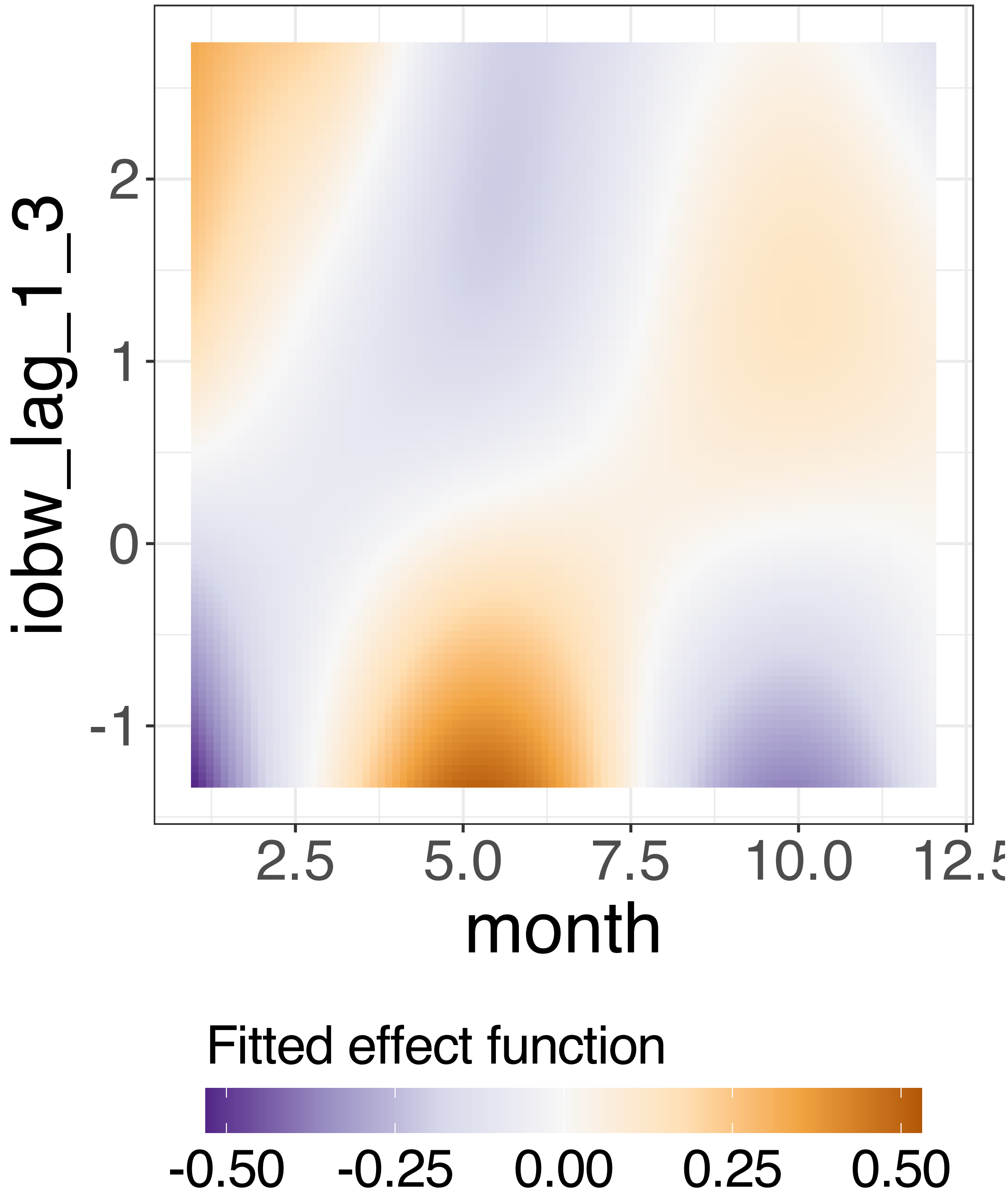


**Supplementary Figure 366.** Decomposition of reconstructed SSANOVA-SVD component function 7.

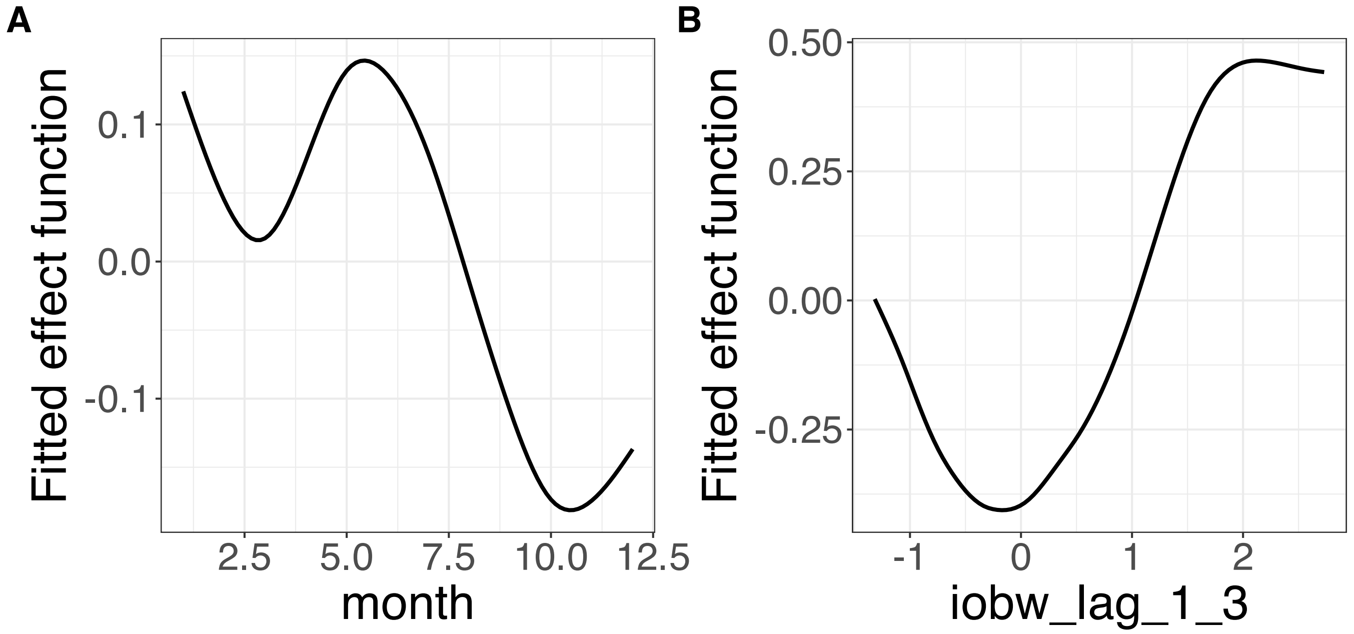


**Supplementary Figure 367.** Decomposition of reconstructed SSANOVA-SVD component function 8. (A) month. (B) iobw_lag_1_3.

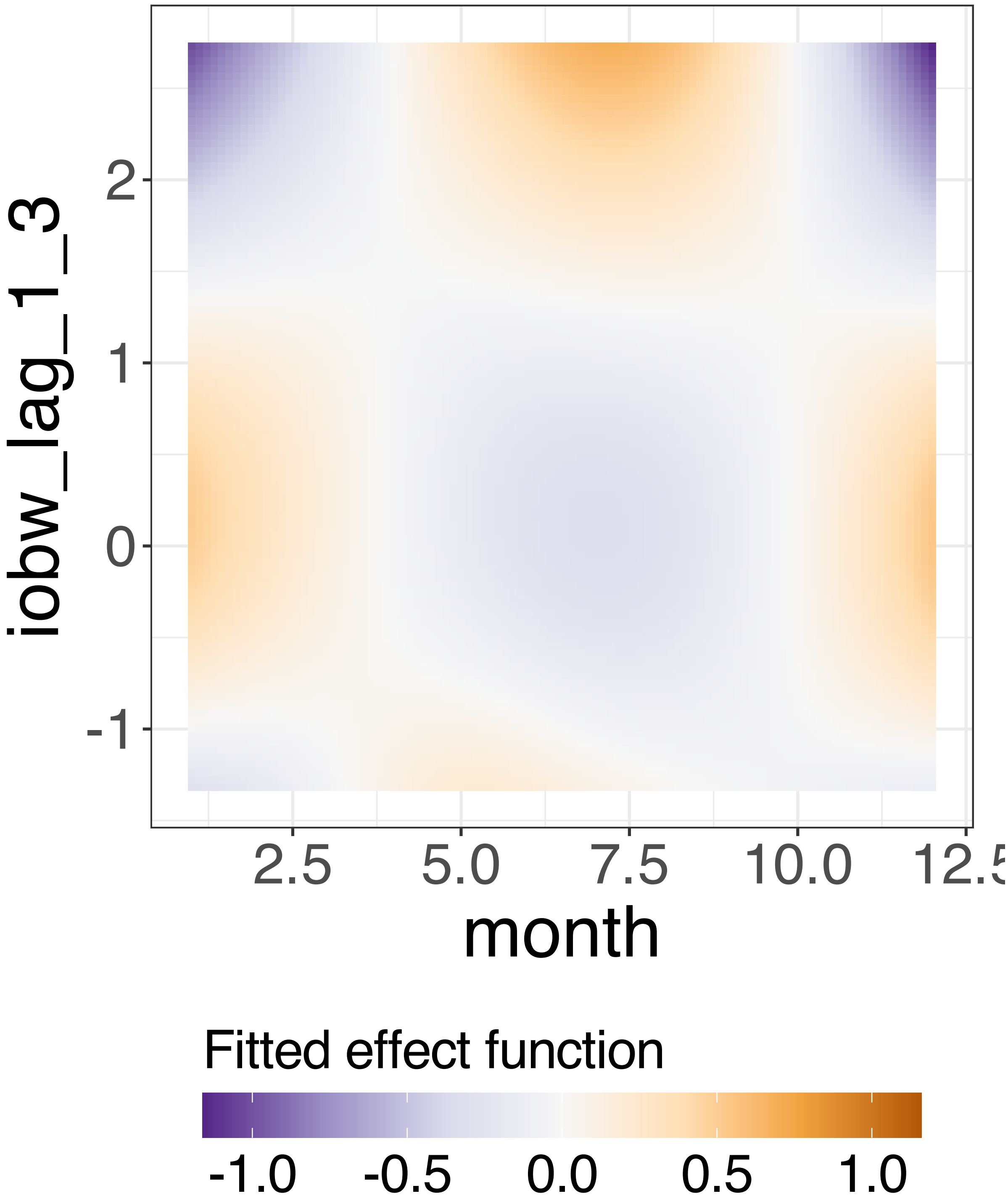


**Supplementary Figure 368.** Decomposition of reconstructed SSANOVA-SVD component function 8.

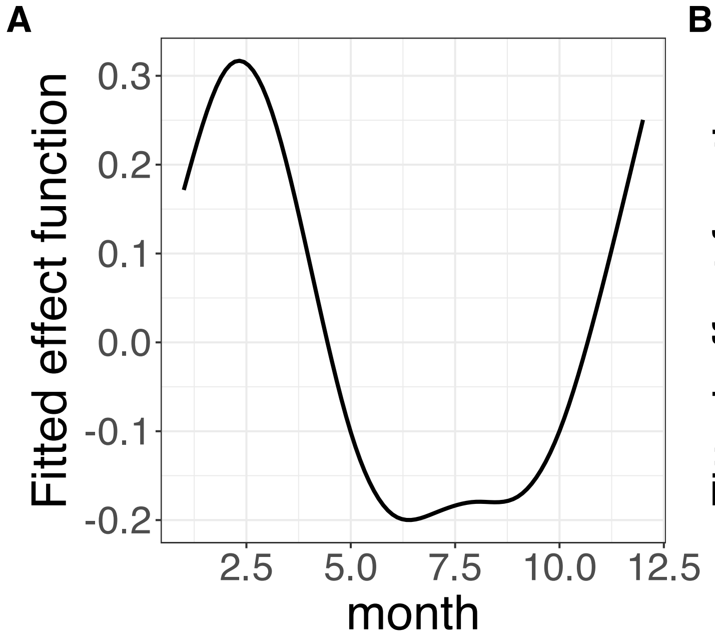


**Supplementary Figure 369.** Decomposition of reconstructed SSANOVA-SVD component function 9. (A) month. (B) iobw_lag_1_3.

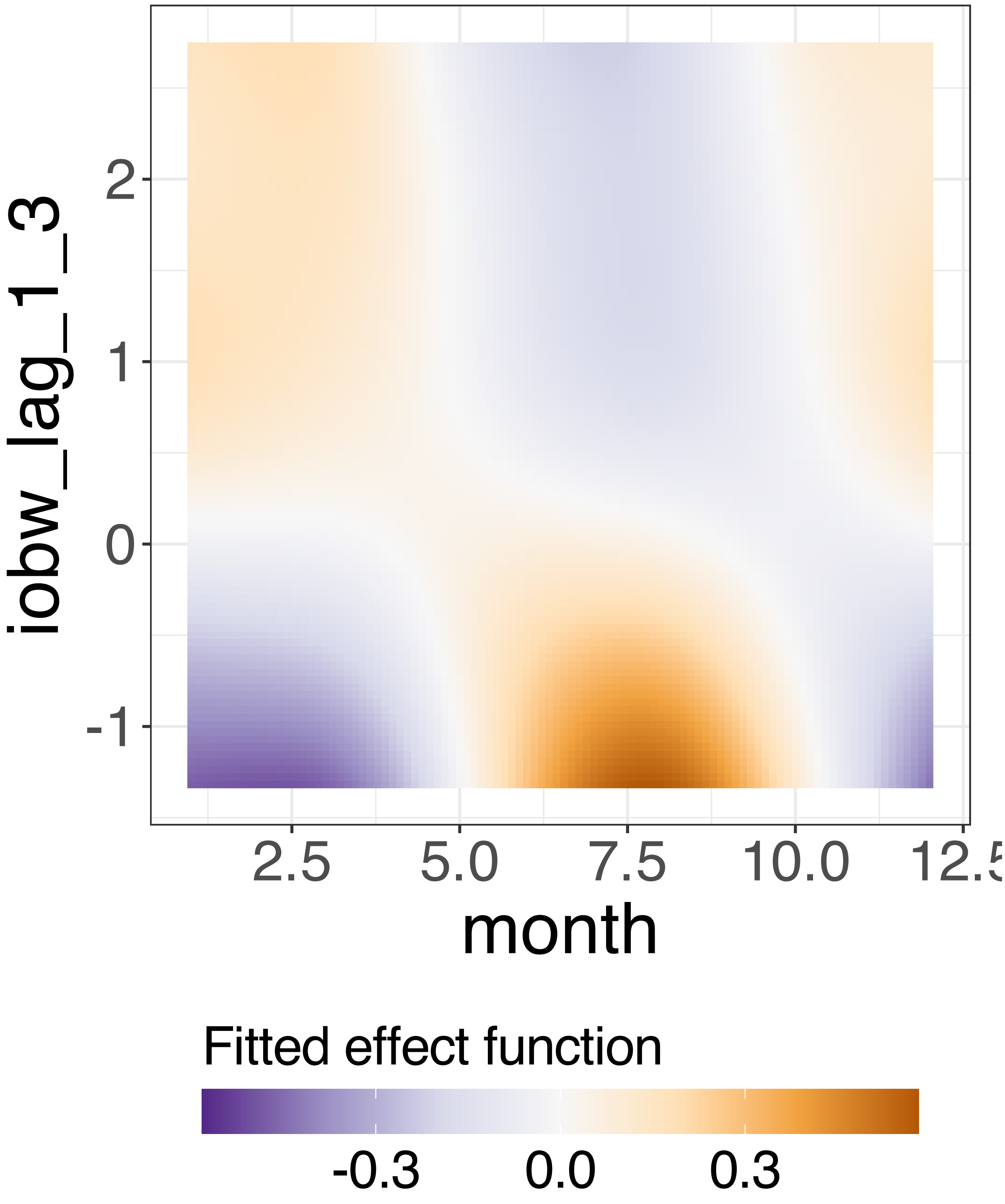


**Supplementary Figure 370.** Decomposition of reconstructed SSANOVA-SVD component function 9.

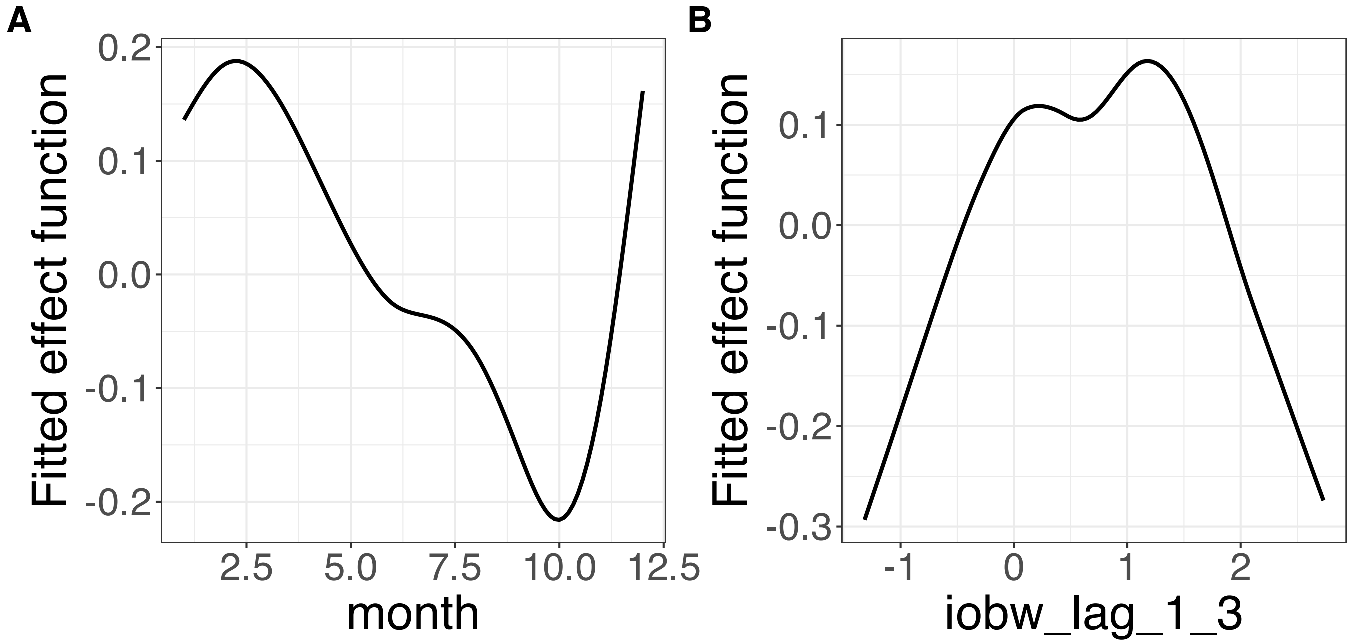


**Supplementary Figure 371.** Decomposition of reconstructed SSANOVA-SVD component function 10. (A) month. (B) iobw_lag_1_3.

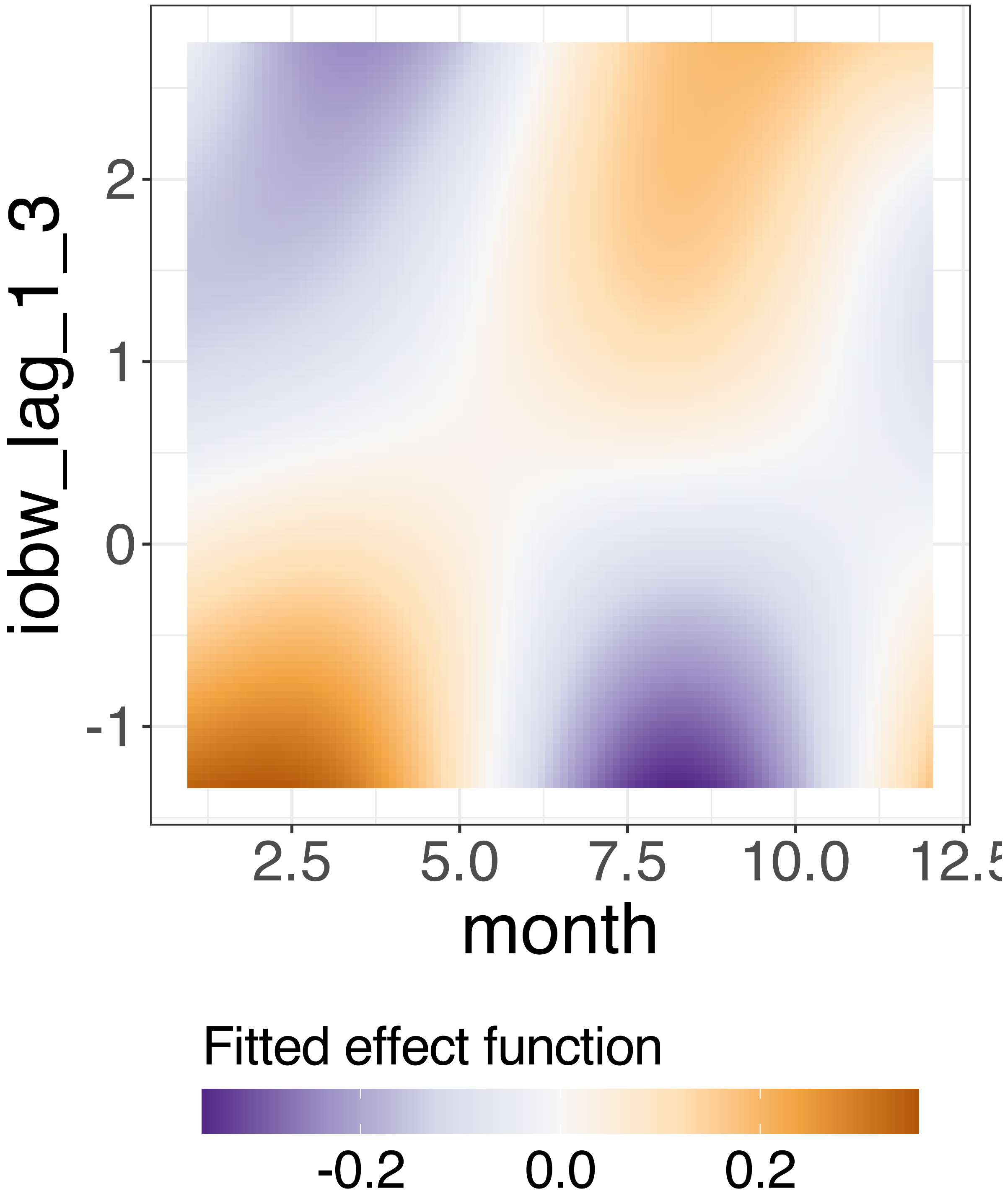


**Supplementary Figure 372.** Decomposition of reconstructed SSANOVA-SVD component function 10.

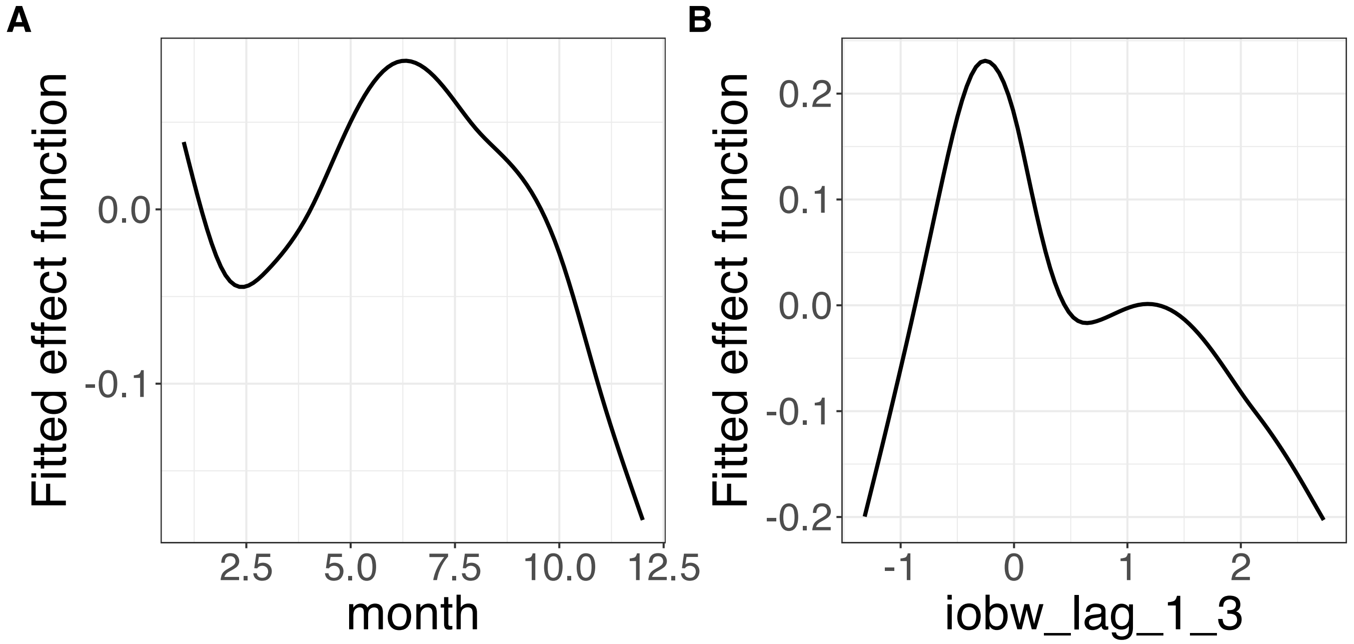


**Supplementary Figure 373.** Decomposition of reconstructed SSANOVA-SVD component function 11. (A) month. (B) iobw_lag_1_3.

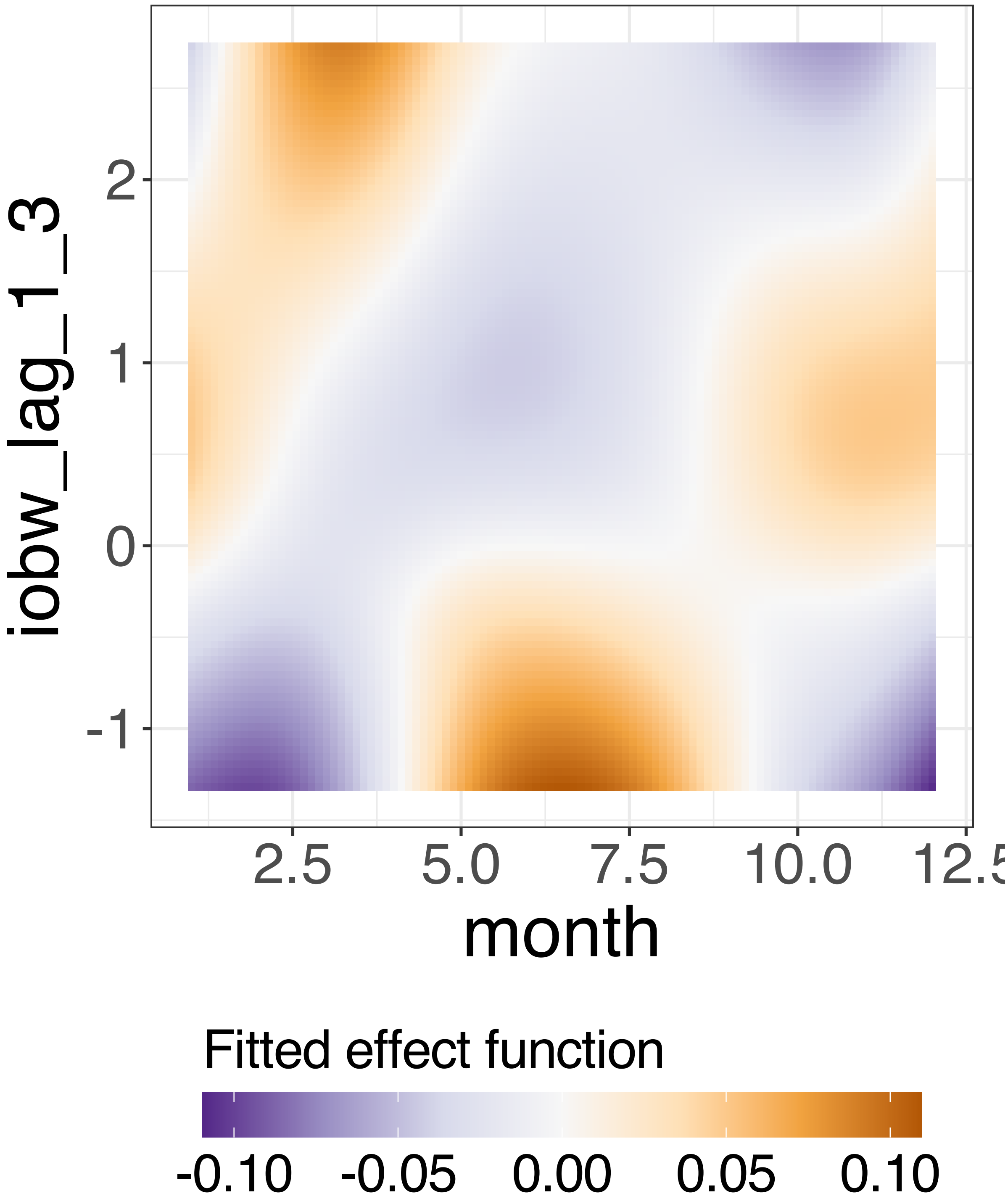


**Supplementary Figure 374.** Decomposition of reconstructed SSANOVA-SVD component function 11.

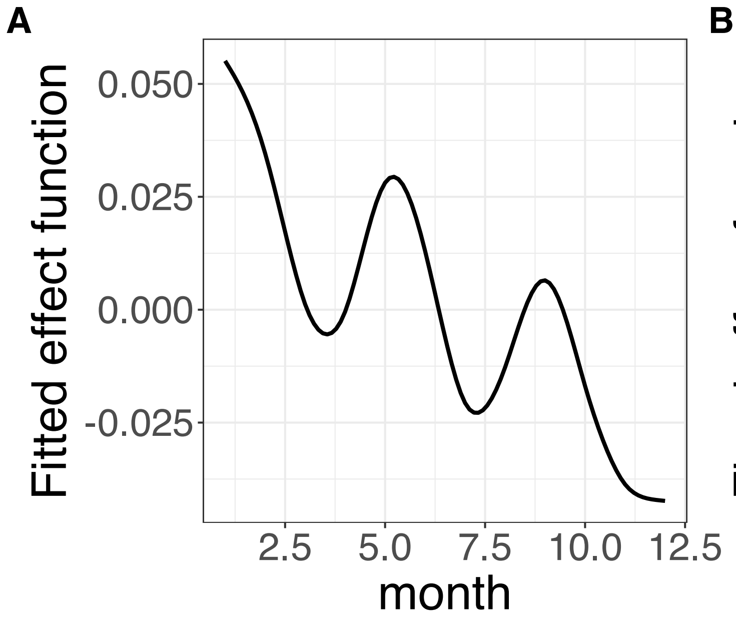


**Supplementary Figure 375.** Decomposition of reconstructed SSANOVA-SVD component function 12. (A) month. (B) iobw_lag_1_3.

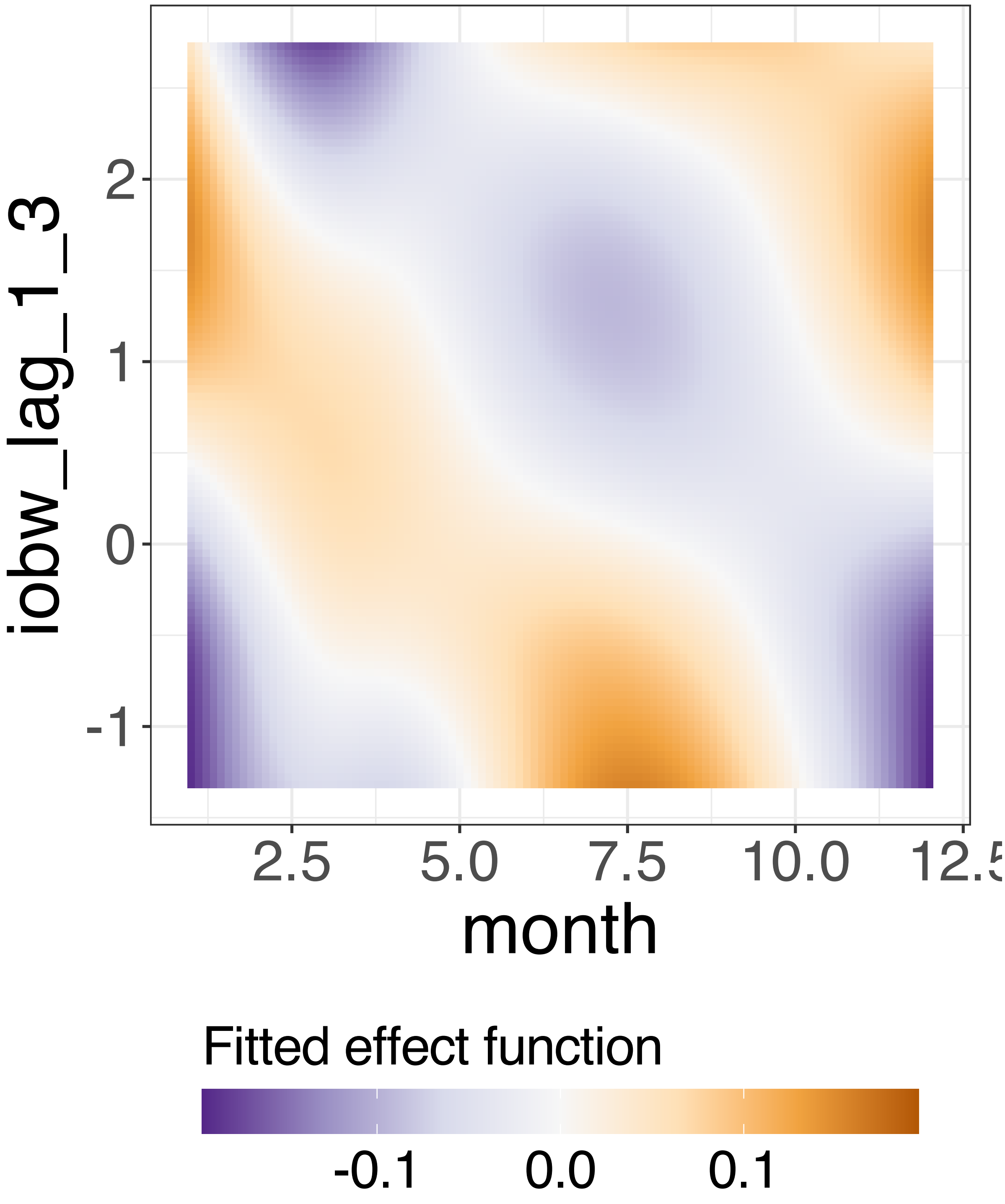


**Supplementary Figure 376.** Decomposition of reconstructed SSANOVA-SVD component function 12.

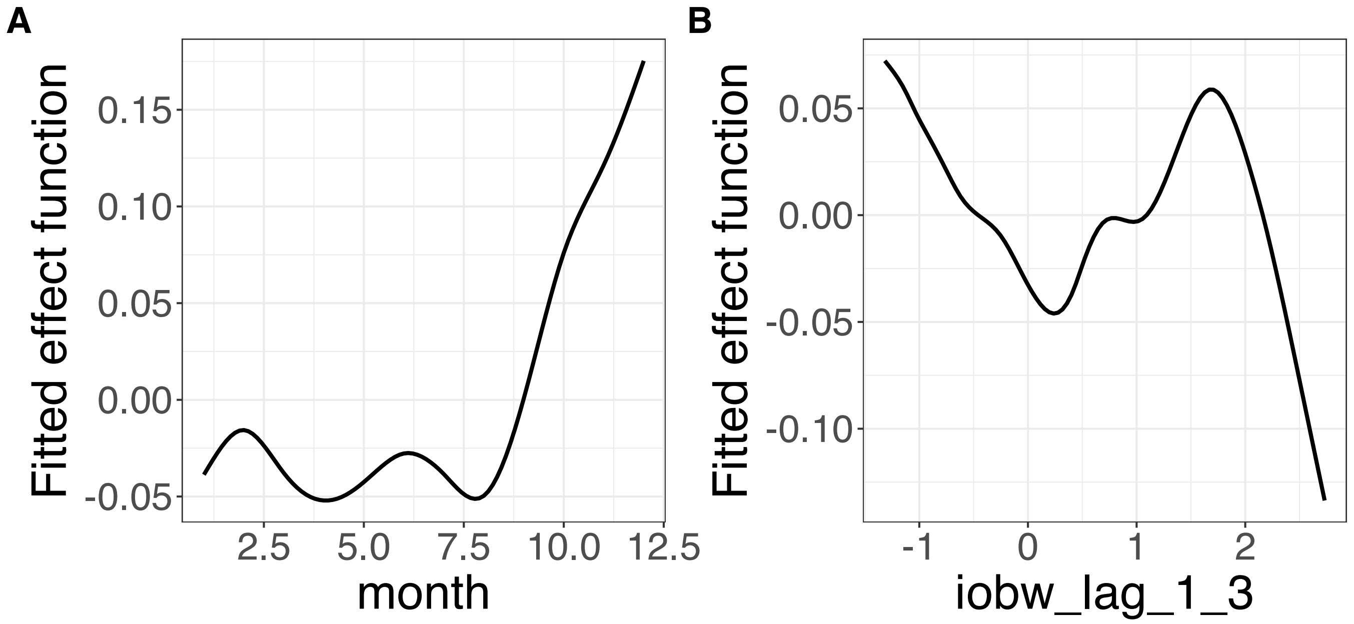


**Supplementary Figure 377.** Decomposition of reconstructed SSANOVA-SVD component function 13. (A) month. (B) iobw_lag_1_3.

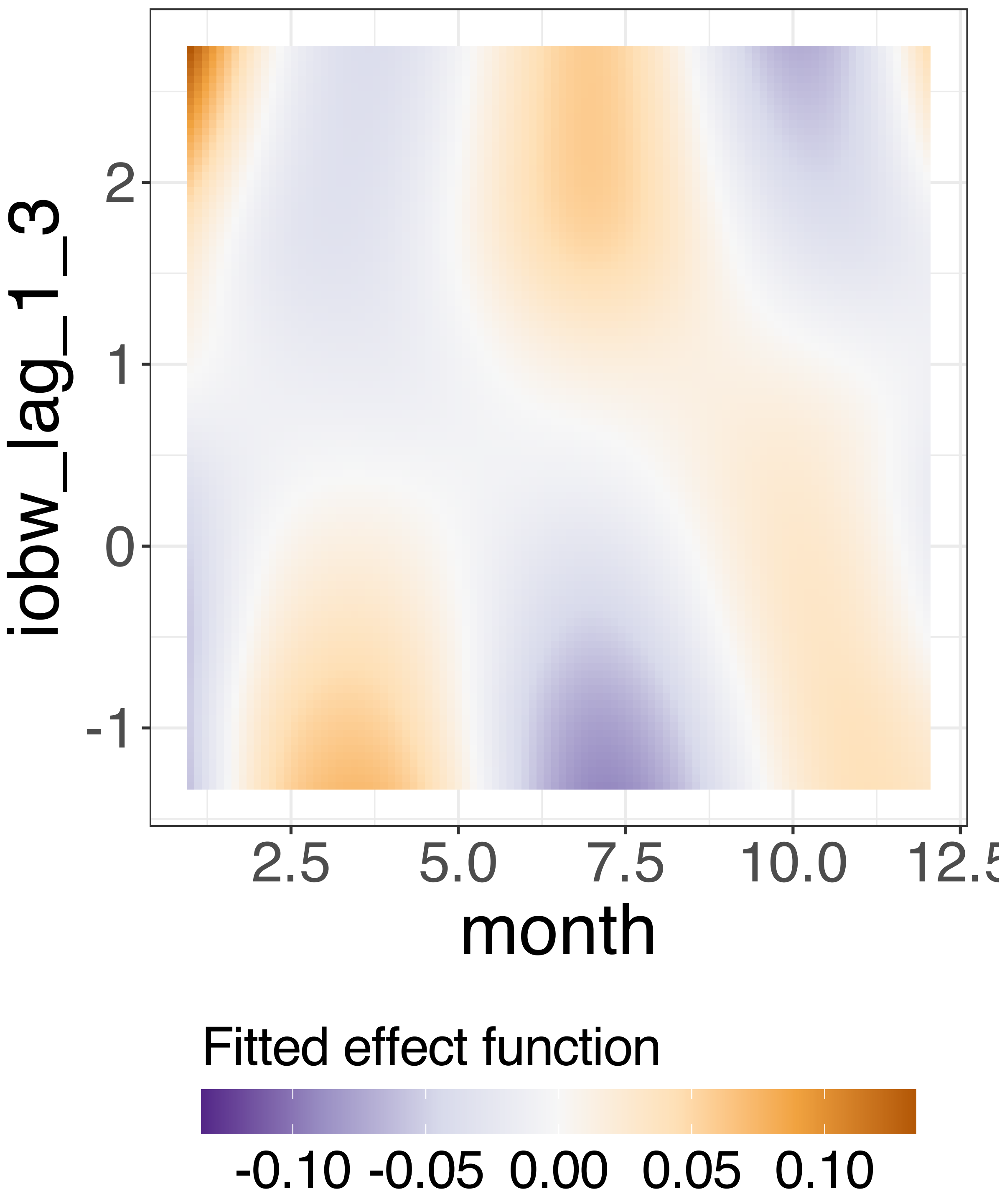


**Supplementary Figure 378.** Decomposition of reconstructed SSANOVA-SVD component function 13.

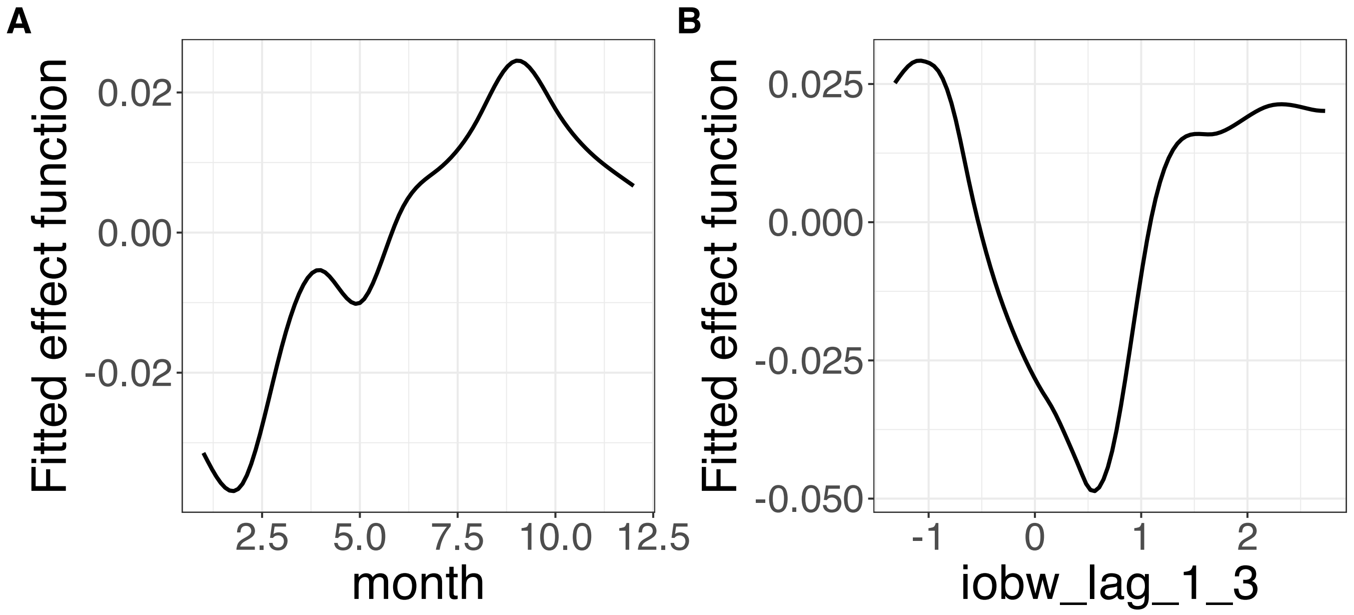


**Supplementary Figure 379.** Decomposition of reconstructed SSANOVA-SVD component function 14. (A) month. (B) iobw_lag_1_3.

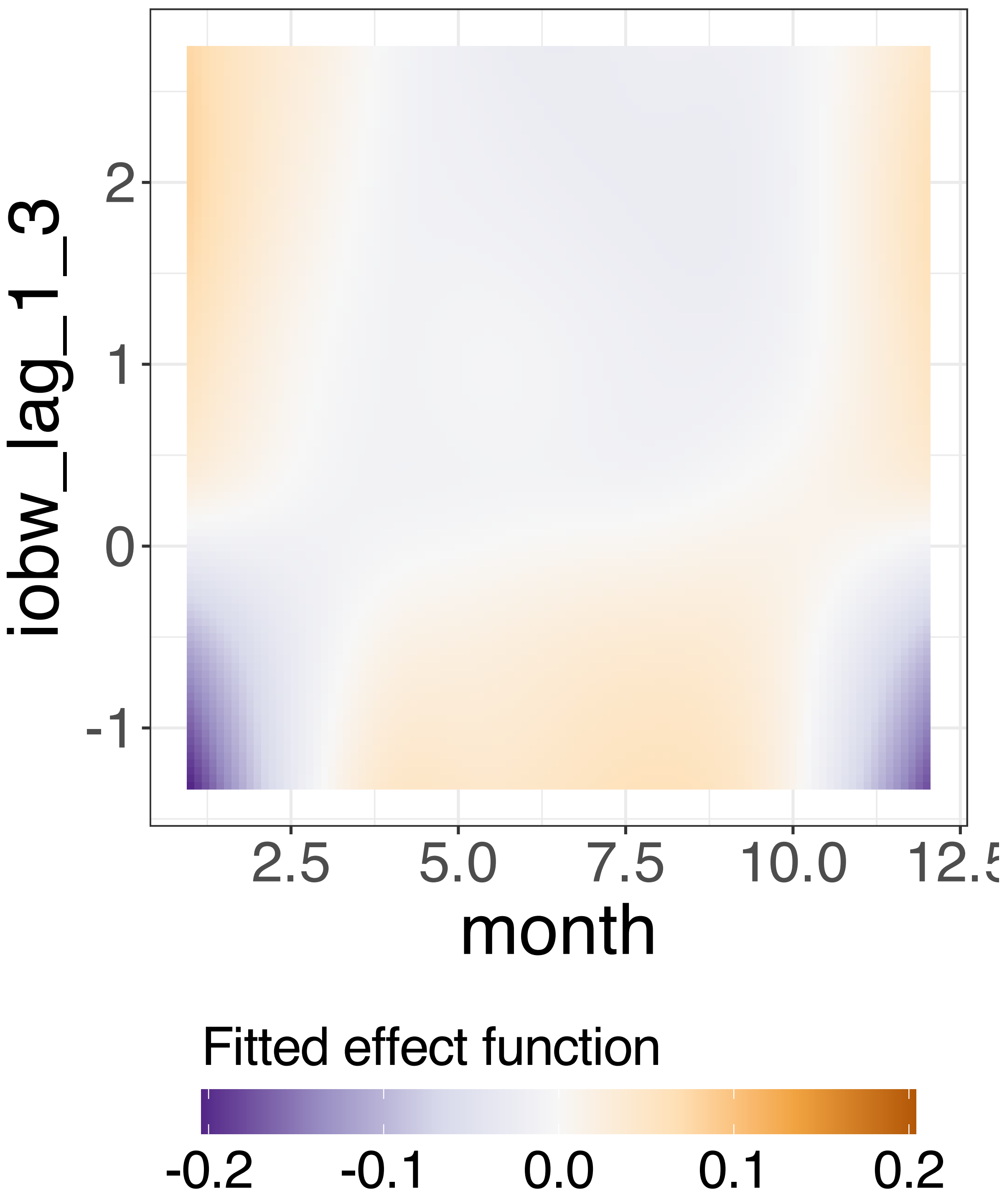


**Supplementary Figure 380.** Decomposition of reconstructed SSANOVA-SVD component function 14.

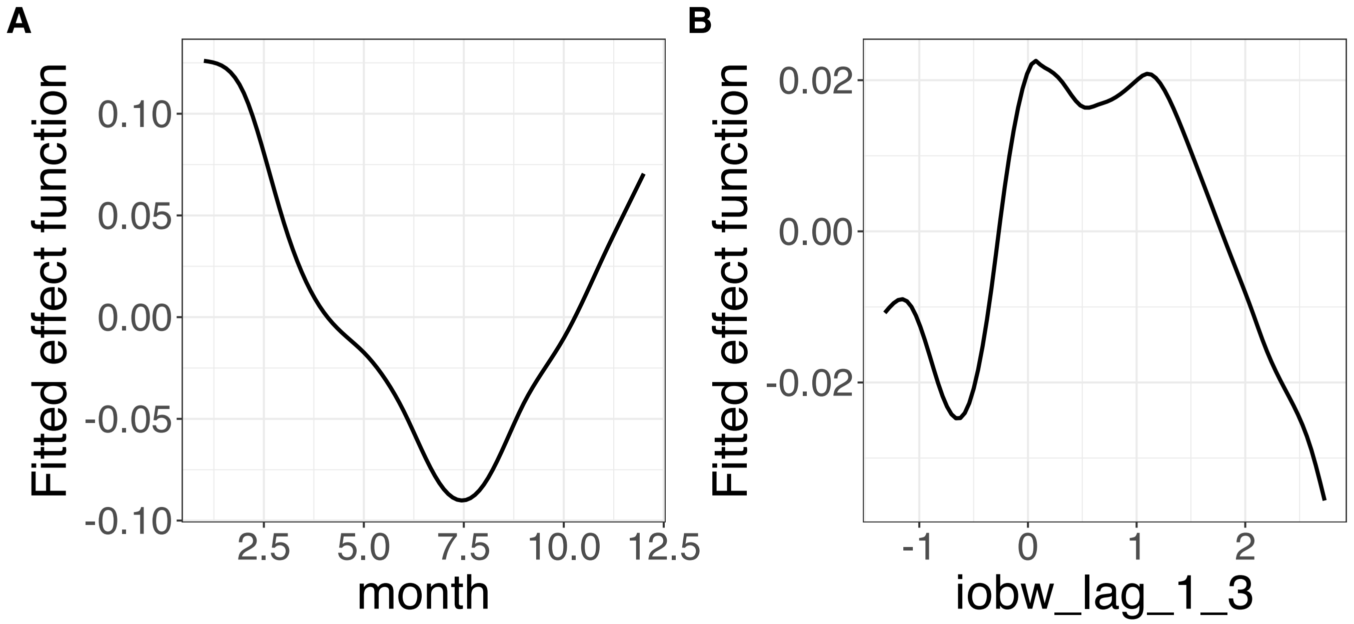


**Supplementary Figure 381.** Decomposition of reconstructed SSANOVA-SVD component function 15. (A) month. (B) iobw_lag_1_3.

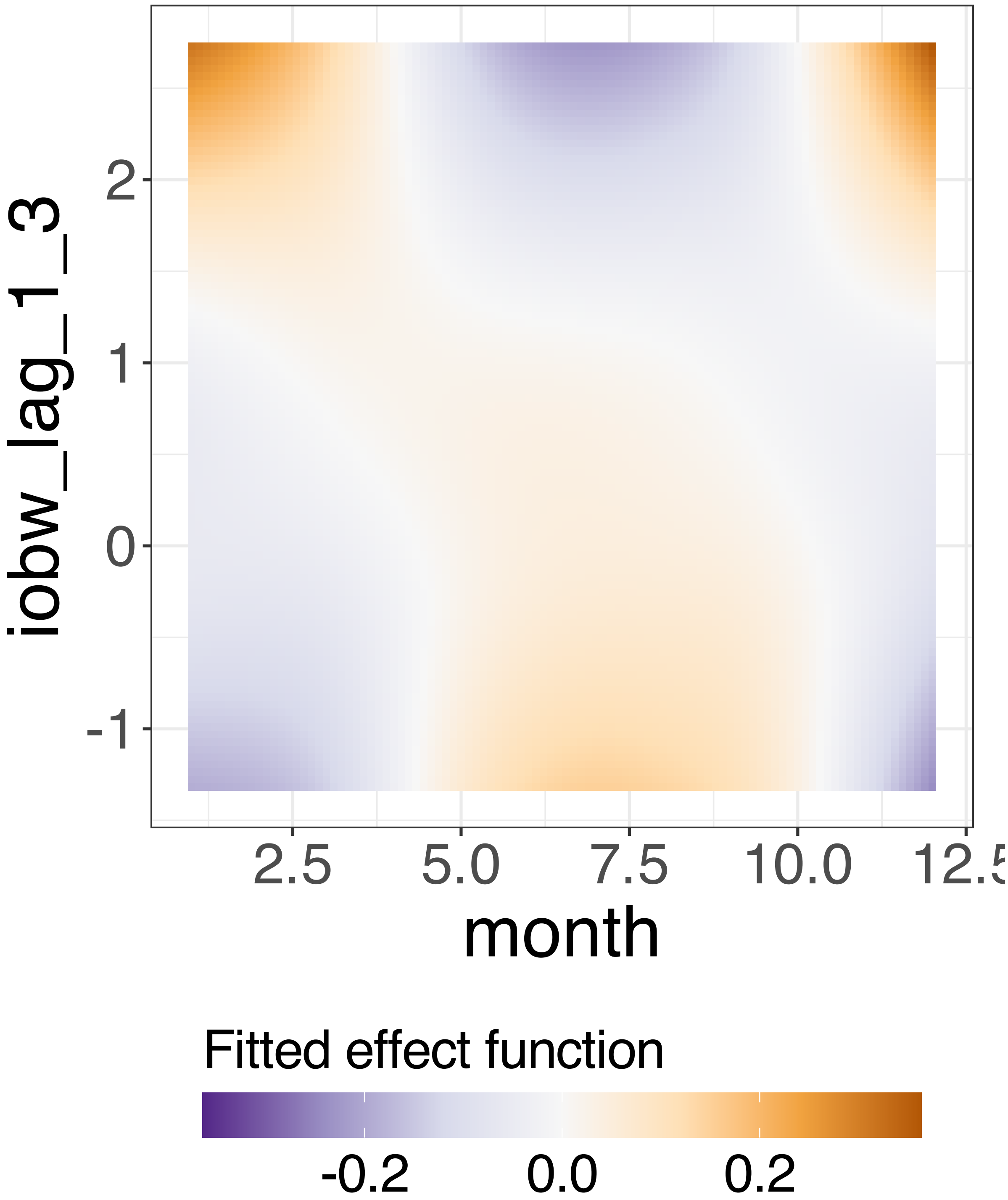


**Supplementary Figure 382.** Decomposition of reconstructed SSANOVA-SVD component function 15.

PCA

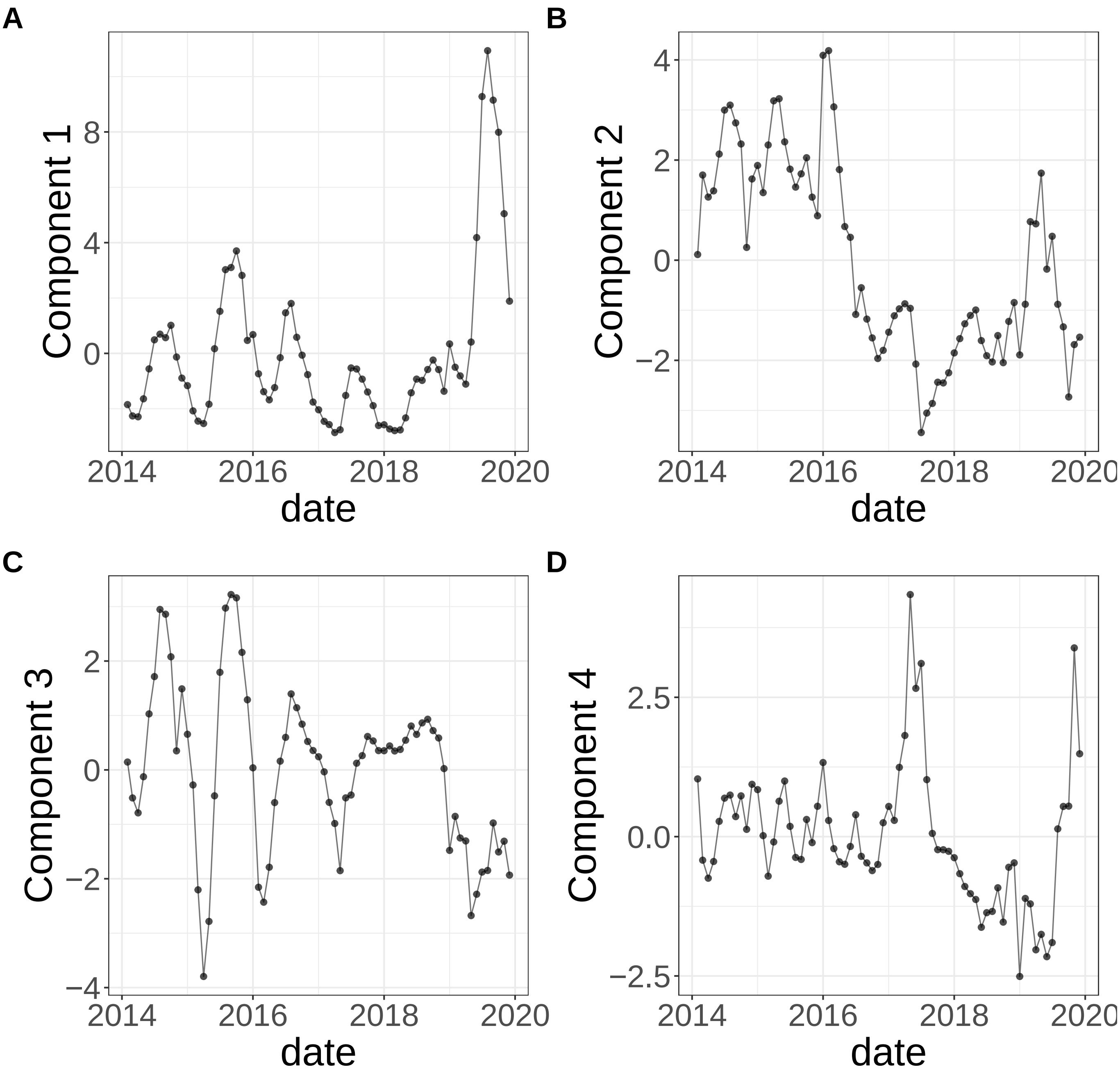


**Supplementary Figure 383.** Components for PCA. (A) Component 1. (B) Component 2. (C) Component 3. (D) Component 4.

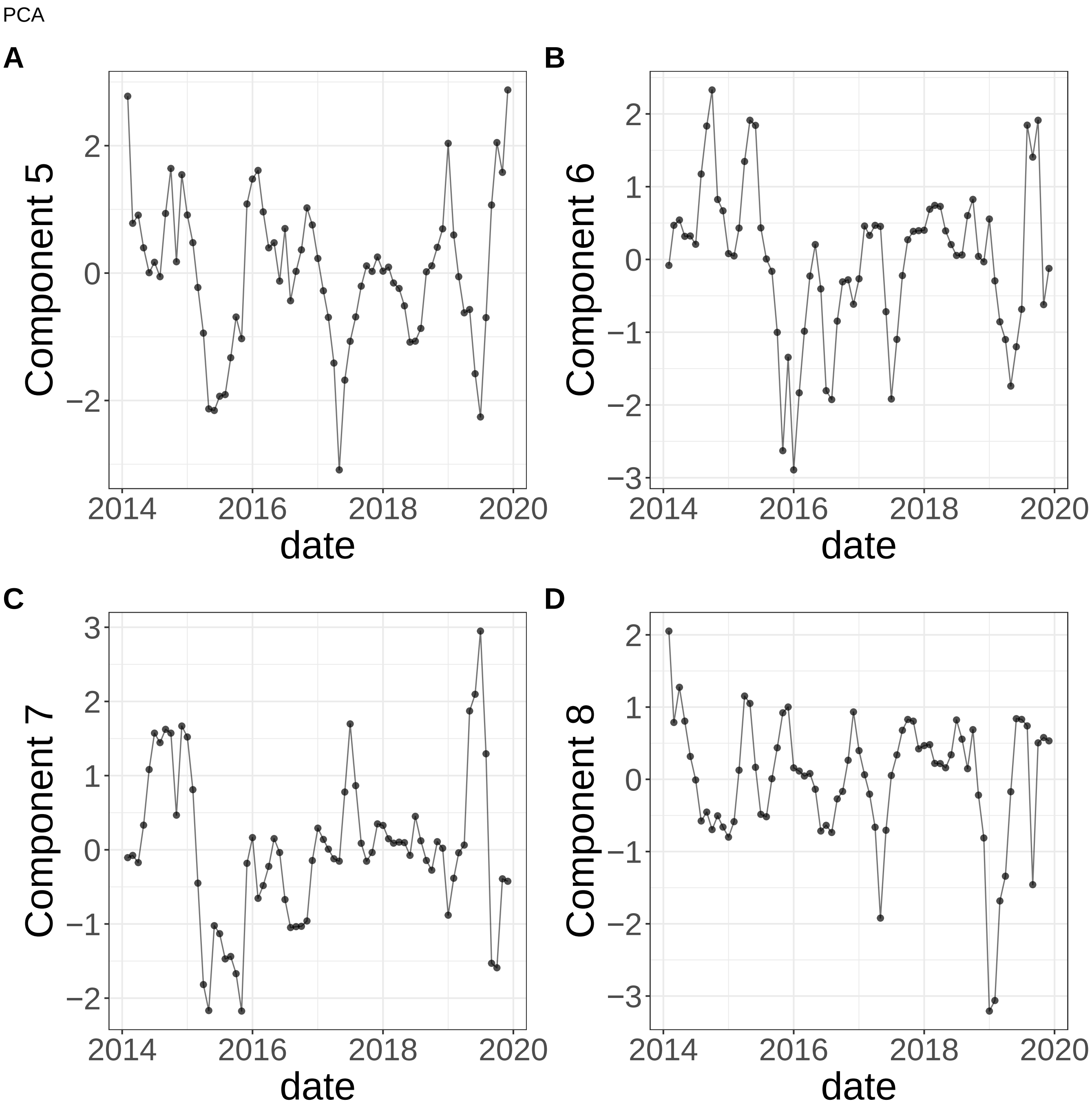


**Supplementary Figure 384.** Components for PCA. (A) Component 5. (B) Component 6. (C) Component 7. (D) Component 8.

PCA

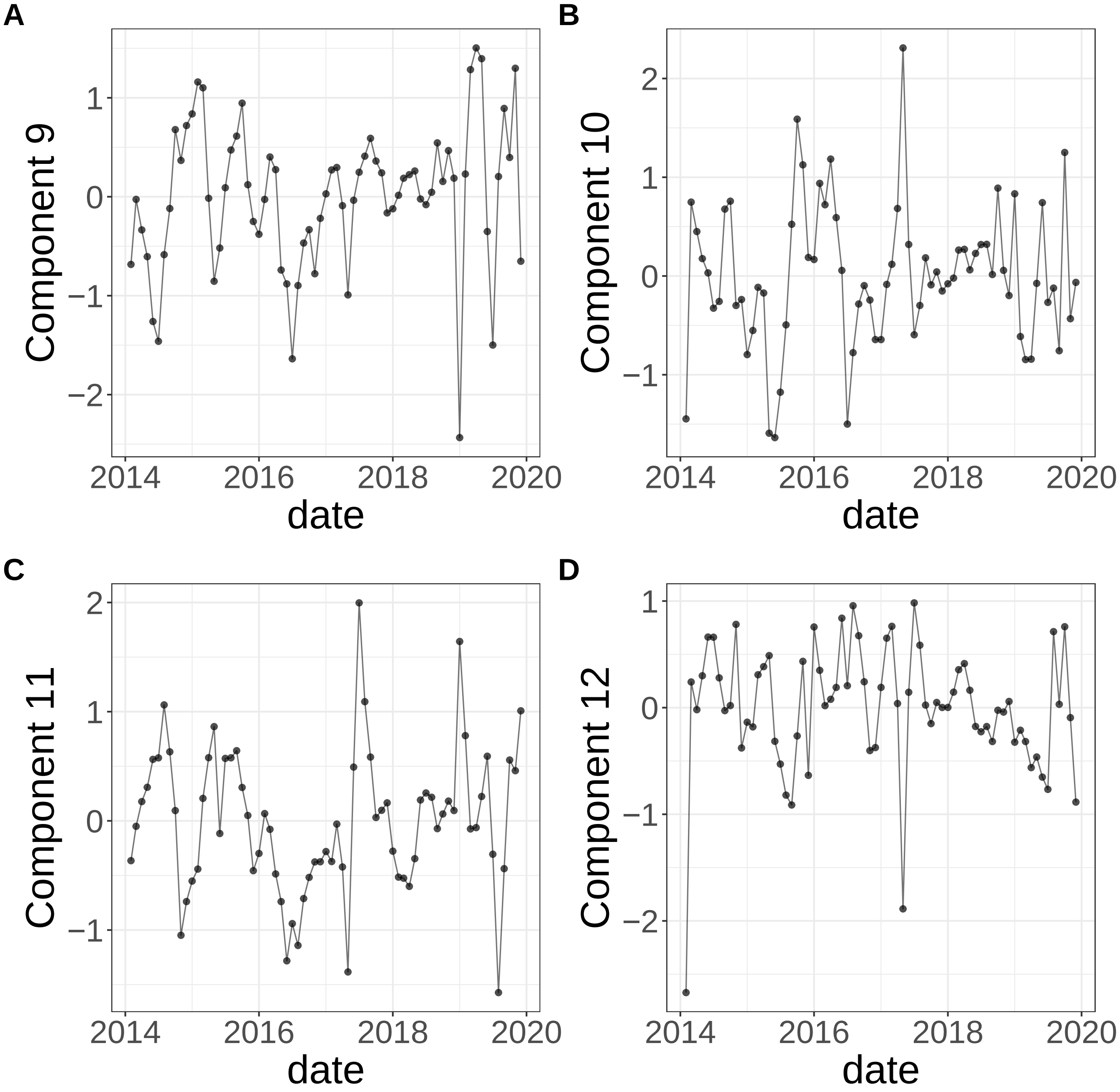


**Supplementary Figure 385.** Components for PCA. (A) Component 9. (B) Component 10. (C) Component 11. (D) Component 12.

PCA

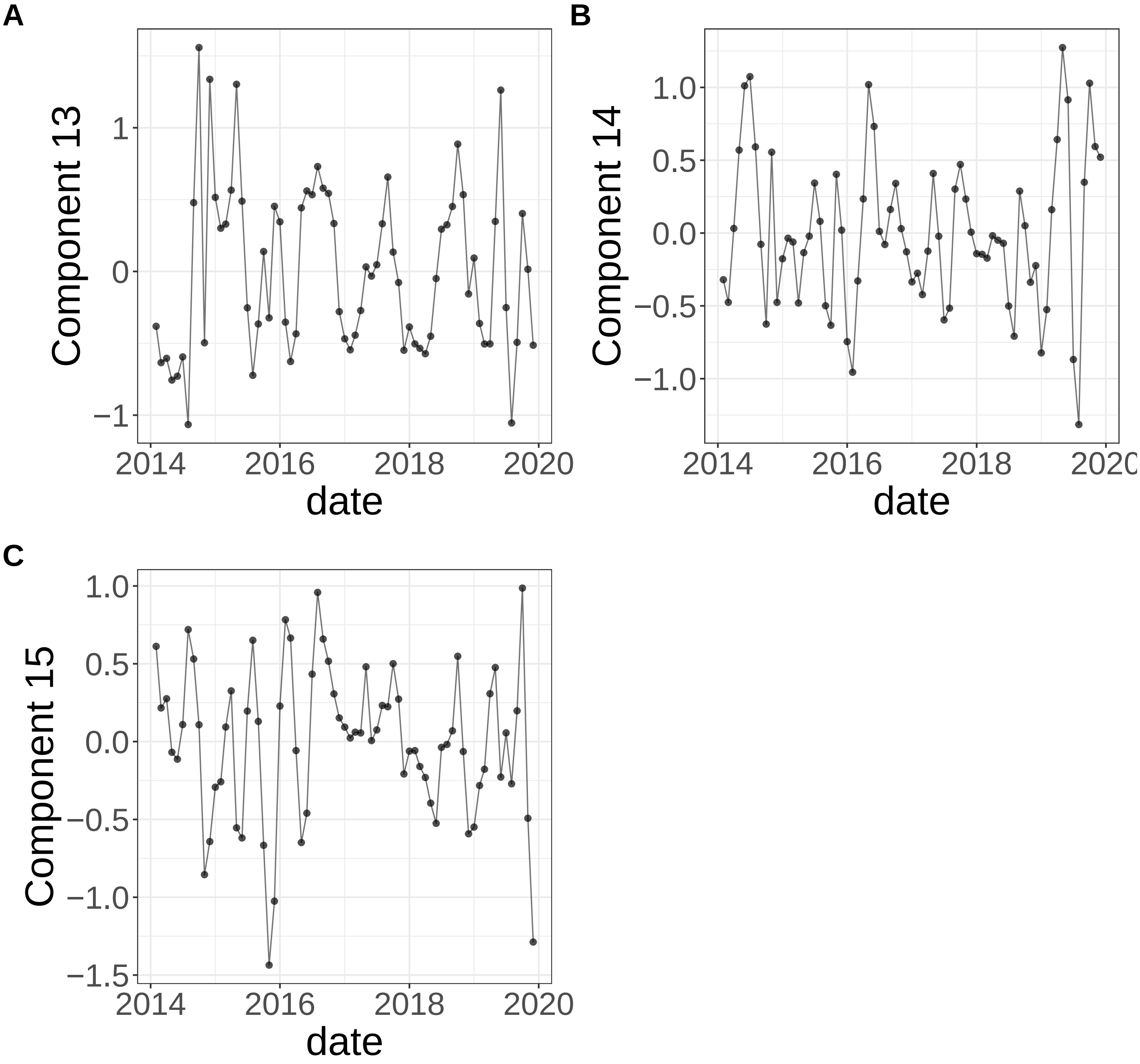


**Supplementary Figure 386.** Components for PCA. (A) Component 13. (B) Component 14. (C) Component 15.

NMF

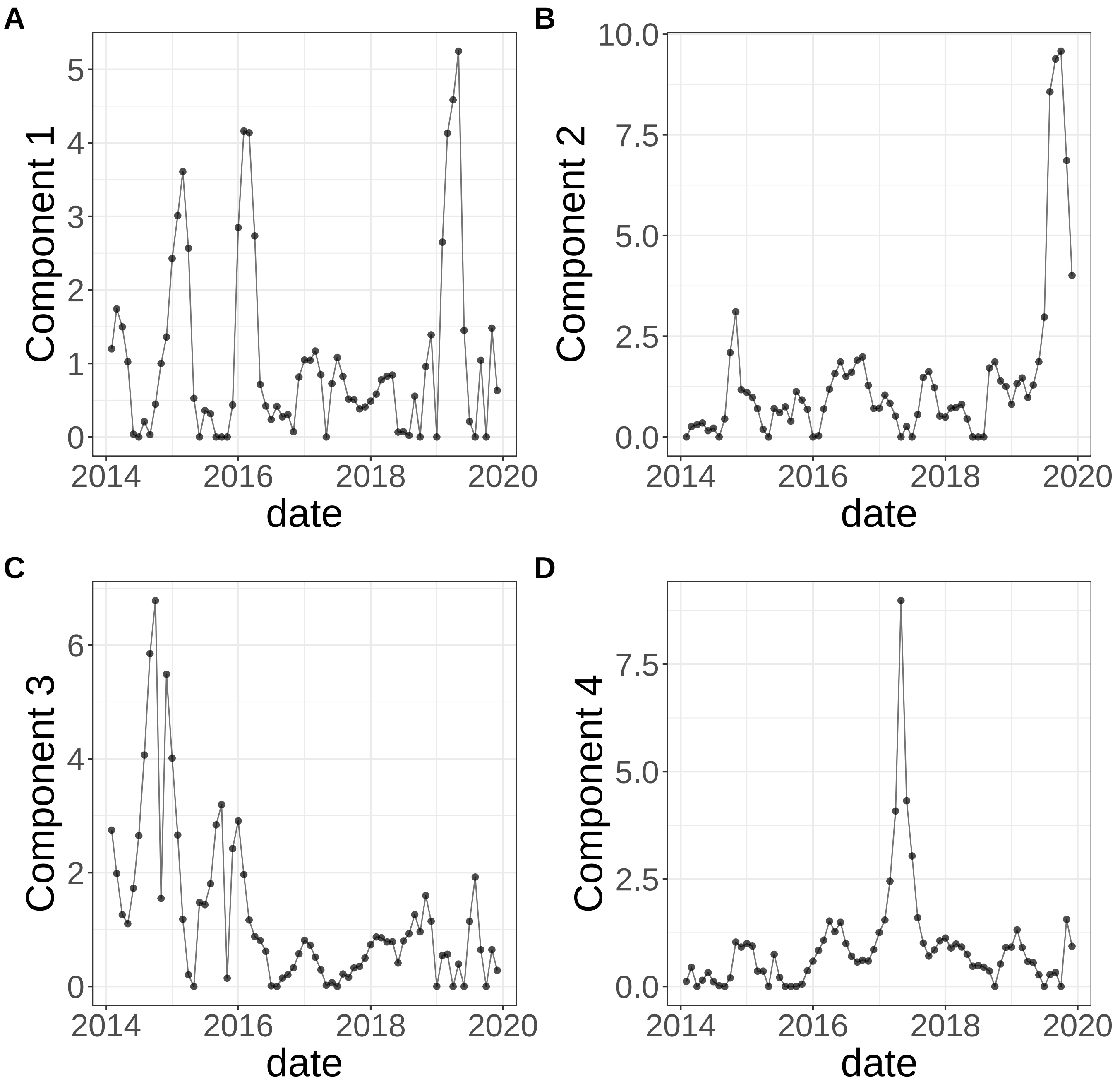


**Supplementary Figure 387.** Components for NMF. (A) Component 1. (B) Component 2. (C) Component 3. (D) Component 4.

NMF

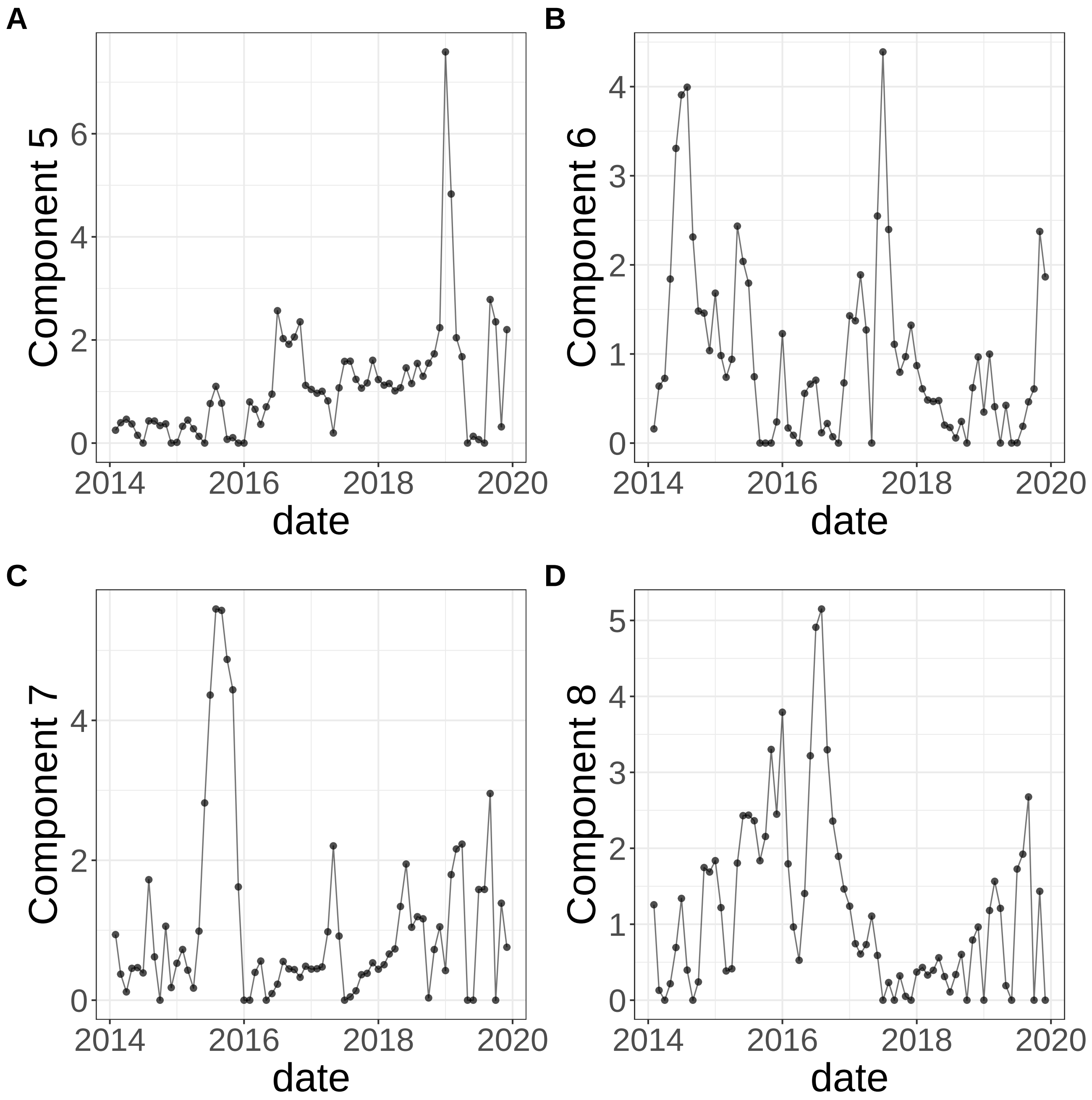


**Supplementary Figure 388.** Components for NMF. (A) Component 5. (B) Component 6. (C) Component 7. (D) Component 8.

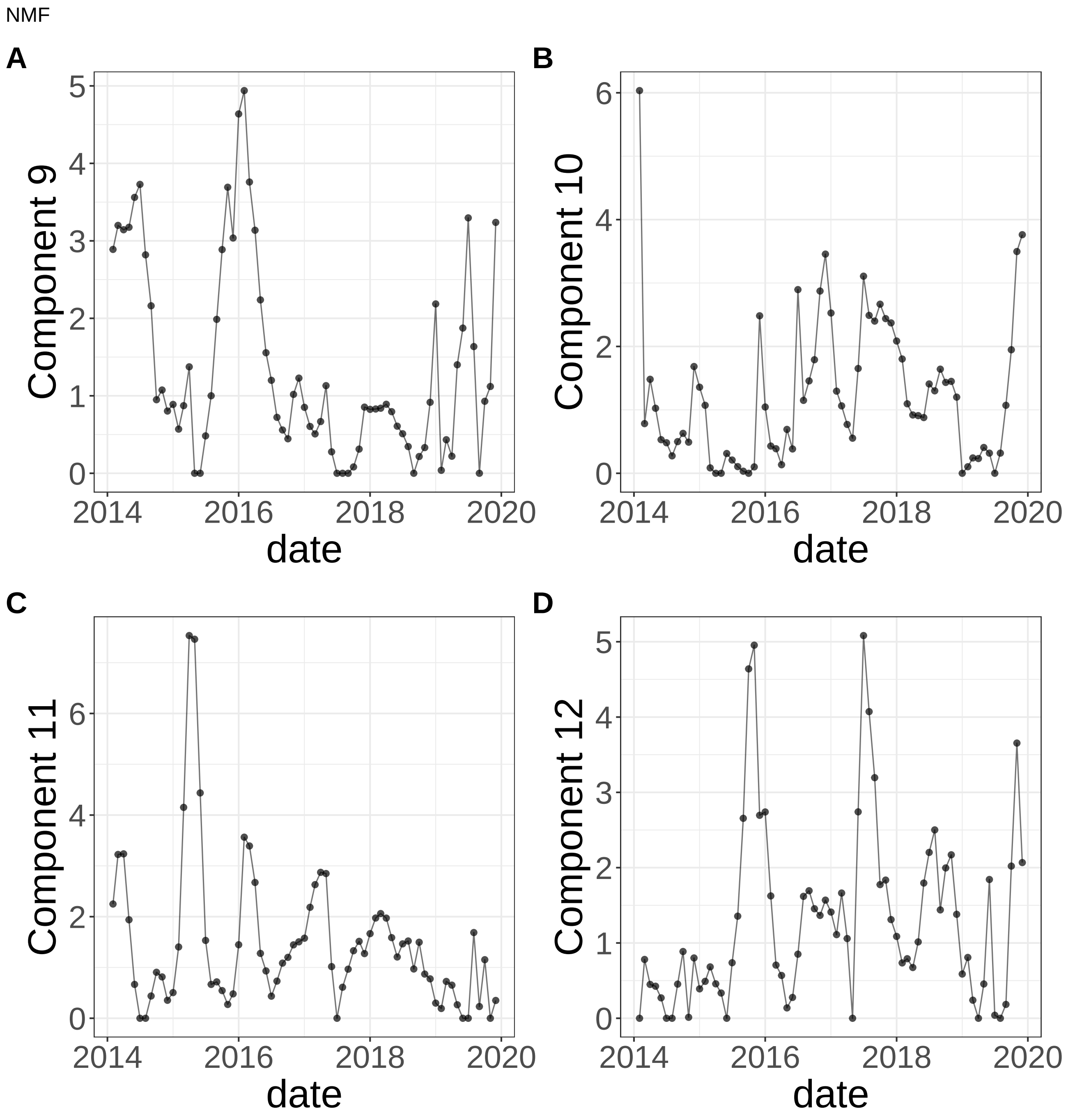


**Supplementary Figure 389.** Components for NMF. (A) Component 9. (B) Component 10. (C) Component 11. (D) Component 12.

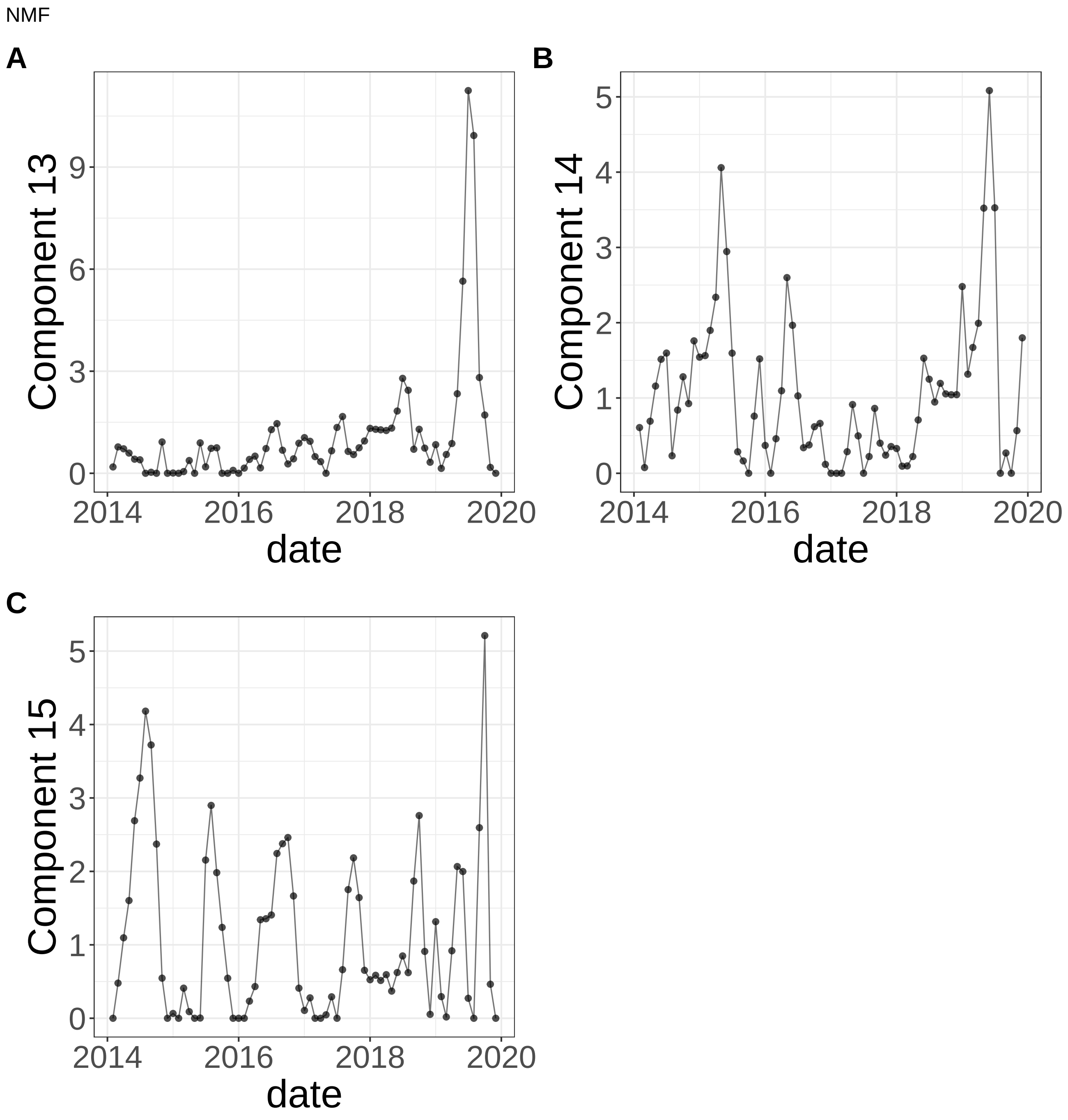


**Supplementary Figure 390.** Components for NMF. (A) Component 13. (B) Component 14. (C) Component 15.

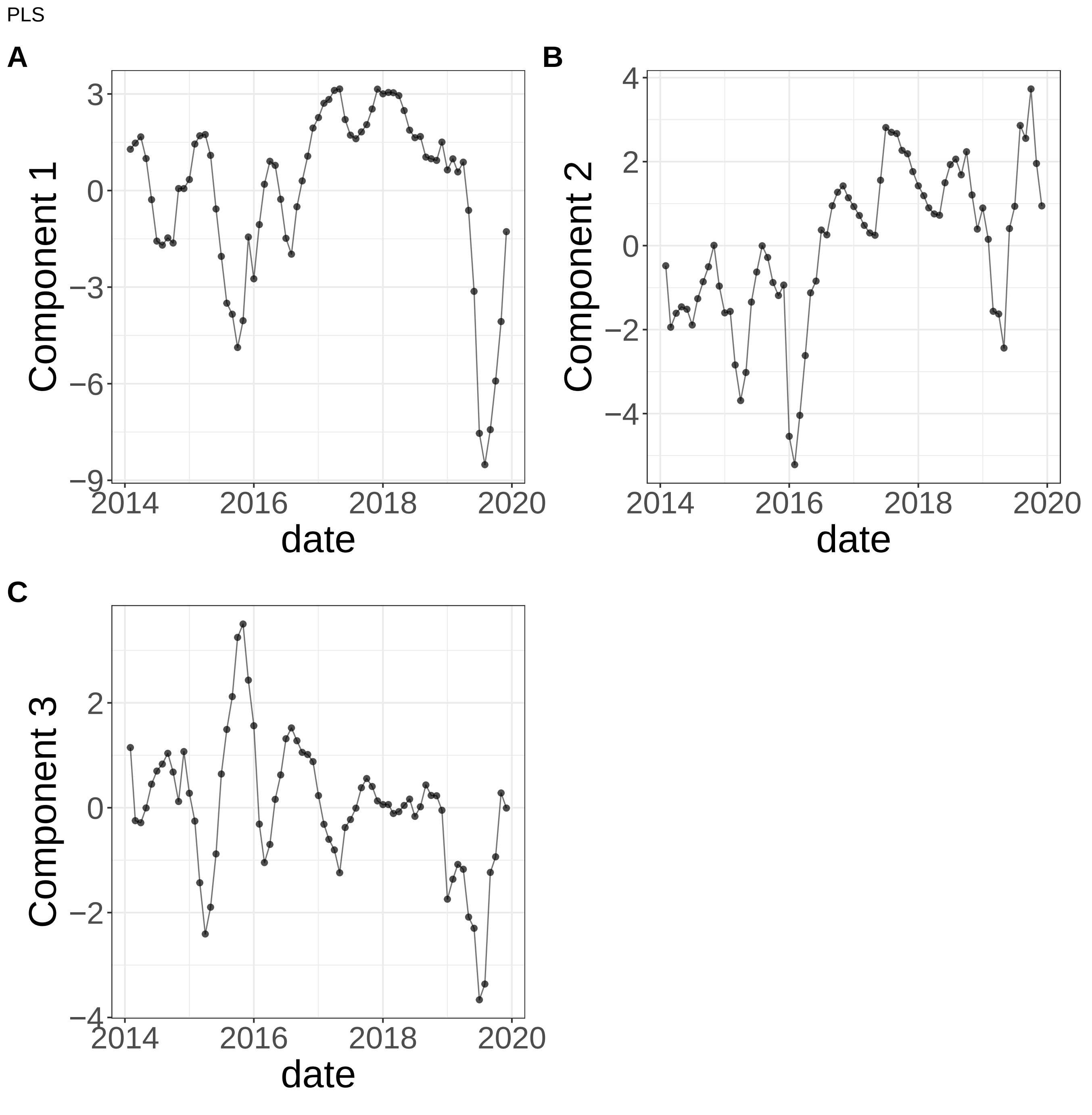


**Supplementary Figure 391.** Components for PLS. (A) Component 1. (B) Component 2. (C) Component 3.

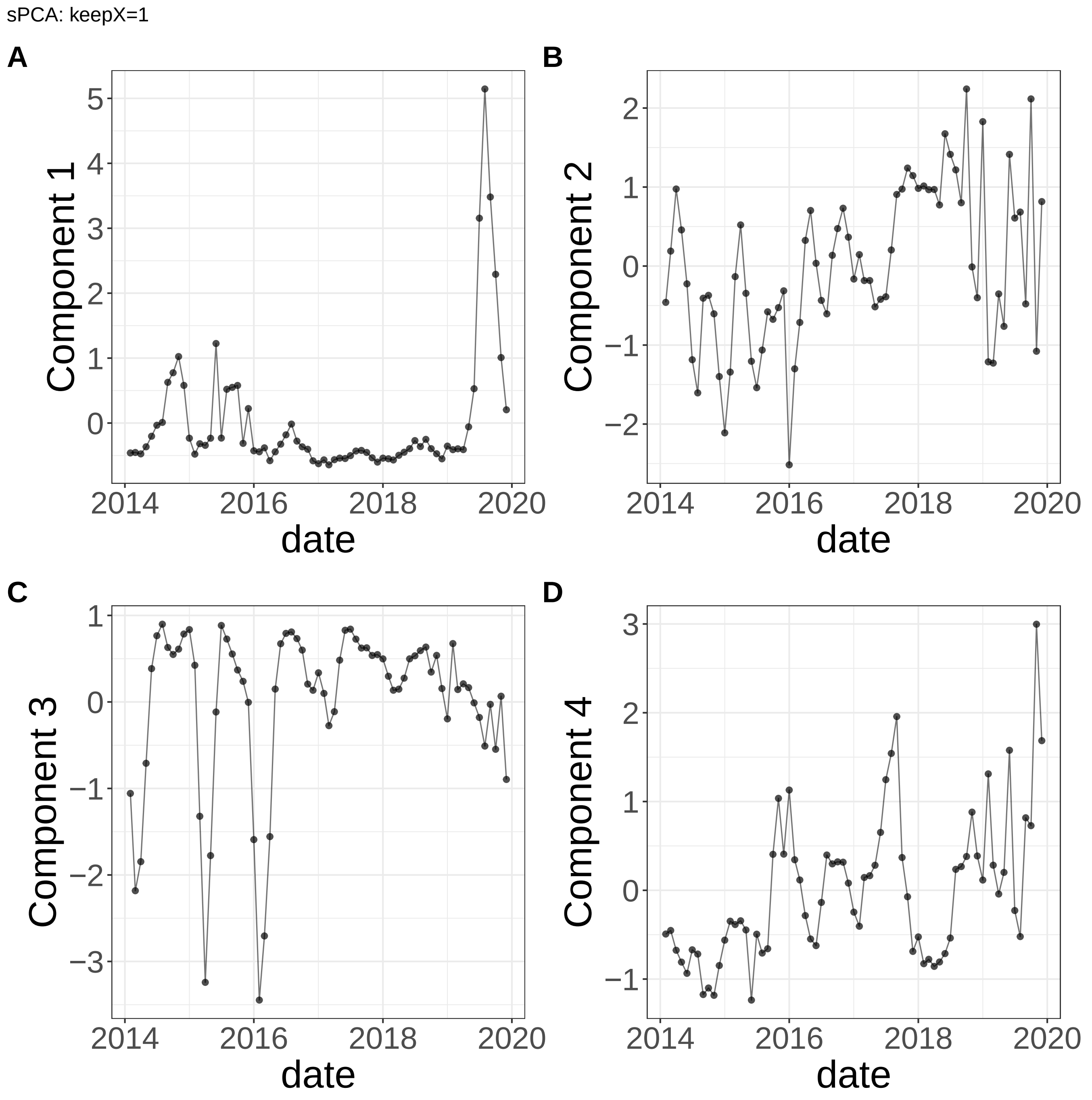


**Supplementary Figure 392.** Components for sPCA: keepX=1. (A) Component 1. (B) Component 2. (C) Component 3. (D) Component 4.

sPCA: keepX=1

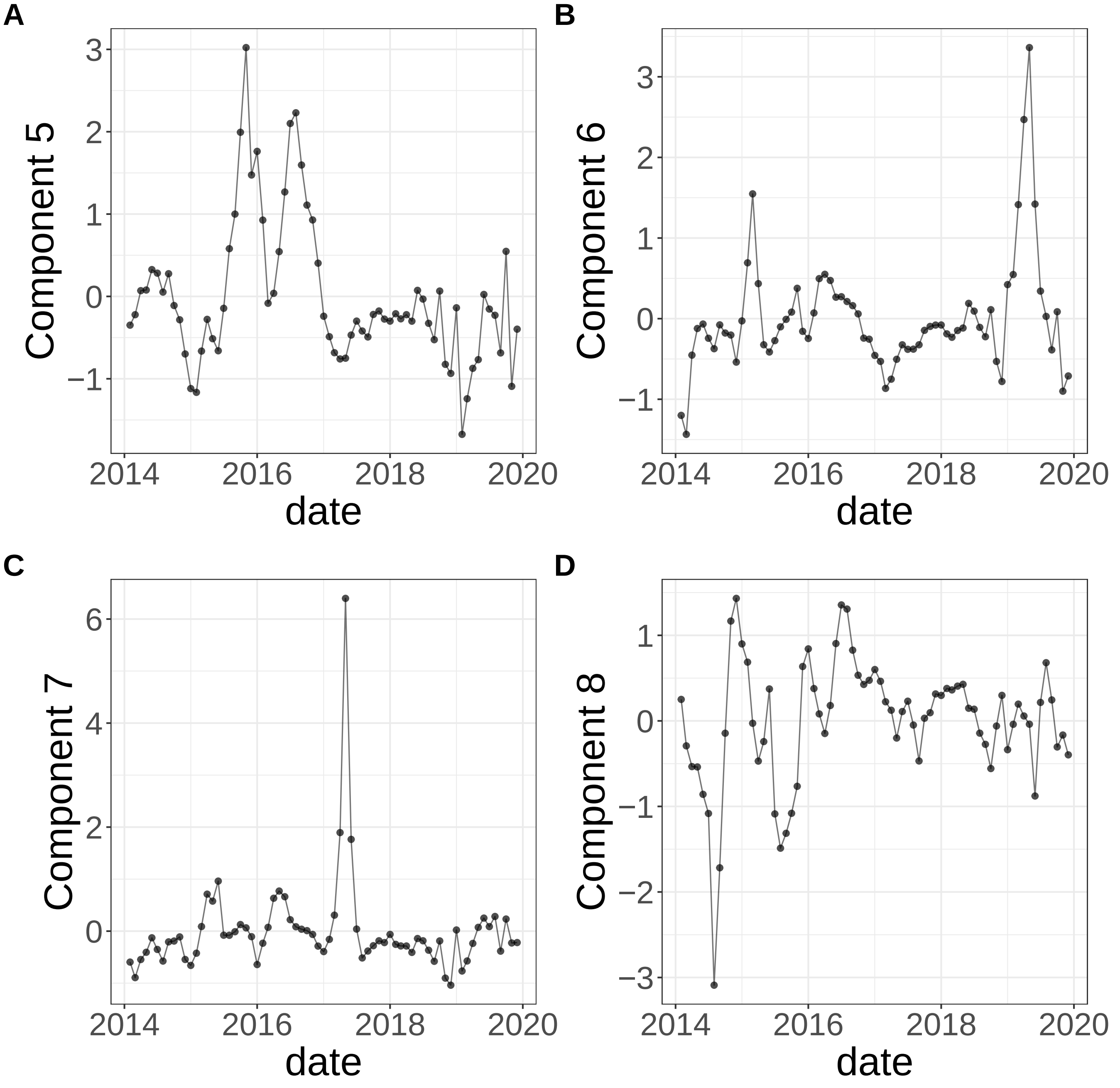


**Supplementary Figure 393.** Components for sPCA: keepX=1. (A) Component 5. (B) Component 6. (C) Component 7. (D) Component 8.

sPCA: keepX=1

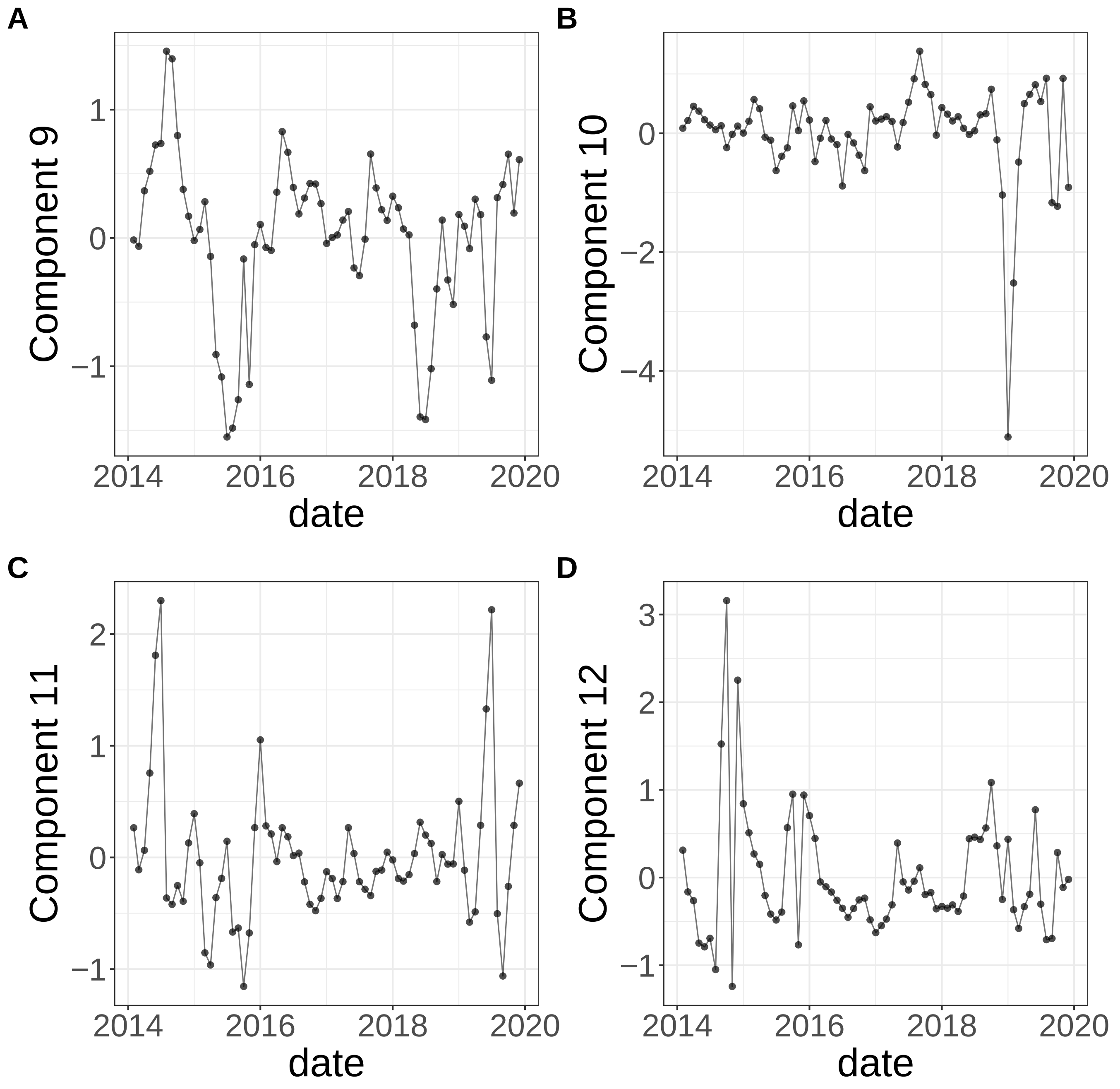


**Supplementary Figure 394.** Components for sPCA: keepX=1. (A) Component 9. (B) Component 10. (C) Component 11. (D) Component 12.

sPCA: keepX=1

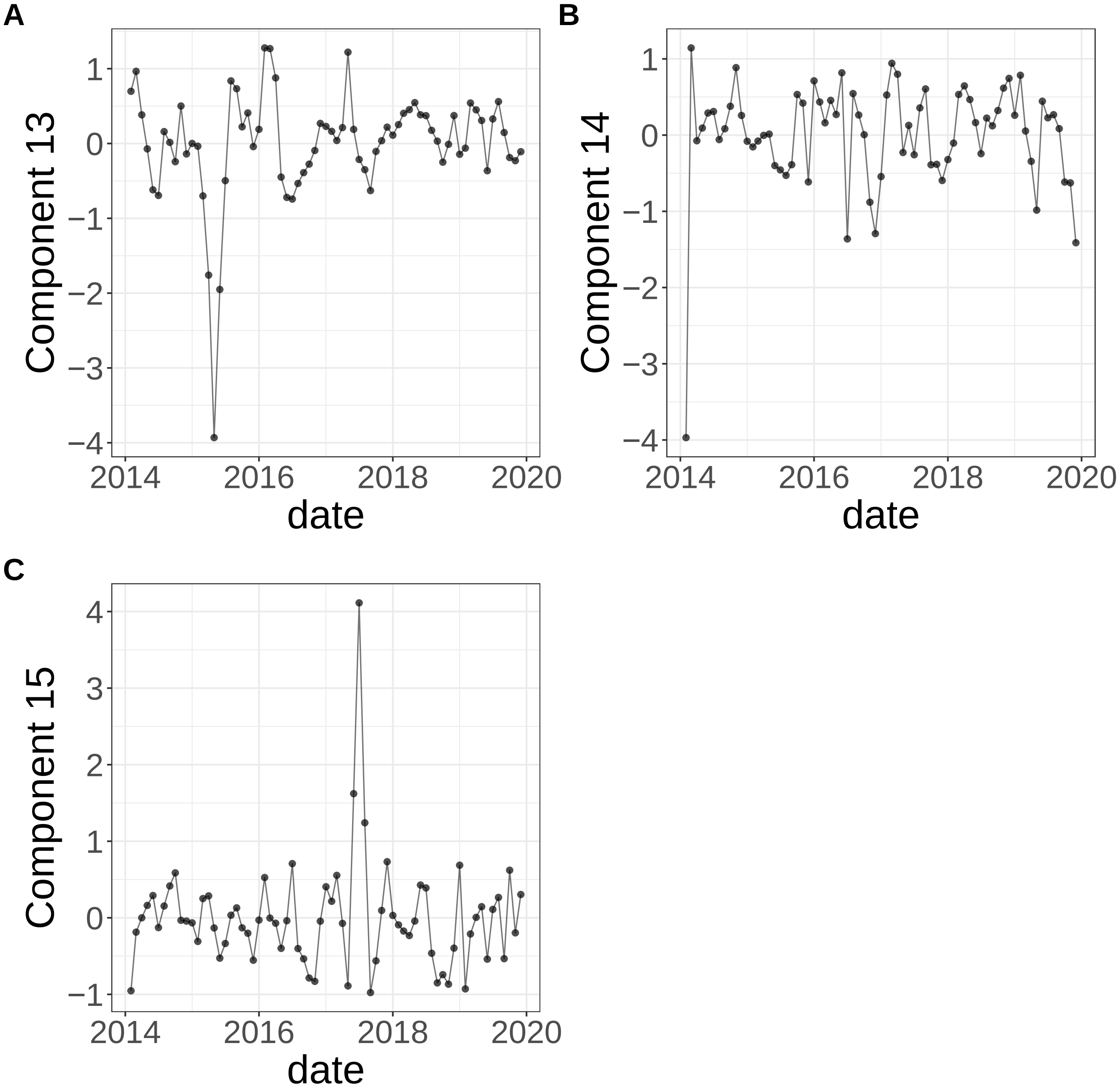


**Supplementary Figure 395.** Components for sPCA: keepX=1. (A) Component 13. (B) Component 14. (C) Component 15.

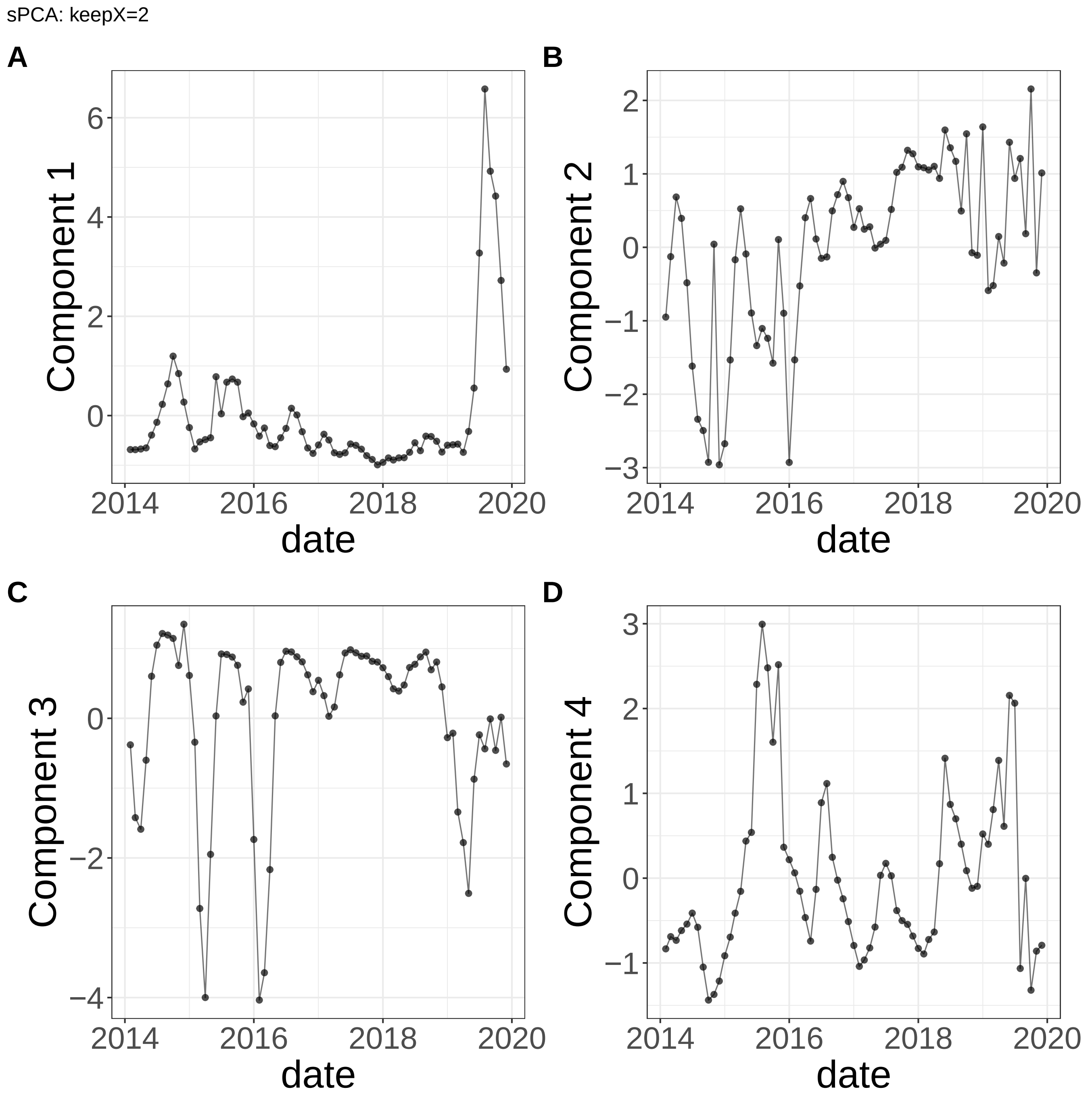


**Supplementary Figure 396.** Components for sPCA: keepX=2. (A) Component 1. (B) Component 2. (C) Component 3. (D) Component 4.

sPCA: keepX=2

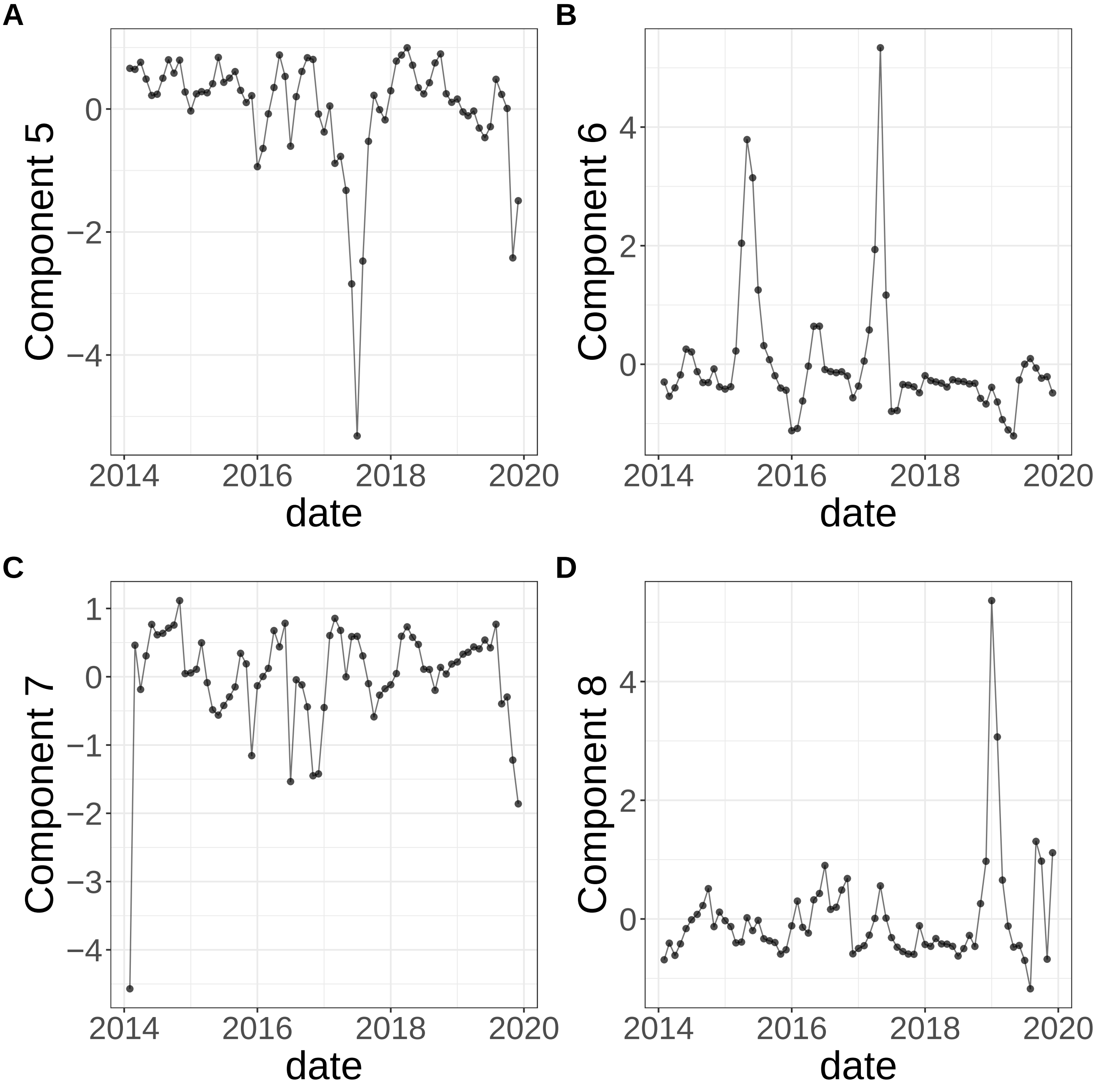


**Supplementary Figure 397.** Components for sPCA: keepX=2. (A) Component 5. (B) Component 6. (C) Component 7. (D) Component 8.

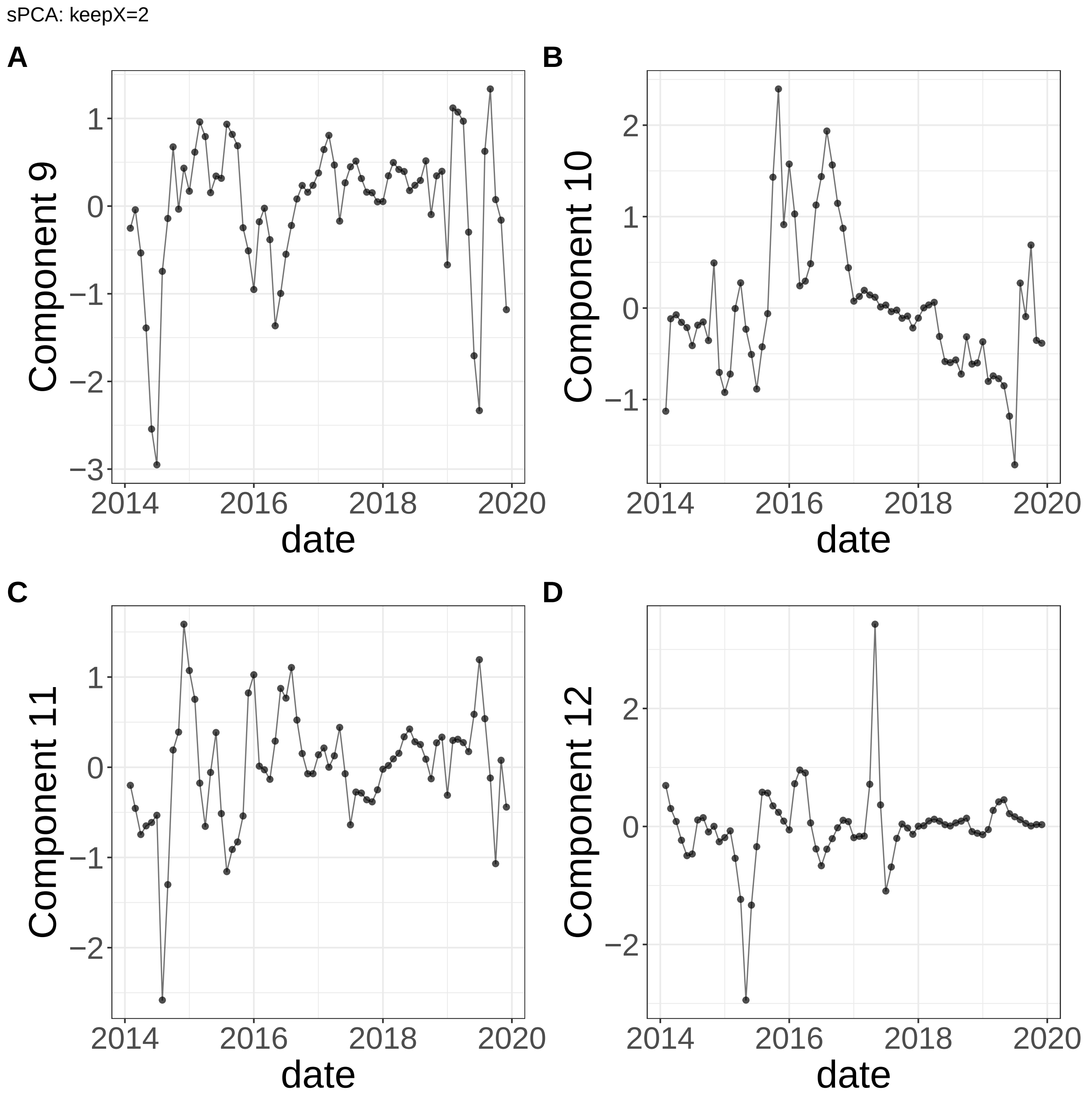


**Supplementary Figure 398.** Components for sPCA: keepX=2. (A) Component 9. (B) Component 10. (C) Component 11. (D) Component 12.

sPCA: keepX=2

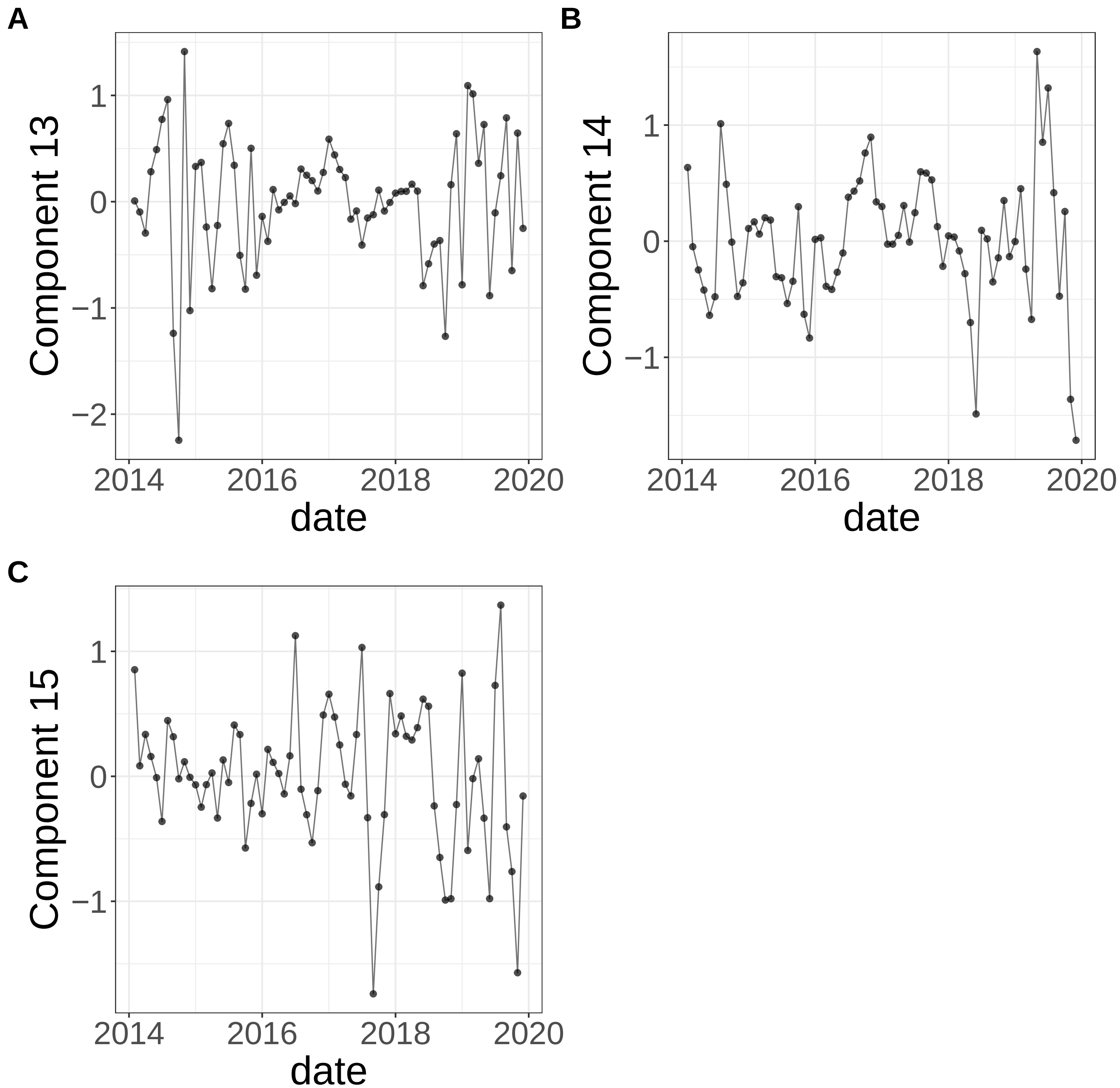


**Supplementary Figure 399.** Components for sPCA: keepX=2. (A) Component 13. (B) Component 14. (C) Component 15.

sPCA: keepX=6

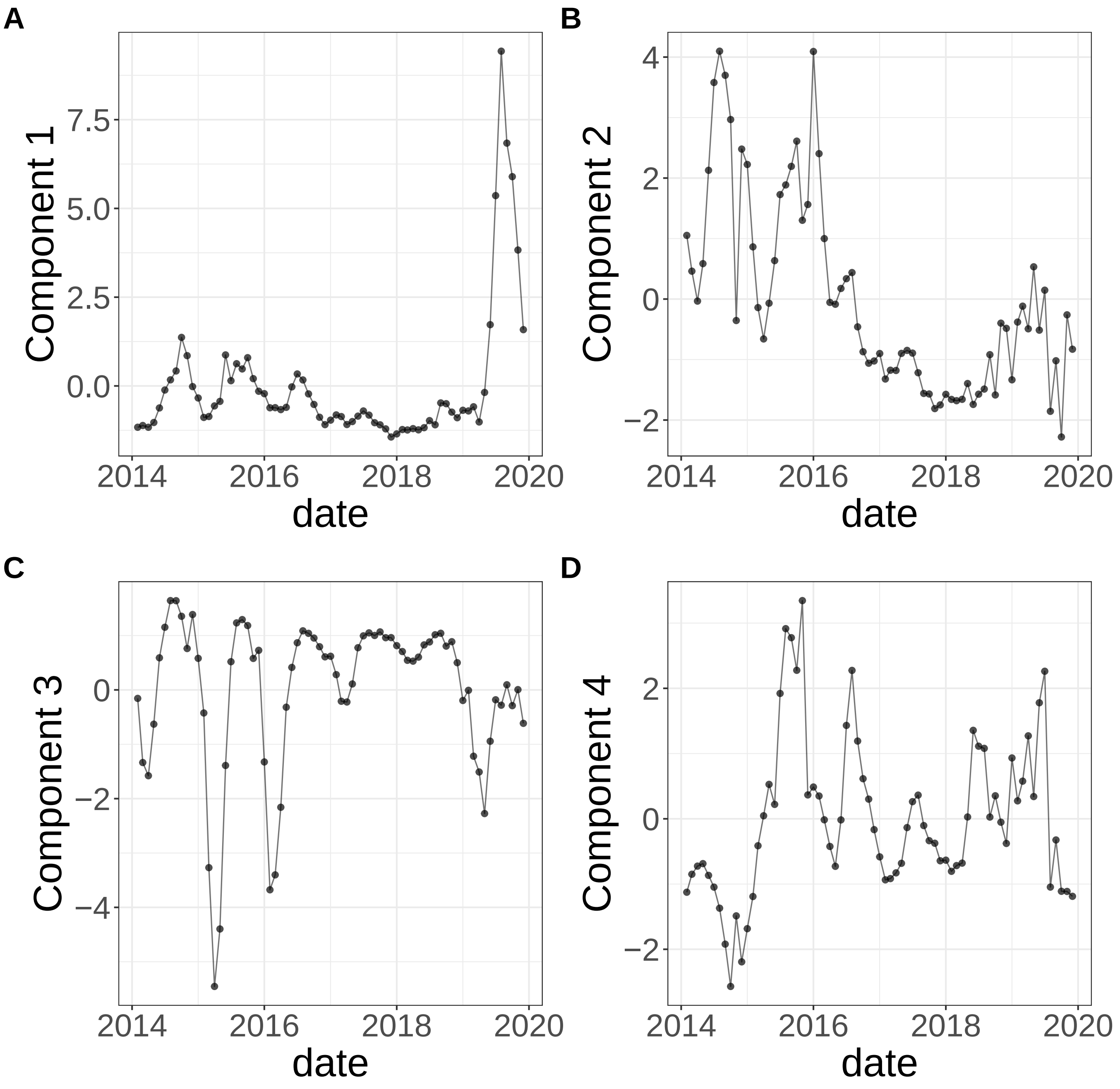


**Supplementary Figure 400.** Components for sPCA: keepX=6. (A) Component 1. (B) Component 2. (C) Component 3. (D) Component 4.

sPCA: keepX=6

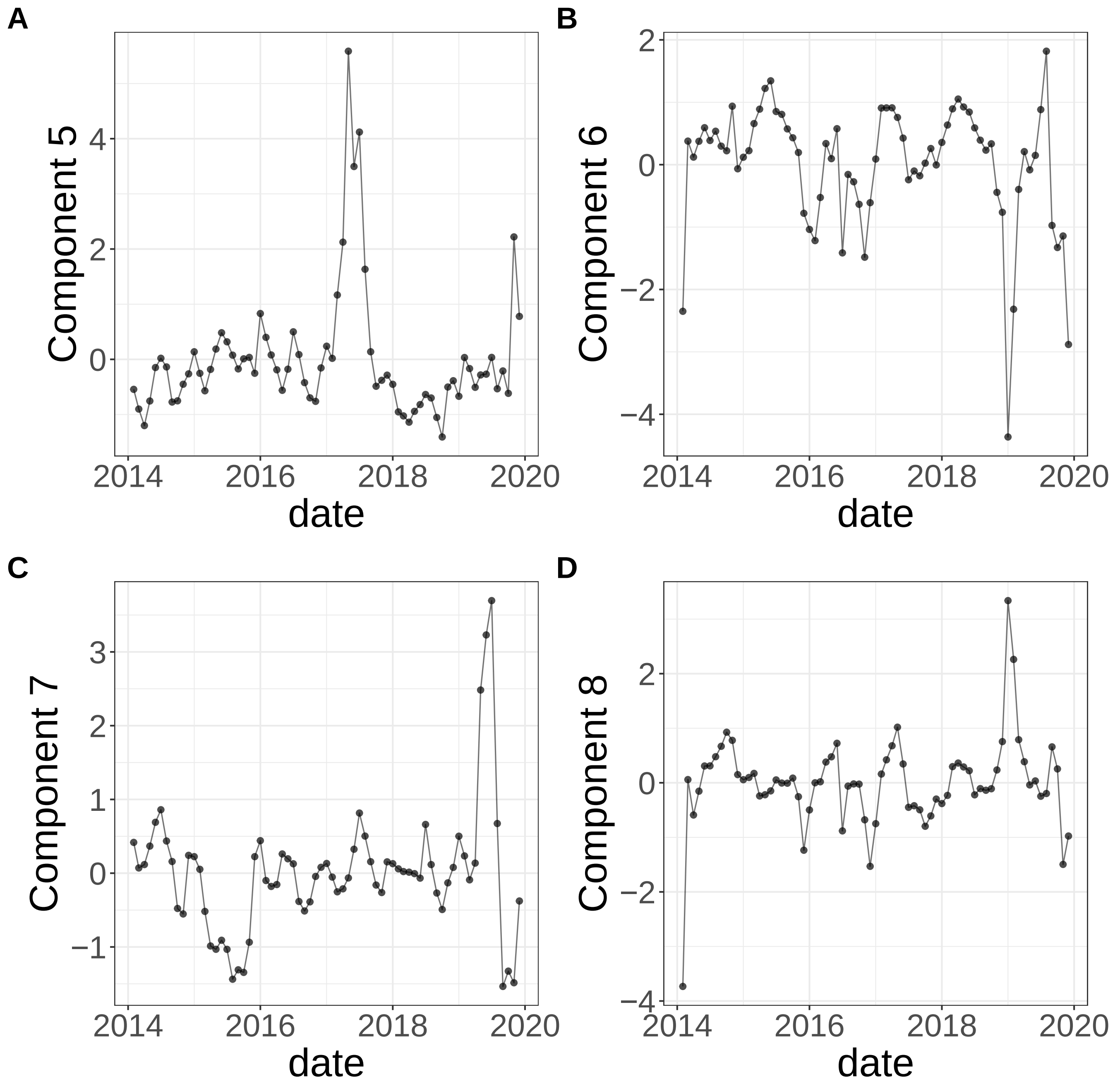


**Supplementary Figure 401.** Components for sPCA: keepX=6. (A) Component 5. (B) Component 6. (C) Component 7. (D) Component 8.

sPCA: keepX=6

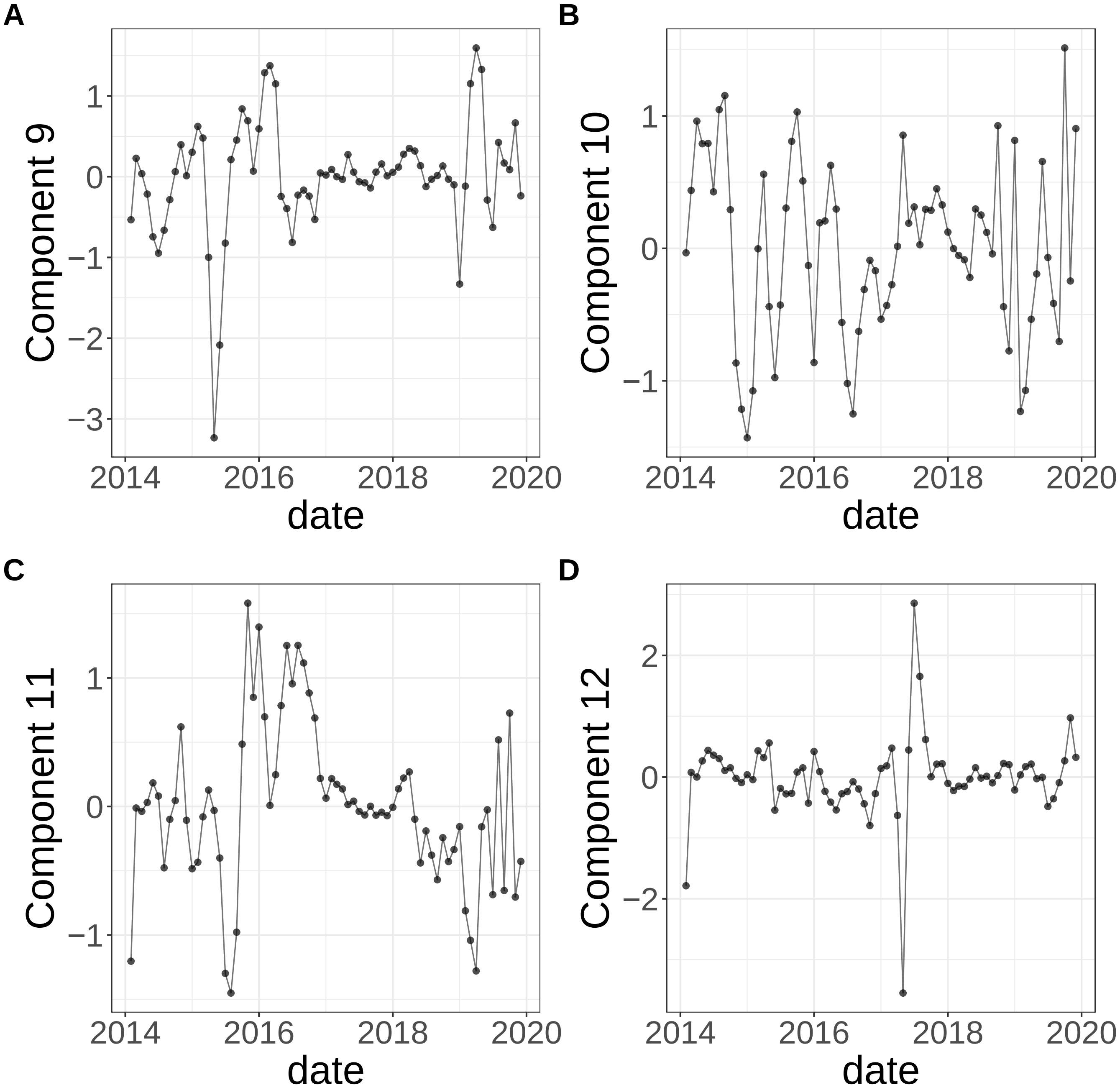


**Supplementary Figure 402.** Components for sPCA: keepX=6. (A) Component 9. (B) Component 10. (C) Component 11. (D) Component 12.

sPCA: keepX=6

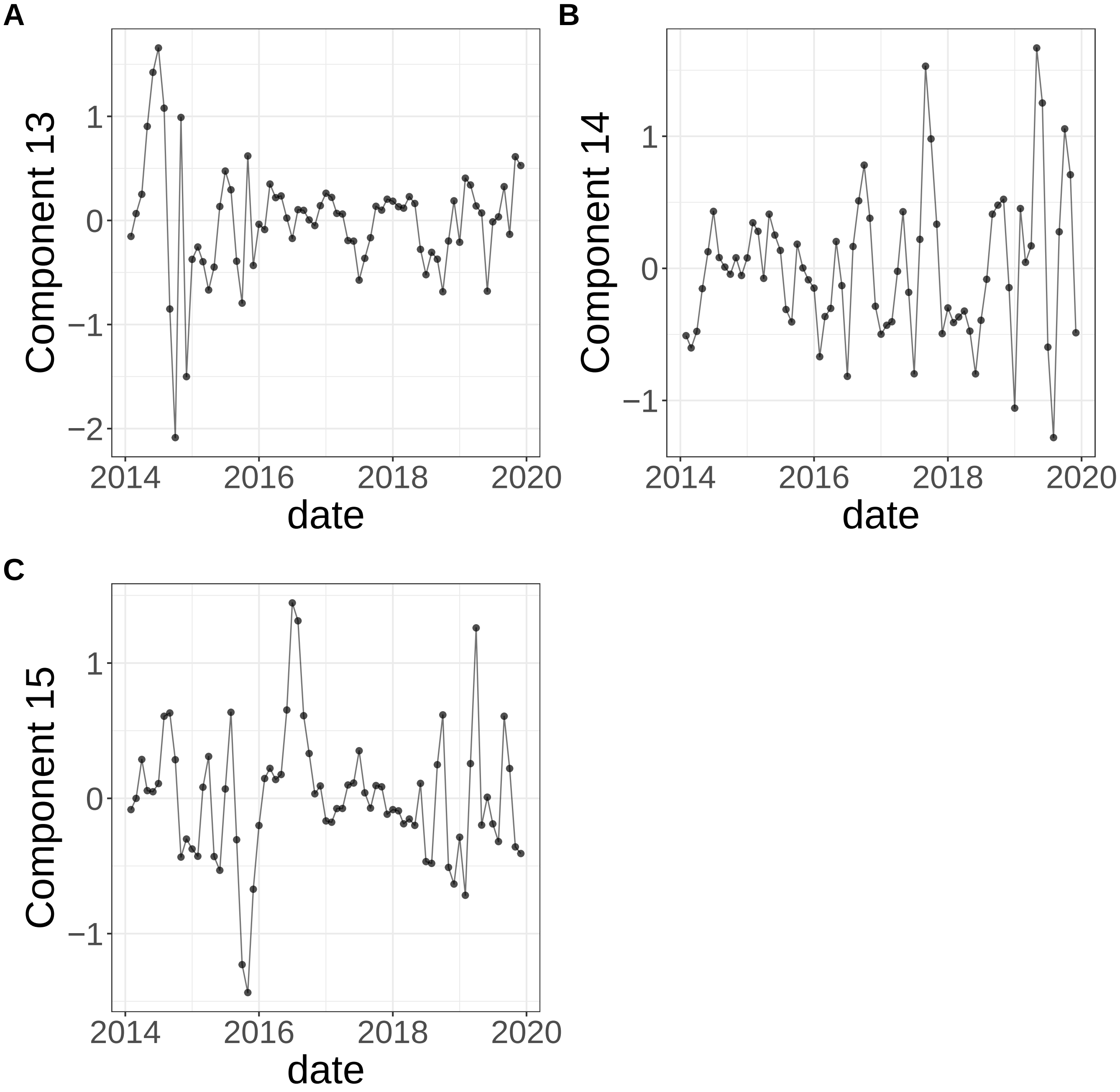


**Supplementary Figure 403.** Components for sPCA: keepX=6. (A) Component 13. (B) Component 14. (C) Component 15.

sPCA: keepX=12

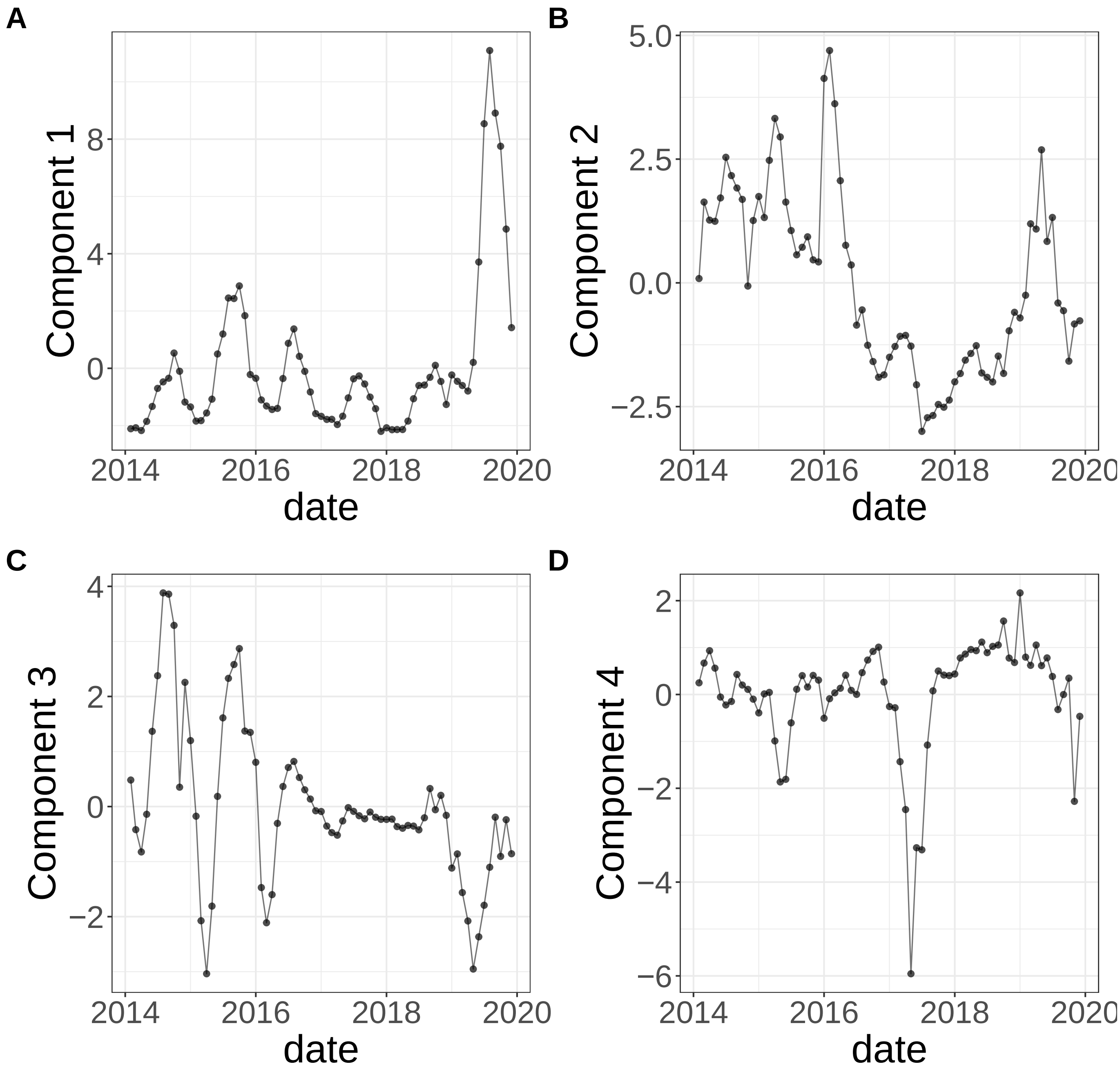


**Supplementary Figure 404.** Components for sPCA: keepX=12. (A) Component 1. (B) Component 2. (C) Component 3. (D) Component 4.

sPCA: keepX=12

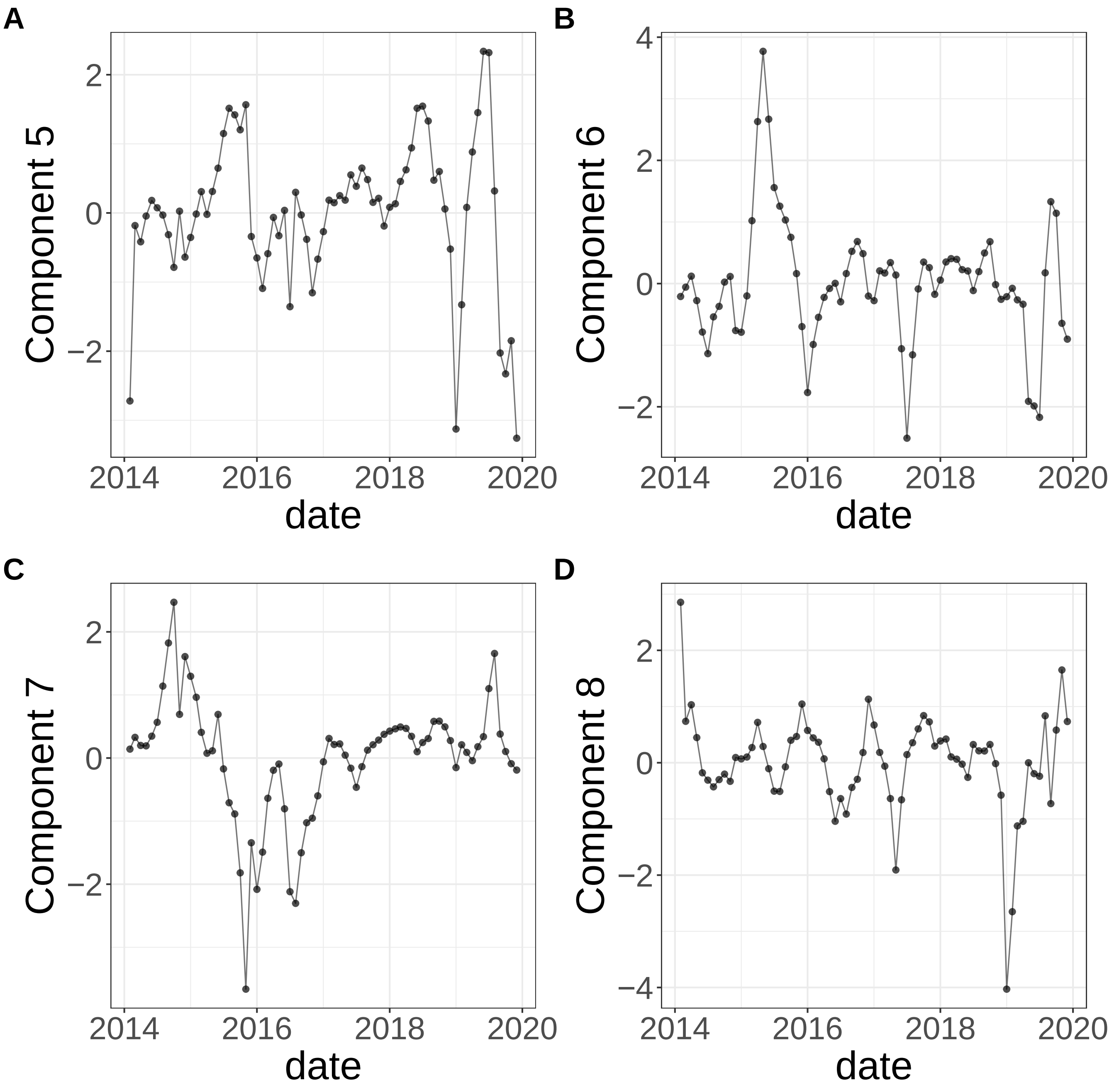


**Supplementary Figure 405.** Components for sPCA: keepX=12. (A) Component 5. (B) Component 6. (C) Component 7. (D) Component 8.

sPCA: keepX=12

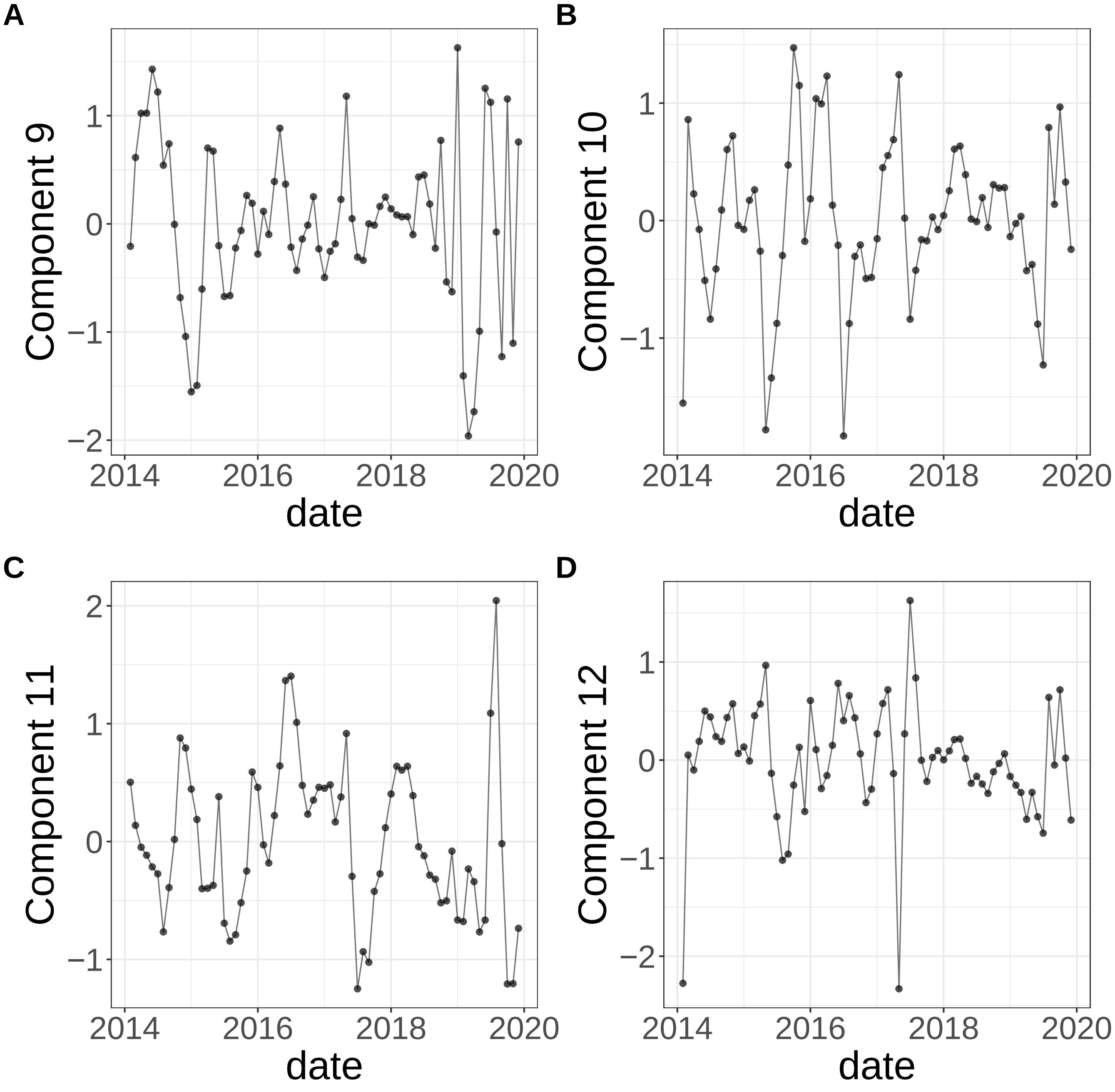


**Supplementary Figure 406.** Components for sPCA: keepX=12. (A) Component 9. (B) Component 10. (C) Component 11. (D) Component 12.

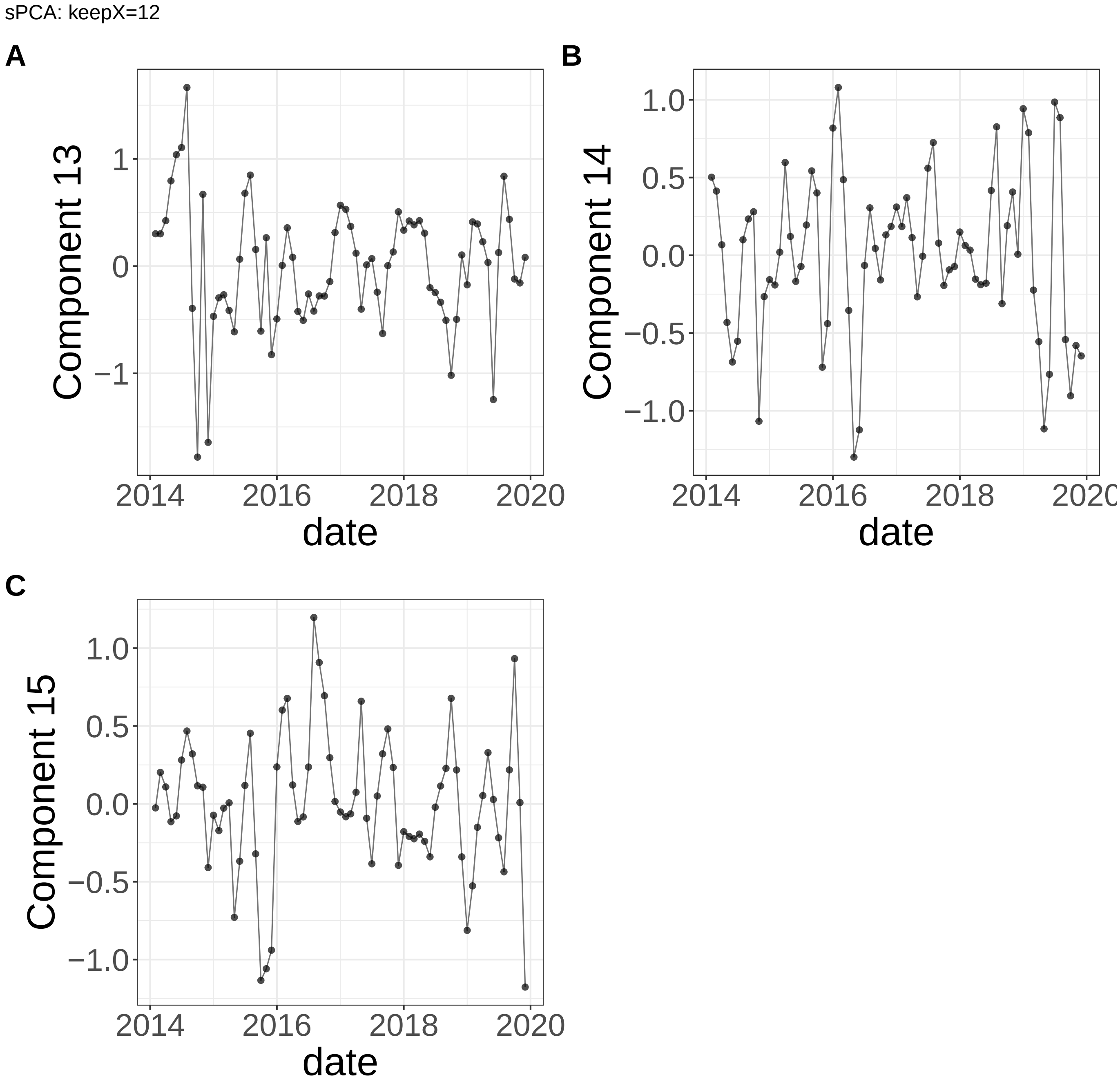


**Supplementary Figure 407.** Components for sPCA: keepX=12. (A) Component 13. (B) Component 14. (C) Component 15.

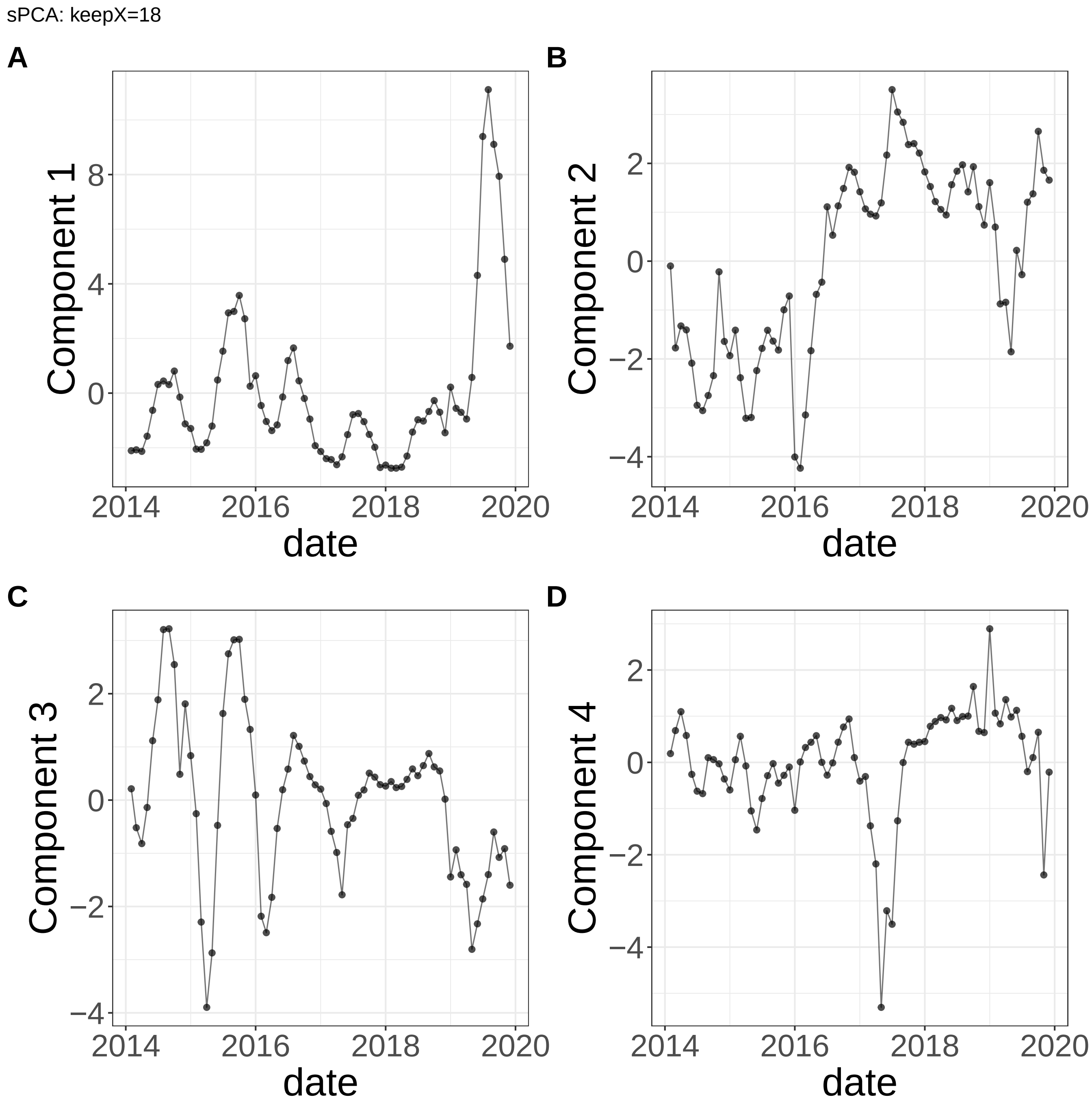


**Supplementary Figure 408.** Components for sPCA: keepX=18. (A) Component 1. (B) Component 2. (C) Component 3. (D) Component 4.

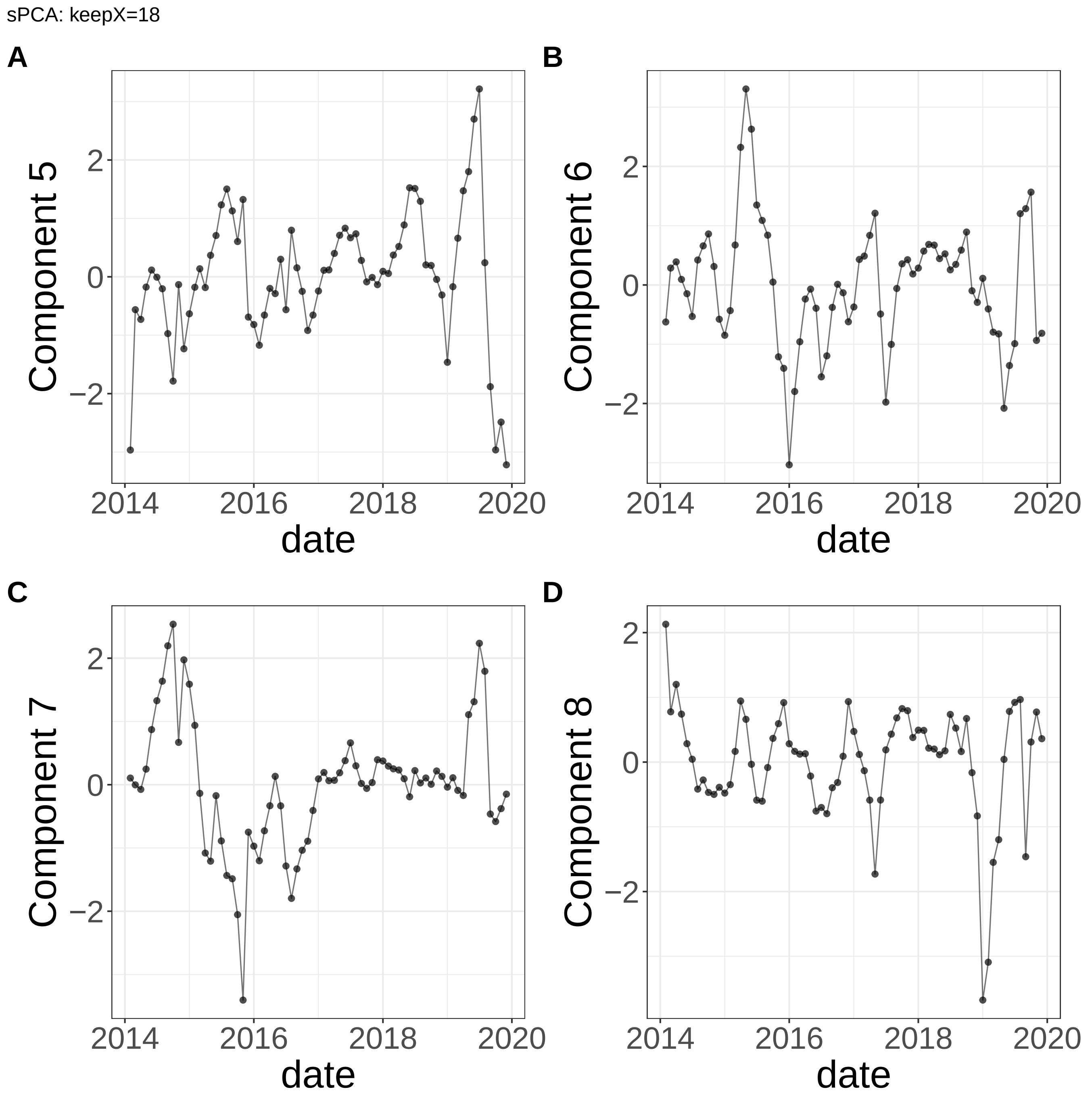


**Supplementary Figure 409.** Components for sPCA: keepX=18. (A) Component 5. (B) Component 6. (C) Component 7. (D) Component 8.

sPCA: keepX=18

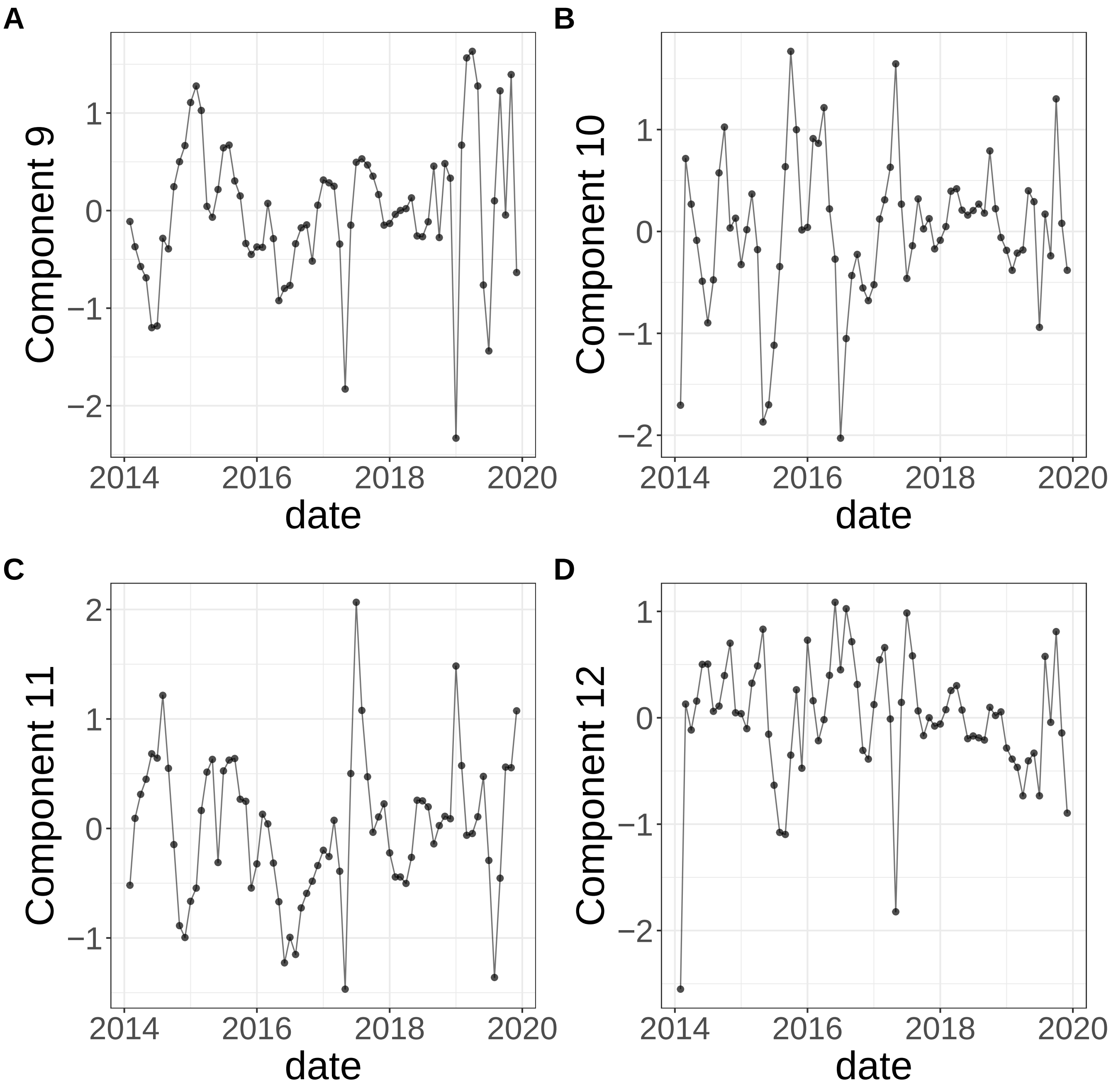


**Supplementary Figure 410.** Components for sPCA: keepX=18. (A) Component 9. (B) Component 10. (C) Component 11. (D) Component 12.

sPCA: keepX=18

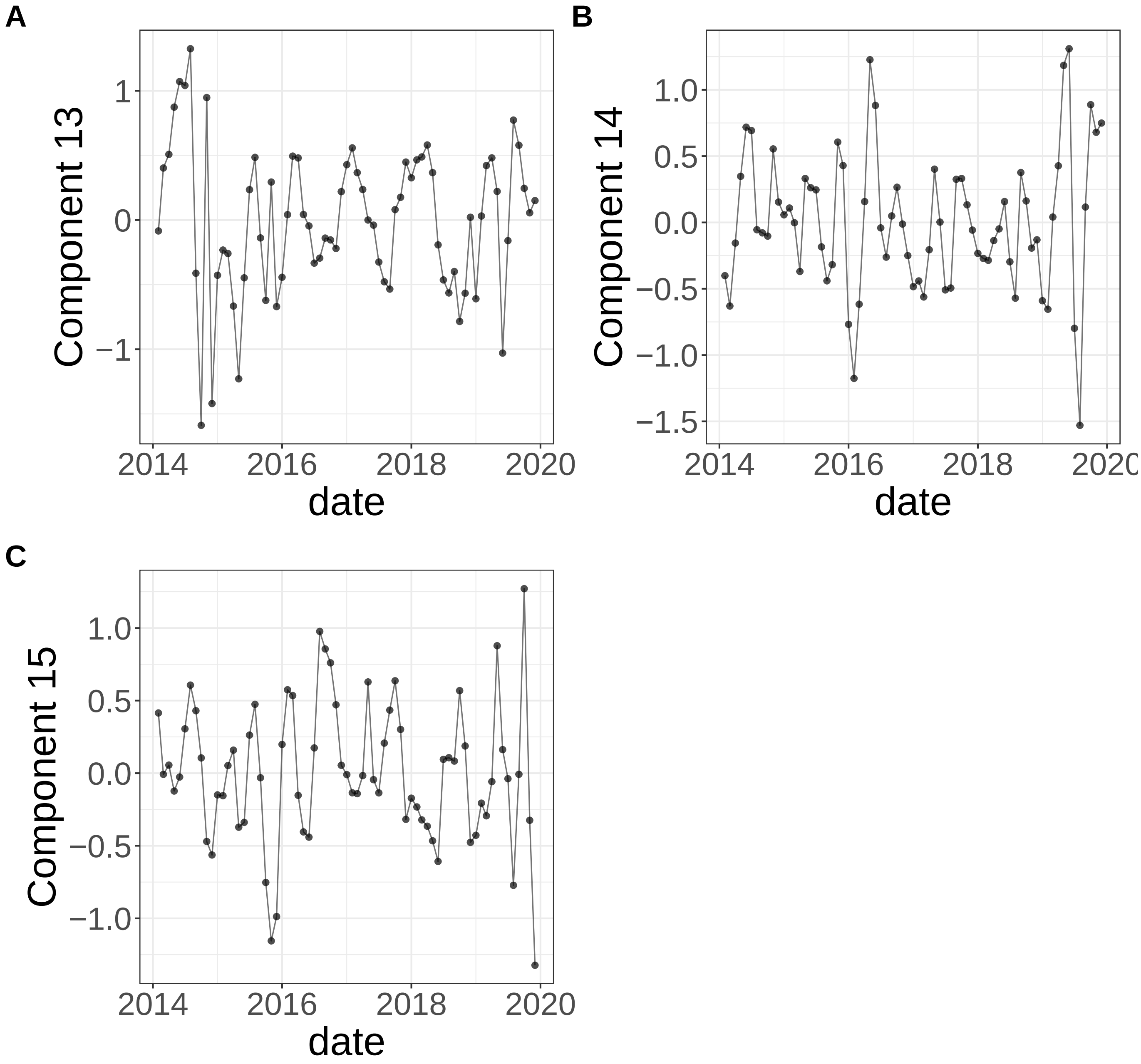


**Supplementary Figure 411.** Components for sPCA: keepX=18. (A) Component 13. (B) Component 14. (C) Component 15.

sNMF: lambda=0.001

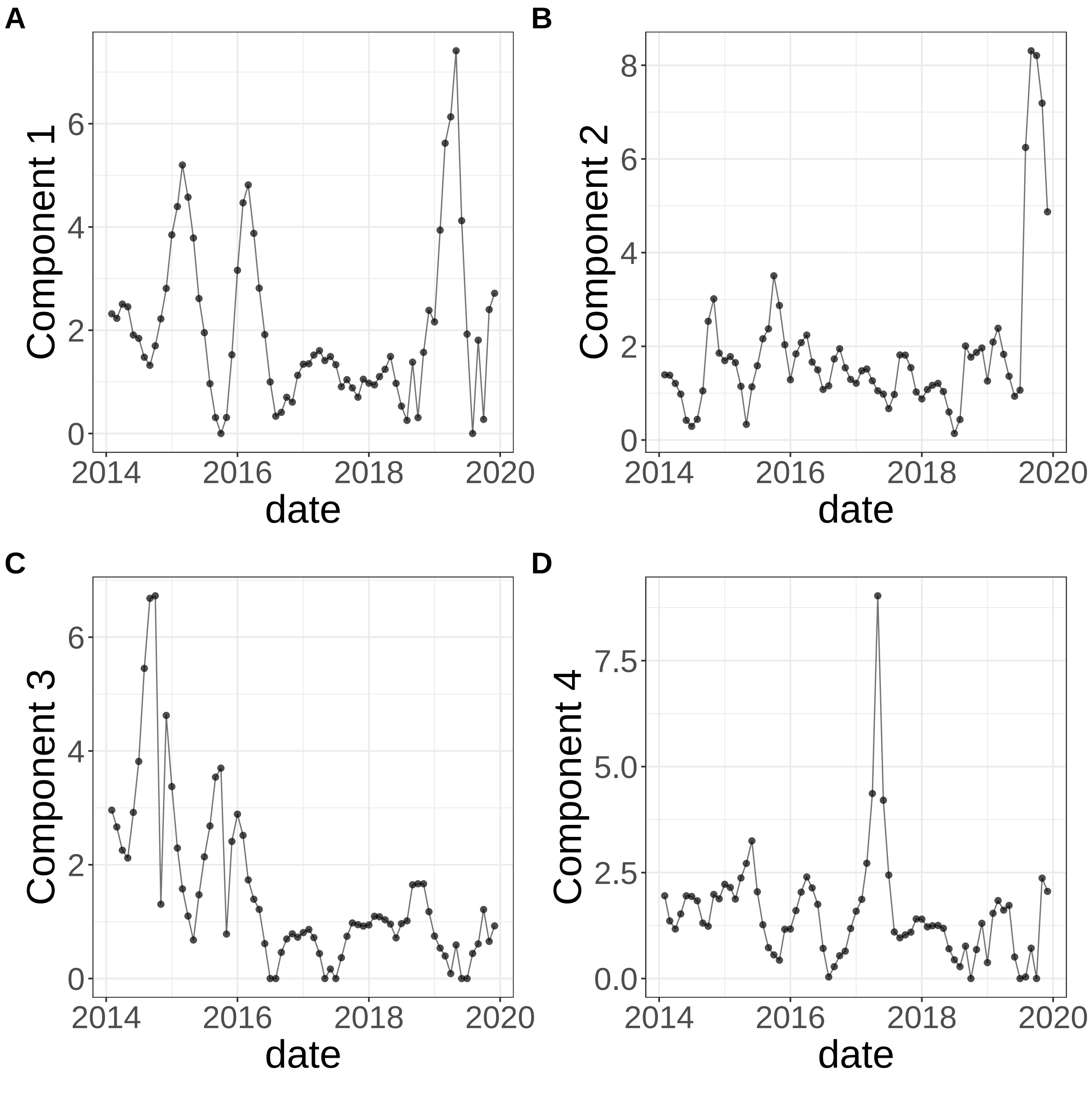


**Supplementary Figure 412.** Components for sNMF: lambda=0.001. (A) Component 1. (B) Component 2. (C) Component 3. (D) Component 4.

sNMF: lambda=0.001

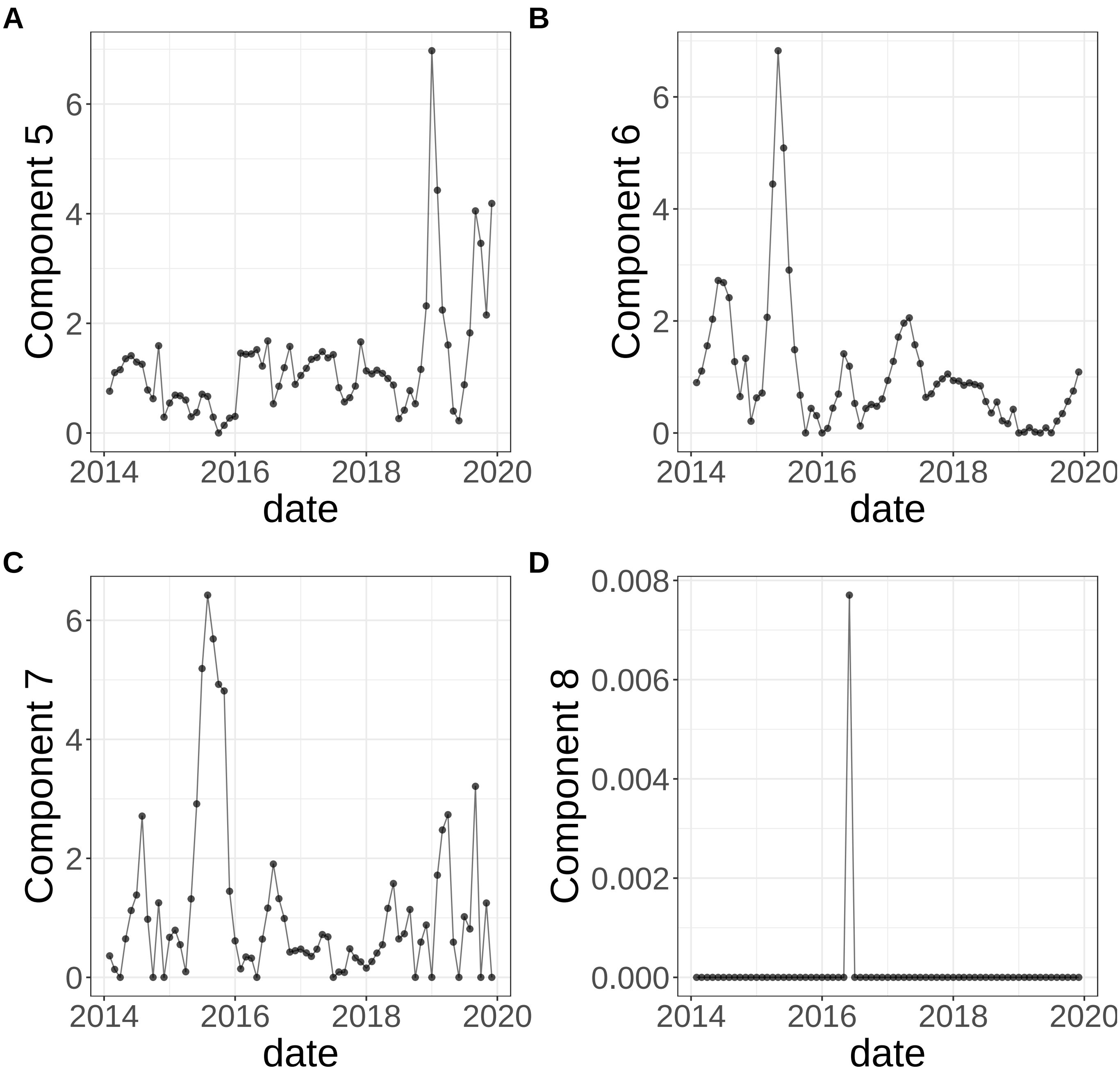


**Supplementary Figure 413.** Components for sNMF: lambda=0.001. (A) Component 5. (B) Component 6. (C) Component 7. (D) Component 8.

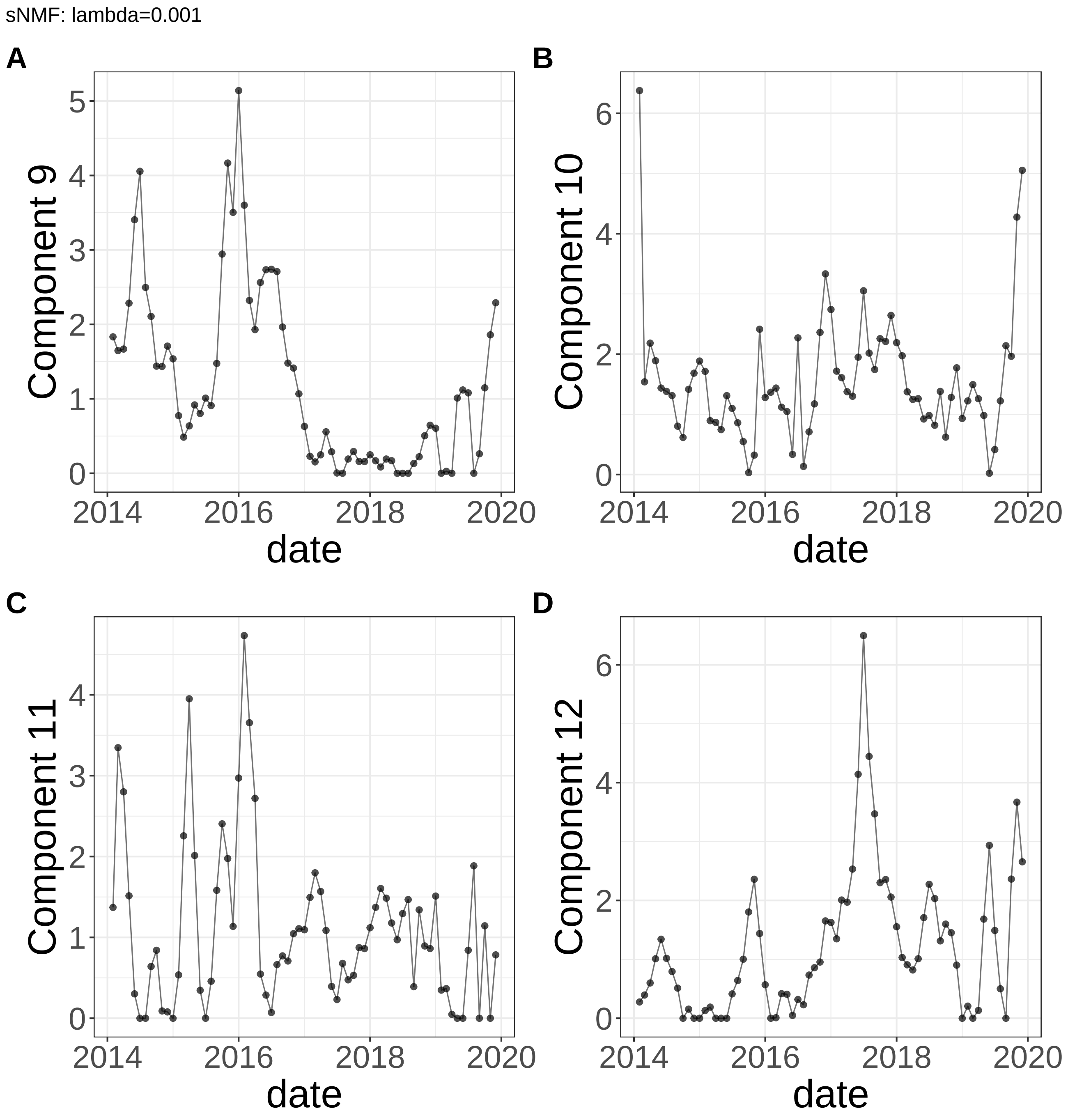


**Supplementary Figure 414.** Components for sNMF: lambda=0.001. (A) Component 9. (B) Component 10. (C) Component 11. (D) Component 12.

sNMF: lambda=0.001

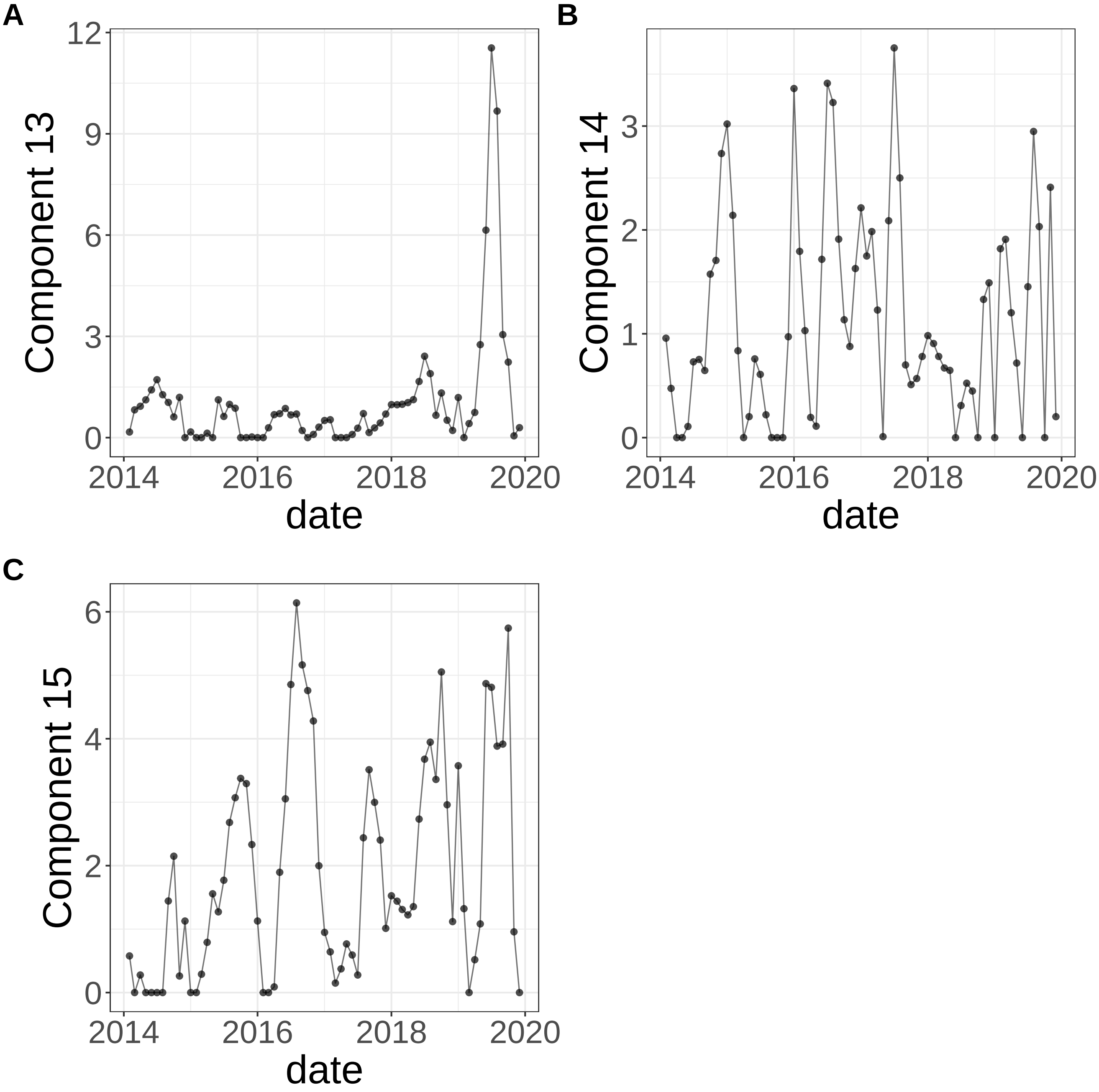


**Supplementary Figure 415.** Components for sNMF: lambda=0.001. (A) Component 13. (B) Component 14. (C) Component 15.

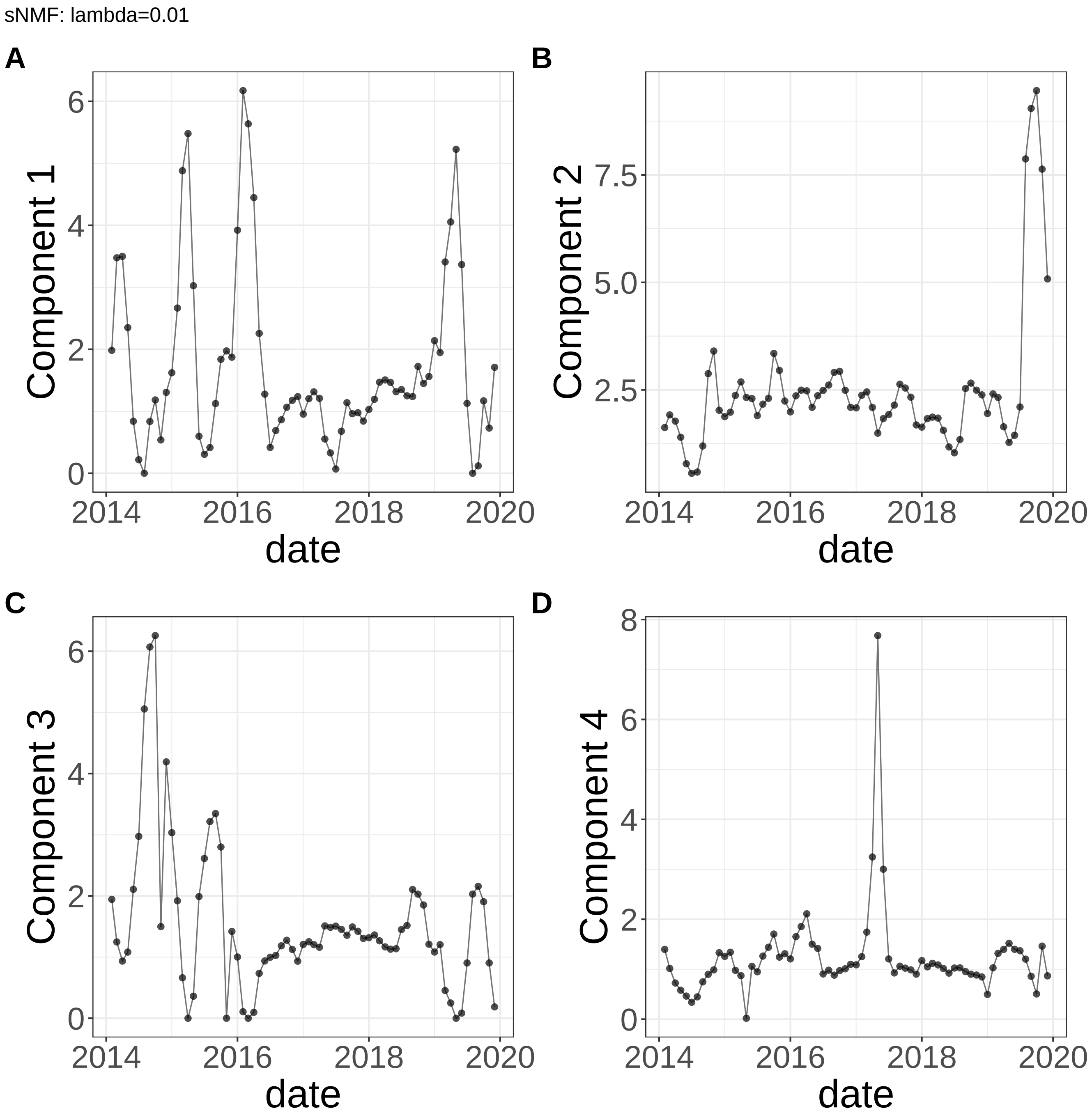


**Supplementary Figure 416.** Components for sNMF: lambda=0.01. (A) Component 1. (B) Component 2. (C) Component 3. (D) Component 4.

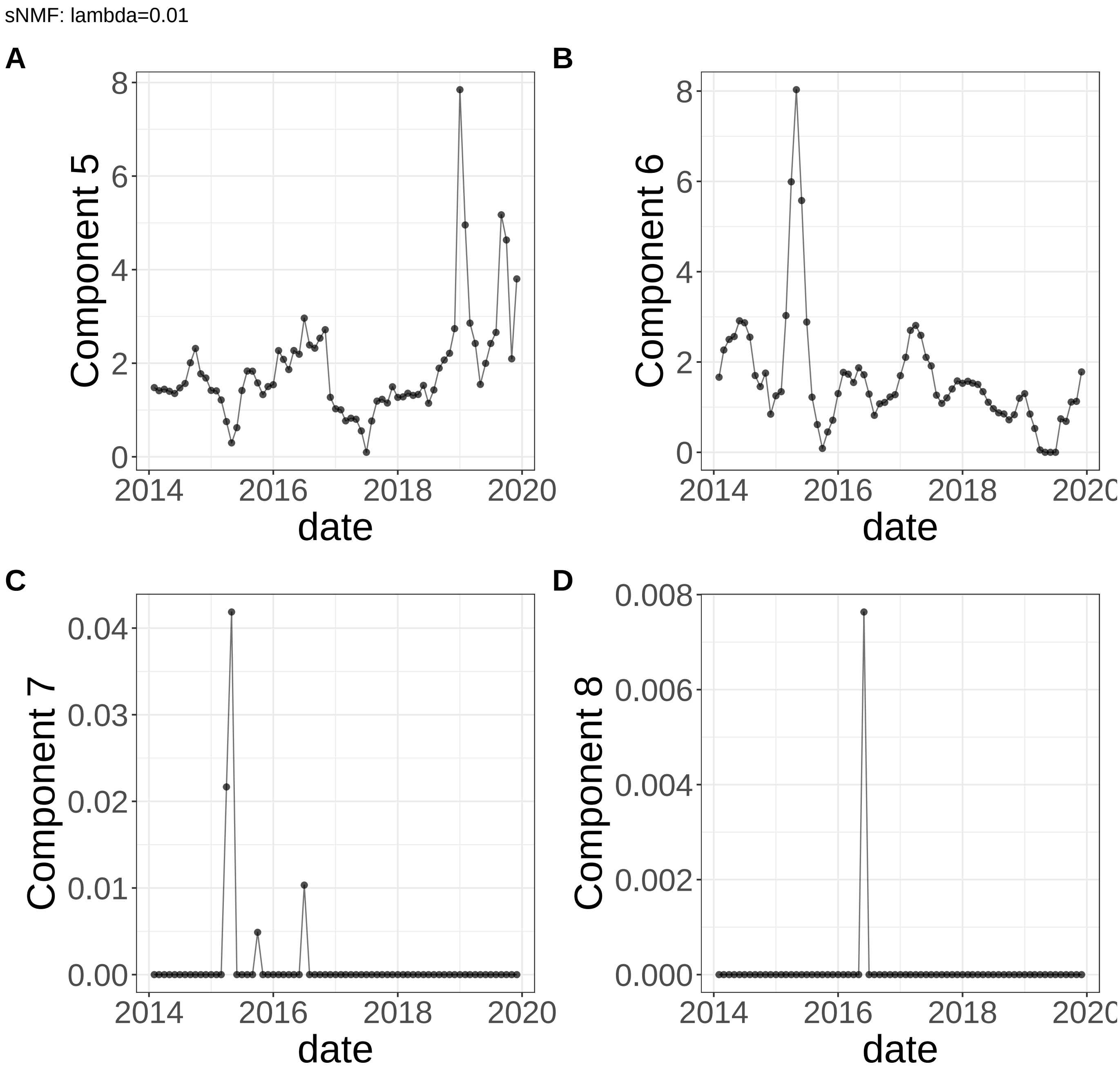


**Supplementary Figure 417.** Components for sNMF: lambda=0.01. (A) Component 5. (B) Component 6. (C) Component 7. (D) Component 8.

sNMF: lambda=0.01

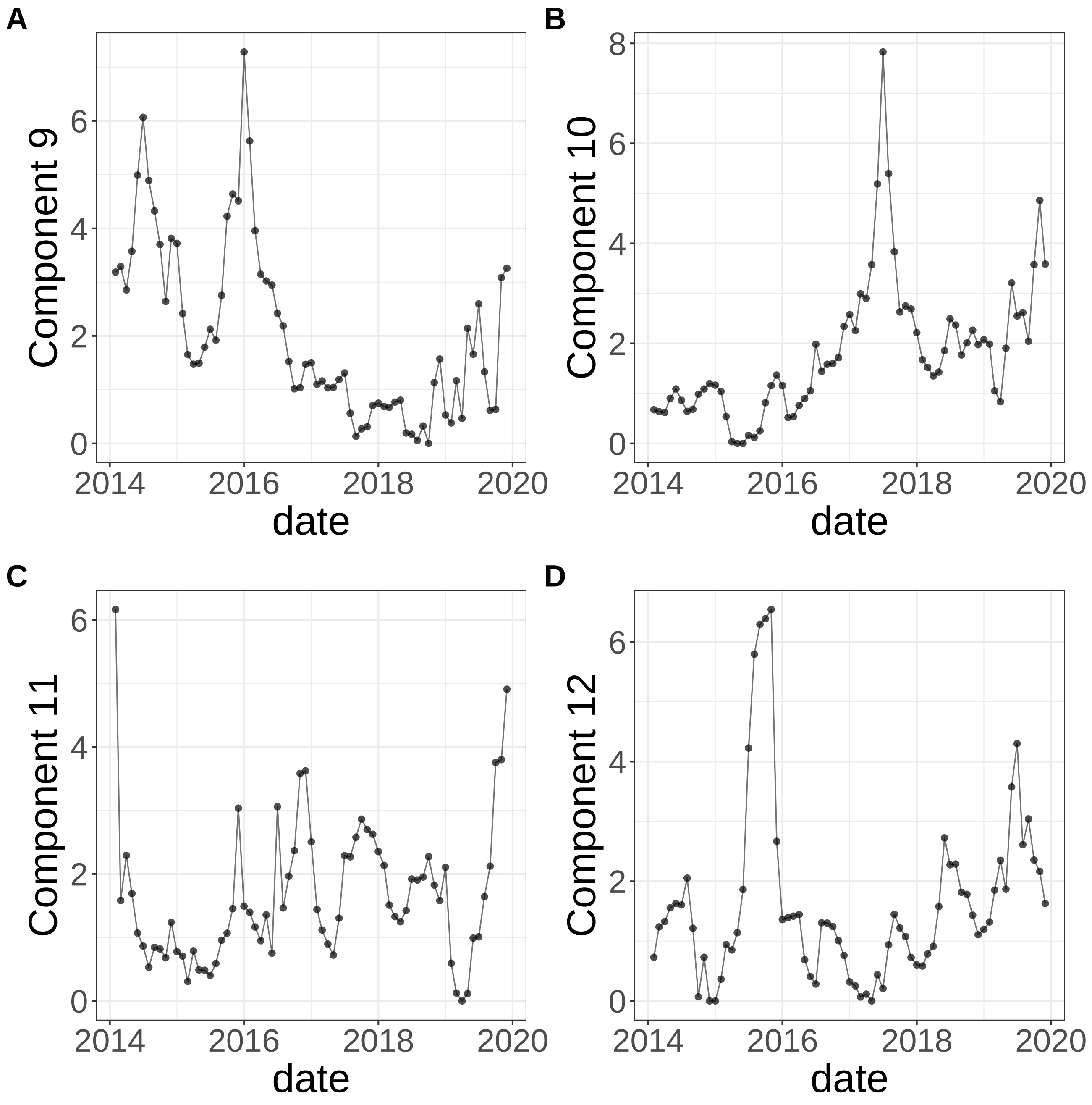


**Supplementary Figure 418.** Components for sNMF: lambda=0.01. (A) Component 9. (B) Component 10. (C) Component 11. (D) Component 12.

sNMF: lambda=0.01

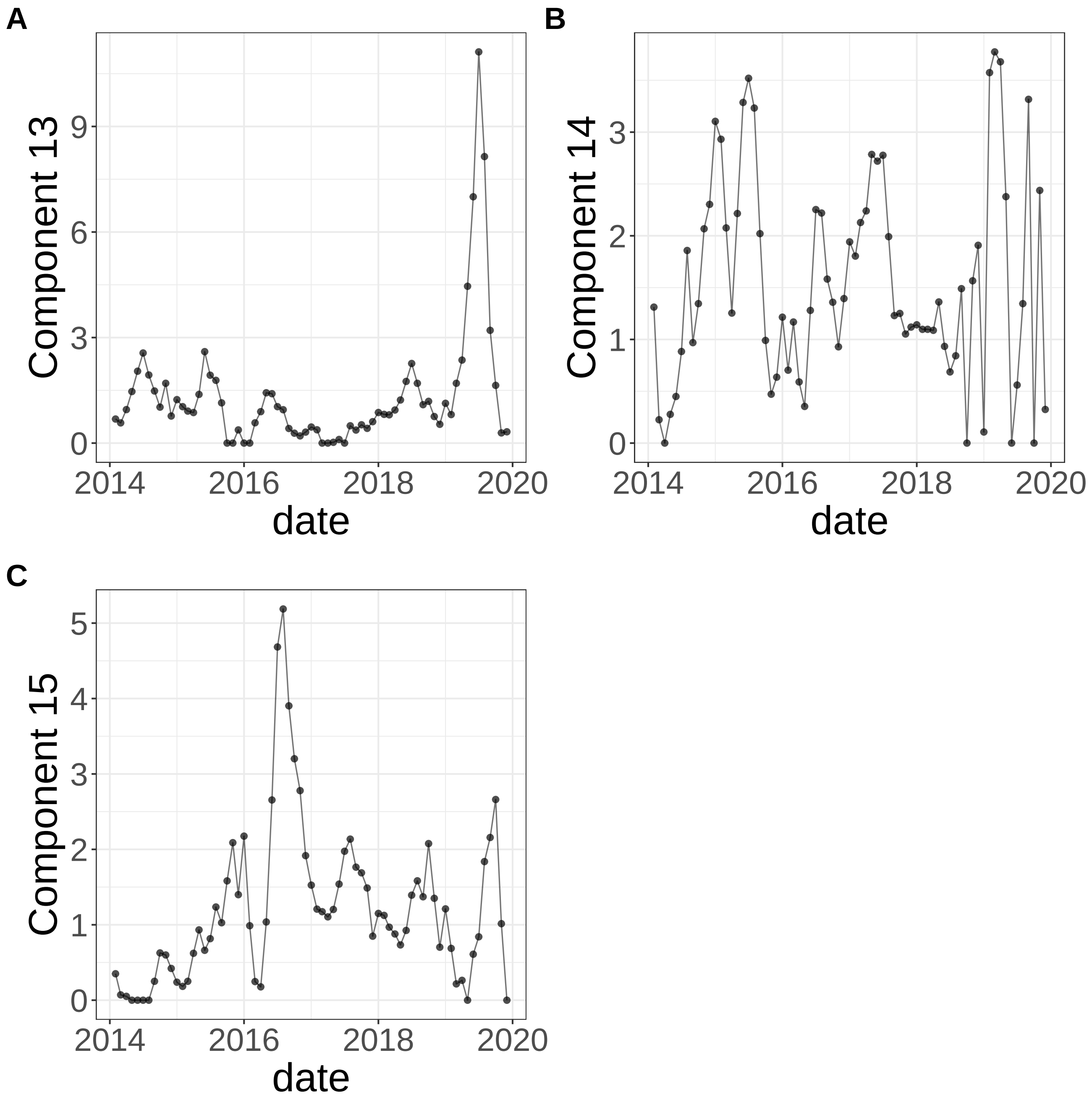


**Supplementary Figure 419.** Components for sNMF: lambda=0.01. (A) Component 13. (B) Component 14. (C) Component 15.

sNMF: lambda=0.1

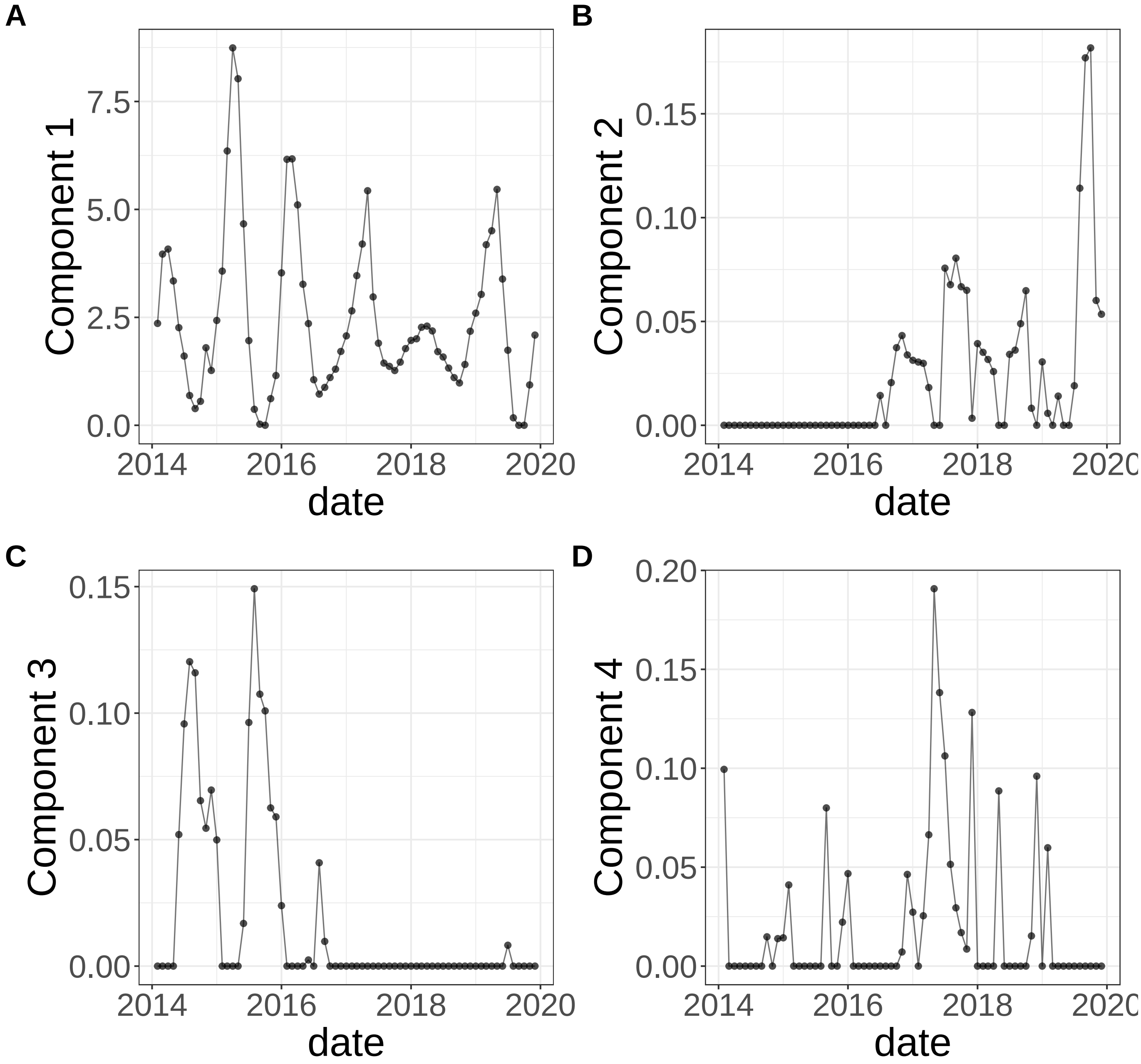


**Supplementary Figure 420.** Components for sNMF: lambda=0.1. (A) Component 1. (B) Component 2. (C) Component 3. (D) Component 4.

sNMF: lambda=0.1

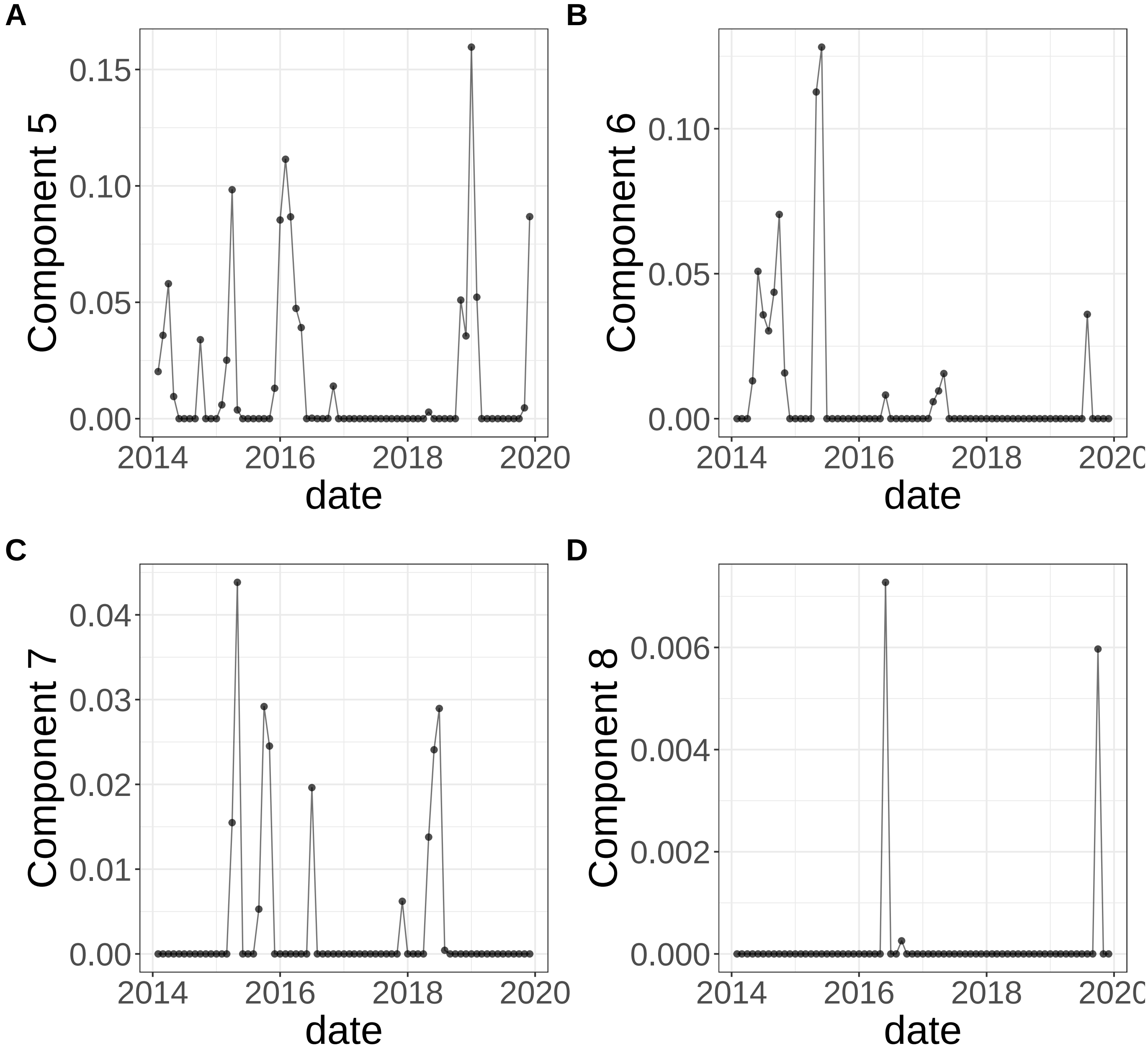


**Supplementary Figure 421.** Components for sNMF: lambda=0.1. (A) Component 5. (B) Component 6. (C) Component 7. (D) Component 8.

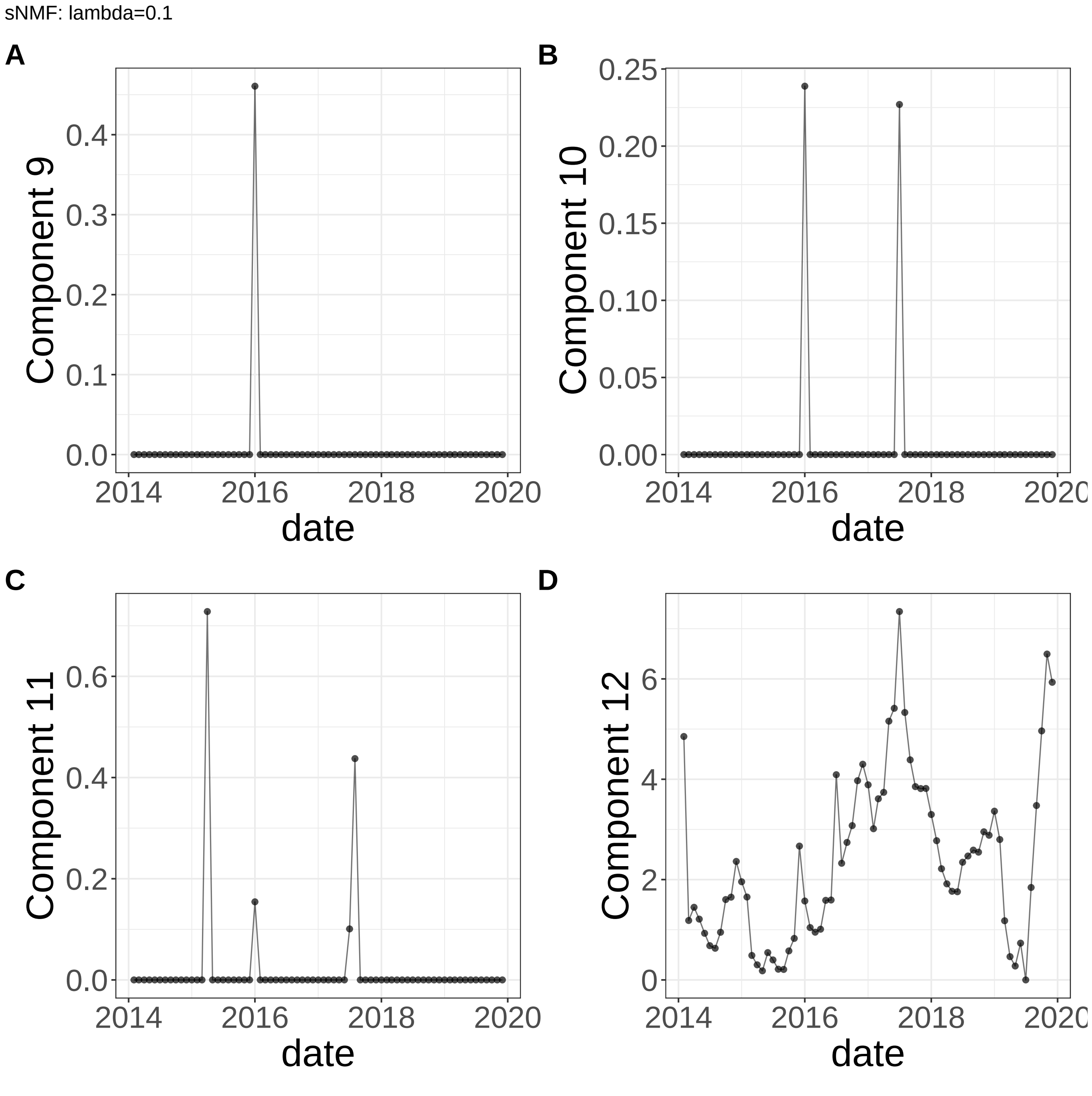


**Supplementary Figure 422.** Components for sNMF: lambda=0.1. (A) Component 9. (B) Component 10. (C) Component 11. (D) Component 12.

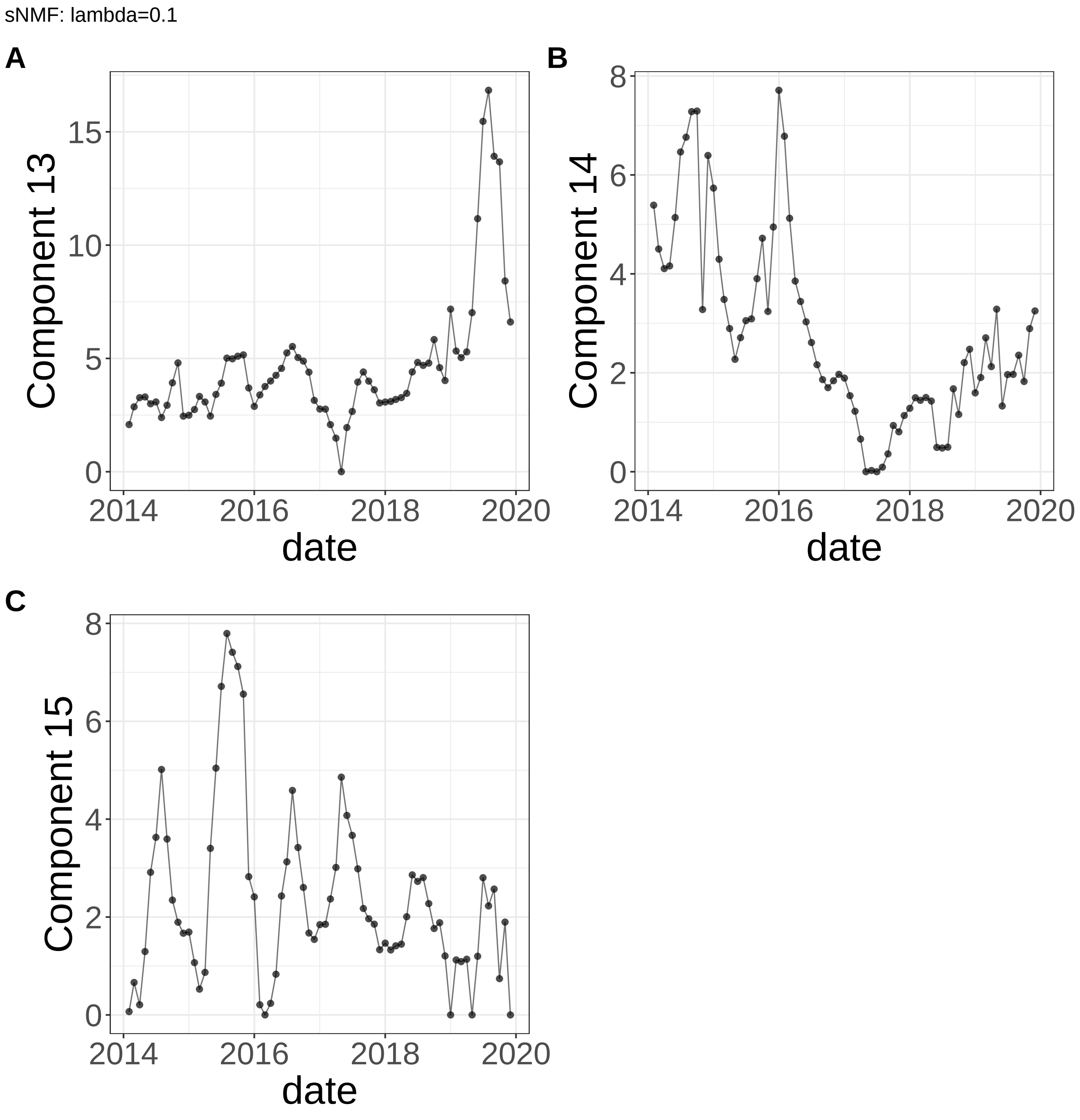


**Supplementary Figure 423.** Components for sNMF: lambda=0.1. (A) Component 13. (B) Component 14. (C) Component 15.

MEFISTO: duplicate views

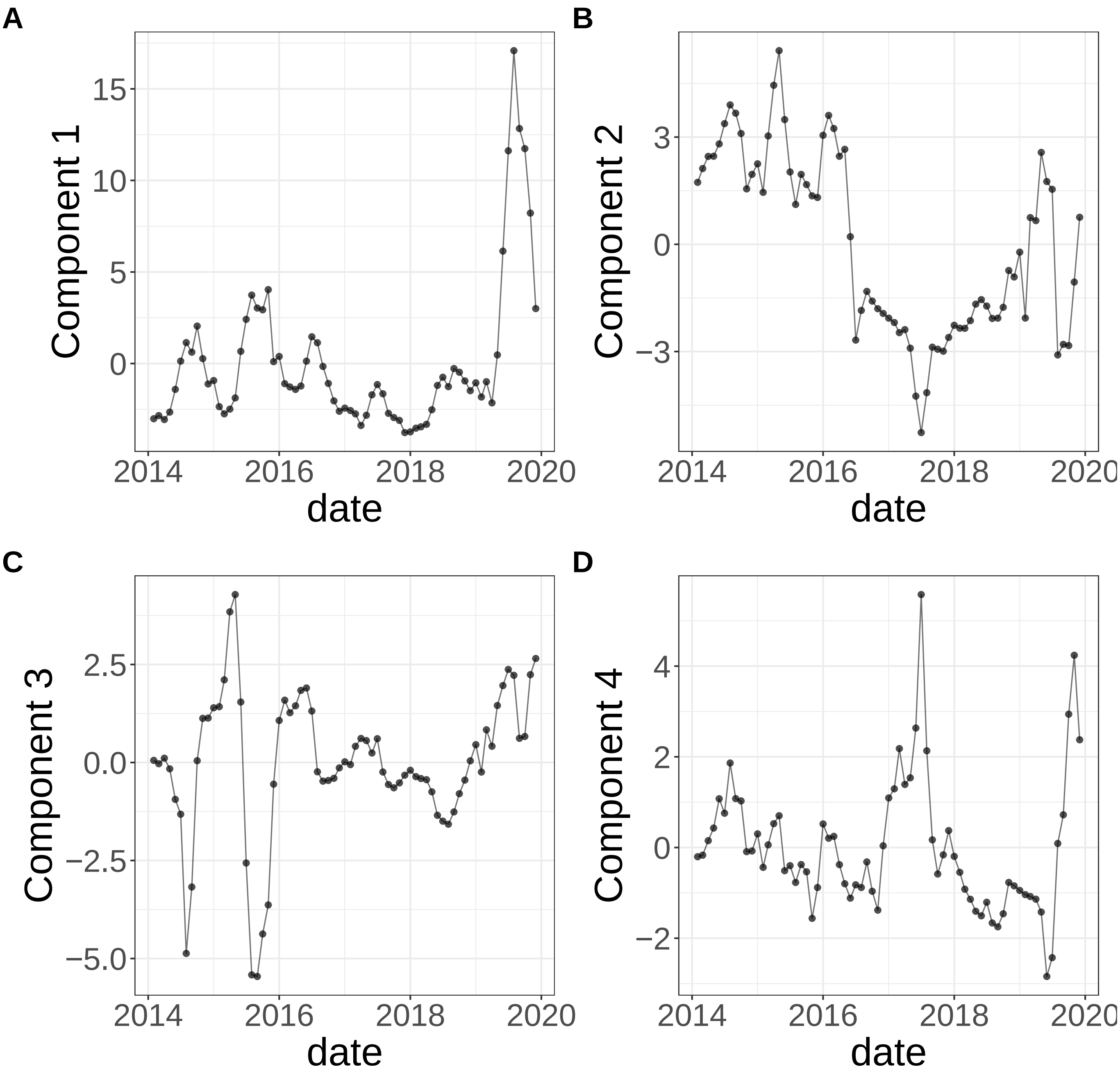


**Supplementary Figure 424.** Components for MEFISTO: duplicate views. (A) Component 1. (B) Component 2. (C) Component 3. (D) Component 4.

MEFISTO: duplicate views

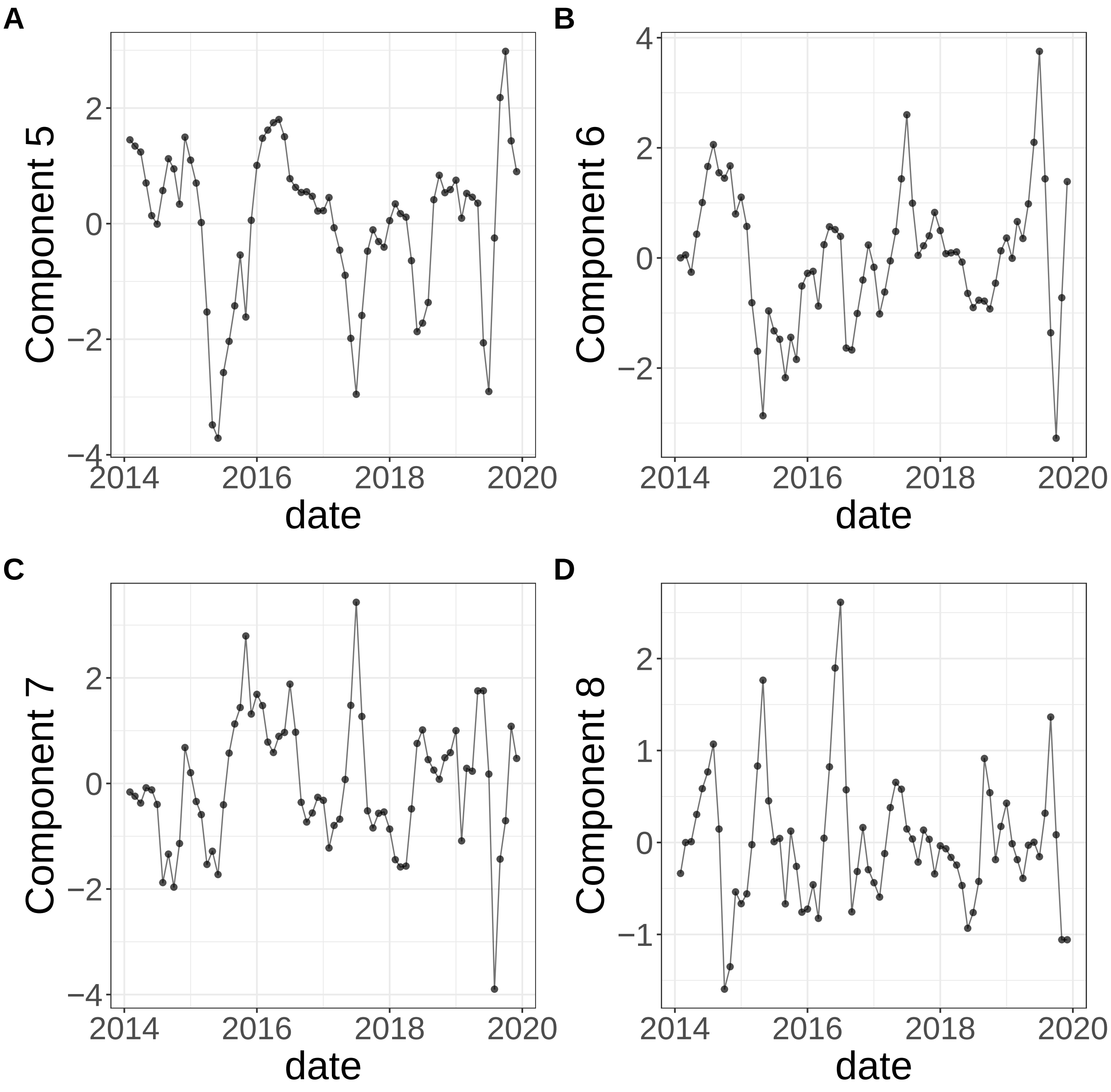


**Supplementary Figure 425.** Components for MEFISTO: duplicate views. (A) Component 5. (B) Component 6. (C) Component 7. (D) Component 8.
