## Supplementary Note 6 : Breeding Bird Survey for "A statistical framework for system-level synthesis"

This note presents the supplementary results for the analysis of Breeding Bird Survey data. The figures include:

- Processes/component functions learned by SSANOVA-SVD (**Supplementary Fig. 222**-**Supplementary Fig. 229**)
- Inference of feature participation from the application of the three-step framework: Volcano plots (**Supplementary Fig. 230**-**Supplementary Fig. 237**), heatmaps of top 50 component-associated features at FDR of 10% (**Supplementary Fig. 238**-**Supplementary Fig. 252**), enrichment for associated birds in predefined ecological groups (**Supplementary Fig. 253**-**Supplementary Fig. 264**)
- Decomposition of the processes in terms of effects due to covariates and their interactions (**Supplementary Fig. 265**-**Supplementary Fig. 294**).
- Components from a representative class of matrix factorization methods (**Supplementary Fig. 295**-**Supplementary Fig. 328**)

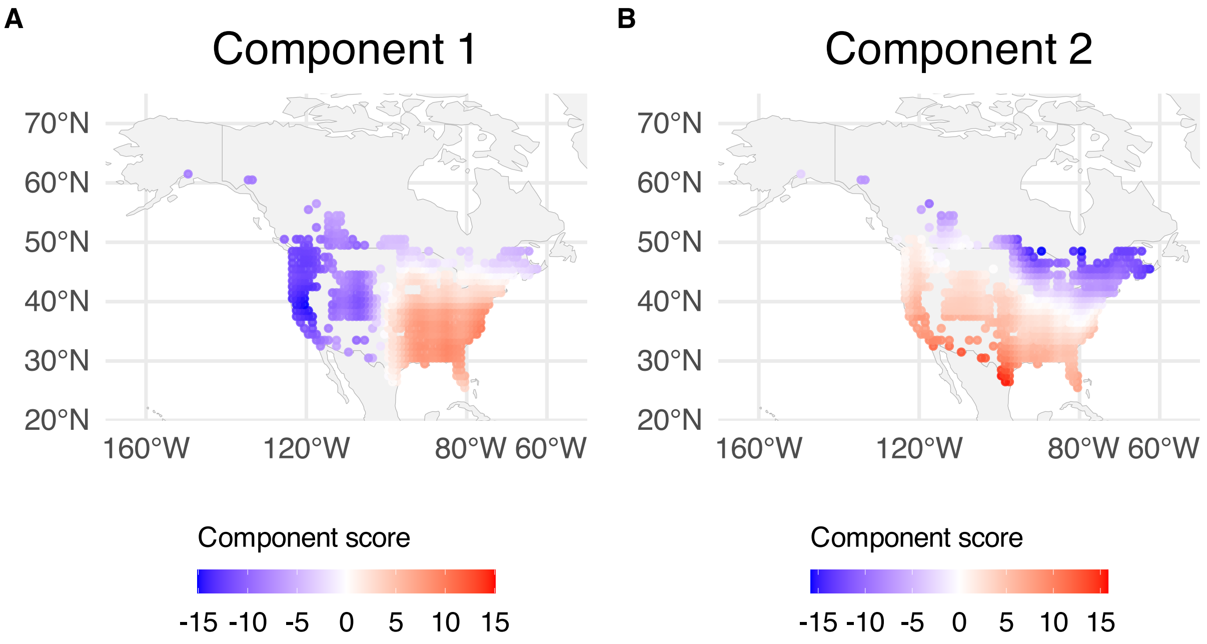


**Supplementary Figure 222.** SSANOVA-SVD reconstructed component functions mapped over BBS sampling locations; component scores are averaged over years within spatial grid. (A) Component 1. (B) Component 2.

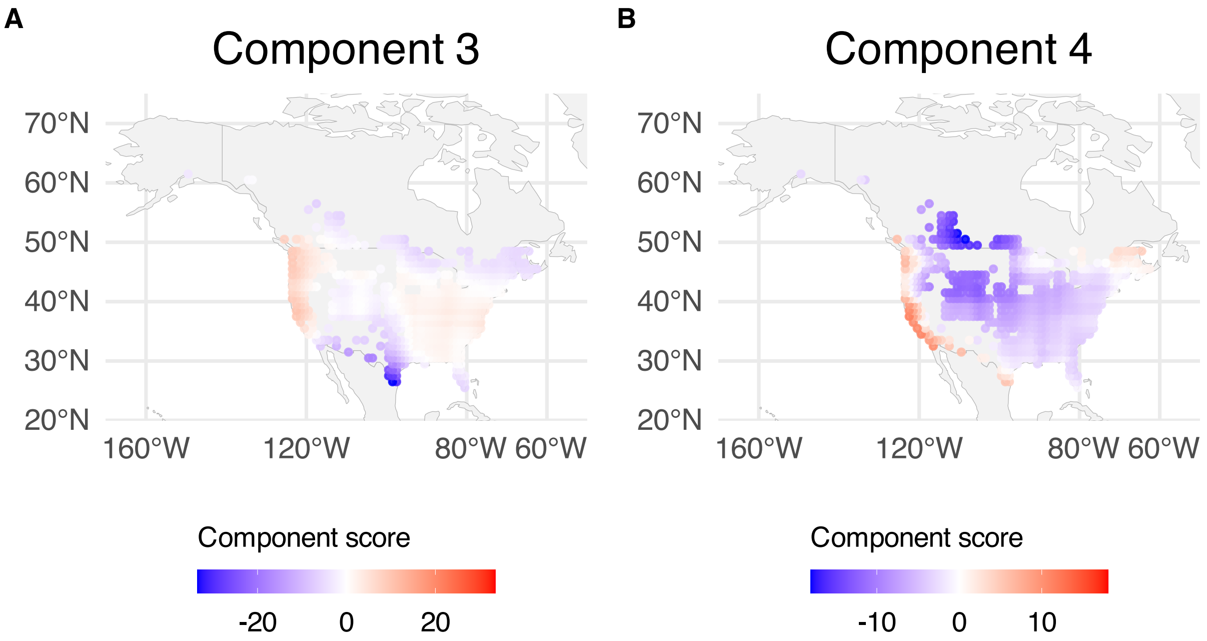


**Supplementary Figure 223.** SSANOVA-SVD reconstructed component functions mapped over BBS sampling locations; component scores are averaged over years within spatial grid. (A) Component 3. (B) Component 4.

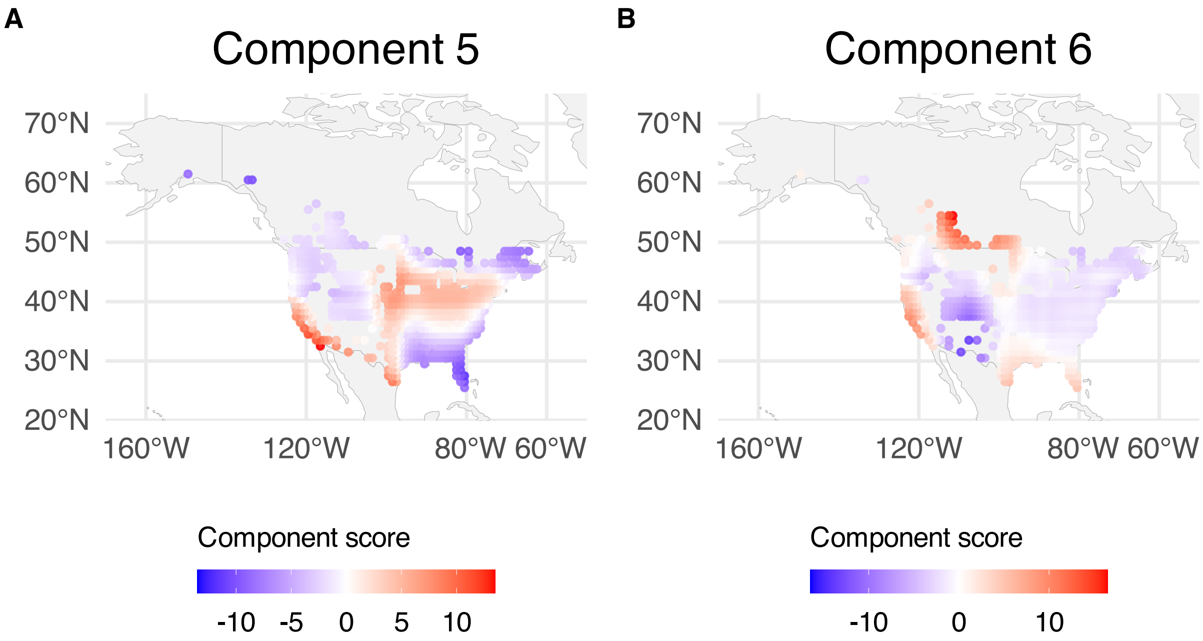


**Supplementary Figure 224.** SSANOVA-SVD reconstructed component functions mapped over BBS sampling locations; component scores are averaged over years within spatial grid. (A) Component 5. (B) Component 6.

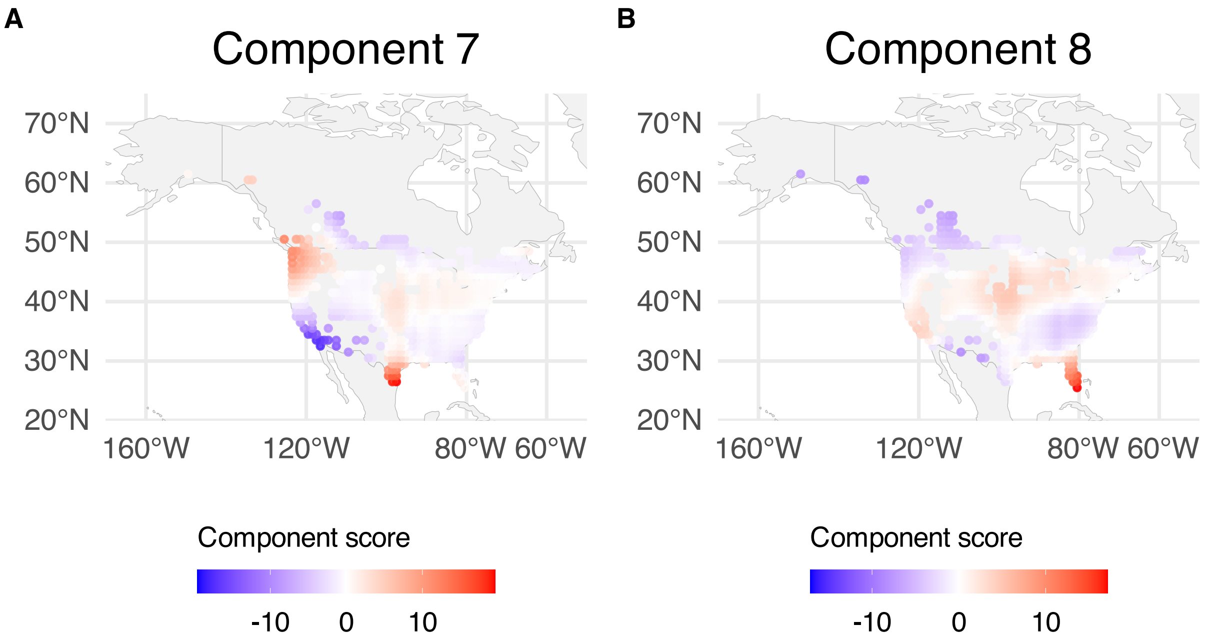


**Supplementary Figure 225.** SSANOVA-SVD reconstructed component functions mapped over BBS sampling locations; component scores are averaged over years within spatial grid. (A) Component 7. (B) Component 8.

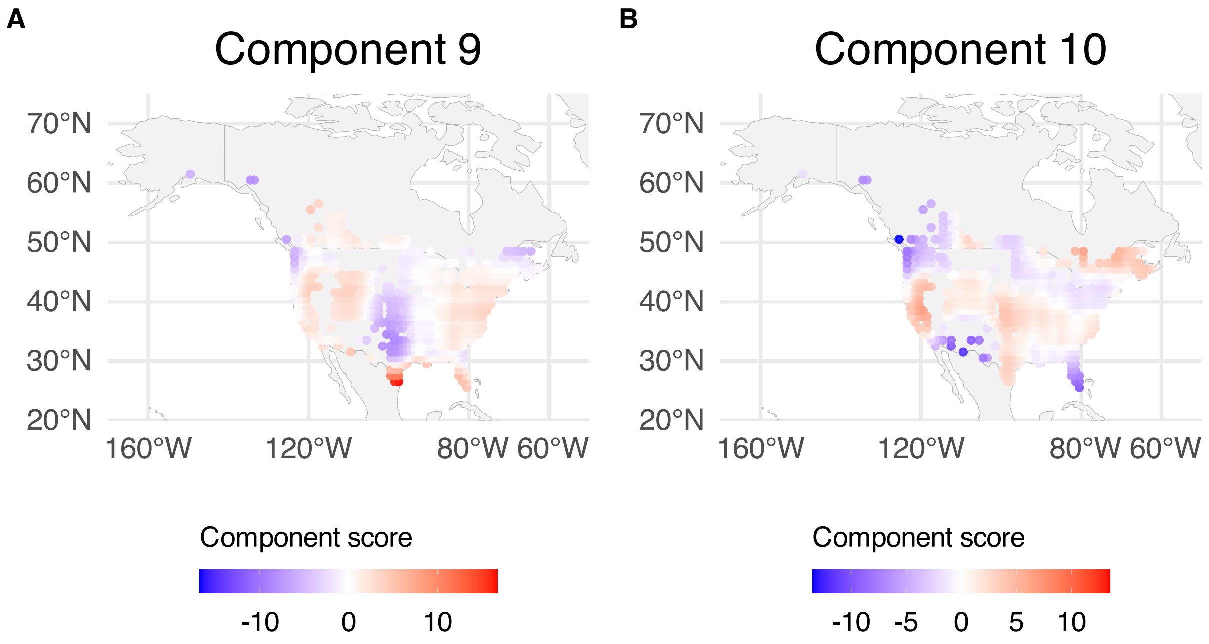


**Supplementary Figure 226.** SSANOVA-SVD reconstructed component functions mapped over BBS sampling locations; component scores are averaged over years within spatial grid. (A) Component 9. (B) Component 10.

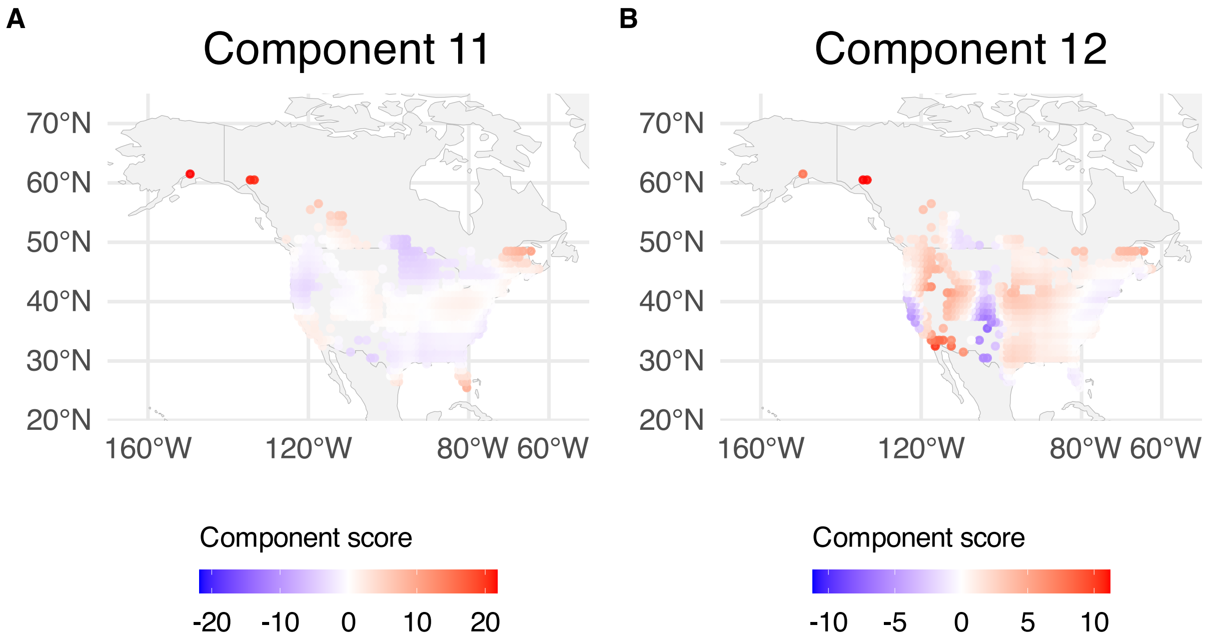


**Supplementary Figure 227.** SSANOVA-SVD reconstructed component functions mapped over BBS sampling locations; component scores are averaged over years within spatial grid. (A) Component 11. (B) Component 12.

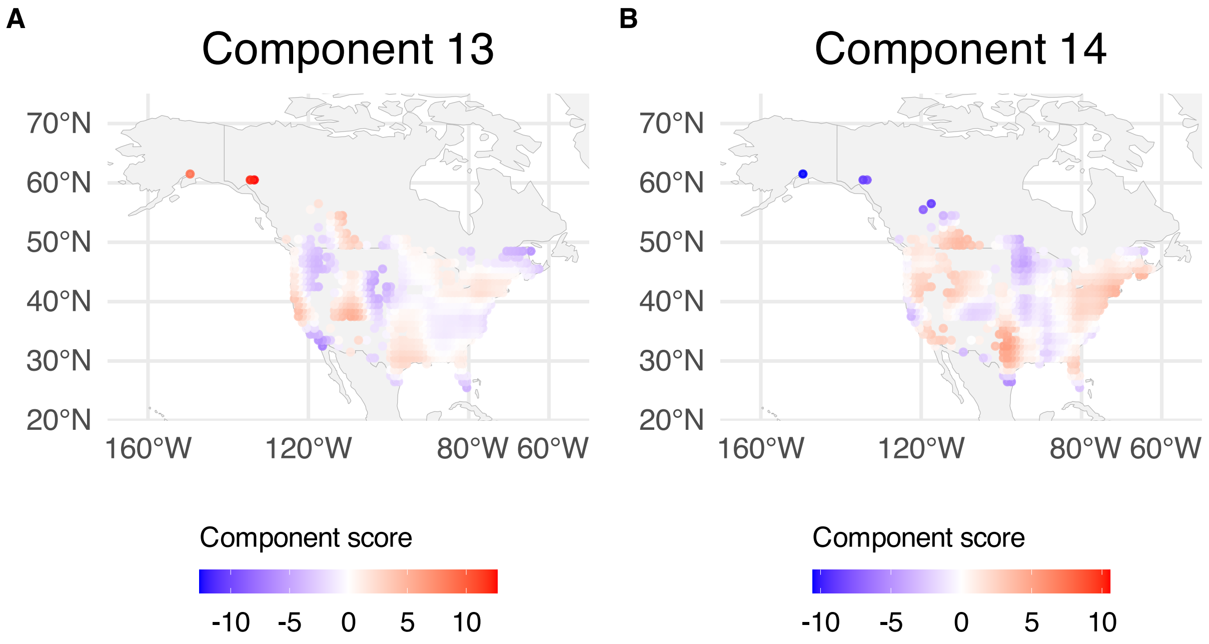


**Supplementary Figure 228.** SSANOVA-SVD reconstructed component functions mapped over BBS sampling locations; component scores are averaged over years within spatial grid. (A) Component 13. (B) Component 14.

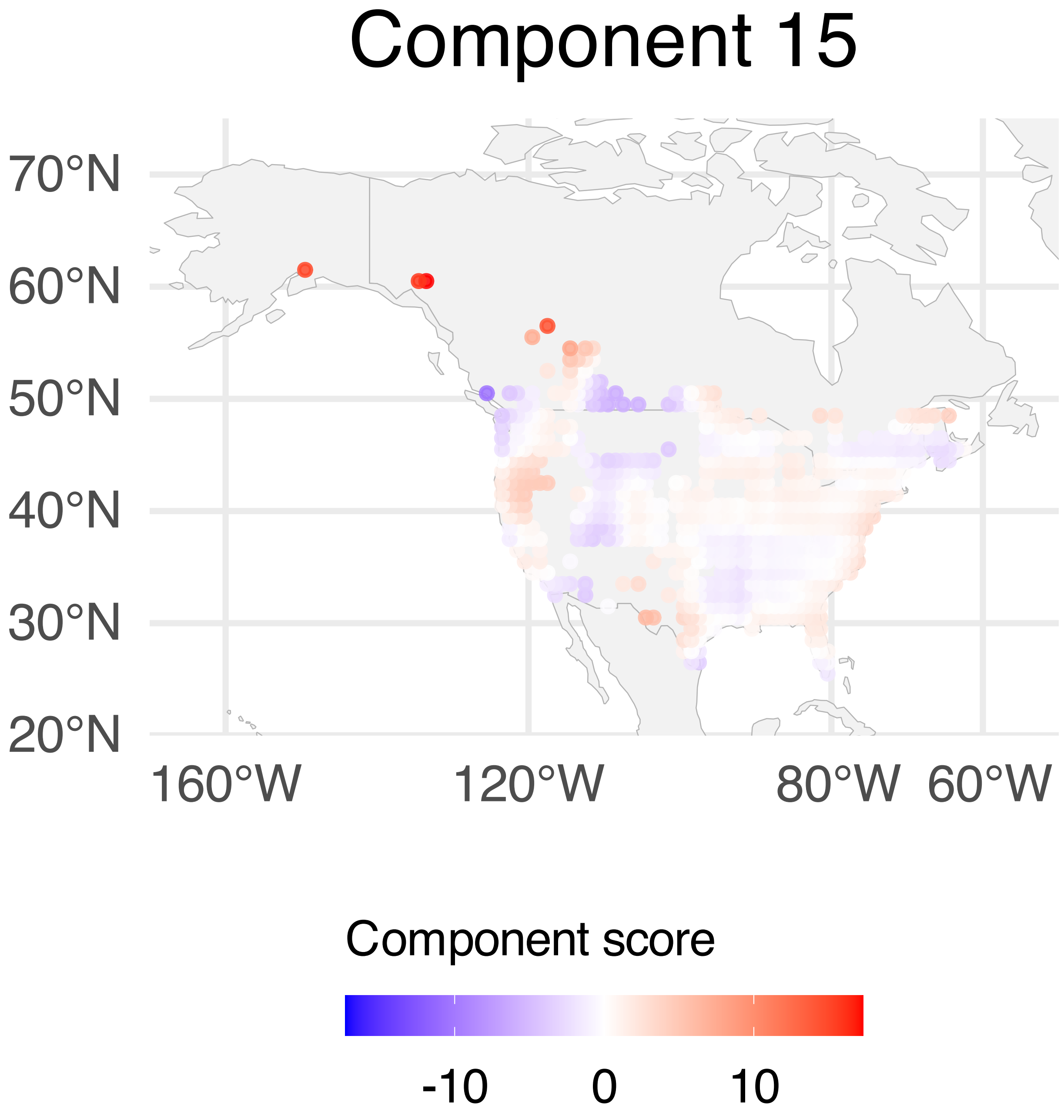


**Supplementary Figure 229.** SSANOVA-SVD reconstructed component functions mapped over BBS sampling locations; component scores are averaged over years within spatial grid. Component 15.

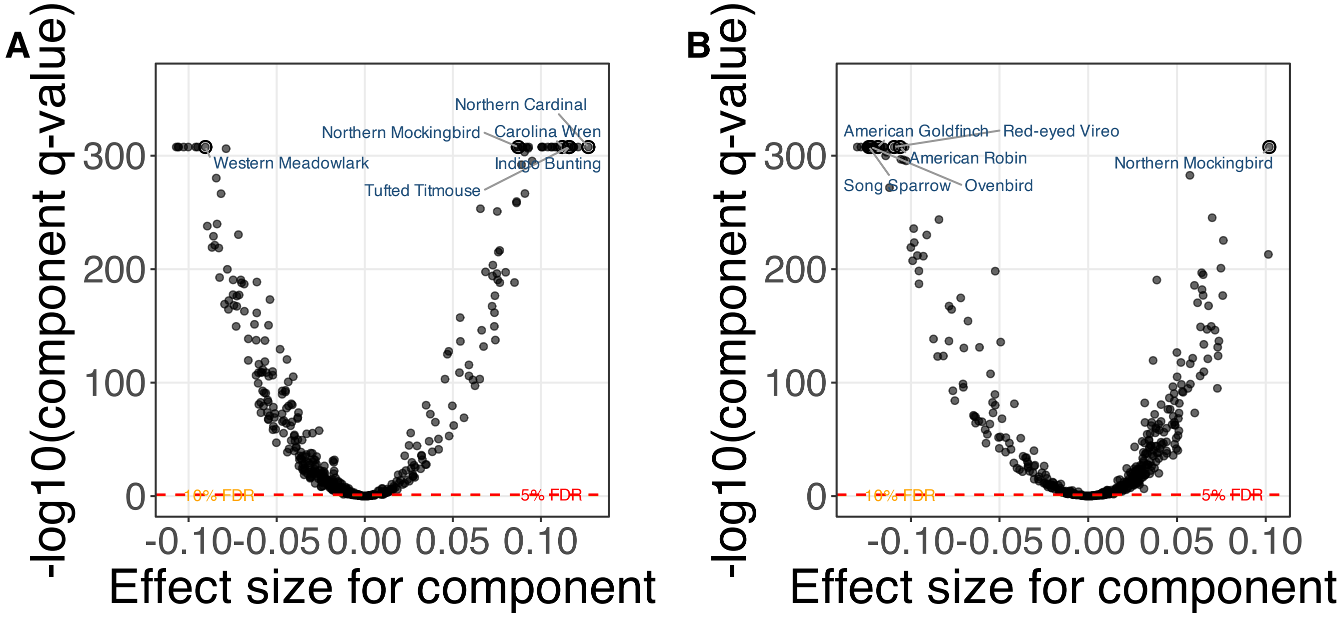


**Supplementary Figure 230.** SSANOVA-SVD based conditional inference for feature participation. (A) Component 1. (B) Component 2.

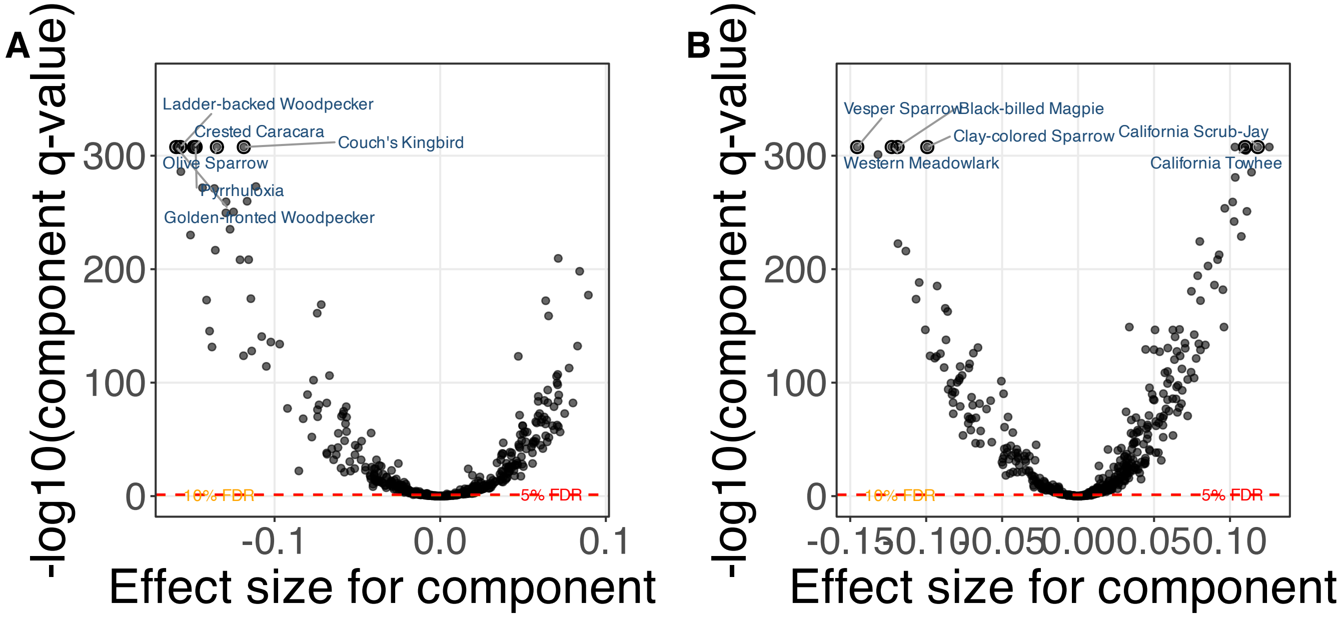


**Supplementary Figure 231.** SSANOVA-SVD based conditional inference for feature participation. (A) Component 3. (B) Component 4.

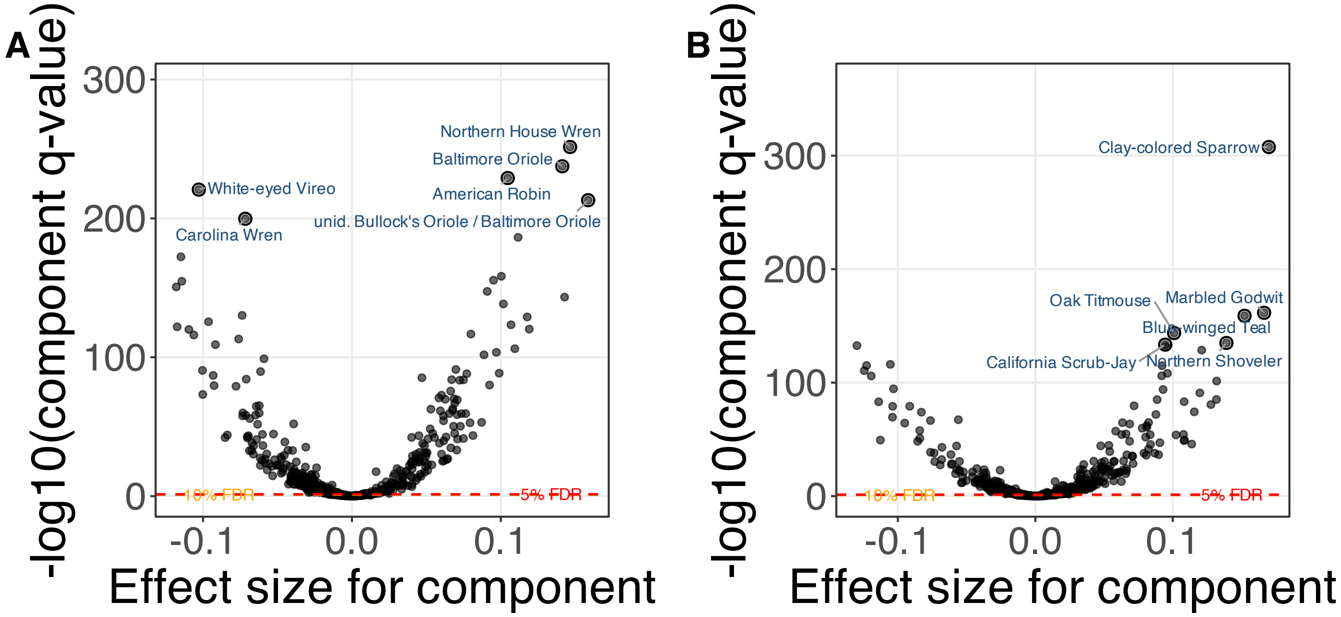


**Supplementary Figure 232.** SSANOVA-SVD based conditional inference for feature participation. (A) Component 5. (B) Component 6.

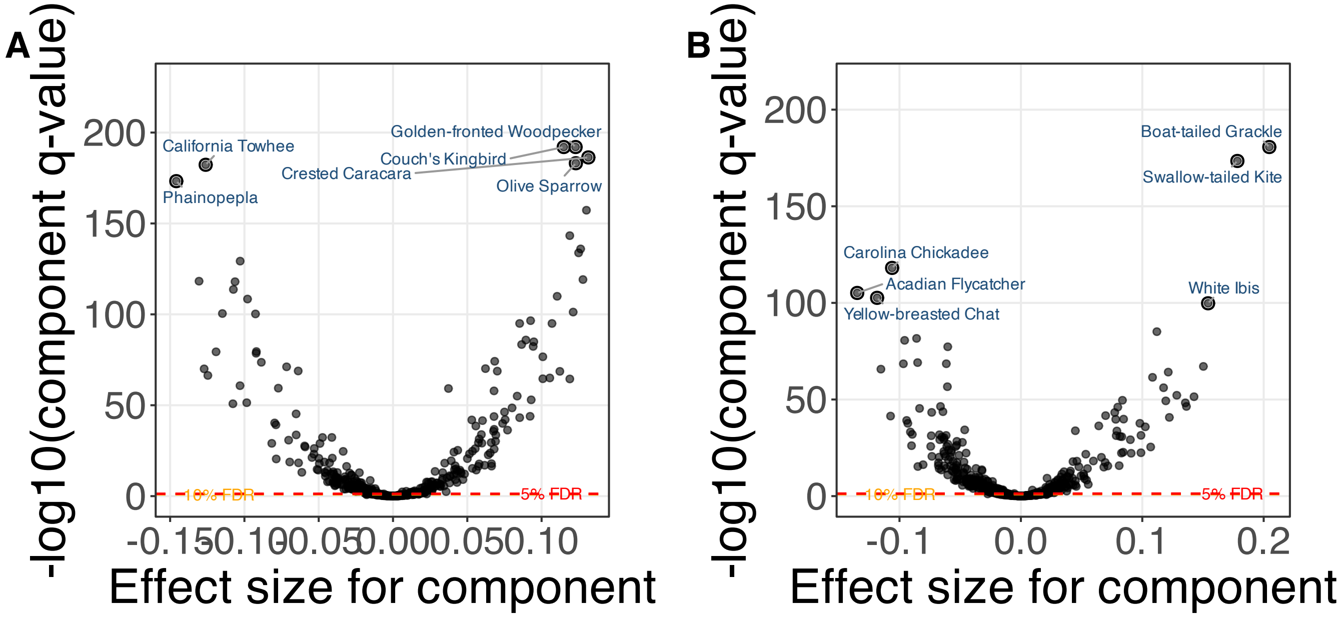


**Supplementary Figure 233.** SSANOVA-SVD based conditional inference for feature participation. (A) Component 7. (B) Component 8.

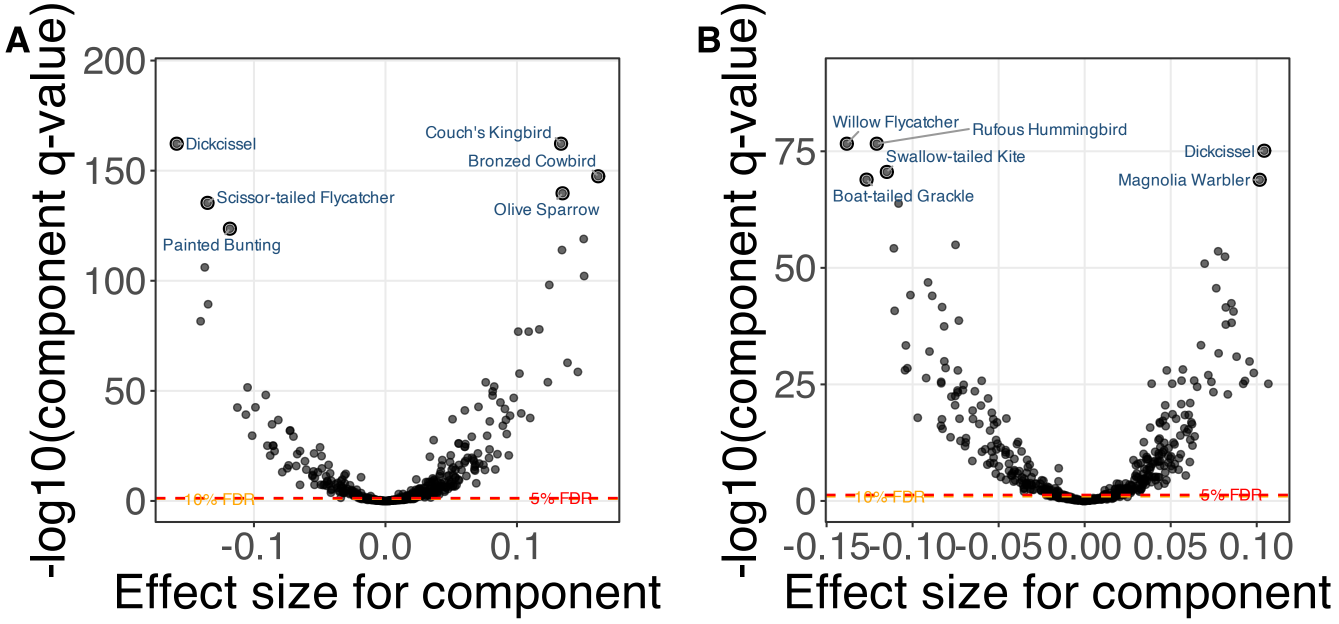


**Supplementary Figure 234.** SSANOVA-SVD based conditional inference for feature participation. (A) Component 9. (B) Component 10.

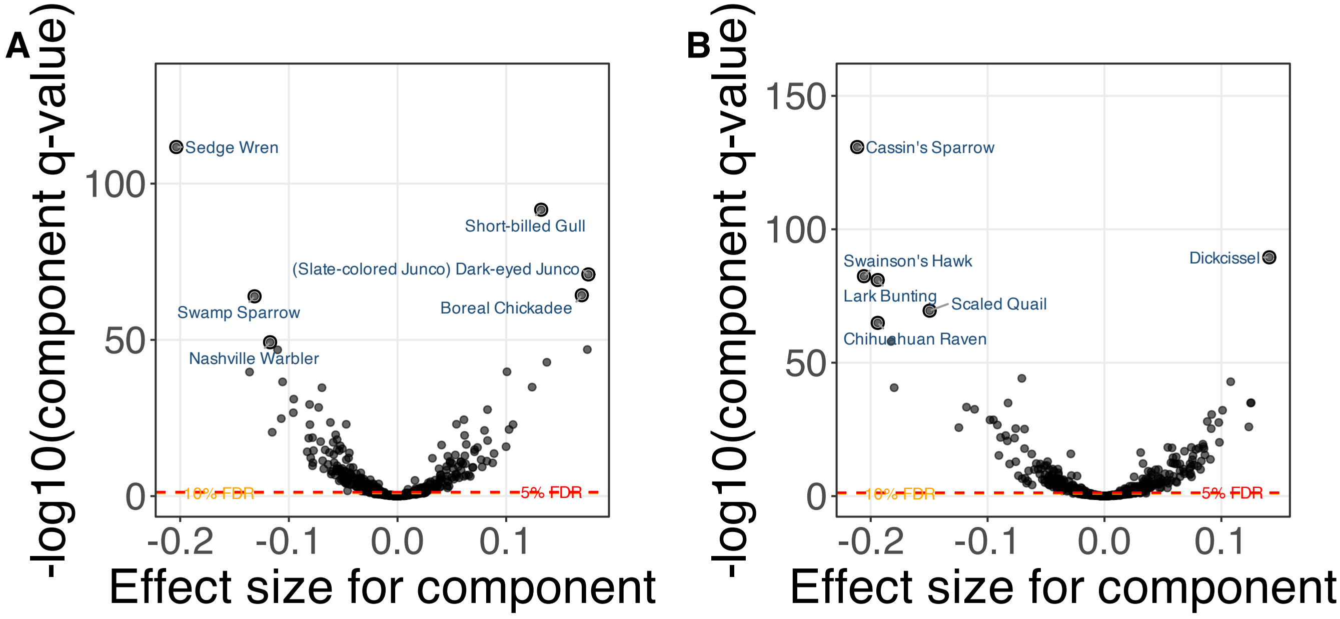


**Supplementary Figure 235.** SSANOVA-SVD based conditional inference for feature participation. (A) Component 11. (B) Component 12.

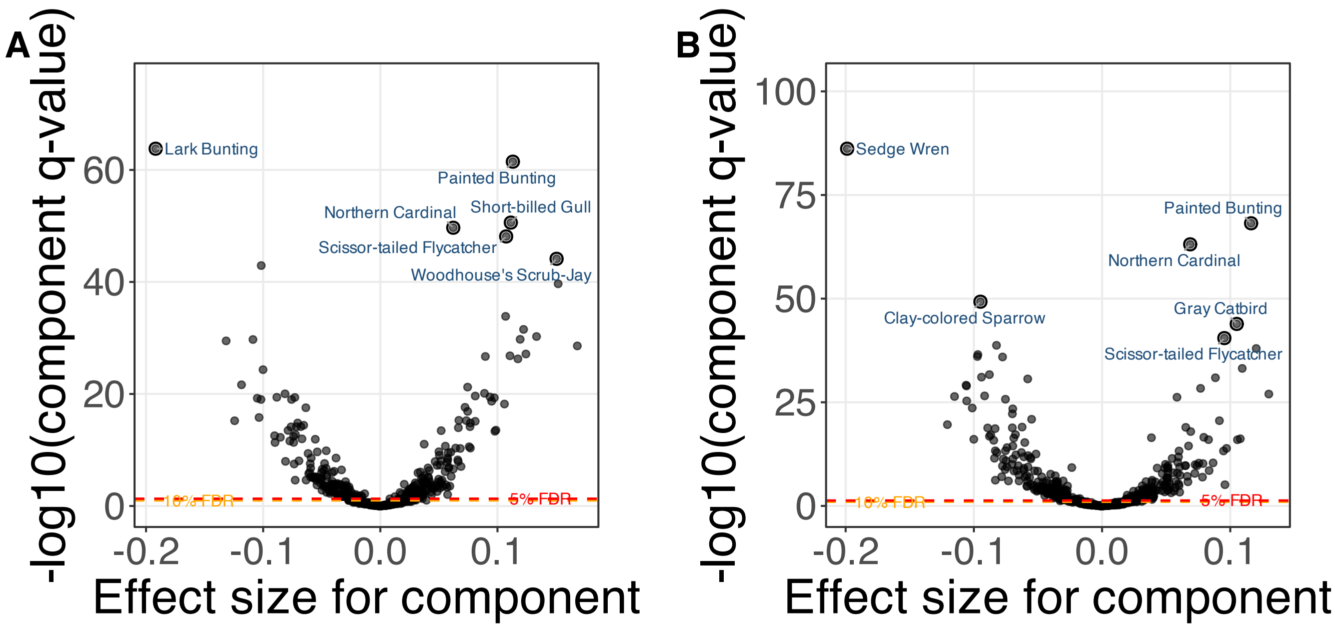


**Supplementary Figure 236.** SSANOVA-SVD based conditional inference for feature participation. (A) Component 13. (B) Component 14.

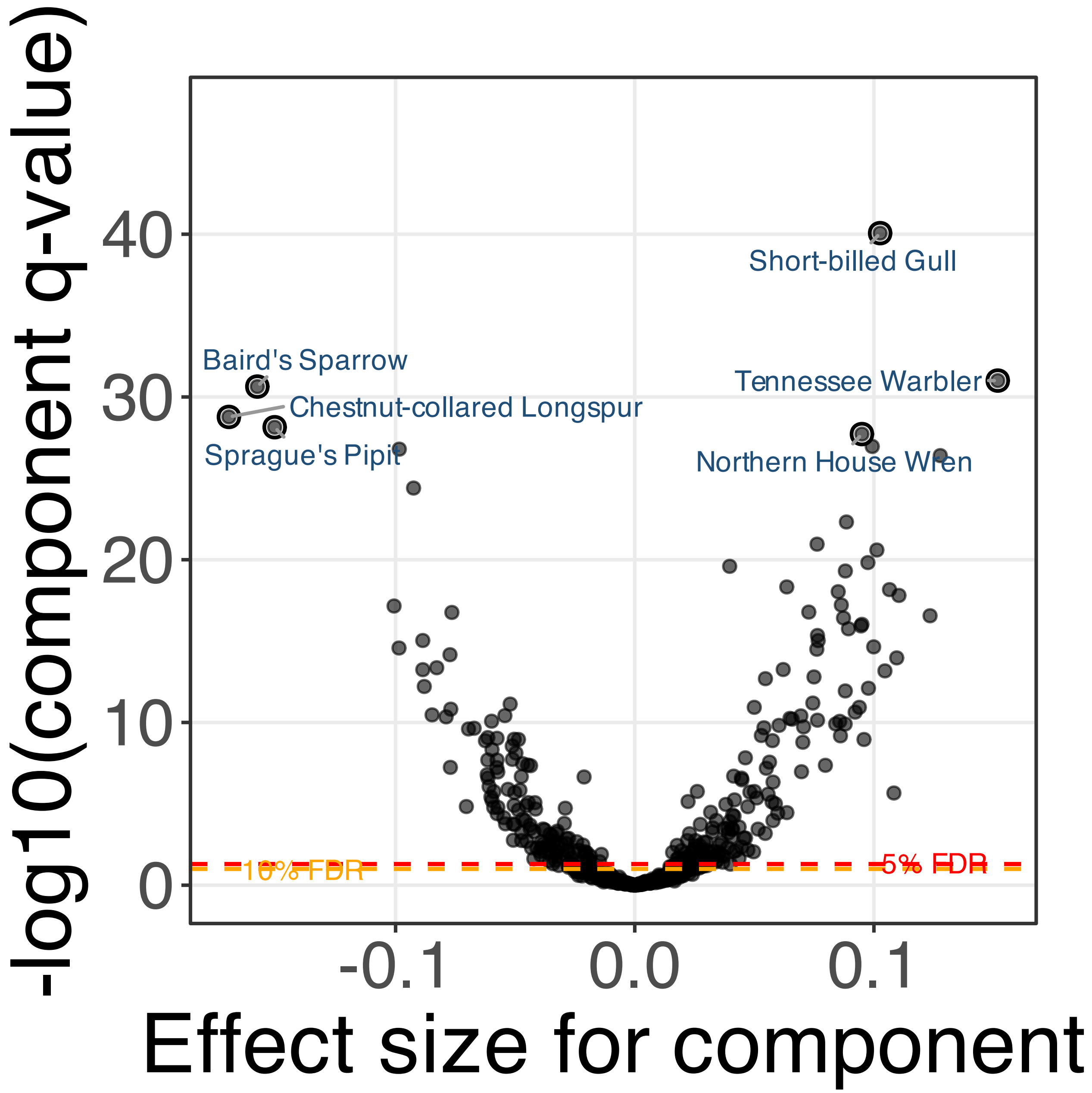


**Supplementary Figure 237.** SSANOVA-SVD based conditional inference for feature participation. Component 15.

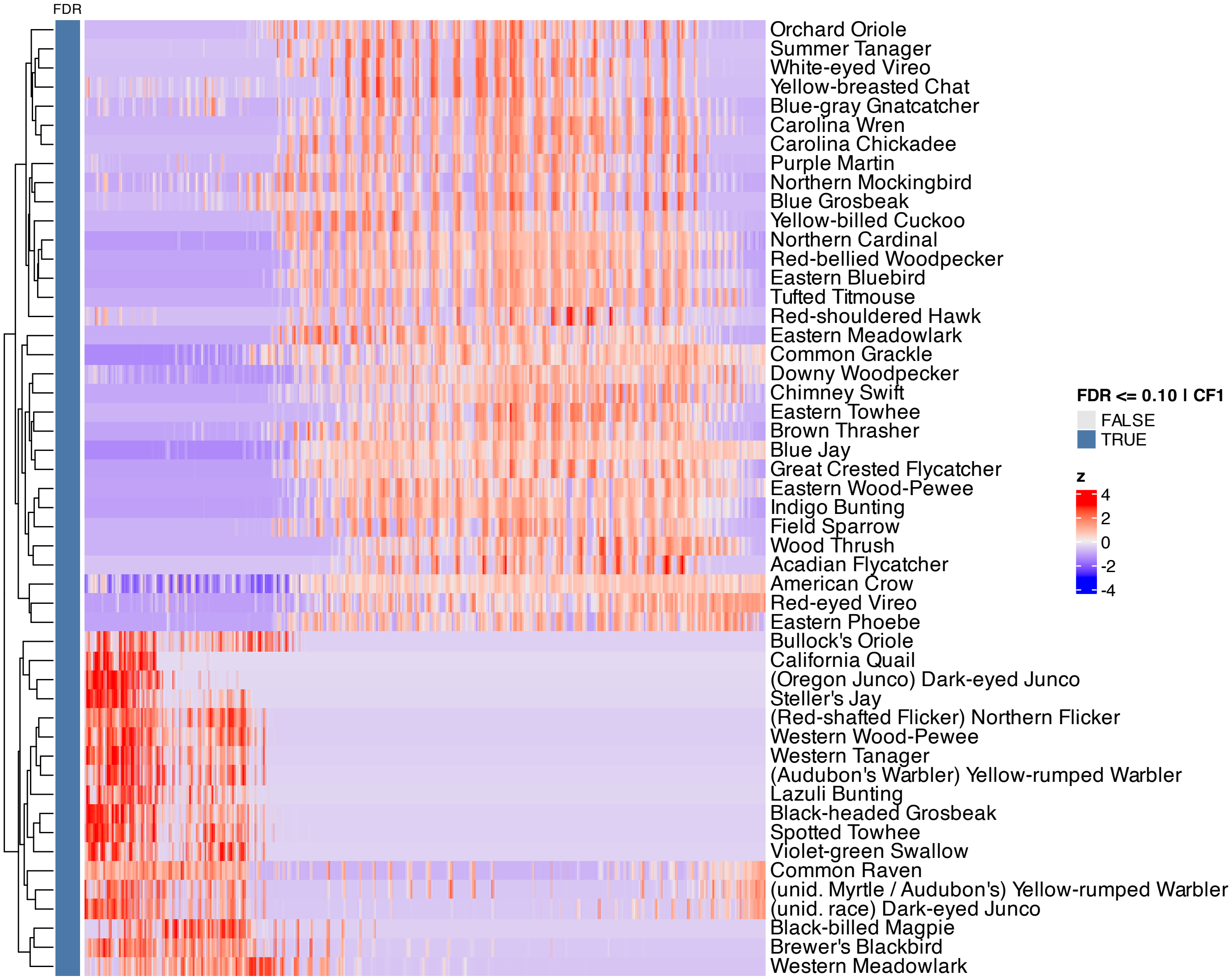


**Supplementary Figure 238.** Heatmap of features inferred to be associated with component 1 by SSANOVA-SVD.

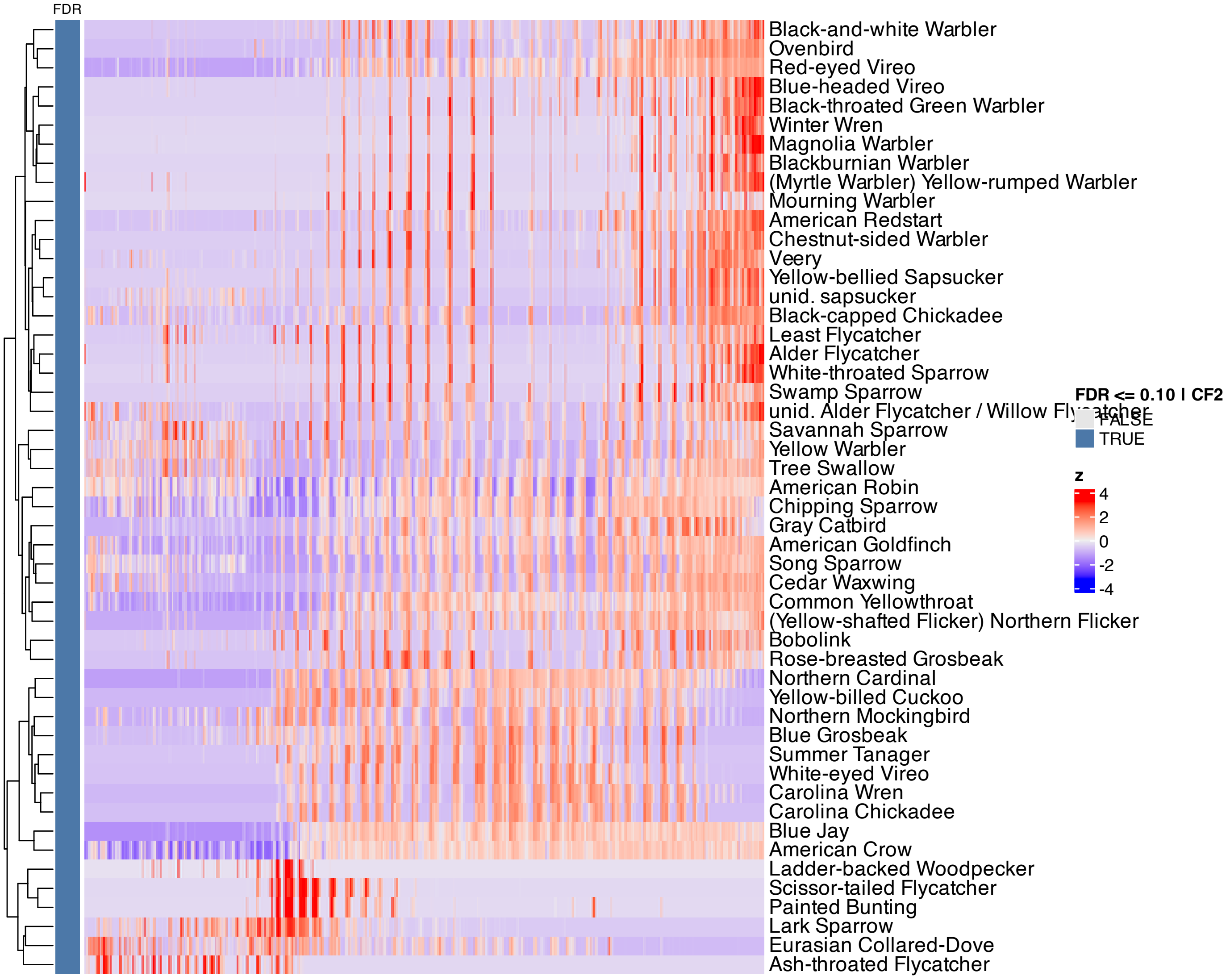


**Supplementary Figure 239.** Heatmap of features inferred to be associated with component 2 by SSANOVA-SVD.

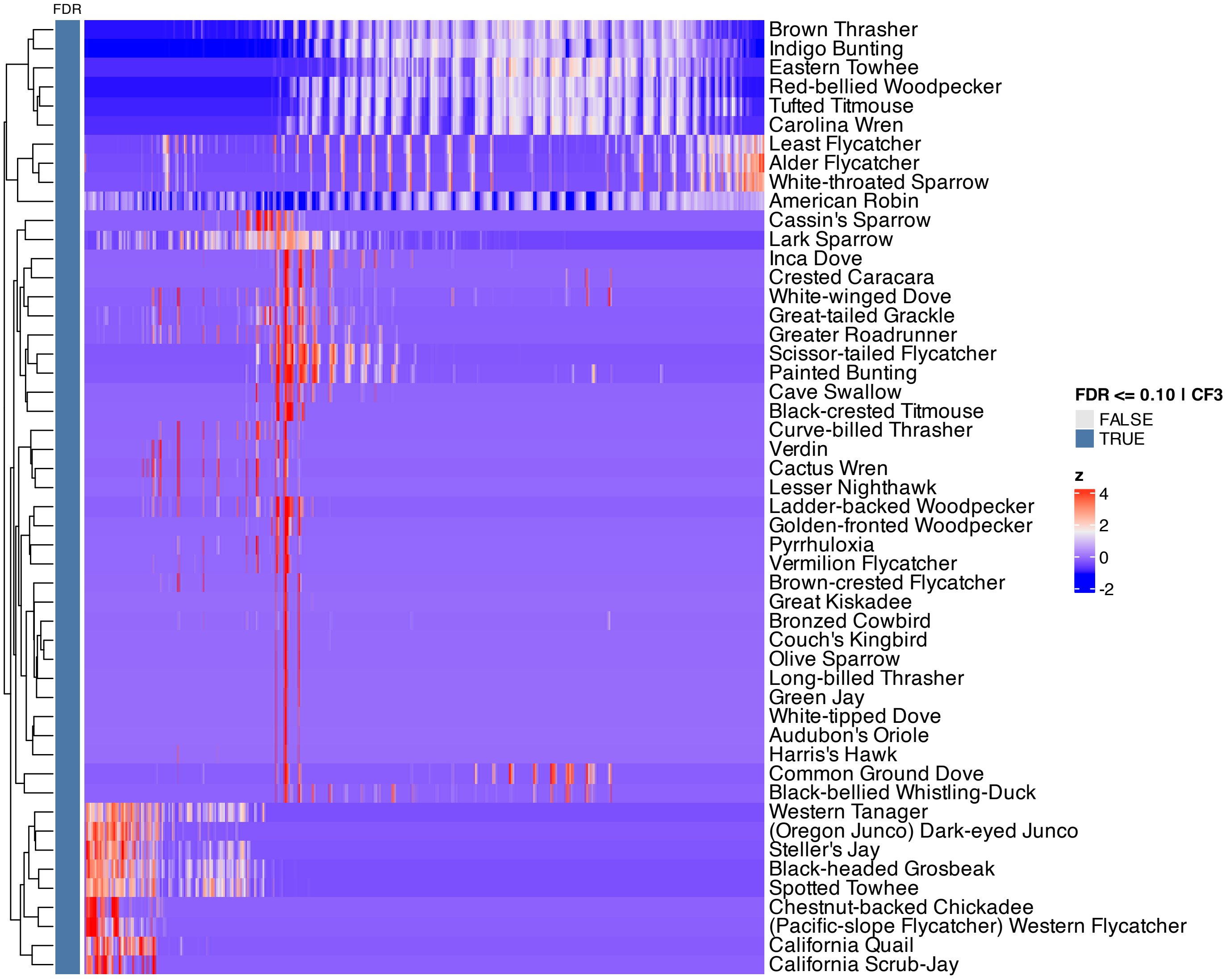


**Supplementary Figure 240.** Heatmap of features inferred to be associated with component 3 by SSANOVA-SVD.

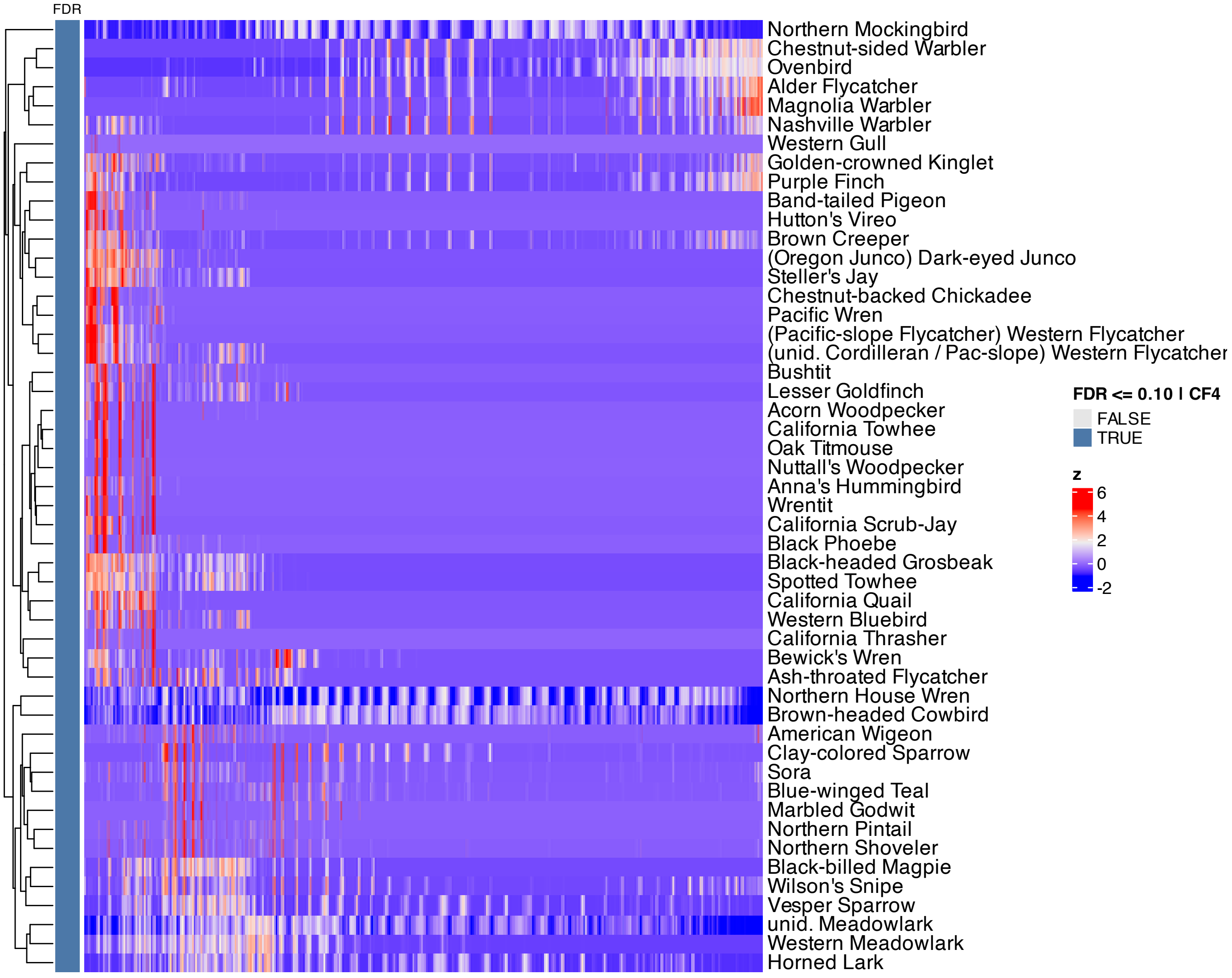


**Supplementary Figure 241.** Heatmap of features inferred to be associated with component 4 by SSANOVA-SVD.

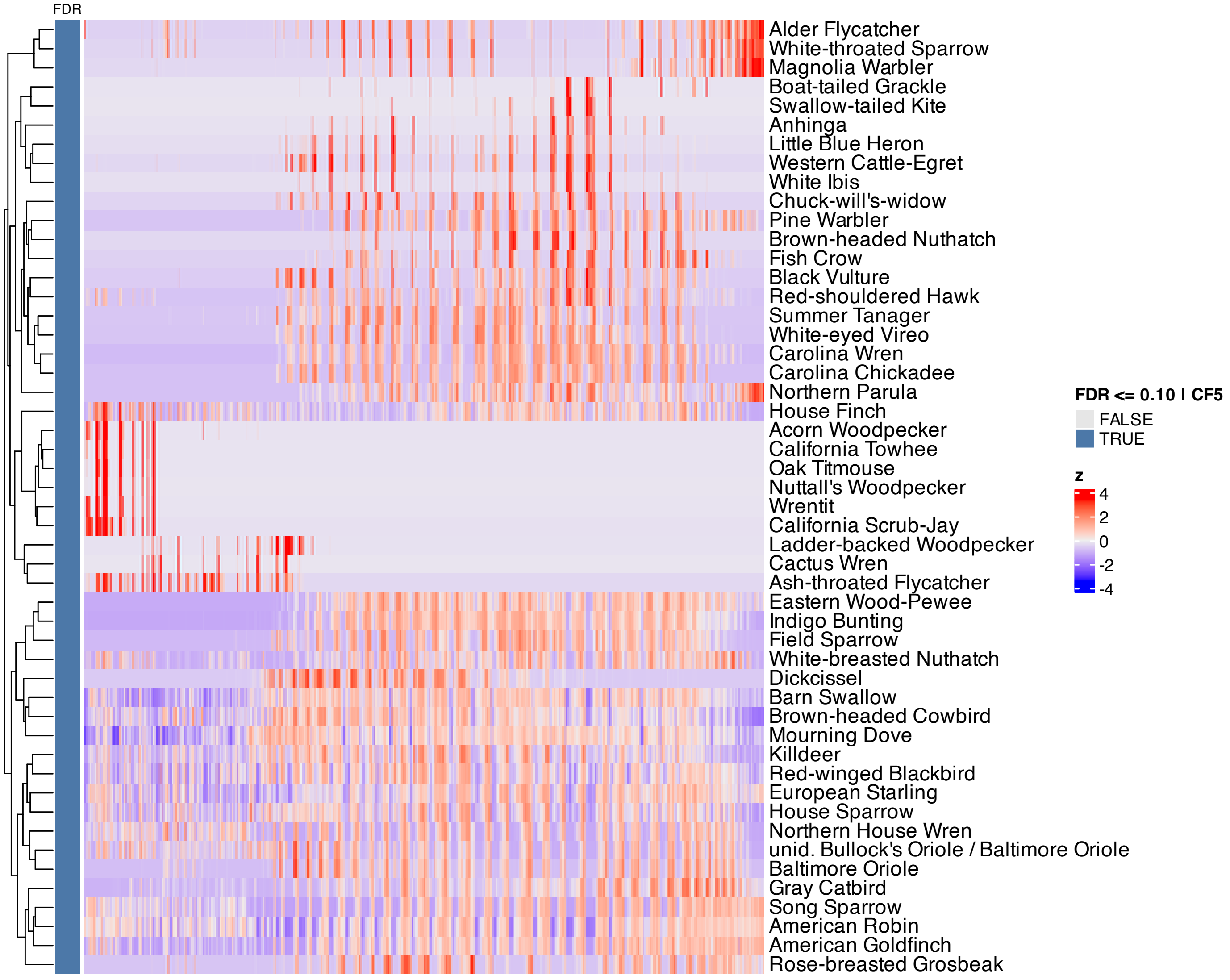


**Supplementary Figure 242.** Heatmap of features inferred to be associated with component 5 by SSANOVA-SVD.

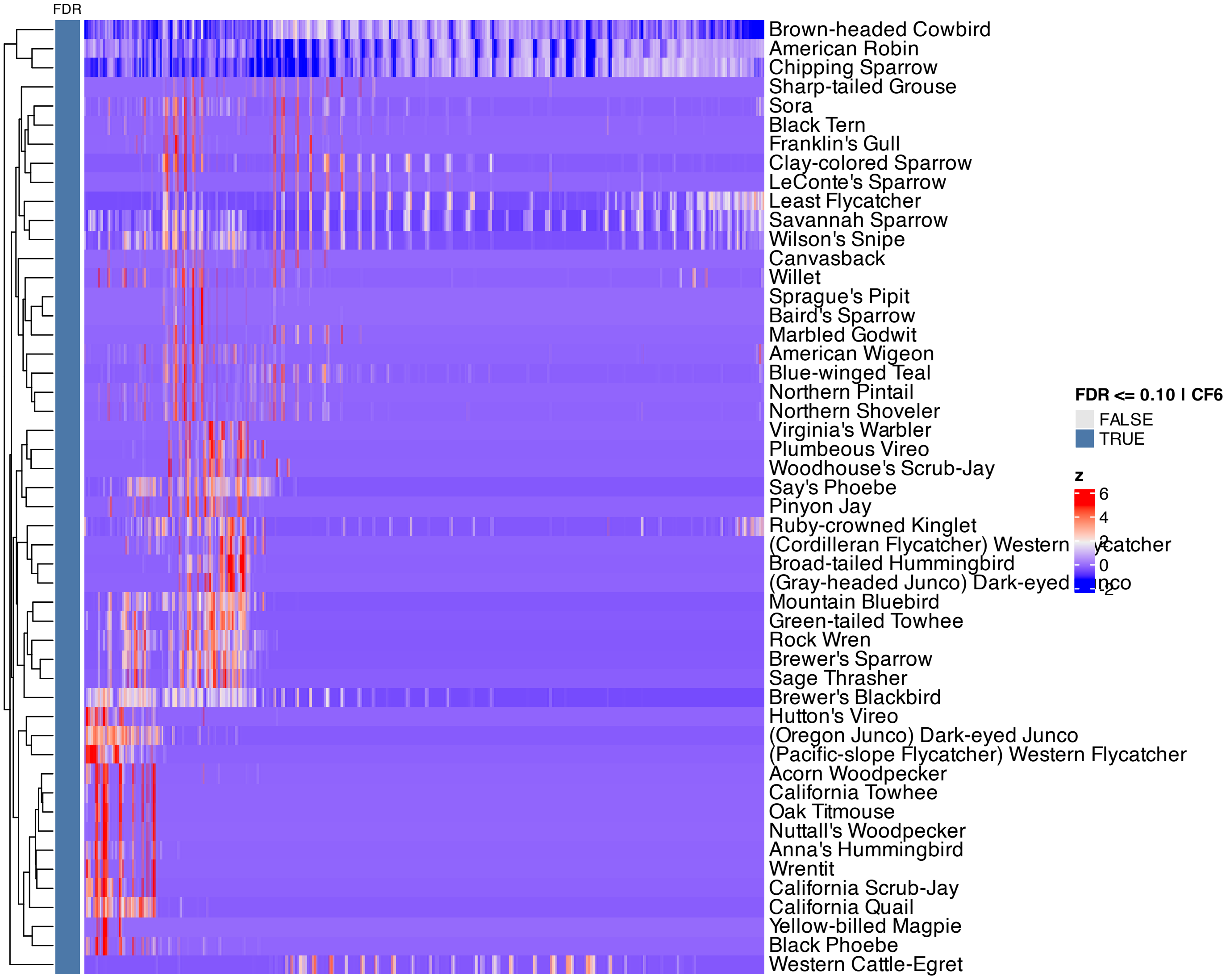


**Supplementary Figure 243.** Heatmap of features inferred to be associated with component 6 by SSANOVA-SVD.

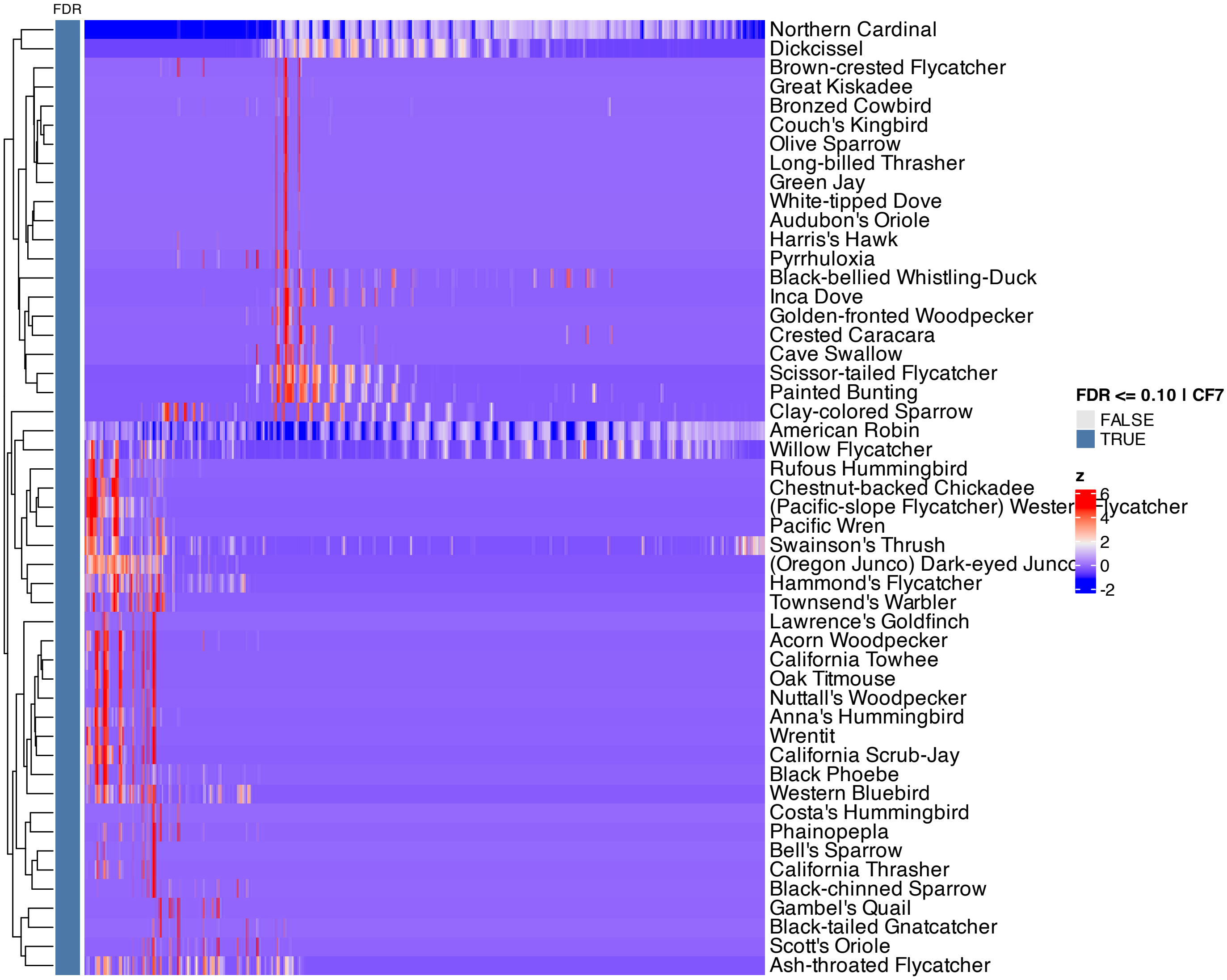


**Supplementary Figure 244.** Heatmap of features inferred to be associated with component 7 by SSANOVA-SVD.

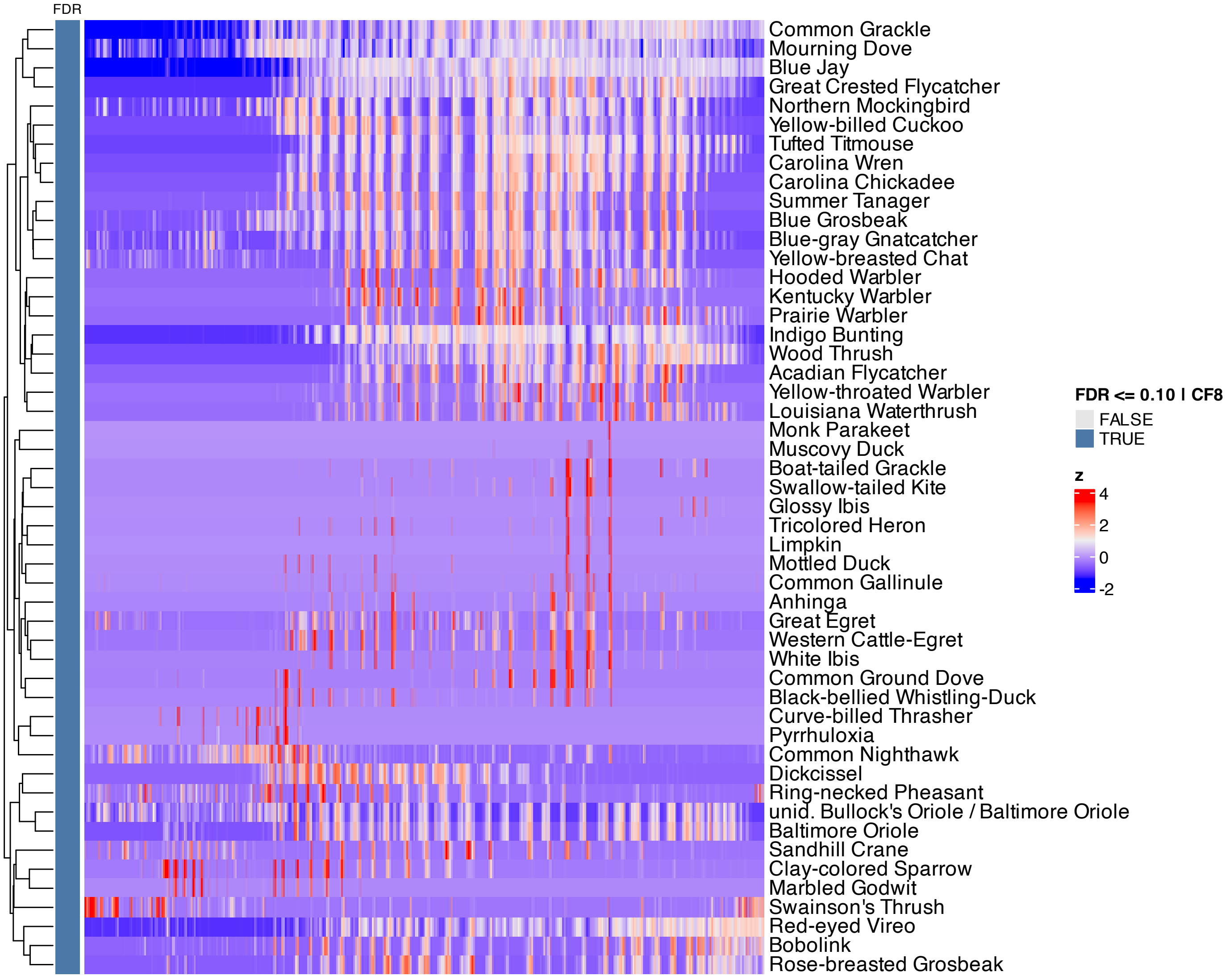


**Supplementary Figure 245.** Heatmap of features inferred to be associated with component 8 by SSANOVA-SVD.

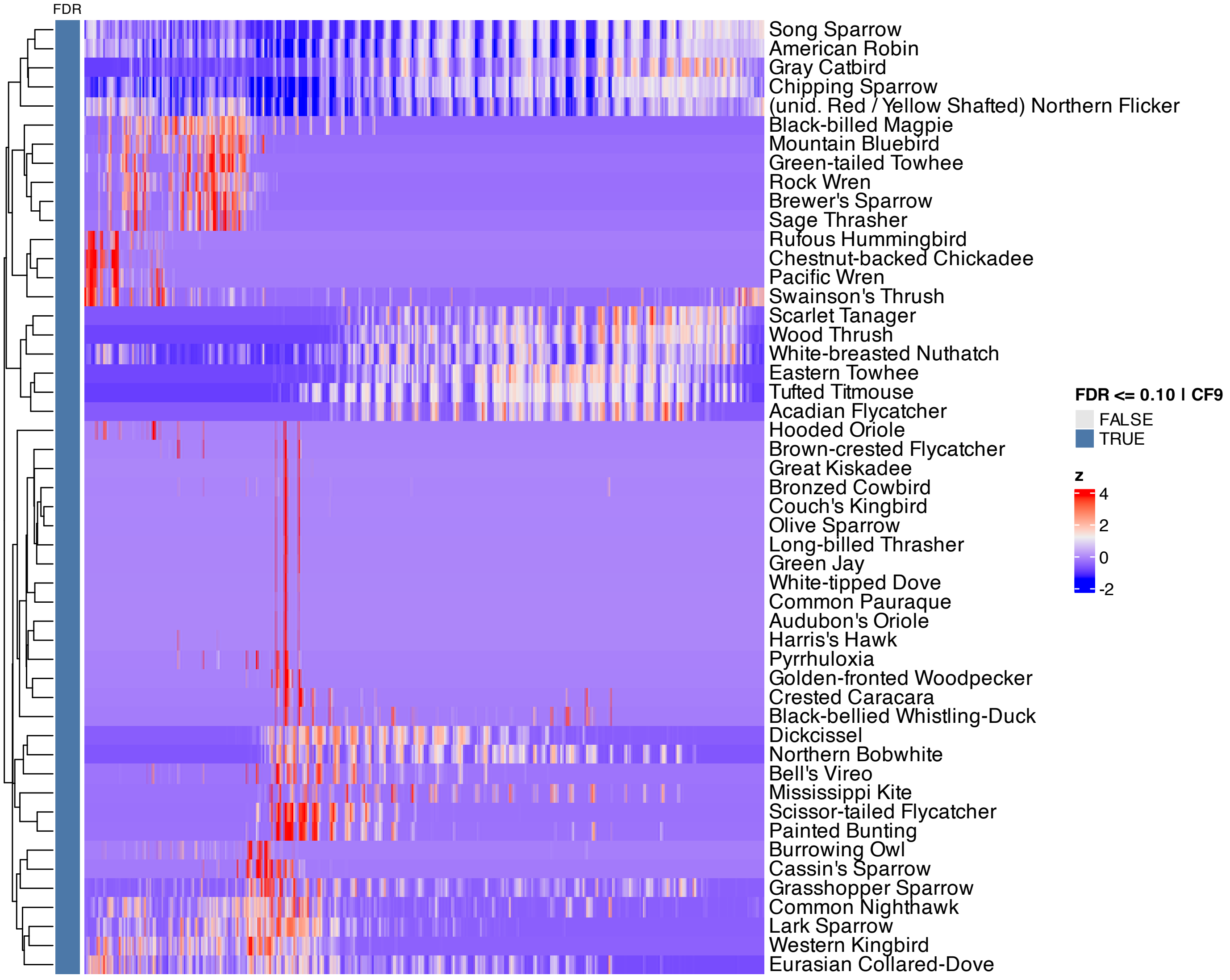


**Supplementary Figure 246.** Heatmap of features inferred to be associated with component 9 by SSANOVA-SVD.

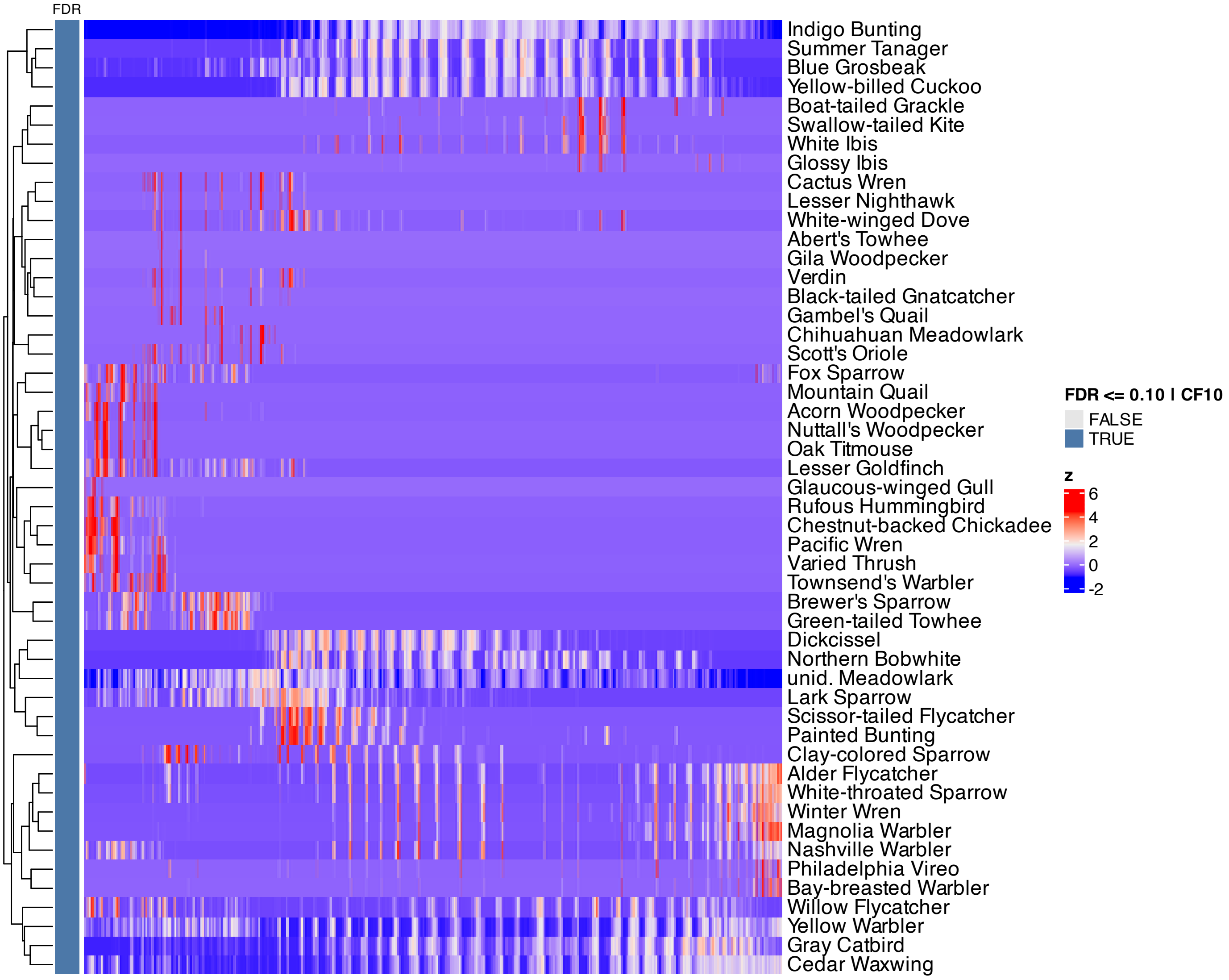


**Supplementary Figure 247.** Heatmap of features inferred to be associated with component 10 by SSANOVA-SVD.

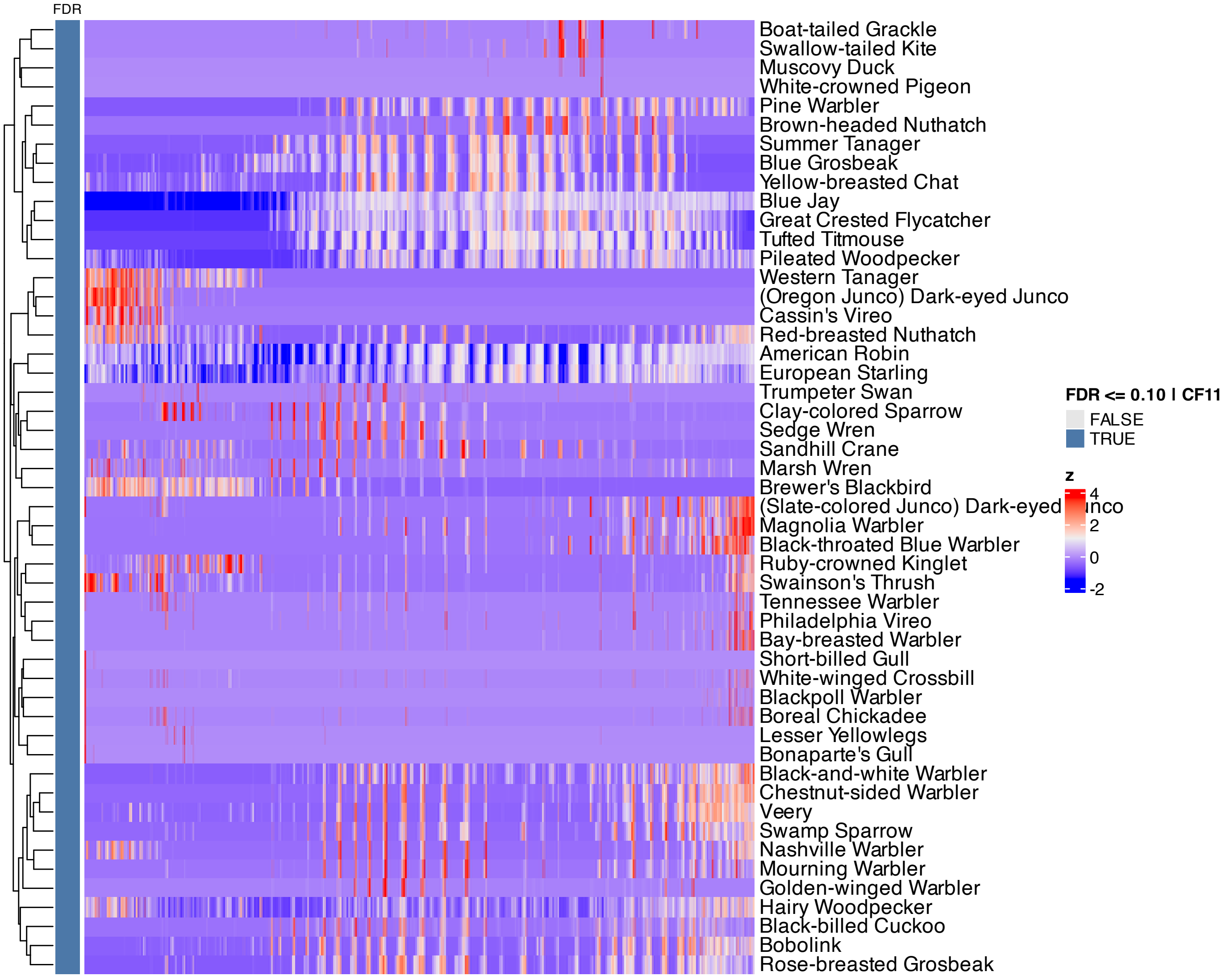


**Supplementary Figure 248.** Heatmap of features inferred to be associated with component 11 by SSANOVA-SVD.

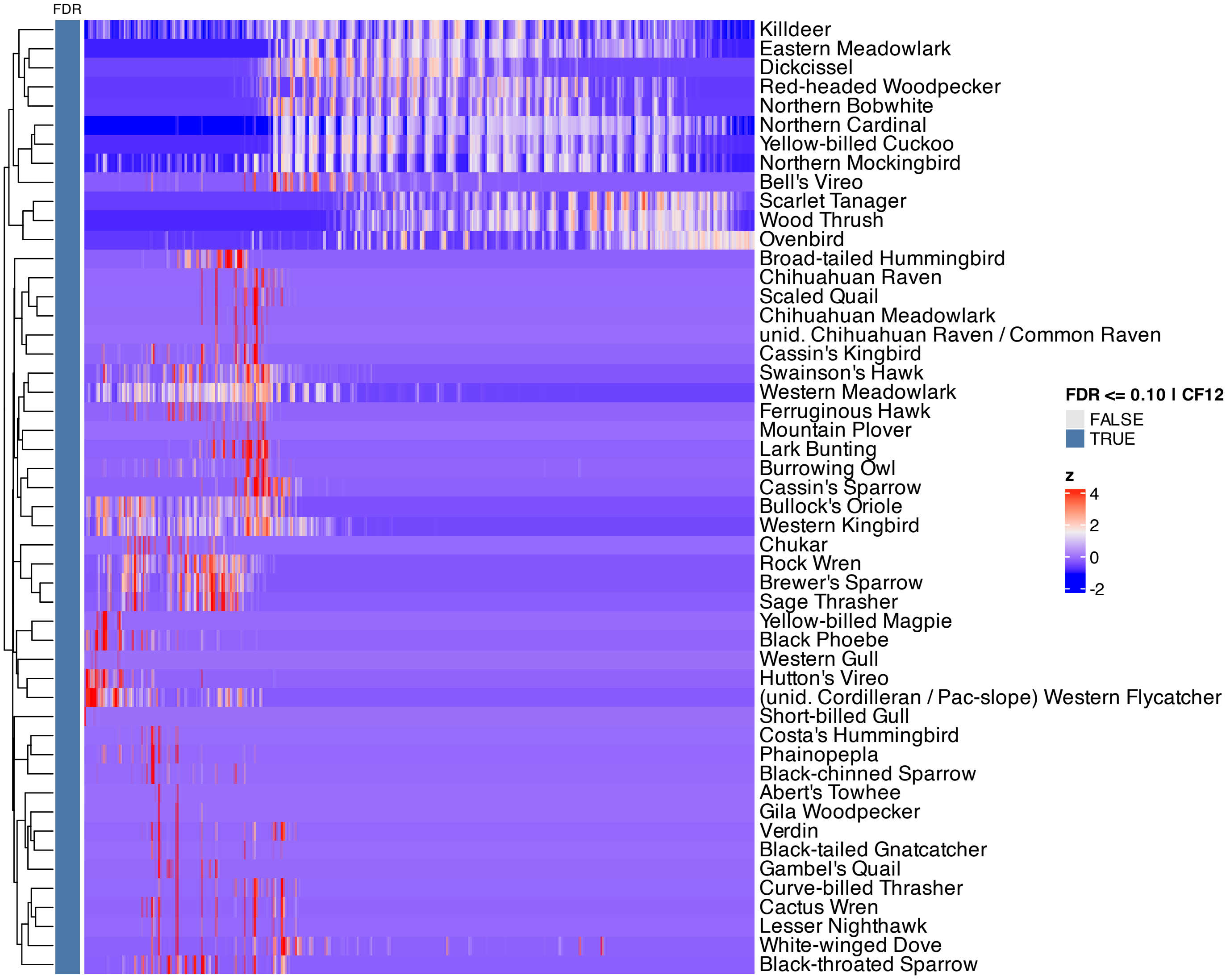


**Supplementary Figure 249.** Heatmap of features inferred to be associated with component 12 by SSANOVA-SVD.

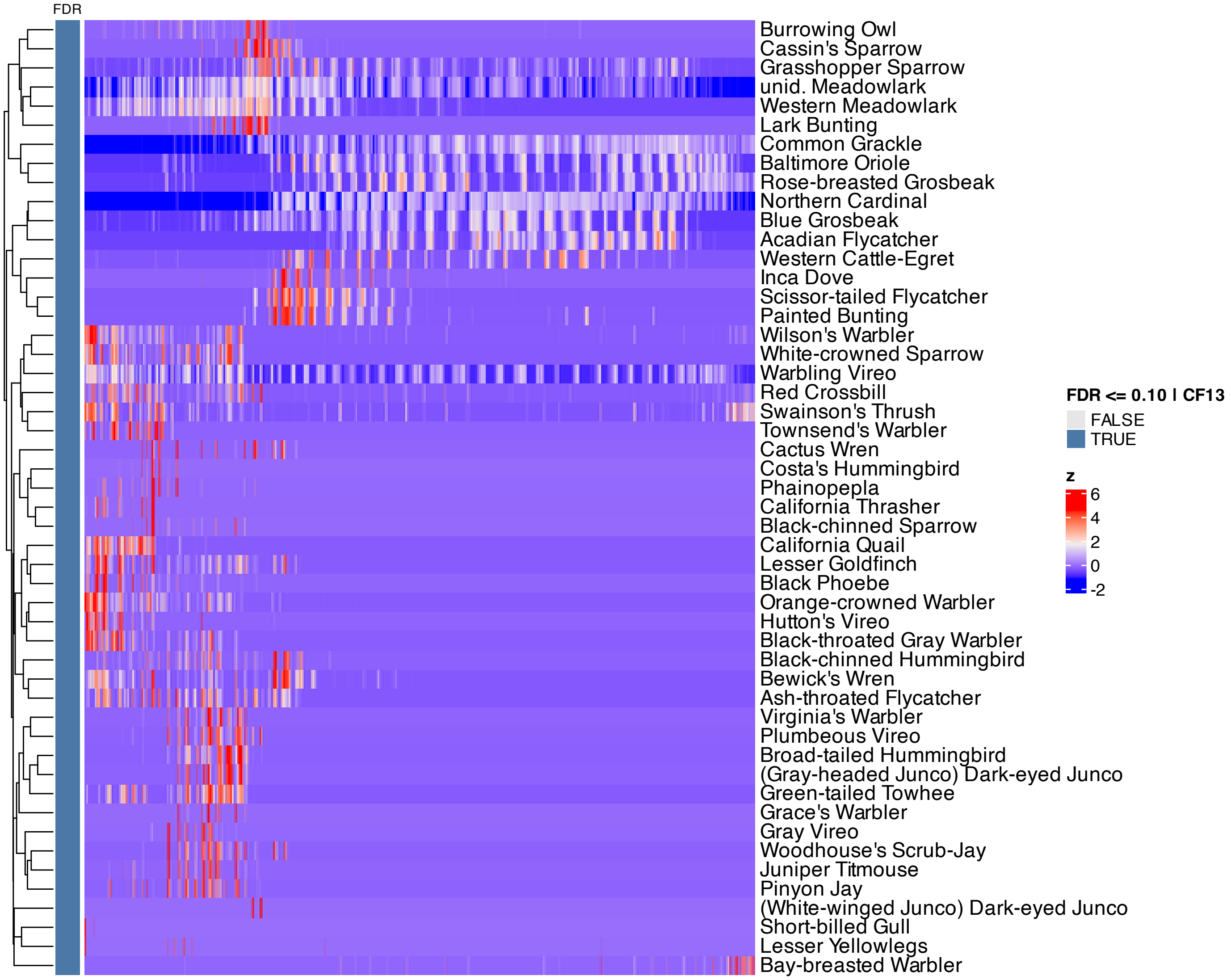


**Supplementary Figure 250.** Heatmap of features inferred to be associated with component 13 by SSANOVA-SVD.

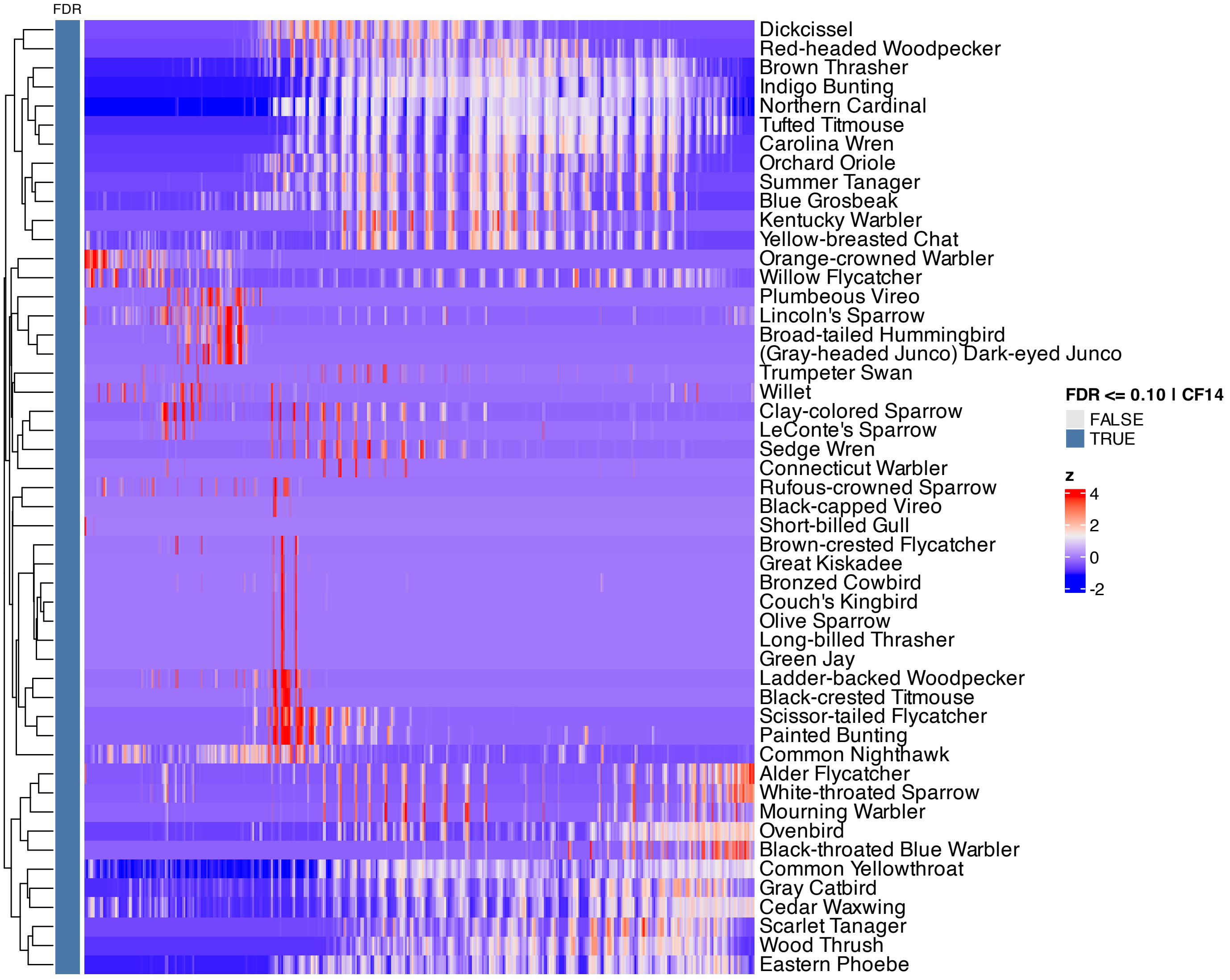


**Supplementary Figure 251.** Heatmap of features inferred to be associated with component 14 by SSANOVA-SVD.

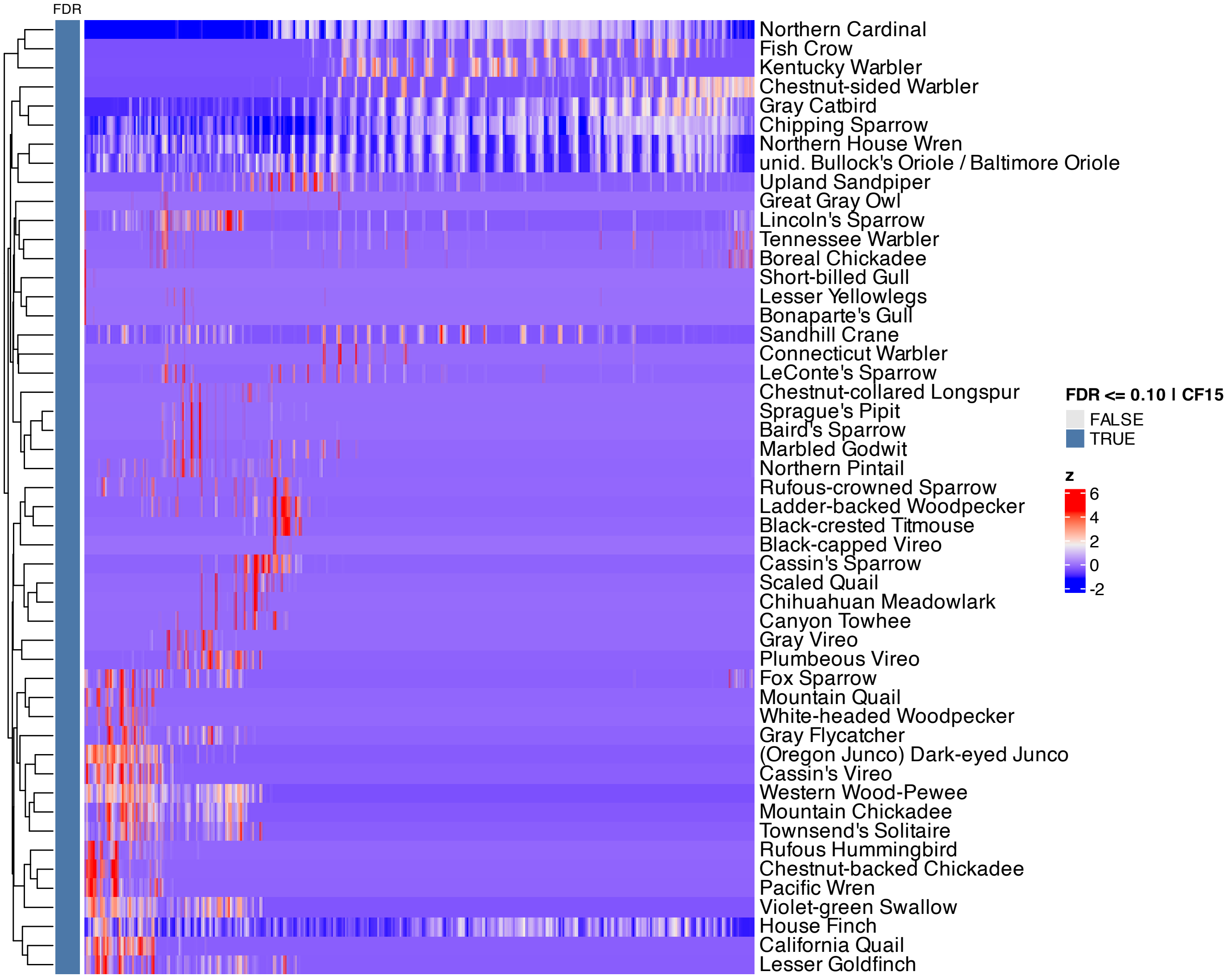


**Supplementary Figure 252.** Heatmap of features inferred to be associated with component 15 by SSANOVA-SVD.

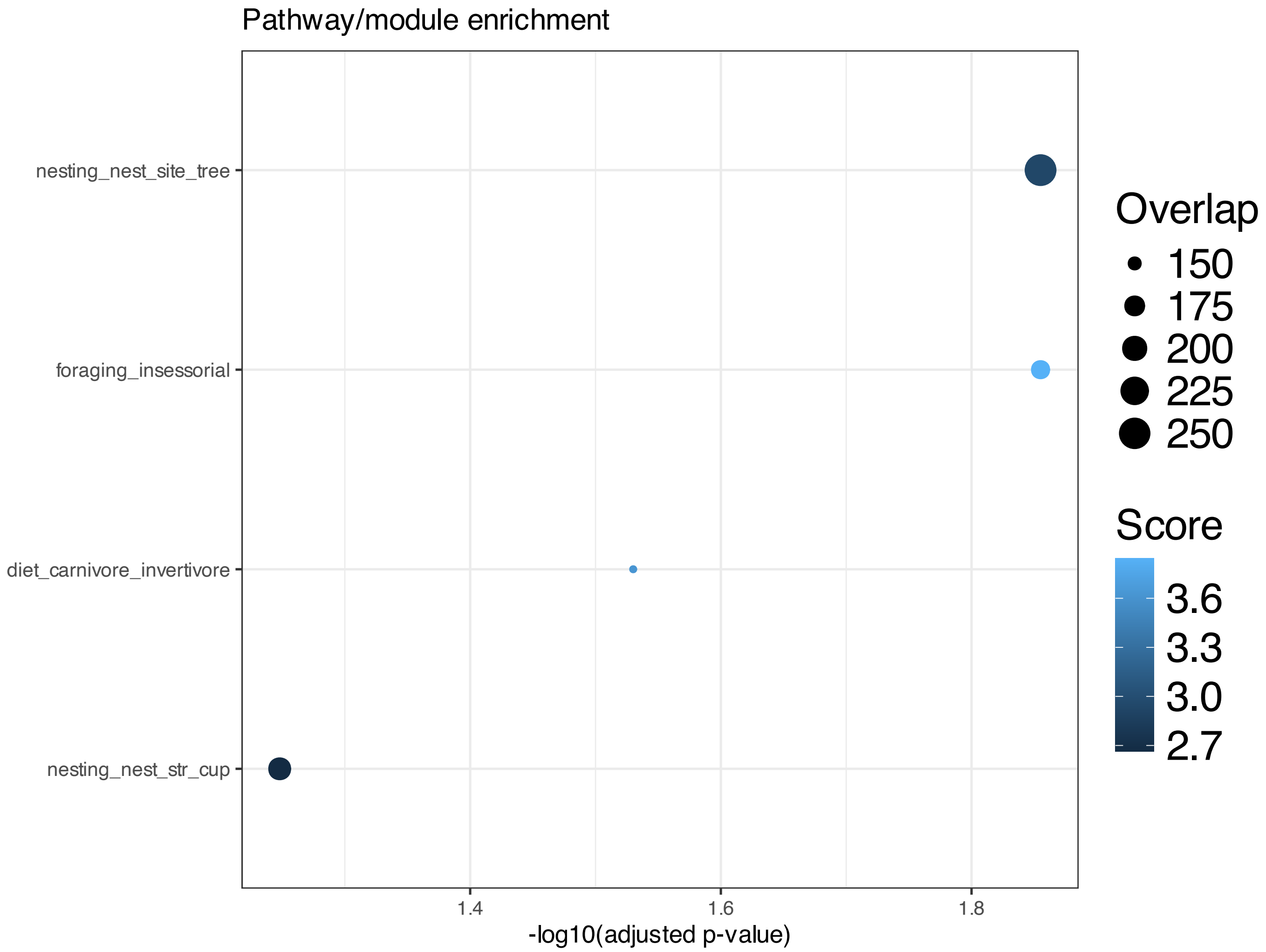


**Supplementary Figure 253.** Enrichment among features inferred to be associated with SSANOVA-SVD component 2; only terms with adjusted P $\leq 0.1$ are shown.

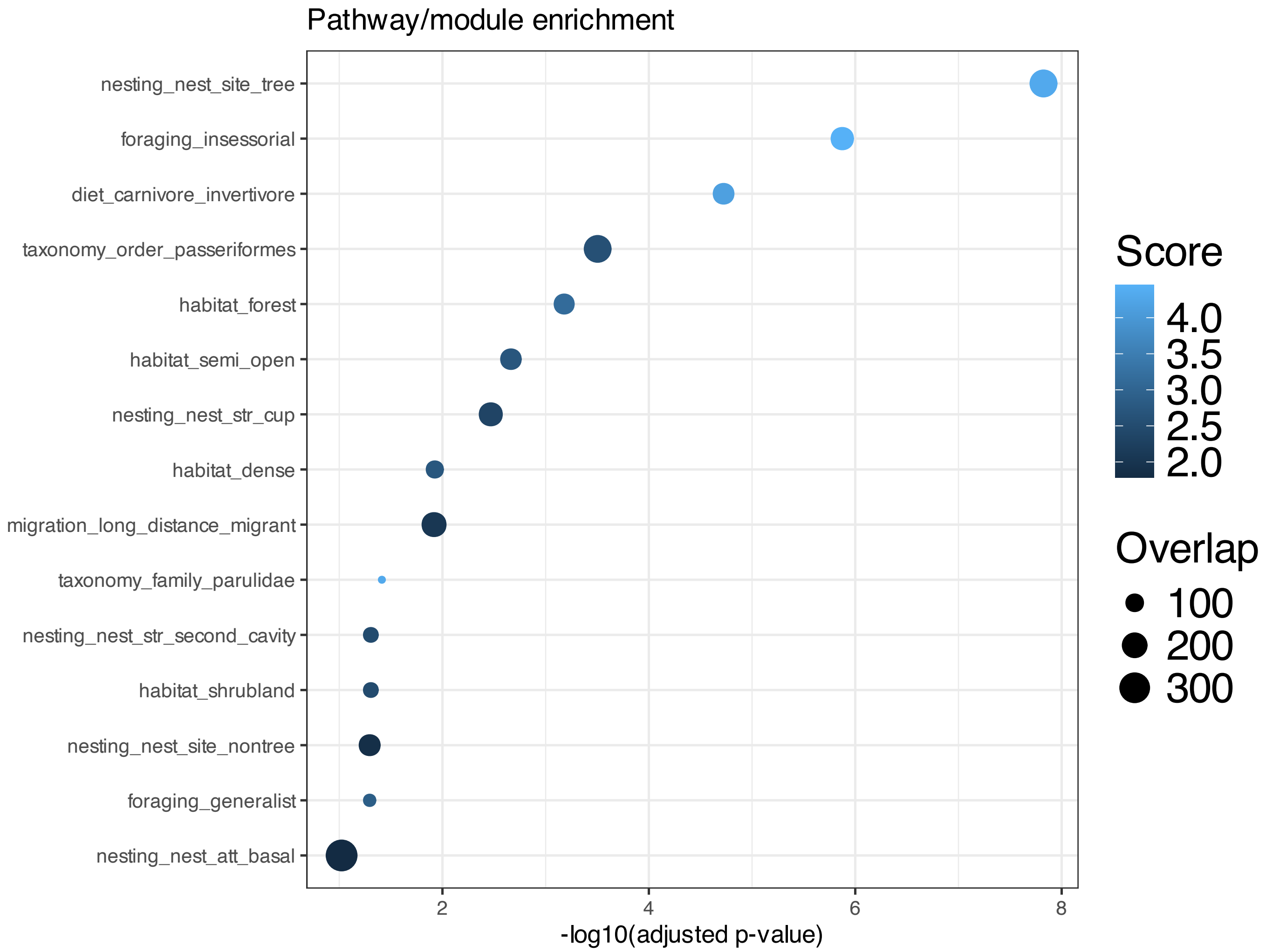


**Supplementary Figure 254.** Enrichment among features inferred to be associated with SSANOVA-SVD component 3; only terms with adjusted P $\leq 0.1$ are shown.

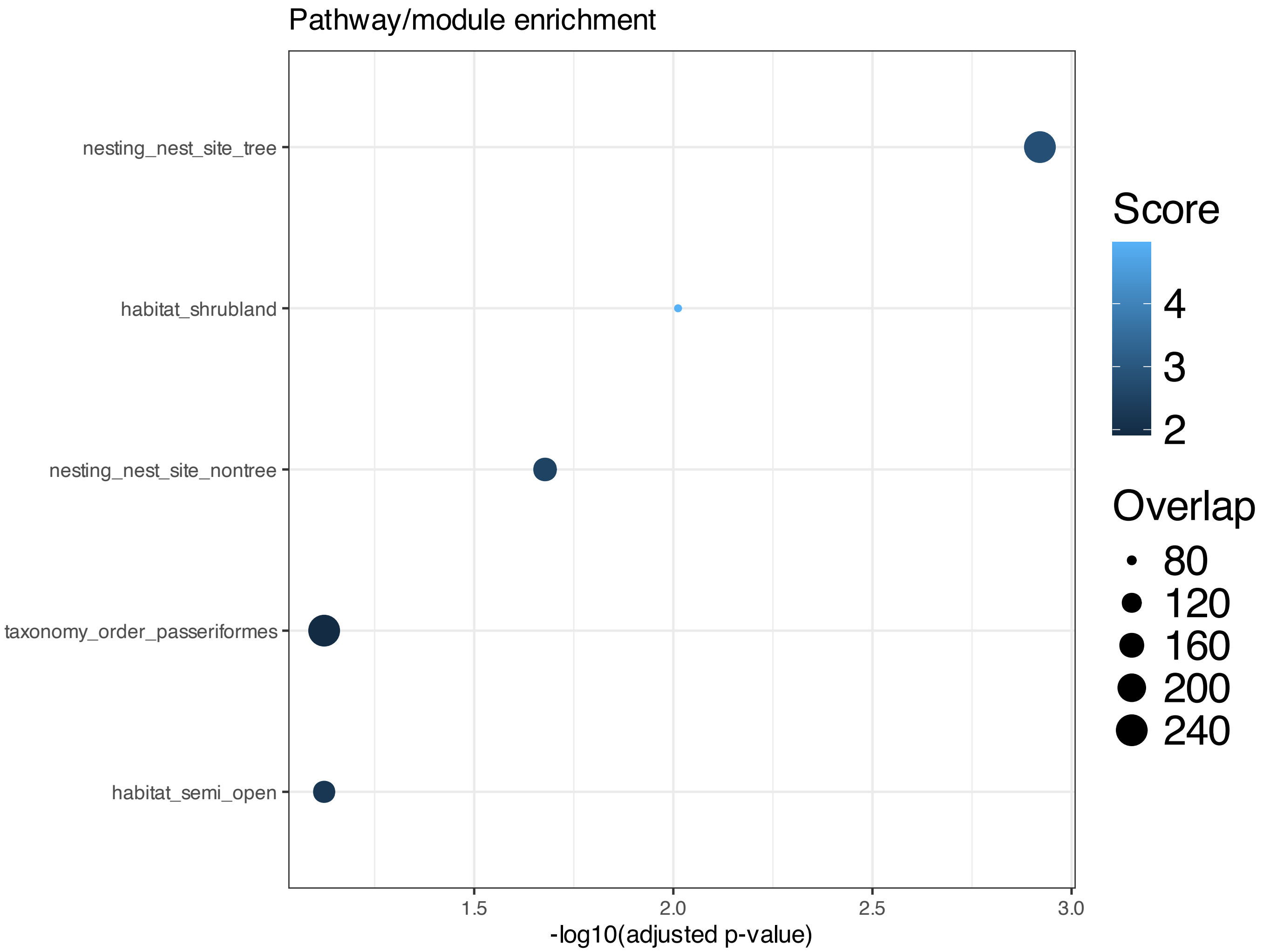


**Supplementary Figure 255.** Enrichment among features inferred to be associated with SSANOVA-SVD component 5; only terms with adjusted P $\leq 0.1$ are shown.

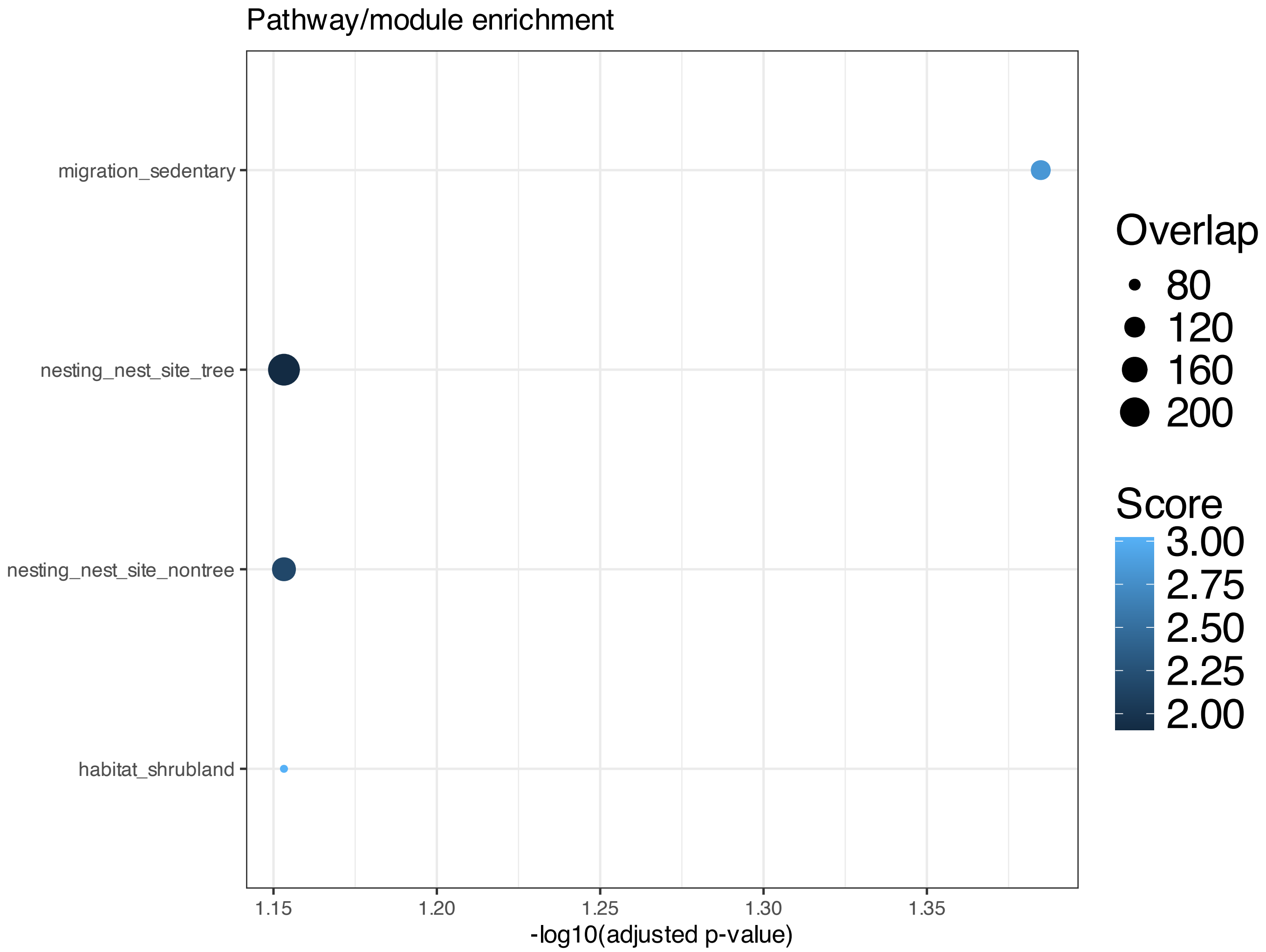


**Supplementary Figure 256.** Enrichment among features inferred to be associated with SSANOVA-SVD component 6; only terms with adjusted P $\leq 0.1$ are shown.

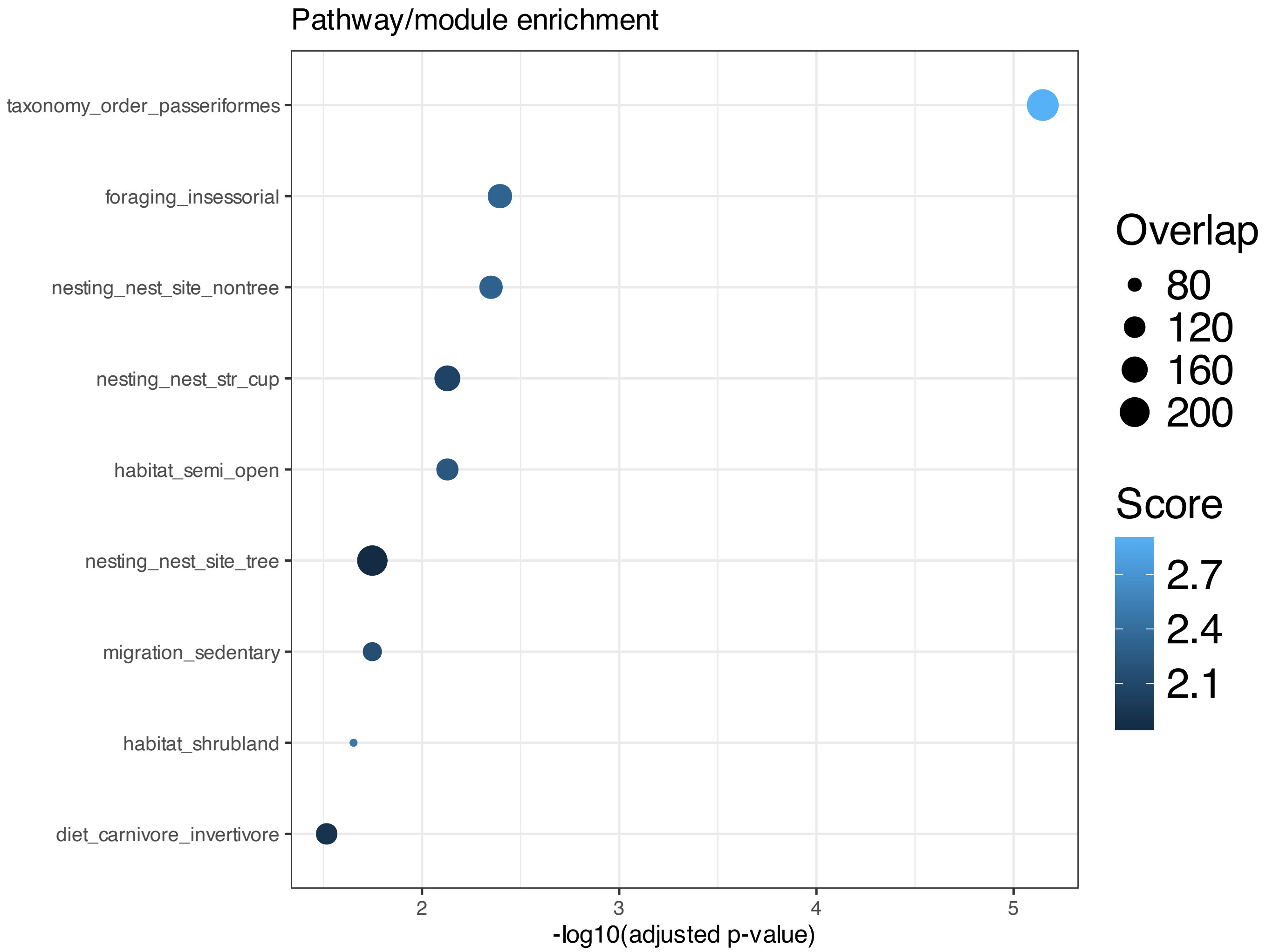


**Supplementary Figure 257.** Enrichment among features inferred to be associated with SSANOVA-SVD component 7; only terms with adjusted P $\leq 0.1$ are shown.

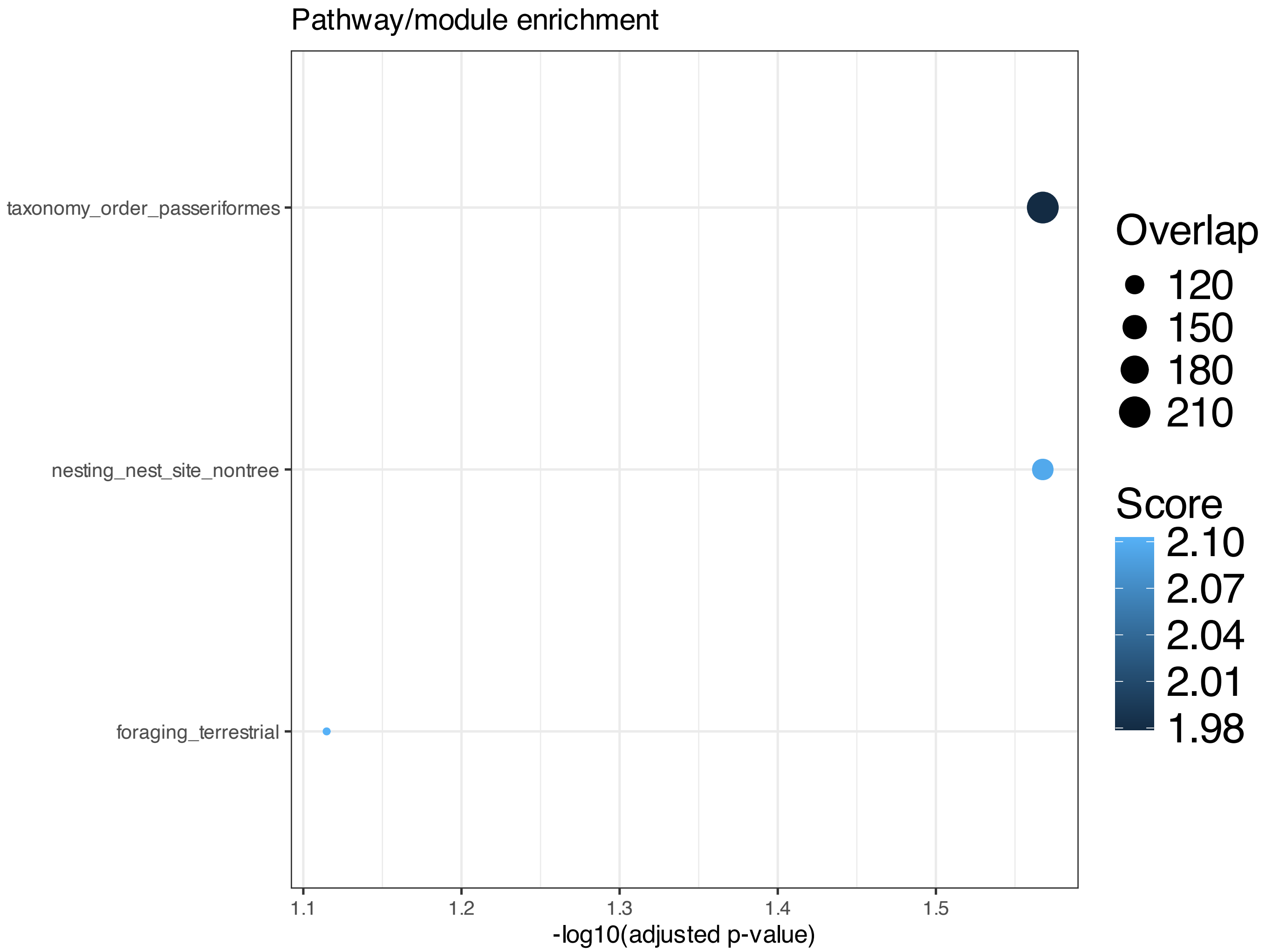


**Supplementary Figure 258.** Enrichment among features inferred to be associated with SSANOVA-SVD component 8; only terms with adjusted P $\leq 0.1$ are shown.

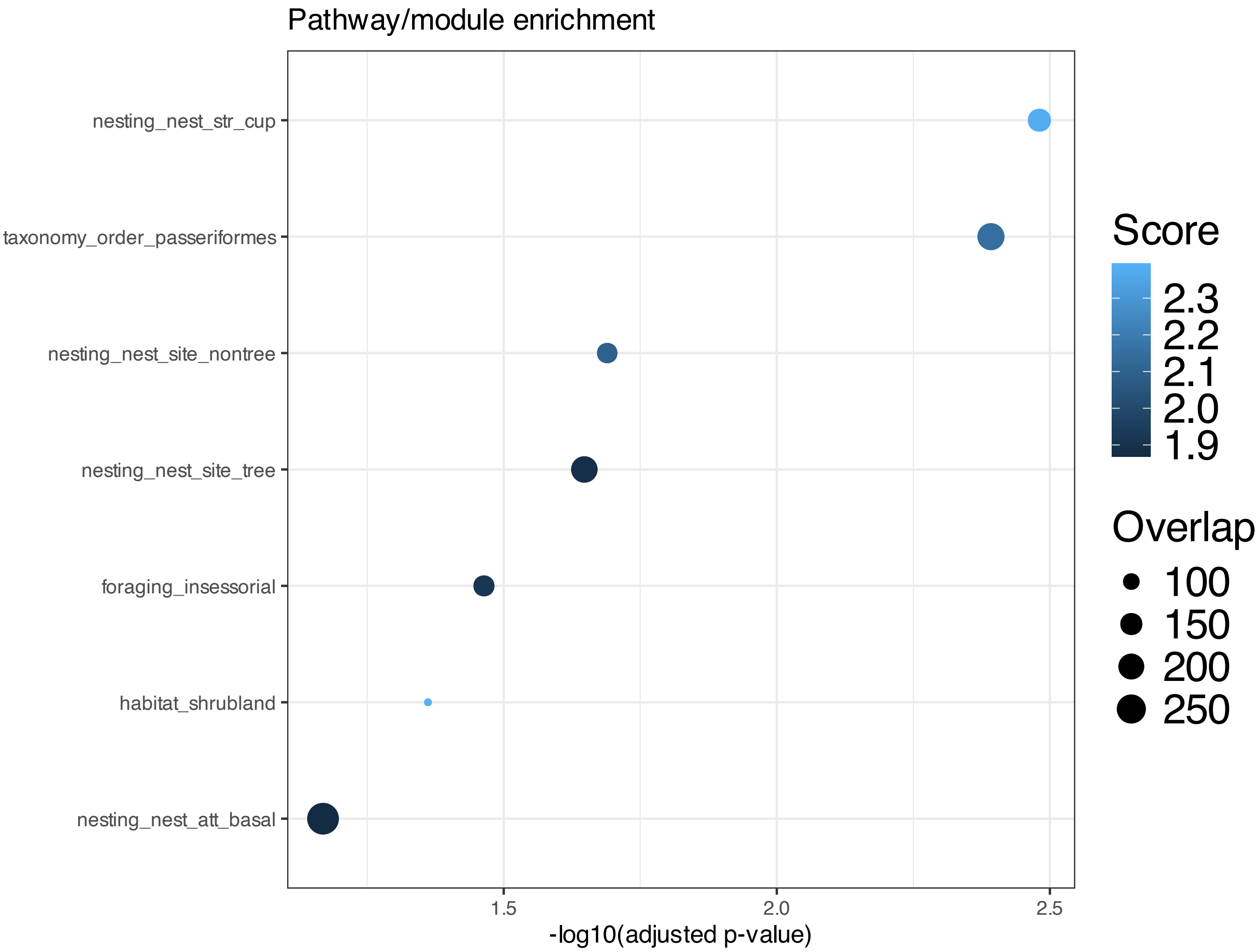


**Supplementary Figure 259.** Enrichment among features inferred to be associated with SSANOVA-SVD component 10; only terms with adjusted P $\leq 0.1$ are shown.

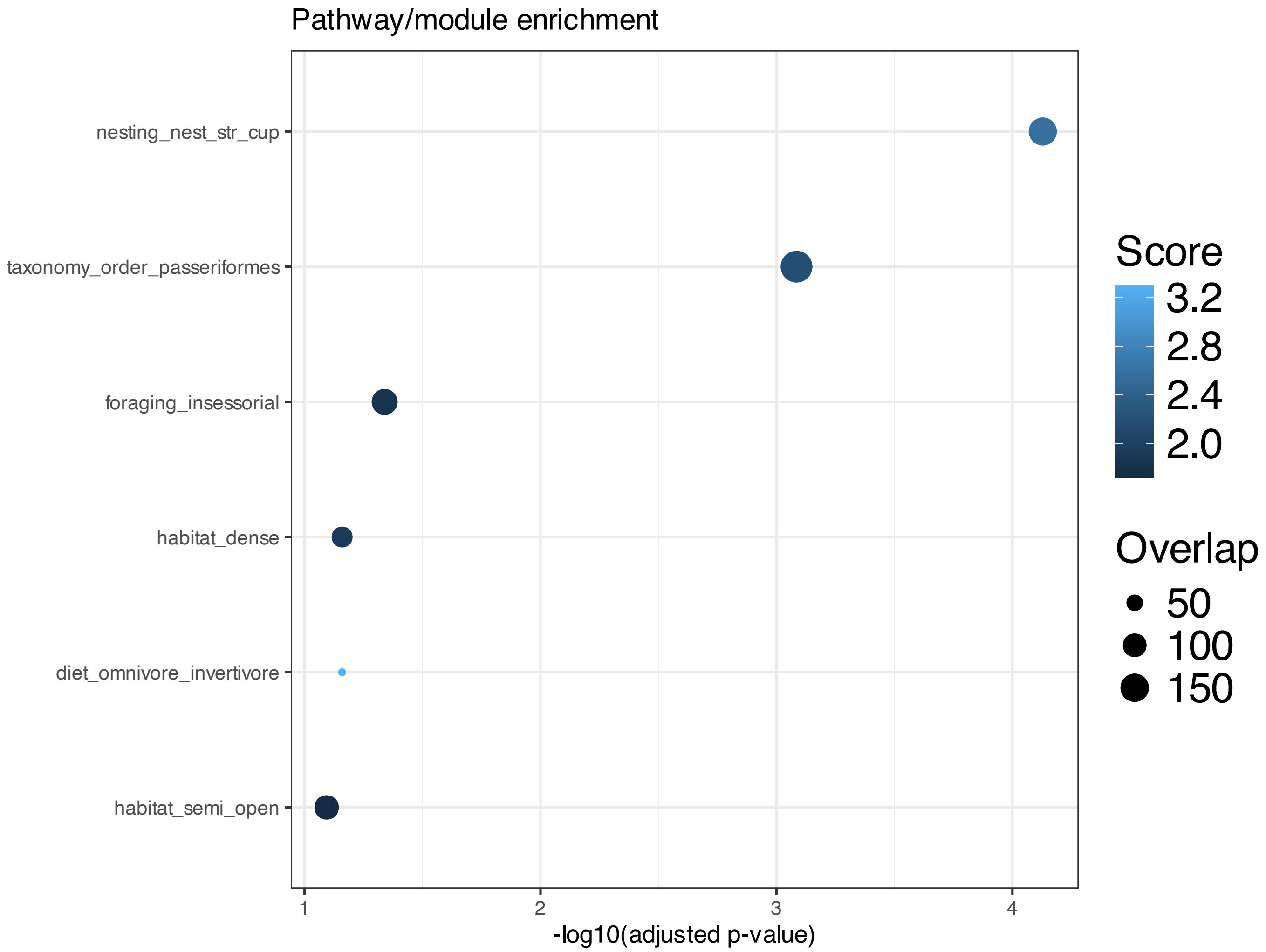


**Supplementary Figure 260.** Enrichment among features inferred to be associated with SSANOVA-SVD component 11; only terms with adjusted P $\leq 0.1$ are shown.

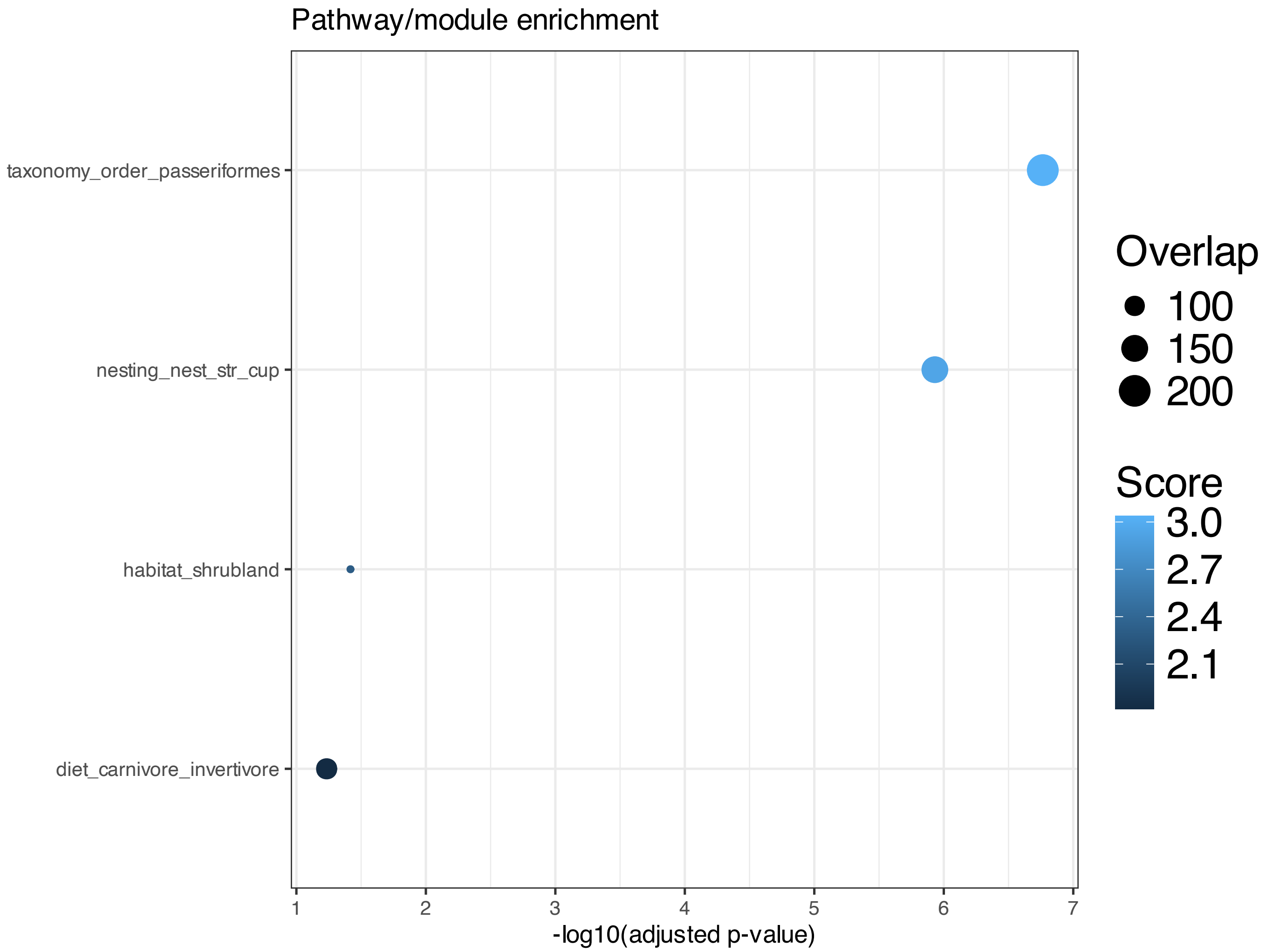


**Supplementary Figure 261.** Enrichment among features inferred to be associated with SSANOVA-SVD component 12; only terms with adjusted P $\leq 0.1$ are shown.

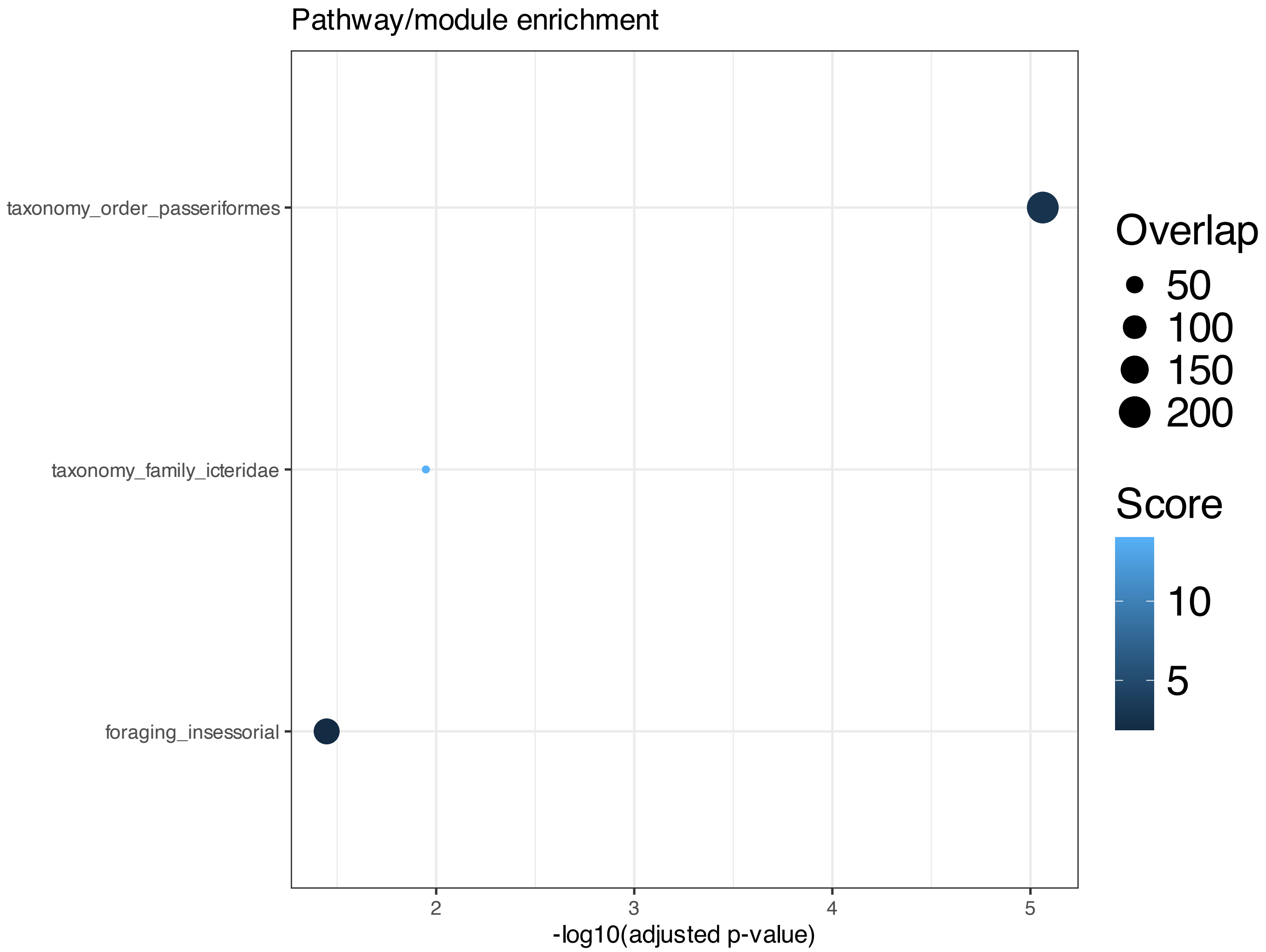


**Supplementary Figure 262.** Enrichment among features inferred to be associated with SSANOVA-SVD component 13; only terms with adjusted P $\leq 0.1$ are shown.

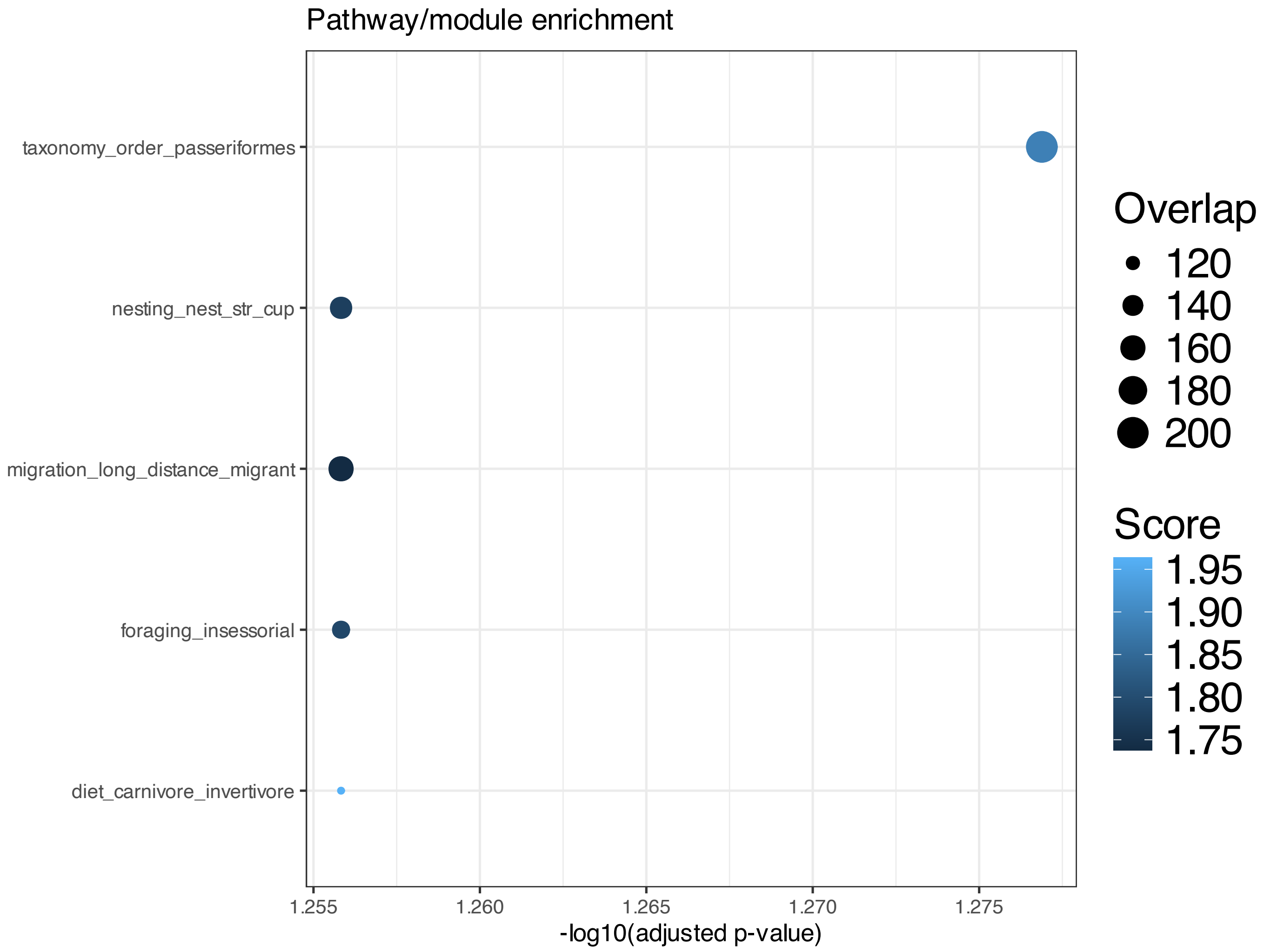


**Supplementary Figure 263.** Enrichment among features inferred to be associated with SSANOVA-SVD component 14; only terms with adjusted P $\leq 0.1$ are shown.

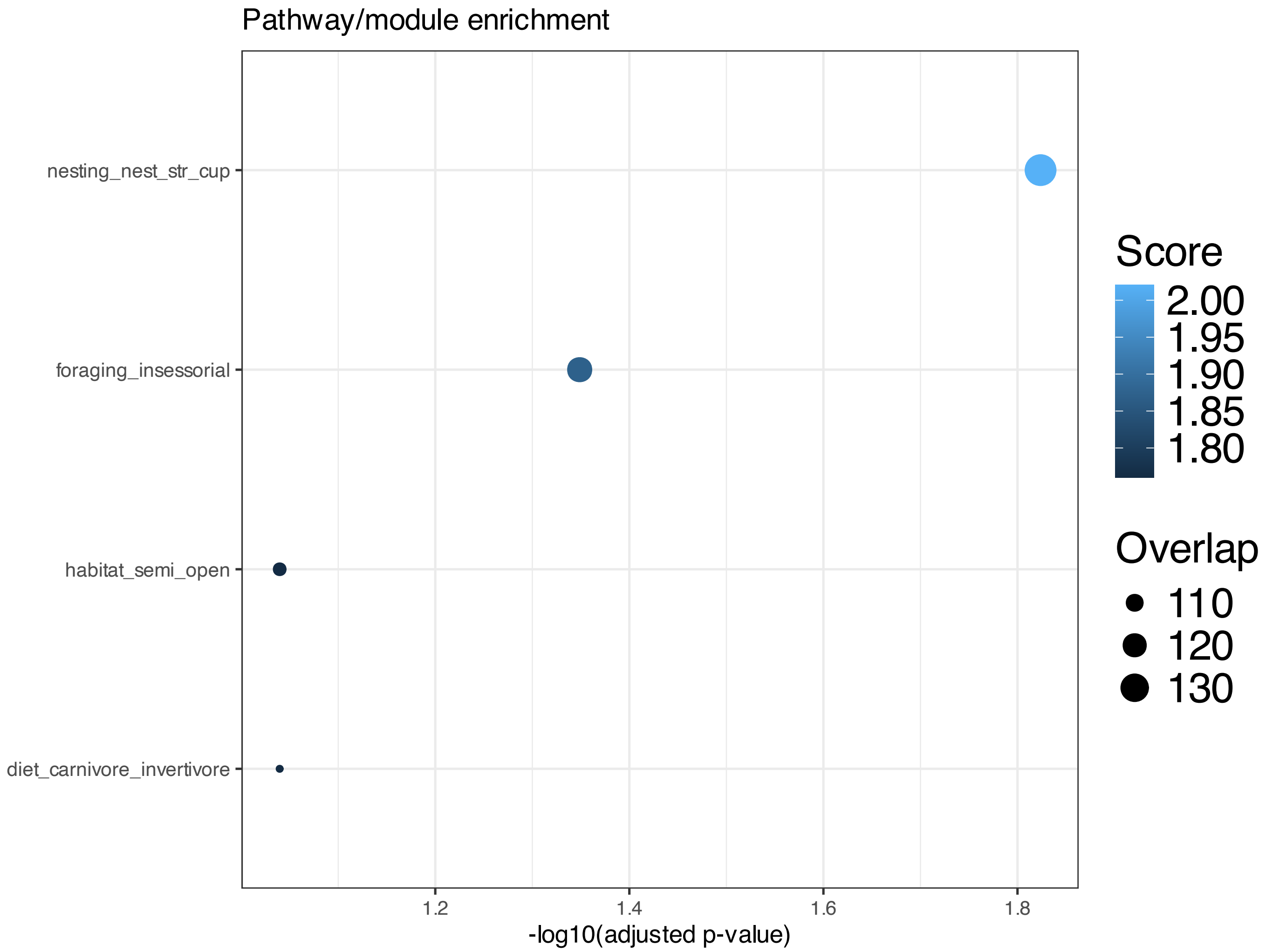


**Supplementary Figure 264.** Enrichment among features inferred to be associated with SSANOVA-SVD component 15; only terms with adjusted P $\leq$ 0.1 are shown.

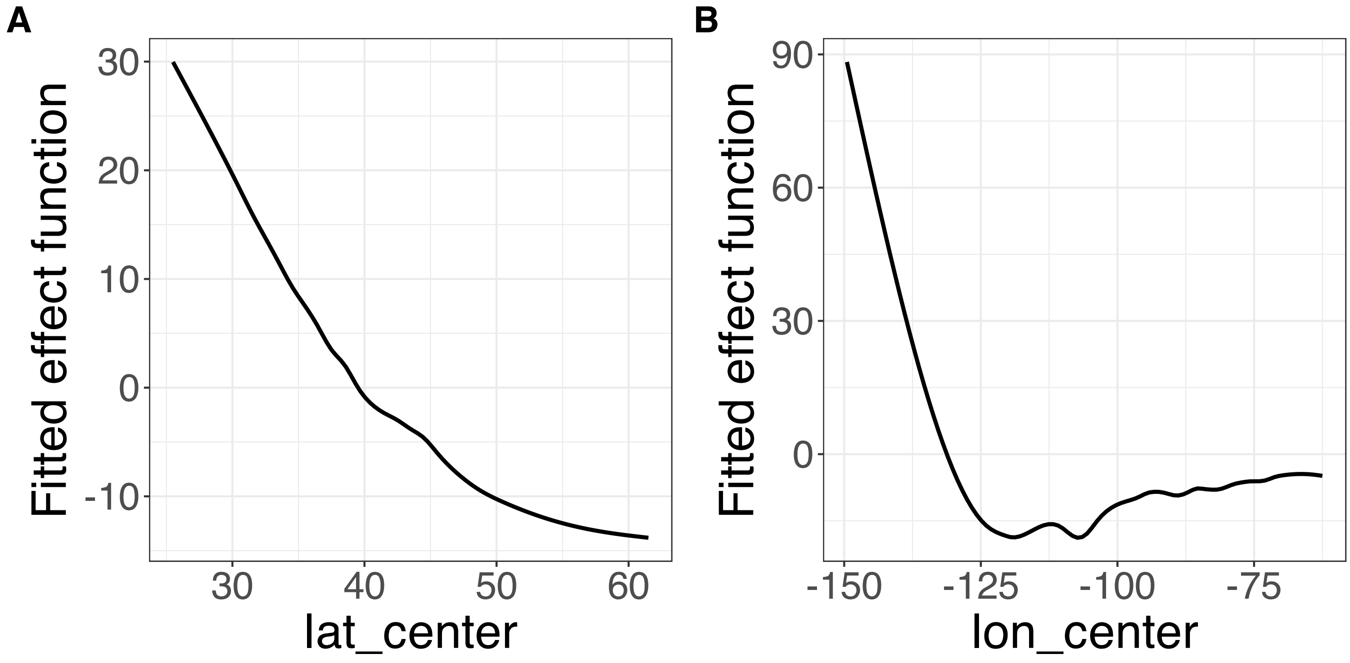


**Supplementary Figure 265.** Decomposition of reconstructed SSANOVA-SVD component function 1. (A) lat_center. (B) lon_center.

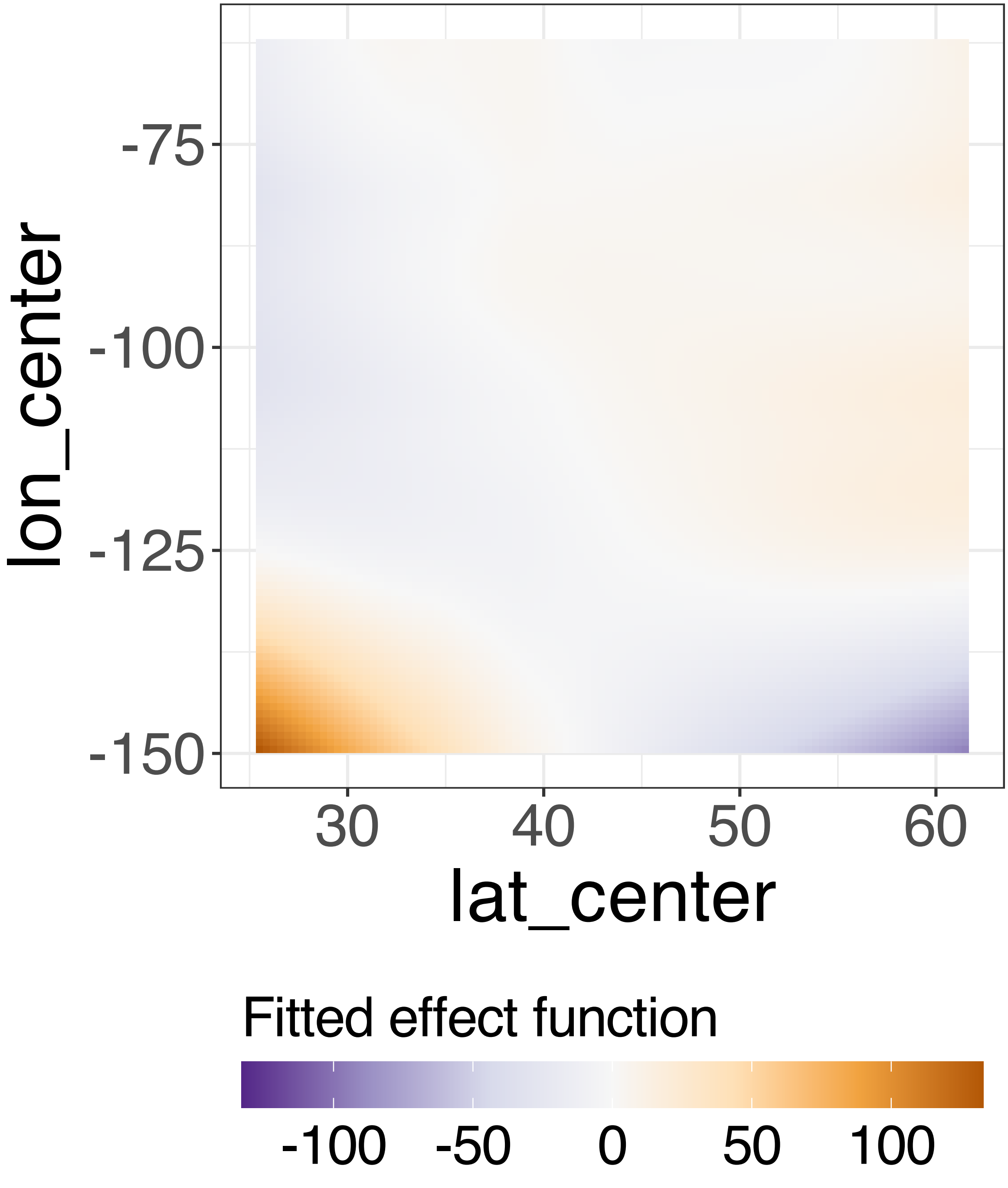


**Supplementary Figure 266.** Decomposition of reconstructed SSANOVA-SVD component function 1.

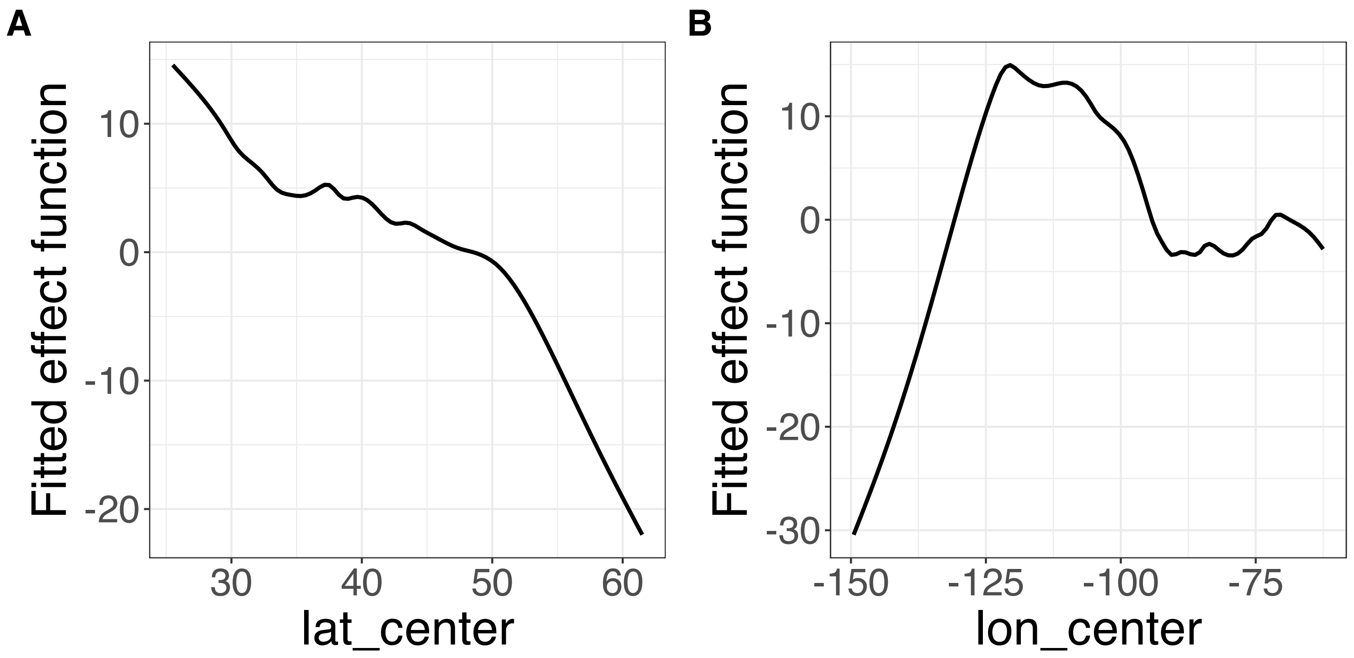


**Supplementary Figure 267.** Decomposition of reconstructed SSANOVA-SVD component function 2. (A) lat_center. (B) lon_center.

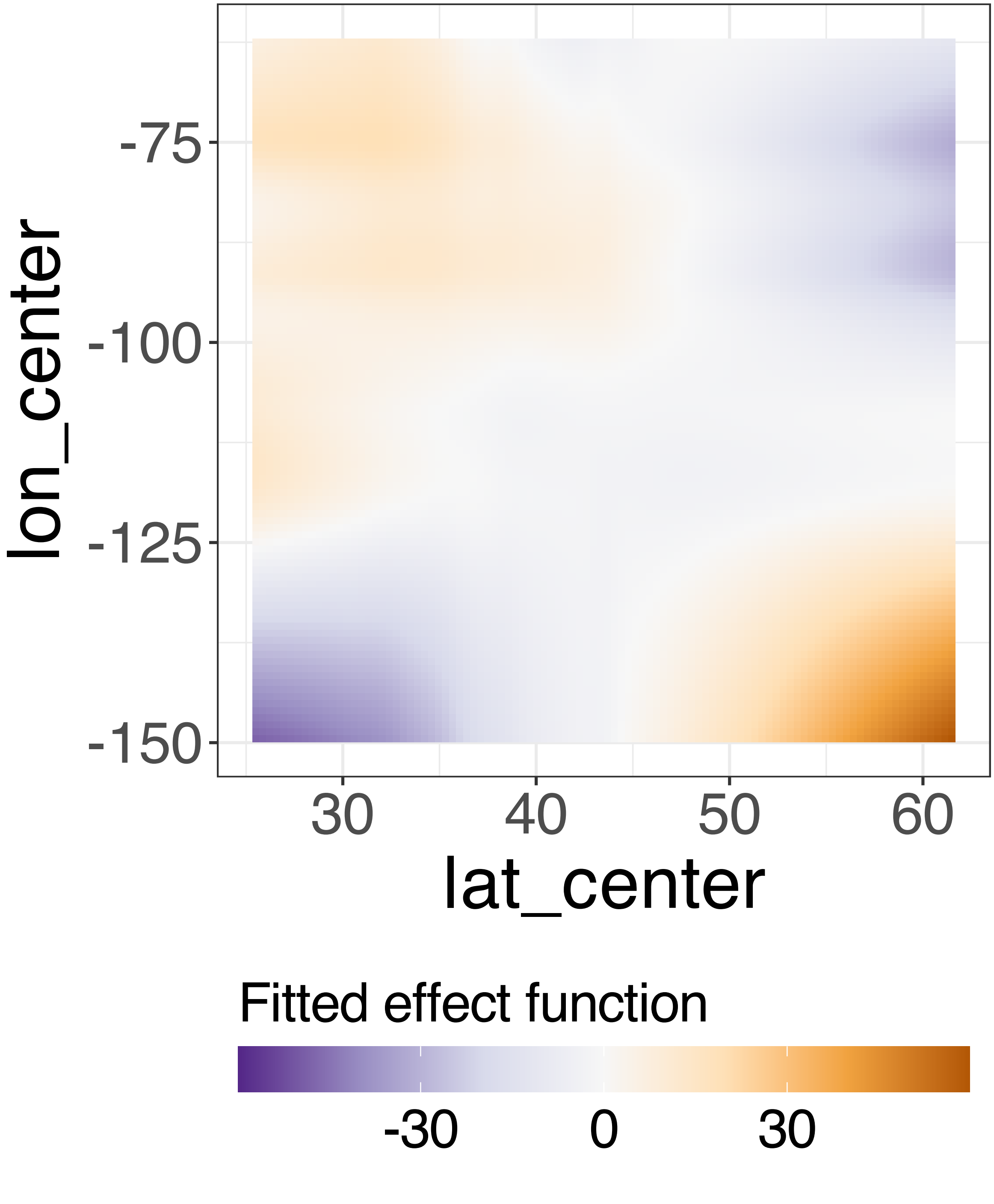


**Supplementary Figure 268.** Decomposition of reconstructed SSANOVA-SVD component function 2.

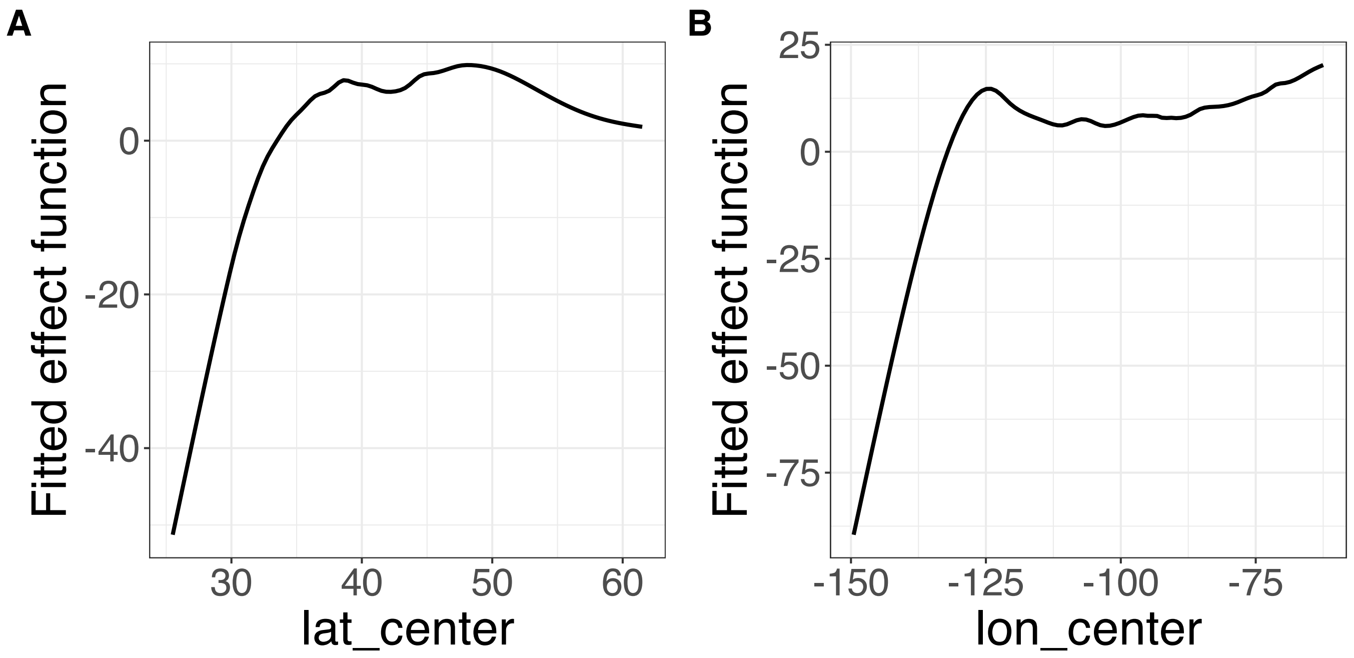


**Supplementary Figure 269.** Decomposition of reconstructed SSANOVA-SVD component function 3. (A) lat_center. (B) lon_center.

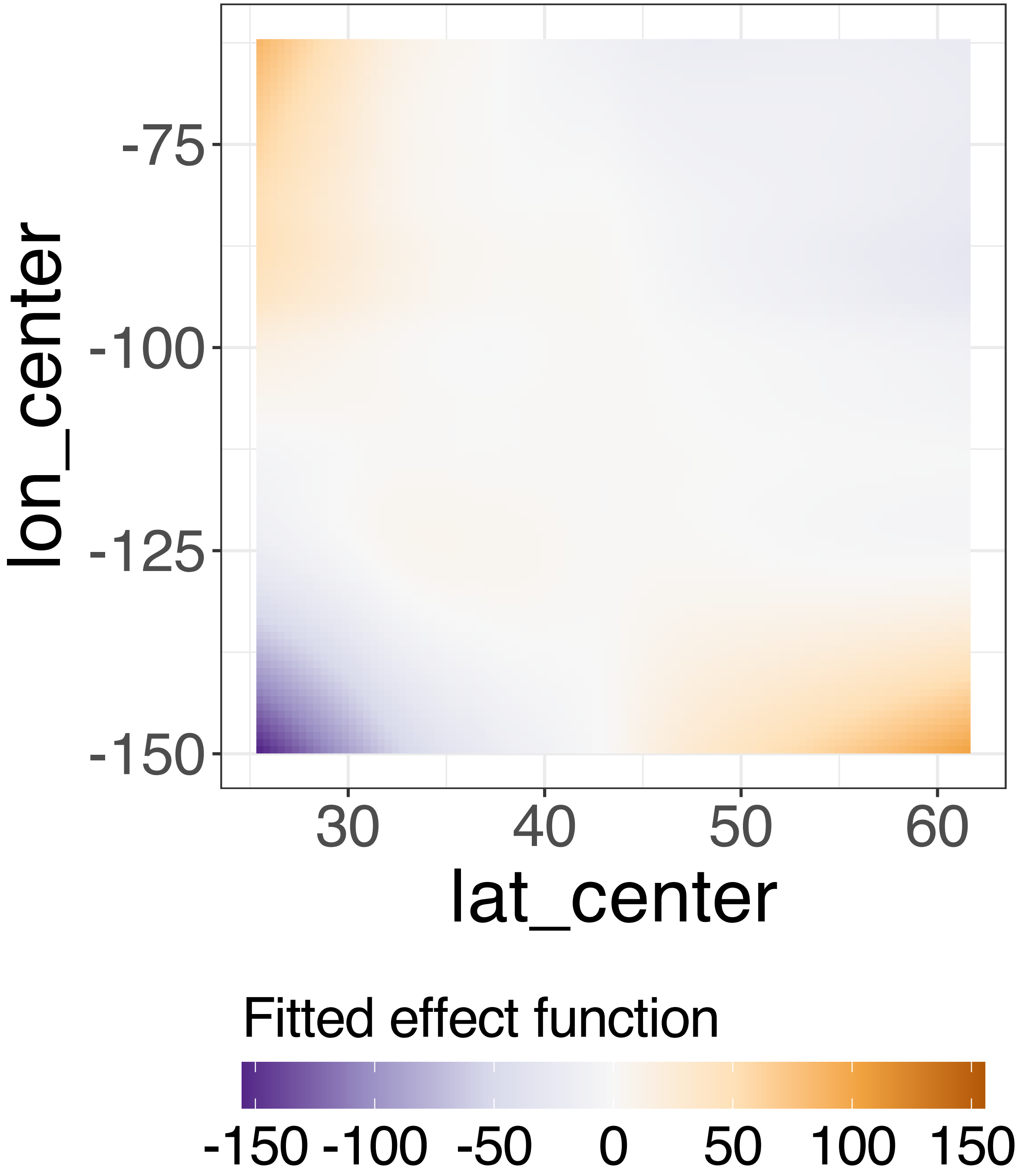


**Supplementary Figure 270.** Decomposition of reconstructed SSANOVA-SVD component function 3.

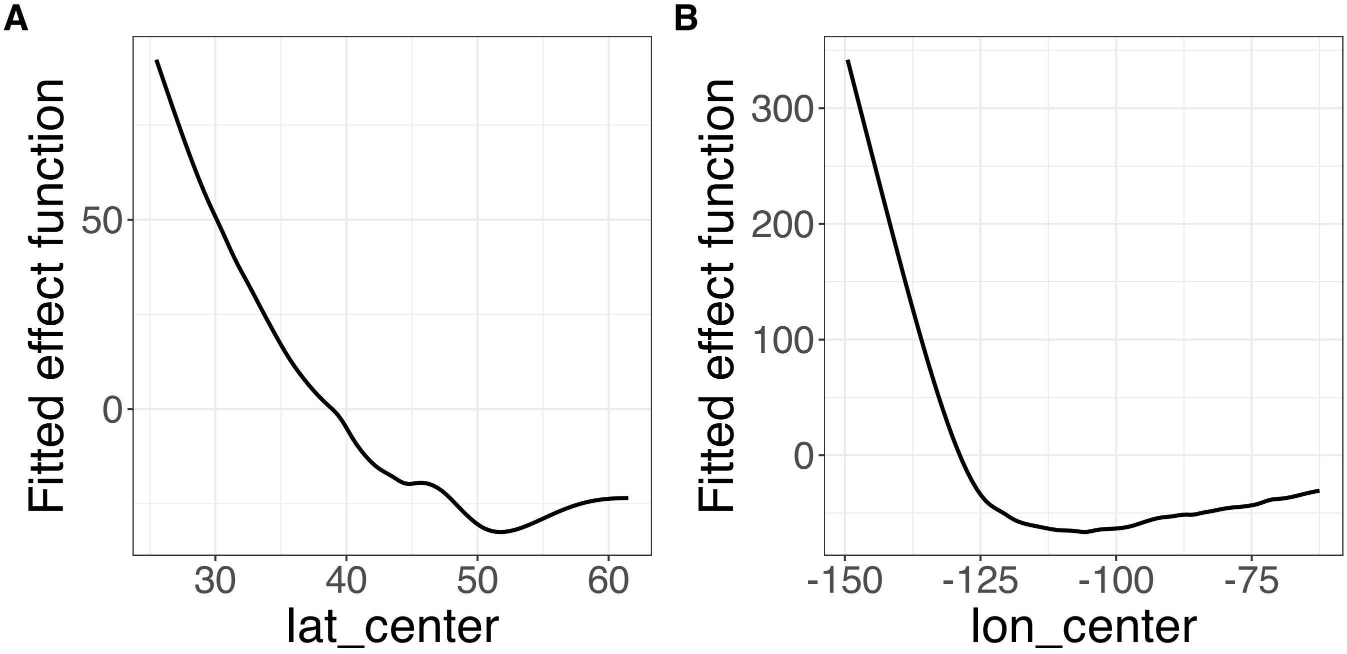


**Supplementary Figure 271.** Decomposition of reconstructed SSANOVA-SVD component function 4. (A) lat_center. (B) lon_center.

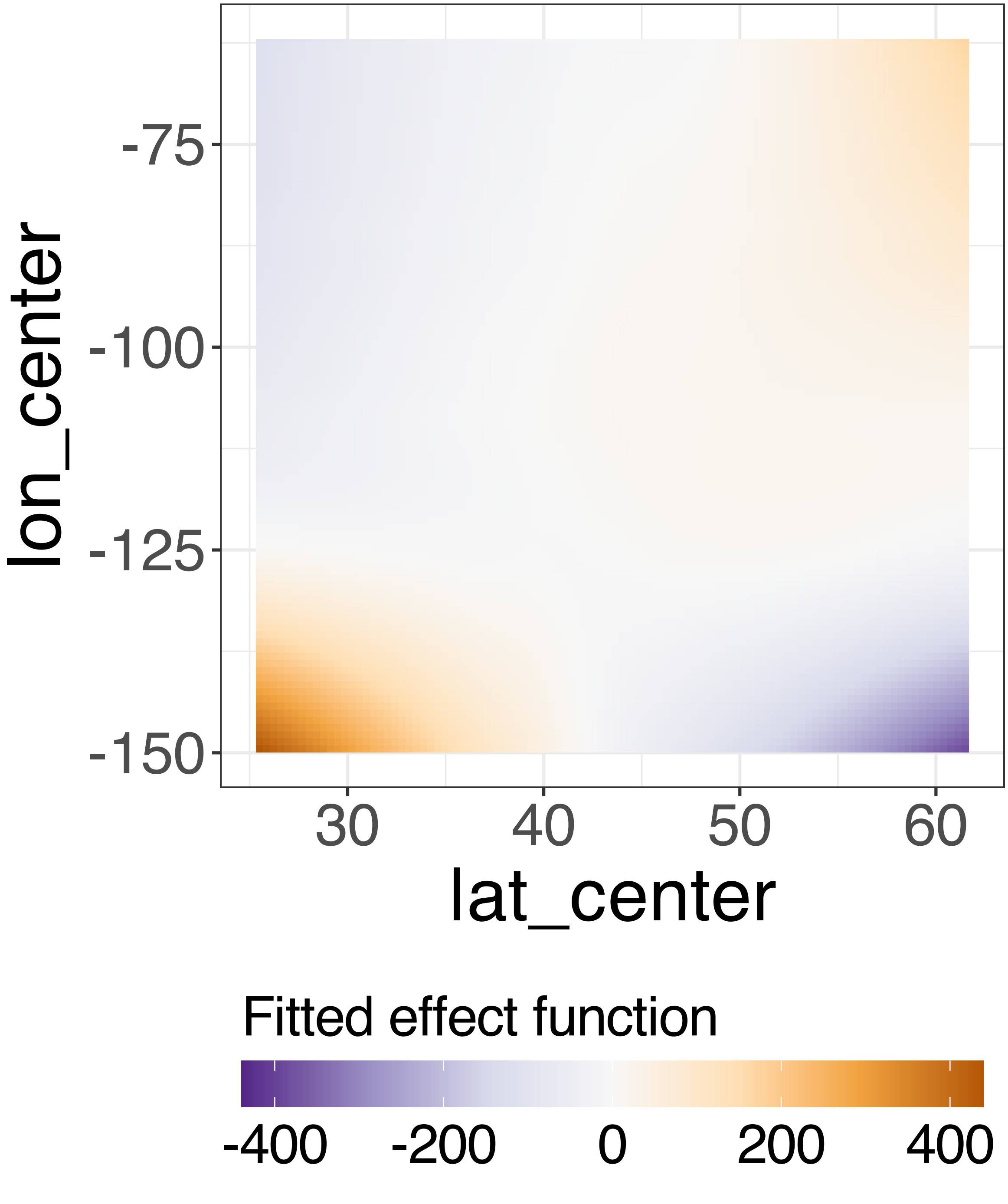


**Supplementary Figure 272.** Decomposition of reconstructed SSANOVA-SVD component function 4.

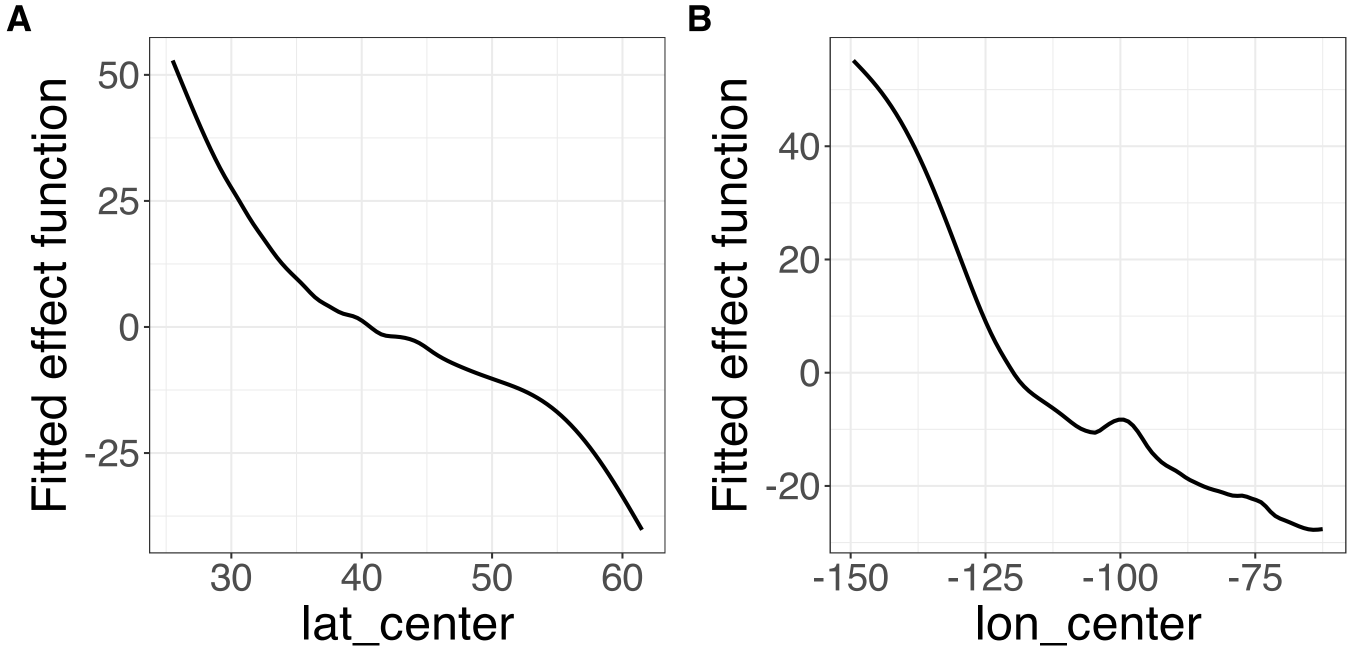


**Supplementary Figure 273.** Decomposition of reconstructed SSANOVA-SVD component function 5. (A) lat_center. (B) lon_center.

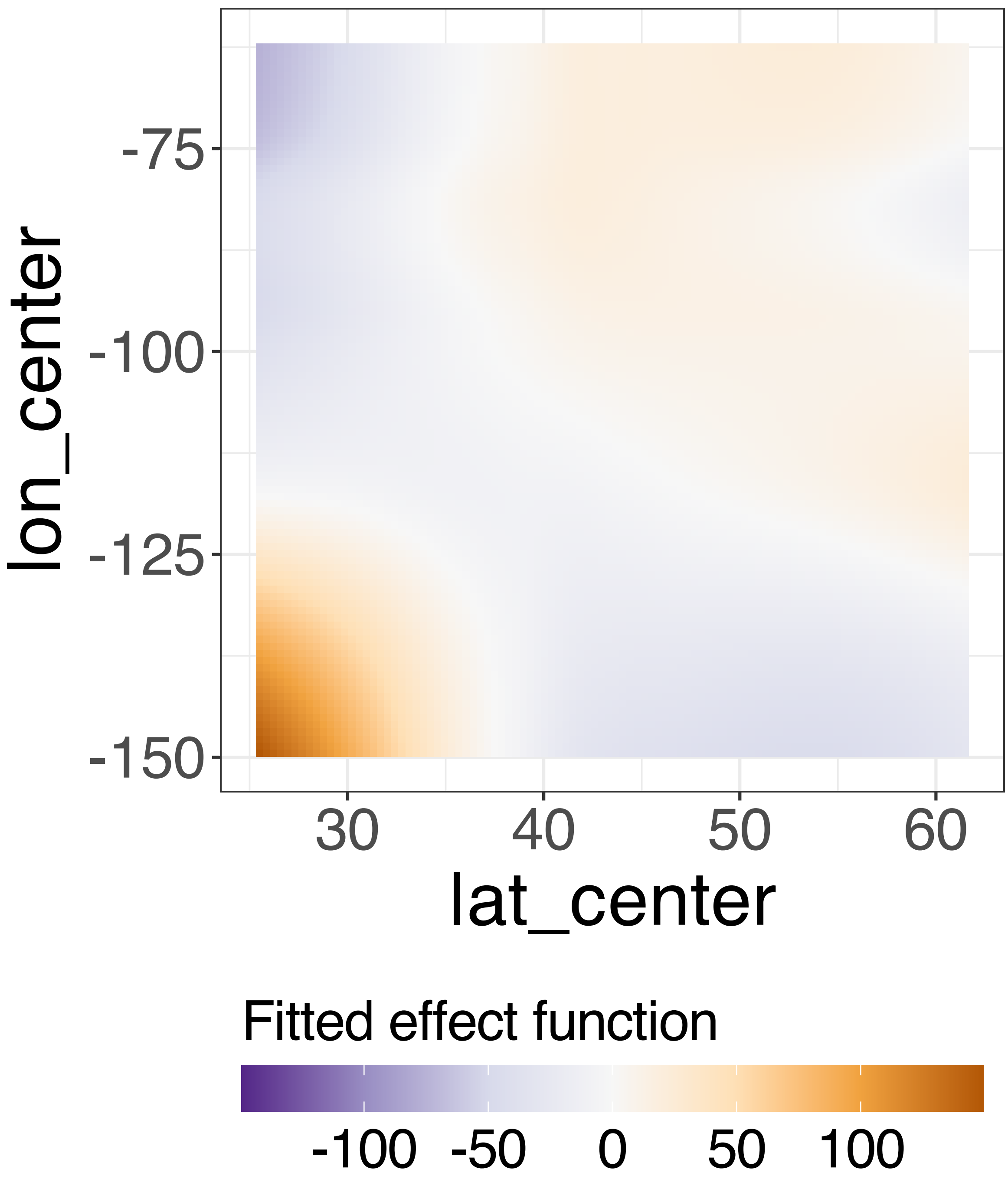


**Supplementary Figure 274.** Decomposition of reconstructed SSANOVA-SVD component function 5.

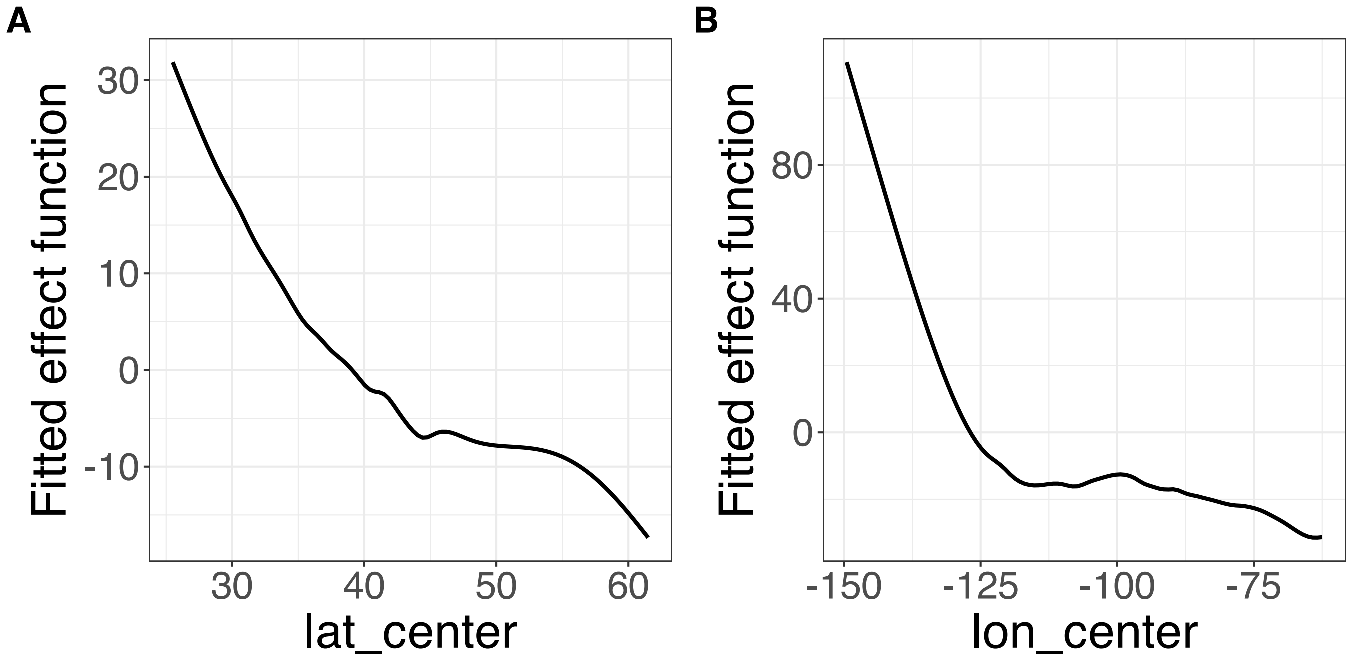


**Supplementary Figure 275.** Decomposition of reconstructed SSANOVA-SVD component function 6. (A) lat_center. (B) lon_center.

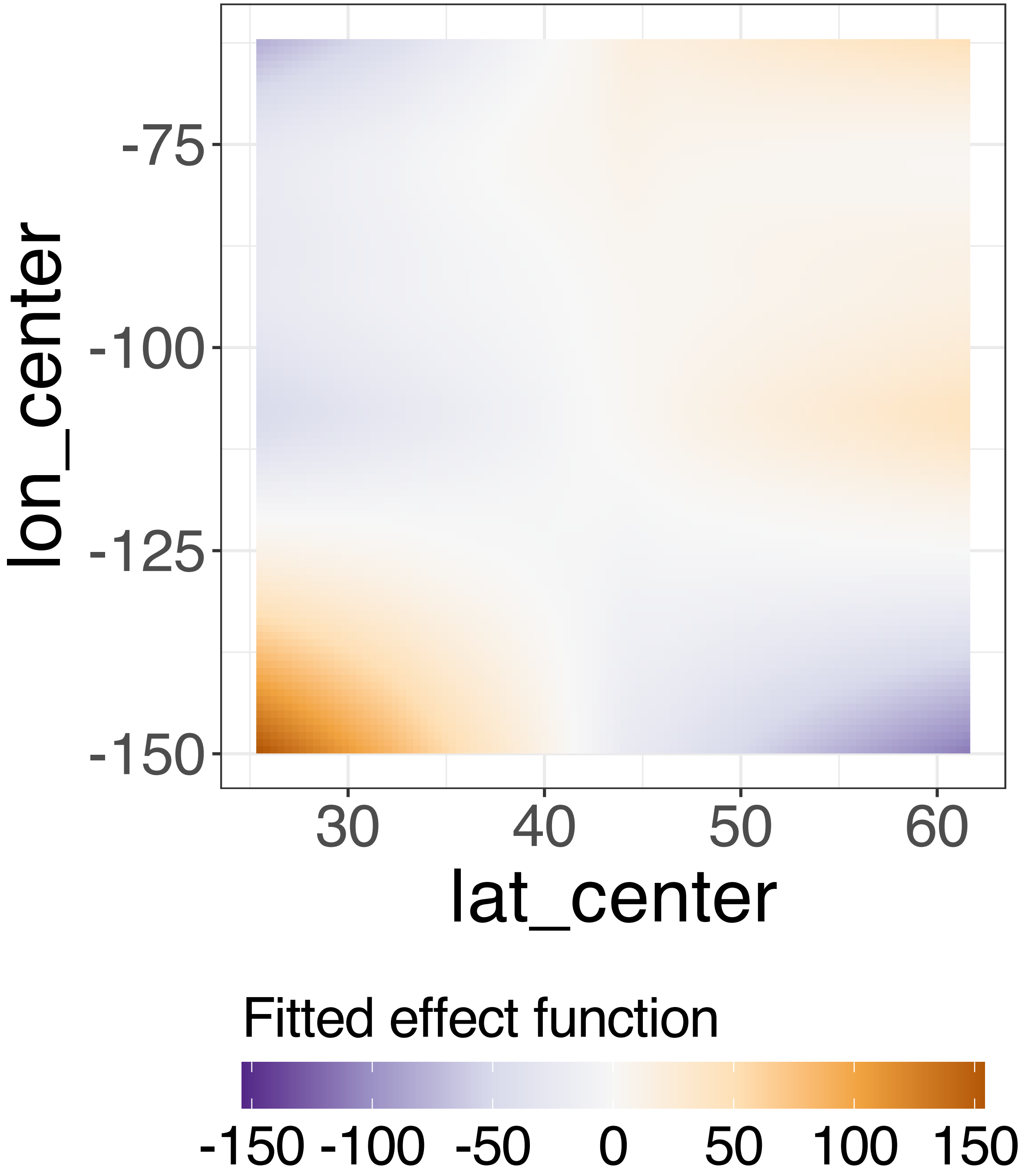


**Supplementary Figure 276.** Decomposition of reconstructed SSANOVA-SVD component function 6.

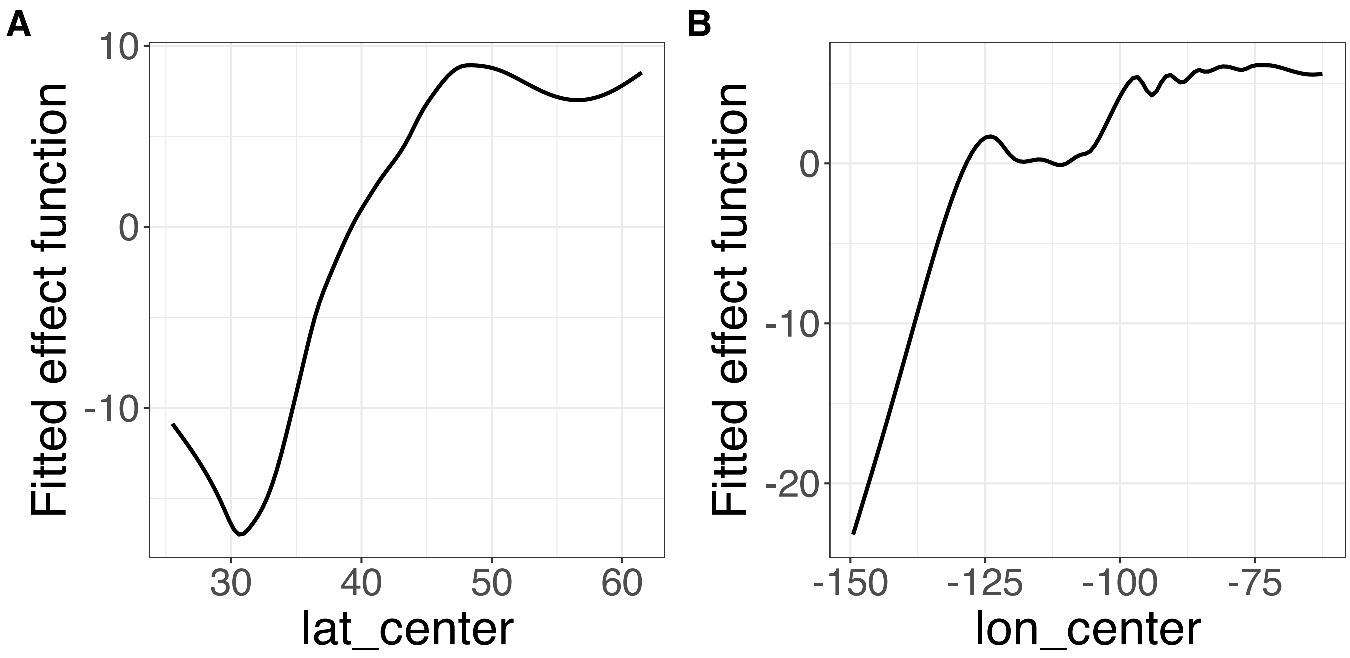


**Supplementary Figure 277.** Decomposition of reconstructed SSANOVA-SVD component function 7. (A) lat_center. (B) lon_center.

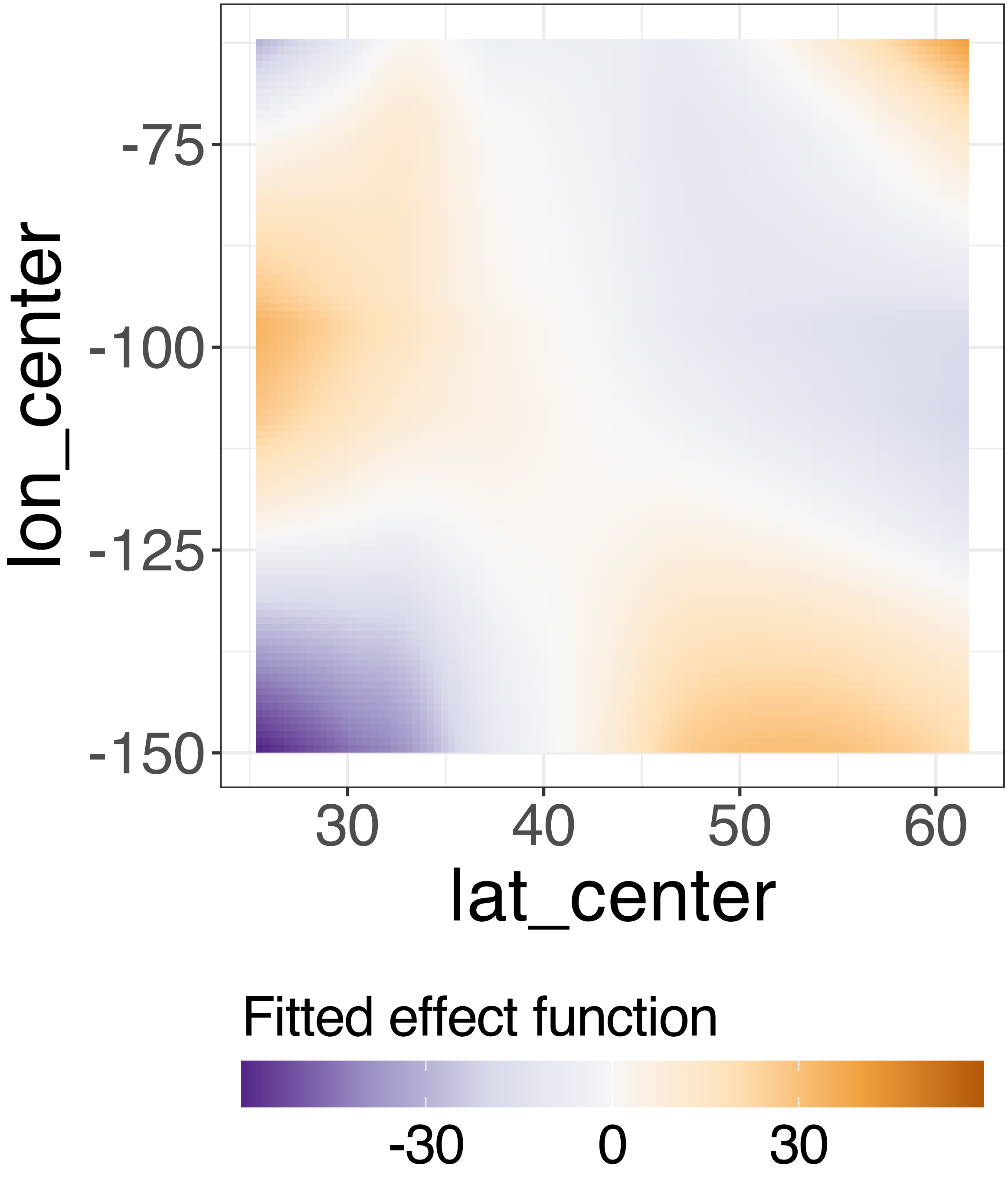


**Supplementary Figure 278.** Decomposition of reconstructed SSANOVA-SVD component function 7.

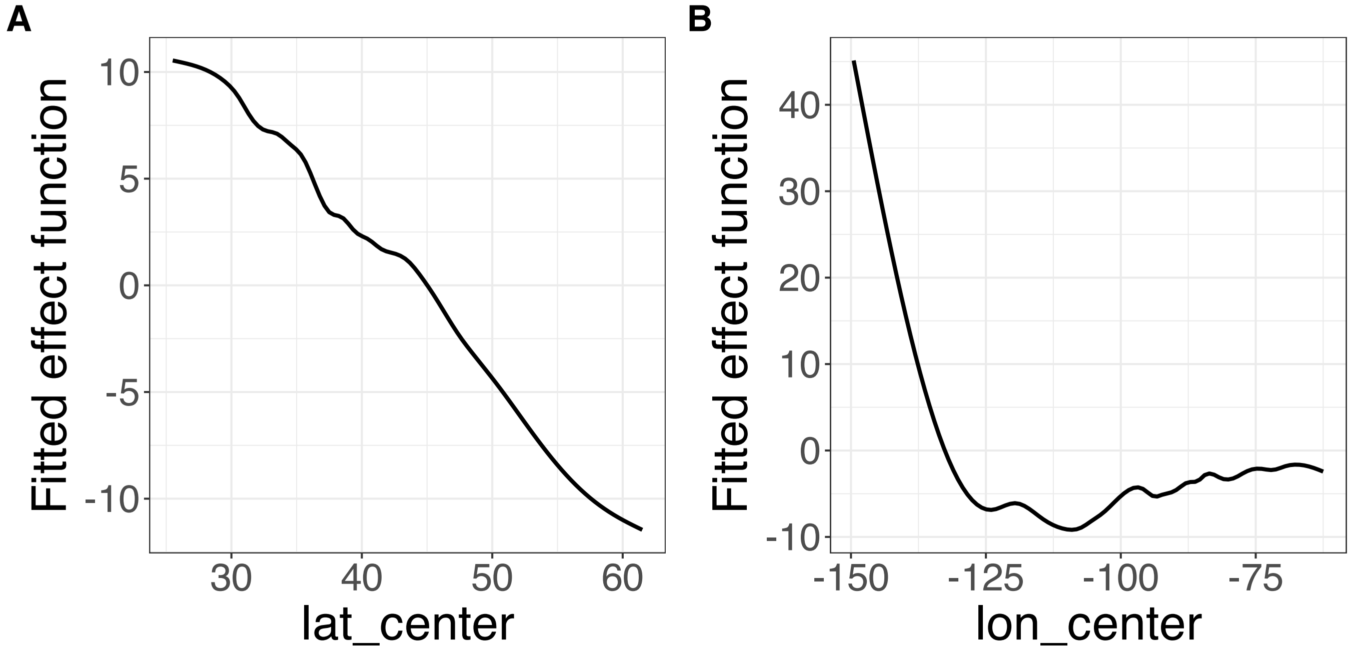


**Supplementary Figure 279.** Decomposition of reconstructed SSANOVA-SVD component function 8. (A) lat_center. (B) lon_center.

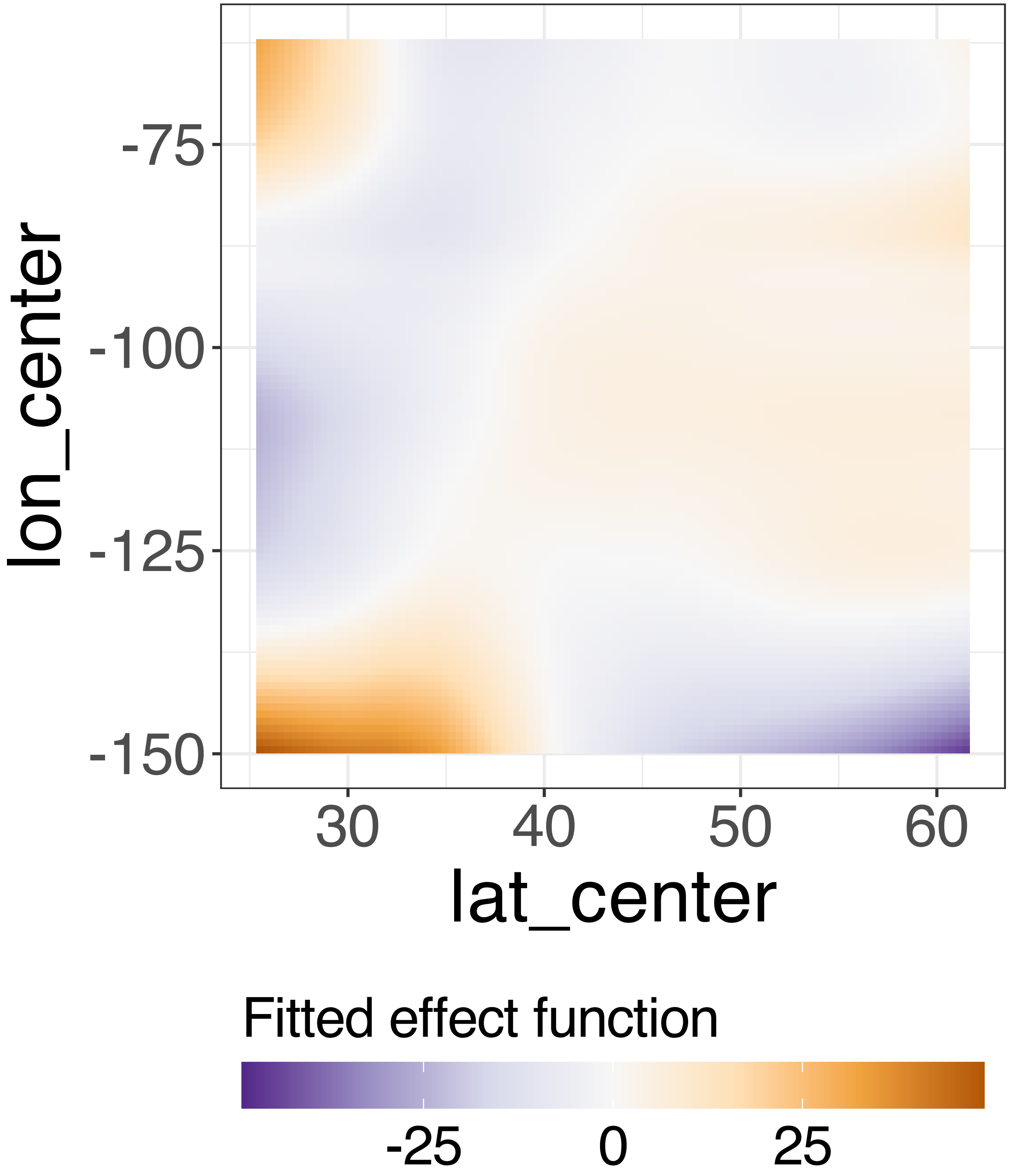


**Supplementary Figure 280.** Decomposition of reconstructed SSANOVA-SVD component function 8.

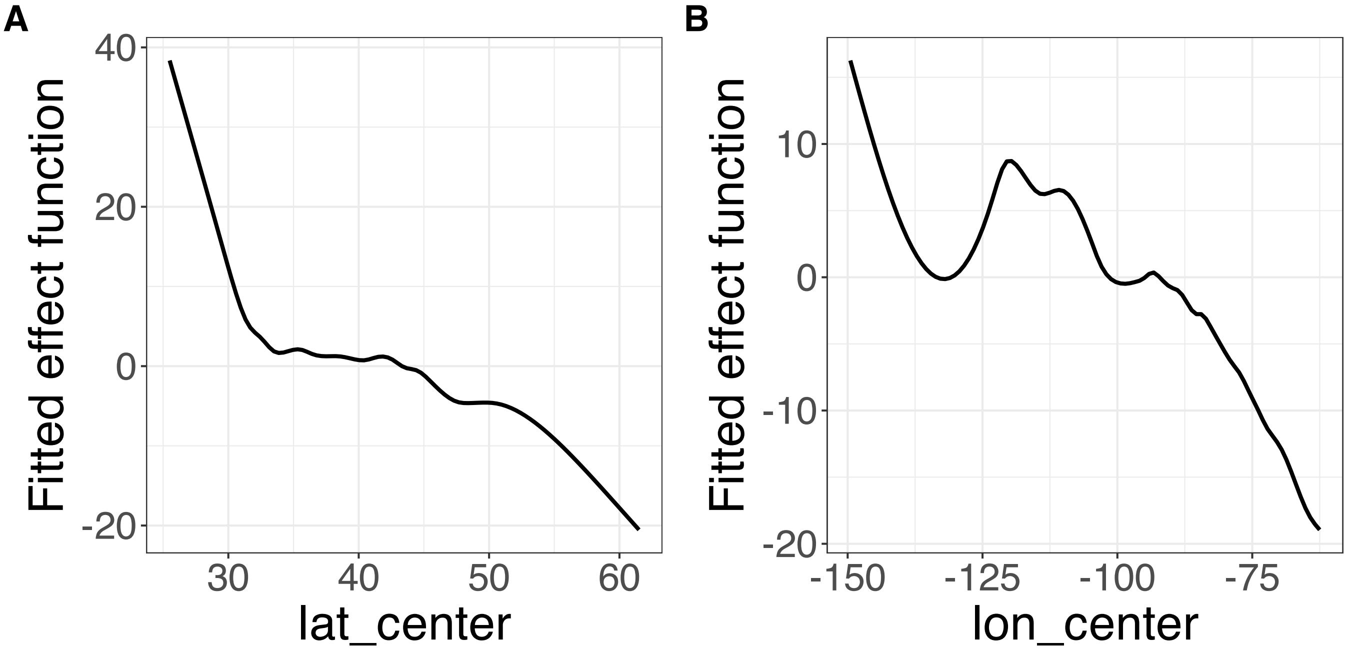


**Supplementary Figure 281.** Decomposition of reconstructed SSANOVA-SVD component function 9. (A) lat_center. (B) lon_center.

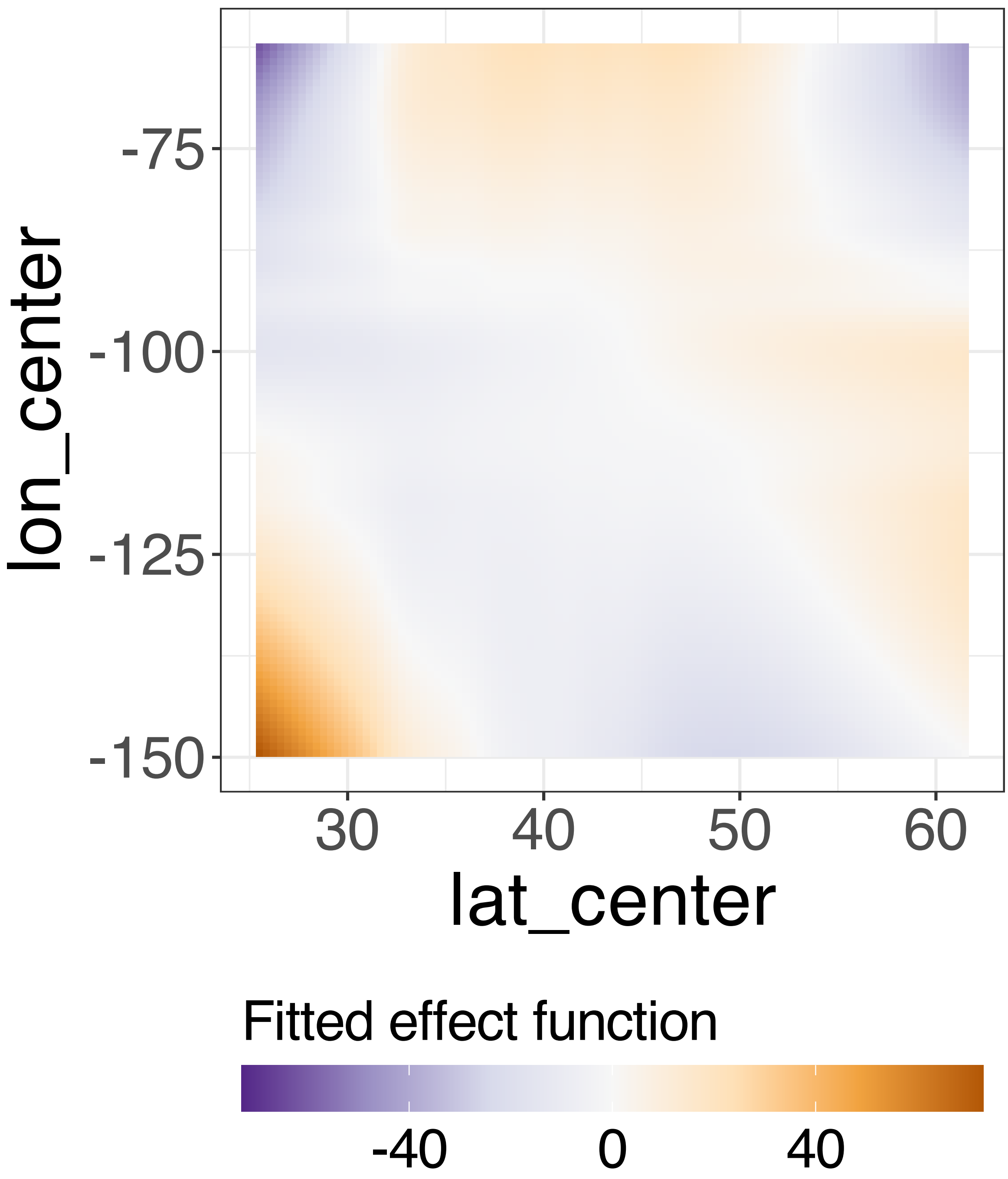


**Supplementary Figure 282.** Decomposition of reconstructed SSANOVA-SVD component function 9.

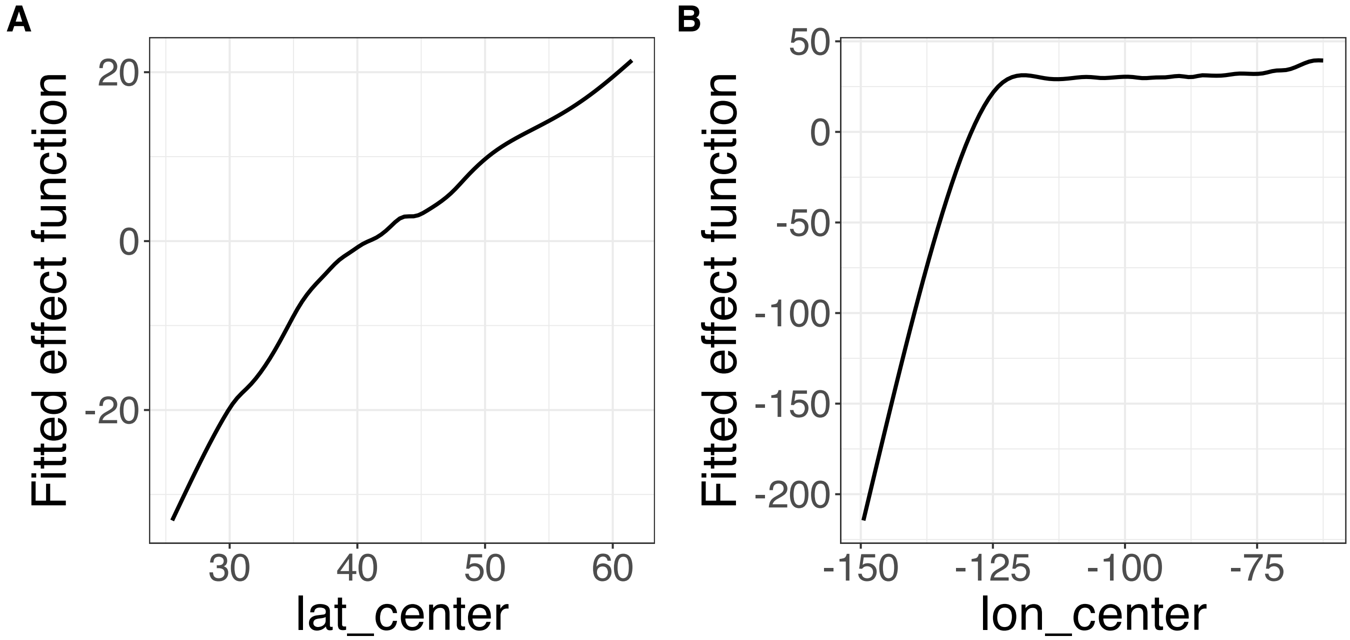


**Supplementary Figure 283.** Decomposition of reconstructed SSANOVA-SVD component function 10. (A) lat_center. (B) lon_center.

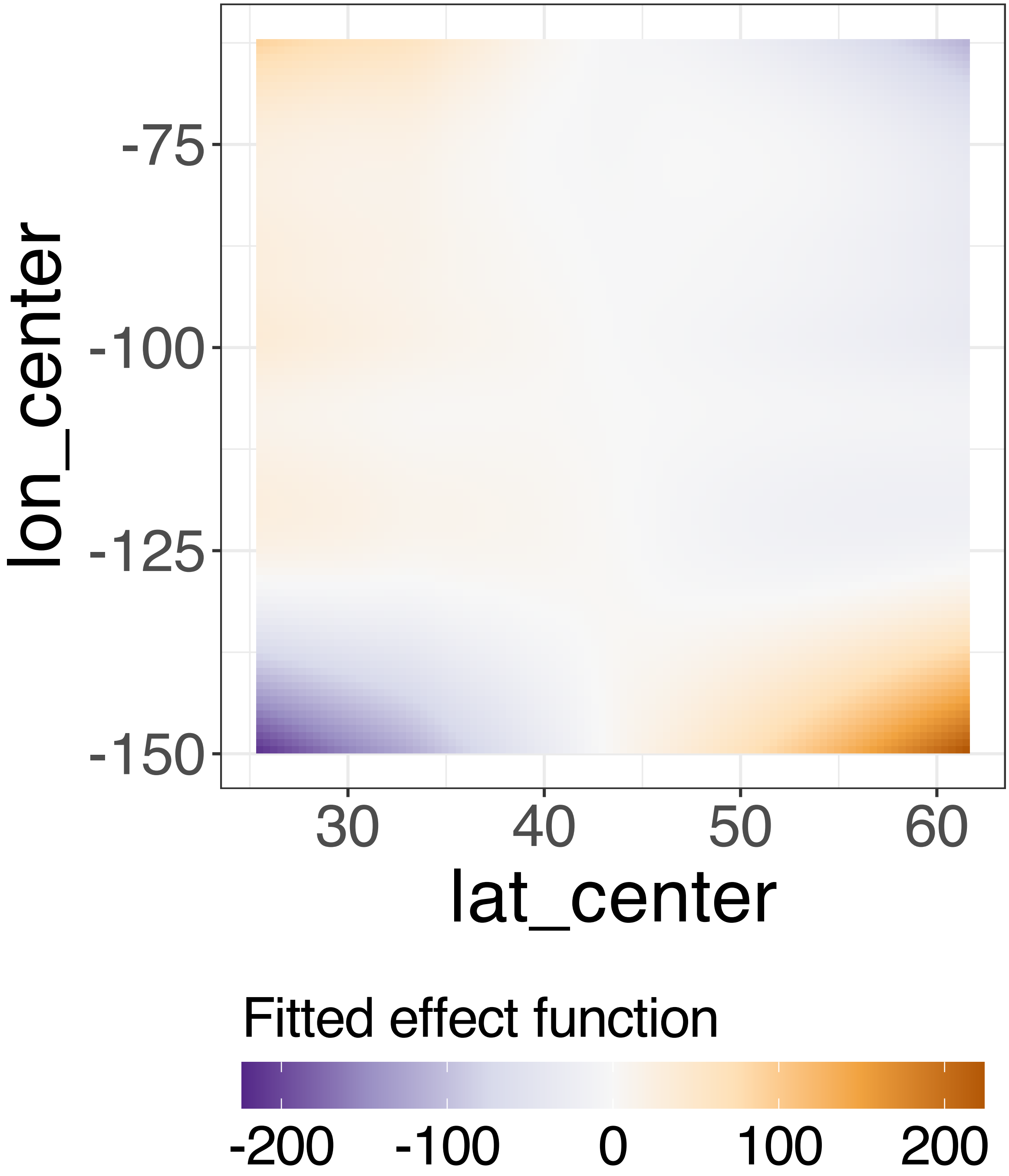


**Supplementary Figure 284.** Decomposition of reconstructed SSANOVA-SVD component function 10.

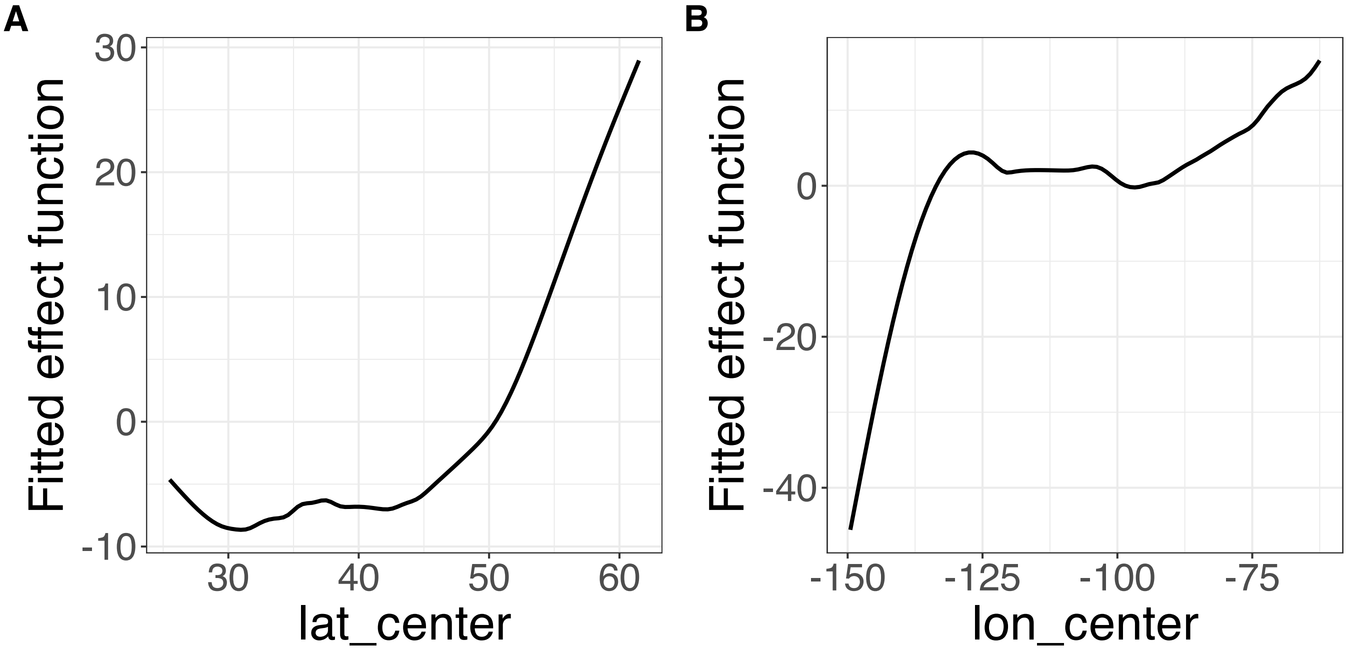


**Supplementary Figure 285.** Decomposition of reconstructed SSANOVA-SVD component function 11. (A) lat_center. (B) lon_center.

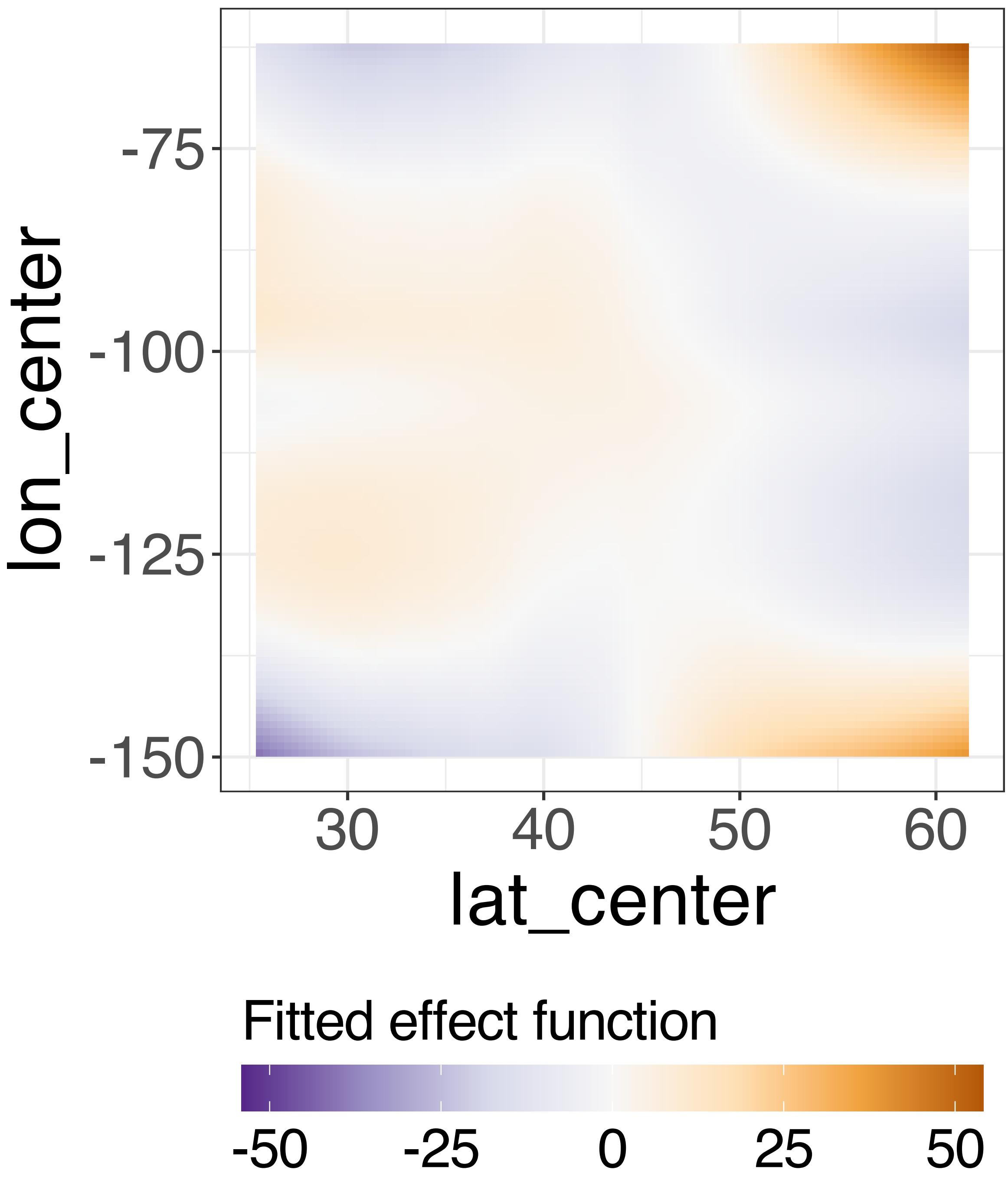


**Supplementary Figure 286.** Decomposition of reconstructed SSANOVA-SVD component function 11.

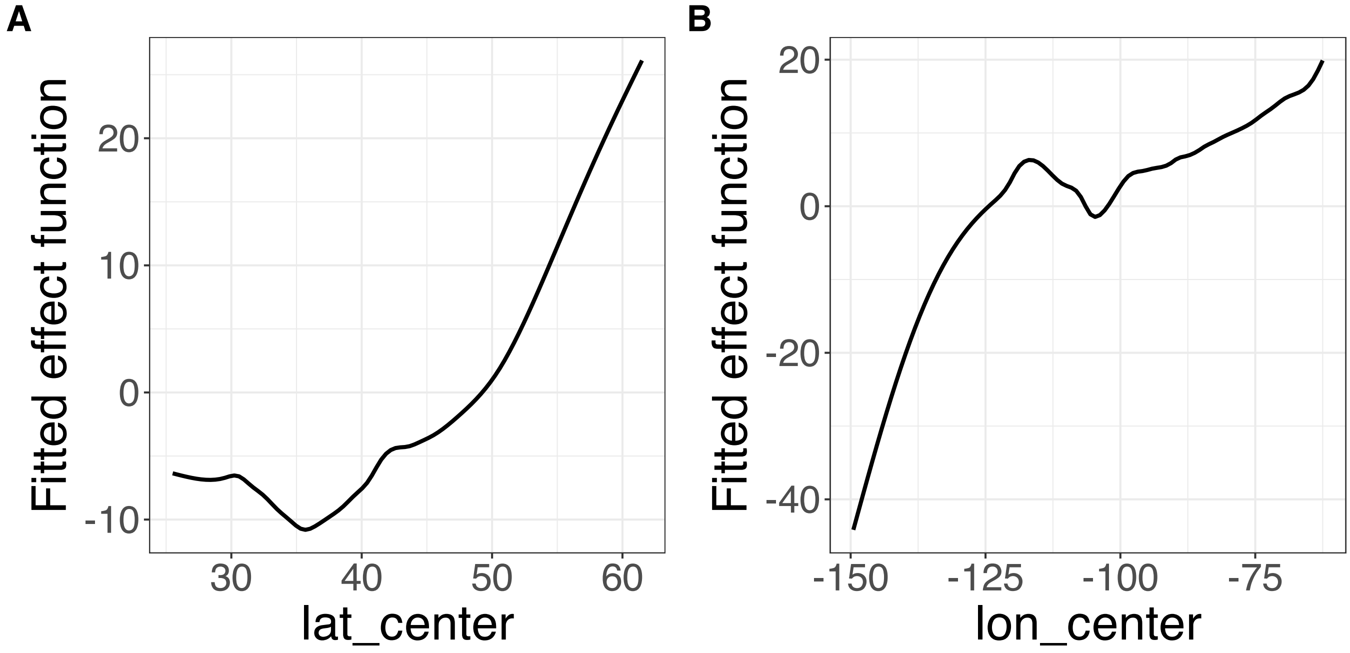


**Supplementary Figure 287.** Decomposition of reconstructed SSANOVA-SVD component function 12. (A) lat_center. (B) lon_center.

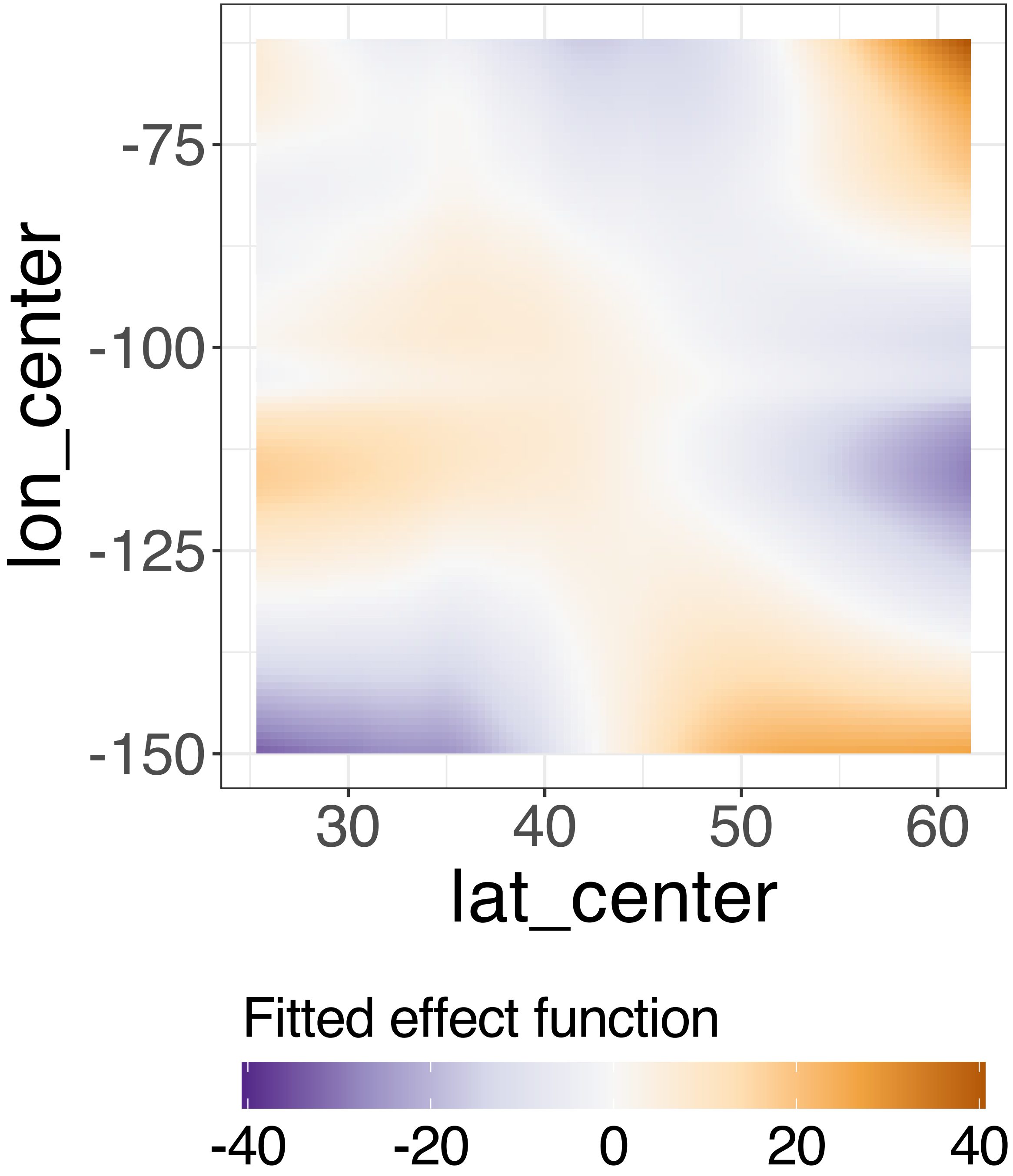


**Supplementary Figure 288.** Decomposition of reconstructed SSANOVA-SVD component function 12.

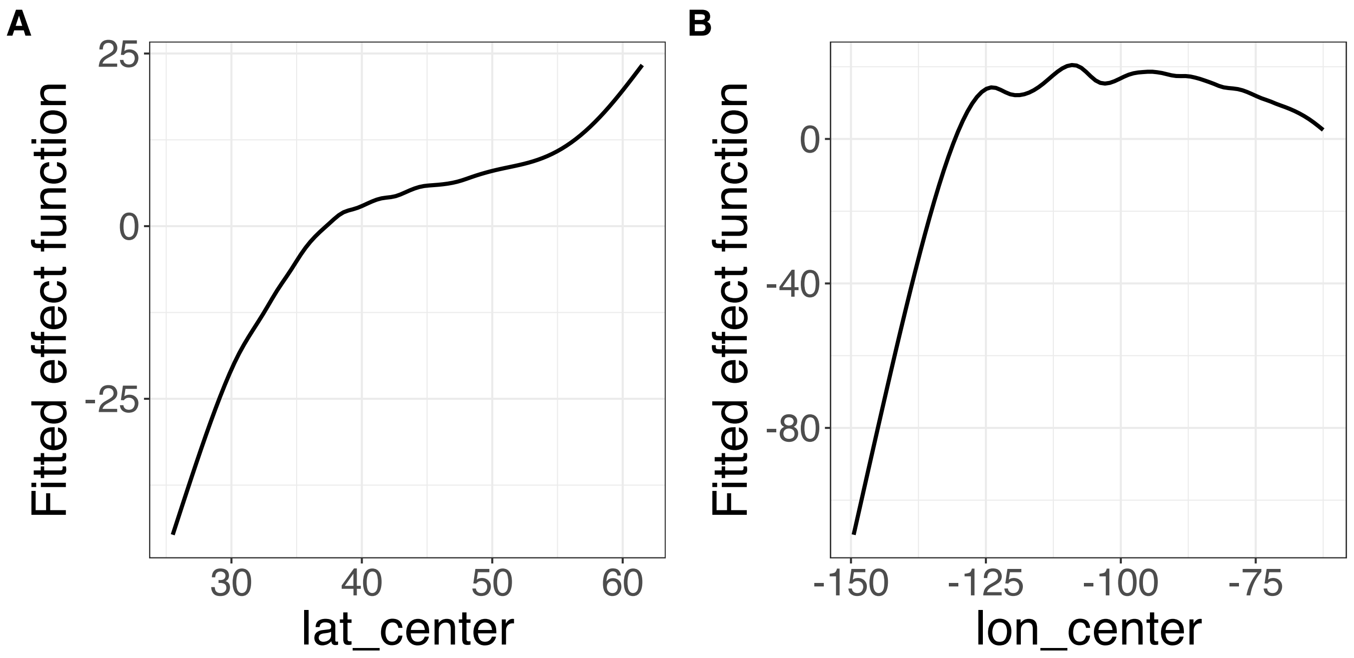


**Supplementary Figure 289.** Decomposition of reconstructed SSANOVA-SVD component function 13. (A) lat_center. (B) lon_center.

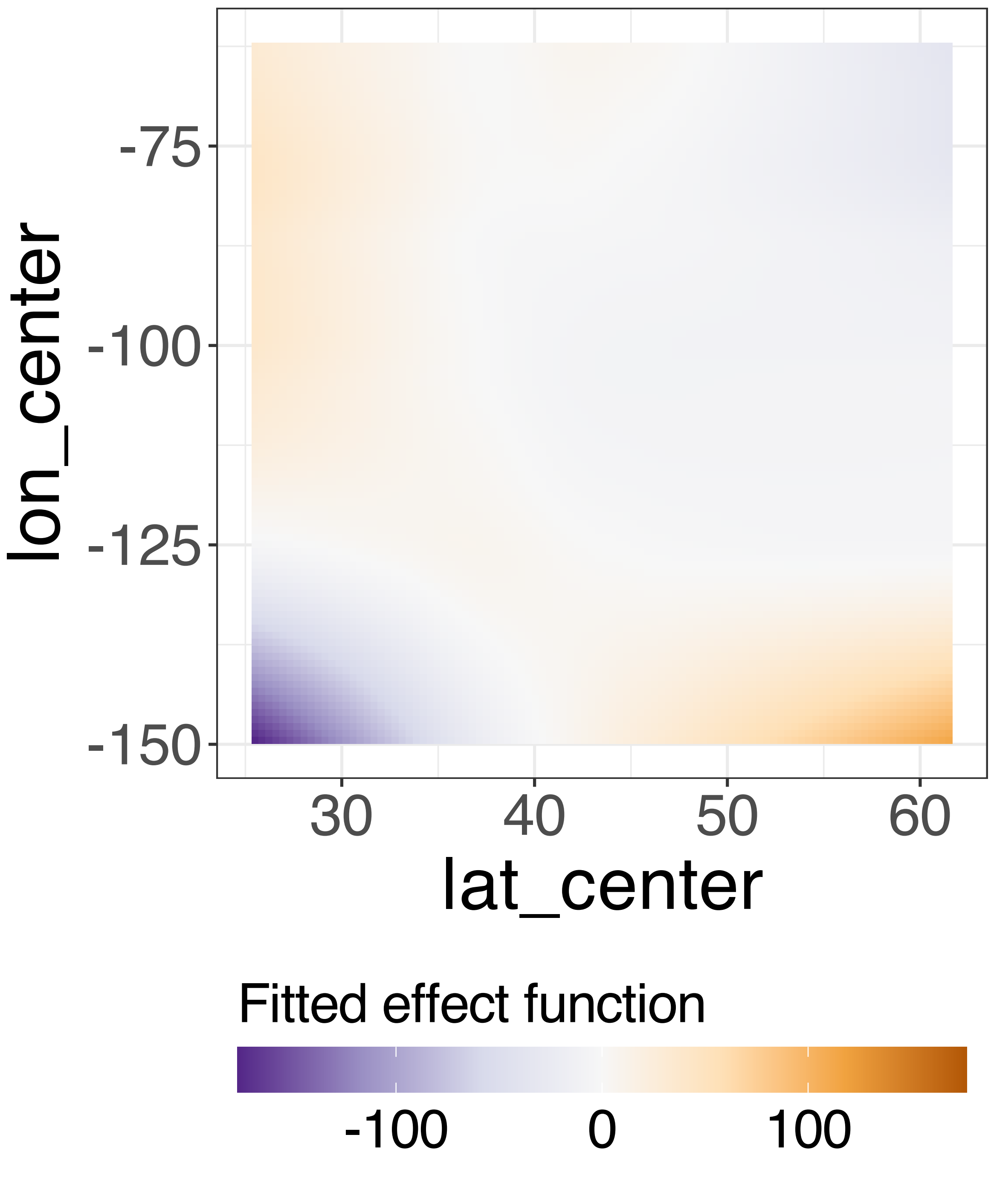


**Supplementary Figure 290.** Decomposition of reconstructed SSANOVA-SVD component function 13.

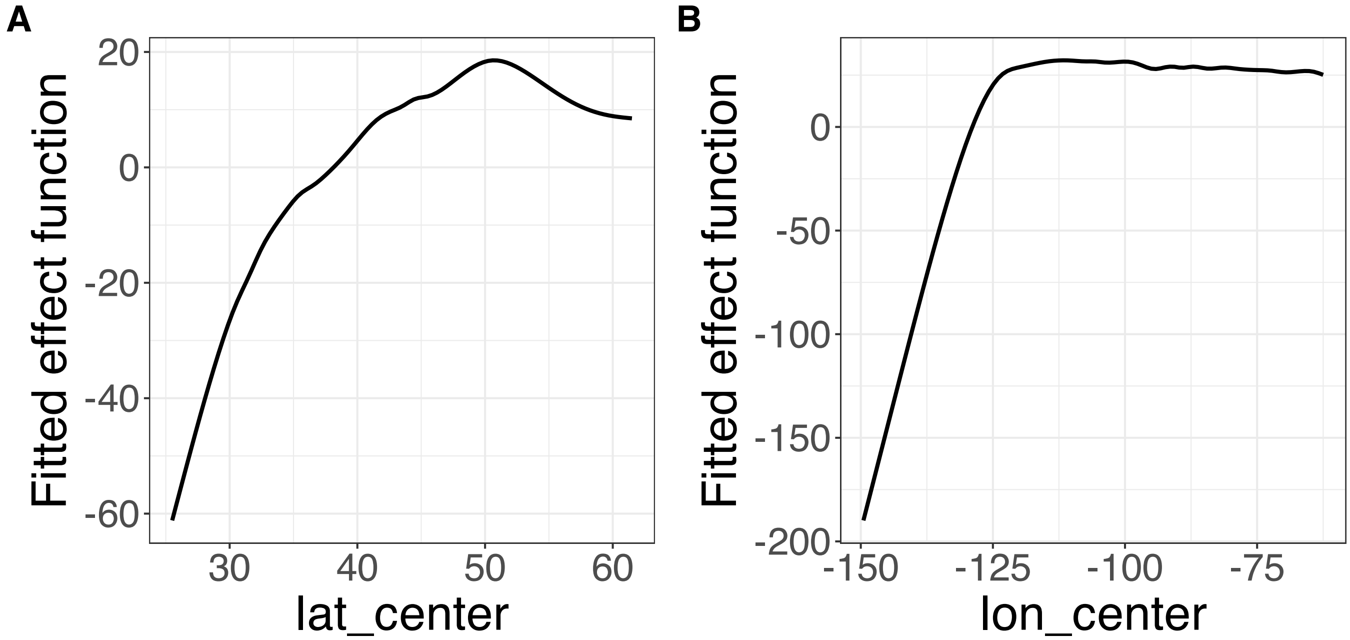


**Supplementary Figure 291.** Decomposition of reconstructed SSANOVA-SVD component function 14. (A) lat_center. (B) lon_center.

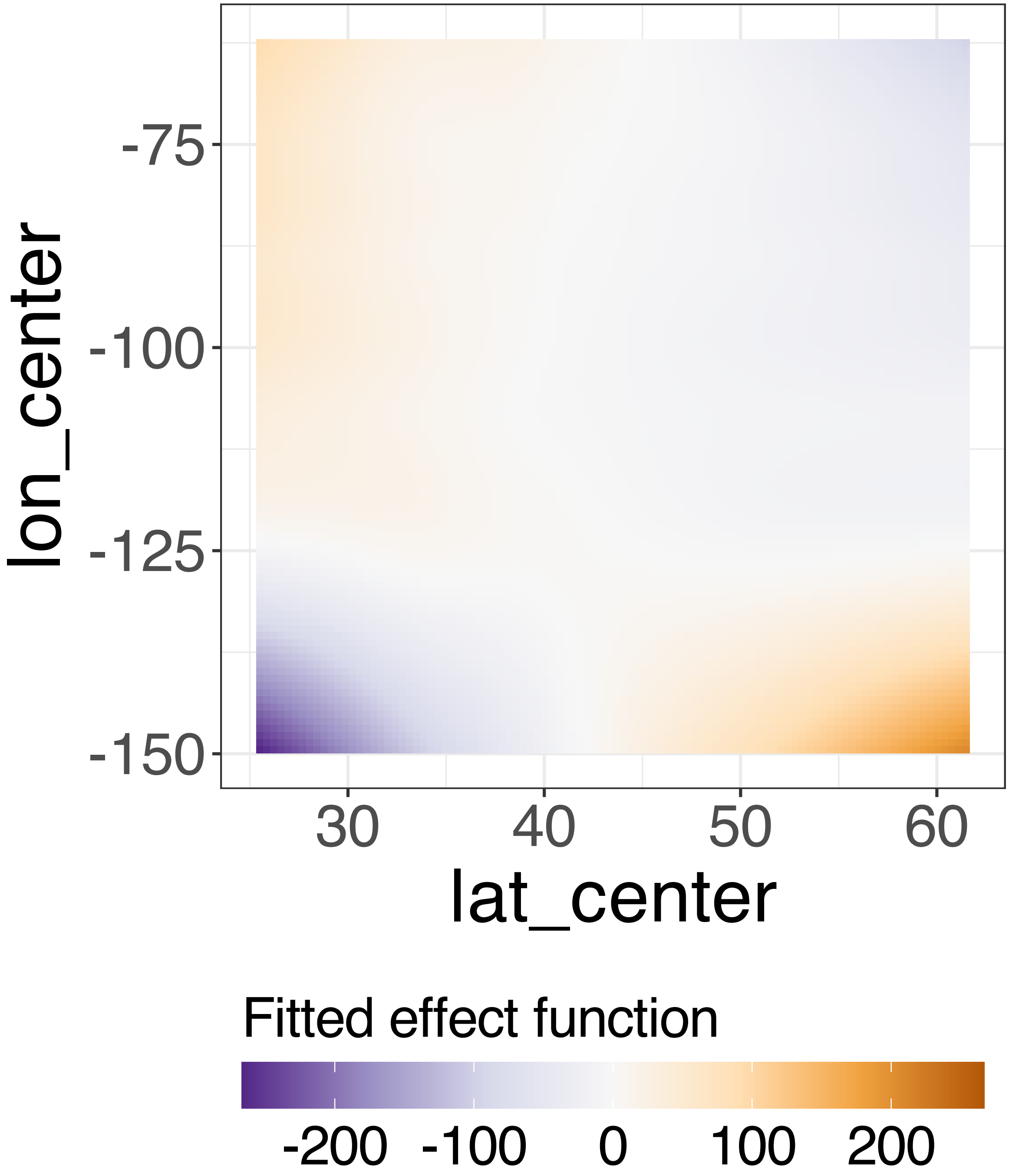


**Supplementary Figure 292.** Decomposition of reconstructed SSANOVA-SVD component function 14.

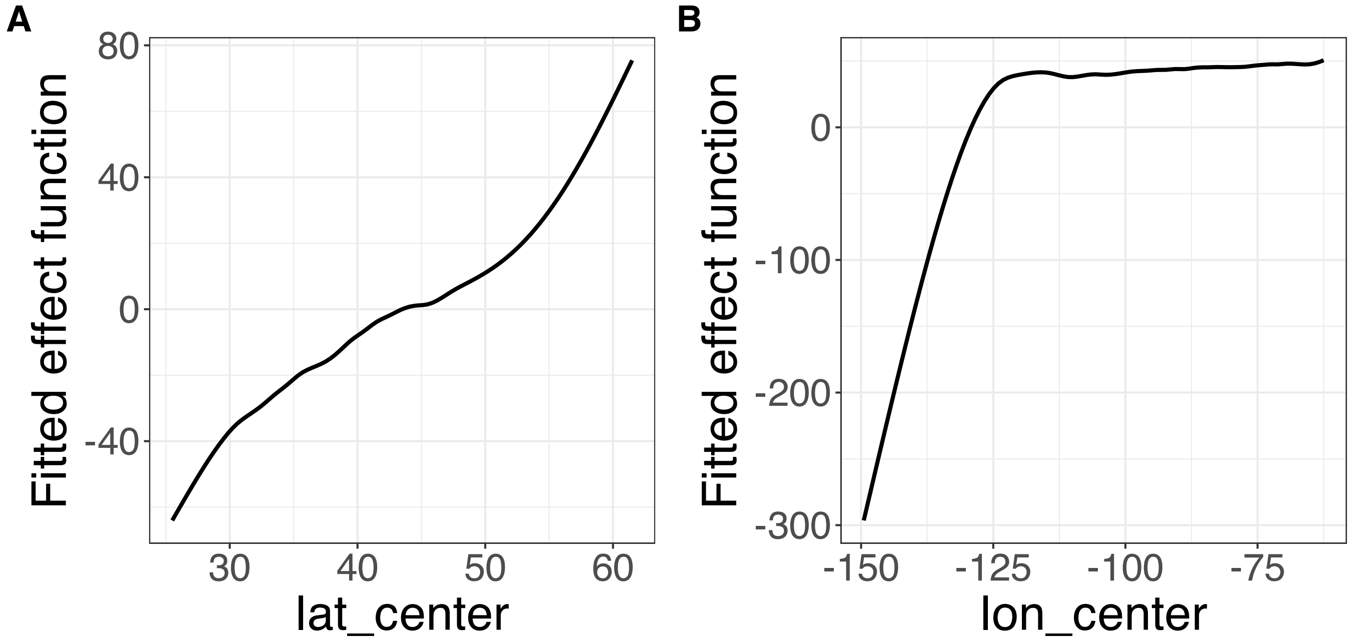


**Supplementary Figure 293.** Decomposition of reconstructed SSANOVA-SVD component function 15. (A) lat_center. (B) lon_center.

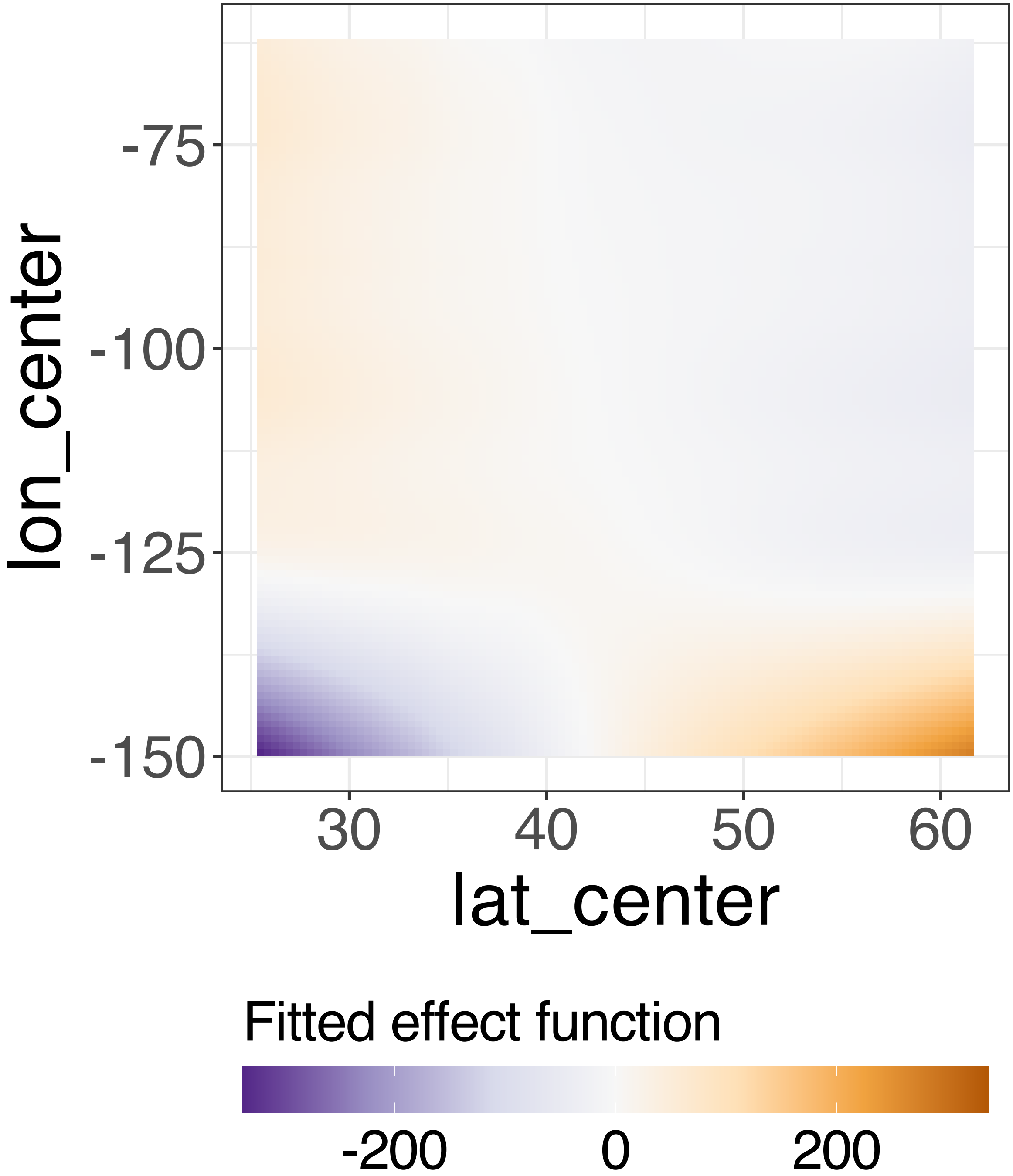


**Supplementary Figure 294.** Decomposition of reconstructed SSANOVA-SVD component function 15.

PCA

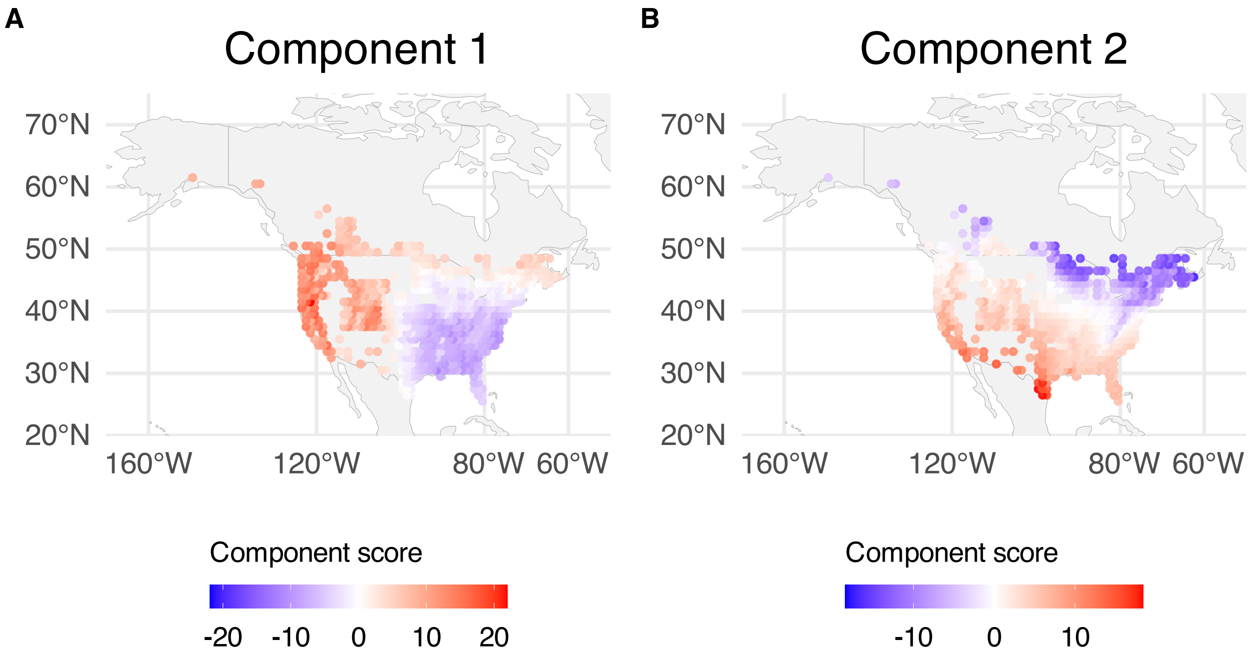


**Supplementary Figure 295.** PCA component scores mapped over BBS sampling locations; scores are averaged over years within spatial grid. (A) Component 1. (B) Component 2.

PCA

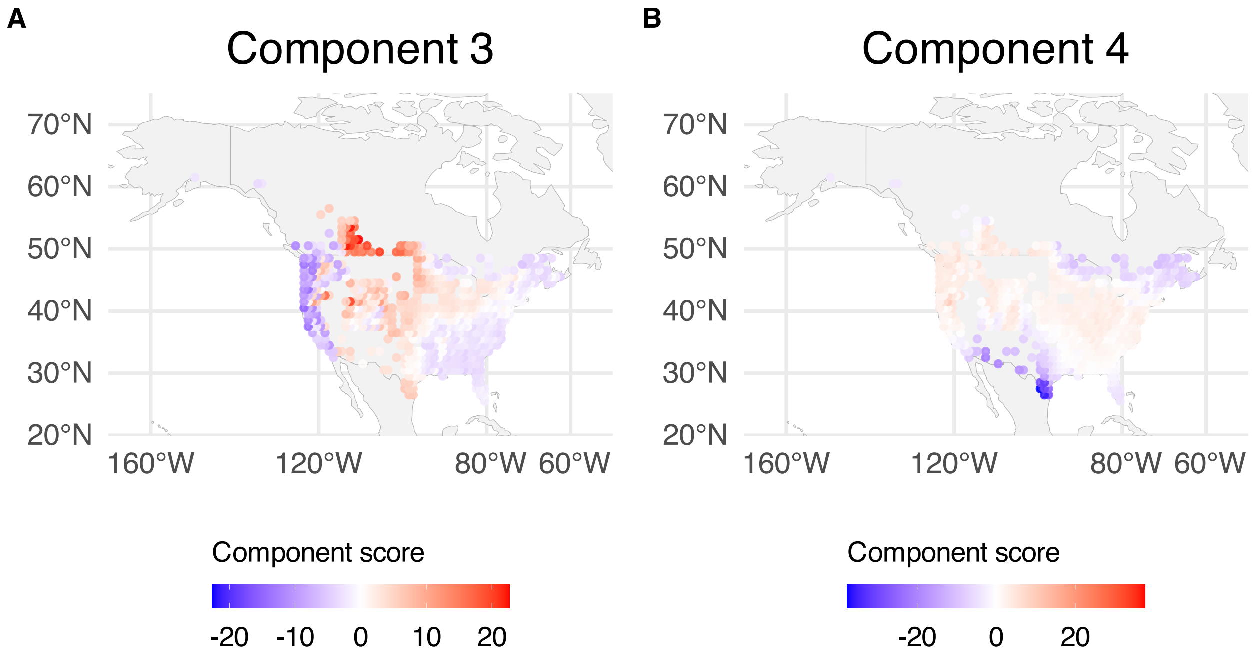


**Supplementary Figure 296.** PCA component scores mapped over BBS sampling locations; scores are averaged over years within spatial grid. (A) Component 3. (B) Component 4.

PCA

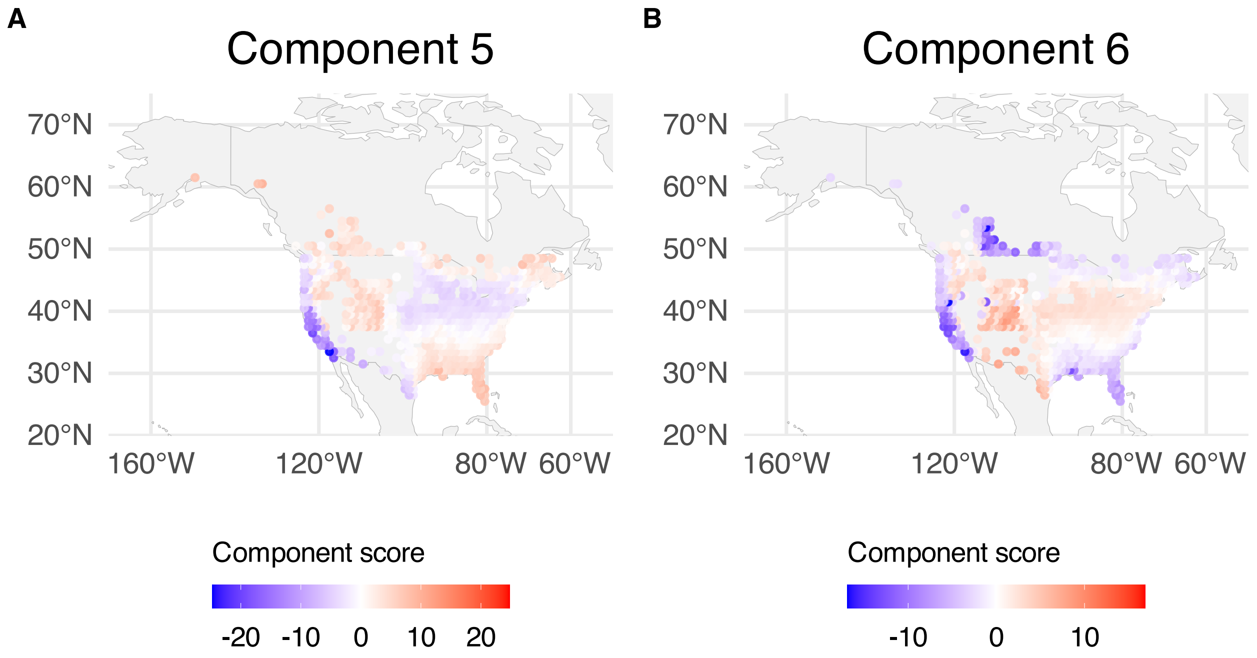


**Supplementary Figure 297.** PCA component scores mapped over BBS sampling locations; scores are averaged over years within spatial grid. (A) Component 5. (B) Component 6.

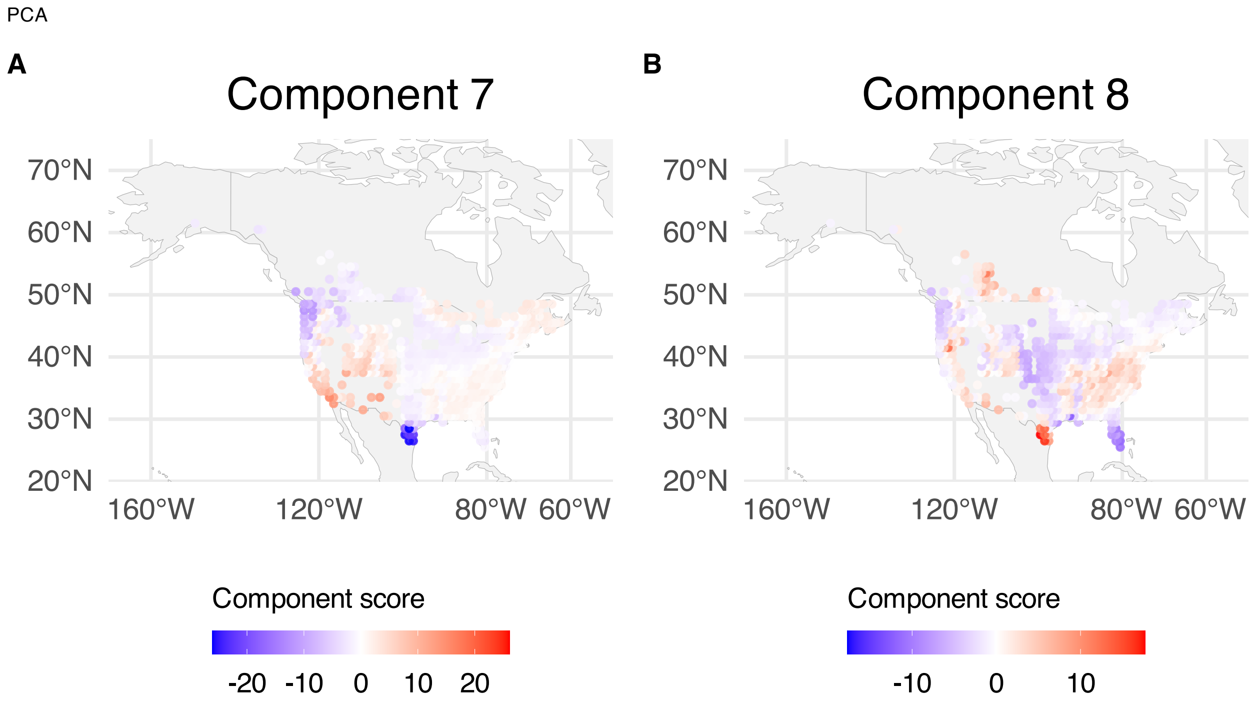


**Supplementary Figure 298.** PCA component scores mapped over BBS sampling locations; scores are averaged over years within spatial grid. (A) Component 7. (B) Component 8.

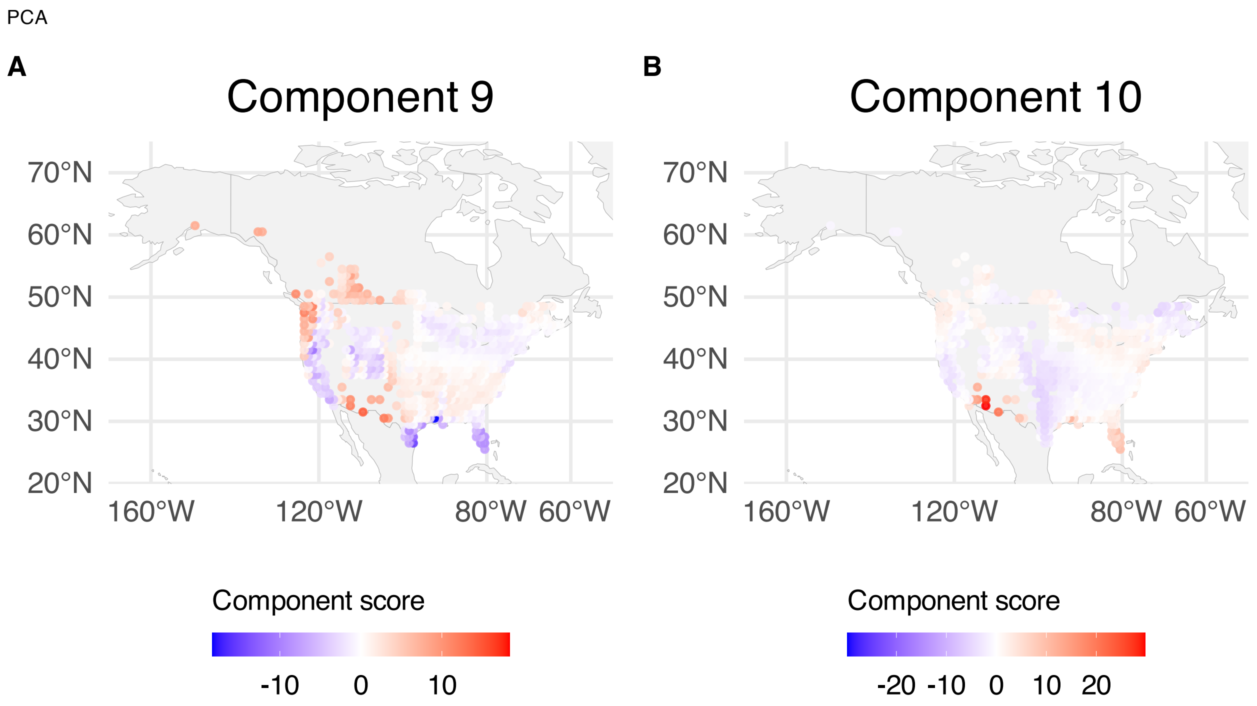


**Supplementary Figure 299.** PCA component scores mapped over BBS sampling locations; scores are averaged over years within spatial grid. (A) Component 9. (B) Component 10.

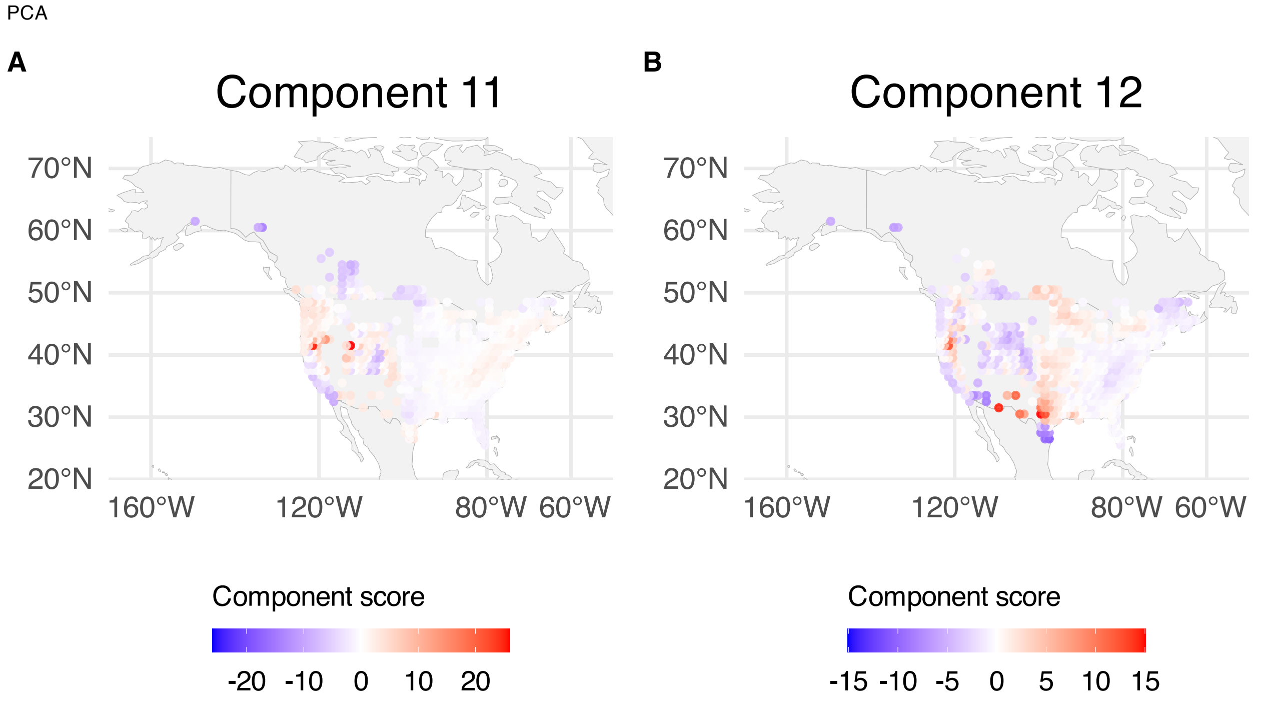


**Supplementary Figure 300.** PCA component scores mapped over BBS sampling locations; scores are averaged over years within spatial grid. (A) Component 11. (B) Component 12.

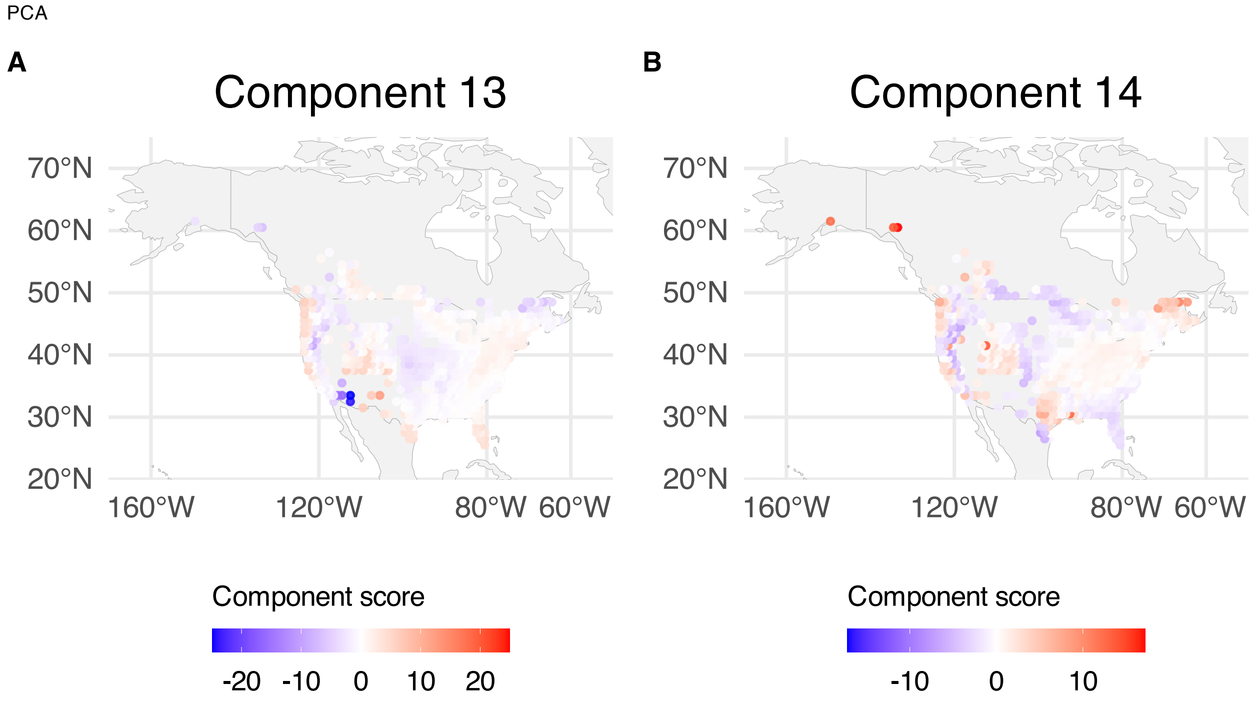


**Supplementary Figure 301.** PCA component scores mapped over BBS sampling locations; scores are averaged over years within spatial grid. (A) Component 13. (B) Component 14.

PCA

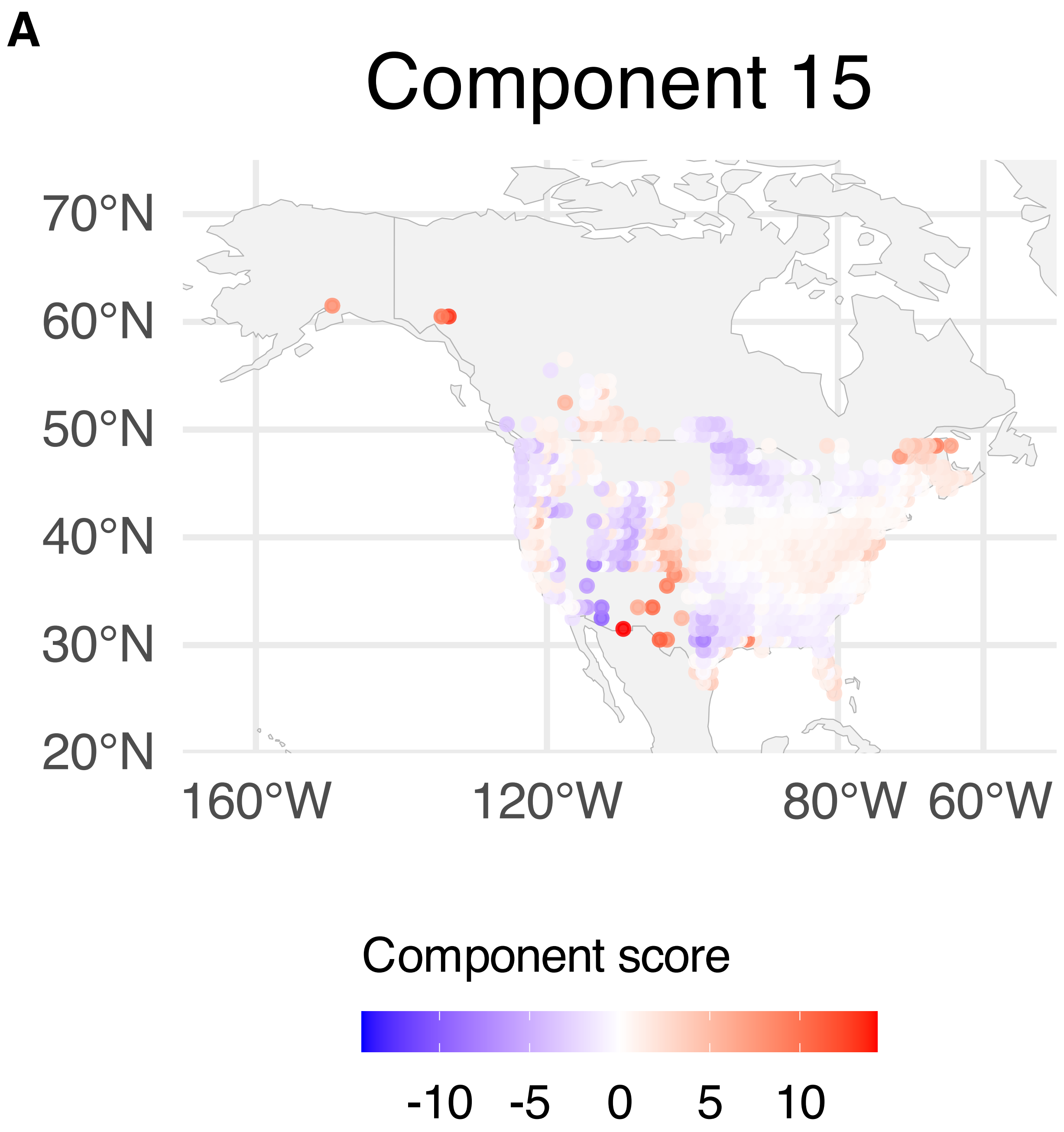


**Supplementary Figure 302.** PCA component scores mapped over BBS sampling locations; scores are averaged over years within spatial grid. Component 15.

PLS

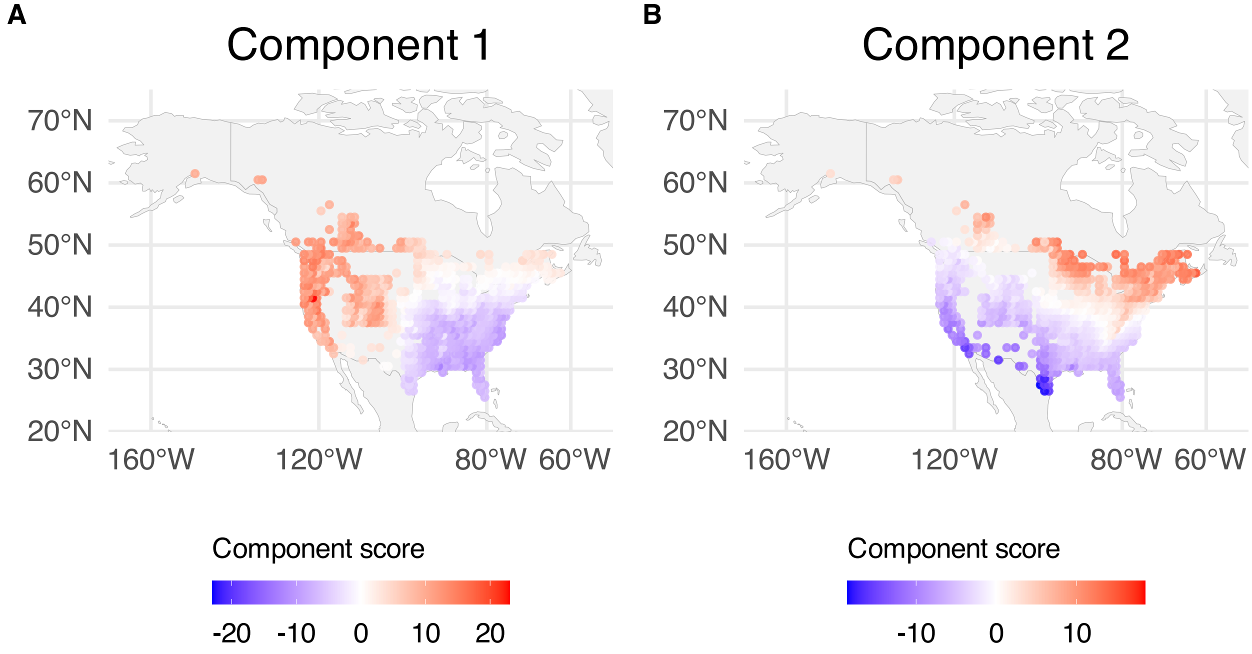


**Supplementary Figure 303.** PLS component scores mapped over BBS sampling locations; scores are averaged over years within spatial grid. (A) Component 1. (B) Component 2.

PLS

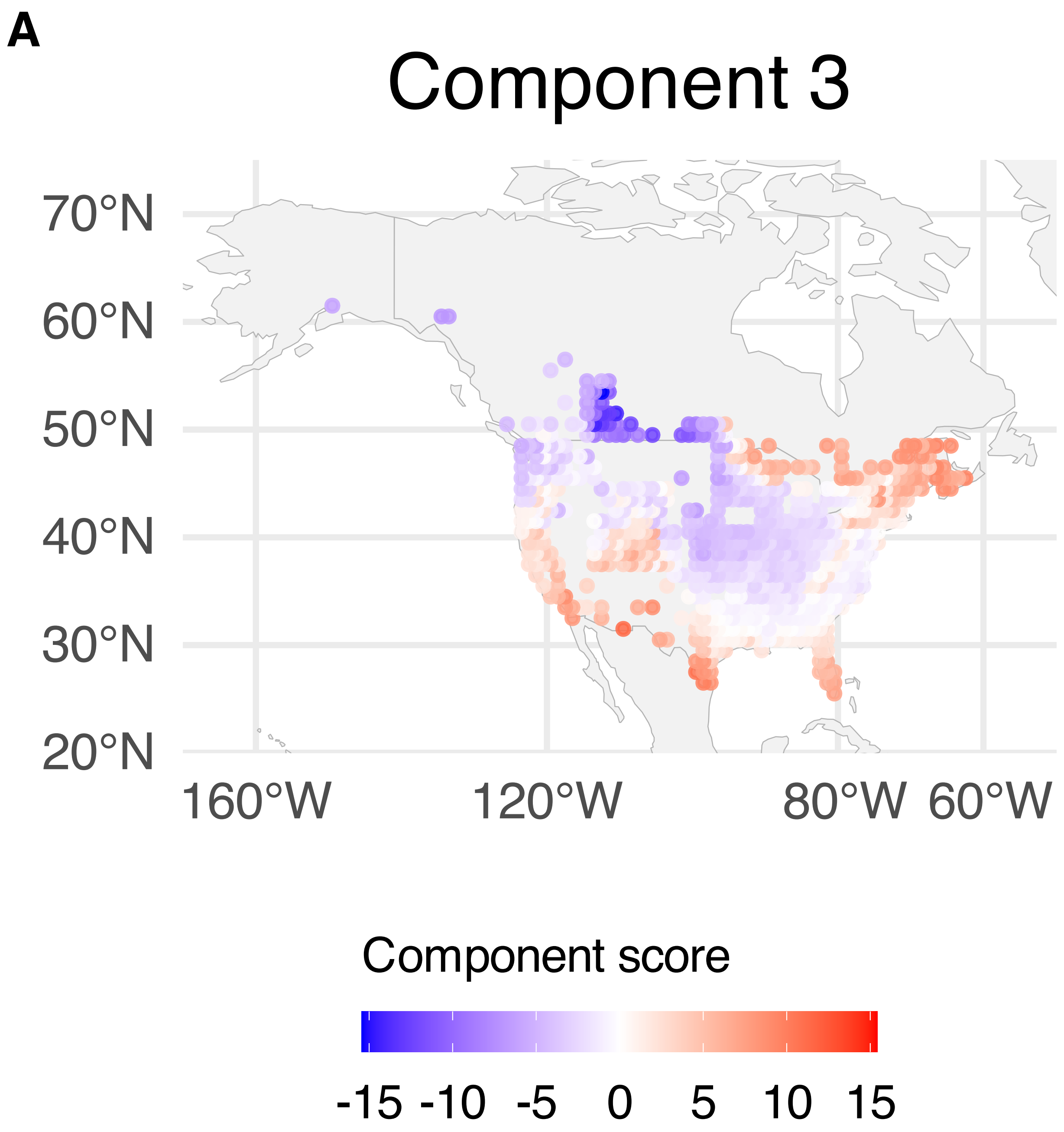


**Supplementary Figure 304.** PLS component scores mapped over BBS sampling locations; scores are averaged over years within spatial grid. Component 3.

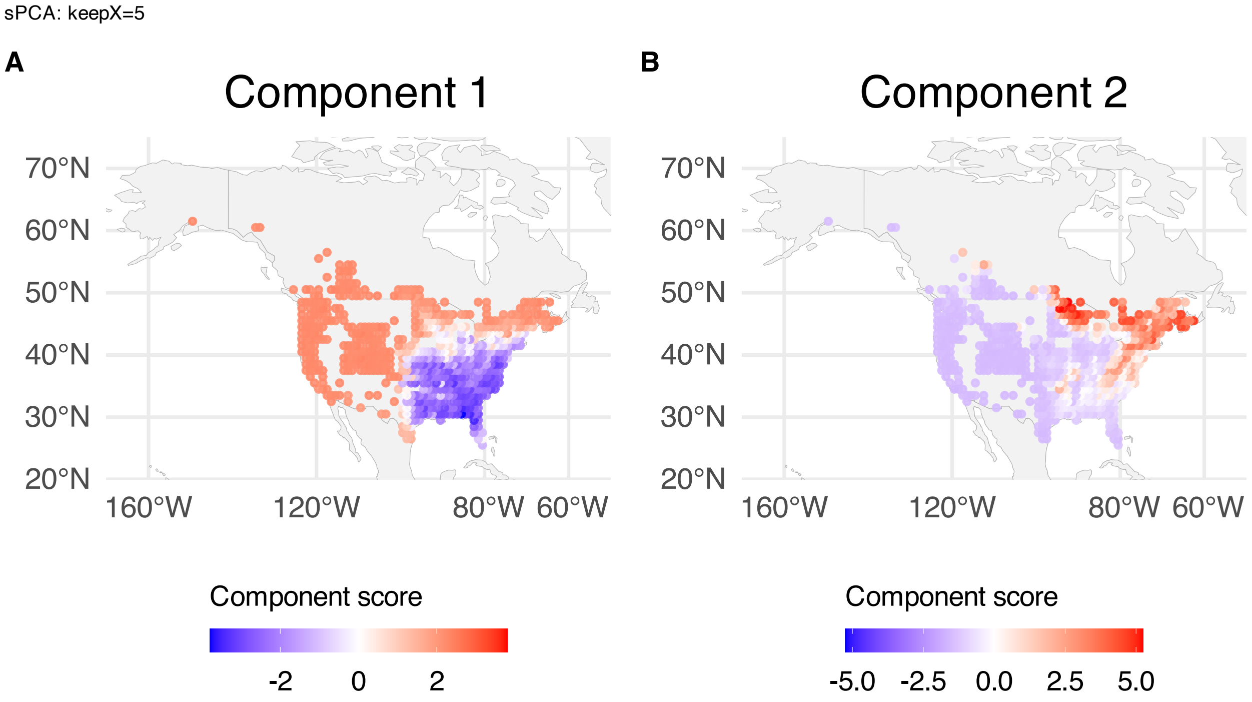


**Supplementary Figure 305.** sPCA: keepX=5 component scores mapped over BBS sampling locations; scores are averaged over years within spatial grid. (A) Component 1. (B) Component 2.

sPCA: keepX=5

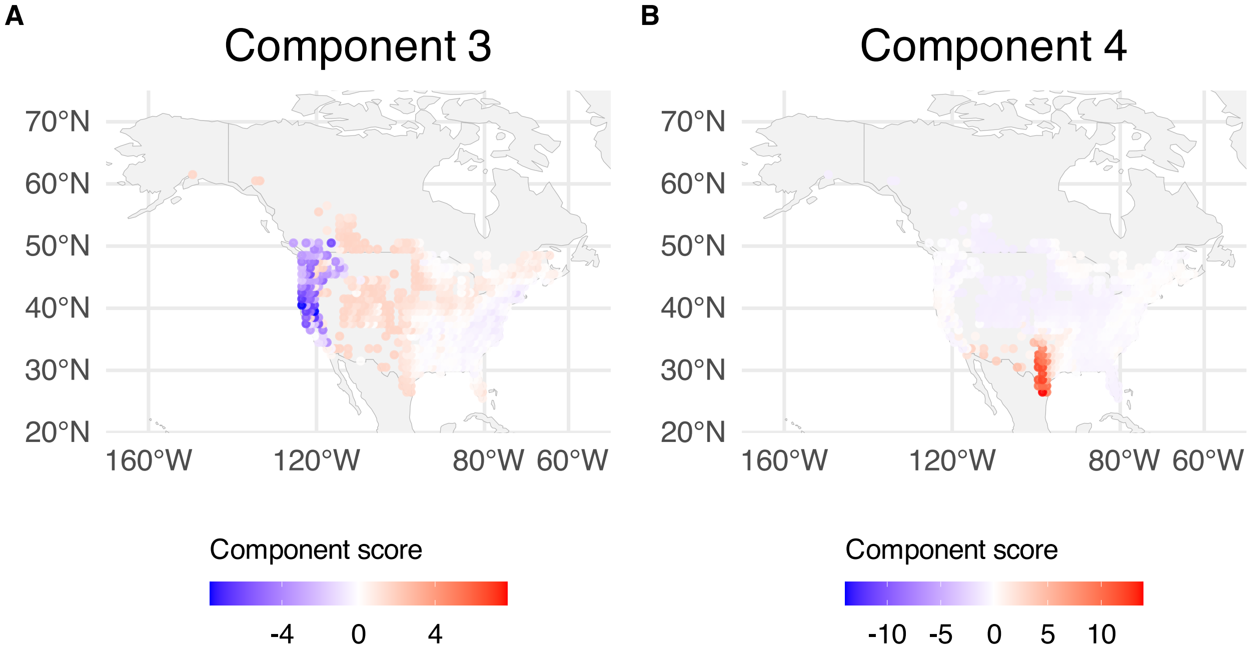


**Supplementary Figure 306.** sPCA: keepX=5 component scores mapped over BBS sampling locations; scores are averaged over years within spatial grid. (A) Component 3. (B) Component 4.

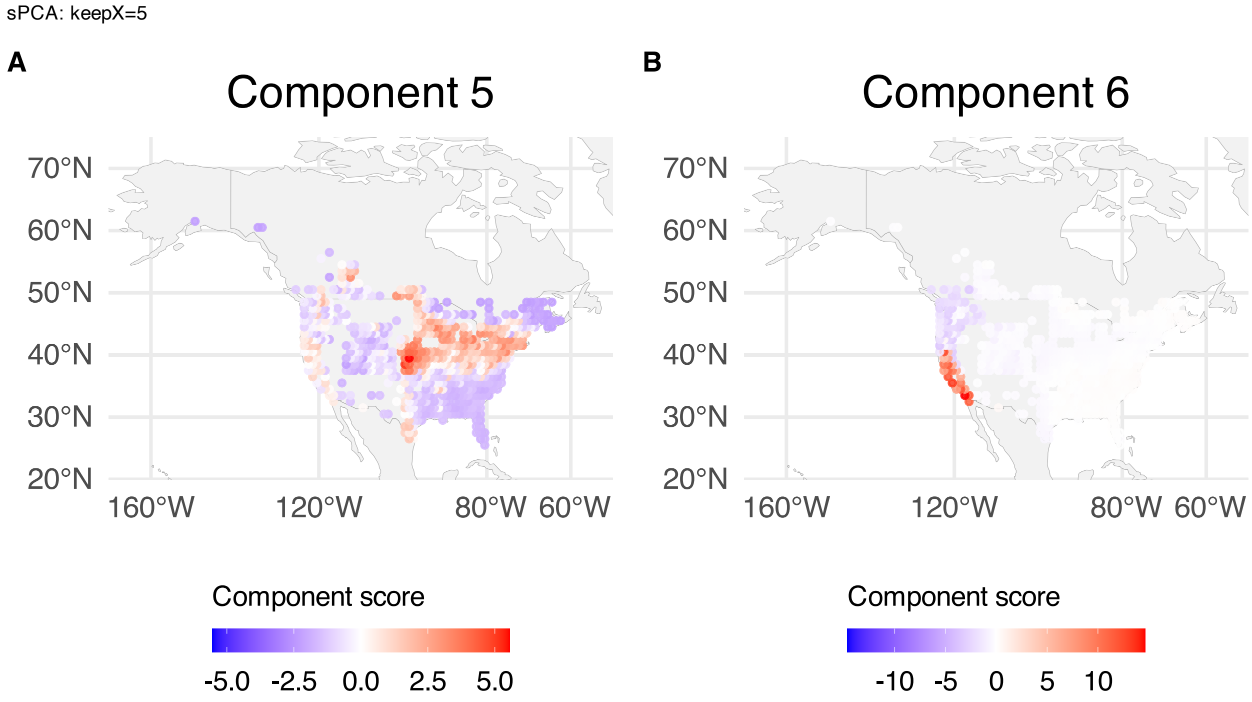


**Supplementary Figure 307.** sPCA: keepX=5 component scores mapped over BBS sampling locations; scores are averaged over years within spatial grid. (A) Component 5. (B) Component 6.

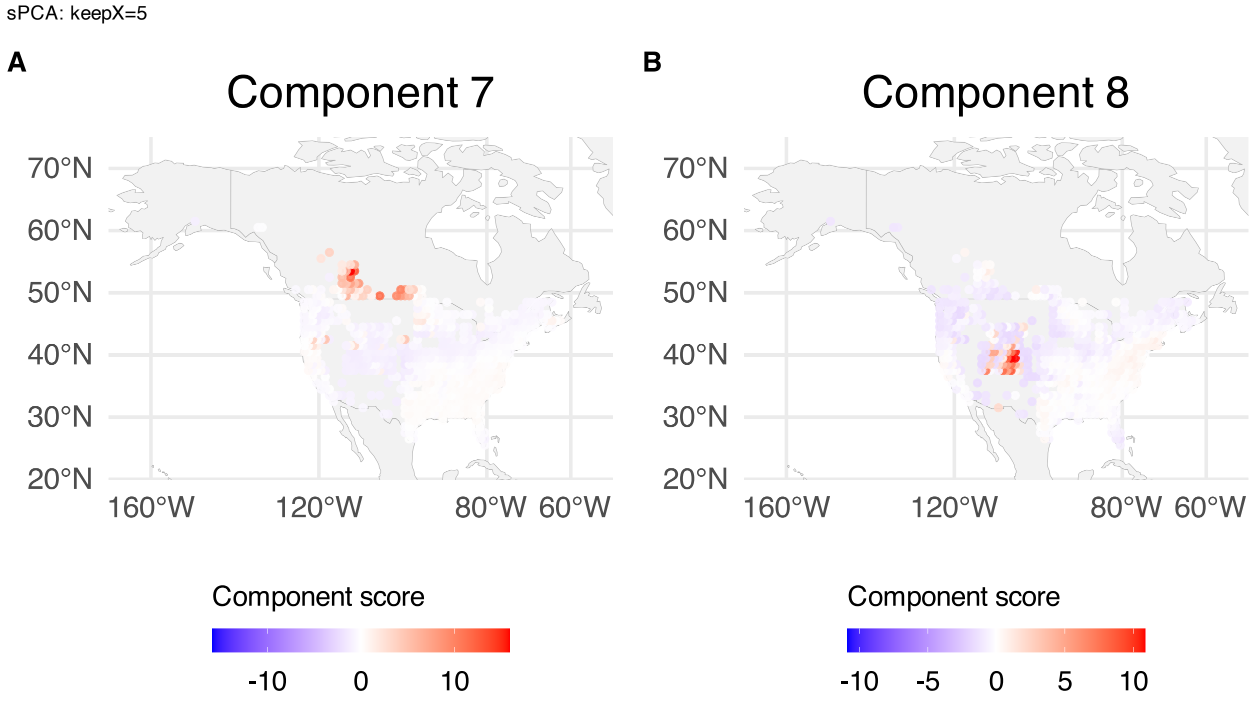


**Supplementary Figure 308.** sPCA: keepX=5 component scores mapped over BBS sampling locations; scores are averaged over years within spatial grid. (A) Component 7. (B) Component 8.

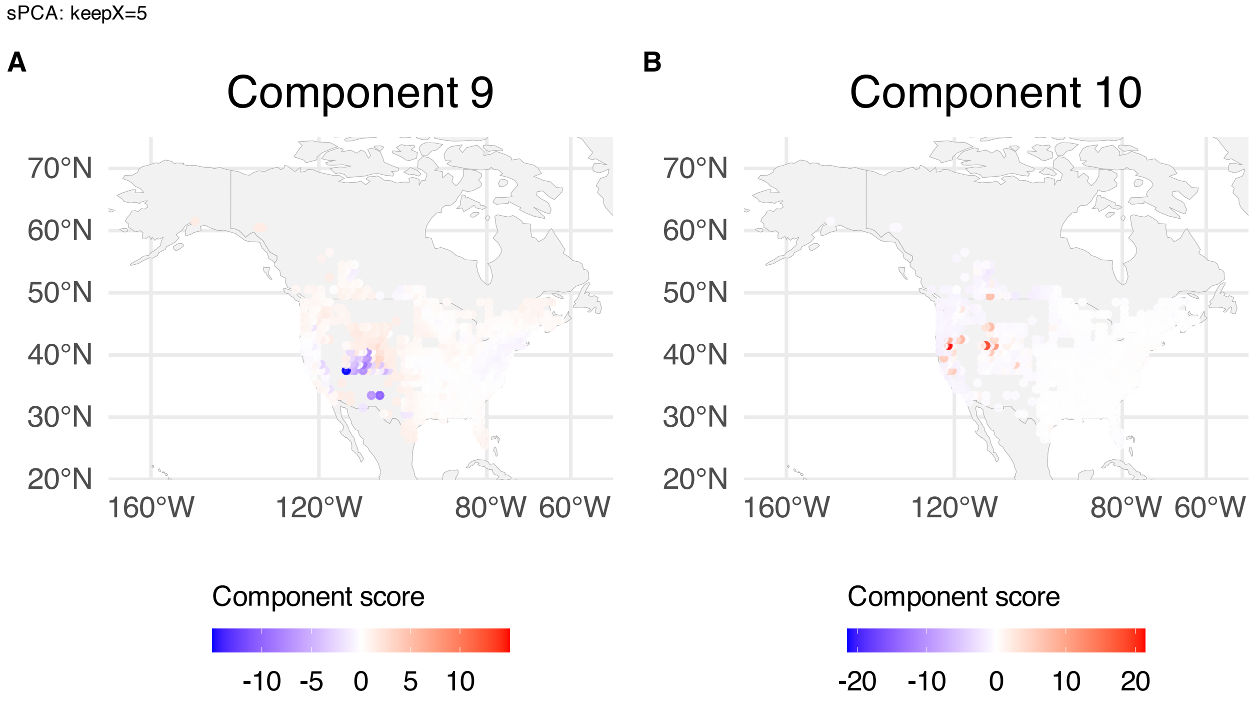


**Supplementary Figure 309.** sPCA: keepX=5 component scores mapped over BBS sampling locations; scores are averaged over years within spatial grid. (A) Component 9. (B) Component 10.

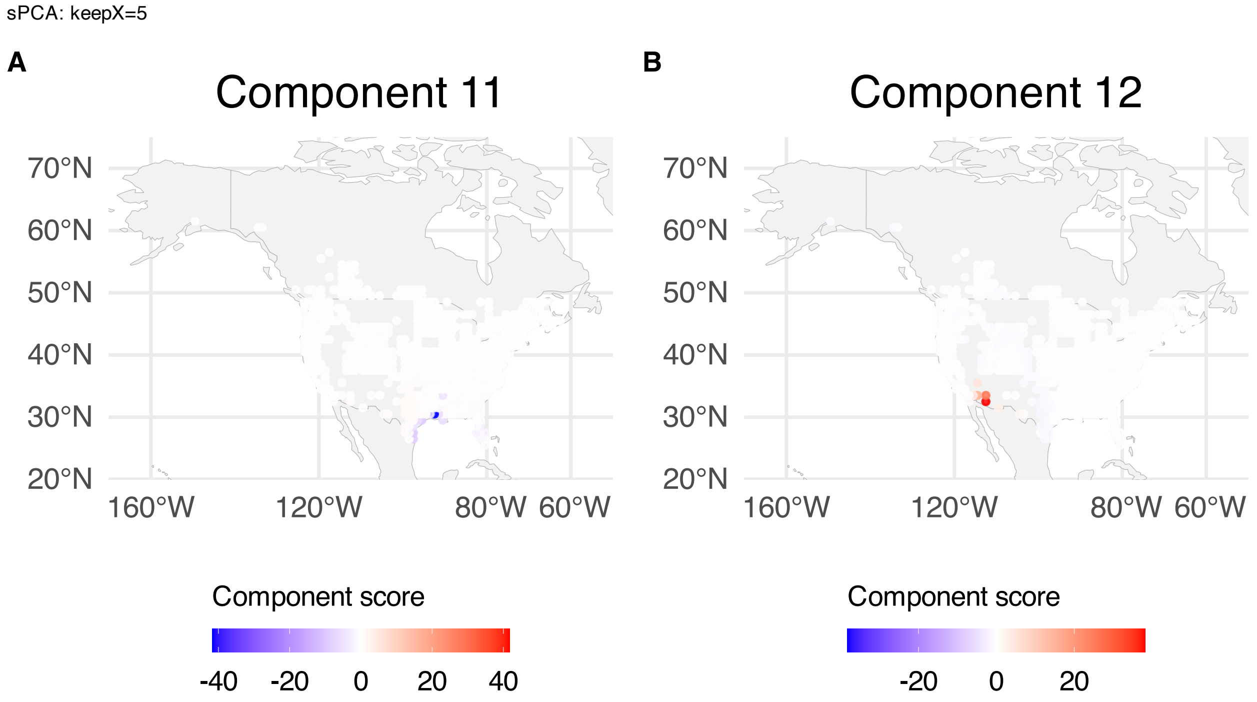


**Supplementary Figure 310.** sPCA: keepX=5 component scores mapped over BBS sampling locations; scores are averaged over years within spatial grid. (A) Component 11. (B) Component 12.

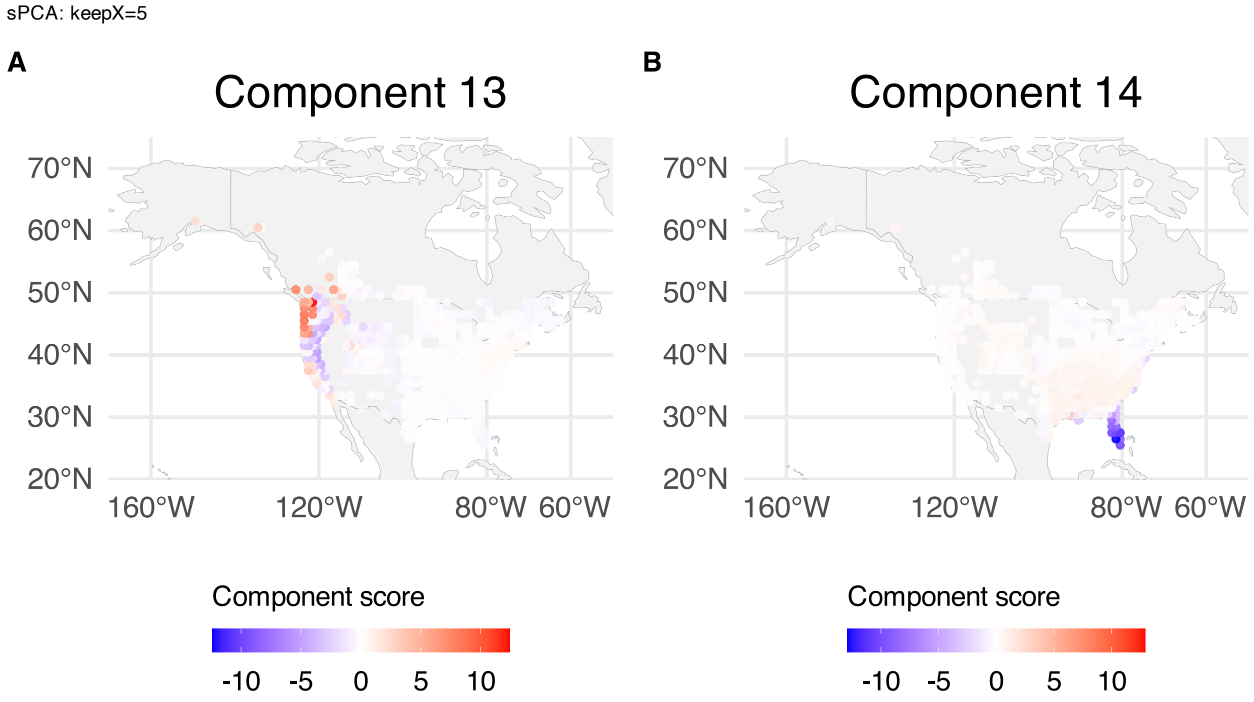


**Supplementary Figure 311.** sPCA: keepX=5 component scores mapped over BBS sampling locations; scores are averaged over years within spatial grid. (A) Component 13. (B) Component 14.

sPCA: keepX=5

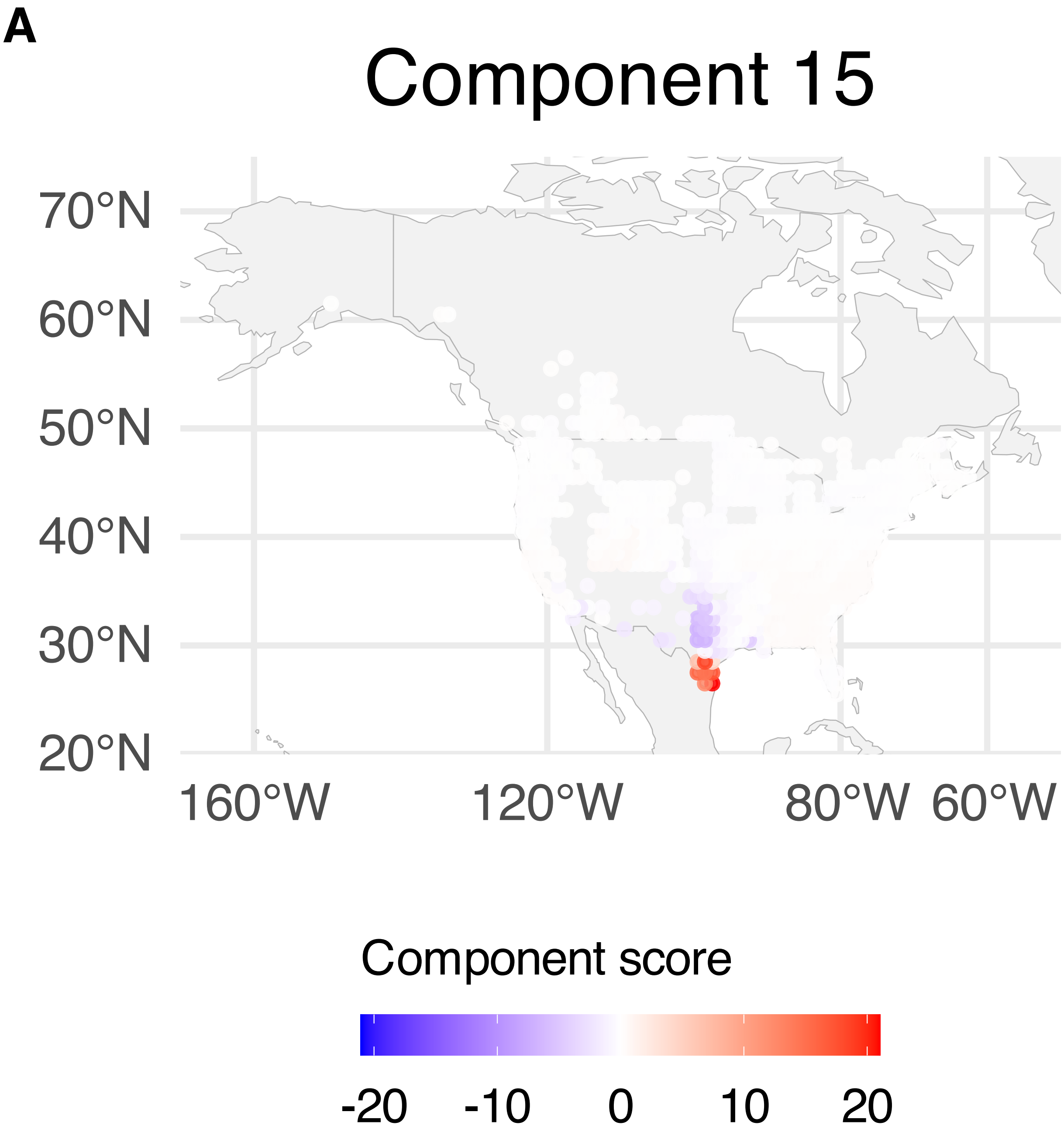


**Supplementary Figure 312.** sPCA: keepX=5 component scores mapped over BBS sampling locations; scores are averaged over years within spatial grid. Component 15.

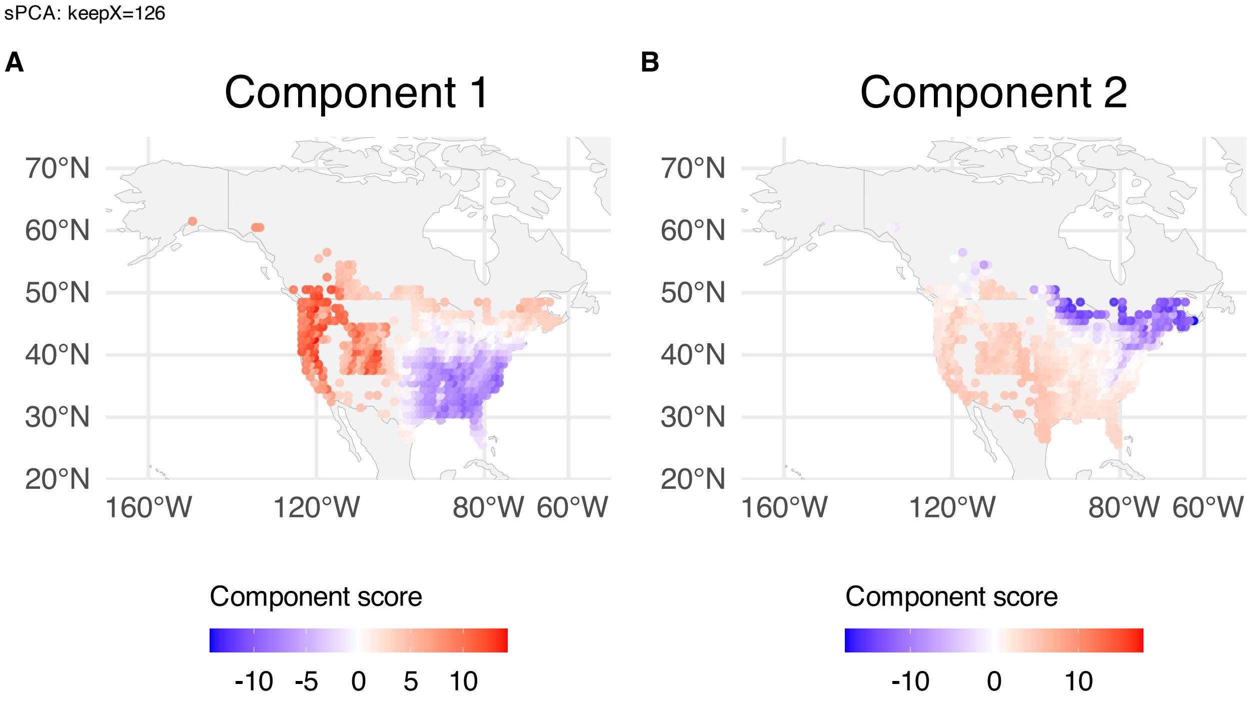


**Supplementary Figure 313.** sPCA: keepX=126 component scores mapped over BBS sampling locations; scores are averaged over years within spatial grid. (A) Component 1. (B) Component 2.

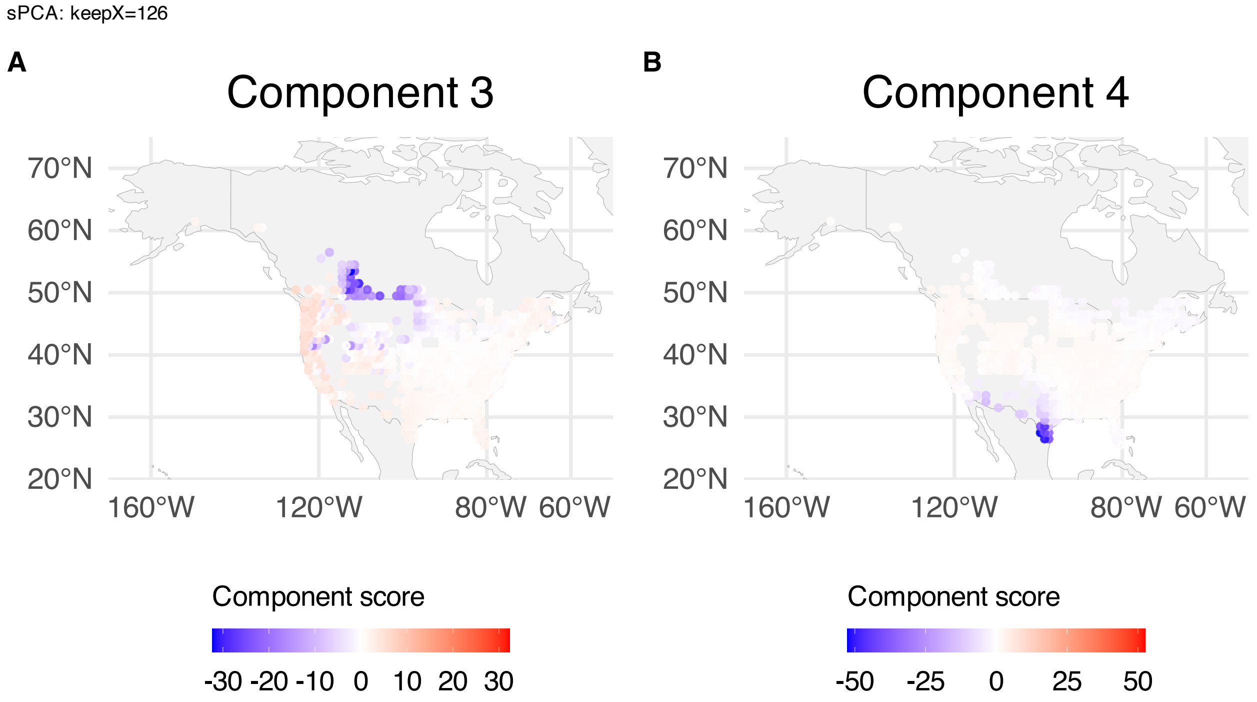


**Supplementary Figure 314.** sPCA: keepX=126 component scores mapped over BBS sampling locations; scores are averaged over years within spatial grid. (A) Component 3. (B) Component 4.

sPCA: keepX=126

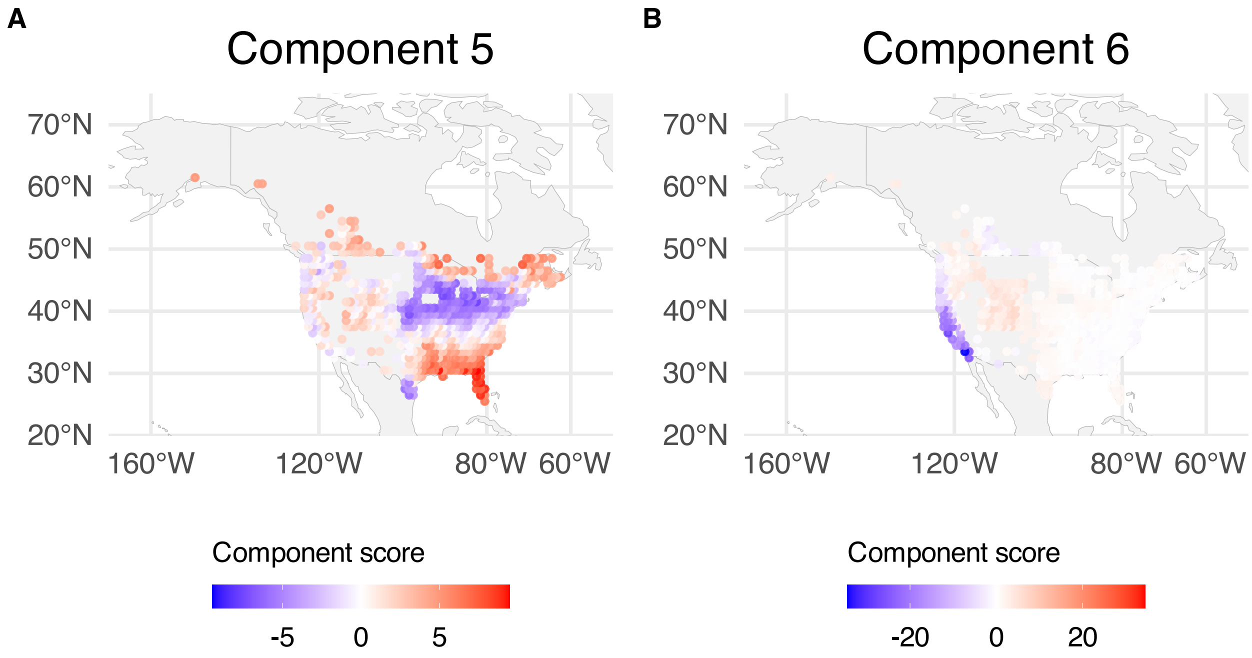


**Supplementary Figure 315.** sPCA: keepX=126 component scores mapped over BBS sampling locations; scores are averaged over years within spatial grid. (A) Component 5. (B) Component 6.

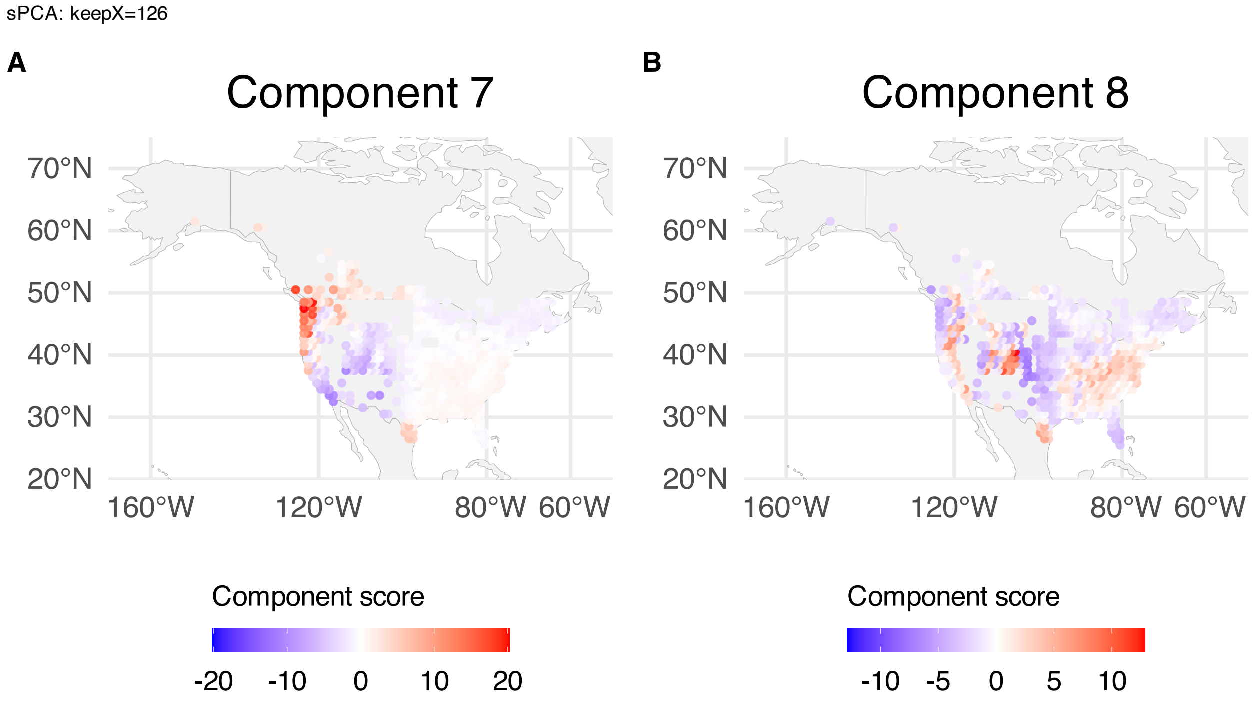


**Supplementary Figure 316.** sPCA: keepX=126 component scores mapped over BBS sampling locations; scores are averaged over years within spatial grid. (A) Component 7. (B) Component 8.

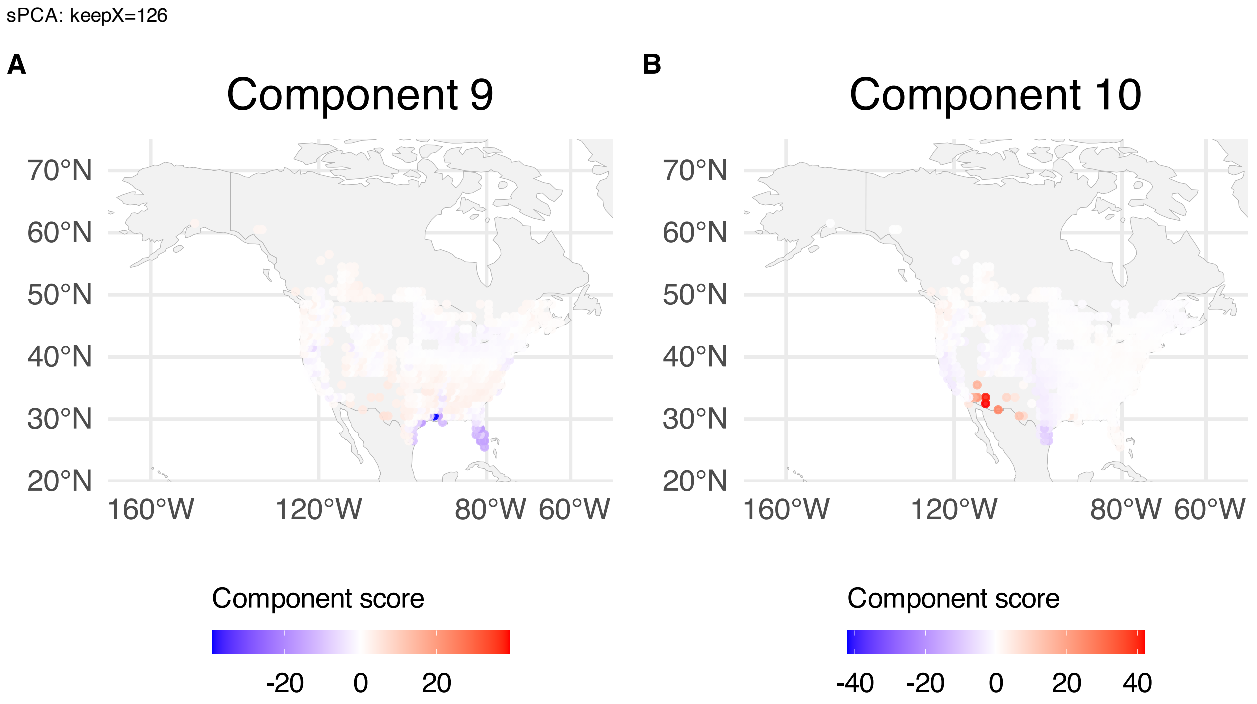


**Supplementary Figure 317.** sPCA: keepX=126 component scores mapped over BBS sampling locations; scores are averaged over years within spatial grid. (A) Component 9. (B) Component 10.

sPCA: keepX=126

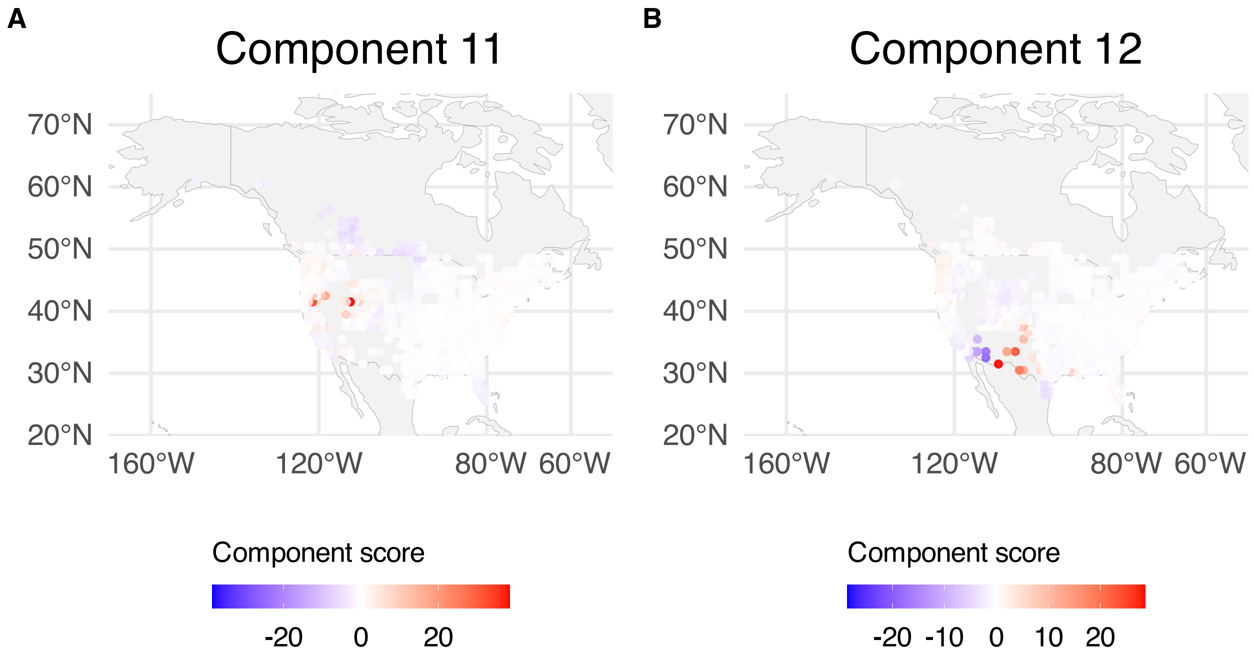


**Supplementary Figure 318.** sPCA: keepX=126 component scores mapped over BBS sampling locations; scores are averaged over years within spatial grid. (A) Component 11. (B) Component 12.

sPCA: keepX=126

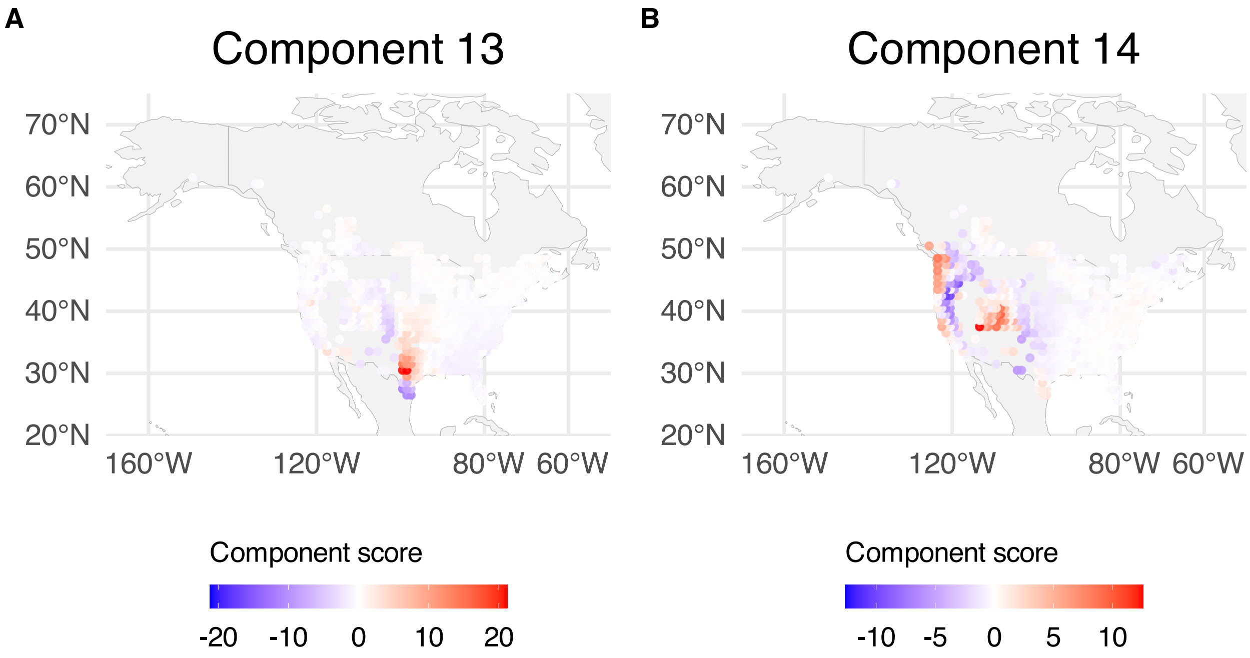


**Supplementary Figure 319.** sPCA: keepX=126 component scores mapped over BBS sampling locations; scores are averaged over years within spatial grid. (A) Component 13. (B) Component 14.

sPCA: keepX=126

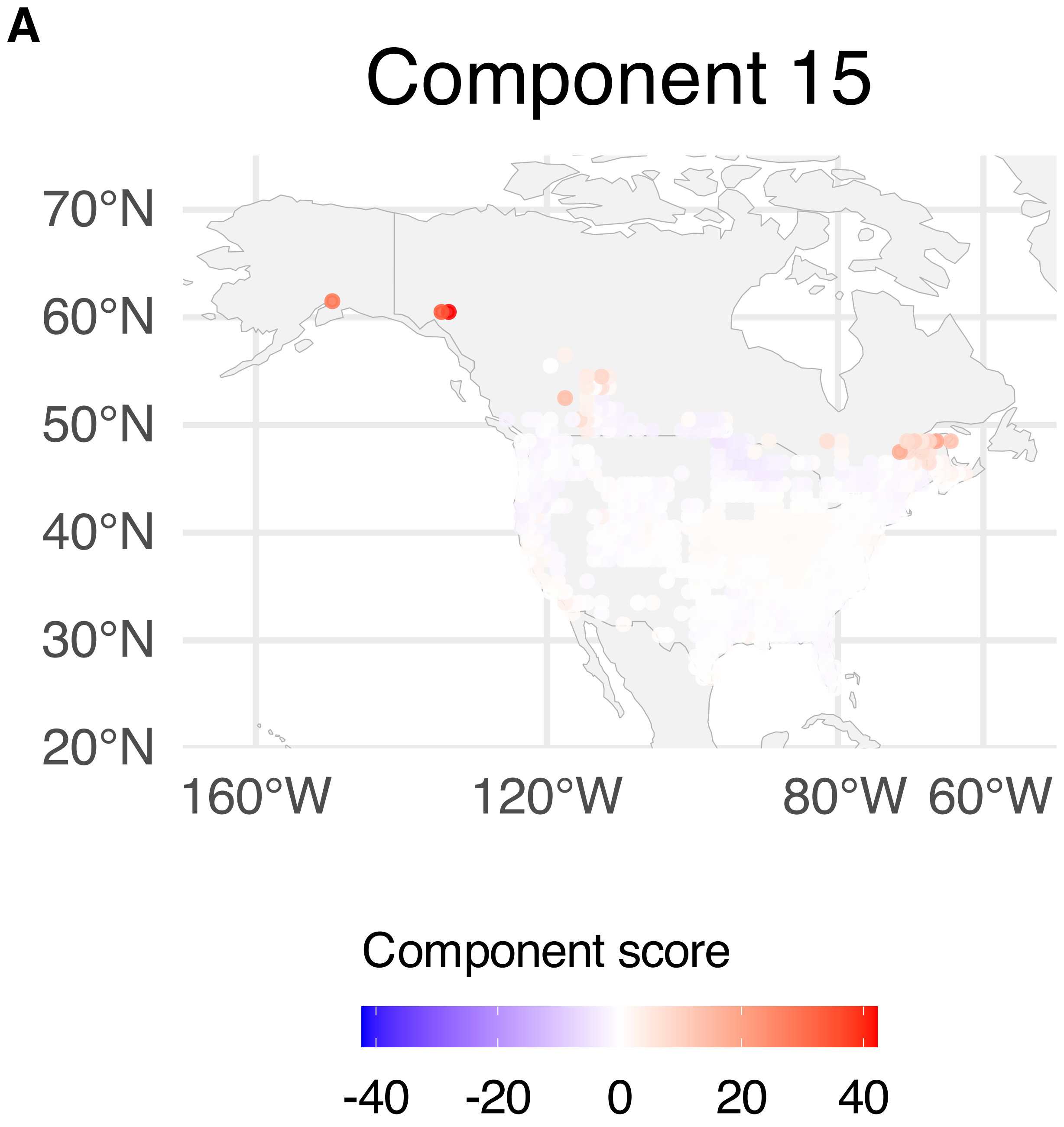


**Supplementary Figure 320.** sPCA: keepX=126 component scores mapped over BBS sampling locations; scores are averaged over years within spatial grid. Component 15.

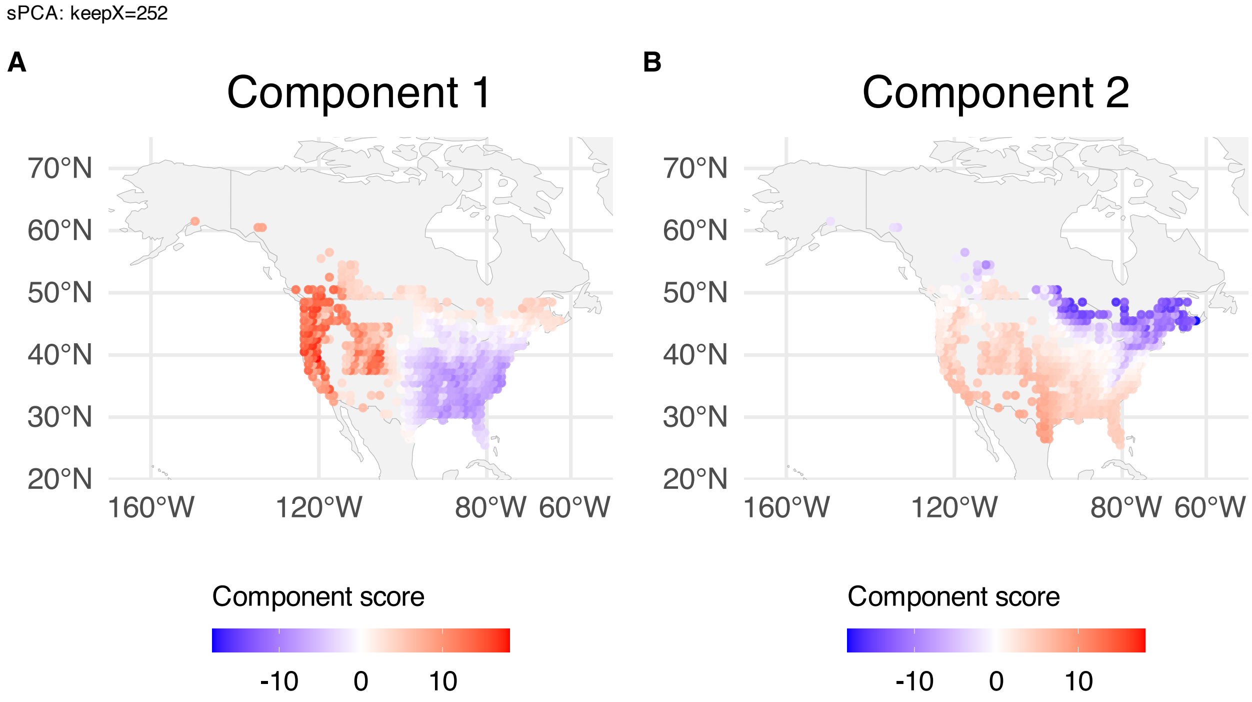


**Supplementary Figure 321.** sPCA: keepX=252 component scores mapped over BBS sampling locations; scores are averaged over years within spatial grid. (A) Component 1. (B) Component 2.

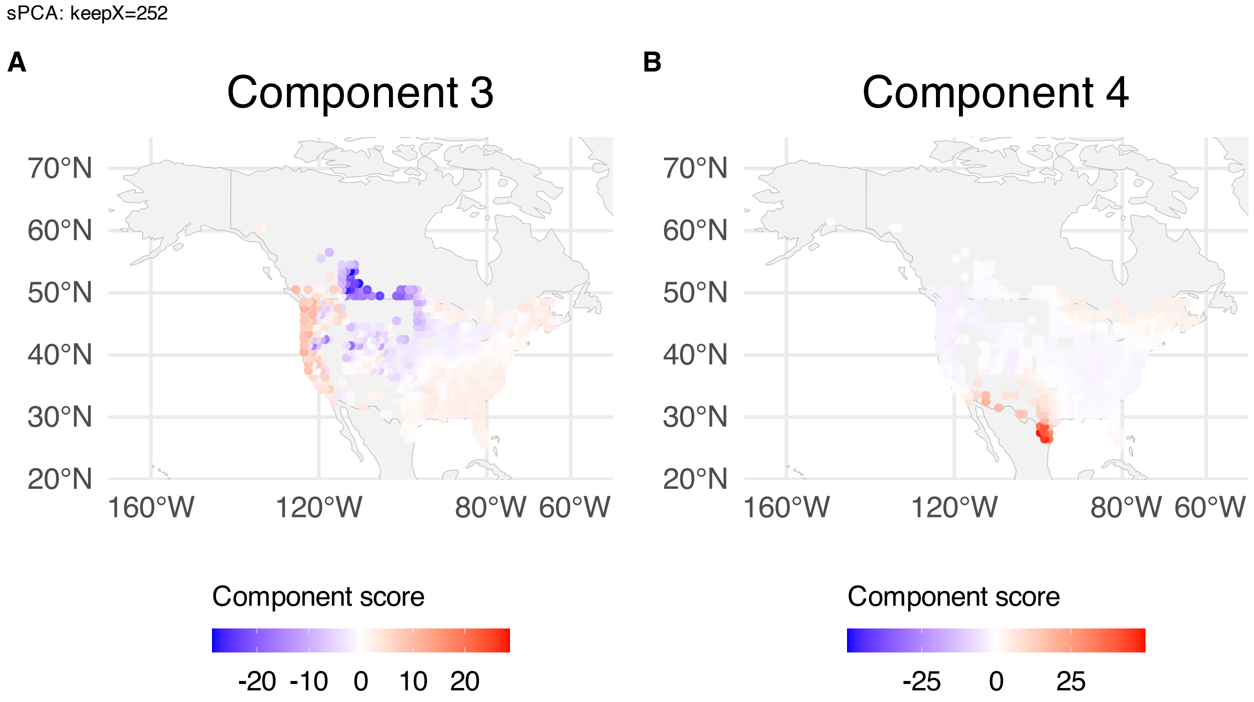


**Supplementary Figure 322.** sPCA: keepX=252 component scores mapped over BBS sampling locations; scores are averaged over years within spatial grid. (A) Component 3. (B) Component 4.

sPCA: keepX=252

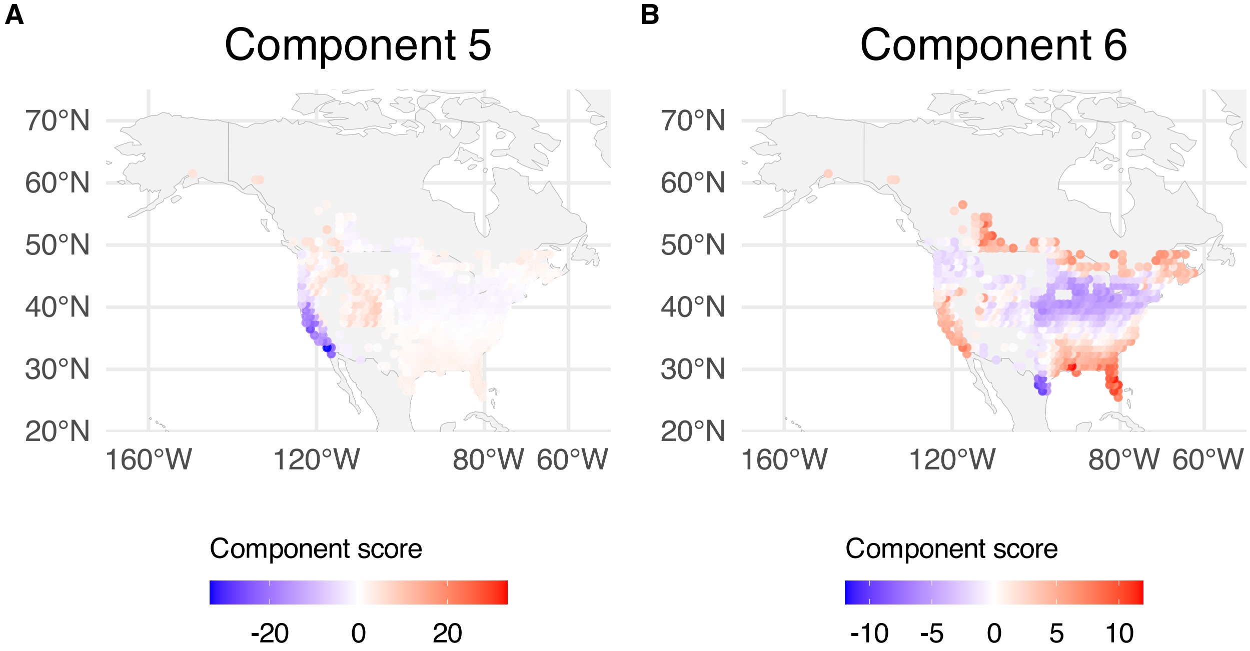


**Supplementary Figure 323.** sPCA: keepX=252 component scores mapped over BBS sampling locations; scores are averaged over years within spatial grid. (A) Component 5. (B) Component 6.

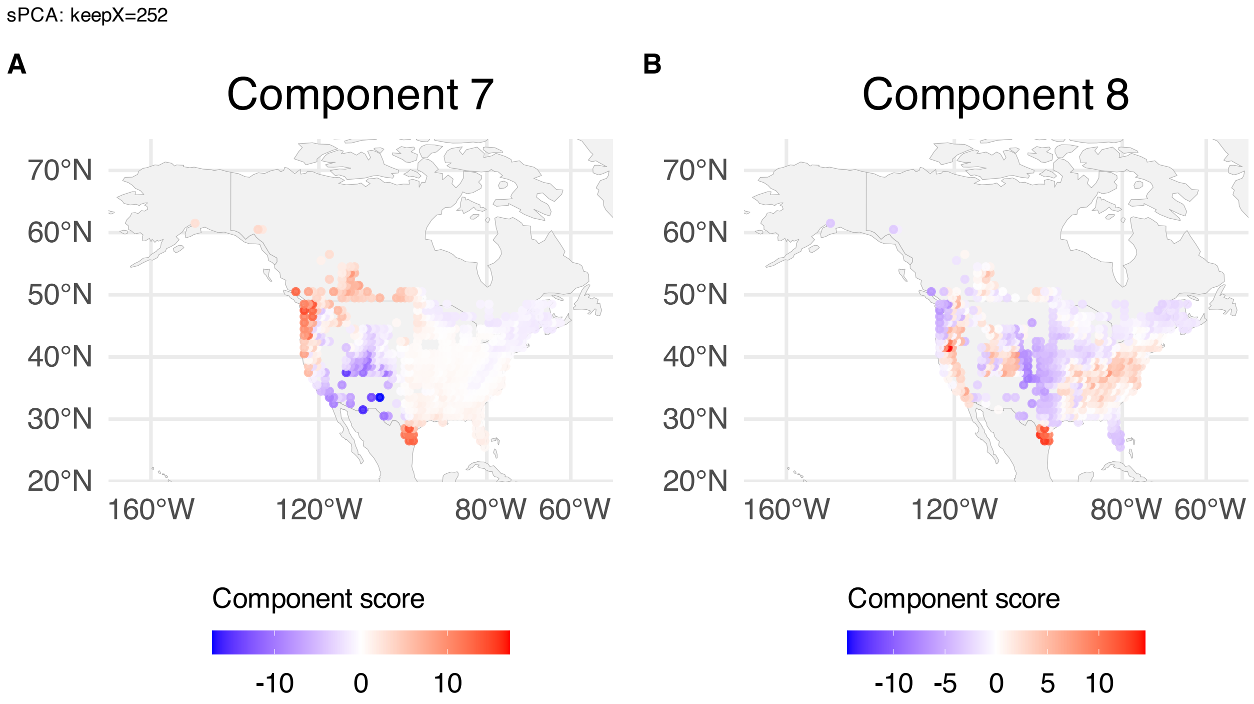


**Supplementary Figure 324.** sPCA: keepX=252 component scores mapped over BBS sampling locations; scores are averaged over years within spatial grid. (A) Component 7. (B) Component 8.

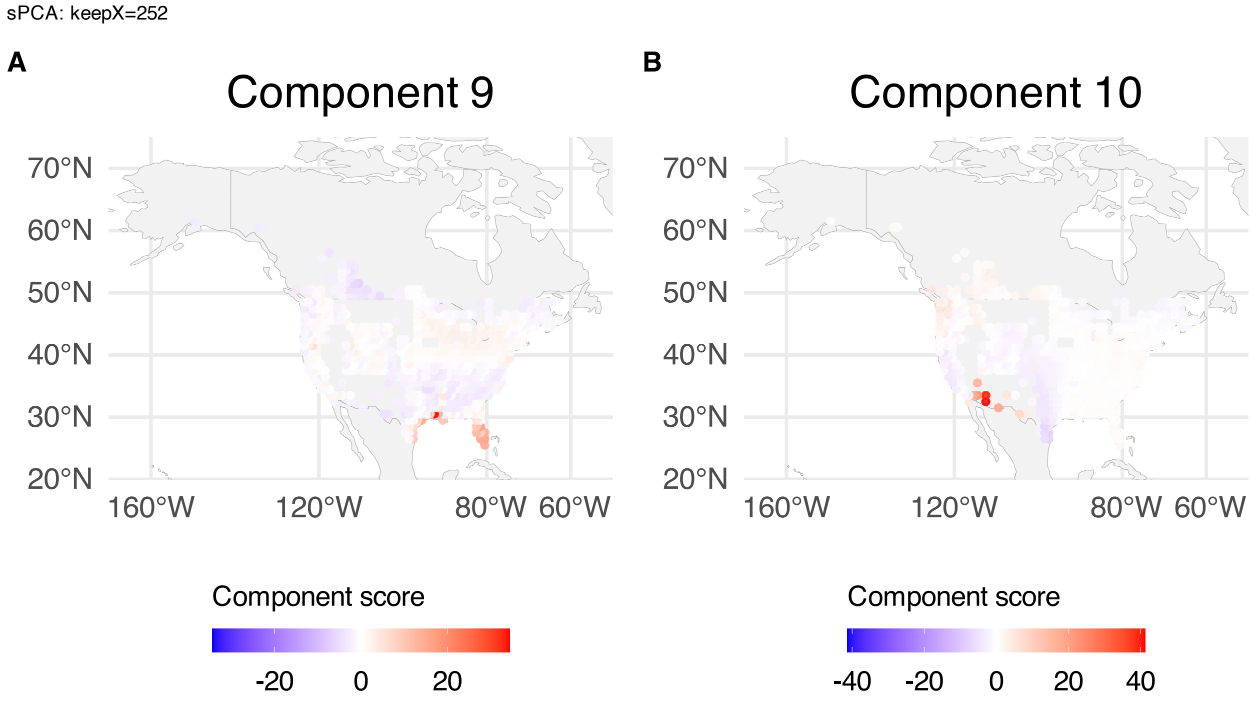


**Supplementary Figure 325.** sPCA: keepX=252 component scores mapped over BBS sampling locations; scores are averaged over years within spatial grid. (A) Component 9. (B) Component 10.

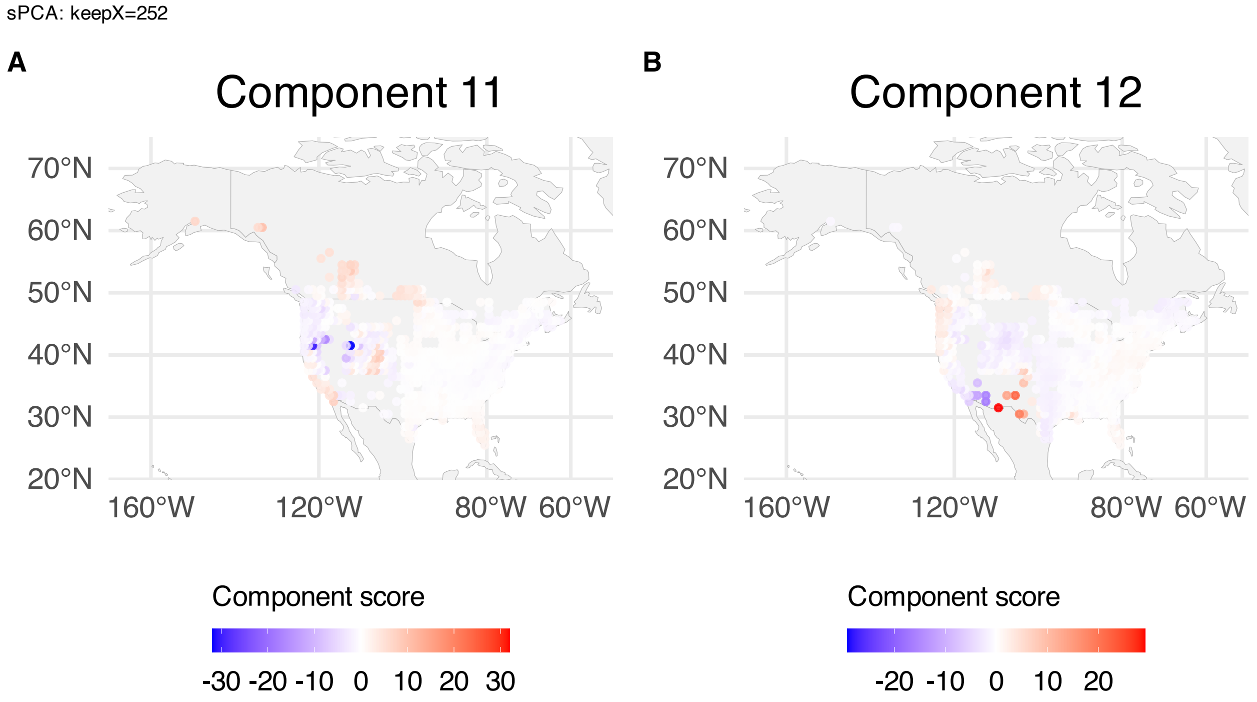


**Supplementary Figure 326.** sPCA: keepX=252 component scores mapped over BBS sampling locations; scores are averaged over years within spatial grid. (A) Component 11. (B) Component 12.

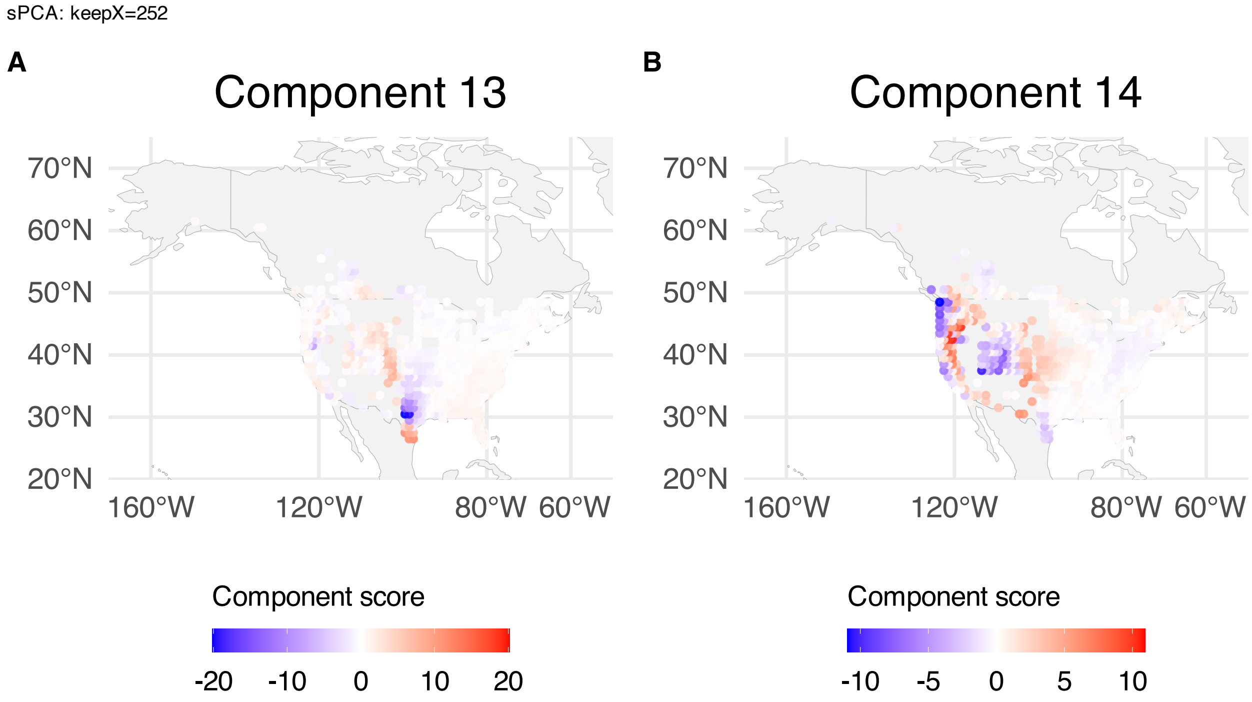


**Supplementary Figure 327.** sPCA: keepX=252 component scores mapped over BBS sampling locations; scores are averaged over years within spatial grid. (A) Component 13. (B) Component 14.

sPCA: keepX=252

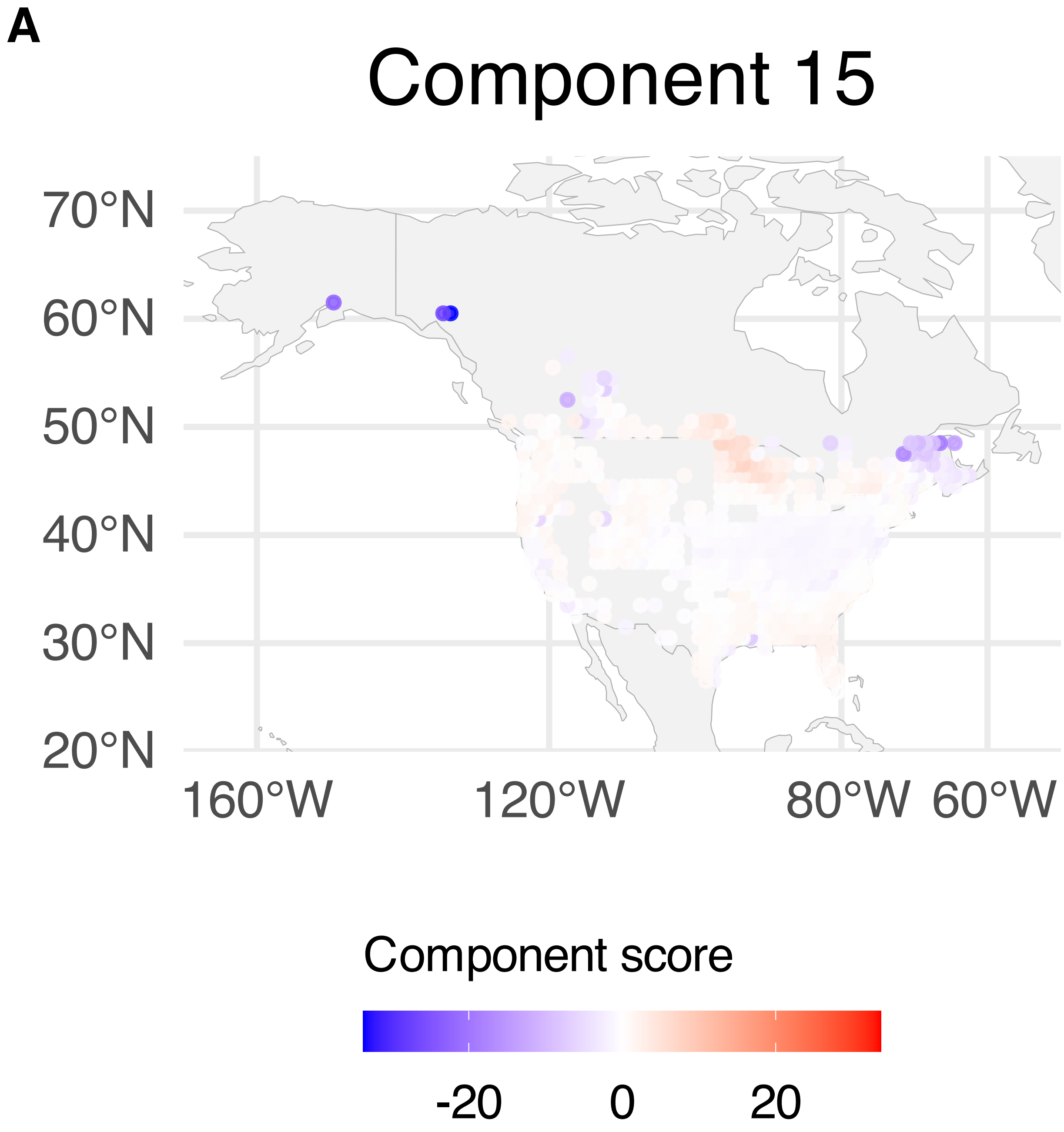


**Supplementary Figure 328.** sPCA: keepX=252 component scores mapped over BBS sampling locations; scores are averaged over years within spatial grid. Component 15.
