## Supplementary Note 5 : Multiple Myeloma for "A statistical framework for system-level synthesis"

This note presents the supplementary results for the analysis of purified Plasma cell data from the multiple myeloma progression study. Given the smaller sample size, only a factorization analysis was done.
The figures include:

- Processes/component functions learned by SSANOVA-SVD (**Supplementary Fig. 177**)
- Heatmaps of the top 50 features with highest absolute loadings (**Supplementary Fig. 178**), gene set enrichment for the top loading features (**Supplementary Fig. 179**)
- Decomposition of the process (**Supplementary Fig. 180**). Because we have a single discrete covariate, the decomposition of the process is identical to the process itself.
- Components from a representative class of matrix factorization methods (**Supplementary Fig. 181**-**Supplementary Fig. 221**)

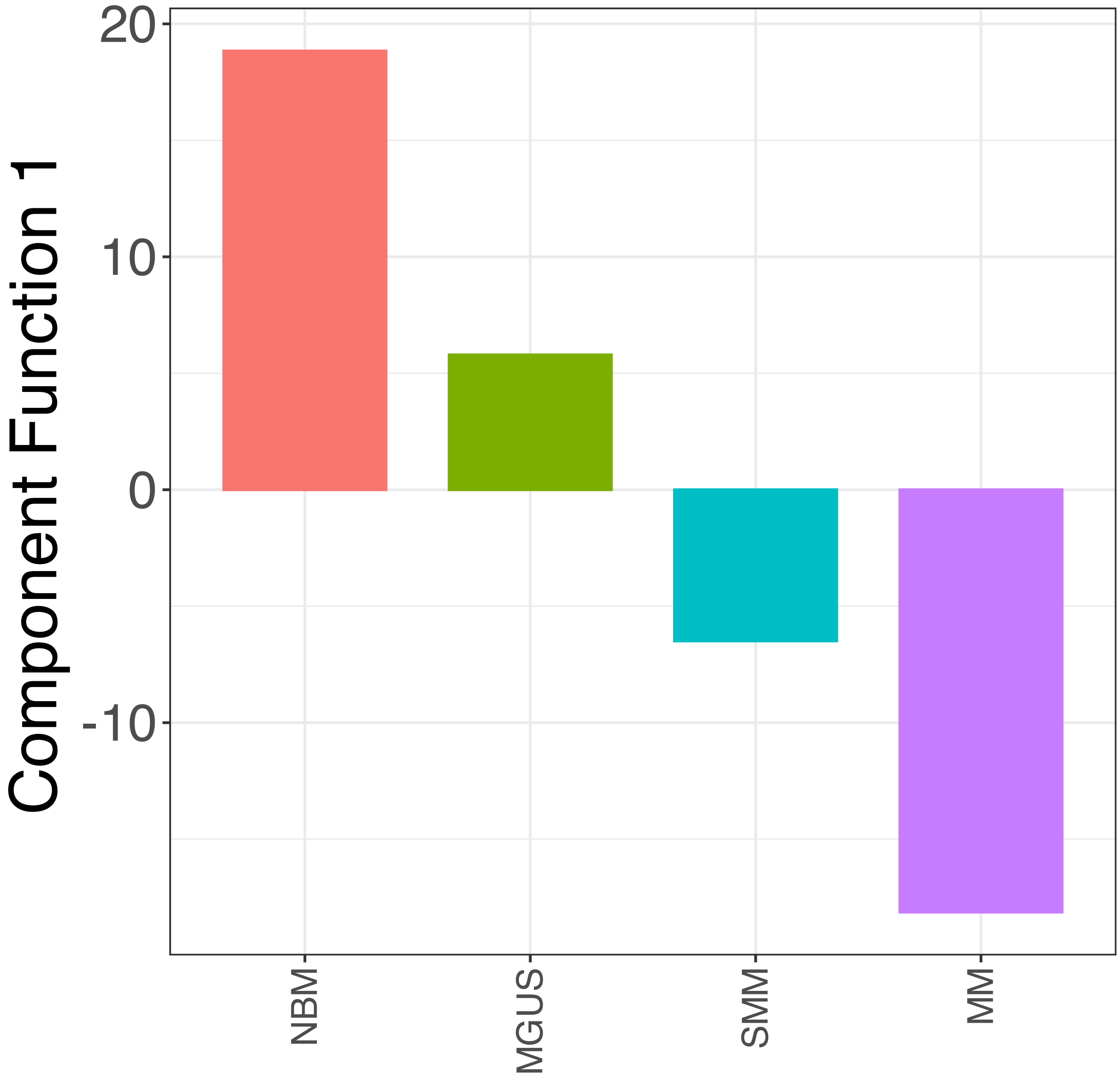


**Supplementary Figure 177.** SSANOVA-SVD reconstructed component functions.Component 1.

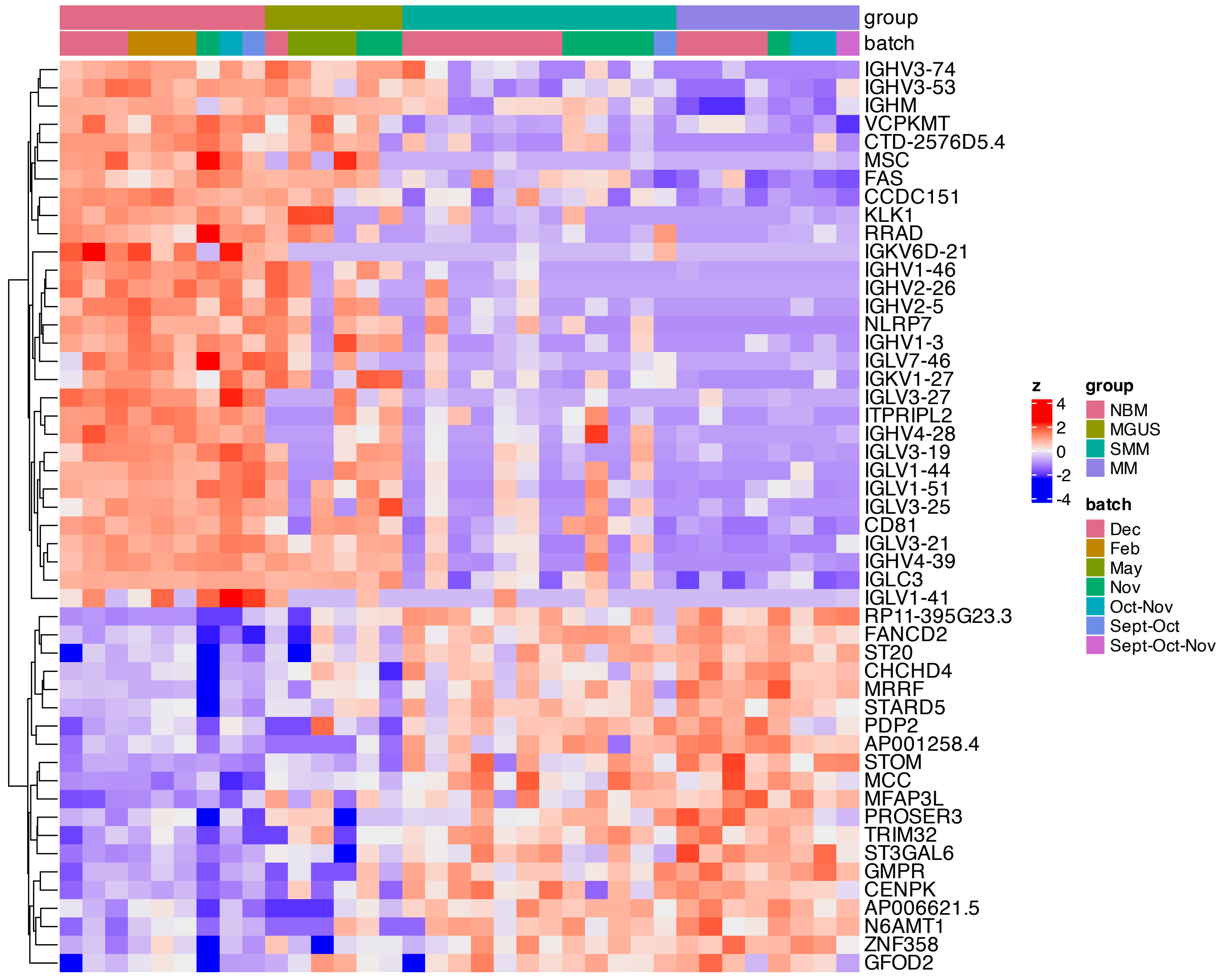


**Supplementary Figure 178.** Heatmap of features with the largest absolute loadings for SSANOVA-SVD component 1.

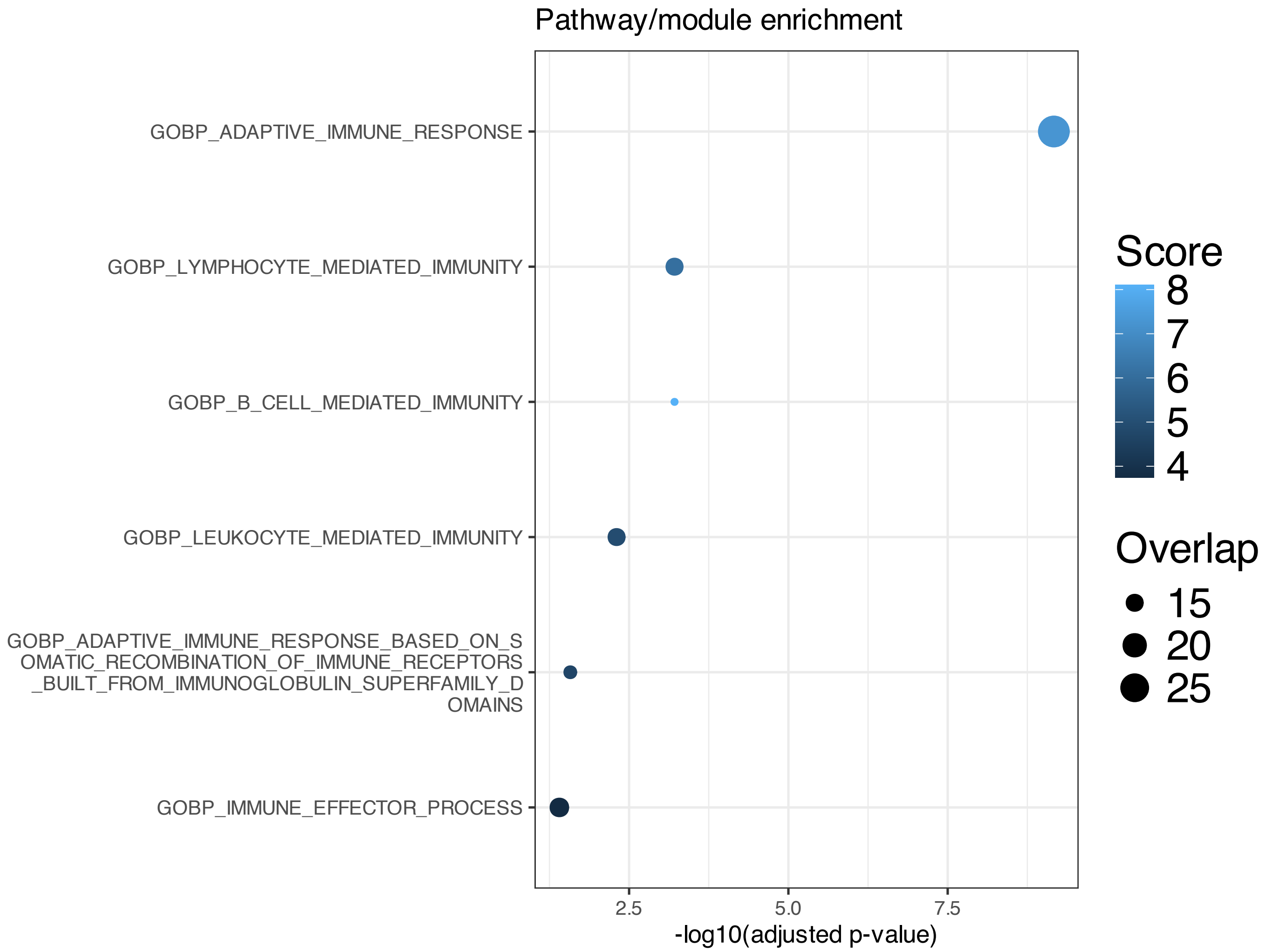


**Supplementary Figure 179.** Enrichment based on features with the largest absolute loadings for SSANOVA-SVD component 1; only terms with adjusted P $\leq 0.1$ are shown.

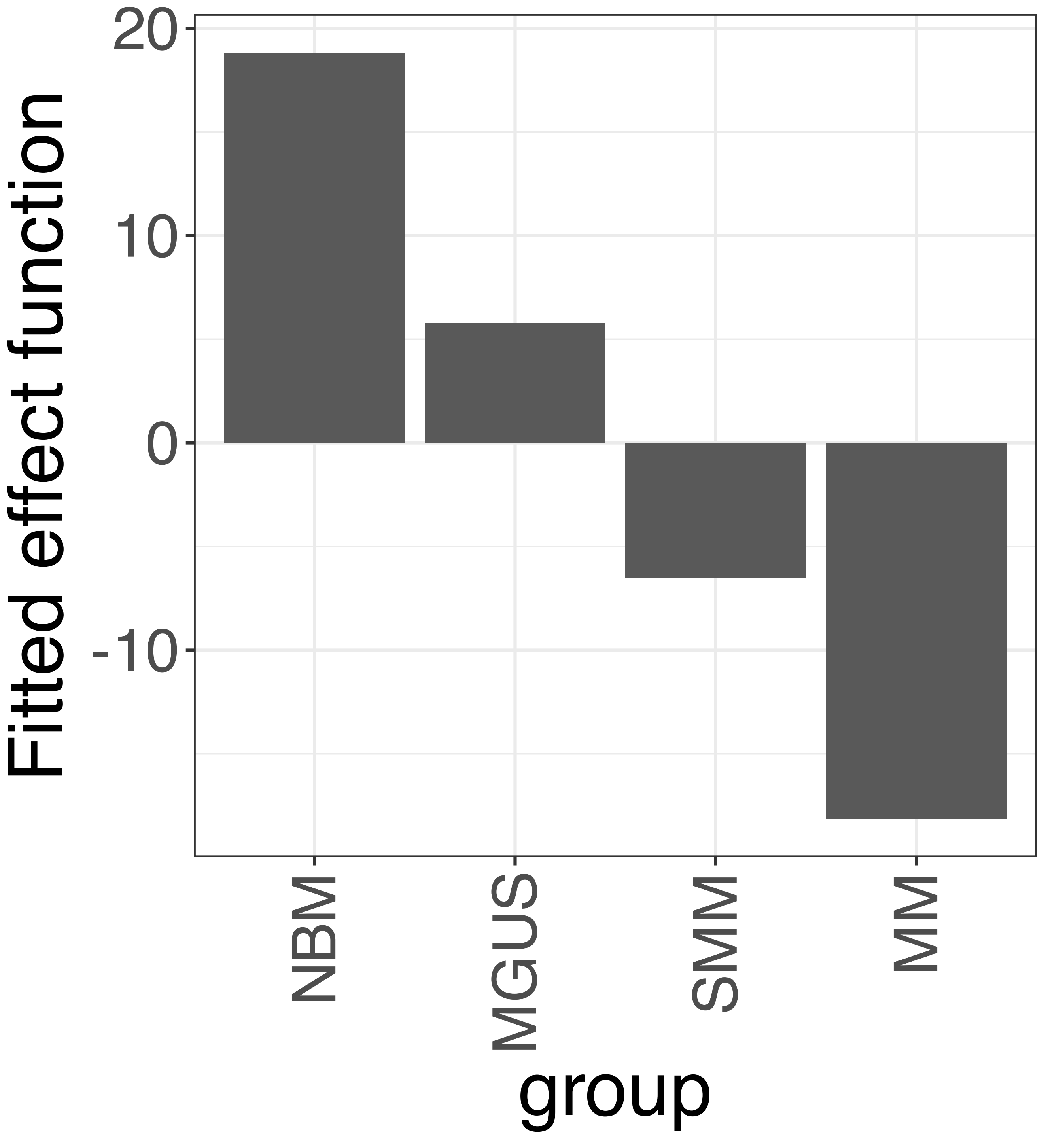


**Supplementary Figure 180.** Decomposition of reconstructed SSANOVA-SVD component function 1.

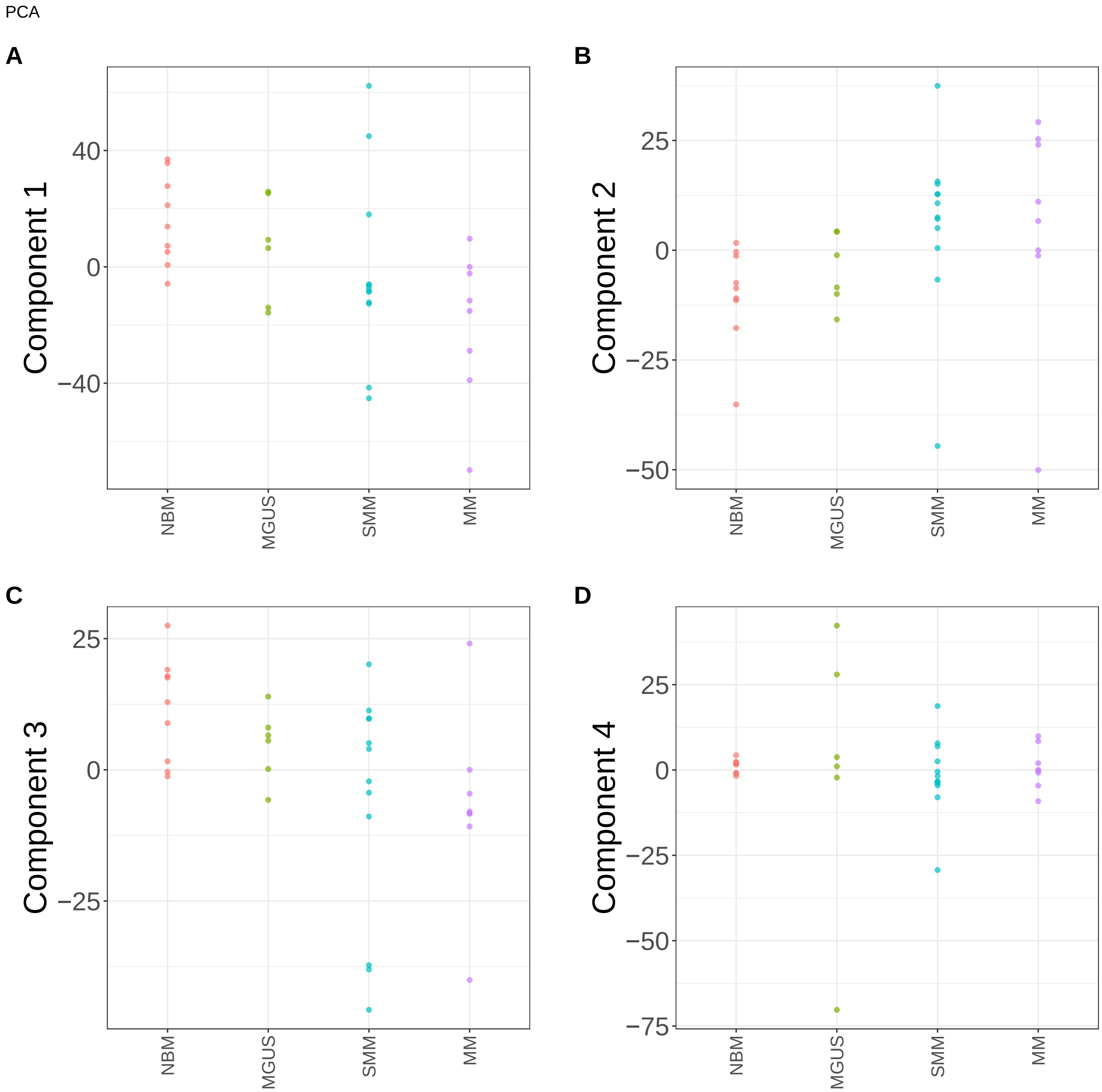


**Supplementary Figure 181.** Components for PCA. (A) Component 1. (B) Component 2. (C) Component 3. (D) Component 4.

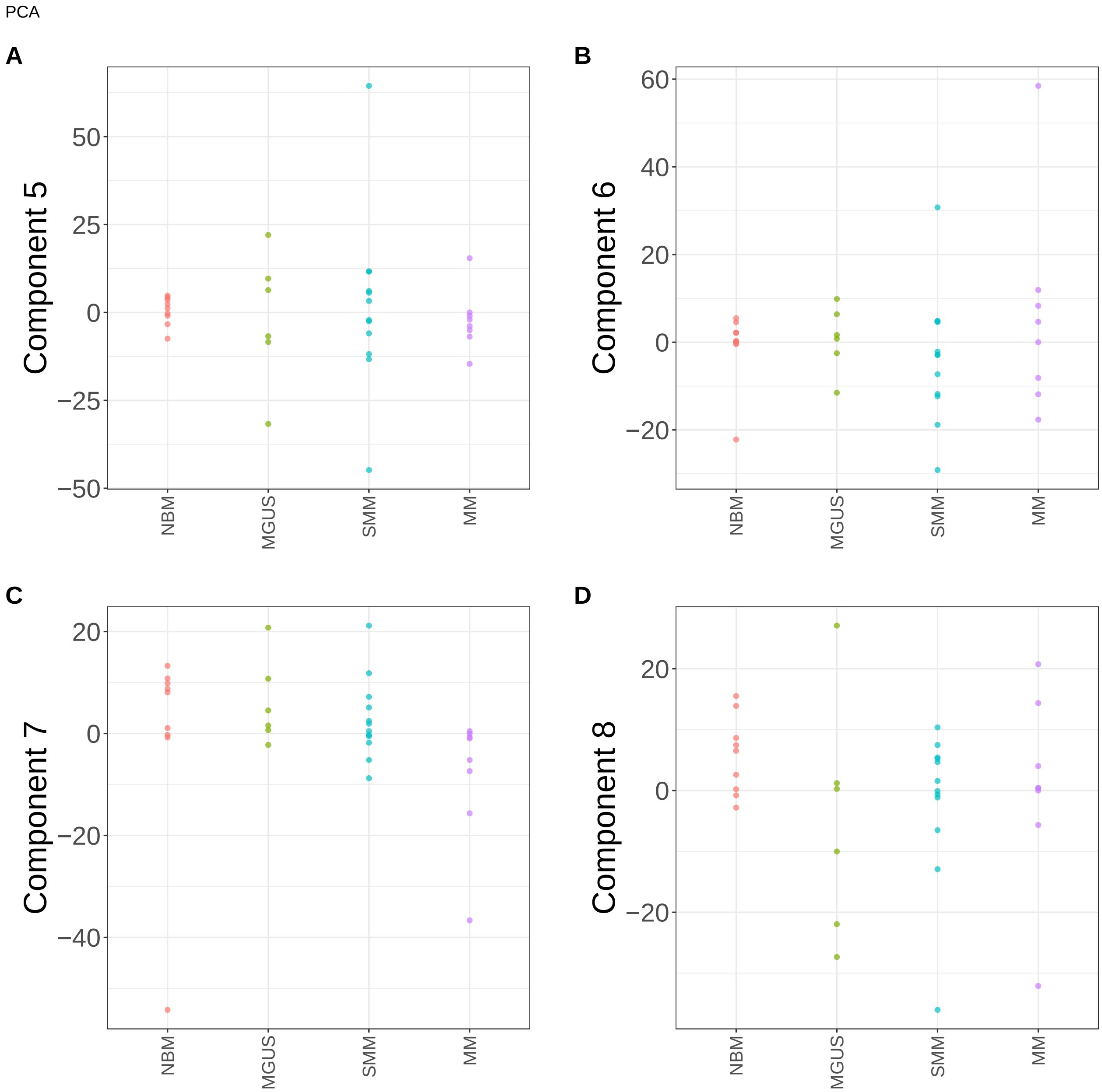


**Supplementary Figure 182.** Components for PCA. (A) Component 5. (B) Component 6. (C) Component 7. (D) Component 8.

PCA

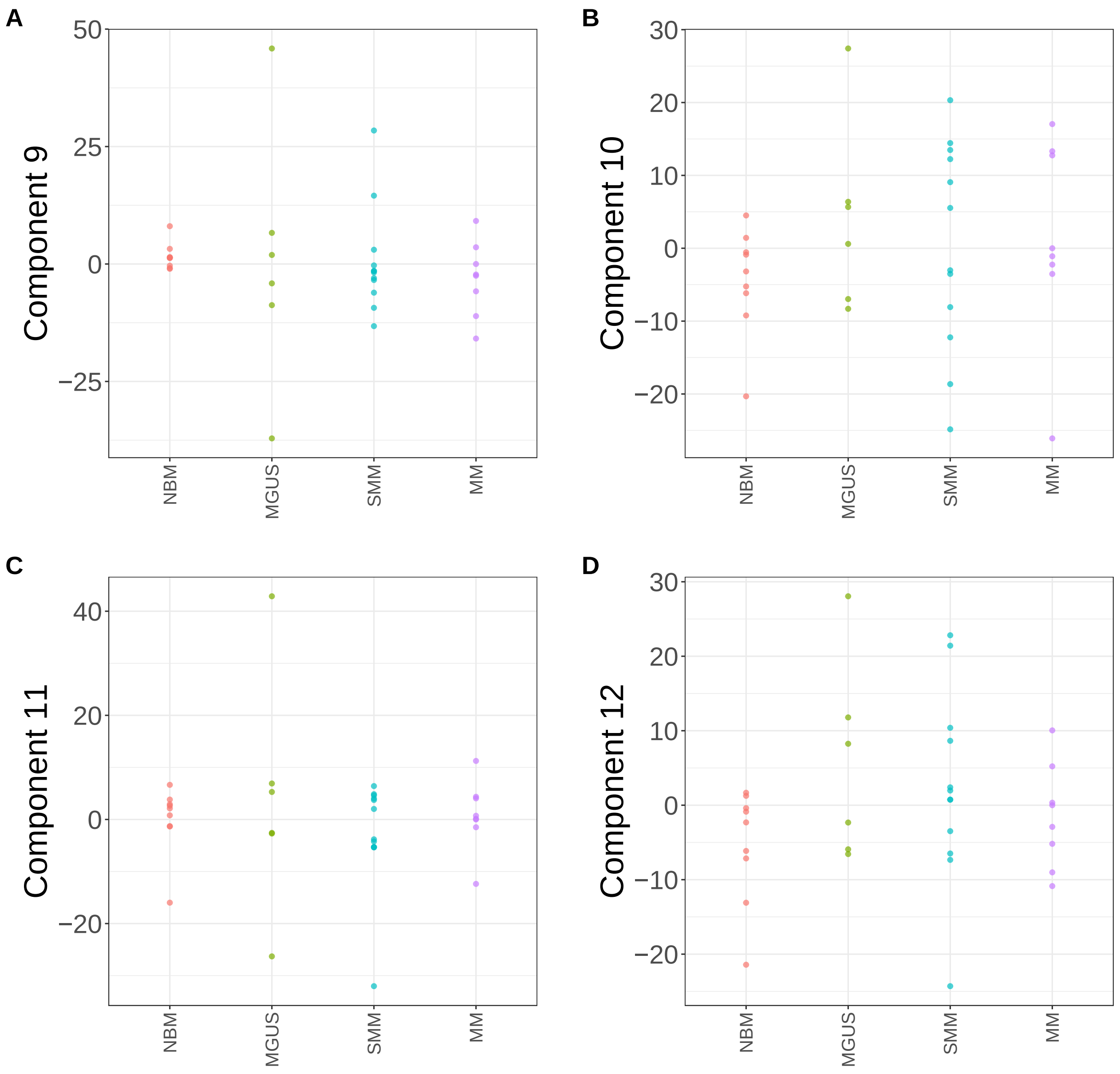


**Supplementary Figure 183.** Components for PCA. (A) Component 9. (B) Component 10. (C) Component 11. (D) Component 12.

PCA

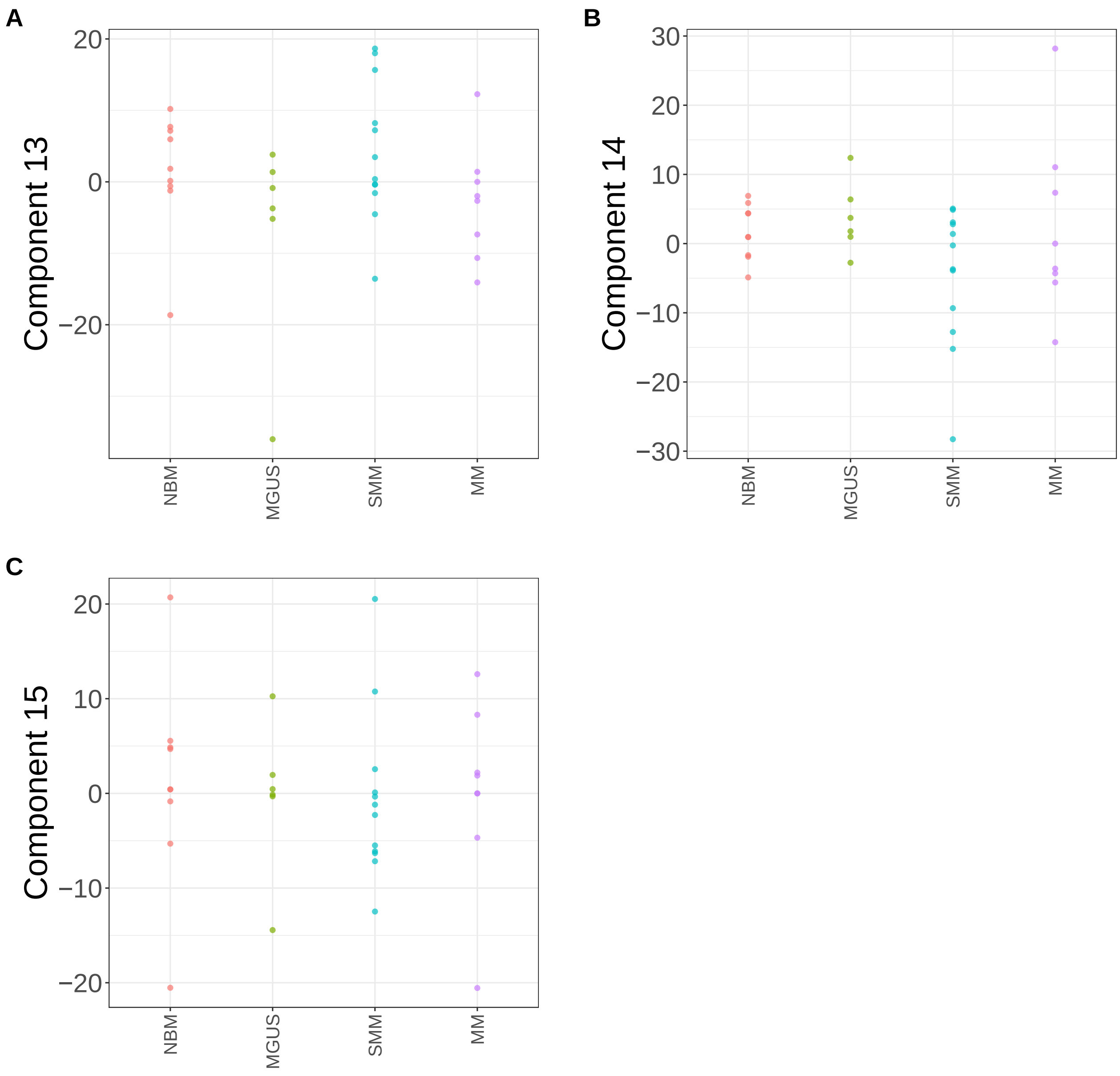


**Supplementary Figure 184.** Components for PCA. (A) Component 13. (B) Component 14. (C) Component 15.

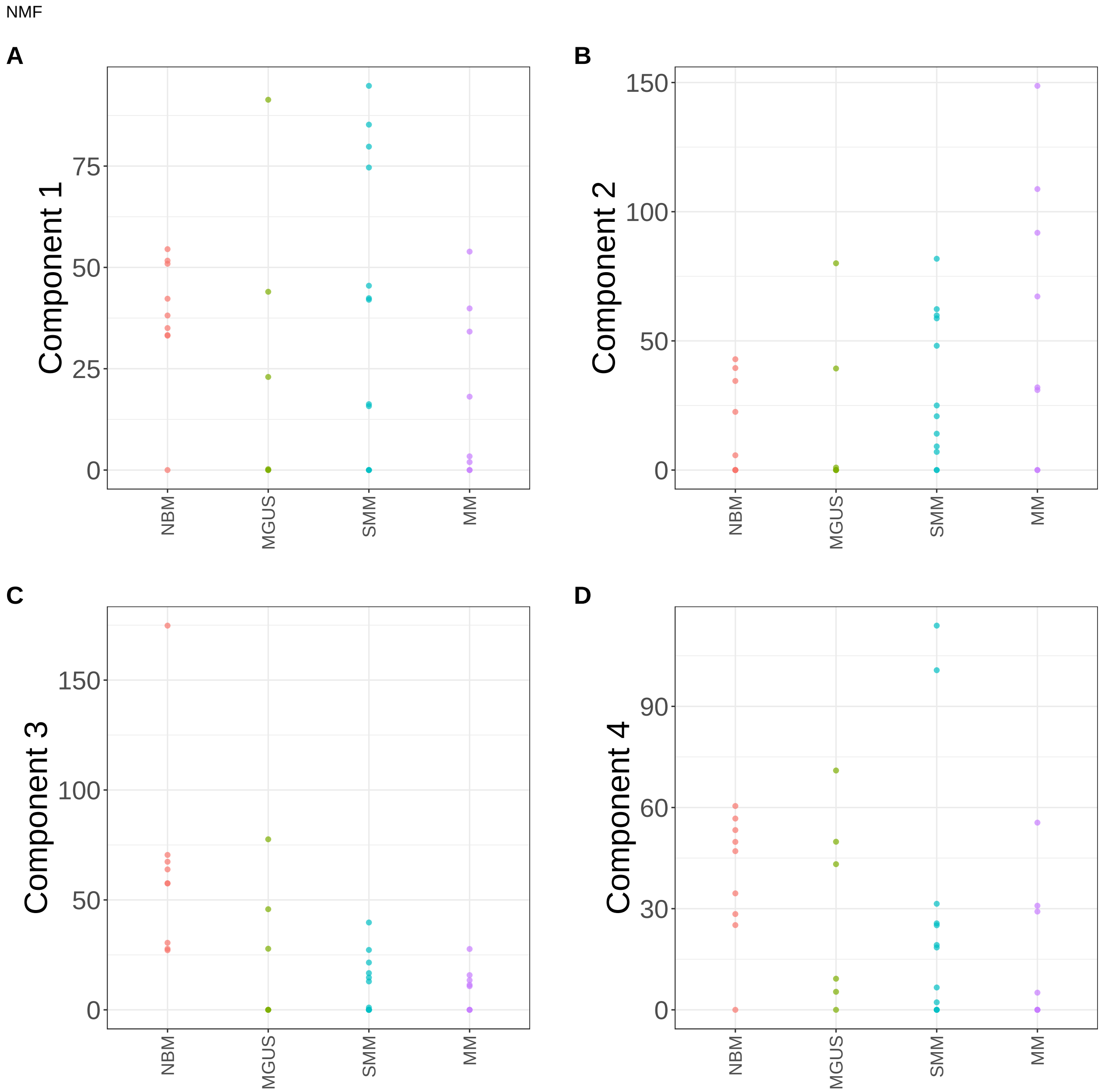


**Supplementary Figure 185.** Components for NMF. (A) Component 1. (B) Component 2. (C) Component 3. (D) Component 4.

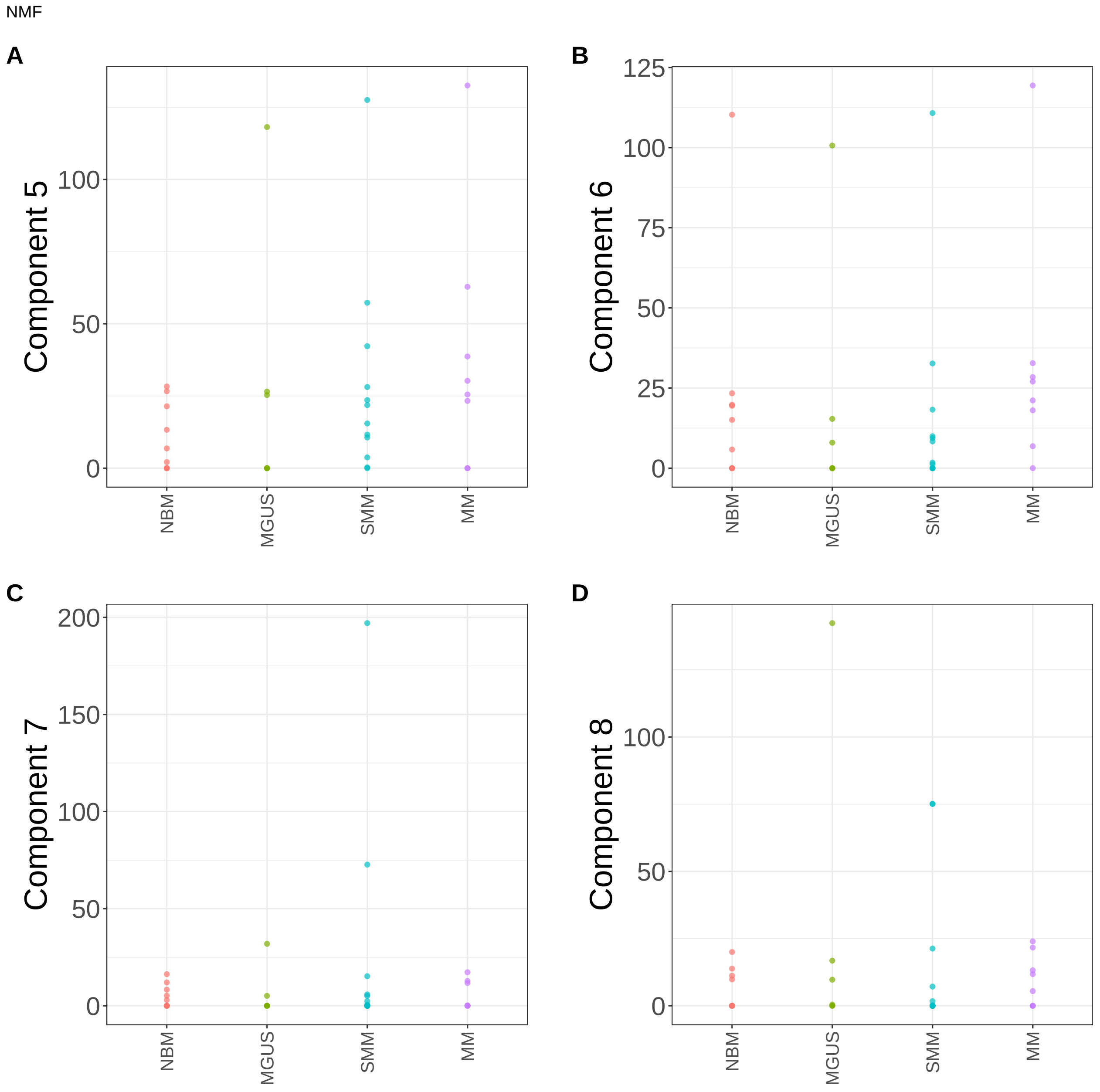


**Supplementary Figure 186.** Components for NMF. (A) Component 5. (B) Component 6. (C) Component 7. (D) Component 8.

NMF

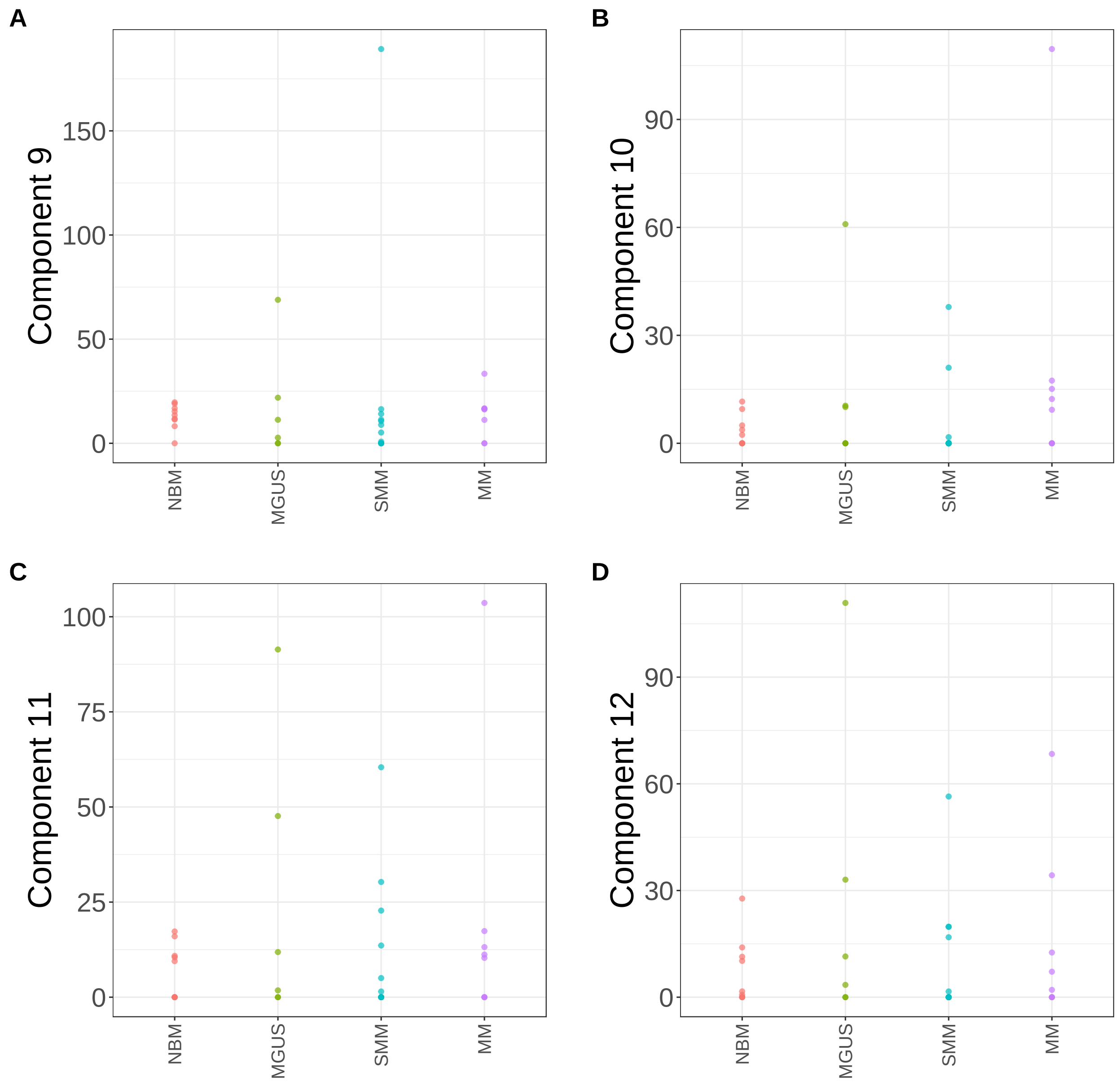


**Supplementary Figure 187.** Components for NMF. (A) Component 9. (B) Component 10. (C) Component 11. (D) Component 12.

NMF

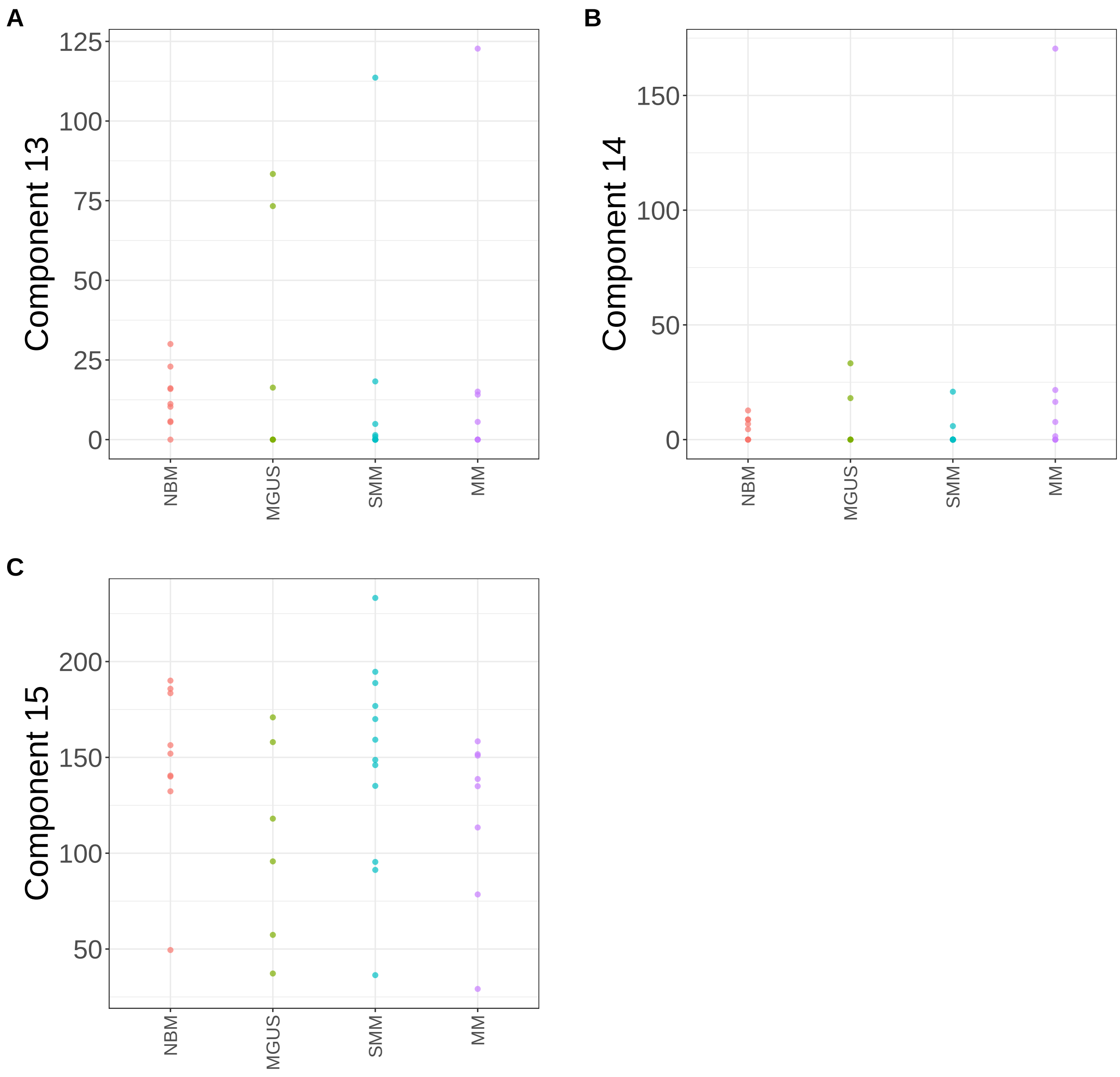


**Supplementary Figure 188.** Components for NMF. (A) Component 13. (B) Component 14. (C) Component 15.

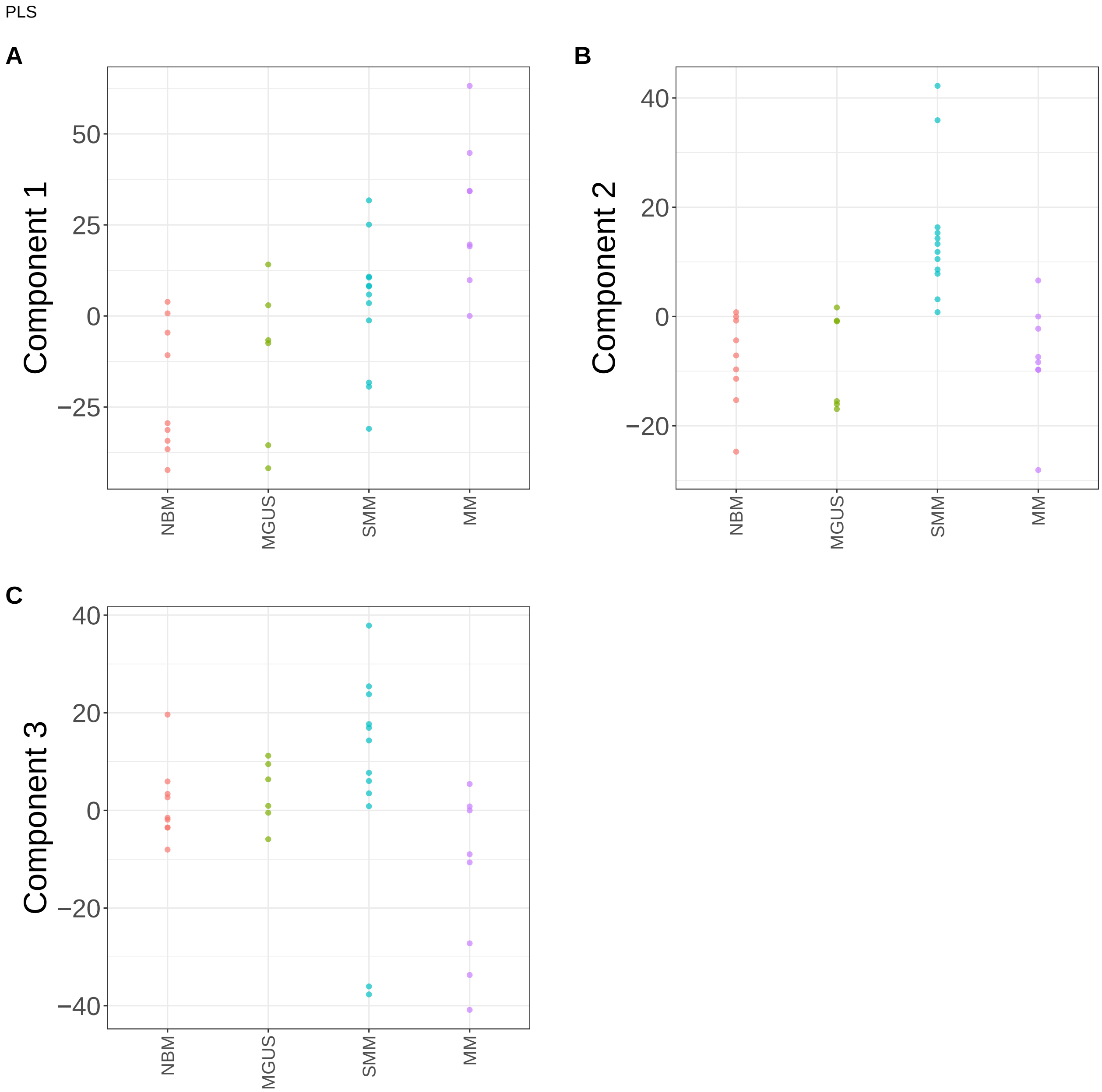


**Supplementary Figure 189.** Components for PLS. (A) Component 1. (B) Component 2. (C) Component 3.

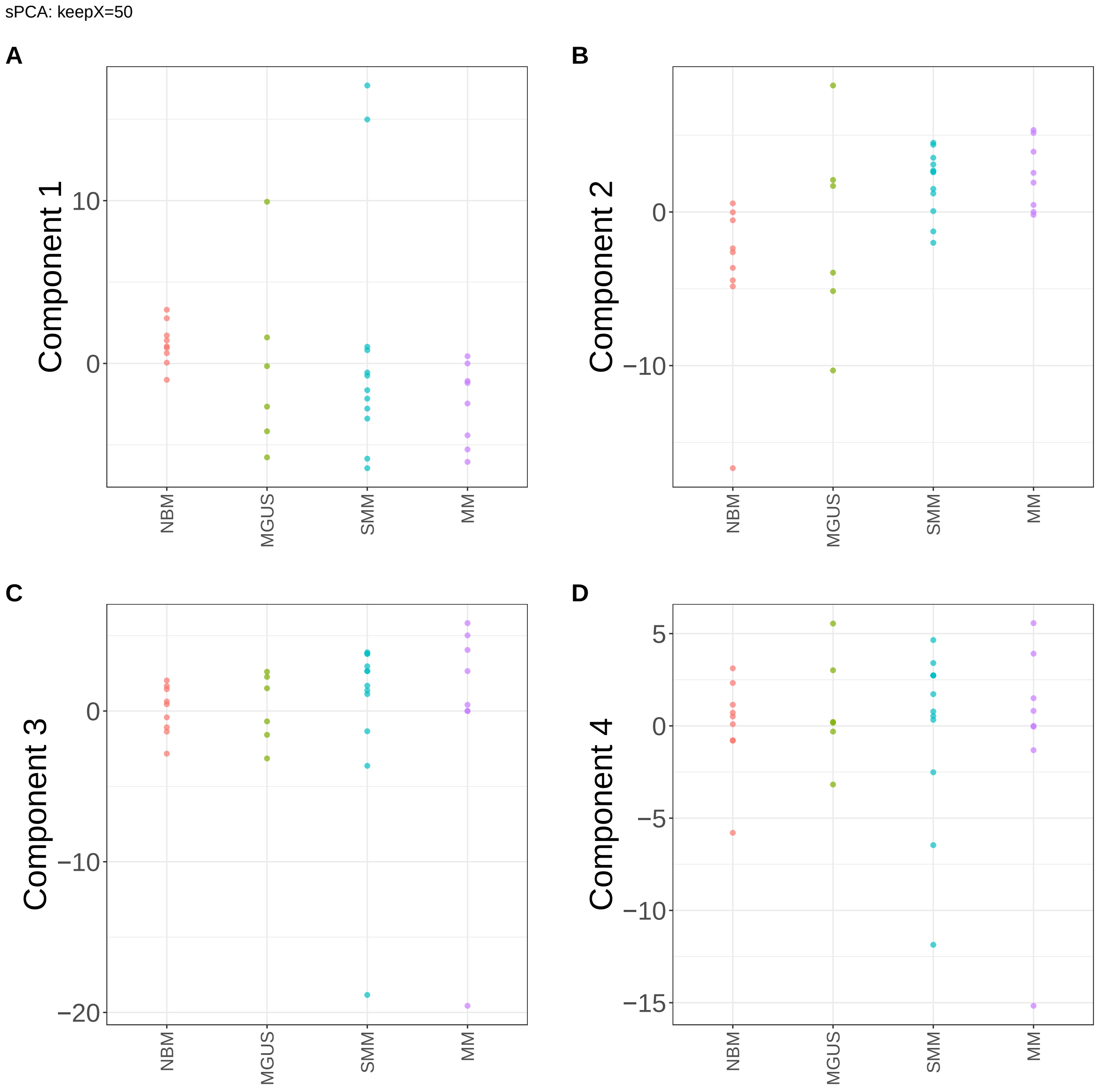


**Supplementary Figure 190.** Components for sPCA: keepX=50. (A) Component 1. (B) Component 2. (C) Component 3. (D) Component 4.

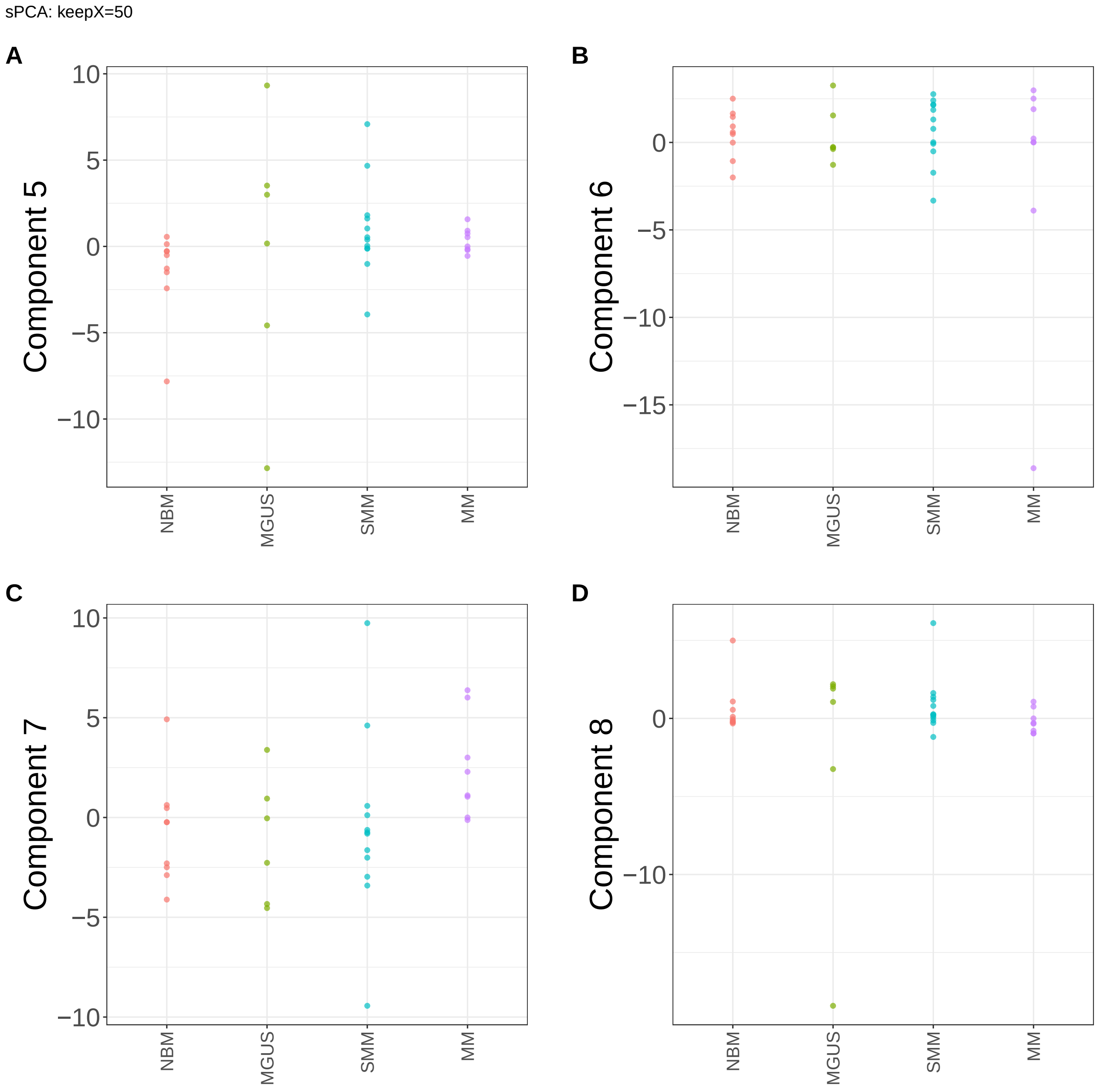


**Supplementary Figure 191.** Components for sPCA: keepX=50. (A) Component 5. (B) Component 6. (C) Component 7. (D) Component 8.

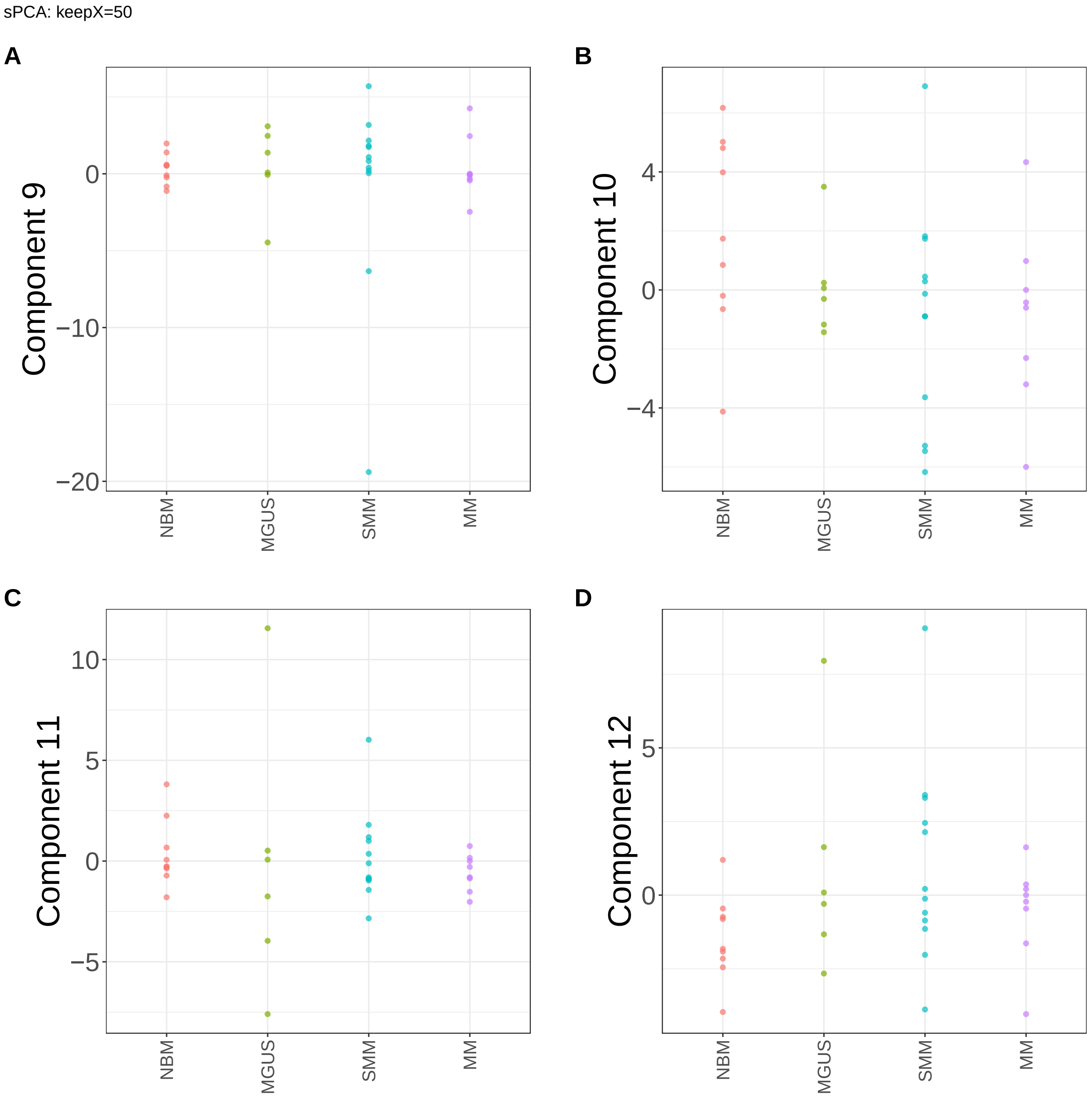


**Supplementary Figure 192.** Components for sPCA: keepX=50. (A) Component 9. (B) Component 10. (C) Component 11. (D) Component 12.

sPCA: keepX=50

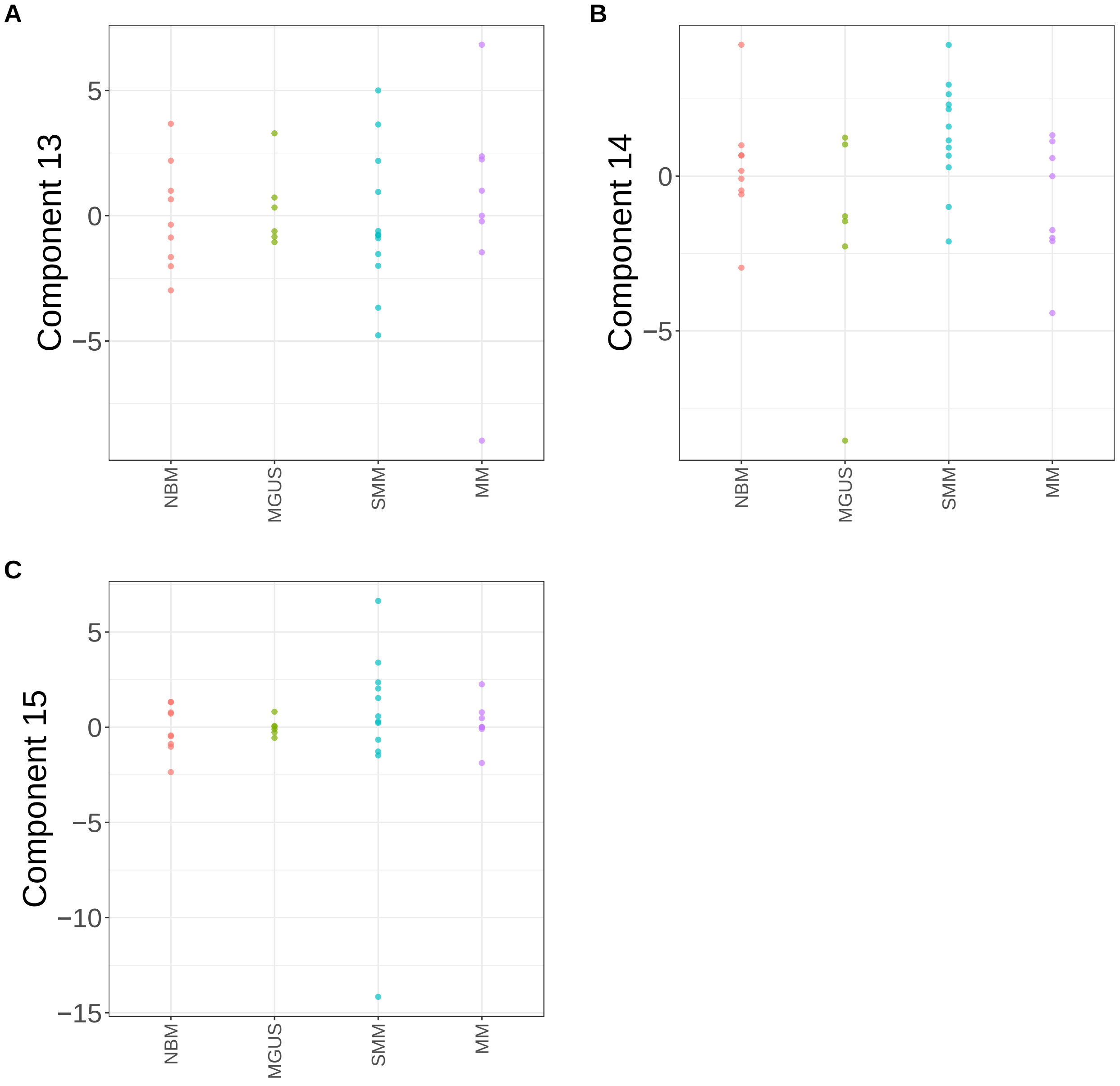


**Supplementary Figure 193.** Components for sPCA: keepX=50. (A) Component 13. (B) Component 14. (C) Component 15.

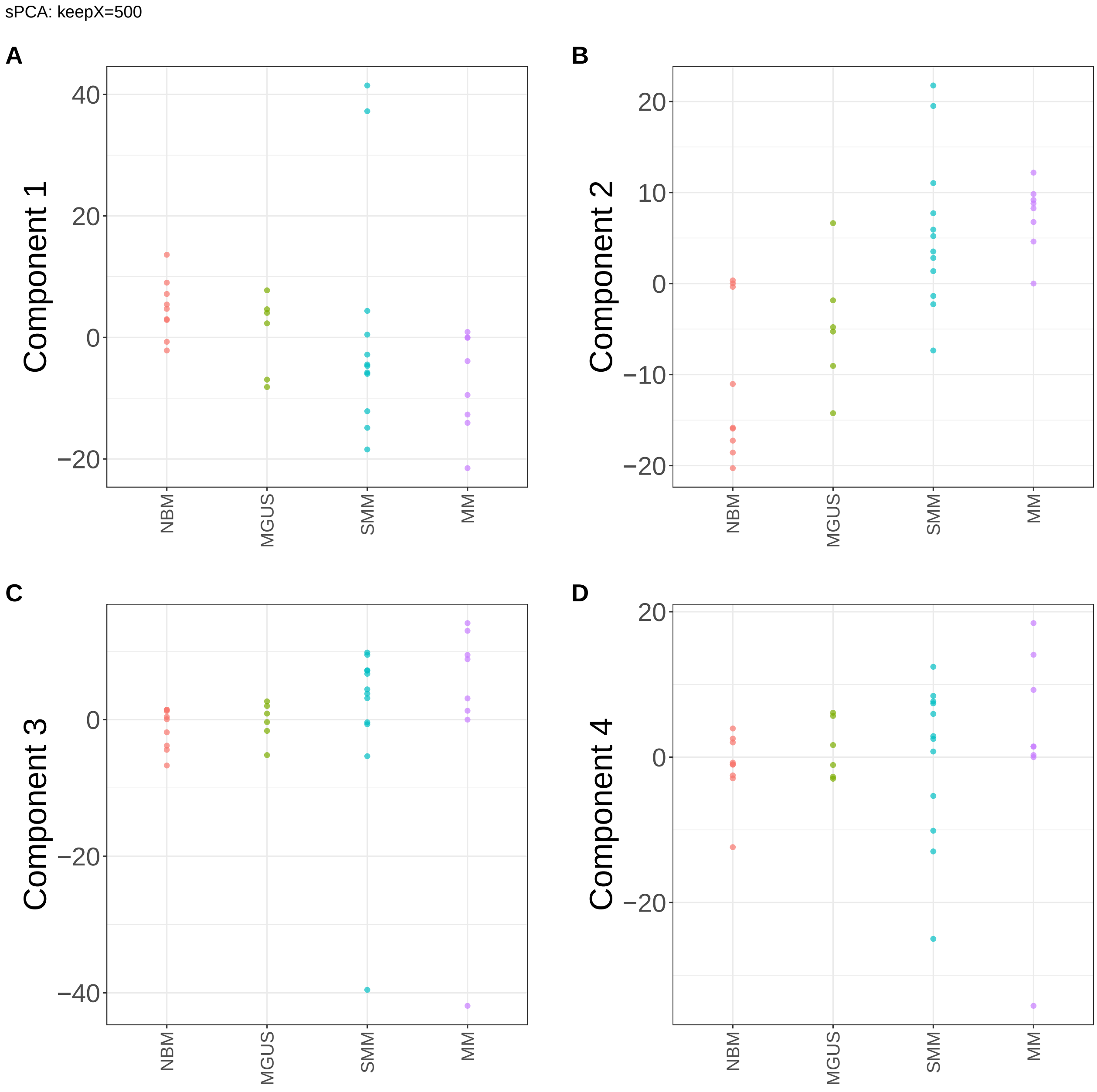


**Supplementary Figure 194.** Components for sPCA: keepX=500. (A) Component 1. (B) Component 2. (C) Component 3. (D) Component 4.

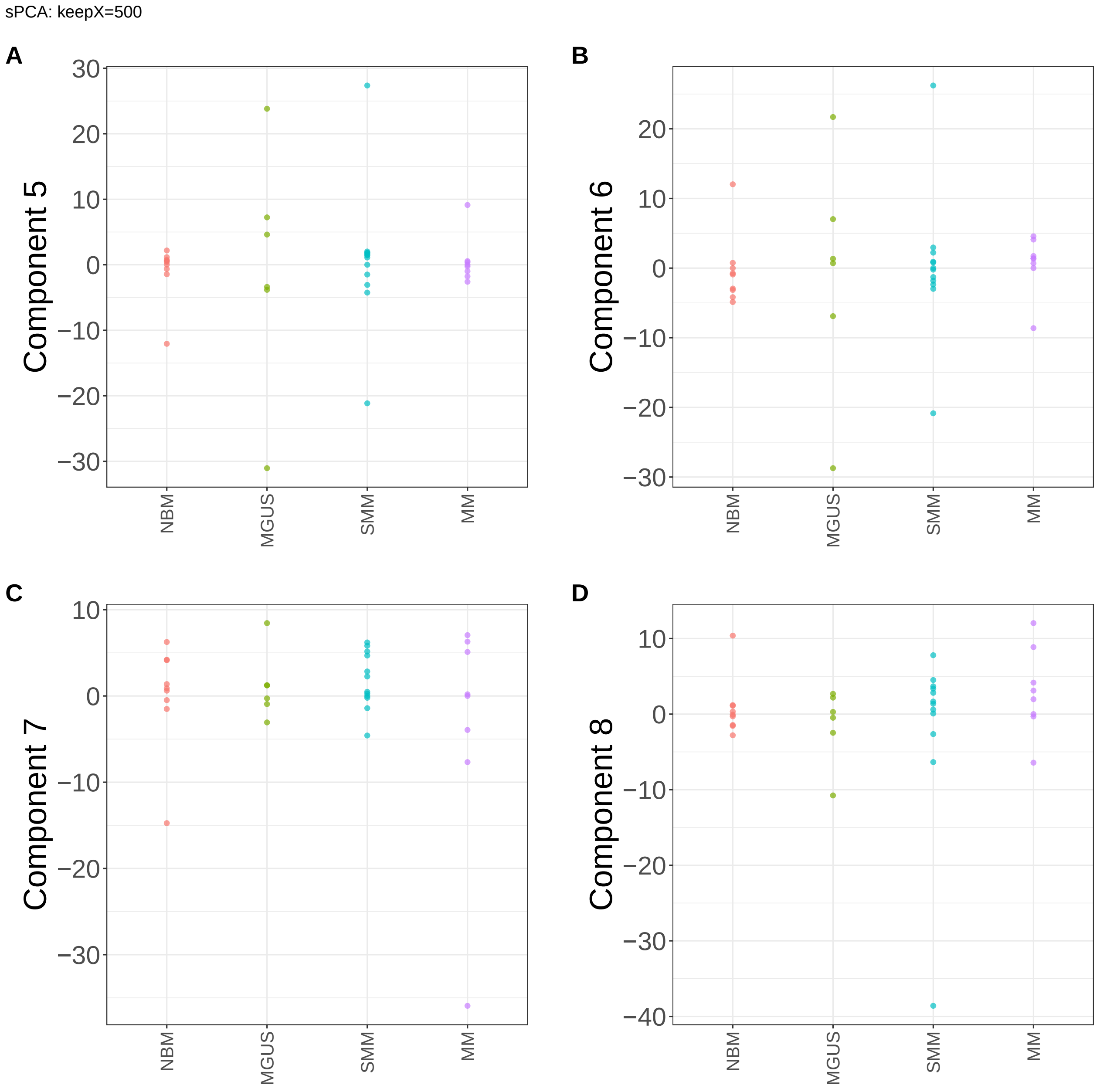


**Supplementary Figure 195.** Components for sPCA: keepX=500. (A) Component 5. (B) Component 6. (C) Component 7. (D) Component 8.

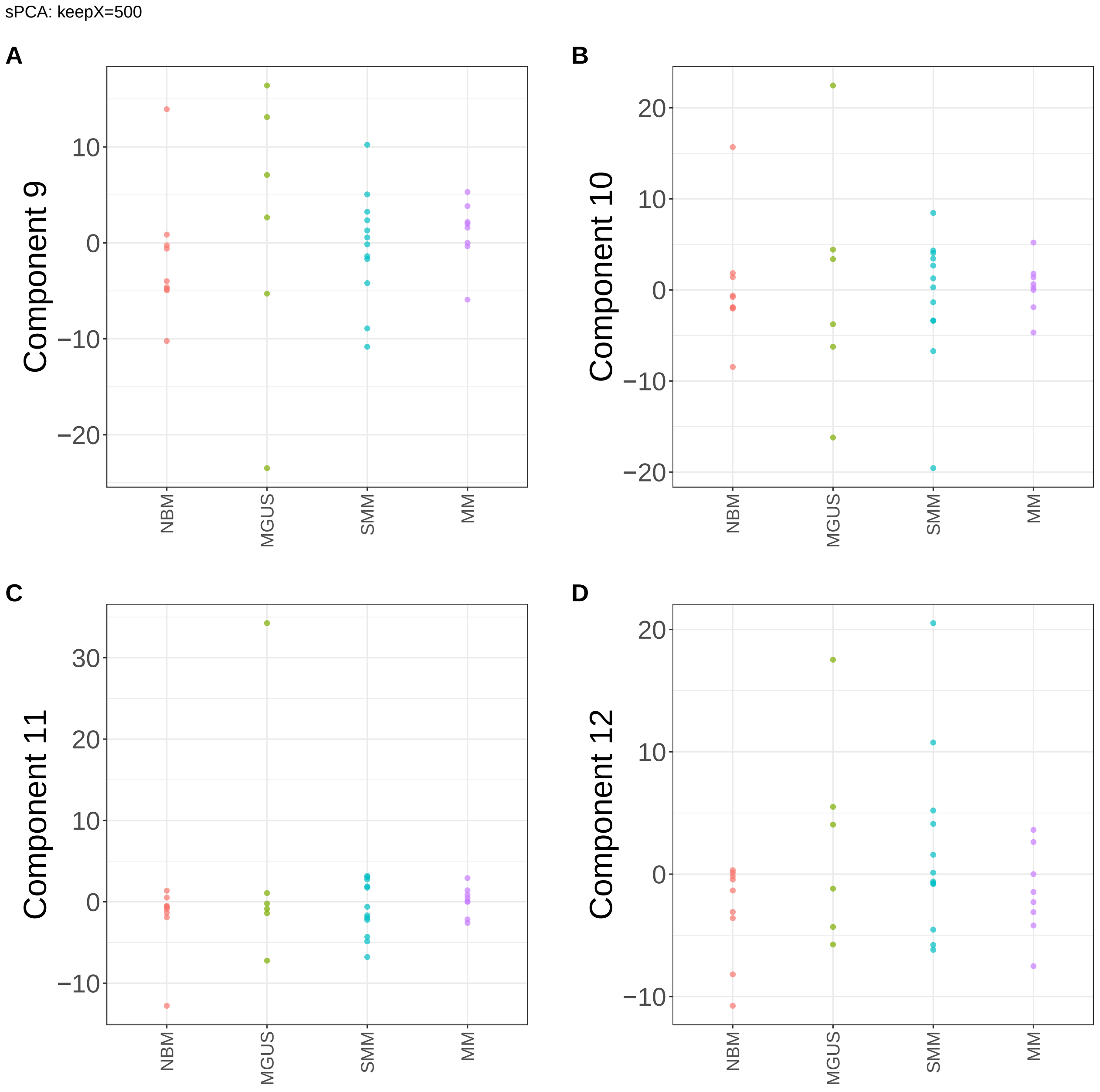


**Supplementary Figure 196.** Components for sPCA: keepX=500. (A) Component 9. (B) Component 10. (C) Component 11. (D) Component 12.

sPCA: keepX=500

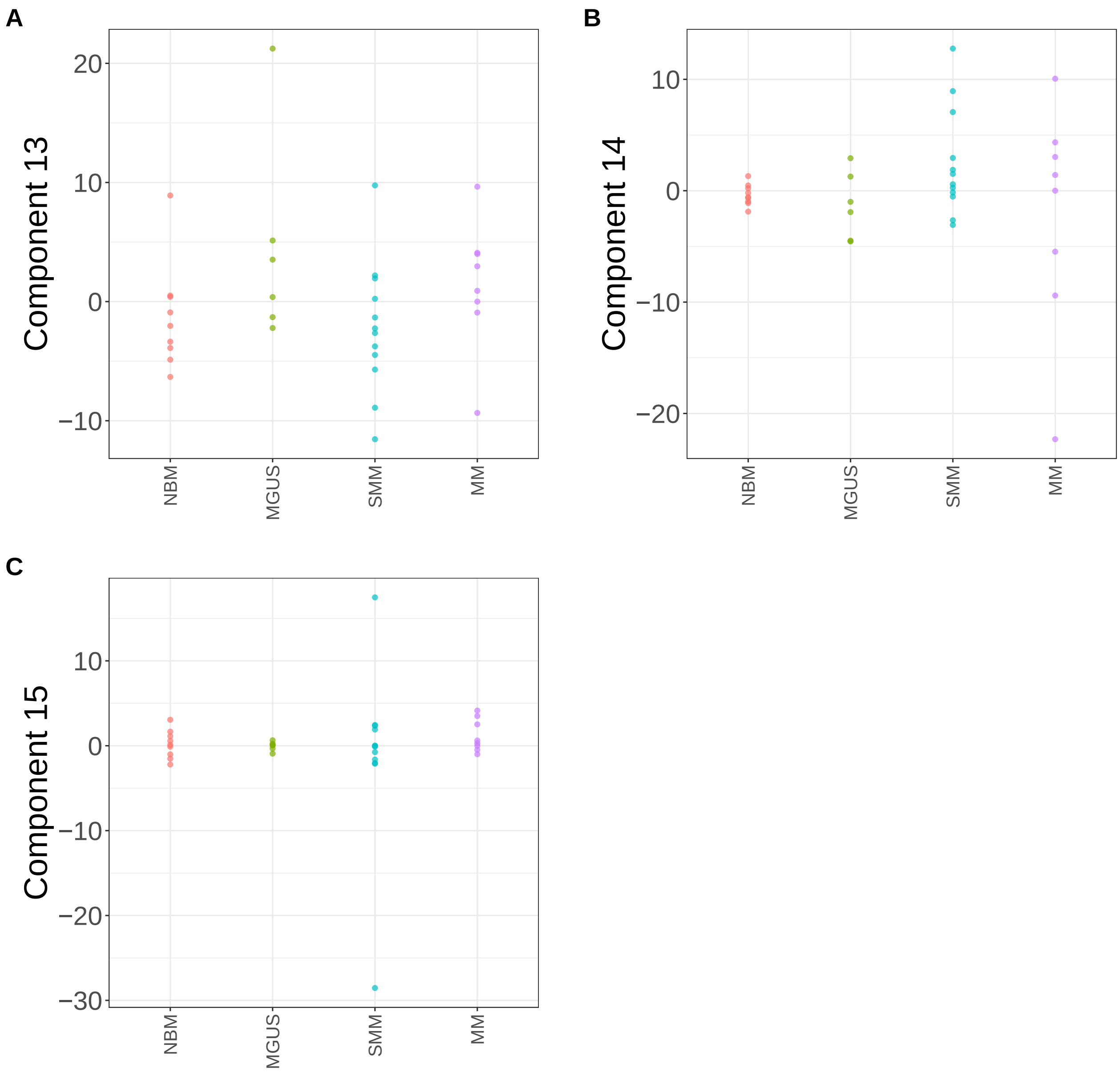


**Supplementary Figure 197.** Components for sPCA: keepX=500. (A) Component 13. (B) Component 14. (C) Component 15.

sPCA: keepX=1250

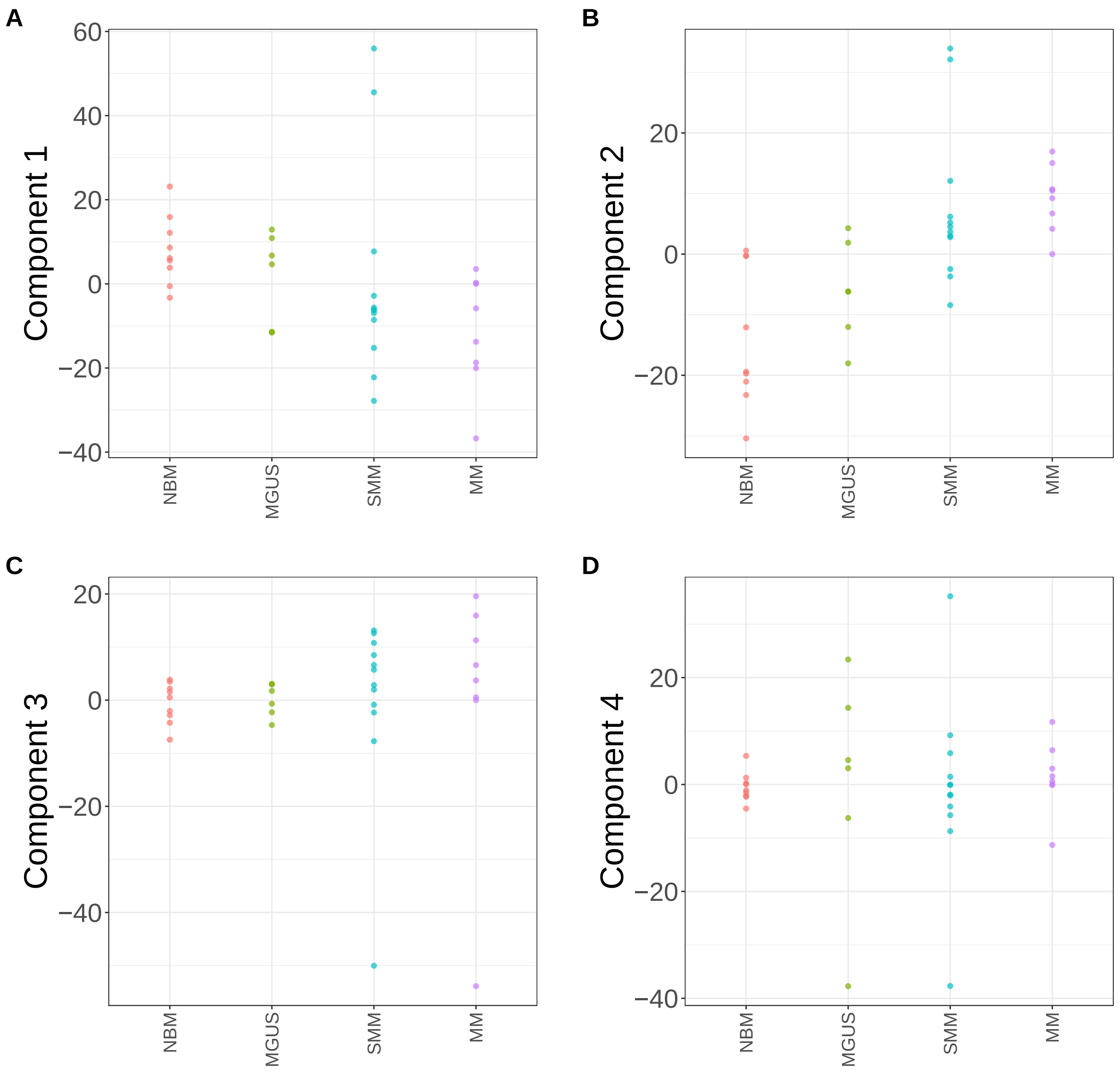


**Supplementary Figure 198.** Components for sPCA: keepX=1250. (A) Component 1. (B) Component 2. (C) Component 3. (D) Component 4.

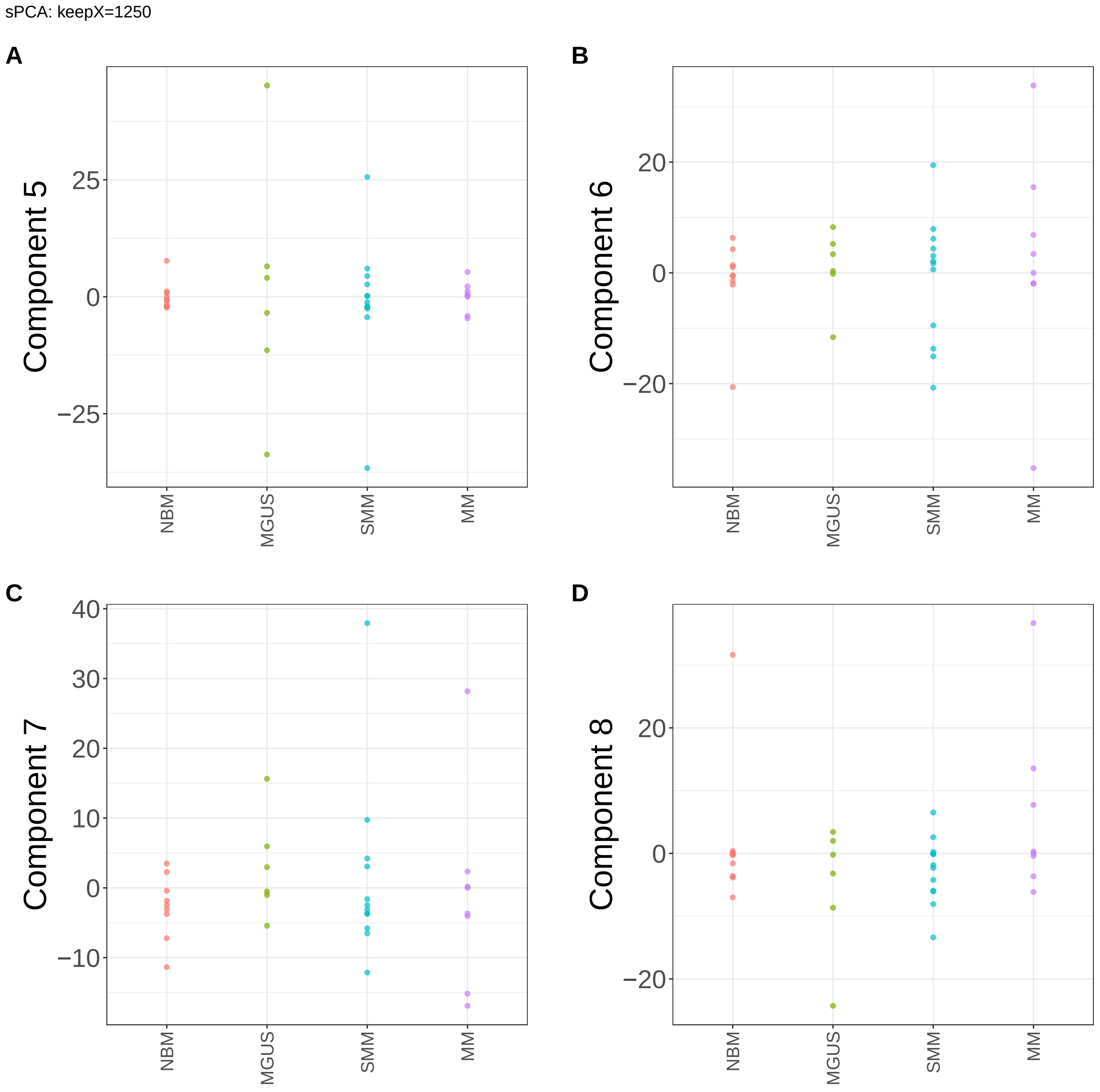


**Supplementary Figure 199.** Components for sPCA: keepX=1250. (A) Component 5. (B) Component 6. (C) Component 7. (D) Component 8.

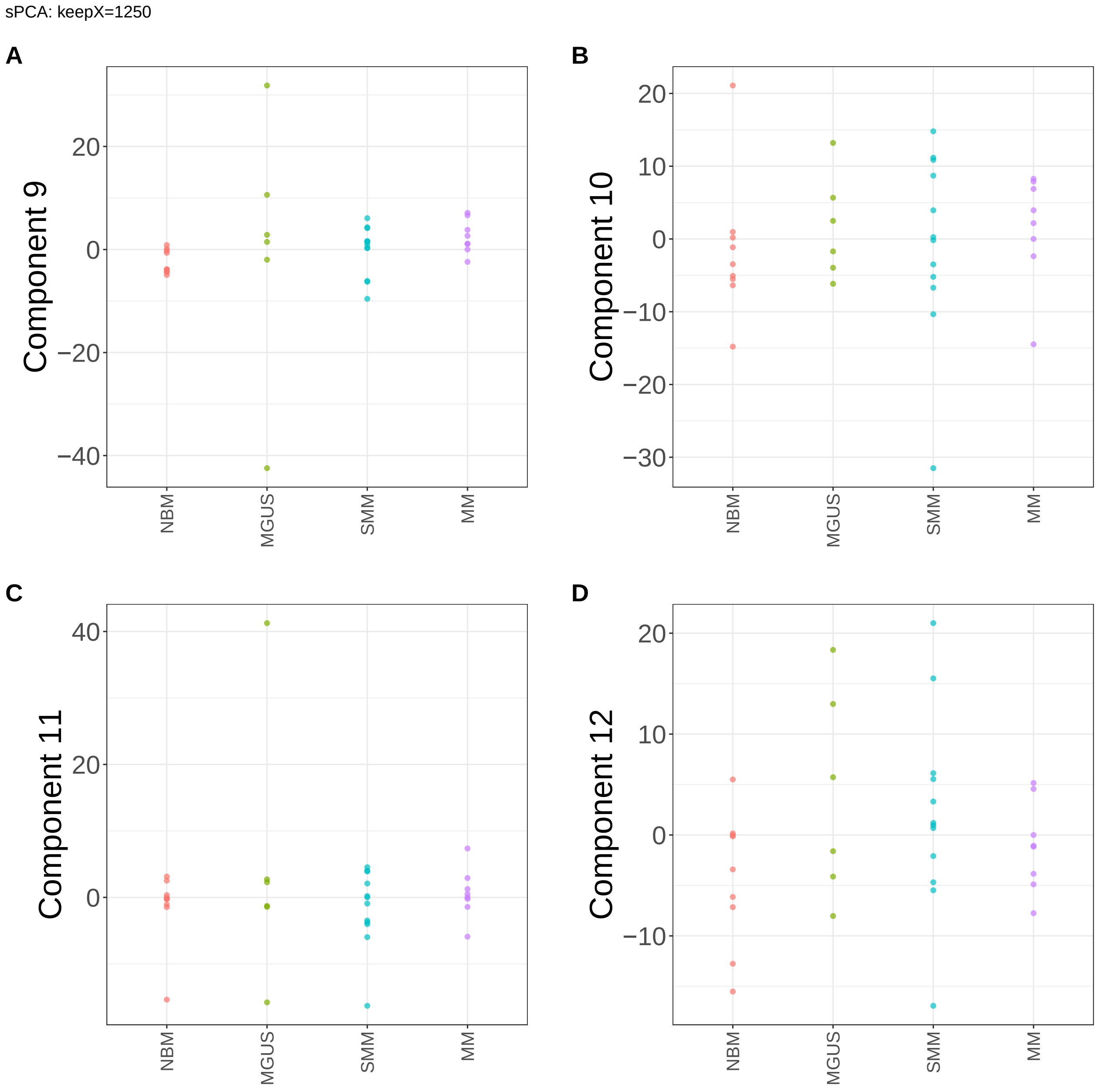


**Supplementary Figure 200.** Components for sPCA: keepX=1250. (A) Component 9. (B) Component 10. (C) Component 11. (D) Component 12.

sPCA: keepX=1250

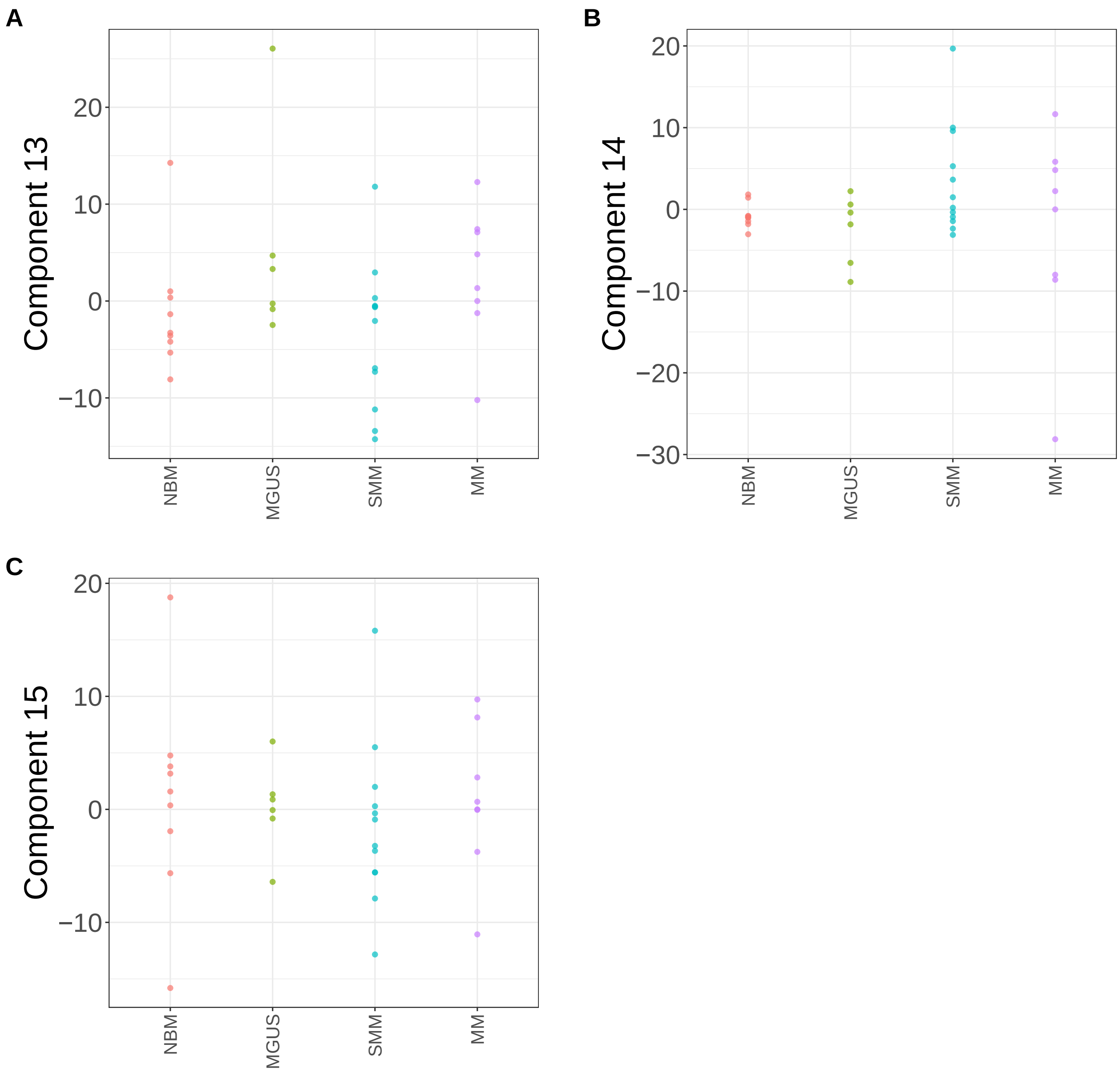


**Supplementary Figure 201.** Components for sPCA: keepX=1250. (A) Component 13. (B) Component 14. (C) Component 15.

sPCA: keepX=2500

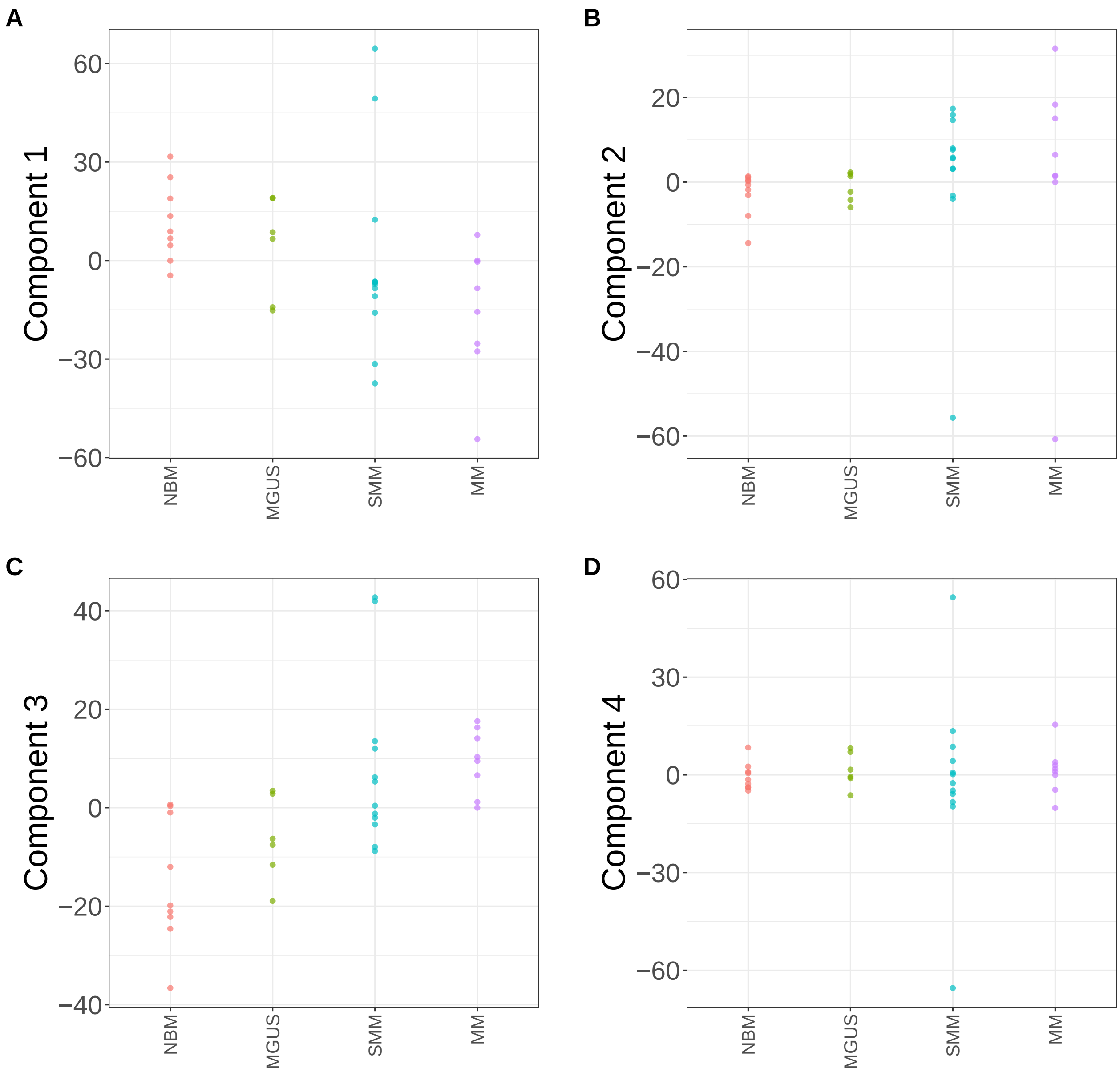


**Supplementary Figure 202.** Components for sPCA: keepX=2500. (A) Component 1. (B) Component 2. (C) Component 3. (D) Component 4.

sPCA: keepX=2500

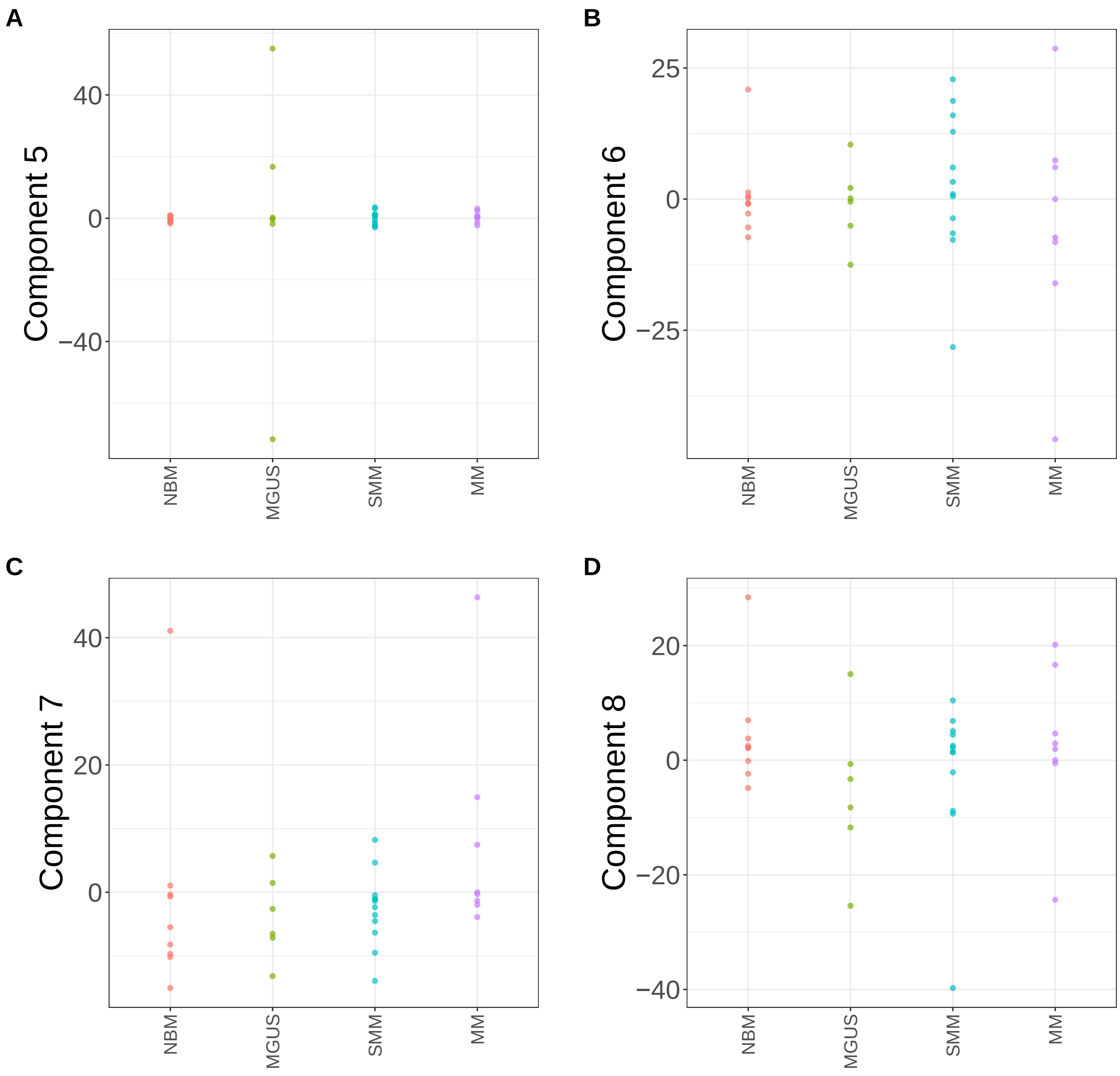


**Supplementary Figure 203.** Components for sPCA: keepX=2500. (A) Component 5. (B) Component 6. (C) Component 7. (D) Component 8.

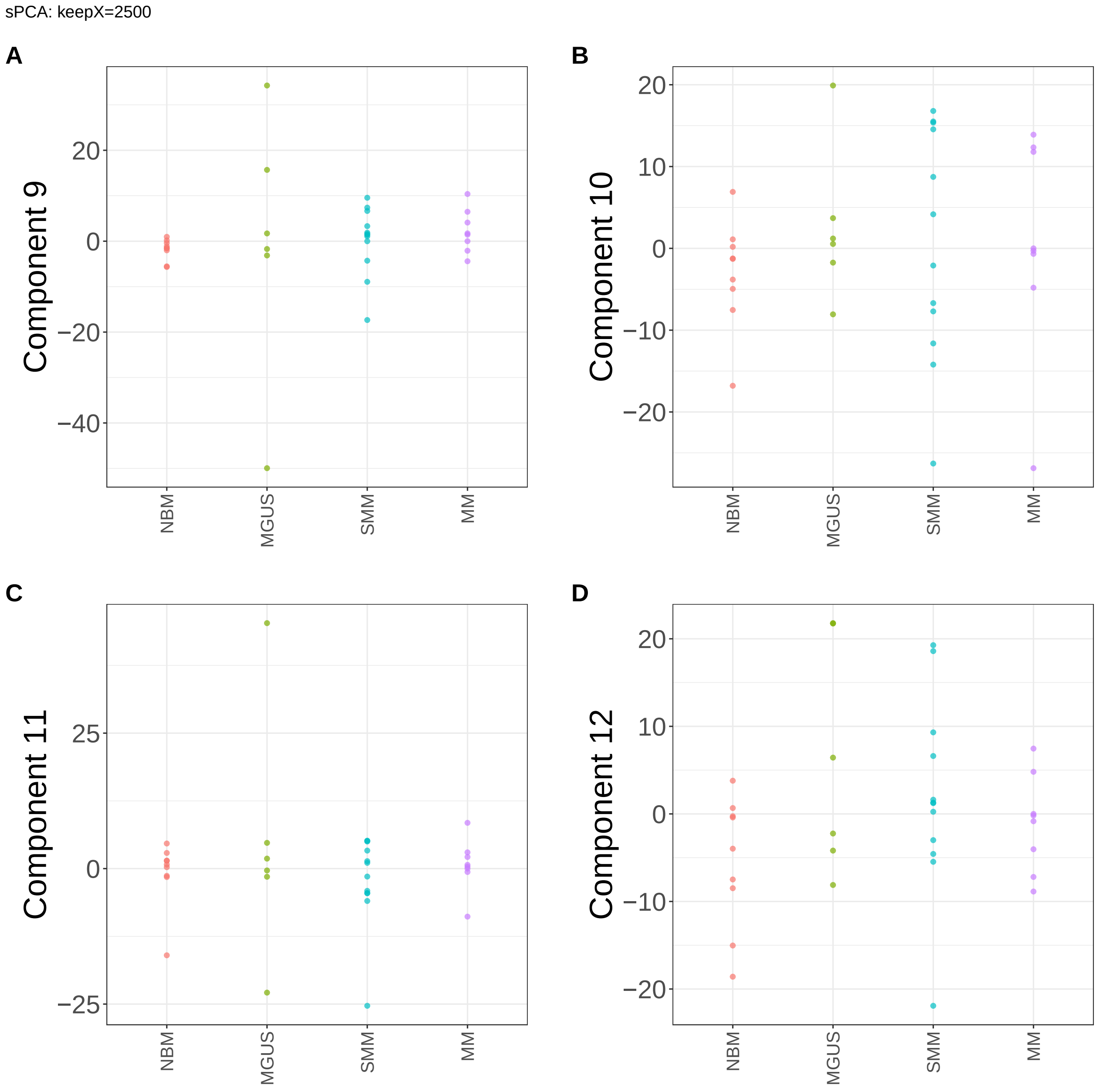


**Supplementary Figure 204.** Components for sPCA: keepX=2500. (A) Component 9. (B) Component 10. (C) Component 11. (D) Component 12.

sPCA: keepX=2500

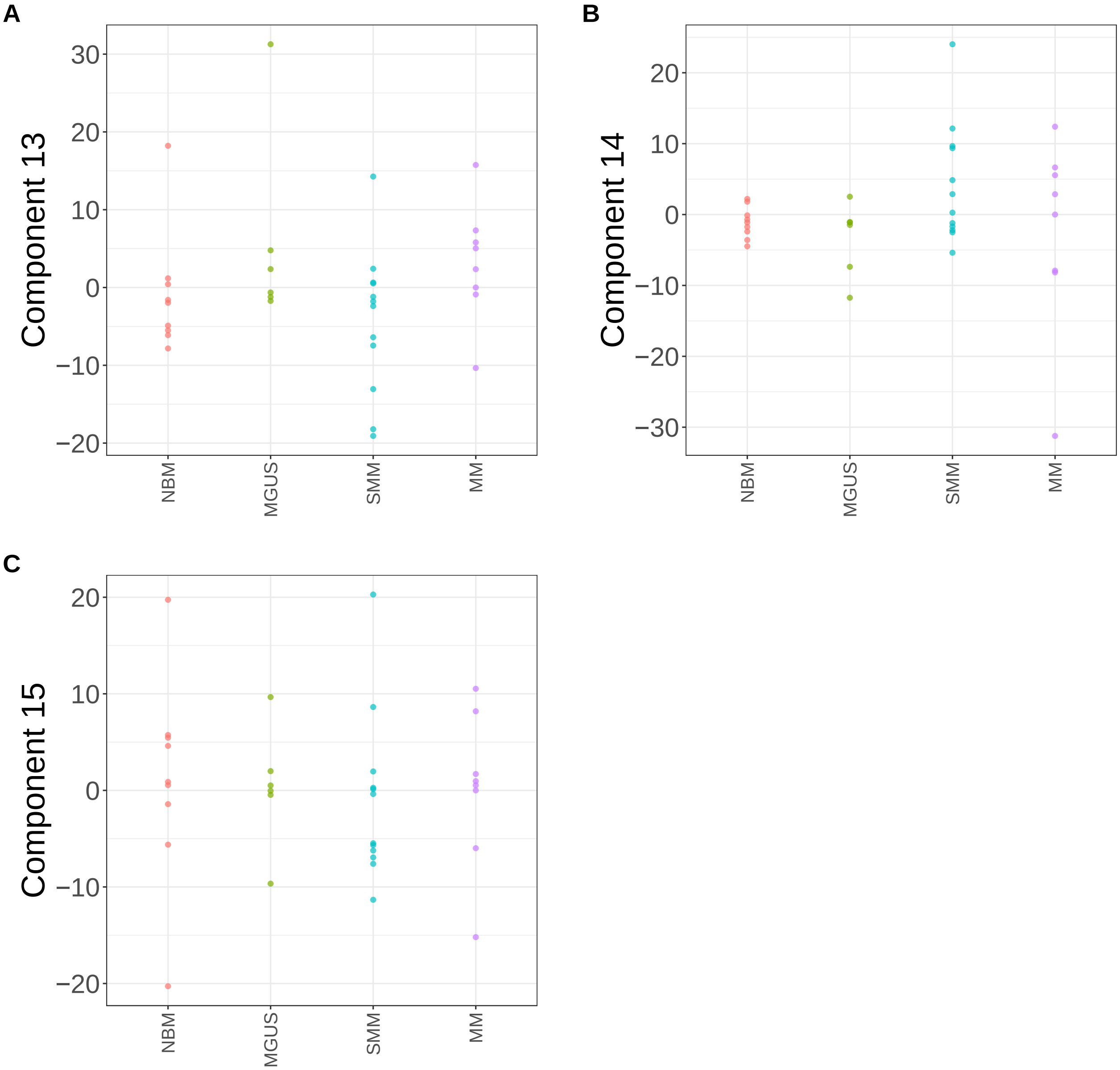


**Supplementary Figure 205.** Components for sPCA: keepX=2500. (A) Component 13. (B) Component 14. (C) Component 15.

sPCA: keepX=3750

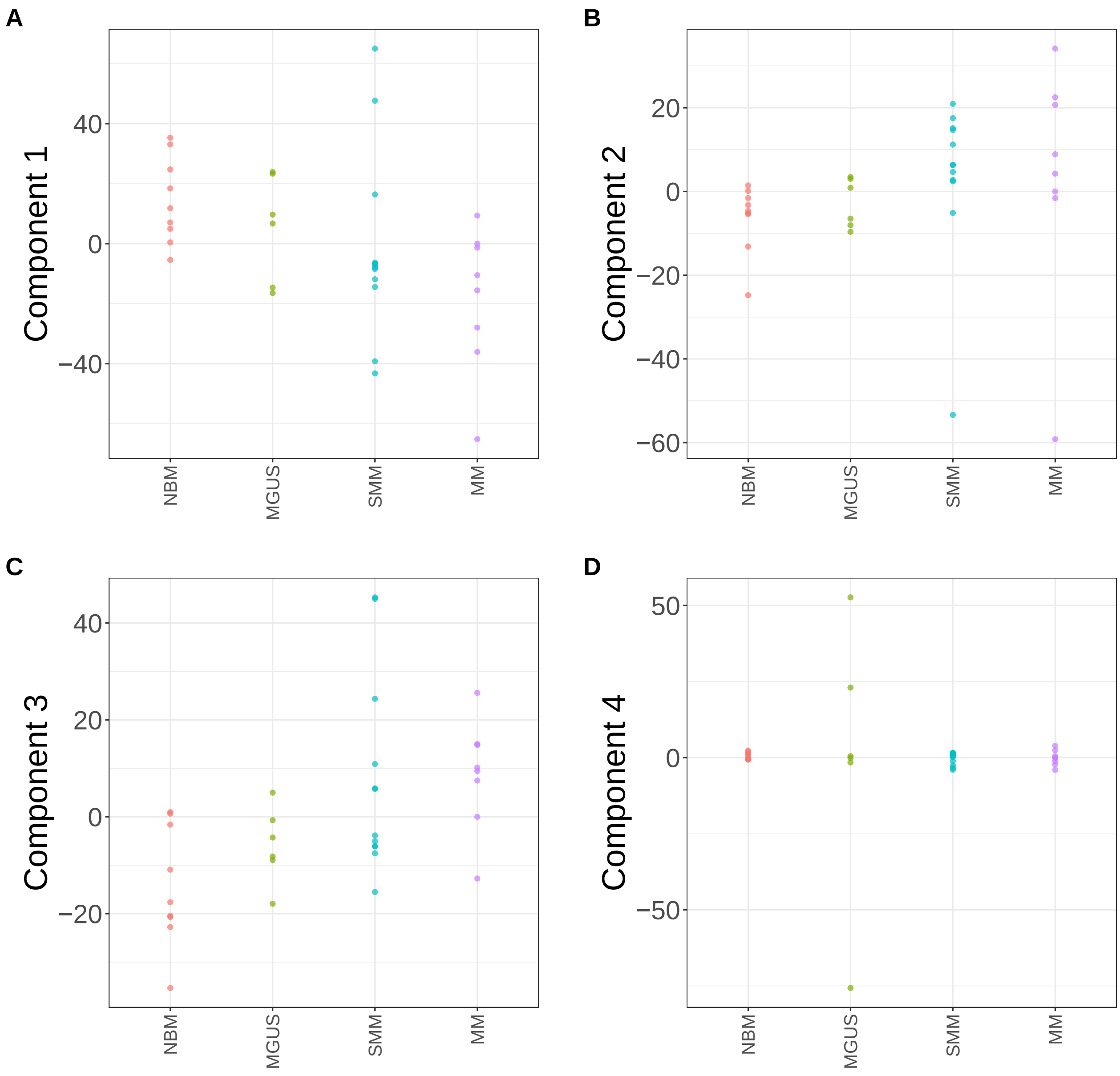


**Supplementary Figure 206.** Components for sPCA: keepX=3750. (A) Component 1. (B) Component 2. (C) Component 3. (D) Component 4.

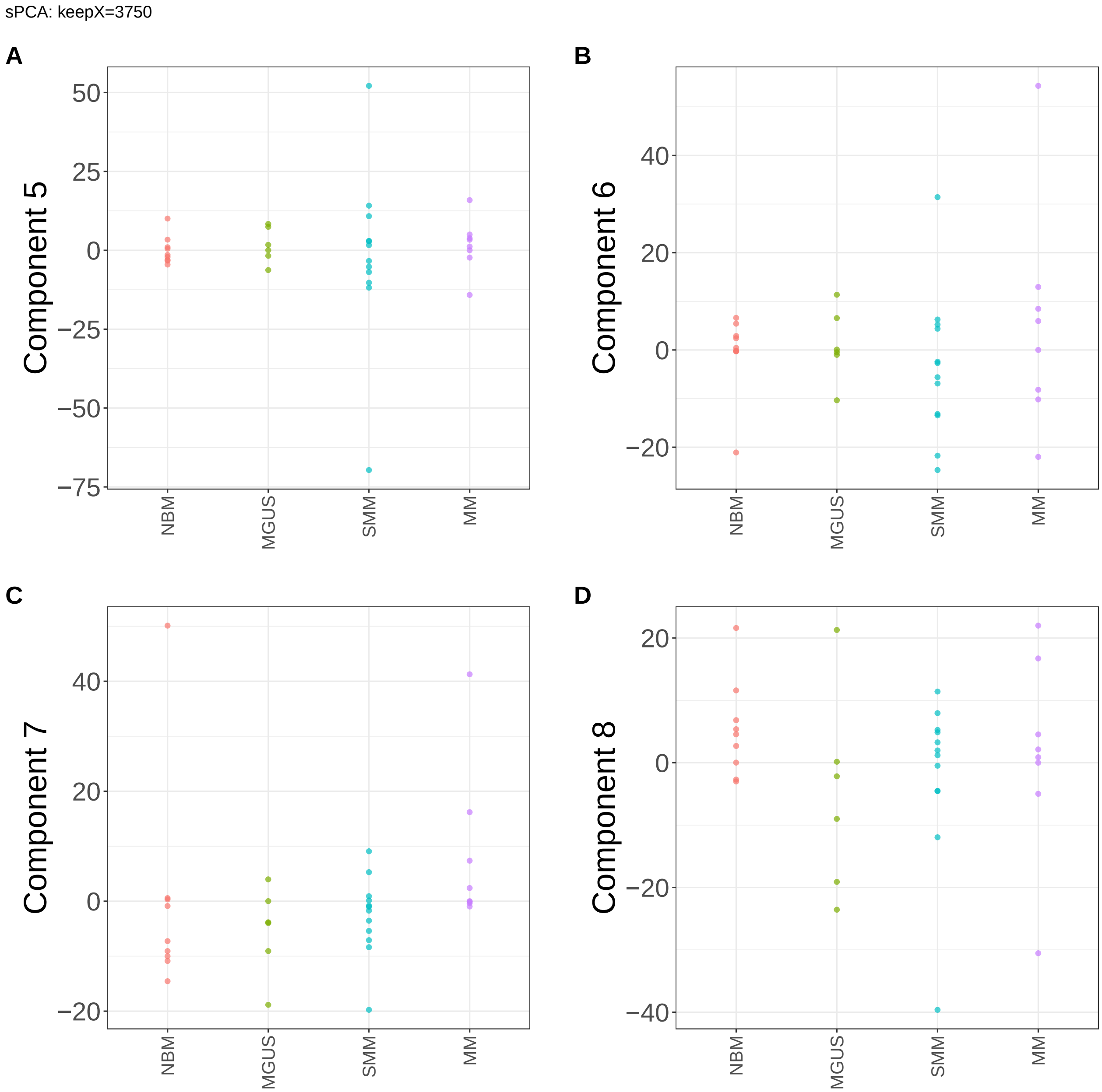


**Supplementary Figure 207.** Components for sPCA: keepX=3750. (A) Component 5. (B) Component 6. (C) Component 7. (D) Component 8.

sPCA: keepX=3750

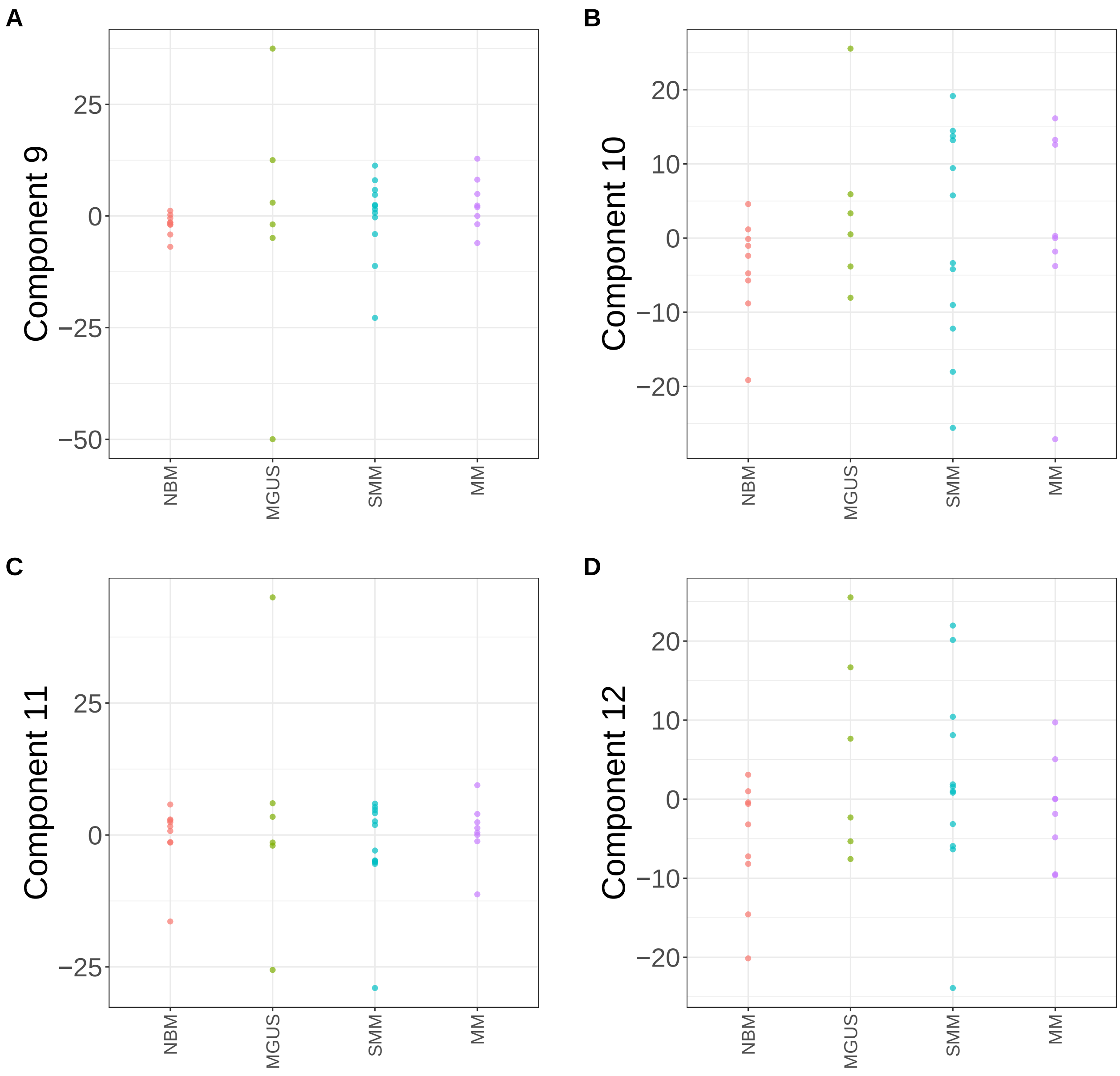


**Supplementary Figure 208.** Components for sPCA: keepX=3750. (A) Component 9. (B) Component 10. (C) Component 11. (D) Component 12.

sPCA: keepX=3750

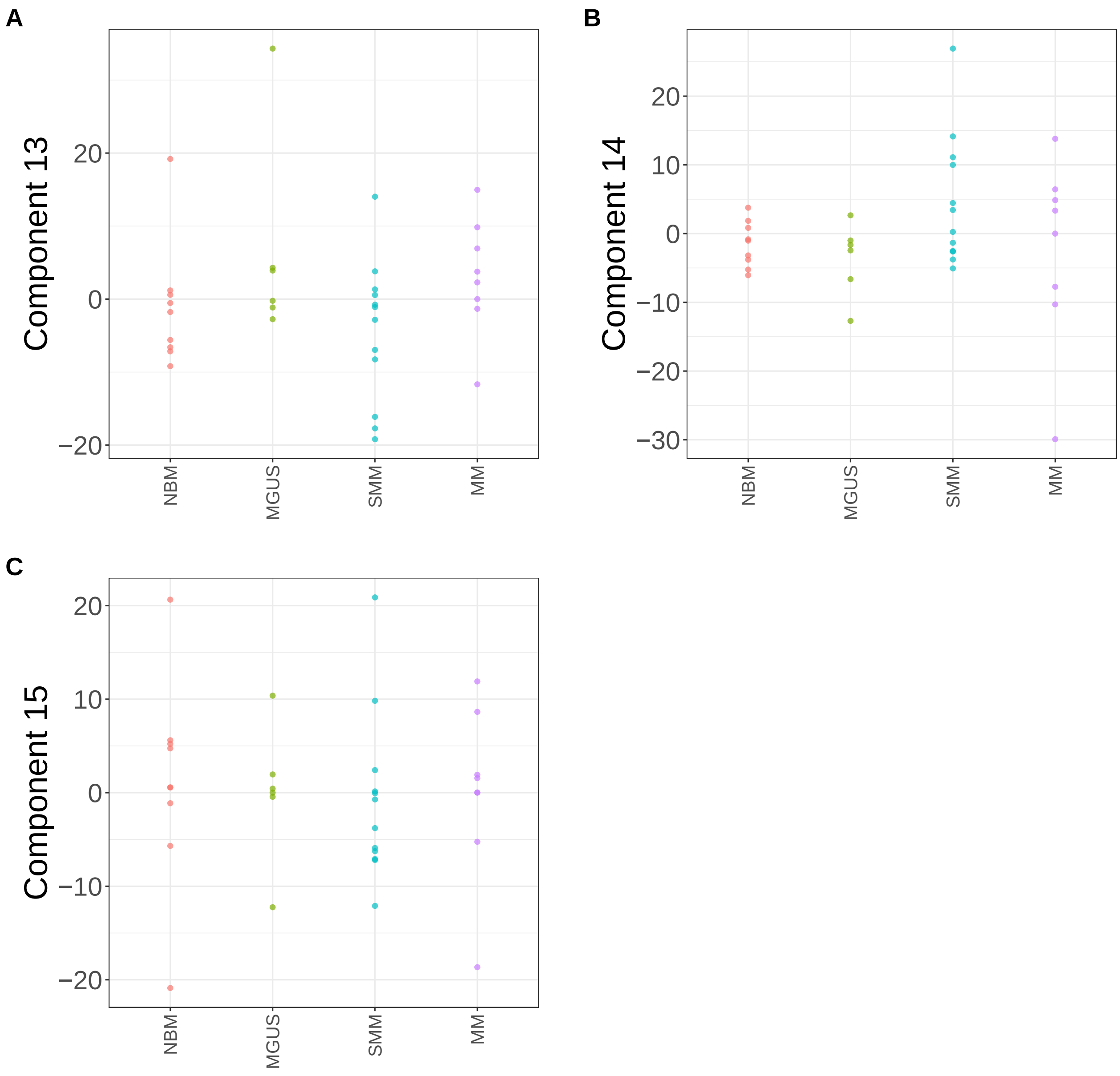


**Supplementary Figure 209.** Components for sPCA: keepX=3750. (A) Component 13. (B) Component 14. (C) Component 15.

sNMF: lambda=0.001

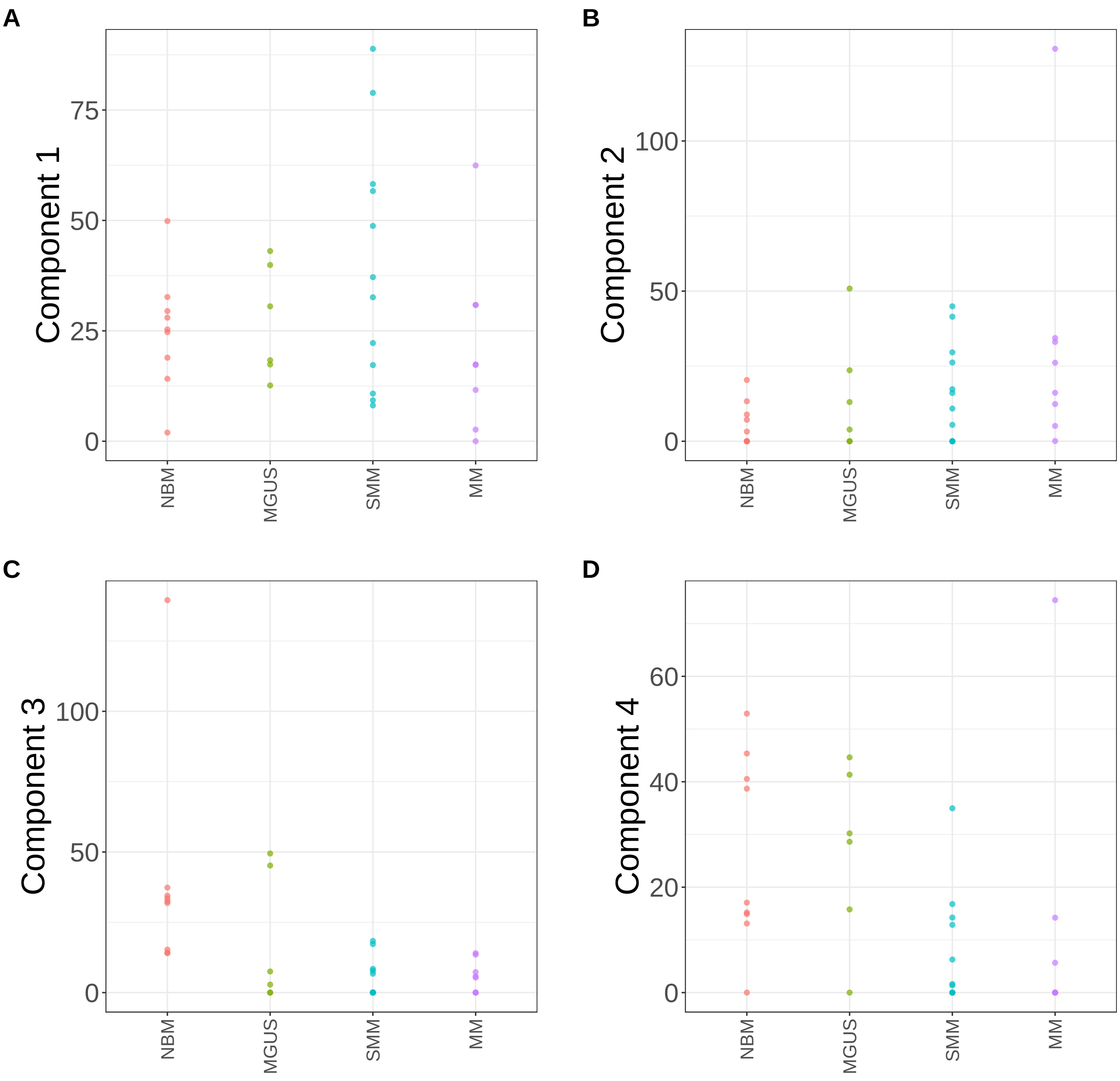


**Supplementary Figure 210.** Components for sNMF: lambda=0.001. (A) Component 1. (B) Component 2. (C) Component 3. (D) Component 4.

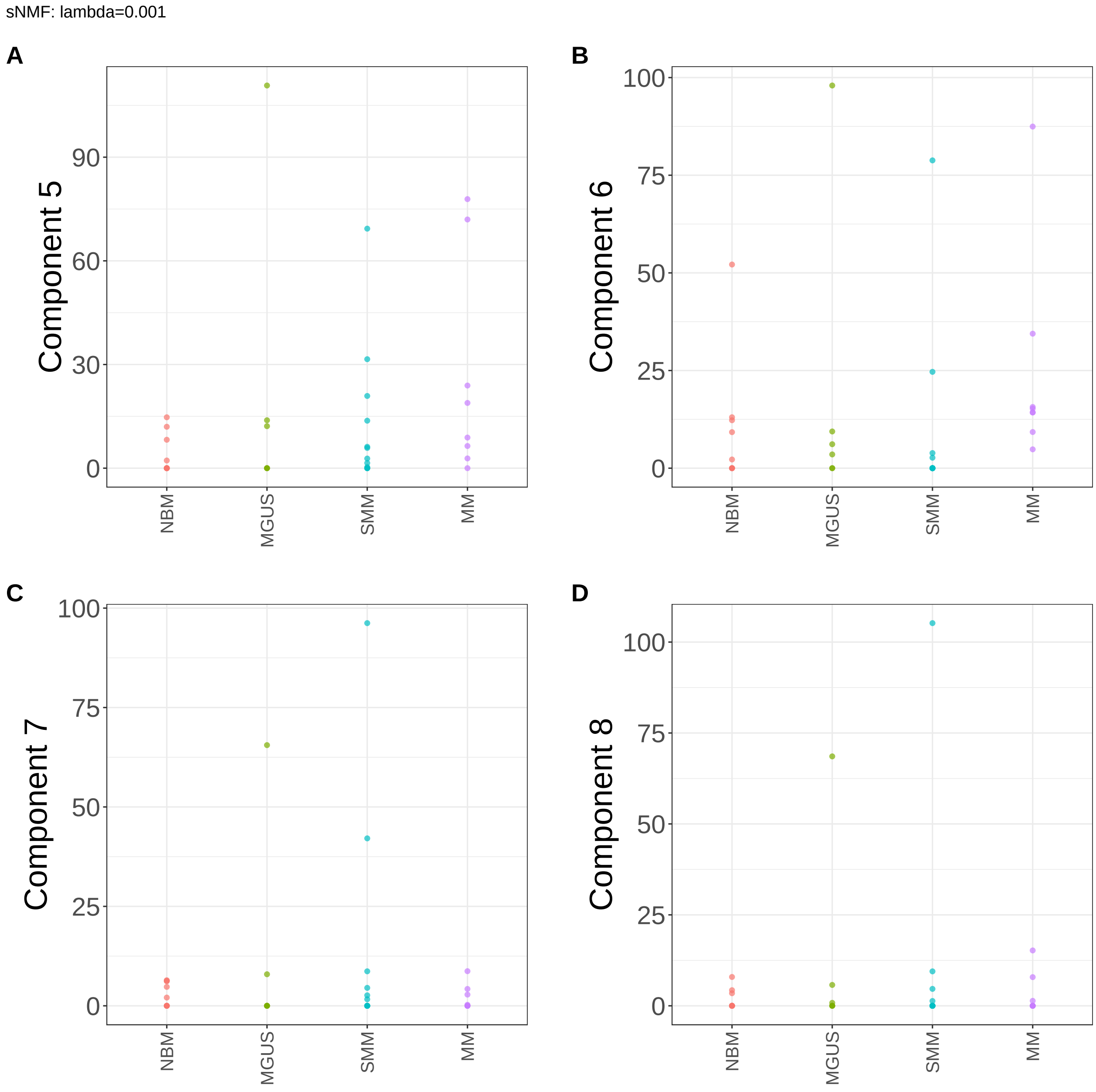


**Supplementary Figure 211.** Components for sNMF: lambda=0.001. (A) Component 5. (B) Component 6. (C) Component 7. (D) Component 8.

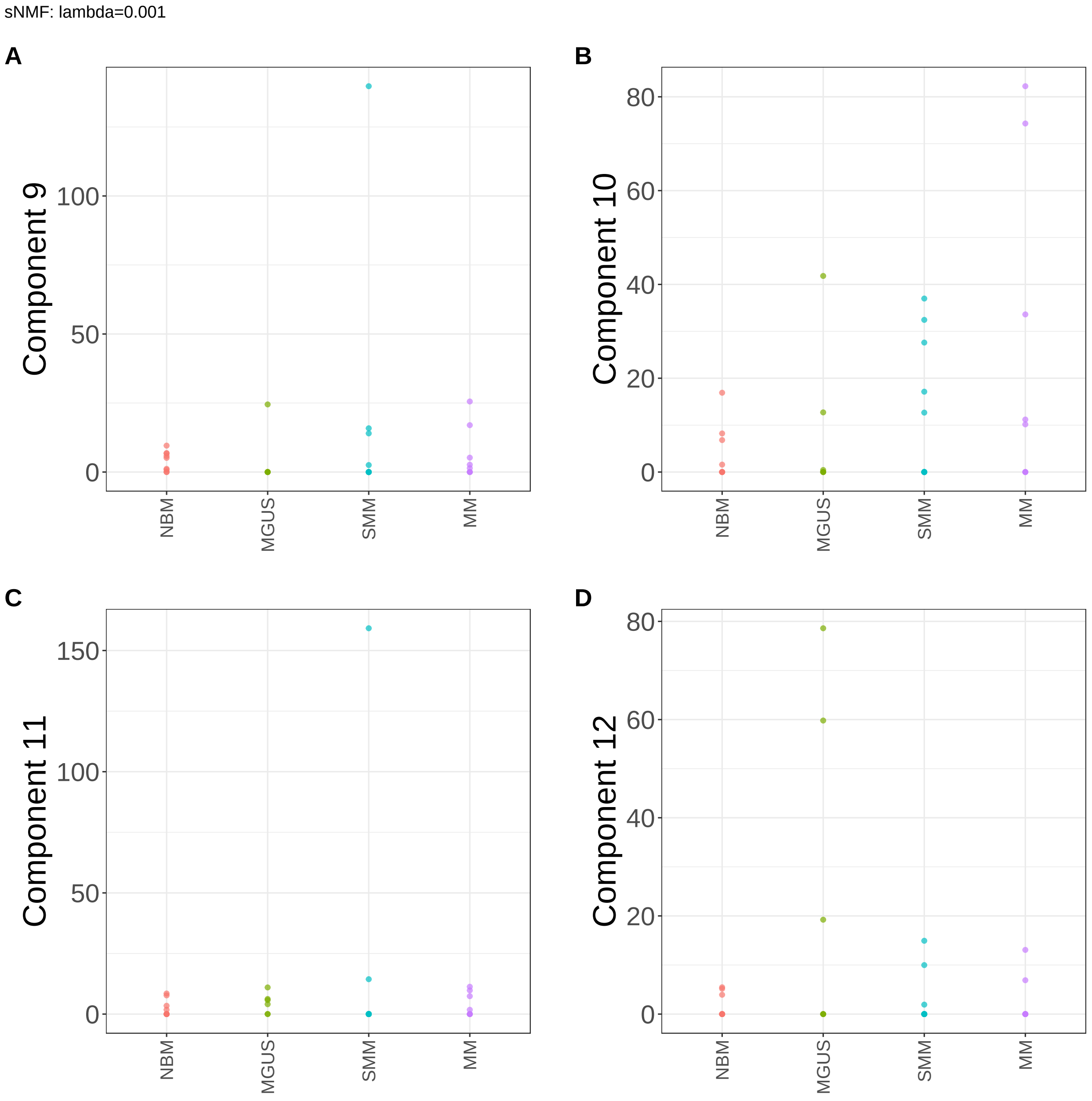


**Supplementary Figure 212.** Components for sNMF: lambda=0.001. (A) Component 9. (B) Component 10. (C) Component 11. (D) Component 12.

sNMF: lambda=0.001

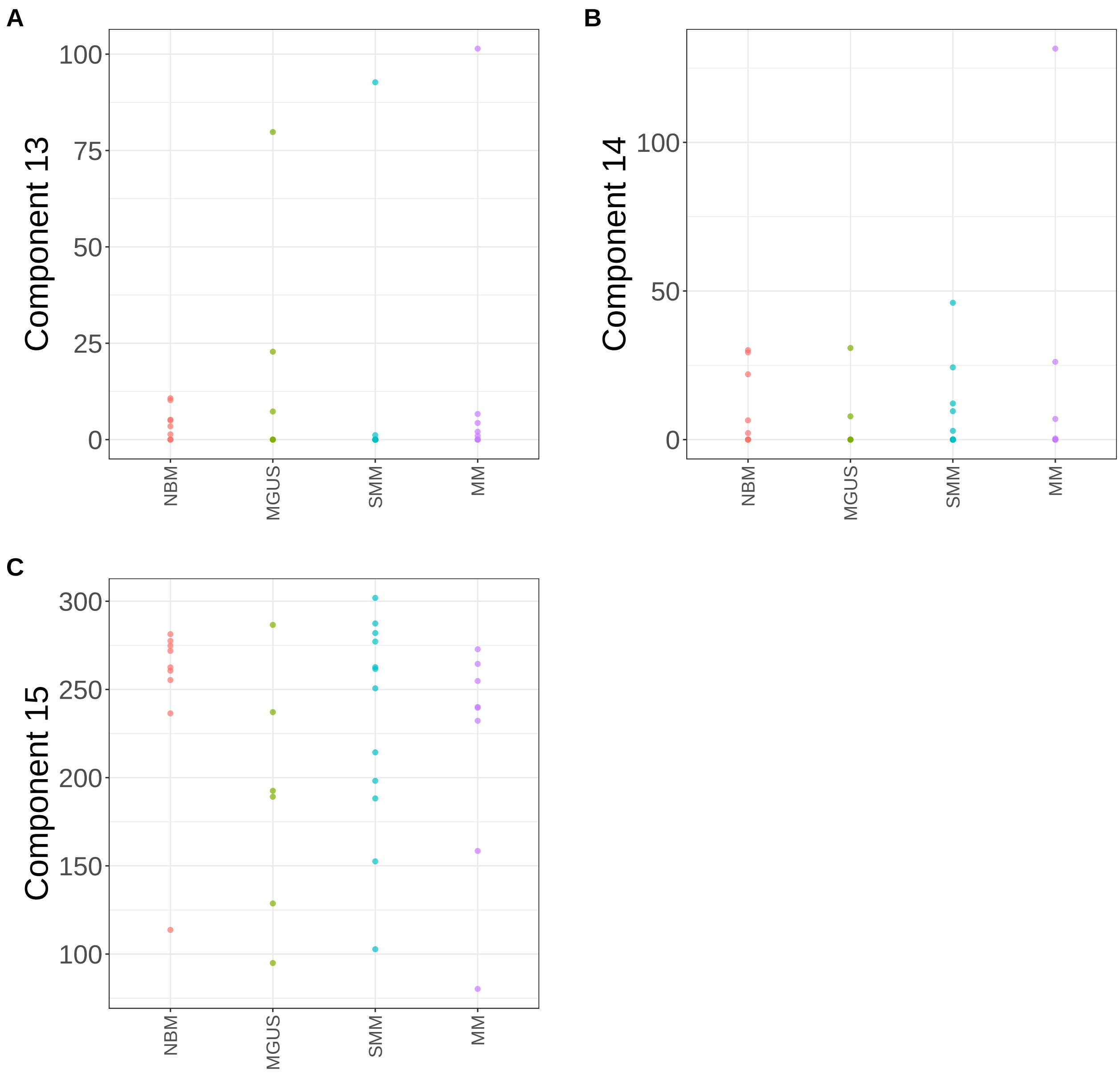


**Supplementary Figure 213.** Components for sNMF: lambda=0.001. (A) Component 13. (B) Component 14. (C) Component 15.

sNMF: lambda=0.01

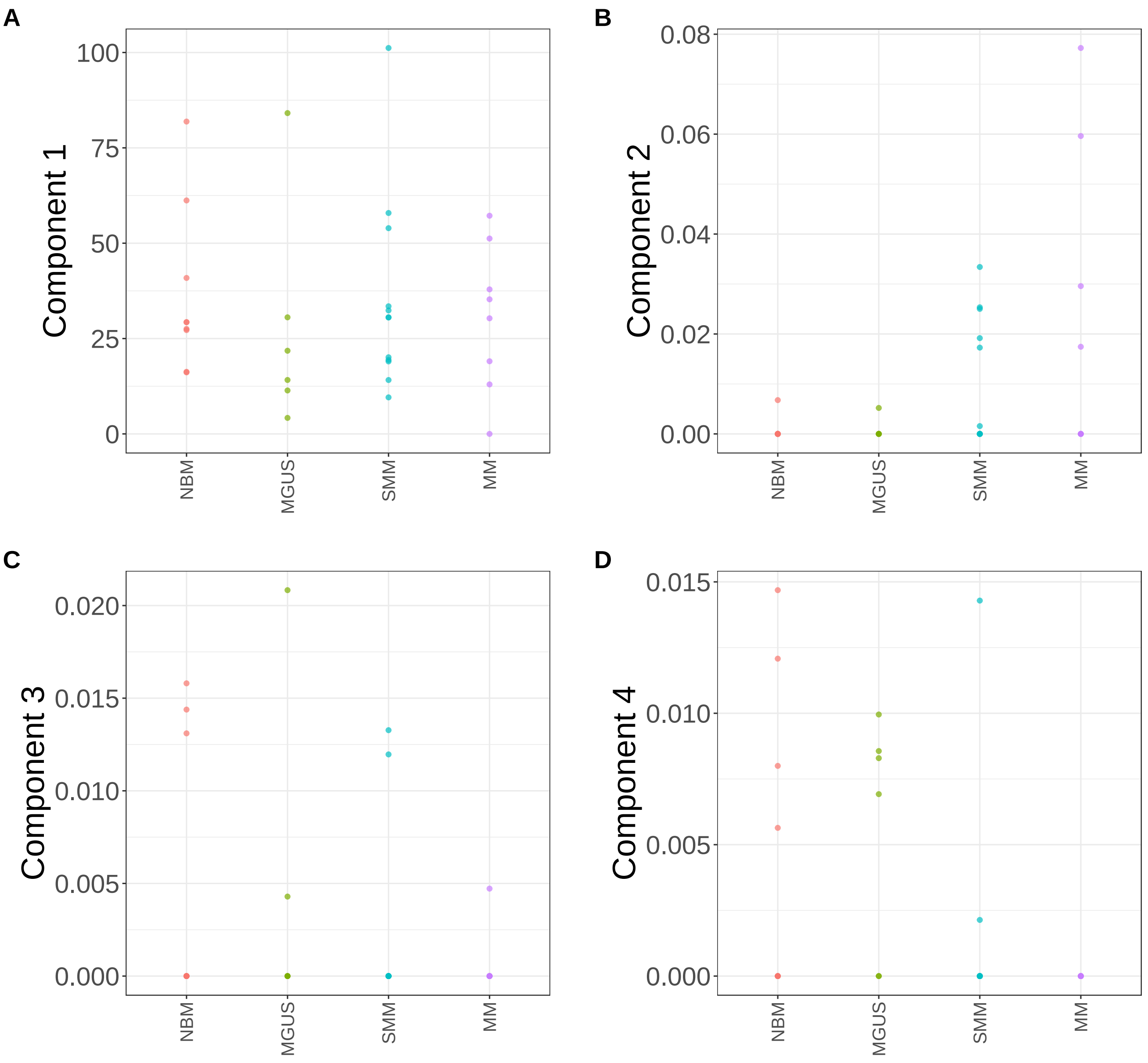


**Supplementary Figure 214.** Components for sNMF: lambda=0.01. (A) Component 1. (B) Component 2. (C) Component 3. (D) Component 4.

sNMF: lambda=0.01

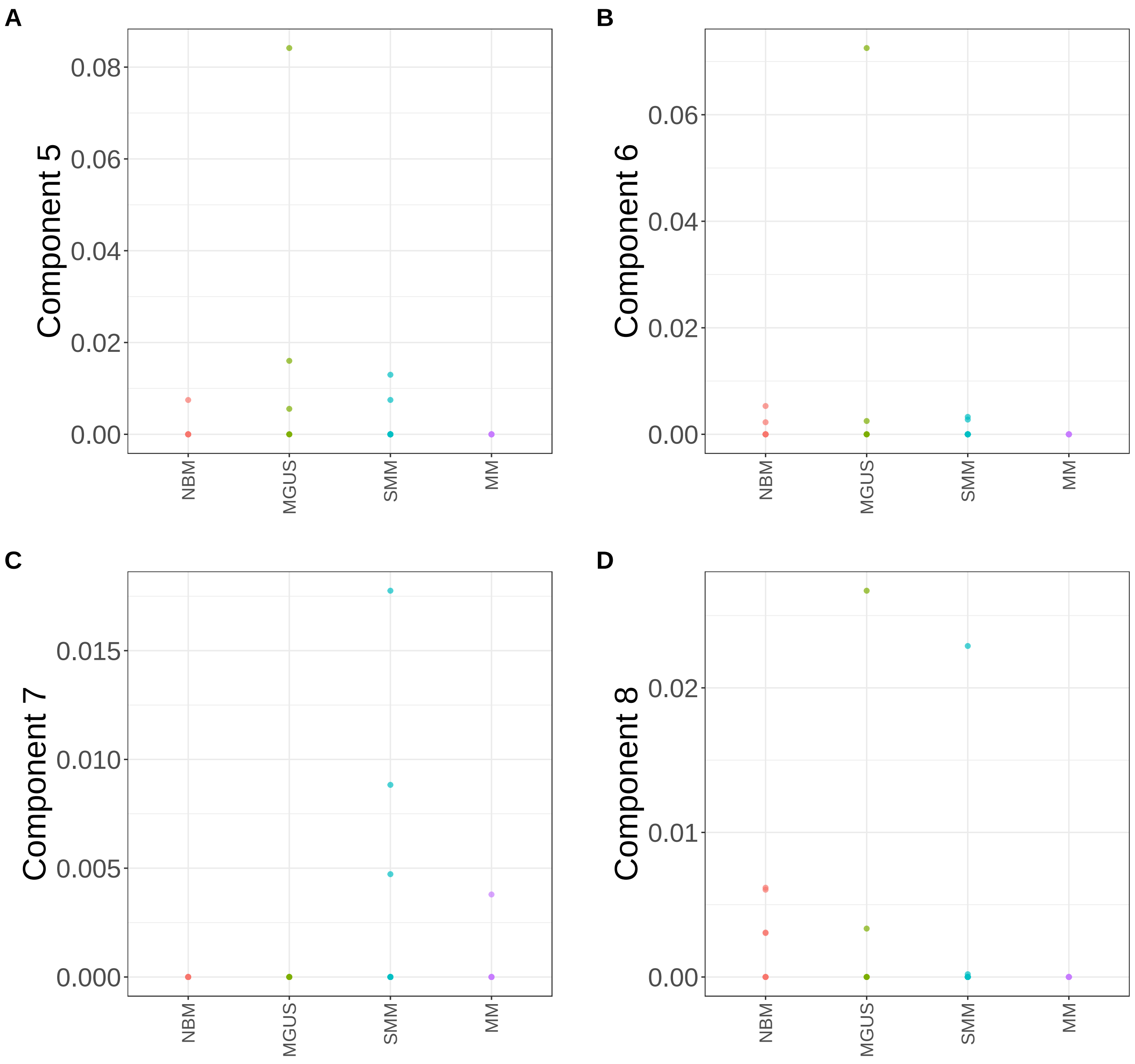


**Supplementary Figure 215.** Components for sNMF: lambda=0.01. (A) Component 5. (B) Component 6. (C) Component 7. (D) Component 8.

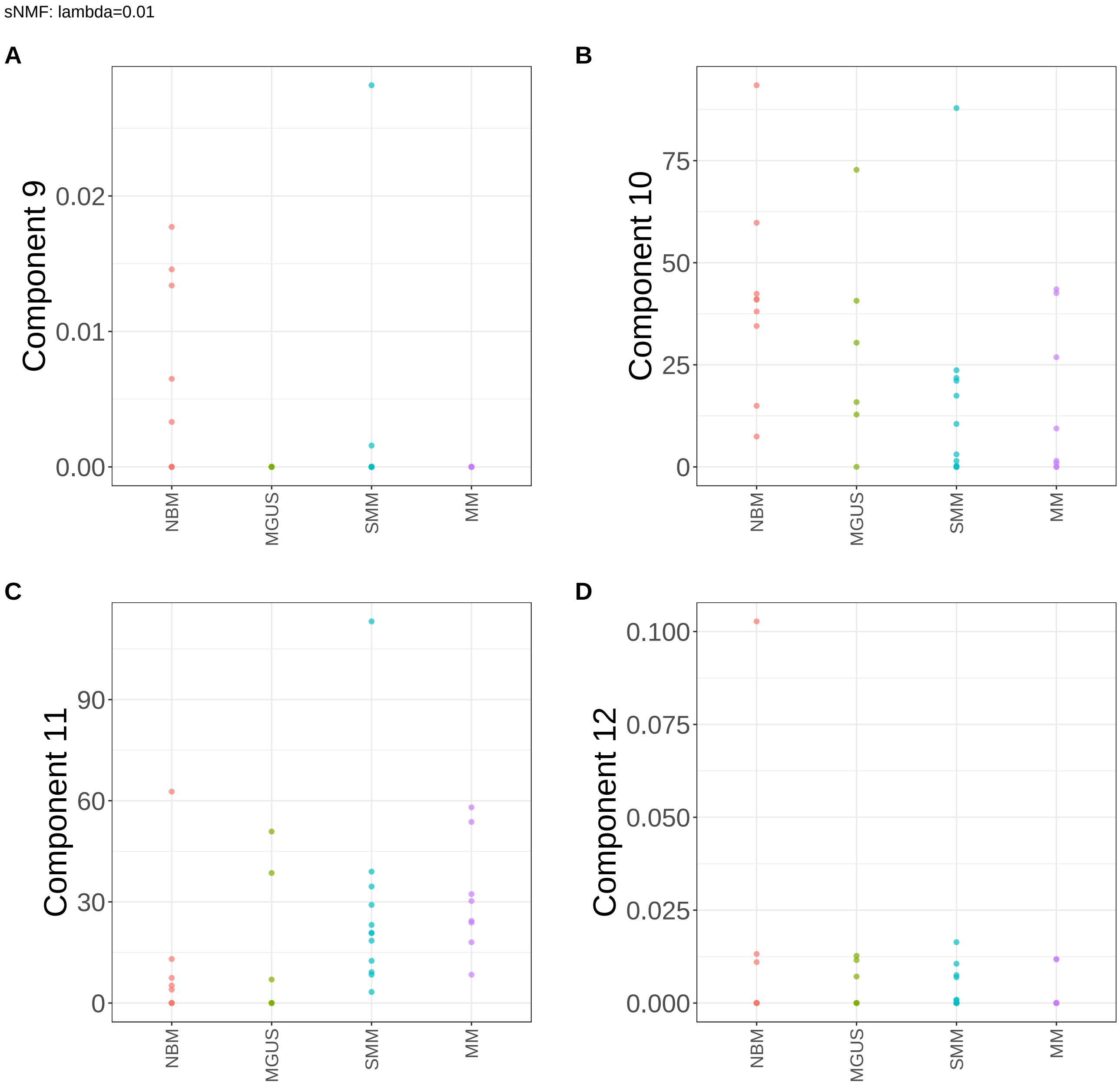


**Supplementary Figure 216.** Components for sNMF: lambda=0.01. (A) Component 9. (B) Component 10. (C) Component 11. (D) Component 12.

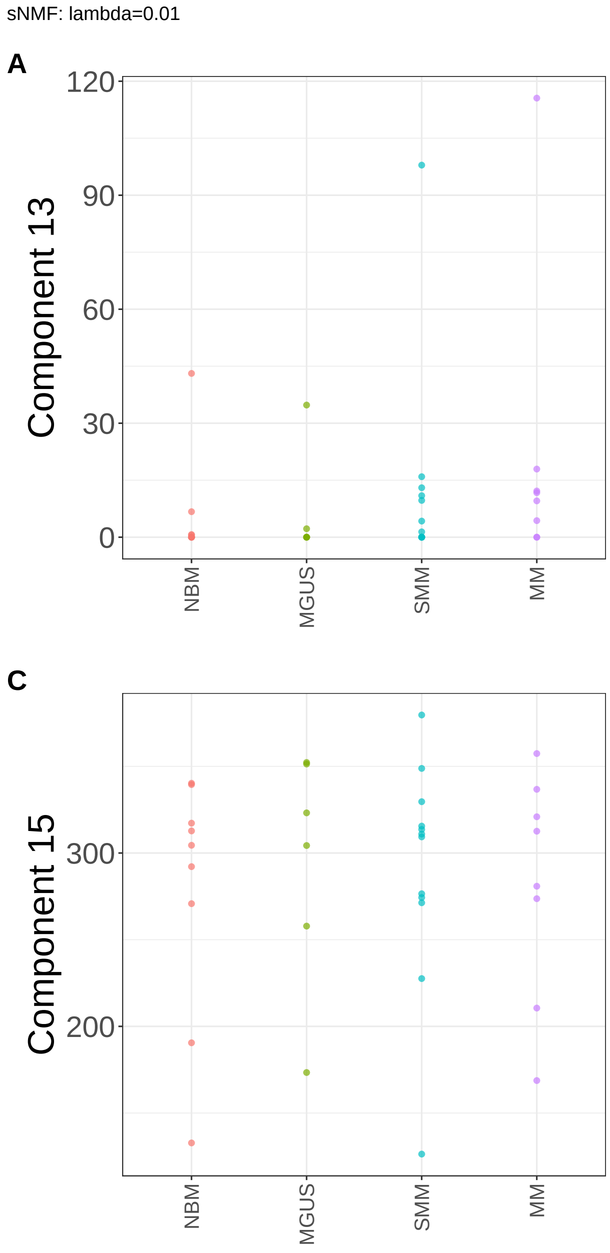


**Supplementary Figure 217.** Components for sNMF: lambda=0.01. (A) Component 13. (B) Component 14. (C) Component 15.

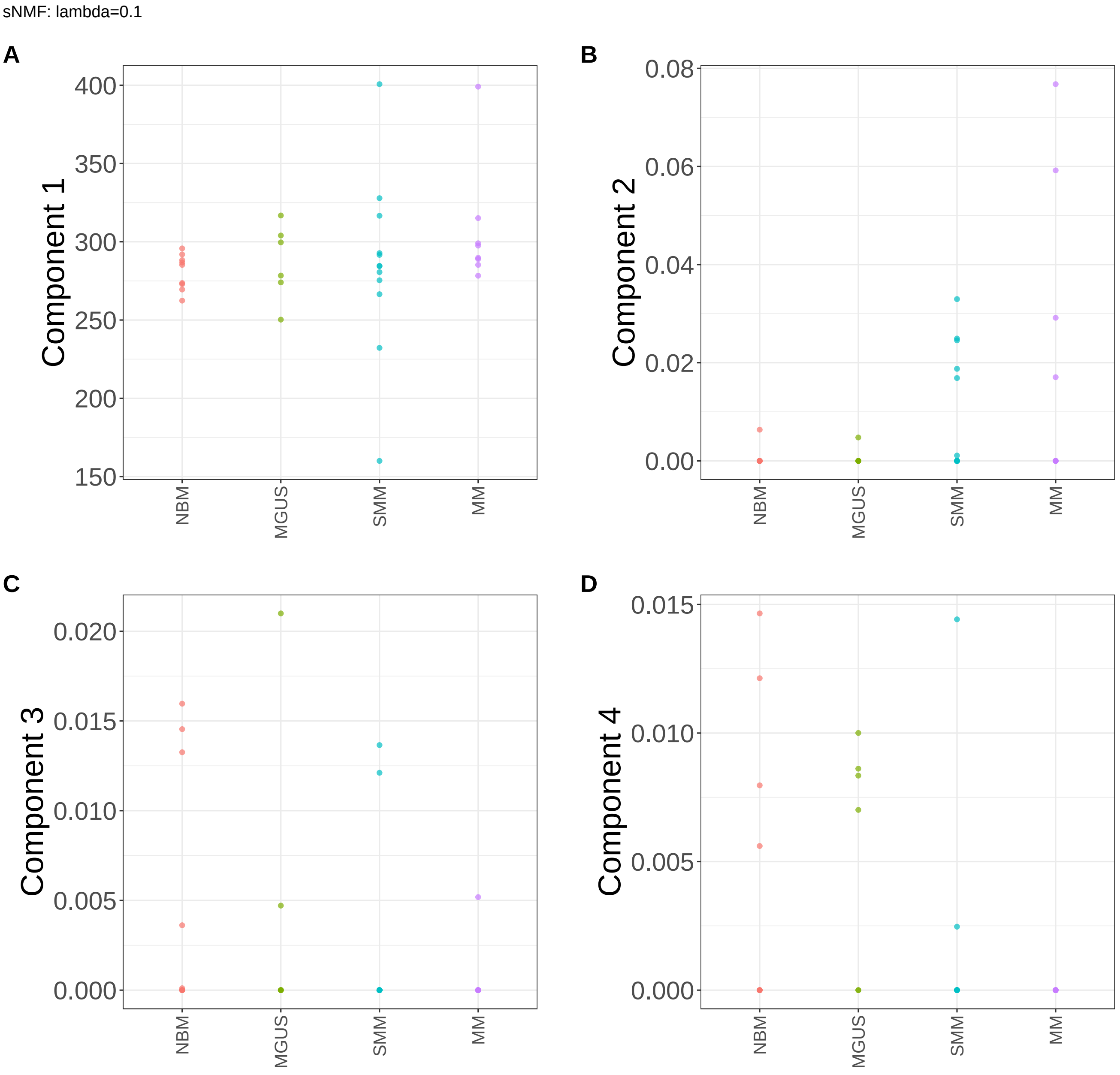


**Supplementary Figure 218.** Components for sNMF: lambda=0.1. (A) Component 1. (B) Component 2. (C) Component 3. (D) Component 4.

sNMF: lambda=0.1

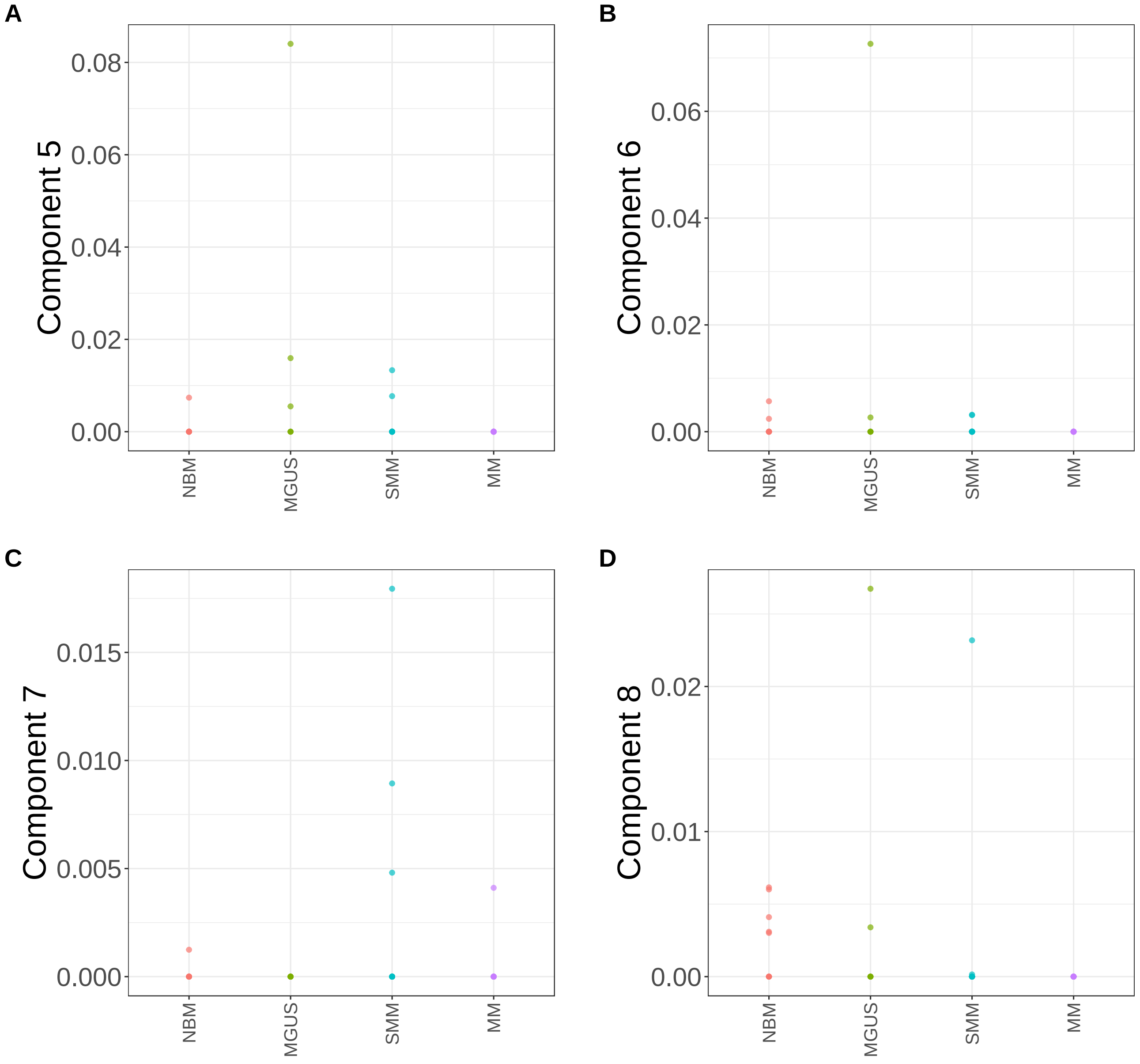


**Supplementary Figure 219.** Components for sNMF: lambda=0.1. (A) Component 5. (B) Component 6. (C) Component 7. (D) Component 8.

sNMF: lambda=0.1

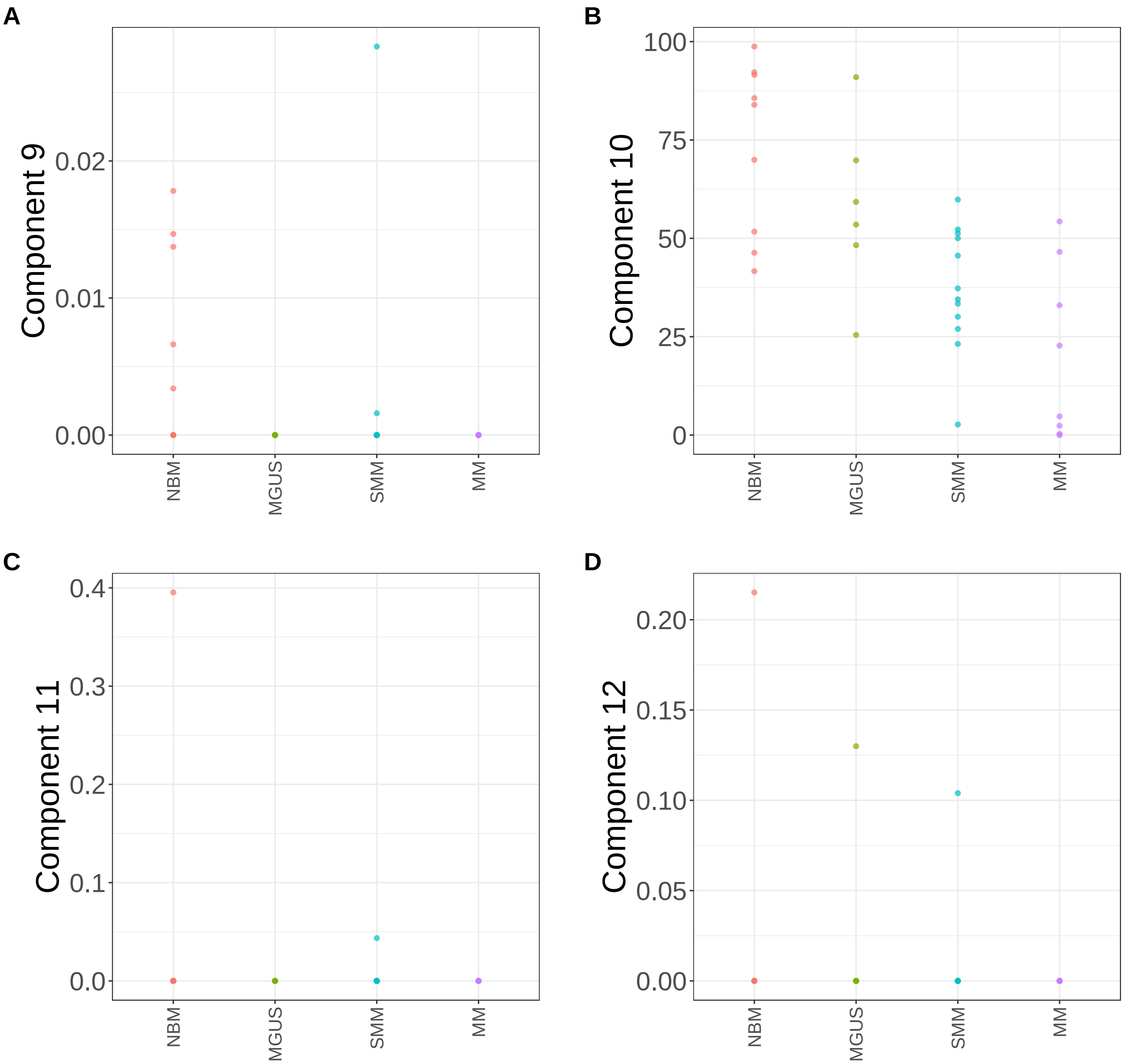


**Supplementary Figure 220.** Components for sNMF: lambda=0.1. (A) Component 9. (B) Component 10. (C) Component 11. (D) Component 12.

sNMF: lambda=0.1

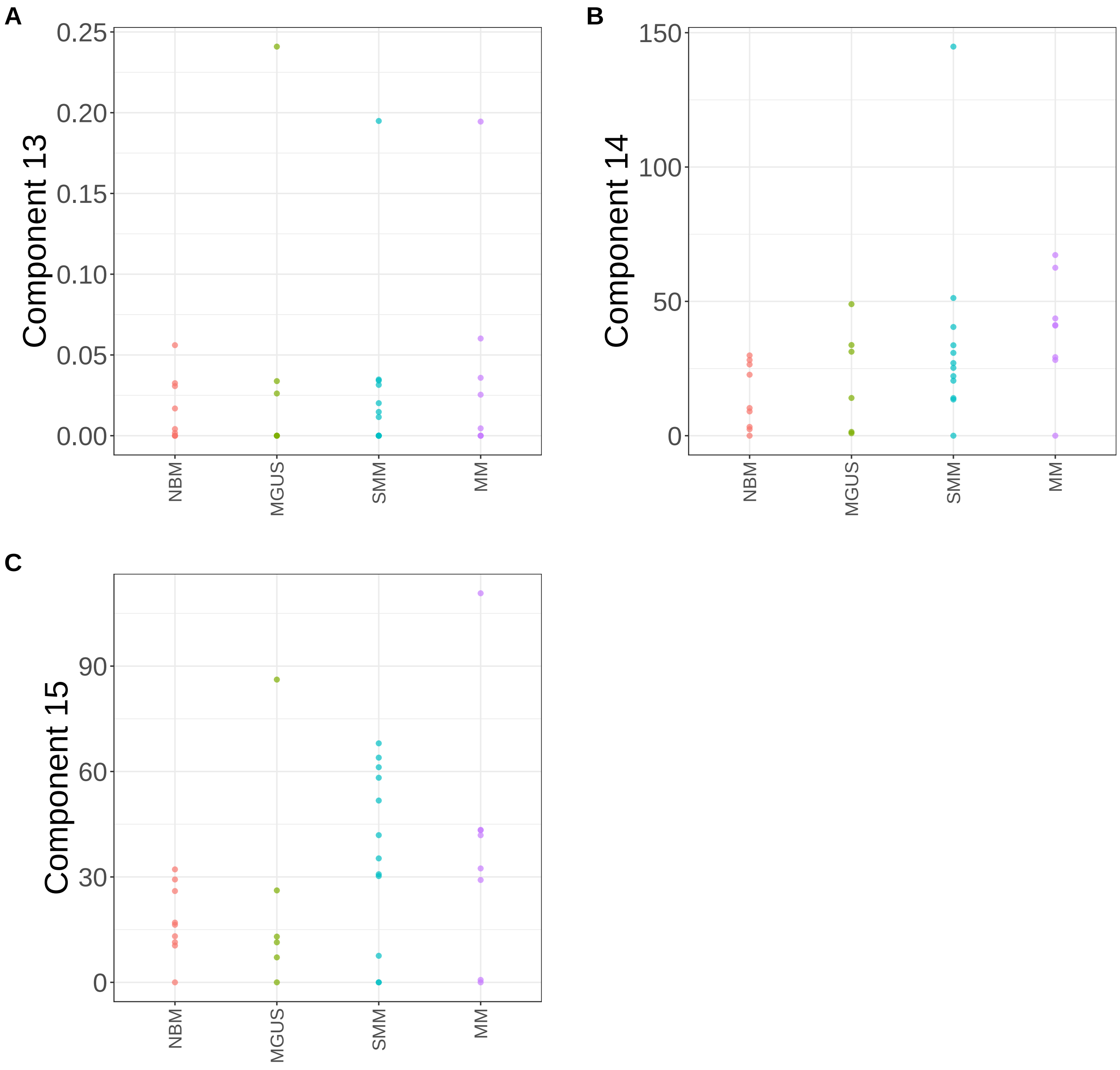


**Supplementary Figure 221.** Components for sNMF: lambda=0.1. (A) Component 13. (B) Component 14. (C) Component 15.
