## Supplementary Note 4 : Malaria for "A statistical framework for system-level synthesis"

This note presents the supplementary results from the analysis of single cell, P. berghei, data. The figures include:

- Processes/component functions learned by SSANOVA-SVD (**Supplementary Fig. 98**-**Supplementary Fig. 103**)
- Inference of feature participation by the application of the three-step framework: Volcano plots (**Supplementary Fig. 104**-**Supplementary Fig. 109**), heatmaps of top 50 component-associated features at FDR of 10% (**Supplementary Fig. 110**-**Supplementary Fig. 120**), geneset pathway enrichment for associated features (**Supplementary Fig. 121**-**Supplementary Fig. 122**)
- Decomposition of the processes (**Supplementary Fig. 123**-**Supplementary Fig. 133**). Because we have a single discrete covariate, the decomposition of the processes are identical to the processes themselves.
- Components from a representative class of matrix factorization methods (**Supplementary Fig. 134**-**Supplementary Fig. 176**)

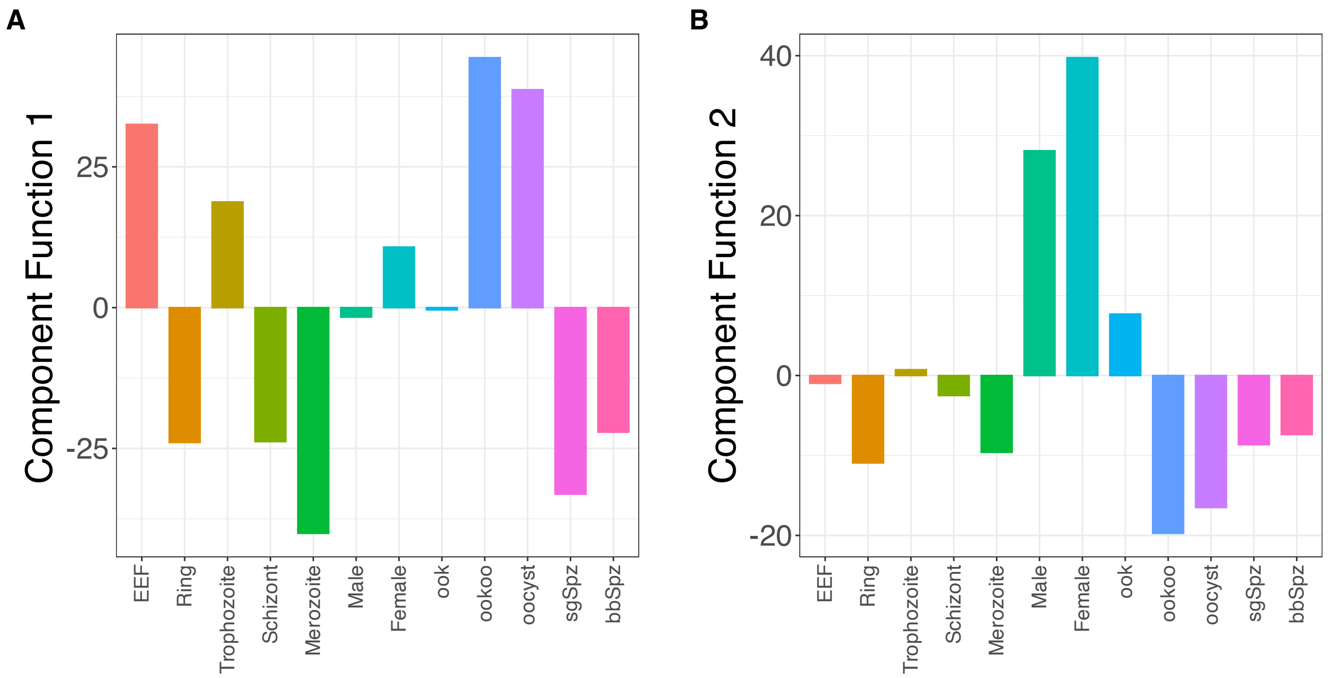


**Supplementary Figure 98.** SSANOVA-SVD reconstructed component functions.(A) Component 1. (B) Component 2.

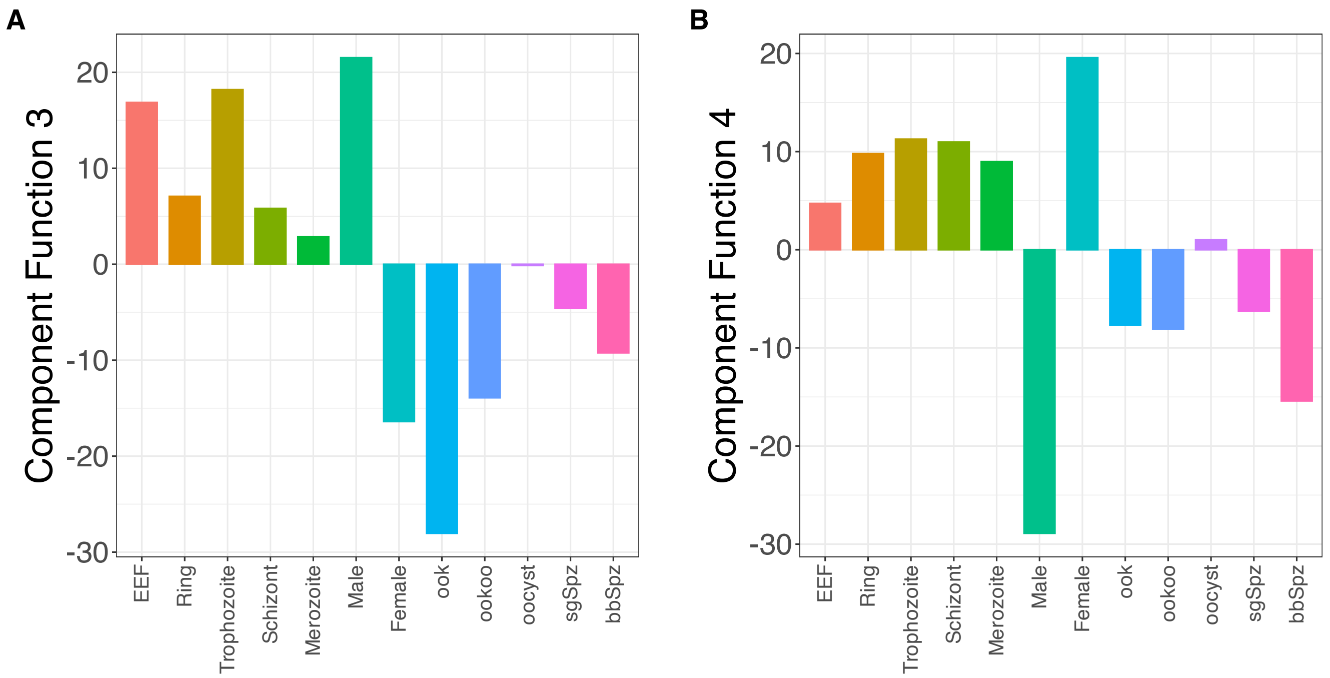


**Supplementary Figure 99.** SSANOVA-SVD reconstructed component functions.(A) Component 3. (B) Component 4.

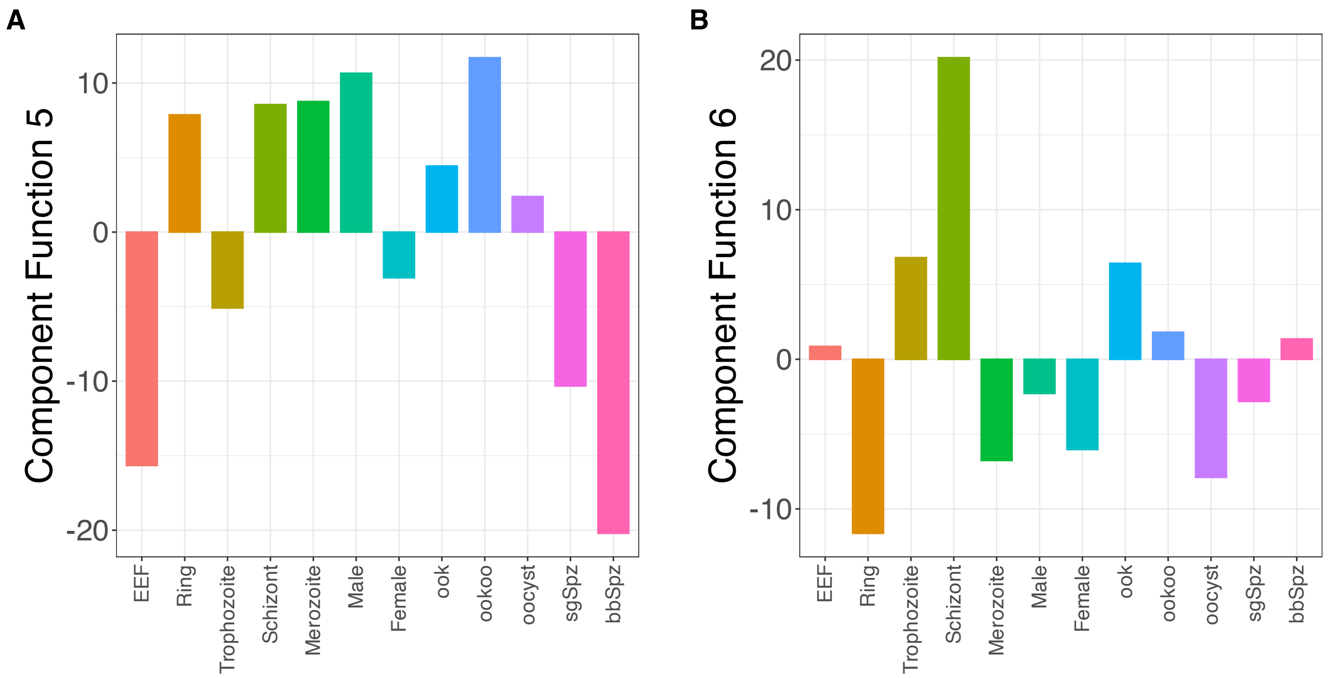


**Supplementary Figure 100.** SSANOVA-SVD reconstructed component functions.(A) Component 5. (B) Component 6.

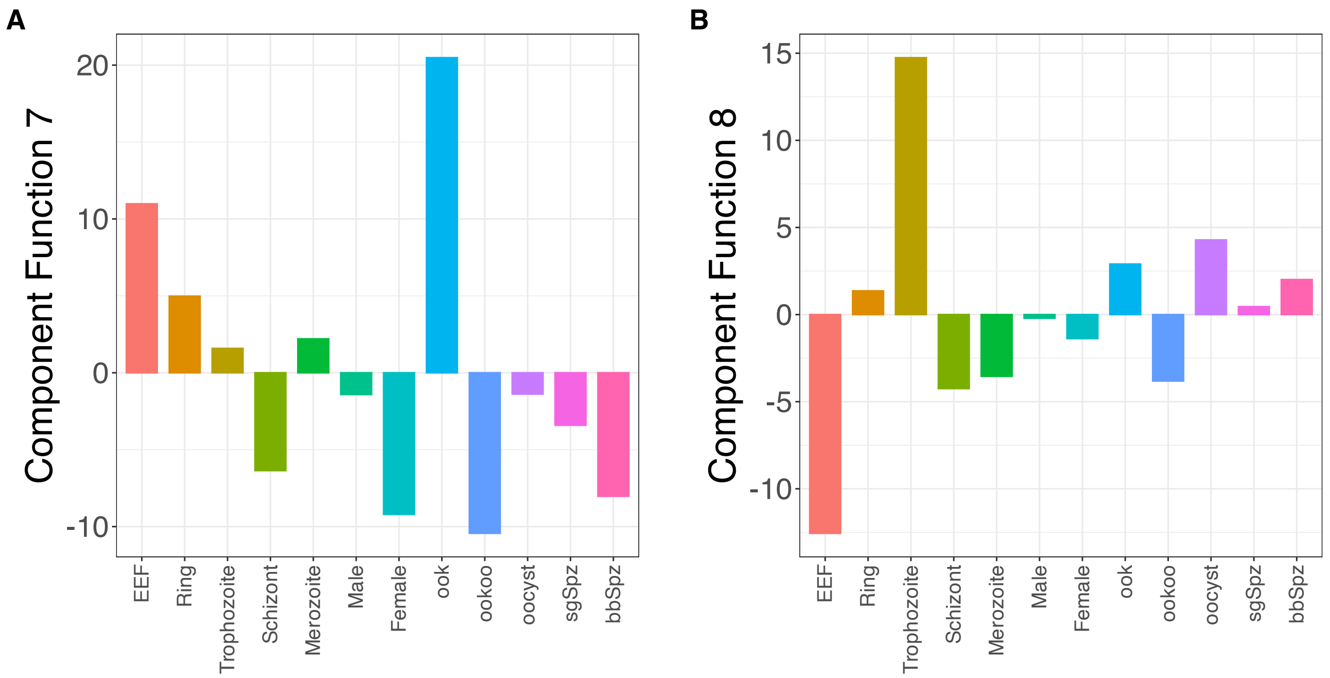


**Supplementary Figure 101.** SSANOVA-SVD reconstructed component functions.(A) Component 7. (B) Component 8.

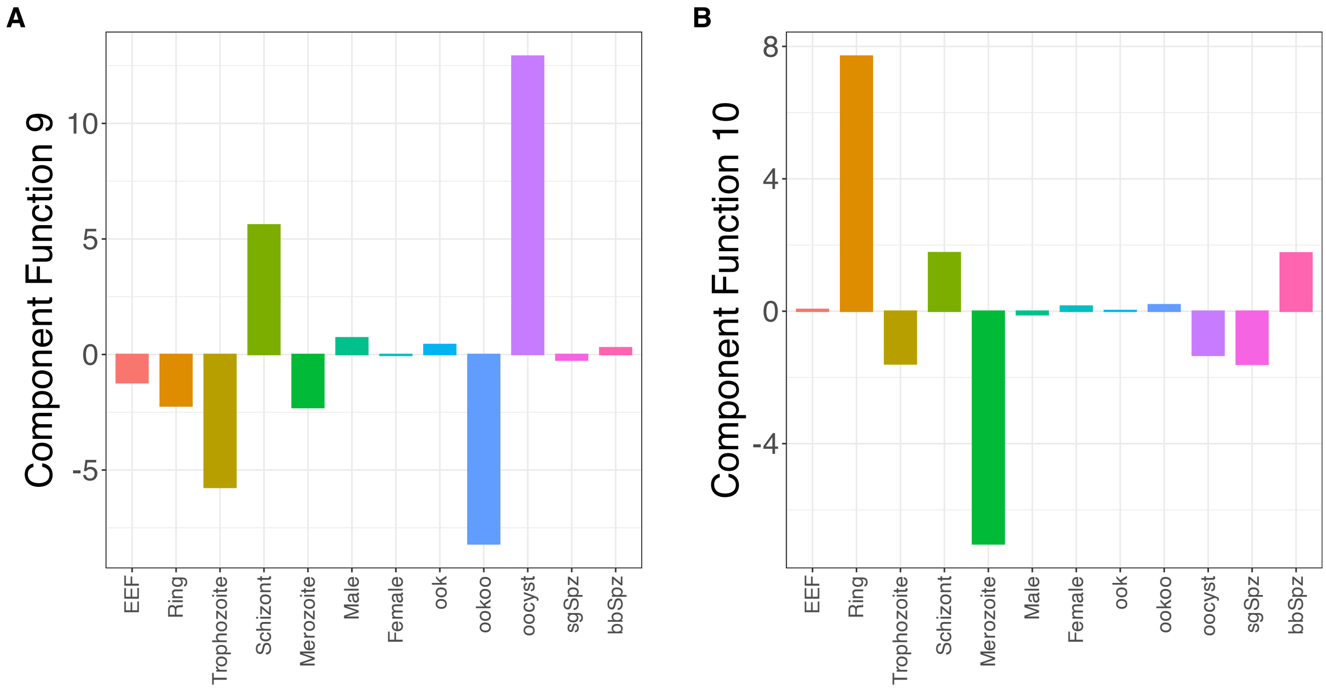


**Supplementary Figure 102.** SSANOVA-SVD reconstructed component functions.(A) Component 9. (B) Component 10.

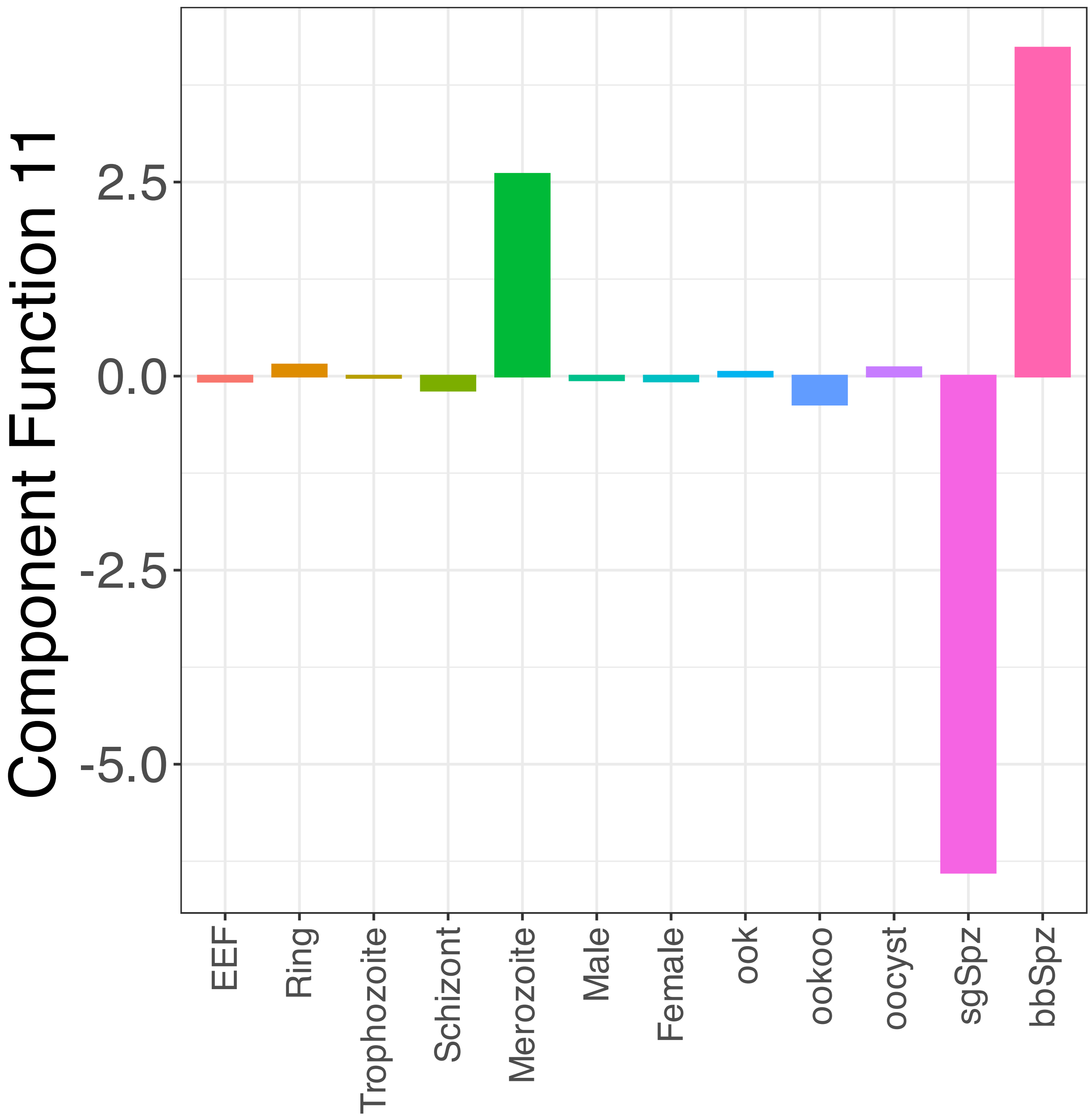


**Supplementary Figure 103.** SSANOVA-SVD reconstructed component functions.Component 11.

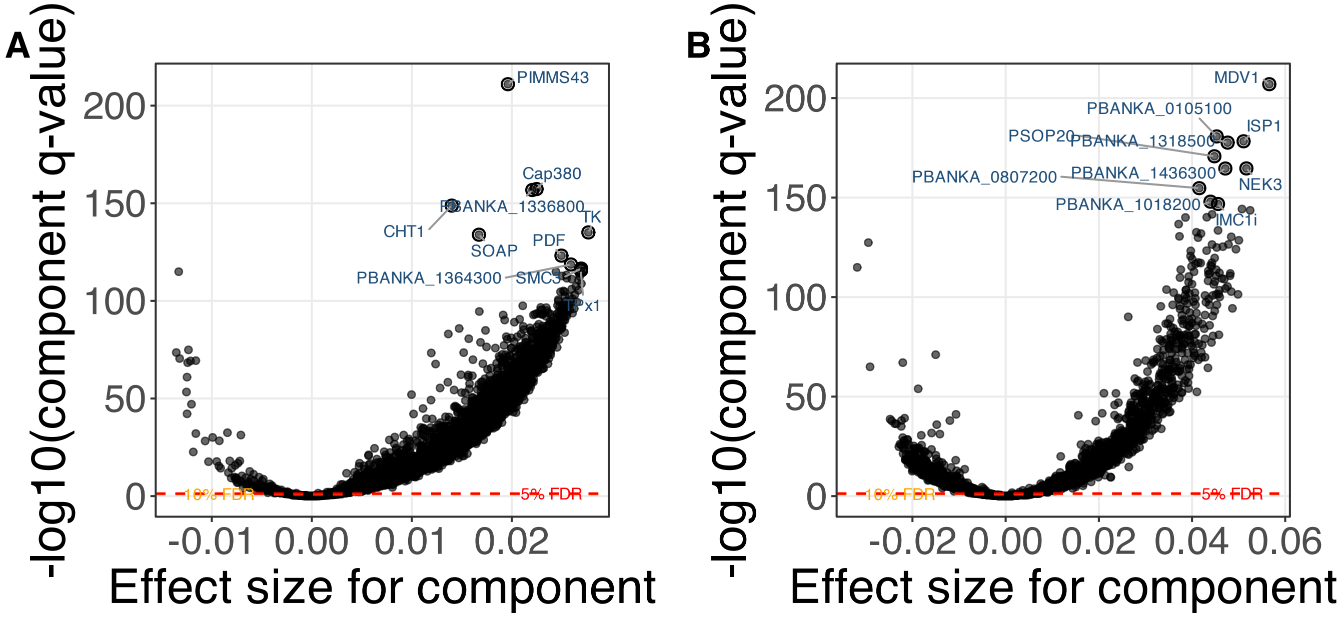


**Supplementary Figure 104.** SSANOVA-SVD based conditional inference for feature participation. (A) Component 1. (B) Component 2.

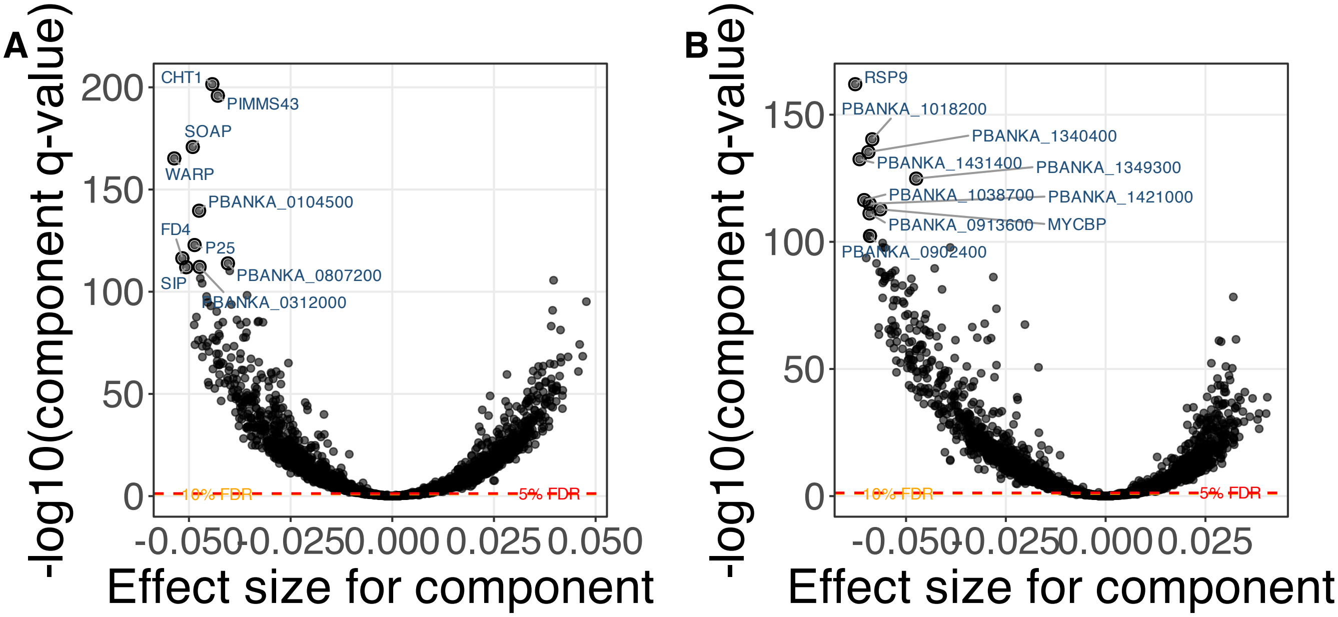


**Supplementary Figure 105.** SSANOVA-SVD based conditional inference for feature participation. (A) Component 3. (B) Component 4.

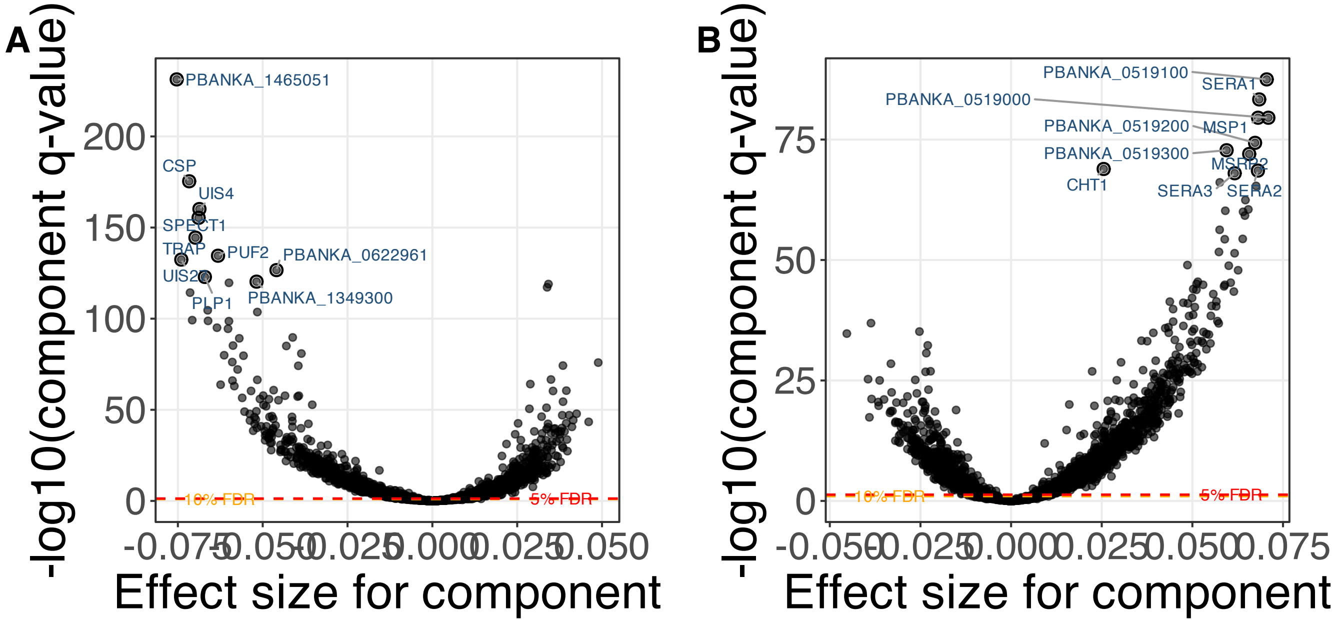


**Supplementary Figure 106.** SSANOVA-SVD based conditional inference for feature participation. (A) Component 5. (B) Component 6.

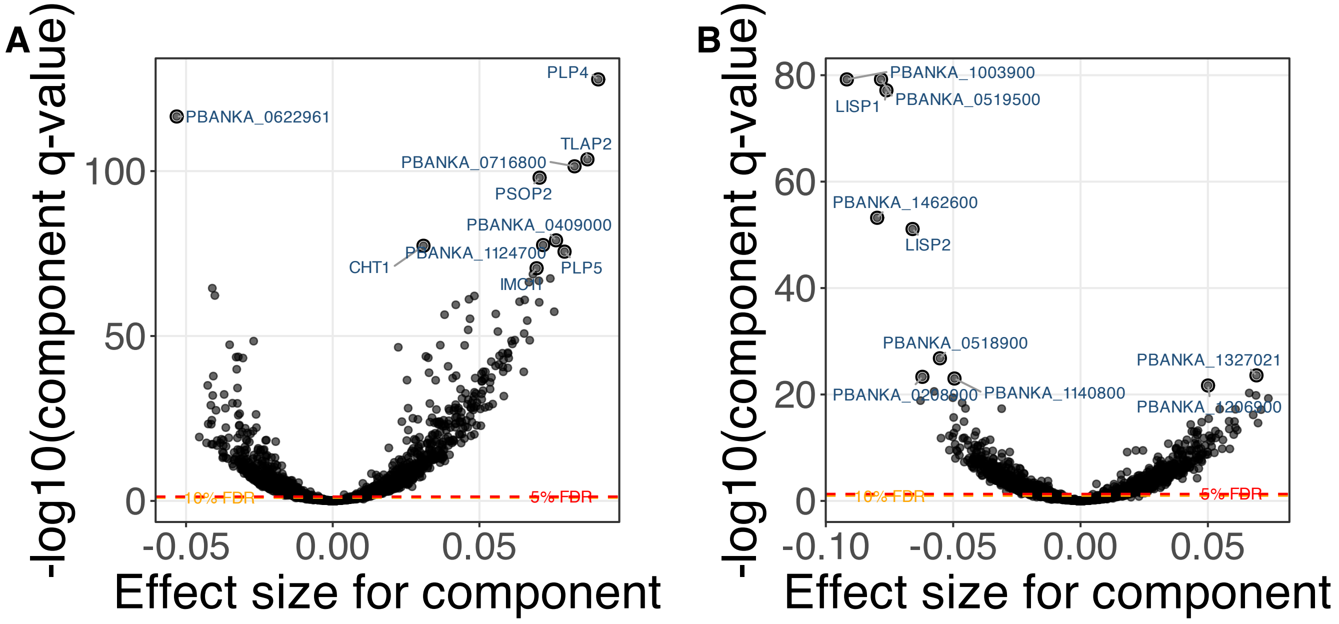


**Supplementary Figure 107.** SSANOVA-SVD based conditional inference for feature participation. (A) Component 7. (B) Component 8.

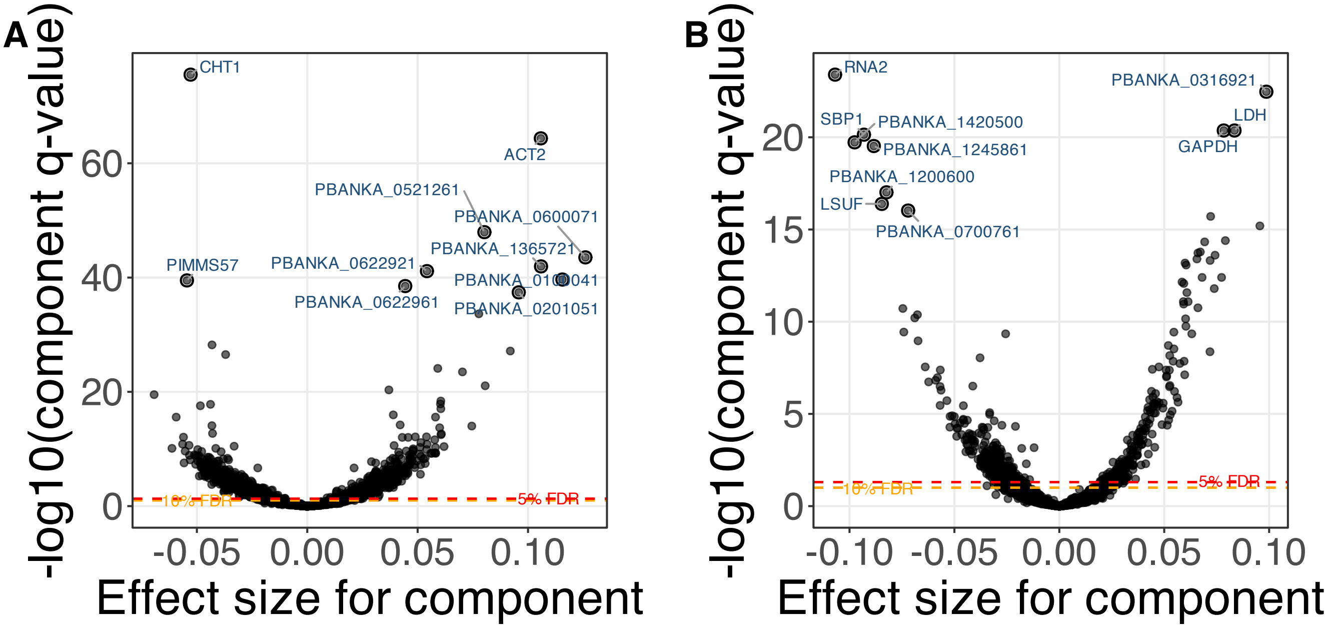


**Supplementary Figure 108.** SSANOVA-SVD based conditional inference for feature participation. (A) Component 9. (B) Component 10.

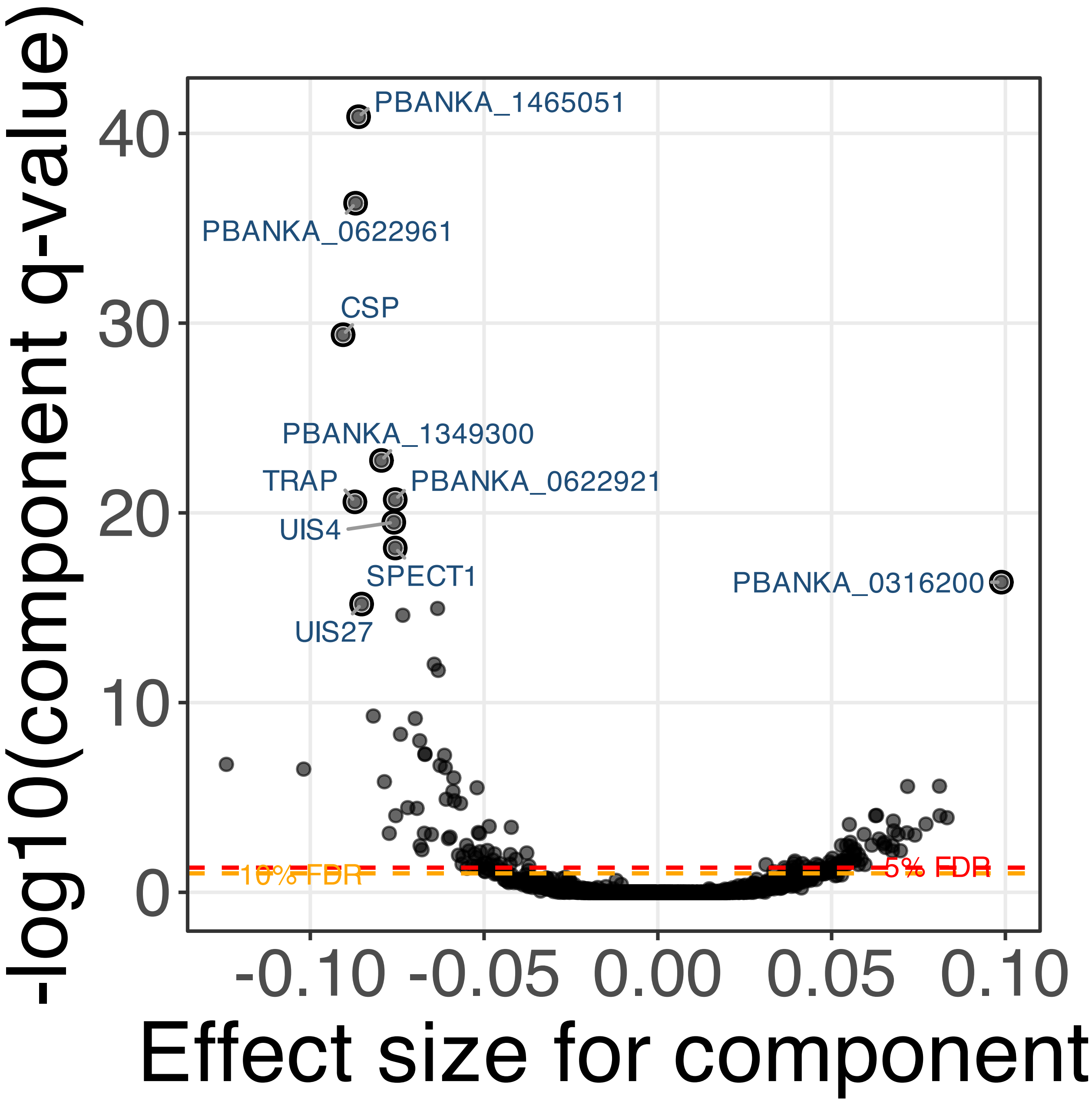


**Supplementary Figure 109.** SSANOVA-SVD based conditional inference for feature participation. Component 11.

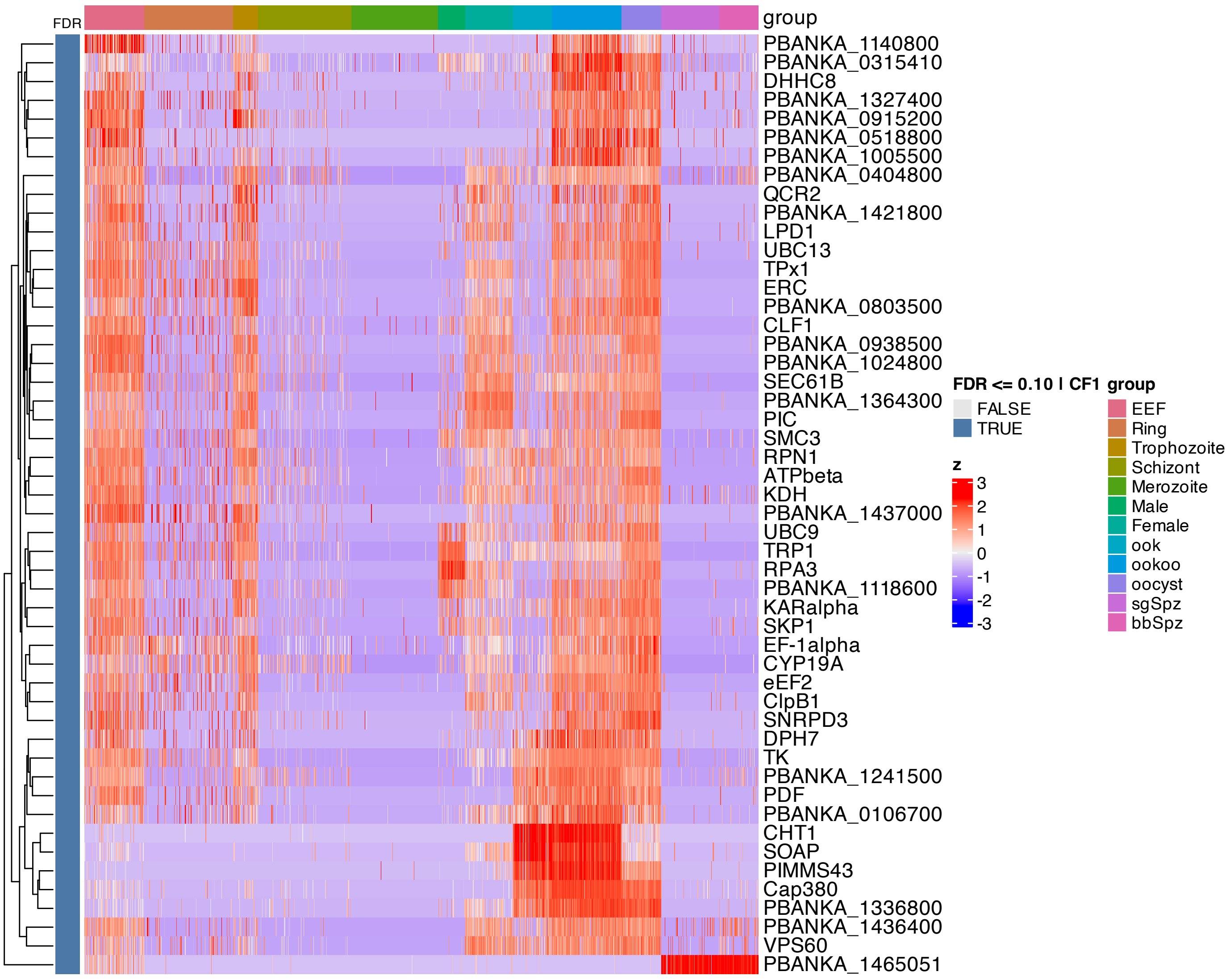


**Supplementary Figure 110.** Heatmap of features inferred to be associated with component 1 by SSANOVA-SVD.

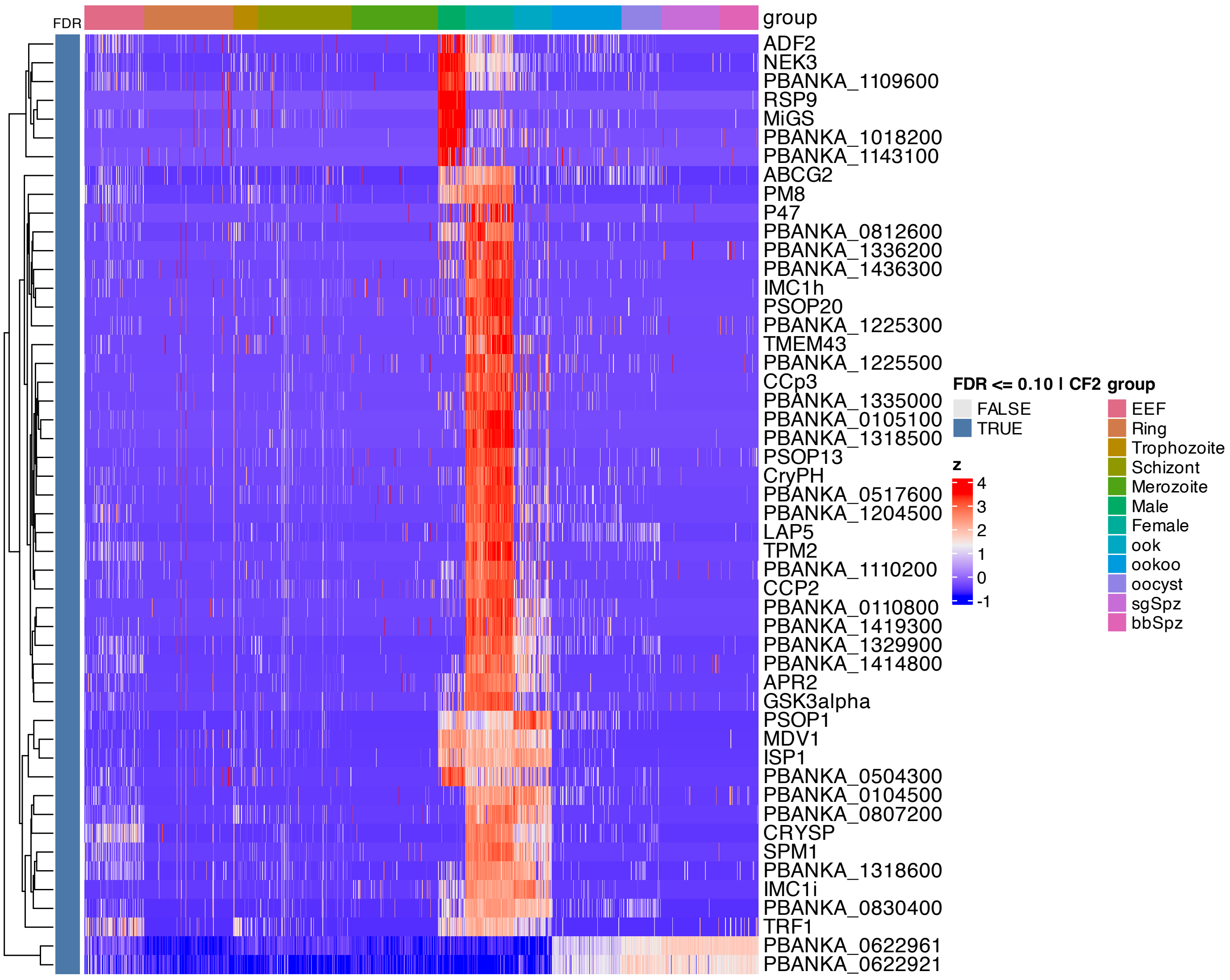


**Supplementary Figure 111.** Heatmap of features inferred to be associated with component 2 by SSANOVA-SVD.

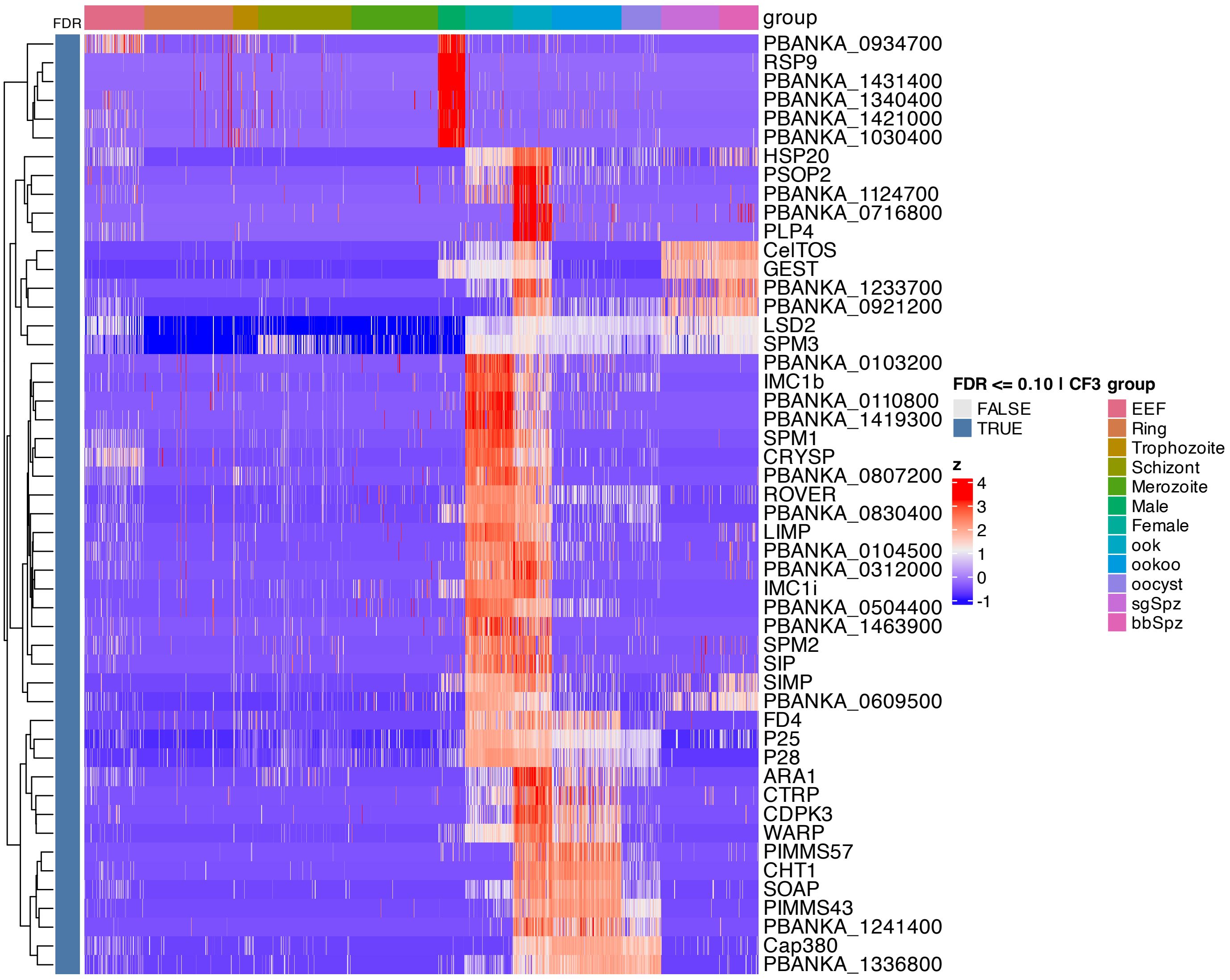


**Supplementary Figure 112.** Heatmap of features inferred to be associated with component 3 by SSANOVA-SVD.

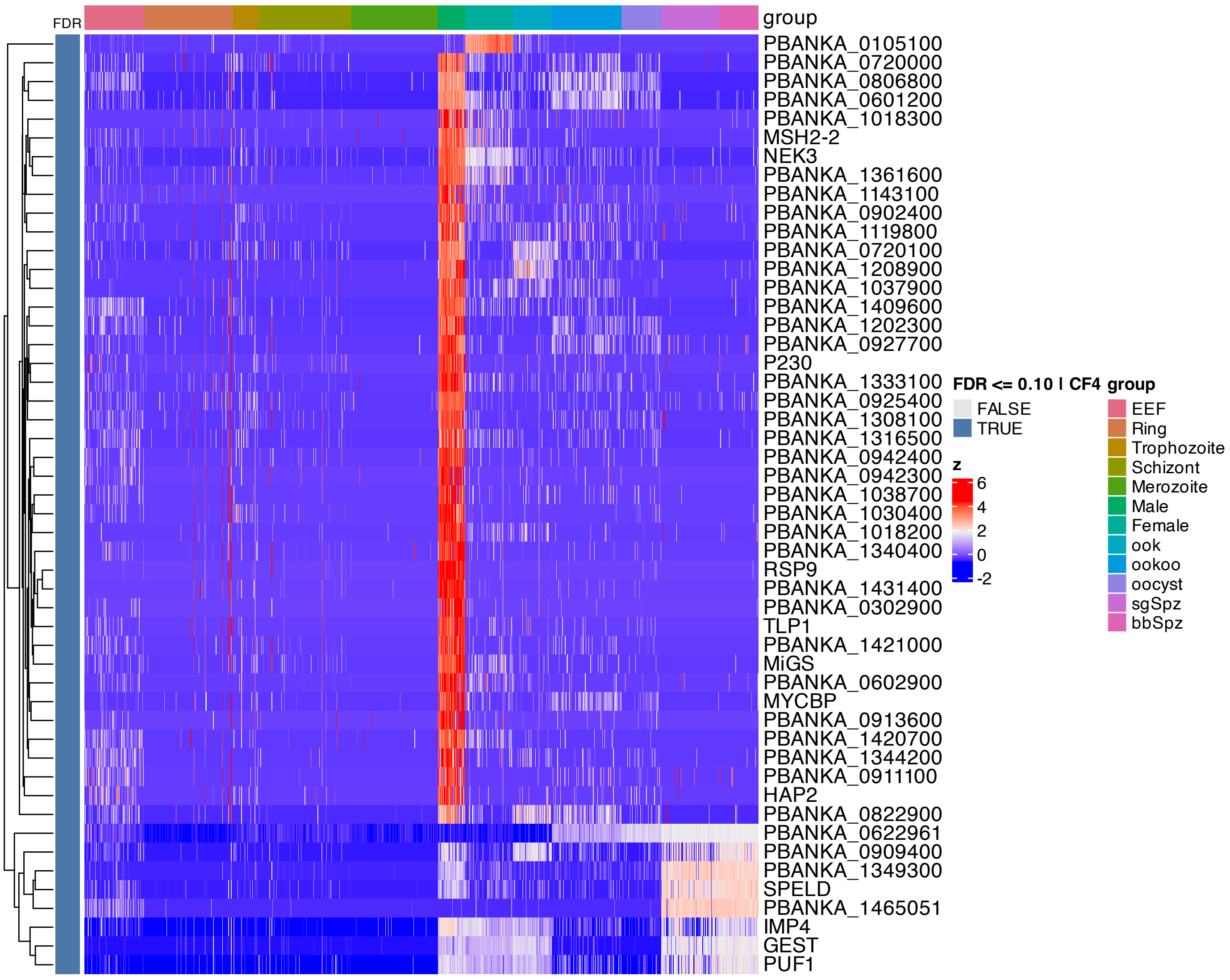


**Supplementary Figure 113.** Heatmap of features inferred to be associated with component 4 by SSANOVA-SVD.

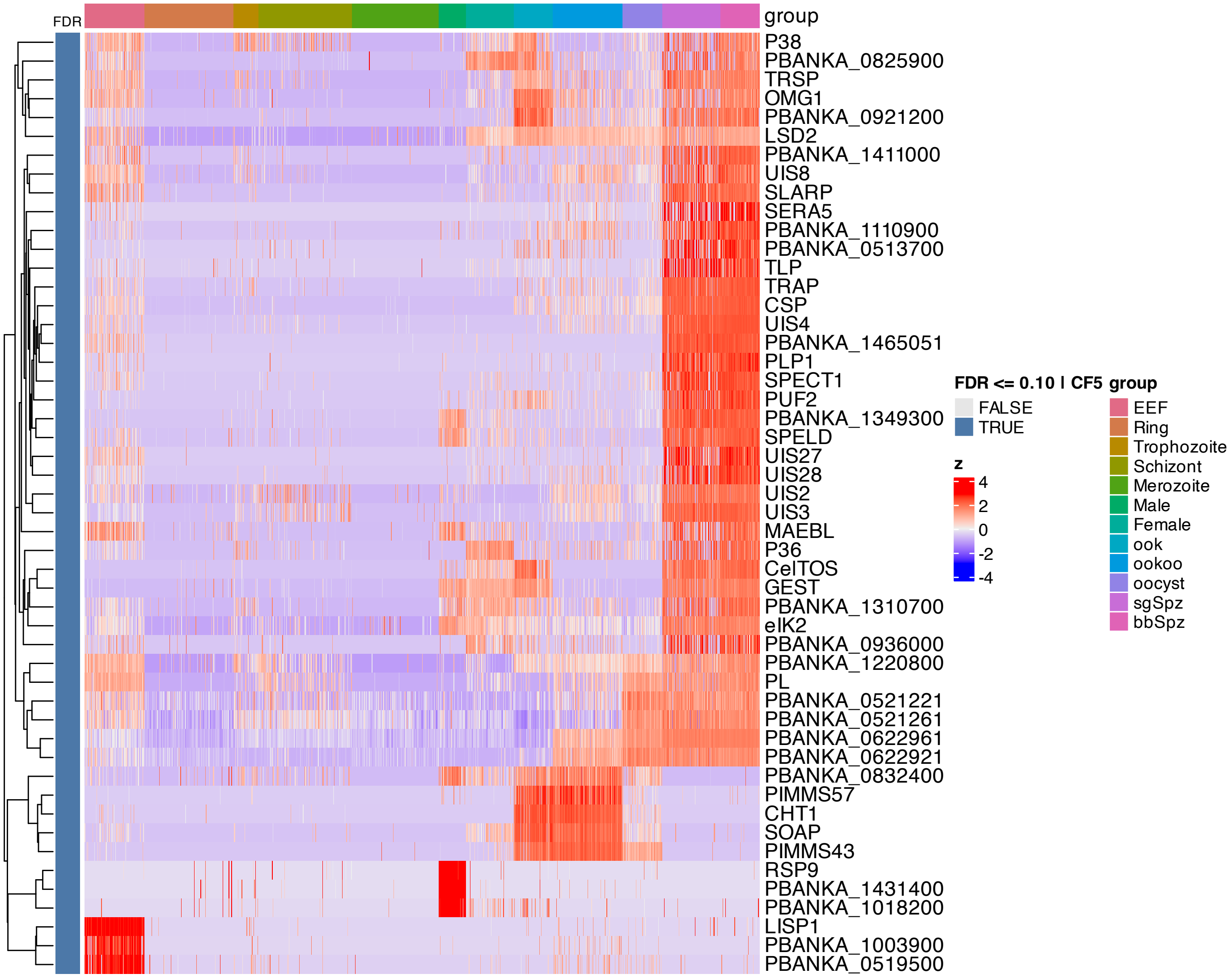


**Supplementary Figure 114.** Heatmap of features inferred to be associated with component 5 by SSANOVA-SVD.

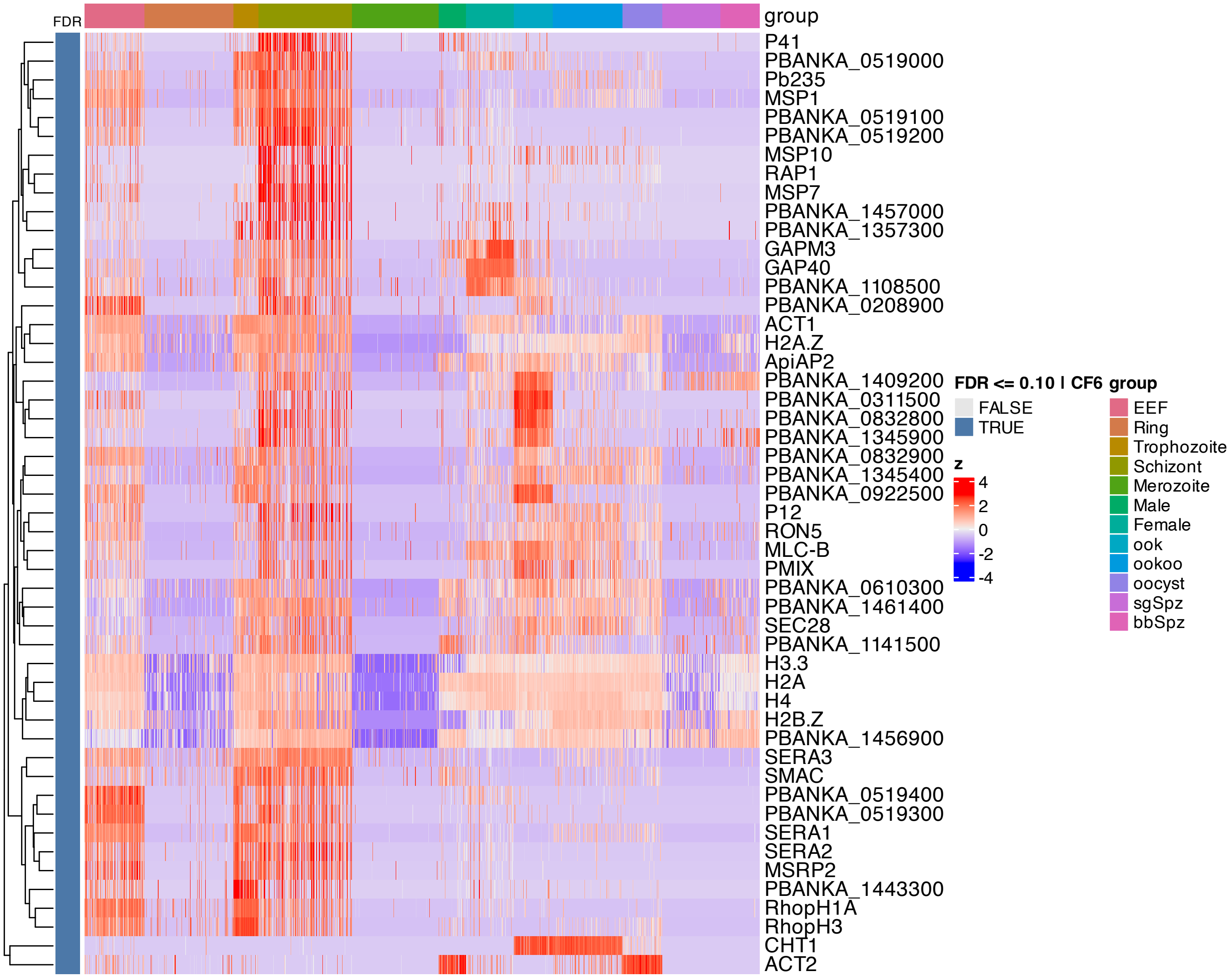


**Supplementary Figure 115.** Heatmap of features inferred to be associated with component 6 by SSANOVA-SVD.

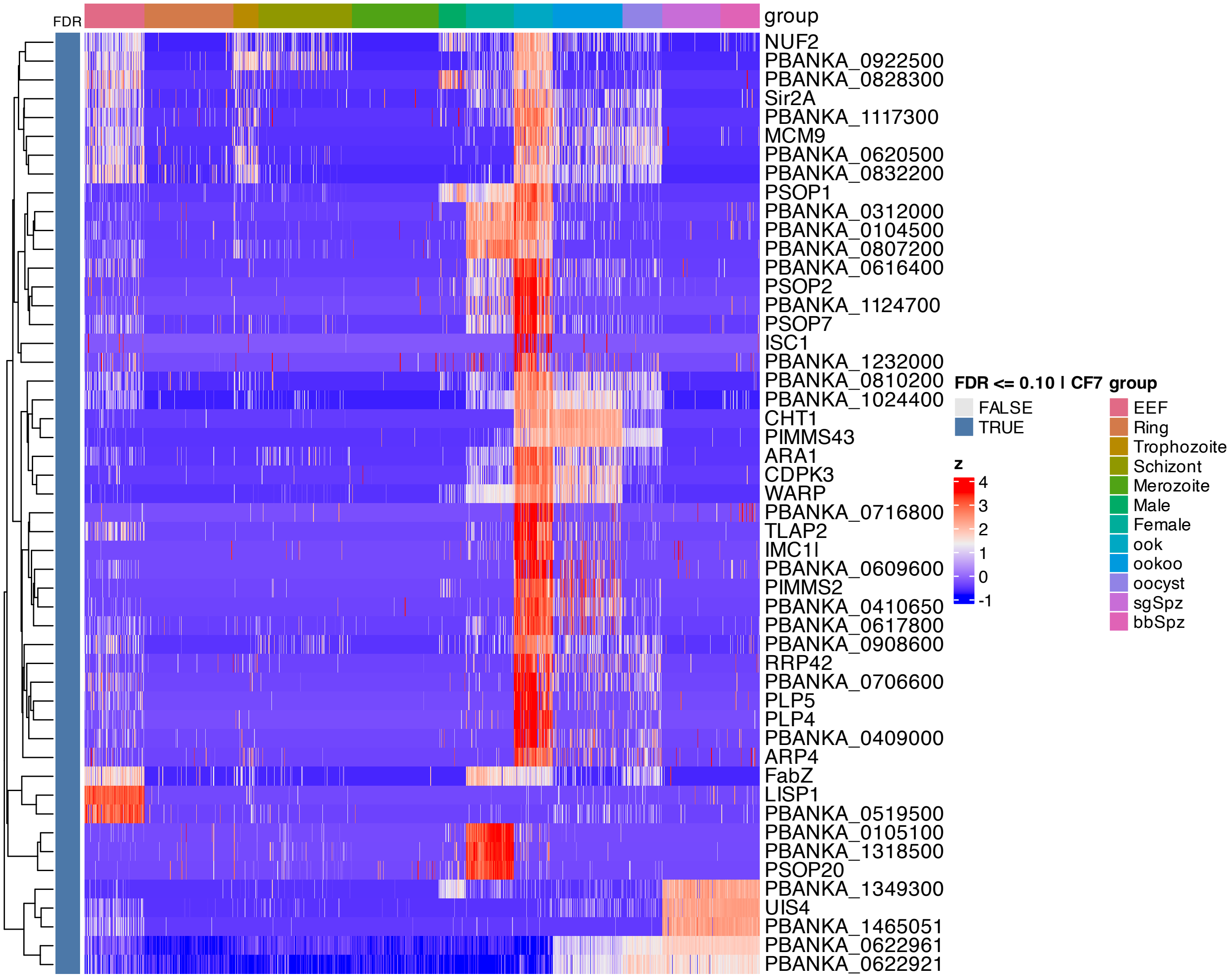


**Supplementary Figure 116.** Heatmap of features inferred to be associated with component 7 by SSANOVA-SVD.

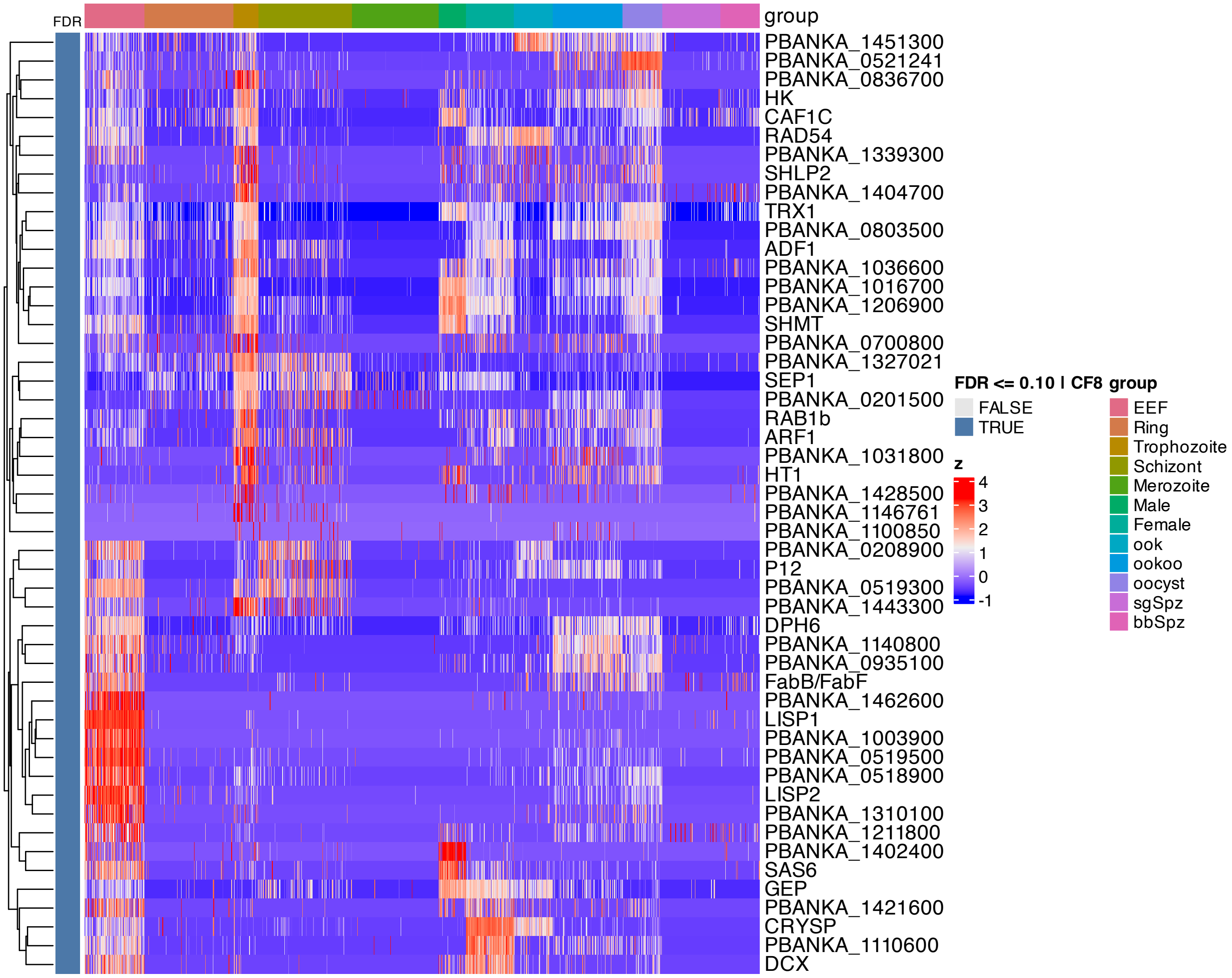


**Supplementary Figure 117.** Heatmap of features inferred to be associated with component 8 by SSANOVA-SVD.

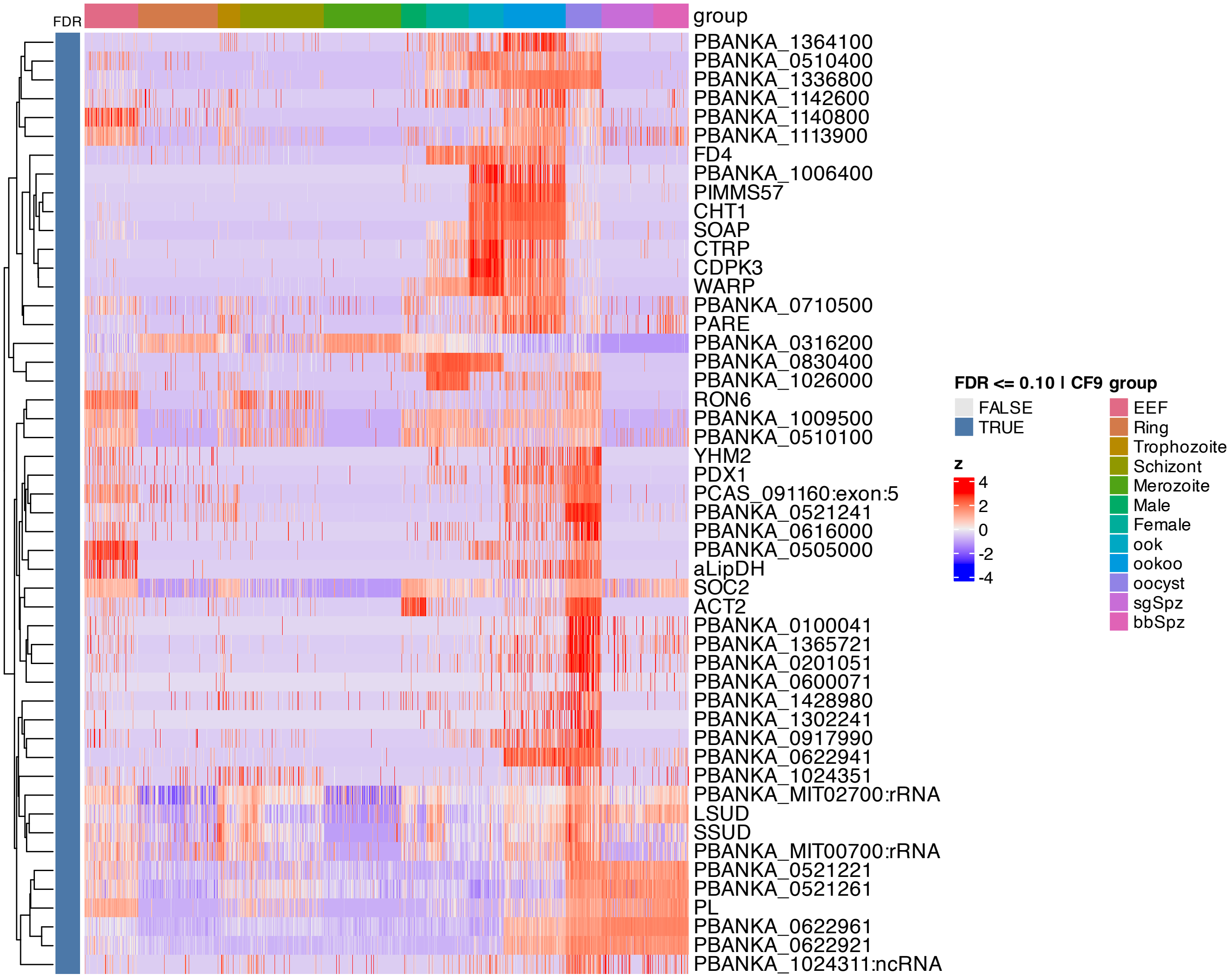


**Supplementary Figure 118.** Heatmap of features inferred to be associated with component 9 by SSANOVA-SVD.

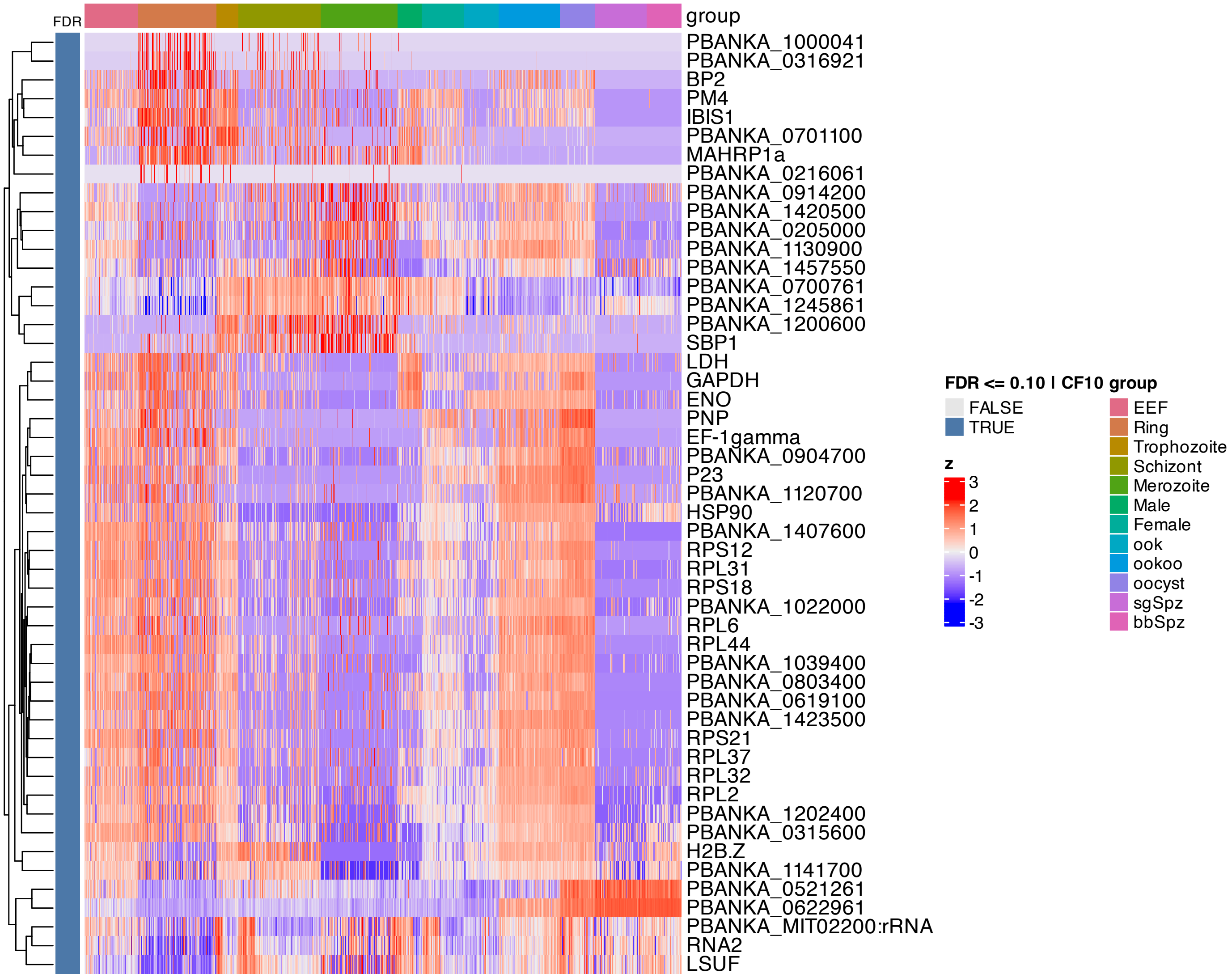


**Supplementary Figure 119.** Heatmap of features inferred to be associated with component 10 by SSANOVA-SVD.

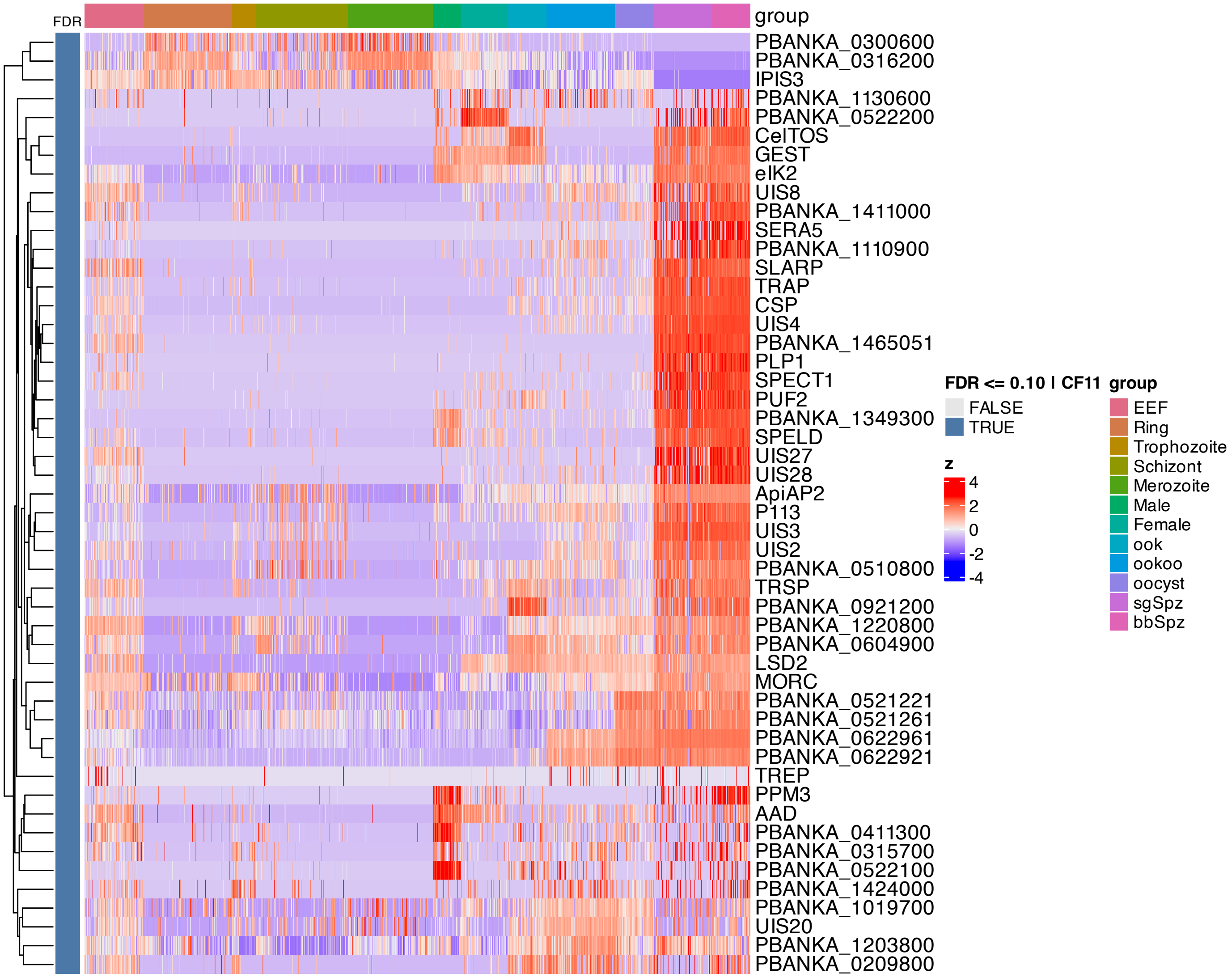


**Supplementary Figure 120.** Heatmap of features inferred to be associated with component 11 by SSANOVA-SVD.

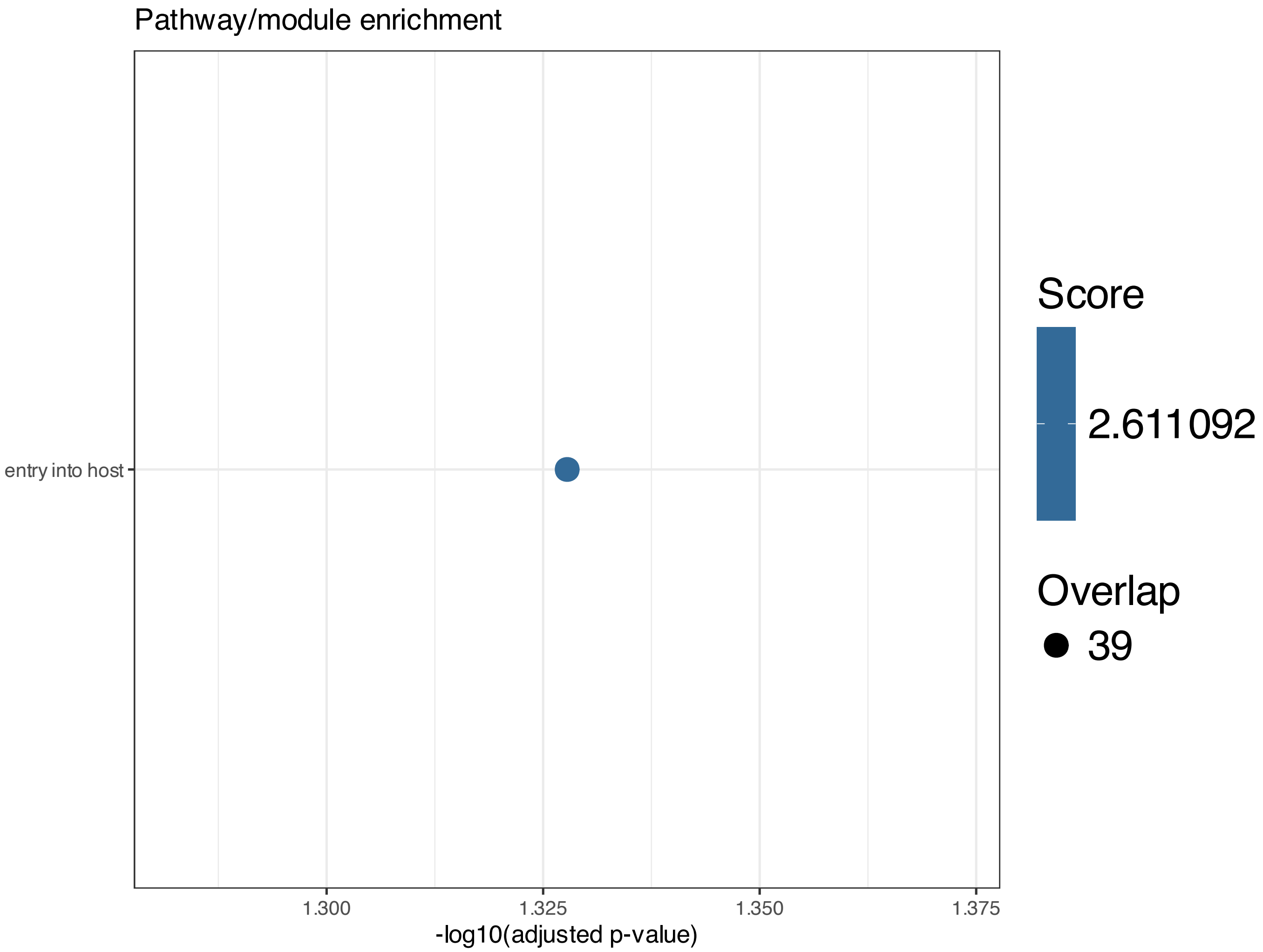


**Supplementary Figure 121.** Enrichment among features inferred to be associated with SSANOVA-SVD component 7; only terms with adjusted P $\leq 0.1$ are shown.

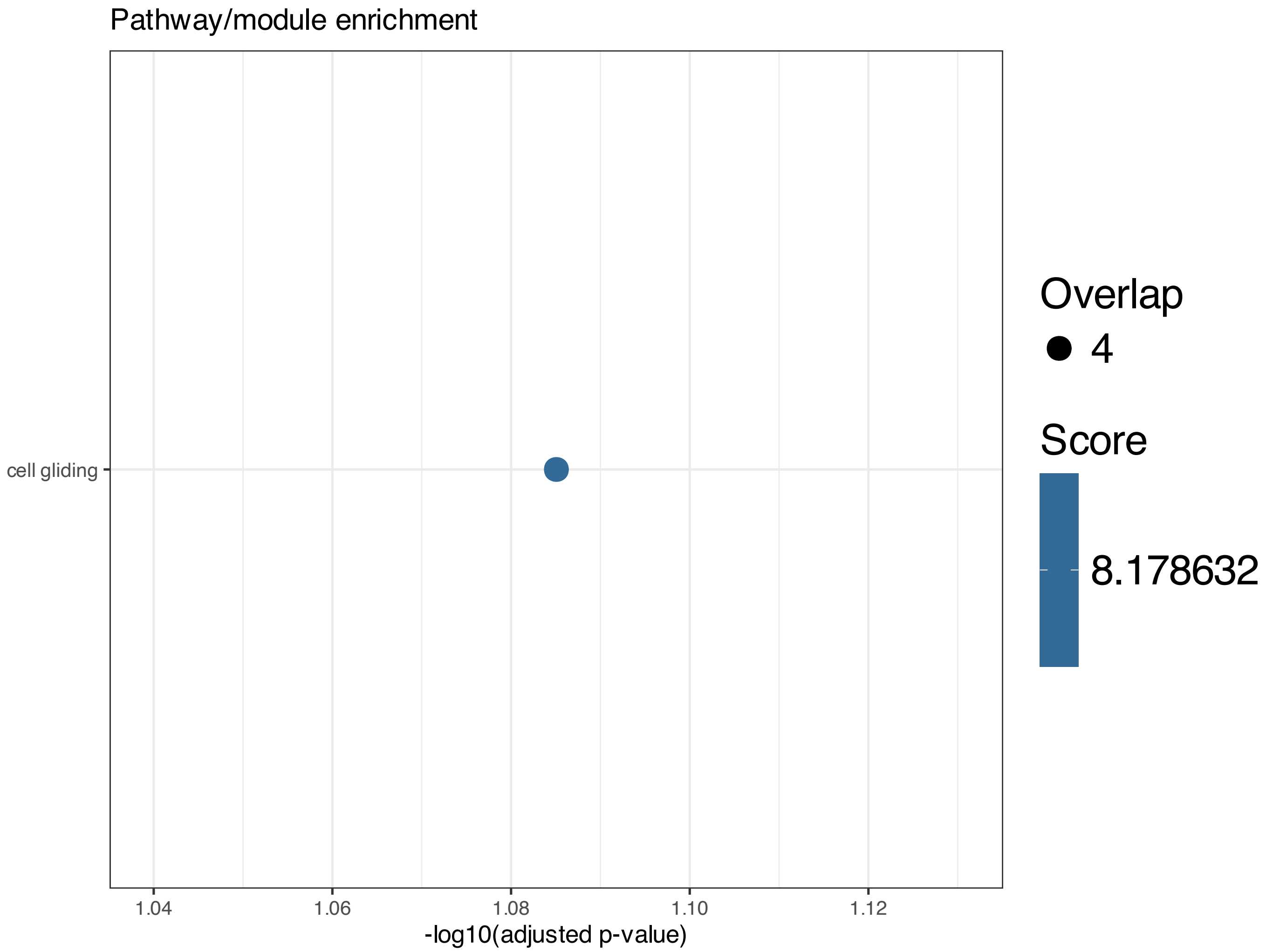


**Supplementary Figure 122.** Enrichment among features inferred to be associated with SSANOVA-SVD component 11; only terms with adjusted P $\leq 0.1$ are shown.

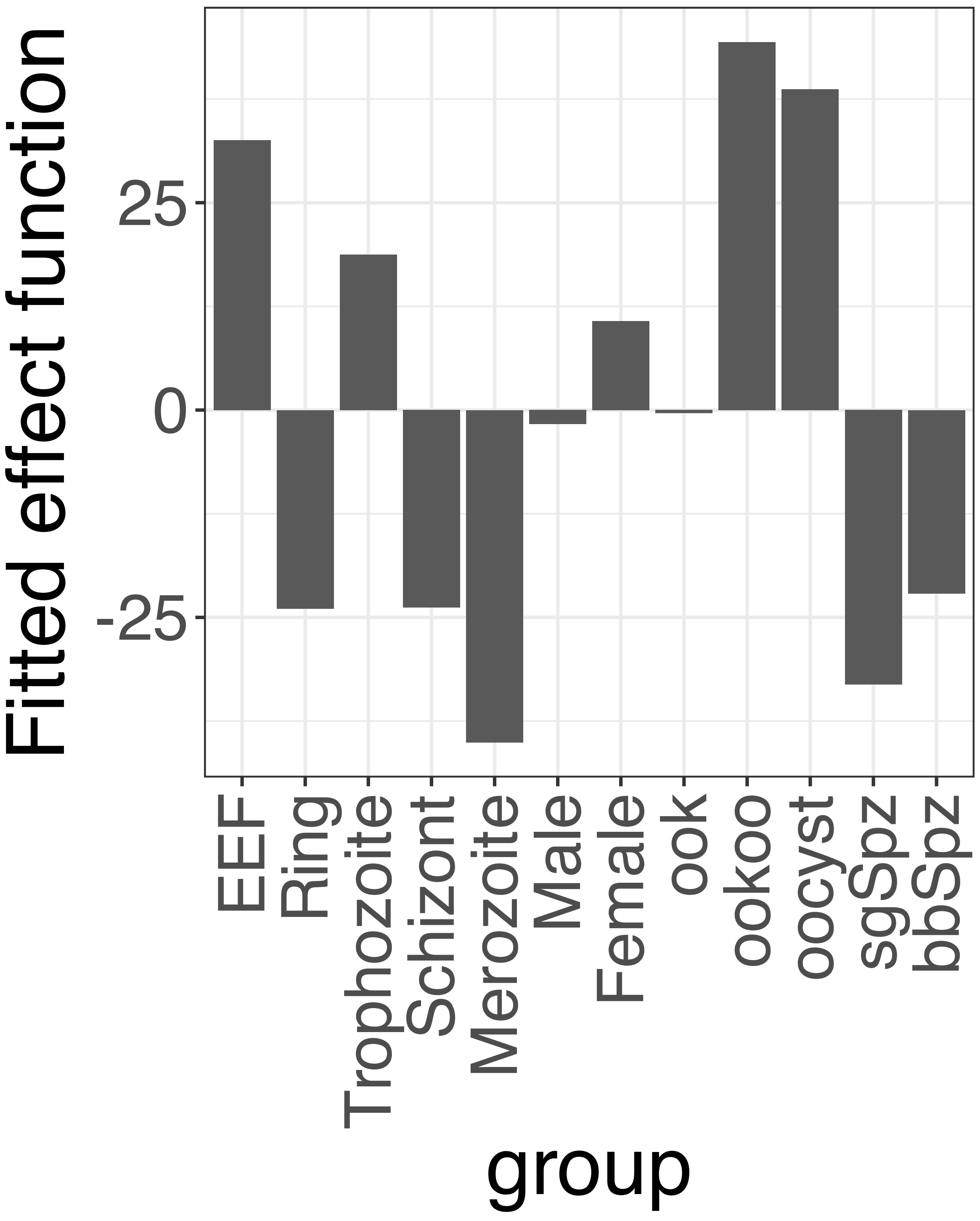


**Supplementary Figure 123.** Decomposition of reconstructed SSANOVA-SVD component function 1.

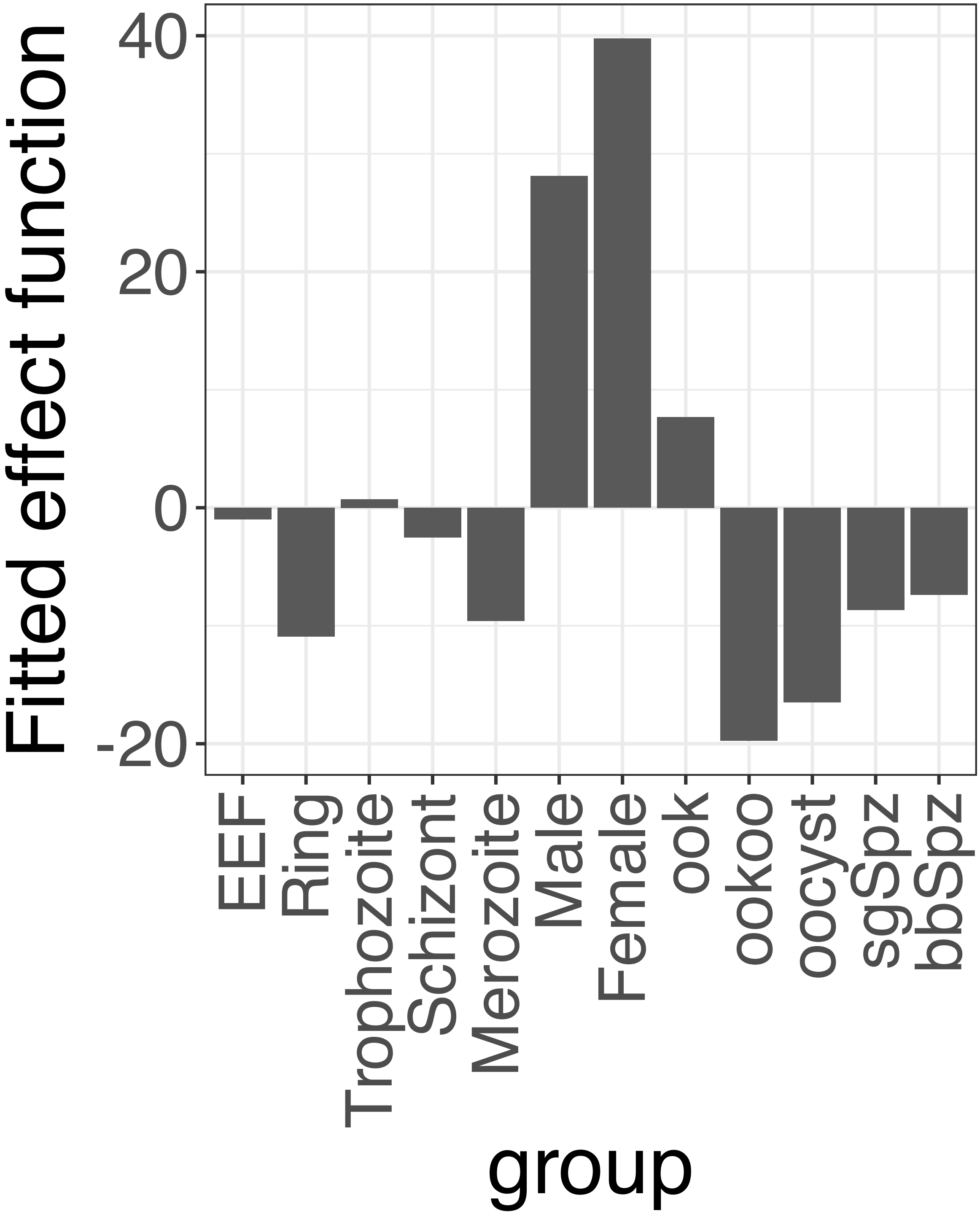


**Supplementary Figure 124.** Decomposition of reconstructed SSANOVA-SVD component function 2.

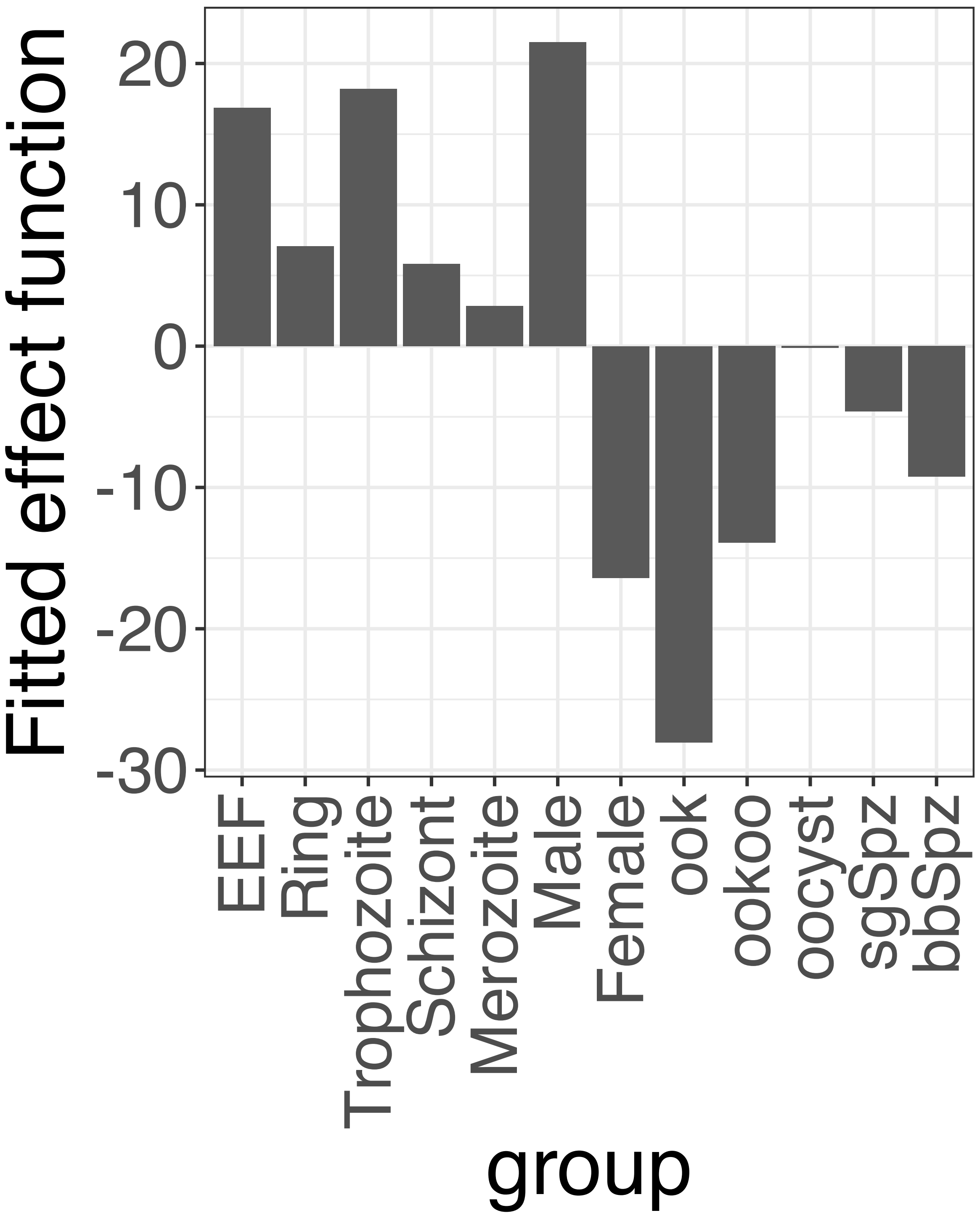


**Supplementary Figure 125.** Decomposition of reconstructed SSANOVA-SVD component function 3.

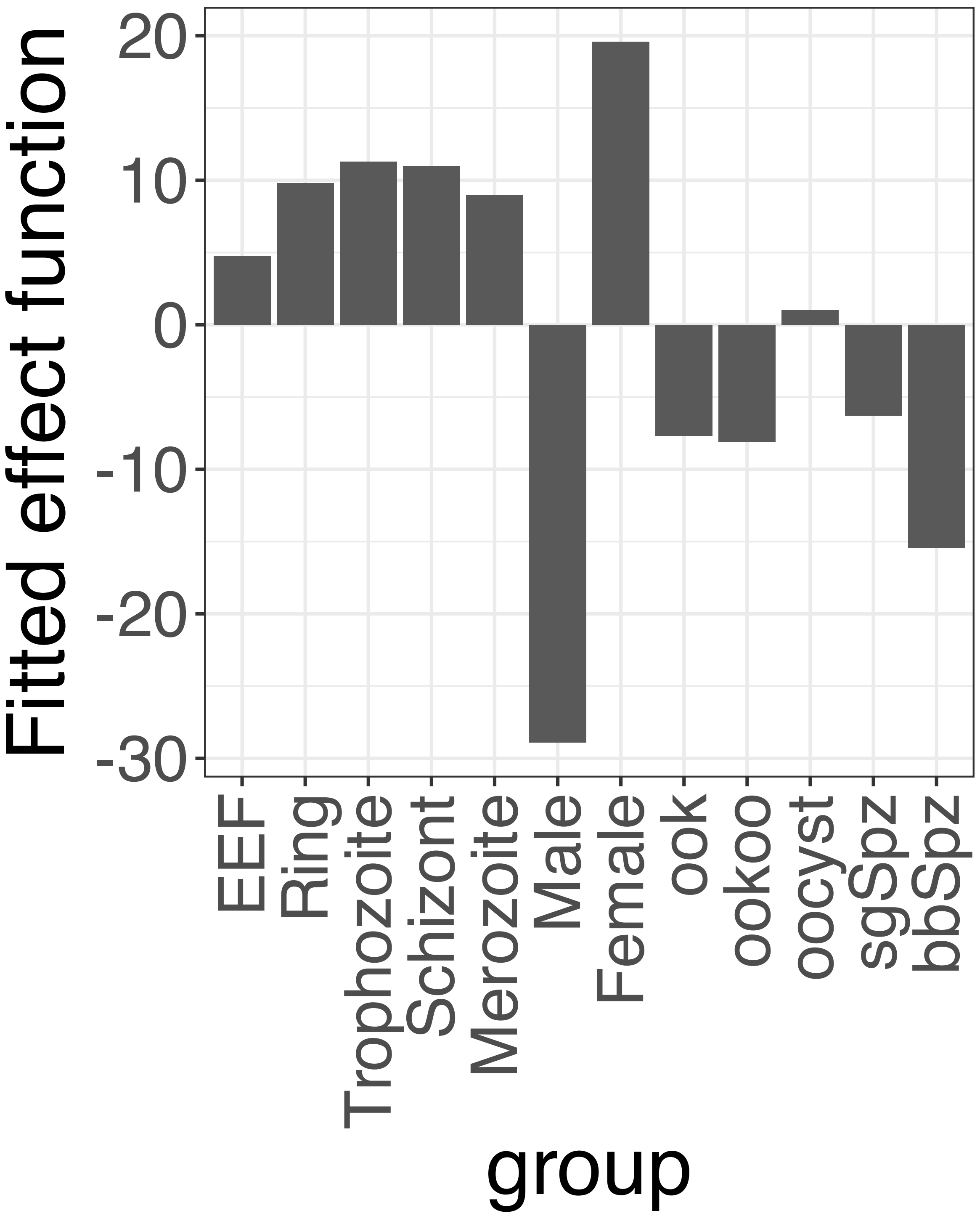


**Supplementary Figure 126.** Decomposition of reconstructed SSANOVA-SVD component function 4.

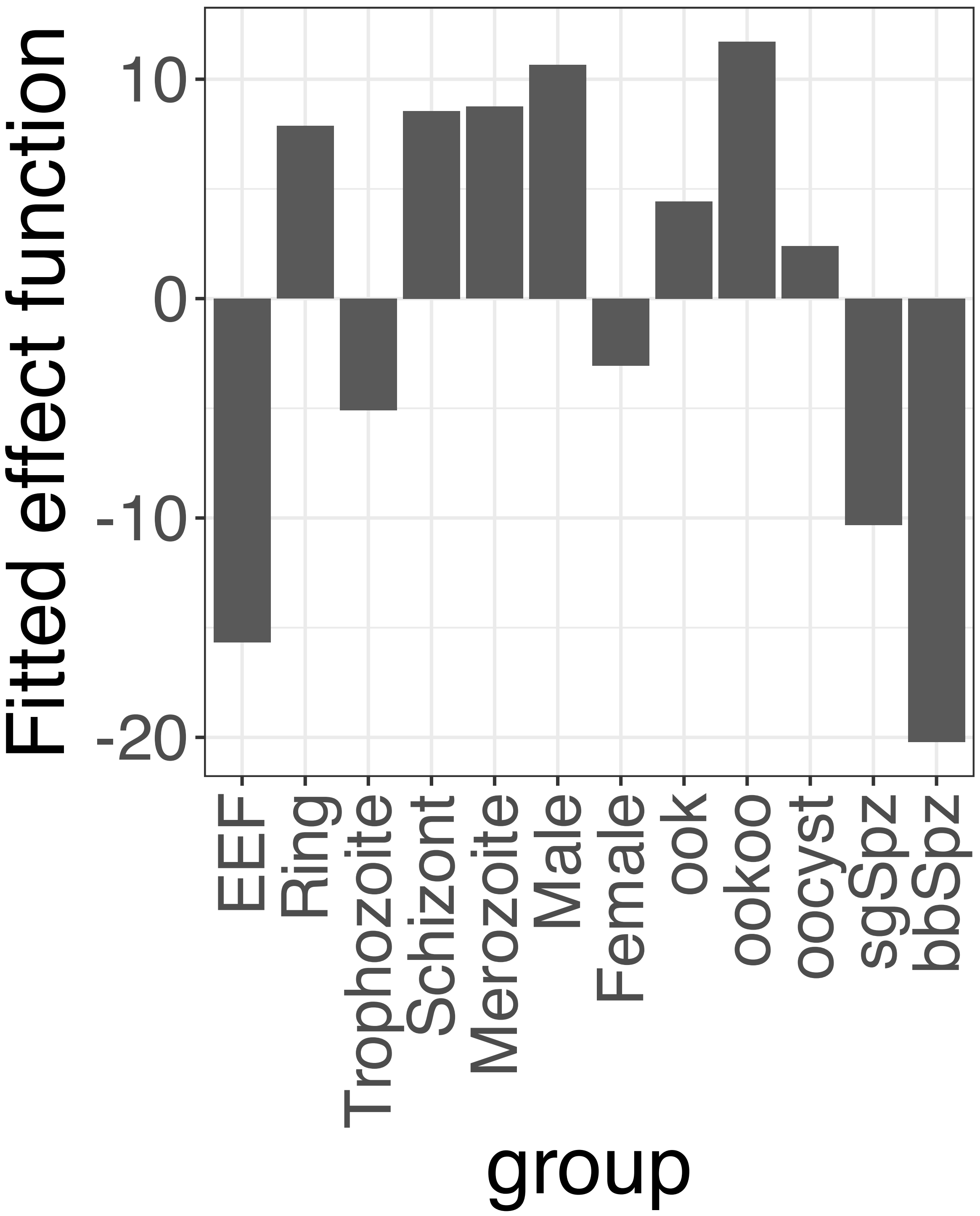


**Supplementary Figure 127.** Decomposition of reconstructed SSANOVA-SVD component function 5.

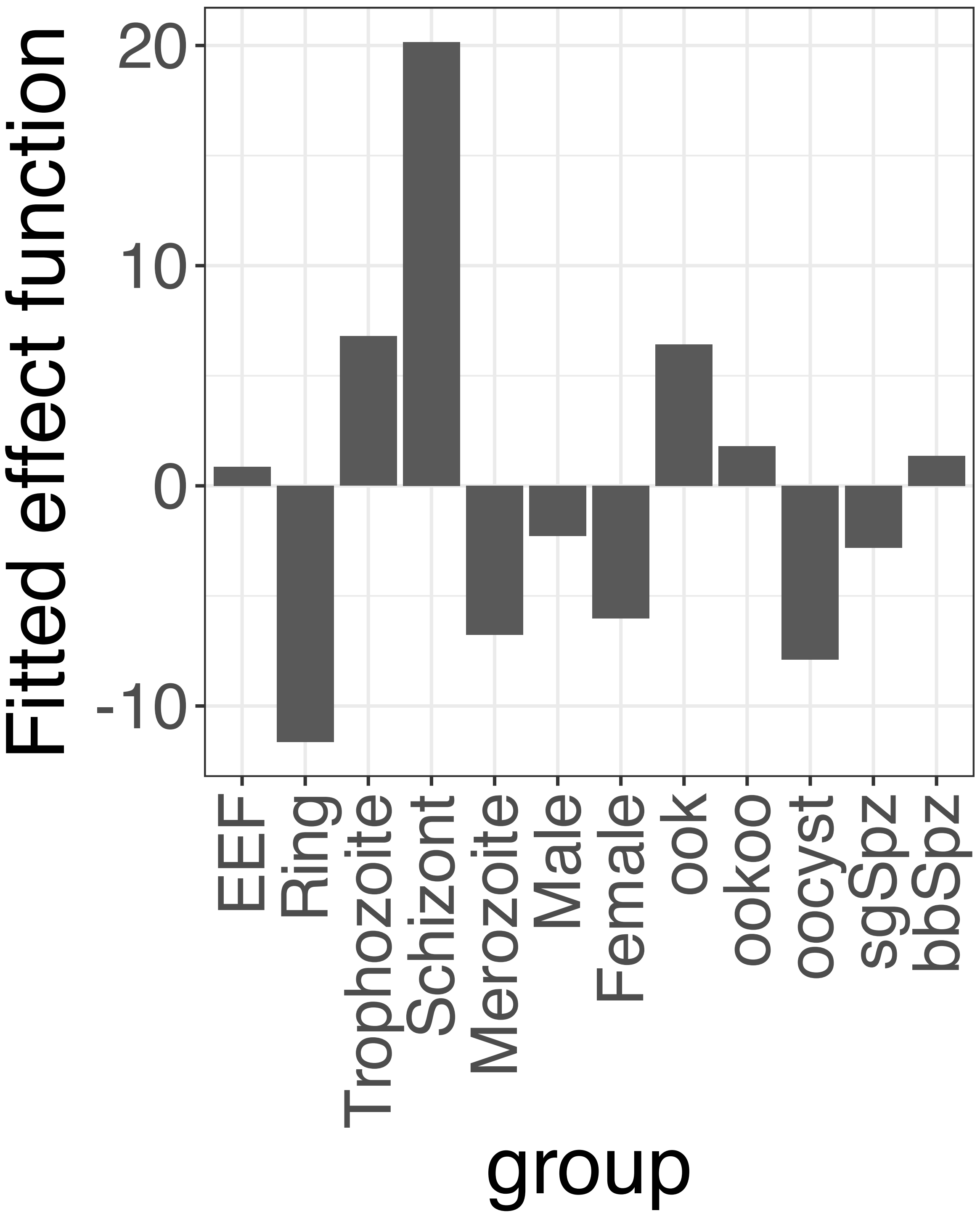


**Supplementary Figure 128.** Decomposition of reconstructed SSANOVA-SVD component function 6.

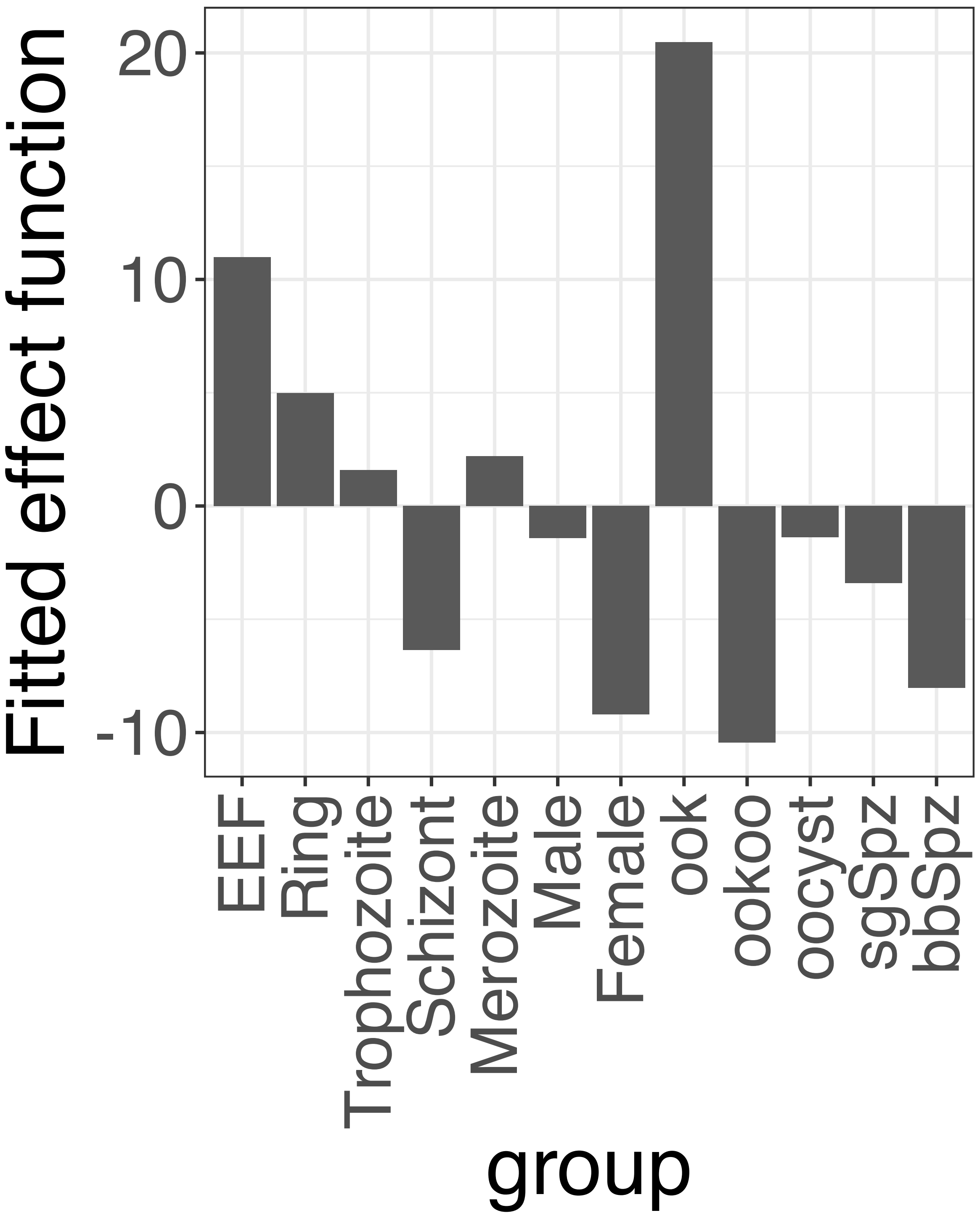


**Supplementary Figure 129.** Decomposition of reconstructed SSANOVA-SVD component function 7.

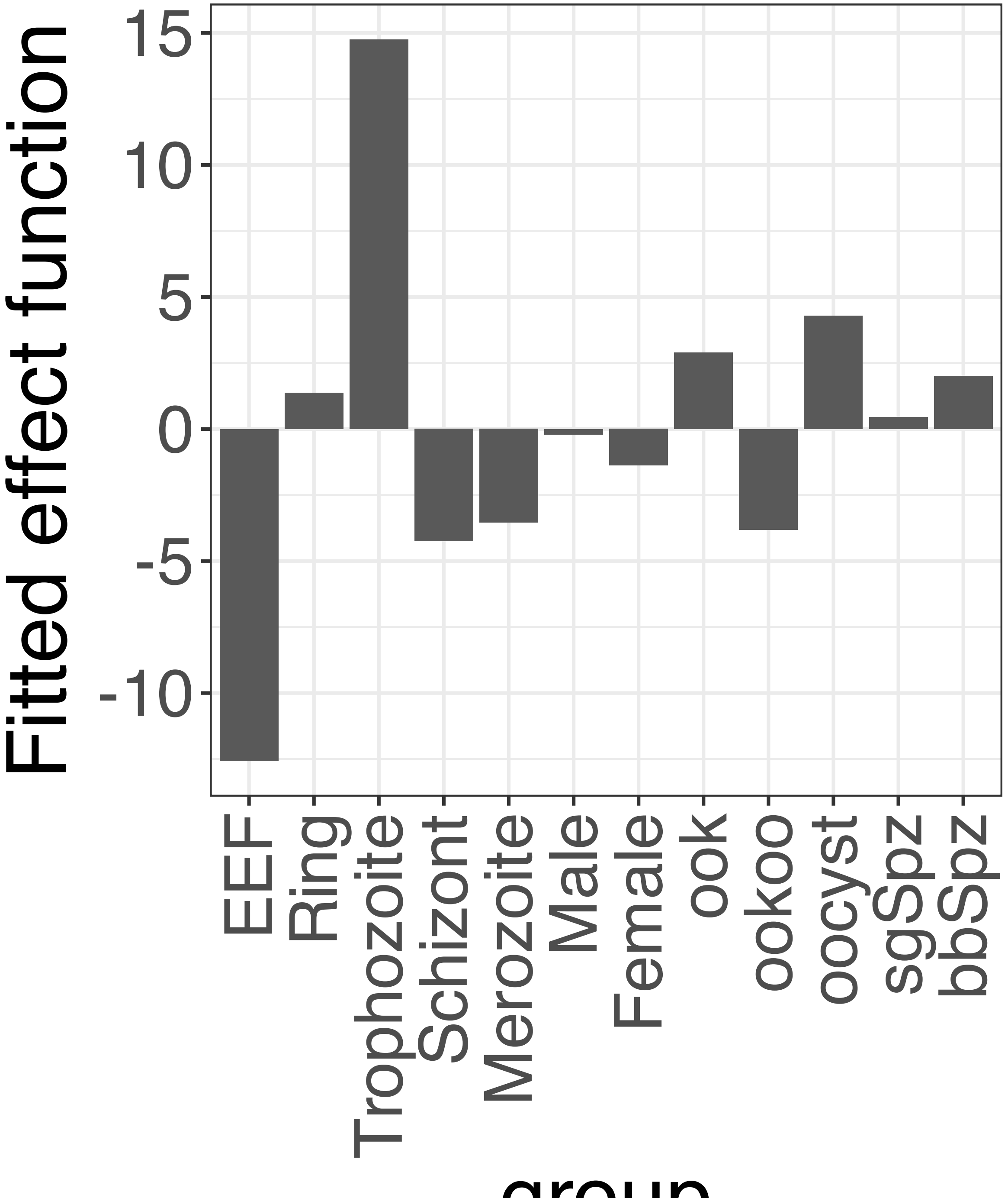


**Supplementary Figure 130.** Decomposition of reconstructed SSANOVA-SVD component function 8.

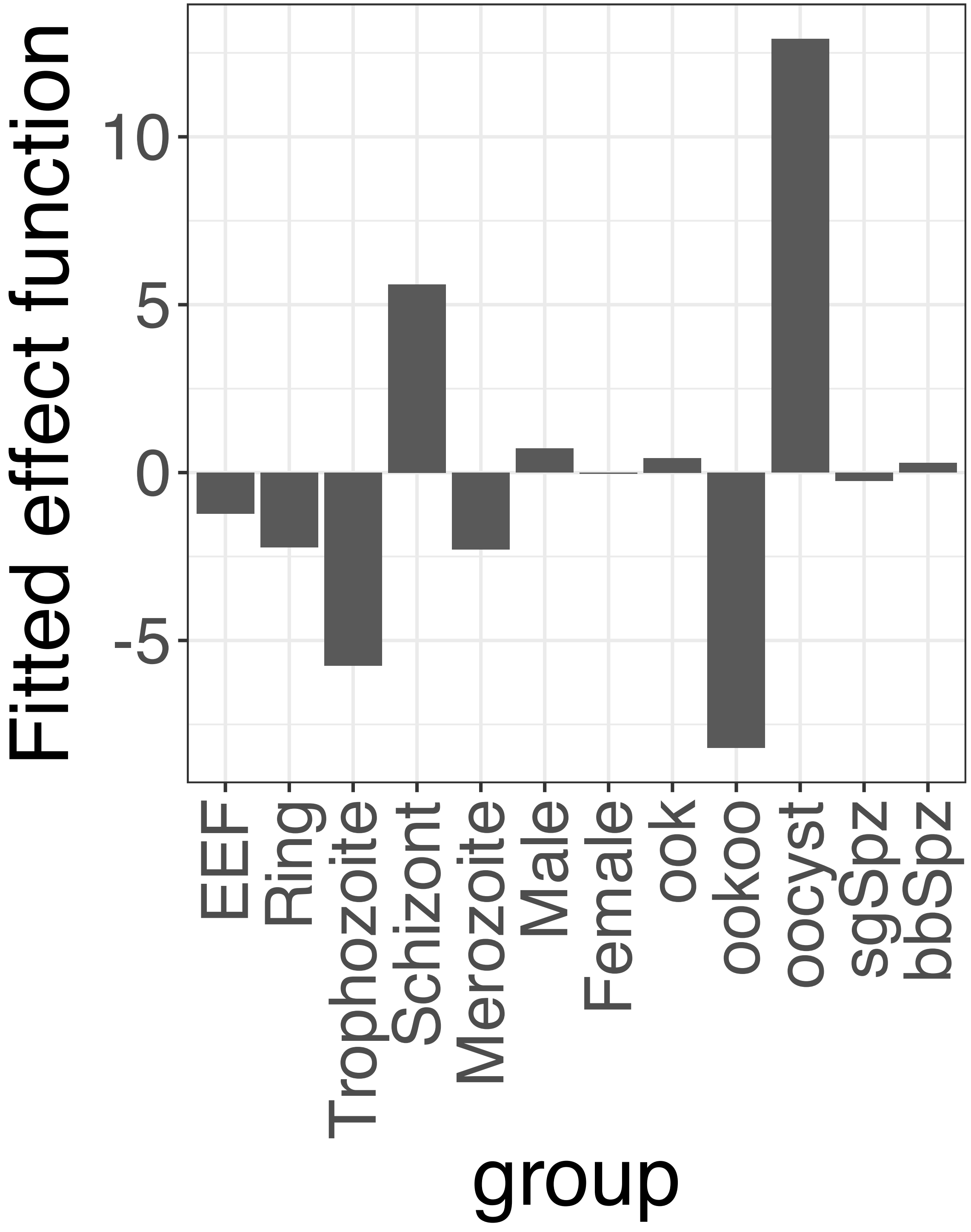


**Supplementary Figure 131.** Decomposition of reconstructed SSANOVA-SVD component function 9.

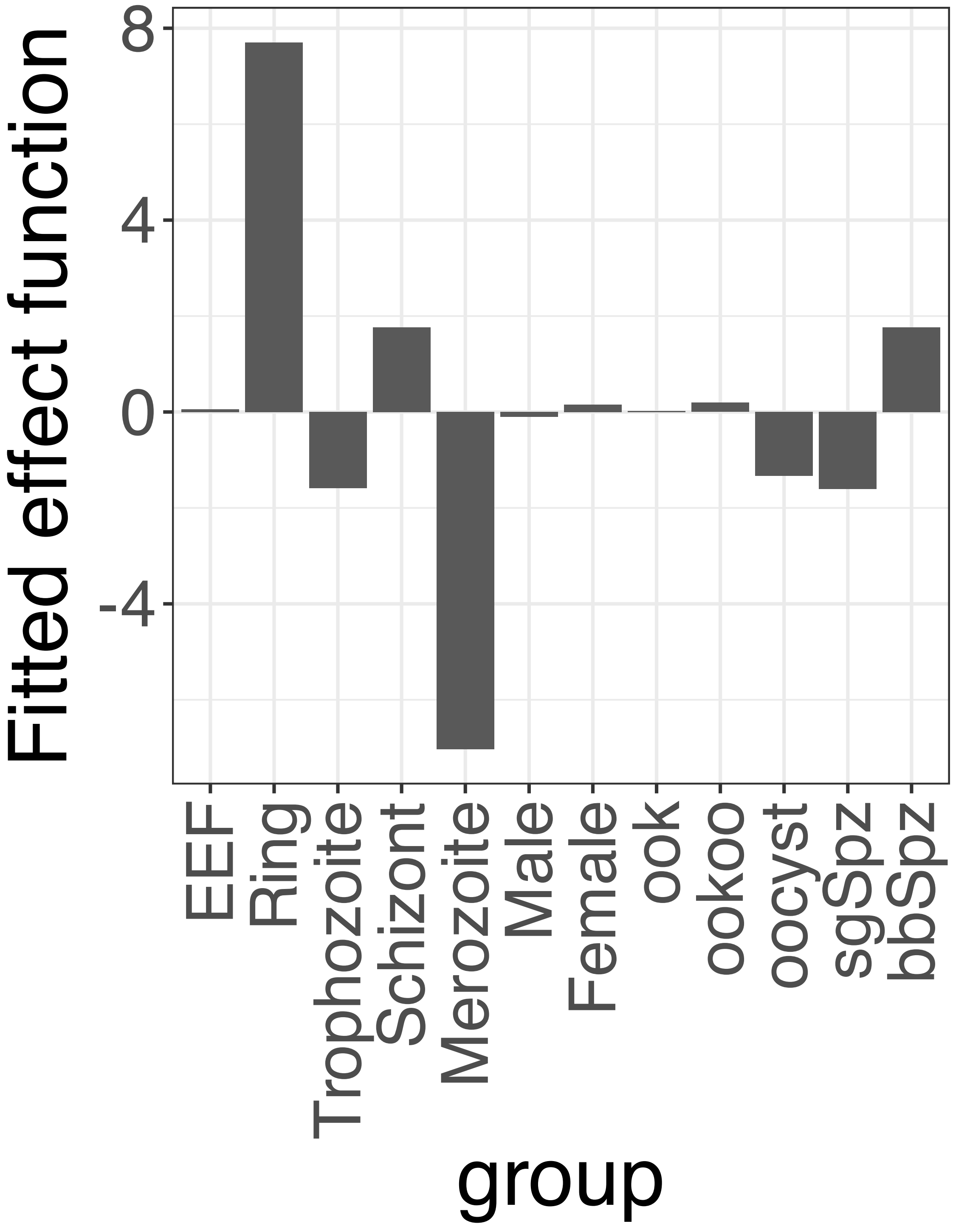


**Supplementary Figure 132.** Decomposition of reconstructed SSANOVA-SVD component function 10.

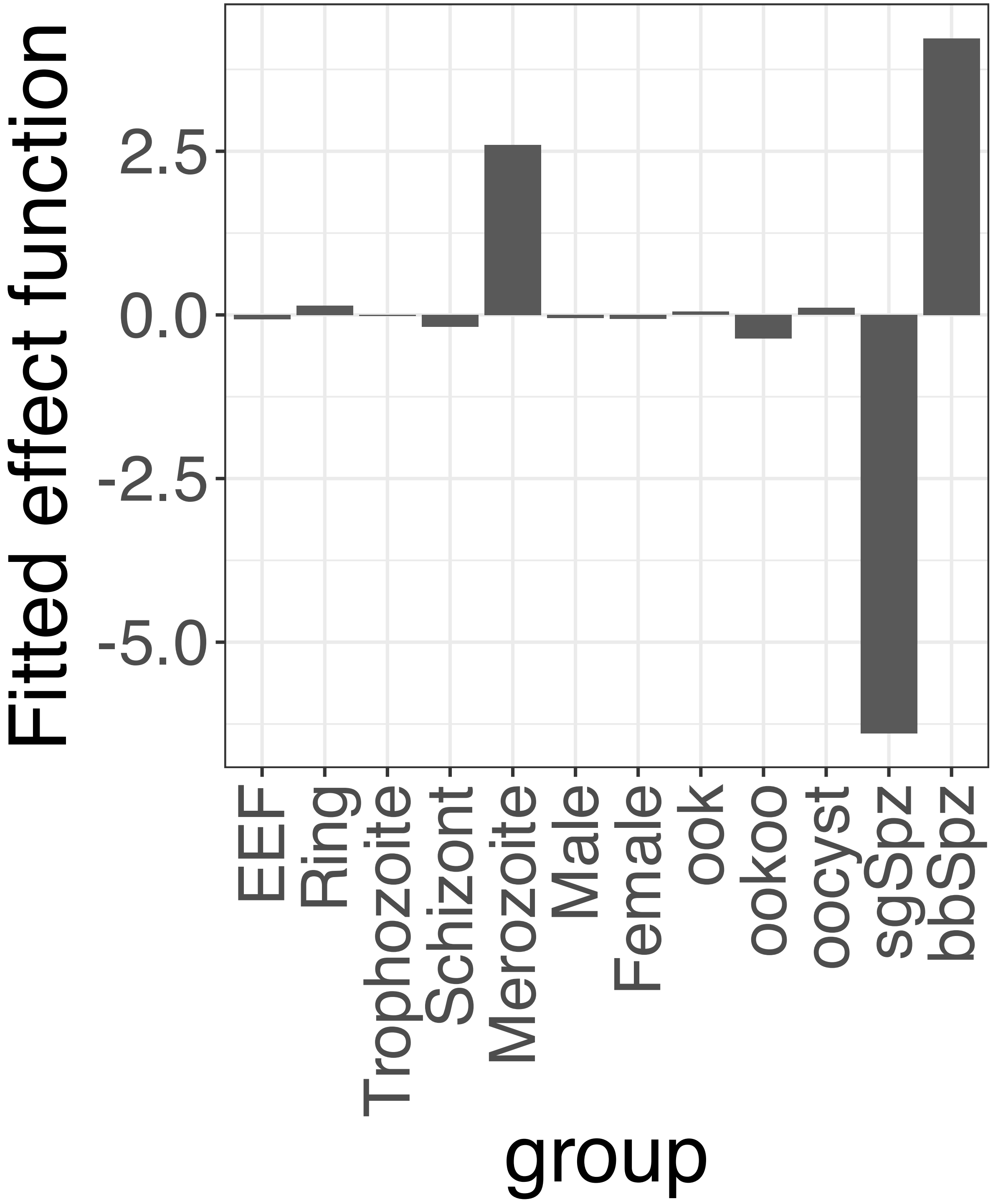


**Supplementary Figure 133.** Decomposition of reconstructed SSANOVA-SVD component function 11.

PCA

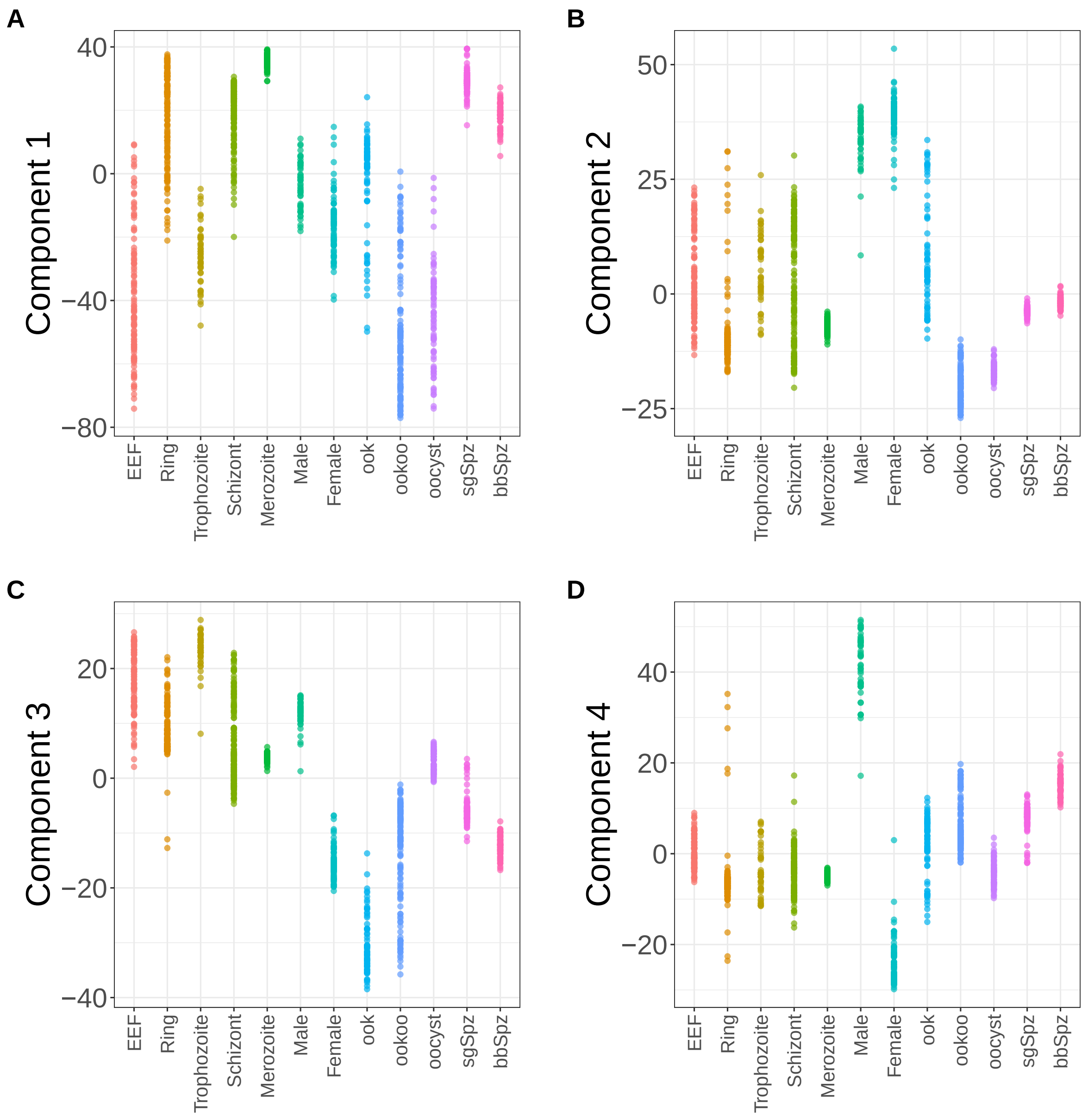


**Supplementary Figure 134.** Components for PCA. (A) Component 1. (B) Component 2. (C) Component 3. (D) Component 4.

PCA

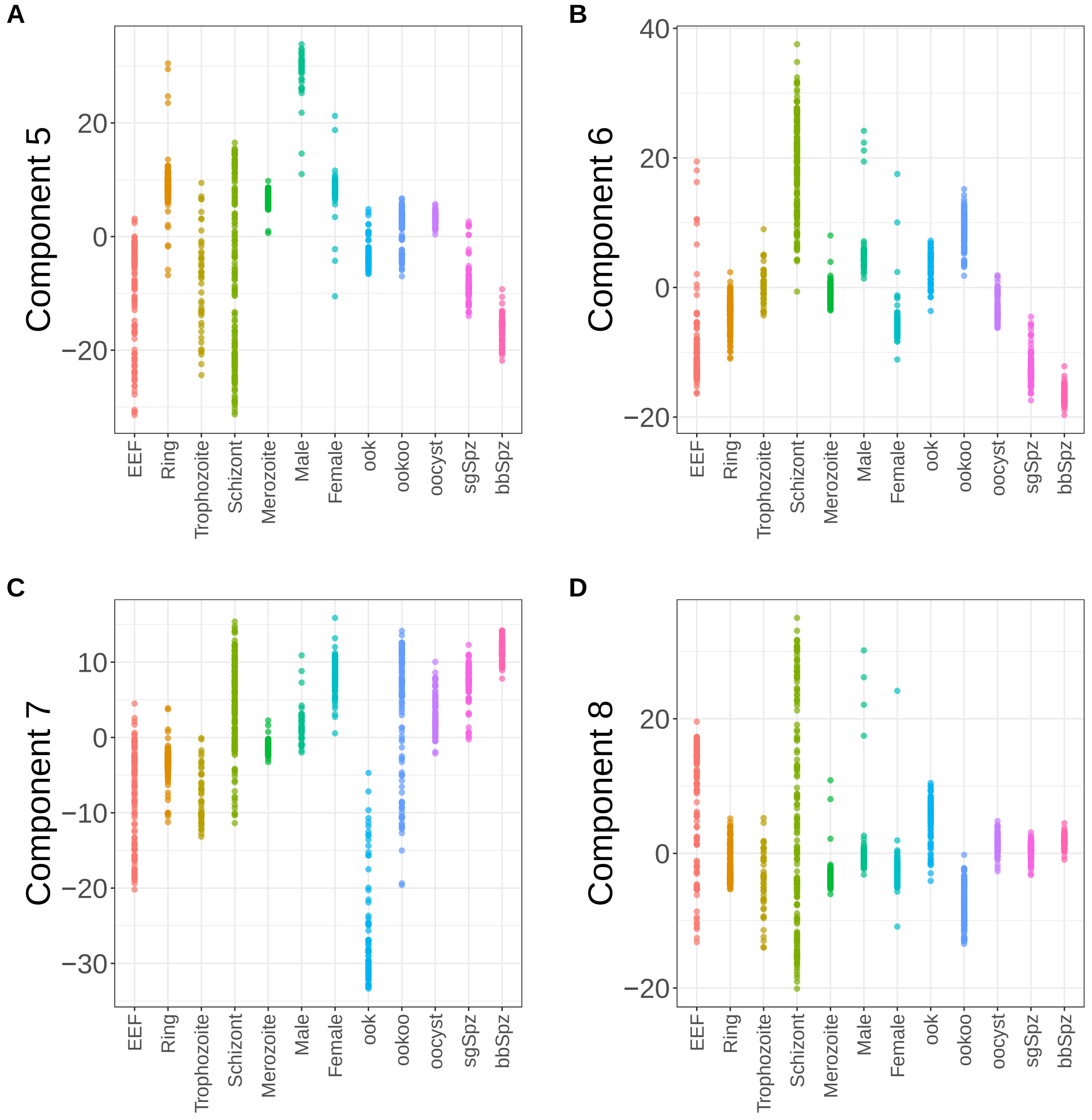


**Supplementary Figure 135.** Components for PCA. (A) Component 5. (B) Component 6. (C) Component 7. (D) Component 8.

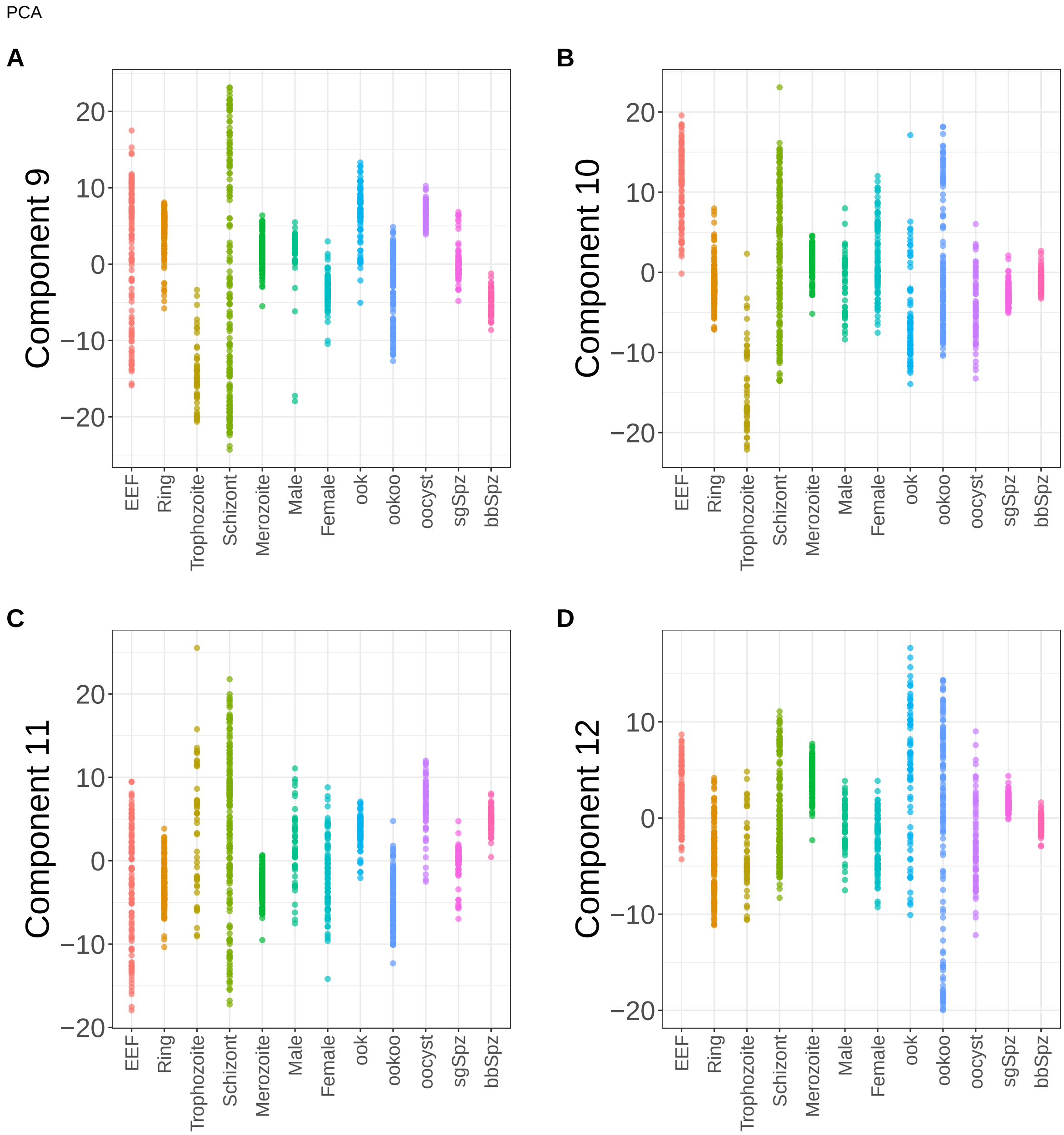


**Supplementary Figure 136.** Components for PCA. (A) Component 9. (B) Component 10. (C) Component 11. (D) Component 12.

PCA

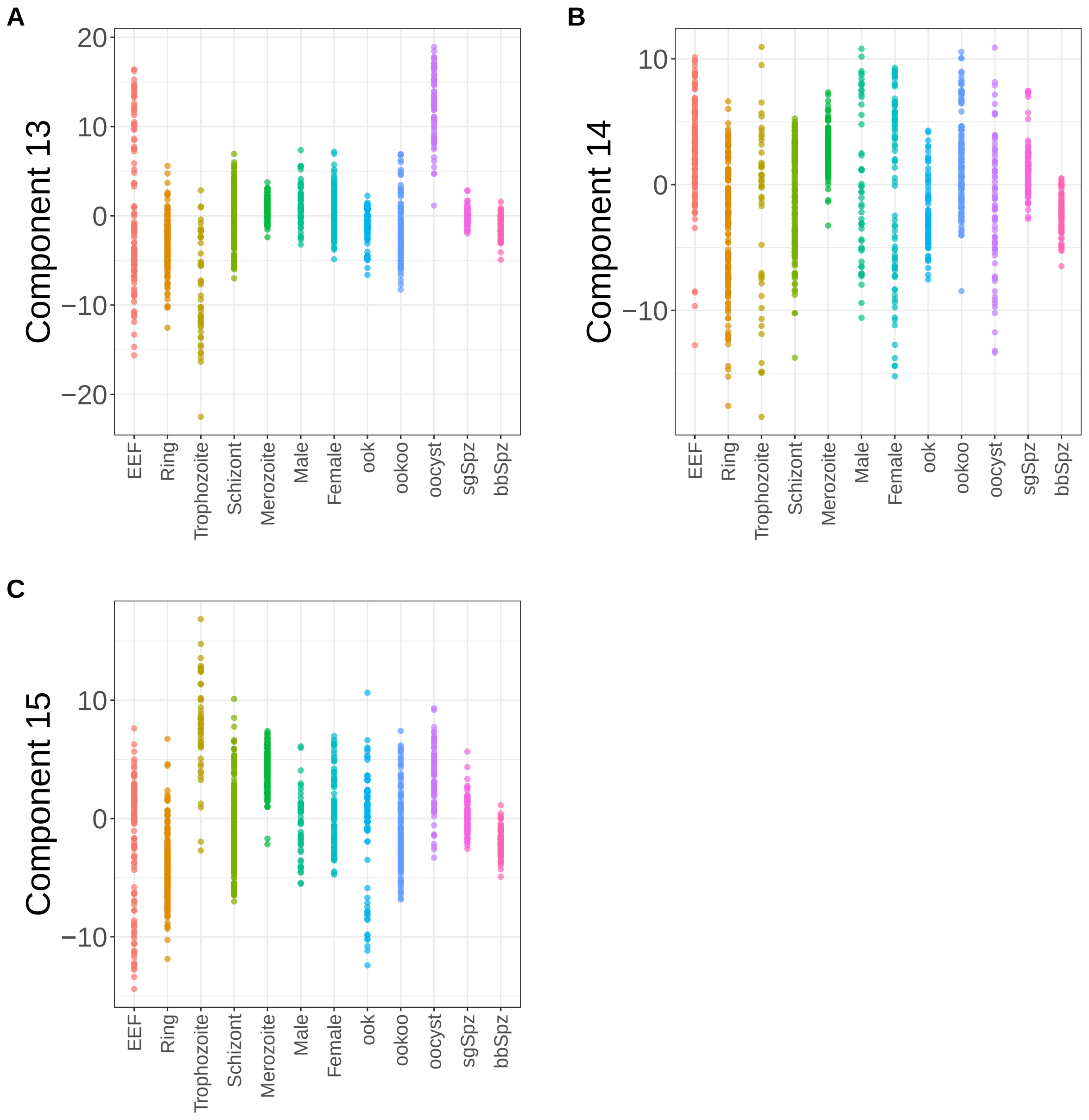


**Supplementary Figure 137.** Components for PCA. (A) Component 13. (B) Component 14. (C) Component 15.

NMF

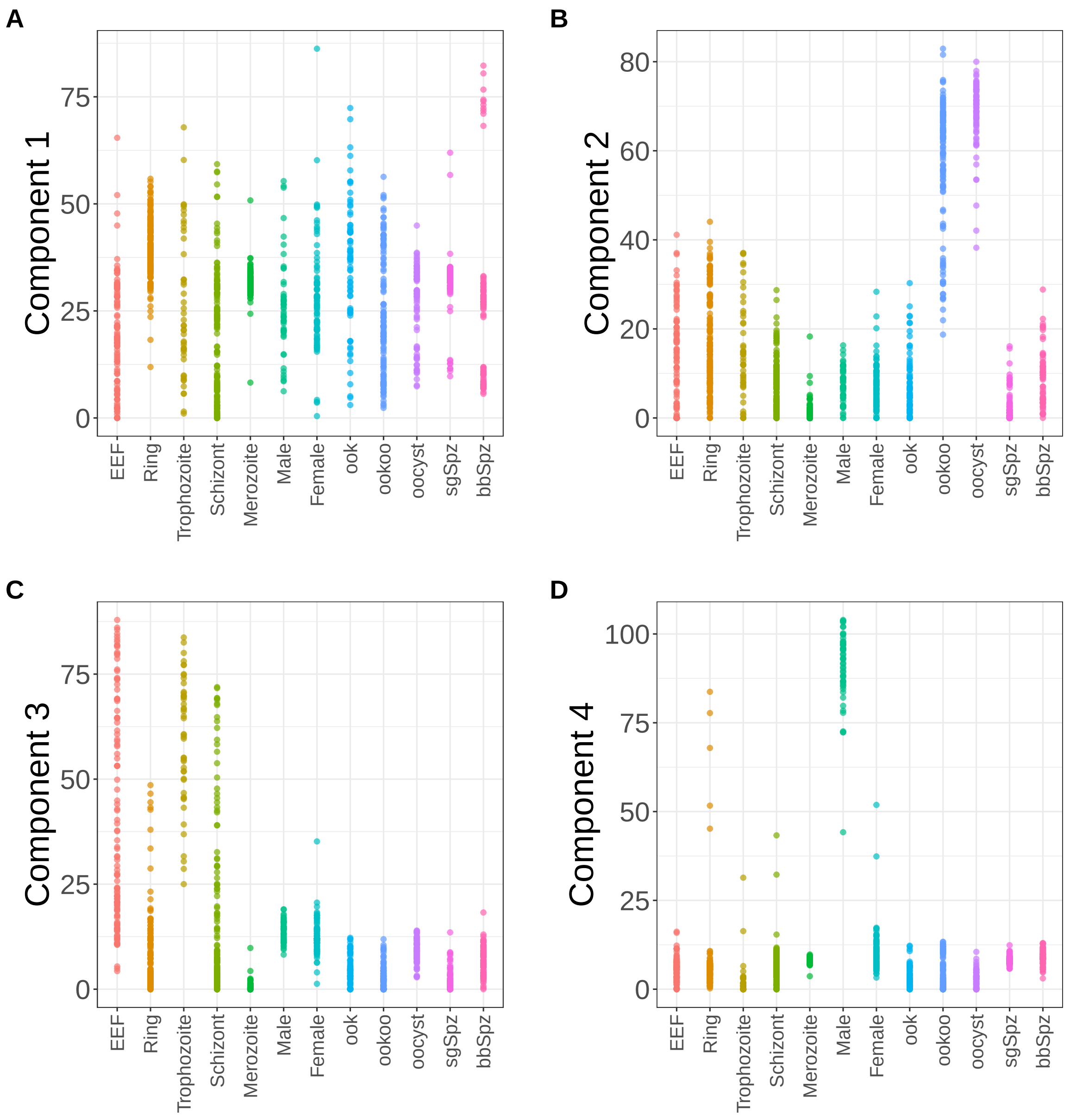


**Supplementary Figure 138.** Components for NMF. (A) Component 1. (B) Component 2. (C) Component 3. (D) Component 4.

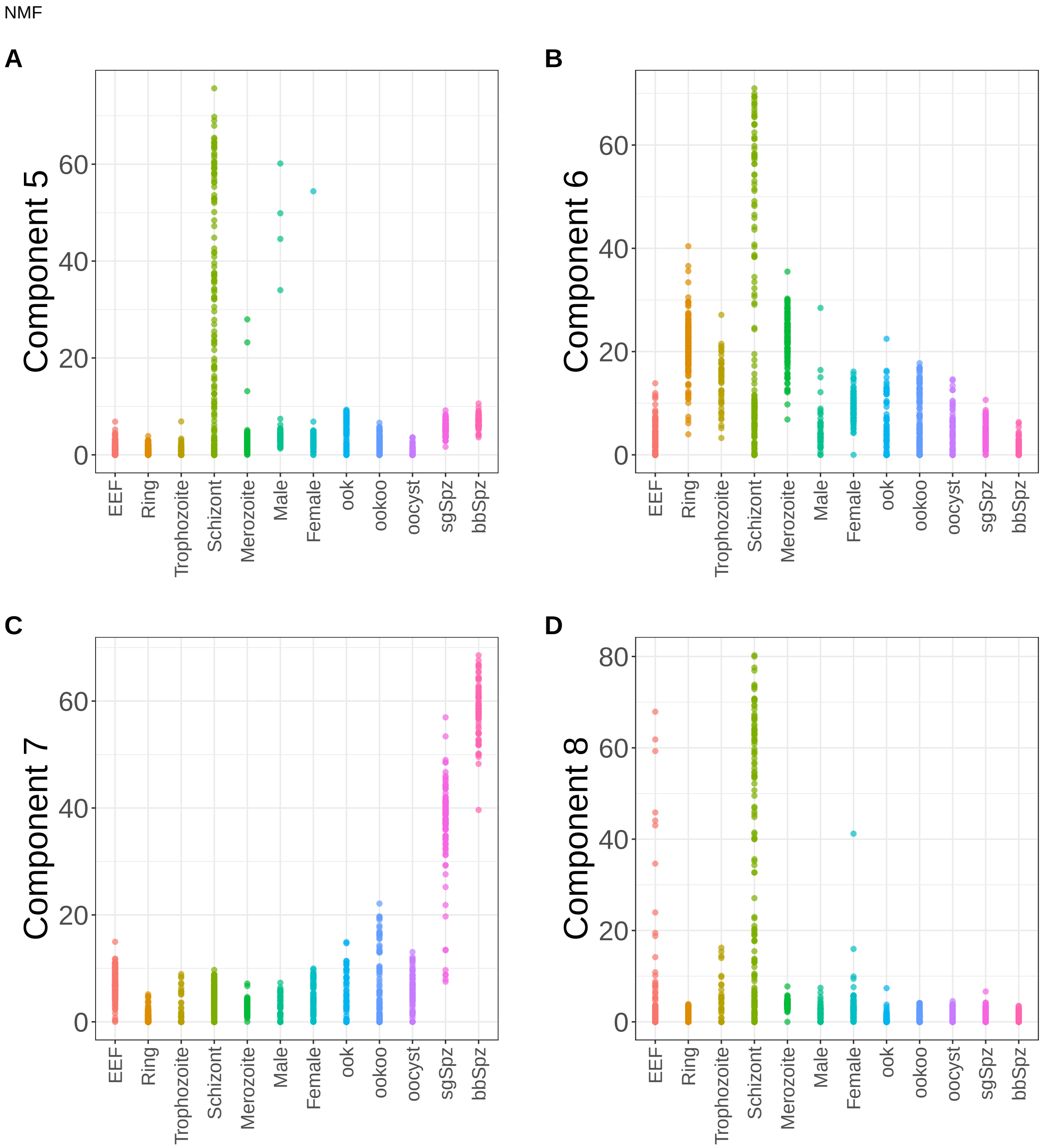


**Supplementary Figure 139.** Components for NMF. (A) Component 5. (B) Component 6. (C) Component 7. (D) Component 8.

NMF

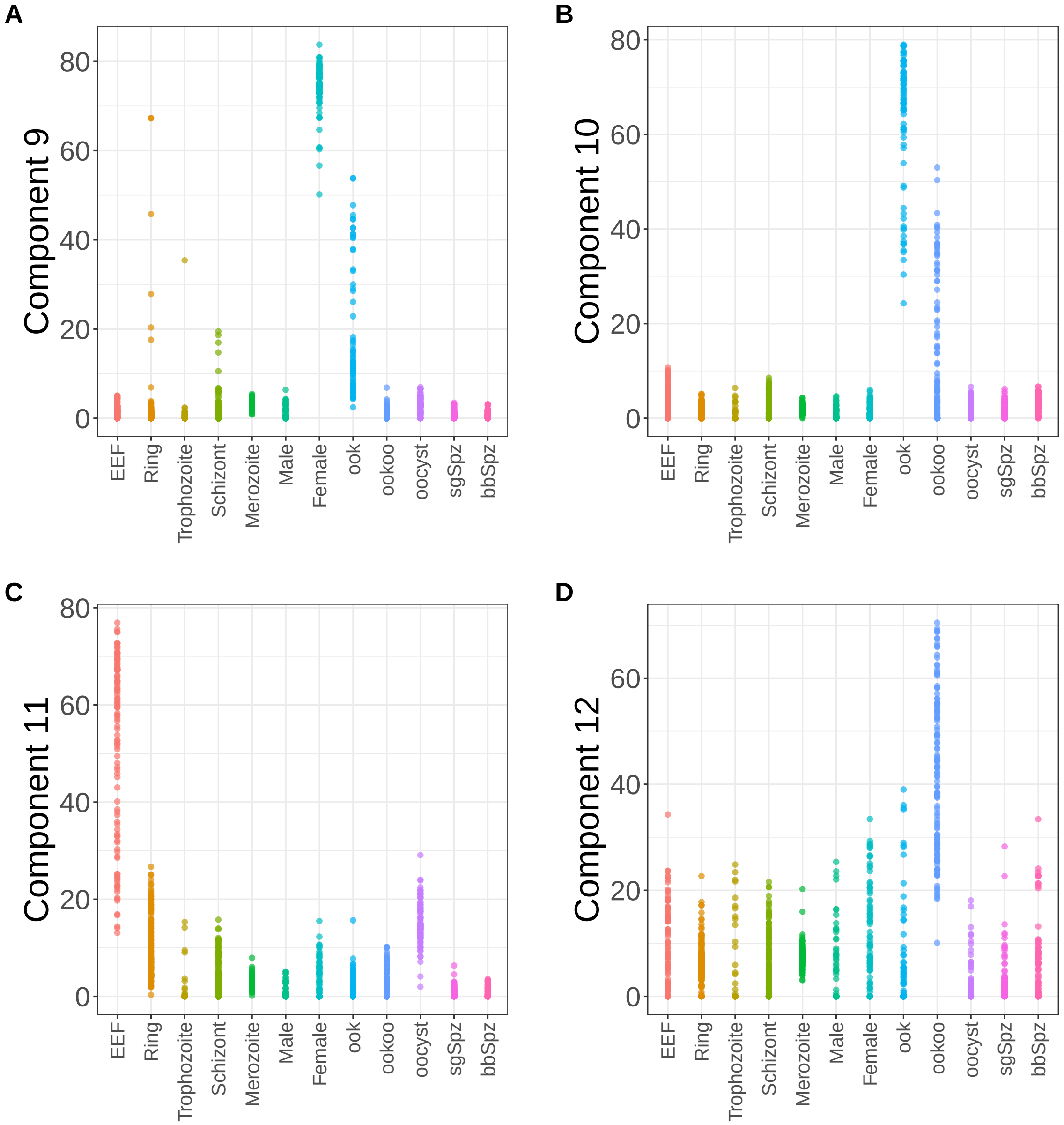


**Supplementary Figure 140.** Components for NMF. (A) Component 9. (B) Component 10. (C) Component 11. (D) Component 12.

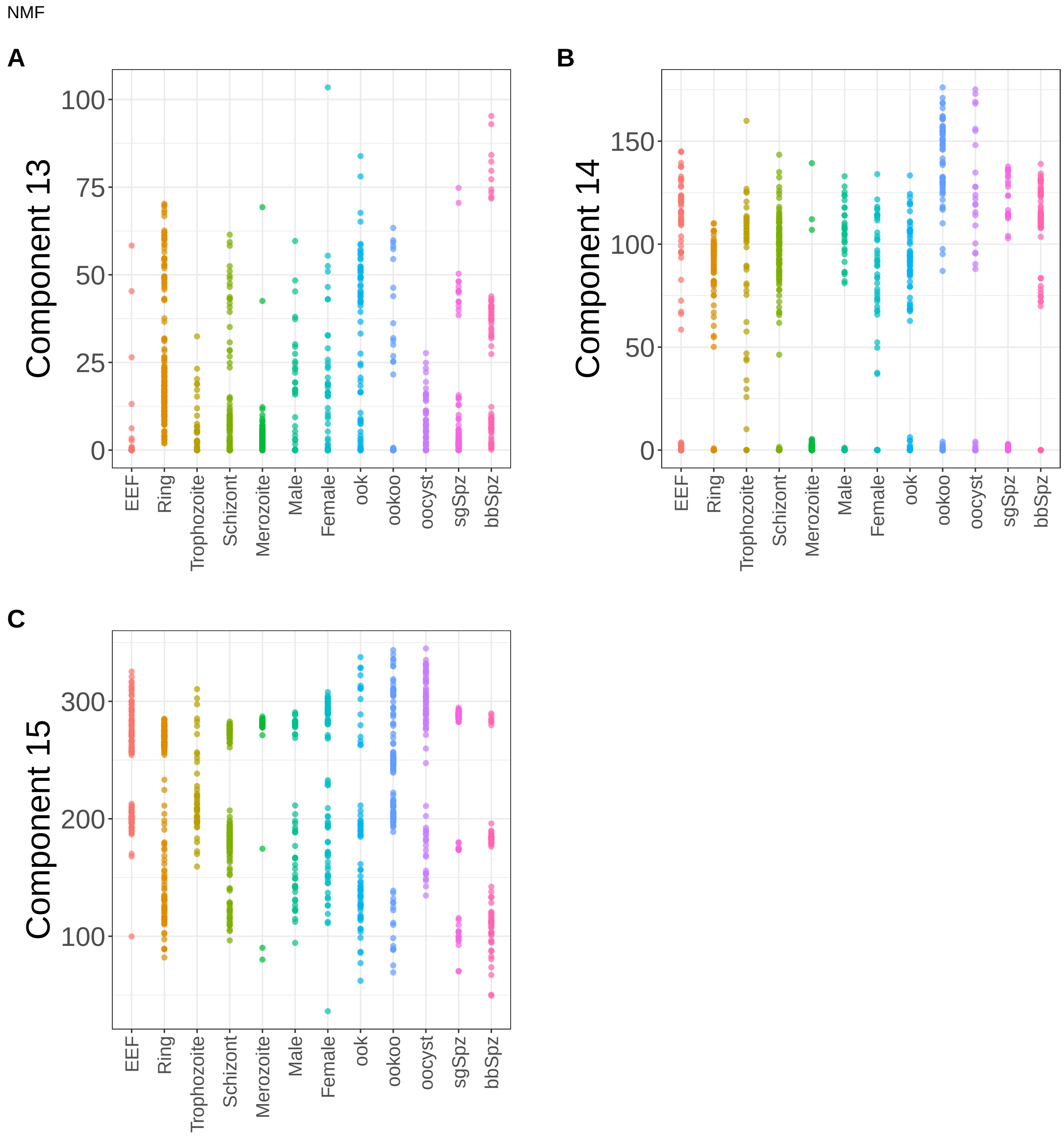


**Supplementary Figure 141.** Components for NMF. (A) Component 13. (B) Component 14. (C) Component 15.

PLS

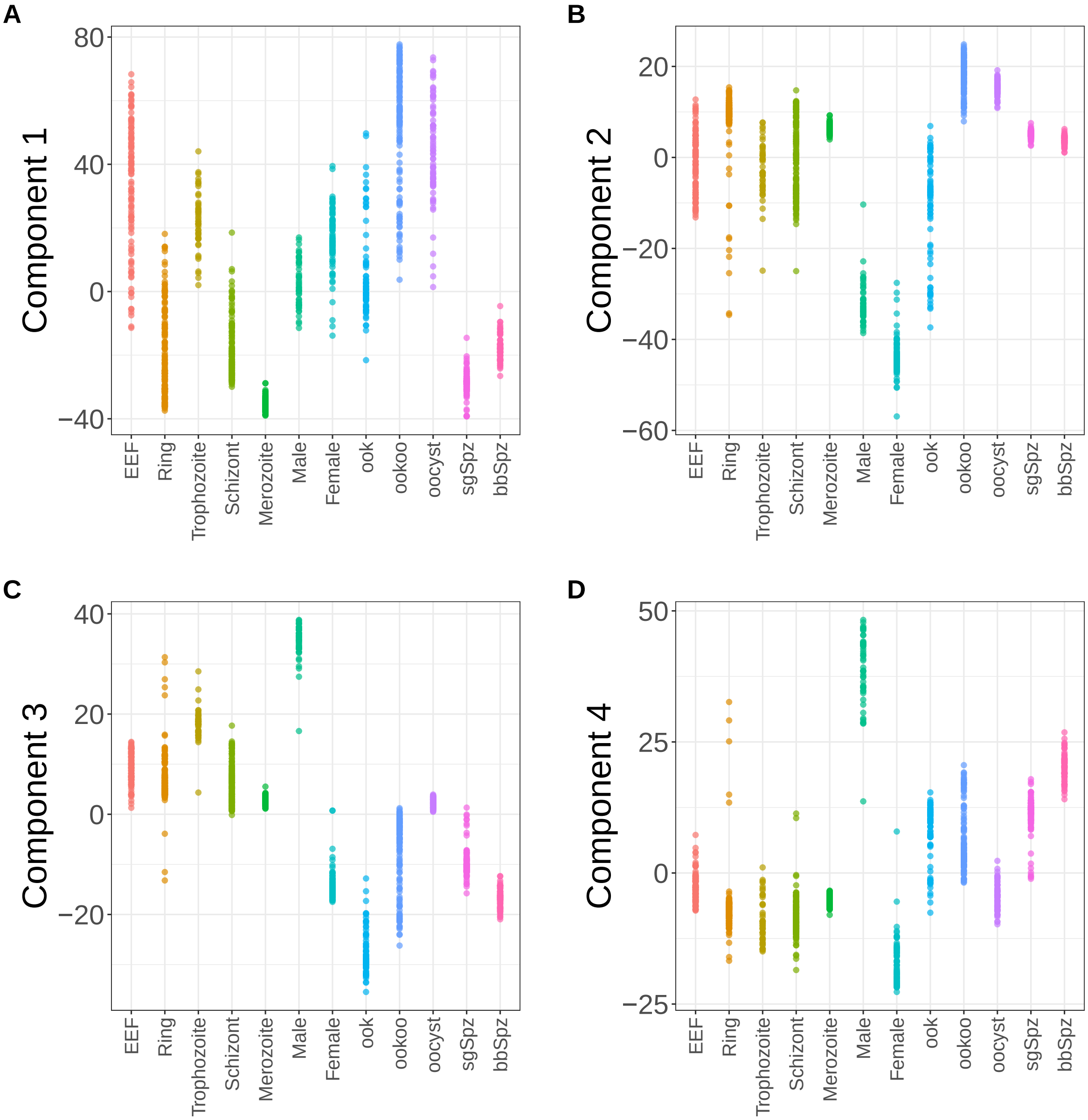


**Supplementary Figure 142.** Components for PLS. (A) Component 1. (B) Component 2. (C) Component 3. (D) Component 4.

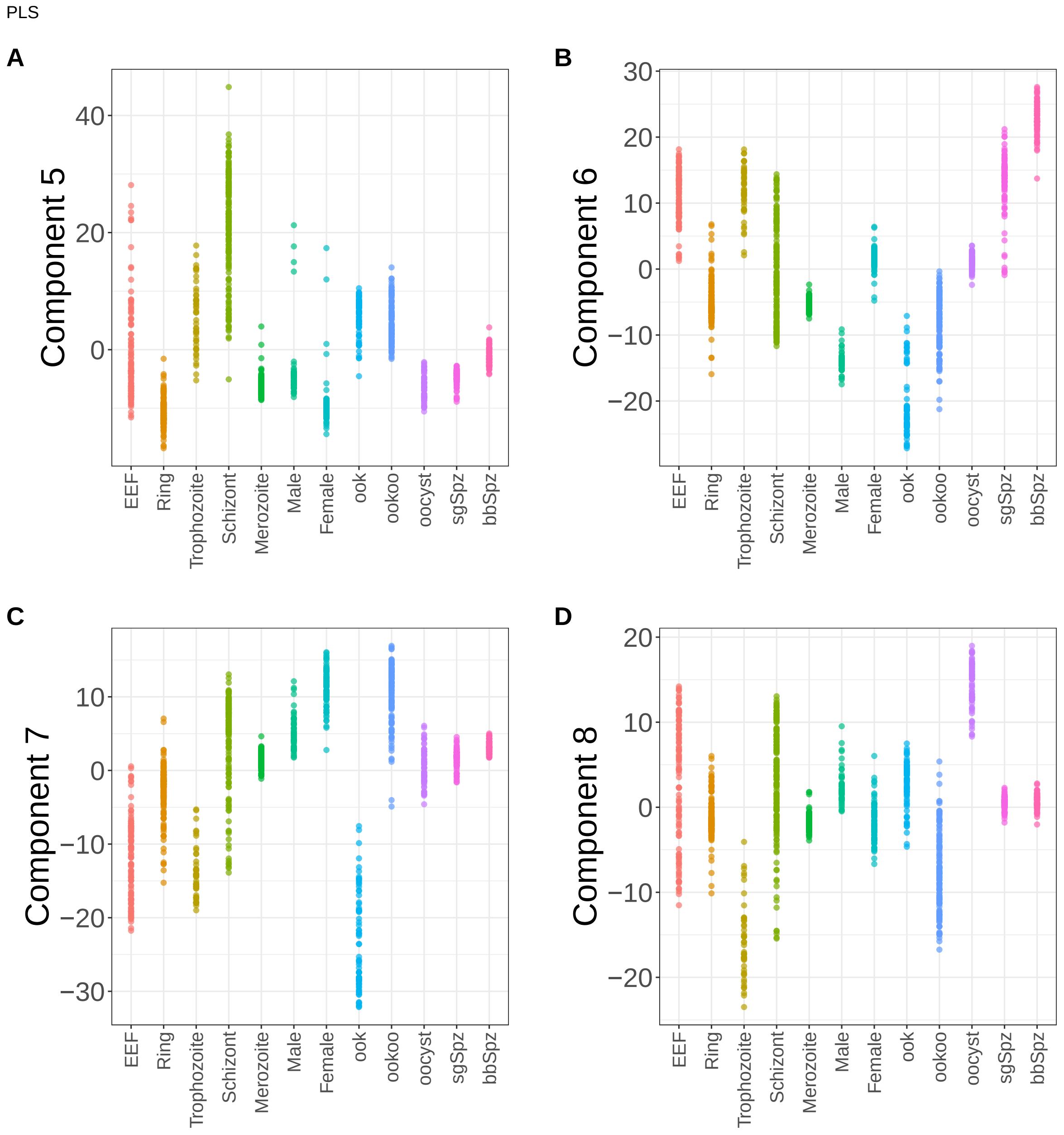


**Supplementary Figure 143.** Components for PLS. (A) Component 5. (B) Component 6. (C) Component 7. (D) Component 8.

PLS

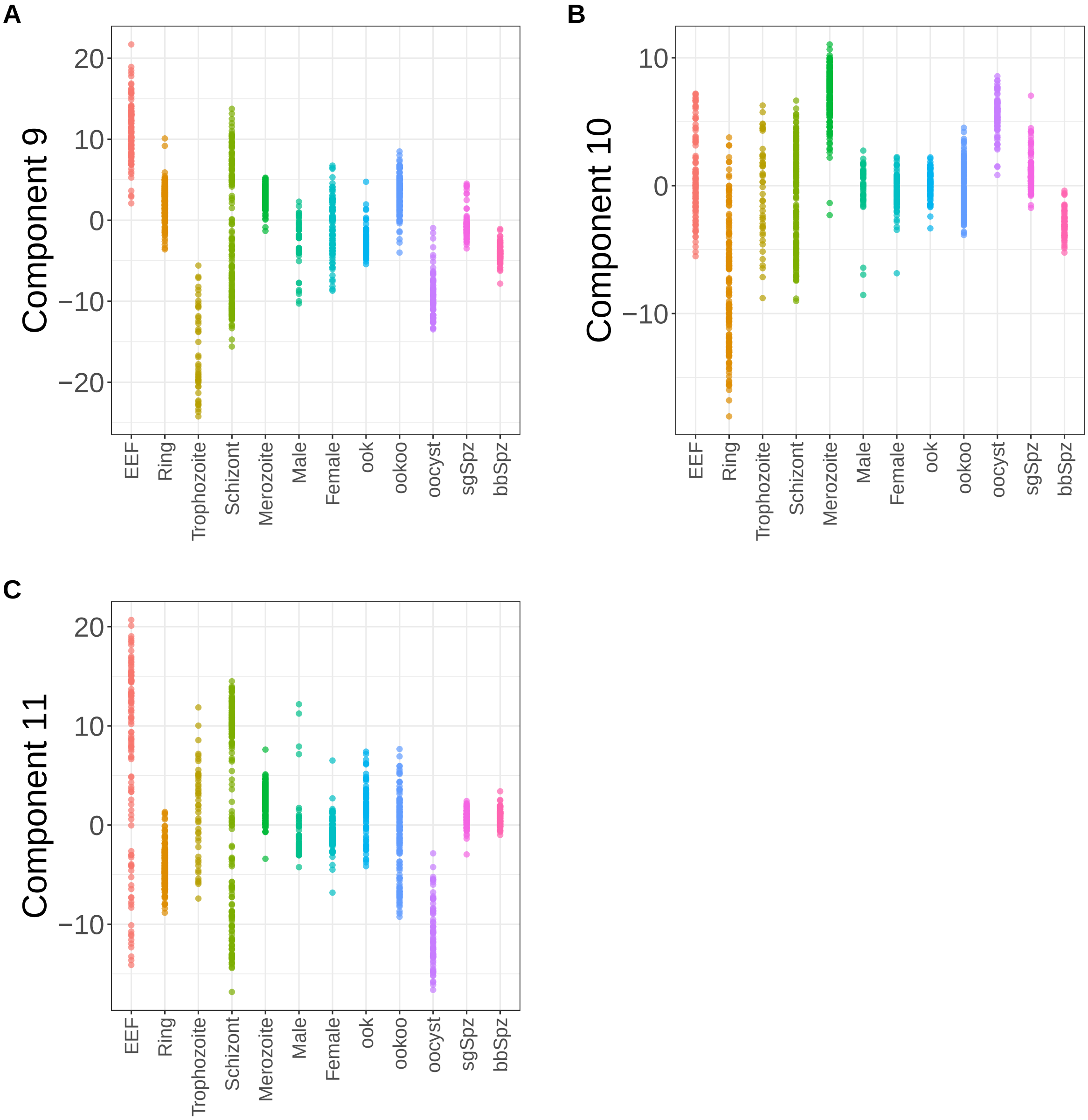


**Supplementary Figure 144.** Components for PLS. (A) Component 9. (B) Component 10. (C) Component 11.

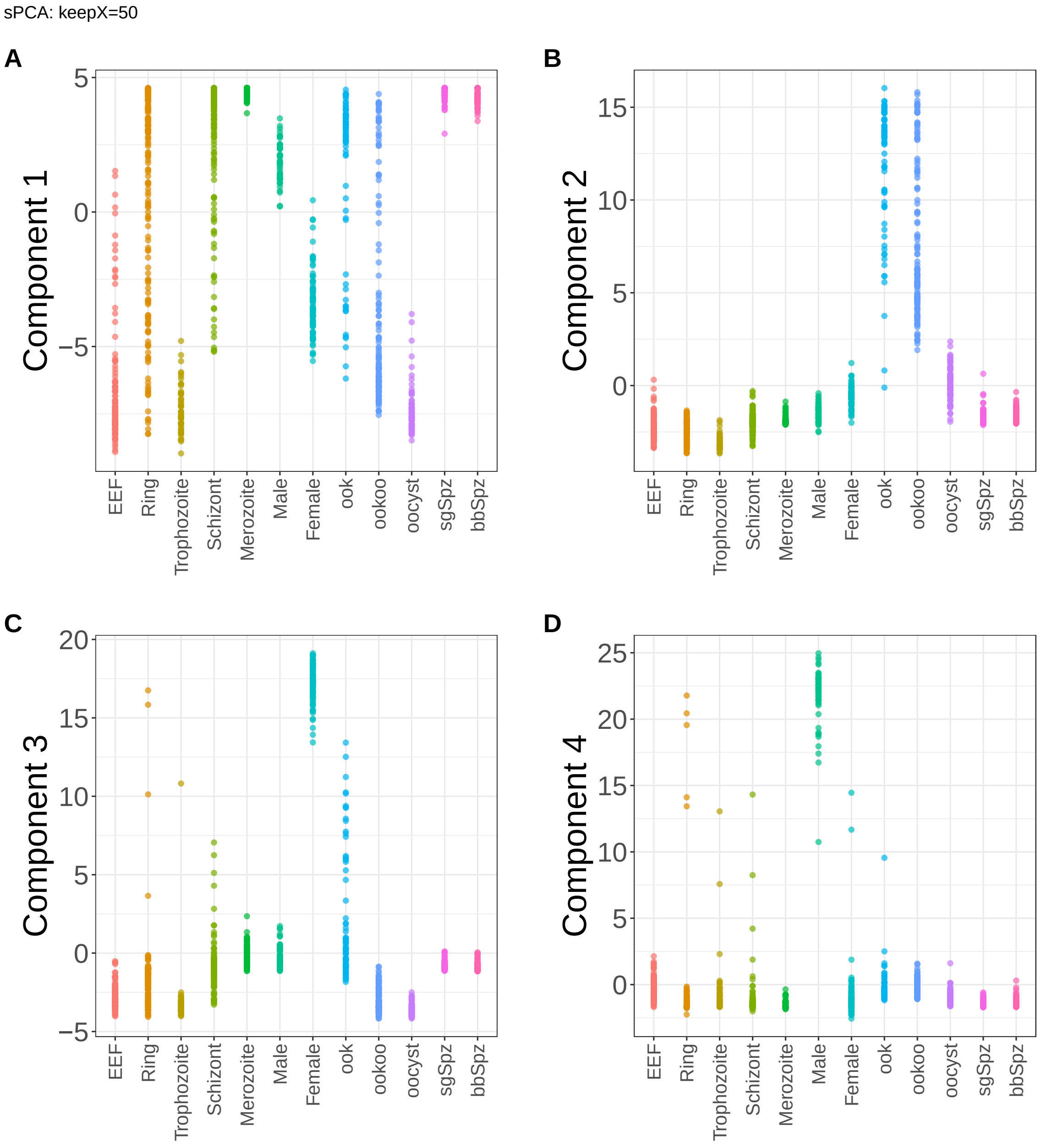


**Supplementary Figure 145.** Components for sPCA: keepX=50. (A) Component 1. (B) Component 2. (C) Component 3. (D) Component 4.

sPCA: keepX=50

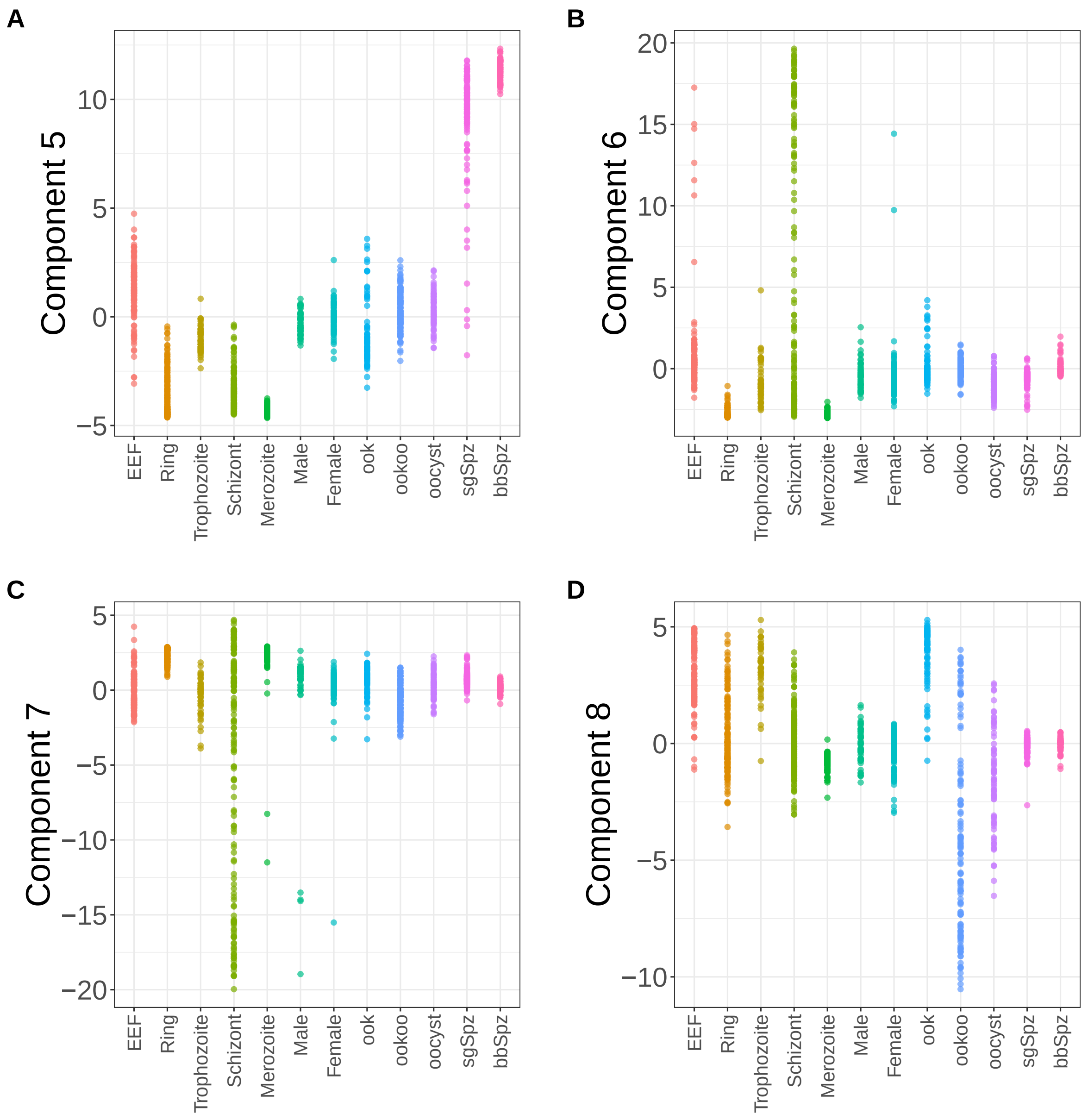


**Supplementary Figure 146.** Components for sPCA: keepX=50. (A) Component 5. (B) Component 6. (C) Component 7. (D) Component 8.

sPCA: keepX=50

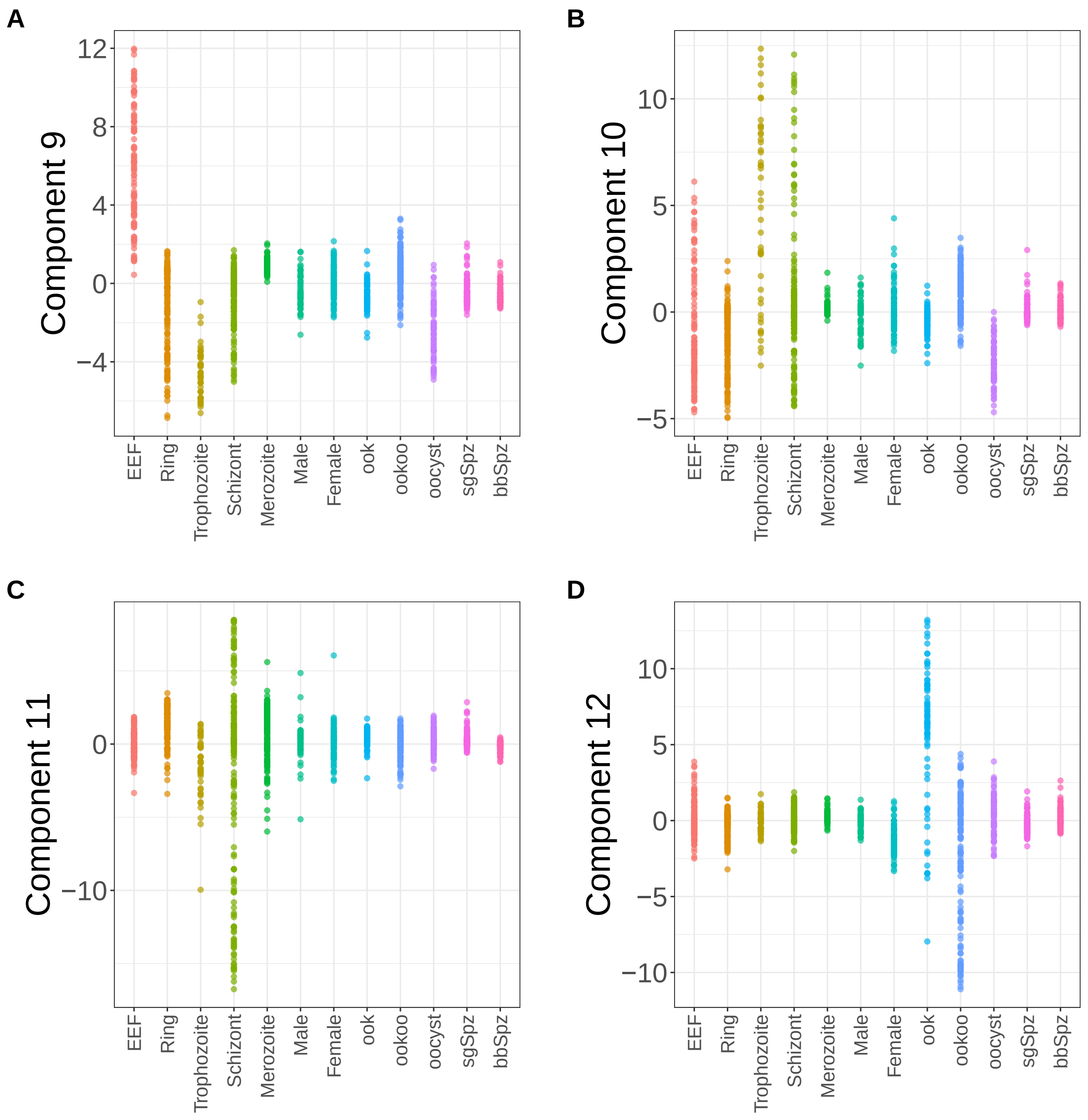


**Supplementary Figure 147.** Components for sPCA: keepX=50. (A) Component 9. (B) Component 10. (C) Component 11. (D) Component 12.

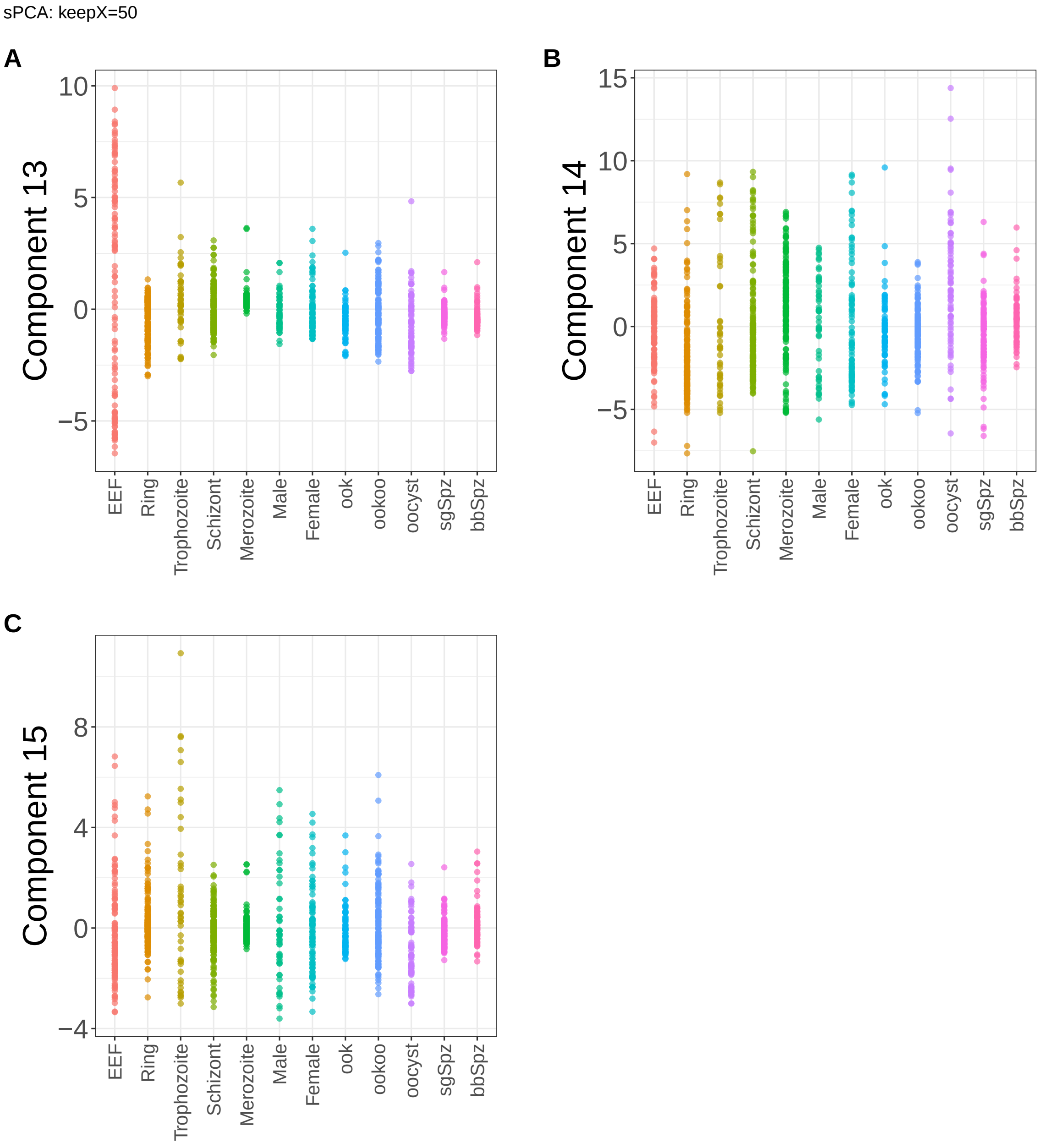


**Supplementary Figure 148.** Components for sPCA: keepX=50. (A) Component 13. (B) Component 14. (C) Component 15.

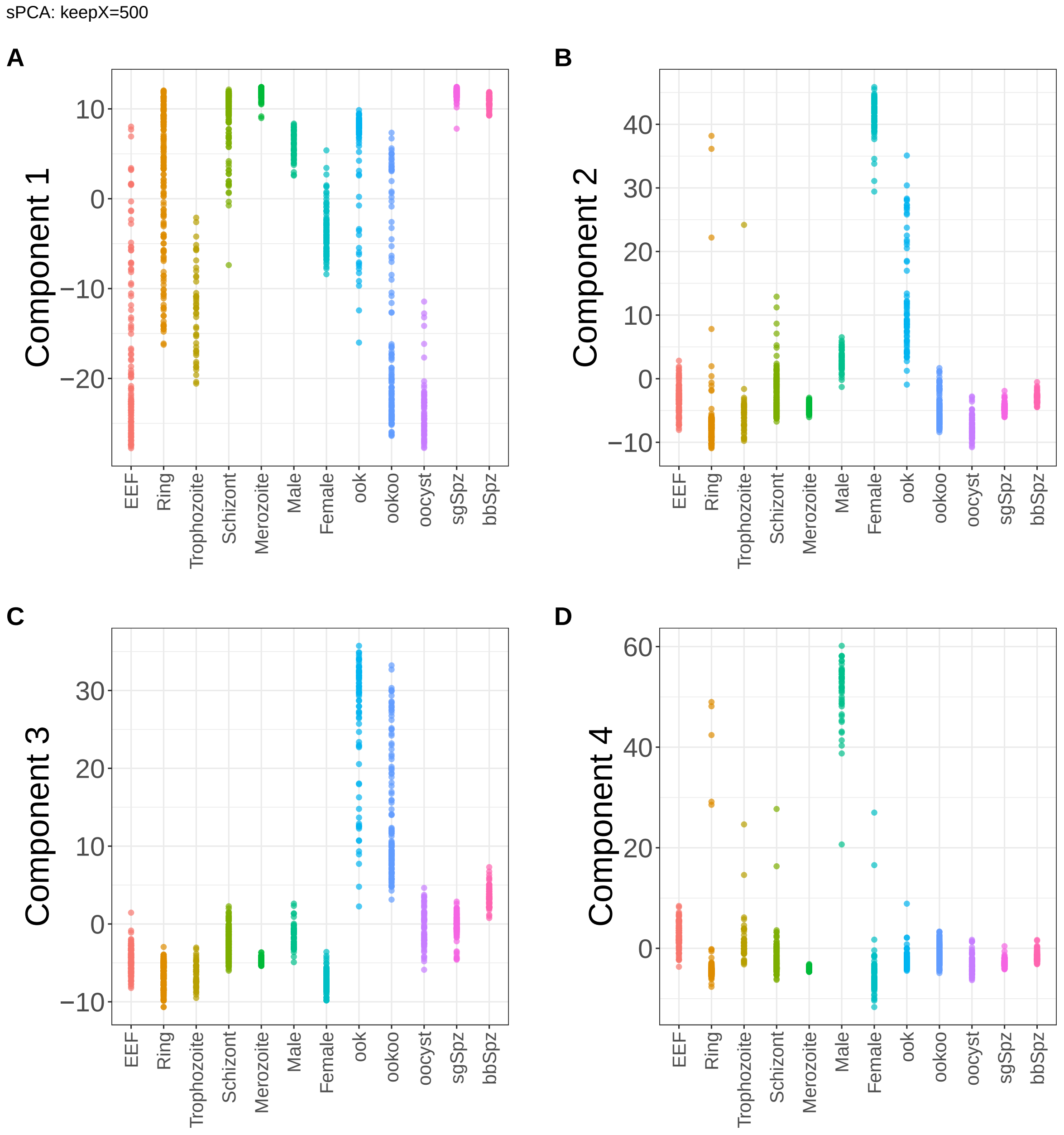


**Supplementary Figure 149.** Components for sPCA: keepX=500. (A) Component 1. (B) Component 2. (C) Component 3. (D) Component 4.

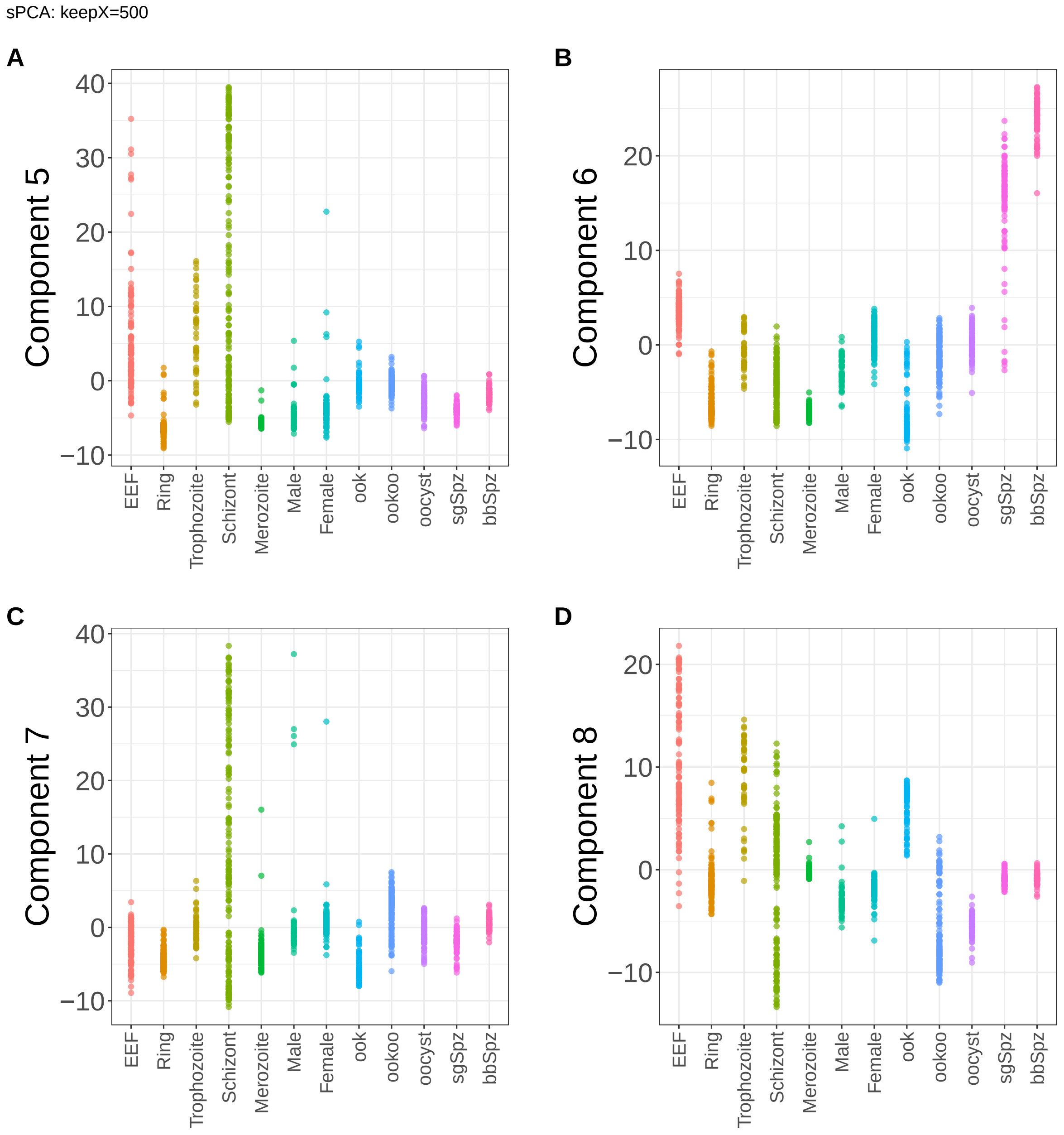


**Supplementary Figure 150.** Components for sPCA: keepX=500. (A) Component 5. (B) Component 6. (C) Component 7. (D) Component 8.

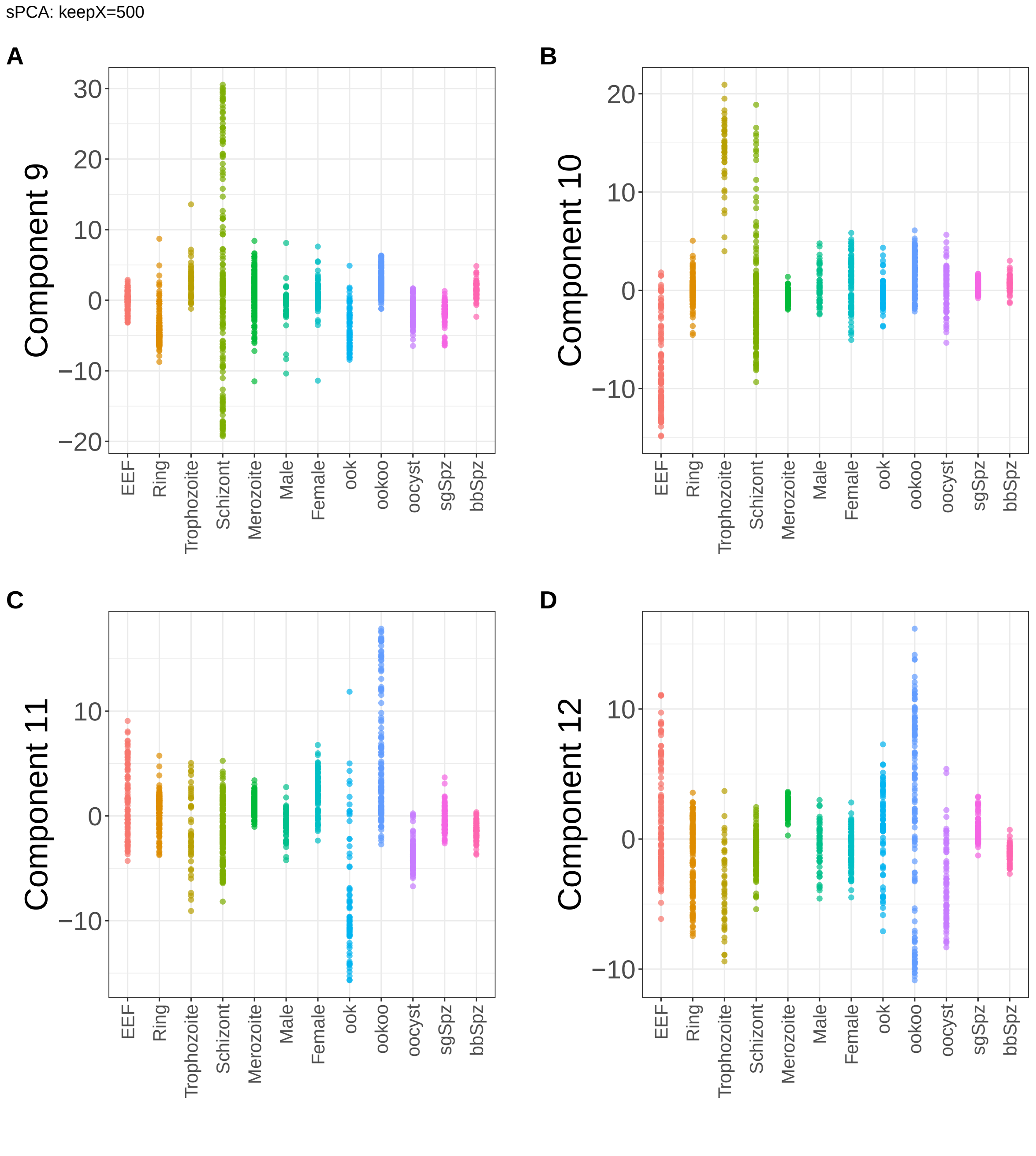


**Supplementary Figure 151.** Components for sPCA: keepX=500. (A) Component 9. (B) Component 10. (C) Component 11. (D) Component 12.

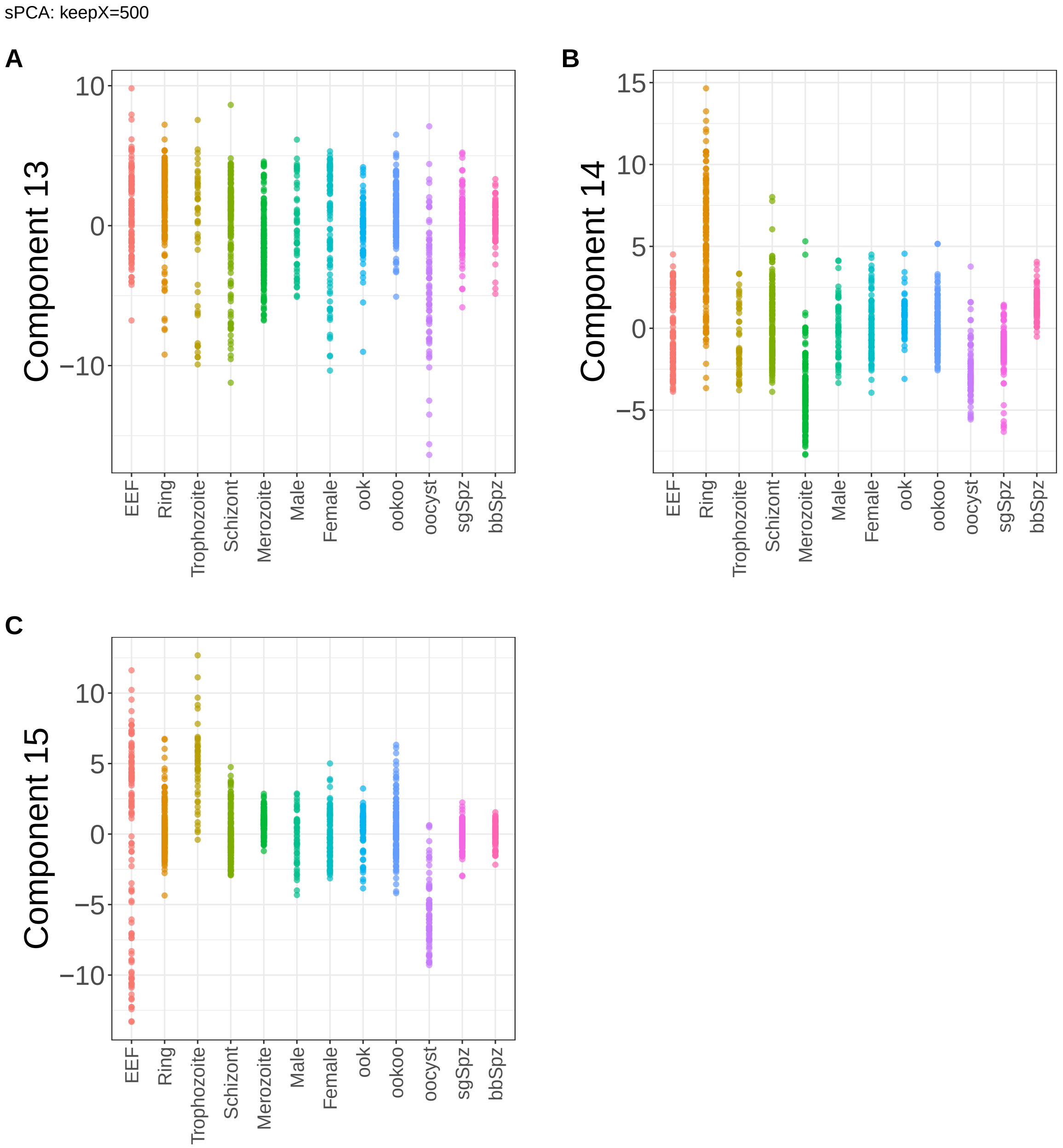


**Supplementary Figure 152.** Components for sPCA: keepX=500. (A) Component 13. (B) Component 14. (C) Component 15.

sPCA: keepX=1250

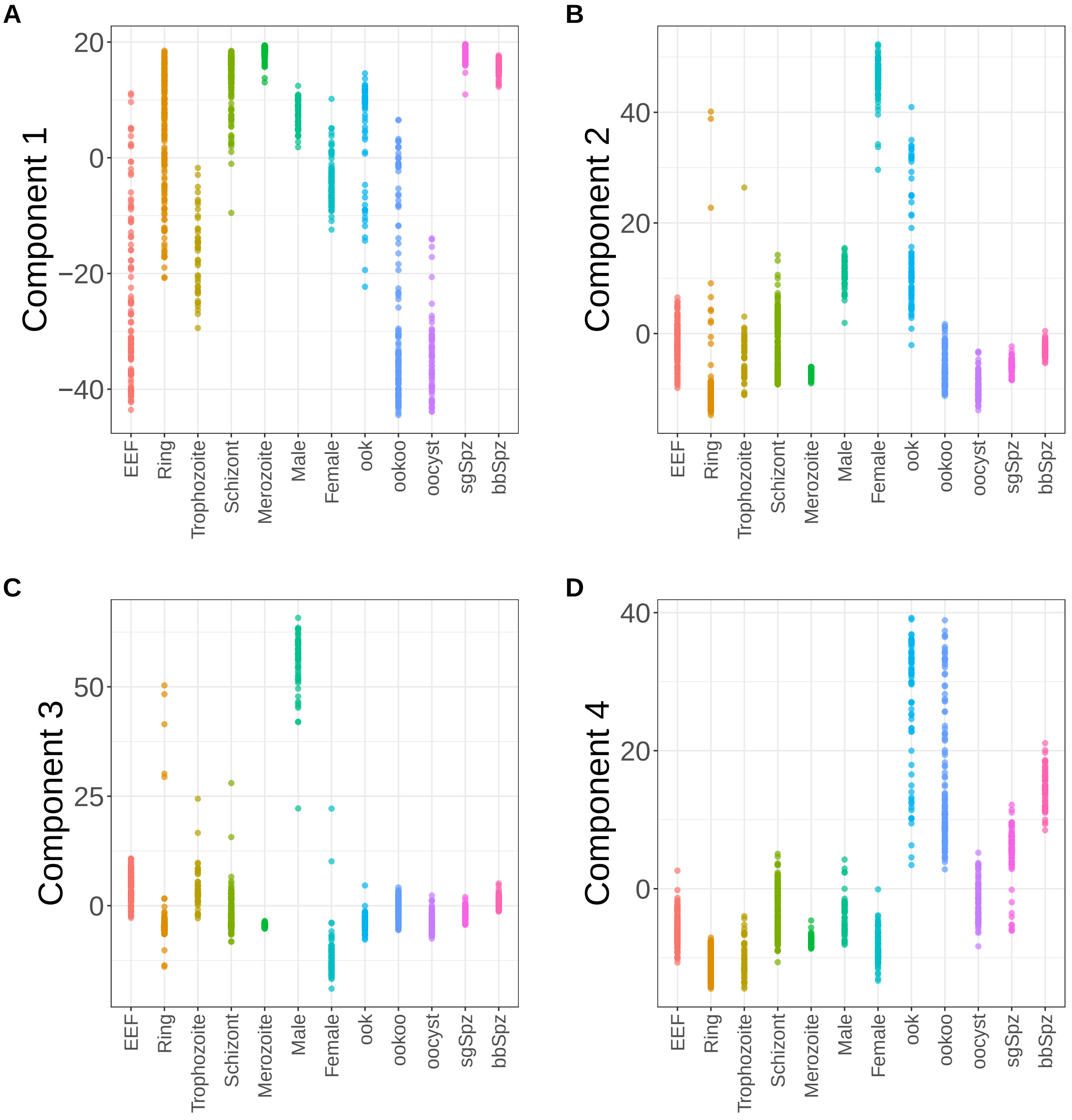


**Supplementary Figure 153.** Components for sPCA: keepX=1250. (A) Component 1. (B) Component 2. (C) Component 3. (D) Component 4.

sPCA: keepX=1250

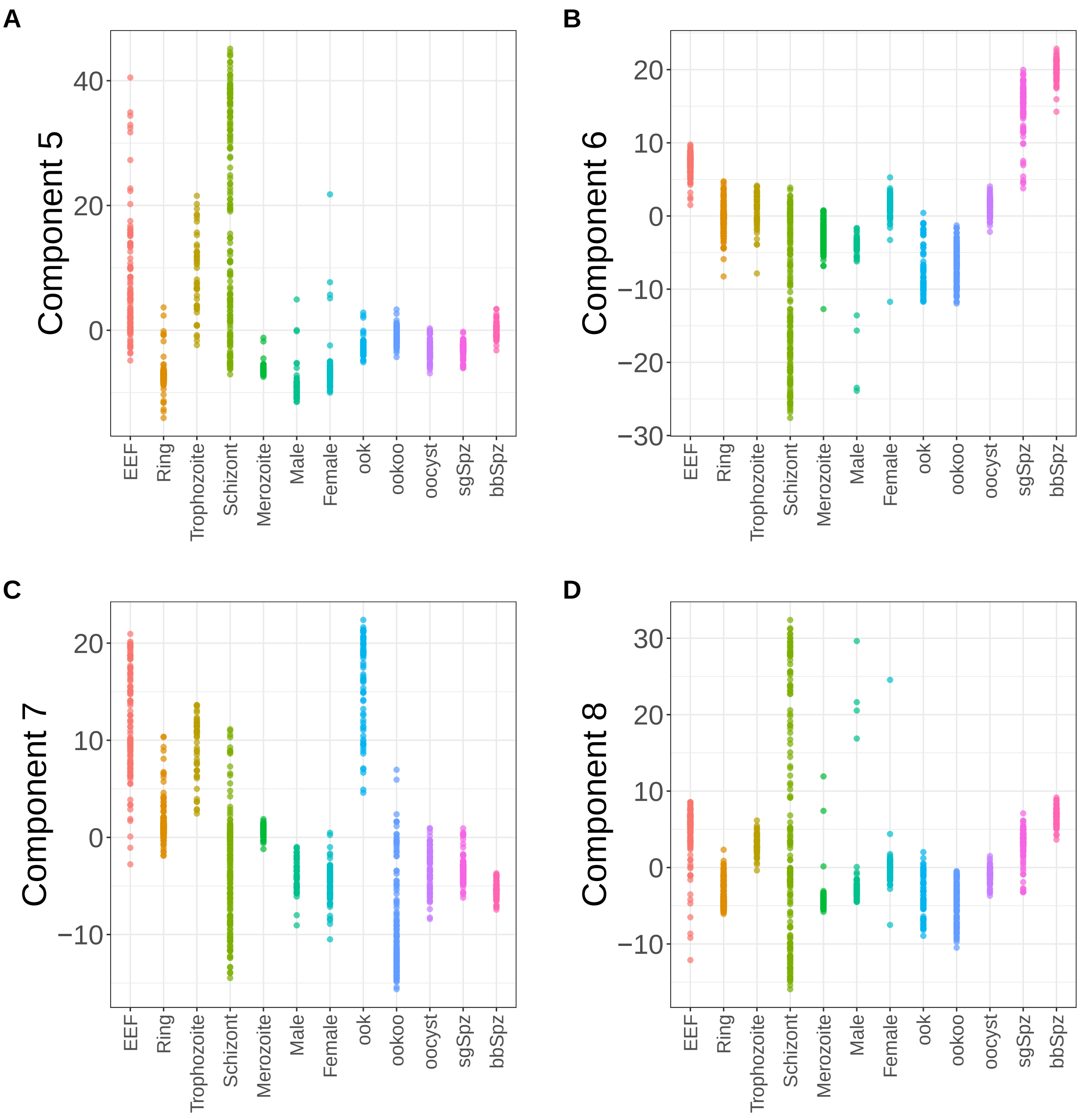


**Supplementary Figure 154.** Components for sPCA: keepX=1250. (A) Component 5. (B) Component 6. (C) Component 7. (D) Component 8.

sPCA: keepX=1250

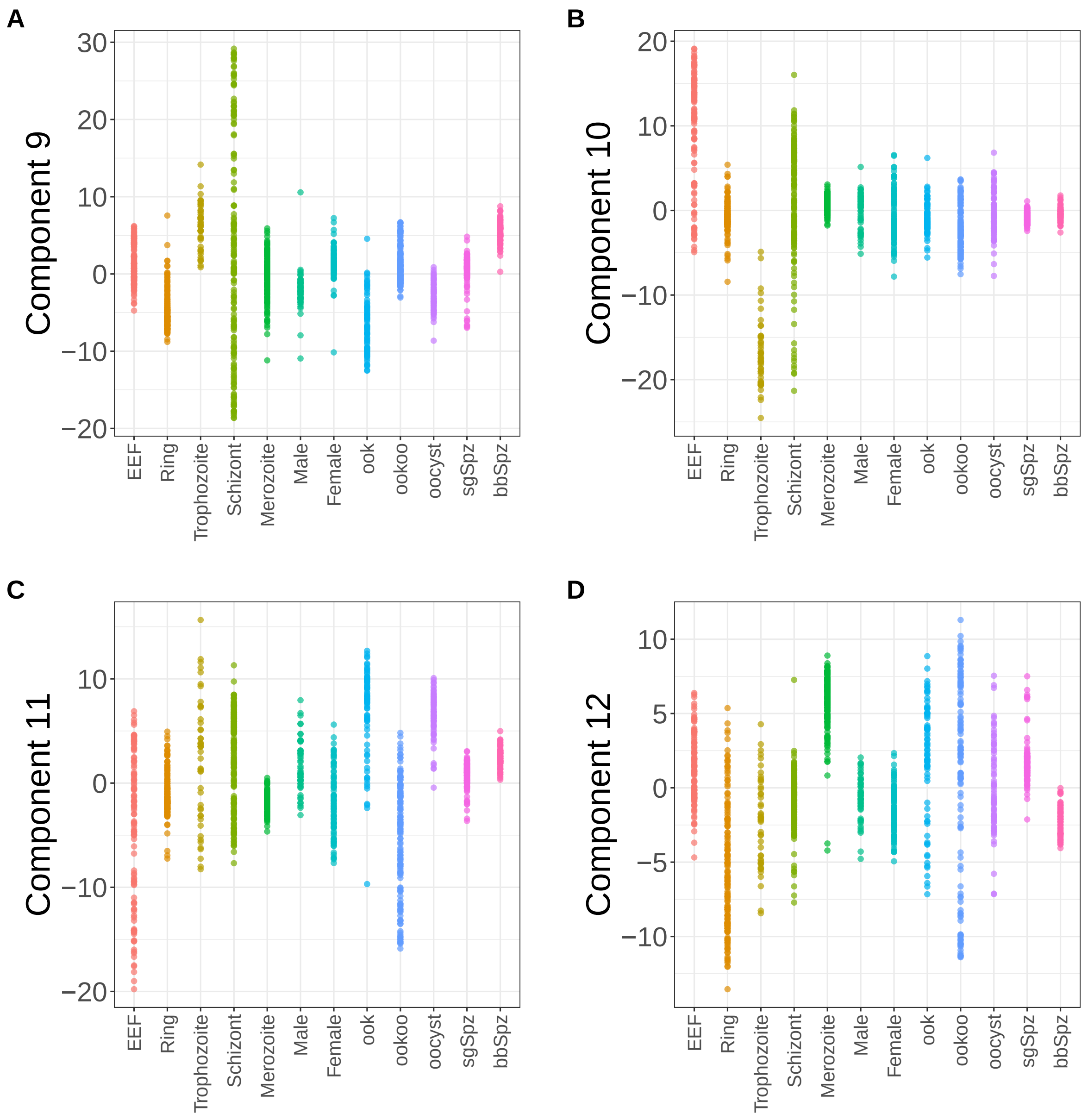


**Supplementary Figure 155.** Components for sPCA: keepX=1250. (A) Component 9. (B) Component 10. (C) Component 11. (D) Component 12.

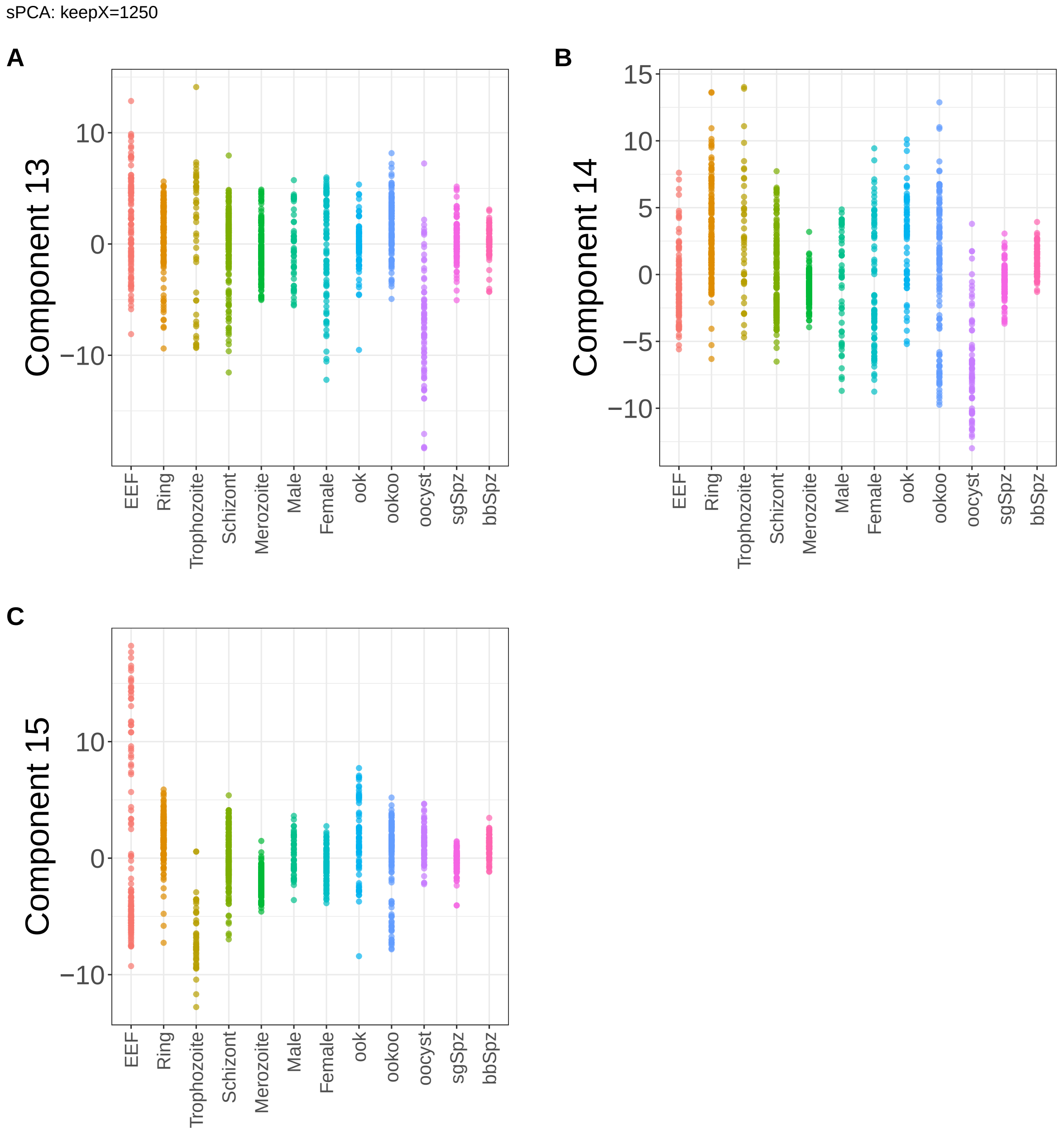


**Supplementary Figure 156.** Components for sPCA: keepX=1250. (A) Component 13. (B) Component 14. (C) Component 15.

sPCA: keepX=2500

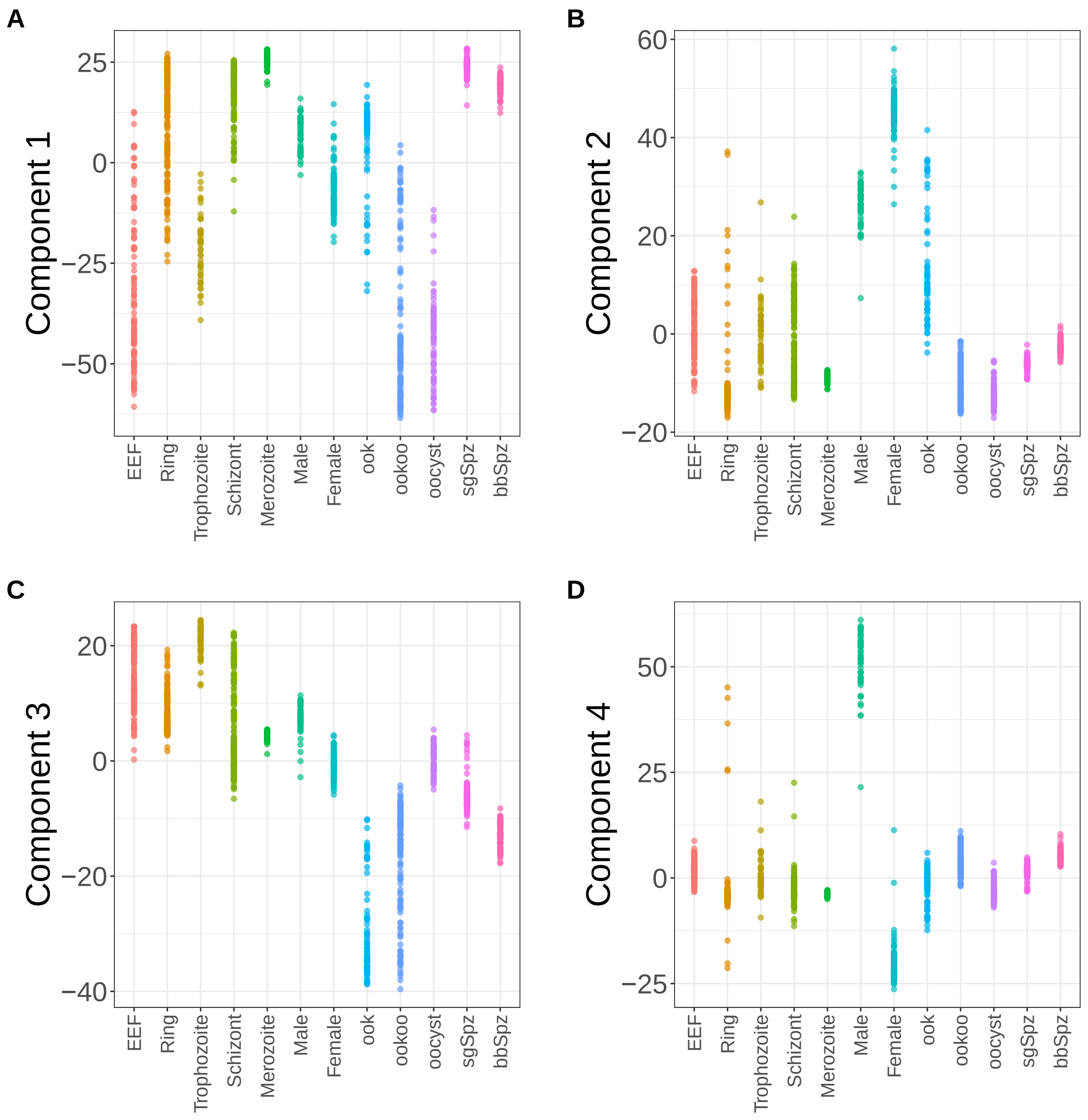


**Supplementary Figure 157.** Components for sPCA: keepX=2500. (A) Component 1. (B) Component 2. (C) Component 3. (D) Component 4.

sPCA: keepX=2500

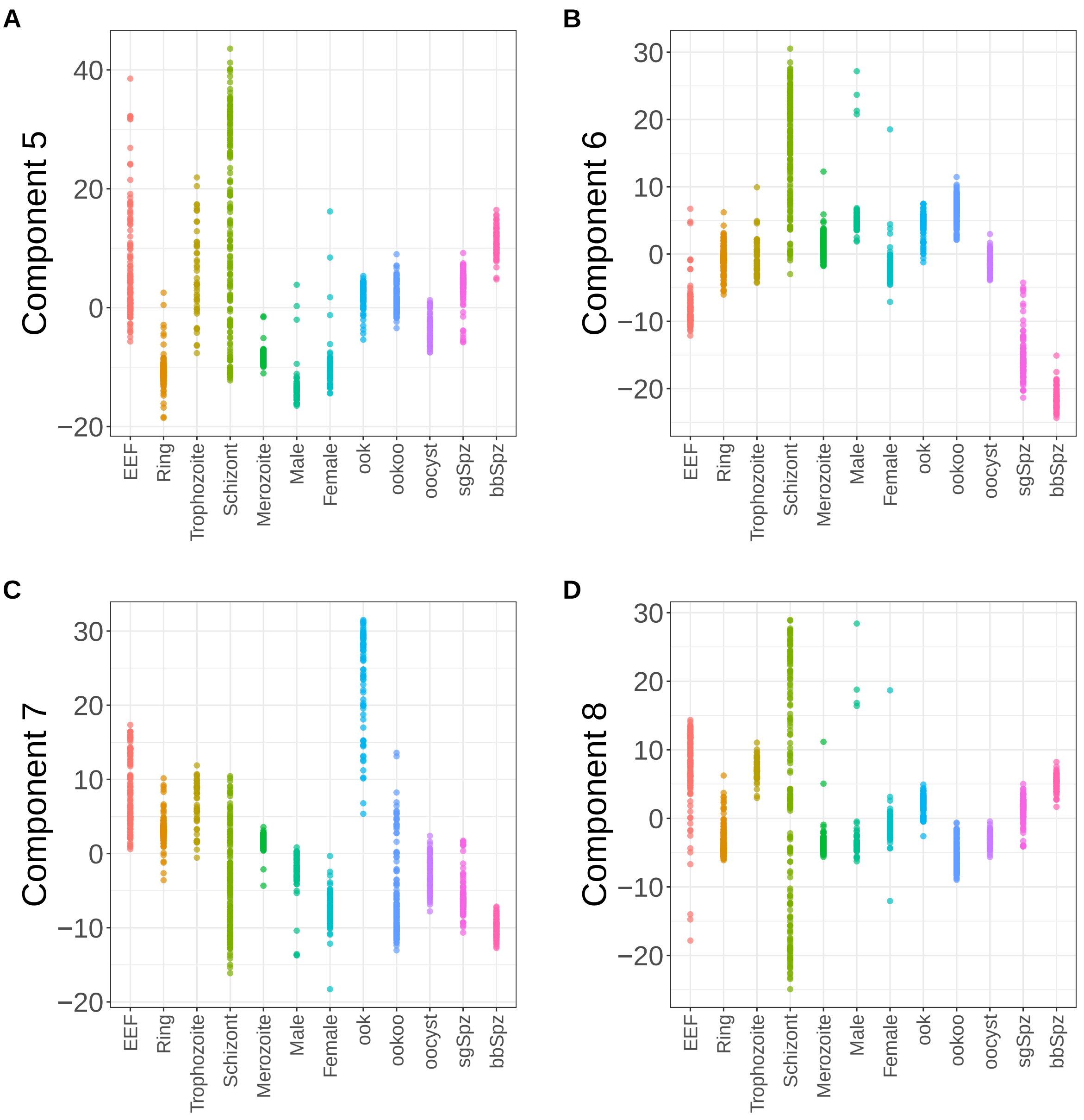


**Supplementary Figure 158.** Components for sPCA: keepX=2500. (A) Component 5. (B) Component 6. (C) Component 7. (D) Component 8.

sPCA: keepX=2500

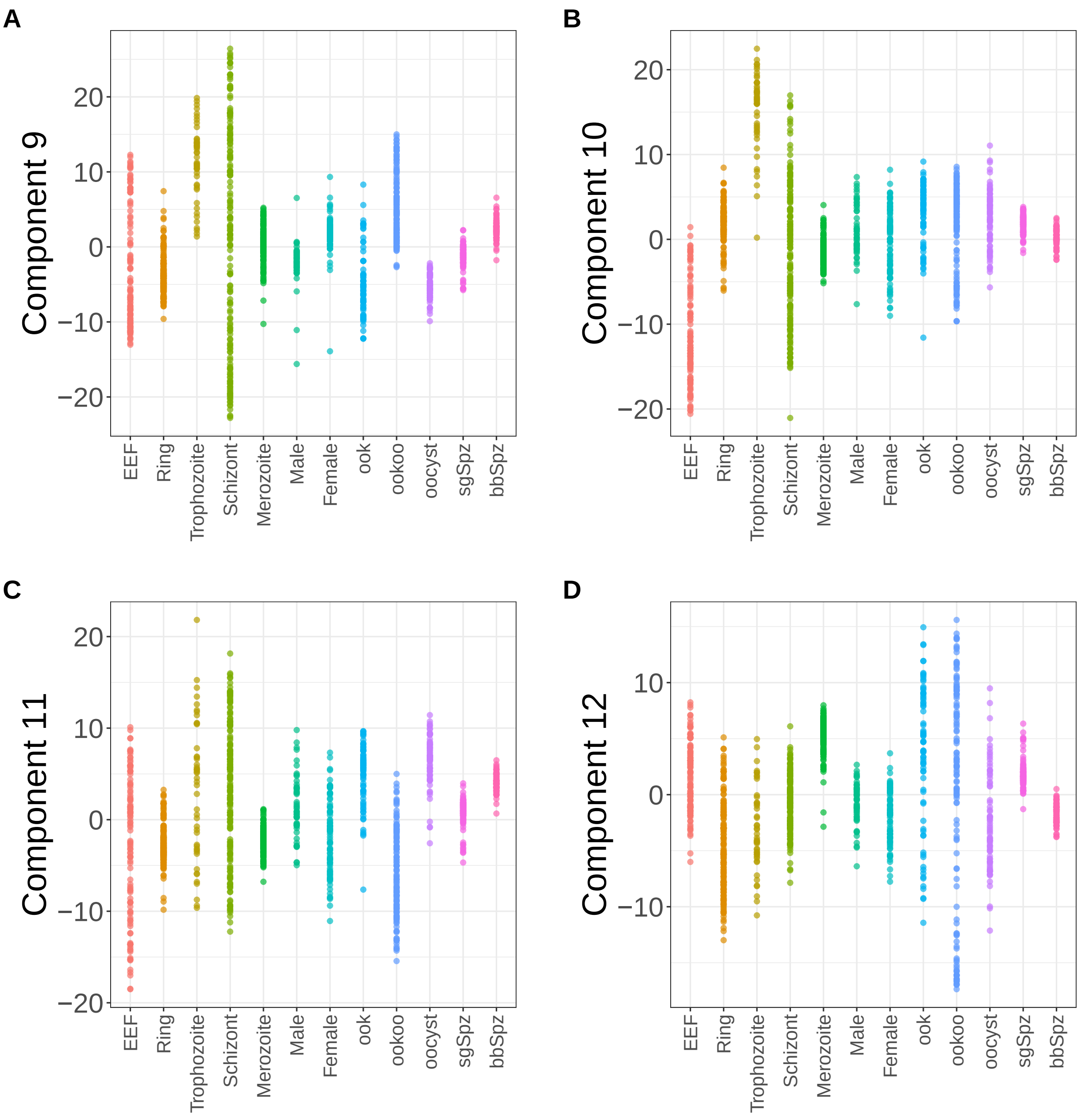


**Supplementary Figure 159.** Components for sPCA: keepX=2500. (A) Component 9. (B) Component 10. (C) Component 11. (D) Component 12.

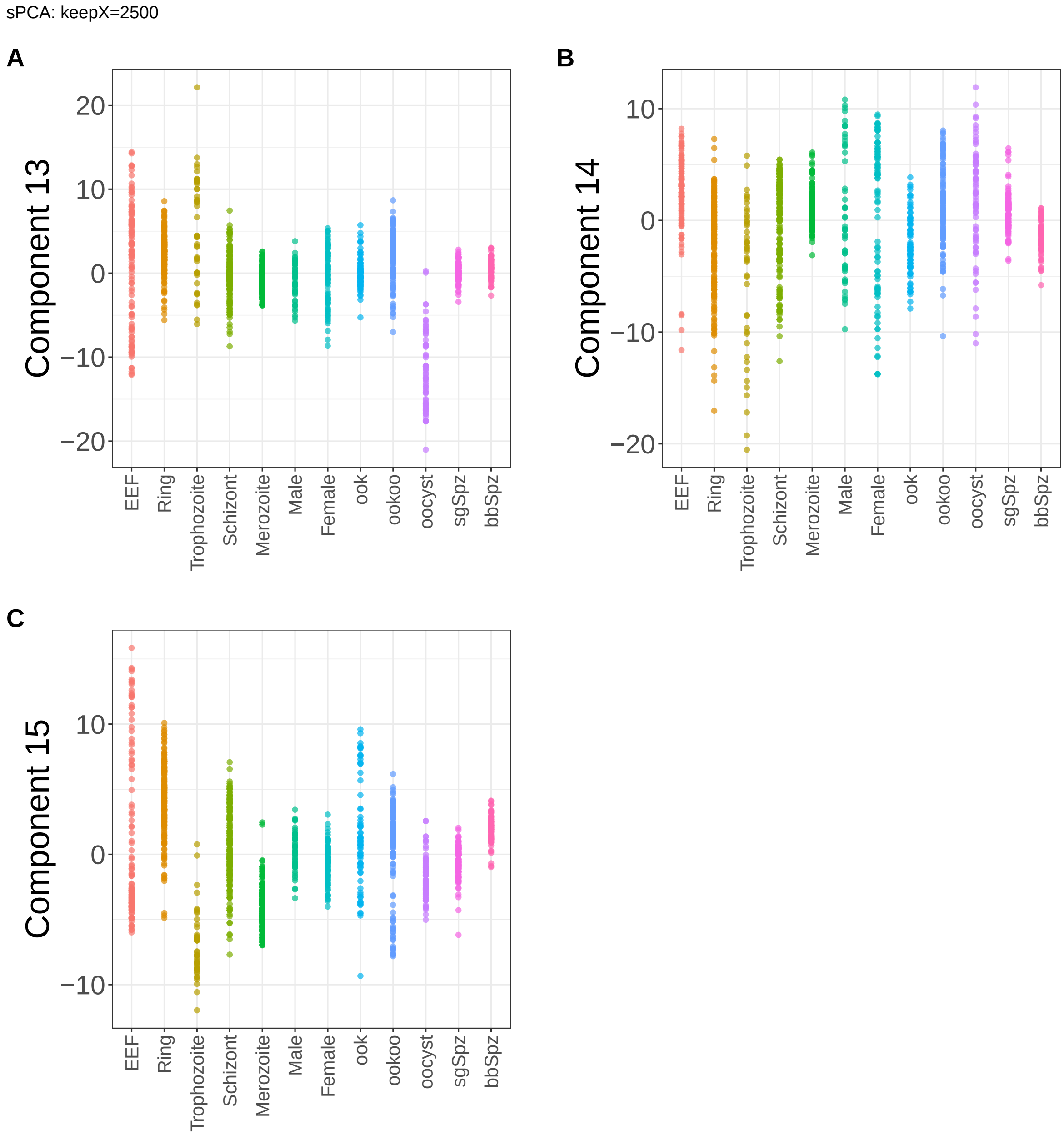


**Supplementary Figure 160.** Components for sPCA: keepX=2500. (A) Component 13. (B) Component 14. (C) Component 15.

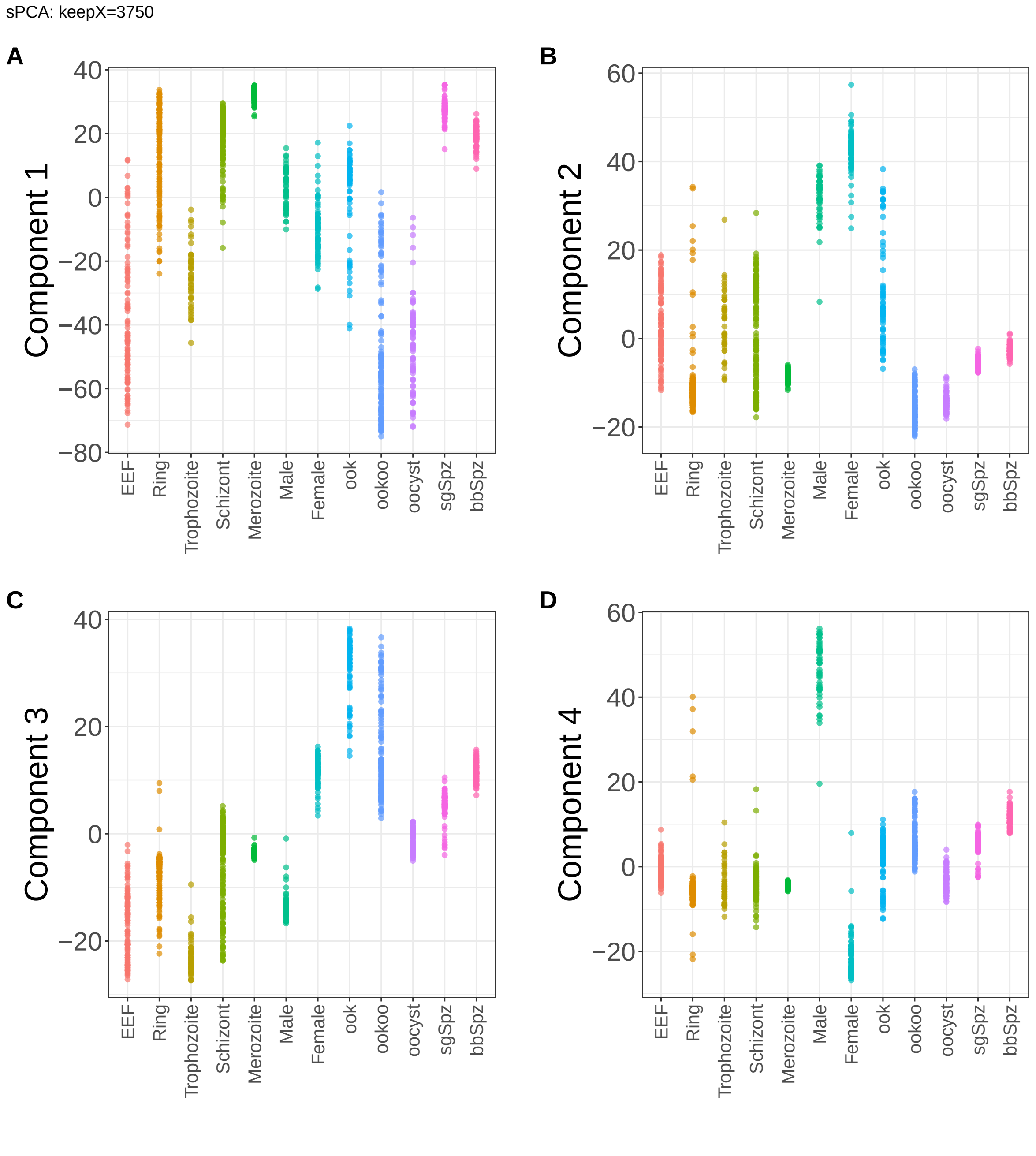


**Supplementary Figure 161.** Components for sPCA: keepX=3750. (A) Component 1. (B) Component 2. (C) Component 3. (D) Component 4.

sPCA: keepX=3750

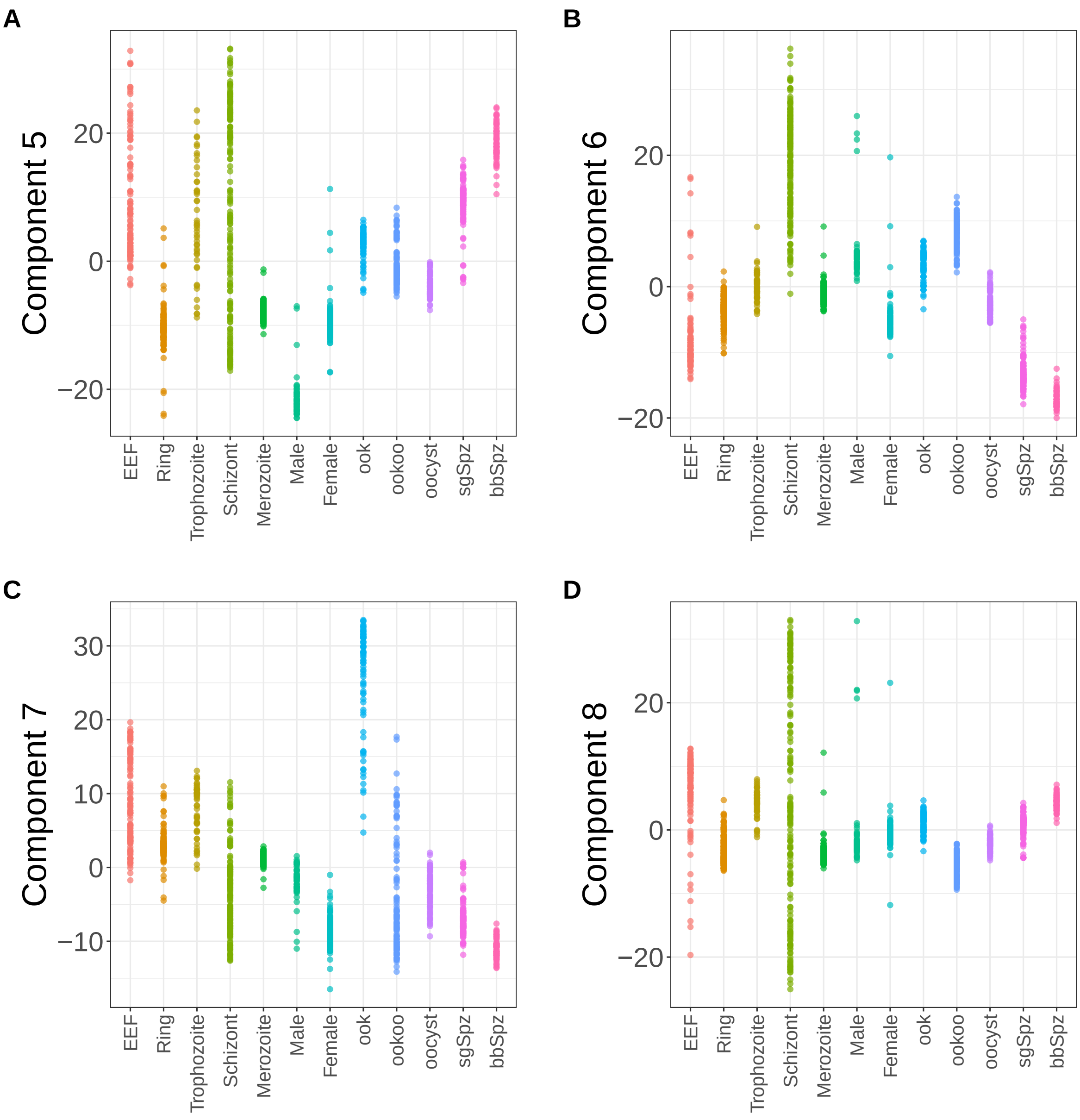


**Supplementary Figure 162.** Components for sPCA: keepX=3750. (A) Component 5. (B) Component 6. (C) Component 7. (D) Component 8.

sPCA: keepX=3750

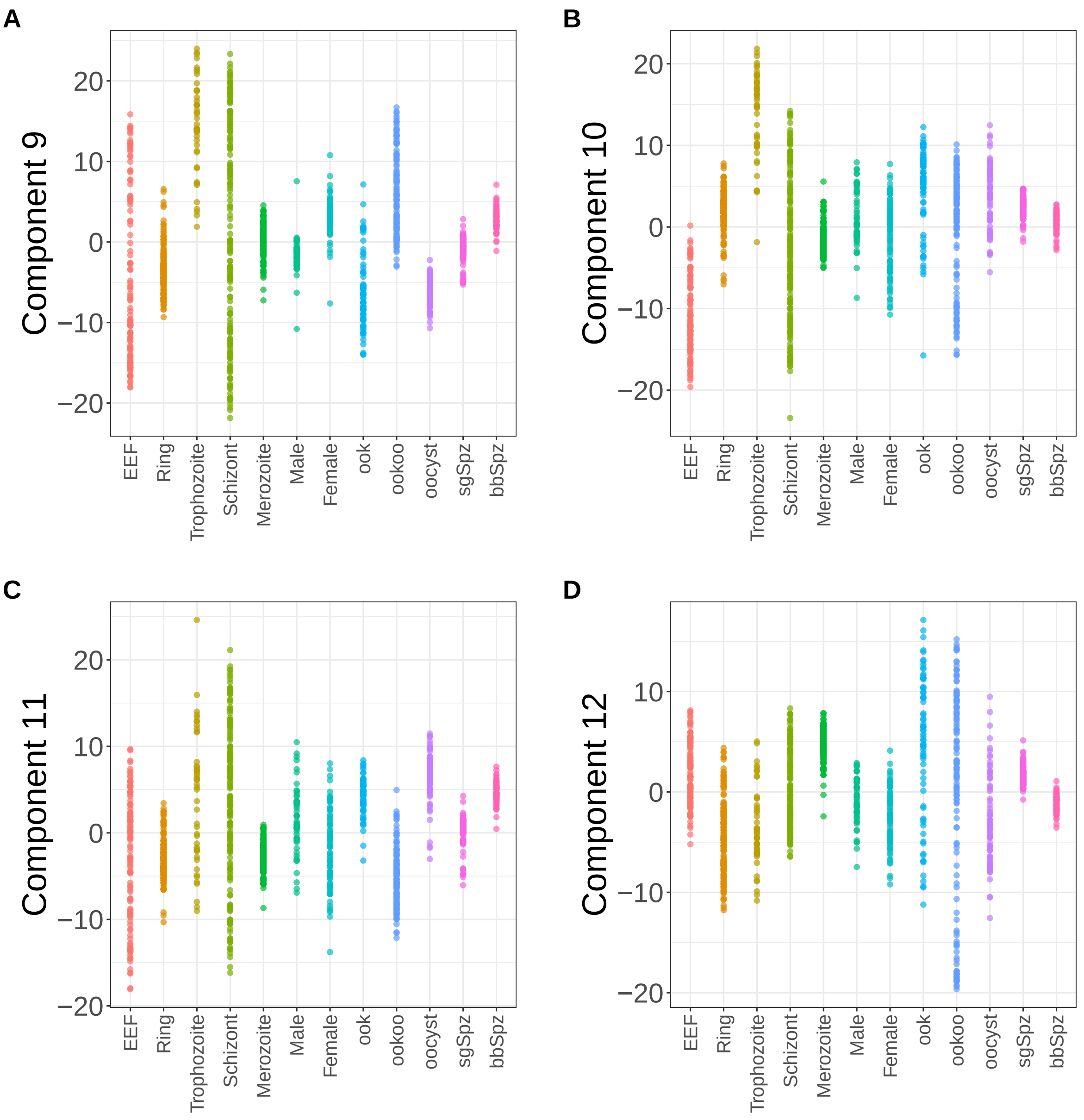


**Supplementary Figure 163.** Components for sPCA: keepX=3750. (A) Component 9. (B) Component 10. (C) Component 11. (D) Component 12.

sPCA: keepX=3750

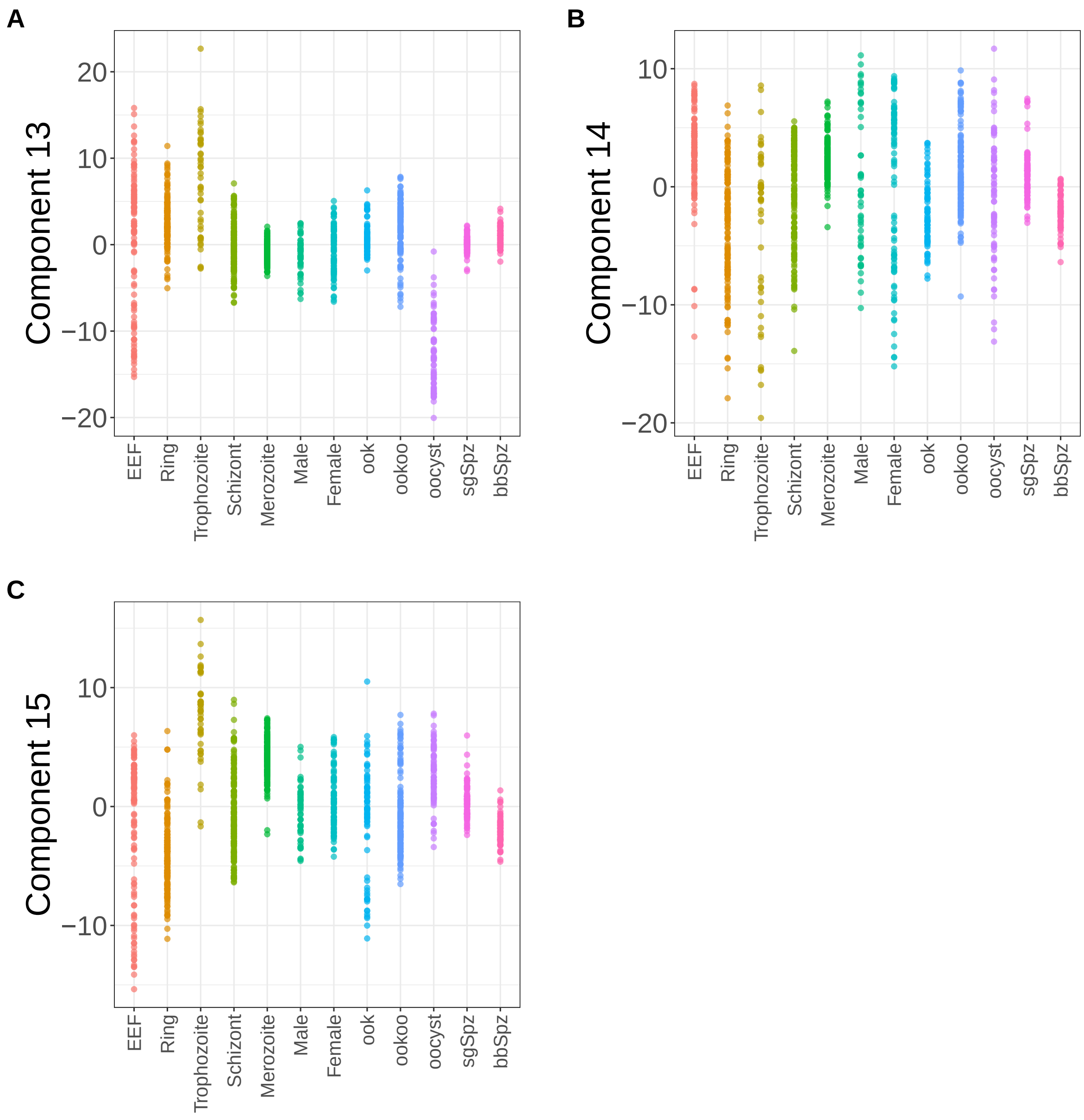


**Supplementary Figure 164.** Components for sPCA: keepX=3750. (A) Component 13. (B) Component 14. (C) Component 15.

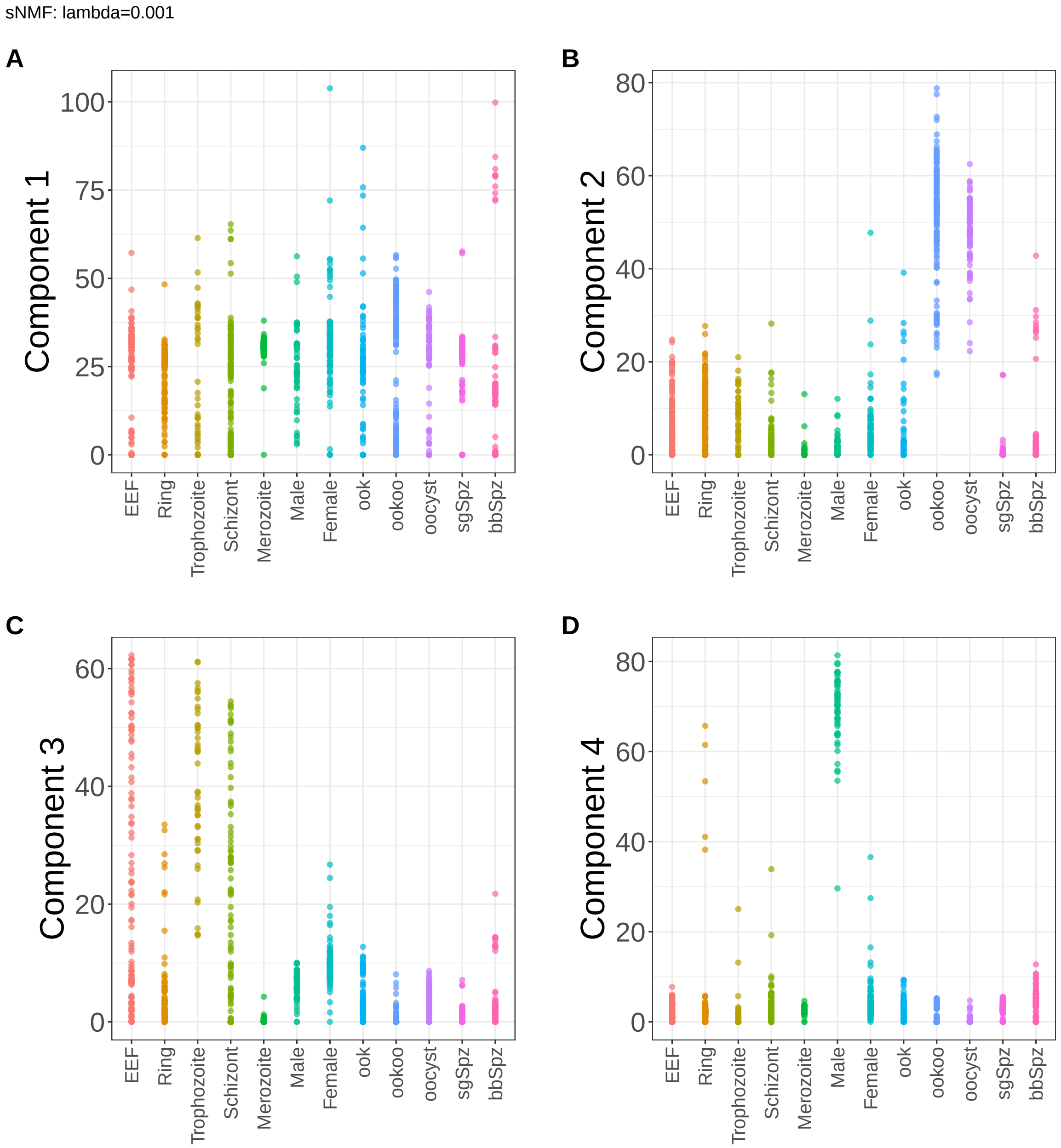


**Supplementary Figure 165.** Components for sNMF: lambda=0.001. (A) Component 1. (B) Component 2. (C) Component 3. (D) Component 4.

sNMF: lambda=0.001

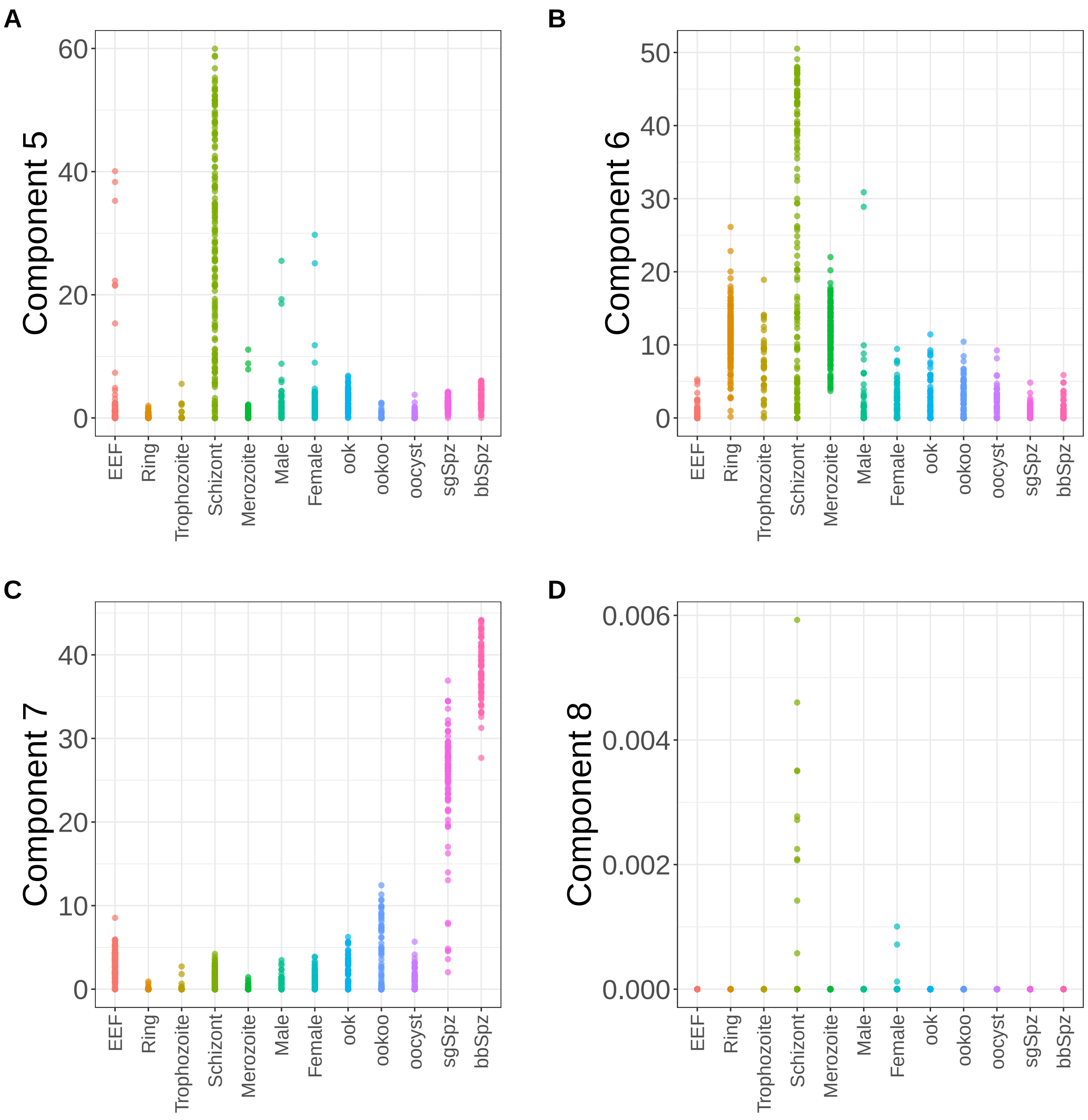


**Supplementary Figure 166.** Components for sNMF: lambda=0.001. (A) Component 5. (B) Component 6. (C) Component 7. (D) Component 8.

sNMF: lambda=0.001

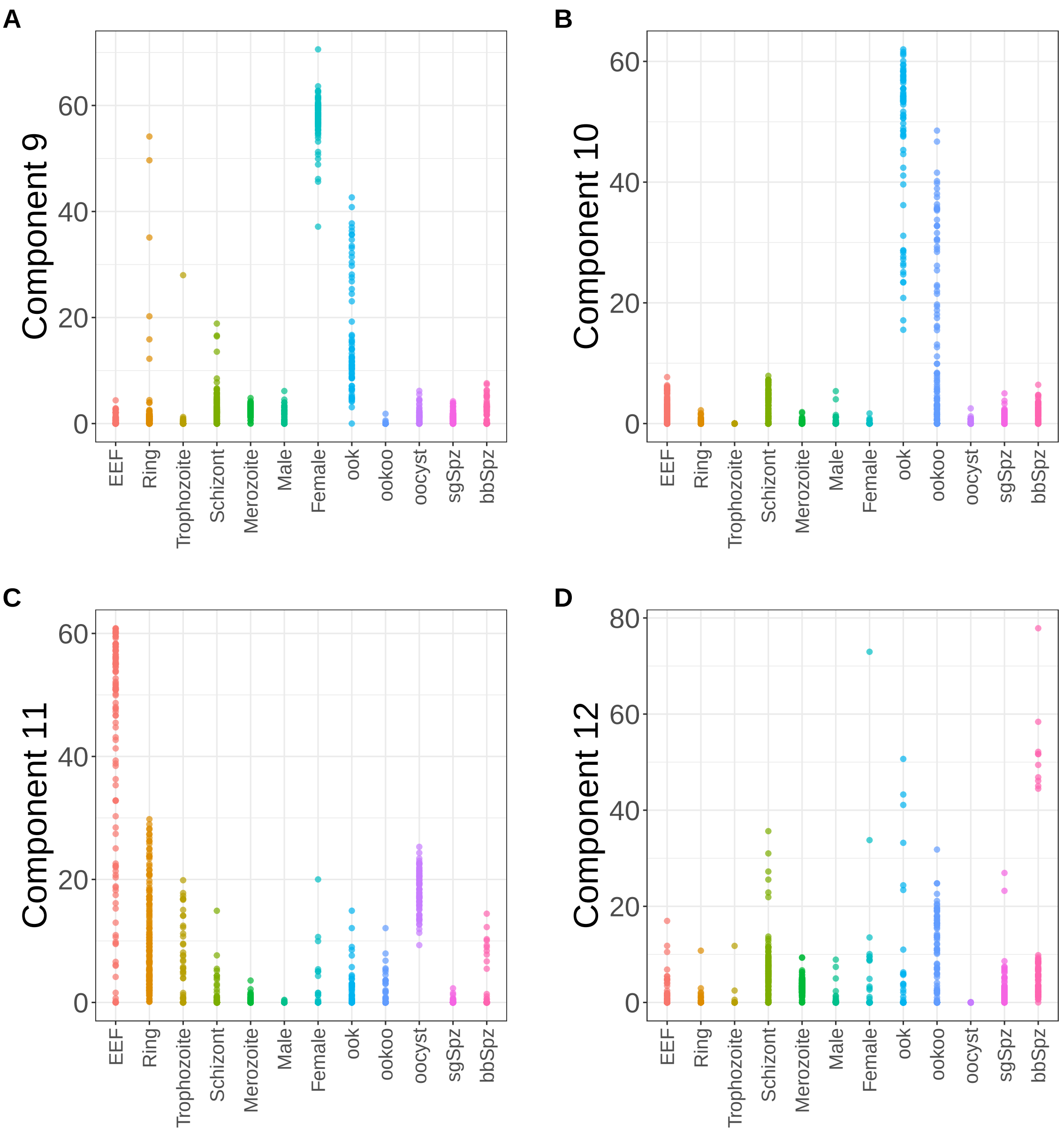


**Supplementary Figure 167.** Components for sNMF: lambda=0.001. (A) Component 9. (B) Component 10. (C) Component 11. (D) Component 12.

sNMF: lambda=0.001

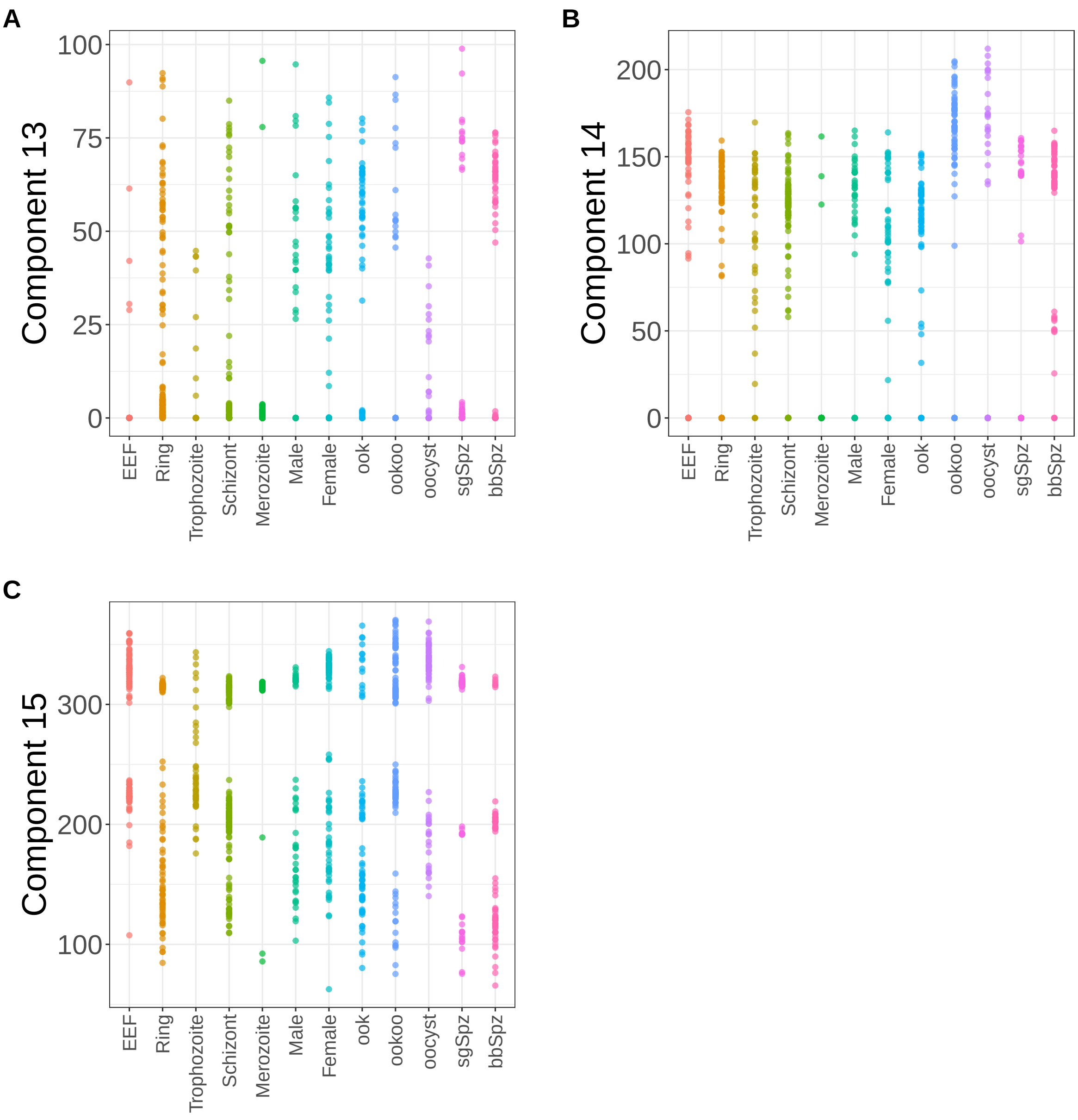


**Supplementary Figure 168.** Components for sNMF: lambda=0.001. (A) Component 13. (B) Component 14. (C) Component 15.

sNMF: lambda=0.01

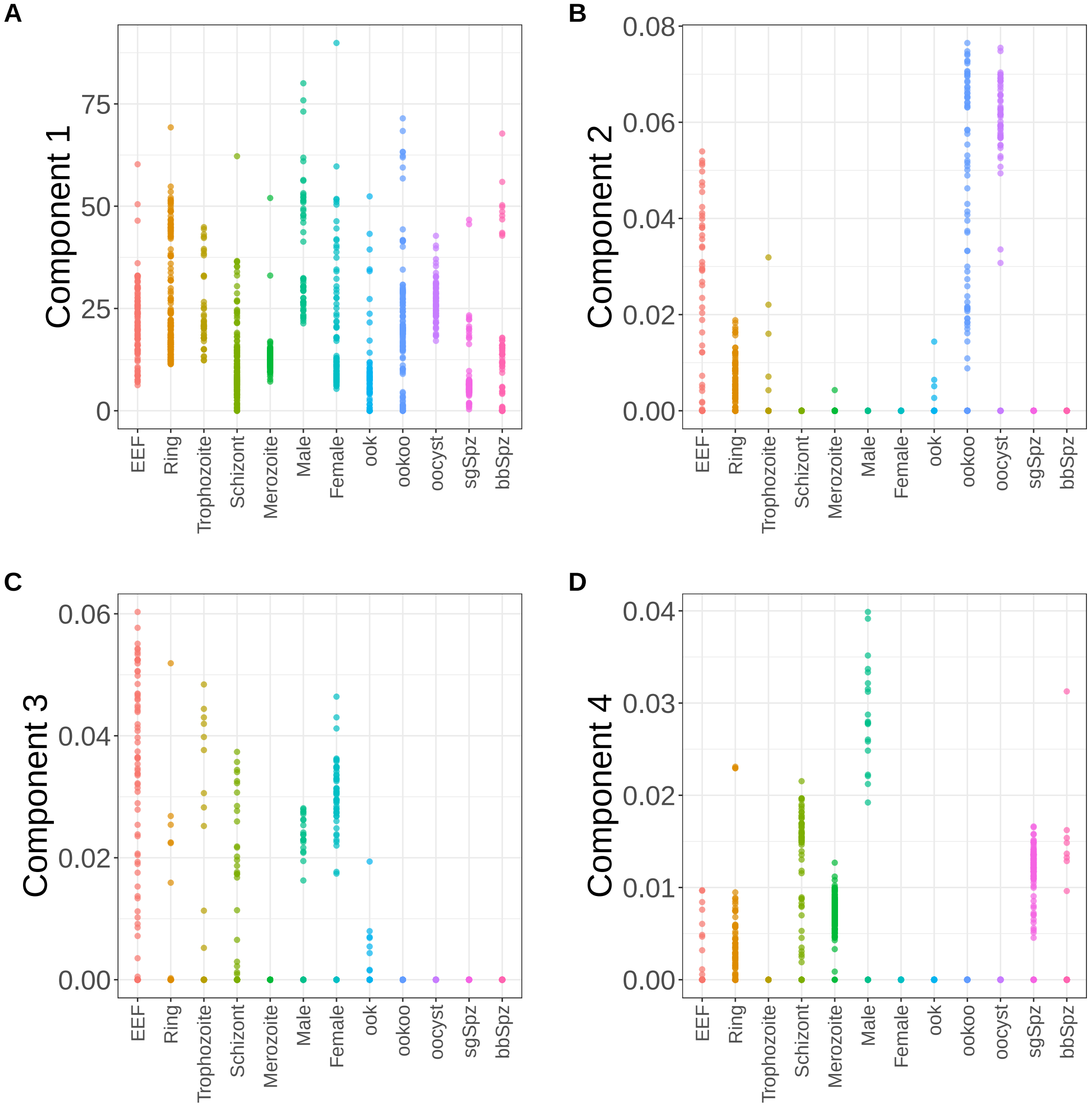


**Supplementary Figure 169.** Components for sNMF: lambda=0.01. (A) Component 1. (B) Component 2. (C) Component 3. (D) Component 4.

sNMF: lambda=0.01

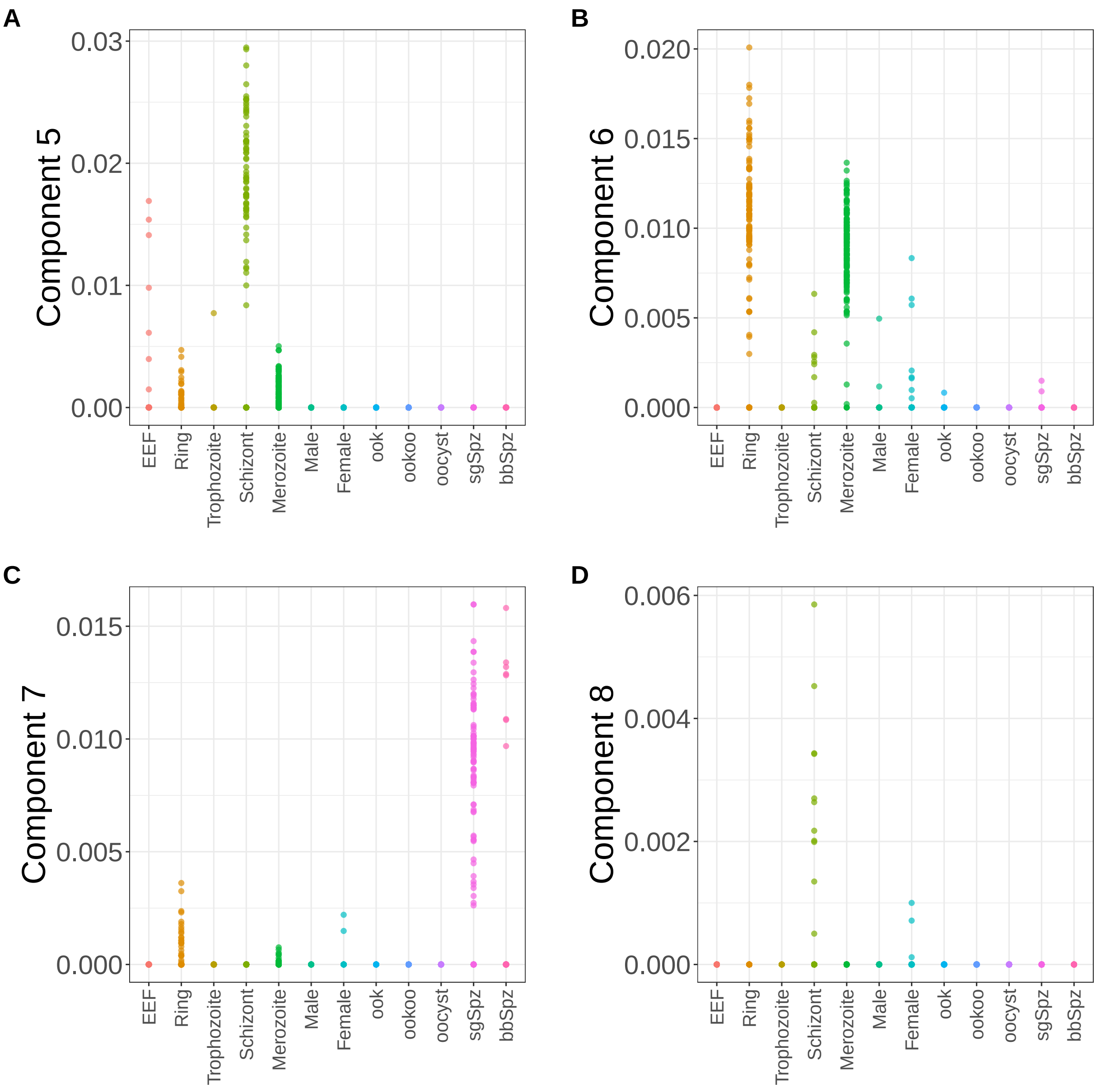


**Supplementary Figure 170.** Components for sNMF: lambda=0.01. (A) Component 5. (B) Component 6. (C) Component 7. (D) Component 8.

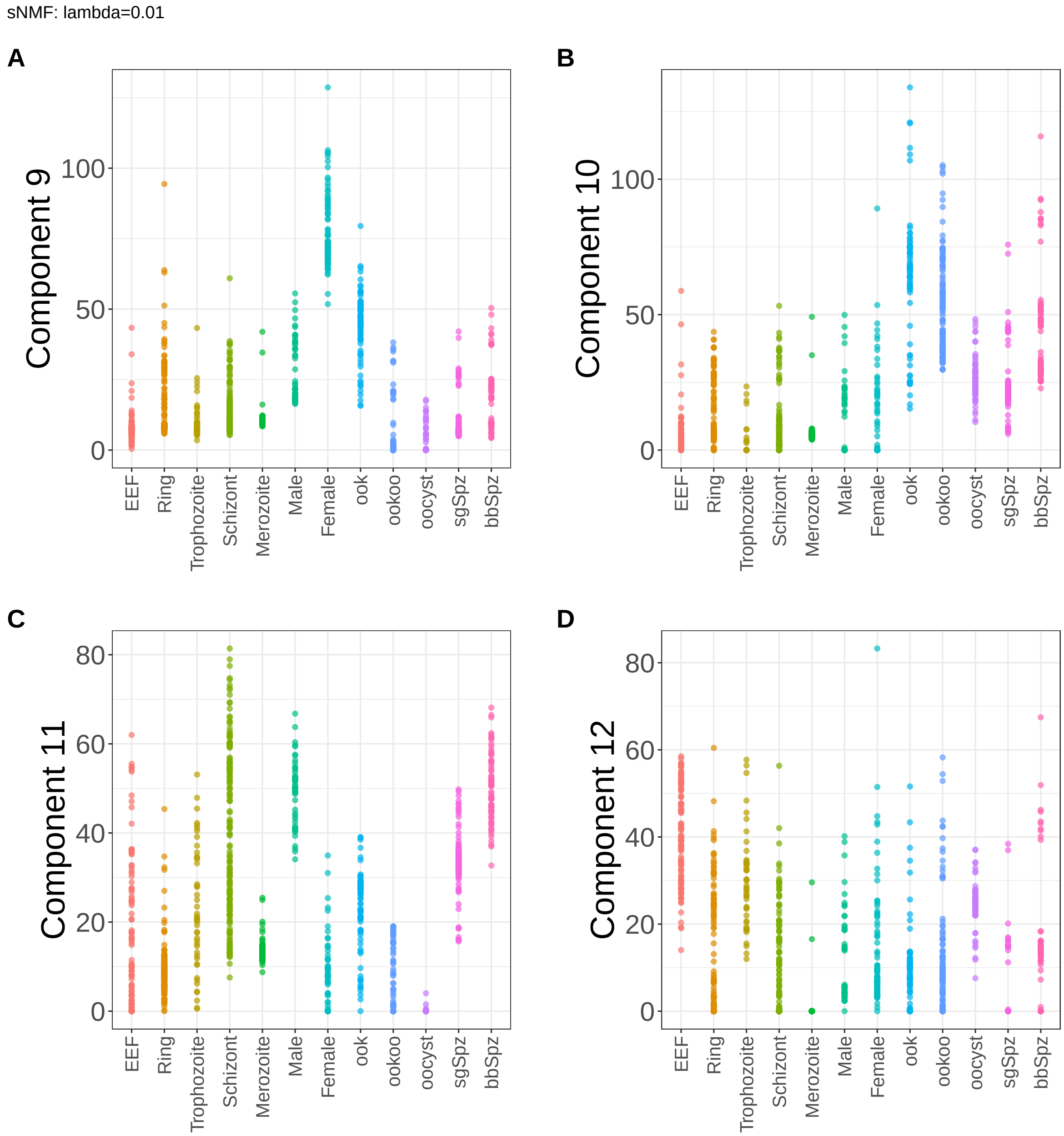


**Supplementary Figure 171.** Components for sNMF: lambda=0.01. (A) Component 9. (B) Component 10. (C) Component 11. (D) Component 12.

sNMF: lambda=0.01

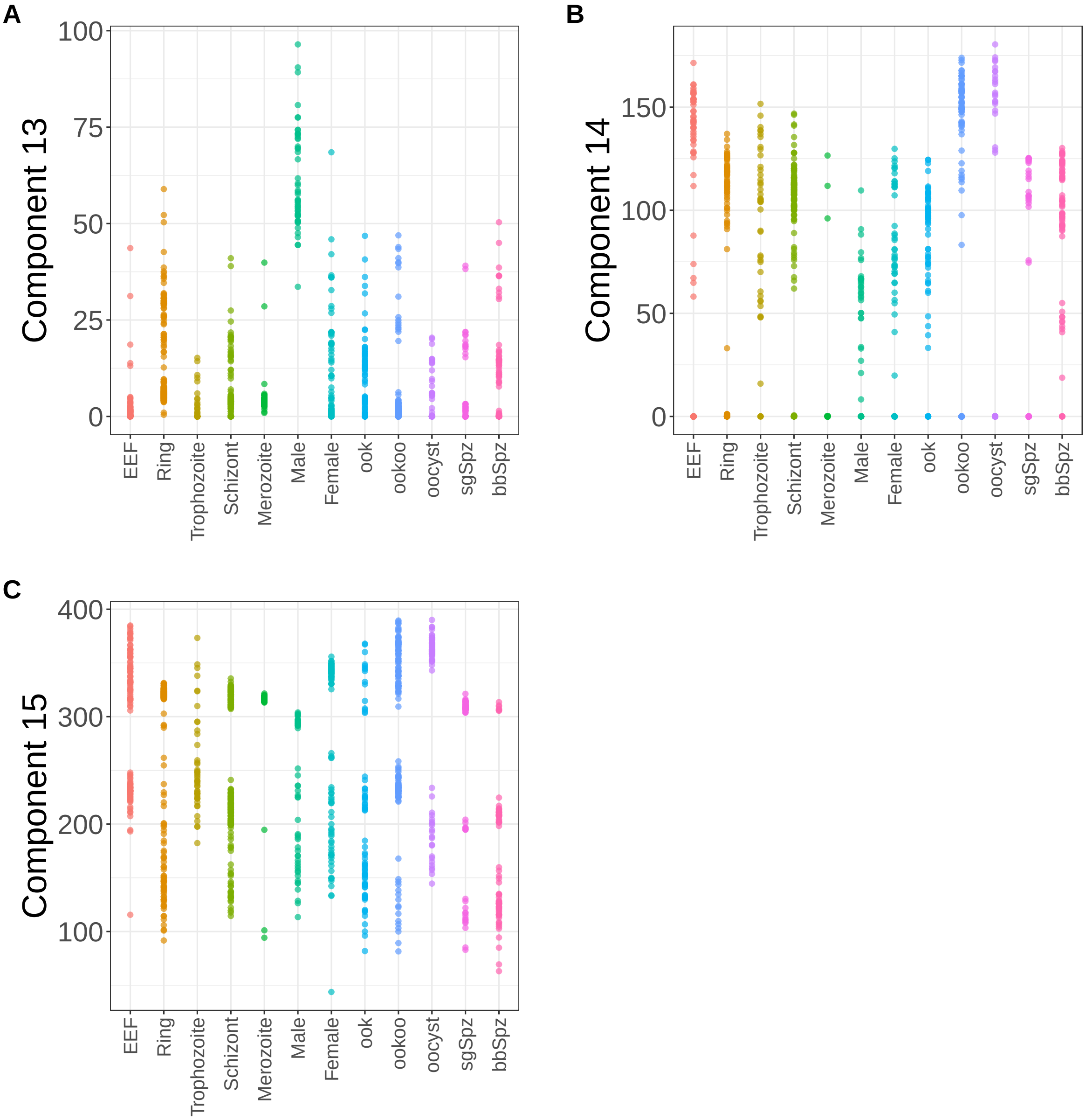


**Supplementary Figure 172.** Components for sNMF: lambda=0.01. (A) Component 13. (B) Component 14. (C) Component 15.

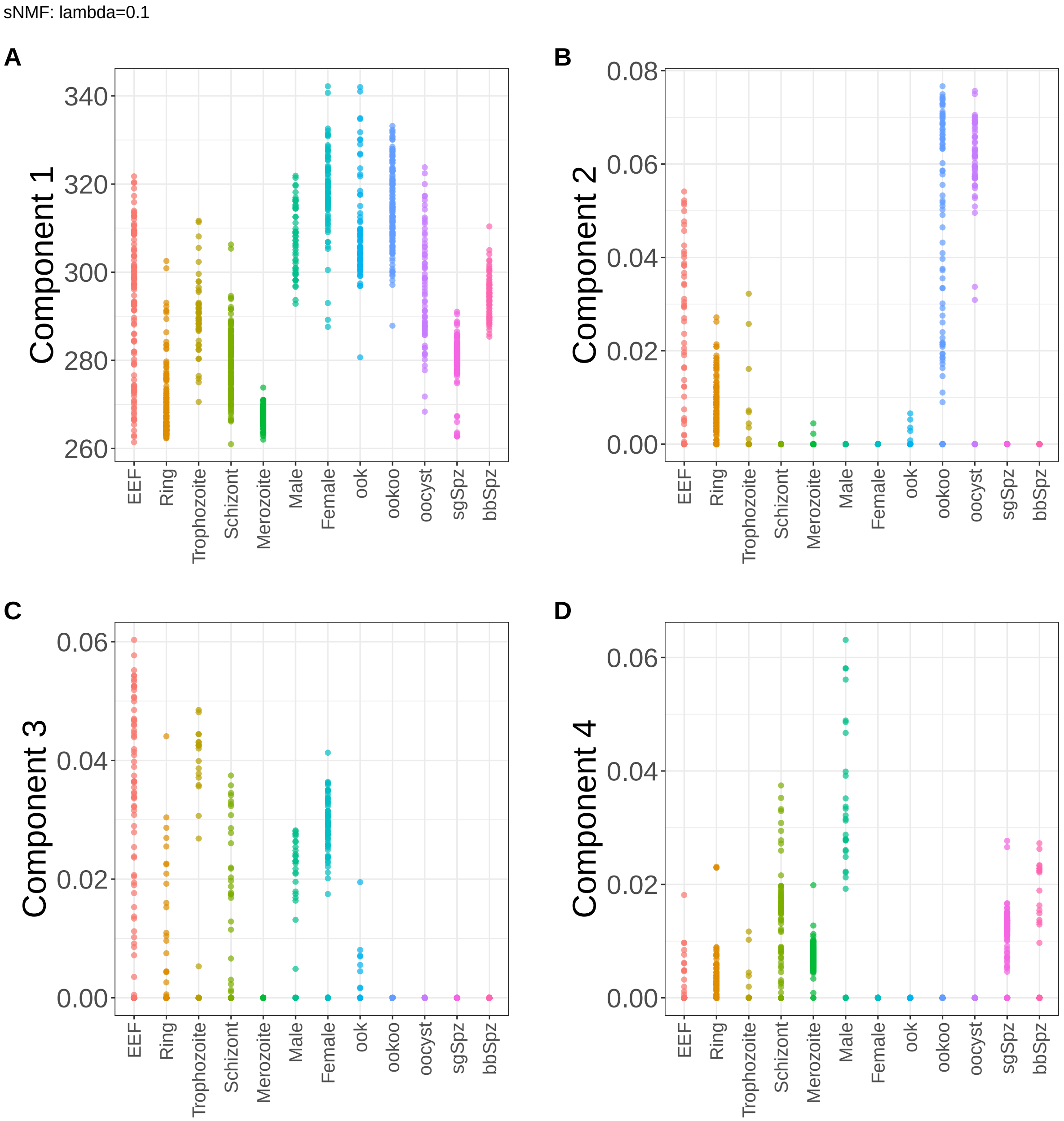


**Supplementary Figure 173.** Components for sNMF: lambda=0.1. (A) Component 1. (B) Component 2. (C) Component 3. (D) Component 4.

sNMF: lambda=0.1

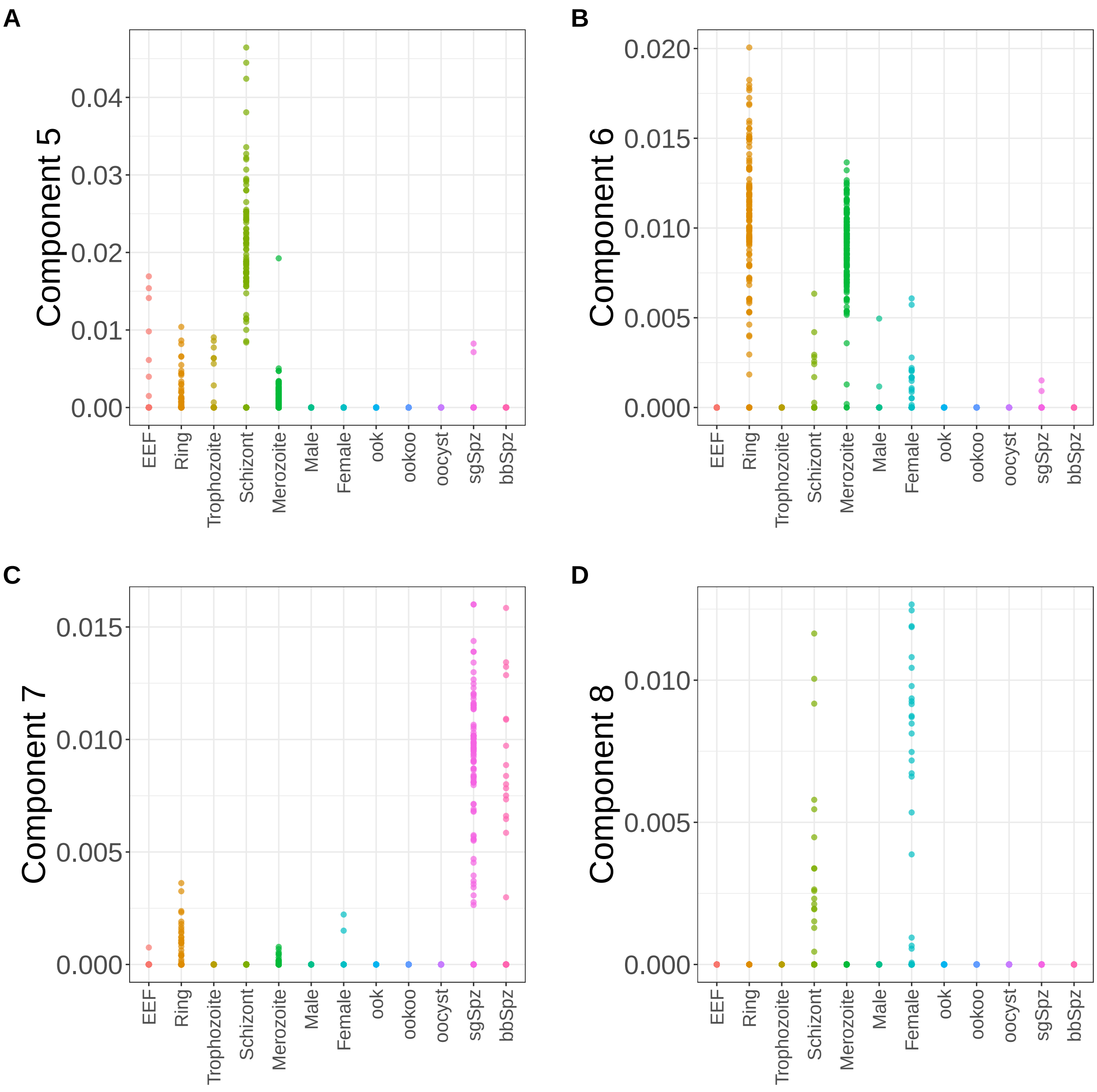


**Supplementary Figure 174.** Components for sNMF: lambda=0.1. (A) Component 5. (B) Component 6. (C) Component 7. (D) Component 8.

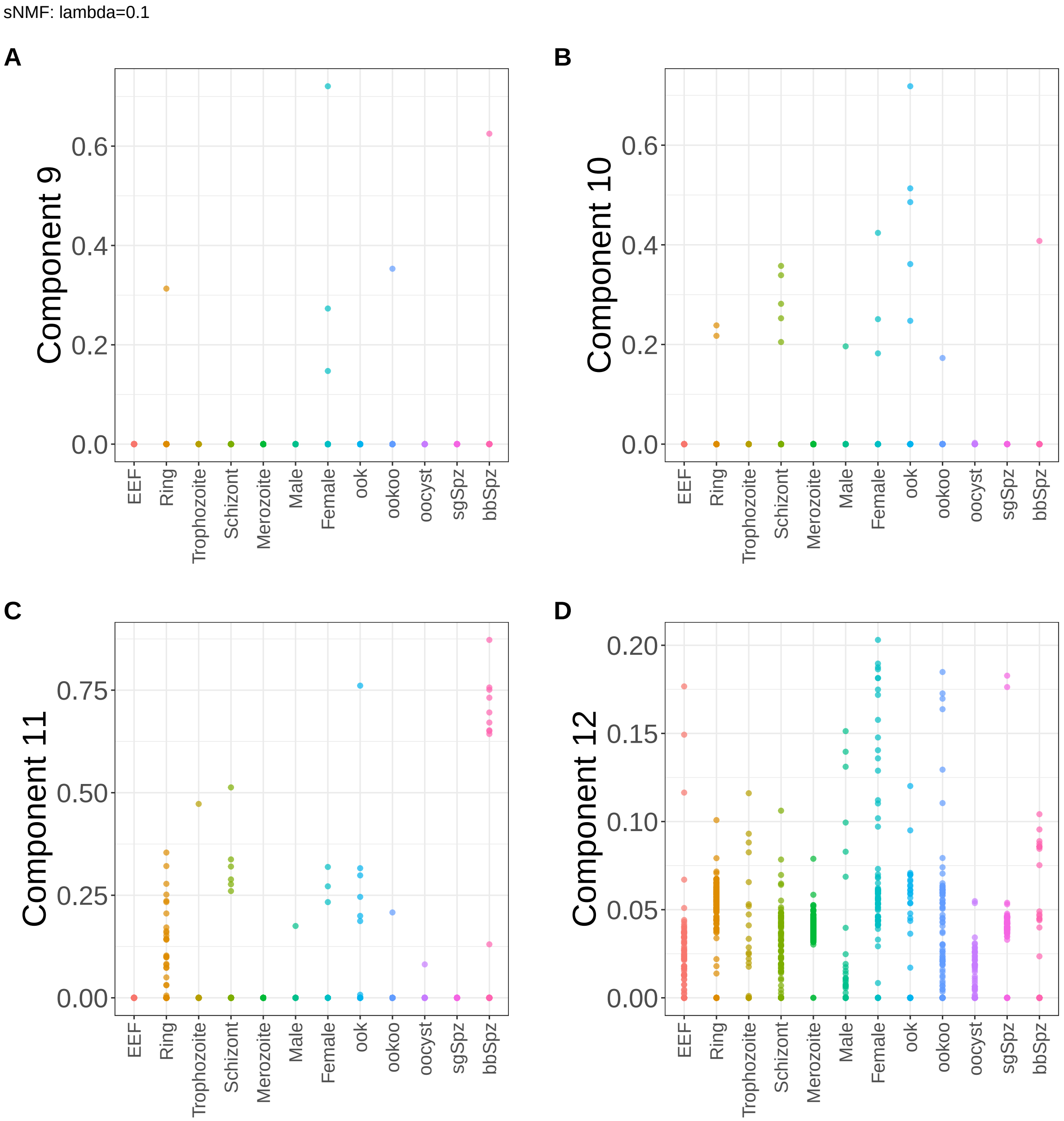


**Supplementary Figure 175.** Components for sNMF: lambda=0.1. (A) Component 9. (B) Component 10. (C) Component 11. (D) Component 12.

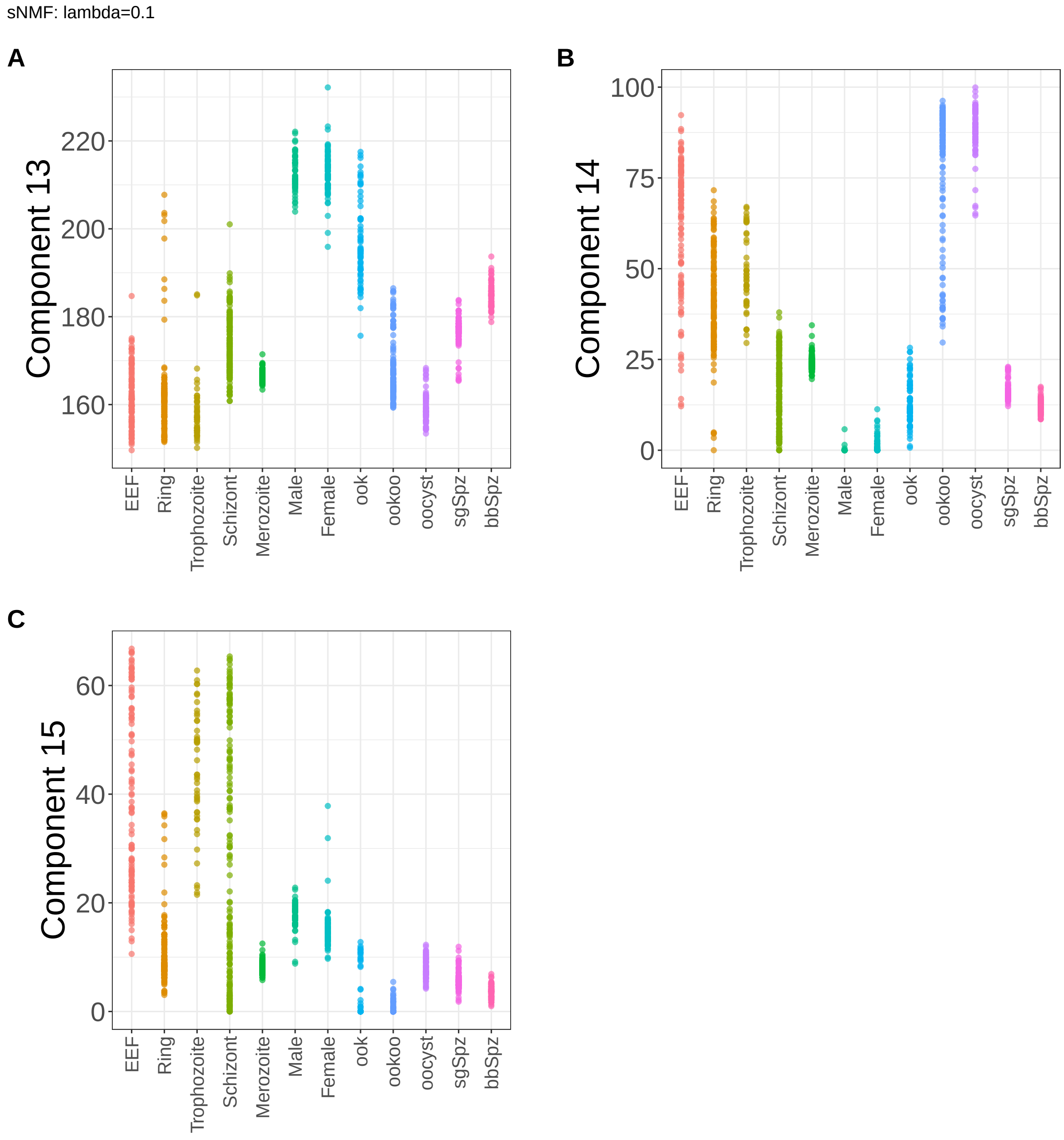


**Supplementary Figure 176.** Components for sNMF: lambda=0.1. (A) Component 13. (B) Component 14. (C) Component 15.
