## Supplementary Note 3 : Bilateral Mouse Models for "A statistical framework for system-level synthesis"

This note presents the supplementary results from the analysis of anti-PD-L1 immune checkpoint blockade data derived from bilateral AB1 and RENCA mouse models of mesothelioma and renal carcinoma, respectively. The mice were categorized as responders (RS) and non-responders (NR) for treatment.

The figures include:

- Processes/component functions learned by SSANOVA-SVD (**Supplementary Fig. 54**)
- Inference of feature participation by the application of the three-step framework: Volcano plots (**Supplementary Fig. 55**), heatmaps of top 50 component-associated features at FDR of 10% (**Supplementary Fig. 56**-**Supplementary Fig. 57**), geneset pathway enrichment for associated features (**Supplementary Fig. 58**)
- Decomposition of the processes in terms of effects due to their covariates and interactions (**Supplementary Fig. 59**-**Supplementary Fig. 60**).
- Components from a representative class of matrix factorization methods (**Supplementary Fig. 61**-**Supplementary Fig. 97**)

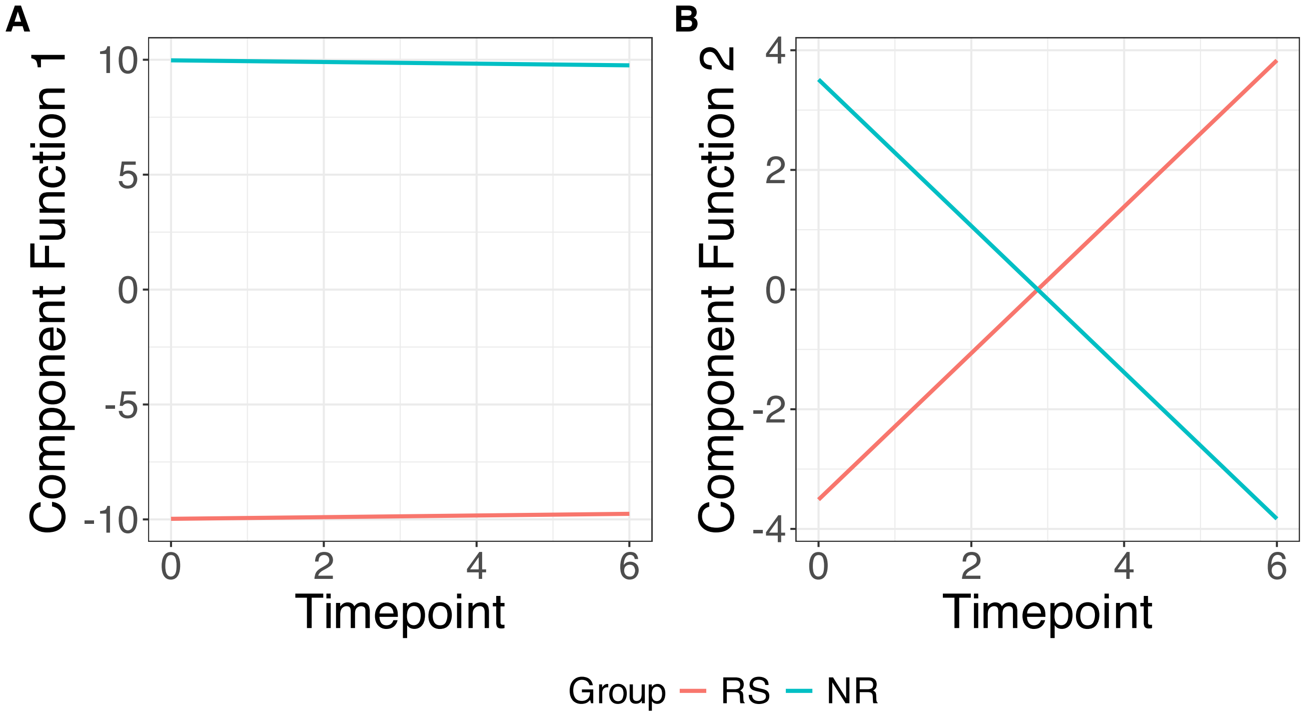


**Supplementary Figure 54.** SSANOVA-SVD reconstructed component functions.(A) Component 1. (B) Component 2.

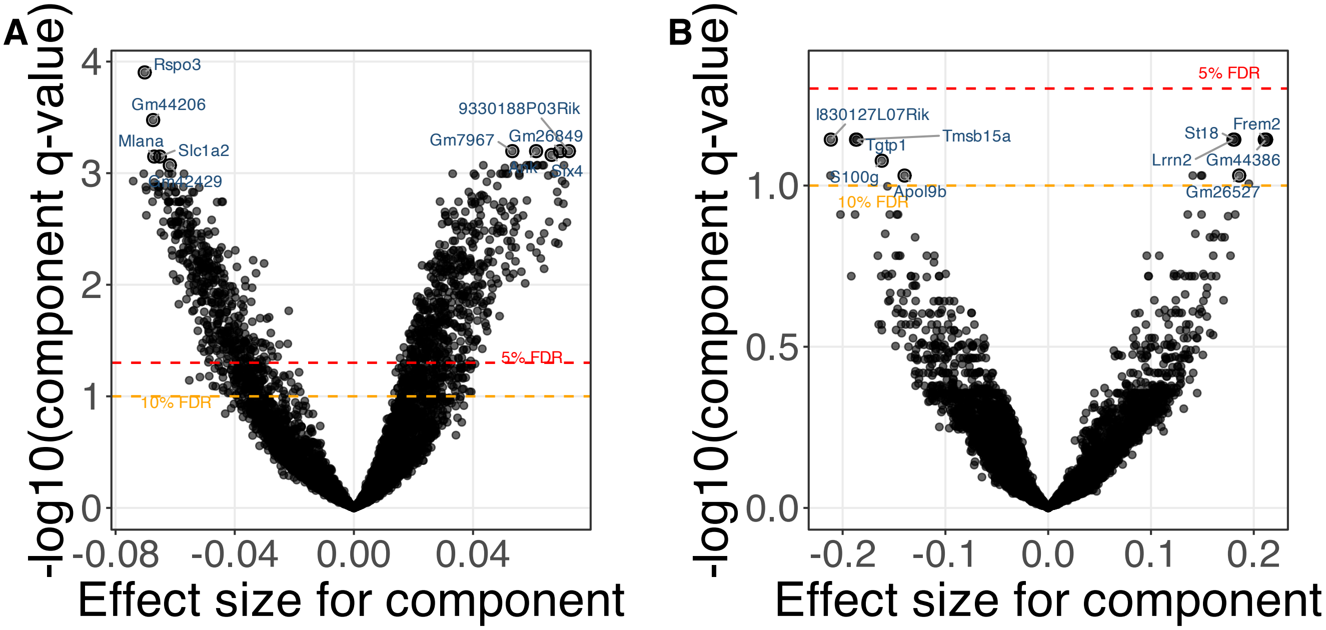


**Supplementary Figure 55.** SSANOVA-SVD based conditional inference for feature participation. (A) Component 1. (B) Component 2.

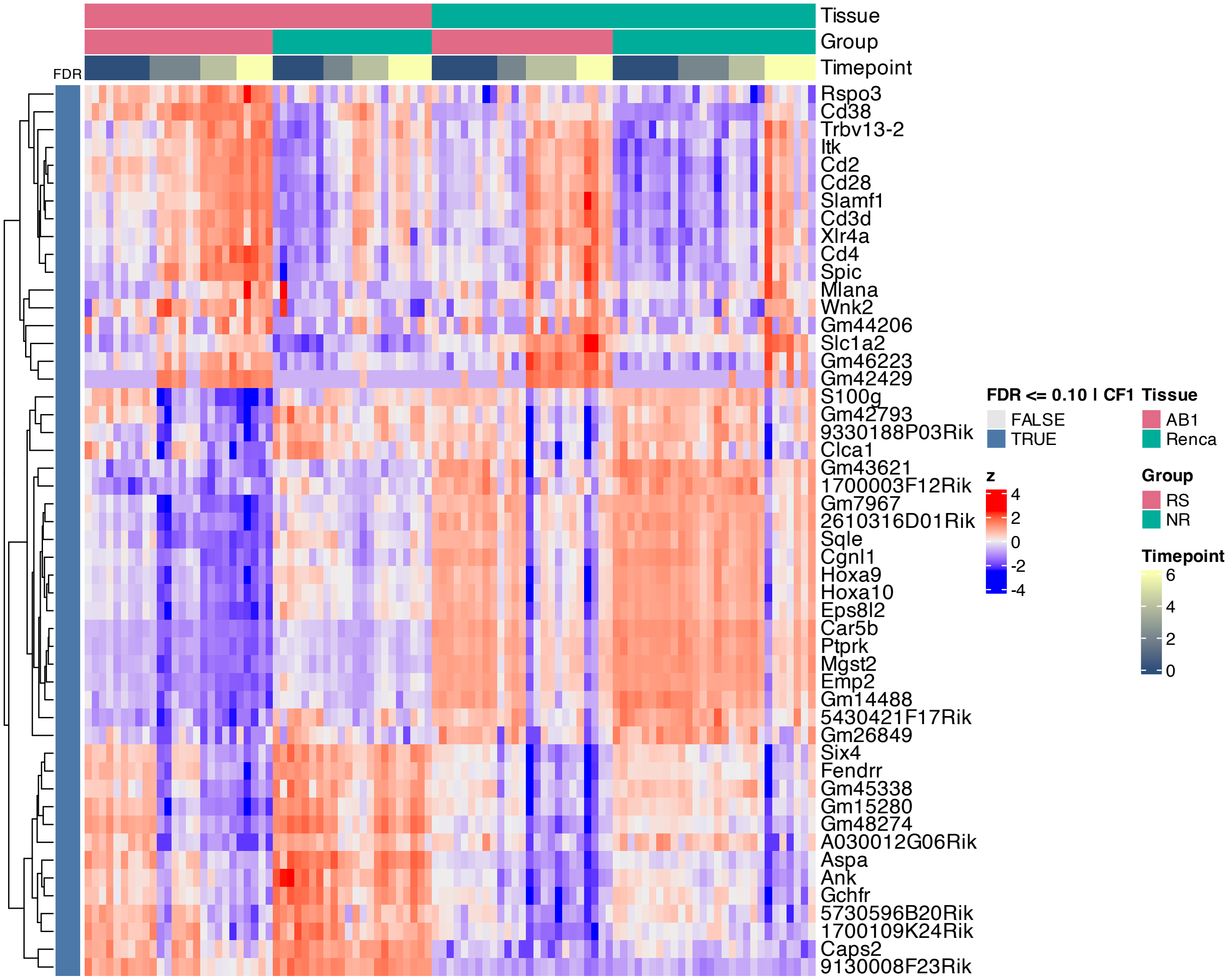


**Supplementary Figure 56.** Heatmap of features inferred to be associated with component 1 by SSANOVA-SVD.

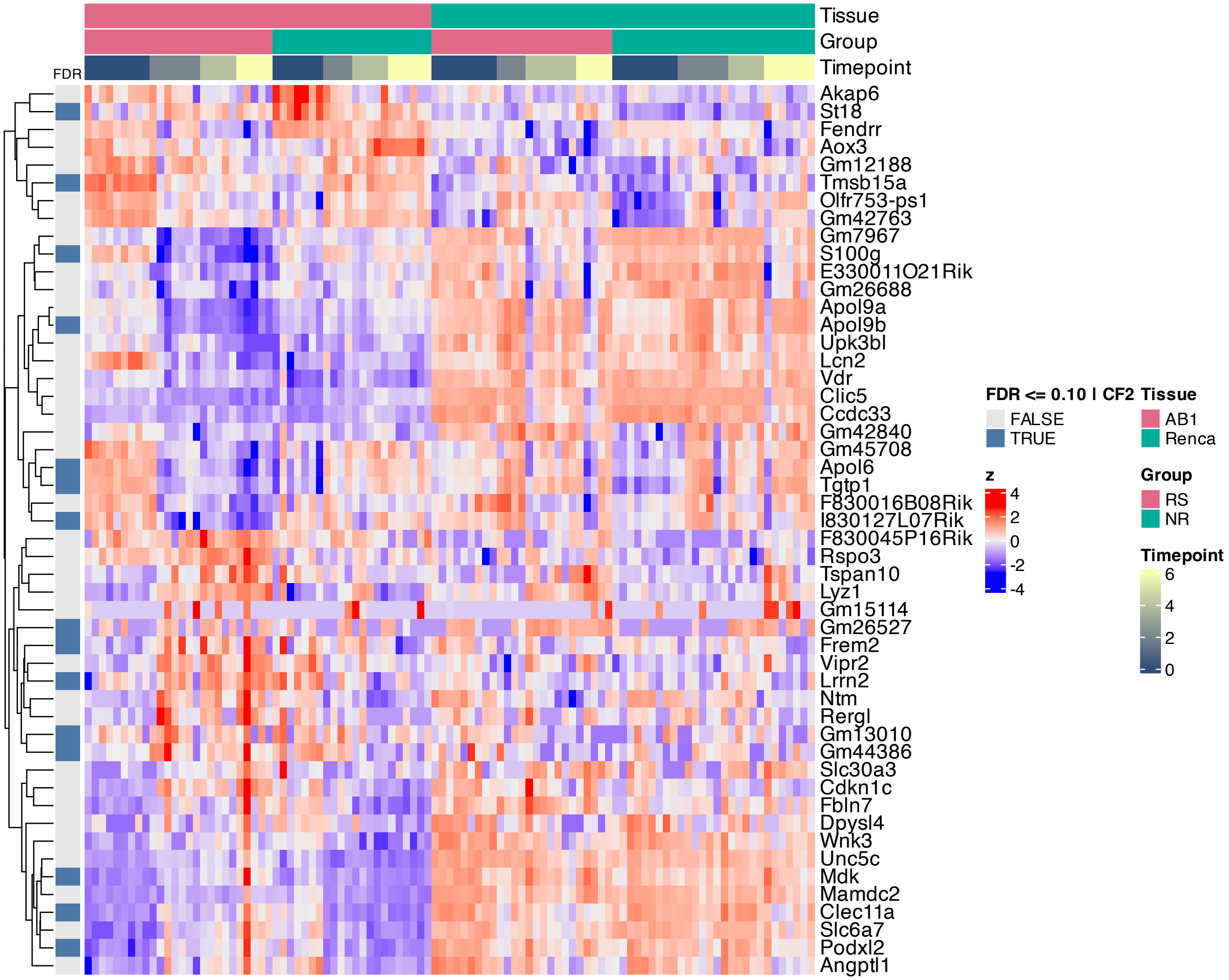


**Supplementary Figure 57.** Heatmap of features inferred to be associated with component 2 by SSANOVA-SVD.

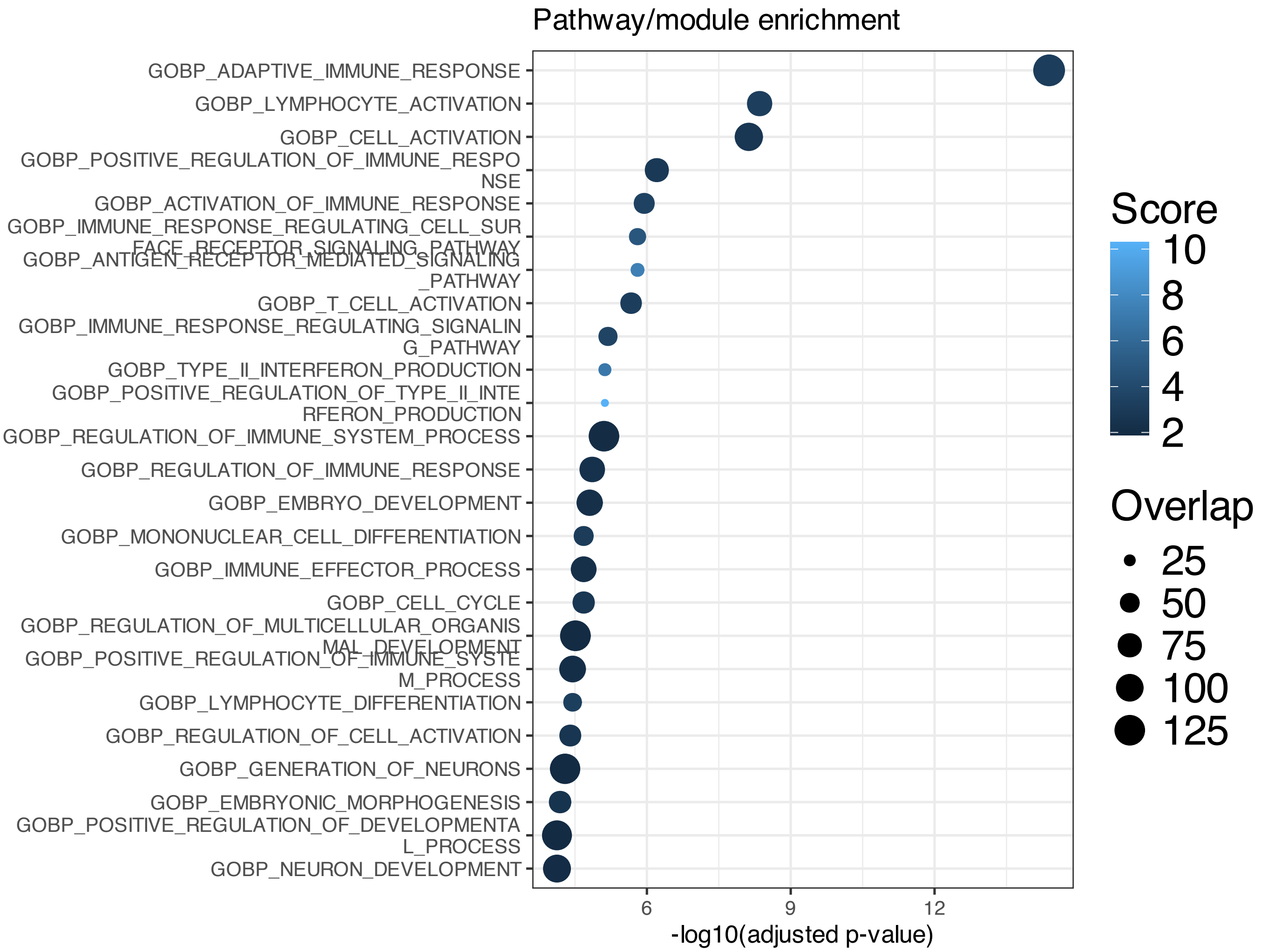


**Supplementary Figure 58.** Enrichment among features inferred to be associated with SSANOVA-SVD component 1; only terms with adjusted P $\leq 0.1$ are shown.

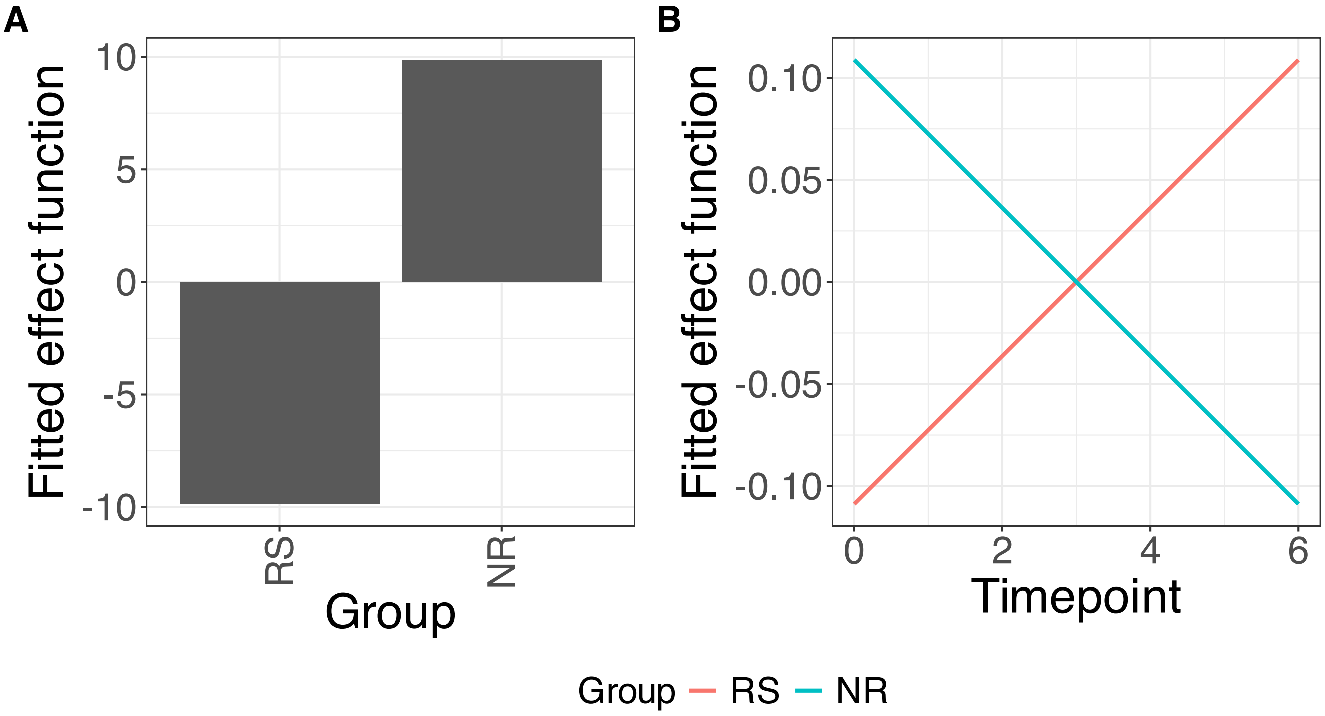


**Supplementary Figure 59.** Decomposition of reconstructed SSANOVA-SVD component function 1. (A) Group. (B) Group:Timepoint.

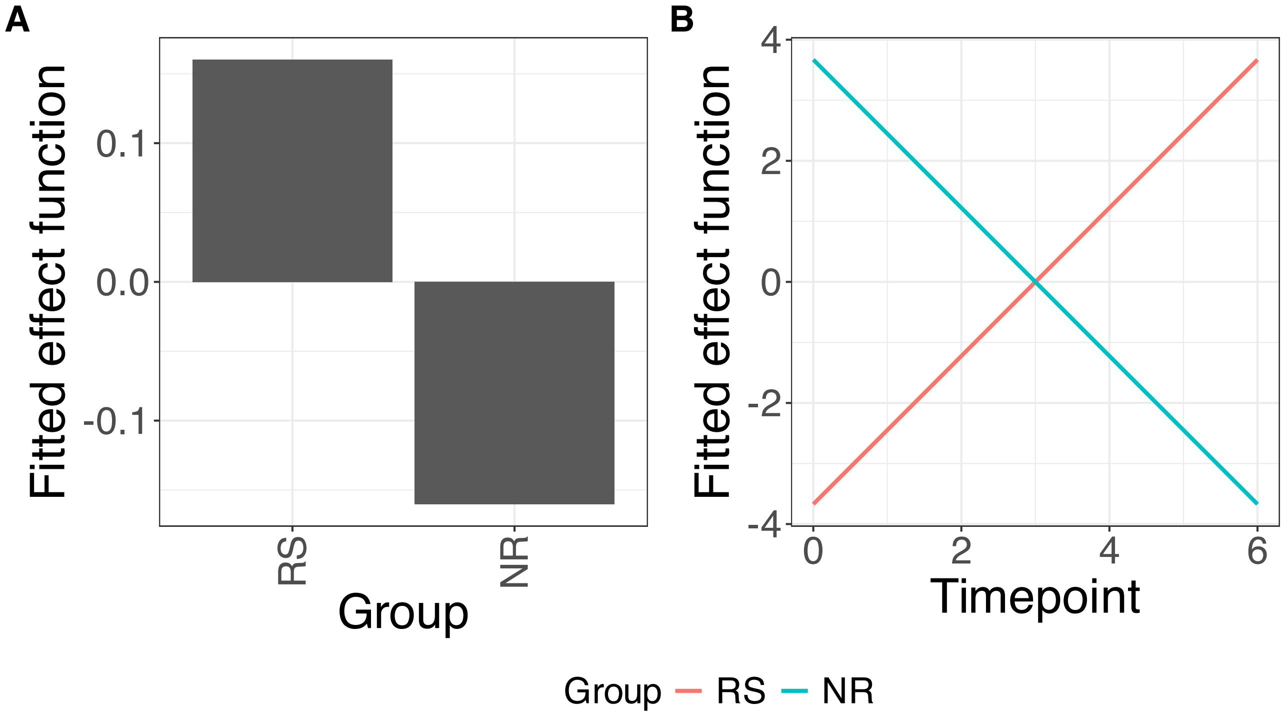


**Supplementary Figure 60.** Decomposition of reconstructed SSANOVA-SVD component function 2. (A) Group. (B) Group:Timepoint.

PCA

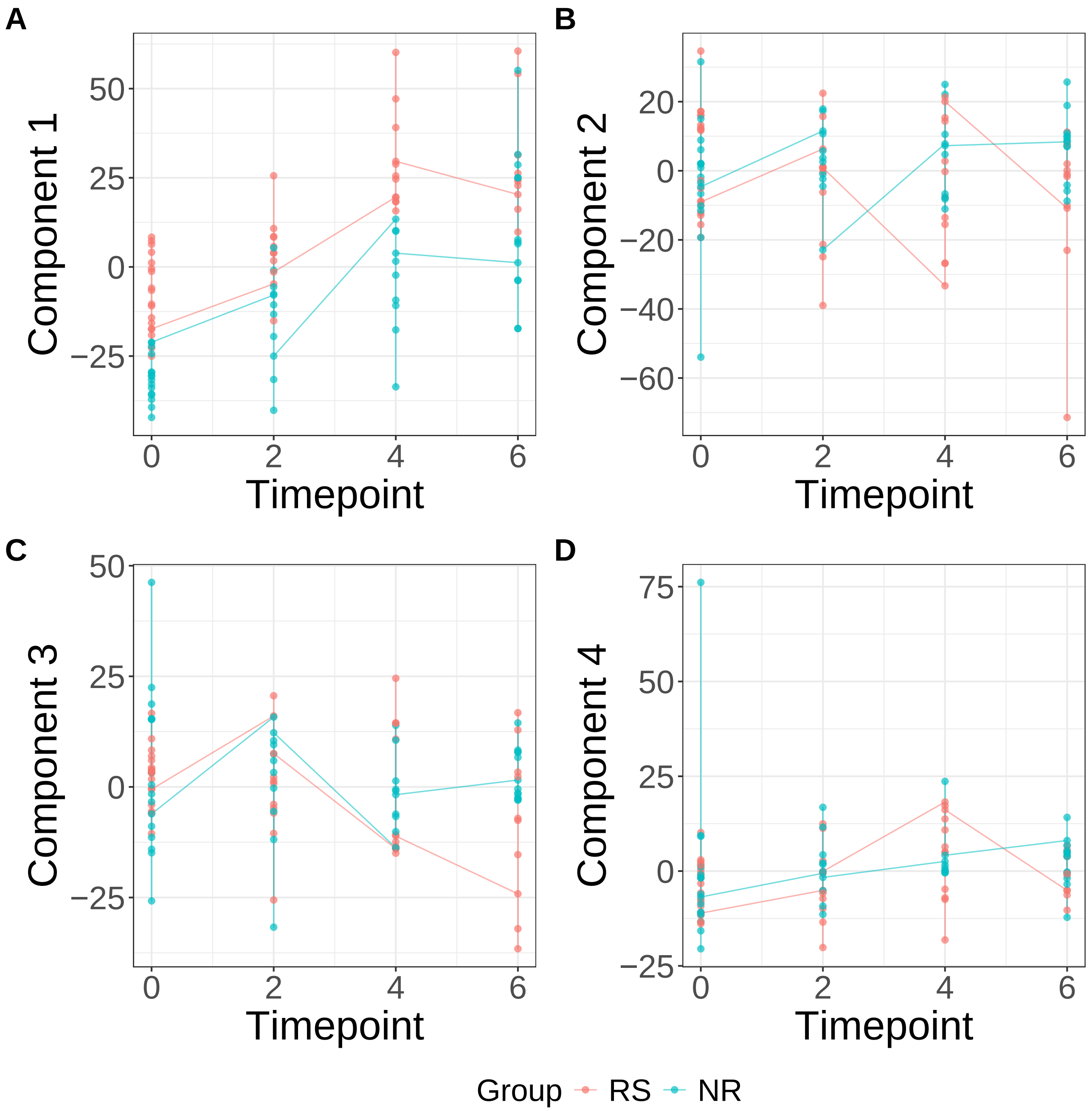


**Supplementary Figure 61.** Components for PCA. (A) Component 1. (B) Component 2. (C) Component 3. (D) Component 4.

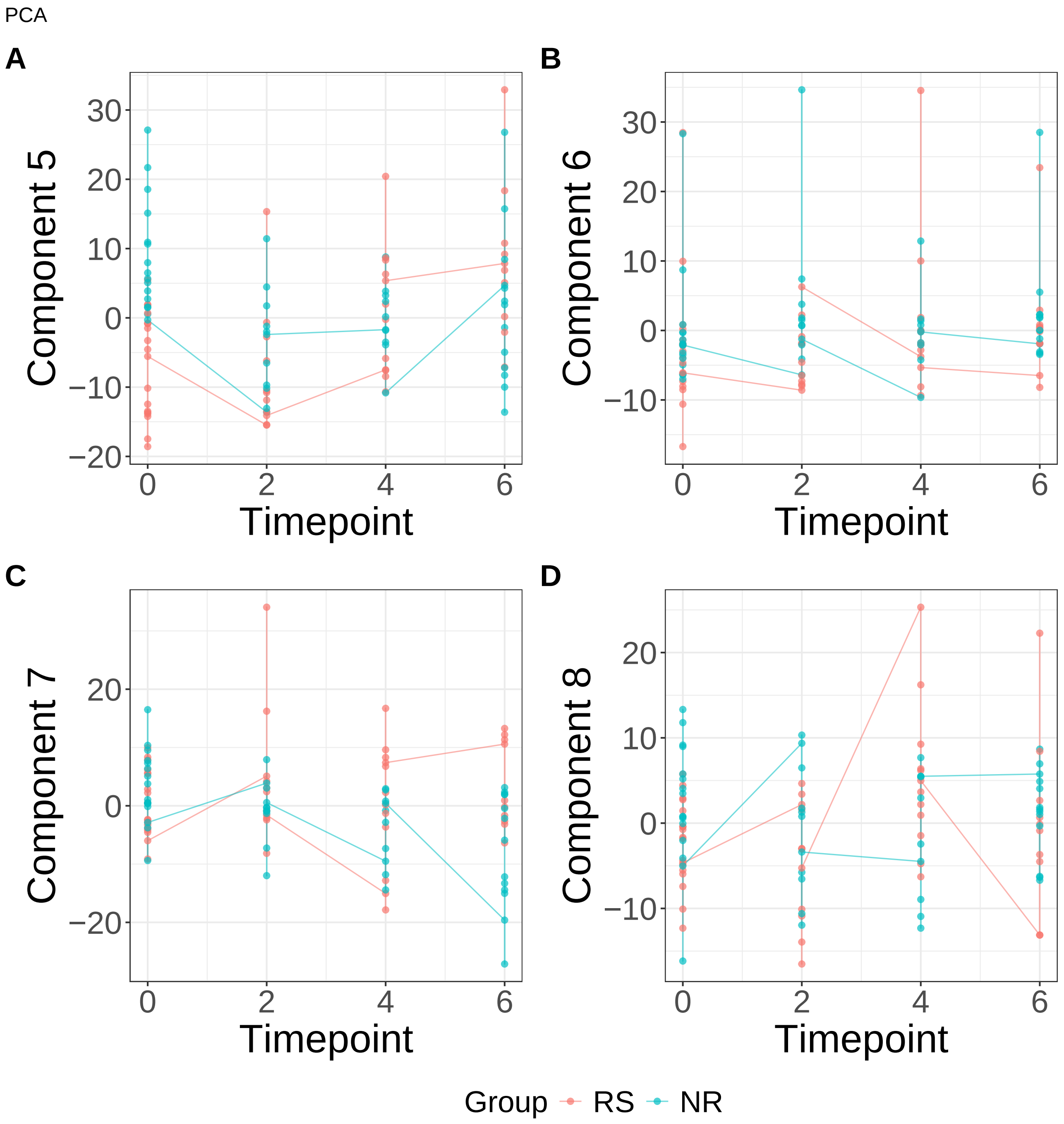


**Supplementary Figure 62.** Components for PCA. (A) Component 5. (B) Component 6. (C) Component 7. (D) Component 8.

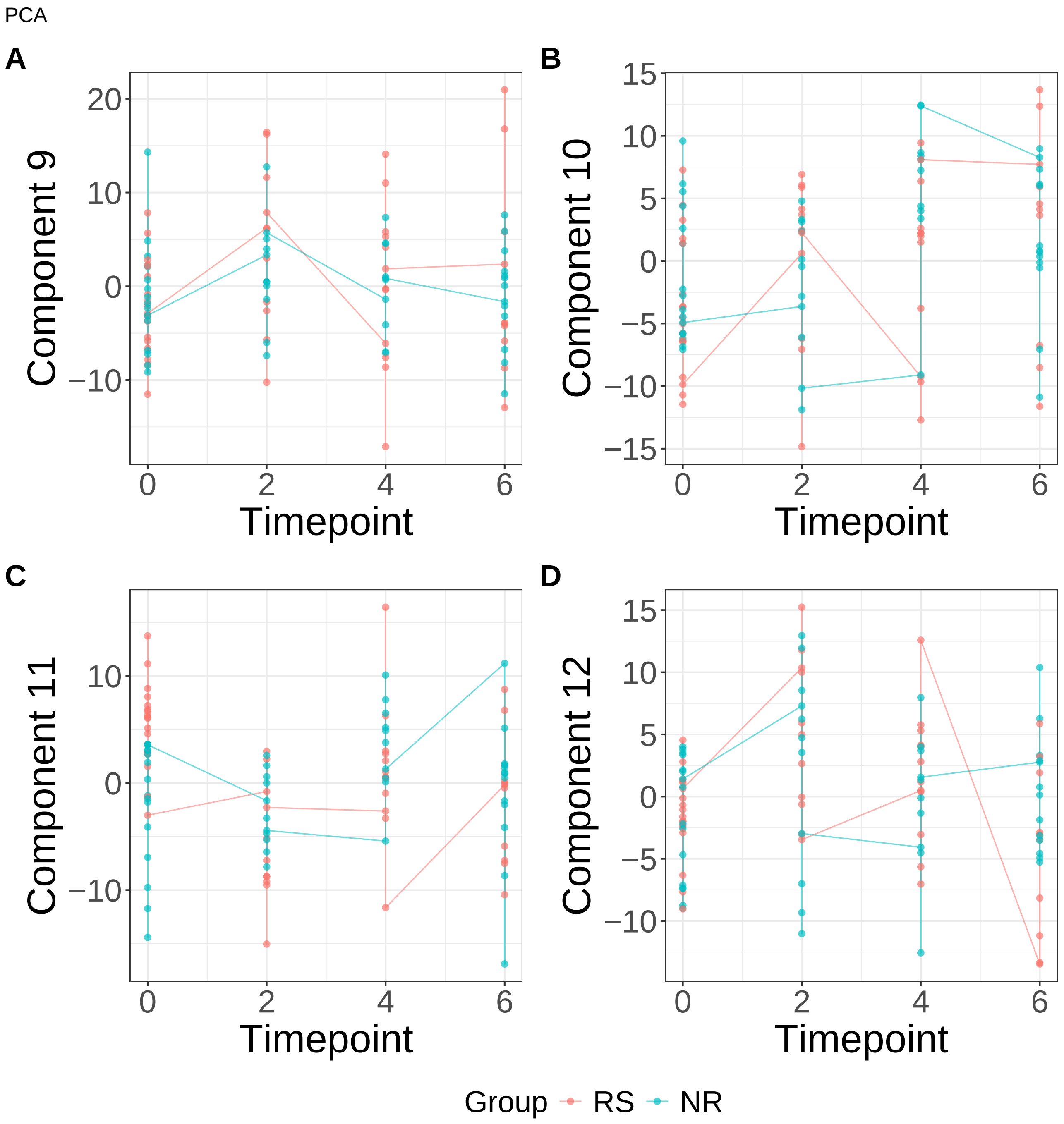


**Supplementary Figure 63.** Components for PCA. (A) Component 9. (B) Component 10. (C) Component 11. (D) Component 12.

PCA

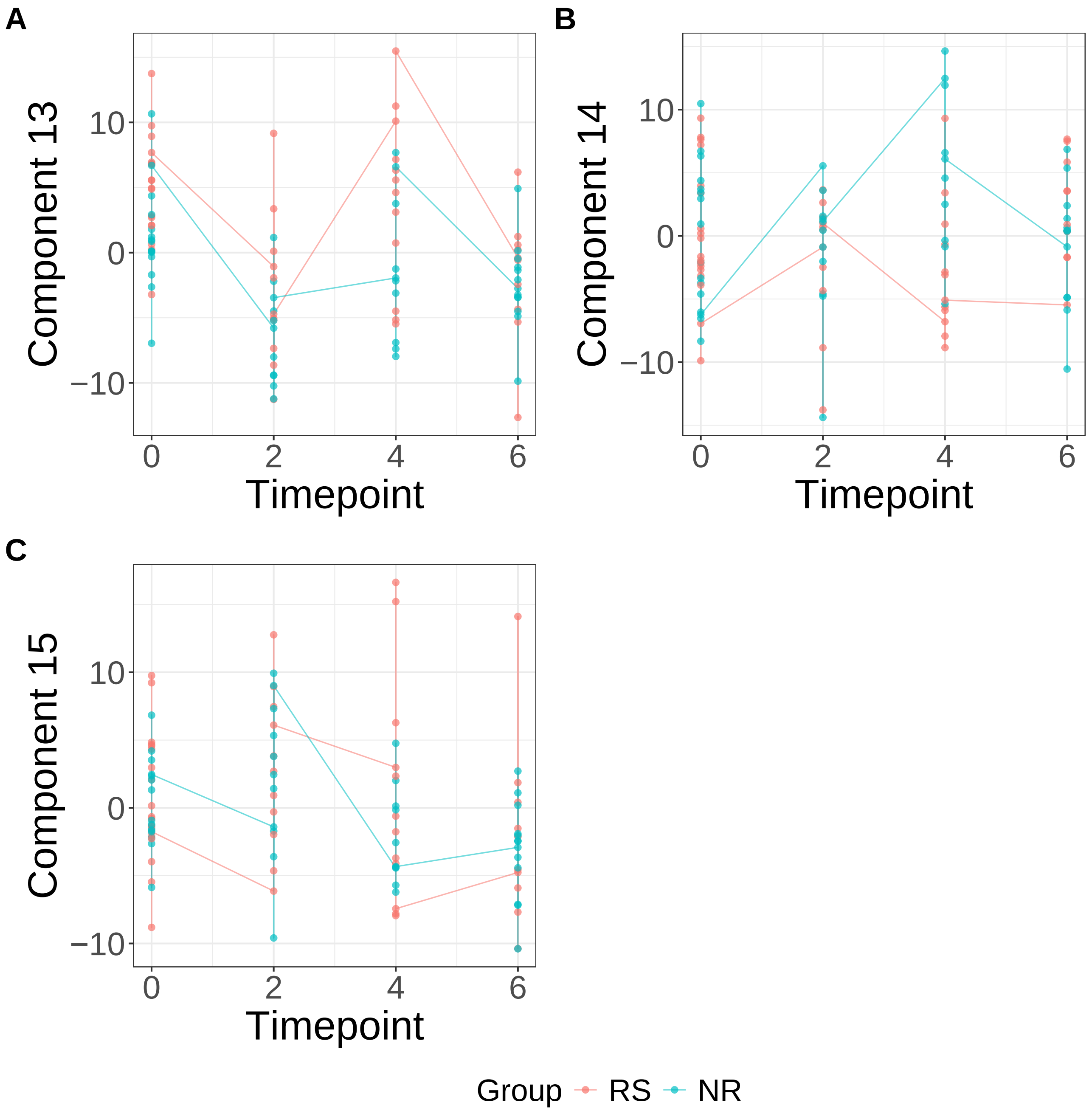


**Supplementary Figure 64.** Components for PCA. (A) Component 13. (B) Component 14. (C) Component 15.

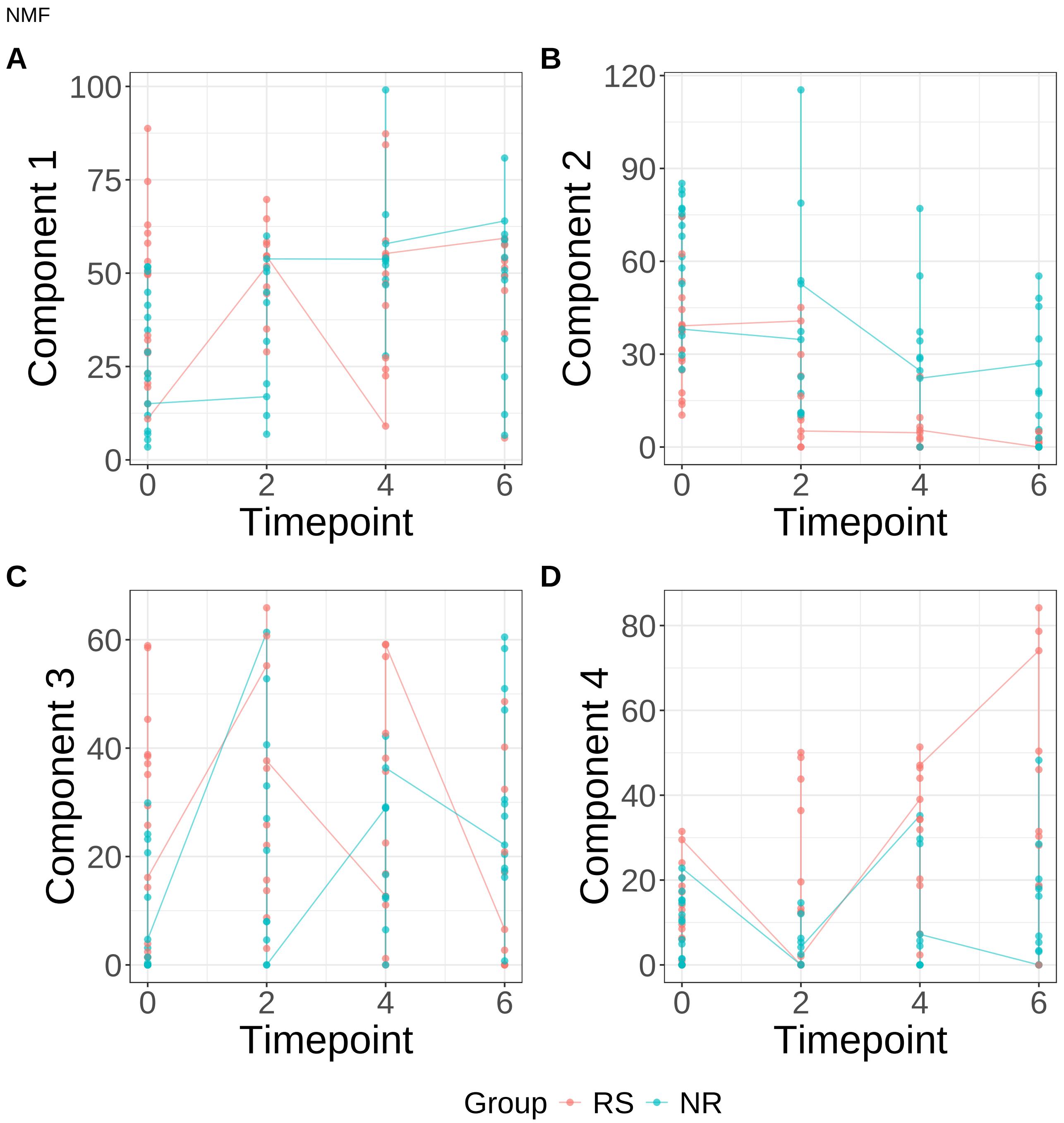


**Supplementary Figure 65.** Components for NMF. (A) Component 1. (B) Component 2. (C) Component 3. (D) Component 4.

NMF

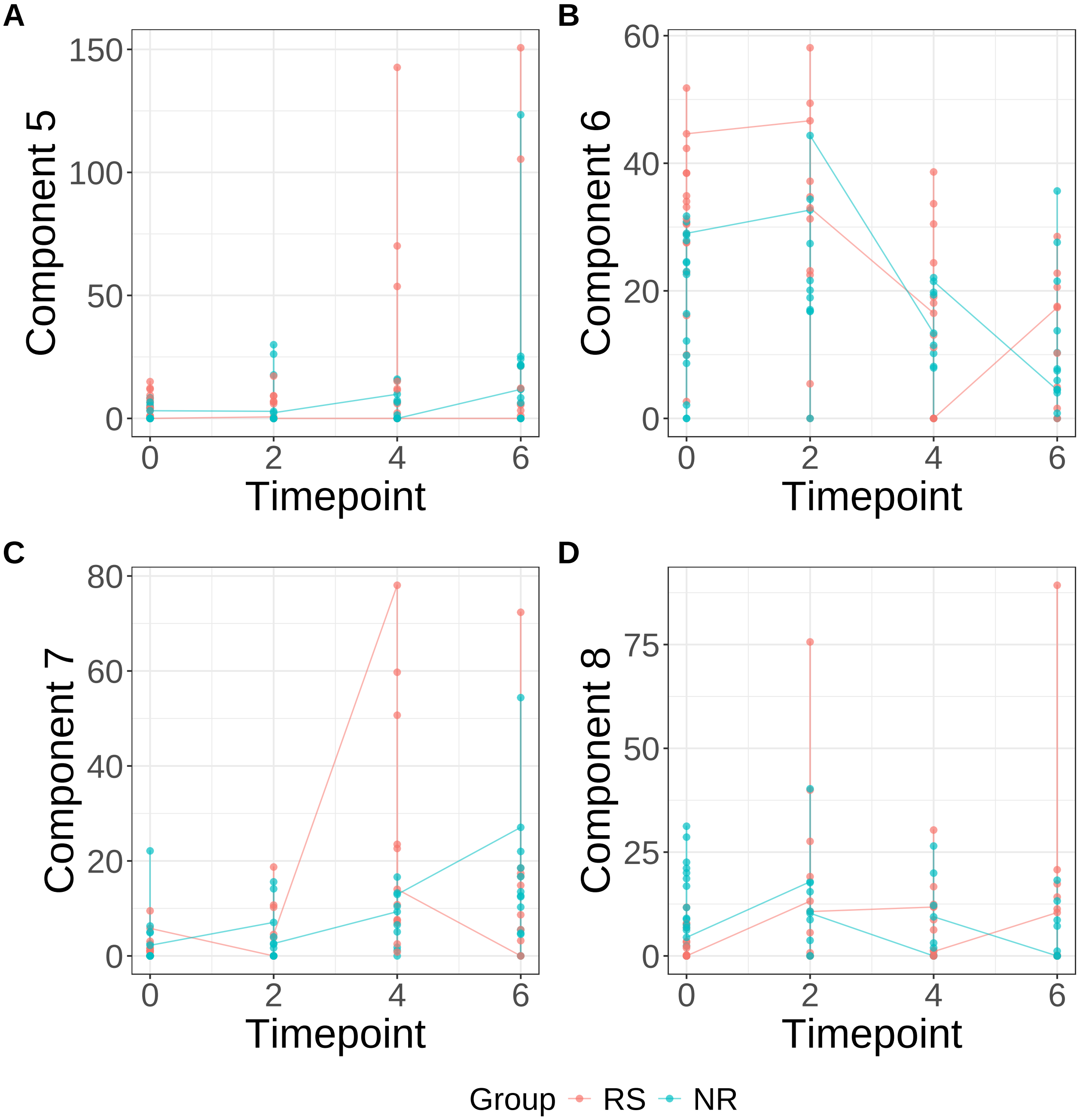


**Supplementary Figure 66.** Components for NMF. (A) Component 5. (B) Component 6. (C) Component 7. (D) Component 8.

NMF

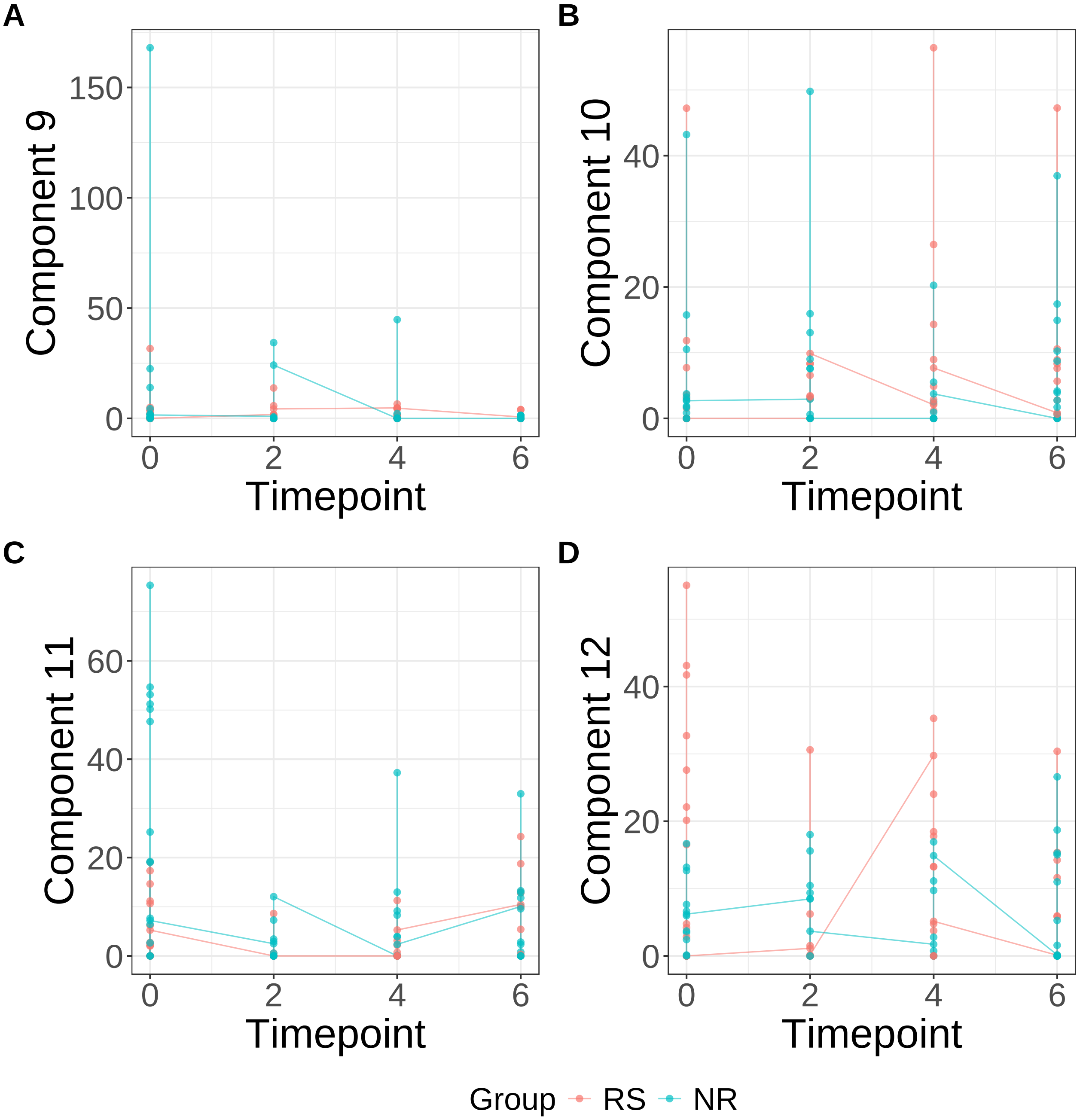


**Supplementary Figure 67.** Components for NMF. (A) Component 9. (B) Component 10. (C) Component 11. (D) Component 12.

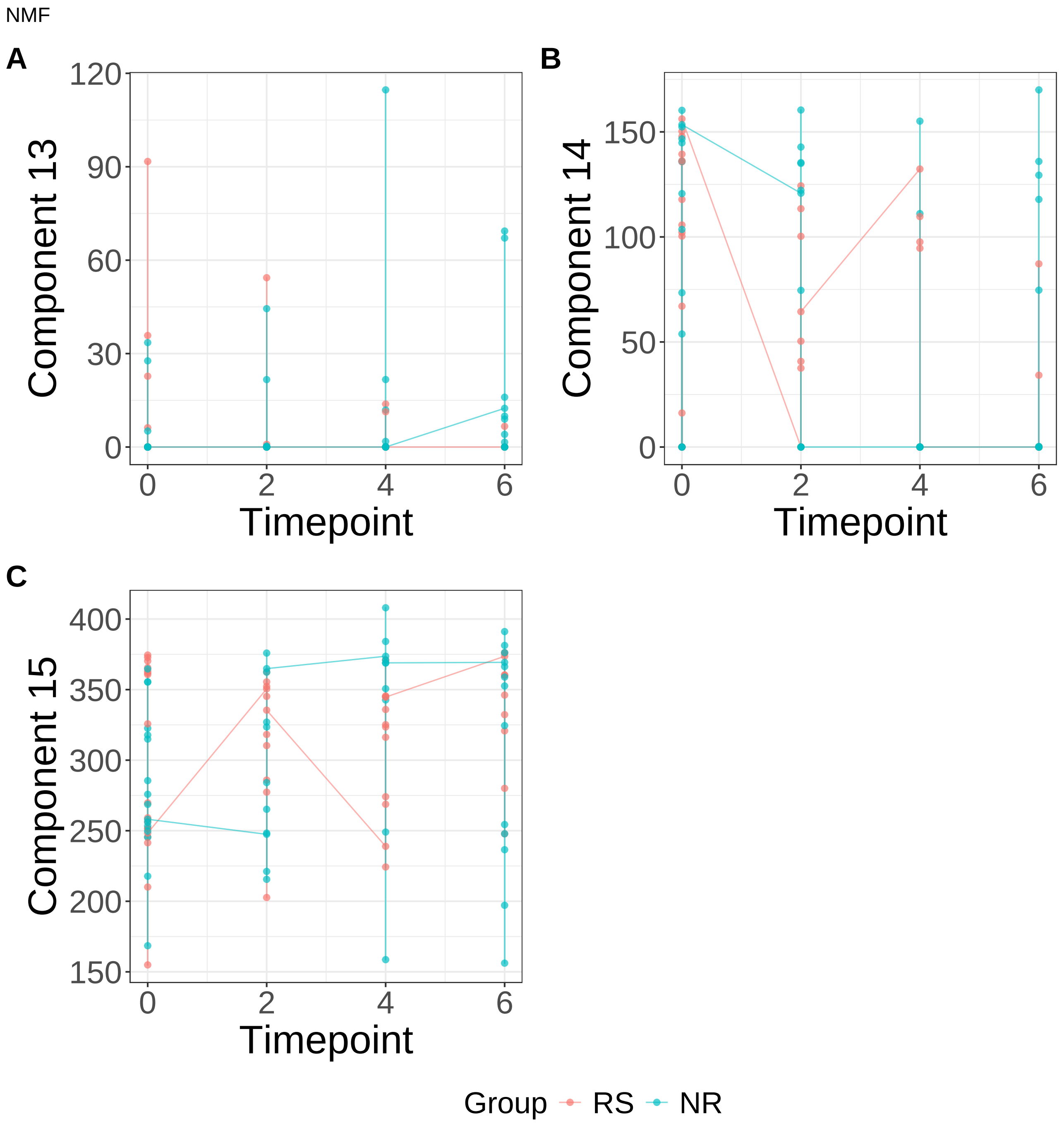


**Supplementary Figure 68.** Components for NMF. (A) Component 13. (B) Component 14. (C) Component 15.

PLS

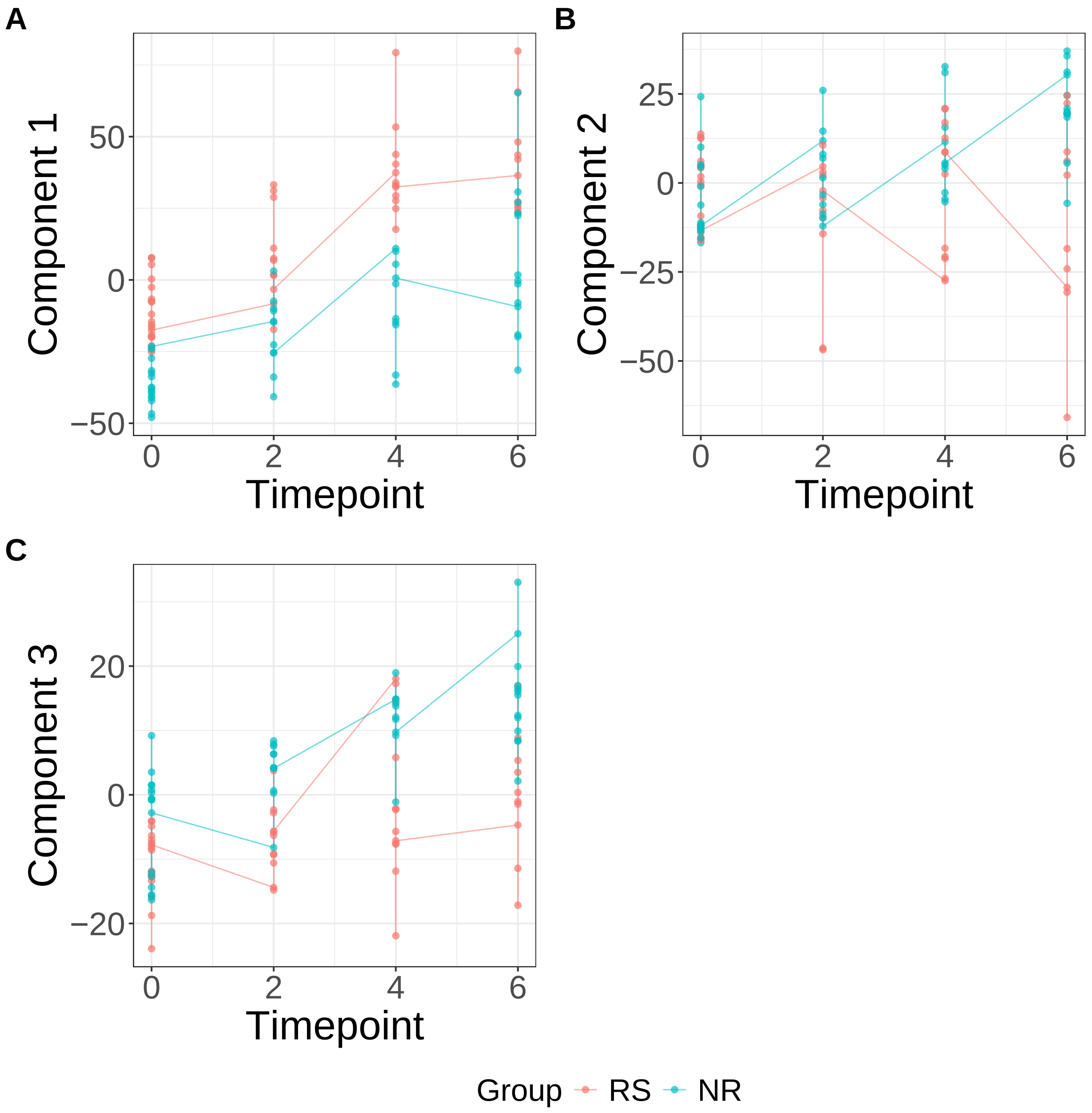


**Supplementary Figure 69.** Components for PLS. (A) Component 1. (B) Component 2. (C) Component 3.

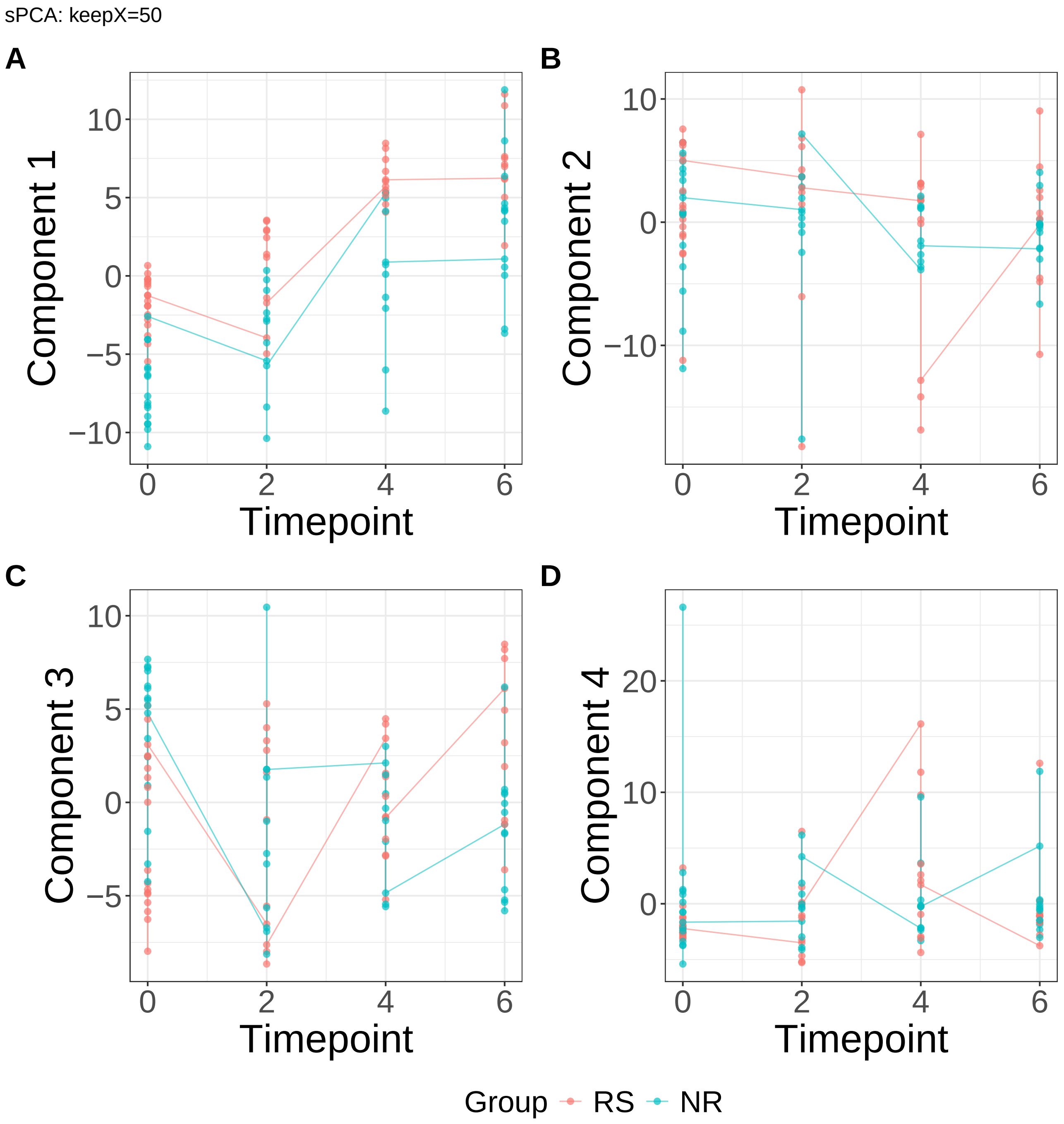


**Supplementary Figure 70.** Components for sPCA: keepX=50. (A) Component 1. (B) Component 2. (C) Component 3. (D) Component 4.

sPCA: keepX=50

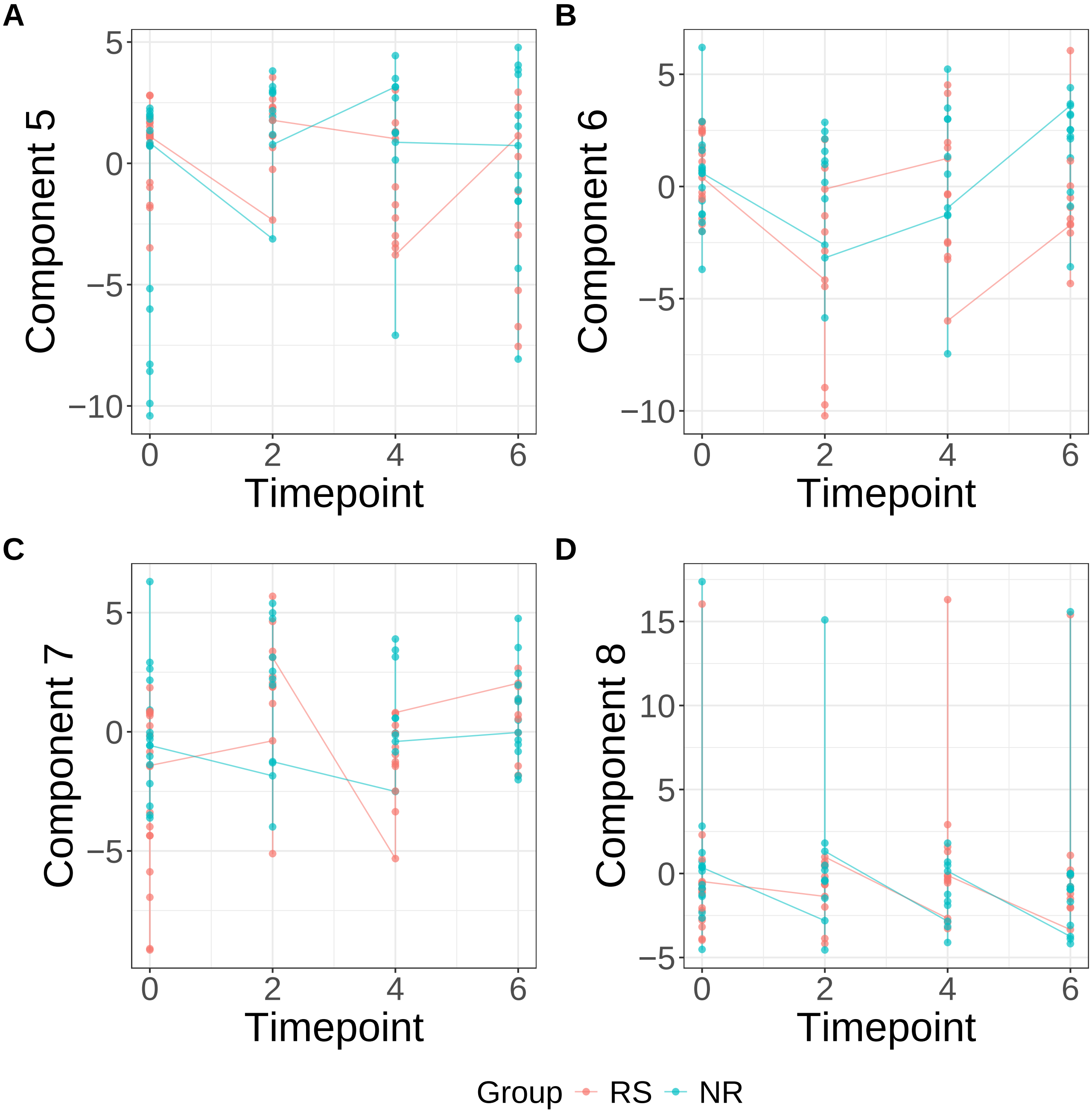


**Supplementary Figure 71.** Components for sPCA: keepX=50. (A) Component 5. (B) Component 6. (C) Component 7. (D) Component 8.

sPCA: keepX=50

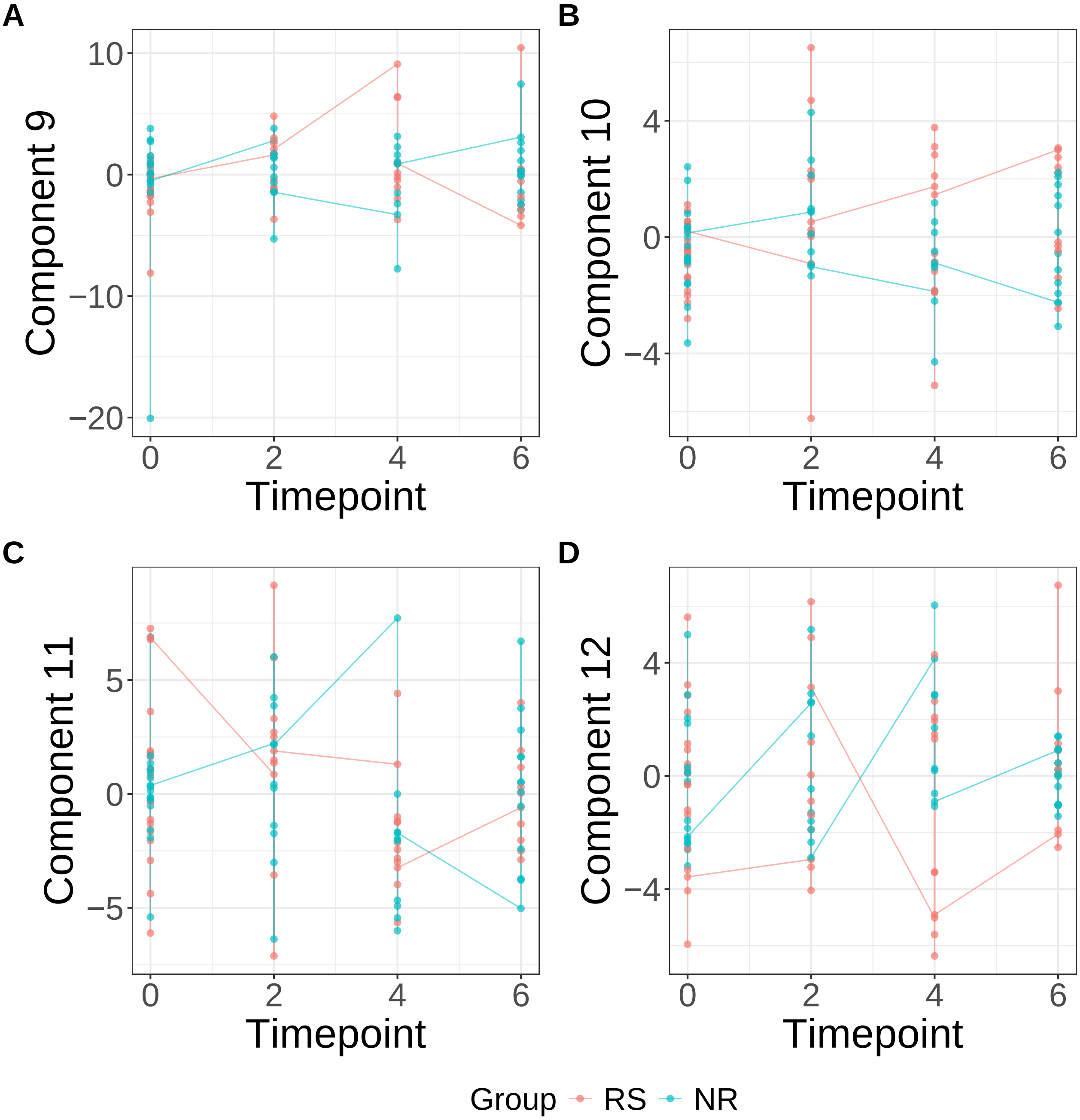


**Supplementary Figure 72.** Components for sPCA: keepX=50. (A) Component 9. (B) Component 10. (C) Component 11. (D) Component 12.

sPCA: keepX=50

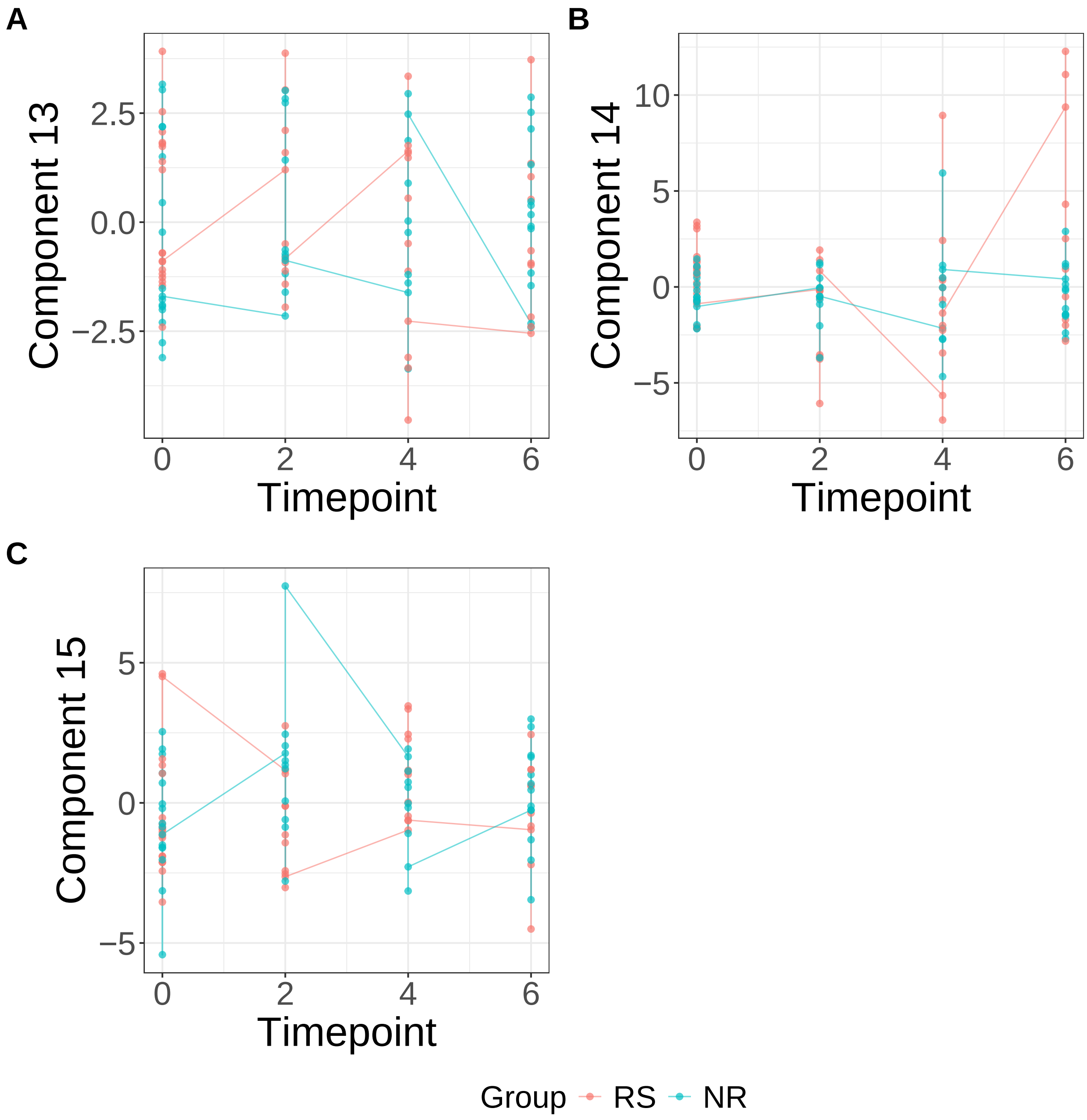


**Supplementary Figure 73.** Components for sPCA: keepX=50. (A) Component 13. (B) Component 14. (C) Component 15.

sPCA: keepX=1250

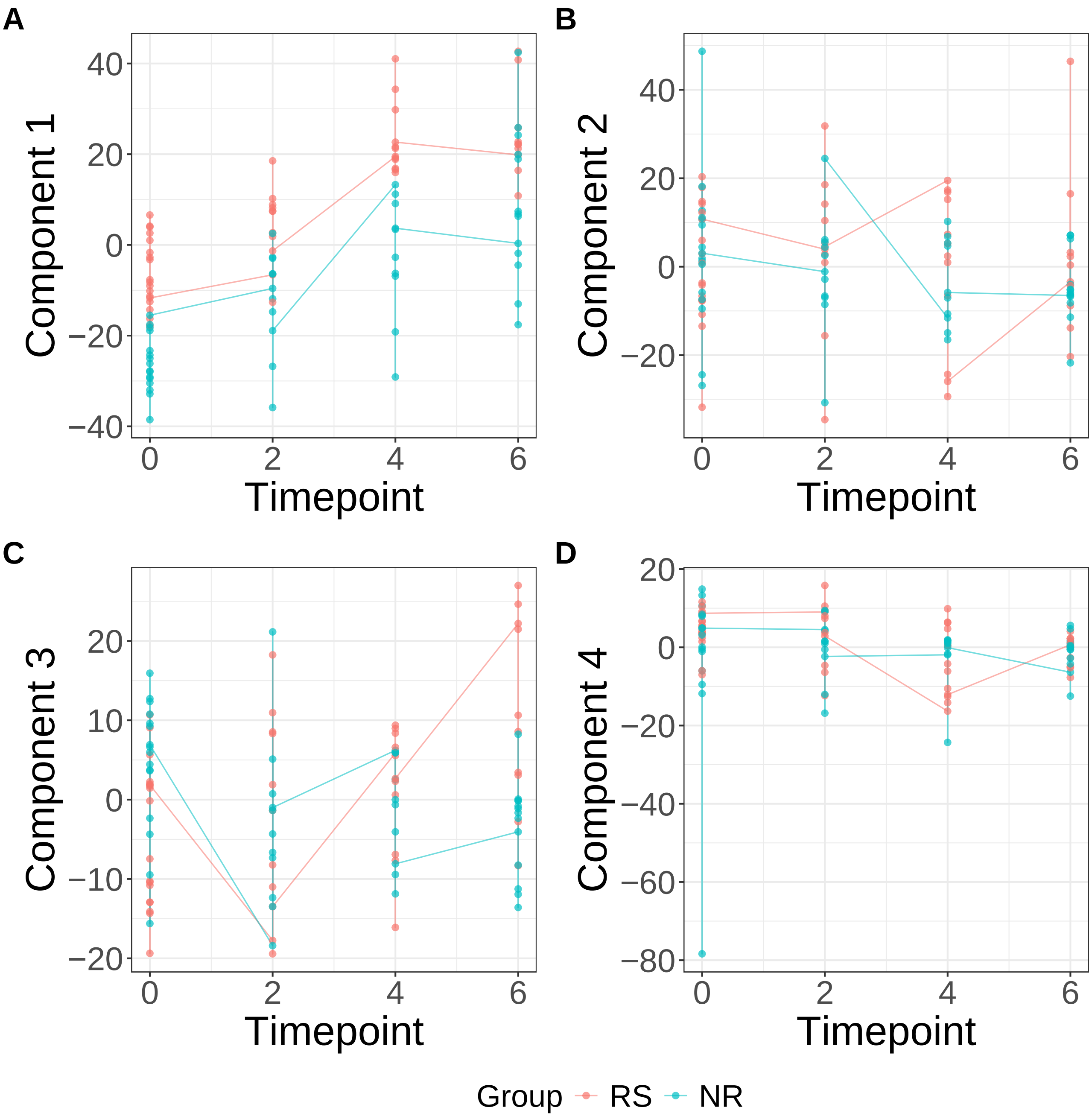


**Supplementary Figure 74.** Components for sPCA: keepX=1250. (A) Component 1. (B) Component 2. (C) Component 3. (D) Component 4.

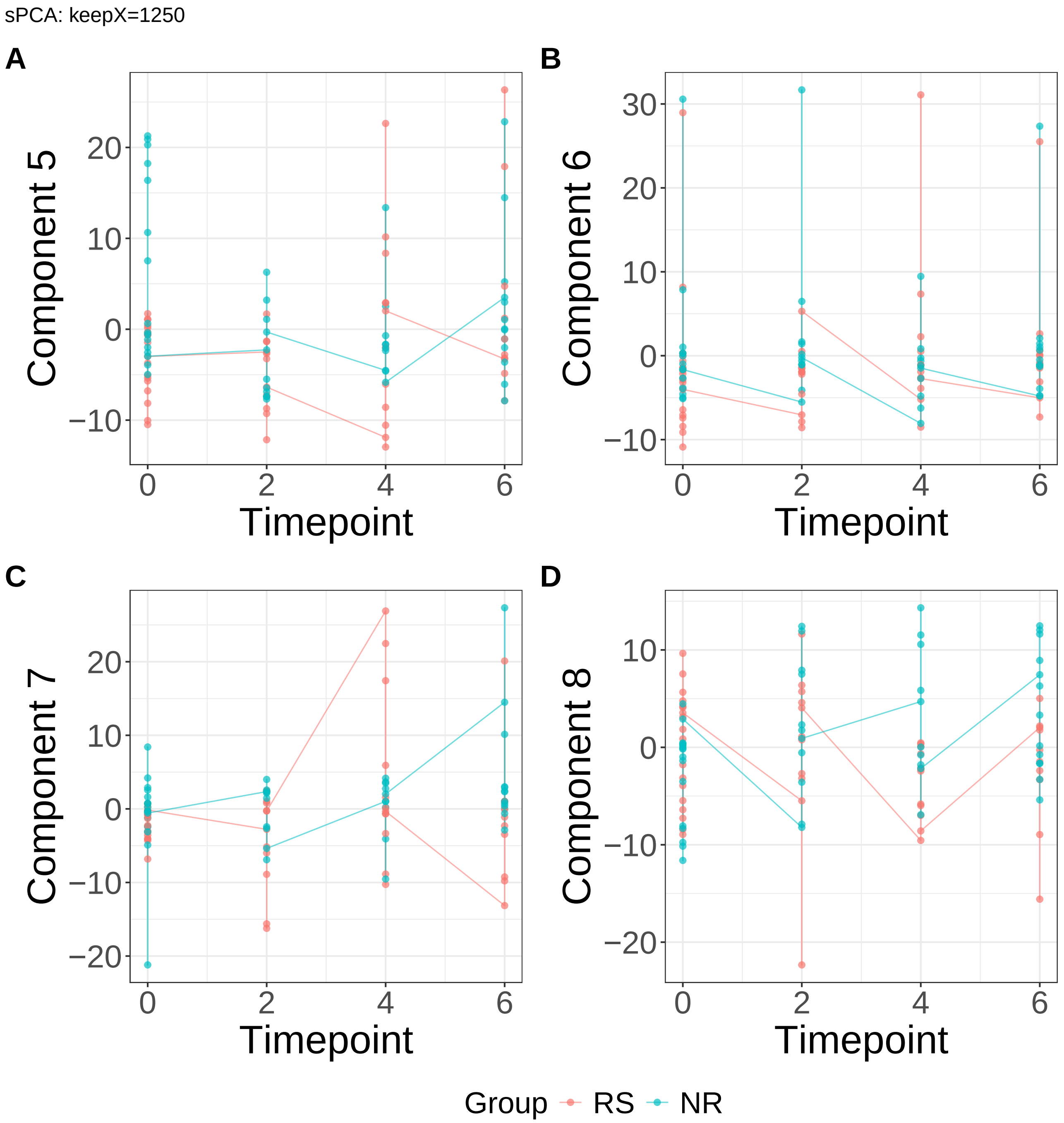


**Supplementary Figure 75.** Components for sPCA: keepX=1250. (A) Component 5. (B) Component 6. (C) Component 7. (D) Component 8.

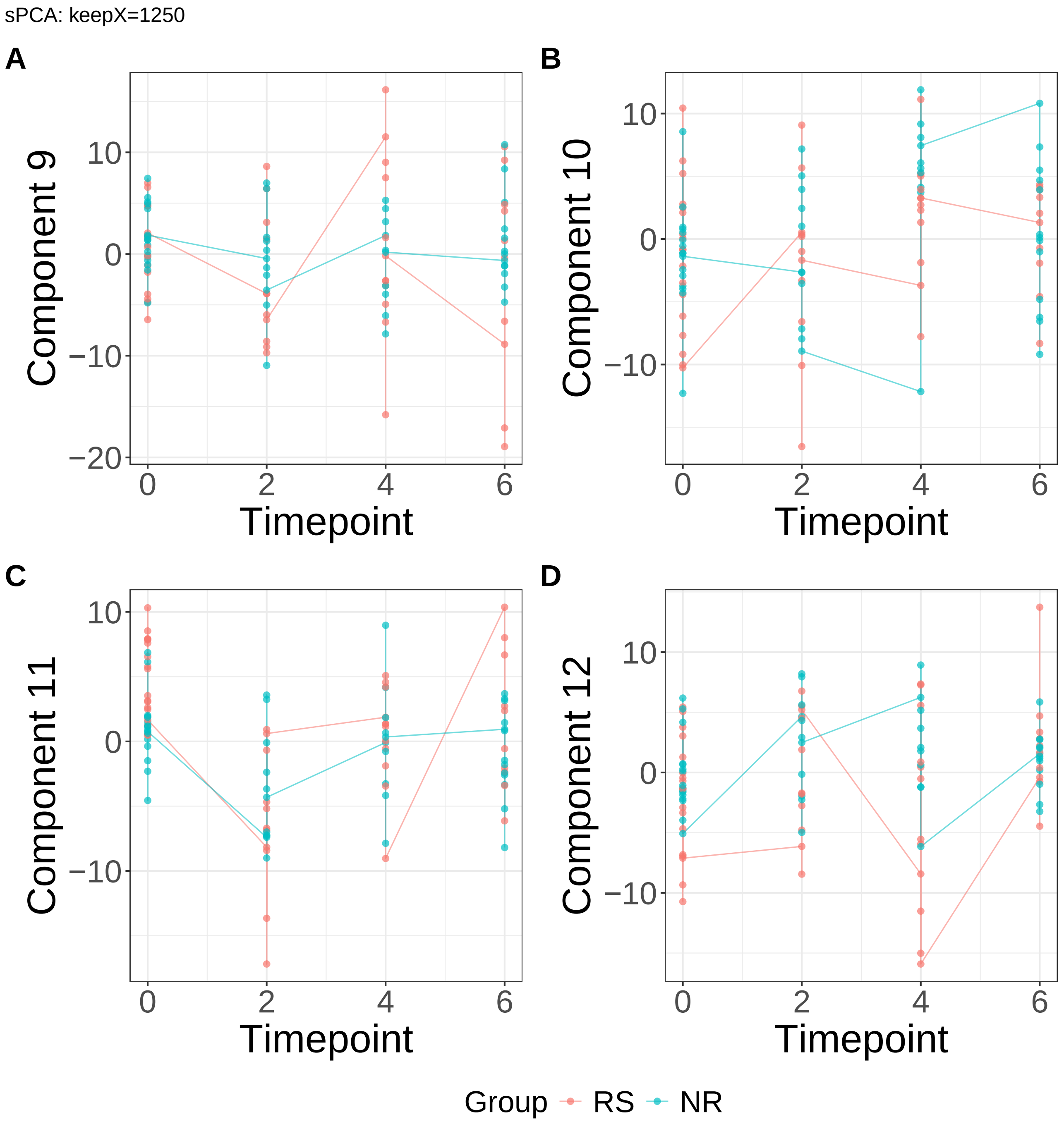


**Supplementary Figure 76.** Components for sPCA: keepX=1250. (A) Component 9. (B) Component 10. (C) Component 11. (D) Component 12.

sPCA: keepX=1250

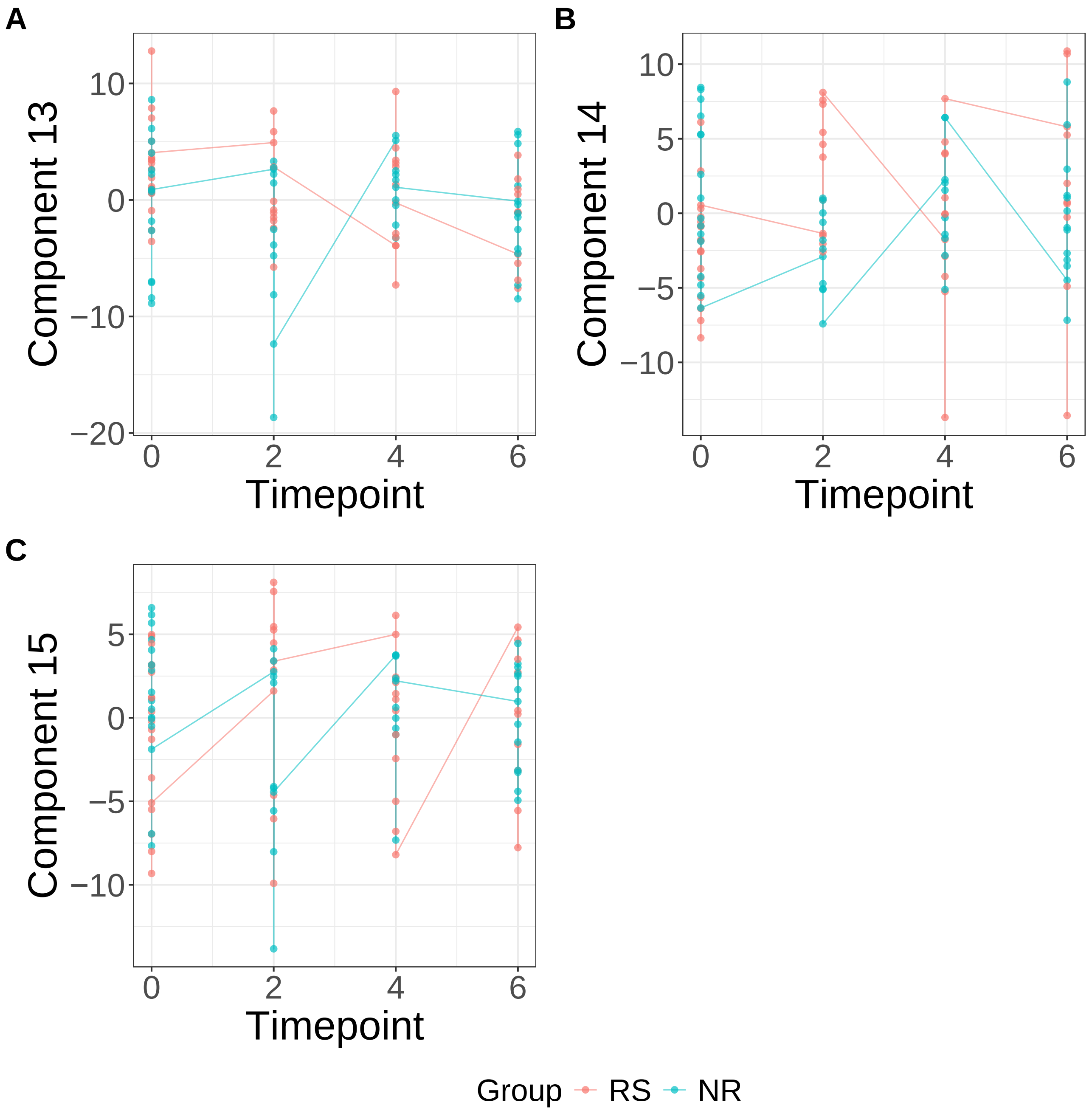


**Supplementary Figure 77.** Components for sPCA: keepX=1250. (A) Component 13. (B) Component 14. (C) Component 15.

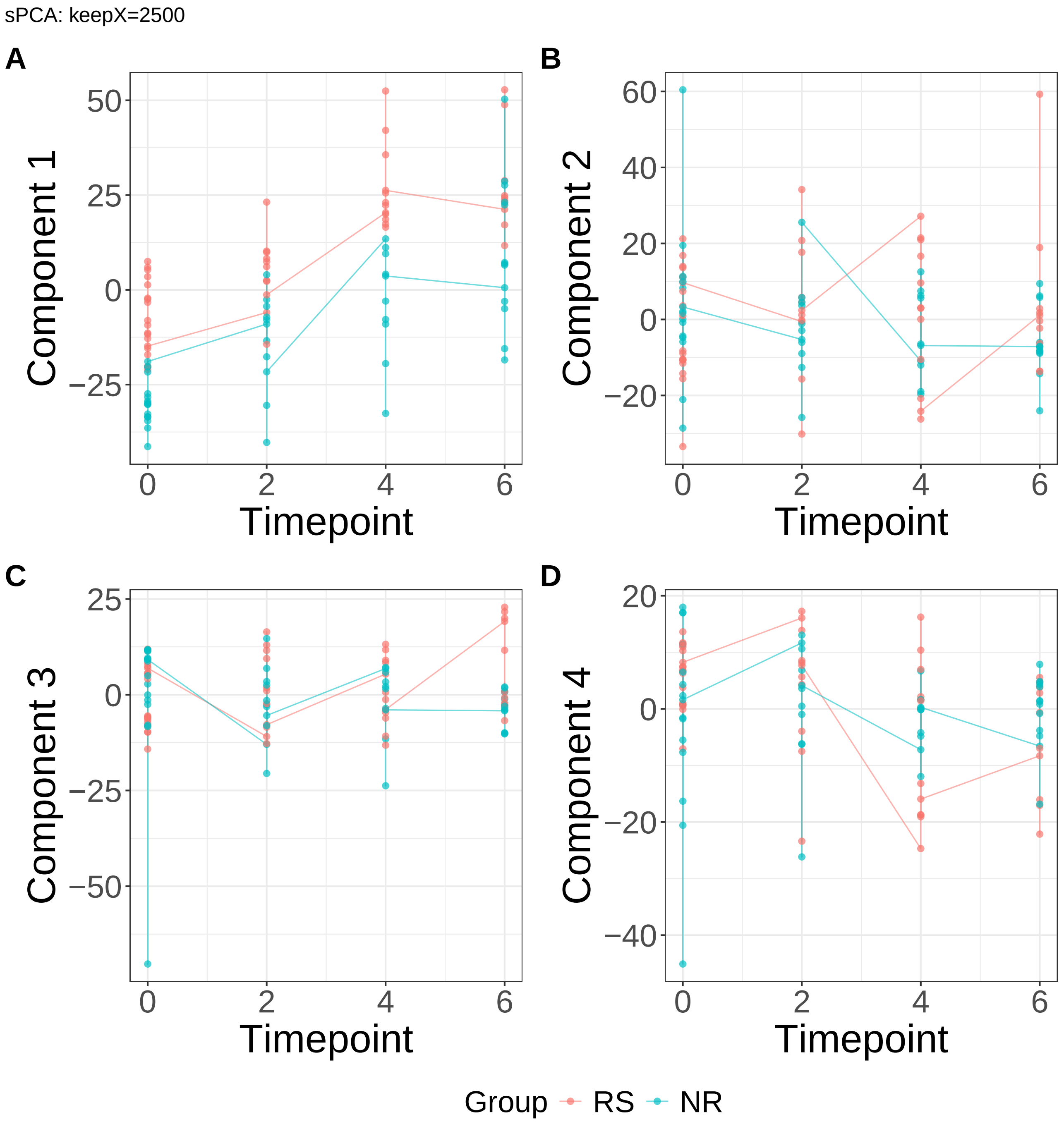


**Supplementary Figure 78.** Components for sPCA: keepX=2500. (A) Component 1. (B) Component 2. (C) Component 3. (D) Component 4.

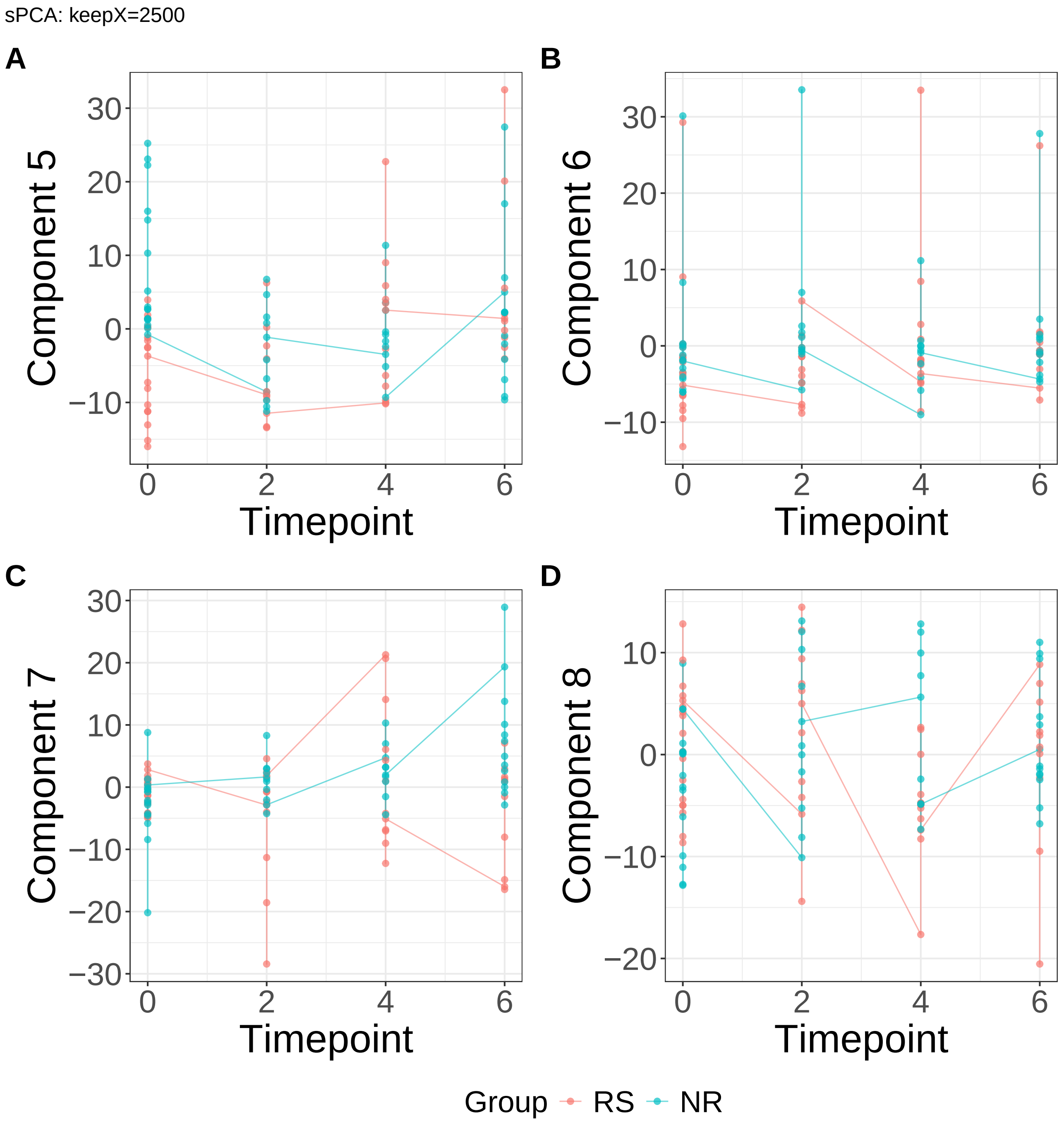


**Supplementary Figure 79.** Components for sPCA: keepX=2500. (A) Component 5. (B) Component 6. (C) Component 7. (D) Component 8.

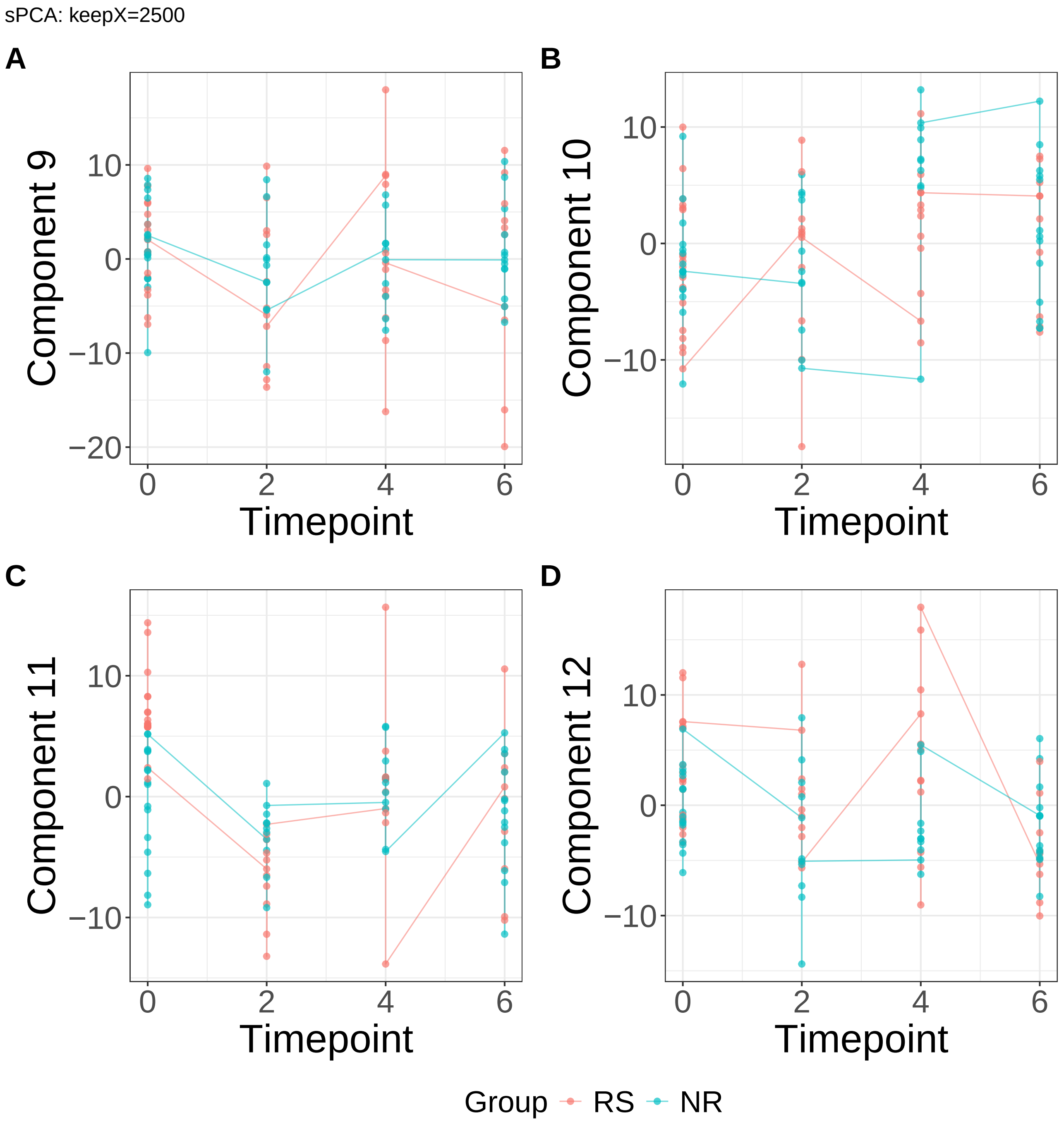


**Supplementary Figure 80.** Components for sPCA: keepX=2500. (A) Component 9. (B) Component 10. (C) Component 11. (D) Component 12.

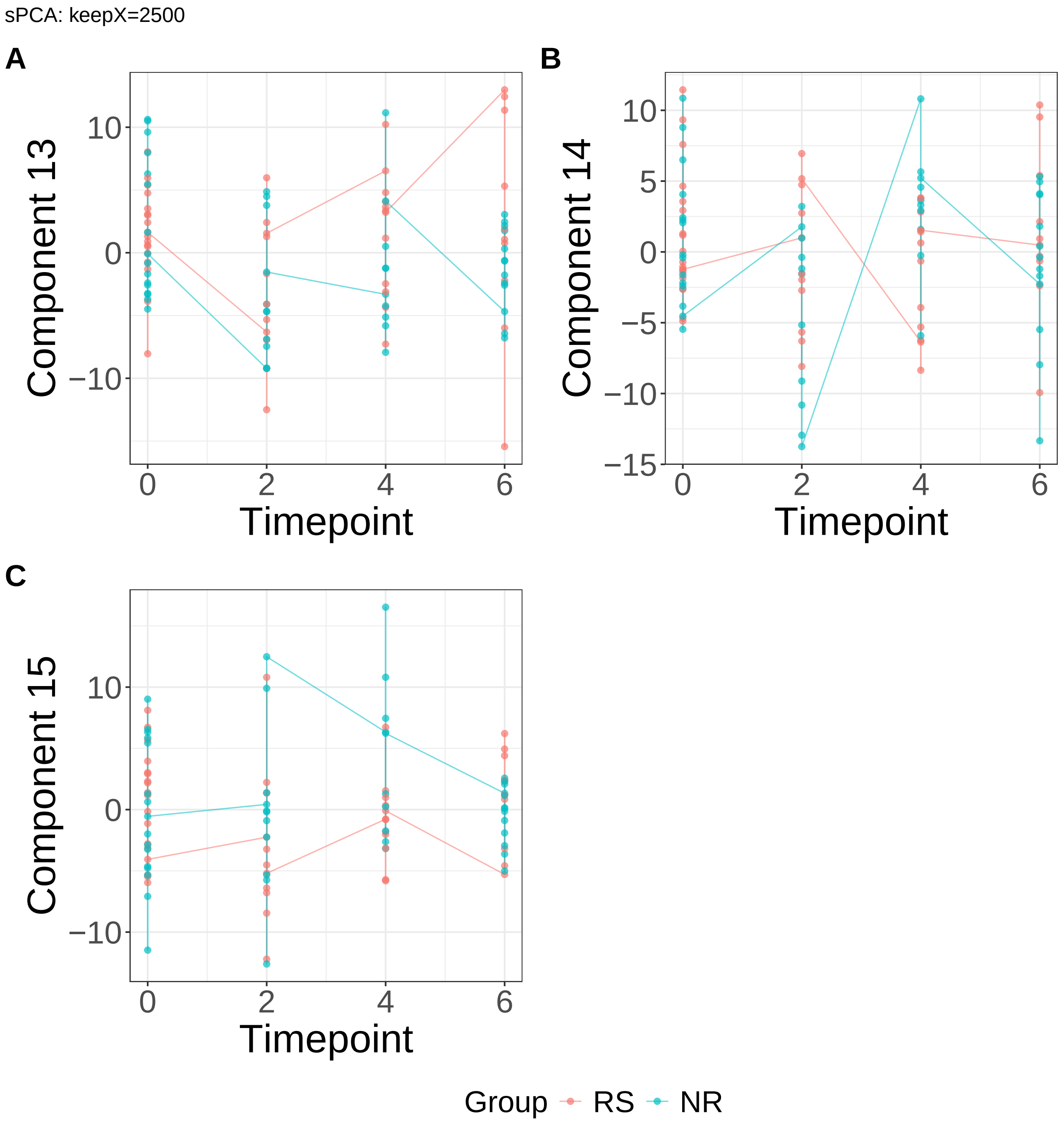


**Supplementary Figure 81.** Components for sPCA: keepX=2500. (A) Component 13. (B) Component 14. (C) Component 15.

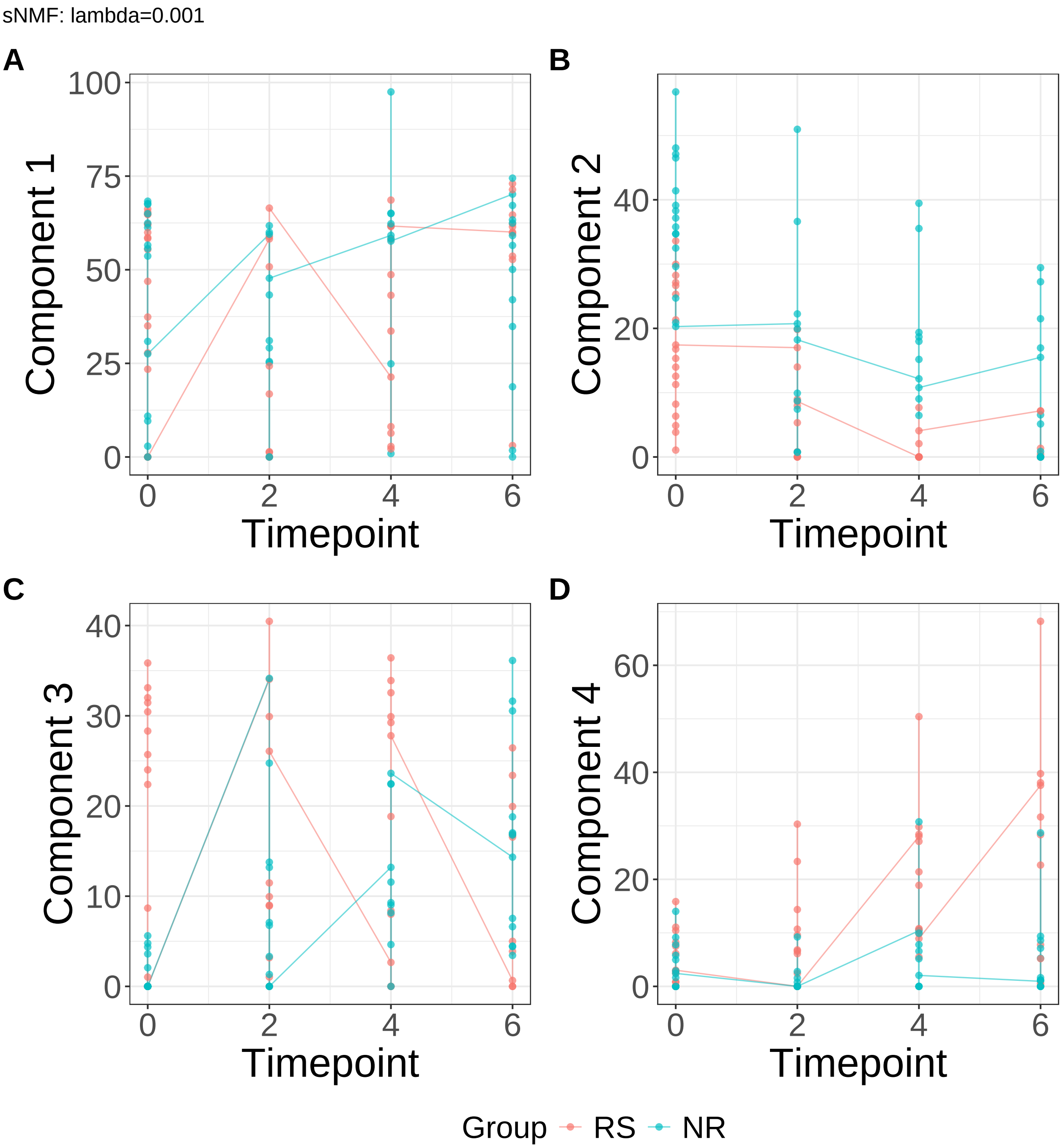


**Supplementary Figure 82.** Components for sNMF: lambda=0.001. (A) Component 1. (B) Component 2. (C) Component 3. (D) Component 4.

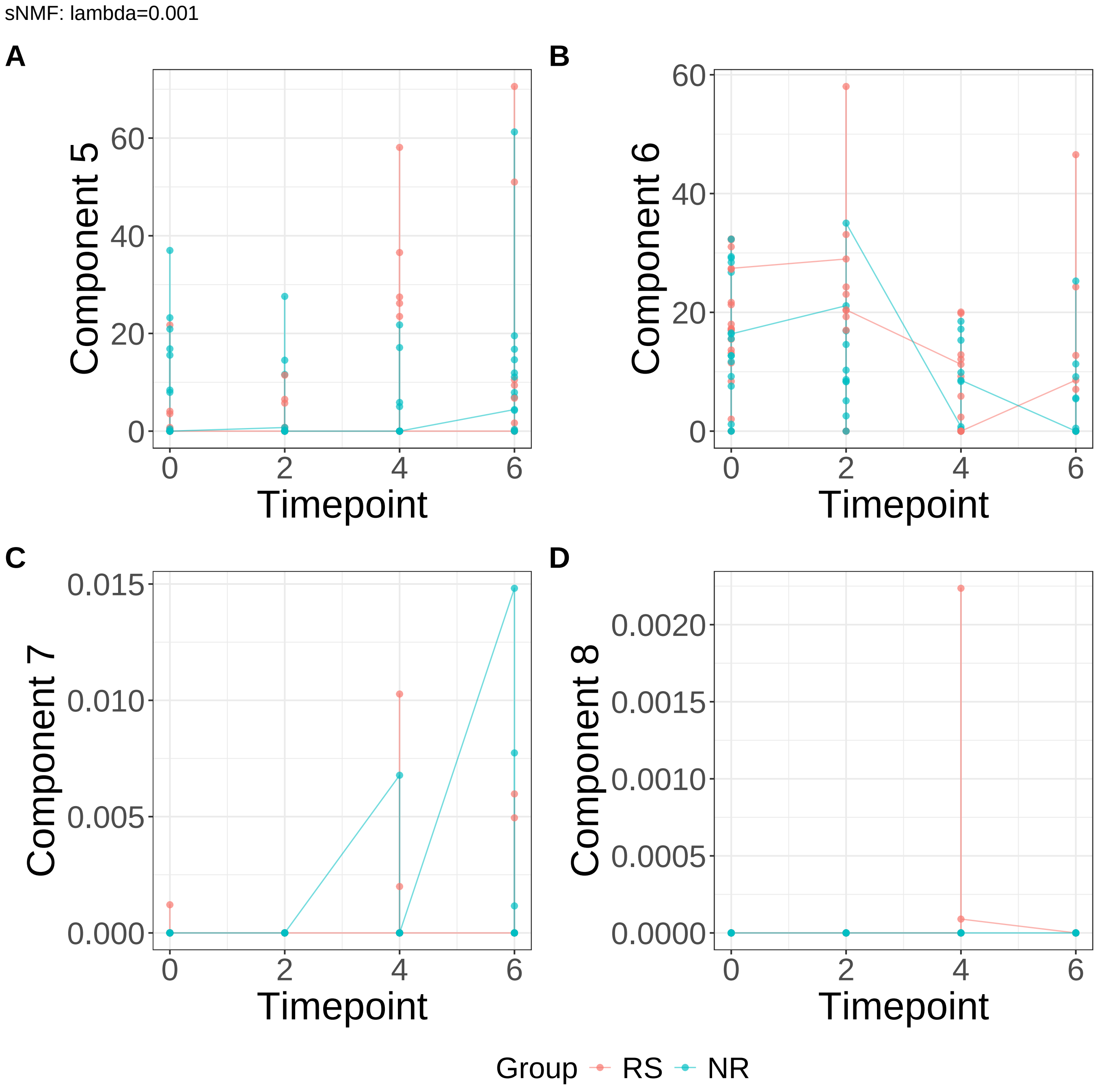


**Supplementary Figure 83.** Components for sNMF: lambda=0.001. (A) Component 5. (B) Component 6. (C) Component 7. (D) Component 8.

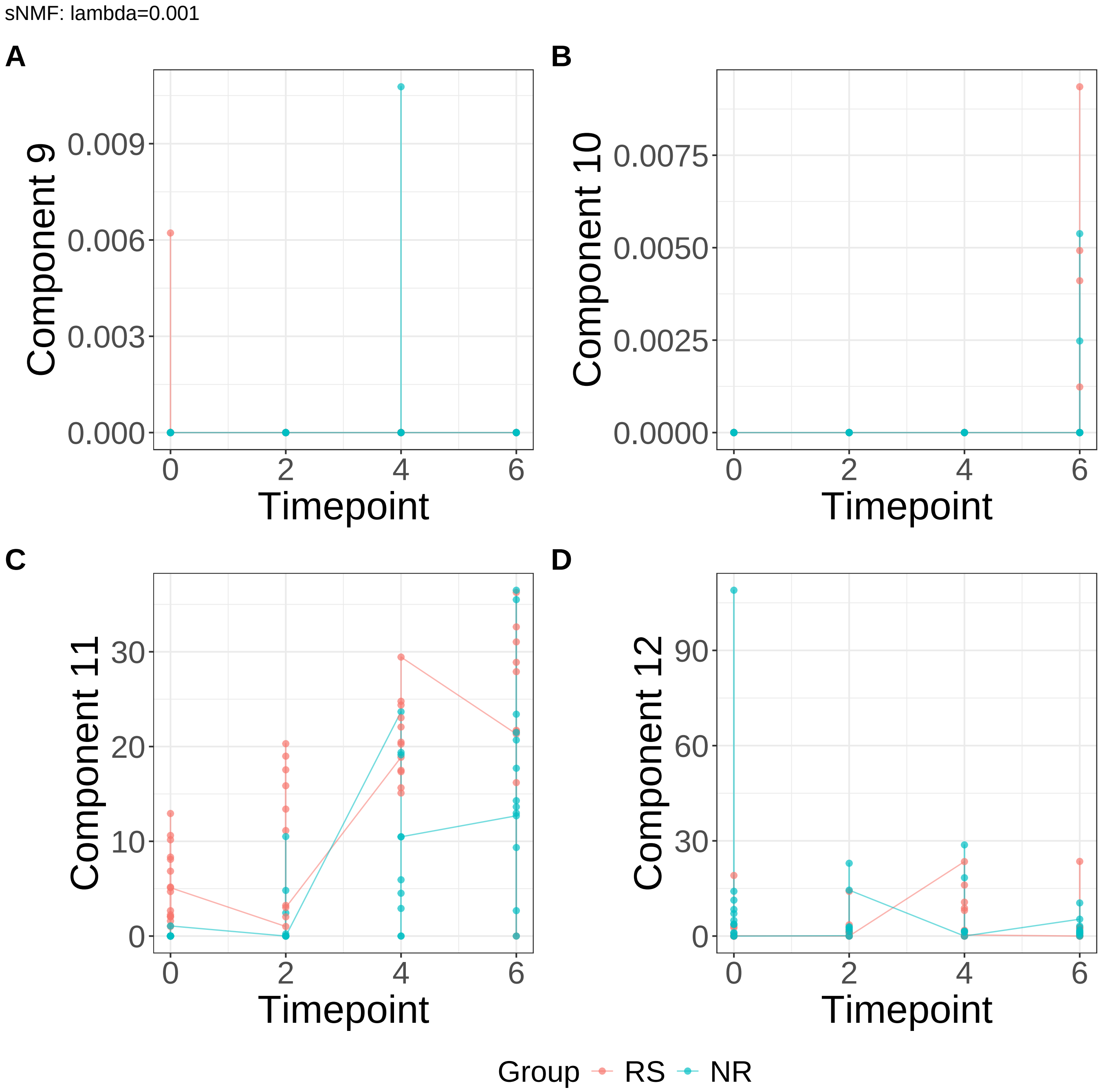


**Supplementary Figure 84.** Components for sNMF: lambda=0.001. (A) Component 9. (B) Component 10. (C) Component 11. (D) Component 12.

sNMF: lambda=0.001

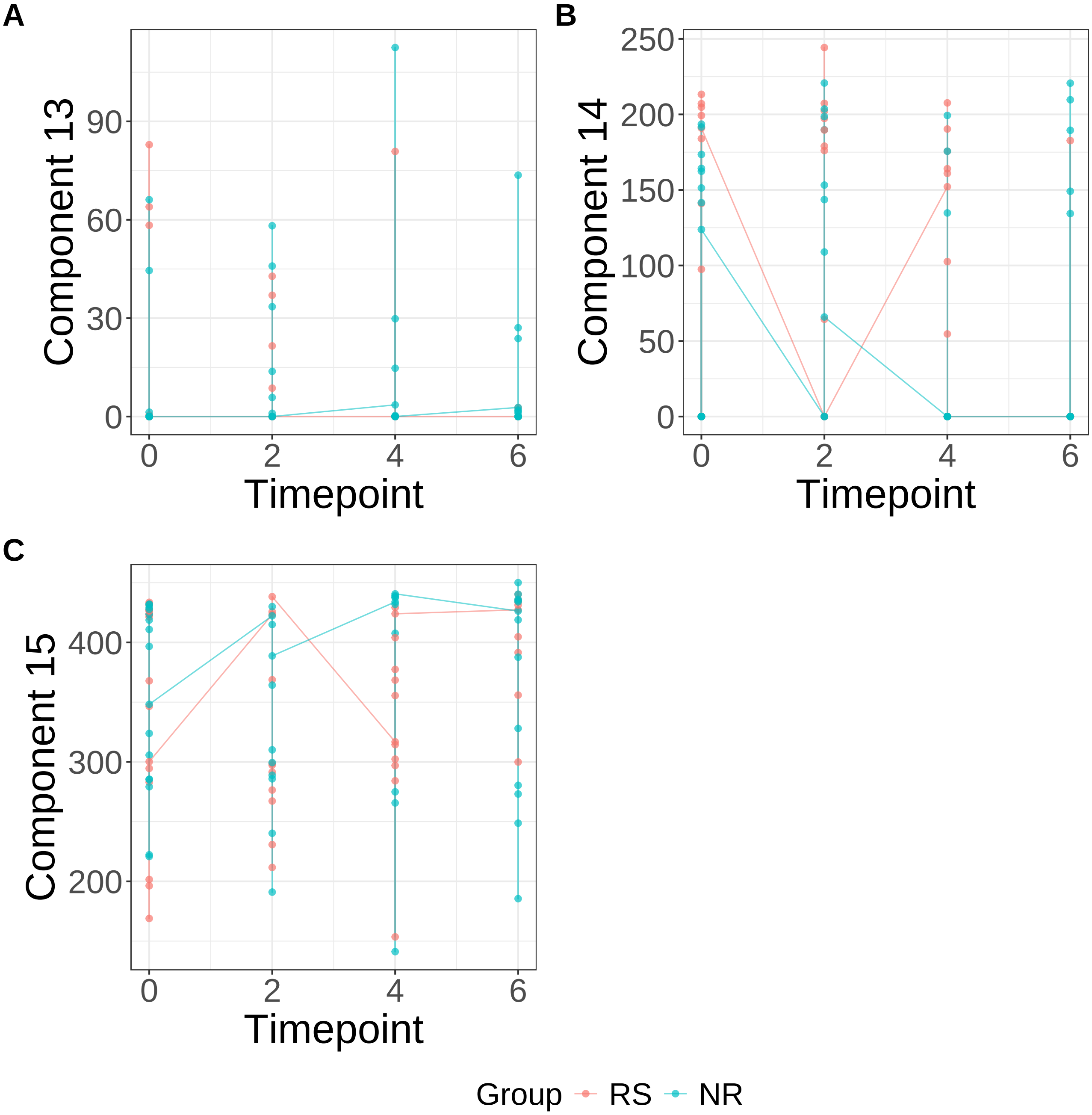


**Supplementary Figure 85.** Components for sNMF: lambda=0.001. (A) Component 13. (B) Component 14. (C) Component 15.

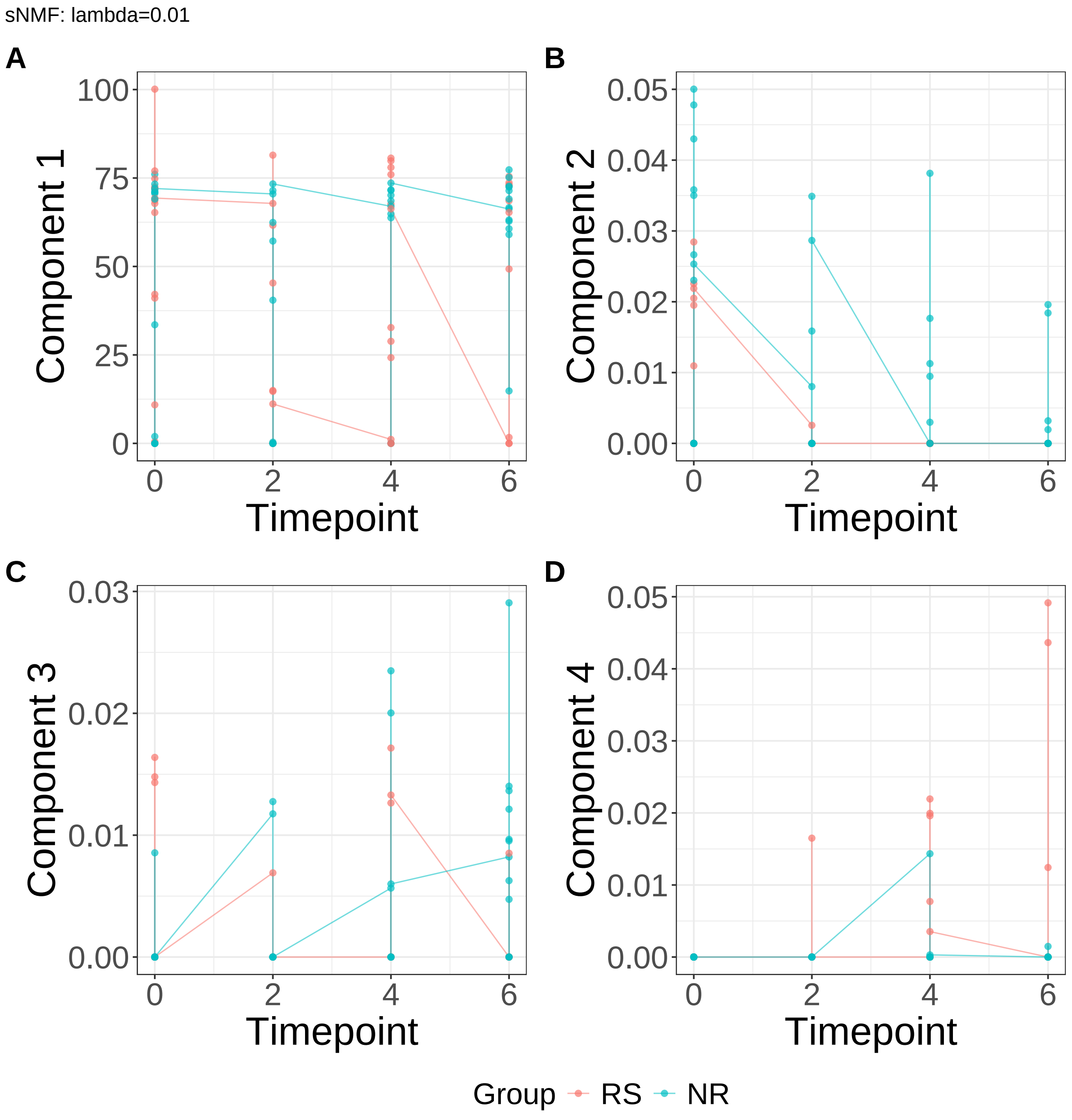


**Supplementary Figure 86.** Components for sNMF: lambda=0.01. (A) Component 1. (B) Component 2. (C) Component 3. (D) Component 4.

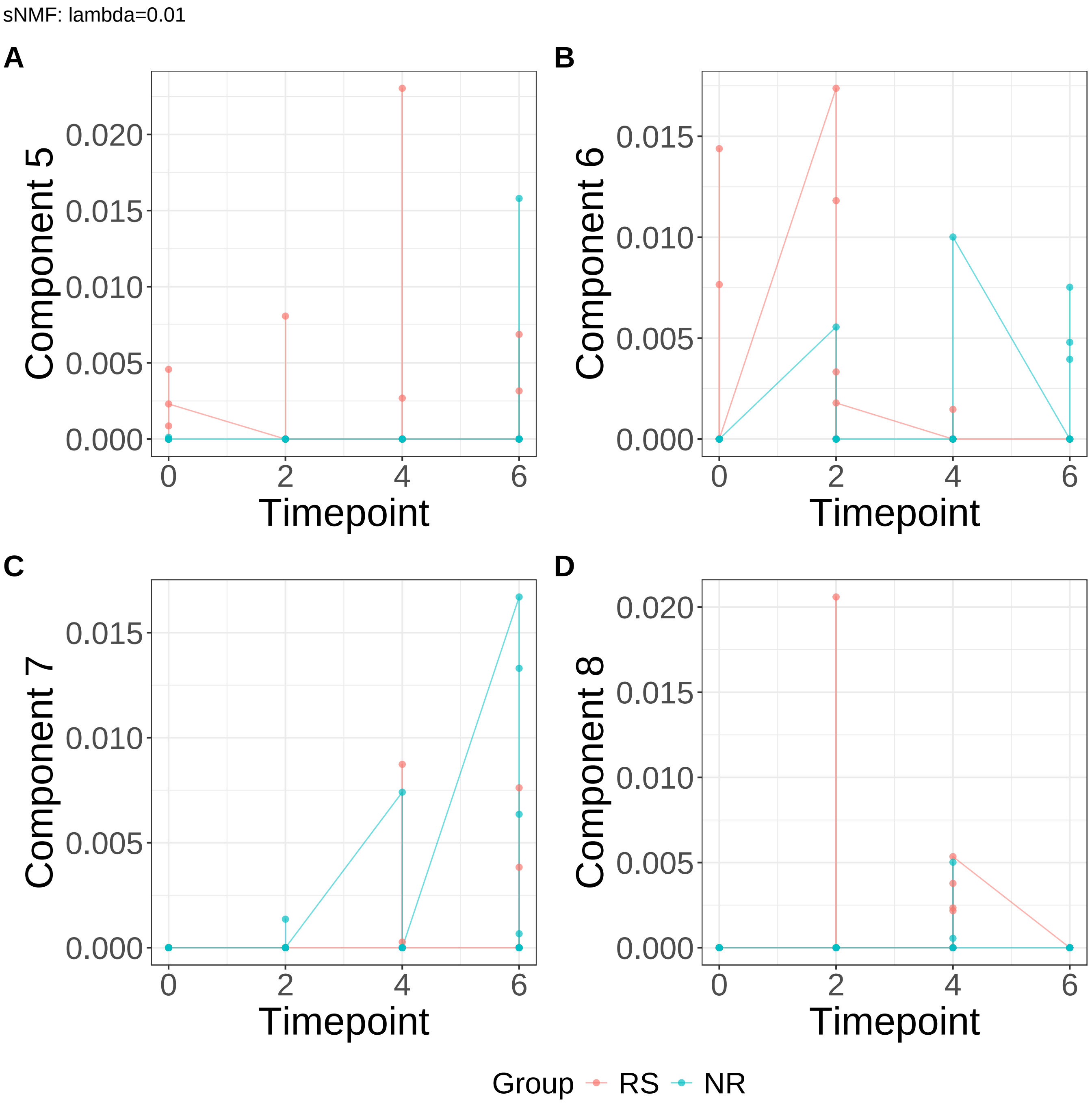


**Supplementary Figure 87.** Components for sNMF: lambda=0.01. (A) Component 5. (B) Component 6. (C) Component 7. (D) Component 8.

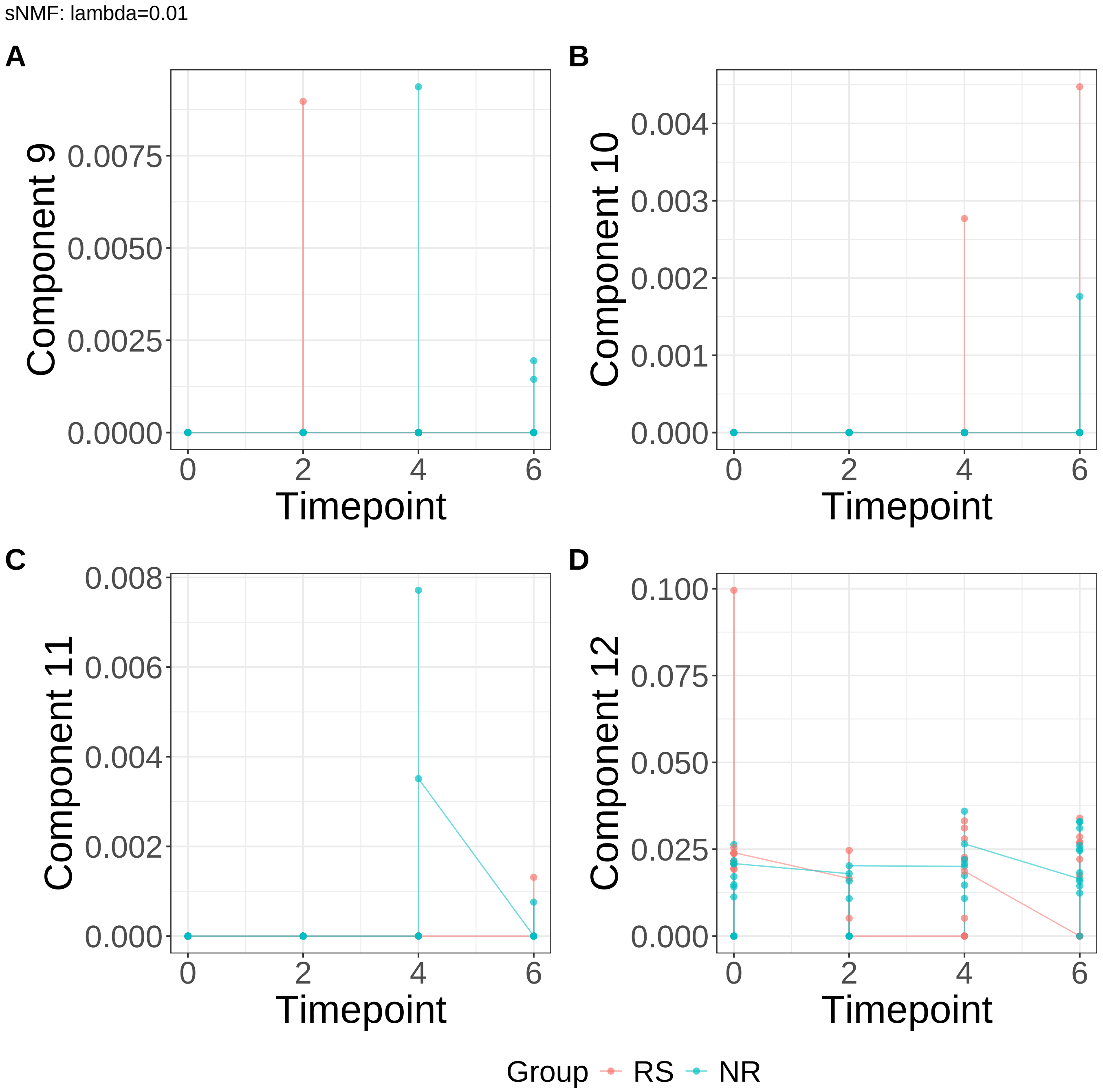


**Supplementary Figure 88.** Components for sNMF: lambda=0.01. (A) Component 9. (B) Component 10. (C) Component 11. (D) Component 12.

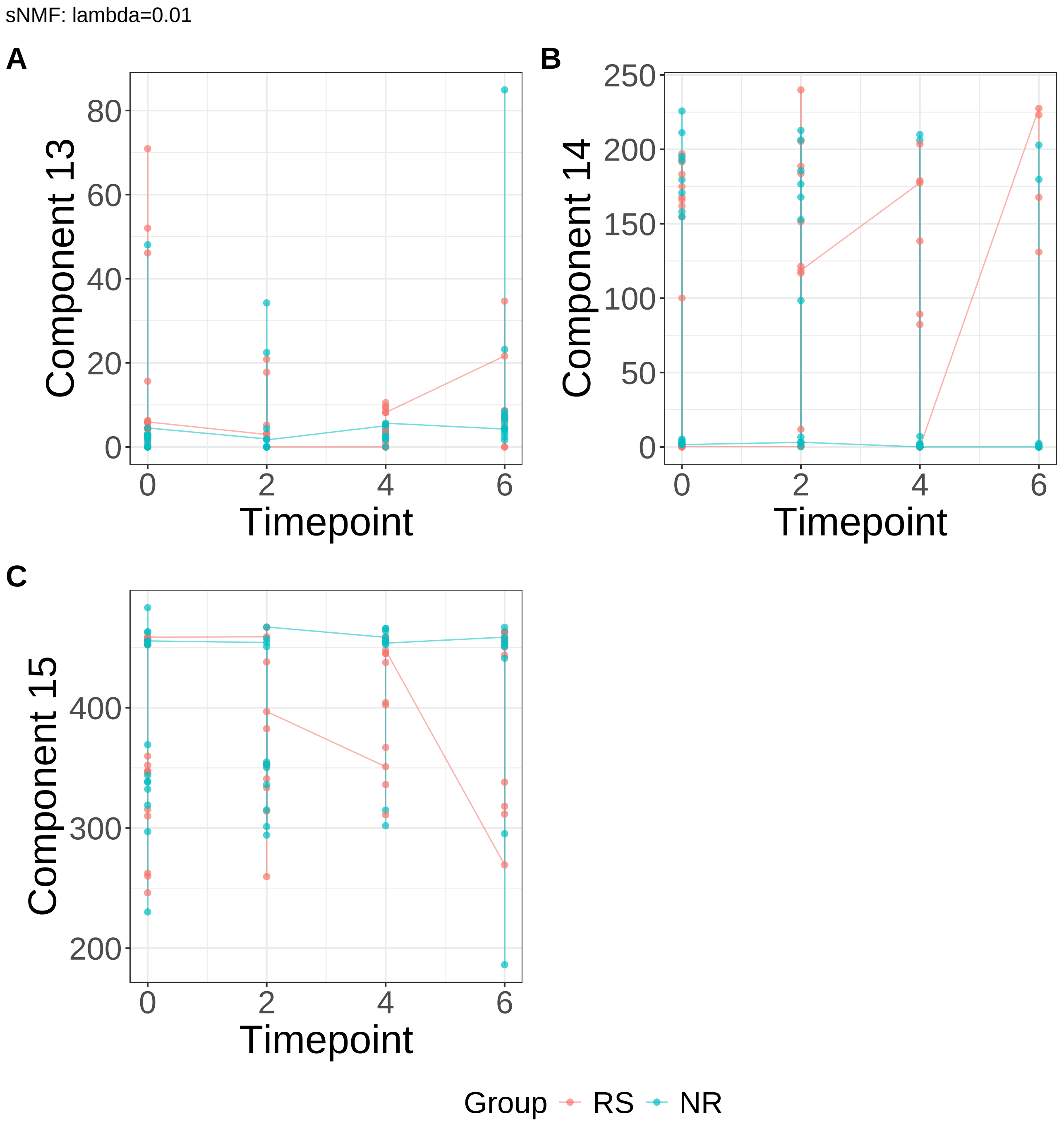


**Supplementary Figure 89.** Components for sNMF: lambda=0.01. (A) Component 13. (B) Component 14. (C) Component 15.

sNMF: lambda=0.1

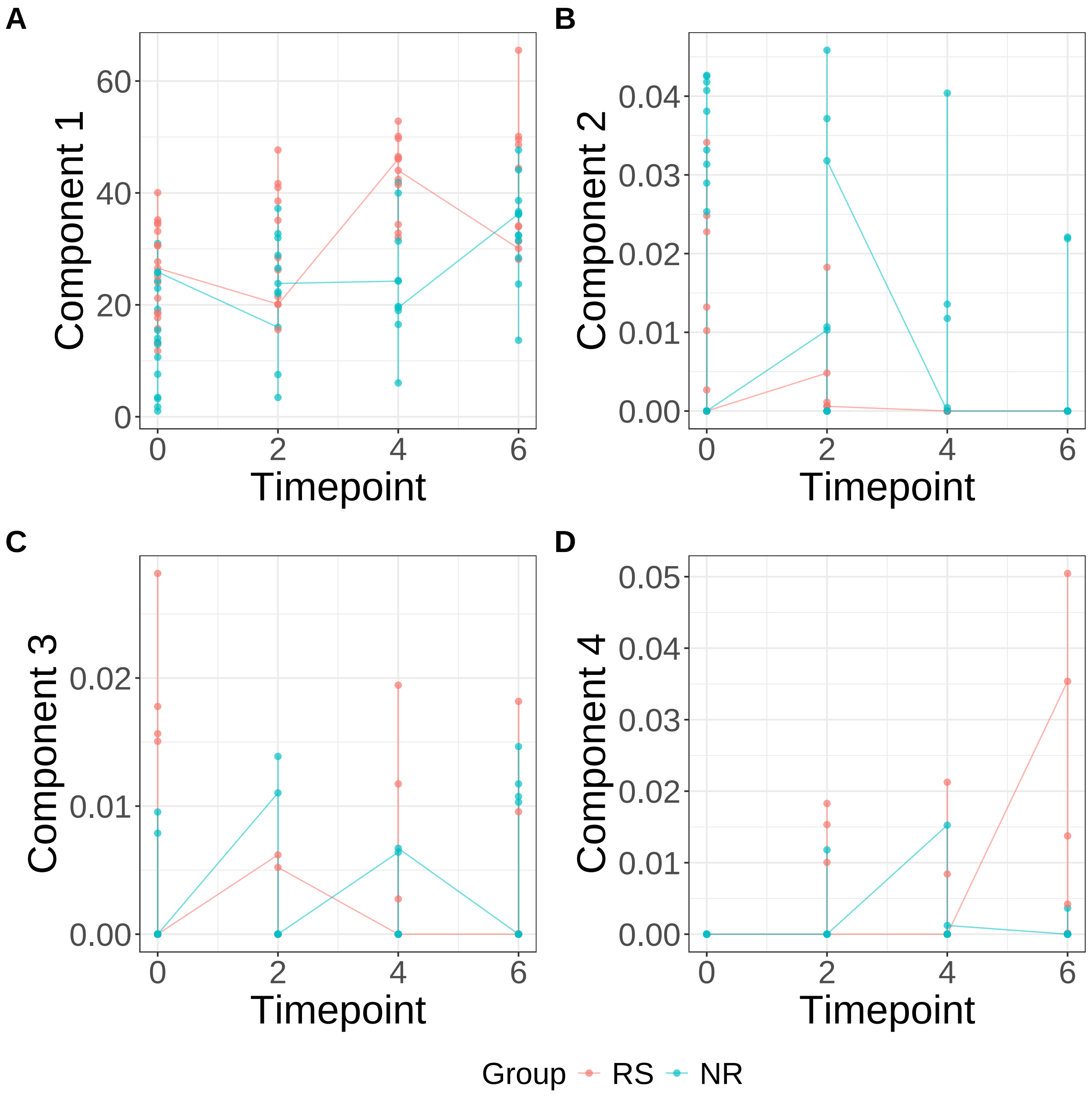


**Supplementary Figure 90.** Components for sNMF: lambda=0.1. (A) Component 1. (B) Component 2. (C) Component 3. (D) Component 4.

sNMF: lambda=0.1

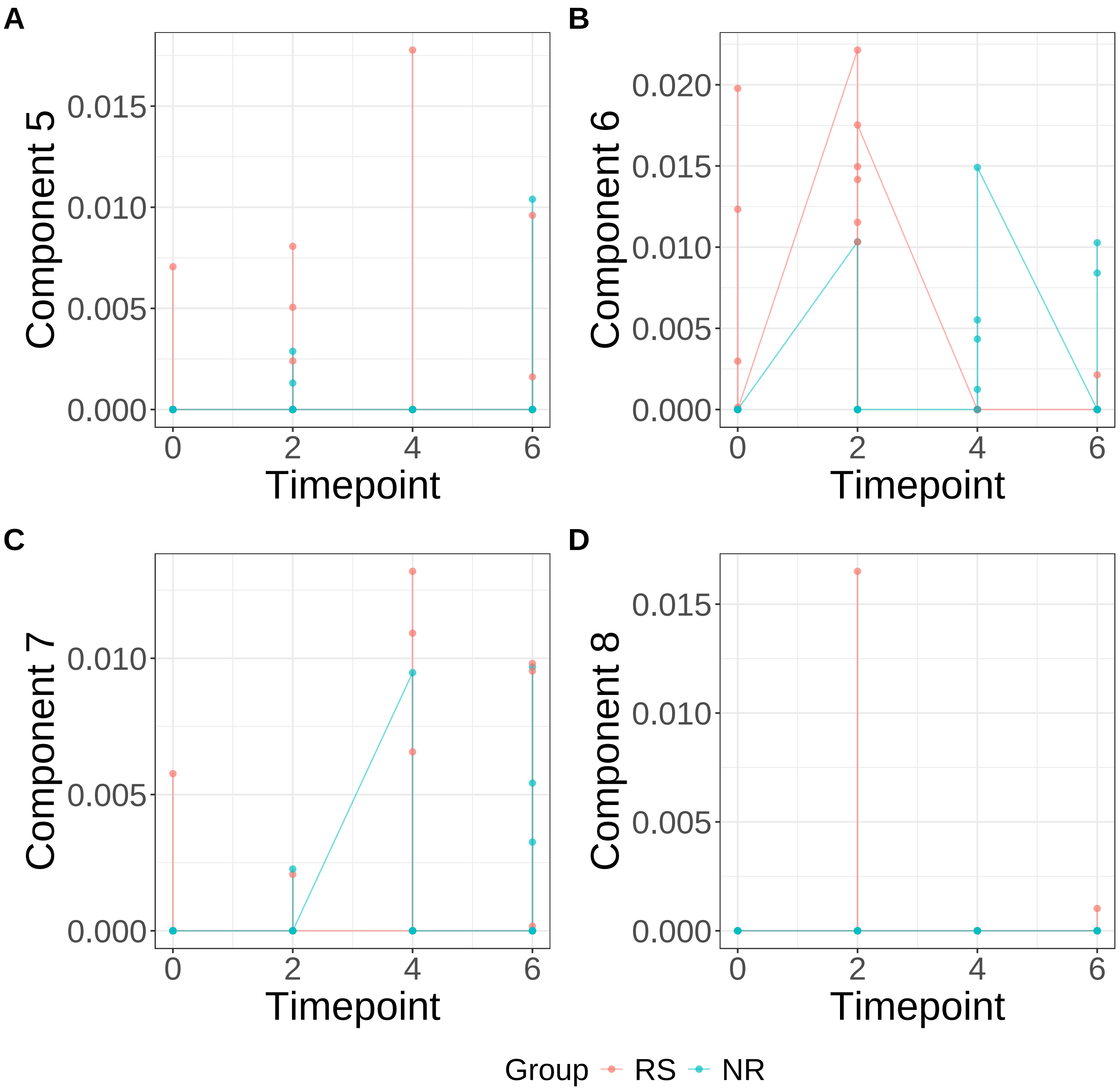


**Supplementary Figure 91.** Components for sNMF: lambda=0.1. (A) Component 5. (B) Component 6. (C) Component 7. (D) Component 8.

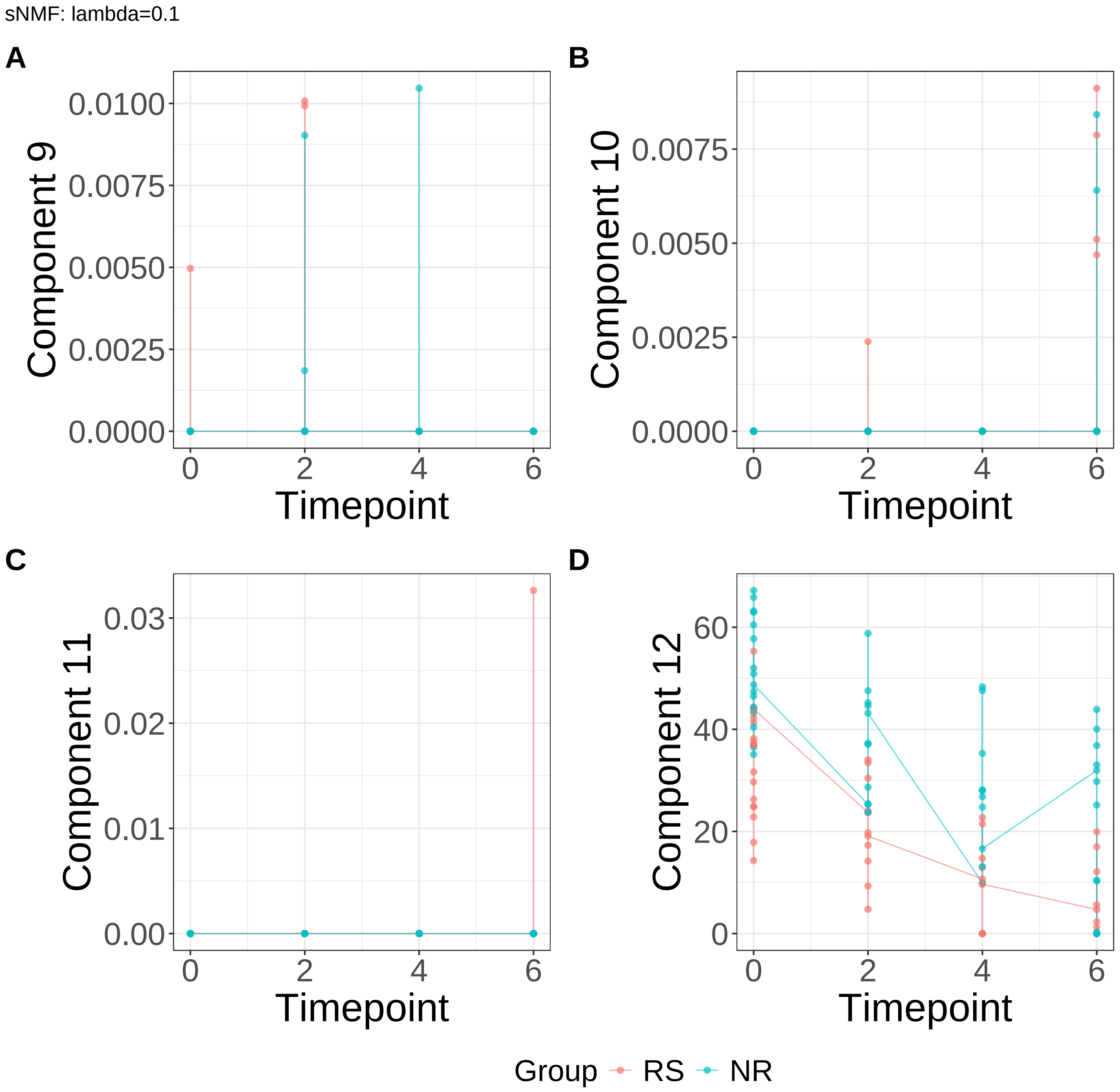


**Supplementary Figure 92.** Components for sNMF: lambda=0.1. (A) Component 9. (B) Component 10. (C) Component 11. (D) Component 12.

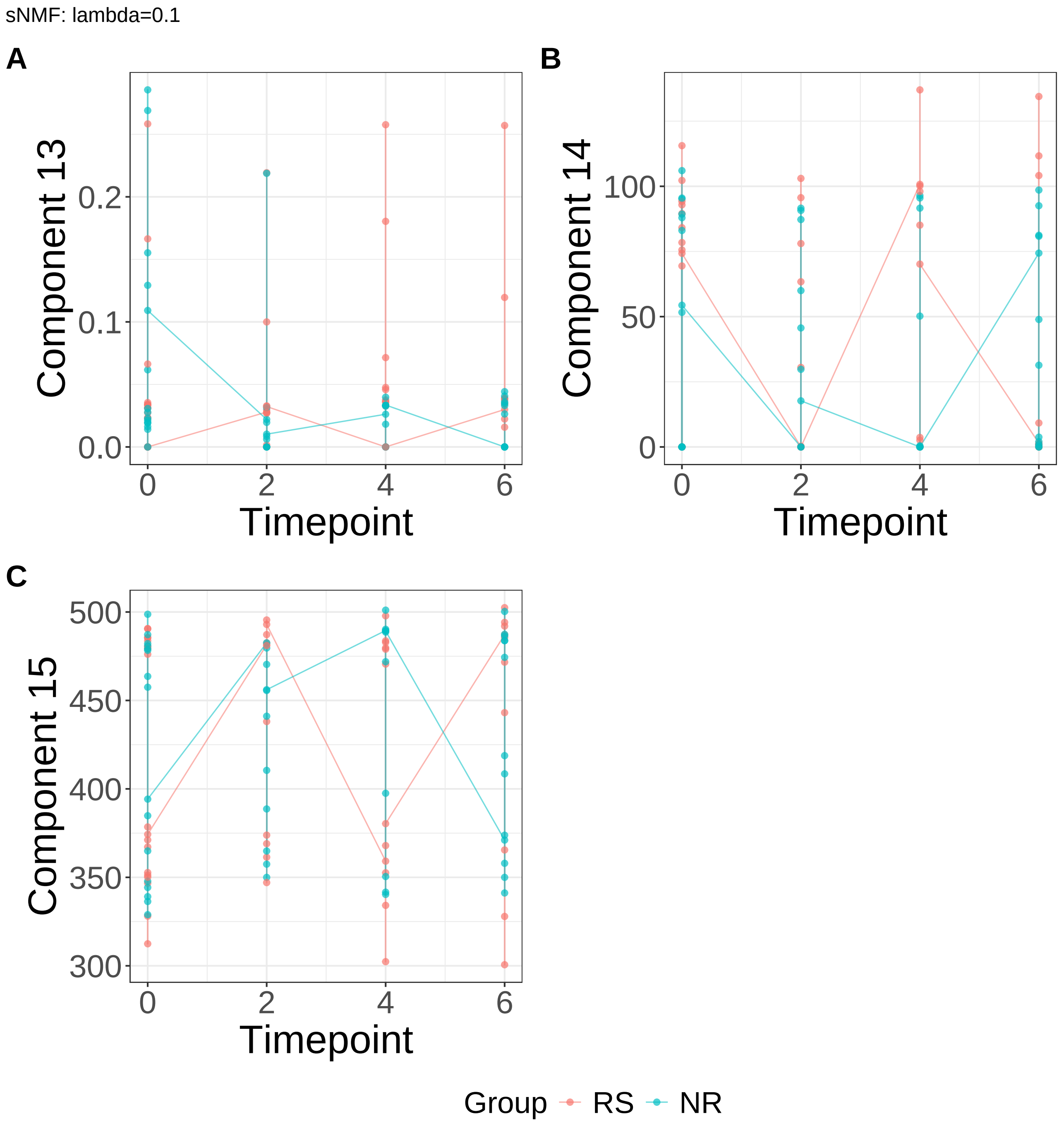


**Supplementary Figure 93.** Components for sNMF: lambda=0.1. (A) Component 13. (B) Component 14. (C) Component 15.

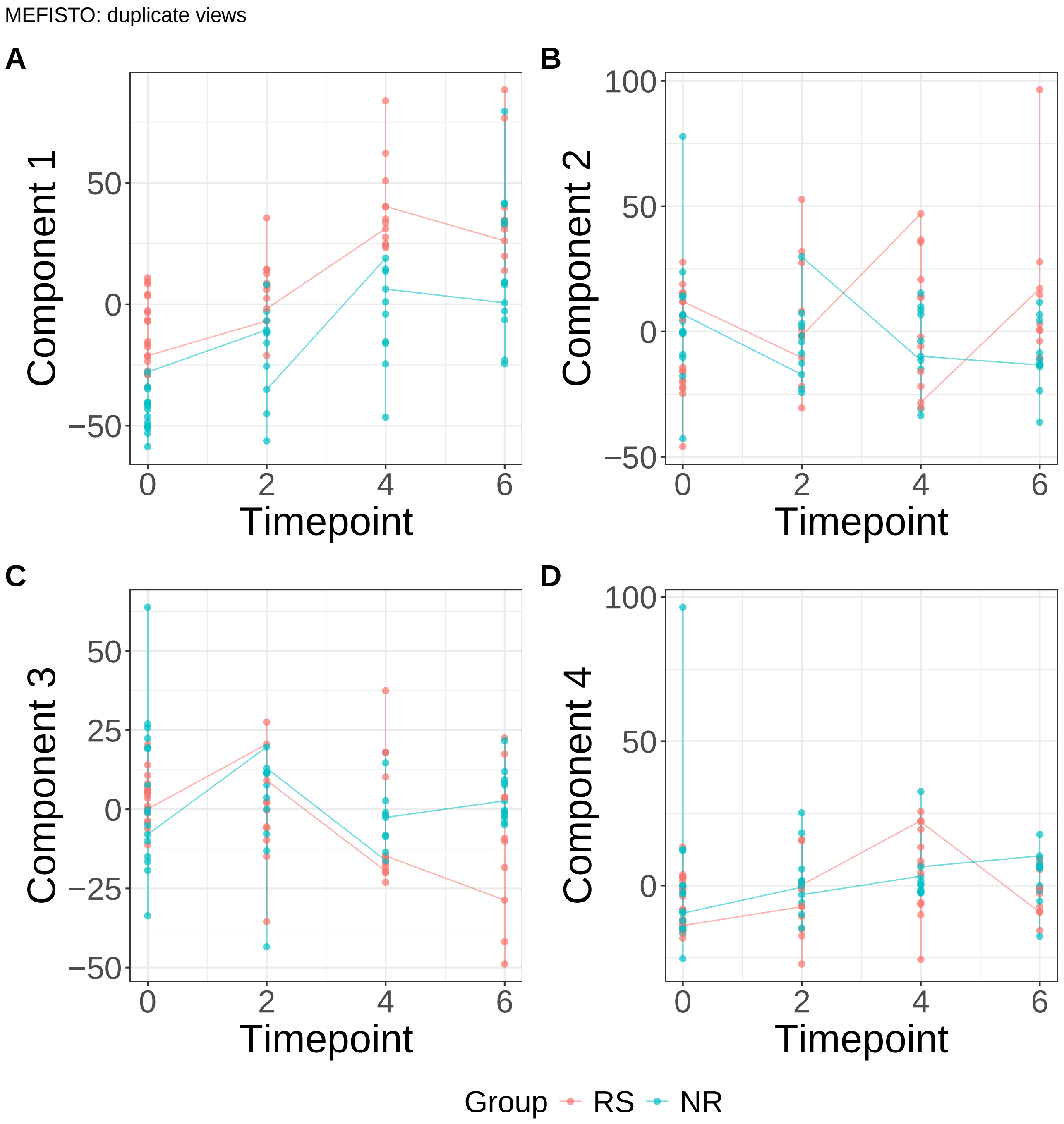


**Supplementary Figure 94.** Components for MEFISTO: duplicate views. (A) Component 1. (B) Component 2. (C) Component 3. (D) Component 4.

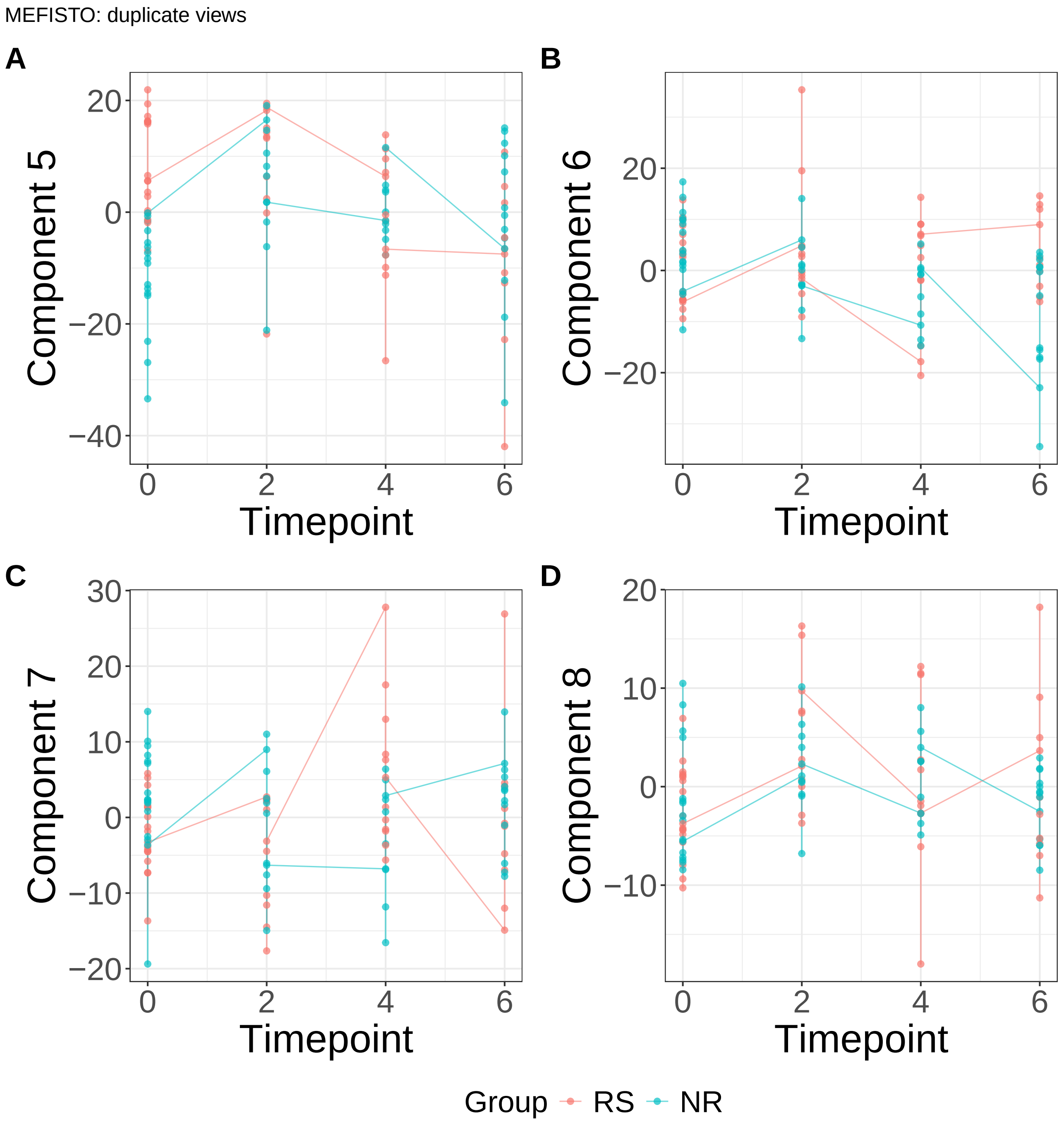


**Supplementary Figure 95.** Components for MEFISTO: duplicate views. (A) Component 5. (B) Component 6. (C) Component 7. (D) Component 8.

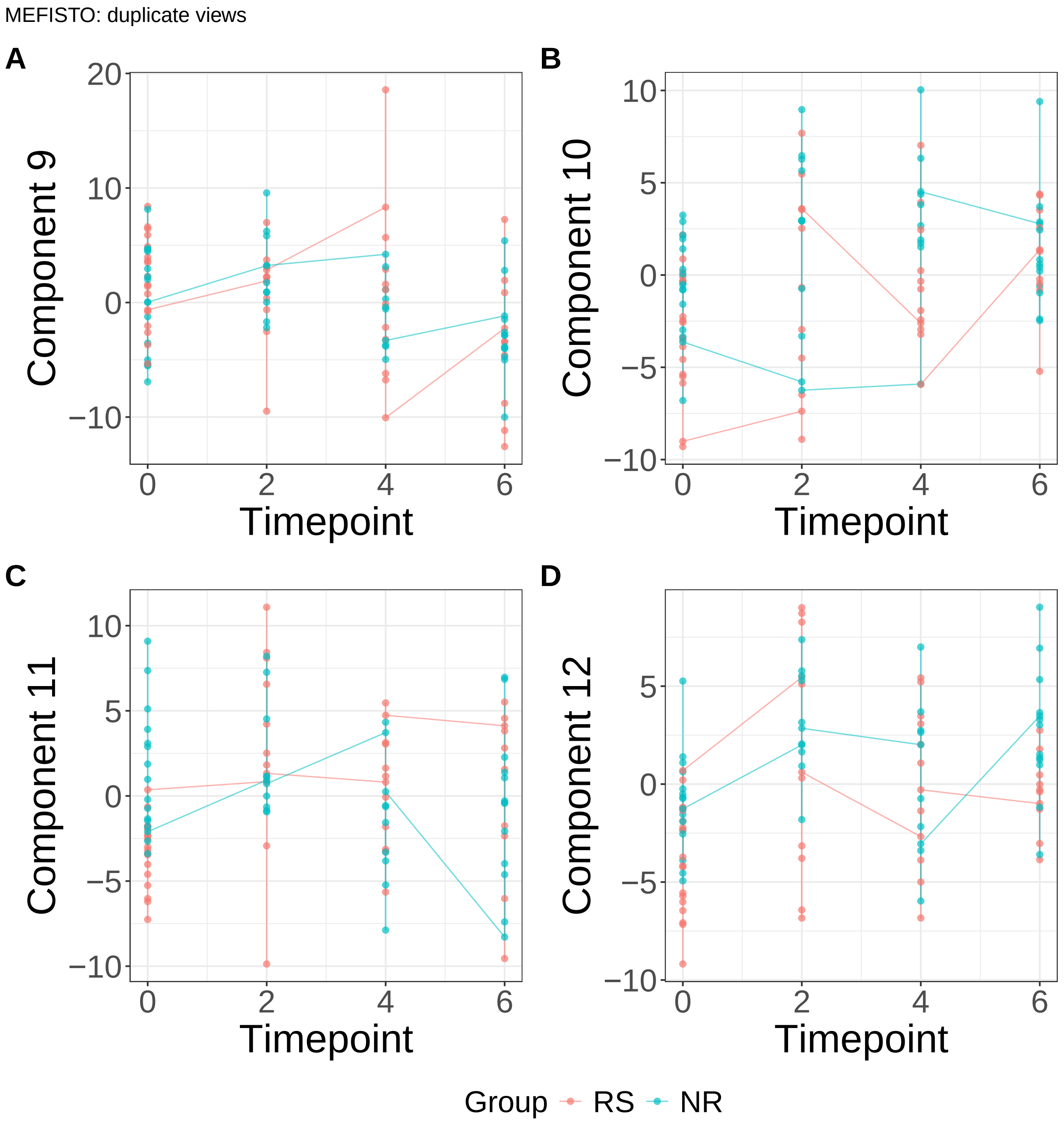


**Supplementary Figure 96.** Components for MEFISTO: duplicate views. (A) Component 9. (B) Component 10. (C) Component 11. (D) Component 12.

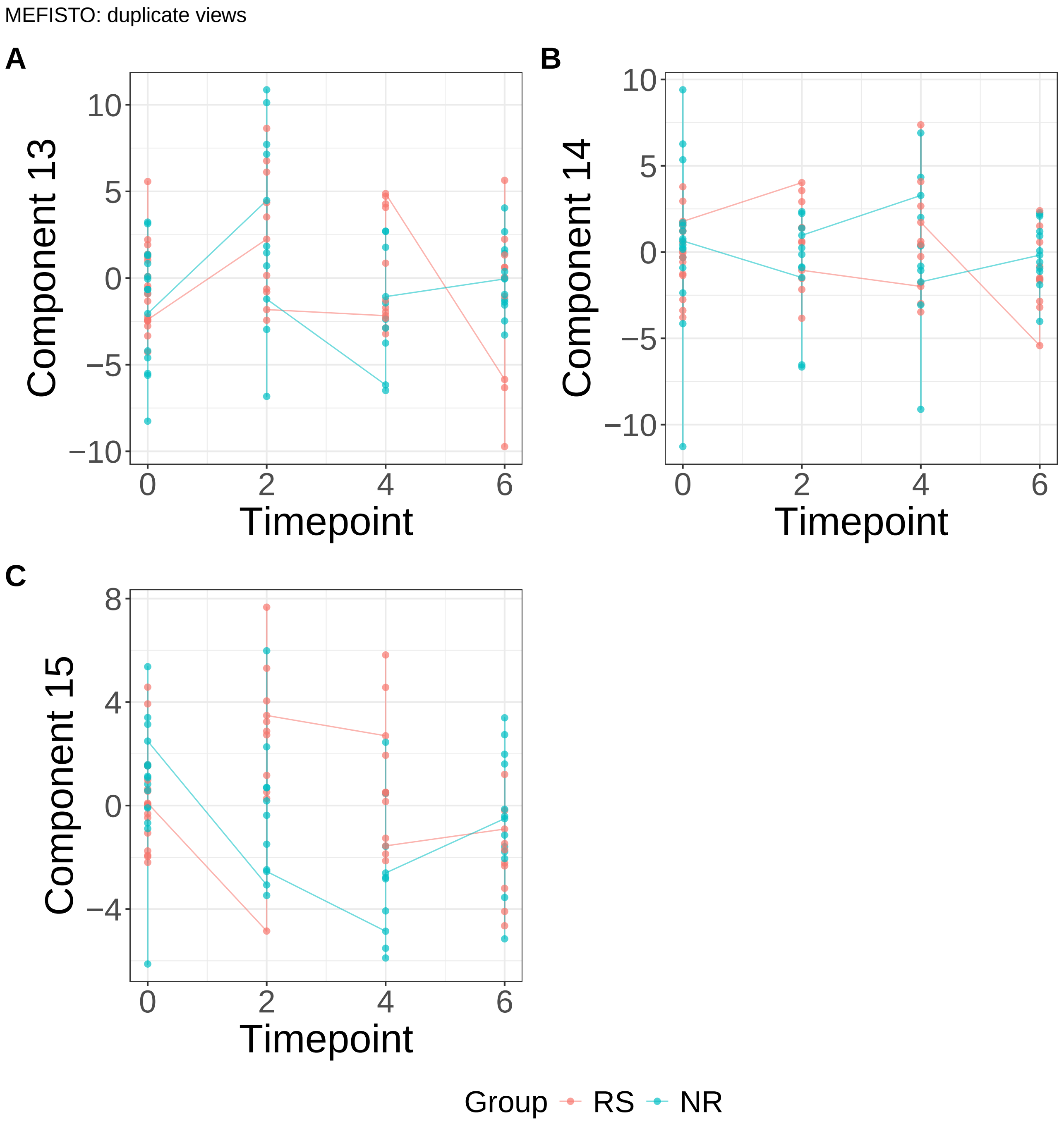


**Supplementary Figure 97.** Components for MEFISTO: duplicate views. (A) Component 13. (B) Component 14. (C) Component 15.
