## Supplementary Note 2 : Circadian Rhythms for "A statistical framework for system-level synthesis"

This note presents the supplementary results from the analysis of circadian rhythms data. The figures include :

- Processes/component functions learned by SSANOVA-SVD (**Supplementary Fig. 10**)
- Inference of feature participation with the processes from the application of the three-step framework: Volcano plots (**Supplementary Fig. 11**), heatmaps of top 50 component-associated features at FDR of 10% (**Supplementary Fig. 12**-**Supplementary Fig. 13**), geneset pathway enrichment for associated features (**Supplementary Fig. 14**)
- Decomposition of the processes with respect to covariates (**Supplementary Fig. 15**-**Supplementary Fig. 16**). Because we have a single term (time), the decomposition is identical to the processes themselves.
- Components from a representative class of matrix factorization methods (**Supplementary Fig. 17**-**Supplementary Fig. 53**)

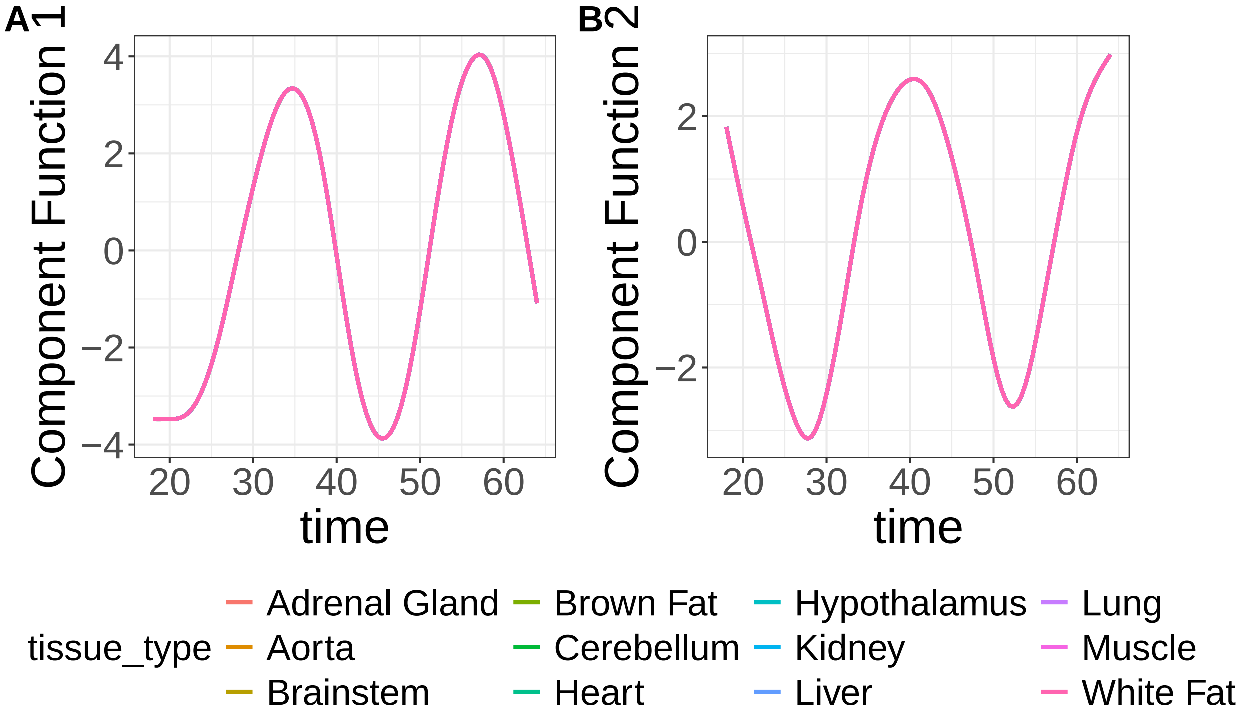


**Supplementary Figure 10.** SSANOVA-SVD reconstructed component functions.(A) Component 1. (B) Component 2.

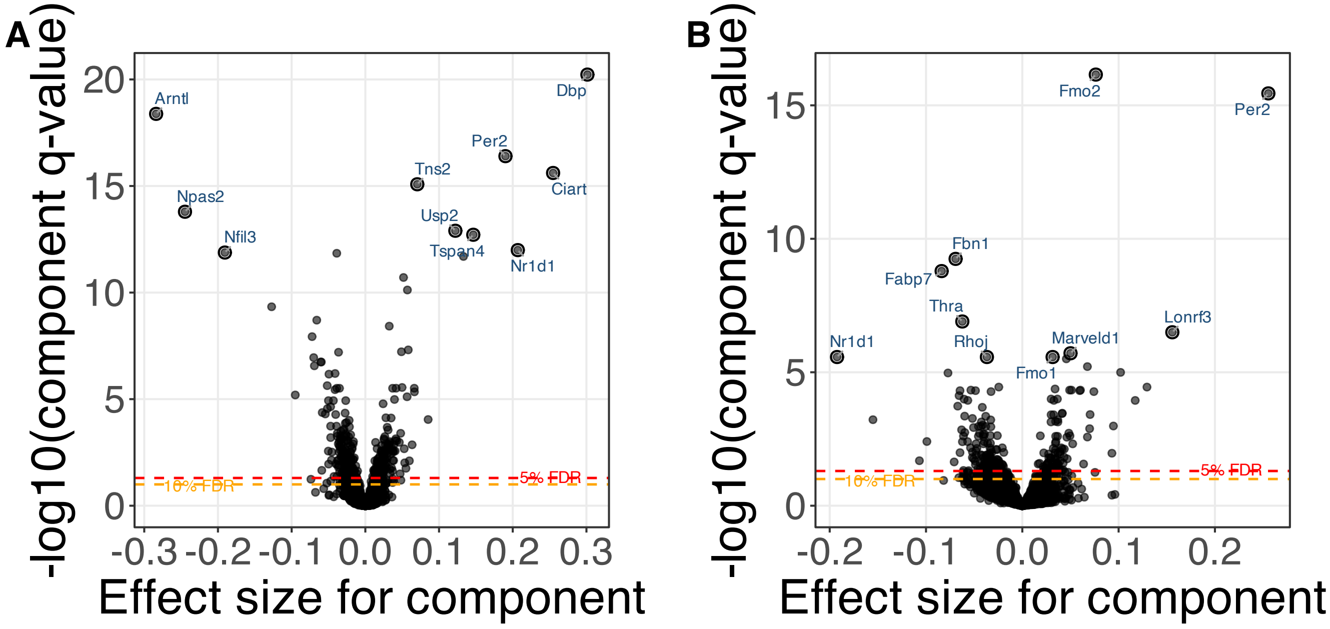


**Supplementary Figure 11.** SSANOVA-SVD based conditional inference for feature participation. (A) Component 1. (B) Component 2.

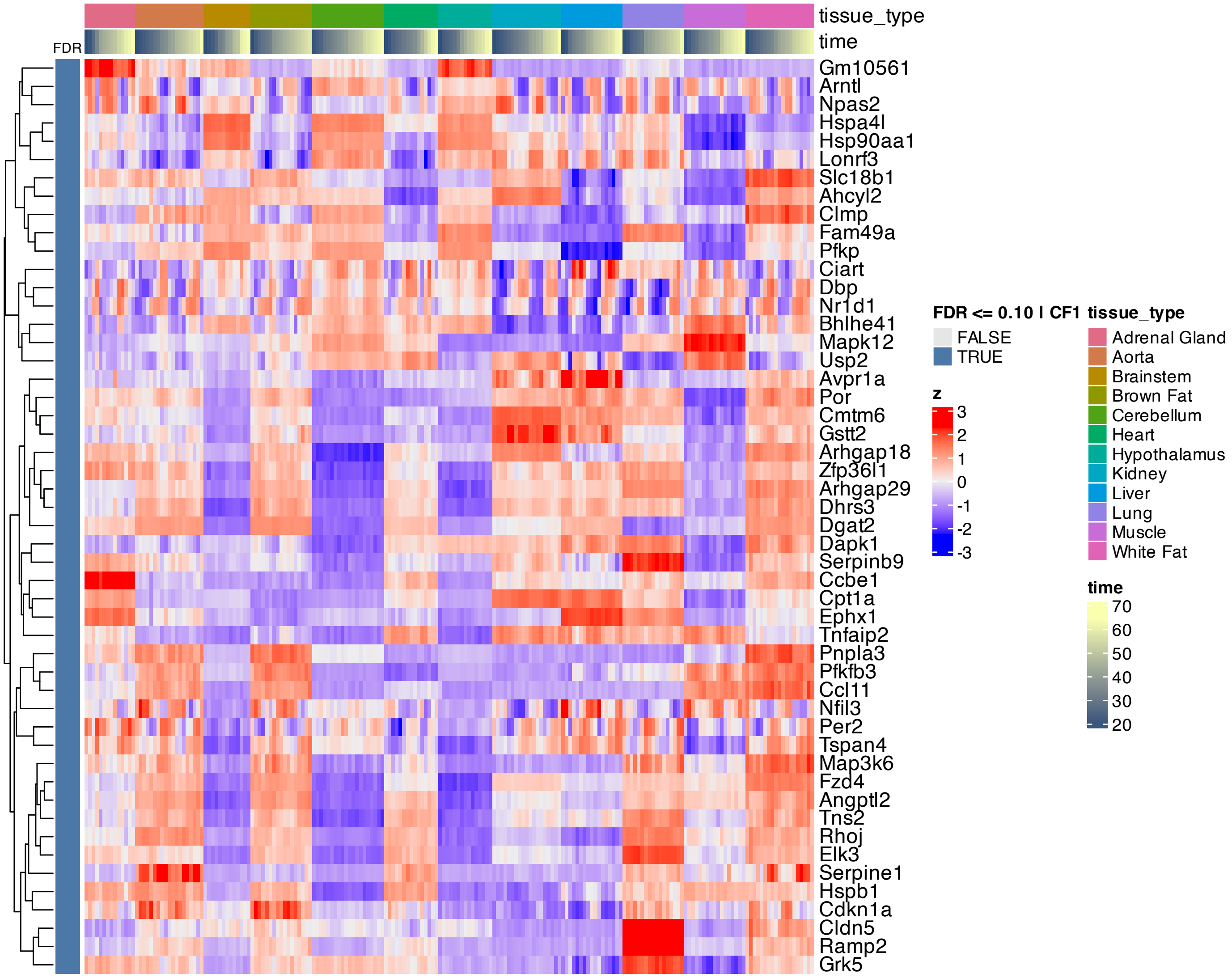


**Supplementary Figure 12.** Heatmap of features inferred to be associated with component 1 by SSANOVA-SVD.

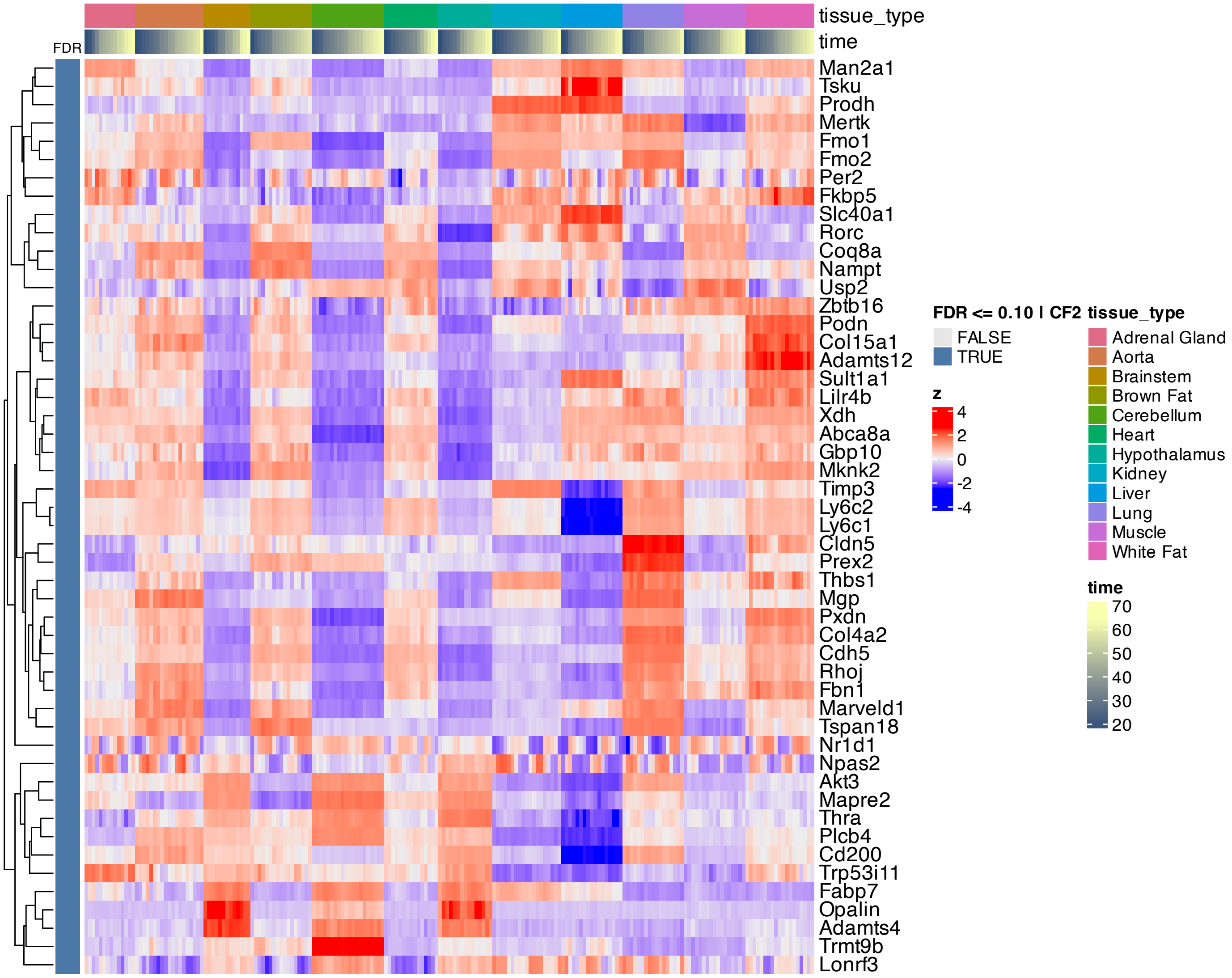


**Supplementary Figure 13.** Heatmap of features inferred to be associated with component 2 by SSANOVA-SVD.

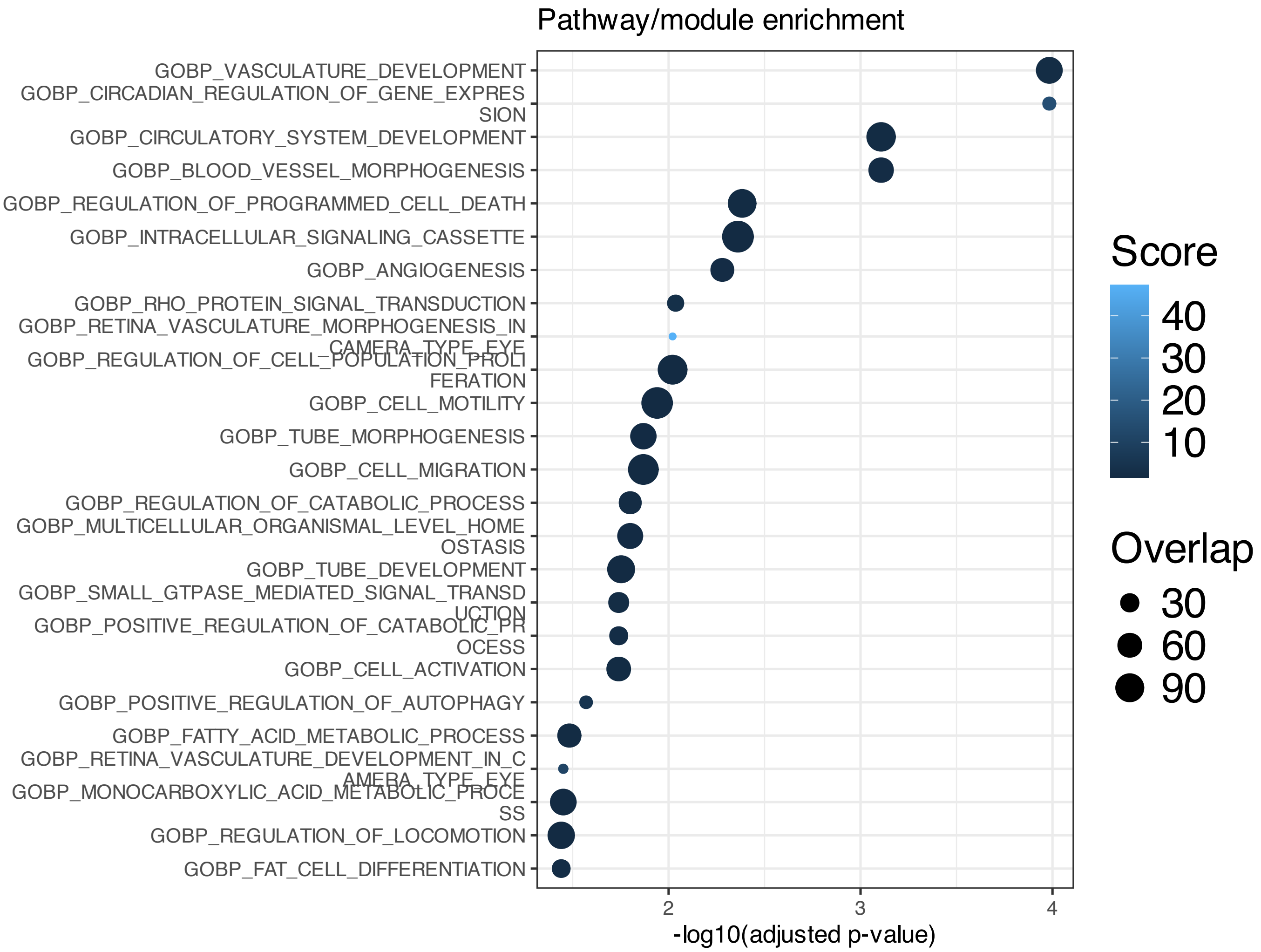


**Supplementary Figure 14.** Enrichment among features inferred to be associated with SSANOVA-SVD component 1; only terms with adjusted P $\leq 0.1$ are shown.

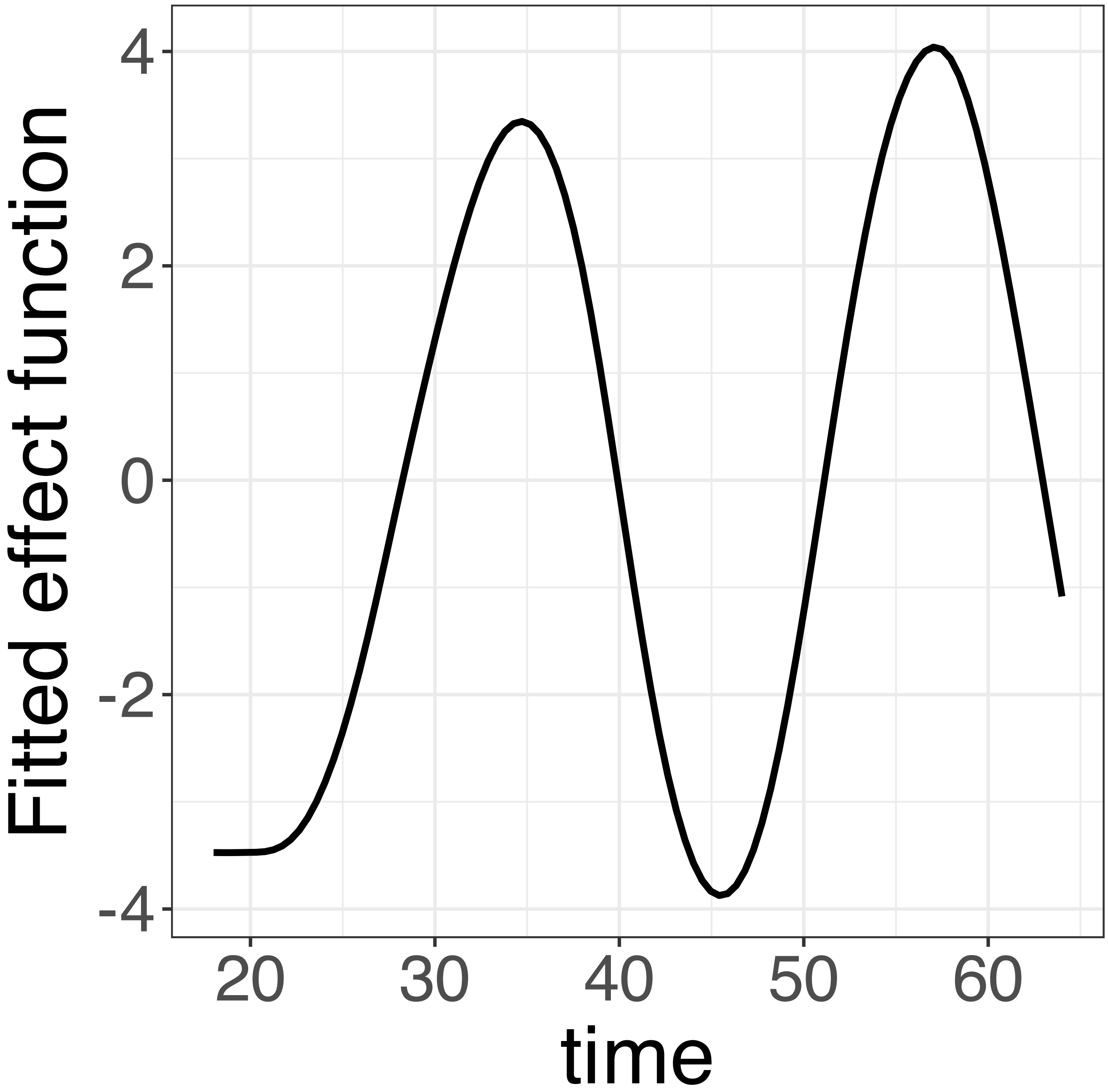


**Supplementary Figure 15.** Decomposition of reconstructed SSANOVA-SVD component function 1.

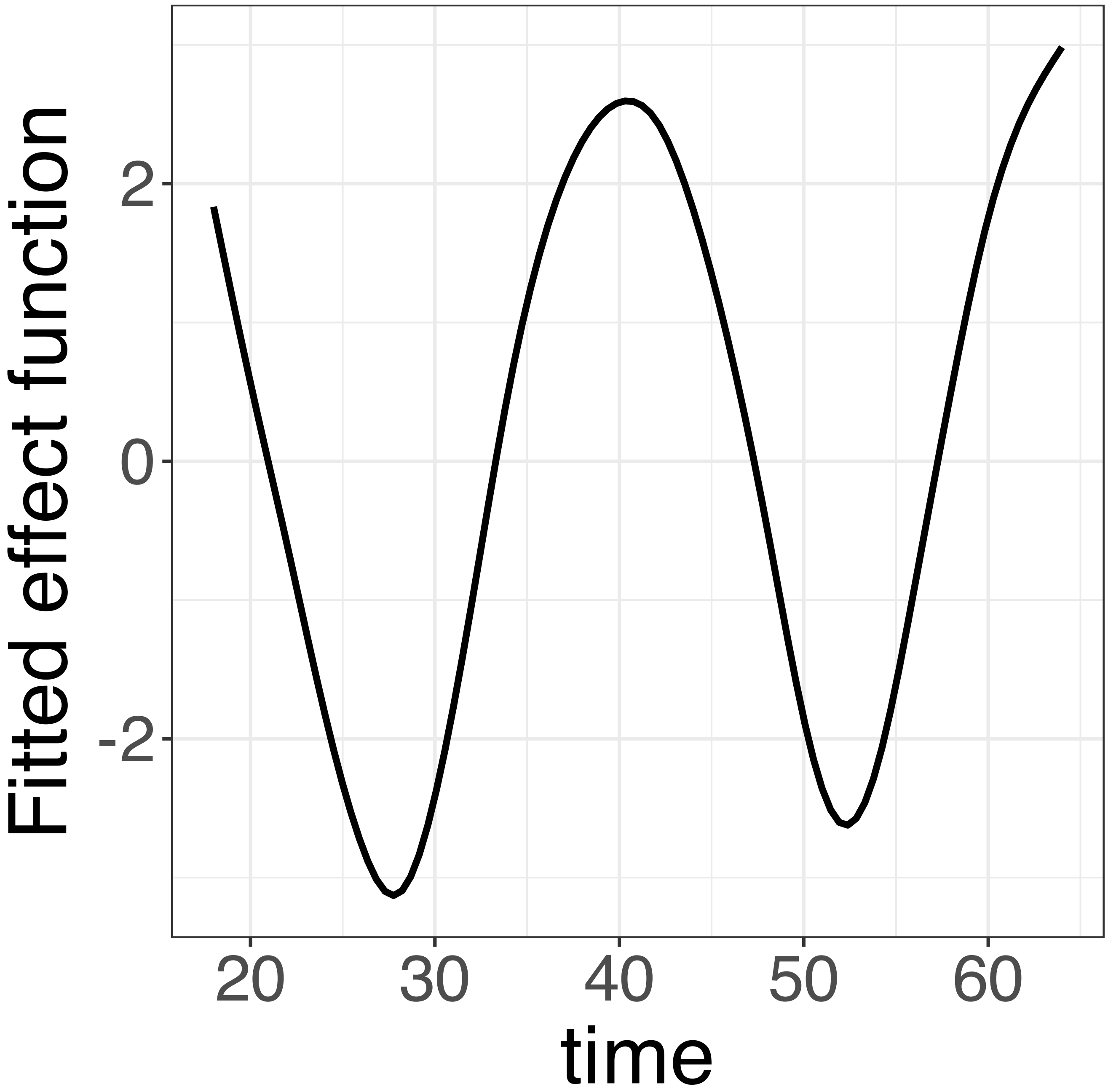


**Supplementary Figure 16.** Decomposition of reconstructed SSANOVA-SVD component function 2.

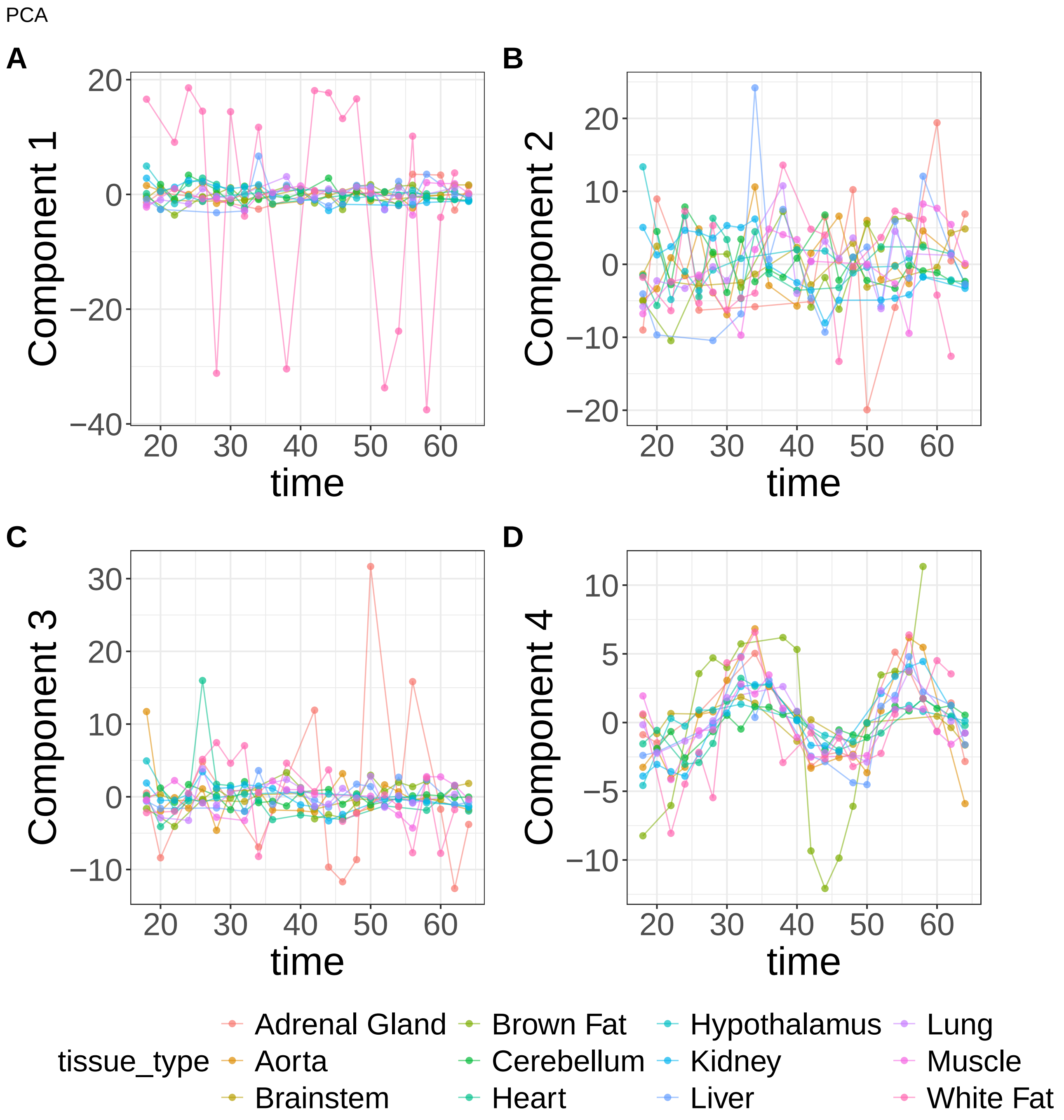


**Supplementary Figure 17.** Components for PCA. (A) Component 1. (B) Component 2. (C) Component 3. (D) Component 4.

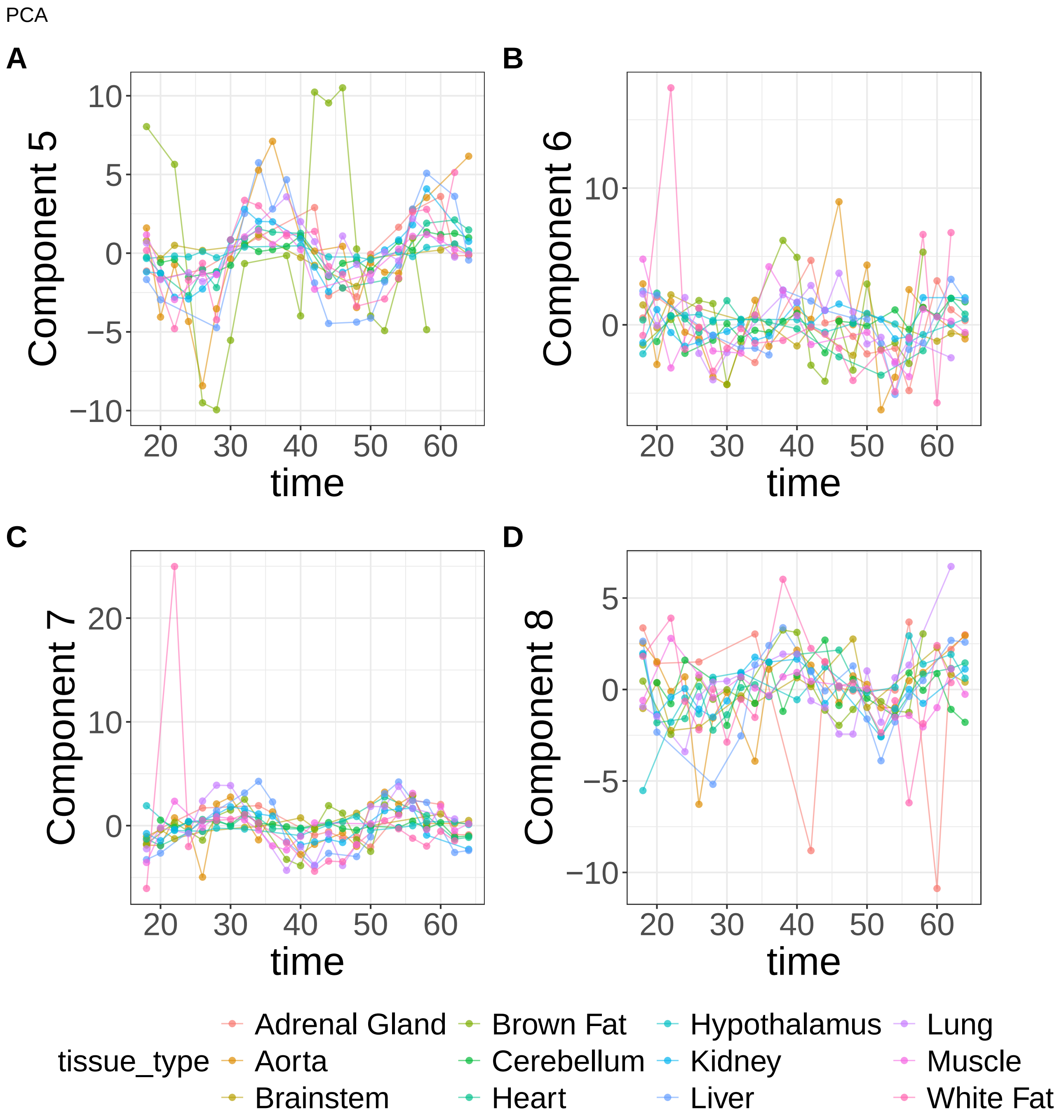


**Supplementary Figure 18.** Components for PCA. (A) Component 5. (B) Component 6. (C) Component 7. (D) Component 8.

PCA

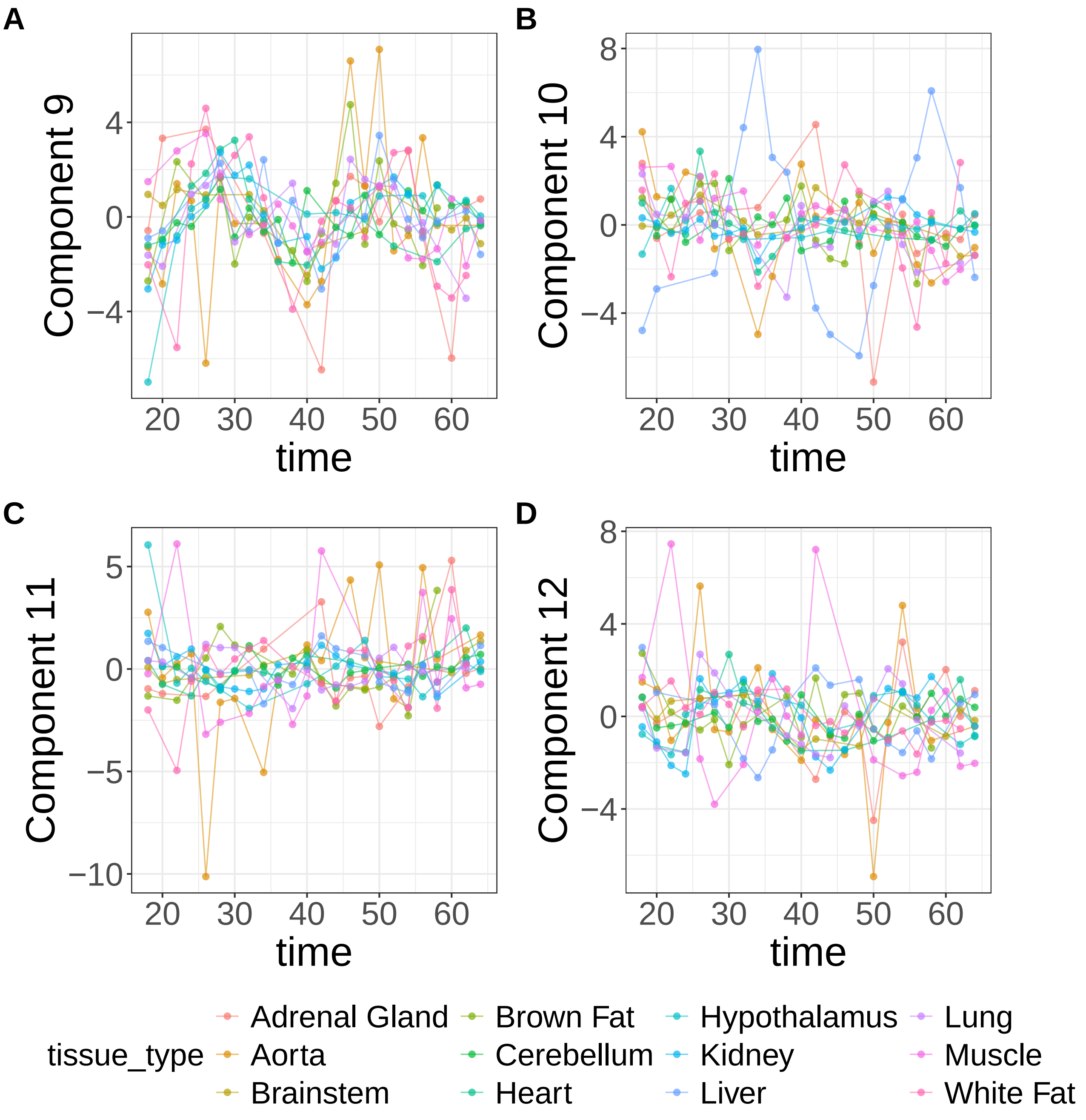


**Supplementary Figure 19.** Components for PCA. (A) Component 9. (B) Component 10. (C) Component 11. (D) Component 12.

PCA

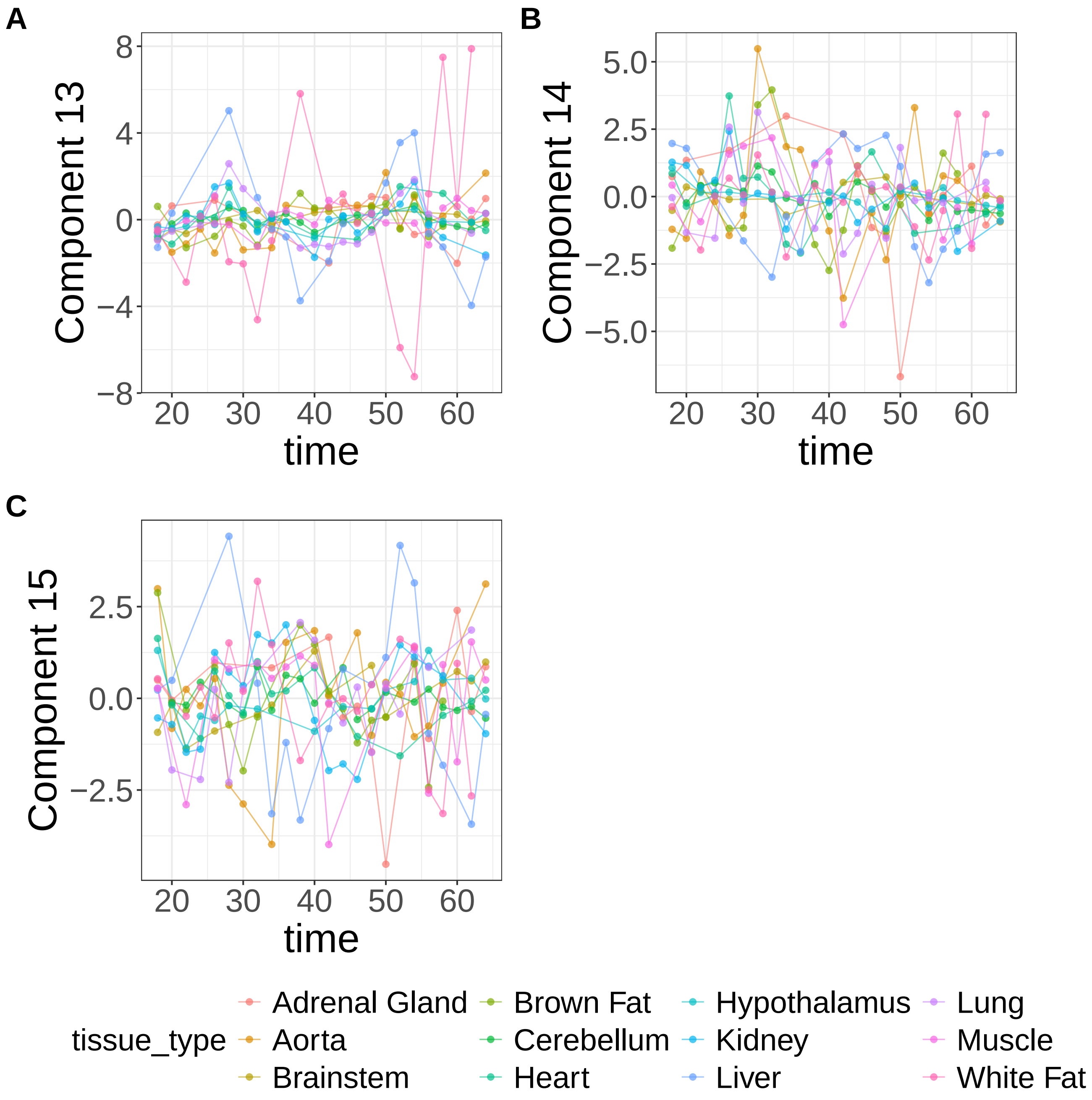


**Supplementary Figure 20.** Components for PCA. (A) Component 13. (B) Component 14. (C) Component 15.

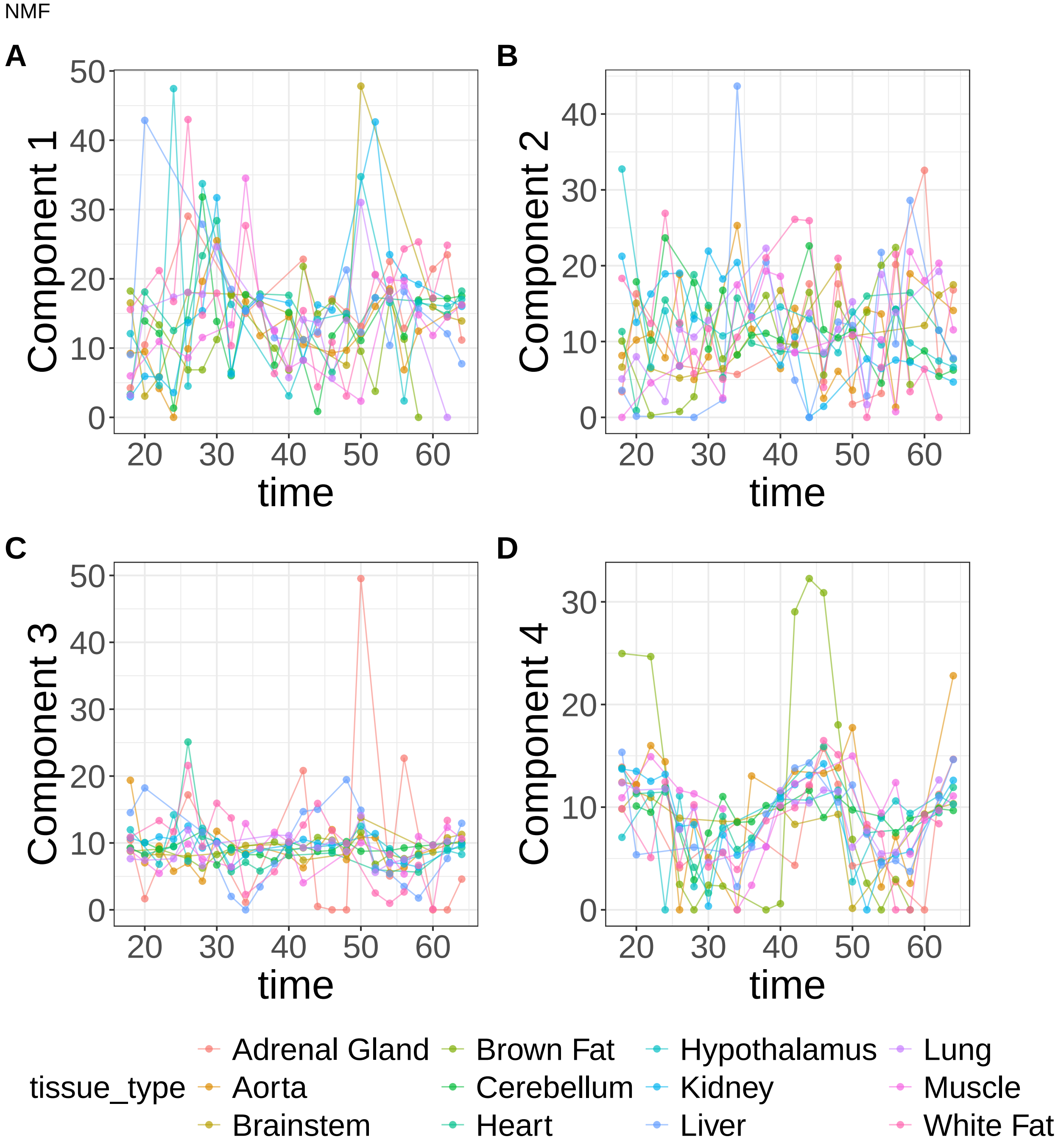


**Supplementary Figure 21.** Components for NMF. (A) Component 1. (B) Component 2. (C) Component 3. (D) Component 4.

NMF

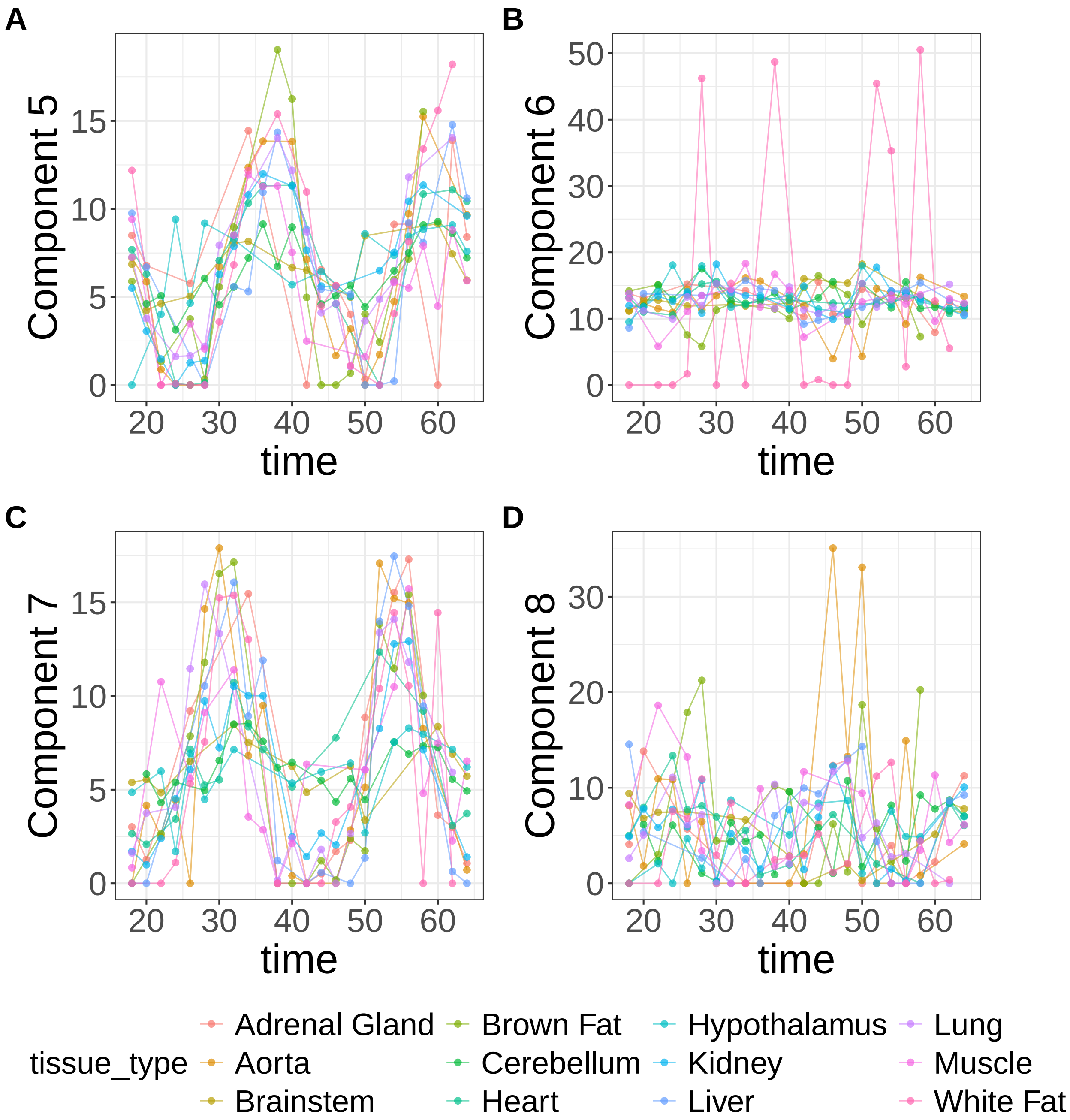


**Supplementary Figure 22.** Components for NMF. (A) Component 5. (B) Component 6. (C) Component 7. (D) Component 8.

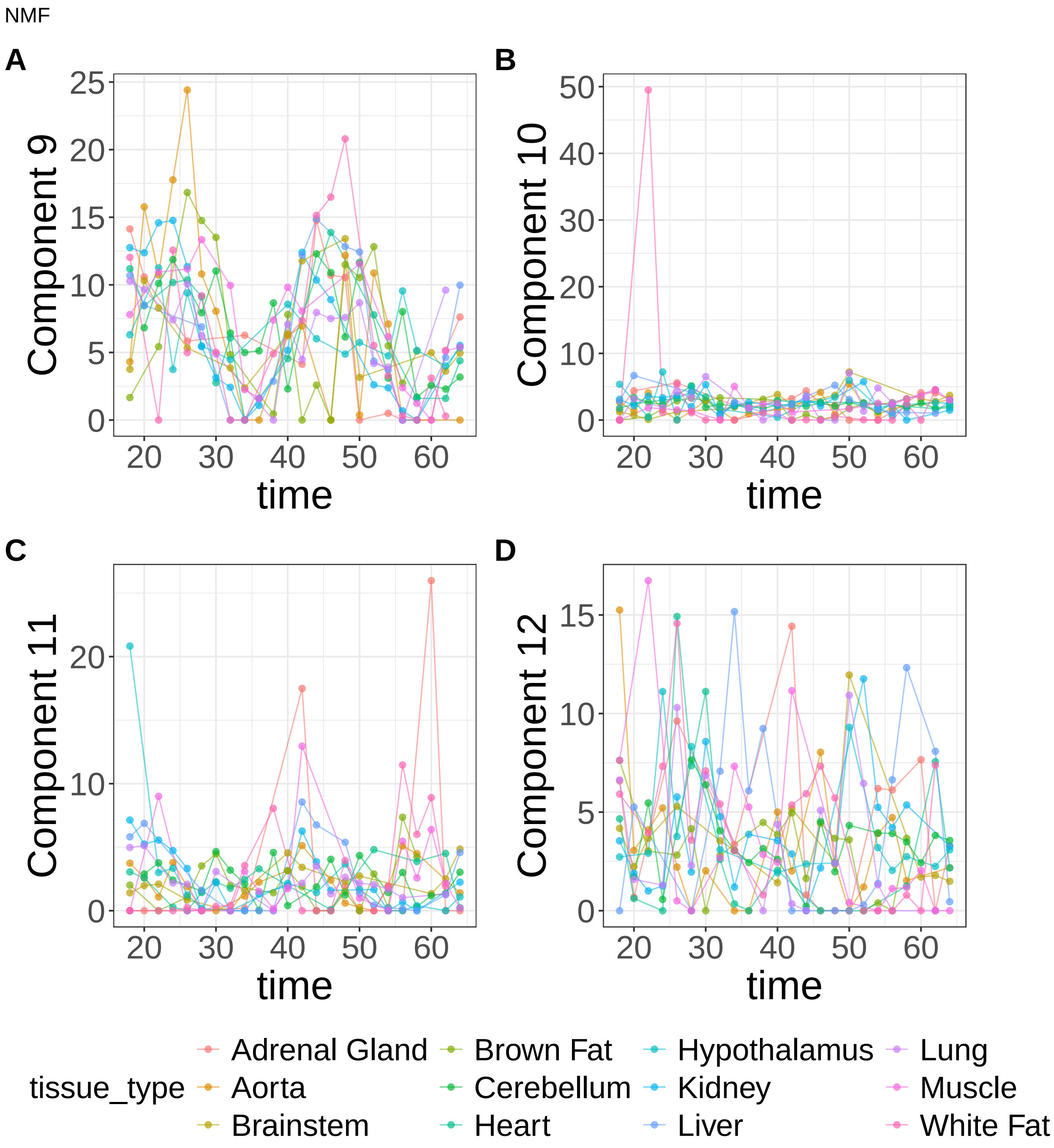


**Supplementary Figure 23.** Components for NMF. (A) Component 9. (B) Component 10. (C) Component 11. (D) Component 12.

NMF

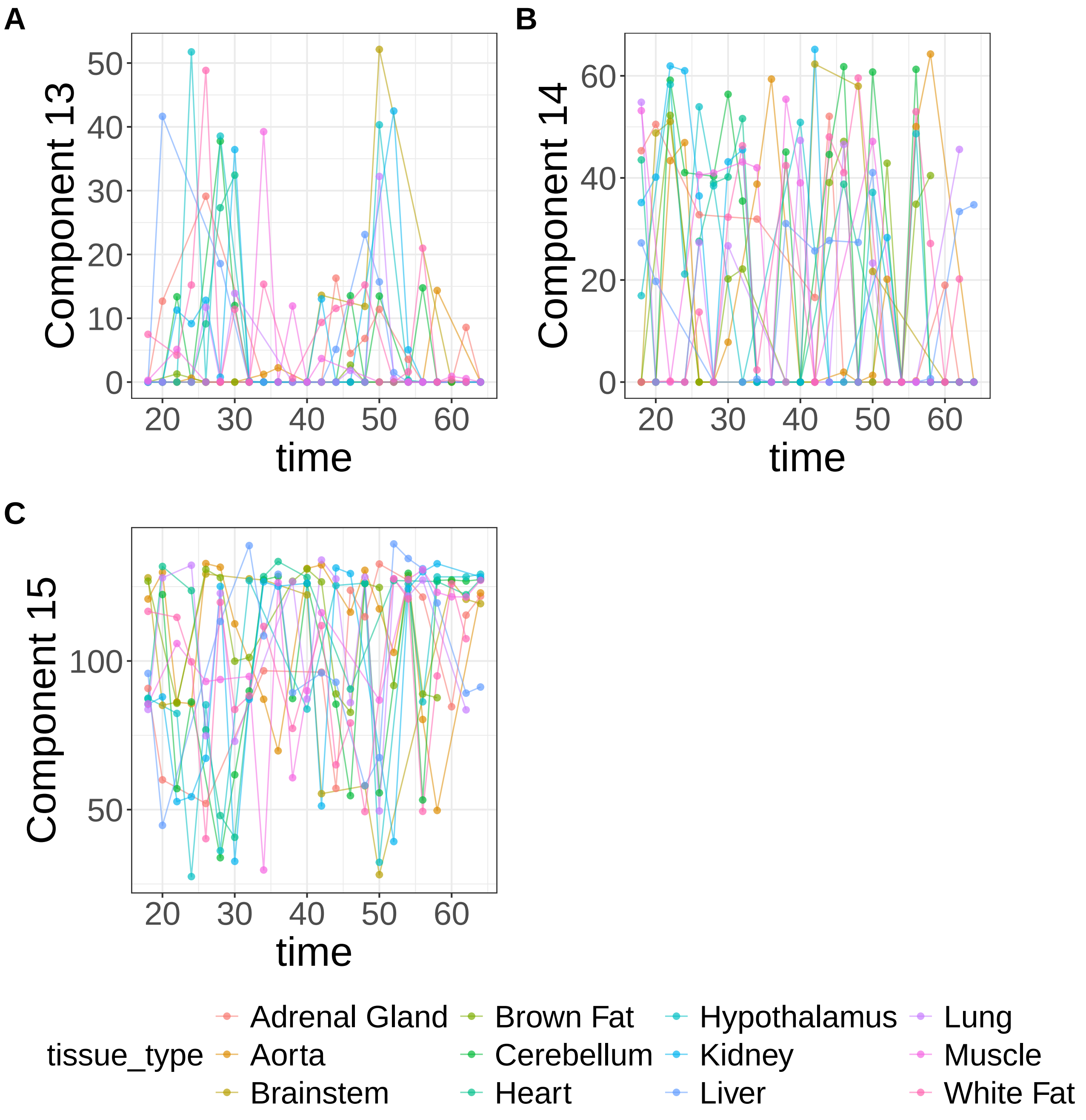


**Supplementary Figure 24.** Components for NMF. (A) Component 13. (B) Component 14. (C) Component 15.

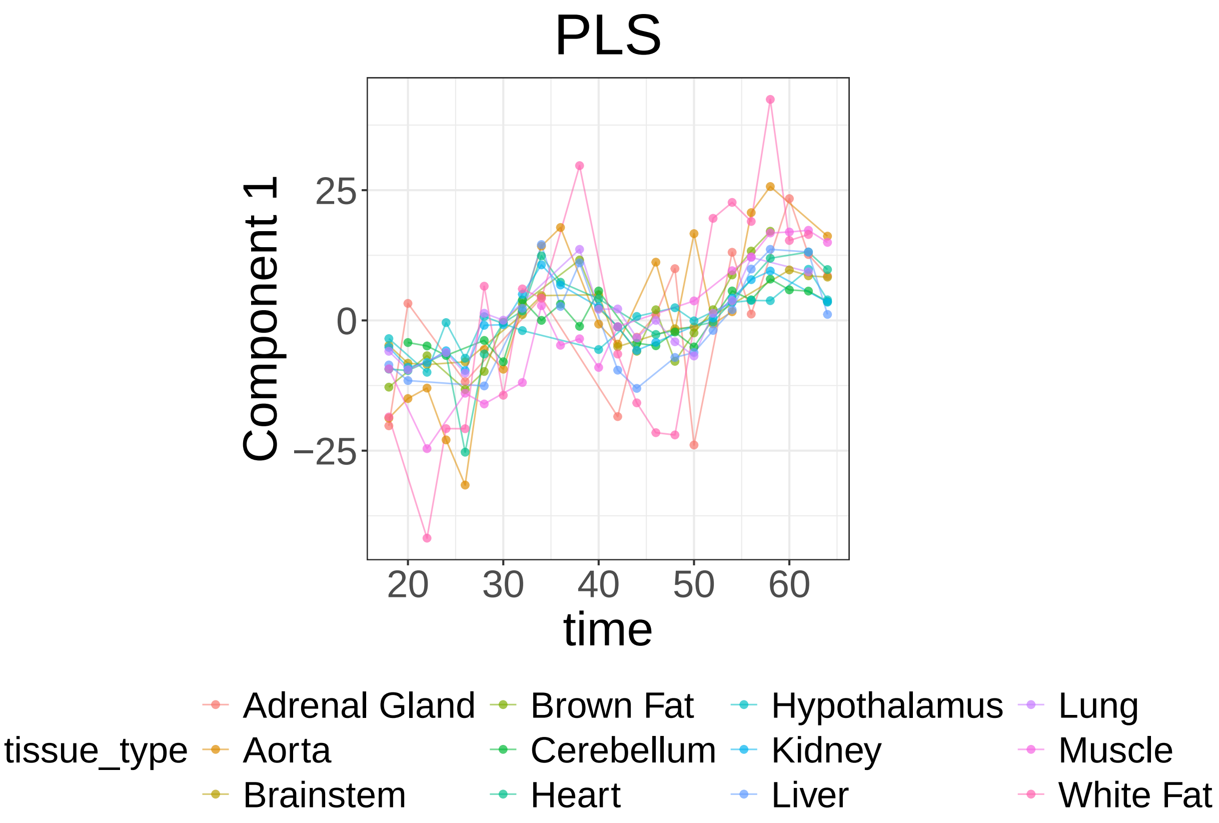


**Supplementary Figure 25.** Components for PLS. Component 1.

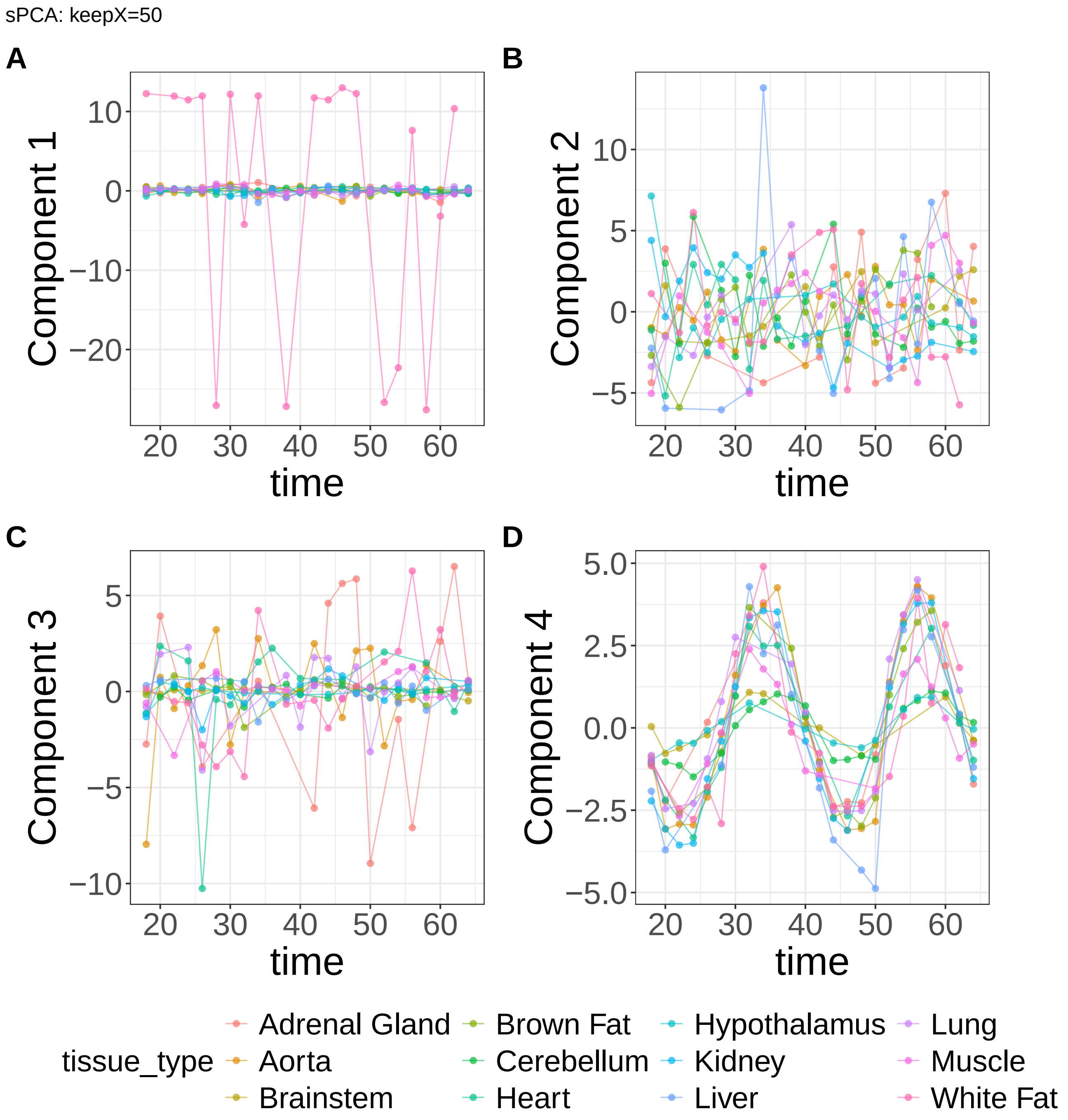


**Supplementary Figure 26.** Components for sPCA: keepX=50. (A) Component 1. (B) Component 2. (C) Component 3. (D) Component 4.

sPCA: keepX=50

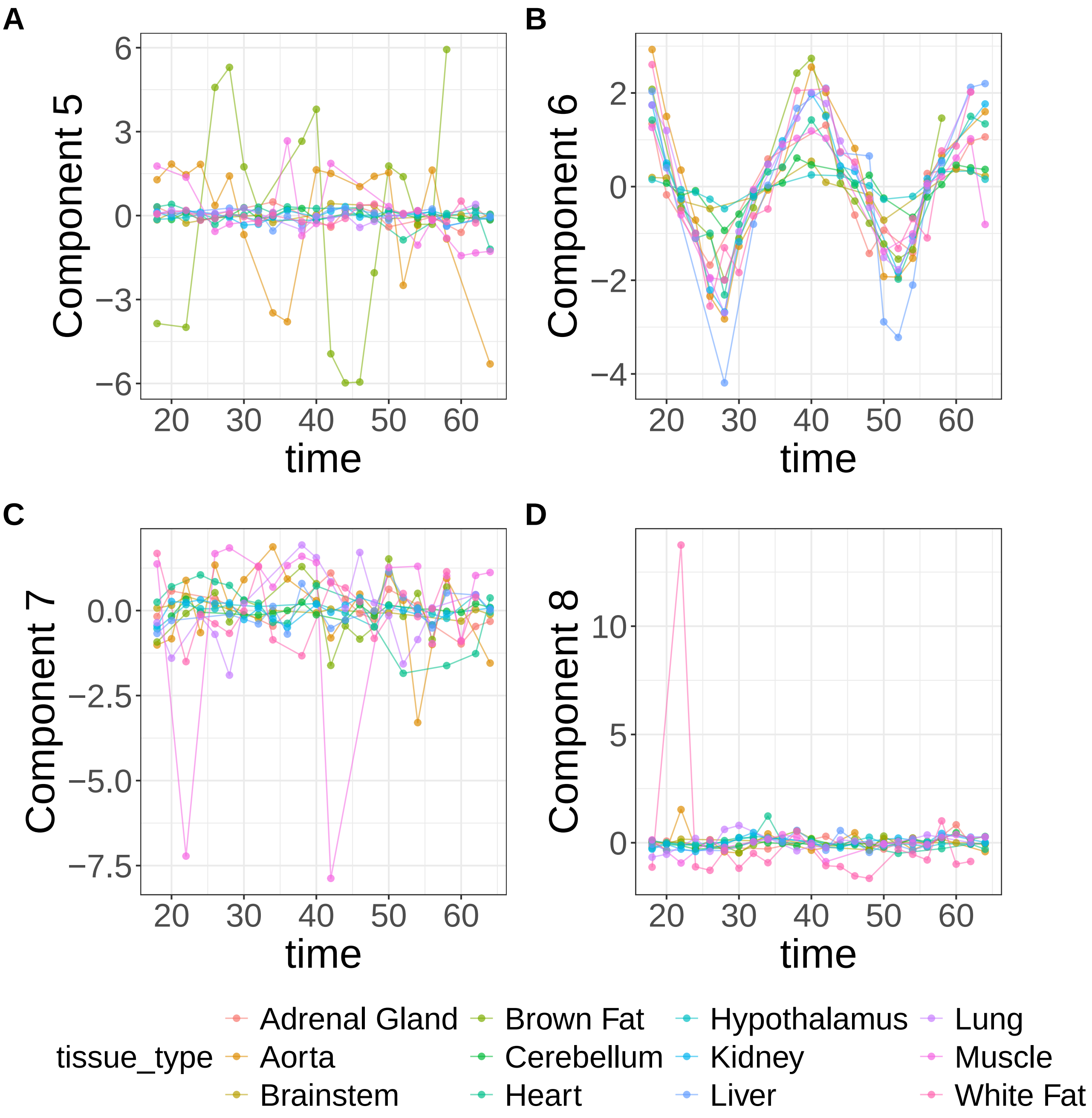


**Supplementary Figure 27.** Components for sPCA: keepX=50. (A) Component 5. (B) Component 6. (C) Component 7. (D) Component 8.

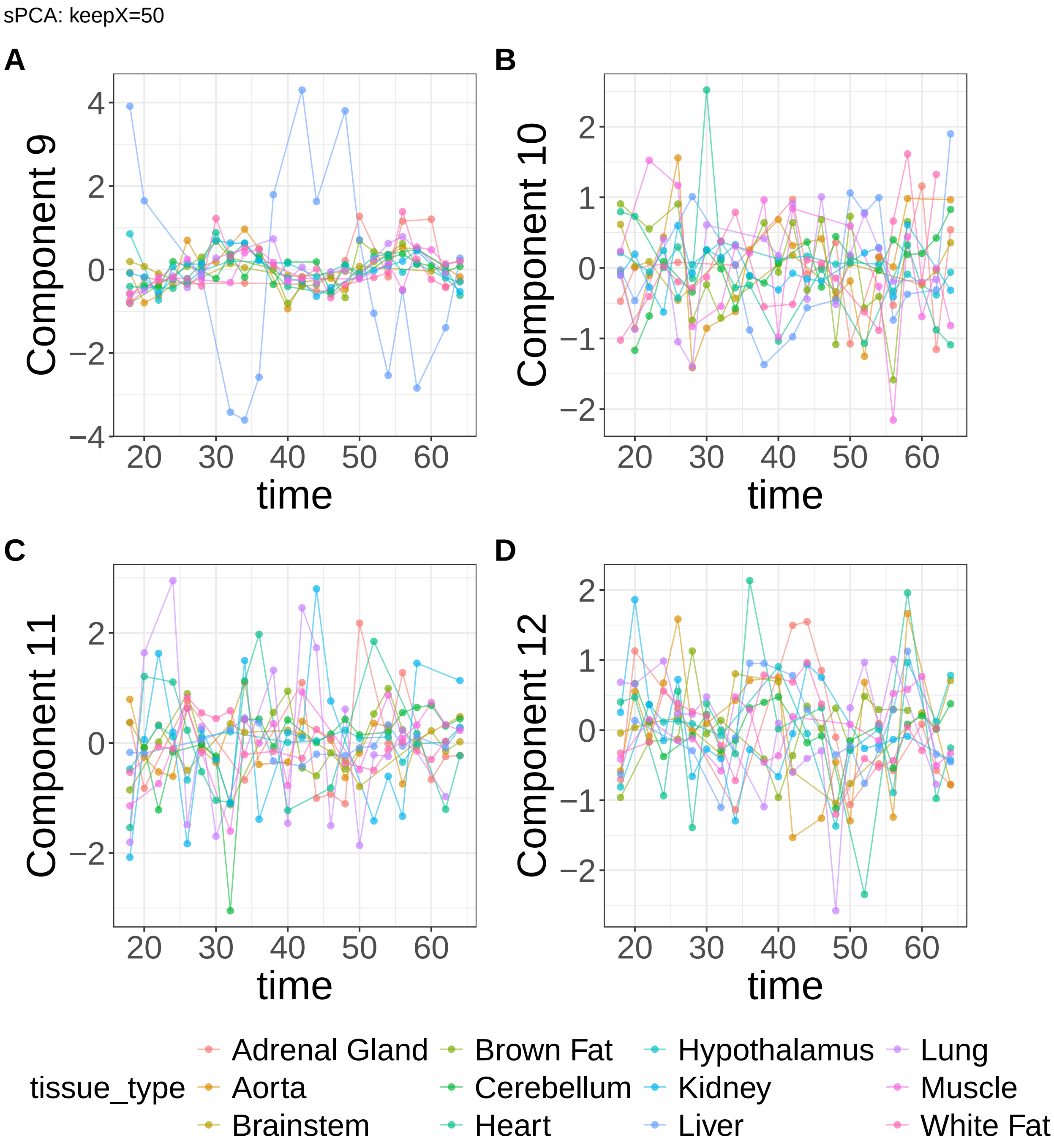


**Supplementary Figure 28.** Components for sPCA: keepX=50. (A) Component 9. (B) Component 10. (C) Component 11. (D) Component 12.

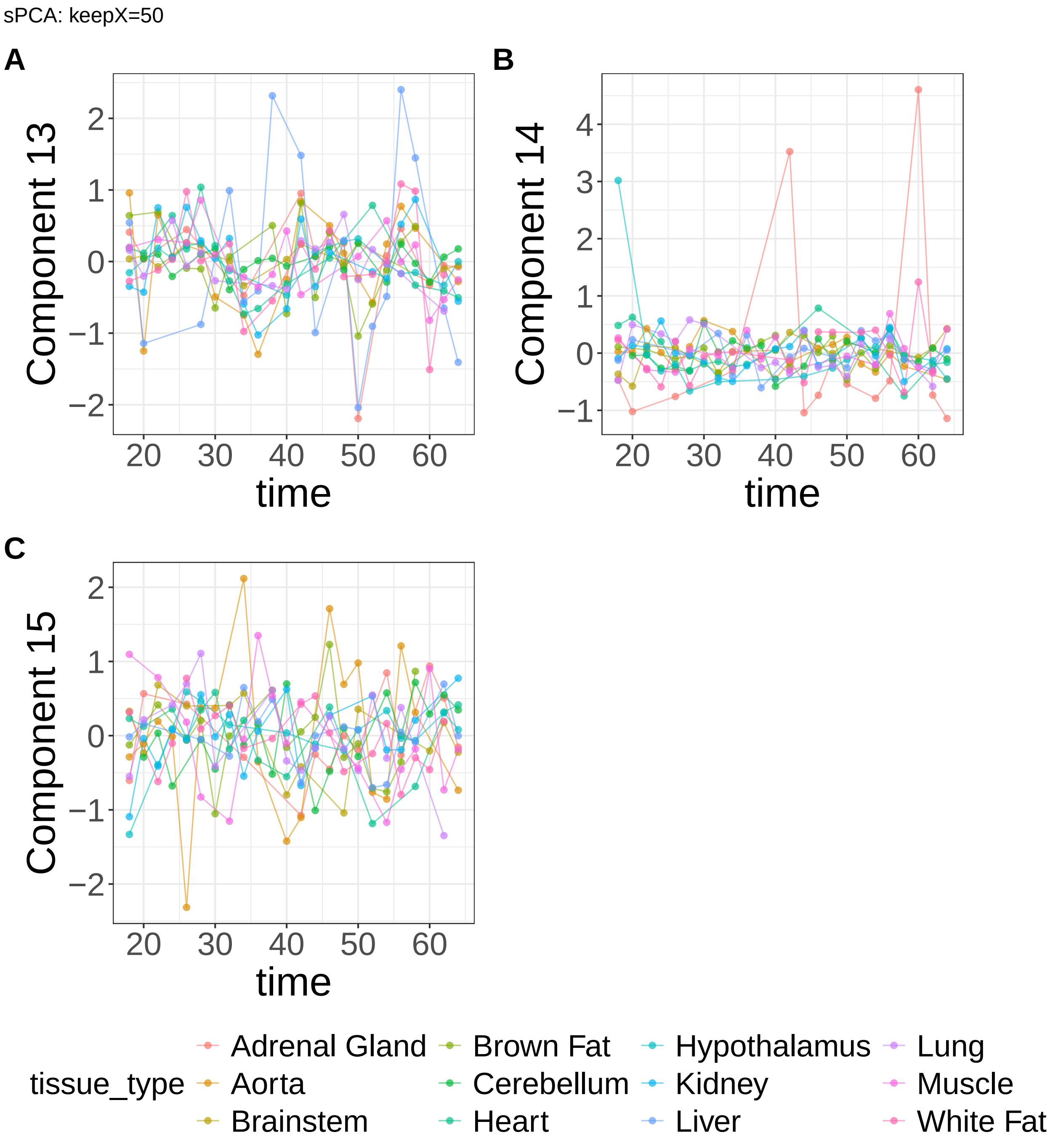


**Supplementary Figure 29.** Components for sPCA: keepX=50. (A) Component 13. (B) Component 14. (C) Component 15.

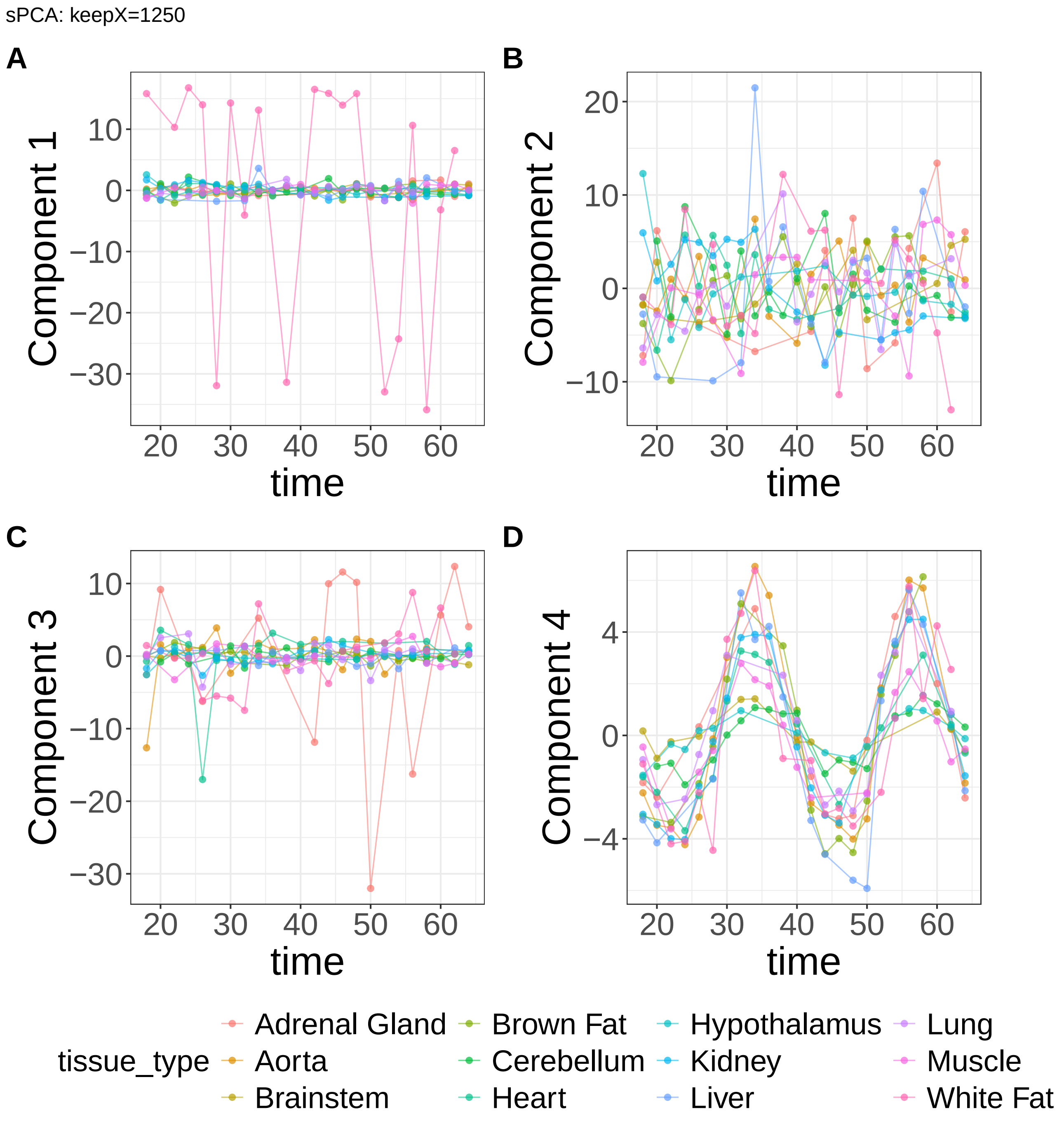


**Supplementary Figure 30.** Components for sPCA: keepX=1250. (A) Component 1. (B) Component 2. (C) Component 3. (D) Component 4.

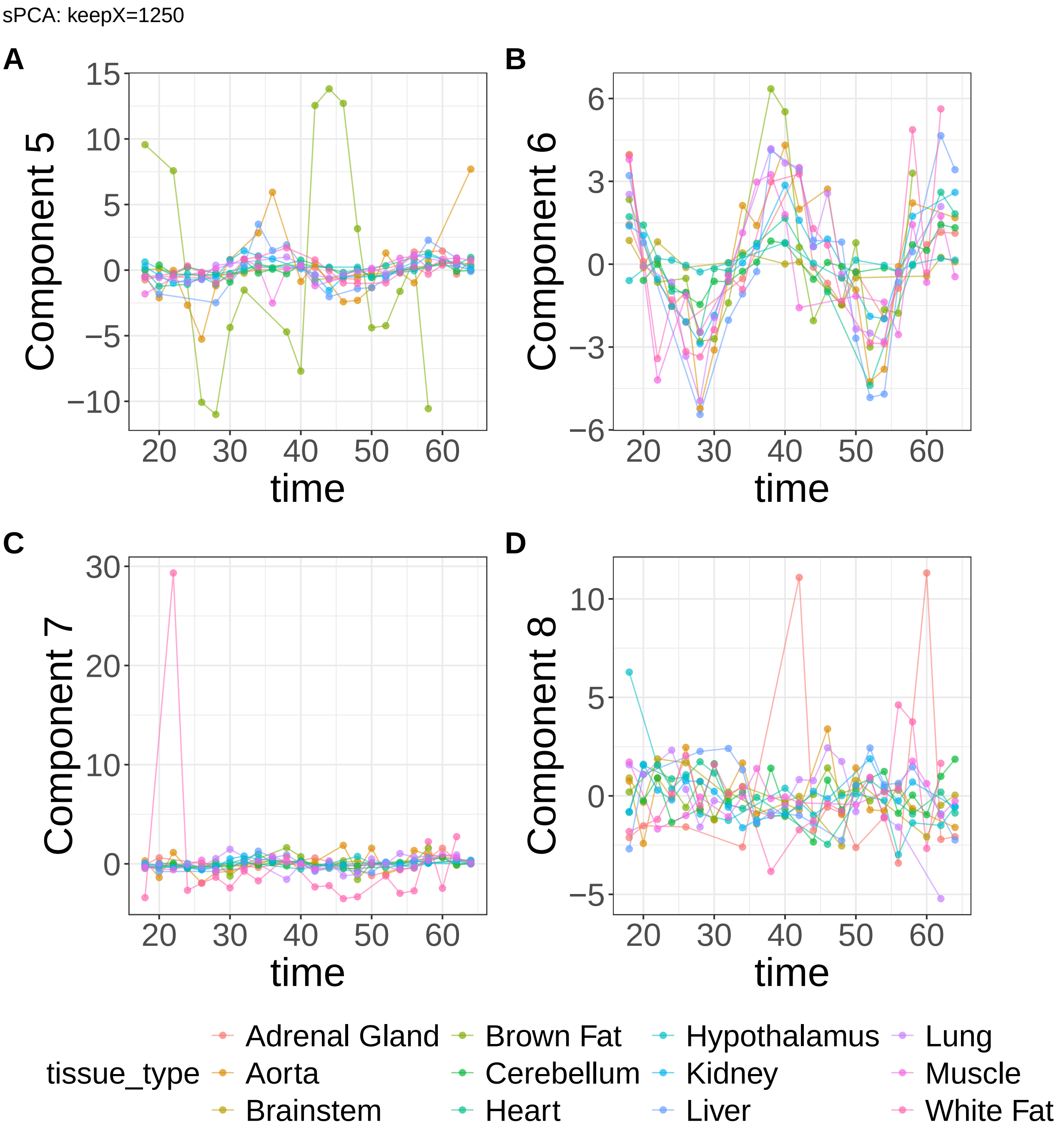


**Supplementary Figure 31.** Components for sPCA: keepX=1250. (A) Component 5. (B) Component 6. (C) Component 7. (D) Component 8.

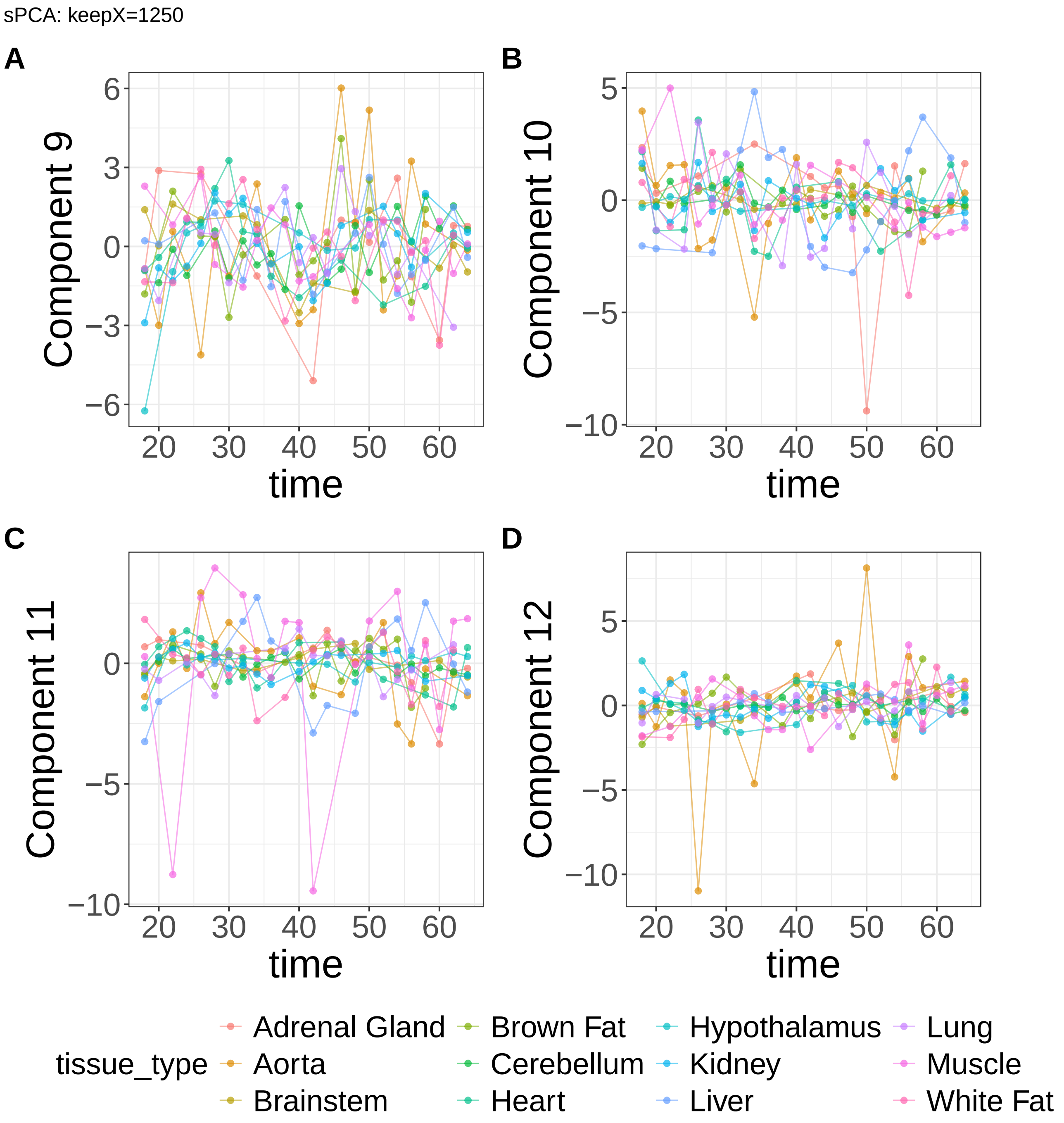


**Supplementary Figure 32.** Components for sPCA: keepX=1250. (A) Component 9. (B) Component 10. (C) Component 11. (D) Component 12.

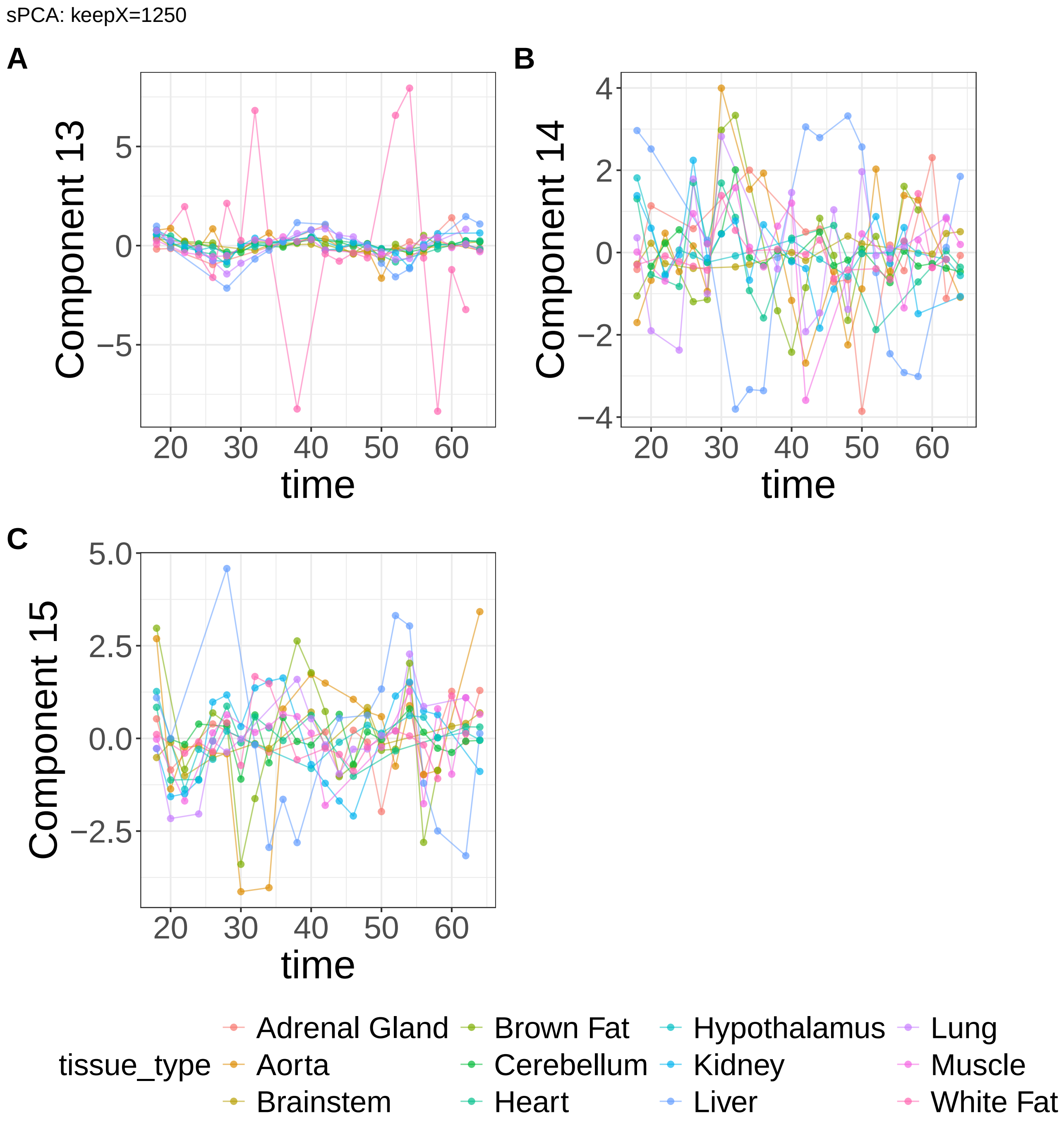


**Supplementary Figure 33.** Components for sPCA: keepX=1250. (A) Component 13. (B) Component 14. (C) Component 15.

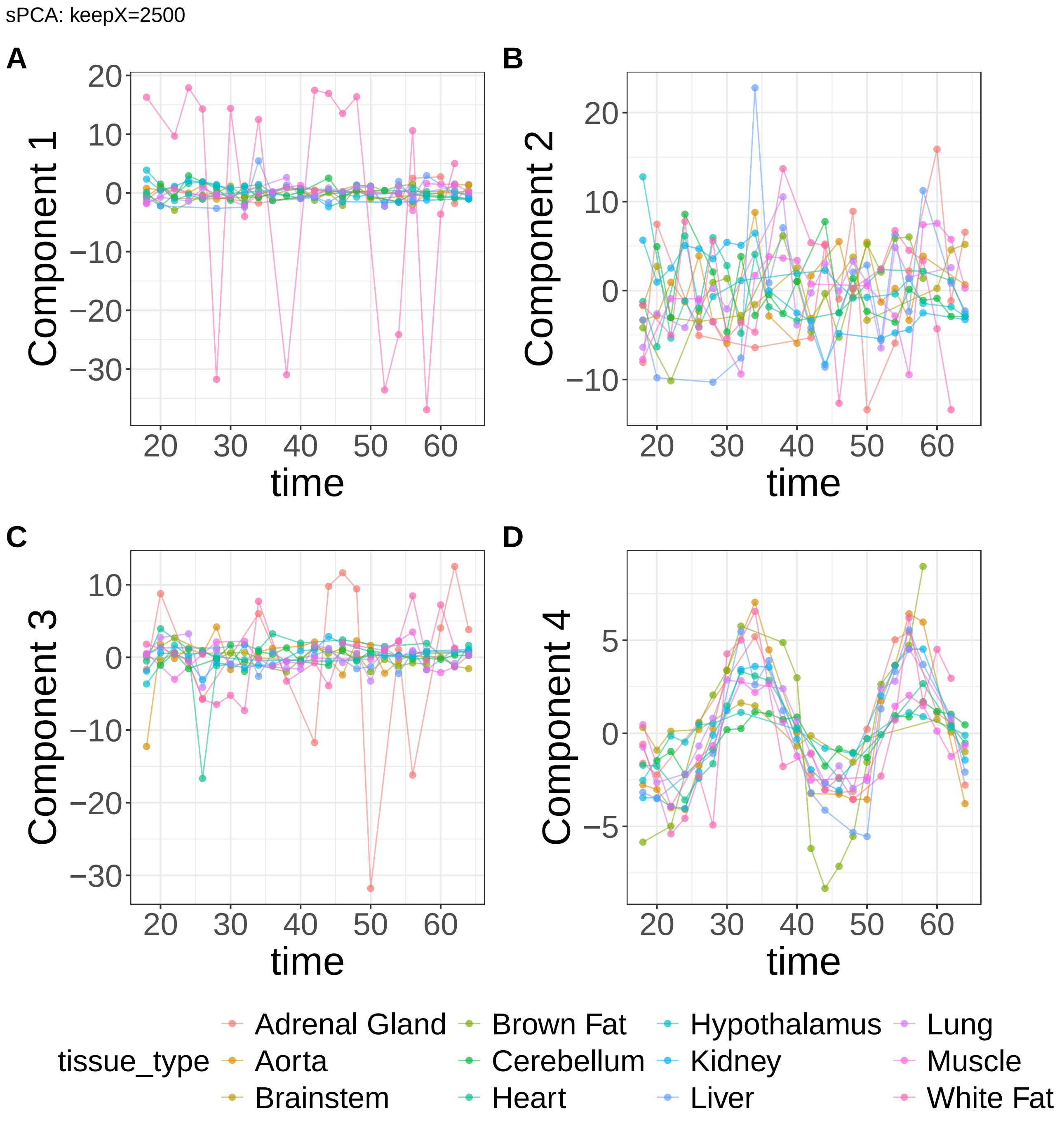


**Supplementary Figure 34.** Components for sPCA: keepX=2500. (A) Component 1. (B) Component 2. (C) Component 3. (D) Component 4.

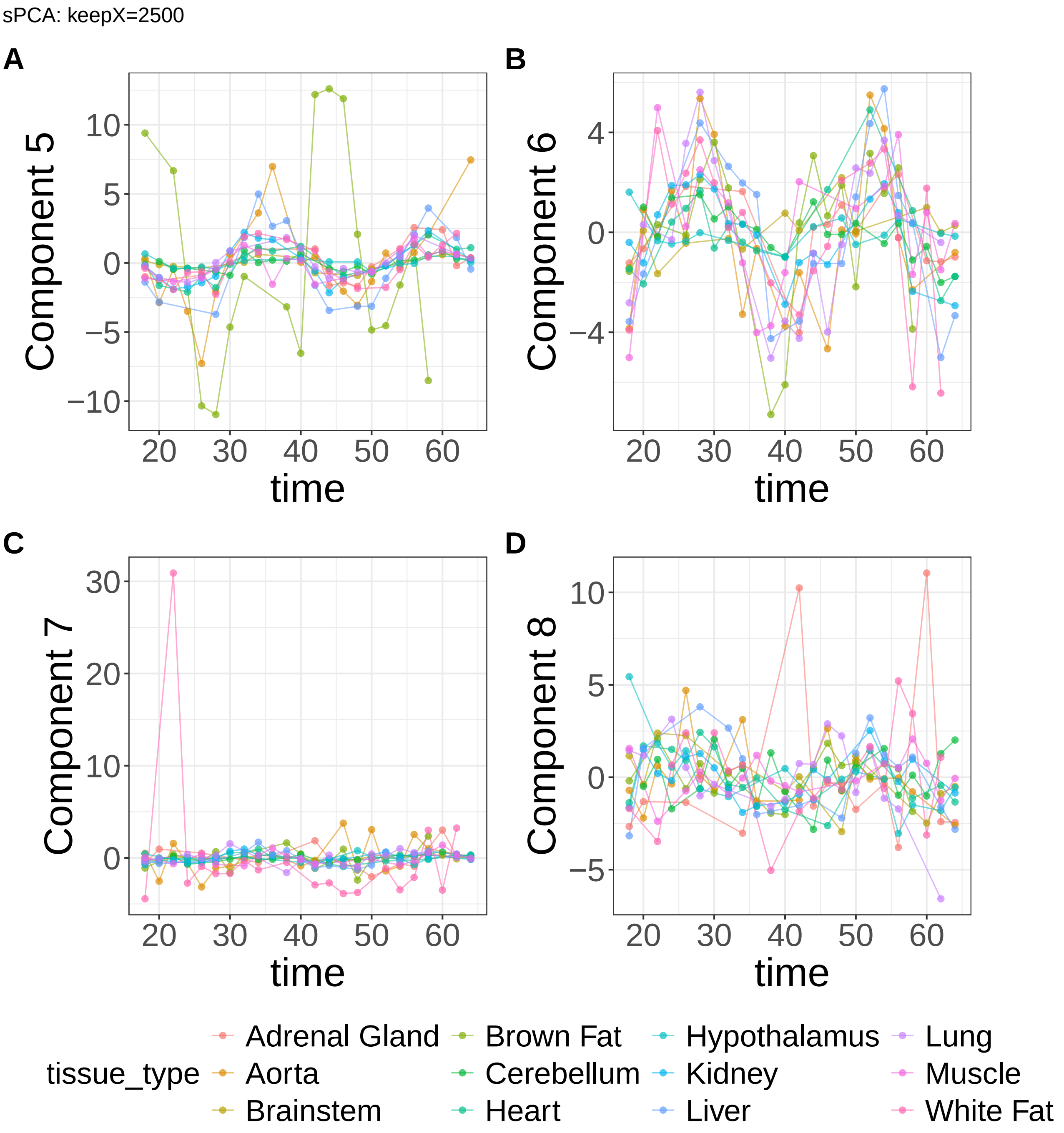


**Supplementary Figure 35.** Components for sPCA: keepX=2500. (A) Component 5. (B) Component 6. (C) Component 7. (D) Component 8.

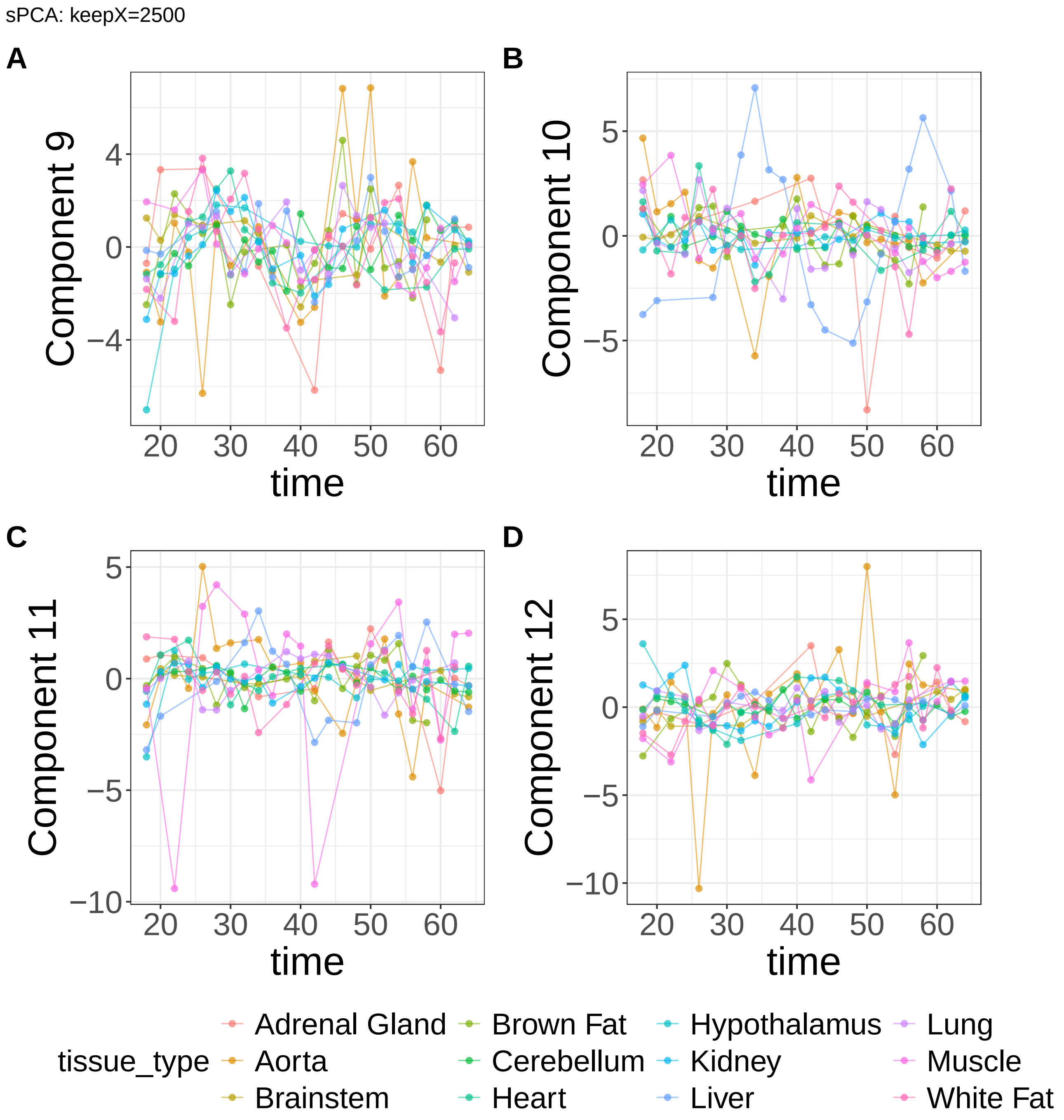


**Supplementary Figure 36.** Components for sPCA: keepX=2500. (A) Component 9. (B) Component 10. (C) Component 11. (D) Component 12.

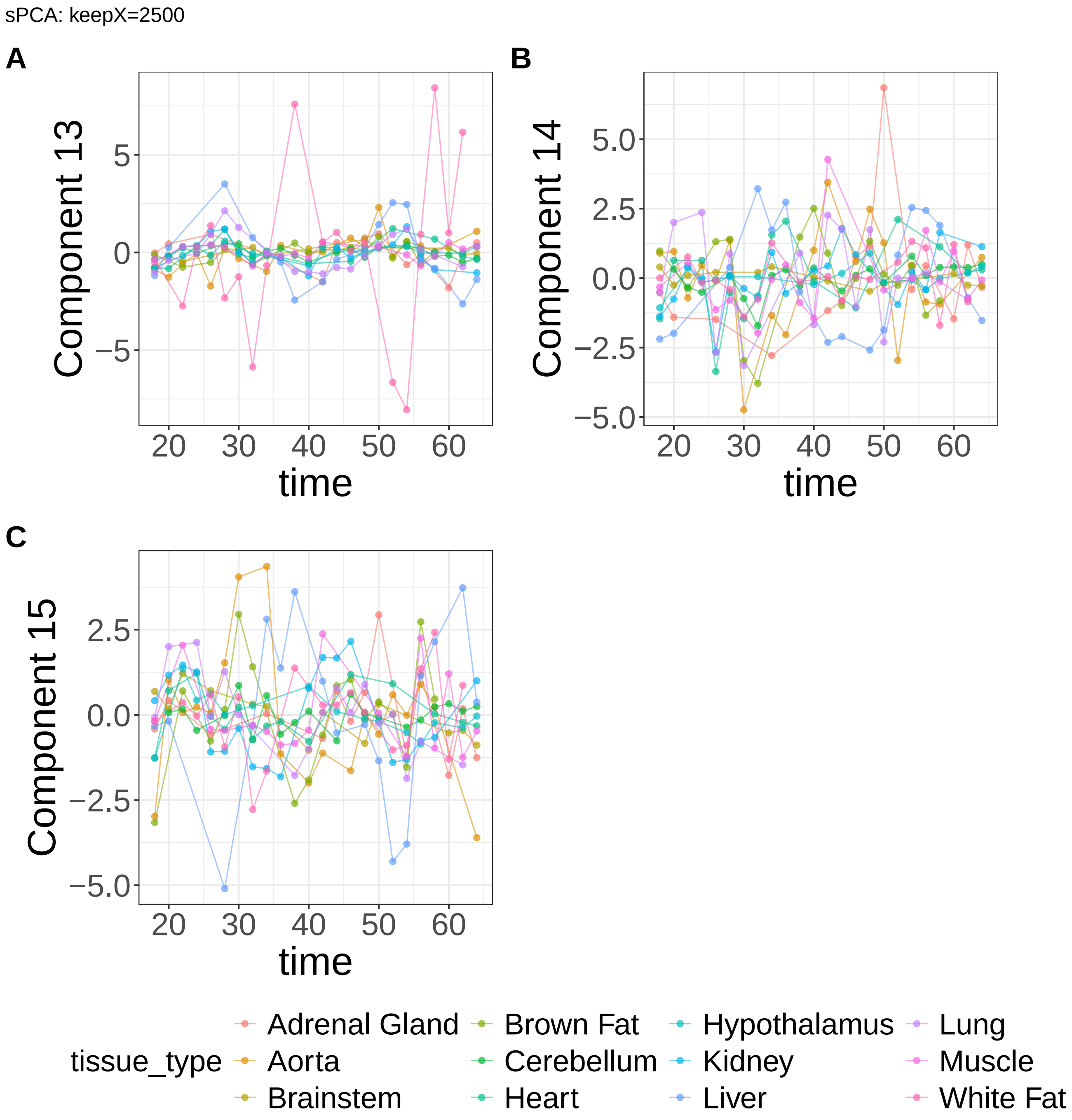


**Supplementary Figure 37.** Components for sPCA: keepX=2500. (A) Component 13. (B) Component 14. (C) Component 15.

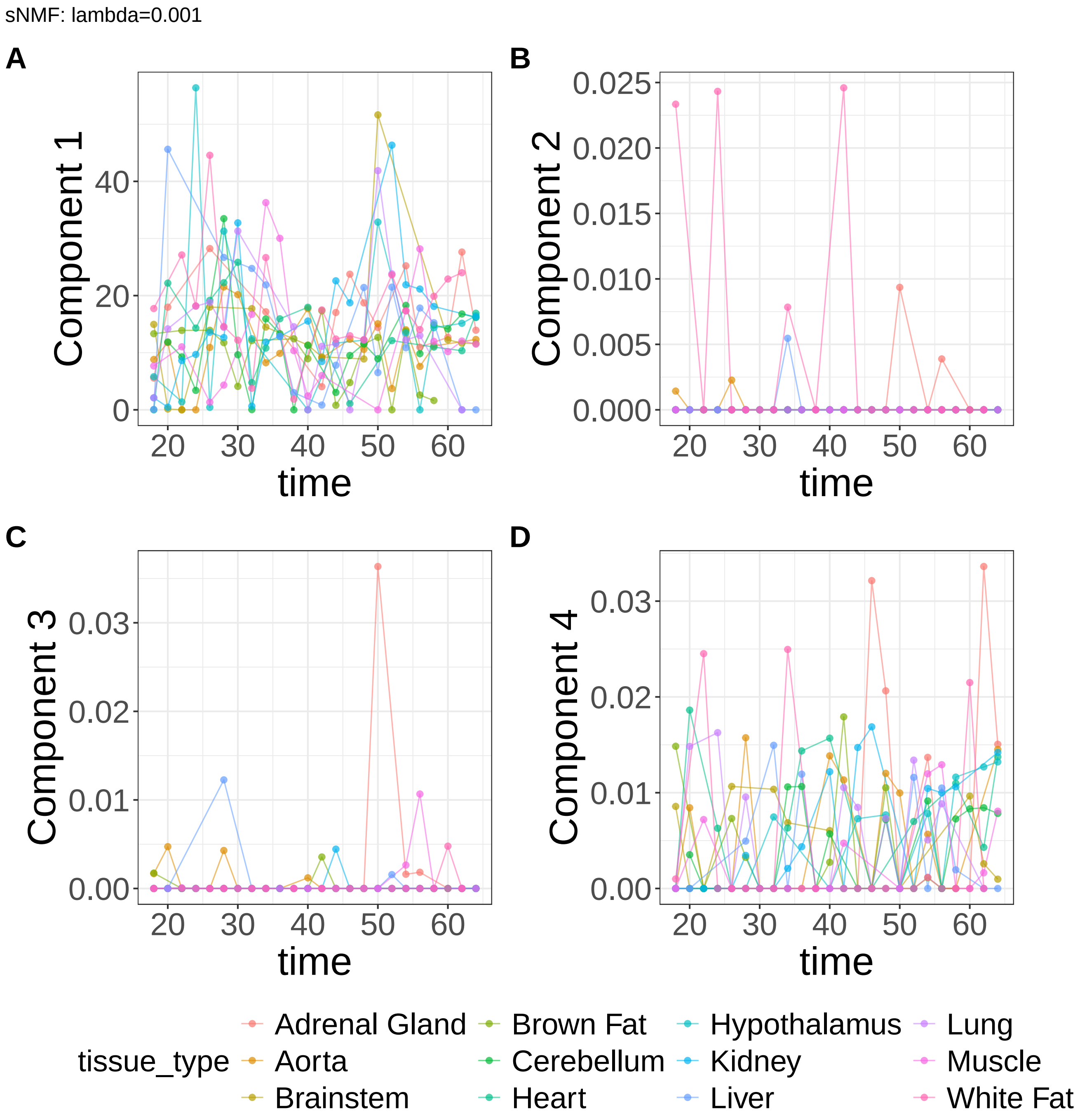


**Supplementary Figure 38.** Components for sNMF: lambda=0.001. (A) Component 1. (B) Component 2. (C) Component 3. (D) Component 4.

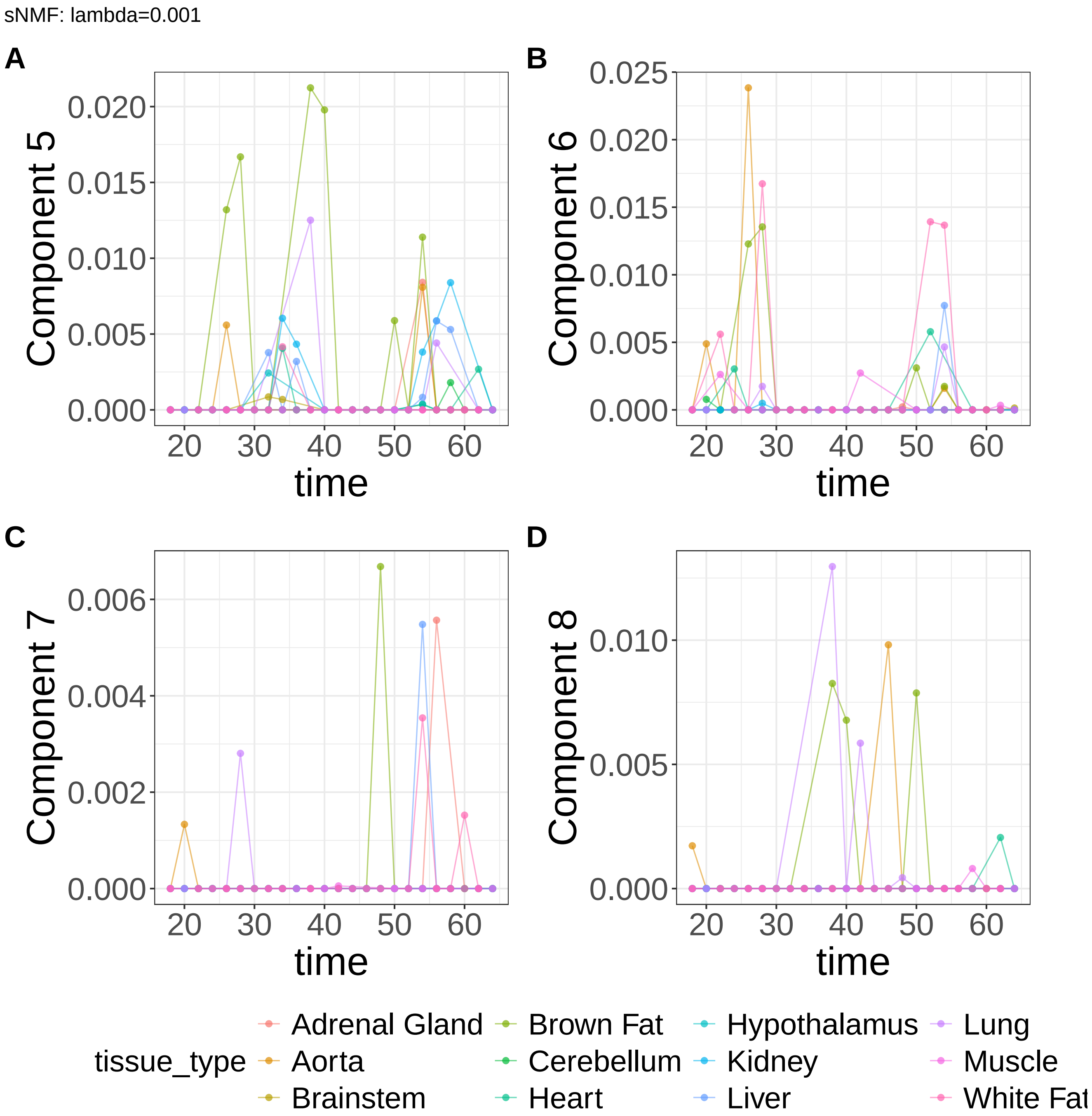


**Supplementary Figure 39.** Components for sNMF: lambda=0.001. (A) Component 5. (B) Component 6. (C) Component 7. (D) Component 8.

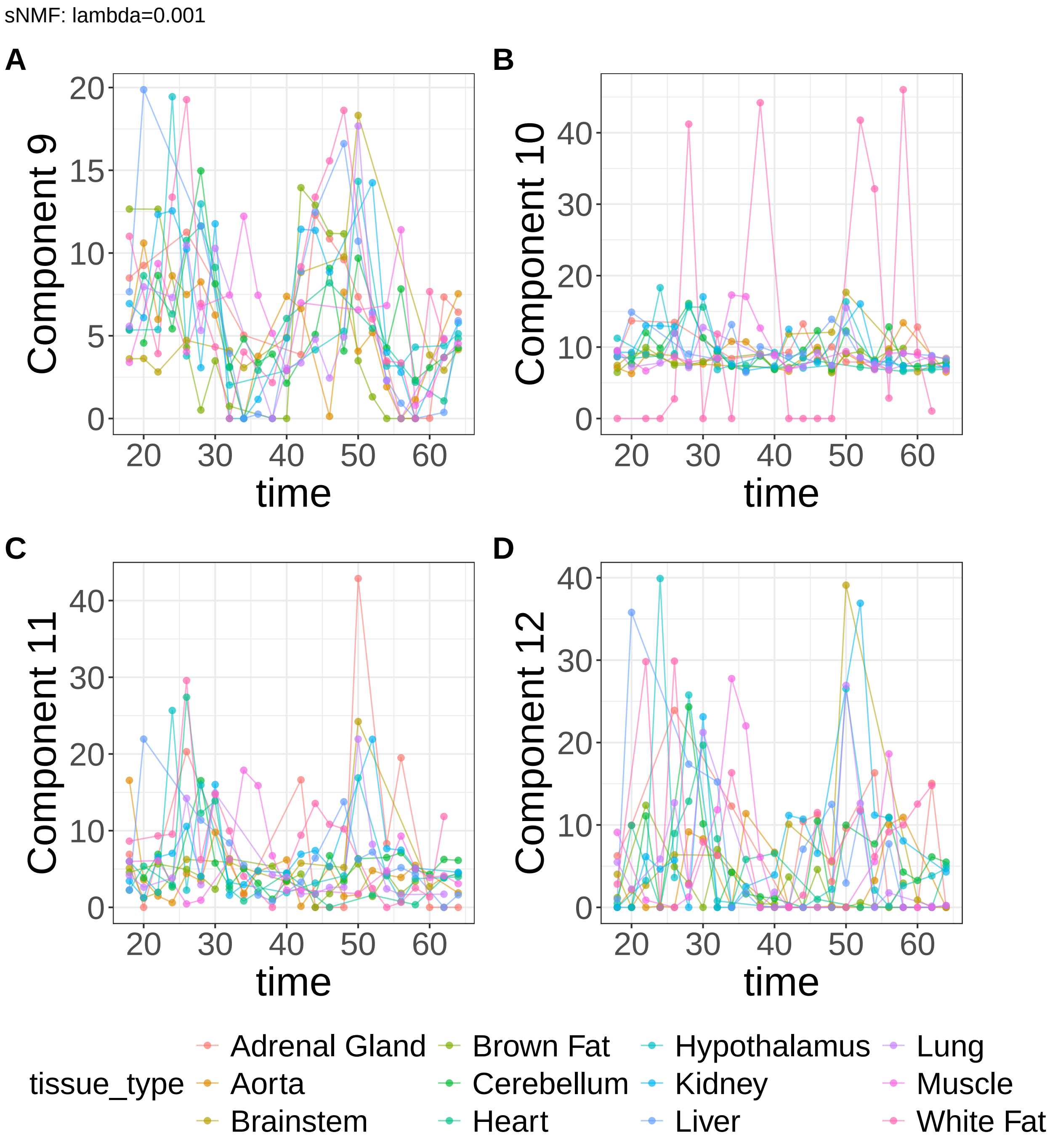


**Supplementary Figure 40.** Components for sNMF: lambda=0.001. (A) Component 9. (B) Component 10. (C) Component 11. (D) Component 12.

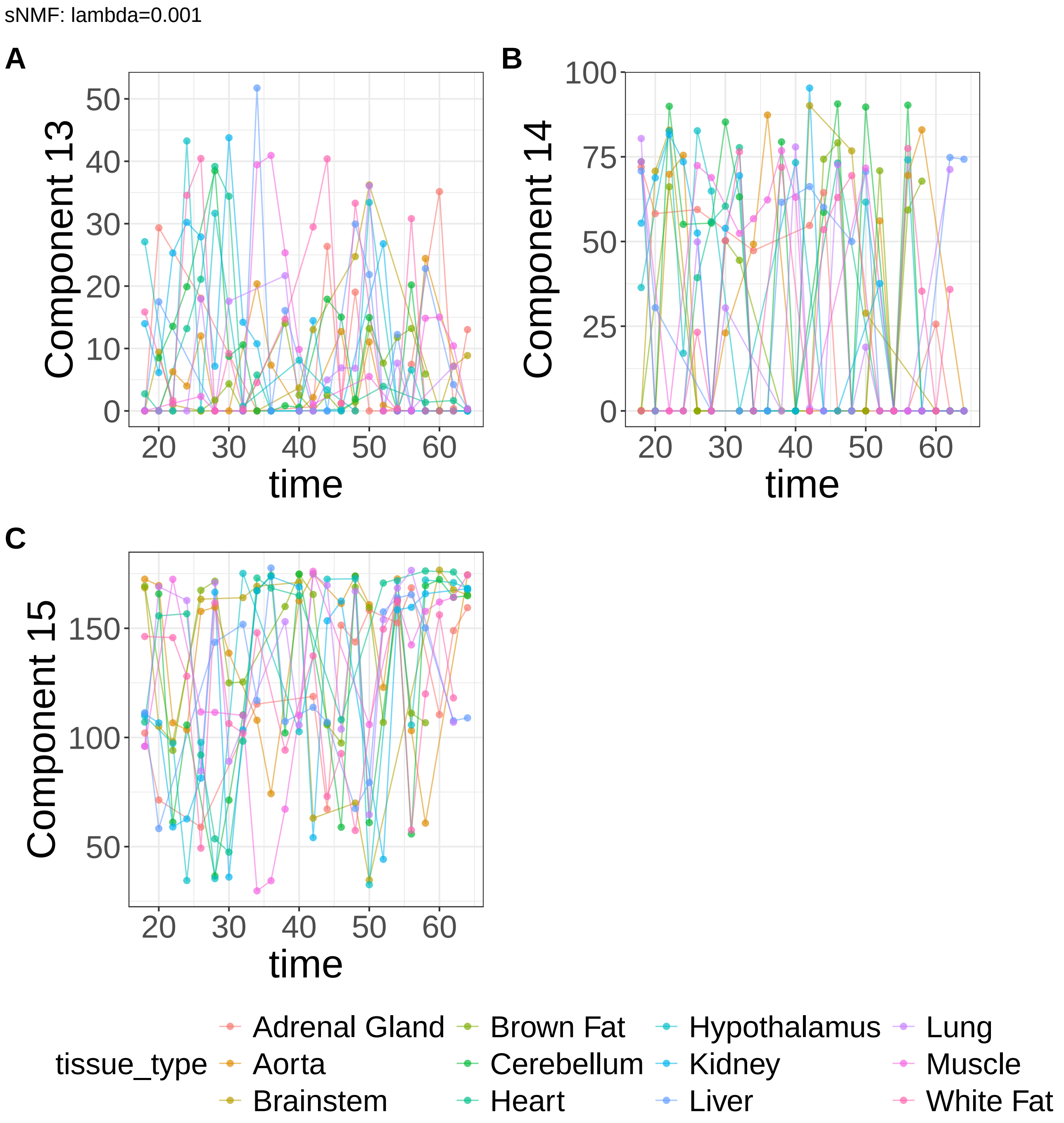


**Supplementary Figure 41.** Components for sNMF: lambda=0.001. (A) Component 13. (B) Component 14. (C) Component 15.

sNMF: lambda=0.01

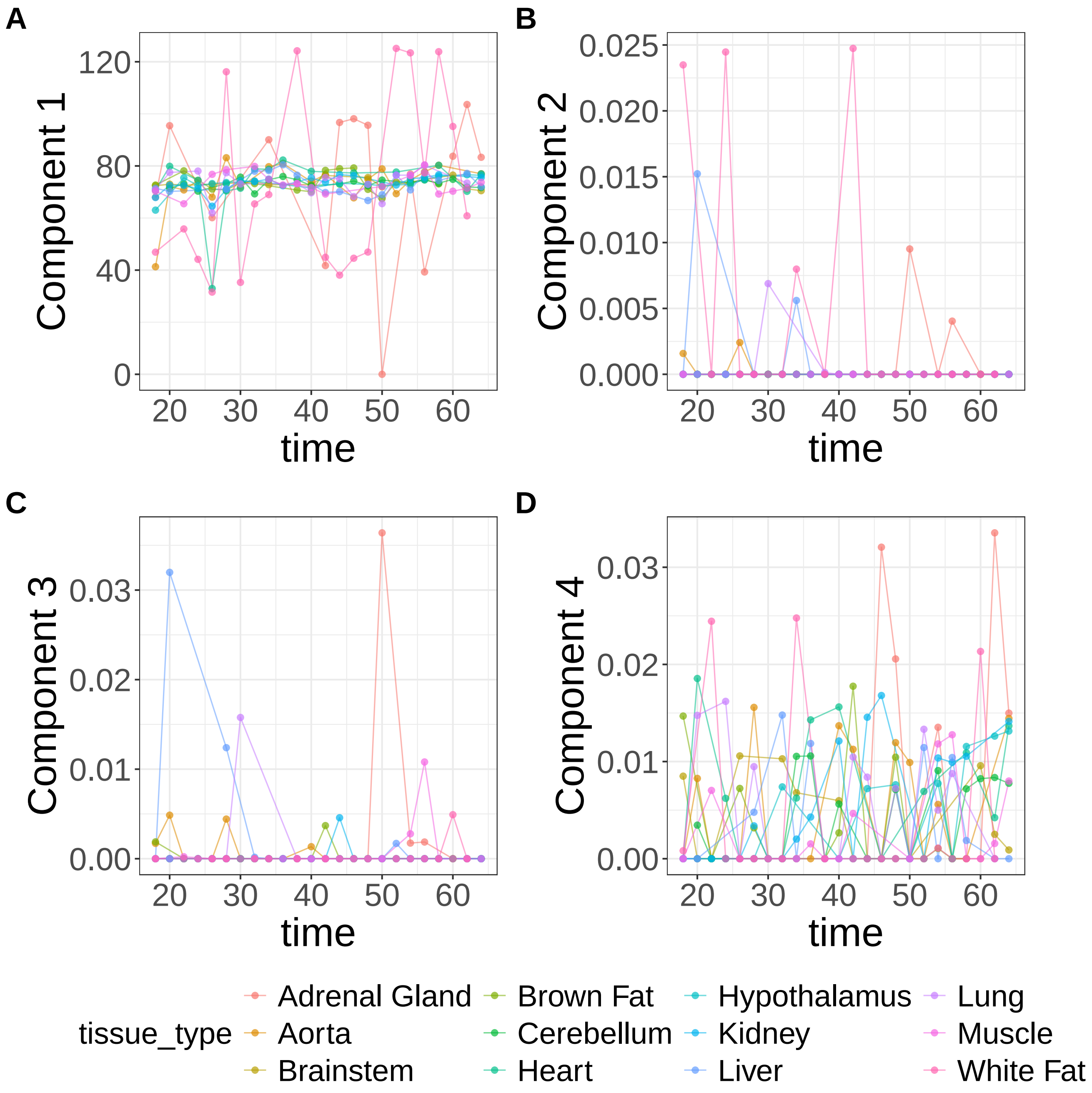


**Supplementary Figure 42.** Components for sNMF: lambda=0.01. (A) Component 1. (B) Component 2. (C) Component 3. (D) Component 4.

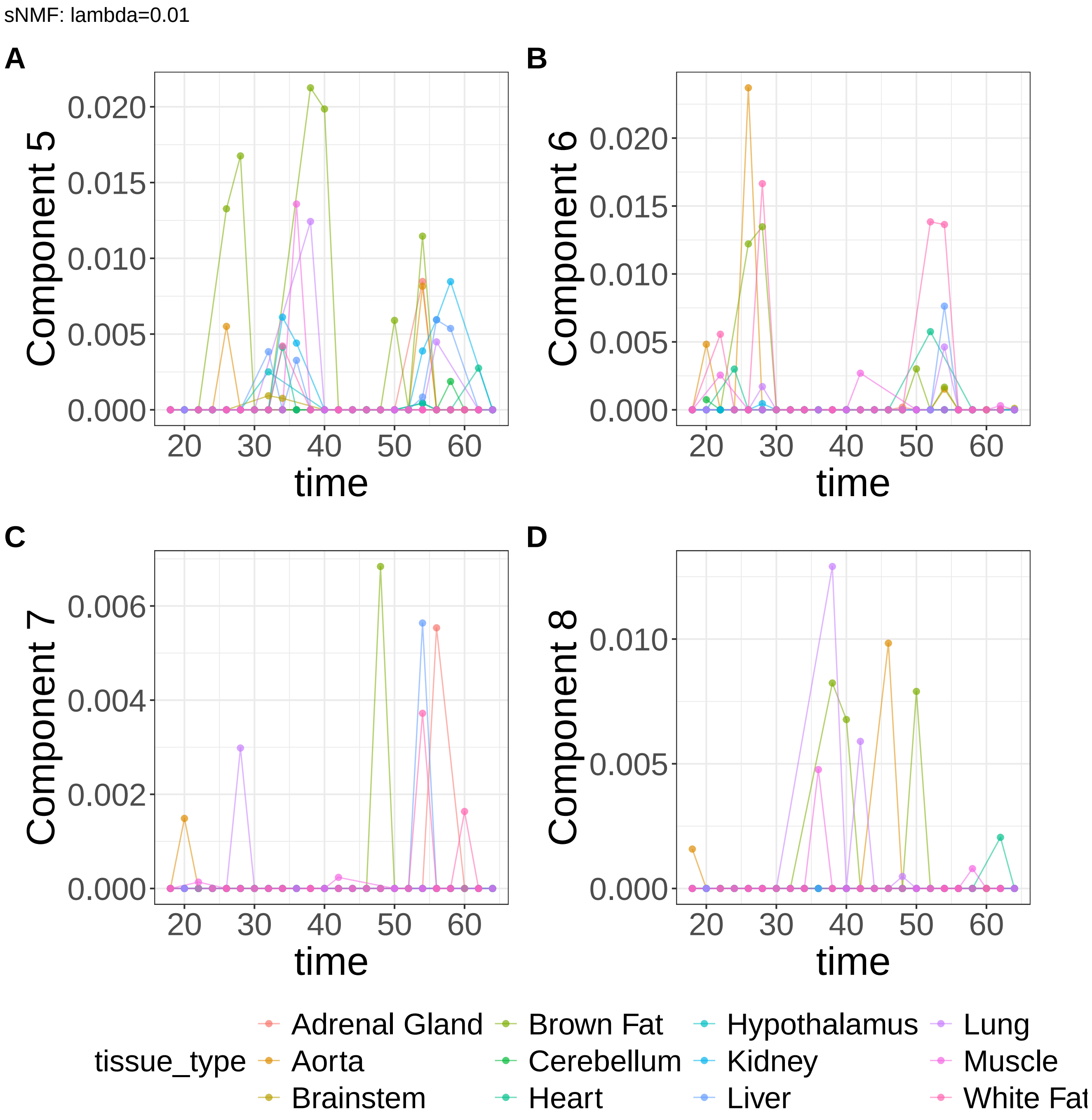


**Supplementary Figure 43.** Components for sNMF: lambda=0.01. (A) Component 5. (B) Component 6. (C) Component 7. (D) Component 8.

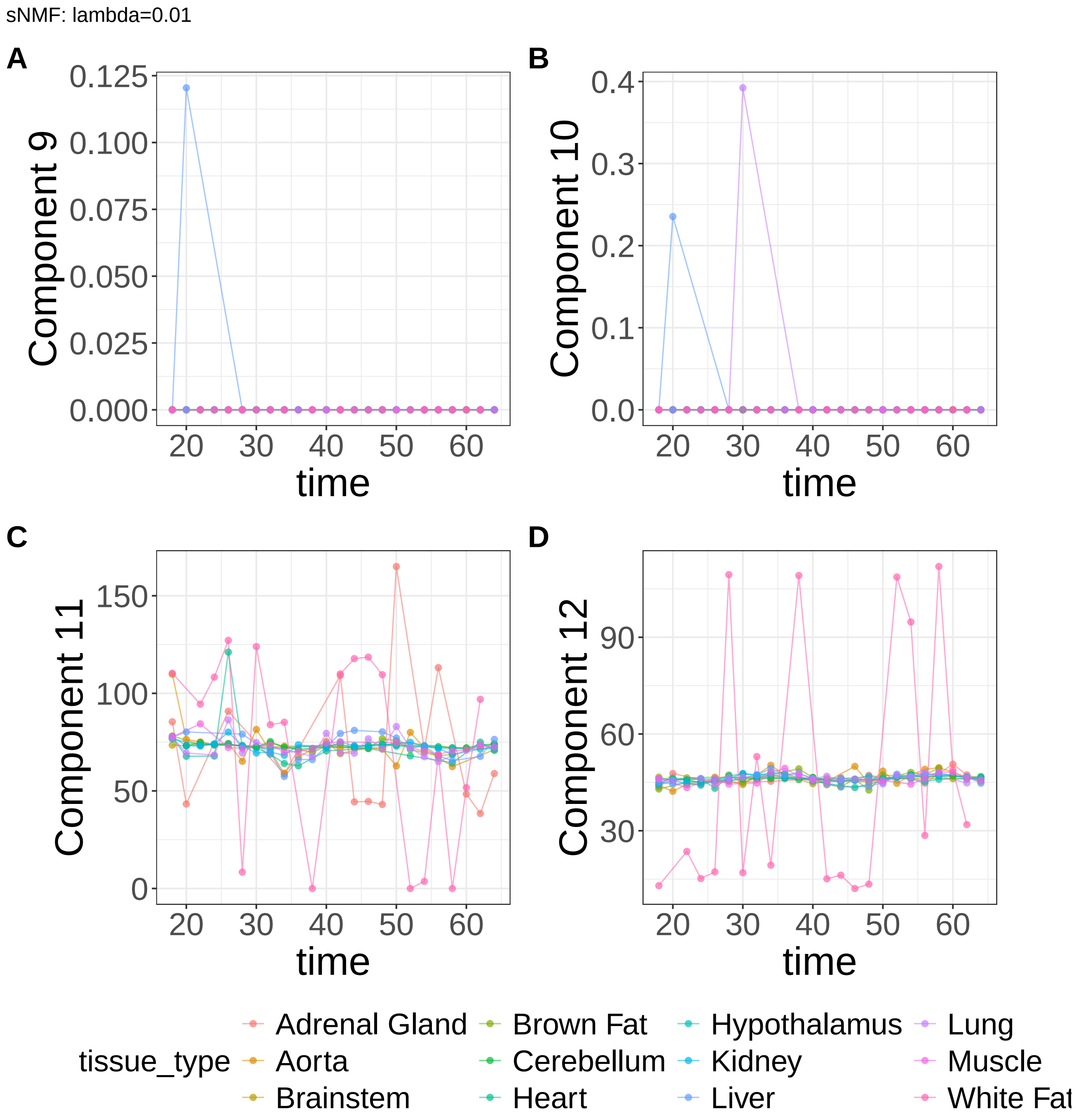


**Supplementary Figure 44.** Components for sNMF: lambda=0.01. (A) Component 9. (B) Component 10. (C) Component 11. (D) Component 12.

sNMF: lambda=0.01

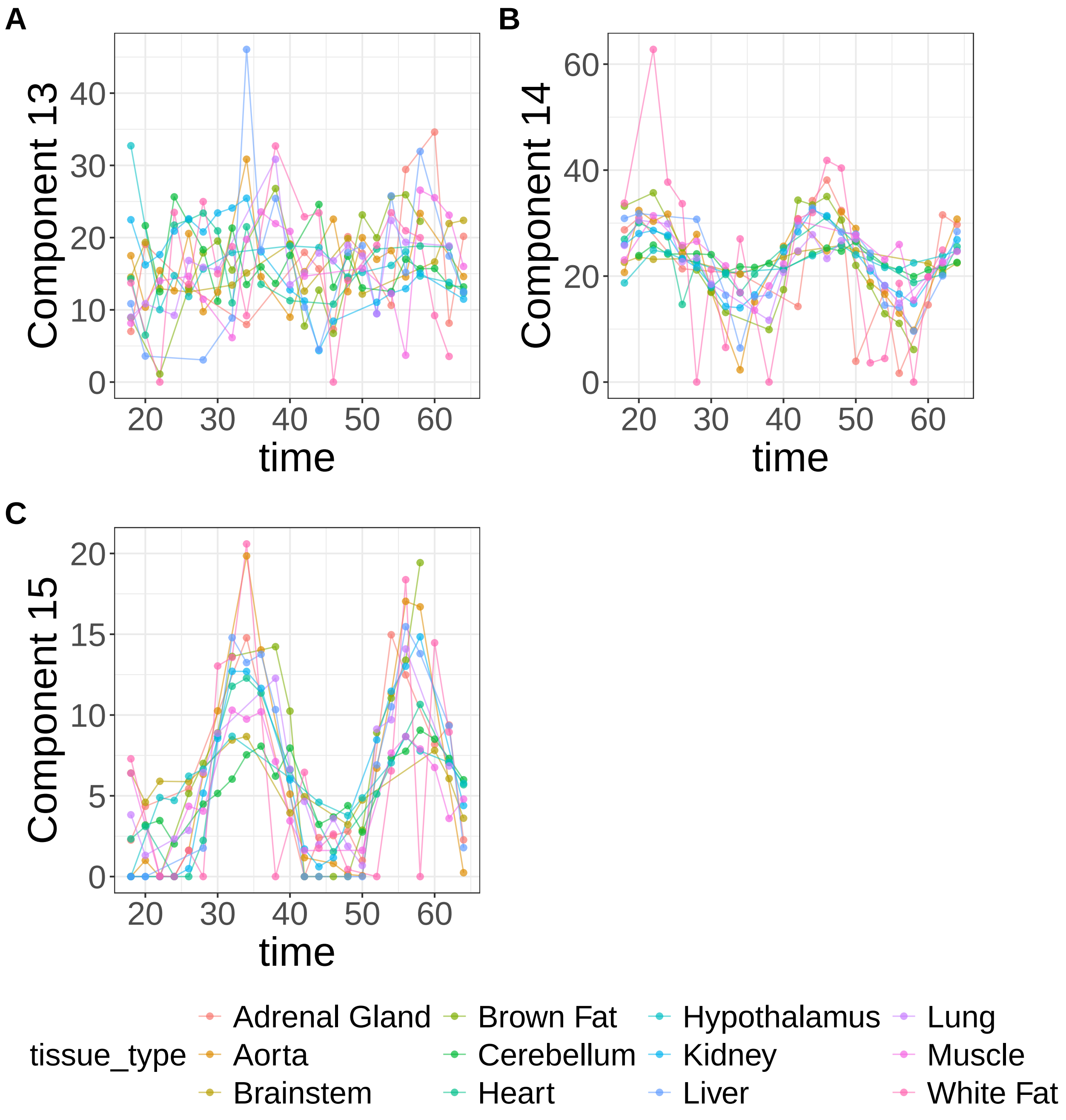


**Supplementary Figure 45.** Components for sNMF: lambda=0.01. (A) Component 13. (B) Component 14. (C) Component 15.

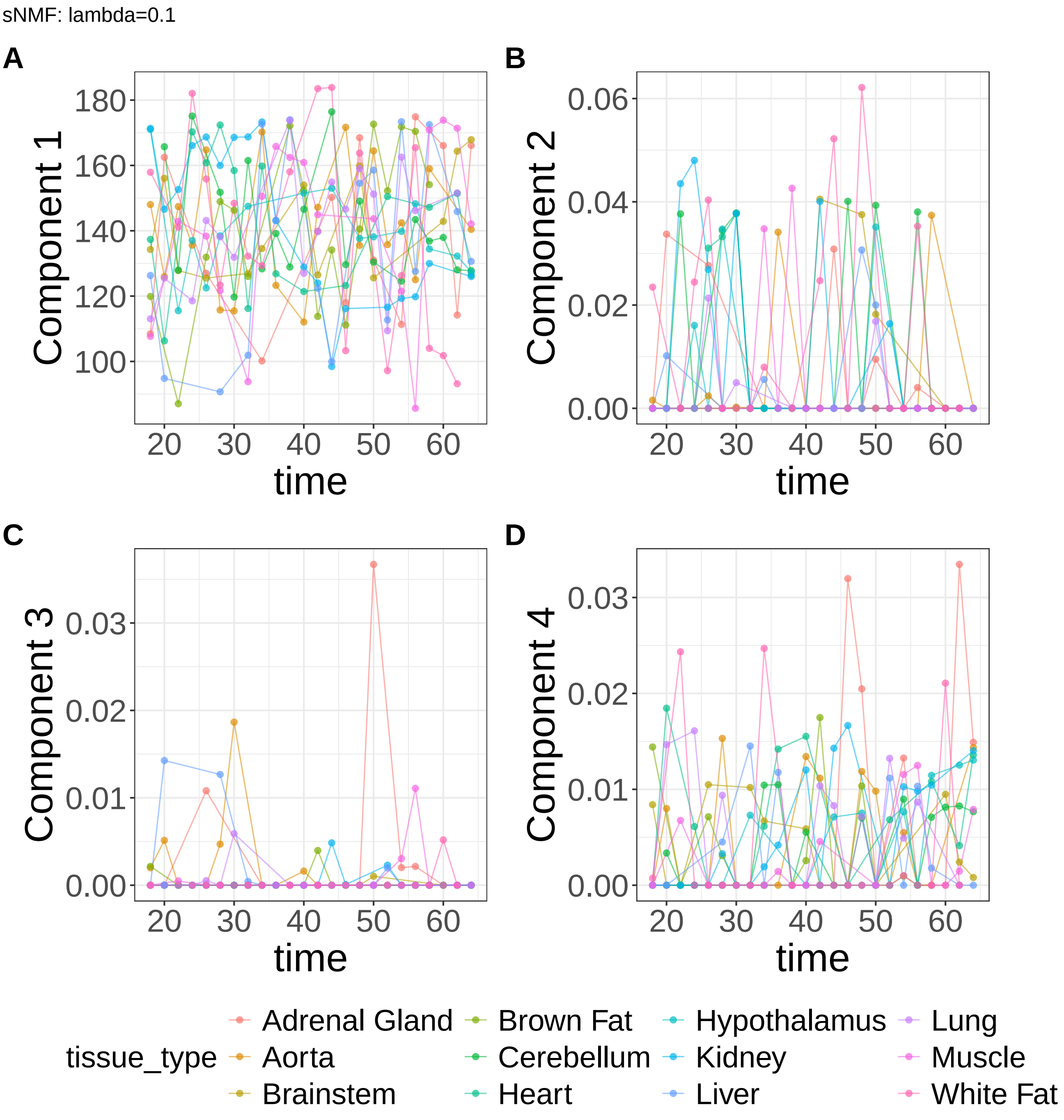


**Supplementary Figure 46.** Components for sNMF: lambda=0.1. (A) Component 1. (B) Component 2. (C) Component 3. (D) Component 4.

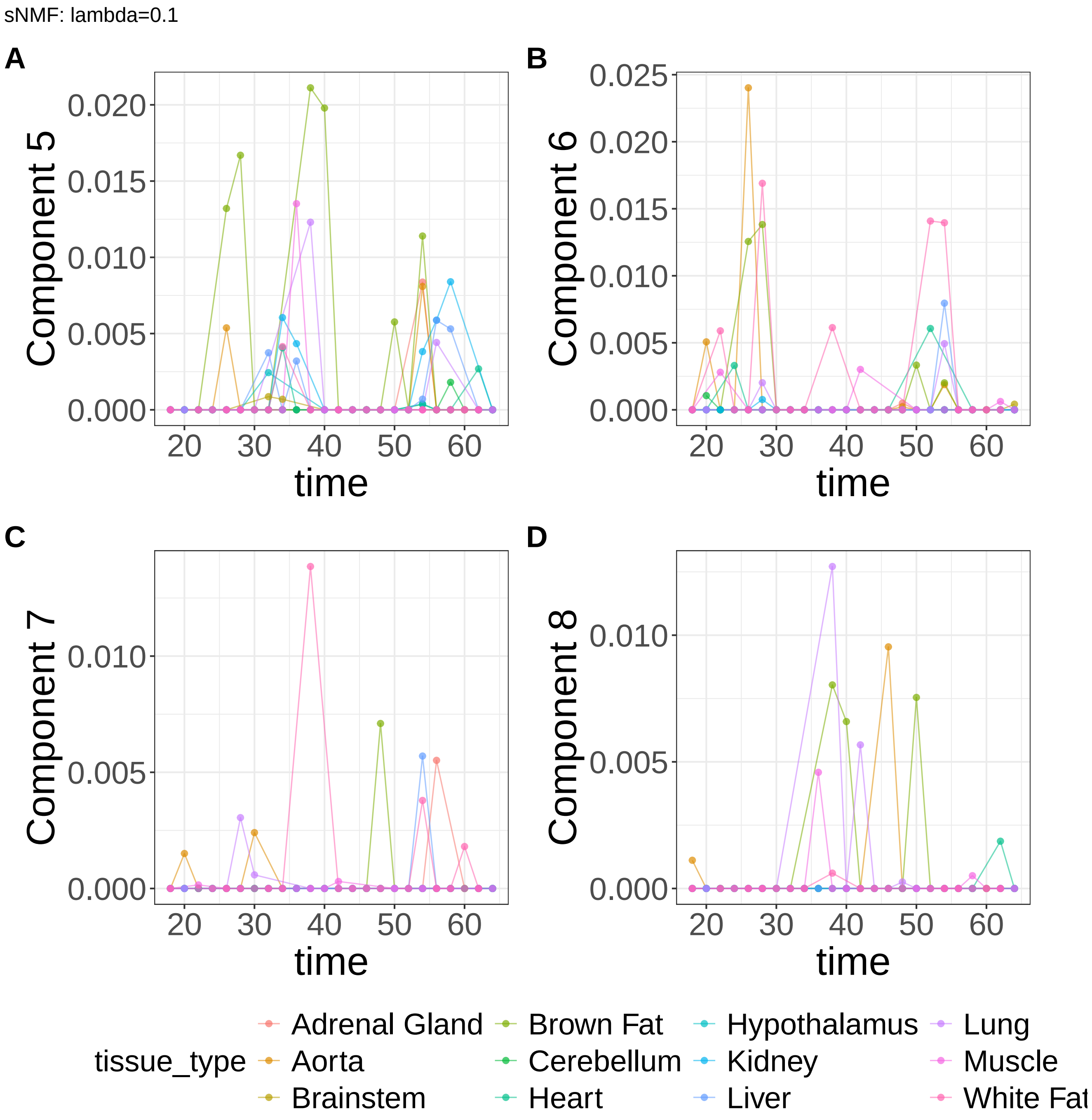


**Supplementary Figure 47.** Components for sNMF: lambda=0.1. (A) Component 5. (B) Component 6. (C) Component 7. (D) Component 8.

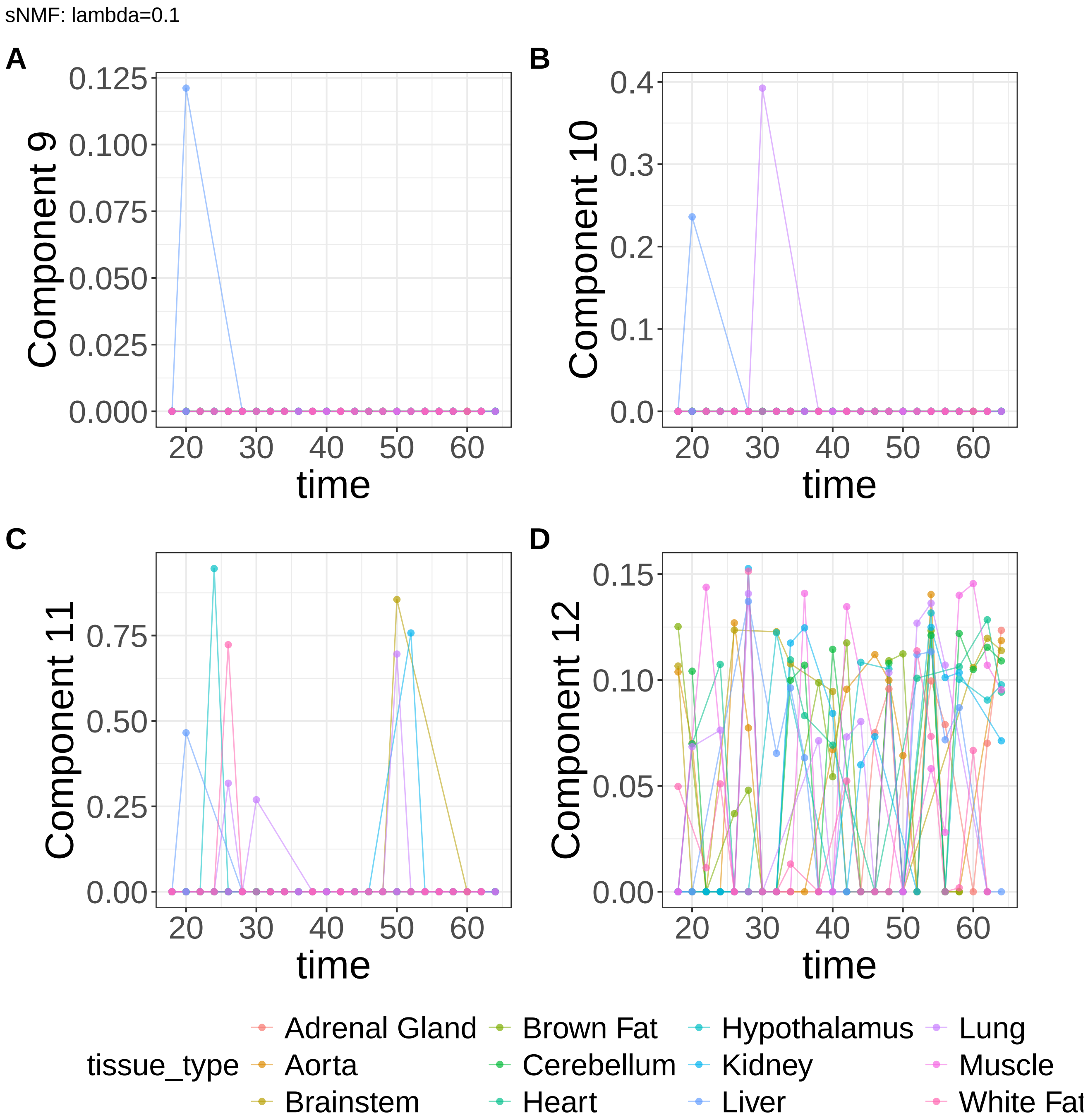


**Supplementary Figure 48.** Components for sNMF: lambda=0.1. (A) Component 9. (B) Component 10. (C) Component 11. (D) Component 12.

sNMF: lambda=0.1

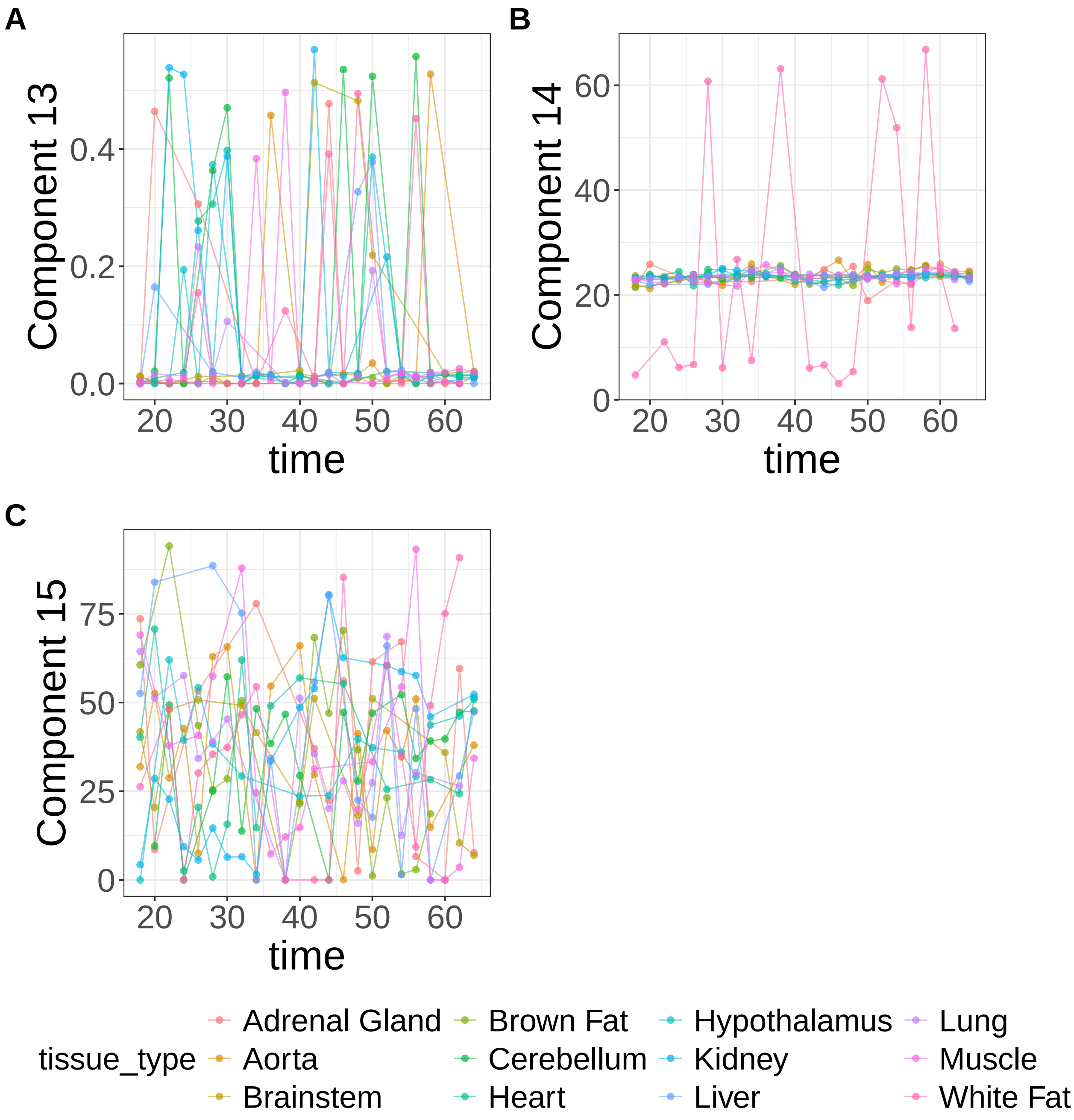


**Supplementary Figure 49.** Components for sNMF: lambda=0.1. (A) Component 13. (B) Component 14. (C) Component 15.

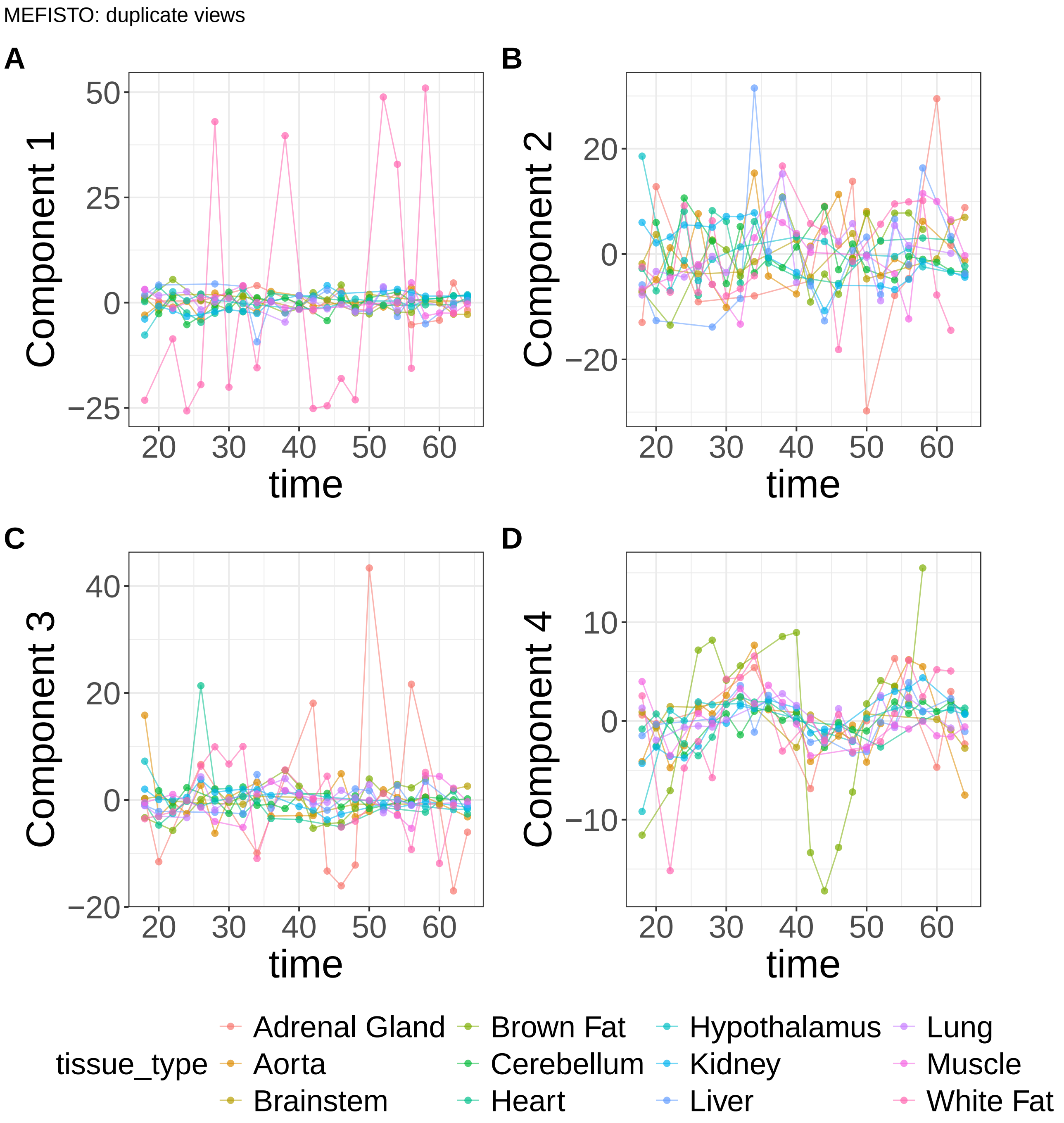


**Supplementary Figure 50.** Components for MEFISTO: duplicate views. (A) Component 1. (B) Component 2. (C) Component 3. (D) Component 4.

MEFISTO: duplicate views

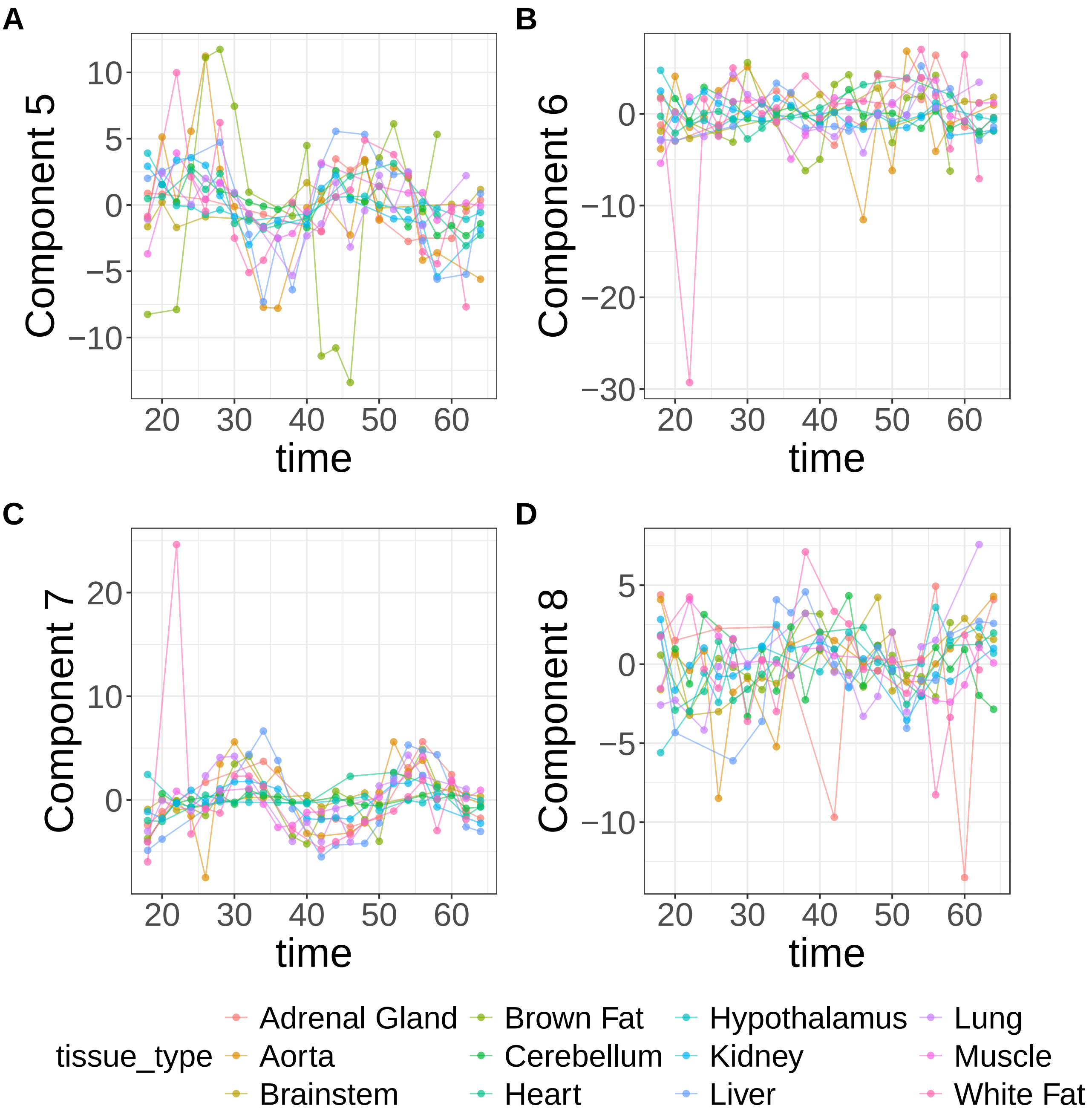


**Supplementary Figure 51.** Components for MEFISTO: duplicate views. (A) Component 5. (B) Component 6. (C) Component 7. (D) Component 8.

MEFISTO: duplicate views

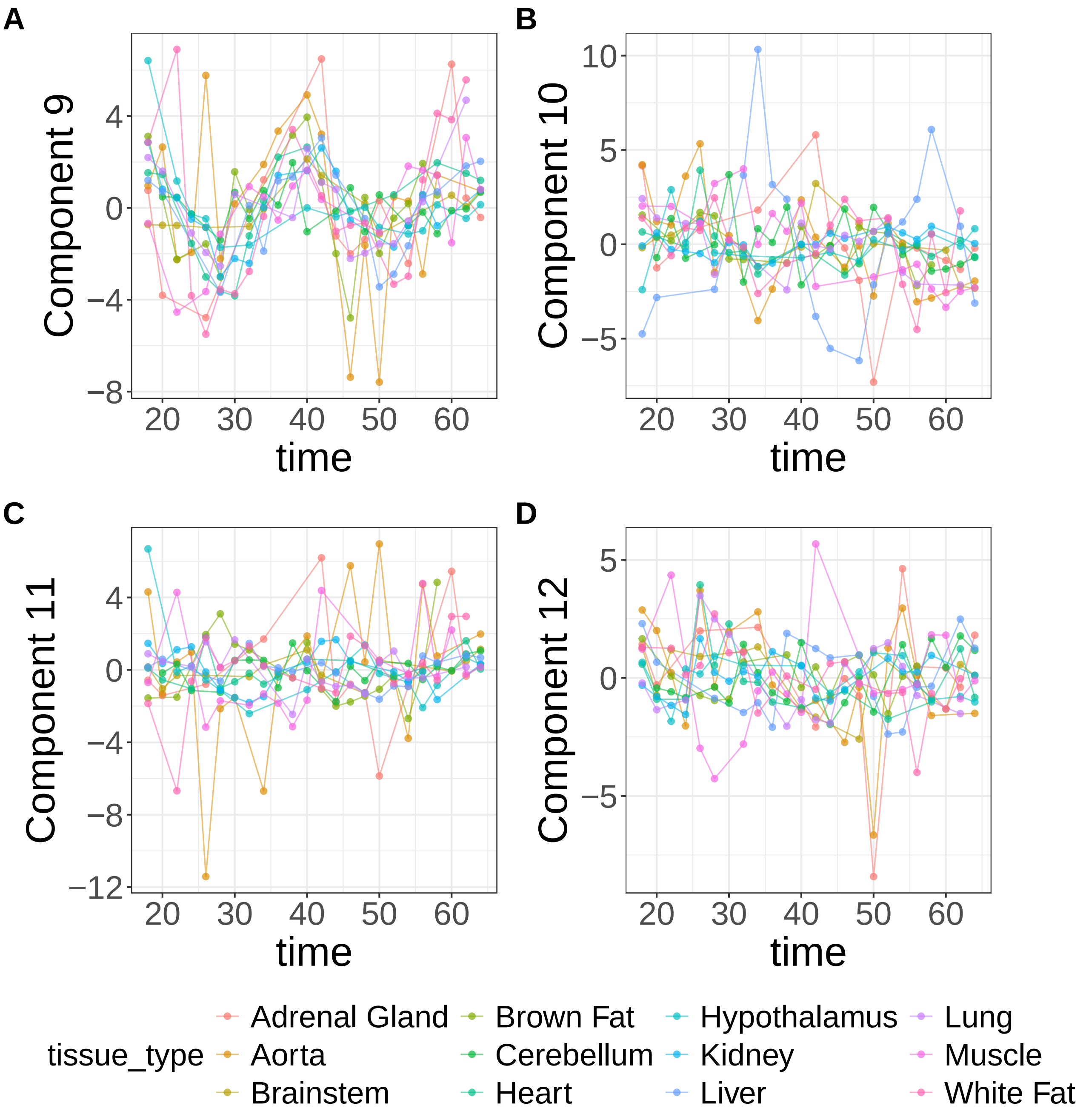


**Supplementary Figure 52.** Components for MEFISTO: duplicate views. (A) Component 9. (B) Component 10. (C) Component 11. (D) Component 12.

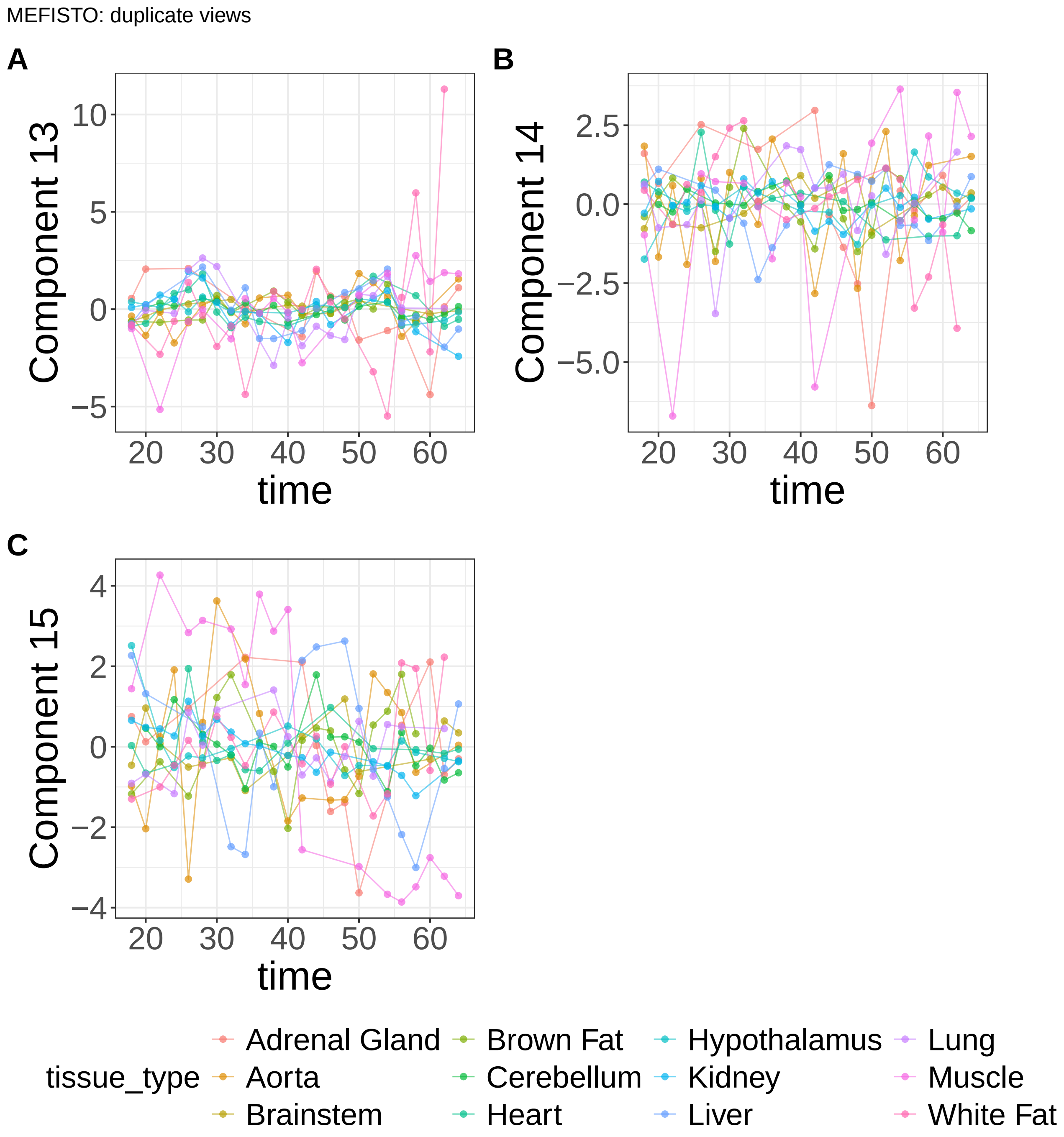


**Supplementary Figure 53.** Components for MEFISTO: duplicate views. (A) Component 13. (B) Component 14. (C) Component 15.
