## Supplementary Note 1 for "A statistical framework for system-level synthesis"

### SSANOVA-SVD coupled with the three-step framework generates reliable system-level synthesis in simulated systems

We have proposed a general three-step framework for system-level synthesis, and developed SSANOVA-SVD to build the system-level processes needed for this purpose. Our proposals, in part, exploit a natural factorization structure that arises by characterizing features as superimpositions of overarching processes/functions, which when evaluated at covariates, yield the component vectors. We therefore sought to assess the advantages of the proposed three-step framework, coupled with SSANOVA-SVD, over conventional matrix factorization methods for system-level synthesis. Unlike conventional matrix factorization methods, SSANOVA-SVD enables prediction of overarching processes in unseen samples from covariates alone and decomposition of the processes into smooth, potentially nonlinear covariate-specific contributions. Within the proposed three-step framework, these predictions further enable conditional inference of feature participation without data reuse. Consequently, prediction and inference metrics, as well as some assessments of interpretation, are available only for SSANOVA-SVD coupled with the three-step framework.

We simulated high-dimensional data containing multiple structured biological processes in the presence of nuisance variation to evaluate the proposed approach to synthesis (**Supplementary Methods: Simulations**). Briefly, samples were simulated over three relevant covariates—two continuous, *time* and *severity*, and one discrete, *group*—together with a nuisance covariate, *batch*. A fraction of features were designated as signal features whose abundances varied with time, severity, and group, while the remaining features were treated as biological nulls. Four biological processes were simulated, with signal features partitioned into four mutually disjoint modules, each participating in exactly one process. The first process induced group-specific shifts in the mean levels of its participating features. The remaining three processes were oscillatory with respect to time: an oscillation shared across groups, an oscillation with group-specific phase shifts, and an oscillation whose amplitude varied with sample-specific severity. Finally, nuisance variation due to batch was overlaid on the resulting biological signal. We varied biological and technical effect sizes and simulation settings to evaluate recovery of biological structure, separation from technical variation, and conditional inference of feature participation in relevant processes.

We evaluated the proposal along complementary aspects of system-level synthesis (**Supplementary Methods: Simulations**). We first assessed whether SSANOVA-SVD more accurately reconstructed the simulated biological signal than competing matrix factorization methods, while avoiding attribution of biological structure to null features. We next evaluated whether feature loadings prioritized signal features i.e., feature participating in the simulated biological processes rather than features associated with nuisance batch variation. Finally, we evaluated the three-step framework by determining whether the reconstructed processes captured the simulated dynamical phenomena, predicted biological signal in held-out data, and enabled conditional inference of feature participation with increasing power while controlling false discoveries. Detailed definitions of the benchmark metrics are provided in **Section 6** of the **Supplementary Methods: Simulations**.

We considered a challenging setting with 125 signal features among 5000 (2.5%). To avoid confounding performance metrics with the number of reconstructed components, all methods were evaluated at a common rank, $K$, chosen via cross-validation by SSANOVA-SVD. Benchmark metrics were therefore calculated using the first $K$ components from each method.

SSANOVA-SVD accurately reconstructed the simulated biological signal while avoiding nuisance variation. Reconstruction error was comparable to that of the best-performing matrix factorization method (**Supplementary Fig. 1A**). Conditional inference based on the reconstructed processes showed the expected increase in power with sample size while controlling type I error and maintaining empirical false discovery rates near the nominal 10% level (**Supplementary Fig. 1B**). The learned feature loadings more accurately distinguished features participating in the simulated biological processes from biological null features, while also separating biological from batch-associated variation (**Supplementary Fig. 1C**). These properties were largely preserved when biological group and batch assignments were partially confounded (**Supplementary Fig. 1** facets), indicating the advantage of jointly modeling nuisance variation. Performance remained robust in a more challenging setting in which batch effects were additionally introduced into a subset of biological signal features (**Supplementary Fig. 2**).

In a densely sampled regime of 100 samples per group (**Supplementary Figs. 3-9**), 2.5% signal features, the processes learned by SSANOVA-SVD explained more than 99.9% of the variation of the four simulated biological processes ($R^2 \geq 0.999$ for each process **Supplementary Fig. 5**A-D) and accurately predicted the simulated biological signals in the signal features in held-out samples ($R^2 = 0.994$) (**Supplementary Fig. 5E**). Decomposition of the component functions into their covariate-specific contributions and their interactions further revealed the simulated structure underlying the process space, distinguishing group-specific mean variation, shared temporal process, group-specific temporal processes, and severity-dependent amplification of temporal process (**Supplementary Fig. 6**-**Supplementary Fig. 9**).

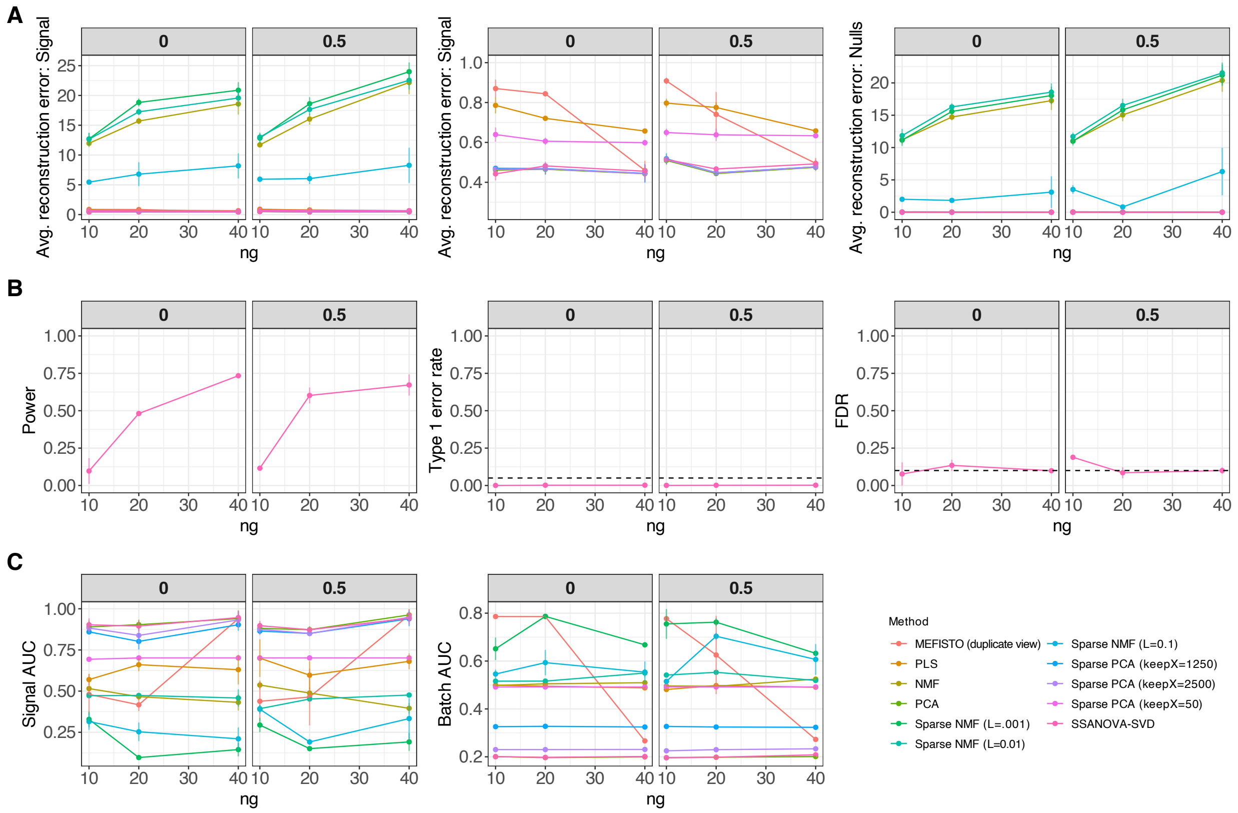


**Supplementary Figure 1: Benchmarking SSANOVA-SVD & the three-step framework in simulated data.** Methods were evaluated for increasing numbers of samples per biological group ($n_g$) under either no confounding (0) or moderate confounding (0.5) of group by batch. SSANOVA-SVD here refers to the implementation wrapping SSANOVA-SVD and the three-step framework. For Sparse NMF and Sparse PCA, the penalty parameters were additionally varied as indicated in the legend. A. Average reconstruction error for signal features (left), the same reconstruction error shown on a zoomed-in scale (middle), and null features (right). B. Power, type I error rate, and false discovery rate (FDR) of the proposed approach for conditional inference of feature participations in simulated relevant processes. Dashed horizontal lines indicate the nominal type I error rate of 0.05 and FDR level of 0.10. C. Area under the receiver operating characteristic curve (AUC) of the proposed approach for prioritizing relevant signal features (left) and irrelevant batch-affected features (middle) in feature loadings. Results are averaged over three simulation replicates. Standard error bars are shown.

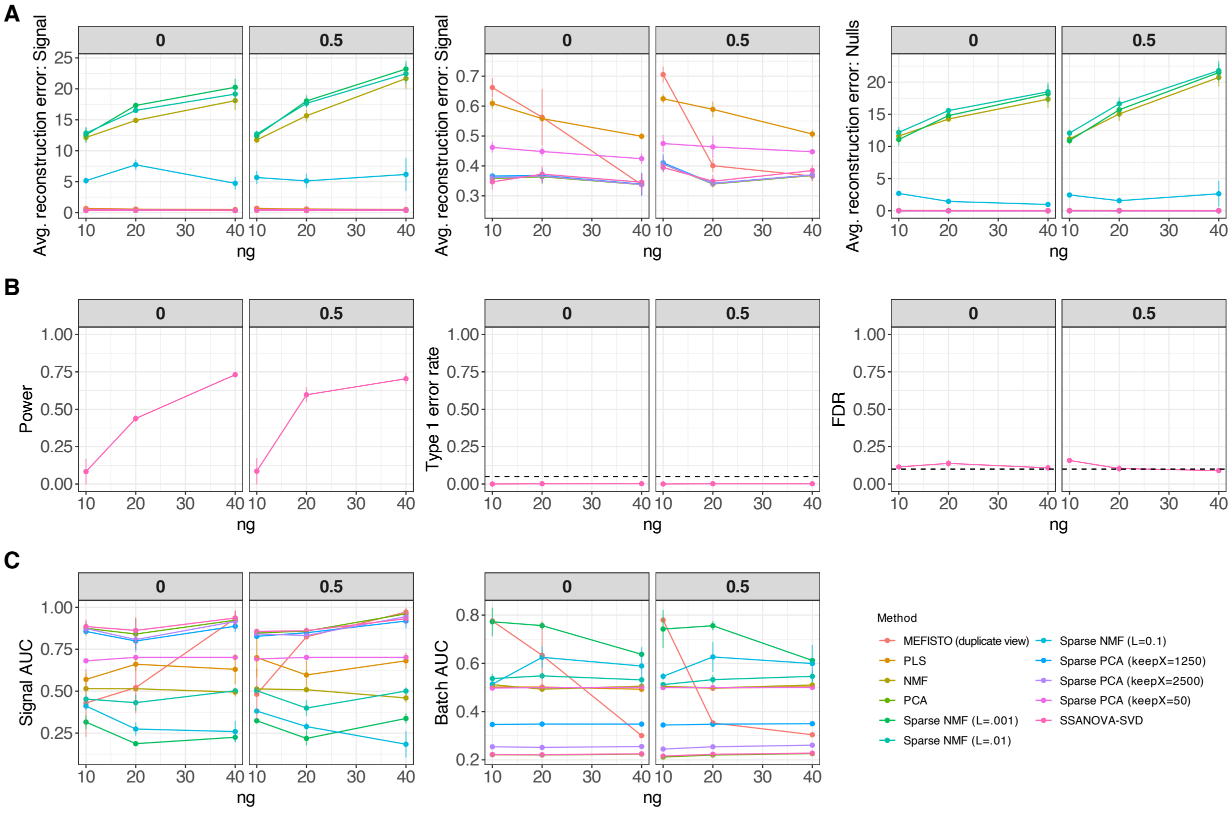


**Supplementary Figure 2: Performance of SSANOVA-SVD and the three-step framework in simulated data when 50% of signal features also experience batch effects.** Methods were evaluated for increasing numbers of samples per biological group ($n_g$) under either no confounding (0) or moderate confounding (0.5) of group by batch. SSANOVA-SVD here refers to the implementation wrapping SSANOVA-SVD and the three-step framework. For Sparse NMF and Sparse PCA, the penalty parameters were additionally varied as indicated in the legend. A. Average reconstruction error for signal features (left), the same reconstruction error shown on a zoomed-in scale (middle), and null features (right). B. Power, type I error rate, and false discovery rate (FDR) of the proposed approach for conditional inference of feature participations in simulated relevant processes. Dashed horizontal lines indicate the nominal type I error rate of 0.05 and FDR level of 0.10. C. Area under the receiver operating characteristic curve (AUC) of the proposed approach for prioritizing relevant signal features (left) and irrelevant batch-affected features (middle) in feature loadings. Results are averaged over three simulation replicates. Standard error bars are shown.

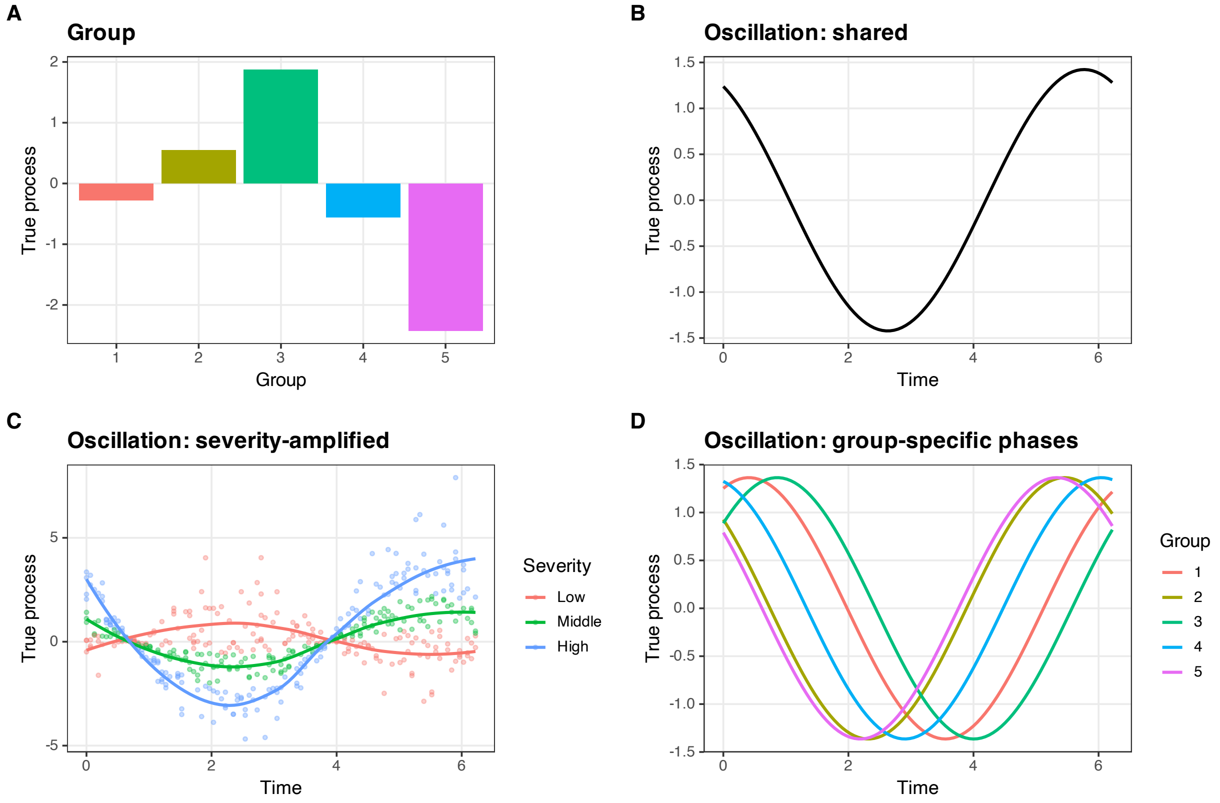


**Supplementary Figure 3: Simulated non-linear processes** True biological processes used in the representative simulation. (A) Group-specific mean shifts. (B) An oscillation shared across groups. (C) A severity-amplified oscillation whose amplitude varies continuously with sample-specific severity. For visualization, samples are divided into low, middle, and high severity strata and smoothed within each stratum; these three curves summarize a continuous severity effect and do not represent three discrete simulated severity states. (D) Oscillations with group-specific phase shifts.

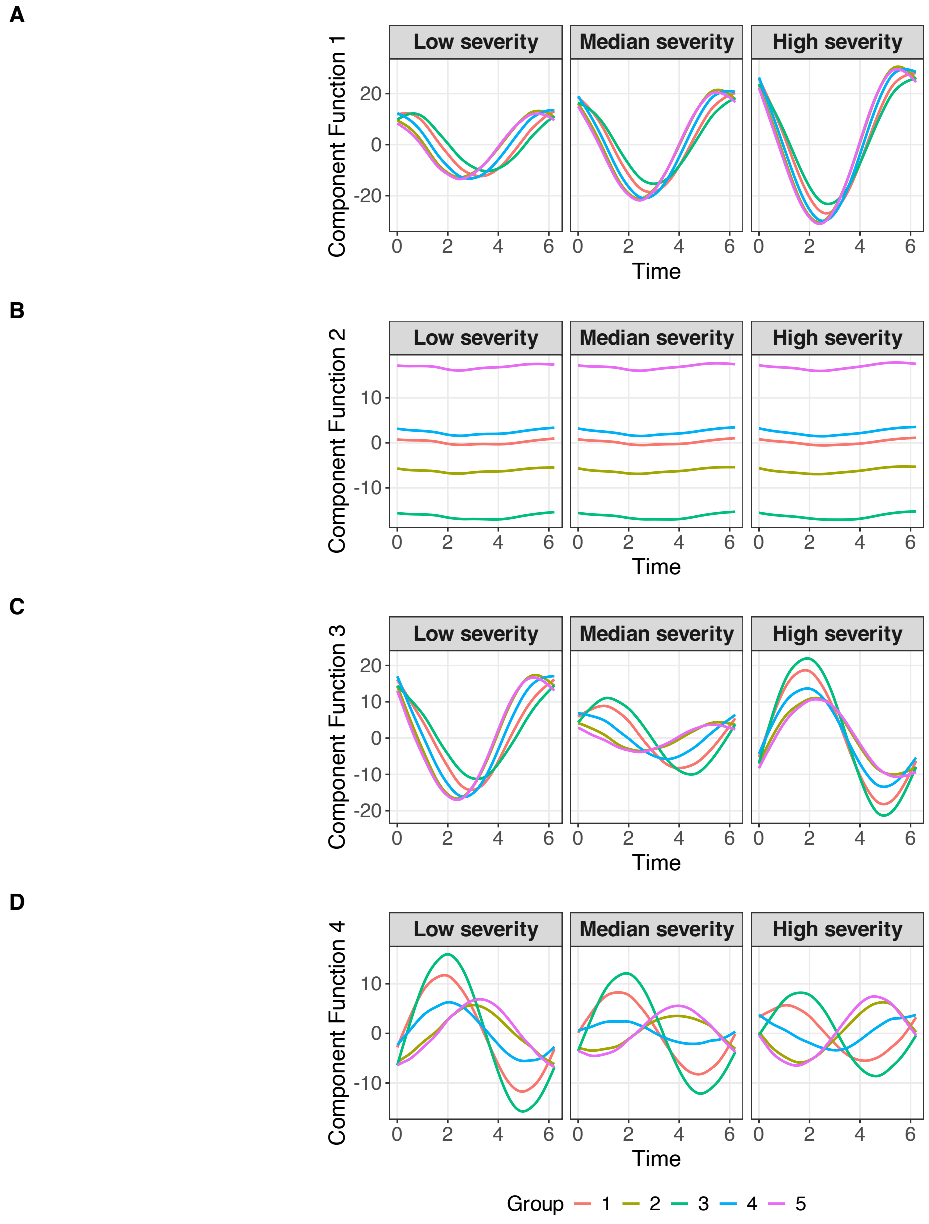


**Supplementary Figure 4: Reconstructed processes** Reconstructed SSANOVA-SVD component functions for the representative simulation. Component functions are shown across the simulated design for low, median, and high severity values.

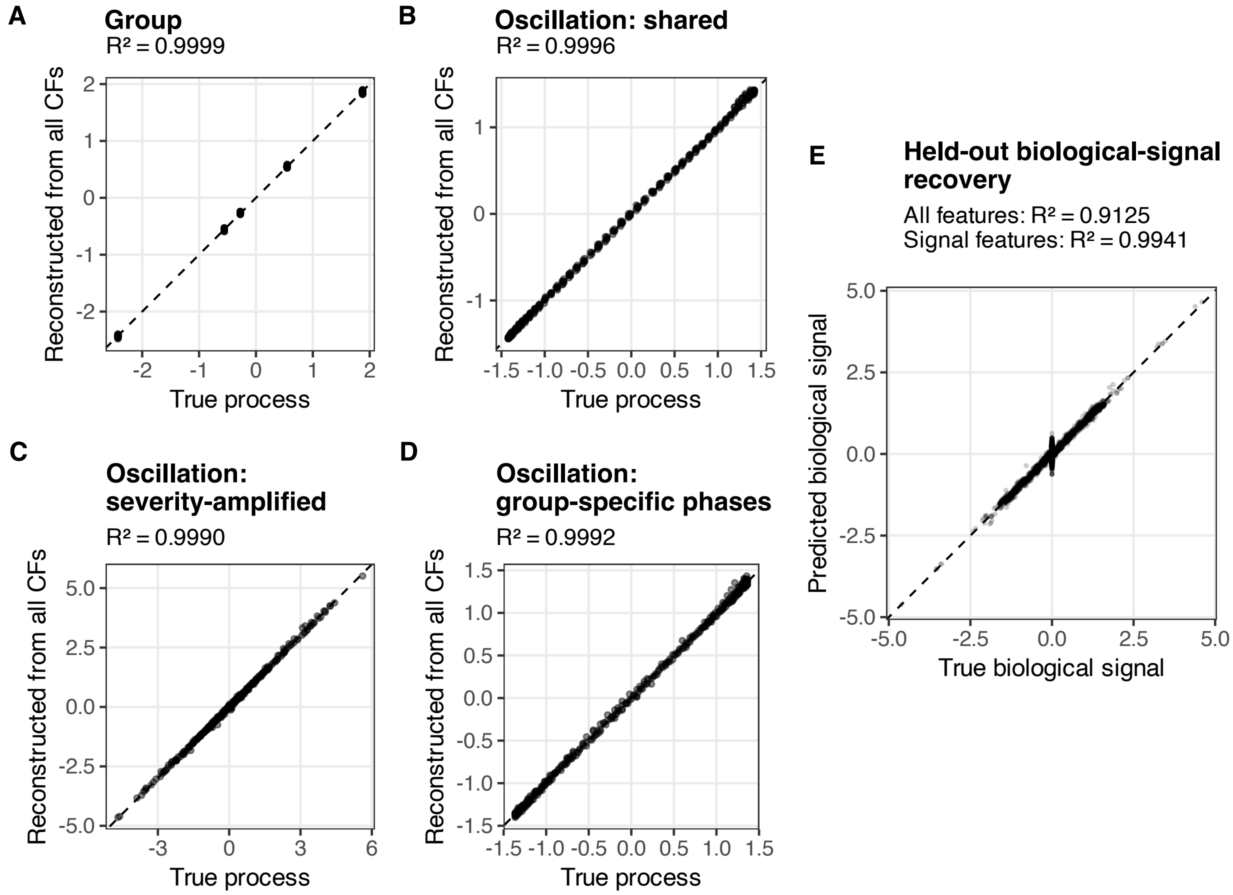


**Supplementary Figure 5: Process recovery** Recovery of the simulated biological structure. Panels A-D compare each true generating process with its reconstruction from the full learned component-function basis; the recovered component space explains 0.9994 of the combined true-process variation. Panel E compares the true and predicted biological signal in held-out samples.

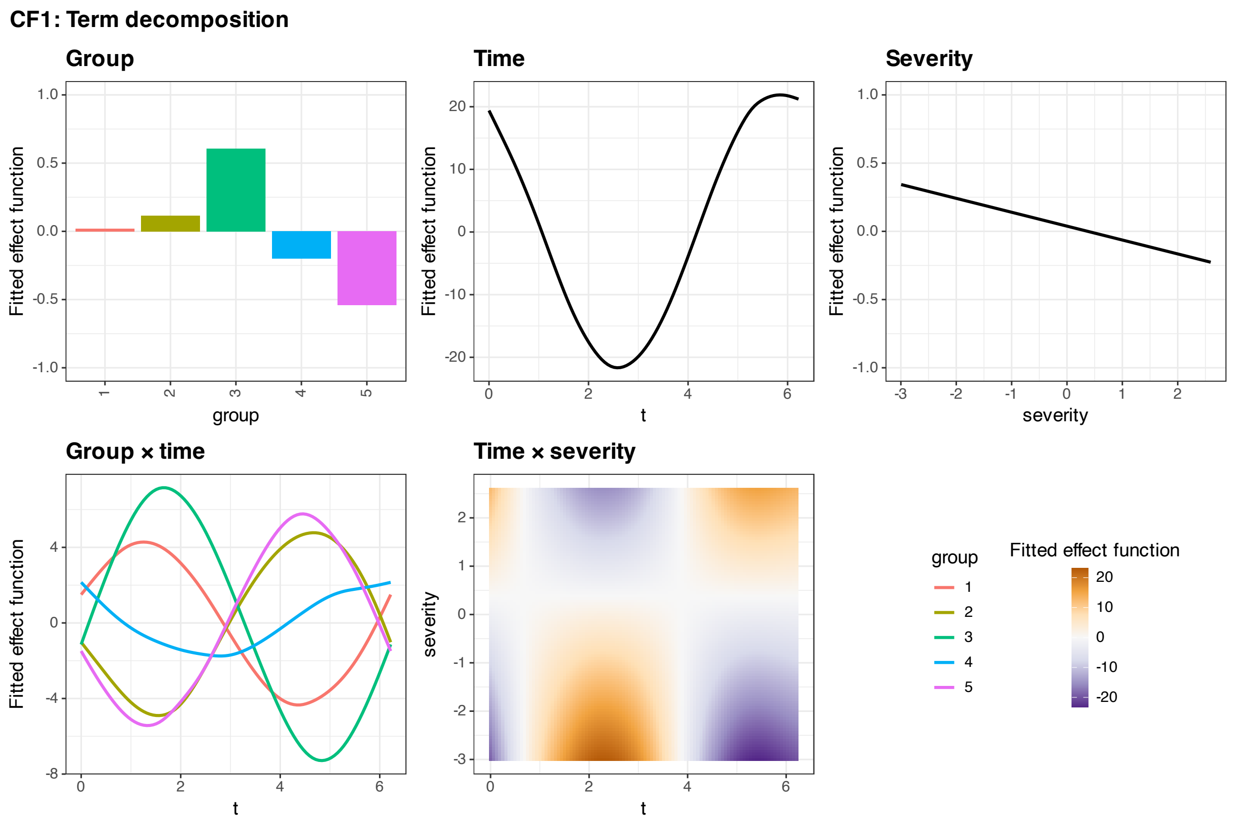


**Supplementary Figure 6: Decomposition of process 1** Term decomposition of reconstructed component function 1 into group, time, severity, group-by-time, and time-by-severity contributions.

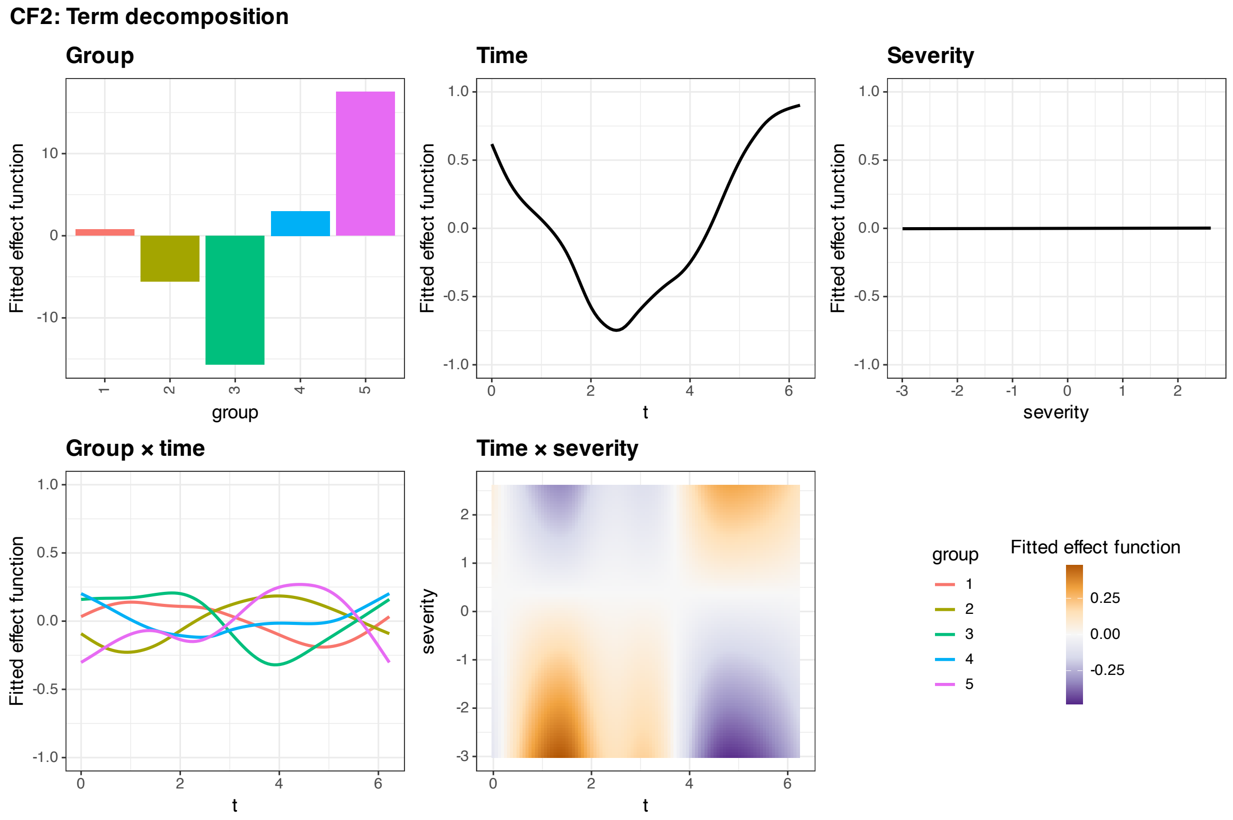


**Supplementary Figure 7: Decomposition of process 2** Term decomposition of reconstructed component function 2 into group, time, severity, group-by-time, and time-by-severity contributions.

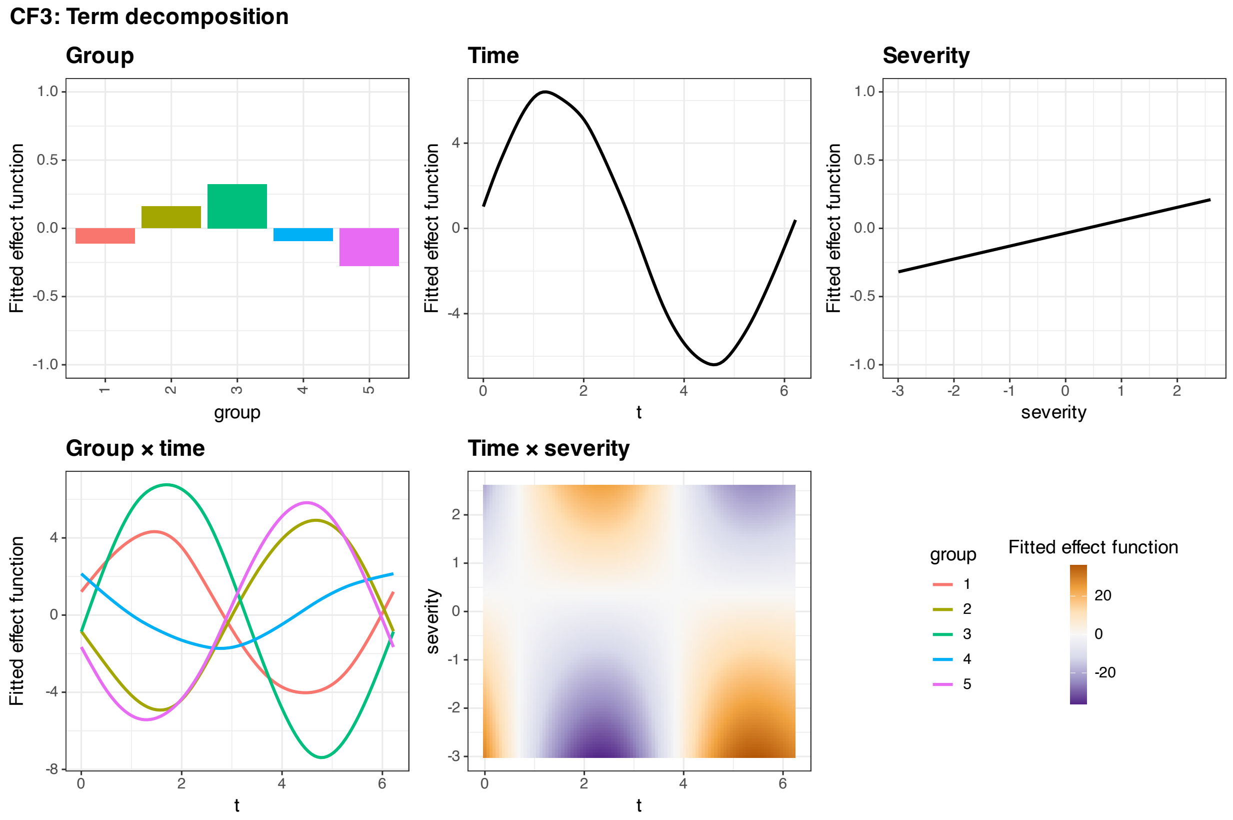


**Supplementary Figure 8: Decomposition of process 3** Term decomposition of reconstructed component function 3 into group, time, severity, group-by-time, and time-by-severity contributions.

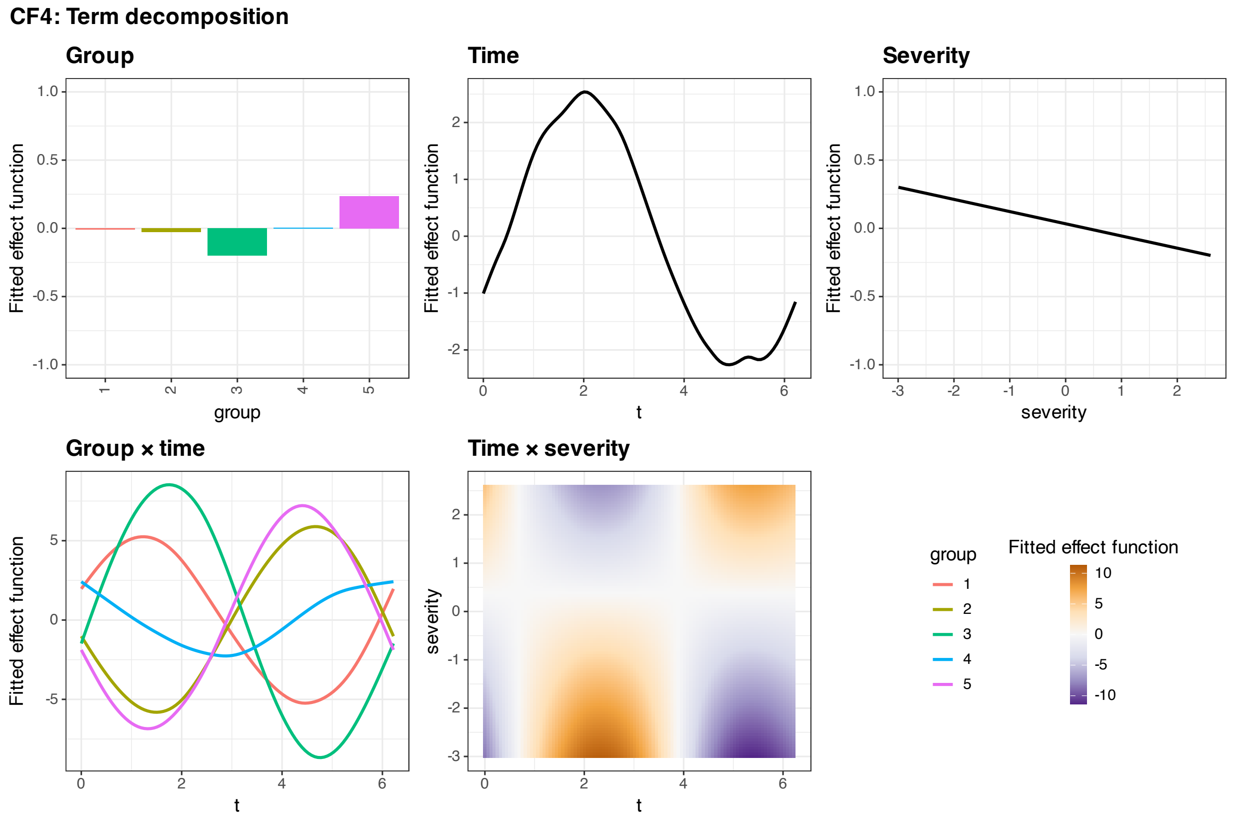


**Supplementary Figure 9: Decomposition of process 4** Term decomposition of reconstructed component function 4 into group, time, severity, group-by-time, and time-by-severity contributions.
