## Supplementary Methods for "A statistical framework for system-level synthesis"

In the main text, we proposed a three step framework for system-level synthesis. We developed SSANOVA-SVD to serve the first step, which involves reconstructing overarching processes as smooth functions parameterized by covariates and their interactions.

**Sections 1-5** of this note presents the details of the SSANOVA-SVD problem, the model-space construction for the overarching processes/functions, and a sequential rank-1 estimation algorithm for solving the problem, together with a representer theorem that enables its implementation by reducing each infinite-dimensional problem in the sequence to a finite-dimensional one.

**Section 6** presents the details of the simulations.

### 1 The SSANOVA-SVD problem

We cast the problem as :

$$(\widehat{U}, \widehat{V}, \widehat{\zeta}) = \underset{U,V,\zeta}{\arg\min} \quad \frac{1}{n} \left\| Y - \left( Z\zeta + \widetilde{U}V^\top \right) \right\|_F^2 + \lambda \sum_{k=1}^{K} J(u_k) \tag{1a}$$

$$\text{subject to} \quad u_k(\cdot) \in \mathcal{H}, \qquad 1 \le k \le K. \tag{1b}$$

$$(u_k, 1)_{\mathcal{H}} = 0 \qquad 1 \le k \le K. \tag{1c}$$

$$(u_k, u_l)_{\mathcal{H}} = 0, \qquad 1 \le k \ne l \le K. \tag{1d}$$

$$v_k^\top v_k = 1, \qquad 1 \le k \le K. \tag{1e}$$

$Y$ is an $n \times p$ data matrix of samples in rows, and features in columns. $Z$ is a design matrix of known confounders, with $n$ rows and $m$ columns. $\zeta$ is a matrix of regression coefficients with $m$ rows and $p$ columns. Here $\widetilde{U}_{n\times K} = (\widetilde{u}_1, \ldots, \widetilde{u}_K)$ is the components matrix obtained by evaluating the overarching processes/functions $U = \{u_1(\cdot), \cdots, u_K(\cdot)\}$ at sample covariates $(x_1, \ldots, x_n)$. Each $x_i$ takes the form $(x_{<1>i}, \ldots, x_{<B>}i) \in \chi$, an input domain for the functions. Here $x_{<t>}$ is a possibly discrete (nominal, ordinal) or continuous real valued covariate. For example, $x_{<1>i}$ may indicate clinical group of observation $i$, and $x_{<2>i}$ the time of measurement of observation $i$. $V_{p\times K} = (v_1 \ldots v_K)$ is the feature loadings matrix.

The set of processes and their corresponding feature loadings $\{u_k(\cdot), v_k\}_{k=1}^K$ define the *system-level model* learned from the data $Y$. The quantity $Z\zeta$ represents nuisance variation specific to the particular set of measurements $Y$.

$\mathcal{H}$ is the model space for the overarching processes. We choose this to be a multiple term smoothing spline ANOVA space of functions[1] (Section 2). $(\cdot,\cdot)_{\mathcal{H}}$ is an inner product for the space, and $J(\cdot)$ is a roughness penalty that enforces smoothness in the $u_k(\cdot)$. In the context of our proposed three-step synthesis framework for system-level synthesis (**Fig. 1** in main text), the space $\mathcal{H}$ allows us to parameterize the overarching processes in terms of covariates, enables prediction in unseen observations with only their covariate information, allows decomposition of the processes for contributions due to covariates and their interactions, and motivates the choice of the roughness penalty $J(\cdot)$. Constraint 1c removes the constant component from $u_k(\cdot)$, allowing feature-specific baselines to be identified via $Z$. Constraint 1d induces orthogonality among the component functions. Constraint 1e identifies the scale of $u_k(\cdot)$.

Numerical estimation is performed via alternating minimization. To compute the rank-$K$ factorization, we sequentially solve constrained rank-one optimization problems, enforcing orthogonality against previously estimated component functions at each step (Sections 3–4). This formulation, together with a representer theorem, reduces each rank-one infinite dimensional optimization problem to a finite-dimensional one (Section 3). The ability to predict the system-level processes in unseen observations without data reuse enables hyperparameters, including the number of components $K$, to be selected by cross-validation based on out-of-sample predictive performance (Section 5).

The following sections describe the model space, sequential algorithm, the representer theorem, and the cross-validation procedure for the estimation of hyperparameters.

### 2 $\mathcal{H}$: A multiple term smoothing spline ANOVA model space for system-level processes $U$

We choose $\mathcal{H}$ to be a reproducing kernel Hilbert space (RKHS) of functions defined on a domain $\chi$, with reproducing kernel $R(\cdot,\cdot)$. The reproducing property yields $u(x) = (R(x,\cdot), u)_{\mathcal{H}}, u \in \mathcal{H},\ x \in \chi$.

The roughness penalty $J(u)$ is a squared seminorm on $\mathcal{H}$, with null space $\mathcal{H}_0 = \{u \in \mathcal{H} : J(u) = 0\}$. The space admits an orthogonal decomposition, $\mathcal{H} = \mathcal{H}_0 \oplus \mathcal{H}_1$, also called as the tensor sum decomposition, where $\mathcal{H}_1$ is the orthogonal complement of $\mathcal{H}_0$. Both subspaces are RKHSs, with reproducing kernels $R_0(\cdot,\cdot)$ and $R_1(\cdot,\cdot)$, respectively, satisfying $R(x,y) = R_0(x,y) + R_1(x,y)$. The null space $\mathcal{H}_0$ is finite-dimensional and is spanned by a set of interpretable basis functions $\{\phi_b\}_{b=1}^{r}$, including the constant function (denoted as 1). Its reproducing kernel is $R_0(x,y) = \sum_{b=1}^{r} \phi_b(x)\phi_b(y)$. Let $\mathcal{P}_1$ denote the orthogonal projection of $u \in \mathcal{H}$ onto $\mathcal{H}_1$. The roughness penalty is defined as

$$J(u) = \|\mathcal{P}_1 u\|_{\mathcal{H}}^2 = (\mathcal{P}_1 u, \mathcal{P}_1 u)_{\mathcal{H}_1}.$$

Thus, the penalty acts only on the component of $u$ belonging to $\mathcal{H}_1$, leaving the null-space component unpenalized.

**Construction of $\mathcal{H}$ from a user-specified R model formula.** We construct $\mathcal{H}$ for the family of functions specified by the end user through an R model formula, following the smoothing spline ANOVA construction of Gu[1] (Chapter 2, Section 2.4).

Suppose the model contains $B$ covariates, with

$$x_{\langle b\rangle} \in \chi_{\langle b\rangle}, \qquad b = 1, \ldots, B.$$

For each covariate, we construct a marginal RKHS $\mathcal{H}_{\langle b\rangle}$ with the orthogonal decomposition

$$\mathcal{H}_{\langle b\rangle} = \mathcal{H}_{0\langle b\rangle} \oplus \mathcal{H}_{1\langle b\rangle},$$

specified through its inner product, roughness penalty, and corresponding reproducing kernels $R_{0\langle b\rangle}$ and $R_{1\langle b\rangle}$.

The full covariate domain is the Cartesian product

$$\chi = \prod_{b=1}^{B} \chi_{\langle b\rangle}.$$

The tensor-product RKHS for functions operating on points $x \in \chi$, is constructed as

$$\mathcal{H} = \bigotimes_{b=1}^{B} \mathcal{H}_{\langle b\rangle} = \bigotimes_{b=1}^{B} \left(\mathcal{H}_{0\langle b\rangle} \oplus \mathcal{H}_{1\langle b\rangle}\right).$$

Expanding this tensor product yields an orthogonal decomposition into subspaces corresponding to constant, main-effect, and interaction terms. The user-specified R model formula determines which of these terms are retained in $\mathcal{H}$.

The inner product, roughness penalty, and reproducing kernels $R_0(\cdot,\cdot)$ and $R_1(\cdot,\cdot)$ for the resulting model space are constructed from the marginal spaces. Because the inner products of the component RKHSs induce metrics that need not be directly comparable, scaling parameters are additionally introduced (Gu, 2013, Section 2.4.5). These scaling parameters are treated as hyperparameters and estimated from the data.

For example, an additive model retains only the constant and main-effect subspaces, whereas a model with interactions additionally includes the corresponding interaction subspaces. The reproducing kernels of these terms are obtained from the marginal kernels through sums and products. Thus, the marginal reproducing kernels provide the building blocks for constructing $\mathcal{H}$ under any supported user-specified R model formula.

We obtain the marginal reproducing kernels for nominal, ordinal, and continuous covariates from the R package *gss*[1].

### 3 Representer theorems

The sequential estimation algorithm presented in the next section exploits the following representer theorems. All the notation from the previous model space section carry over. Tilde notation over a function name (e.g., $\tilde{u}$) is used to represent evaluations at specified points (usually $x_i, i = 1, \dots n$ ).

**Theorem 1.** *Let $y_1, \dots y_n$ be scalar observations at points $x_1, \dots, x_n$. Let $\mathcal{H}$ be an RKHS with reproducing kernel $R$. Let $\mathcal{H} = \mathcal{H}_0 \oplus \mathcal{H}_1$ define a tensor sum decomposition of $\mathcal{H}$, with $\mathcal{H}_0$ containing simple to interpret basis functions, including constant functions. Let $R_1$ be the reproducing kernel of $\mathcal{H}_1$, and $J(u) = (u, u)_{\mathcal{H}_1}$. For $u \in \mathcal{H}$, let $\tilde{u} = (u(x_1), \dots, u(x_n))^T$. Suppose $q \in \mathcal{H}_f = \mathcal{H}_0 \oplus \text{Span}\{R_1(x_i), i = 1, \dots n\}$. Then the solution to the following infinite dimensional problem, $\hat{u}(\cdot)$, also lies in the finite dimensional space $\mathcal{H}_f$.*

$$\widehat{u} = \arg\min_{u} \frac{1}{n} \|y - \widetilde{u}\|_2^2 + \lambda J(u), \tag{2a}$$

$$\text{subject to} \quad u \in \mathcal{H}. \tag{2b}$$

$$(q, u)_{\mathcal{H}} = 0. \tag{2c}$$

*Proof.* For any $u \in \mathcal{H}$, $u = u_f + \rho$, where $u_f \in \mathcal{H}_f$, and $\rho \in \mathcal{H} \ominus \mathcal{H}_f$, so $(u_f, \rho) = 0$.

The inner product term $(q, u)_{\mathcal{H}} = (q, u_f)_{\mathcal{H}} + (q, \rho)_{\mathcal{H}} = (q, u_f)_{\mathcal{H}} + 0$ as $q \in \mathcal{H}_f$.

Let $R_{x_i} = R(x_i, \cdot)$ be the representer of evaluation at $x_i$. Then the first term in the objective:

$$\begin{aligned}
\|y - \tilde{u}\|_2^2 &= \sum_{i=1}^{n} (y_i - u(x_i))^2 \\
&= \sum_{i=1}^{n} (y_i - (R_{x_i}, u))^2 \\
&= \sum_{i=1}^{n} (y_i - (R_{x_i}, u_f + \rho))^2 \\
&= \sum_{i=1}^{n} (y_i - (R_{x_i}, u_f))^2 \text{ as } R_{x_i} \in \mathcal{H}_f \\
&= \sum_{i=1}^{n} (y_i - u_f(x_i))^2 \\
&= \|y - \tilde{u}_f\|_2^2
\end{aligned}$$

The penalty term:

$$\begin{aligned} J(u) &= (u,u)_{\mathcal{H}_1} \\ &= (u_f,u_f)_{\mathcal{H}_1} + (\rho,\rho)_{\mathcal{H}_1} \\ &\geq J(u_f) \end{aligned}$$

Thus, $\rho$ drops out of the minimization problem 2, $u(\cdot)$ lies in finite dimensions, and the problem can be written as:

$$\widehat{u} = \arg\min_{u} \frac{1}{n} \|y - \widetilde{u}\|_2^2 + \lambda J(u), \tag{3a}$$

$$\text{subject to} \qquad u \in \mathcal{H}_f. \tag{3b}$$

$$(q,u)_{\mathcal{H}} = 0. \tag{3c}$$

□

**Computation of the solution** Because $u \in \mathcal{H}_f$,

$$\begin{aligned} u(x_i) &= \underbrace{[\phi_1(x_i),\ldots\phi_r(x_i)]d}_{Projection\ onto\ \mathcal{H}_0} + \underbrace{[R_1(x_1,x_i),\ldots,R_1(x_n,x_i)]c}_{Projection\ onto\ \mathrm{Span}\{R_1(x_1,\cdot),\ldots,R_1(x_n,\cdot)\}} \\ &:= s(x_i)^T d + \tilde{r}_1(x_i)^T c \end{aligned} \tag{4}$$

Thus

$$\tilde{u} = Sd + \tilde{R}_1 c$$

where the $i^{th}$ row of $S$ is $s(x_i)^T$ and the $i^{th}$ row of $\tilde{R}_1$ is $\tilde{r}_1(x_i)^T$. Following the multiple term RKHS formulation of section 2.4.5 in Gu[1], we set $J(u) = (u,u)_{\mathcal{H}_1} = c^T \tilde{R}_1 c$, $(q,u)_{\mathcal{H}} = \tilde{q}^T \tilde{R}^{+} \tilde{u} = \tilde{q}^T \tilde{R}^{+}(Sd + \tilde{R}_1 c)$, where $\tilde{q}_i = q(x_i)$, and $\tilde{R}^{+}$ is the pseudoinverse of $\tilde{R}$ with $\tilde{R}(i,j) = R(x_i,x_j)$, $i,j \in \{1,\ldots n\}$.

Problem 2 is then solved as:

$$\hat{u}(x) = s(x)^T \hat{d} + \tilde{r}_1(x)^T \hat{c} \tag{5}$$

where $\hat{c}$ and $\hat{d}$ are obtained by solving the reduced problem :

$$\begin{aligned} (\widehat{c},\widehat{d}) &= \arg\min_{c,d} \frac{1}{n} \left\| y - (Sd + \tilde{R}_1 c) \right\|_2^2 + \lambda c^T \tilde{R}_1 c, \\ \text{subject to} \qquad & \tilde{q}^T \tilde{R}^{+}(Sd + \tilde{R}_1 c) = 0. \end{aligned} \tag{6}$$

Solution based on the Lagrangian is then straightforwardly obtained.

**Theorem 2.** *Let $y_1,\ldots y_n$ be scalar observations at points $x_1,\ldots,x_n$. Let $\mathcal{H}$ be an RKHS with reproducing kernel $R$. Let $\mathcal{H} = \mathcal{H}_0 \oplus \mathcal{H}_1$ define a tensor sum decomposition of $\mathcal{H}$, with $\mathcal{H}_0$ containing simple to interpret basis functions, including constant functions. Let $R_1$ be the reproducing kernel of $\mathcal{H}_1$, and $J(u) = (u,u)_{H_1}$. Let $\tilde{u} = (u(x_1),\ldots,u(x_n))^T$. Suppose $q_1 \ldots q_k \in \mathcal{H}_f = \mathcal{H}_0 \oplus \mathrm{Span}\{R_1(x_i), i = 1,\ldots n\}$. Then the solution to the following infinite dimensional problem, $\hat{u}(\cdot)$, also lies in the finite dimensional space $\mathcal{H}_f$.*

$$\widehat{u} = \arg\min_{u} \frac{1}{n} \|y - \widetilde{u}\|_2^2 + \lambda J(u), \tag{7a}$$

$$\text{subject to} \qquad u \in \mathcal{H}. \tag{7b}$$

$$(q_t,u)_{\mathcal{H}} = 0, \quad 1 \leq t \leq k. \tag{7c}$$

*Proof.* $u \in \mathcal{H} \implies u = u_f + \rho$, $u_f \in H_f$, and $\rho \in \mathcal{H} \ominus \mathcal{H}_f$, with $(u_f, \rho) = 0$. Following the same arguments of theorem 1, we have:

$$
\begin{aligned}
&(q_t, u)_{\mathcal{H}} = (q_t, u_f)_{\mathcal{H}} \\
&\|y - \tilde{u}\|_2^2 = \|y - \tilde{u_f}\|_2^2 \\
&J(u) \geq J(u_f)
\end{aligned}
$$

So $\rho$ drops out of the minimization problem 7, $u(\cdot)$ lies in finite dimensions, and the problem reduces to:

$$
\widehat{u} = \arg\min_{u} \frac{1}{n} \|y - \widetilde{u}\|_2^2 + \lambda J(u), \tag{8a}
$$

$$
\text{subject to} \quad u \in \mathcal{H}_f. \tag{8b}
$$

$$
(q_t, u) = 0, \quad 1 \leq t \leq k. \tag{8c}
$$

□

**Computation of the solution** Same arguments following equations 4 and 6 allows us to write the solution as:

$$
u(x) = s(x)^T \hat{d} + \tilde{r}_1(x)^T \hat{c} \tag{9}
$$

where $\hat{c}$ and $\hat{d}$ are solutions of the following problem :

$$
\begin{aligned}
(\widehat{c}, \widehat{d}) = \arg\min_{c,d} \frac{1}{n} \left\| y - (Sd + \tilde{R}_1 c) \right\|_2^2 + \lambda c^T \tilde{R}_1 c, \\
\text{subject to} \qquad Q^T \tilde{R}^+ (Sd + \tilde{R}_1 c) = 0.
\end{aligned} \tag{10}
$$

where the $t^{th}$ column of $Q$ is $\tilde{q}_t$. Solution based on the Lagrangian is then straightforwardly available.

**Remark** Similar to smoothing spline ANOVA models, for computational efficiency, we might approximate the solution $u(x)$ due to theorems 1 and 2, over a subset of $l$ basis points as $s(x)^T d + \tilde{\tilde{r}}_1(x)^T c$ where $\tilde{\tilde{r}}_1(x) = [R_1(x_{z_1}, x), \ldots, R_1(x_{z_l}, x)]$. For simplicity, we reuse the notation $c$. $z_1 \ldots z_l \in \{1, \ldots n\}$. In this case, we have $\tilde{u} = Sd + \tilde{\tilde{R}}_1 c$, where the $i^{th}$ row of $\tilde{\tilde{R}}_1$ is $\tilde{\tilde{r}}_1(x_i)^T$. We set $J(u) = (u, u)_{\mathcal{H}_1} = c^T \tilde{\tilde{G}}_1 c$, where $\tilde{\tilde{G}}_1(i,j) = R_1(x_{z_i}, x_{z_j})$, $i, j \in \{1, \ldots, l\}$. We set $(q, u)_{\mathcal{H}} = \tilde{q}^T A^T \tilde{\tilde{G}}^+ A \tilde{u} = \tilde{q}^T A^T \tilde{\tilde{G}}^+ A(Sd + \tilde{\tilde{R}}_1 c)$, where $A$ extracts the $l$ basis point indices from $n$ dimensional vectors. $\tilde{q}_i = q(x_i)$, and $\tilde{\tilde{G}}^+$ is the pseudoinverse of $\tilde{\tilde{G}}$ with $\tilde{\tilde{G}}(i,j) = R(x_{z_i}, x_{z_j})$, $i, j \in \{1, \ldots l\}$.

### 4 Sequential estimation algorithm for the SSANOVA-SVD problem

To fit the component functions $U$ in problem 1, we consider an efficient sequential estimation algorithm in which we estimate $u_k$ conditional on the previously estimated functions $u_t$, $t < k$. Under this estimation strategy, we have the rank-1 SSANOVA-SVD problem :

$$
(\widehat{\zeta}, \widehat{u}(\cdot), \widehat{v}) = \arg\min_{\zeta \in \mathbb{R}^{m \times p},\, v \in \mathbb{R}^p,\, u} \frac{1}{n} \left\| \left( Y - \sum_{t=1}^{k-1} \widehat{\tilde{u}}_t \widehat{v}_t^T \right) - Z\zeta - \tilde{u} v^\top \right\|_F^2 + \lambda J(u) \tag{11a}
$$

$$
\text{subject to} \quad u \in \mathcal{H} \tag{11b}
$$

$$
(u, 1)_{\mathcal{H}} = 0 \tag{11c}
$$

$$
v^\top v = 1 \tag{11d}
$$

$$
(\widehat{u}_t, u)_{\mathcal{H}} = 0, \quad t = 1, \ldots, k-1 \tag{11e}
$$

We consider two cases. Estimation of $u_1$ and estimation of $u_k, k > 1$. In both cases, the problems are non-convex. An alternating minimization algorithm is used for estimation:

1. Fit $\zeta$, given $\widehat{u}(\cdot)$, $\widehat{v}$.
2. Fit $u(\cdot)$, given $\widehat{\zeta}$ and $\widehat{v}$.
3. Fit $v$, given $\widehat{\zeta}$ and $\widehat{u}(\cdot)$.

We describe the two cases now.

#### 4.1 Case 1: Estimation of $u_1(\cdot)$

When solving for $u_1(\cdot)$, no prior components exist, therefore constraint 11e is dropped. We have:

$$
\begin{aligned}
(\widehat{\zeta}, \widehat{u}(\cdot), \widehat{v}) = & \underset{\zeta \in \mathbb{R}^{m \times p},\, v \in \mathbb{R}^p,\, u}{\arg\min} \frac{1}{n} \left\| Y - Z\zeta - \tilde{u}v^\top \right\|_F^2 + \lambda J(u) && \text{(12a)} \\
\text{subject to} \quad & u \in \mathcal{H} && \text{(12b)} \\
& (u, 1)_{\mathcal{H}} = 0 && \text{(12c)} \\
& v^\top v = 1 && \text{(12d)}
\end{aligned}
$$

##### 4.1.1 Fit $\zeta$, given $\widehat{u}(\cdot)$, $\widehat{v}$

Let $\widehat{\widetilde{u}} = [\widehat{u}(x_1), \ \ldots, \ \widehat{u}(x_n)]^T$. With $Y_1 = Y - \widehat{\widetilde{u}}\widehat{v}^T$, the problem of estimating $\zeta$ reduces to

$$
argmin_{\zeta \in \mathcal{R}^{m \times p}} \ \frac{1}{n} \left\| Y_1 - Z\zeta \right\|_F^2
$$

$\widehat{\zeta}$ is available as the ordinary least squares estimator:

$$
\widehat{\zeta} = \left( Z^T Z \right)^+ Z^T Y_1 \tag{13}
$$

where the superscript + represents the pseudoinverse.

##### 4.1.2 Fit $u(\cdot)$, given $\hat{\zeta}$, $\hat{v}$

Let $Y_2 = Y - Z\widehat{\zeta}$. The problem reduces to:

$$
\begin{aligned}
\widehat{u} = & \underset{u}{\arg\min} \frac{1}{n} \left\| Y_2 - \widetilde{u}\widehat{v}^\top \right\|_F^2 + \lambda J(u), && \text{(14a)} \\
\text{subject to} \quad & u \in \mathcal{H} && \text{(14b)} \\
& (u, 1) = 0 && \text{(14c)}
\end{aligned}
$$

Let $\widehat{w} = Y_2 \widehat{v} \in \mathcal{R}^n$. Then: $\left\| Y_2 - \widetilde{u}\widehat{v}^T \right\|_F^2 = \widetilde{u}^T \widetilde{u} + 2\widehat{w}^T \widetilde{u} + Constant$. The problem is equivalent to minimizing: $\frac{1}{n} \left\| \widehat{w} - \widetilde{u} \right\|^2 + \lambda \cdot J(u)$ under the constraints. We have the equivalent problem:

$$
\begin{aligned}
\widehat{u} = & \underset{u}{\arg\min} \frac{1}{n} \left\| \widehat{w} - \widetilde{u} \right\|_2^2 + \lambda J(u), && \text{(15a)} \\
\text{subject to} \quad & u \in \mathcal{H} && \text{(15b)} \\
& (u, 1) = 0 && \text{(15c)}
\end{aligned}
$$

Because the constant function $1 \in \mathcal{H}_f$, problem 15 is an instance of the problem covered by Theorem 1, $u \in \mathcal{H}_f$, and the solution $\hat{u}$ is available from 5.

###### 4.1.3 Fit $v$, given $\widehat{u}(\cdot)$, $\widehat{\zeta}$

The problem reduces to:

$$\underset{v \in \mathbb{R}^p}{\arg\min} \ \frac{1}{n} \left\| Y_2 - \widehat{\widetilde{u}} v^\top \right\|_F^2 \tag{16}$$

$$\text{subject to } v^T v = 1 \tag{17}$$

The solution is obtained as:

$$\widehat{v} = Y_2 \widehat{\widetilde{u}} / \| Y_2 \widehat{\widetilde{u}} \| \tag{18}$$

##### 4.2 Case 2: Estimation of $u_k(\cdot)$ given $\hat{u}_t(\cdot), 1 \le t < k$

In this case, constraint 11e is active, and we have the full problem 12 to solve.

###### 4.2.1 Fit $\zeta$, given $\widehat{u}(\cdot)$, $\widehat{v}$

Let $\widehat{\widetilde{u}} = [\widehat{u}(x_1), \ \ldots, \ \widehat{u}(x_n)]^T$. With $Y_1^* = Y - \sum_{t=1}^{k-1} \widehat{\widetilde{u}}_t \widehat{v}_t^T - \widehat{\widetilde{u}} \widehat{v}^T$ , the problem of estimating $\zeta$ reduces to

$$argmin_{\zeta \in \mathcal{R}^{m \times p}} \ \ \frac{1}{n} \| Y_1^* - Z\zeta \|_F^2$$

$\widehat{\zeta}$ is available as the ordinary least squares estimator:

$$\widehat{\zeta} = \left( Z^T Z \right)^+ Z^T Y_1^* \tag{19}$$

where the superscript + represents the pseudoinverse.

###### 4.2.2 Fit $u(\cdot)$, given $\hat{\zeta}$, $\hat{v}$

Let $Y_2^* = Y - \sum_{t=1}^{k-1} \widehat{\widetilde{u}}_t \widehat{v}_t^T - Z\widehat{\zeta}$. The problem reduces to:

$$\widehat{u} = \underset{u}{\arg\min} \ \frac{1}{n} \left\| Y_2^* - \widetilde{u} \widehat{v}^\top \right\|_F^2 + \lambda J(u), \tag{20a}$$

$$\text{subject to} \qquad u \in \mathcal{H} \tag{20b}$$

$$(u, 1) = 0 \tag{20c}$$

$$(u_t, u) = 0, 1 \le t \le k-1 \tag{20d}$$

Let $\widehat{w} = Y_2^* \widehat{v} \in \mathcal{R}^n$. Then: $\left\| Y_2^* - \widetilde{u} \widehat{v}^T \right\|_F^2 = \widetilde{u}^T \widetilde{u} + 2 \widehat{w}^T \widetilde{u} + Constant$. The problem is equivalent to minimizing: $\frac{1}{n} \| \widehat{w} - \widetilde{u} \|^2 + \lambda \cdot J(u)$ under the constraints. We have the equivalent problem:

$$\widehat{u} = \underset{u}{\arg\min} \ \frac{1}{n} \| \widehat{w} - \widetilde{u} \|_2^2 + \lambda J(u), \tag{21a}$$

$$\text{subject to} \qquad u \in \mathcal{H} \tag{21b}$$

$$(u, 1) = 0 \tag{21c}$$

$$(u_t, u) = 0, 1 \le t \le k-1 \tag{21d}$$

Because $1 \in \mathcal{H}_f, u_t \in \mathcal{H}_f, 1 \le t \le k-1$, problem 21 is an instance of the problem covered by Theorem 2, $u \in \mathcal{H}_f$, and the solution $\hat{u}$ is available from 9.

##### 4.2.3 Fit $v$, given $\widehat{u}(\cdot)$, $\widehat{\zeta}$

The problem reduces to:

$$\underset{v\in\mathbb{R}^p}{\arg\min}\ \frac{1}{n}\left\|Y_2^* - \widehat{\widehat{u}}v^\top\right\|_F^2 \tag{22}$$

$$\text{subject to } v^Tv = 1 \tag{23}$$

The solution is obtained as:

$$\widehat{v} = Y_2^*\widehat{\widehat{u}}/\|Y_2^*\widehat{\widehat{u}}\| \tag{24}$$

This completes the derivation of the sequential estimation algorithm for SSANOVA-SVD.

#### 5 Cross-validation by scoring out-of-sample predictive capacity

Model hyperparameters were selected using cross-validation based on prediction of held-out feature measurements. The procedure adapts standard cross-validation procedures for fitting smoothing spline ANOVA models to the SSANOVA-SVD problem.

**Strategy** For a required number of components, $K$, the sequential estimation algorithm of the previous section estimates the component functions $U = (u_1(\cdot), \ldots, u_K(\cdot))$. For any given set of covariate values from the input domain $\chi$, the component functions can be straightforwardly evaluated using 9. Thus, given a train and test split with their corresponding train and test covariates, we can train SSANOVA-SVD using the training set and its covariates, predict the entire test data matrix from the covariates for test data, and evaluate the mean squared error between the predicted test matrix and the observed test matrix, without double dipping onto the test data matrix. This score is then numerically optimized over to select the hyperparameters including the number of components $K$, smoothing penalty $\lambda$, and the term-wise normalization scales $\theta$ that are implicit in the reproducing kernels $R$ and $R_1$(section 2.4.5 in Gu[1]).

Briefly, $K$ was selected from a grid of $\{1 \ldots K_{max}\}$, where $K_{max}$ was input by the user. $\lambda$ was selected from a grid of $\{10^{-7}, 10^{-6}, \ldots, 1\}$. $\theta$ was numerically optimized over the CV score.

##### 5.1 Algorithm

Let $Y \in \mathbb{R}^{n\times p}$ denote the feature-by-sample data matrix, where $p$ is the number of features and $n$ is the number of samples. Let $\mathcal{I}_1, \ldots, \mathcal{I}_F$ denote a partition of the samples into $F$ cross-validation folds, with $\mathcal{I}_{-f}$ denoting the training samples obtained by excluding fold $f$ and $n_f = |\mathcal{I}_f|$ denoting the number of held-out samples in fold $f$.

For a candidate number of components $K$, penalty parameter $\lambda$, and vector of term-wise normalization parameters for the reproducing kernel, $\theta = (\theta_1, \ldots, \theta_L)^T$, where $L$ is the number of penalized SSANOVA components, SSANOVA-SVD is fitted using the samples in $\mathcal{I}_{-f}$. The fitted processes are then evaluated at the covariates of the held-out samples to obtain $\widehat{U}_f$. The corresponding held-out feature measurements are predicted as

$$\widehat{Y}_f = \widehat{U}_f\widehat{V}^\top + Z_f\widehat{\zeta},$$

where $\widehat{V}$ denotes the estimated feature-loading matrix, $Z_f$ denotes the measured confounder design matrix for the held-out samples, and $\widehat{\zeta}$ denotes the corresponding estimated adjustment coefficients. Thus, the process values of the held-out samples are predicted from their covariates without using their feature measurements.

For fold $f$, prediction error is measured by

$$\text{MSE}_f(K, \lambda, \theta) = \frac{1}{n_f p}\left\|Y_f - \widehat{Y}_f\right\|_F^2,$$

where $\|\cdot\|_F$ denotes the Frobenius norm. The cross-validation criterion is the mean of the fold-specific prediction errors,

$$\mathrm{CV}(K,\lambda,\theta)=\frac{1}{F}\sum_{f=1}^{F}\mathrm{MSE}_f(K,\lambda,\theta).$$

**Initialization.** Let $K^{\mathrm{in}}$, $\lambda^{\mathrm{in}}$, and $\theta^{\mathrm{in}}$ denote the initial hyperparameter values, and let $\mathcal{K}$ and $\mathcal{L}$ denote the candidate grids for $K$ and $\lambda$, respectively. $\mathcal{K}$ was set to $\{1\ldots K_{max}\}$ , where $K_{max}$ was input by the user. $\lambda$ was selected from a grid of $\{10^{-7},10^{-6},\ldots,1\}$. $\theta$ was numerically optimized. Initialization of the hyperparameters followed a strategy similar to Gu[1].

First, an initial rank is selected by minimizing the prediction criterion while holding $\lambda$ and $\theta$ at their initial values,

$$K^{(0)}=\arg\min_{K\in\mathcal{K}}\mathrm{CV}\left(K,\lambda^{\mathrm{in}},\theta^{\mathrm{in}}\right).$$

For cross-validation fold $f$, let $G_{\beta f}\in\mathbb{R}^{q_f\times q_f}$ denote the (possibly reduced-basis) kernel matrix for penalized SSANOVA term $\beta$, evaluated at the $q_f$ basis points used for the training in that fold. For $\beta=1,\ldots,L$, define

$$\widetilde{\theta}_\beta=\left\{\frac{1}{F}\sum_{f=1}^{F}\mathrm{tr}\left(G_{\beta f}\right)\right\}^{-1}.$$

The relative smoothing parameters are then placed on a common scale. Let $S_f\in\mathbb{R}^{n_{-f}\times r}$ denote the null-space design matrix in training fold $f$, where $n_{-f}=|\mathcal{I}_{-f}|$ and $r$ is the dimension of the null space. Let $R_{1f}(\theta)\in\mathbb{R}^{n_{-f}\times q_f}$ denote the combined penalized reduced-basis representation constructed using $\theta$, with $q_f$ columns. Define

$$s=\frac{1}{F}\sum_{f=1}^{F}\frac{\|S_f\|_F^2}{r}$$

and

$$r_\theta=\frac{1}{F}\sum_{f=1}^{F}\frac{\left\|R_{1f}(\widetilde{\theta})\right\|_F^2}{q_f}.$$

The common scaling factor is

$$a=\left(\frac{s}{r_\theta}\right)^{1/2},$$

and

$$\widetilde{\theta}\leftarrow a\widetilde{\theta}.$$

When the SSANOVA representation contains no null-space component, this common rescaling is omitted.

Holding $K=K^{(0)}$ and $\theta=\widetilde{\theta}$ fixed, an initial value of $\lambda$ is selected by

$$\widetilde{\lambda}=\arg\min_{\lambda\in\mathcal{L}}\mathrm{CV}\left(K^{(0)},\lambda,\widetilde{\theta}\right).$$

SSANOVA-SVD is subsequently fitted to the training samples of each cross-validation fold using $(K^{(0)},\widetilde{\lambda},\widetilde{\theta})$. For reconstructed process $k$ in fold $f$, let $c_{kf}\in\mathbb{R}^{q_f}$ denote the fitted coefficients associated with the penalized reduced-basis representation. For each SSANOVA component $\beta$, define its average fitted contribution to the penalty term by

$$E_\beta=\frac{1}{F}\sum_{f=1}^{F}\left\{\frac{1}{K^{(0)}}\sum_{k=1}^{K^{(0)}}c_{kf}^{\mathsf{T}}G_{\beta f}c_{kf}\right\}.$$

The signal-adaptive initial smoothing parameters are then

$$\theta_\beta^{(0)} = \widetilde{\theta}_\beta^2 E_\beta, \qquad \beta = 1, \ldots, L.$$

A common rescaling is applied once more to $\theta^{(0)}$. Specifically, define

$$r_\theta^{(0)} = \frac{1}{F}\sum_{f=1}^{F} \frac{\left\|R_{1f}(\theta^{(0)})\right\|_F^2}{q_f}, \qquad a_0 = \left(\frac{s}{r_\theta^{(0)}}\right)^{1/2},$$

and set

$$\theta^{(0)} \leftarrow a_0 \theta^{(0)}.$$

As above, this rescaling is omitted in the absence of a null-space component.

Finally, $\lambda$ is re-optimized at the initialized rank and smoothing parameters,

$$\lambda^{(0)} = \arg\min_{\lambda \in \mathcal{L}} \mathrm{CV}\left(K^{(0)}, \lambda, \theta^{(0)}\right).$$

The resulting $\lambda^{(0)}$ is held fixed during the subsequent coordinate-wise optimization (as only the product $\lambda\theta_\beta^{-1}$ enters the penalty).

**Coordinate-wise optimization.** Starting from $(K^{(0)}, \lambda^{(0)}, \theta^{(0)})$, the rank and smoothing parameters are iteratively updated while $\lambda^{(0)}$ remains fixed. At iteration $t = 1, 2, \ldots$, the rank is first updated over its candidate grid,

$$K^{(t)} = \arg\min_{K \in \mathcal{K}} \mathrm{CV}\left(K, \lambda^{(0)}, \theta^{(t-1)}\right).$$

The smoothing parameters are then jointly optimized on the $\log_{10}$ scale while holding $K = K^{(t)}$ and $\lambda = \lambda^{(0)}$ fixed. Writing $\eta = \log_{10}\theta$,

$$\eta^{(t)} = \arg\min_{\eta} \mathrm{CV}\left(K^{(t)}, \lambda^{(0)}, 10^{\eta}\right), \qquad \theta^{(t)} = 10^{\eta^{(t)}}.$$

The continuous optimization over $\eta$ is performed using the `BOBYQA` algorithm[2] implemented in `nlopt`[3].

After each complete iteration, convergence is assessed from both the change in the smoothing parameters and the change in prediction error. The parameter-change criterion is

$$\Delta_x^{(t)} = \left[\frac{1}{L}\sum_{\beta=1}^{L}\left\{\log_{10}\theta_\beta^{(t)} - \log_{10}\theta_\beta^{(t-1)}\right\}^2\right]^{1/2},$$

and the objective-change criterion is

$$\Delta_f^{(t)} = \left|\mathrm{CV}\left(K^{(t)}, \lambda^{(0)}, \theta^{(t)}\right) - \mathrm{CV}\left(K^{(t-1)}, \lambda^{(0)}, \theta^{(t-1)}\right)\right|.$$

Optimization terminates when either

$$\Delta_x^{(t)} \le \varepsilon_x \qquad \text{or} \qquad \Delta_f^{(t)} \le \varepsilon_f,$$

where $\varepsilon_x$ and $\varepsilon_f$ are the prescribed parameter- and objective-convergence tolerances, respectively, or when the maximum number of coordinate-wise iterations is reached.

### 6 Simulations

We evaluated SSANOVA-SVD using simulated high-dimensional datasets containing multiple structured biological processes, along with nuisance batch variation. The simulation was designed to provide known ground truth at both the sample and feature levels, permitting evaluation of recovery of biological variation, and inference of feature participations.

Briefly, samples were simulated over three relevant covariates – two continuous, *time* and *severity*, and one discrete, *group* – together with a nuisance covariate, *batch*. A fraction of features were designated as signal features and allowed to vary over the time ×severity× group design, while the remaining features were treated as nulls. Four biological processes were simulated, and correspondingly, the signal features were partitioned into four mutually disjoint modules, each participating in exactly one process. The first process induced group-specific shifts in the mean levels of its participating features. The remaining three processes were oscillatory with respect to time: an oscillation shared across groups, an oscillation with group-specific phase shifts, and an oscillation whose amplitude varied with sample-specific severity. Finally, nuisance variation due to batch was overlaid on the resulting biological signal.

#### 6.1 Sampling design

**Group** We simulated $p$ features across $G$ groups, with $n_g$ samples per group, for a total number of samples, $n = Gn_g$.

**Time** Samples were assigned to a regular temporal grid, $t^*_{gr}$ , subsequently jittered for simulating time imbalance to yield the final sampling times $t_{gr}$. Let

$$t^*_{gr} = \frac{2\pi(r-1)}{n_g - 1}, \qquad r = 1, \dots, n_g. \tag{25}$$

To induce departures from balanced temporal sampling, independent Gaussian perturbations were added to the sampled time points while retaining support over the same temporal domain, $t_{gr} = \left(t^*_{gr} + \epsilon_{gr}\right) \bmod 2\pi$, $\epsilon_{gr} \sim N(0, \sigma_t^2)$. The resulting time values were sorted within each group, and were assigned as sampling times for each sample within the group.

**Severity** Each sample was assigned a continuous severity covariate as follows. For each sample $i$, let $s^*_i \sim N(0, 1)$. The resulting $s^*_i$ values were standardized across all samples to have mean zero and unit variance, yielding the final severity variable $s_i$.

**Batch** Each sample was assigned to one of $N_{batch}$ technical batches. Two candidate batch assignments were first generated, and then mixed for the degree of confounding desired.

First, a random assignment was sampled independently for each sample,

$$b_i^{(R)} \sim \text{Uniform}\{1, \dots, N_{batch}\}, \tag{26}$$

Second, a confounded batch assignment was defined deterministically by mapping groups cyclically to the batches, as

$$b_i^{(C)} = [(g_i - 1) \bmod N_{batch}] + 1. \tag{27}$$

For each sample, an indicator

$$C_i \sim \text{Bernoulli}(\gamma) \tag{28}$$

was then generated, where $\gamma$ determines the rate of confounding. The final batch assignment was set as:

$$b_i = \begin{cases} b_i^{(C)}, & C_i = 1, \\ b_i^{(R)}, & C_i = 0. \end{cases} \tag{29}$$

Thus, $\gamma = 0$ corresponds to randomized design over batches, whereas $\gamma = 1$ assigns batch deterministically from biological group. Intermediate values give the probability that an individual sample receives a confounded batch assignment.

#### 6.2 Relevant processes, and feature participations

Relevant signal was distributed across $M = 4$ processes and feature modules.

##### 6.2.1 Relevant processes

The **first** process induces group-specific differences in the average levels of the participating features. Let $\mu_g \sim N(0, \sigma_M^2)$, $g = 1, \ldots, G$. Let $g_i$ denote the group to which sample $i$ belongs. We set

$$f_1(i) = \mu_{g_i} \tag{30}$$

The **second** process is a module-specific oscillatory process parameterized by time, and shared across biological groups. For sample $i$,

$$f_2(i) = A_2 \cos(t_i + \phi_2). \tag{31}$$

where the amplitude $A_2 = |a_2|$ with $a_2 \sim N(\sigma_M, 0.25^2)$, and the phase $\phi_2 \sim \text{Uniform}(-\pi, \pi)$.

The **third** process is an oscillation with group-specific phases. For sample $i$,

$$f_3(i) = A_3 \cos\left(t_i + \phi_{3,g_i}\right). \tag{32}$$

where $A_3$ is sampled as before, and group-specific phase shifts were drawn independently as $\phi_{3g} \sim N(0, \sigma_\phi^2)$, $g = 1, \ldots, G$.

The **fourth** process represented a temporal oscillation, the amplitude of which depended continuously on severity. For sample $i$,

$$f_4(i) = (A_{04} + A_{14} s_i) \cos(t_i + \phi_4). \tag{33}$$

where the amplitude baseline, severity-dependent amplitude coefficient, and phase were generated as

$$A_{04} = |a|\,, a \sim N(\sigma_M, 0.25^2) \tag{34}$$
$$A_{14} \sim N(\sigma_M, 0.25^2), \tag{35}$$
$$\phi_4 \sim \text{Uniform}(-\pi, \pi), \tag{36}$$

The severity-modulation coefficient $A_{14}$ was centered at $\sigma_M$ rather than zero, producing an appreciable severity-dependent change in amplitude.

We note that although cosine functions were used to generate the temporal processes, these functional forms were not supplied to SSANOVA-SVD during model fitting. Temporal and interaction effects were estimated using the same nonparametric SSANOVA representation used in the data applications.

##### 6.2.2 Feature participations

Let $\pi_{\text{signal}}$ denote the fraction of features affected by the relevant processes. The number of features in each module was set to $p_m = \left\lfloor \frac{p\pi_{\text{signal}}}{M} \right\rfloor$. Participating features in each module were sampled sequentially without replacement, so the modules were disjoint. The union of the $M$ modules defined the relevant signal set, while all remaining measured features were nulls.

The simulation contained exactly one realization each of four process types mentioned in the previous subsection: a group-specific mean effect, a shared temporal oscillation, a temporal oscillation with group-specific phase shifts, and a severity-modulated temporal oscillation. For feature $j$ assigned to module $m$, $m \in \{1, \ldots 4\}$, a feature-specific loading magnitude was generated as

$$a_{jm} \sim N(1, 0.15^2). \tag{37}$$

An independent random sign

$$r_{jm} \in \{-1, +1\}, \qquad P(r_{jm} = 1) = P(r_{jm} = -1) = \frac{1}{2}, \tag{38}$$

was then applied, giving the final loading

$$\ell_{jm} = r_{jm} a_{jm}. \tag{39}$$

###### 6.2.3 Mapping biological processes to measured features

For each biological module $m$, $m \in \{1, \ldots, 4\}$, the corresponding sample-level process vector is available as:

$$\mathbf{f}_m = (f_m(1), \ldots, f_m(n))^\top. \tag{40}$$

For feature $j$ belonging to module $m$, its unscaled biological contribution to sample $i$ was $B_{ij}^{(0)} = \ell_{jm} f_m(i)$. Because the biological modules were disjoint, every biological feature loads on to exactly one relevant process. Features outside all modules experience no relevant effects: $B_{ij}^{(0)} = 0$.

Finally, the complete biological signal matrix was multiplied by a global signal-strength parameter,

$$B_{ij} = c_{\text{bio}} B_{ij}^{(0)}. \tag{41}$$

##### 6.3 Nuisance variation

Batch effects were applied to a specified fraction of measured features, $\pi_B$. The total number of batch-affected features was $p_B = \text{round}(p\pi_B)$. Overlap between biological and batch-affected features was controlled by an overlap fraction $\xi$. Within each biological module, $\xi$ fraction of features was sampled without replacement and marked as batch affected. The residual number of batch-affected features were sampled from the null features. So $\xi = 0$ produced disjoint signal and batch-affected feature sets, whereas larger values introduced technical contamination of signal features.

For every batch-affected feature $j$ and batch $b$, its unscaled batch effect was generated as

$$\beta_{jb} \sim N(0, \sigma_B^2) \tag{42}$$

For simplicity, we set $\sigma_B = \sigma_M$ defined in the previous subsection, and controlled the overall strength with a scale parameter $c_{\text{batch}}$, so that the final technical contribution for feature $j$ and sample $i$ was

$$T_{ij} = c_{\text{batch}} \beta_{j, b_i} \tag{43}$$

for batch-affected features, and zero otherwise.

##### 6.4 Feature-specific Gaussian noise

For feature $j$, a variance multiplier was sampled as

$$o_j \sim \text{Gamma}(a, a), \tag{44}$$

using the shape–rate parameterization. The feature-specific residual variance was then set to:

$$\sigma_j^2 = \sigma^2 o_j. \tag{45}$$

Because

$$E(o_j) = 1, \tag{46}$$

the average feature-specific residual variance was $\sigma^2$, while $a$ controlled heterogeneity among features. Conditional on $\sigma_j^2$, residual errors were generated independently as

$$\epsilon_{ij} \sim N(0, \sigma_j^2). \tag{47}$$

For example, when $a = 1$, $o_j \sim \text{Gamma}(1, 1)$, corresponding to an exponential distribution with mean one, resulting in substantial variation in residual variance across features.

##### 6.4.1 Training and held-out test data

After simulation, samples were randomly divided into training and held-out test sets. The test set contained

$$n_{\text{test}} = \text{round}(0.30n) \tag{48}$$

samples drawn uniformly without replacement from all simulated samples, and the remaining samples formed the training set. The split was not stratified by biological group, technical batch, time, or severity.

#### 6.5 Default feature configuration

Simulations contained $p = 5{,}000$ measured features, five biological groups, five technical batches, and four biological modules. When the total biological-signal fraction was set to $\pi_{\text{signal}} = 0.10/4$, each of the four biological modules contained

$$\left\lfloor \frac{5000 \times 0.10/4}{4} \right\rfloor = 31 \tag{49}$$

features, giving 125 signal features in total, with rest nulls.

When the batch fraction $\pi_B = 0.40$, 2,000 of the 5,000 measured features were affected by technical batch variation. Setting the overlap fraction $\xi = 0$ made these technical features disjoint from the signal features.

#### 6.6 Factorization benchmark

Each simulated dataset was analyzed using SSANOVA-SVD and the competing factorization methods. For methods that do not allow measured confounders to be separately input, mirroring practice, the data were apriori adjusted for the confounder design. The generating biological processes, biological-module assignments, technical effects, and other ground-truth quantities were retained for benchmark evaluation and were not supplied to the model fitting routines.

Factorization methods were evaluated at a common rank $K$, where $K$ was the maximum number of components returned by SSANOVA-SVD. Let $\mathbf{B} \in \mathbb{R}^{n \times p}$ denote the true simulated biological contribution and let $\widehat{\mathbf{B}}$ denote the biological contribution reconstructed by a factorization method using its first $K$ components. For a set of features $\mathcal{S}$, reconstruction error was defined as the mean squared error per feature and sample,

$$\text{RE}(\mathcal{S}) = \frac{\left\| \mathbf{B}_{\cdot,\mathcal{S}} - \widehat{\mathbf{B}}_{\cdot,\mathcal{S}} \right\|_F^2}{|\mathcal{S}|\, n}.$$

Reconstruction error was calculated separately for biological signal features, $\mathcal{S}_{\text{bio}}$, and biological null features, $\mathcal{S}_0$. Because $\mathbf{B}_{\cdot,\mathcal{S}_0} = \mathbf{0}$ by construction, null-feature reconstruction error reduces to

$$\text{RE}(\mathcal{S}_0) = \frac{\left\| \widehat{\mathbf{B}}_{\cdot,\mathcal{S}_0} \right\|_F^2}{|\mathcal{S}_0|\, n},$$

and measures biological structure spuriously assigned to null features.

Feature prioritization was evaluated from the estimated feature loadings. For feature $j$, we defined its loading score across the first $K$ components as

$$s_j = \max_{1 \le k \le K} |\widehat{v}_{jk}|.$$

Biological-signal AUC was the area under the receiver operating characteristic curve obtained using $s_j$ to distinguish biological signal features from non-signal features. Equivalently,

$$\mathrm{AUC}_{\mathrm{bio}} = \Pr(s_{j_1} > s_{j_0}) + \frac{1}{2}\Pr(s_{j_1} = s_{j_0}), \qquad j_1 \in \mathcal{S}_{\mathrm{bio}}, \quad j_0 \notin \mathcal{S}_{\mathrm{bio}}.$$

Batch-feature AUC was defined analogously,

$$\mathrm{AUC}_{\mathrm{batch}} = \Pr(s_{j_1} > s_{j_0}) + \frac{1}{2}\Pr(s_{j_1} = s_{j_0}), \qquad j_1 \in \mathcal{S}_{\mathrm{batch}}, \quad j_0 \notin \mathcal{S}_{\mathrm{batch}},$$

where $\mathcal{S}_{\mathrm{batch}}$ denotes the set of features affected by batch. An AUC of 0.5 corresponds to chance-level discrimination, whereas an AUC of 1 corresponds to complete separation.

##### 6.7 Inference benchmark for the 3-step synthesis framework

Conditional inference on feature participations from SSANOVA-SVD reconstructed components was evaluated using the known biological module membership of every measured feature. Features belonging to any of the simulated biological modules were treated as true biological alternatives, whereas all remaining biological-null features were treated as nulls for inference evaluation.

Inference was performed on held-out observations, preserving the separation between factorization and conditional inference used throughout the study. For each feature, limma was used to regress measurements from the unseen test data on the predicted component scores. To test for association with any of the reconstructed components, we used the omnibus $F$-test. Discoveries were defined using Benjamini–Hochberg-adjusted $p$-values at a false discovery rate threshold of 10%.

Let $\mathcal{D}$ denote the discovered feature set, $\mathcal{S}$ the set of truly biological features, and $\mathcal{N}$ the set of biological null features. Empirical power was calculated as

$$\mathrm{Power} = \frac{|\mathcal{D} \cap \mathcal{S}|}{|\mathcal{S}|}, \tag{50}$$

and empirical type-I error as

$$\text{Type I error} = \frac{|\mathcal{D} \cap \mathcal{N}|}{|\mathcal{N}|}. \tag{51}$$

For simulations with at least one discovery, the false discovery proportion was

$$\mathrm{FDP} = \frac{|\mathcal{D} \cap \mathcal{N}|}{|\mathcal{D}|}, \tag{52}$$

which was summarized across simulation replicates to assess empirical false discovery behavior. Discrimination between biological and null features was additionally quantified using the area under the receiver-operating-characteristic curve derived from the corresponding feature-level inference statistics.

##### 6.8 Benchmark configurations

For the benchmarking experiments, we simulated $p$ = 5,000 features across five biological groups and five batches, with four biological processes and 2.5% (125) of features designated as biological signal features. We varied the number of samples per group ($n_g \in \{10, 20, 40\}$), the degree of confounding between biological group and batch ($\gamma \in \{0, 0.5\}$), and the fraction of biological signal features also affected by batch ($\xi \in \{0, 0.5\}$). Batch effects were introduced into 40% of all features. Biological and batch effect scales were fixed at $c_{\mathrm{bio}} = c_{\mathrm{batch}} = 2$, the feature-noise shape parameter was fixed at $a = 1$, and the time-imbalance parameter was fixed at 0.5. Severity was simulated independently of biological group. Each combination of simulation parameters was evaluated across three independently simulated replicates. The primary benchmark used $\xi = 0$, while the corresponding supplementary benchmark used $\xi = 0.5$ to evaluate performance when biological signal features were additionally affected by batch.

### 7 Benchmark methods

We compared SSANOVA-SVD with matrix factorization methods spanning unsupervised, supervised, sparse and covariate-aware approaches, including principal component analysis (PCA), non-negative matrix factorization (NMF), partial least squares (PLS), sparse PCA, sparse NMF and MEFISTO. All benchmark methods were fitted to the same simulated datasets. Features were standardized before model fitting. SSANOVA-SVD modeled nuisance batch variation directly through its nuisance design. For methods without an explicit nuisance component, known batch-associated variation was first removed by feature-wise regression on the known nuisance design, and the residualized measurements were supplied to the factorization method.

To avoid confounding differences in performance from differences in model rank, all methods were evaluated using a common representational budget. The number of processes $K$ was selected by cross-validation using SSANOVA-SVD, and each benchmark method was evaluated using its first $K$ components. Components were retained in their native order without matching or reordering using the simulated ground truth. For methods with scale-indeterminate factorizations, feature-loading vectors were normalized to unit Euclidean norm and the corresponding scale was transferred to the component scores, preserving the reconstructed measurements.

PCA was fitted using `stats::prcomp` in R. NMF was fitted using `sklearn.decomposition.NMF` in `scikit-learn`. Because NMF requires non-negative input, the preprocessed measurements were shifted to non-negative values before fitting and transformed back to the original benchmark scale for reconstruction. PLS was fitted using mixOmics::pls in regression mode[4], with the data matrix supplied as the predictor block and the model matrix corresponding to the biological covariates used by SSANOVA-SVD supplied as the response block. Sparse PCA was fitted using mixOmics::spca[4], retaining 1%, 25% or 75% of the measured features in each component. Sparse NMF was fitted using `sklearn.decomposition.NMF` with an $L_1$ penalty on the feature-loading matrix, using regularization parameters $\lambda \in \{10^{-3}, 10^{-2}, 10^{-1}\}$.

MEFISTO[5] was included as a covariate-aware factorization method. Because the MEFISTO implementation operates on a multi-view representation, the single simulated measurement matrix was supplied as two identical views, and the representation from one view was retained for evaluation. Numeric continuous covariates represented in the SSANOVA-SVD biological model were supplied as MEFISTO covariates, while categorical covariates could not be incorporated into its covariate model.
